# Biological Systematic of the genus *Notodiaptomus* Kiefer 1936 (Copepoda: Calanoida)

**DOI:** 10.1101/2024.07.23.604319

**Authors:** Luis Geraldes-Primeiro, Mauro J. Cavalcanti, Edinaldo N. Santos-Silva

## Abstract

The present effort is a systematic review of the most diverse diaptomid copepods from the South America, the *Notodiaptomus* Kiefer 1936. We proposed to review and identify new morphological characteristics for 37 currently accepted species, assess nomenclatural status, and to examine their relatedness relations. The identification of 2700 morphological hypotheses for males and females are encoded in DEscription Language for TAxonomy (DELTA), used for a historical evaluation of the genus, and arrive at the first parsimony-based phylogeny for *Notodiaptomus*. We present synonymy of all species, status of type material, updated record of occurrences, taxonomic remarks, and dichotomous and interactive key for males and females. The rescue of the common and exclusive ancestor based on 80 synapomorphic characters recovers the monophyly of the organisms of the considered species, supports the discussion of the exclusive conditions and the historically indicated taxonomic relationships. Through this we wish to help establish an additional step on the consistency of the taxonomy of these species, contributing to answer important questions about the origin of some of the characteristics presented.

## Introduction

The genus *Notodiaptomus* Kiefer 1936 represents one of the most significant groups of Neotropical aquatic biodiversity, the Calanoids (Santos-Silva *et al*., 1999). The order Calanoida Sars 1903 constitutes the most numerous basis of metazoans (Walter & Boxshall, 2023) and is associated with the mechanisms of energy transfer, nutrient transport, biomass regulation of primary producers and stocks of planktophagous organisms (Gaviria & Aranguren-Riaño, 2019). Among the calanoids, *Notodiaptomus* is the most widely distributed and diverse set in the Neotropical region (Santos-Silva *et al*., 2013), therefore, it is often related to these mentioned importance (Melo *et al*., 2006; De-Carli *et al*., 2018).

However, the confusing taxonomic history and subjective knowledge about the morphology of this set of organisms have made it difficult to recognize its actual biological diversity. This problem represents a common thread for inaccurate answers about *who they are*, *where they are*, *why they are*, and *how to preserve them*. Otherwise, this scenario not only compromises the documentation of biological diversity but results in the precarious provision of evidence that supports molecular, ecological, biogeographic understandings and conservation strategies of the aquatic ecosystems to which they are related.

Although the problem associated with the morphological knowledge of *Notodiaptomus* is current, its origin dates to before its creation by Kiefer in 1936, with *N. gibber* (Poppe 1889) being, currently, an older accepted species and founded 47 years earlier. Over time, questions about the evidence that supported this grouping (Wright, 1936; 1937) called into question its taxonomic validity. Even with this, new organisms continued to be added to the group, some with more careful descriptions (*i.e*., with objective and clear attributes), detailed illustrations, but most with serious deficiencies in this aspect.

The effects of these inconsistencies have contributed to the heterogeneity of the grouping and caused problems to the taxonomic stability of the genus. Additionally, the inclusion of new species from nomenclatural recombination without valid grounds (*i.e.*, the presentation of unequivocal scientific evidence) accentuates these unintended effects. Paggi (2001) also adds that the fragmented and incomplete character of most of the available descriptions render several of these taxonomic problems irresolute. Nevertheless, interpretations of intra- and inter-population variability of diagnostic characters are overestimated and mistakenly based distinctions, although the effect is reversed in the face of underestimates (Paggi, 2001).

Despite these observations related to the inconsistencies of *Notodiaptomus*, the efforts of Kiefer (1936; 1956) still represent the most current scientific system to the present day. Unfortunately, since then, any comprehensive dedication to the taxonomy of all members of the genus has been published, with the designation of the type-species (Santos-Silva *et al*., 1999) and the review of the *nordestinus* complex (Wright 1935) (Santos-Silva *et al*., 2015) the greatest approximations currently offered. The weight of the lack of other collaborations on the systematics of this genus keeps its morphological limits obscure, which increases the uncertainties about its real variability, originality, and true distribution among ecosystems.

Thus, the study presented below starts from the morphological prism to analyze the biological diversity of the genus *Notodiaptomus.* Through taxonomy and phylogeny, we seek to offer a proposal of objective, integral, consistent taxonomic identity based on formal nomenclatural guidelines. To this end, we offer a taxonomic review, starting from the nomenclatural examination in the light of the ICZN (1999), morphological redescription, taxonomic comments, updated geographic distribution list, and dichotomous identification keys for males and females. Finally, a hypothesis of the relatedness relations between the organisms of the species considered is presented and discussed.

## Materials and methods

### Approaching object

The taxonomic history of the genus *Notodiaptomus* is supported since before its founding by Kiefer in 1936 and is confused with the trajectory of the first diaptomids described for the Neotropical region. Until 1933 all the Diaptomidae Baird 1850 described from South America were called *Diaptomus* Westwood, 1836. This later turned out to be a problem, as those organisms belonging to this genus does not actually occur in the Neotropical region (Santos-Silva *et al*., 2008). Despite this, even today some taxonomists use the artifice of including in “Diaptomus” (sensu lato) all those Diaptomidae for which it is not possible to define the genus with certainty, such is the uncertainty and imprecision existing in the taxonomic determination of this group.

The first copepod described from South America was *Diaptomus* (s.l.) *brasiliensis* Lubbock 1855, from Patagonia, later recombined to *Pseudoboeckella* Mrázek 1901 from Centropagidae Giesbrecht 1893. Absolutely, the first legitimate Diaptomidae was described from Brazil by Poppe as *Diaptomus* (s.l.) *gibber* Poppe 1889. Poppe described another species, *D. deitersi* Poppe 1891 for the same region. The first Calanoida copepod of the Amazonia was *Diaptomus* (s.l.) *henseni* Dahl 1894. In 1897, Richard described *Diaptomus* (s.l.) *bergi* Richard 1897, and Mrazek *Diaptomus* (s.l.) *michaelsoni* Mrazek 1901. In the same year, Sars described other three *Diaptomus* (s.l.): *D. furcatus*, *D. conifer*, and *D. coronatus*. More two other species were described by Van Douwe: *D.* (s.l.) *gracilipes* Van Douwe 1911, and *D.* (s.l.) *aculeatus* Van Douwe 1911. Wright (1927) thought the latter species was identical to *D. furcatus* Sars 1901, probably. Tollinger (1911) published a study on the distribution of the Diaptomidae, which then totaled ten species of them. *Diaptomus* (s.l.) *marshi* Marsh 1913 was described from Guatemala, thus, twelve species were already known for the Neotropical region.

Wright (1927) reviewed the species of *Diaptomus* (s.l.) from South America and described nine new species. From this work, until 1937, he described other new species of *Diaptomus* and, in 1938, summarized all knowledge about this group in South America. In 1933, Brehm proposed the genus *Argyrodiaptomus* to include *D. bergi*, *D. furcatus*, *D. aculeatus*, *D. denticulatus* Pesta 1927, and a new species that he designated as the genus type, *A. granulosus* Brehm 1933. Three years forward, Kiefer (1936) proposed the division of several *Diaptomus* (s.l.) from South America through the founding six new genera: *Notodiaptomus*, *Rhacodiaptomus*, *Dactylodiaptomus*, *Calodiaptomus*, *Odontodiaptomus*, and *Idiodiaptomus*. This decision was contrary to Wright (1927), that considered inappropriate the formal division of South American species into taxonomic groups. During his study of the diaptomids of South America (Wright, 1927), the author considered the existence of closely relatable organisms, but others with insufficient morphological information for non-monospecific groups.

Despiteful, Wright had already verified some related groups were related, some being more homogeneous than others. In 1935, Wright more clearly admitted the existence of these groups to which he referred in the 1927 work and proposed to gather some *Diaptomus* (sensu lato) into a complex of forms denominated “*nordestinus* complex.” The morphological convergence identified by Wright (1935) was for the fifth swimming leg, exclusively. For males: (1) right basis with conspicuous internal proximal angle “like a rounded shoulder” (*i.e.*, intumescence), with or without a small prominence; (2) right endopod small and conical; (3) right exopod 2 with inner margin with an angular or rounded prominence (*i.e.*, curved ridge posteriorly), best present in lateral view; (4) left exopod with two closely fused segments, each of them carries a pad with hair (*i.e.*, inner lobe); (5) left exopod 2 ending in a digitiform process with small “spiniform arrow“; and (6) left endopod unisegmented. For female fifth swimming legs: (1) basis with outer seta inserted on prominence; and (2) basis with distal angular expansion posteriorly.

Originally, the grouping consisted of *D. nordestinus*, *D. henseni*, *D. iheringi*, *and D. deitersi.* Wright (1936) expanded the group to 11 species, considering the integration of: *D. amazonicus*, *D. cearensis*, *D. dahli*, *D. isabelae*, *D. jatobensis*, *D. conifer*, and *D. inflatus.* Even so, the author thought it premature to propose new formal genus for these group.

Notwithstanding, Kiefer (1936) used the “*nordestinus*” group as a basis for proposing the creation of the genus *Notodiaptomus.* In the proposal, Kiefer included seven species from Wright’s group and another fourth species of *Diaptomus* (sensu lato): *D. santaremensis* Wright 1927, *D. carteri* Lowndes 1934, *D. anisitsi* Daday 1905, and *D. incompositus* Brian 1926. Although never accepted this proposal, Wright (1937) agreed to include *D. anisitsi*, and *D. incompositus* to the “*nordestinus*” complex. Thenceforth, the genus had been formed for 13 species, all currently recognized as *Notodiaptomus* and integrated by three additional characteristics: (1) “nature” of the male fifth left leg exopod 2 distally; (2) male right antennule with “spiniform” protuberances in segments 10, 11, 13, 15 and 16 (*i.e.*, modified seta on actual segments 10, 11, and 13; spinous process on actual segments 15 and 16); and (3) male right antennule actual segment 20 with hyaline lamella.

In the following year, Wright (1937) verified, at that time, that 47 species of *Diaptomus* (sensu lato) were already known for South America. In practice, the author understood that his opposition to decision of Kiefer (1936) was ineffective but reinforced the need for complete and objective descriptions as presuppositions for the acceptance of safe proposals. For Wright, it was imprudent to accept Kiefer’s genus because of the unavailability of adequate diagnoses.

A gap of 19 years lasted until the inclusion of a new species in *Notodiaptomus*. Kiefer introduced to science *N. maracaibensis* Kiefer 1954, and two years later expanded the genus as four more species of *Diaptomus* (s.l.): *D. conifer* Sars 1901, *D. dahli* Wright 1936, *D. isabelae* Wright 1936, and *D. jatobensis* Wright 1936. At the time, a new morphological set was added to the diagnosis of the genus: (1) female fifth swimming legs with endopod unisegmented; (2) female fifth swimming legs endopod with row of oblique spinules plus 1 or 2 setae; (3) male fifth swimming legs coxa with rounded lobule and “hyaline spine” (*i.e.*, sensilla); (4) male fifth right swimming leg basis with structure in varied shape and size, lamella posteriorly (*i.e.*, posterior protrusion); (5) male fifth right swimming leg exopod 1 with length greater than width; (6) male fifth right swimming leg exopod 2 with rounded-broad hyaline protuberance on inner “caudal surface” (*i.e.*, inner distal curved ridge posteriorly); (7) male right swimming leg exopod 2 with outer spine on base of the terminal claw generally; (8) male left swimming leg exopod 2 with distal portion digitiform; (9) male left swimming leg exopod 2 with short broad seta, directed inwards, on distal portion; (10) male right antennule without “spiniform protuberance” on segment 14 (*i.e.*, spinous process); and (11) male right antennule with narrow hyaline membrane on segment 20 generally (*i.e.*, hyaline spinous process).

Although the efforts of Kiefer (1936; 1956) contain numerous shortcomings, it is the only one available to date. Any in-depth incursion into *Notodiaptomus* has been published since his effort and the weight of the lack of a work dedicated to the revision of the genus keeps its morphological limits obscure. Undoubtedly, this has increased the inconsistency about the real variability of the grouping, sustaining uncertainties about its originality and distribution among ecosystems faithfully.

Brandorff (1976) carried out important contribution about the geographical distribution of Diaptomidae in South America. Undoubtedly this was a milestone in the study of these organisms. However, we must be sure of the need to make a new attempt to study the distribution of Diaptomidae, determine the existing patterns and try to explain them with objective scientific foundations.

Over time several new species were added to the genus, but no one made any revisions or more fully defined what the genus *Notodiaptomus* was. Therefore, this group apparently became less and less homogeneous, which led Dussart (1985) to propose four subgenus: *Notodiaptomus*, *Wrightius*, *Caleodiaptomus*, and *Amazonius*. However, he reproduced the same mistake as Kiefer (1936) in not making any diagnosis of these subgenus. Reid (1987) had already reported this deficiency and indicated the non-conformity of Dussart’s proposal (1985), through Article 13 of the International Code of Zoological Nomenclature (ICZN, 1985; 1999).

In this same contribution, Reid (1987) commented on the unique diagnosis given by Dussart (1985), subgenus *Notodiaptomus*: “The entire diagnosis of the subgenus is: with exopod article 2 of the left leg 5 of male *’à soie spiniforme droite ou à peine courbée, dressée et court’*” (Reid, 1987, p. 378). Such a diagnosis does not seem to be sufficient to separate this group from the other species of the genus. For Santos-Silva *et al*. (2015), perhaps, one way forward might be to recover the diagnoses made by Wright (1935) for the *nordestinus* complex and by Kiefer (1936; 1956) for *Notodiaptomus*.

Santos-Silva *et al*. (1999) when verifying exactly these inconsistencies for the *Notodiaptomus* grouping, carried out a detailed study on *Notodiaptomus deitersi* (Poppe 1891) and designated it type-species. In this study, the authors provided an amendment to the description of the species under standards of modern accuracy, designating the neotype and bringing consistent improvements to the knowledge about the genus. A few years later, Santos-Silva (2008) again offered a new contribution to the clarification of knowledge about the group to proposing the distribution and history of genus into Diaptomidae, Pseudodiaptomidae and Centropagidae from Brazil, where 24 species of *Notodiaptomus* are presented through a complete bibliographical review, geographical distribution, and pertinent observations. Another relevant contribution come from Perbiche-Neves *et al*. (2015), who analyzed the diaptomids of the Prata River basin, the second largest in the Neotropical region (South America). In this report, the author presented a morphological approach and discussed 13 species of *Notodiaptomus* with diagnoses, scientific illustrations, and identification keys.

Recently, Santos-Silva *et al*. (2015) brought new advances to the taxonomic status of *Notodiaptomus* through reviewing the history of the species contained in the *nordestinus* complex (Wright 1935), Kiefer’s stated basis for the founding of the genus. In this study, new descriptions were offered for the eleven species, except *N. inflatus*, and *N. dahli*, which could not be reanalyzed due to the lack of the type-material. According to the authors, the successive addition of several species – “probably unrelated” – and the inconsistency in the description of the genus, have contributed to the heterogeneity of *Notodiaptomus* and originated problems related to its taxonomic stability, thus making dedicated studies on the genus indispensable.

Over time, other species were included in *Notodiaptomus*, through taxonomic recombination or original description. The most recent are *N. simillimus* Cicchino, Santos Silva & Robertson 2001, *N. dentatus* Paggi 2001, *N. cannarensis* Alonso, Santos-Silva & Jaume 2017, and *N. nelsoni* Previattelli, Perbiche-Neves & Rocha 2017. Currently, there are 37 species accepted in *Notodiaptomus* (Walter & Boxshall, 2023), which are distributed from Antigua and Barbuda (Netherlands Antilles, Central America) to Argentina, 24 of them occur in Brazil (Santos-Silva, 2008), some with more careful descriptions, well-done and detailed illustrations, but the vast majority still with serious deficiencies in this aspect. In the present, there is already an accumulation of knowledge and information that, together with access to modern techniques and methodologies, have made studies on these organisms increasingly possible and enlightening.

### Specimen sources

The specimens examined during this study were obtained through duplicates of the type-material stored in the collection of the Plankton Laboratory of the Instituto Nacional de Pesquisas da Amazônia, collections or visits to the original biological collection in museum. Priority was given to the study of the type-material of each species, considered in the order holotypes, paratypes, neotypes, syntypes and in another case topotypes deposited in public collections and accessible or collected *in nature*. Prior to the examinations all specimens were confirmed with the support of the original literature, redescriptions, and dichotomous keys were adopted when necessary (*e.g.*, Santos-Silva *et al*., 2015; Perbiche-Neves *et al*., 2015). The state of the type-material, material examined, information of the type-locality, deposit of the testimonial material and synonymous history are offered before the diagnosis and redescription of each species presented, with the addition of the updated geographical records and subsequent notes. Therefore, the specimens considered came from the following collections:

**INPA** Instituto Nacional de Pesquisas da Amazônia (Brazil). Non-insecta Invertebrates Collection.

**MNK** Staatliches Museum für Naturkunde Karlsruhe (Karlsruhe, German). Kiefer Collection.

**MZUSP** Museu de Zoologia da Universidade de São Paulo (Brazil); Carcinologic Collection.

**MNHN** Muséum National d’Histoire Naturelle (Paris, France).

**NHMUK** Natural History Museum (London, England)

**USNM** National Museum of Natural History, Smithsonian Institution (Washington, D.C., USA)

Initially, 37 taxa of *Notodiaptomus* were considered for the predefined analysis stages of the study: (1) evaluation of nomenclatural validity and scientific evidence, (2) morphological redescription of male and female, and (3) relationship test of the organisms of each species. Exception done for *N. difficilis* Dussart & Frutos 1986, *N. dilatatus* Dussart 1984, *N. dahli* (Wright 1936), and *N. anceps* Brehm 1958, with nomenclatural validity checked, but type-material or other references unavailable for morphological redescriptions. The taxa inquirendum *N. spiniger* (Brian 1926), *N. kieferi* Brandorff 1973, *N. ohlei* (Brandorff 1978), and *N. santafesinus* Ringuelet & Ferrato 1967 were re-approximated for further investigation.

All individuals of each species were dissected or factored according to the morphology *sensu* Huys & Boxshall (1991) and mounted on 16 slides between the sex equally divided. The preparation of the slides followed the method presented in Kihara & Rocha (2009), for the semi-permanent assemblies lactophenol was adopted as a medium and Entellan® for permanent sealing. All procedures and analyses were conducted from the Plankton Laboratory (INPA), Carcinofauna Laboratory (MZUSP), and Meiofauna Laboratory (USP), where Leica MZ12.5 trinocular optical stereomicroscope or Leica MZ95 for dissection, and Leica DM 2500 10x-100x or Olympus BH-2 with differential interference contrast for morphological recognized were used.

### Morphological obtaining method

The procedure for obtaining the data followed the morphological factorization of 12 primary parts of each organism: (1) body; (2) antennules; (3) antennae; (4) mandible; (5) maxillulae; (6) maxillae; (7) maxilliped; (8) first swimming legs; (9) second swimming legs; (10) third swimming legs; (l1) fourth swimming legs; and (12) fifth swimming legs. For each factor, its respective segmentation pattern (*sensu* Huys & Boxshall, 1991) was considered as secondary parts, where ornamental and sensory structures (*sensu* Garm & Watling, 2013), shapes, total and relative size, positioning and insertion origin could be characterized attributes. For the morphological interpretation, observation planes were standardized based on the habitus of the organism, determining the anterior, posterior, dorsal, ventral, distal, proximal, inner, and outer planes (Fig. 1).

**FIGURE 1.**
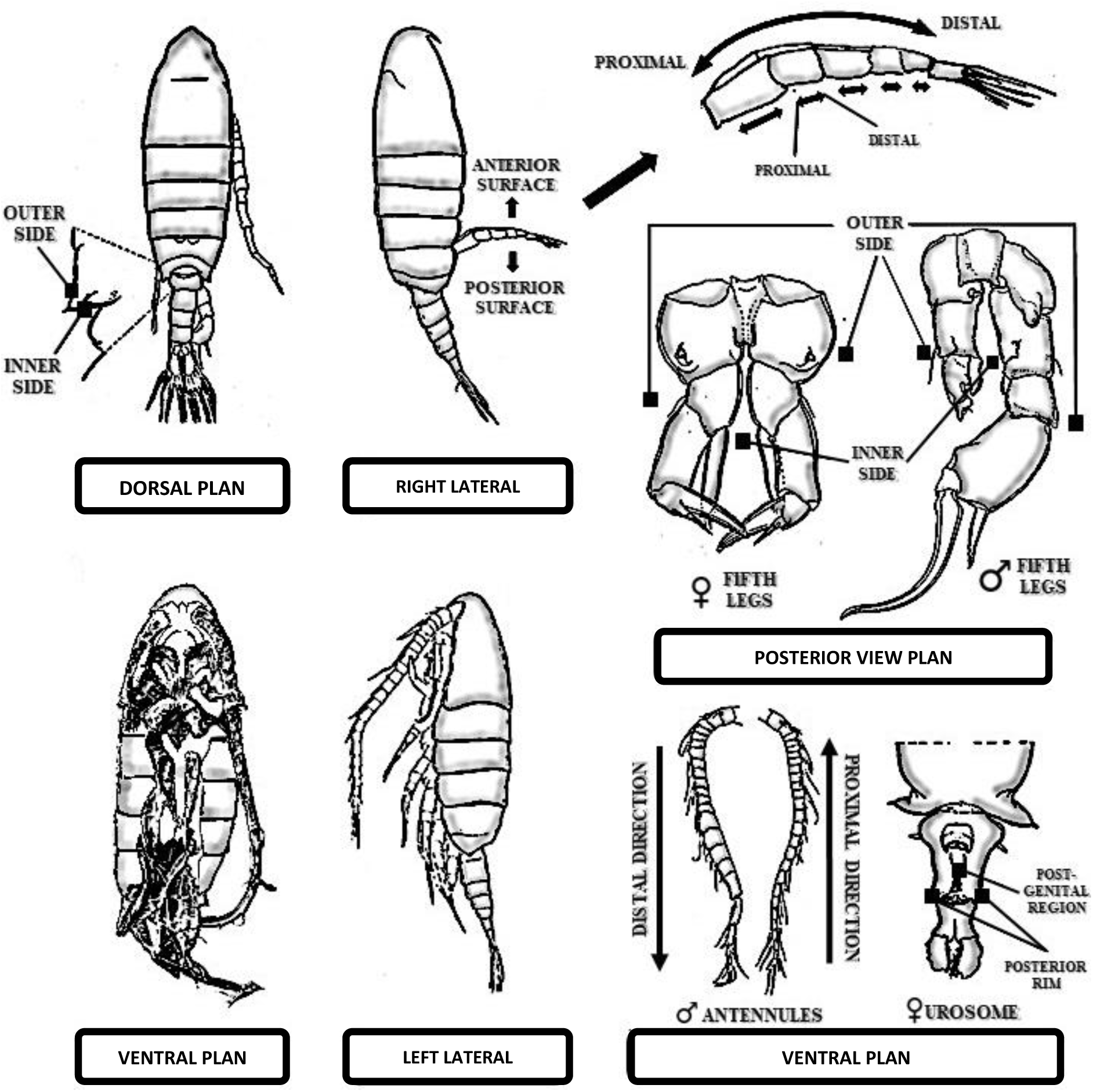
Schematic illustration of the observational patterning (viewing planes) assumed for the morphological interpretations. Adaptation from the originals by D. Previattelli, and E.N. Santos-Silva.

The data obtained through the examinations were coded in DELTA (DEscription Language for TAxonomy), format (Dallwitz, 1980), using the FreeDelta Editor 2.9.7.0 software (Cavalcanti & Santos-Silva, 2009). DELTA-coded data can be translated into several formats for generation of natural-language descriptions, key construction, interactive identification and information retrieval, and phylogenetic analyses (Askevold & O’Brien, 1994; Coleman et al., 2010). The morphological parts and their attributes were encoded as characters and character states, compiled on the basis of ancestral copepod attributes and terminology in Huys & Boxshall (1991). From the data coded in DELTA, we then generated morphological description in natural language and constructed dichotomous identification keys for males and females, as well as creating interactive identification keys using the INTIMATE 1.10 and INTKEY 5.12T programs (Dallwitz, 1993; Dallwitz *et al*., 2010), examined character list, and characters matrix (supplementary material S1, and S2). For editing the schematic illustrations presented to characters with relative size and form, the GIMP 2.10.34 software was used with support of Wacon Intuous Pro-Paper Edition PHT660P creative pen tablet. All schematic illustrations were identified with the number corresponding to the attribute and sequence of states according to the list of characters and considering the morphology of the type-species of *Notodiaptomus* completely illustrated in Santos-Silva *et al*. (1999).

### Analytical methods

The nomenclatural analysis of the species considered was carried out from the investigation of the taxonomic trajectory and the nomenclatural acts submitted. We assume as a basis the prerogatives of the International Code of Zoological Nomenclature (1999) for taxonomic availability and, subsequently, the criteria and scientific evidence presented in each publication analyzed. The taxonomic validation and the verified scientific foundations supported the continuation of the morphological and relatedness relations analyses of the organisms of each species.

For the phylogenetic analysis we considered a database of 37 taxa, including 30 species of *Notodiaptomus* (table 1), the outgroup, and 2700 morphological characters (supplementary material, S1). *N. caperatus* Bowman 1979 was included in this stage of the research and had its information abducted from the original literature, with complete description and objective attributes. Initially, a tree of characters was created using the FreeDelta Editor 2.9.7.0, and output as a nexus file. Mainly of characters were binary, although thirty-two characters has three states. Inapplicable or missing characters were coded ‘?’. Characters were unordered and equally weighted. Despite the number of multistate characters, the non-ordering renders the transition effects between states (1, 2, 3) ineffective and does not imply about polarity or order. The phylogenetic analysis under maximum parsimony was performed in the PAUP 4.0b10 program (Swofford, 2002) and visualized in the FigTree v1.4.4 software (Rambaut, 2016).

**TABLE 1.**
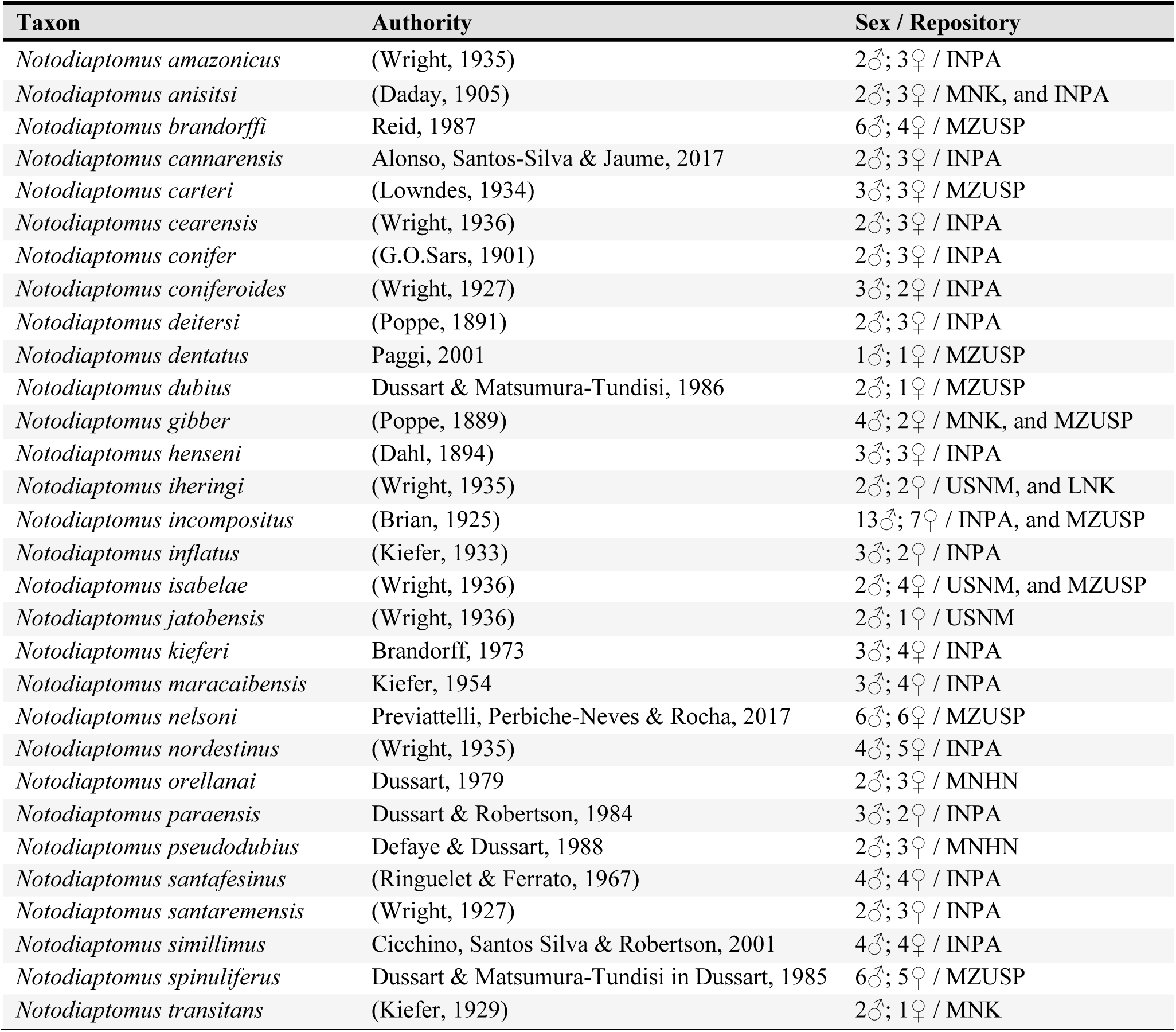
Comparative taxa examined to establish diagnostic features of the genus *Notodiaptomus* Kiefer 1936.

| **Taxon** | **Authority** | **Sex / Repository** |
| --- | --- | --- |
| *Notodiaptomus amazonicus* | (Wright, 1935) | 2♂; 3♀ / INPA |
| *Notodiaptomus anisitsi* | (Daday, 1905) | 2♂; 3♀ / MNK, and INPA |
| *Notodiaptomus brandorffi* | Reid, 1987 | 6♂; 4♀ / MZUSP |
| *Notodiaptomus cannarensis* | Alonso, Santos-Silva & Jaume, 2017 | 2♂; 3♀ / INPA |
| *Notodiaptomus carteri* | (Lowndes, 1934) | 3♂; 3♀ / MZUSP |
| *Notodiaptomus cearensis* | (Wright, 1936) | 2♂; 3♀ / INPA |
| *Notodiaptomus conifer* | (G.O.Sars, 1901) | 2♂; 3♀ / INPA |
| *Notodiaptomus coniferoides* | (Wright, 1927) | 3♂; 2♀ / INPA |
| *Notodiaptomus deitersi* | (Poppe, 1891) | 2♂; 3♀ / INPA |
| *Notodiaptomus dentatus* | Paggi, 2001 | 1♂; 1♀ / MZUSP |
| *Notodiaptomus dubius* | Dussart & Matsumura-Tundisi, 1986 | 2♂; 1♀ / MZUSP |
| *Notodiaptomus gibber* | (Poppe, 1889) | 4♂; 2♀ / MNK, and MZUSP |
| *Notodiaptomus henseni* | (Dahl, 1894) | 3♂; 3♀ / INPA |
| *Notodiaptomus iheringi* | (Wright, 1935) | 2♂; 2♀ / USNM, and LNK |
| *Notodiaptomus incompositus* | (Brian, 1925) | 13♂; 7♀ / INPA, and MZUSP |
| *Notodiaptomus inflatus* | (Kiefer, 1933) | 3♂; 2♀ / INPA |
| *Notodiaptomus isabelae* | (Wright, 1936) | 2♂; 4♀ / USNM, and MZUSP |
| *Notodiaptomus jatobensis* | (Wright, 1936) | 2♂; 1♀ / USNM |
| *Notodiaptomus kieferi* | Brandorff, 1973 | 3♂; 4♀ / INPA |
| *Notodiaptomus maracaibensis* | Kiefer, 1954 | 3♂; 4♀ / INPA |
| *Notodiaptomus nelsoni* | Previattelli, Perbiche-Neves & Rocha, 2017 | 6♂; 6♀ / MZUSP |
| *Notodiaptomus nordestinus* | (Wright, 1935) | 4♂; 5♀ / INPA |
| *Notodiaptomus orellanai* | Dussart, 1979 | 2♂; 3♀ / MNHN |
| *Notodiaptomus paraensis* | Dussart & Robertson, 1984 | 3♂; 2♀ / INPA |
| *Notodiaptomus pseudodubius* | Defaye & Dussart, 1988 | 2♂; 3♀ / MNHN |
| *Notodiaptomus santafesinus* | (Ringuelet & Ferrato, 1967) | 4♂; 4♀ / INPA |
| *Notodiaptomus santaremensis* | (Wright, 1927) | 2♂; 3♀ / INPA |
| *Notodiaptomus simillimus* | Cicchino, Santos Silva & Robertson, 2001 | 4♂; 4♀ / INPA |
| *Notodiaptomus spinuliferus* | Dussart & Matsumura-Tundisi in Dussart, 1985 | 6♂; 5♀ / MZUSP |
| *Notodiaptomus transitans* | (Kiefer, 1929) | 2♂; 1♀ / MNK |

Analyzes were conducted using heuristic search (with 1000 replicates with random input order; branch swapping: tree-bisection-reconnection), character-state optimization through accelerated transformation (ACCTRAN) and a prior rooting. The outgroup was selected in the context of current phylogenetic hypotheses and systematic propositions about the order Calanoida. Pseudodiaptomidae Sars 1902 and Centropagidae Giesbrecht 1893 were deduced sister groups of Diaptomidae in Bradford-Grieve *et al*. (2010) and, respectively, represented by *Pseudodiaptomus* sp. and *Boeckella bergi* Richard 1897. The genus *Rhacodiaptomus* is exclusive cluster for the Amazonia, hypothetically not comprehensive to the inner group with a wide Neotropical distribution. As well as *Rhacodiaptomus*, the genus *Argyrodiaptomus* was contained in the objective of separating the *Diaptomus* (sensu lato) from South America (Brehm, 1933), a hypothesis disputed (Wright, 1936; 1937) and considered for testing. *Diaptomus susanae* Paggi 1976 was cited as close to *Notodiaptomus* (Paggi, 1976) and represents the grouping of Neotropical diaptomids artificially included as *Diaptomus* (sensu lato). *Diaptomus castor* (Jurine 1820), a true representative of the European genus *Diaptomus* Westwood 1836 was defined for polarization of the characters in the goal to test the relationship between the groups historically confused.

## Results and discussion

### Nomenclatural evaluation and taxonomic evidence

Amid the *Notodiaptomus* species evaluated for nomenclatural consistency and taxonomic evidence, six species proposed for the genus *Notodiaptomus* Kiefer 1936 were refuted. They are the taxa *Notodiaptomus echinatus* (Lowndes 1934), *Notodiaptomus lobifer* (Pesta 1927), *Notodiaptomus susanae* (Paggi 1976), *Notodiaptomus ohlei* (Brandorff 1978), *Notodiaptomus oliveirai* Matsumura-Tundisi, Espindola, Tundisi, Souza-Soares & Degani 2010, and *Notodiaptomus spiniger* (Brian 1926). All proposals were initially considered for *Notodiaptomus*, with inconsistencies identified and presented hereafter.

*Notodiaptomus echinatus* (Lowndes 1934) is an attempt at nomenclatural recombination resulting from the proposal to synonymize *Diaptomus* (sensu lato) *echinatus* Lowndes 1934 to the *Notodiaptomus kieferi* Brandorff 1973 (Defaye & Dussart, 1988). For the goals of this proposal, Defaye & Dussart (1988) officially transferred *D.* (s.l.) *echinatus* to *Notodiaptomus echinatus*, without presenting valid evidence that would support both denominations for a same set of biological characteristics. Within the research, it is possible to observe only partial illustrations of the right antennule and fifth male legs and posterior dorsal region of the female habit, through specimens collected in Venezuela. The unavailability of the examined material or other material from the location indicated by the authors make it impossible to confirm the significant morphological variations that can be identified between the illustrative interpretation of Defaye & Dussart (1988) and those of Brandorff (1978) to *N. kieferi*. However, the proposed synonymization between the species is scientifically unfounded as long as it does not present adequate evidence and the transfer of *D.* (s.l.) *echinatus* to *Notodiaptomus* is invalid. Thus, it is natural that the taxon returns to its original taxonomic shelter and is recognized as *Diaptomus* (sensu lato) *echinatus* Lowndes 1934.

*Notodiaptomus lobifer* (Pesta 1927), incurs similar basis to the previous case. This species was presented to science by Pesta from organism from Argentina and recombined for the first time to the genus *Notodiaptomus* by Ringuelet (1958), without offering evidence to justify the proposal. Later, Dussart (1984) when dealing with *Notodiaptomus coniferoides* (Wright 1927) from Venezuela (which would come to be recognized as *Notodiaptomos simillimus* Cicchino, Santos-Silva & Robertson, 2001), mentioned *Diaptomus* (s.l.) *lobifer* as a junior synonym of *N. coniferoides*, again in a scientifically unfounded nomenclatural act. Perbiche-Neves *et al*. (2015) seems to have agreed with Dussart’s proposal, although they also do not offer valid scientific elements. Apparently, all those represent involuntary errors that make the suggested synonymizations and the permanence of the taxon in the genus *Notodiaptomus* unfeasible. Therefore, the species should return to its original temporary shelter, and be recognized as *Diaptomus* (s.l.) *lobifer* Pest 1927.

*Notodiaptomus susanae* (Paggi 1976) was founded through organisms from the lentic water bodies North Argentina and as *Diaptomus* (s.l.) *susanae* Paggi 1976. The inaugural taxonomic presentation has a solid morphological foundation and represents a nomenclatural act that satisfies the requirements of the ICZN (1999). Originally the species was indicated with close proximity to *Notodiaptomus* but inserted into *Diaptomus* sensu lato officially. There have been several attempts to recombine the species as *Notodiaptomus* (Battistoni, 1998; Dussart & Defaye, 2002; Perbiche-Neves *et al*., 2015; Perbiche-Neves *et al*., 2020), all unfounded by the absence of objective and valid scientific evidence. Some of these reports indicate the existence of the academic thesis of Juan Cesar Paggi (1995) as the basis for these attempts, unfortunately an invalid effort as a scientific record in the criteria of the ICZN (1999, Art. 7 and 8). The recent citation of the species as *D.* (s.l.) *susanae* (Paggi, 2011, p. 420) reinforces the author’s original position. For the present effort, we accessed topotypes of the taxon and reaffirmed its morphological proximity to *Notodiaptomus*, to be proven and formally presented through valid scientific evidence.

*Notodiaptomus ohlei* (Brandorff 1978) is a nomenclatural recombination proposed by Dussart (1985) from *Diaptomus* (s.l.) *ohlei* Brandorff 1978. At the time, the author proposed four subgenera for *Notodiaptomus*, among which *Notodiaptomus* (*Amazonius*) was typified by organisms of *D.* (s.l.) *ohlei*. The presentation of Dussart’s new nominal taxa does not contain a formal presentation of diagnosis or any description of differential characters, a direct violation to the International Code of Zoological Nomenclature, article 13.1 (recommendation 13a) for names published after 1930 (ICZN, 1999). It is inevitable, therefore, to recognize a taxonomic proposal surrounded by serious inconsistencies and scientifically invalid. In this way, the taxa defined as types for each subgenus should return to their original taxonomic grouping, such as *N. ohlei* (Brandorff 1978), which should be correctly named *Diaptomus* (s.l.) *ohlei* Brandorff 1978.

*Notodiaptomus oliveirai* Matsumura-Tundisi, Espindola, Tundisi, Souza-Soares & Degani 2010 is a taxonomic proposal with nomenclatural and morphological misinterpretations. It was presented to science based on organisms from the Barra Bonita Reservoir, Tietê River, southeastern Brazil. Perbiche-Neves *et al*. (2015) proposed its recognition as a junior-synonym of *Notodiaptomus henseni* (Dahl, 1894) contesting the variability in the shape of the outer margin of exopod 2 of the fifth right leg of males of both species, an effect exclusively dependent on the observation plan of the morphological fraction. Undoubtedly, morphological descriptions not based on clear and objective references expose taxonomic proposals to subjectivation and scientific irreplicability, unfortunately an increasingly common practice.

Nevertheless, the arguments offered by Perbiche-Neves *et al*. (2015) for the case are consistent and based on robust evidential observations, which presuppose acceptance to the taxonomic proposal in Matsumura-Tundisi *et al*. (2010). However, we believe that the character set presented is subjective, incomplete, and supported by inconsistent evidence throughout its nomenclatural act. The illustrations provided do not correspond to the images presented (*vide* Matsumura-Tundisi *et al*., 2010, figs. 3a and 3b, and 5a and 5b), without mentioning the observation plan or illustration used. Advisedly, this violates ICZN Articles 13.1.1 and Recommendation 13a (1999) and, therefore, the desire for consistency and taxonomic universality. Unfortunately, the proposal is affected by the non-objectivity and clarity of the presentation of characters and other evidence, which truly compromises scientific reproducibility. Thus, even before its recognition as a synonym of *N. henseni*, we assessed the proposed taxa as unsustainable evidently for its acceptance and, therefore, as an invalid name.

*Diaptomus* (s.l.) *spiniger* Brian 1926 was a species founded with organisms from Argentina and has a taxonomic trajectory based on confusions regarding its true identity. Brehm (1933) when creating the genus *Argyrodiaptomus* and based on the presence of “tiny chitinous spikes covering the inner side of the base of the fifth left leg of males, and in many cases also the distal inner side of the right base” (Brehm, 1933: 283, free translation), transferred the species to this group. However, Kiefer (1936) considered this evidence insufficient to maintain it as a true *Argyrodiaptomus*, which, in practice, was only portrayed by Ringuelet & Ferrato (1967) and Dussart & Frutos (1985), when listing *Diaptomus* (s.l.) *spiniger* as a senior-synonym of *Diaptomus* (s.l.) *toldti* Pesta 1927 and *Diaptomus* (s.l.) *birabeni* Brehm 1933, respectively.

Dussart & Defaye (2002) returned to the nomenclature proposed by Brehm and listed *Argyrodiaptomus spiniger* as *incertae sedis*, not attributing minimal scientific evidence that would justify this decision. More recently, Perbiche-Neves *et al*. (2015) redescribed specimens from the Prata River Basin (Southern Neotropical) as *Notodiaptomus spiniger*, alleging as the basis of the comments by Kiefer (1936) and Dussart & Frutos (1985). Apparently, this is an involuntary error, as Kiefer (1936) does not mention the species to *Notodiaptomus*, and in Dussart & Frutos (1985), even though the illustrations are labelled as “*Notodiaptomus spiniger*”, no morphological evidence is presented to support the transference of the species to this genus, an additional fact to its citation as *Diaptomus* (s.l.) throughout the entire discussion in the account.

Notwithstanding this, in all these cases there are direct infractions of the ICZN (1999), especially as stated in Art. 13.1 regarding the need to objectively present characters set that differentiate associated organisms from related taxonomic names, and Art. 16.1 regarding the importance of making explicit the intention to present new taxonomic acts through the use of abbreviations of latinized terms. Thus, supported by this trajectory, we did not consider, for this research, the inclusion of *Diaptomus* s.l. *spiniger* Brian, 1926 as a member of *Notodiaptomus*, which should remain in the original taxonomic shelter until new scientific efforts address its issues.

However, based on the analysis of the nomenclatural validity of the species recognized by science as belonging to the genus *Notodiaptomus*, as well as the availability and access to type specimens of each species, we considered 31 specific hypotheses for the morphological review study, which are the basis for the presentation of the redescription and diagnosis of the genus considering 2700 characters identified between male and female, examined in each species according to origin (table 1).

### Taxonomy

**Order Calanoida Sars G.O., 1903**

**Family Diaptomidae Baird, 1850**

**Subfamily Diaptominae Kiefer, 1932**

#### Genus *Notodiaptomus* Kiefer, 1936

Type-species: ***Notodiaptomus deitersi*** (Poppe 1891), designated in Santos-Silva *et al*. (1999).

##### Diagnosis

**(1)** Male cephalosome with dorsal suture, separated from first metasome segment, and incomplete mostly; **(2)** antennules with vestigial seta on segments 2, 3 and 5; **(3)** male right antennule actual 22-segmented; **(4)** male right antennule actual segment 3, 7, 9 and 14 with one seta surpassing to distal margin beyond three sequential segments, straight and blunt apex, mostly; **(5)** male right antennule with modified seta as a process on segments 10, 11, 13; **(6)** male right antennule segments 15 and 16 with spinous process, mostly; **(7)** male right antennule with modified seta on segment 13 larger than those on 10 and 11; **(9)** male left antennule segment 13 e 15 without aesthetasc; **(10)** left antennule with setae on segment 23 posteriorly; **(11)** left antennule with setae on segment 24 ventrally; **(12)** antenna endopod 2-segmented, mostly; **(13)** antenna first segment with row of spinules and pore, mostly; **(14)** antenna coxa with seta reaching to the endopod 1, mostly; **(15)** first swimming legs endopod 2-segmented, and exopod 2 without outer spine; **(16)** second to fourth swimming leg with double distal row of spinules on endopod 3 anteriorly, mostly; **(17)** male fifth right swimming leg endopodal lobe with spinules on distal-inner rim; **(18)** male fifth right swimming leg with rudimentary praecoxa; **(19)** male fifth right swimming leg coxa with conical process projecting over base, mostly; **(20)** male fifth right swimming leg basis with posterior protrusion, mostly; **(21)** male fifth right swimming leg basis with oblique deep groove, mostly; **(22)** male fifth right swimming leg exopod 1 longer than broad, mostly; **(23)** male fifth right swimming leg with exopod 1 with triangular outer process posteriorly, mostly; **(24)** male fifth right swimming leg exopod 1 with distal process inserted at posterior or marginal surface, mostly; **(25)** male fifth right swimming leg with exopod 2 with distal-internal curve on posterior surface, mostly; **(26)** fifth right leg exopod 2 with angular or rounded prominence on inner margin, generally; **(27)** male fifth right swimming leg exopod 2 with terminal claw curved proximally, mostly; **(28)** male fifth left swimming leg coxa with conical process bearing apex sensilla; **(29)** male fifth left swimming leg exopod 2 predominantly with digitiform distal process, mostly; **(30)** male fifth left swimming leg exopod 2 without transverse row of denticles on distal process, mostly; **(31)** female urosome 3-segmented, mostly; **(32)** female antennules without aesthetics on segments 13 and 15; **(33)** female fifth swimming legs basis with expansion distal inner rim posteriorly, mostly; (**34)** female fifth swimming legs basis with outer seta longer 2x than original segment, possibly; **(35)** female fifth swimming legs with endopod unsegmented, mostly; **(36)** female fifth legs endopod 1-segmented, mostly; **(37)** female fifth legs endopod with two setae, mostly; **(38)** female fifth swimming legs exopod 2 with lateral spine, equal size or larger than next segment usually.

Confirmatory characters to **Male.**

1. Right epimeral plate reduced, as rounded distal corner segment limit ..................................................... **2** Right epimeral plate prominent, as projections ...................................................................................... **21**
2(1). Right antennule length surpassing to genital segment .............................................................................. **3** Right antennule length not surpassing to genital segment ...................................................................... **19**
3(2). Right antennule length extending beyond caudal rami ............................................................................. **4** Right antennule length not extending beyond caudal rami ..................................................................... **11**
4(3). Fifth left swimming leg exopod 2 with spiniform seta length not surpassing the distal-point of the original segment ................................................................................................................................ **5** Fifth left swimming leg exopod 2 with spiniform seta length surpassing the distal-point of the original segment ............................................................................................................................................. **8**
5(4). Fifth right swimming leg endopod separated from the basis .................................................................... **6** Fifth right swimming leg endopod fused to basis ..................................................................................... **7**
6(5). Fifth left swimming leg length reaching first right exopod segment, medially; fifth left swimming leg basis presence of inner protuberance doubly; fifth right swimming leg basis with inner protuberance proximally; fifth right swimming leg exopod 2 with inner-posterior process, sub­triangular, and inserted distally .................................................................... ***Notodiaptomus henseni*** Fifth left swimming leg length surpassing first right exopod segment; fifth left and right swimming legs basis without inner protuberance; fifth right swimming leg exopod 2 without inner-posterior process ................................................................................................... ***Notodiaptomus nordestinus***
7(5). Limit between fourth and fifth metasome segments without ornamentation; left antennule actual segment 11 with single seta; fifth right swimming leg exopod 2 with outer spine inserted sub­distally, and rectilinear form ......................................................................... ***Notodiaptomus deitersi*** Limit between fourth and fifth metasome segments ornamented with spinules row, lateral and dorsal doubly; left antennule actual segment 11 with double seta; fifth right swimming leg exopod 2 with outer spine inserted medially, and arched inward .........................................***Notodiaptomus anisitsi***
8(4). Limit between fourth and fifth metasome segments without ornamentation; right antennule actual segment 1 without spinules; right antennule actual segment 8 with conical seta not reaching to middle-point of the sequent segment; right antennule actual segment 12 with conical seta length not smaller than to conical seta of the segment 8.............................................................................. **9** Limit between fourth and fifth metasome segments ornamented; right antennule actual segment 1 with spinules; right antennule actual segment 8 with conical seta reaching to middle-point of the sequent segment; right antennule actual segment 12 with conical seta length smaller than to conical seta of the segment 8.......................................................................................................... **10**
9(8). Fifth right swimming leg endopod separated from the basis; right antennule actual segment 14 without spinous process; left antennule actual segment 11 with single seta; fifth left swimming leg length reaching first right exopod segment proximally.............................. ***Notodiaptomus caperatus*** Fifth right swimming leg endopod fused to basis; right antennule actual segment 14 with spinous process; left antennule actual segment 11 with double seta; fifth left swimming leg length reaching first right exopod segment medially ............................................................... ***Notodiaptomus gibber***
10(8). Fifth right swimming leg endopod separated from the basis; limit between fourth and fifth metasome segments with setules; right antennule actual segment 13 with modified seta length reaching to the distal-point of the sequence segment; fifth left swimming leg length reaching first right exopod segment medially ................................................................................. ***Notodiaptomus santafesinus*** Fifth right swimming leg endopod fused to basis; limit between fourth and fifth metasome segments with spinules row, double dorsally, and single laterally; right antennule actual segment 13 with modified seta length reaching to the middle-point of the sequence segment; fifth left swimming leg length reaching first right exopod segment proximally .............................. ***Notodiaptomus iheringi***
11(3). Fifth right swimming leg endopod separated from the basis .................................................................. **12** Fifth right swimming leg endopod fused to basis ................................................................................... **16**
12(11). Fifth left swimming leg exopod 2 with spiniform seta length not surpassing the distal-point of the original segment; right antennule actual segment 13 with modified seta length reaching to the distal-point of the sequence segment; fifth right swimming leg exopod 2 with outer spine arched; right antennule actual segment 12 with conical seta length smaller than to conical seta of the segment 8........................................................................................................................................ **13** Fifth left swimming leg exopod 2 with spiniform seta length surpassing the distal-point of the segment; right antennule actual segment 13 with modified seta length reaching to the middle-point of the sequence segment; fifth right swimming leg exopod 2 with outer spine rectilinear; right antennule actual segment 12 with conical seta length not smaller than to conical seta of the segment 8 ....... **15**
13(12). Limit between fourth and fifth metasome segments without ornamentation; fifth right swimming leg coxa with distal process projecting over basis not beyond the first third posteriorly; fifth right swimming leg exopod 2 with outer spine length 2x lesser than the length of the second exopod.. **14** Limit between fourth and fifth metasome segments ornamented with spinules row, double dorsally, and single laterally; fifth right swimming leg coxa with distal process projecting over basis until the medial surface posteriorly; fifth right swimming leg exopod 2 with outer spine length 1.5x lesser than the length of the second exopod ............................................................ ***Notodiaptomus nelsoni***
14(13). Right antennule actual segment 15 without spinous process; fifth right swimming leg endopod 1- segmented; fifth left swimming leg length reaching first right exopod segment distally; fifth right swimming leg exopod 2 with outer spine directed internally ........... ***Notodiaptomus santaremensis*** Right antennule actual segment 15 with spinous process; fifth right swimming leg endopod 2- segmented; fifth left swimming leg length surpassing first right exopod segment; fifth right swimming leg exopod 2 with outer spine directed externally ........................ ***Notodiaptomus carteri***
15(12). Fifth right swimming leg exopod 1 longer than broad; fifth left swimming leg length reaching first right exopod segment medially; fifth right swimming leg coxa with distal process projecting over basis not beyond the first third posteriorly; fifth right swimming leg exopod 1 with internal prominence in acute form ............................................................................................ ***Notodiaptomus transitans*** Fifth right swimming leg exopod 1 broader than long; fifth left swimming leg length surpassing first right exopod segment; fifth right swimming leg coxa with distal process projecting over basis beyond the first third, until medial surface posteriorly; fifth right swimming leg exopod 1 without internal prominence ................................................................................... ***Notodiaptomus paraensis***
16(11). Limit between fourth and fifth metasome segments without ornamentation; fifth left swimming leg exopod 2 with spiniform seta length not surpassing the distal-point of the original segment; fifth right swimming leg exopod 1 without internal prominence; caudal rami without reticulated main axis .................................................................................................................................................. **17** Limit between fourth and fifth metasome segments ornamented with spinules row, single dorsally, and laterally; fifth left swimming leg exopod 2 with spiniform seta length surpassing the distal-point of the original segment; fifth right swimming leg exopod 1 with internal prominence in acute form; caudal rami with reticulated main axis ........................................ ***Notodiaptomus incompositus***
17(16). Left antennule actual segment 11 with single seta; fifth left swimming leg length reaching to first right exopod segment; fifth right swimming leg exopod 2 with outer spine rectilinear, and more than 2x lesser than the length of exopod 2................................................................................................... **18** Left antennule actual segment 11 with double seta; fifth left swimming leg length surpassing to first right exopod segment; fifth right swimming leg exopod 2 with outer spine arched outwards, and 2x lesser than length exopod 2 ................................................................... ***Notodiaptomus amazonicus***
18(17). Right antennule actual segment 13 with modified seta length reaching to the middle-point of the sequence segment; fifth left swimming leg length reaching to first right exopod segment distally; fifth right swimming leg coxa with distal process projecting over basis beyond the first third, until medial surface; fifth right swimming leg exopod 2 with outer spine 3x lesser than length of exopod 2........................................................................................... ***Notodiaptomus conifer*** Right antennule actual segment 13 with modified seta length reaching to the distal-point of the sequence segment; fifth left swimming leg length reaching first right exopod segment proximally; fifth right swimming leg coxa with distal process projecting over basis not beyond the first third; fifth right swimming leg exopod 2 with outer spine 6x lesser than length of exopod 2.................................................................................................... ***Notodiaptomus isabelae***
19(2). Fifth left swimming leg exopod 2 with spiniform seta length not surpassing the distal-point of the original segment; fifth left swimming leg length reaching to first right exopod segment proximally; fifth right swimming leg basis without inner protuberance ............................................................ **20** Fifth left swimming leg exopod 2 with spiniform seta length surpassing the distal-point of the original segment; fifth left swimming leg length reaching to first right exopod segment medially; fifth right swimming leg basis with inner protuberance, double and inserted medially .................................................................................... ***Notodiaptomus pseudodubius***
20(19). Fifth right swimming leg endopod separated from the basis; right antennule actual segment 13 with modified seta length reaching to the middle-point of the sequence segment; fifth right swimming leg coxa with distal process projecting over basis until the medial surface posteriorly; fifth right swimming leg exopod 2 with outer spine rectilinear form ...................... ***Notodiaptomus brandorffi*** Fifth right swimming leg endopod fused to basis; right antennule actual segment 13 with modified seta length reaching to the distal-point of the sequence segment; fifth right swimming leg coxa with distal process projecting over basis until the distal surface posteriorly; fifth right swimming leg exopod 2 with outer spine arched inward ..................................................... ***Notodiaptomus dubius***
21(1). Right antennule extending beyond caudal rami ...................................................................................... **22** Right antennule not extending beyond caudal rami ................................................................................. **27**
22(21). Limit between fourth and fifth metasome segments without ornamentation .......................................... **23** Limit between fourth and fifth metasome segments ornamented ........................................................... **25**
23(22). Fifth left swimming leg exopod 2 with spiniform seta length not surpassing the distal-point of the original segment; fifth right swimming leg exopod 1 with internal prominence; fifth right swimming leg exopod 2 with outer spine inserted sub-distally ...................................................... **24** Fifth left swimming leg exopod 2 with spiniform seta length surpassing the distal-point of the original segment; fifth right swimming leg exopod 1 with internal prominence in acute form; fifth right swimming leg exopod 2 with outer spine inserted medially .......................... ***Notodiaptomus kieferi***
24(23). Fifth right swimming leg exopod 1 longer than broad; fifth left swimming leg length surpassing first right exopod segment; right antennule actual segment 13 with modified seta reaching to the middle-point of the sequent segment; fifth right swimming leg exopod 2 with outer spine lesser 1.5x than the length of exopod 2....................................................... ***Notodiaptomus maracaibensis*** Fifth right swimming leg exopod 1 broader than long; fifth left swimming leg length reaching first right exopod segment distally; right antennule actual segment 13 with modified seta reaching to the middle-point of the sequence segment; fifth right swimming leg exopod 2 with outer spine lesser 2x than the length of exopod 2..................................................................... ***Notodiaptomus inflatus***
25(22). Limit between fourth and fifth metasome segments ornamented with spinules row, single dorsally, and laterally; fifth right swimming leg exopod 1 longer than broad; fifth right swimming leg endopod 1-segmented ............................................................................................... **26** Limit between fourth and fifth metasome segments ornamented with spinules row, double dorsally, and laterally; fifth right swimming leg exopod 1 broader than long; fifth right swimming leg endopod 2-segmented ............................................................................................... ***Notodiaptomus orellanai***
26(25). Fifth left swimming leg length reaching first right exopod segment proximally; right antennule actual segment 13 with modified seta length reaching to the middle-point of the sequence segment; fifth right swimming leg coxa with distal process projecting over basis not beyond the first third posteriorly; fifth right swimming leg exopod 2 with outer spine inserted sub-distally ....................................................................................................................................... ***Notodiaptomus dentatus*** Fifth left swimming leg length surpassing first right exopod segment; right antennule actual segment 13 with modified seta reaching to the distal-point of the sequence segment; fifth right swimming leg coxa with distal process projecting over basis beyond the first third, until medial surface posteriorly; fifth right swimming leg exopod 2 with outer spine inserted medially ..........................................................................................................................................***Notodiaptomus jatobensis***
27(21). Fifth left swimming leg exopod 2 with spiniform seta length not surpassing the distal-point of the segment; fifth right swimming leg exopod 2 with outer spine lesser than original segment up to 2x its length .......................................................................................................................................... **28** Fifth left swimming leg exopod 2 with spiniform seta length surpassing the distal-point of the segment; fifth right swimming leg exopod 2 with outer spine lesser than original segment beyond to 2x its length ............................................................................................................................................... **30**
28(27). Limit between fourth and fifth metasome segments without ornamentation; left antennule actual segment 11 with single seta; fifth right swimming leg exopod 2 with outer spine arched, and length 1.5x lesser than length original segment ......................................................................................... **29** Limit between fourth and fifth metasome segments ornamented with spinules row, double dorsally, and single laterally; left antennule actual segment 11 with double seta; fifth right swimming leg exopod 2 with outer spine in rectilinear form, and length 2x lesser than the length of exopod 2.................................................................................... ***Notodiaptomus spinuliferus***
29(28). Fifth right swimming leg exopod 1 longer than broad; fifth right swimming leg endopod 2-segmented; fifth left swimming leg length reaching first right exopod segment proximally; fourth urosome segment without sub-conical blunt dorsal-lateral process ......................... ***Diaptomus* (s.l.) *susanae*** Fifth right swimming leg exopod 1 broader than long; fifth right swimming leg endopod 1-segmented; fifth left swimming leg length surpassing first right exopod segment; fourth urosome segment with sub-conical blunt dorsal-lateral process................................................ ***Notodiaptomus cannarensis***
30(27). Fifth right swimming leg endopod separated from the basis; fifth right swimming leg coxa with distal process not projecting over basis; fifth right swimming leg exopod 2 without curved ridge on distal posterior surface ....................................................................................... ***Notodiaptomus simillimus*** Fifth right swimming leg endopod fused to basis; fifth right swimming leg coxa with distal process projecting over basis posteriorly; fifth right swimming leg exopod 2 with curved ridge on distal posterior surface .............................................................................................................................. **31**
31(30). Fifth right swimming leg exopod 1 longer than broad; left antennule actual segment 11 with double seta; right antennule actual segment 13 with modified seta length reaching to the middle-point of the sequence segment; fifth left swimming leg with length reaching to first right exopod segment distally ........................................................................................................ ***Notodiaptomus cearensis*** Fifth right swimming leg exopod 1 broader than long; left antennule actual segment 11 with single seta; right antennule actual segment 13 with modified seta length reaching to the distal-point of the sequence segment; fifth left swimming leg with length reaching to first right exopod segment medially ................................................................................................. ***Notodiaptomus coniferoides***

Confirmatory characters to **Female.**

1. Limit between fourth and fifth metasome segments without ornamentation .......................................... **2** Limit between fourth and fifth metasome segments with ornamentation ............................................. **22**
2(1). Epimeral plates asymmetrical ................................................................................................................. **3** Epimeral plates symmetrical ................................................................................................................. **19**
3(2). Fourth metasome segment absence of dorsal protuberance .................................................................... **4** Fourth metasome segment presence of dorsal protuberance .................................................................**11**
4(3). Fourth and fifth metasome segments separated; genital double-somite with disto-ventral tumescence; genital double-somite with ventral vertical folds .....................,,,,........ ***Notodiaptomus caperatus*** Fourth and fifth metasome segments fused; genital double-somite without disto-ventral tumescence; genital double-somite without ventral vertical folds ........................................................................... **5**
5(4). Fourth and fifth metasome segments fused totally .................................................................................. **6** Fourth and fifth metasome segments fused partially ................................................................................ **7**
6(5). Genital double-somite with sensilla inserted not on lobular base; right epimeral plate reaching half length of the genital segment ............................................................................ ***Notodiaptomus kieferi*** Genital double-somite with sensilla inserted on lobular base; right epimeral plate not reaching half length of the genital segment .................................................................. ***Notodiaptomus inflatus***
7(5). Right antennule extending beyond caudal rami ...................................................................................... **8** Right antennule not extending beyond caudal rami .............................................................................. **10**
8(7). Fifth swimming legs endopod 1-segmented; genital double-somite with sensilla inserted not on lobular base; prosome segments without spinules at some prosomal segment; right epimeral plate not reaching half length of the genital segment ........................................................................................ **9** Fifth swimming legs endopod 2-segmented; genital double-somite with sensilla inserted on lobular base; prosome segments with spinules at least at one prosomal segment; right epimeral plate reaching half length of the genital segment ................................................. ***Notodiaptomus simillimus***
9(8). Fifth swimming legs basis with seta no longer 2x than origin segment; genital double-somite without lateral protuberance ............................................................... ***Notodiaptomus amazonicus*** Fifth swimming legs basis with seta longer 2x than origin segment; genital double-somite with lateral protuberance, wrinkled ............................................................ ***Notodiaptomus nordestinus***
10(7). Fifth swimming legs basis with seta no longer 2x than origin segment; second urosome segment with ventral fusion to anal segment; fifth swimming legs exopod 3 with outer seta 1x lesser than the inner seta; second urosome segment with elongated spiniform process on right side distally ................................................................................. ***Notodiaptomus paraensis*** Fifth swimming legs basis with seta longer 2x than origin segment; second urosome segment without ventral fusion to anal segment; fifth swimming legs exopod 3 with outer seta 3x lesser than the inner seta; second urosome segment without elongated spiniform process ......... ***Notodiaptomus cearensis***
11(3). Fourth and fifth metasome segments fused totally ............................................................................... **12** Fourth and fifth metasome segments fused partially ............................................................................ **14**
12(11). Right antennule extending beyond caudal rami; right epimeral plate not reaching half length of the genital segment .............................................................................................................................. **13** Right antennule not extending beyond caudal rami; right epimeral plate reaching half length of the genital segment ............................................................... ***Notodiaptomus santaremensis***
13(12). Fifth swimming legs endopod 1-segmented; right epimeral plate with one projection of each side; prosome segments with spinules at least at one prosomal segment; second urosome segment with ventral fusion to anal segment ................................................ ***Notodiaptomus santafesinus*** Fifth swimming legs endopod 2-segmented; right epimeral plate with two projection of each side; prosome segments without spinules at prosomal segments; second urosome segment without ventral fusion to anal segment ....................................................................................... ***Notodiaptomus gibber***
14(11). Fourth and fifth metasome segments fused partially on lateral surface............ ***Notodiaptomus dubius*** Fourth and fifth metasome segments fused partially on dorsal surface ................................................ **15**
15(14). Fifth swimming legs endopod 1-segmented ......................................................................................... **16** Fifth swimming legs endopod 2-segmented .......................................................................................... **17**
16(15). Right antennule surpassing to genital double-segment; fifth swimming legs basis with outer seta longer 2x than origin segment; genital double-somite with ventral vertical folds ............................................................................ ***Notodiaptomus conifer*** Right antennule not surpassing to genital double-segment; fifth swimming legs basis with outer seta no longer 2x than origin segment; genital double-somite without ventral vertical folds ............................................................................ ***Notodiaptomus pseudodubius***
17(15). Right antennule extending beyond caudal rami; prosome segments without spinules at prosomal segments; right epimeral plate not reaching half length of the genital segment ............................ **18** Right antennule not extending beyond caudal rami; prosome segments with spinules at least at one prosomal segment; right epimeral plate reaching half length of the genital segment........................................................................... ***Notodiaptomus coniferoides***
18(17). Fourth metasome segment with dorsal protuberance, digitiform; right epimeral plate with two projections of each side; right antennule exceeding the caudal setae; genital double-somite with sensilla inserted not on lobular base ......................................................................... ***Notodiaptomus transitans*** Fourth metasome segment with dorsal protuberance, rounded; right epimeral plate with one projection of each side; right antennule not exceeding the caudal setae; genital double-somite with sensilla inserted on lobular base ......................................................................... ***Notodiaptomus carteri***
19(2). Fourth and fifth metasome segments separated ....................................... ***Notodiaptomus deitersi*** Fourth and fifth metasome segments fused ........................................................................................... **20**
20(19). Fourth and fifth metasome segments fused totally; right antennule not extending beyond caudal rami; fifth swimming legs endopod 2-segmented; Fifth swimming legs exopod 3 with outer seta 1x lesser than the inner seta ................................................................... ***Diaptomus*** (s.l.) ***susanae*** Fourth and fifth metasome segments fused partially; right antennule extending beyond caudal rami; fifth swimming legs endopod 1-segmented; fifth swimming legs exopod 3 with outer seta 3x lesser than the inner seta ........................................................................................................................... **21**
21(20). Right epimeral plate with one projection of each side; urosome 3-segmented; fifth swimming legs basis with outer seta longer 2x than origin segment; prosome segments with spinules at least at one prosomal segment .................................................................... ***Notodiaptomus henseni*** Right epimeral plate with two projections of each side; urosome 2-segmented; fifth swimming legs basis with outer seta no longer 2x than origin segment; prosome segments without spinules at prosomal segments ................................................................ ***Notodiaptomus cannarensis***
22(1). Fourth and fifth metasome segments separated .................................................................................... **23** Fourth and fifth metasome segments fused ........................................................................................... **25**
23(22). Limit between fourth and fifth metasome segments ornamented with single spinules row; right antennule not exceeding the caudal setae ....................................................................................... **24** Limit between fourth and fifth metasome segments ornamented with double spinules row; right antennule exceeding the caudal setae .......................................................... ***Notodiaptomus iheringi***
24(23). Right epimeral plate not reaching half length of the genital segment; fifth swimming legs basis with outer seta no longer 2x than origin segment; fifth swimming legs exopod 3 with outer seta 1x lesser than the inner seta ............................................................................ ***Notodiaptomus spinuliferus*** Right epimeral plate reaching half length of the genital segment; fifth swimming legs basis with outer seta longer 2x than origin segment; fifth swimming legs exopod 3 with outer seta 3x lesser than the inner seta ....................................................................... ***Notodiaptomus incompositus***
25(22). Fourth and fifth metasome segments fused totally .............................................................................. **26** Fourth and fifth metasome segments fused partially ............................................................................ **28**
26(25). Limit between fourth and fifth metasome segments ornamented with single spinules row; epimeral plates symmetrical; fourth metasome segment with dorsal protuberance .... ***Notodiaptomus isabelae*** Limit between fourth and fifth metasome segments ornamented with double spinules row; epimeral plates asymmetrical; fourth metasome segment without dorsal protuberance ............................... **27**
27(26). Right antennule not exceeding the caudal setae; fifth swimming legs basis with outer seta no longer 2x than origin segment; genital double-somite without ventral vertical folds ....................................................................................... ***Notodiaptomus maracaibensis*** Right antennule exceeding the caudal setae; fifth swimming legs basis with outer seta longer 2x than origin segment; genital double-somite with ventral vertical folds .......... ***Notodiaptomus jatobensis***
28(25). Fourth and fifth metasome segments fused partially on lateral surface .......... ***Notodiaptomus nelsoni*** Fourth and fifth metasome segments fused partially on dorsal surface ................................................ **29**
29(28). Limit between fourth and fifth metasome segments ornamented with spinules as a row; fourth metasome segment without dorsal protuberance ........................................................................... **30** Limit between fourth and fifth metasome segments ornamented with spinules as a patch; fourth metasome segment with dorsal protuberance ................................... ***Notodiaptomus orellanai***
30(29). Limit between fourth and fifth metasome segments ornamented with single spinules row; second urosome segment without ventral fusion to anal segment; genital double-somite without posterior-dorsal process; genital double-somite without ventral vertical folds ............................................. **31** Limit between fourth and fifth metasome segments ornamented with double spinules row; second urosome segment with ventral fusion to anal segment; genital double-somite with posterior-dorsal process; genital double-somite with ventral vertical folds ........................... ***Notodiaptomus anisitsi***
31(30). Right antennule extending beyond caudal rami; fifth swimming legs endopod 1-segmented; fifth swimming legs basis with outer seta longer 2x than origin segment; prosomal segments with spinules at least at one prosomal segment posteriorly .......................... ***Notodiaptomus dentatus*** Right antennule not extending beyond caudal rami; fifth swimming legs endopod 2-segmented; fifth swimming legs basis with outer seta no longer 2x than origin segment; prosomal segments without spinules at prosomal segment posteriorly ................................... ***Notodiaptomus brandorffi***

### Diagnosis and species redescription

#### Notodiaptomus deitersi (Poppe, 1891)

##### Synonymy

*Diaptomus deitersi* Poppe, 1891: 248, figs. 1–3; De Guerne & Richard, 1892: 2, pl. 10–12; Richard, 1897b: 298; Giesbrecht & Schmeil, 1898: 81; Sars 1901: 10, 12; Daday, 1905: 151, 152; Tollinger, 1911: 69, 270, 271, fig. E; Brian, 1926: 182, 183; Spandl, 1926: 104, figs. 7a-d; Pesta, 1927: 80; Wright, 1927: 73, 75, 95, 100, 102, pl. 8, figs. 5–6; 1935: 213, 219, 220; 1937: 76; 1938b: 562; Lowndes, 1934: 89, 90, 91, 96–98, pl. 2; figs. 2a-b; Brehm, 1959: 511, 514, 515, 516, figs. 15–22; Dussart & Matsumura-Tundisi, 1986: 250. *Notodiaptomus deitersi*; Kiefer, 1936a: 197; 1954: 173; 1956: 242; Brehm, 1956: 413, 414; 1958a: 168; 1959: 514, 515, 516, figs. 15–22; Ringuelet, 1958a: 50; Ringuelet & Martínez de Ferrato, 1967: 411, 417, pl. 2, figs. 11–14; Brandorff, 1972: 44; 1976: 616, fig. 2; Löffler, 1981: 15; Dussart & Defaye, 1983: 134; Dussart & Frutos, 1985: 306; Matsumura-Tundisi, 1986: 537, 100, figs. 26–33; Reid, 1987:377; Sendacz, 1993: 35; Battistoni, 1995: 959; Rocha *et al*., 1995: 155, 156; Lansac-Tôha *et al*., 1997: 140, tab. 3; Santos-Silva, 1998: 208; Santos-Silva *et al*., 1999: 114–128, figs. 1–8, tabs. 1–2; Santos-Silva *et al*., 2015: 16–21, figs. 1–8, tabs. 1–2, identification key to male, and female; Perbiche-Neves *et al*., 2020: 696-697, key to the Neotropical diaptomid, fig. 21.15 E; Geraldes-Primeiro *et al*., 2021: 2. *Neodiaptomus deitersi;* Brehm, 1959: 510, 511, 514, 515, 517, figs. 15–22. *Notodiaptomus (Notodiaptomus) deitersi*; Dussart, 1985a: 208.

**FIGURE 2.**
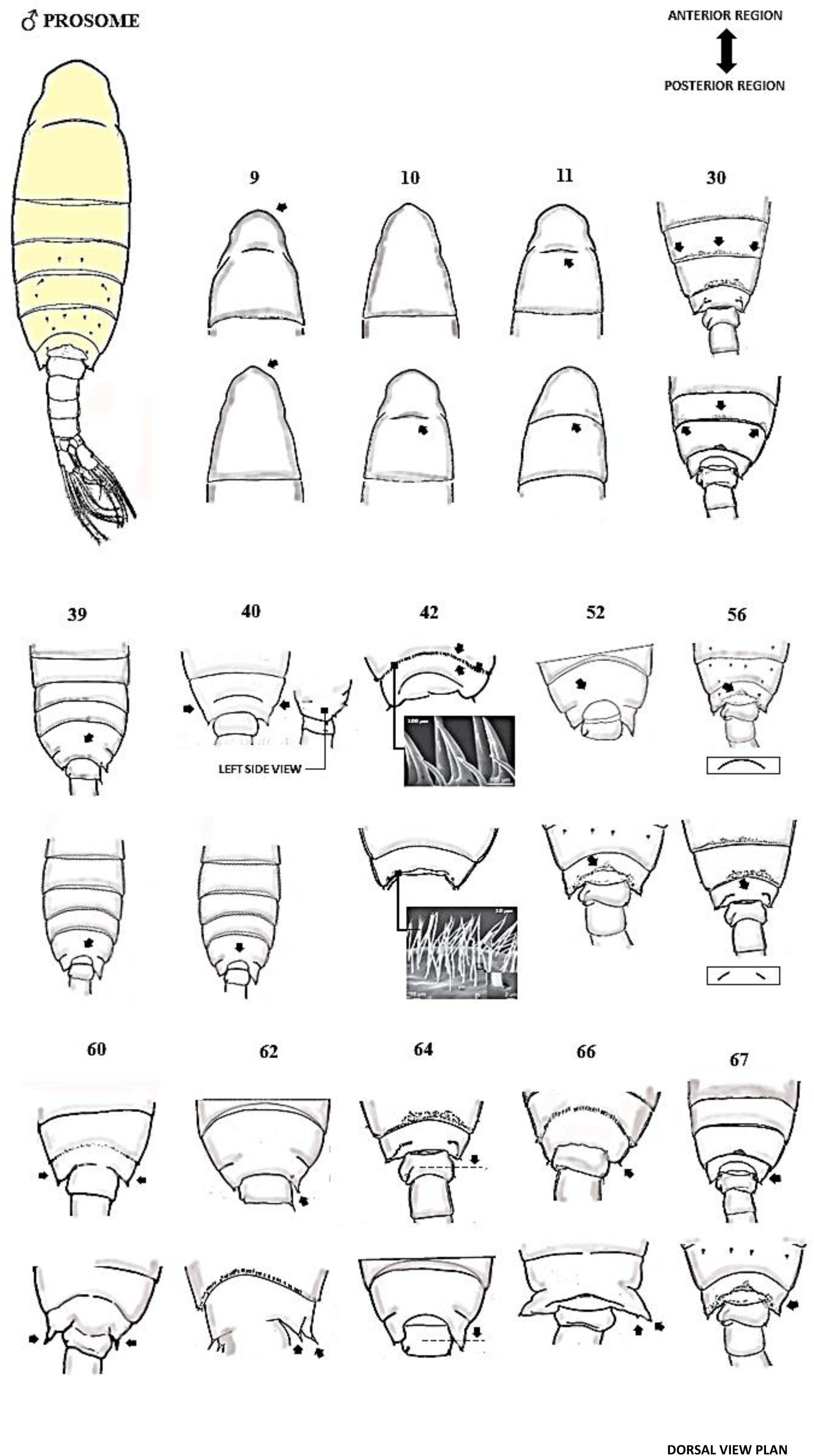
Attributes of the male prosome respectively: 9-67 chars.

**FIGURE 3.**
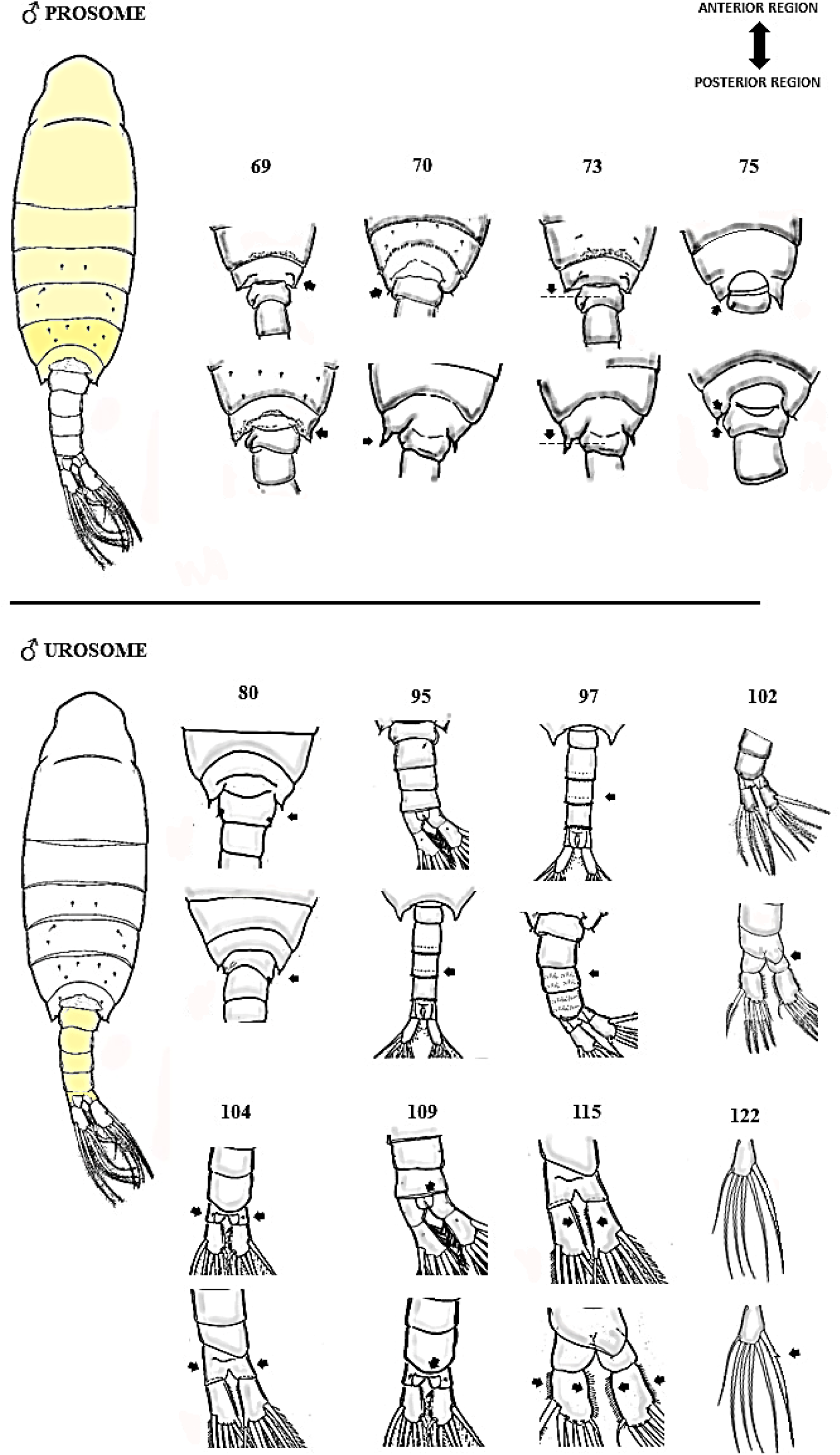
Attributes of the male prosome, and urosome, respectively: 69-75, and 80-122 chars. Dorsal view plan.

**FIGURE 4.**
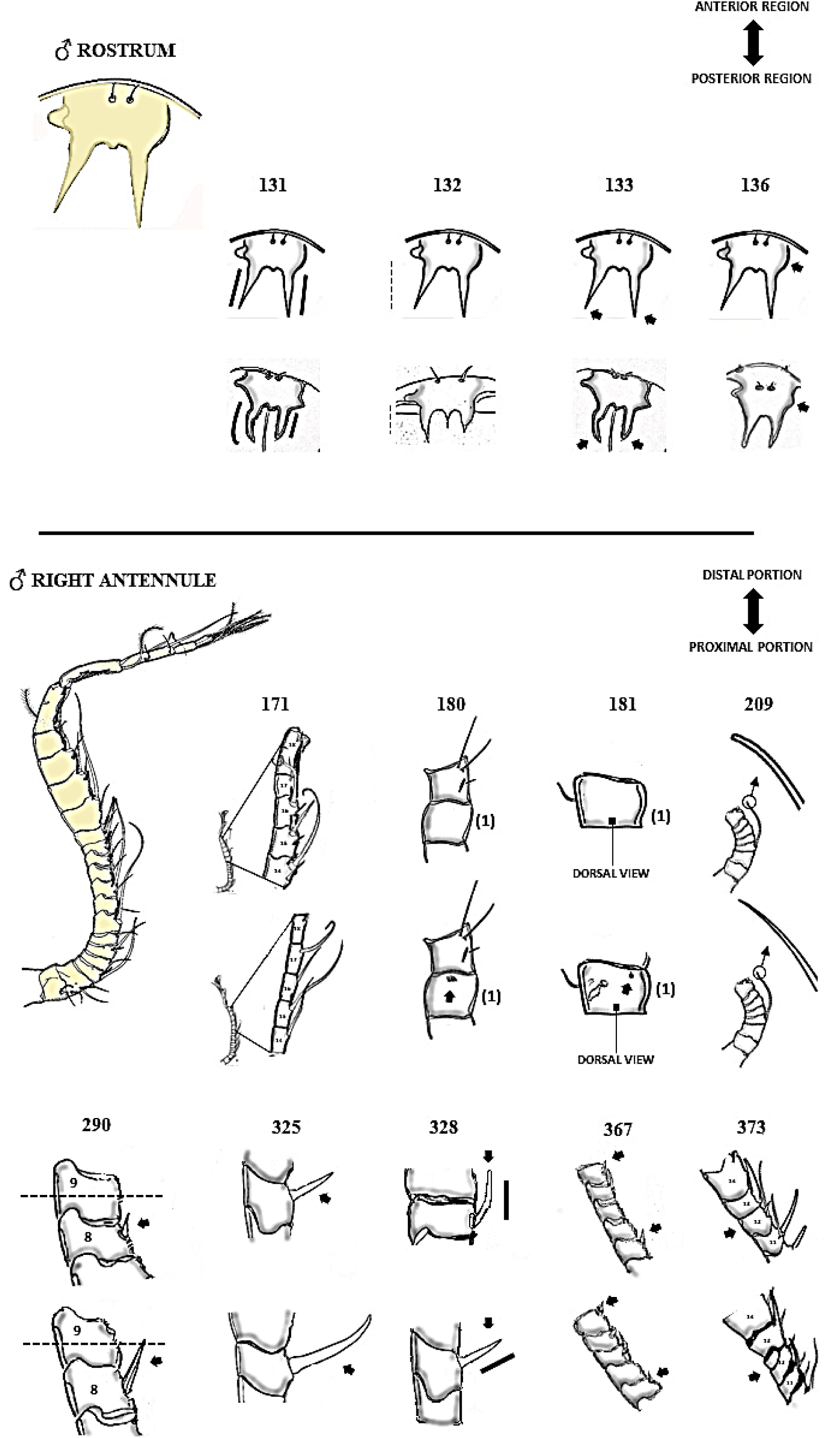
Attributes of the male rostrum, and right antennule, respectively: 131-136, and 171-373 chars. Ventral view plan.

**FIGURE 5.**
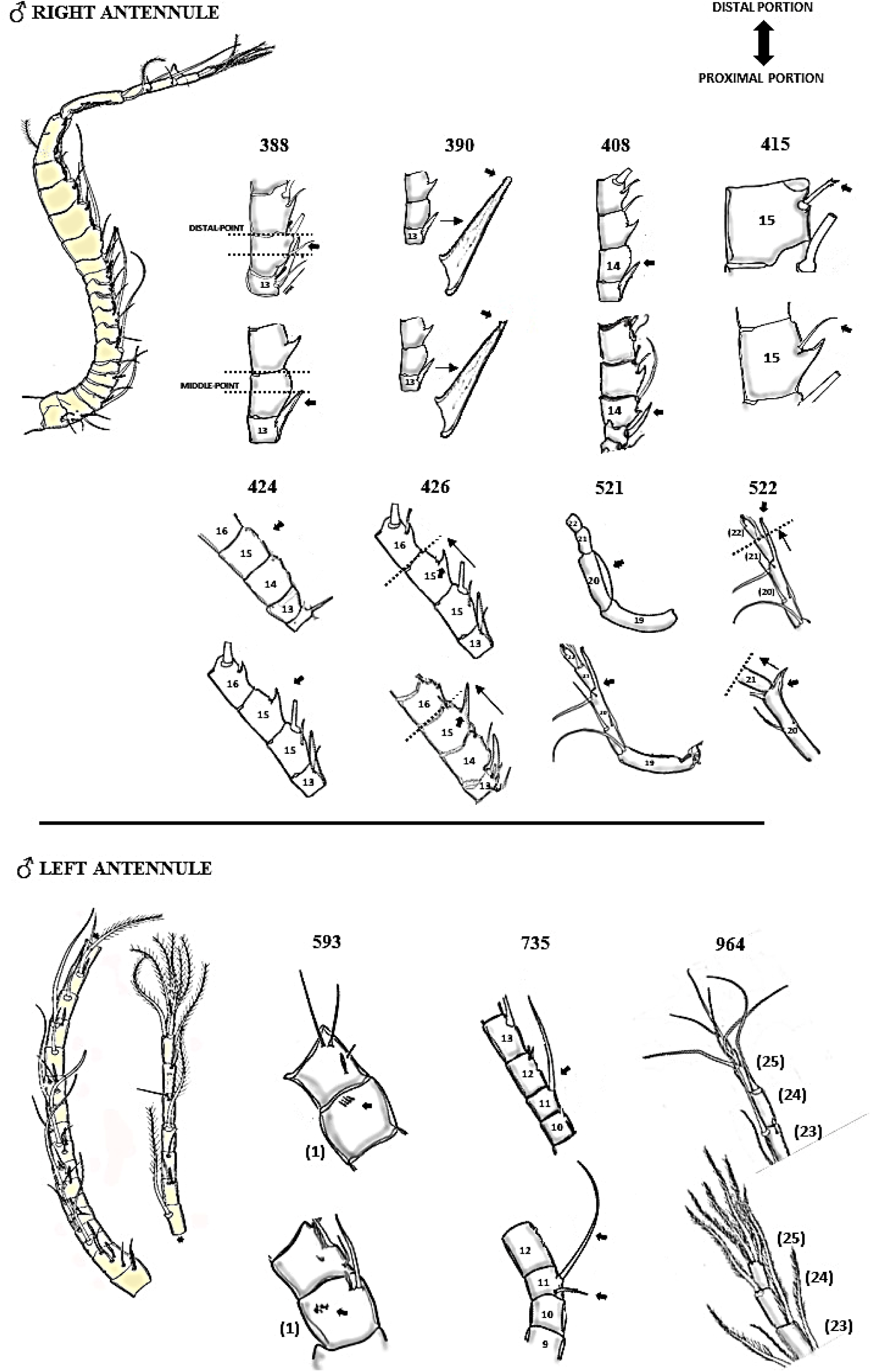
Attributes of the male right antennule, and left antennulae, respectively: 388-522, and 593-964 chars. Ventral view plan.

**FIGURE 6.**
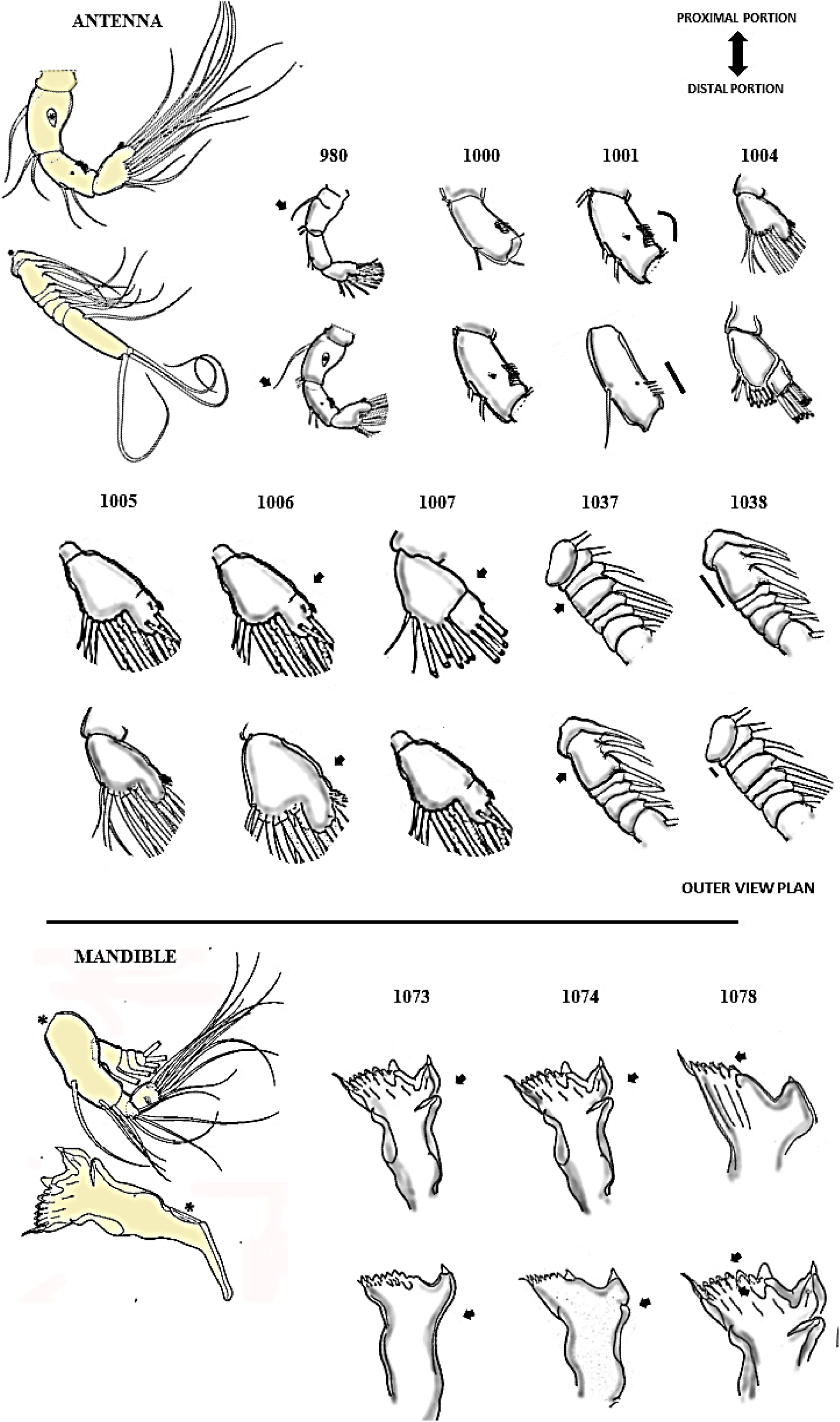
Attributes of the antenna, and mandible, respectively: 980-1038, and 1073-1078 (inner view) chars.

**FIGURE 7.**
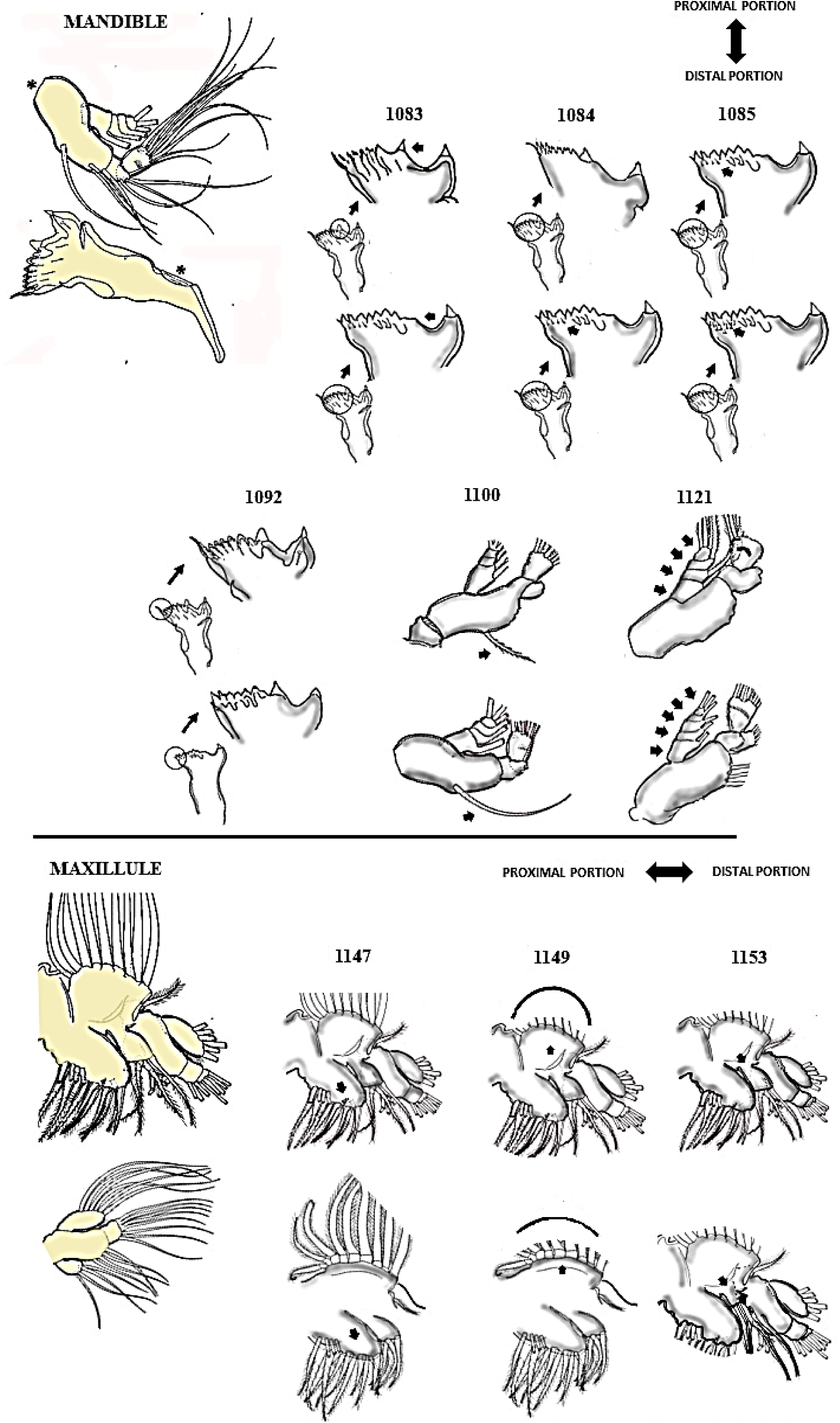
Attributes of the mandible, and maxillule respectively: 1083-1121, and 1147-1153 chars. Inner view

**FIGURE 8.**
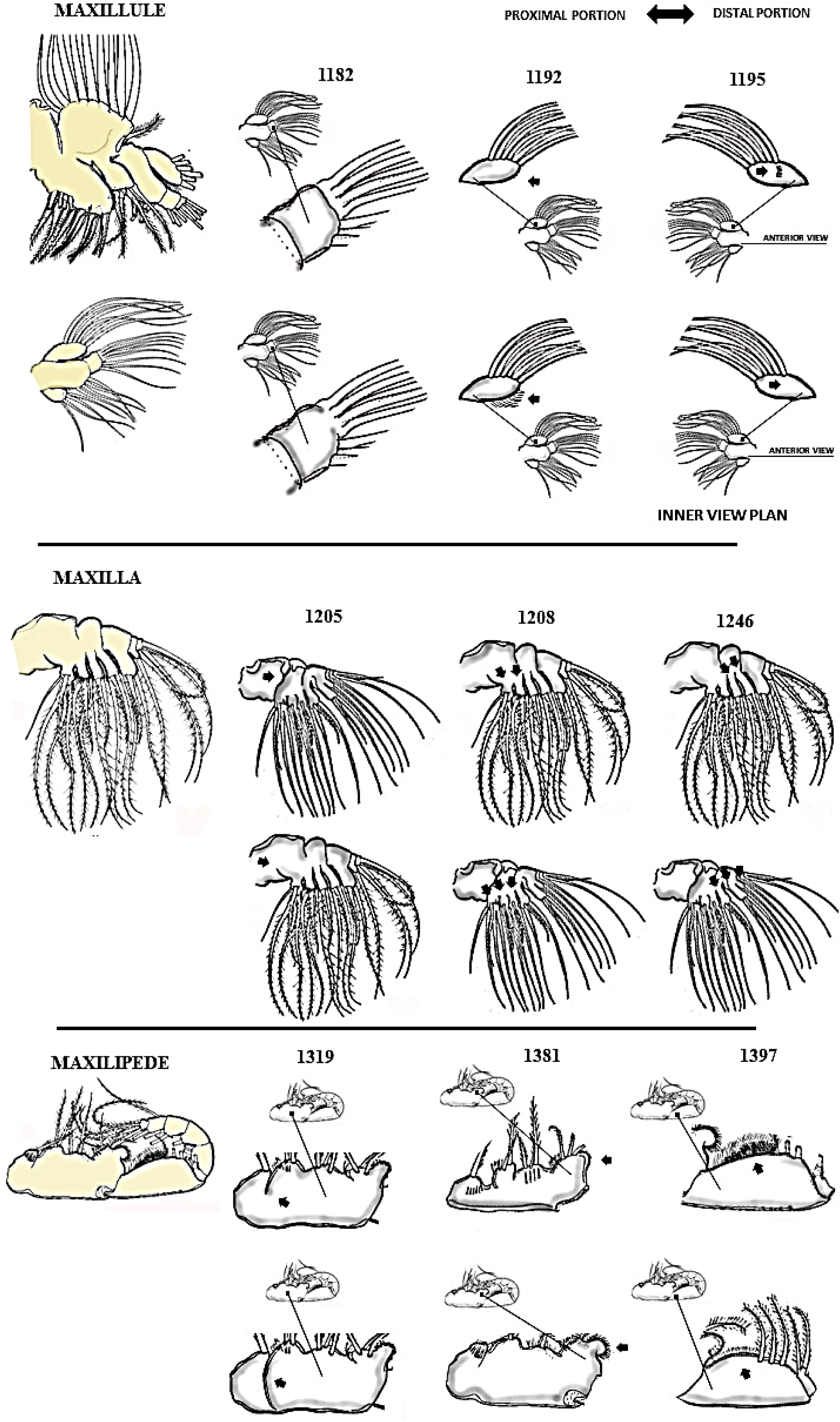
Attributes of the maxillule, maxila, and maxilipede, respectively: 1182-1195, 1205-1246, and 1319-1397

**FIGURE 9.**
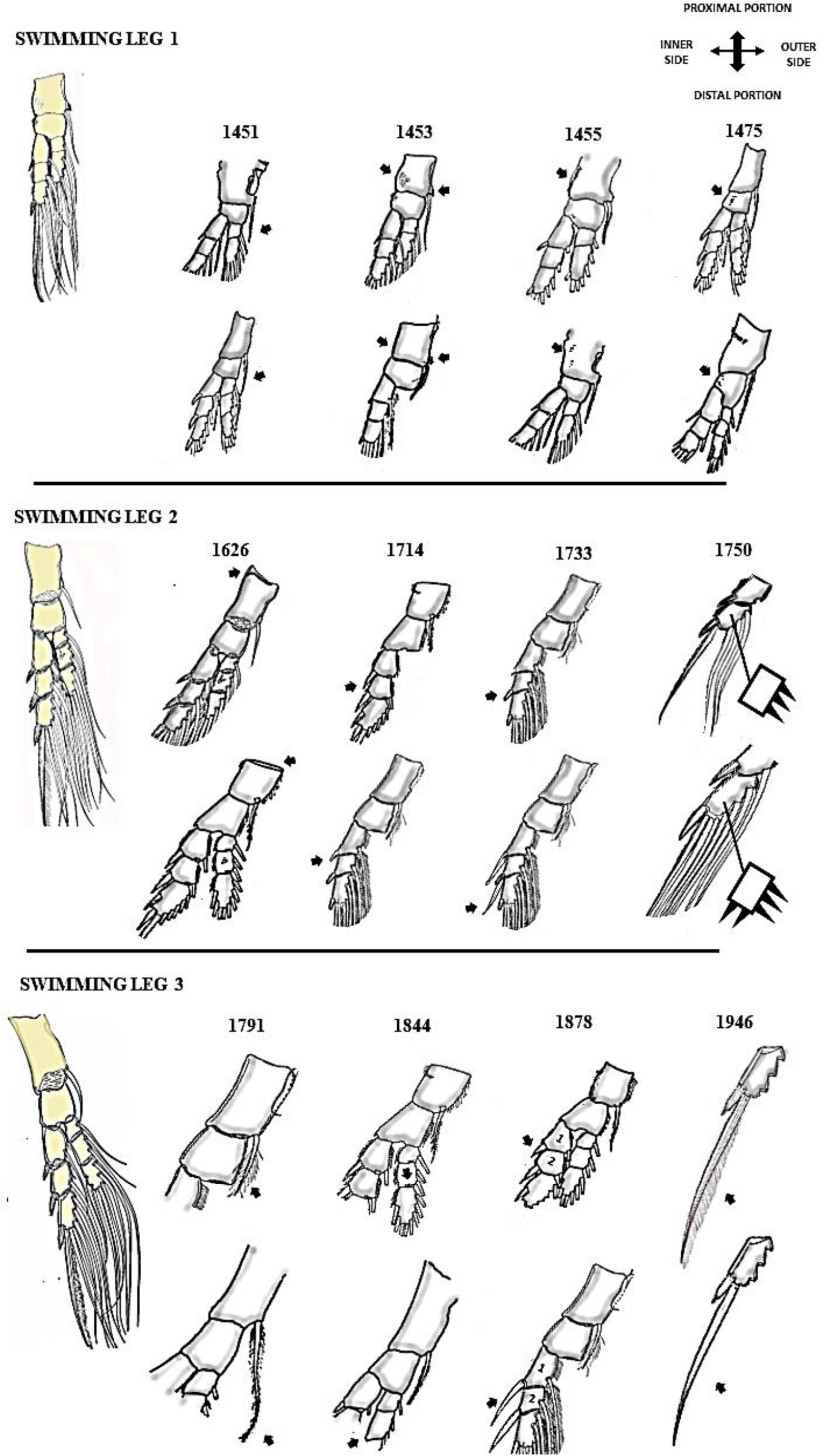
Attributes of the swimming legs 1, 2 and 3 respectively: 1451-1475, 1626-1750, and 1791-1946

**FIGURE 10.**
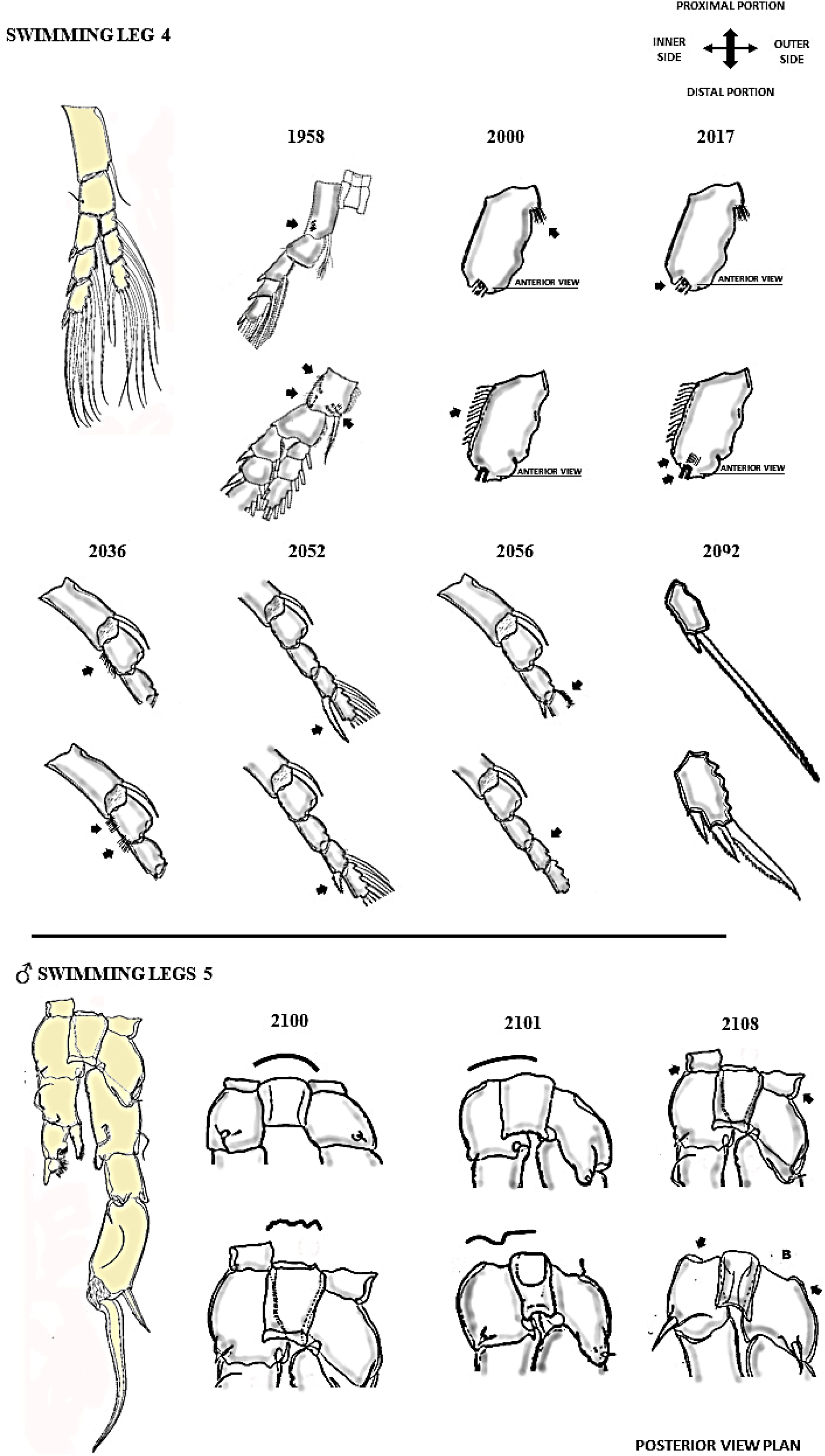
Attributes of the swimming legs 4 and 5 (male) respectively: 1958-2092 and 2100-2108 chars.

**FIGURE 11.**
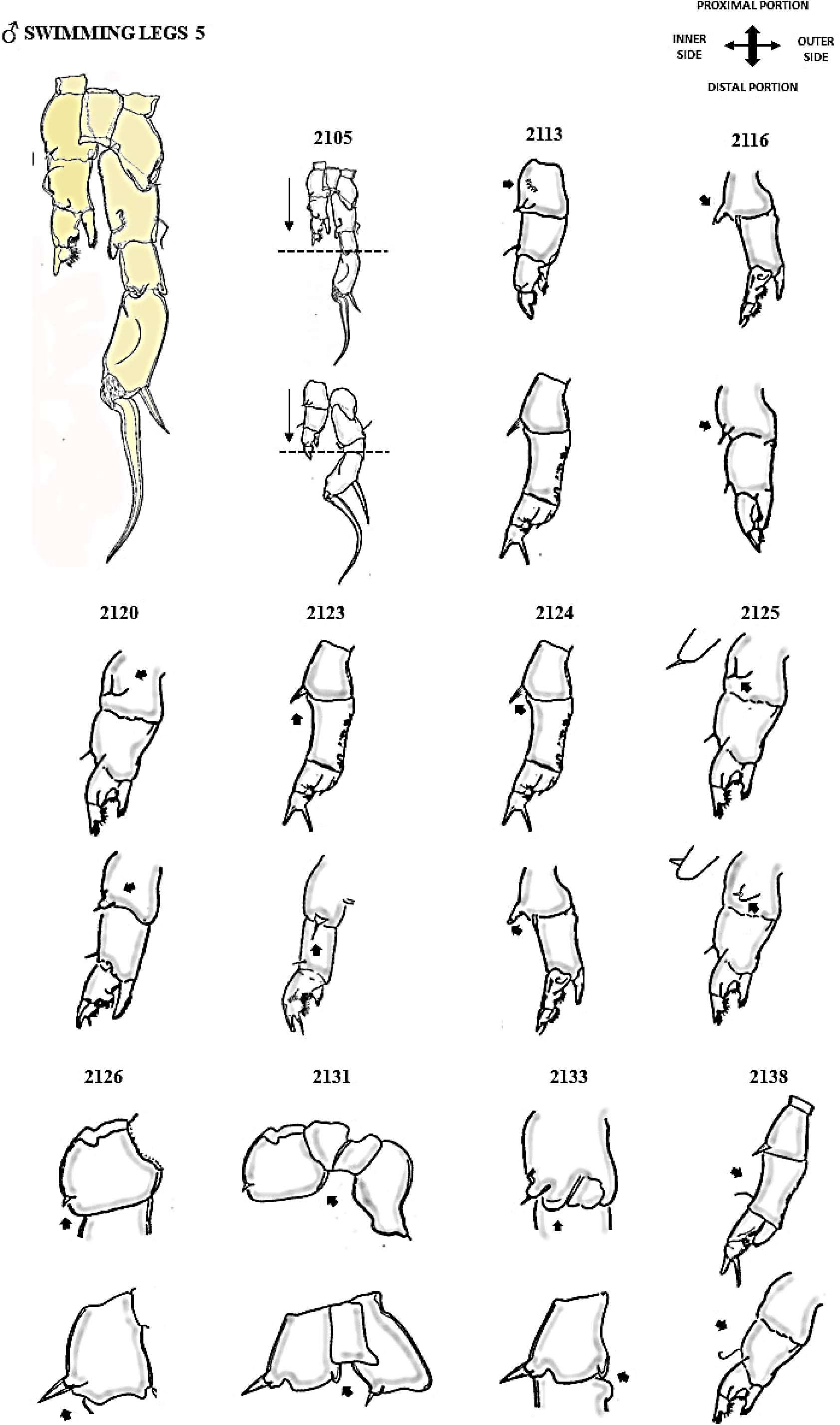
Attributes of the male swimming legs 5 respectively: 2105-2138 chars. Posterior view plan.

**FIGURE 12.**
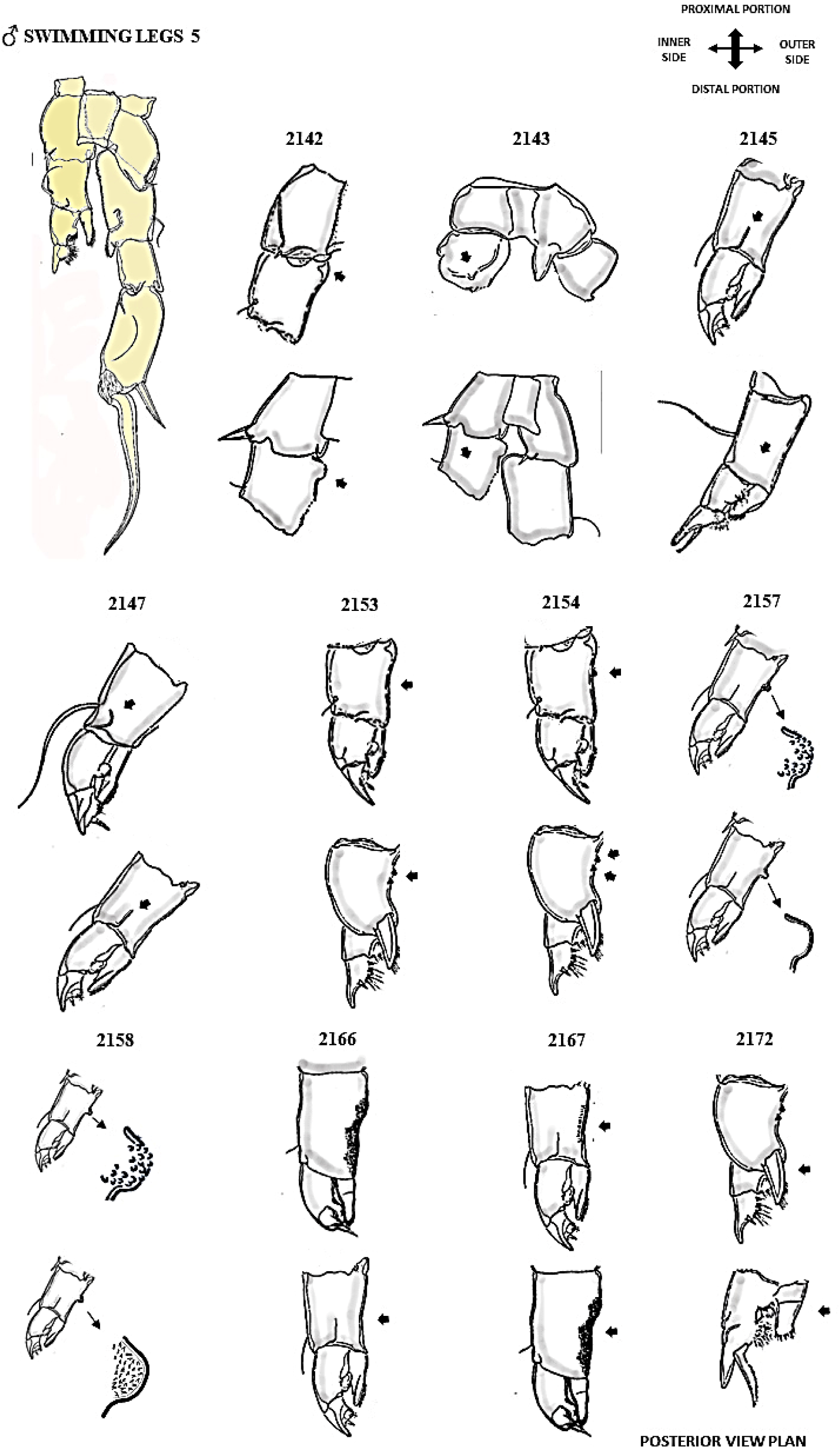
Attributes of the male swimming legs 5 respectively: 2142-2172 chars.

**FIGURE 13.**
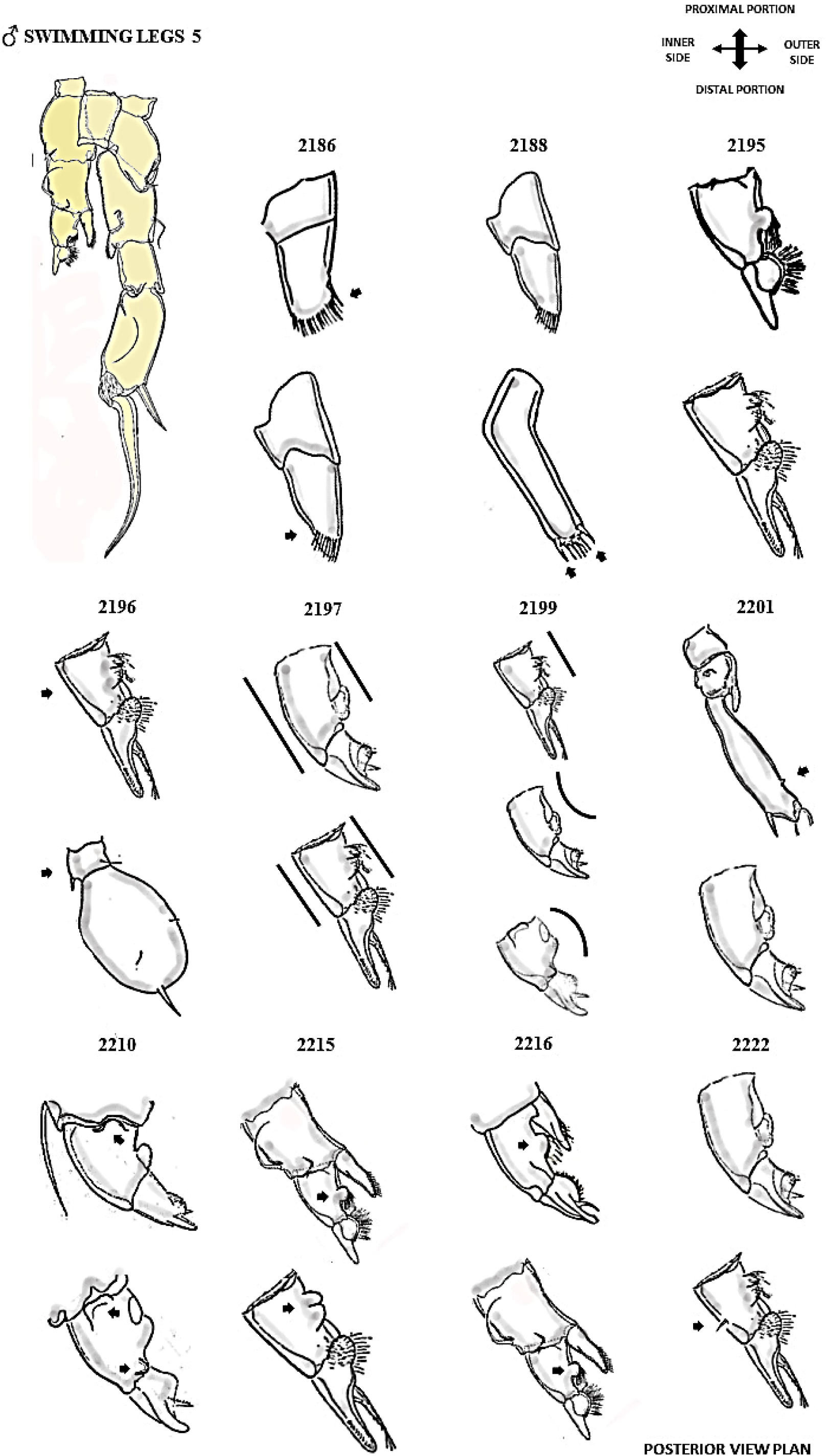
Attributes of the male swimming legs 5 respectively: 2186-2222 chars.

**FIGURE 14.**
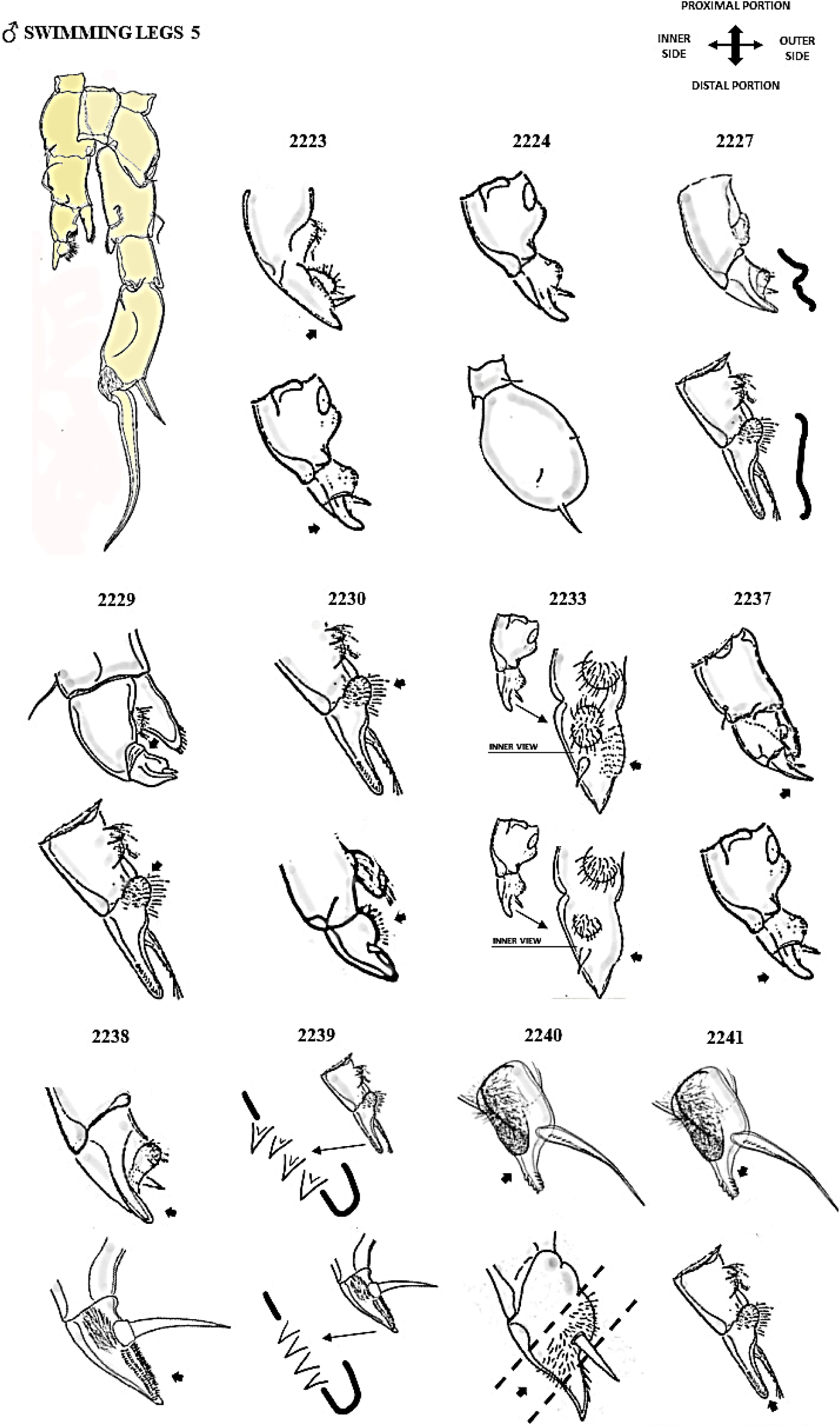
Attributes of the male swimming legs 5 respectively: 2223-2241 chars. Posterior view plan.

**FIGURE 15.**
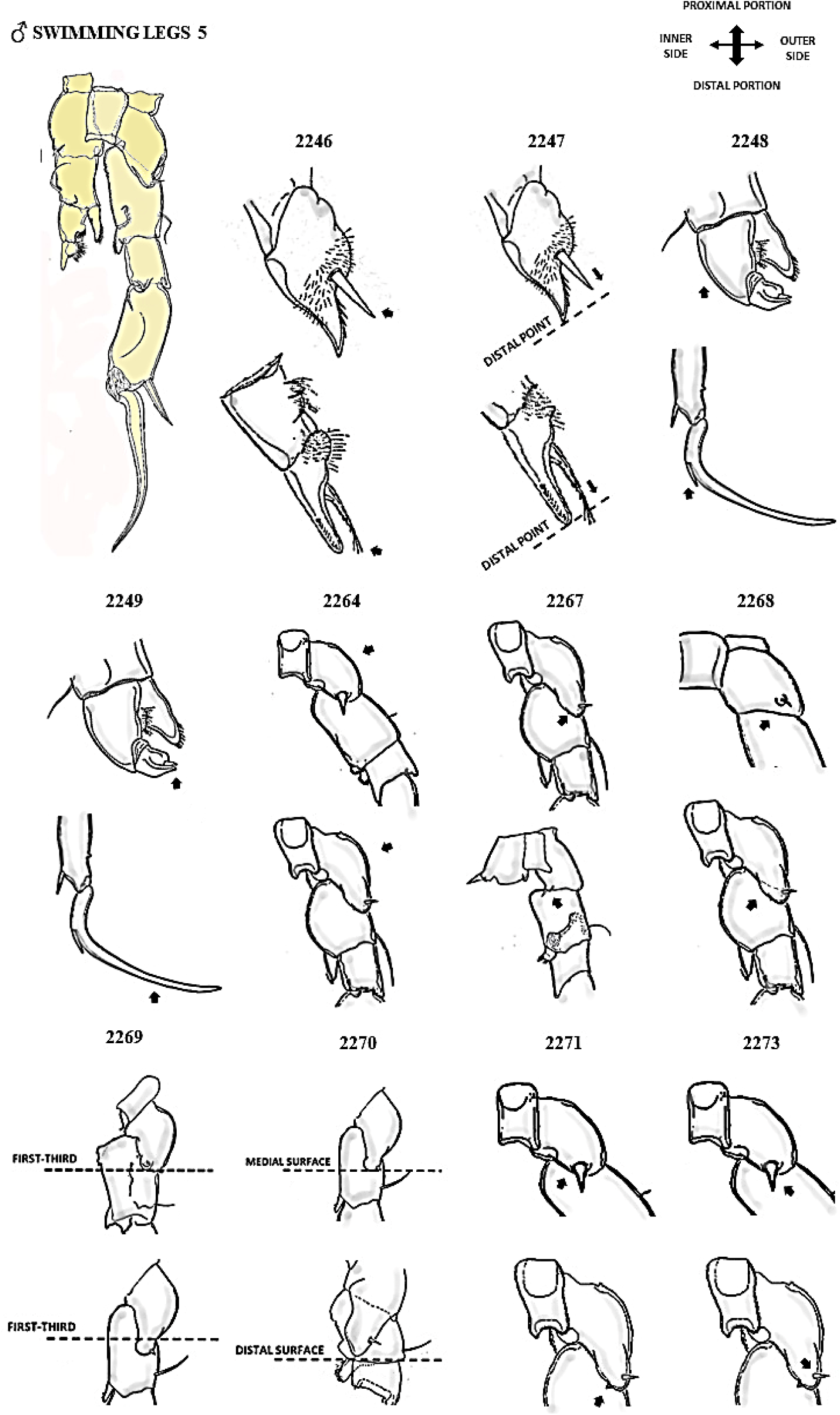
Attributes of the male swimming legs 5 respectively: 2246-2273 chars. Posterior view plan.

**FIGURE 16.**
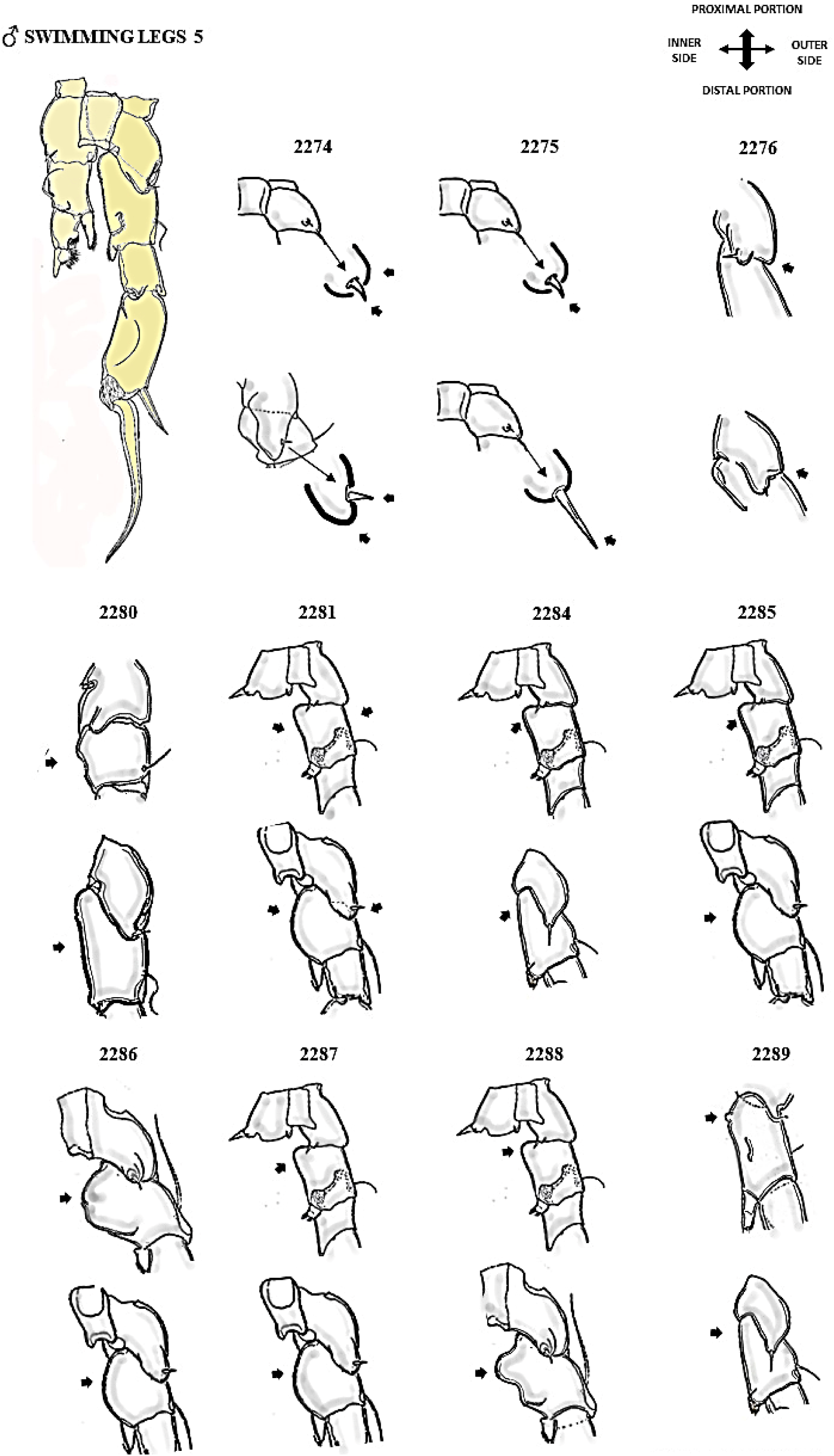
Attributes of the male swimming legs 5 respectively: 2274-2289 chars. Posterior view plan.

**FIGURE 17.**
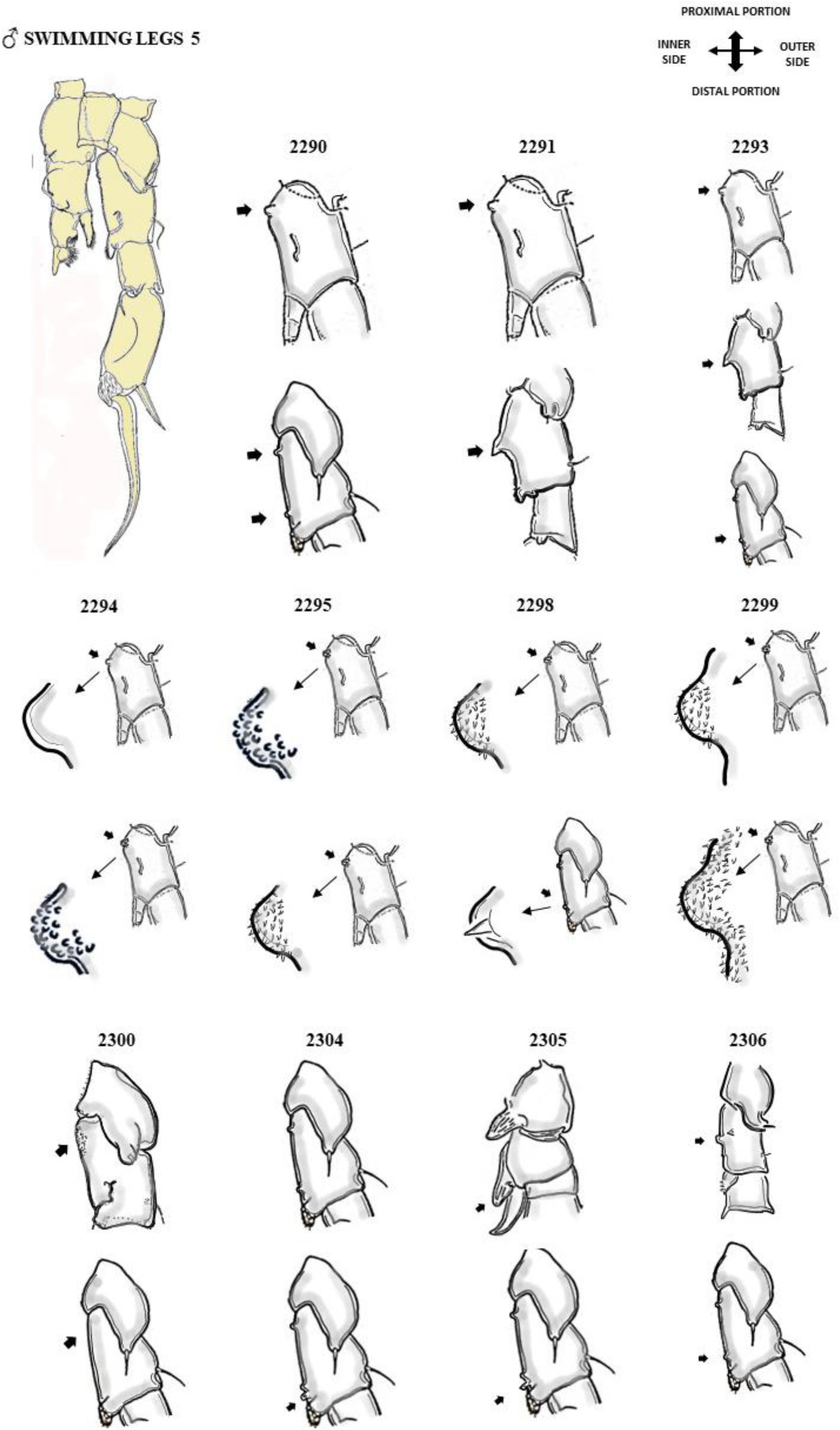
Attributes of the male swimming legs 5 respectively: 2290-2306 chars. Posterior view plan.

**FIGURE 18.**
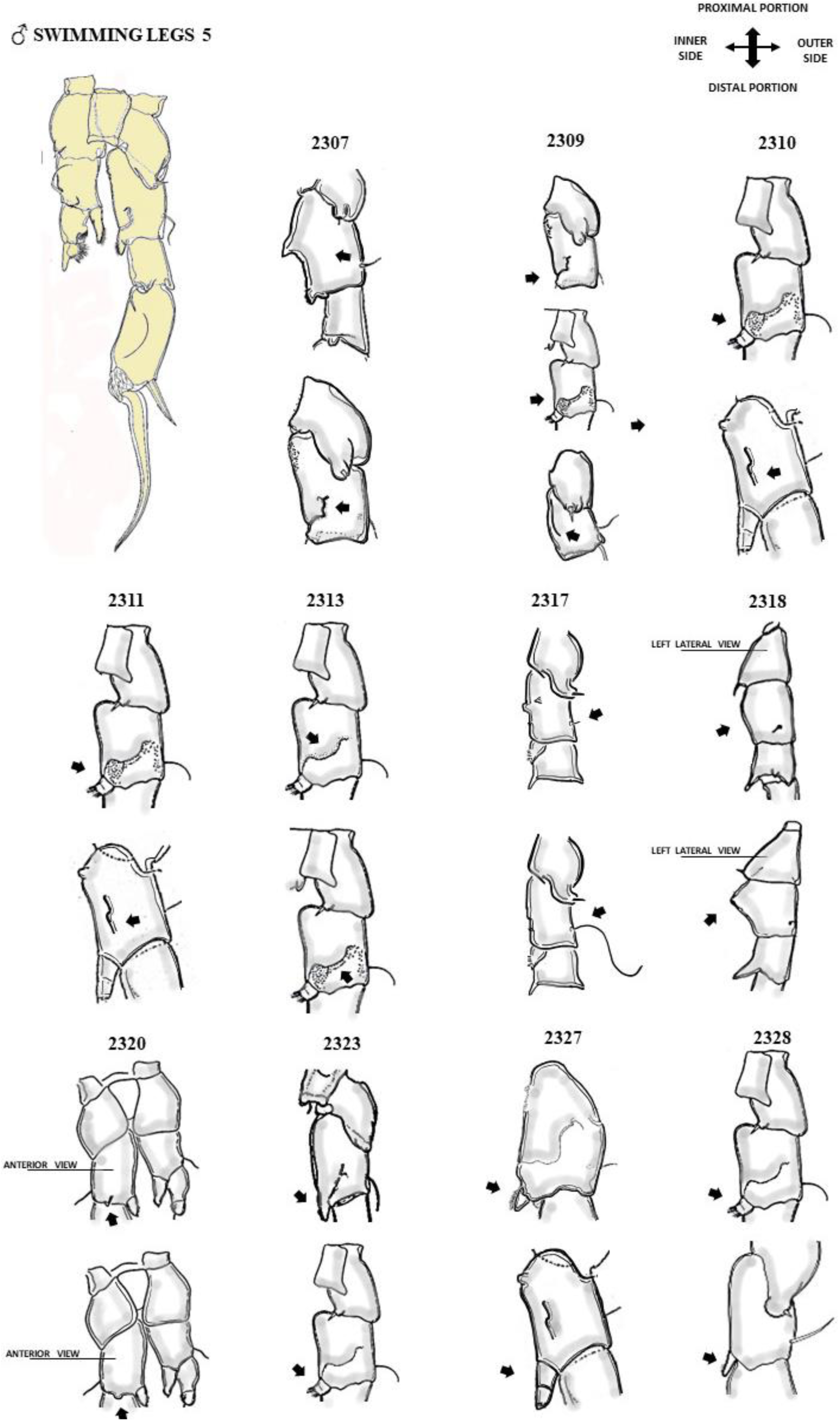
Attributes of the male swimming legs 5 respectively: 2307-2328 chars. Posterior view plan.

**FIGURE 19.**
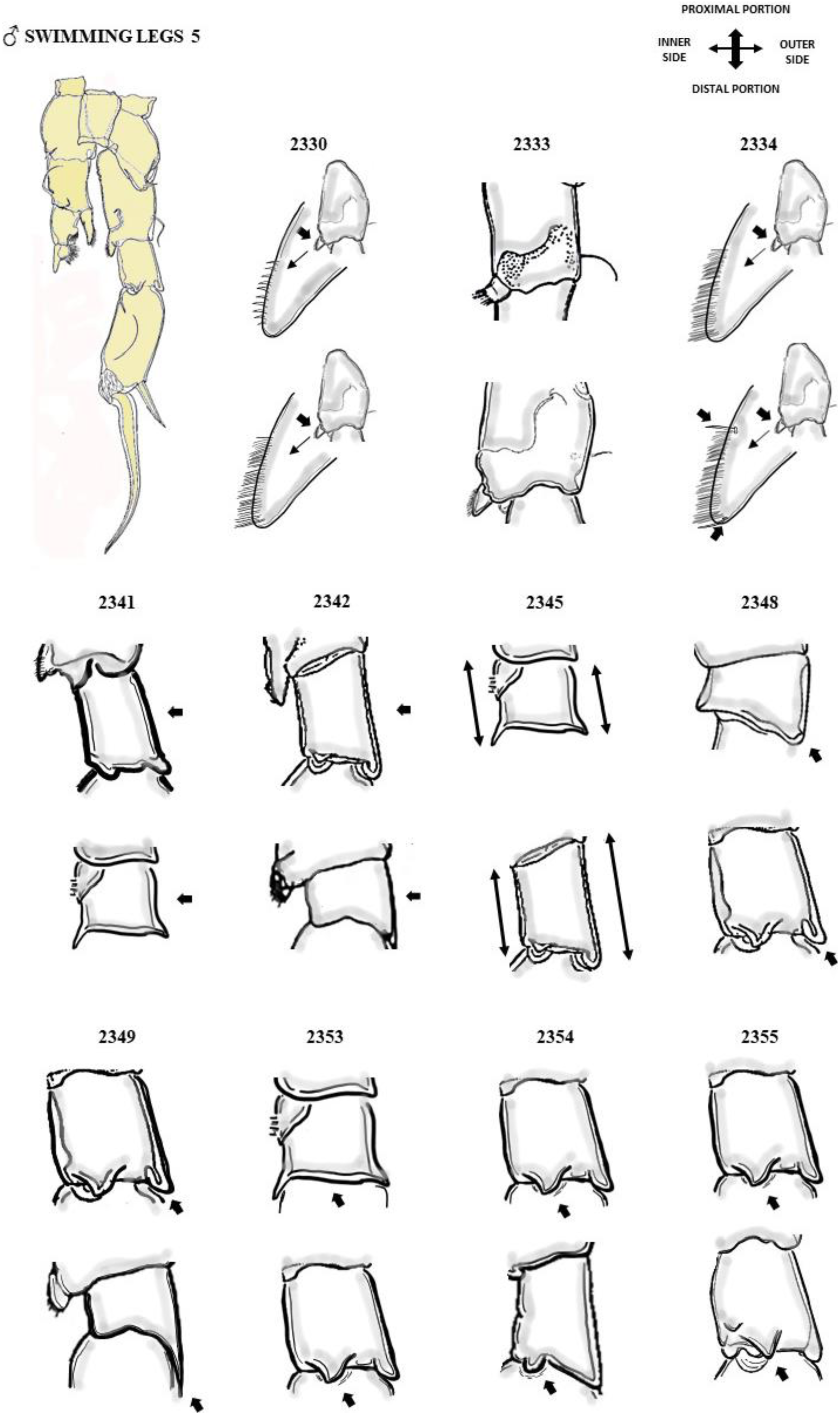
Attributes of the male swimming legs 5 respectively: 2330-2355 chars. Posterior view plan.

**FIGURE 20.**
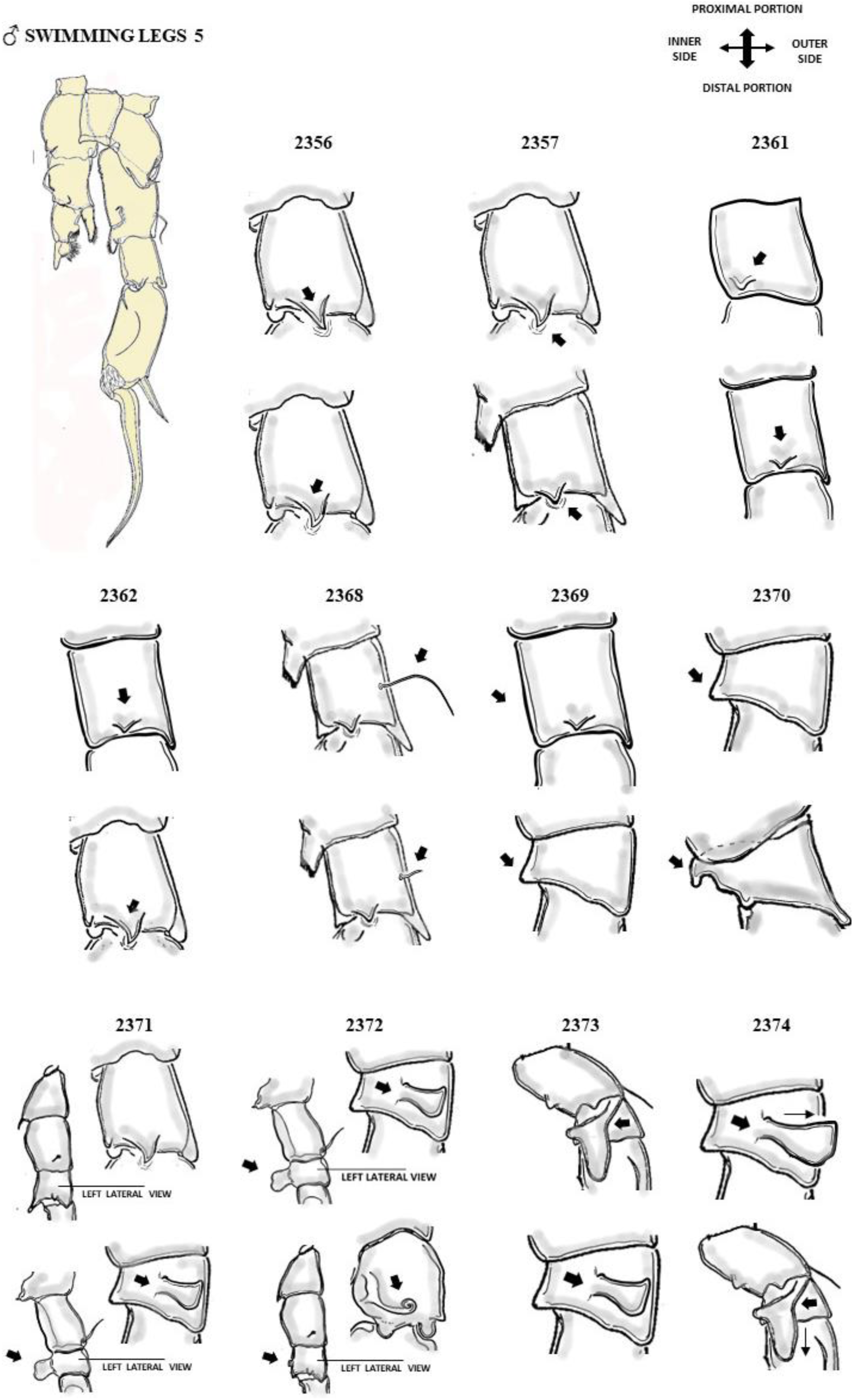
Attributes of the male swimming legs 5 respectively: 2356-2374 chars. Posterior view plan.

**FIGURE 21.**
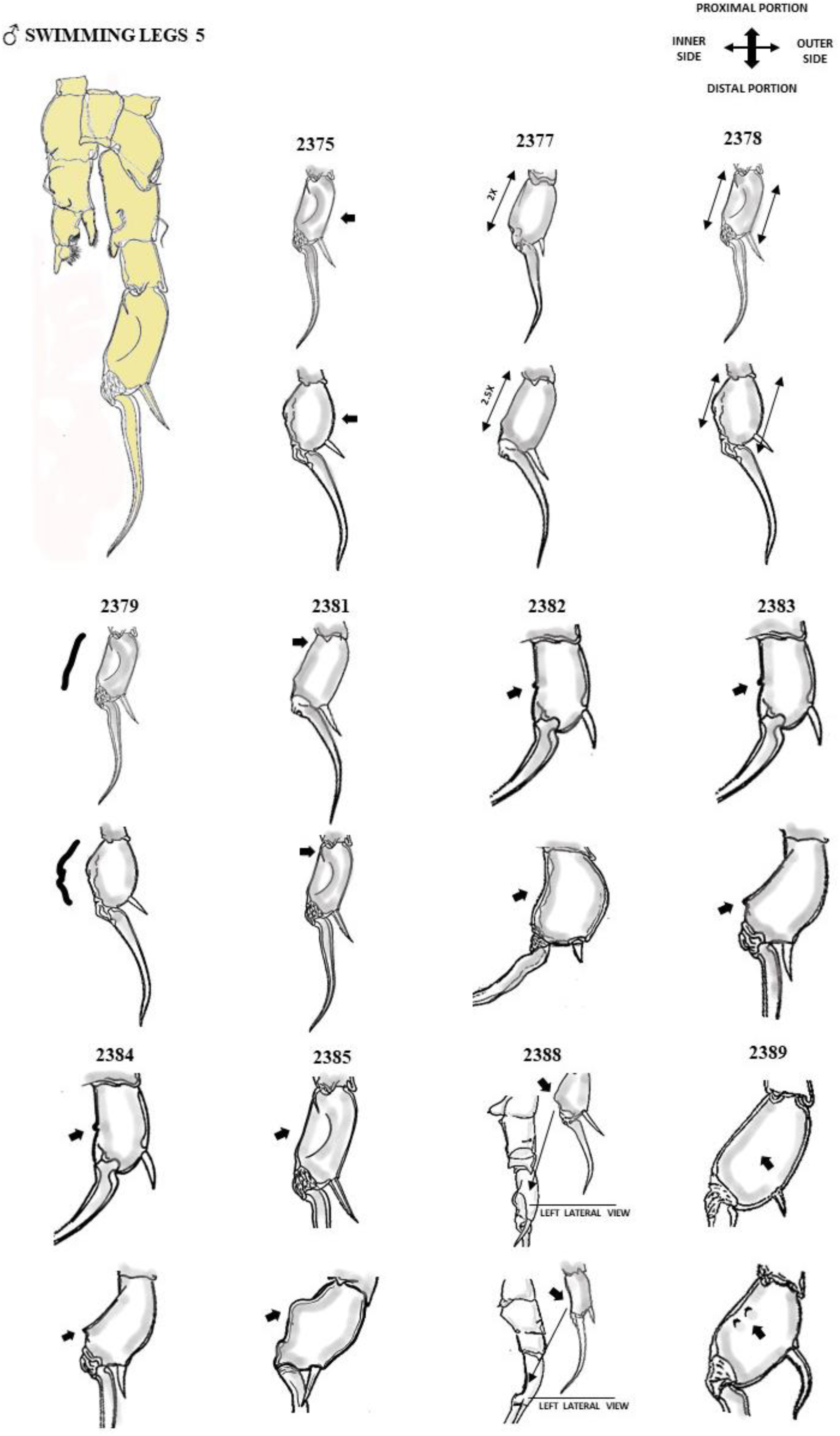
Attributes of the male swimming legs 5 respectively: 2375-2389 chars. Posterior view plan.

**FIGURE 22.**
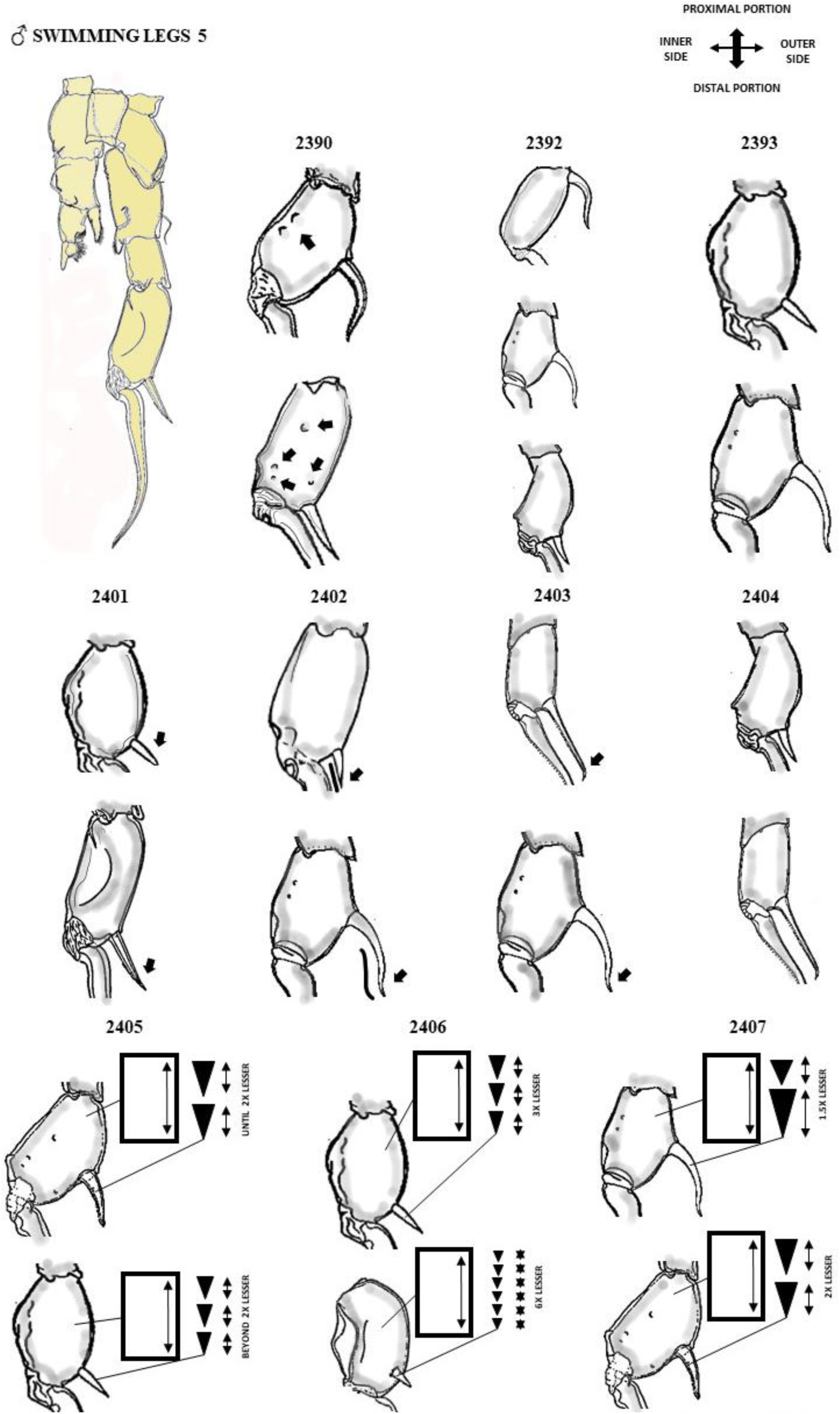
Attributes of the male swimming legs 5 respectively: 2390-2407 chars. Posterior view plan.

**FIGURE 23.**
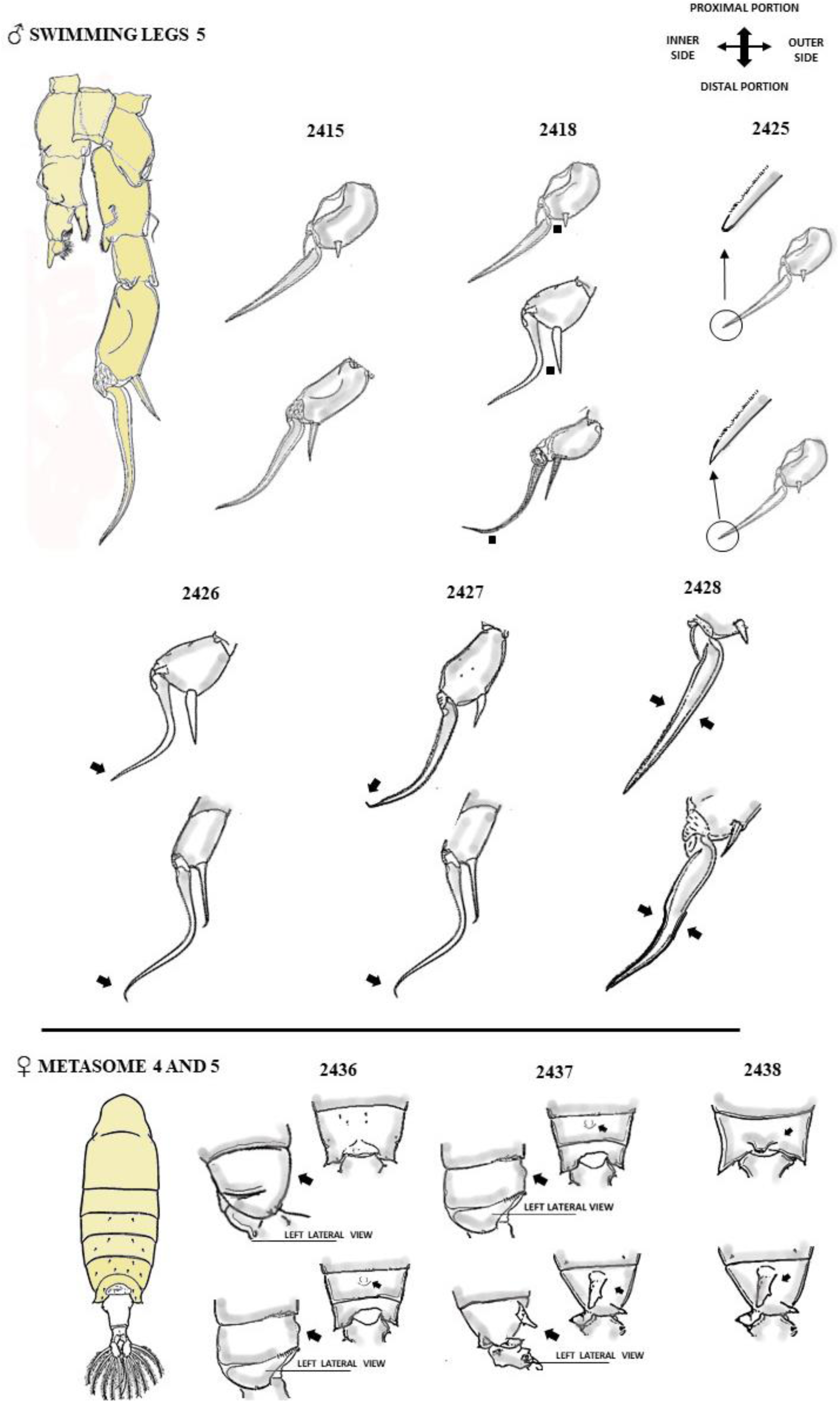
Attributes of the male swimming legs 5, and female metasome 5, respectively: 2415-2428 chars, and 2436-2438 chars (anterior-posterior in dorsal view plan). Posterior view plan.

**FIGURE 24.**
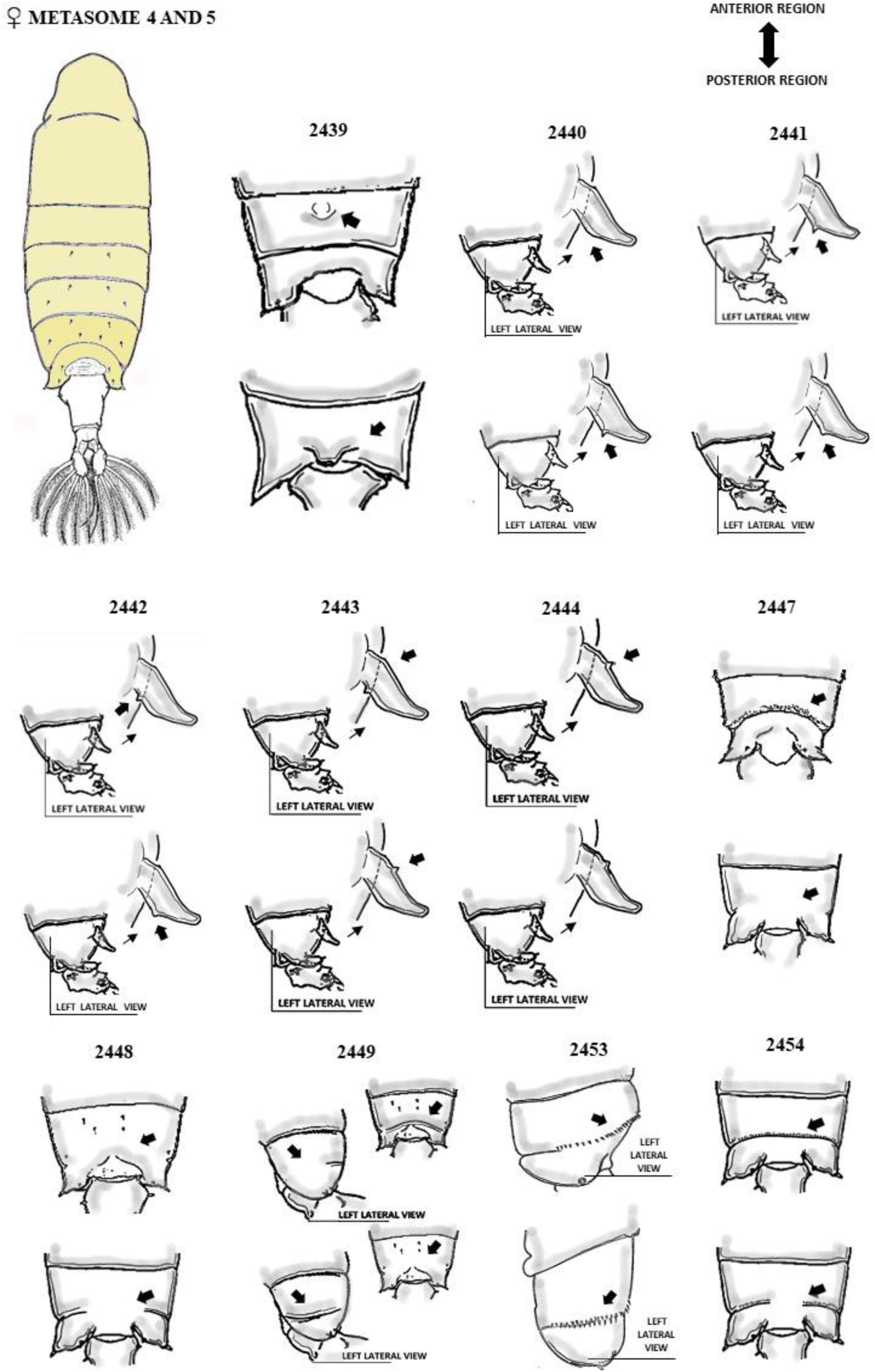
Attributes of the female metasome 4 and 5, respectively: 2439-2454 chars. Dorsal view plan.

**FIGURE 25.**
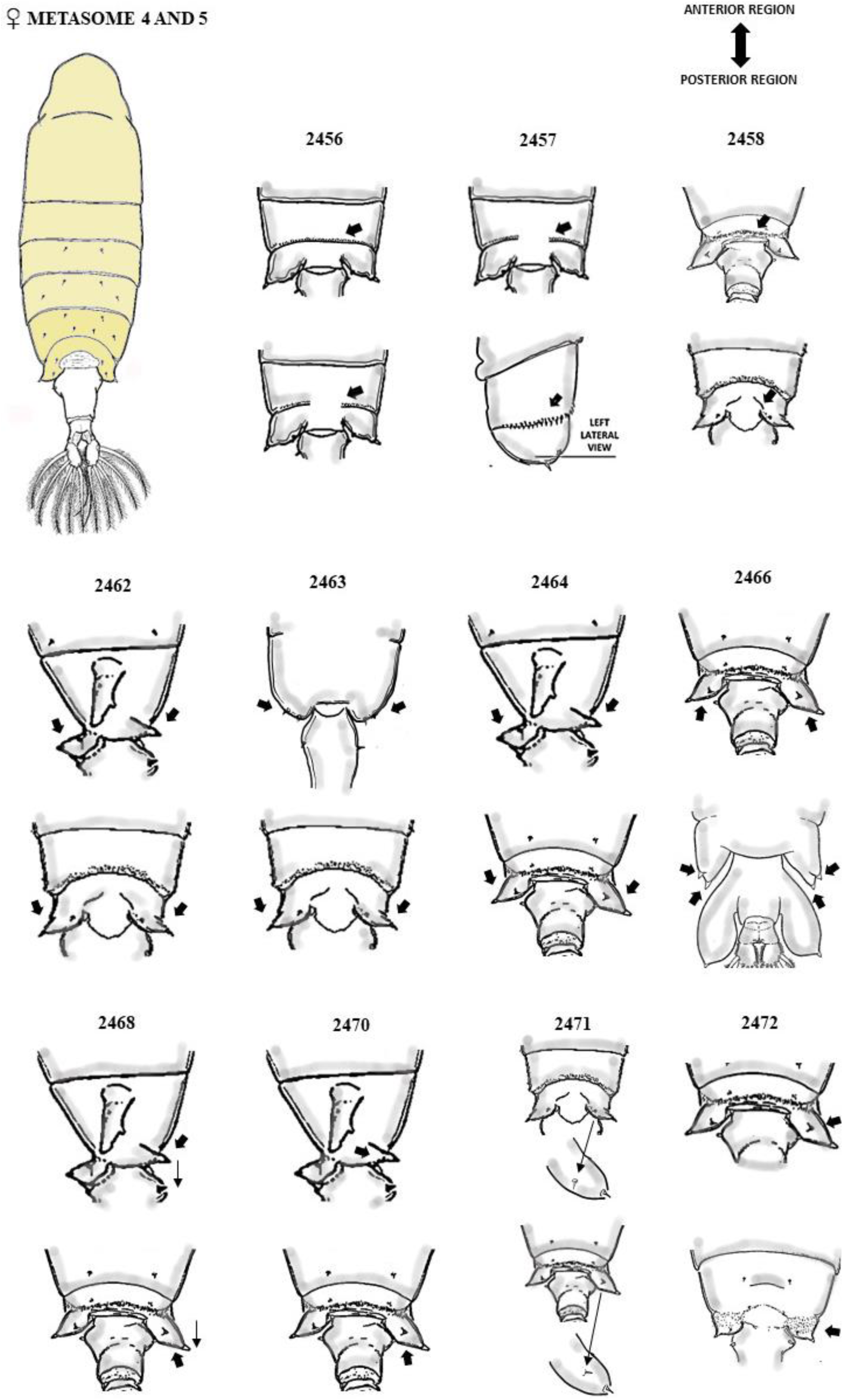
Attributes of the female metasome 4 and 5, respectively: 2456-2472 chars. Dorsal view plan.

**FIGURE 26.**
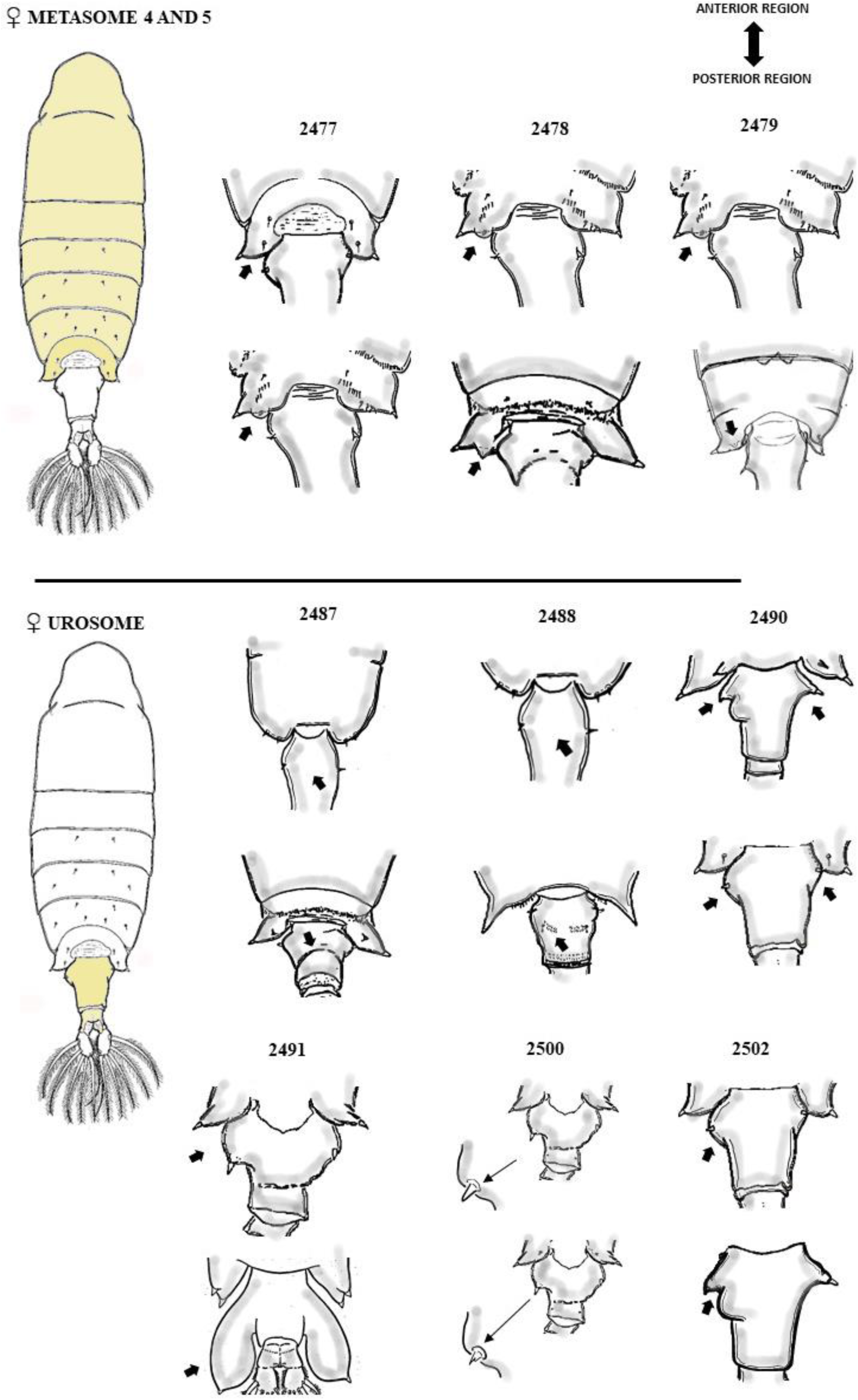
Attributes of the female metasome 5, and urosome, respectively: 2477-2502 chars. Dorsal view plan.

**FIGURE 27.**
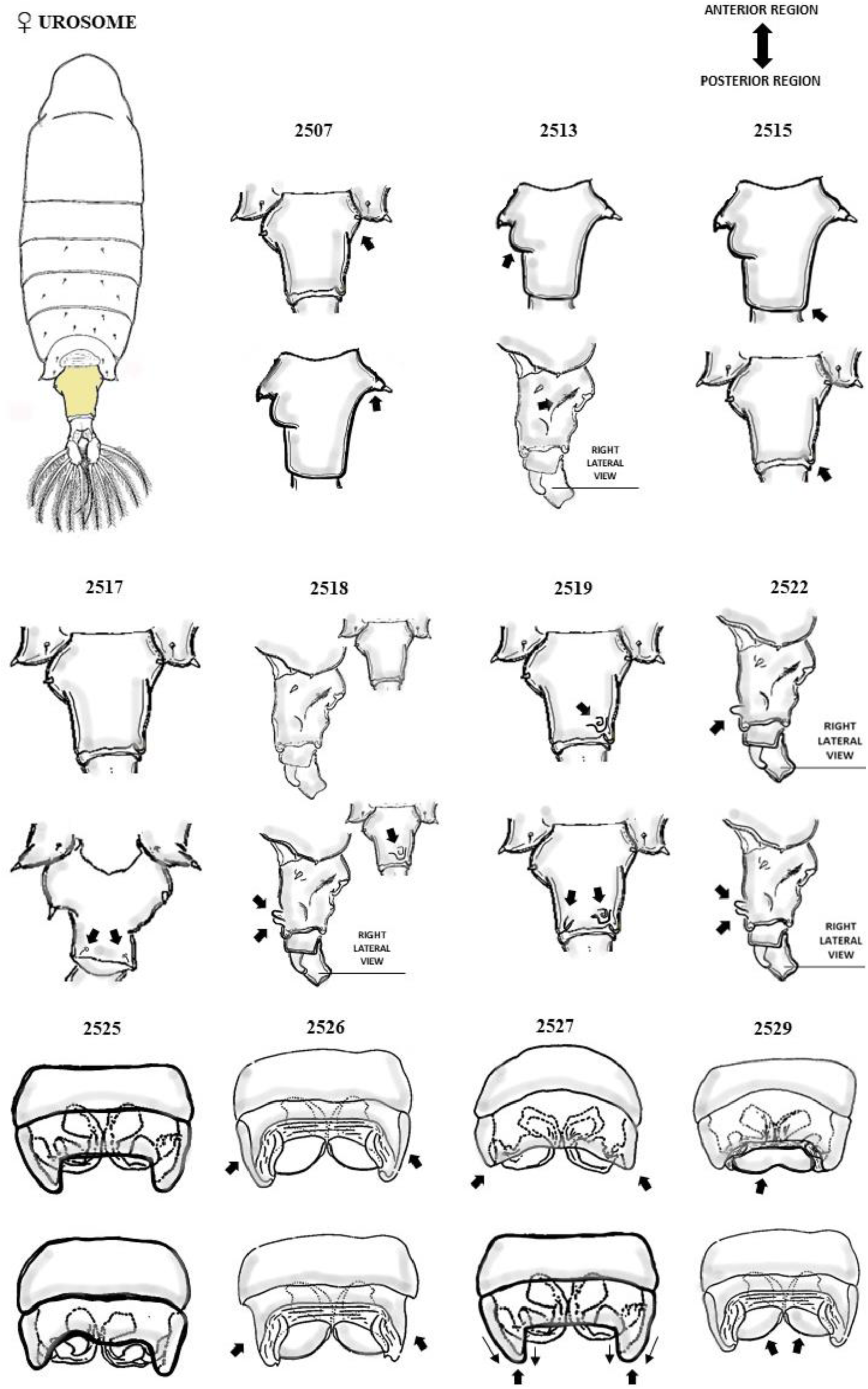
Attributes of the female urosome, and genital double-somite, respectively: 2507-2522 chars (dorsal view plan), and 2525-2529 chars (ventral view plan).

**FIGURE 28.**
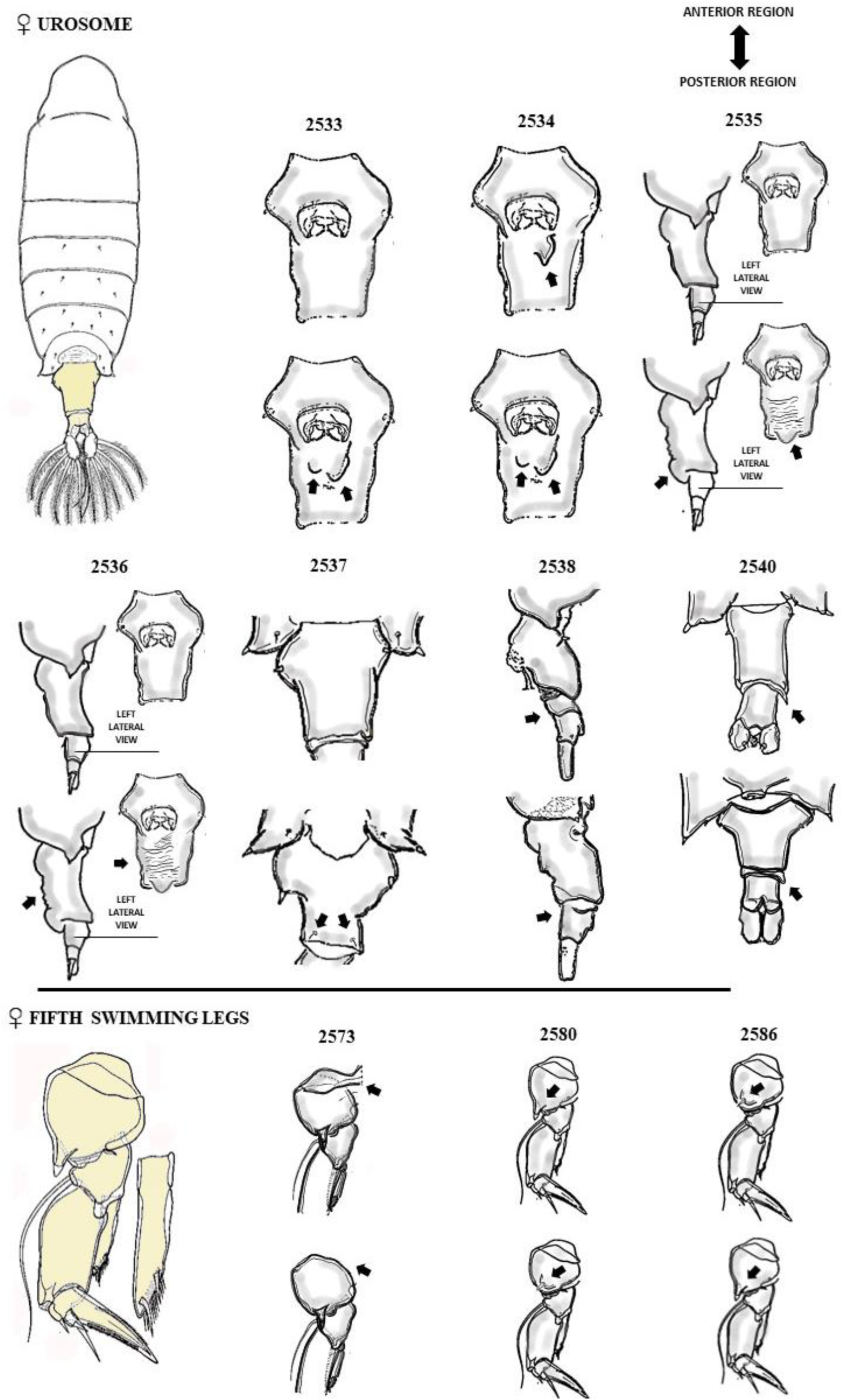
Attributes of the female urosome, and swimming leg 5, respectively: 2533-2540 chars in ventral view plan (2537 and 2540 in dorsal view, and 2538 in left lateral view), and 2573-2589 chars in posterior view plan.

**FIGURE 29.**
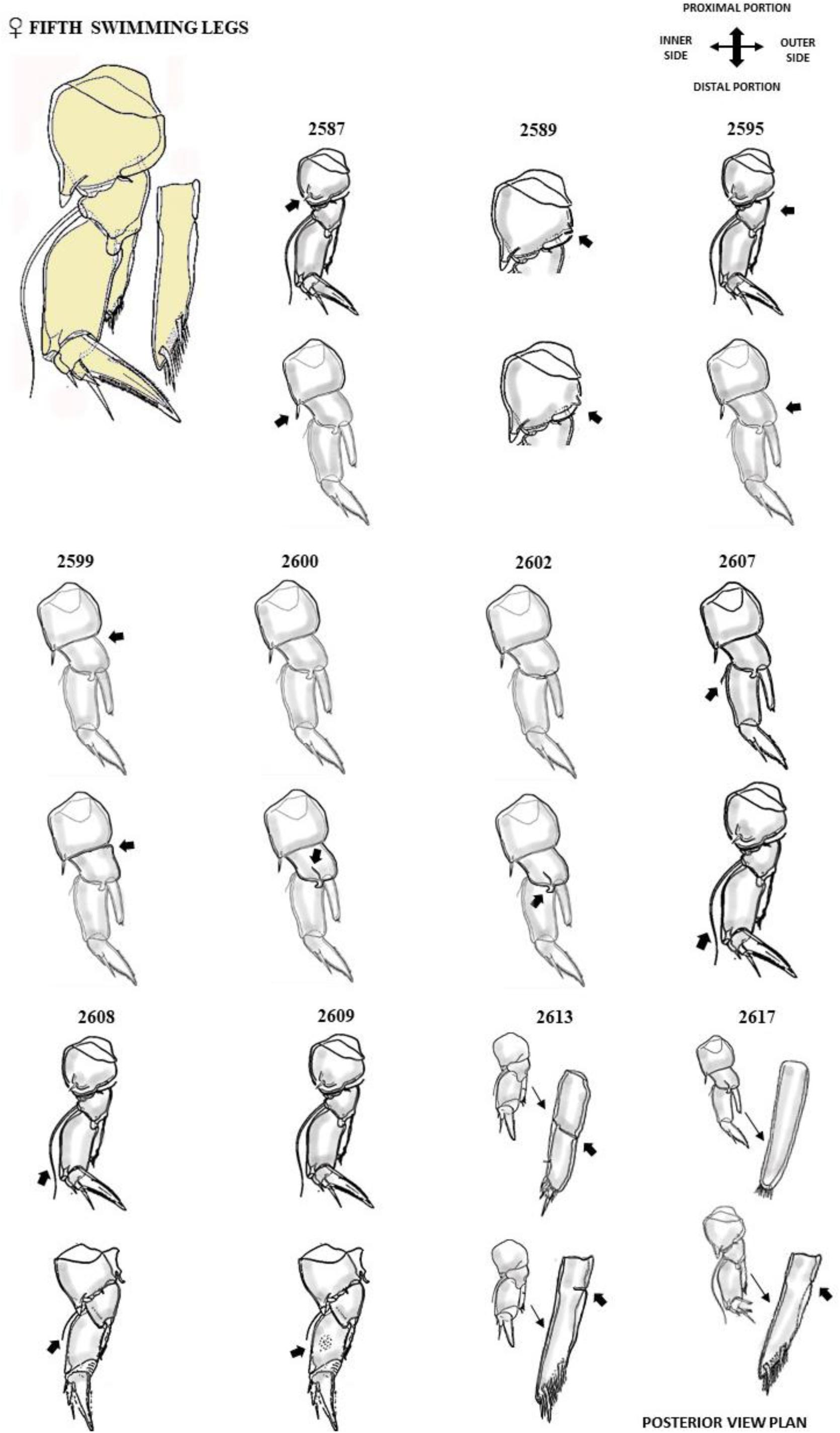
Attributes of the female swimming leg 5, respectively: 2587-2617 chars. Posterior view plan.

**FIGURE 30.**
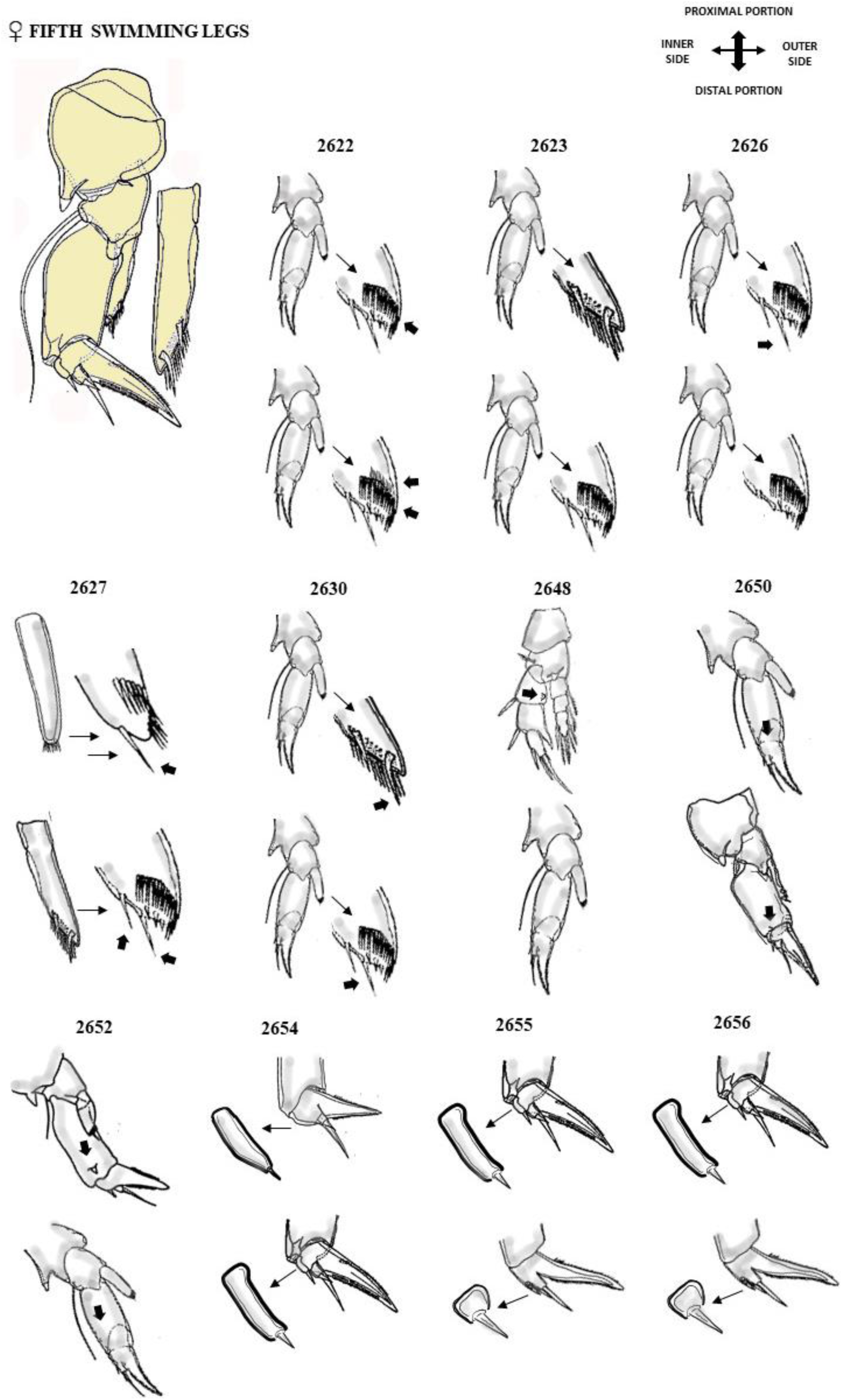
Attributes of the swimming legs 5 respectively: 2622-2656 chars. Posterior view plan.

**FIGURE 31.**
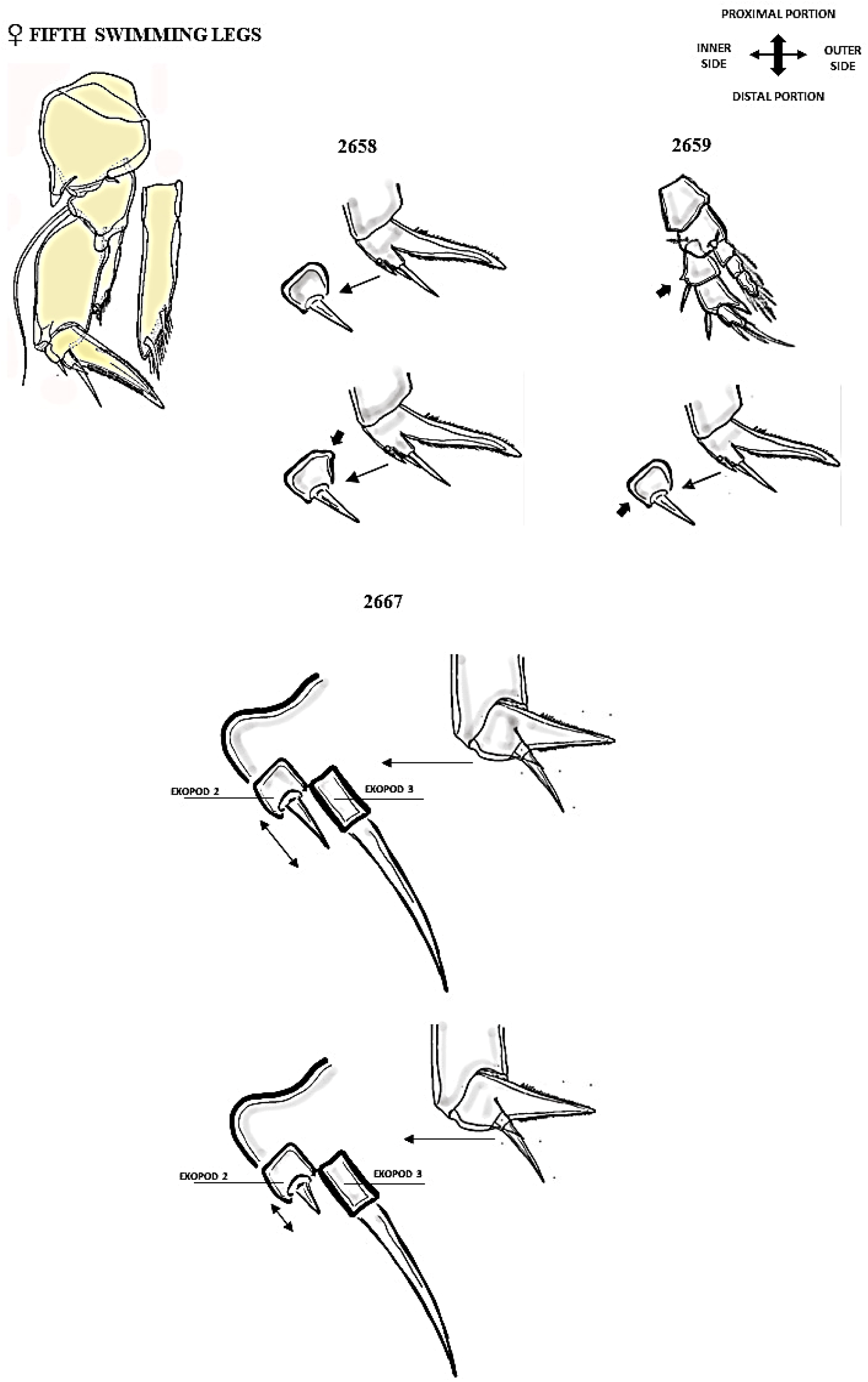
Attributes of the swimming legs 5 respectively: 2658-2667 chars. Posterior view plan.

**FIGURE 32.**
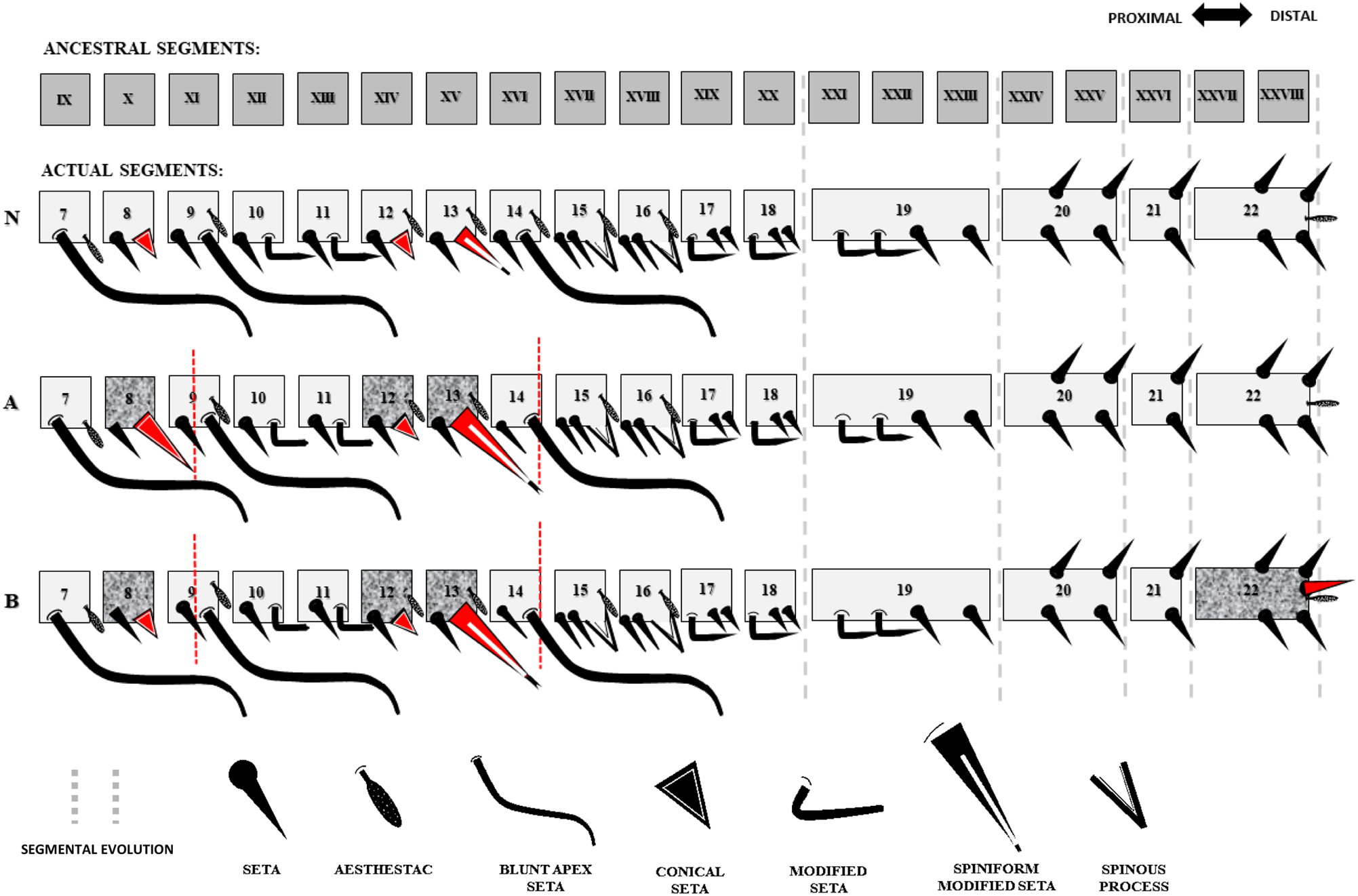
Schematization of the ornamental pattern variability between segment 7 to 22 of the male right antennule in *Notodiaptomus*. **Legend. “N”** to *Notodiaptomus* antennule pattern; **“A”** antennule variation A; **“B”** antennule variation B; segments bearing variable elements in shading; variable elements, and its range limits in red.

**FIGURE 33.**
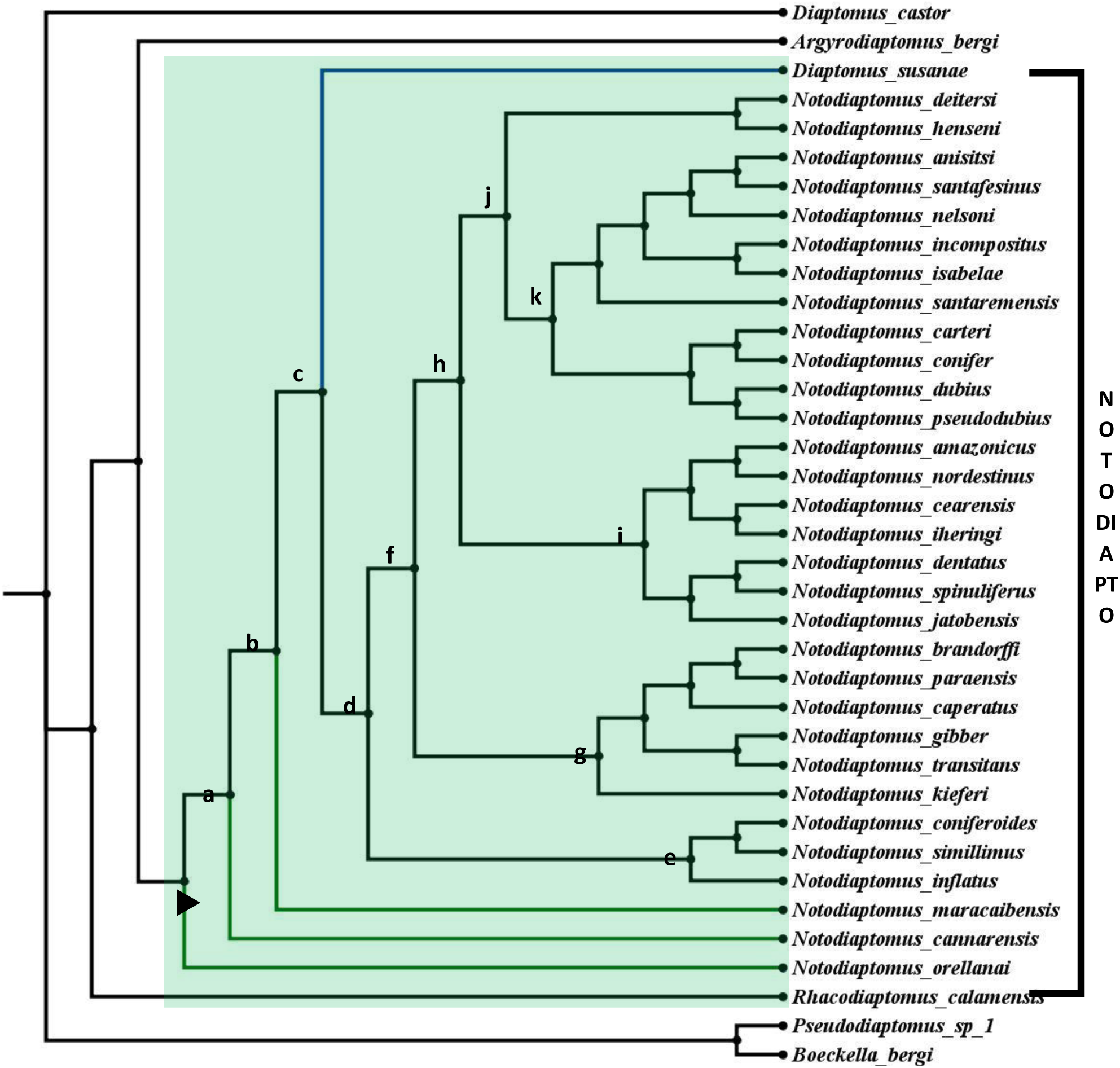
Graphical representation of the phylogenetic relationships obtained. Common and exclusive ancestor *Notodiaptomus* represented by ►; green field covers the cladograms of the inner group, partially represented by the letters a – k.

**TABLE 2.** Historical series of characteristics attributed to the genus *Notodiaptomus* Kiefer 1936. List of characters in Wright (1935), Kiefer (1936), Kiefer (1956) and Santos-Silva *et al*. (1999). Relative sharing frequency of each attribute analyzed between the congeners considering (% sharing), respectively.

| **A) In Wright, 1935. Original proposal to *nordestinus* complex** |  | **% Sharing** |
| --- | --- | --- |
| **1a.** | Male. 5th right leg basis with conspicuous proximal inner angle, rounded "shoulder", with or without small prominence | 68,75% |
| **2a.** | Male. 5th right leg "small" and conical endopod | subjective |
| **3a.** | Male. 5th right leg exopod 2 with angular or rounded prominence on inner margin <sup>(D)</sup> | 65,6% |
| **4a.** | Male. 5th left leg with exopods fused "narrowly", with presence of "hair" pad | 100% |
| **5a.** | Male. 5th left leg exopod with distal segment with "little spine" and ending in a finger-like process | 87,5% |
| **6a.** | Male. 5th left leg with endopod unisegmented | 87,5% |
| **7a.** | Female. 5th legs basis with outer seta on proeminence. | subjective |
| **8a.** | Female. 5th swimming legs basis with expanded distal inner rim posteriorly <sup>(D)</sup> | 87,5% |
| **B) In Kiefer, 1936. Original proposal of the genus *Notodiaptomus*** |  | **% Sharing** |
| **1b.** | Male. "The nature of the final segment of the left exopod of the male fifth leg" | subjective |
| **2b.** | Male. Right antennule with conspicuous spines on segments 10, 11, 13, 15 and 16 <sup>(D)</sup> | 96,87% |
| **3b.** | Male. Right antennule with hyaline lamella on 20 segments ("at most one") | high variation |
| **C) In Kiefer, 1956. Proposal for expansion to the genus *Notodiaptomus*** |  | **% Sharing** |
| **1c.** | Female. 5th swimming legs with endopod unisegmented <sup>(D)</sup> | 68,75% |
| **2c.** | Female. 5th swimming legs endopod with row of oblique spinules + 1-2 setae | 12,5% |
| **3c.** | Male. 5th swimming legs coxa with rounded lobule and "hyaline" spine | 100% |
| **4c.** | Male. 5th right swimming leg basis with structure in varied shape and size, lamella posteriorly <sup>(D)</sup> | 68,75% |
| **5c.** | Male. 5th right swimming leg exopod 1 with length greater than width, mostly <sup>(D)</sup> | 84,37% |
| **6c.** | Male. 5th right leg exopod 2 with rounded-broad hyaline protuberance on inner "caudal surface" <sup>(D)</sup> | 65,6% |
| 7c. | Male. 5th right swimming leg exopod 2 with outer spine on base of the terminal claw generally | 84,37% |
| 8c. | Male. 5th left swimming leg exopod 2 with distal portion digitiform | 84,37% |
| 9c. | Male. 5th left swimming leg exopod 2 with short broad seta, directed inwards, on distal portion | 59,37% |
| 10c. | Male. Right antennule without “spineform protuberance” on segment 14 <sup>(D)</sup> | 96,87% |
| 11c. | Male. Right antennule with narrow hyaline membrane on segment 20 generally | high variation |
| **D) In Santos-Silva *et al.*, 1999 (reassessed in Santos-Silva *et al.*, 2015)** |  | **% Sharing** |
| 1d. | Dimorphic rostrum | 100% |
| 2d. | Female. Antennules without aesthetasc on segments 13 and 15 | 100% |
| 3d. | Male. Left antennule with seta on segments 13, 14, 15, 16, 17 and 18 | 100% |
| 4d. | Male. Antennules with vestigial seta on segments 2, 3 and 5 <sup>(*)</sup> <sup>(D)</sup> | 100% |
| 5d. | Antennules with elongated seta on segments 3, 7, 9 and 14, and blunt apex <sup>(*)</sup> <sup>(D)</sup> | 90,60% |
| 6d. | Antennules with conical seta on segments 8 and 12 | 100% |
| 7d. | Antenna. Endopod 2-segmented | 93,70% |
| 8d. | Antenna. First segment with row of spinules and pore <sup>(D)</sup> | 81,25% |
| 9d. | Antenna. Exopod 7-segmented | 96,87% |
| 10d. | Swimming legs 1-4. “Poorly ornamented” | subjective |
| 11d. | Swimming leg 1. Endopod 2-segmented | 100% |
| 12d. | Swimming leg 1. Without external spine on exopod 2 | 100% |
| 13d. | Swimming leg 2. Schmeil's organ on endopod 2 posteriorly | 100% |
| 14d. | Swimming legs 2-4. Outer spine on each exopod | 100% |
| 15d. | Swimming legs 2-4. Distal row of spinules on exopod 3 anteriorly | 90,60% |
| 16d. | Swimming legs 2-4. Double distal row of spinules on endopod 3 anteriorly <sup>(D)</sup> | 93,75% |
| 17d. | Swimming leg 4. Posterior seta near outer margin of base | 100% |
| 18d. | Male. Cephalosome with incomplete suture dorsally <sup>(D)</sup> | 71,87% |
| 19d. | Male. Metasoma 5 with symmetrical epimeral plates “nearly” | subjective |
| 20d. | Male. Urosome 5-segmented | 100% |
| 21d. | Male. Symmetrical caudal rami | 100% |
| 22d. | Male. Caudal rami longer than wide | 100% |
| 23d. | Male. Caudal rami with setules on inner margin <sup>(*)</sup> | 100% |
| 24d. | Male. Right antennule geniculate 22-segmented | 100% |
| 25d. | Male. Right antennule with two setae on segment 20 posteriorly | 100% |
| 26d. | Male. Right antennule with seta on segment 21 ventrally | 100% |
| 27d. | Male. Right antennule with spinous process on segments 15 and 16 | 69,87% |
| 28d. | Male. Right antennule with modified setae on segments 10, 11, 13, 17, 18 and 19 | 93,75% |
| 29d. | Male. Right antennule with setae smaller than the others in segment 19 | 100% |
| 30d. | Male. Right antennule with modified setae as process on segments 10, 11 and 13 | 100% |
| 31d. | Male. Right antennule with modified seta on segment 13 larger than those on 10 and 11 <sup>(D)</sup> | 100% |
| 32d. | Male. Right antennule with modified setae on segs. 17, 18 and 19 different from those on 10 and 11 | 93,75% |
| 33d. | Left antennule. Second and last composite segment | 100% |
| 34d. | Left antennule. Setae on segment 22 posteriorly | 100% |
| 35d. | Left antennule. Setae on segment 23 posteriorly <sup>(D)</sup> | 100% |
| 36d. | Left antennule. Setae on segment 24 ventrally <sup>(D)</sup> | 100% |
| 37d. | Left antennule. Absence aestetasc on segment 13 | 100% |
| 38d. | Left antennule. Absence aestetasc on segment 15 | 100% |
| 39d. | Male. 5th swimming legs asymmetrical | 100% |
| 40d. | Male. 5th right swimming leg with rudimentary precoxa <sup>(*)</sup> (D) | 100% |
| 41d. | Male. 5th right swimming leg coxa with conical process projecting over base (D) | 90,60% |
| 42d. | Male. 5th right swimming leg coxa with conical process with sharp apex sensillae | 100% |
| 43d. | Male. 5th right swimming leg basis with posterior medial growth (“posterior protrusion”) <sup>(*)</sup> (D) | 68,75% |
| 44d. | Male. 5th right swimming leg basis with oblique deep groove <sup>(*)</sup> (D) | 59,37% |
| 45d. | Male. 5th right swimming leg basis with tubercles over margin of oblique groove <sup>(*)</sup> | 43,75% |
| 46d. | Male. 5th right swimming leg basis with semicircular inner lamella | 31,25% |
| 47d. | Male. 5th right swimming leg basis with tubercles covering semicircular lamella and adjacent area | 12,50% |
| 48d. | Male. 5th right swimming leg exopod 2-segmented | 100% |
| 49d. | Male. 5th right swimming leg exopod 1 with acute distal process (“outgrowth”) <sup>(*)</sup> | 40,62% |
| 50d. | Male. 5th right swimming leg exopod 1 with triangular outer process posteriorly (D) | 93,75% |
| 51d. | Male. 5th right swimming leg exopod 2 with distal-internal curve on posterior surface <sup>(*)</sup> (D) | 65,62% |
| 52d. | Male. 5th right swimming leg exopod 2 with outer spine on the “distal third” | 83,30% |
| 53d. | Male. 5th right swimming leg exopod 2 with curved terminal claw proximally (D) | 78,12% |
| 54d. | Male. 5th right swimming leg terminal claw with a row of spinules internally | 81,25% |
| 55d. | Male. 5th right swimming leg endopod with spinules on distal inner rim of lobe | 100% |
| 56d. | Male. 5th right swimming leg endopod with spinules on inner-anterior surface | 100% |
| 57d. | Male. 5th left swimming leg coxa with conical process, and sensillae on apex | 100% |
| 58d. | Male. 5th left swimming leg basis with tubercles group on inner-proximal surface | 28,12% |
| 59d. | Female. Rostrum symetrical and without basal process | 100% |
| 60d. | Female. Metasome 5 with symetrical epimeral plates | 12,50% |
| 61d. | Female. Urosome 3-segmented | 96,87% |
| 62d. | Female. Urosome with symetrical genital double-somite and inflated anteriorly <sup>(*)</sup> | 3,12% |
| 63d. | Female. Urosome with not fully developed operculum on anal segment | 21,87% |
| 64d. | Female. Symetrical caudal rami with setules innerly <sup>(*)</sup> | 96,87% |
| 65d. | Female. 5th legs with sub-triangular basis | 100% |
| 66d. | Female. 5th legs basis with seta reaching beyond of the distal margin of the exopod 1 | 21,87% |
| 67d. | Female. 5th legs exopod 1 with length greater than length of the exopod 2 | 100% |
| 68d. | Female. 5th legs exopod 2 with outer spine (lateral spine) <sup>(*)</sup> | 100% |
| 69d. | Female. 5th legs exopod 2 with denticles on terminal claw medially and “laterally” (D) | 100% |
| 70d. | Female. 5th legs endopod 1-segmented (D) | 68,75% |
| 71d. | Female. 5th legs endopod with two setae (D) | 84,37% |
| 72d. | Female. 5th legs endopod with oblique row of spinules sub-terminally and anteriorly | 100% |

<sup>(\*)</sup> Differential characteristic to *nordestinus* complex diagnosis by Santos-Silva *et al.* (2015). <sup>(D)</sup> Diagnostic attributes identified to genus *Notodiaptomus*.

**TABLE 3.**
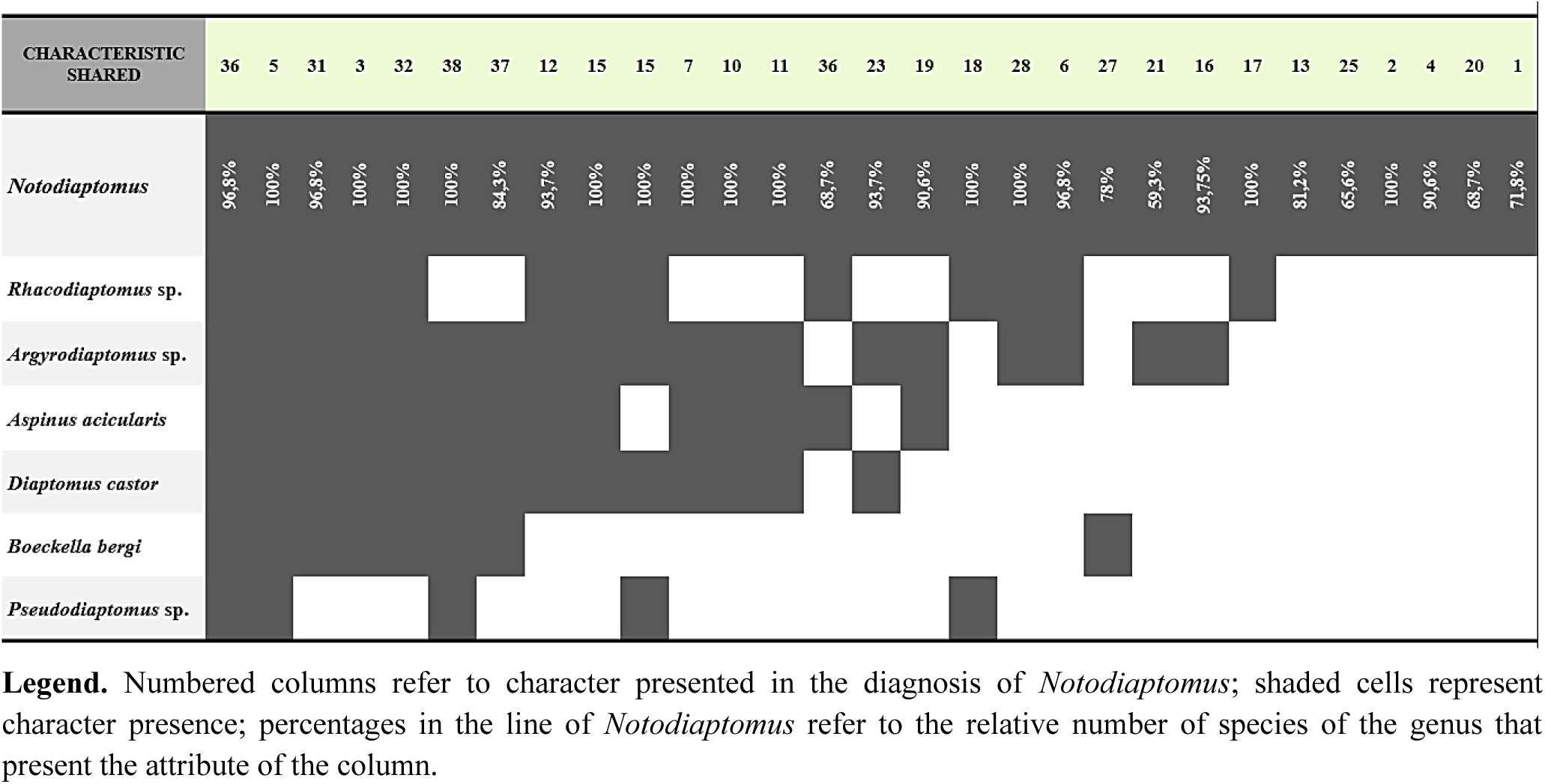
Morphological basis and sharing of diagnostic attributes of *Notodiaptomus* abducted in literature, focus on Santos-Silva *et al*. (1999).

##### Type locality

Lake in Cuiaba city, Mato Grosso, Brazil. The referred lake does not exist more. Neotype: BRAZIL: Mato Grosso, Pedra Branca Bay.

##### Type material

Neotype: 1 male, adult dissected, built on 20 slides, X.30.1996, collected by Vangil Pinto da Silva (INPA - CR 550a-t), and designated by Santos-Silva *et al*. (2015). Additional material of the same place, date e collector: 1 female adult dissected on twenty slides; 231 females, and 57 males in alcohol (INPA - CR 551a-t, 552 e 676, specially); four pack of 20 male, and 20 females in alcohol, stored in MZUSP.

##### Material examined

Neotype: 1 male adult (INPA - CR 550a-t); 1 female (INPA - CR 551a-t, 552 and 676, specially). Additional material: 2 males, and 3 females, entire in alcohol, deposited in collection of the Plankton Laboratory in the Instituto Nacional de Pesquisas da Amazônia; 1 male, and 1 female, entire in alcohol, stored in Zoology Museum of Sao Paulo (no code). 1 male (INPA-COP001, slides a-h) and 1 female (INPA-COP002, slides a-h) were selected to be dissection on eight slides each and deposited in the Zoological Collection of the INPA, Brazil.

##### Diagnosis

**(1)** first swimming legs exopod 3 with presence of spinules; **(2)** male fifth left swimming leg basis with presence of minutely granular as a patch innerly; **(3)** male fifth left swimming leg exopod 2 with setulose pad not rounded prominently; **(4)** male fifth right swimming leg exopod 2 with curved ridge on distal posterior surface closer to innerly; **(5)** male fifth right swimming leg exopod 2 with outer spine in rectilinear form; **(6)** female fourth and fifth metasome segments separated; **(7)** female right epimeral plate without dorsal-posterior sensilla; **(8)** female right antennule not similarity to male full, absence of setae on actual segments 13, 14, 15, 16, 17, 18, and absence of aestestasc on actual segments 13, 14, 15, and 16; **(9)** female fifth swimming legs basis with outer seta reaching to exopod 1 distally; **(10)** female fifth swimming legs exopod 3 with outer seta smaller than inner nearly 1 time.

##### Redescription

###### MALE

Body 1166 micrometers excluding caudal setae. Male body smaller and slenderer than female. Nerve axons myelinated. Prosome 6-segmented; widest at first metasome segment; without one line of setules at posterior margin; without spinules at segments. Cephalosome anterior margin rounded; with dorsal suture; incomplete; separate from first metasome segment. First metasome segment without sensilla. Second metasome segment with sensilla; 2 dorsally; of equal size. Third metasome segment with sensillae; 4 laterally; of unequal size; non-ornamented posterior margin. Fourth metasome segment with sensillae; 2 dorsally; 4 laterally; of equal size; separated from the fifth metasome. Limit between fourth and fifth metasome segments without ornamentation. Fifth metasome segment with sensilla; 4 laterally; fifth metasome segment equal size; fifth metasome segment without ornamentation; Fifth metasome segment without dorsal conical process; with epimeral plates. Epimeral plates symmetrical. Right epimeral plates reduced, as rounded distal corner segment limit; with sensilla; at the apex of projection; without ornamentation.

##### Urosome

5-segmented; Urosome 5 - free segments. Genital somite asymmetrical in dorsal view; with single aperture; located on left side; ventrolaterally on posterior rim; with sensillae; on both sides; one; at left lateral; posteriorly; one; at right rim; posteriorly; of equal size between then. Third urosome segment without spinules; without external seta. Fourth urosome segment without spinules; without sub-conical blunt dorsal-lateral process. Anal segment presence of dorsal sensillae; one on each side; medially inserted; presence of operculum; convex; covering the anal aperture fully. Caudal rami symmetrical; separated from anal segment; longer than wide; with setules; continuous on; inner side; each ramus bearing 6 caudal setae; 5 marginals; plumose; and 1 internal dorsally; straight; not reticulated main axis; outermost seta with outer spiniform process absent.

##### Appendices features

Rostrum symmetrical; separated from dorsal cephalic shield; by complete suture; sensillae present; one pair; anteriorly inserted on surface tegument; with rostral filament; double; paired; extended; into point; with basal process; in ventral view, rounded on left side; without a smaller basal expansion on the right side.

##### Antennules

Asymmetrical. **Right antennules**. Uniramous; right antennule surpassing to genital segment; right antennule extending beyond caudal rami.

Right antennule ancestral segment I and II separated. Ancestral segment II and III fused. Ancestral segment III and IV fused. Ancestral segment IV and V separated. Ancestral segment V and VI separated. Ancestral segment VI and VII separated. Ancestral segment VII and VIII separated. Ancestral segment VIII and IX separated. Ancestral segment IX and X separated. Ancestral segment X and XI separated. Ancestral segment XI and XII separated. Ancestral segment XII and XIII separated. Ancestral segment XIII and XIV separated. Ancestral segment XIV and XV separated. Ancestral segment XV and XVI separated. Ancestral segment XVI and XVII separated. Ancestral segment XVII and XVIII separated. Ancestral segment XVIII and XIX separated. Ancestral segment XIX and XX separated. Ancestral segment XX and XXI separated. Ancestral segment XXI and XXII fused. Ancestral segment XXII and XXIII fused. Ancestral segment XXIII and XXIV separated. Ancestral segment XXIV and XXV fused. Ancestral segment XXV and XXVI separated. Ancestral segment XXVI and XXVII separated. Ancestral segment XXVII and XXVIII fused.

Right antennule actual 22-segmented; geniculated; between the segment 18 and segment 19; with swollen and modified region; formed by 5 segments; between 13 and 17 segments. Actual segment 1 with seta; one element; straight; none larger than segment; without spinules; without vestigial seta; without conical seta; without modified seta; without spinous process; with aesthetasc; one element. Actual segment 2 with seta; three elements; of unequal size; straight; none larger than segment; without spinules; with vestigial seta; one element; without conical seta; without modified seta; without spinous process; with aesthetasc; one element. Actual segment 3 with seta; one element; one larger than segment; surpassing to distal margin; beyond three sequential segments; straight; blunt apex; without spinules; with vestigial seta; one element; without conical seta; without modified seta; without spinous process; with aesthetasc. Actual segment 4 with seta; one element; one larger than segment; surpassing to distal margin; straight; not beyond three sequential segments; without spinules; without vestigial seta; without conical seta; without modified seta; without spinous process; without aesthetasc. Actual segment 5 with seta; one element; straight; one larger than segment; surpassing to distal margin; not beyond three sequential segments; without spinules; with vestigial seta; one element; without conical seta; without modified seta; without spinous process; with aesthetasc; one element. Actual segment 6 with seta; one element; none larger than segment; straight; without spinules; without vestigial seta; without conical seta; without modified seta; without spinous process; without aesthetasc. Actual segment 7 with seta; one element; straight; one larger than segment; surpassing to distal margin; beyond three sequential segments; blunt apex; without spinules; without vestigial seta; without conical seta; without modified seta; without spinous process; with aesthetasc; one element. Actual segment 8 with seta; one element; straight; none larger than segment; without spinules; without vestigial seta; with conical seta; one element; not reaching to middle-point of the sequent segment; without modified seta; without spinous process; without aesthetasc. Actual segment 9 with seta; two elements; of unequal size; straight; one larger than segment; surpassing to distal margin; beyond three sequential segments; blunt apex; without spinules; without vestigial seta; without conical seta; without modified seta; without spinous process; with aesthetasc; one element. Actual segment 10 with seta; one element; straight; none larger than segment; without spinules; without vestigial seta; without conical seta; with modified seta; presenting blunt apex; slender form; surpassing to distal margin; beyond of the sequential segment; parallel to antennule direction; without spinous process; without aesthetasc. Actual segment 11 with seta; one element; straight; one larger than segment; surpassing to distal margin; not beyond three sequential segments; without spinules; without vestigial seta; without conical seta; with modified seta; slender form; presenting blunt apex; surpassing to distal margin; beyond of the sequential segment; parallel to antennule direction; shorter length than homologous of actual segment 13; without spinous process; without aesthetasc. Actual segment 12 with seta; one element; straight; one larger than segment; surpassing to distal margin; not beyond three sequential segments; without spinules; without vestigial seta; with conical seta; one element; not smaller than to segment 8; without modified seta; without spinous process; with aesthetasc; one element; absent internal perpendicular fission. Actual segment 13 with seta; one element; straight; one larger than segment; surpassing to distal margin; not beyond three sequential segments; without spinules; without vestigial seta; without conical seta; with modified seta; stout form; surpassing to distal margin; to the middle-point of the sequence segment; perpendicular to antennule direction; presenting bifid apex; without spinous process; with aesthetasc; one element. Actual segment 14 with seta; two elements; of unequal size; straight; one larger than segment; surpassing to distal margin; beyond three sequential segments; blunt apex; without spinules; without vestigial seta; without conical seta; without modified seta; without spinous process; with aesthetasc; one element. Actual segment 15 with seta; two elements; of unequal size; straight; not bifidform; none larger than segment; without spinules; without vestigial seta; without conical seta; without modified seta; with spinous process; on outer margin; surpassing distal margin; with aesthetasc; one element. Actual segment 16 with seta; two elements; of unequal size; plumose; one larger than segment; surpassing to distal margin; not beyond three sequential segments; not bifidform; without spinules; without vestigial seta; without conical seta; without modified seta; with spinous process; on outer margin; surpassing distal margin; unequal size to process on preceding segment; with aesthetasc; one element. Actual segment 17 with seta; two elements; of unequal size; straight; none larger than segment; bifidform; without spinules; without vestigial seta; without conical seta; with modified seta; one element; stout form; surpassing to distal margin; not beyond of the sequential segment; parallel to antennule direction; without spinous process; without aesthetasc. Actual segment 18 with seta; two elements; of equal size; straight; none larger than segment; without spinules; without vestigial seta; without conical seta; with modified seta; one element; stout form; surpassing distal margin; parallel to antennule direction; without spinous process; without aesthetasc. Actual segment 19 with seta; two elements; of unequal size; plumose; none larger than segment; without spinules; without vestigial seta; without conical seta; with modified seta; two elements; stout form; at least one bifid form; surpassing distal margin; parallel to antennule direction; without spinous process; with aesthetasc; one element. Actual segment 20 with seta; four elements; of unequal size; straight; one larger than segment; surpassing to distal margin; beyond three sequential segments; without spinules; without vestigial seta; without conical seta; without modified seta; without spinous process; without aesthetasc. Actual segment 21 with seta; two elements; of equal size; plumose; one larger than segment; surpassing to distal margin; greater 3x than original segment; without spinules; without vestigial seta; without conical seta; without modified seta; without spinous process; without aesthetasc. Actual segment 22 with seta; four elements; of equal size; one larger than segment; plumose; surpassing to distal margin; greater 3x than original segment; without spinules; without vestigial seta; without conical seta; without modified seta; without spinous process; with aesthetasc; one element.

##### Left antennules

Uniramous; Left antennule surpassing to prosome; Left antennule extending beyond caudal rami. Ancestral segment I and II separated. Ancestral segment II and III fused. Ancestral segment III and IV fused. Ancestral segment IV and V separated. Ancestral segment V and VI separated. Ancestral segment VI and VII separated. Ancestral segment VII and VIII separated. Ancestral segment VIII and IX separated. Ancestral segment IX and X separated. Ancestral segment X and XI separated. Ancestral segment XI and XII separated. Ancestral segment XII and XIII separated. Ancestral segment XIII and XIV separated. Ancestral segment XIV and XV separated. Ancestral segment XV and XVI separated. Ancestral segment XVI and XVII separated. Ancestral segment XVII and XVIII separated. Ancestral segment XVIII and XIX separated. Ancestral segment XIX and XX separated. Ancestral segment XX and XXI separated. Ancestral segment XXI and XXII separated. Ancestral segment XXII and XXIII separated. Ancestral segment XXIII and XXIV separated. Ancestral segment XXIV and XXV separated. Ancestral segment XXV and XXVI separated. Ancestral segment XXVI and XXVII separated. Ancestral segment XXVII and XXVIII fused.

Left antennule actual 25-segmented; not-geniculated. Actual segment 1 with seta; one element; none larger than segment; straight; without spinules; without vestigial seta; without conical seta; without modified seta; without spinous process; with aesthetasc; one element. Actual segment 2 with seta; three elements; of equal size; none larger than segment; straight; without spinules; with vestigial seta; one element; without conical seta; without modified seta; without spinous process; with aesthetasc; one element. Actual segment 3 with seta; one element; one larger than segment; straight; surpassing to distal margin; beyond three sequential segments; without spinules; with vestigial seta; one element; without conical seta; without modified seta; without spinous process; with aesthetasc. Actual segment 4 with seta; one element; none larger than segment; straight; without spinules; without vestigial seta; without conical seta; without modified seta; without spinous process; without aesthetasc. Actual segment 5 with seta; one element; one larger than segment; straight; surpassing to distal margin; not beyond three sequential segments; without spinules; with vestigial seta; one element; without conical seta; without modified seta; without spinous process; with aesthetasc; one element. Actual segment 6 with seta; one element; none larger than segment; straight; without spinules; without vestigial seta; without conical seta; without modified seta; without spinous process; without aesthetasc. Actual segment 7 with seta; one element; one larger than segment; straight; surpassing to distal margin; beyond three sequential segments; without spinules; without vestigial seta; without conical seta; without modified seta; without spinous process; with aesthetasc; one element. Actual segment 8 with seta; one element; one larger than segment; straight; surpassing distal margin; without spinules; without vestigial seta; with conical seta; without modified seta; without spinous process; without aesthetasc. Actual segment 9 with seta; two elements; of unequal size; one larger than segment; straight; surpassing to distal margin; beyond three sequential segments; without spinules; without vestigial seta; without conical seta; without modified seta; without spinous process; with aesthetasc; one element. Actual segment 10 with seta; one element; none larger than segment; straight; without spinules; without vestigial seta; without conical seta; without modified seta; without spinous process; without aesthetasc. Actual segment 11 with seta; one element; one larger than segment; straight; surpassing to distal margin; beyond three sequential segments; without spinules; without vestigial seta; without conical seta; without modified seta; without spinous process; without aesthetasc. Actual segment 12 with seta; one element; one larger than segment; straight; surpassing distal margin; without spinules; without vestigial seta; with conical seta; without modified seta; without spinous process; with aesthetasc; one element. Actual segment 13 with seta; one element; none elongated; straight; surpassing distal margin; without spinules; without vestigial seta; without conical seta; without modified seta; without spinous process; without aesthetasc. Actual segment 14 with seta; one element; elongated; straight; surpassing to distal margin; beyond three sequential segments; without spinules; without vestigial seta; without conical seta; without modified seta; without spinous process; with aesthetasc; one element. Actual segment 15 with seta; one element; larger than segment; straight; surpassing to distal margin; not beyond three sequential segments; without spinules; without vestigial seta; without conical seta; without modified seta; without spinous process; without aesthetasc. Actual segment 16 with seta; one element; larger than segment; plumose; surpassing to distal margin; not beyond three sequential segments; without spinules; without vestigial seta; without conical seta; without modified seta; without spinous process; with aesthetasc; one element. Actual segment 17 with seta; one element; not larger than segment; straight; without spinules; without vestigial seta; without conical seta; without modified seta; without spinous process; without aesthetasc. Actual segment 18 with seta; one element; larger than segment; straight; surpassing to distal margin; beyond three sequential segments; without spinules; without vestigial seta; without conical seta; without modified seta; without spinous process; without aesthetasc. Actual segment 19 with seta; one element; not larger than segment; straight; surpassing distal margin; without spinules; without vestigial seta; without conical seta; without modified seta; without spinous process; with aesthetasc; one element. Actual segment 20 with seta; one element; not larger than segment; straight; surpassing distal margin; without spinules; without vestigial seta; without conical seta; without modified seta; without spinous process; without aesthetasc. Actual segment 21 with seta; one element; larger than segment; plumose; surpassing to distal margin; beyond three sequential segments; without spinules; without vestigial seta; without conical seta; without modified seta; without spinous process; without aesthetasc. Actual segment 22 with seta; two elements; of unequal size; one of them elongated; plumose; surpassing to distal margin; without spinules; without vestigial seta; without conical seta; without modified seta; without spinous process; without aesthetasc. Actual segment 23 with seta; two elements; of unequal size; one larger than segment; plumose; surpassing to distal margin; greater 3x than original segment; without spinules; without vestigial seta; without conical seta; without modified seta; without spinous process; without aesthetasc. Actual segment 24 with seta; two elements; of equal size; one larger than segment; plumose; surpassing to distal margin; greater 3x than original segment; without spinules; without vestigial seta; without conical seta; without modified seta; without spinous process; without aesthetasc. Actual segment 25 with seta; four elements; of equal size; elongated; plumose; surpassing to distal margin; 4 times larger than segment; without spinules; without vestigial seta; without conical seta; without modified seta; without spinous process; with aesthetasc; one element.

##### Antenna

Biramous. Antenna coxa separated from the basis; bearing seta; 1; on inner surface; at distal corner; reaching to the endopod 1. Antenna basis (fusion) separated from the endopodal segment; bearing seta; 2; on inner surface; at distal corner. Endopodal ancestral segment I and II separated. Ancestral segment II and III fused. Ancestral segment III and IV fused. Ancestral segment III and IV fully. Antenna endopod actual 2-segmented. Actual segment 1 not bilobate; with seta; two; on inner margin; with spinules; as a row; obliquely; on outer surface; with pore. Actual segment 2 bilobate; with discontinuity on outer cuticle; not developed as a suture; inner lobe bearing 8 setae; distally; outer lobe bearing 7 setae; distally; with spinules; as a patch; on outer surface. Antenna exopod ancestral segment I and II separated. Ancestral segment II and III fused. Ancestral segment III and IV fused. Ancestral segment IV and V separated. Ancestral segment V and VI separated. Ancestral segment VI and VII separated. Ancestral segment VII and VIII separated. Ancestral segment VIII and IX separated. Ancestral segment IX and X fused. Antenna exopod actual 7-segmented. Actual segment 1 single; elongated (width-length, equal or larger ratio 2:1); with seta; one; at inner surface. Actual segment 2 compound; elongated (larger width-length ratio 2:1); with seta; three; at inner surface. Actual segment 3 single; not elongated (lesser width-length ratio 2:1); with seta; one; at inner surface. Actual segment 4 single; not elongated (lesser width-length ratio 2:1); with seta; one; at inner surface. Actual segment 5 single; not elongated (lesser width-length ratio 2:1); with seta; one; at inner surface. Actual segment 6 single; not elongated (lesser width-length ratio 2:1); with seta; one; at inner surface. Actual segment 7 compound; elongated (larger or equal width-length ratio 2:1); with seta; one; at inner surface; and three; at distal surface.

##### Oral features. Mandible

Coxal gnathobase sclerotized; with lobe; prominent; on caudal margin; presence of cutting blade; with tooth-like prominence; two, distinctly; 1 acute; on caudal margin; and 1 triangular; on sub-caudal margin; without acute projection between the prominences; with additional spinules; as a row; on dorsal surface; with seta; 1; dorsally; on apical surface; with spinules; apicalmost. Mandible palps biramous; comprising the basis; with seta; four; differently inserted; first medially; reaching to beyond the endopod 1; second distally; third distally; fourth distally; on inner margin; none with setulose ornamentation. Mandible endopod 2-segmented. Mandible endopod 1 with lobe; bearing seta; four; distally inserted; without spinules. Mandible endopod 2 without lobe; bearing setae; nine elements; distally inserted; with spinules; as a row; double. Mandible exopod 4-segmented. Mandible exopod 1 with seta; one element; distally; on inner margin. Mandible exopod 2 with seta; one element; distally; on inner side. Mandible exopod 3 with seta; one element; distally; on inner side. Mandible exopod 4 with setae; three elements; on terminal region. **Maxillule**. Birramous. Maxillule 3-segmented. Maxillule praecoxa with praecoxal arthrite; bearing spines; fifteen elements; ten marginally; plus, five sub-marginally; with spinules; as a patch; on sub-marginal surface. Maxillule coxa with coxal epipodite; with conspicuous outer lobe; bearing setae; nine elements; with coxal endite; elongated (larger or equal width-length ratio 2:1); bearing setae; four elements. Maxillule basis with basal endite; double; first proximal; elongated (larger width-length ratio 2:1; separated from basis; with setae; four elements; distally inserted; second distal; fused to basis; not elongated (lesser width-length ratio 2:1); with setae; four elements; distally inserted; with setules; as a row; on inner side; basal exite present; with setae; one element; on outer surface. Maxillule endopod 1-segmented. Endopod 1 bilobate; first proximal; with setae; three elements; second distal; with setae; five elements. Maxillule exopod 1-segmented. Exopod 1 with setae; six elements; with setules; as a row; on inner side; spinules absent. **Maxilla**. Uniramous. Maxilla 5-segmented. Maxilla praecoxa fused to coxa; incompletely; distinct externally; with praecoxal endite; double; first elongated endite (larger or equal width length ratio 2:1); proximally inserted; with seta; straight, or plumose; 1 straight; 4 plumose; with spine; single; without spinules; without setule; second elongated endite (larger or equal width length ratio 2:1); distally inserted; with seta; plumose; 3 plumose; without spine; with spinules; as a row; on distal margin; with setule; as a row; on distal margin; absence of outer seta. Maxilla coxa with coxal endite; double; first elongated endite (larger or equal width); proximally inserted; with seta; plumose; 3 plumose; without spine; without spinules; with setules; as a row; on proximal margin; second elongated endite (larger or equal width); distally inserted; with seta; plumose; 3 plumose; without spine; without spinules; with setules; as a row; on proximal margin; absence of outer seta. Maxilla basis with basal endite; single; elongated (larger or equal width-length ratio 2:1); with seta; plumose; 3 plumose; without spinules; absence of outer seta. Maxilla endopod 2-segmented. Endopod 1 with seta; 2 plumose; without spine; without spinules; without setules. Maxilla endopod 2 with seta; 2 plumose; without spine; without spinules; without setules. **Maxilliped**. Uniramous; Maxilliped 8-segmented. Maxilliped praecoxa fused to coxa; incompletely; distinct internally; with praecoxal endite; not elongated (lesser width-length ratio 2:1); distally inserted; with seta; 1 straight; with spinules; as a row; single; on basal surface; without setules. Maxilliped coxa with coxal endite; three coxal endite; first elongated (larger or equal width); proximally inserted; with seta; 2 plumose; with spinules; as a patch; single; on apical surface; without setules; second not elongated (lesser width-length ratio 2:1); medially inserted; with seta; 3 plumose; with spinules; as a row; single; on medial surface; without setules; third elongated (larger or equal width length ratio 2:1); distally inserted; with seta; 3 plumose; none reaching to beyond of the basis; with spinules; as a row; single; on basal surface; without setules; with lobe; prominence; at inner distal angle; ornamented; with spinules; continuously on margin. Maxilliped basis without basal endite; with seta; 3 plumose; with spinules; as a row; single; on medial surface; with setules; as a row; single; on inner margin. Maxilliped endopod segment 6-segmented. Endopod 1 with seta; 2 plumose; on inner surface. Endopod 2 with seta; 3 plumose; on inner surface. Endopod 3 with seta; 2 plumose; on inner surface. Endopod 4 with seta; 2 plumose; on inner surface. Endopod 5 with seta; 2 plumose; on inner surface, or on outer surface; outer seta absent. Endopod 6 with seta; 4 plumose; on inner surface, or on outer surface.

##### Swimming legs features

**First swimming legs.** Symmetrical; biramous. First swimming legs intercoxal plate without seta. First swimming legs praecoxa absent. First swimming legs coxa with seta; one; straight; distally inserted; on inner surface; surpassing to first endopodal segment; with setules; two group; as a patch; on inner margin; and as a row; double; on anterior surface; outerly; without spinules; without spine. First swimming legs basis without seta; with setules; as a patch; single; on outer surface; without spinules; without spine. First swimming legs endopod 2-segmented. Endopod 1 with seta; straight; restricted; to inner surface; one element; without spine; with setules; as a row; single; continuously; on outer surface; without spinules; absence of Schmeil’s organ. Endopod 2 with seta; unrestricted; three on inner surface; one on outer surface; two on distal surface; straight; without spine; with setules; as a row; single; continuously; on outer surface; without spinules; absence of Schmeil’s organ. Endopod 3 absence. First swimming legs exopod 1 with seta; restricted; 1 on inner surface; with spine; 1; stout; smaller than original segment; serrated; on inner side; continuously; with setules; as a row; single; as a row; innerly. First swimming legs exopod 2 with seta; restricted; 1 on inner surface; straight; without spine; with setules; as a row; single; continuously; on inner margin, or on outer margin; without spinules. First swimming legs exopod 3 with setule; as a row; single; continuously; on outer surface; with spinules; with seta; unrestricted; 2 on inner surface; 2 on terminal surface; with spine; 2; unequal size; first no longer 2x than origin segment; stout; serrated; on inner side, or on outer side; equally; second longer 3x than origin segment; slender; serrated; on outer side; with ornamentation on non-serrated side; by setules. **Second swimming legs**. Symmetrical; Second swimming legs biramous. Second swimming legs intercoxal plate without seta. Second swimming legs praecoxa present; located laterally. Second swimming legs coxa with seta; straight; distally inserted; on inner surface; surpassing to basal segment; without setules; without spinules; without spine. Second swimming legs basis without seta; without setules; without spinules; without spine. Second swimming legs endopod 3-segmented. Endopod 1 with seta; straight; restricted; one on inner surface; without spine; with setules; as a row; single; continuously; on outer surface; without spinules; absence of Schmeil’s organ. Endopod 2 with seta; straight; unrestricted; two on inner surface; without spine; with setules; as a row; single; continuously; on outer side; without spinules; presence of Schmeil’s organ; on posterior surface. Endopod 3 with seta; straight; unrestricted; three on inner surface; two on outer surface; two on distal surface; without spine; without setules; with spinules; as a row; double; distally inserted; at anterior surface; absence of Schmeil’s organ. Second swimming legs exopod 1 with seta; restricted; one on inner surface; with spine; 1; stout; not reaching to distal-third of the exopod 2; serrated; on inner side, or on outer side; with setules; as a row; single; continuously; on inner side; without spinules; absence of Schmeil’s organ. Exopod 2 with seta; unrestricted; one on inner surface; with spine; 1; stout; not surpassing the exopod 3; serrated; on inner side, or on outer side; with setules; as a row; single; continuously; on inner surface; without spinules; absence of Schmeil’s organ. Exopod 3 with seta; plurimarginal; three on inner surface; two on terminal surface; with spine; 2; unequal size; first no longer 2x than origin segment; stout; serrated; on inner side, or on outer side; equally; second longer 2x than origin segment; slender; serrated; on outer side; with ornamentation on non-serrated side; of setules; setules on outer surface; as a row; single; continuously; on inner surface; with spinules; as a row; single; distally inserted; at anterior surface; absence of Schmeil’s organ. **Third swimming legs**. Symmetrical; Third swimming legs biramous. Third swimming legs intercoxal plate without seta. Third swimming legs praecoxa present; not laterally located. Third swimming legs coxa with seta; straight; distally inserted; on inner surface; surpassing to first endopodal segment; without setules; without spinules; without spine. Third swimming legs basis without seta; without setules; without spinules; without spine. Third swimming legs endopod 3-segmented. Endopod 1 with seta; restricted; one on inner surface; without spine; without setules; without spinules; absence of Schmeil’s organ. Endopod 2 with seta; restricted; two on inner surface; straight; without spine; without setules; without spinules; absence of Schmeil’s organ. Endopod 3 with seta; straight; plurimarginal; two on inner surface; two on outer surface; three on terminal surface; without spine; without setules; with spinules; as a row; distally inserted; double; at anterior surface; absence of Schmeil’s organ. Third swimming legs exopod 1 with seta; restricted; straight; one on inner surface; with spine; 1; stout; not reaching to the distal-third of the exopod 2; serrated; equally; on inner surface, or on outer surface; with setules; as a row; single; continuously; on inner surface; without spinules; absence of Schmeil’s organ. Exopod 2 with seta; straight; restricted; one on inner surface; with spine; 1; stout; not reaching out to exopod 3; serrated; on inner side, or on outer side; equally; with setules; as a row; single; continuously; on inner side; without spinules; absence of Schmeil’s organ. Exopod 3 without setules; with spinules; as a row; single; distally inserted; at anterior surface; with seta; straight; unrestricted; three on inner surface; two on terminal surface; with spine; 2; unequal size; first no longer 2x than origin segment; stout; serrated; on inner side, or on outer side; equally; second longer 2x than origin segment; slender; serrated; on outer side; with ornamentation on non-serrated side; of setules; absence of Schmeil’s organ. **Fourth swimming legs**. Symmetrical; biramous. Intercoxal plate without sensilla. Praecoxa present. Coxa with seta; distally inserted; on inner margin; reaching out to endopod 1; without spinules; setules absent. Basis with seta; one; medially inserted; on posterior surface; smaller than the original segment; without setules; without spinules; without spine. Fourth swimming legs endopod 3-segmented. Endopod 1 with seta; one; restricted; on inner surface; without spine; without setules; without spinules; absence of Schmeil’s organ. Endopod 2 with seta; restricted; two on inner side; without spine; with setules; as a row; single; continuously; on outer surface; without spinules; absence of Schmeil’s organ. Endopod 3 with seta; unrestricted; two on inner surface; two on outer surface; three on distal surface; without spine; without setules; with spinules; as a row; double; distally inserted; at anterior surface; absence of Schmeil’s organ. Fourth swimming legs exopod 1 with seta; restricted; one on inner surface; with spine; 1; stout; not reaching out to distal-third of the exopod 2; serrated; on inner side, or on outer side; equally; with setules; as a row; single; continuously; on inner surface; without spinules; absence of Schmeil’s organ. Exopod 2 with seta; restricted; one on inner surface; with spine; 1; stout; not reaching the end of exopod 3; serrated; on inner side, or on outer side; equally; with setules; as a row; single; continuously; on inner surface; without spinules; absence of Schmeil’s organ. Exopod 3 without setules; with spinules; as a row; single; distally inserted; at anterior surface; with seta; unrestricted; three on inner surface; two on distal surface; with spine; 2; unequal size; first no longer 2x than origin segment; stout; serrated; on inner side, or on outer side; equally; second longer 2x than origin segment; slender; serrated; on outer side; without ornamentation on non-serrated side; absence of Schmeil’s organ.

##### Fifth swimming legs features

Asymmetrical. Fifth swimming leg intercoxal plate with length not equal or greater than width on 1.5x; with irregular proximal margin; discontinuous to; the anterior margin of the left coxa, or the anterior margin of the right coxa; posterior sensilla on the right lateral absent. **Fifth left swimming leg**. Fifth left swimming leg biramous; leg reaching first right exopod segment; proximally. Fifth left swimming leg praecoxa present; rudimentary; separated from the coxae; without ornamentation. Fifth left swimming leg coxa concave inner side; without teeth-like structures; with process; conical; on posterior surface; outer side; distally inserted; not projecting over basis; with sensilla; stout; triangular; at apex; no longer 2x than insertion basis; without swelling; without seta; without spinules. Fifth left swimming leg basis sub-cylindrical; unequal size between inner and outer side; shorter outer than inner side; with concave inner side; rounded internal proximal expansion absent; without outgrowth; with groove; deep; obliquely; on posterior surface; not reaching the endopodal lobe; not ornamented; absence of protuberance; with seta; outerly inserted; no longer 2x than origin segment; presence of minutely granular; as a patch; innerly. Fifth left swimming leg endopod segments 1 and 2 fused; segments 2 and 3 fused; 1-segmented; stout; separated from the basis; ornamented; on inner side; with spinules; more than four elements; as a row; terminally; row of setules absent; without seta. Fifth left swimming leg exopod segments 1 and 2 separated; segments 2 and 3 fused; 2-segmented; stout; separated from the basis. Fifth left swimming leg exopod 1 sub-triangular; longer than broad; equal size between inner and outer side; rectilinear inner side; convex outer side; without swelling; without marginal extension; without process; with lobe; single; circular; medially inserted; on inner side; covered; by setules; without outer spine; absence seta. Fifth left swimming leg exopod 2 digitiform; longer than broad; equal size between inner and outer side; disform inner side; with rectilinear outer side; setulose pad present; not prominently rounded; proximally; on inner side; inflated medial region absent; distal process present; digitiform; non denticulate; without transverse row of denticles; none oblique row of 5 denticles; not innerly directed; with seta; spiniform; ornamented by spinules; not surpassing the distal-point of the segment; without outer spine; terminal claw absent.

##### Fifth right swimming leg

Biramous. Fifth right swimming leg praecoxa present; separated from the coxae; without ornamentation. Fifth right swimming leg coxa convex inner side; without teeth-like structures; with process; rounded; distally inserted; on posterior surface; closest to the outer rim; projecting over basis; beyond the first third; until the medial surface; without triangular protuberance innerly; with sensilla; slender; at apex; no longer 2x than basal insertion; without marginal extension; without seta; without spinules. Fifth right swimming leg basis cylindrical; unequal size between inner and outer side; shorter outer than inner side; rectilinear inner side; tumescence present; not inflated; restricted on inner surface; proximally; with protuberance; single; semicircular; on inner side; proximally; ornamented; with tubercles; as a patch; minutely granular; covering the element; until to adjacent surface; absence of distinct minutely granular; additional inner process absent; with posterior groove; deep; obliquely; reaching the endopodal lobe; ornamented; with tubercles; throughout of the outer border; with seta; outerly inserted; on anterior surface; no longer 2x than origin segment; posterior protrusion present; distal process absent. Fifth right swimming leg with endopodite present; fused to basis; on anterior surface; ancestral segments 1 and 2 fused; ancestral segments 2 and 3 fused; stout; ornamented; with setules; as a row; on inner side; terminally; without seta. Fifth right swimming leg exopod segments 1 and 2 separated; segments 2 and 3 fused; 2-segmented; stout; separated from the basis. Fifth right swimming leg exopod 1 sub-cylindrical; longer than broad; nearly 1.25 times; unequal size between both sides; shorter inner than outer side; convex inner side; rectilinear outer side; with marginal extension; sub-triangular; distally inserted; at outer rim; spinules absent; with process; triangular; arched; internally directed; blunt tip; sclerotized; without ornamentation; distally inserted; at posterior surface; projecting over next segment; without outer spine; without seta; internal prominence absent; lamella on posterior surface absent. Fifth right swimming leg exopod 2 cylindrical; longer than broad; nearly 2 times; equal size between both sides; disform inner side; convex outer side; without posterior proximal swelling; inner-posterior process absent; without marginal expansion; curved ridge on distal posterior surface present; chitinous knobs absent; with outer spine; inserted sub-distally; rectilinear; ornamented innerly; by spinules; as a row; not ornamented outerly; sharp tip; without apparent curve; lesser than the length of the exopod 2; until to 2 times its size; 2x; sensilla absent; terminal claw present; equal or longer 1.5 times than insertion segment; sclerotized; arched; inward; with conspicuous curve; proximally; ornamented innerly; by spinules; as a row; partially on extension; medially, or distally; not ornamented outerly; sharp tip; not curved tip; without medial constriction; hyaline process absent.

##### FEMALE

Body longer and wider than male; Female body 1168 micrometers excluding caudal setae. Widest at first metasome segment. Distal margin of the prosomal segments without one line of setules at posterior margin. Prosome segments without spinules at prosomal segments. Fourth metasome segment absence of dorsal protuberance. Fourth and fifth metasome segments separated. Limit between fourth and fifth metasome segments without ornamentation. **Fifth metasome segment**. Fifth metasome segment with sensilla; dorsally; 2 elements; with epimeral plates. Epimeral plates symmetrical. Right epimeral plates prominent, as projections; one posterior-laterally directed; not reaching half length of the genital segment; with sensilla at the apex; dorsal-posterior sensilla absent; without ornamentation. Left epimeral plate without expansion.

##### Urosome

3-segmented. **Genital double-somite**. Asymmetrical in dorsal view; longer than broad; longer than other urosomites combined; dorsal suture at mid-length absent; not covered by spinules; with swelling; rounded; unequal size; greater left than right; anteriorly; with sensillae; on both sides; one; stout; with robust apex; at left lateral; not on lobular base; medially; one; stout; at right lateral; not on lobular base; anteriorly; with robust apex; of equal size between then; lateral protuberance absent; with right posterior rim expanded; over next segment; without slender sensilla on each posterior rim; without posterior-dorsal process. Genital double-somite opercular pad present; broader than longer; symmetrical; development laterally; expanded posteriorly; covering partially; double gonoporal slit; located ventrally; with arthrodial membrane; inserted anteriorly; post-genital process absent; disto-ventral tumescence absent; ventral vertical folds absent; dorsal sensilla absent. Second urosome segment without ventral fusion to anal segment; right distal process absent. Caudal rami patch of setules on outer surface absent; patch of spinules on outer surface absent.

##### Appendices features

Rostrum basal process absent. **Antennules**. Symmetrical. Right antennule surpassing to genital double-segment; extending beyond caudal rami. Right antennule not exceeding the caudal setae. Right antennule ornamentation pattern equals to male left antennule; mostly. Actual segment 13 without seta; without aesthetasc. Actual segment 14 without seta; without aesthetasc. Actual segment 15 without seta; without aesthetasc. Actual segment 16 without seta; without aesthetasc. Actual segment 17 without seta. Actual segment 18 without seta.

##### Fifth swimming legs

Symmetrical; Fifth swimming legs biramous. Fifth swimming legs intercoxal plate longer than wide; separated from the legs. Fifth swimming legs praecoxa with sclerite praecoxal; separated from the coxae; without ornamentation. Fifth swimming legs coxa with process; conical; at the outer rim; distally; sensilla present; stout; at apex; projecting over basal segment; no longer 2x than basal insertion; marginal extension absent; without swelling; without seta; without spinules. Fifth swimming legs basis sub-triangular; unequal size between inner and outer sides; shorter outer than inner side; with convex inner side; without proximal inner outgrowth; without groove; with distal extension; on posterior surface; with seta; outerly inserted; on anterior surface; longer 2x than origin segment; reaching to exopod 1 distally. Fifth swimming legs endopod segments 1 and 2 fused; segments 2 and 3 fused; 1-segmented; stout; separated from the basis; present discontinuity cuticle; on inner side; with spinules; as a row; single; non-oblique; sub-terminally; at anterior surface; with seta; double; one medially; on posterior surface; rectilinear; one distally; on posterior surface; arched; of unequal size; distal seta longer than medial seta. Fifth swimming legs exopod segments 1 and 2 separated; segments 2 and 3 separated; 3-segmented; separated from the basis. Fifth swimming legs exopod 1 sub-cylindrical; longer than wide; longer or equal than 2 times; with unequal size between inner and outer side; shorter inner than outer side; with convex inner side; with rectilinear outer side; without swelling; without marginal extension; without posterior process; without spine; without seta. Fifth swimming legs exopod 2 sub-cylindrical; longer than broad; longer or equal than 2 times; without swelling; without marginal extension; without process; without lobe; with spine; inserted laterally; rectilinear; without ornamentation; sharp tip; equal size or larger than next segment; without seta. Fifth swimming legs exopod 3 cylindrical; longer than wide; without swelling; without process; without lobe; without spine; with seta; double; inserted terminally; unequal size between them; outer seta smaller than inner; nearly 1 time; outer seta not ornamented by setules; without ornamentation; presence of terminal claw; sclerotized; arched; externally directed; convex inner side; with ornamentation; of denticles; as a row; on surface partially; at medial region; concave outer side; with ornamentation; of denticles; as a row; on surface partially; at medial region; blunt tip; 6 times longer than origin segment.

##### Distribution records

###### BRAZIL

**Piauí**: Paranaguá Lake (Spandl, 1926). **Mato Grosso**: pond in Cuiabá City (Poppe, 1891); Sá Mariana, and Recreio Lagoons (Matsumura-Tundisi, 1986); Pedra Branca Bay (Santos-Silva *et al*., 1999). **Mato Grosso do Sul**: Guaraná Lake, Pousada das Garças, and Paraná River (Lansac-Tôha *et al*., 1997). PARAGUAY. Samples of several localities on Makthlawaiya region, 23°25’S, 58°19W, and Nanahua, 32°30’S, 59°30’W (Lowndes, 1934). ARGENTINA. **Corrientes**: Ibera Lake; Merces (Brehm, 1959). **Missiones**: San Ignacio (Brehm, 1959). **Santa Fé**: Los Espejos, and Madrejón Dom Felipe Lakes (Ringuelet & Martinez de Ferrato, 1967).

##### Habitat

Habitat in freshwaters: lakes, shallow lakes, littoral zones of lakes.

##### Remarks

The organisms bearing the name were summarily described from a lagoon in the city of Mato Grosso, Brazil, a type locality that does not currently exist. Santos-Silva *et al*., (1999) when re-examining the species found that the holotype and paratypes had disappeared and added neotype and additional material from Pedra Branca Bay, Mato Grosso, Brazil in the INPA collection. This material was the basis for redescription of the species, which has some morphological confusions discussed below.

Popper originally described the species as *Diaptomus* (s.l.) *deitersi*. Kiefer (1936) when founding *Notodiaptomus* recombined the species, that also went through other proposals for nomenclatural recombination by Brehm (1959) and Dussart (1985), the first to genus *Neodiaptomus* and second to subgenus *Notodiaptomus* (*Notodiaptomus*) *deitersi*. None of these proposals have been accepted over time, due to non-compliance with the ICZN (1999) and/or fragile evidential bases.

Santos-Silva *et al*. (2015), when addressing the taxon to the *nordestinus* complex (Wright 1935) offered comments on the presence of “pyramidal projection covered by tiny spinules”, “hyaline membrane” on the antepenultimate segment of the male’s right antennule”, “conical seta on actual segment 8 of male A1R as long as the modified setae on segments 10 and 11” and “female with bisegmented endopod”. All these were observations of Lowndes (1934), of which were not corroborated by Brehm (1959) or in the review of *nordestinus*. In contrast, Ringuelet & Martinez de Ferrato (1967), and Matsumura (1986) not only recorded the presence of the “pyramidal projection”, but, for the latter, it was illustrated for the female fourth metasome. However, bisegmentation for the female endopod was not maintained by Matsumura (1986). For Santos-Silva *et al*. (2015) these observations are justified by the lack of morphological details in the description of the species, which resulted in the various inconsistencies in later studies.

For the present study, none of these observations were corroborated. In contrast, all the morphological characteristics indicated for *Notodiaptomus*, in creation (Kiefer, 1936) and magnification (Kiefer, 1956), could be present. The same can be concluded for the abductions performed in the original record of the Wright’s complex (1935), having seen this to be the first taxonomic shelter of the species before its current recombination.

*N. deitersi* was designated type carrying the name *Notodiaptomus* in a nomenclatural act of Santos-Silva *et al*. (1999). There is previous mention that defines the species as a “genotype” of the genus (*sensu* Ringuelet, 1958), a term of common use, not recommended, but recognized by the ICZN (1999) as a type species possibly (recommendation 67A). Apparently Dussart & Defaye (1983) disagreed with this typification when they proposed *N. gibber* as a type species, although they formally did not present evidence that defined a diagnostic morphological set for this. However, although currently the oldest, *N. gibber* is not a nominal species included in the foundation of the genus originally, a fact that invalidates its election as a type species of *Notodiaptomus* as it is contrary to what was established in the ICZN (1999) for type fixation (Art. 67.2.1).

#### Notodiaptomus amazonicus (Wright, 1935)

##### Synonymy

*Diaptomus henseni*; Wright, 1927(nec. Dahl): 73, 75, 96, 100, 102, pl. 8, figs. 7–11. *Diaptomus amazonicus* Wright, 1935: 214, 219, 220, 221, 222, 225, 228, pl. 1, figs. 2, 5, 9, 14, 16; 1936a: 80; 1937: 73, 76; 1938b: 562; Brehm, 1960: 50; Reid, 1991: 737, 738, 740. *Notodiaptomus amazonicus*; Kiefer, 1936a: 197, fig. 6; 1956: 242; Brehm, 1958a: 168; Löffler, 1963: 208; Ringuelet & Martinez de Ferrato, 1967: 411, 414, pl. 1, figs. 7–11; Brandorff, 1972: 4, 5, 10, 18, 25, 38, 43, figs. 29–32; 1973b: 205, 206; 1976: 614, 616, fig. 2; Andrade & Brandorff, 1975: 97; Hardy, 1980: 594, 596, 603, 604; Löffler, 1981: 15; Carvalho,1983: 717; Dussart & Defaye, 1983: 136; Dussart, 1984a: 34, 35, 39, 48, 51, 53, fig. 5A; Robertson & Hardy, 1984: 347, tab. 3; Arcifa, 1984: 143, tab. 7; Dussart & Frutos, 1985: 307; Matsumura-Tundisi, 1986: 537, 547, figs. 22– 25, 100; Montú & Gloeden, 1986: 6, 83, fig. 25k-m; Cicchino *et al*., 1989: 101; Santos-Silva *et al*., 1989: 726, 727, figs. 47–68; Reid & Moreno, 1990: 731; Reid, 1991: 737, 738, 740; Santos-Silva, 1991: 33, 34, fig. 10; 1998: 206; Sendacz & Melo Costa, 1991: 468; Bozelli, 1992: 254, tab. 6; Santos-Silva & Robertson, 1993: 101; Sendacz, 1993: 35; Battistoni, 1995: 958; Rocha *et al*., 1995: 156; Santos-Silva *et al*., 1999: 127; Bohrer & Araújo, 1999: 92, 94; Santos-Silva *et al*., 2015: 41–45, figs. 21–23, identification key to male and female; Perbiche-Neves *et al*., 2015: 63, figs. 53–54; Perbiche-Neves *et al*., 2020: 697-698, key to the Neotropical diaptomid, fig. 21.15 A; Geraldes-Primeiro *et al*. (2017). *Notodiaptomus* (*Notodiaptomus*) *amazonicus*; Dussart, 1985a: 208. *Notodiaptomus* cf. *amazonicus*; Lima *et al*. 1996: 114, 115, fig. 3; Lansac-Tôha *et al*., 1997: 140, 141, Tab. 3.

##### Type locality

Not specified in the original species description. Syntypes located in the USMN, USA, allow us to deduce Lake Arari, Ilha de Marajó, Pará, Brazil as the type locality. Other locations are related to specie by Wright (1935) originally: Tapajós River, and Arama River, Pará, Brazil, and Essequibo River, and Georgetown, both in British Guiana.

##### Type material

Holotype is not specified originally. Examples of the type-series, 1 male, and 1 female, are listed in the National Museum of Natural History at the Smithsonian, USA, under codes USNM 59515 and USNM 59817, respectively.

##### Material examined

Topotype: 3 males, and 2 females from the Calado Lake (final stretch), collected in 22.XI.1983; 2 males, and 2 females from the Amanã Lake (middle stretch), Amazonas River, Brazil, uninformed collector, 07.XI.1979. Each material is stored entire in alcohol in the collection of Plankton Laboratory, INPA, Brazil. 1 male (INPA-COP003, slides a-h) and 1 female (INPA-COP004, slides a-h) were selected to be dissection on eight slides each and deposited in the Zoological Collection of the INPA, Brazil.

##### Diagnosis

**(1)** Male actual segment 13 with modified seta presenting rounded apex; **(2)** male actual segment 20 with spinous process reaching beyond of distal-point segment 21, when present (variable condition); **(3)** male fifth left swimming leg exopod 2 with distal process in acute-form; **(4)** male fifth right swimming leg basis with inner protuberance, single, semicircular, proximally; **(5)** male fifth right swimming leg exopod 2 with 1-2 chitinous knobs posteriorly; **(6)** female caudal rami with presence of patch of setules on outer surface; **(7)** female fifth swimming legs basis with outer seta no longer 2x than origin segment; **(8)** female fifth swimming legs endopod without discontinuity cuticle.

##### Redescription

###### MALE

Body 998 micrometers excluding caudal setae. Male body smaller and slenderer than female. Nerve axons myelinated. Prosome 6-segmented; widest at first metasome segment; without one line of setules at posterior margin; without spinules at segments. Cephalosome anterior margin rounded; with dorsal suture; incomplete; separate from first metasome segment. First metasome segment without sensilla. Second metasome segment without sensilla. Third metasome segment without sensillae; non-ornamented posterior margin. Fourth metasome segment without sensillae; separated from the fifth metasome. Limit between fourth and fifth metasome segments without ornamentation. Fifth metasome segment without sensilla; Fifth metasome segment without ornamentation; Fifth metasome segment without dorsal conical process; with epimeral plates. Epimeral plates symmetrical. Right epimeral plates reduced, as rounded distal corner segment limit; with sensilla; at the apex of projection; without ornamentation.

##### Urosome

5-segmented; Urosome 5 - free segments. Genital somite symmetrical in dorsal view; with single aperture; located on left side; ventrolaterally on posterior rim; with sensillae; on both sides; one; at left lateral; posteriorly; one; at right rim; posteriorly; of equal size between then. Third urosome segment without spinules; without external seta. Fourth urosome segment without spinules; without sub-conical blunt dorsal-lateral process. Anal segment absence of dorsal sensillae; presence of operculum; convex; not covering the anal aperture fully. Caudal rami symmetrical; separated from anal segment; longer than wide; with setules; continuous on; inner side; each ramus bearing 6 caudal setae; 5 marginals; plumose; and 1 internal dorsally; straight; not reticulated main axis; outermost seta with outer spiniform process absent.

##### Appendices features

Rostrum symmetrical; separated from dorsal cephalic shield; by complete suture; sensillae present; one pair; anteriorly inserted on surface tegument; with rostral filament; double; paired; extended; into point; with basal process; in ventral view, rounded on left side; without a smaller basal expansion on the right side.

##### Antennules

Asymmetrical. **Right antennules**. Uniramous; right antennule surpassing to genital segment; right antennule not extending beyond caudal rami.

Right antennule ancestral segment I and II separated. Ancestral segment II and III fused. Ancestral segment III and IV fused. Ancestral segment IV and V separated. Ancestral segment V and VI separated. Ancestral segment VI and VII separated. Ancestral segment VII and VIII separated. Ancestral segment VIII and IX separated. Ancestral segment IX and X separated. Ancestral segment X and XI separated. Ancestral segment XI and XII separated. Ancestral segment XII and XIII separated. Ancestral segment XIII and XIV separated. Ancestral segment XIV and XV separated. Ancestral segment XV and XVI separated. Ancestral segment XVI and XVII separated. Ancestral segment XVII and XVIII separated. Ancestral segment XVIII and XIX separated. Ancestral segment XIX and XX separated. Ancestral segment XX and XXI separated. Ancestral segment XXI and XXII fused. Ancestral segment XXII and XXIII fused. Ancestral segment XXIII and XXIV separated. Ancestral segment XXIV and XXV fused. Ancestral segment XXV and XXVI separated. Ancestral segment XXVI and XXVII separated. Ancestral segment XXVII and XXVIII fused.

Right antennule actual 22-segmented; geniculated; between the segment 18 and segment 19; with swollen and modified region; formed by 5 segments; between 13 and 17 segments. Actual segment 1 with seta; one element; straight; none larger than segment; without spinules; without vestigial seta; without conical seta; without modified seta; without spinous process; with aesthetasc; one element. Actual segment 2 with seta; three elements; of unequal size; straight; none larger than segment; without spinules; with vestigial seta; one element; without conical seta; without modified seta; without spinous process; with aesthetasc; one element. Actual segment 3 with seta; one element; one larger than segment; surpassing to distal margin; beyond three sequential segments; straight; blunt apex; without spinules; with vestigial seta; one element; without conical seta; without modified seta; without spinous process; with aesthetasc. Actual segment 4 with seta; one element; one larger than segment; surpassing to distal margin; straight; not beyond three sequential segments; without spinules; without vestigial seta; without conical seta; without modified seta; without spinous process; without aesthetasc. Actual segment 5 with seta; one element; straight; one larger than segment; surpassing to distal margin; not beyond three sequential segments; without spinules; with vestigial seta; one element; without conical seta; without modified seta; without spinous process; with aesthetasc; one element. Actual segment 6 with seta; one element; none larger than segment; straight; without spinules; without vestigial seta; without conical seta; without modified seta; without spinous process; without aesthetasc. Actual segment 7 with seta; one element; straight; one larger than segment; surpassing to distal margin; beyond three sequential segments; blunt apex; without spinules; without vestigial seta; without conical seta; without modified seta; without spinous process; with aesthetasc; one element. Actual segment 8 with seta; one element; straight; none larger than segment; without spinules; without vestigial seta; with conical seta; one element; not reaching to middle-point of the sequent segment; without modified seta; without spinous process; without aesthetasc. Actual segment 9 with seta; two elements; of unequal size; straight; one larger than segment; surpassing to distal margin; beyond three sequential segments; blunt apex; without spinules; without vestigial seta; without conical seta; without modified seta; without spinous process; with aesthetasc; one element. Actual segment 10 with seta; one element; straight; none larger than segment; without spinules; without vestigial seta; without conical seta; with modified seta; presenting blunt apex; slender form; surpassing to distal margin; beyond of the sequential segment; parallel to antennule direction; without spinous process; without aesthetasc. Actual segment 11 with seta; one element; straight; one larger than segment; surpassing to distal margin; not beyond three sequential segments; without spinules; without vestigial seta; without conical seta; with modified seta; slender form; presenting blunt apex; surpassing to distal margin; beyond of the sequential segment; parallel to antennule direction; shorter length than homologous of actual segment 13; without spinous process; without aesthetasc. Actual segment 12 with seta; one element; straight; one larger than segment; surpassing to distal margin; not beyond three sequential segments; without spinules; without vestigial seta; with conical seta; one element; not smaller than to segment 8; without modified seta; without spinous process; with aesthetasc; one element; absent internal perpendicular fission. Actual segment 13 with seta; one element; straight; one larger than segment; surpassing to distal margin; not beyond three sequential segments; without spinules; without vestigial seta; without conical seta; with modified seta; stout form; surpassing to distal margin; to the middle-point of the sequence segment; parallel to antennule direction; presenting rounded apex; without spinous process; with aesthetasc; one element. Actual segment 14 with seta; two elements; of unequal size; straight; one larger than segment; surpassing to distal margin; beyond three sequential segments; blunt apex; without spinules; without vestigial seta; without conical seta; without modified seta; without spinous process; with aesthetasc; one element. Actual segment 15 with seta; two elements; of unequal size; straight; not bifidform; none larger than segment; without spinules; without vestigial seta; without conical seta; without modified seta; with spinous process; on outer margin; surpassing distal margin; with aesthetasc; one element. Actual segment 16 with seta; two elements; of unequal size; plumose; one larger than segment; surpassing to distal margin; not beyond three sequential segments; not bifidform; without spinules; without vestigial seta; without conical seta; without modified seta; with spinous process; on outer margin; surpassing distal margin; unequal size to process on preceding segment; with aesthetasc; one element. Actual segment 17 with seta; two elements; of unequal size; straight; none larger than segment; bifidform; without spinules; without vestigial seta; without conical seta; with modified seta; one element; stout form; surpassing to distal margin; not beyond of the sequential segment; parallel to antennule direction; without spinous process; without aesthetasc. Actual segment 18 with seta; two elements; of equal size; straight; none larger than segment; without spinules; without vestigial seta; without conical seta; with modified seta; one element; stout form; surpassing distal margin; parallel to antennule direction; without spinous process; without aesthetasc. Actual segment 19 with seta; two elements; of unequal size; plumose; none larger than segment; without spinules; without vestigial seta; without conical seta; with modified seta; two elements; stout form; at least one bifid form; surpassing distal margin; parallel to antennule direction; without spinous process; with aesthetasc; one element. Actual segment 20 with seta; four elements; of unequal size; straight; one larger than segment; surpassing to distal margin; beyond three sequential segments; without spinules; without vestigial seta; without conical seta; without modified seta; with spinous process; distally; reaching beyond of distal-point segment 21; without aesthetasc. Actual segment 21 with seta; two elements; of equal size; plumose; one larger than segment; surpassing to distal margin; greater 3x than original segment; without spinules; without vestigial seta; without conical seta; without modified seta; without spinous process; without aesthetasc. Actual segment 22 with seta; four elements; of equal size; one larger than segment; plumose; surpassing to distal margin; greater 3x than original segment; without spinules; without vestigial seta; without conical seta; without modified seta; without spinous process; with aesthetasc; one element.

##### Left antennules

Uniramous; Left antennule surpassing to prosome; Left antennule not extending beyond caudal rami. Ancestral segment I and II separated. Ancestral segment II and III fused. Ancestral segment III and IV fused. Ancestral segment IV and V separated. Ancestral segment V and VI separated. Ancestral segment VI and VII separated. Ancestral segment VII and VIII separated. Ancestral segment VIII and IX separated. Ancestral segment IX and X separated. Ancestral segment X and XI separated. Ancestral segment XI and XII separated. Ancestral segment XII and XIII separated. Ancestral segment XIII and XIV separated. Ancestral segment XIV and XV separated. Ancestral segment XV and XVI separated. Ancestral segment XVI and XVII separated. Ancestral segment XVII and XVIII separated. Ancestral segment XVIII and XIX separated. Ancestral segment XIX and XX separated. Ancestral segment XX and XXI separated. Ancestral segment XXI and XXII separated. Ancestral segment XXII and XXIII separated. Ancestral segment XXIII and XXIV separated. Ancestral segment XXIV and XXV separated. Ancestral segment XXV and XXVI separated. Ancestral segment XXVI and XXVII separated. Ancestral segment XXVII and XXVIII fused.

Left antennule actual 25-segmented; not-geniculated. Actual segment 1 with seta; one element; none larger than segment; straight; without spinules; without vestigial seta; without conical seta; without modified seta; without spinous process; with aesthetasc; one element. Actual segment 2 with seta; three elements; of equal size; none larger than segment; straight; without spinules; with vestigial seta; one element; without conical seta; without modified seta; without spinous process; with aesthetasc; one element. Actual segment 3 with seta; one element; one larger than segment; straight; surpassing to distal margin; beyond three sequential segments; without spinules; with vestigial seta; one element; without conical seta; without modified seta; without spinous process; with aesthetasc. Actual segment 4 with seta; one element; none larger than segment; straight; without spinules; without vestigial seta; without conical seta; without modified seta; without spinous process; without aesthetasc. Actual segment 5 with seta; one element; one larger than segment; straight; surpassing to distal margin; not beyond three sequential segments; without spinules; with vestigial seta; one element; without conical seta; without modified seta; without spinous process; with aesthetasc; one element. Actual segment 6 with seta; one element; none larger than segment; straight; without spinules; without vestigial seta; without conical seta; without modified seta; without spinous process; without aesthetasc. Actual segment 7 with seta; one element; one larger than segment; straight; surpassing to distal margin; beyond three sequential segments; without spinules; without vestigial seta; without conical seta; without modified seta; without spinous process; with aesthetasc; one element. Actual segment 8 with seta; one element; one larger than segment; straight; surpassing distal margin; without spinules; without vestigial seta; with conical seta; without modified seta; without spinous process; without aesthetasc. Actual segment 9 with seta; two elements; of unequal size; one larger than segment; straight; surpassing to distal margin; beyond three sequential segments; without spinules; without vestigial seta; without conical seta; without modified seta; without spinous process; with aesthetasc; one element. Actual segment 10 with seta; one element; none larger than segment; straight; without spinules; without vestigial seta; without conical seta; without modified seta; without spinous process; without aesthetasc. Actual segment 11 with seta; two elements; one larger than segment; straight; surpassing to distal margin; beyond three sequential segments; without spinules; without vestigial seta; without conical seta; without modified seta; without spinous process; without aesthetasc. Actual segment 12 with seta; one element; one larger than segment; straight; surpassing distal margin; without spinules; without vestigial seta; with conical seta; without modified seta; without spinous process; with aesthetasc; one element. Actual segment 13 with seta; one element; none elongated; straight; surpassing distal margin; without spinules; without vestigial seta; without conical seta; without modified seta; without spinous process; without aesthetasc. Actual segment 14 with seta; one element; elongated; straight; surpassing to distal margin; beyond three sequential segments; without spinules; without vestigial seta; without conical seta; without modified seta; without spinous process; with aesthetasc; one element. Actual segment 15 with seta; one element; larger than segment; straight; surpassing to distal margin; not beyond three sequential segments; without spinules; without vestigial seta; without conical seta; without modified seta; without spinous process; without aesthetasc. Actual segment 16 with seta; one element; larger than segment; plumose; surpassing to distal margin; not beyond three sequential segments; without spinules; without vestigial seta; without conical seta; without modified seta; without spinous process; with aesthetasc; one element. Actual segment 17 with seta; one element; not larger than segment; straight; without spinules; without vestigial seta; without conical seta; without modified seta; without spinous process; without aesthetasc. Actual segment 18 with seta; one element; larger than segment; straight; surpassing to distal margin; beyond three sequential segments; without spinules; without vestigial seta; without conical seta; without modified seta; without spinous process; without aesthetasc. Actual segment 19 with seta; one element; not larger than segment; straight; surpassing distal margin; without spinules; without vestigial seta; without conical seta; without modified seta; without spinous process; with aesthetasc; one element. Actual segment 20 with seta; one element; not larger than segment; straight; surpassing distal margin; without spinules; without vestigial seta; without conical seta; without modified seta; without spinous process; without aesthetasc. Actual segment 21 with seta; one element; larger than segment; plumose; surpassing to distal margin; beyond three sequential segments; without spinules; without vestigial seta; without conical seta; without modified seta; without spinous process; without aesthetasc. Actual segment 22 with seta; two elements; of unequal size; one of them elongated; plumose; surpassing to distal margin; without spinules; without vestigial seta; without conical seta; without modified seta; without spinous process; without aesthetasc. Actual segment 23 with seta; two elements; of unequal size; one larger than segment; plumose; surpassing to distal margin; greater 3x than original segment; without spinules; without vestigial seta; without conical seta; without modified seta; without spinous process; without aesthetasc. Actual segment 24 with seta; two elements; of equal size; one larger than segment; plumose; surpassing to distal margin; greater 3x than original segment; without spinules; without vestigial seta; without conical seta; without modified seta; without spinous process; without aesthetasc. Actual segment 25 with seta; four elements; of equal size; elongated; plumose; surpassing to distal margin; 4 times larger than segment; without spinules; without vestigial seta; without conical seta; without modified seta; without spinous process; with aesthetasc; one element.

##### Antenna

Biramous. Antenna coxa separated from the basis; bearing seta; 1; on inner surface; at distal corner; reaching to the endopod 1. Antenna basis (fusion) separated from the endopodal segment; bearing seta; 2; on inner surface; at distal corner. Endopodal ancestral segment I and II separated. Ancestral segment II and III fused. Ancestral segment III and IV fused. Ancestral segment III and IV fully. Antenna endopod actual 2-segmented. Actual segment 1 not bilobate; with seta; two; on inner margin; with spinules; as a row; obliquely; on outer surface; with pore. Actual segment 2 bilobate; with discontinuity on outer cuticle; not developed as a suture; inner lobe bearing 8 setae; distally; outer lobe bearing 7 setae; distally; with spinules; as a patch; on outer surface. Antenna exopod ancestral segment I and II separated. Ancestral segment II and III fused. Ancestral segment III and IV fused. Ancestral segment IV and V separated. Ancestral segment V and VI separated. Ancestral segment VI and VII separated. Ancestral segment VII and VIII separated. Ancestral segment VIII and IX separated. Ancestral segment IX and X fused. Antenna exopod actual 7-segmented. Actual segment 1 single; elongated (width-length, equal or larger ratio 2:1); with seta; one; at inner surface. Actual segment 2 compound; elongated (larger width-length ratio 2:1); with seta; three; at inner surface. Actual segment 3 single; not elongated (lesser width-length ratio 2:1); with seta; one; at inner surface. Actual segment 4 single; not elongated (lesser width-length ratio 2:1); with seta; one; at inner surface. Actual segment 5 single; not elongated (lesser width-length ratio 2:1); with seta; one; at inner surface. Actual segment 6 single; not elongated (lesser width-length ratio 2:1); with seta; one; at inner surface. Actual segment 7 compound; elongated (larger or equal width-length ratio 2:1); with seta; one; at inner surface; and three; at distal surface.

##### Oral features

**Mandible**. Coxal gnathobase sclerotized; with lobe; prominent; on caudal margin; presence of cutting blade; with tooth-like prominence; two, distinctly; 1 acute; on caudal margin; and 1 triangular; on sub-caudal margin; without acute projection between the prominences; with additional spinules; as a row; on dorsal surface; with seta; 1; dorsally; on apical surface; with spinules; apicalmost. Mandible palps biramous; comprising the basis; with seta; four; differently inserted; first medially; reaching to beyond the endopod 1; second distally; third distally; fourth distally; on inner margin; none with setulose ornamentation. Mandible endopod 2-segmented. Mandible endopod 1 with lobe; bearing seta; four; distally inserted; without spinules. Mandible endopod 2 without lobe; bearing setae; nine elements; distally inserted; with spinules; as a row; double. Mandible exopod 4-segmented. Mandible exopod 1 with seta; one element; distally; on inner margin. Mandible exopod 2 with seta; one element; distally; on inner side. Mandible exopod 3 with seta; one element; distally; on inner side. Mandible exopod 4 with setae; three elements; on terminal region. **Maxillule**. Birramous. Maxillule 3-segmented. Maxillule praecoxa with praecoxal arthrite; bearing spines; fifteen elements; ten marginally; plus, five sub-marginally; with spinules; as a patch; on sub-marginal surface. Maxillule coxa with coxal epipodite; with conspicuous outer lobe; bearing setae; nine elements; with coxal endite; elongated (larger or equal width-length ratio 2:1); bearing setae; four elements. Maxillule basis with basal endite; double; first proximal; elongated (larger width-length ratio 2:1; separated from basis; with setae; four elements; distally inserted; second distal; fused to basis; not elongated (lesser width-length ratio 2:1); with setae; four elements; distally inserted; with setules; as a row; on inner side; basal exite present; with setae; one element; on outer surface. Maxillule endopod 1-segmented. Endopod 1 bilobate; first proximal; with setae; three elements; second distal; with setae; five elements. Maxillule exopod 1-segmented. Exopod 1 with setae; six elements; with setules; as a row; on inner side; spinules absent. **Maxilla**. Uniramous. Maxilla 5-segmented. Maxilla praecoxa fused to coxa; incompletely; distinct externally; with praecoxal endite; double; first elongated endite (larger or equal width length ratio 2:1); proximally inserted; with seta; straight, or plumose; 1 straight; 4 plumose; with spine; single; without spinules; without setule; second elongated endite (larger or equal width length ratio 2:1); distally inserted; with seta; plumose; 3 plumose; without spine; with spinules; as a row; on distal margin; with setule; as a row; on distal margin; absence of outer seta. Maxilla coxa with coxal endite; double; first elongated endite (larger or equal width); proximally inserted; with seta; plumose; 3 plumose; without spine; without spinules; with setules; as a row; on proximal margin; second elongated endite (larger or equal width); distally inserted; with seta; plumose; 3 plumose; without spine; without spinules; with setules; as a row; on proximal margin; absence of outer seta. Maxilla basis with basal endite; single; elongated (larger or equal width-length ratio 2:1); with seta; plumose; 3 plumose; without spinules; absence of outer seta. Maxilla endopod 2-segmented. Endopod 1 with seta; 2 plumose; without spine; without spinules; without setules. Maxilla endopod 2 with seta; 2 plumose; without spine; without spinules; without setules. **Maxilliped**. Uniramous; Maxilliped 8-segmented. Maxilliped praecoxa fused to coxa; incompletely; distinct internally; with praecoxal endite; not elongated (lesser width-length ratio 2:1); distally inserted; with seta; 1 straight; with spinules; as a row; single; on basal surface; without setules. Maxilliped coxa with coxal endite; three coxal endite; first elongated (larger or equal width); proximally inserted; with seta; 2 plumose; with spinules; as a patch; single; on apical surface; without setules; second not elongated (lesser width-length ratio 2:1); medially inserted; with seta; 3 plumose; with spinules; as a row; single; on medial surface; without setules; third elongated (larger or equal width length ratio 2:1); distally inserted; with seta; 3 plumose; none reaching to beyond of the basis; with spinules; as a row; single; on basal surface; without setules; with lobe; prominence; at inner distal angle; ornamented; with spinules; continuously on margin. Maxilliped basis without basal endite; with seta; 3 plumose; with spinules; as a row; single; on medial surface; with setules; as a row; single; on inner margin. Maxilliped endopod segment 6-segmented. Endopod 1 with seta; 2 plumose; on inner surface. Endopod 2 with seta; 3 plumose; on inner surface. Endopod 3 with seta; 2 plumose; on inner surface. Endopod 4 with seta; 2 plumose; on inner surface. Endopod 5 with seta; 2 plumose; on inner surface, or on outer surface; outer seta absent. Endopod 6 with seta; 4 plumose; on inner surface, or on outer surface.

##### Swimming legs features

**First swimming legs.** Symmetrical; biramous. First swimming legs intercoxal plate without seta. First swimming legs praecoxa absent. First swimming legs coxa with seta; one; straight; distally inserted; on inner surface; surpassing to first endopodal segment; with setules; two group; as a patch; on inner margin; and as a row; double; on anterior surface; outerly; without spinules; without spine. First swimming legs basis without seta; with setules; as a patch; single; on outer surface; without spinules; without spine. First swimming legs endopod 2-segmented. Endopod 1 with seta; straight; restricted; to inner surface; one element; without spine; with setules; as a row; single; continuously; on outer surface; without spinules; absence of Schmeil’s organ. Endopod 2 with seta; unrestricted; three on inner surface; one on outer surface; two on distal surface; straight; without spine; with setules; as a row; single; continuously; on outer surface; without spinules; absence of Schmeil’s organ. Endopod 3 absence. First swimming legs exopod 1 with seta; restricted; 1 on inner surface; with spine; 1; stout; smaller than original segment; serrated; on inner side; continuously; with setules; as a row; single; as a row; innerly. First swimming legs exopod 2 with seta; restricted; 1 on inner surface; straight; without spine; with setules; as a row; single; continuously; on inner margin, or on outer margin; without spinules. First swimming legs exopod 3 with setule; as a row; single; continuously; on outer surface; without spinules; with seta; unrestricted; 2 on inner surface; 2 on terminal surface; with spine; 2; unequal size; first no longer 2x than origin segment; stout; serrated; on inner side, or on outer side; equally; second longer 3x than origin segment; slender; serrated; on outer side; with ornamentation on non-serrated side; by setules. **Second swimming legs**. Symmetrical; Second swimming legs biramous. Second swimming legs intercoxal plate without seta. Second swimming legs praecoxa present; located laterally. Second swimming legs coxa with seta; straight; distally inserted; on inner surface; surpassing to basal segment; without setules; without spinules; without spine. Second swimming legs basis without seta; without setules; without spinules; without spine. Second swimming legs endopod 3-segmented. Endopod 1 with seta; straight; restricted; one on inner surface; without spine; with setules; as a row; single; continuously; on outer surface; without spinules; absence of Schmeil’s organ. Endopod 2 with seta; straight; unrestricted; two on inner surface; without spine; with setules; as a row; single; continuously; on outer side; without spinules; presence of Schmeil’s organ; on posterior surface. Endopod 3 with seta; straight; unrestricted; three on inner surface; two on outer surface; two on distal surface; without spine; without setules; with spinules; as a row; double; distally inserted; at anterior surface; absence of Schmeil’s organ. Second swimming legs exopod 1 with seta; restricted; one on inner surface; with spine; 1; stout; not reaching to distal-third of the exopod 2; serrated; on inner side, or on outer side; with setules; as a row; single; continuously; on inner side; without spinules; absence of Schmeil’s organ. Exopod 2 with seta; unrestricted; one on inner surface; with spine; 1; stout; not surpassing the exopod 3; serrated; on inner side, or on outer side; with setules; as a row; single; continuously; on inner surface; without spinules; absence of Schmeil’s organ. Exopod 3 with seta; plurimarginal; three on inner surface; two on terminal surface; with spine; 2; unequal size; first no longer 2x than origin segment; stout; serrated; on inner side, or on outer side; equally; second longer 2x than origin segment; slender; serrated; on outer side; with ornamentation on non-serrated side; of setules; setules on outer surface; as a row; single; continuously; on inner surface; with spinules; as a row; single; distally inserted; at anterior surface; absence of Schmeil’s organ. **Third swimming legs**. Symmetrical; Third swimming legs biramous. Third swimming legs intercoxal plate without seta. Third swimming legs praecoxa present; not laterally located. Third swimming legs coxa with seta; straight; distally inserted; on inner surface; surpassing to first endopodal segment; without setules; without spinules; without spine. Third swimming legs basis without seta; without setules; without spinules; without spine. Third swimming legs endopod 3-segmented. Endopod 1 with seta; restricted; one on inner surface; without spine; without setules; without spinules; absence of Schmeil’s organ. Endopod 2 with seta; restricted; two on inner surface; straight; without spine; without setules; without spinules; absence of Schmeil’s organ. Endopod 3 with seta; straight; plurimarginal; two on inner surface; two on outer surface; three on terminal surface; without spine; without setules; with spinules; as a row; distally inserted; double; at anterior surface; absence of Schmeil’s organ. Third swimming legs exopod 1 with seta; restricted; straight; one on inner surface; with spine; 1; stout; not reaching to the distal-third of the exopod 2; serrated; equally; on inner surface, or on outer surface; with setules; as a row; single; continuously; on inner surface; without spinules; absence of Schmeil’s organ. Exopod 2 with seta; straight; restricted; one on inner surface; with spine; 1; stout; not reaching out to exopod 3; serrated; on inner side, or on outer side; equally; with setules; as a row; single; continuously; on inner side; without spinules; absence of Schmeil’s organ. Exopod 3 without setules; with spinules; as a row; single; distally inserted; at anterior surface; with seta; straight; unrestricted; three on inner surface; two on terminal surface; with spine; 2; unequal size; first no longer 2x than origin segment; stout; serrated; on inner side, or on outer side; equally; second longer 2x than origin segment; slender; serrated; on outer side; with ornamentation on non-serrated side; of setules; absence of Schmeil’s organ. **Fourth swimming legs**. Symmetrical; biramous. Intercoxal plate without sensilla. Praecoxa present. Coxa with seta; distally inserted; on inner margin; reaching out to endopod 1; without spinules; setules absent. Basis with seta; one; medially inserted; on posterior surface; smaller than the original segment; without setules; without spinules; without spine. Fourth swimming legs endopod 3-segmented. Endopod 1 with seta; one; restricted; on inner surface; without spine; without setules; without spinules; absence of Schmeil’s organ. Endopod 2 with seta; restricted; two on inner side; without spine; with setules; as a row; single; continuously; on outer surface; without spinules; absence of Schmeil’s organ. Endopod 3 with seta; unrestricted; two on inner surface; two on outer surface; three on distal surface; without spine; without setules; with spinules; as a row; double; distally inserted; at anterior surface; absence of Schmeil’s organ. Fourth swimming legs exopod 1 with seta; restricted; one on inner surface; with spine; 1; stout; not reaching out to distal-third of the exopod 2; serrated; on inner side, or on outer side; equally; with setules; as a row; single; continuously; on inner surface; without spinules; absence of Schmeil’s organ. Exopod 2 with seta; restricted; one on inner surface; with spine; 1; stout; not reaching the end of exopod 3; serrated; on inner side, or on outer side; equally; with setules; as a row; single; continuously; on inner surface; without spinules; absence of Schmeil’s organ. Exopod 3 without setules; with spinules; as a row; single; distally inserted; at anterior surface; with seta; unrestricted; three on inner surface; two on distal surface; with spine; 2; unequal size; first no longer 2x than origin segment; stout; serrated; on inner side, or on outer side; equally; second longer 2x than origin segment; slender; serrated; on outer side; without ornamentation on non-serrated side; absence of Schmeil’s organ.

##### Fifth swimming legs features

Asymmetrical. Fifth swimming leg intercoxal plate with length not equal or greater than width on 1.5x; with irregular proximal margin; discontinuous to; the anterior margin of the left coxa, or the anterior margin of the right coxa; posterior sensilla on the right lateral absent. **Fifth left swimming leg**. Fifth left swimming leg biramous; leg surpassing first right exopod segment. Fifth left swimming leg praecoxa present; rudimentary; separated from the coxae; without ornamentation. Fifth left swimming leg coxa concave inner side; without teeth-like structures; with process; conical; on posterior surface; outer side; distally inserted; not projecting over basis; with sensilla; stout; triangular; at apex; no longer 2x than insertion basis; without swelling; without seta; without spinules. Fifth left swimming leg basis sub-cylindrical; unequal size between inner and outer side; shorter outer than inner side; with concave inner side; rounded internal proximal expansion absent; without outgrowth; with groove; deep; obliquely; on posterior surface; not reaching the endopodal lobe; not ornamented; absence of protuberance; with seta; outerly inserted; no longer 2x than origin segment; absence minutely granular. Fifth left swimming leg endopod segments 1 and 2 fused; segments 2 and 3 fused; 1-segmented; stout; separated from the basis; ornamented; on inner side; with spinules; more than four elements; as a row; terminally; row of setules absent; without seta. Fifth left swimming leg exopod segments 1 and 2 separated; segments 2 and 3 fused; 2-segmented; stout; separated from the basis. Fifth left swimming leg exopod 1 sub-cylindrical; longer than broad; unequal size between inner and outer side; shorter inner than outer side; concave inner side; convex outer side; without swelling; without marginal extension; without process; with lobe; single; circular; medially inserted; on inner side; covered; by setules; without outer spine; absence seta. Fifth left swimming leg exopod 2 sub-triangular; longer than broad; equal size between inner and outer side; disform inner side; with rectilinear outer side; setulose pad present; prominently rounded; proximally; on inner side; inflated medial region absent; distal process present; acute-form; non denticulate; without transverse row of denticles; none oblique row of 5 denticles; not innerly directed; with seta; spiniform; ornamented by spinules; not surpassing the distal-point of the segment; without outer spine; terminal claw absent.

##### Fifth right swimming leg

Biramous. Fifth right swimming leg praecoxa present; separated from the coxae; without ornamentation. Fifth right swimming leg coxa convex inner side; without teeth-like structures; with process; rounded; distally inserted; on posterior surface; closest to the outer rim; projecting over basis; beyond the first third; until the medial surface; without triangular protuberance innerly; with sensilla; slender; at apex; no longer 2x than basal insertion; without marginal extension; without seta; without spinules. Fifth right swimming leg basis cylindrical; unequal size between inner and outer side; shorter outer than inner side; rectilinear inner side; tumescence present; not inflated; restricted on inner surface; proximally; with protuberance; single; semicircular; on inner side; proximally; ornamented; with tubercles; as a patch; minutely granular; covering the element; not until to adjacent surface; absence of distinct minutely granular; additional inner process absent; with posterior groove; deep; obliquely; not reaching the endopodal lobe; ornamented; with tubercles; throughout of the outer border; with seta; outerly inserted; on anterior surface; no longer 2x than origin segment; posterior protrusion present; distal process absent. Fifth right swimming leg with endopodite present; fused to basis; on anterior surface; ancestral segments 1 and 2 fused; ancestral segments 2 and 3 fused; stout; ornamented; with setules; as a row; on inner side; terminally; without seta. Fifth right swimming leg exopod segments 1 and 2 separated; segments 2 and 3 fused; 2-segmented; stout; separated from the basis. Fifth right swimming leg exopod 1 trapezium; longer than broad; nearly 1.25 times; unequal size between both sides; shorter inner than outer side; convex inner side; rectilinear outer side; with marginal extension; sub-triangular; distally inserted; at outer rim; spinules absent; with process; triangular; arched; internally directed; blunt tip; sclerotized; without ornamentation; distally inserted; at posterior surface; projecting over next segment; without outer spine; without seta; internal prominence absent; lamella on posterior surface absent. Fifth right swimming leg exopod 2 elliptical; longer than broad; nearly 2.5 times; equal size between both sides; disform inner side; convex outer side; without posterior proximal swelling; inner-posterior process absent; without marginal expansion; curved ridge on distal posterior surface present; chitinous knobs present; with 1–2 posteriorly; with outer spine; inserted sub-distally; arched; externally directed; ornamented innerly; by spinules; as a row; not ornamented outerly; sharp tip; without apparent curve; lesser than the length of the exopod 2; until to 2 times its size; 2x; sensilla absent; terminal claw present; equal or longer 1.5 times than insertion segment; sclerotized; arched; inward; with conspicuous curve; medially; ornamented innerly; by spinules; as a row; partially on extension; medially, or distally; not ornamented outerly; sharp tip; not curved tip; without medial constriction; hyaline process absent.

##### FEMALE

Body longer and wider than male; Female body 1471 micrometers excluding caudal setae. Widest at first metasome segment. Distal margin of the prosomal segments without one line of setules at posterior margin. Prosome segments without spinules at prosomal segments. Fourth metasome segment absence of dorsal protuberance. Fourth and fifth metasome segments fused; partially; on dorsal surface. Limit between fourth and fifth metasome segments without ornamentation. **Fifth metasome segment**. Fifth metasome segment without sensilla; with epimeral plates. Epimeral plates asymmetrical. Right epimeral plates prominent, as projections; thinner than the left; one posterior-dorsally directed; not reaching half length of the genital segment; with sensilla at the apex; dorsal-posterior sensilla present; slender; without ornamentation. Left epimeral plate without expansion.

##### Urosome

3-segmented. **Genital double-somite**. Asymmetrical in dorsal view; longer than broad; longer than other urosomites combined; dorsal suture at mid-length absent; not covered by spinules; with swelling; rounded; unequal size; greater left than right; anteriorly; with sensillae; on both sides; one; stout; with robust apex; at left lateral; not on lobular base; medially; one; stout; at right lateral; not on lobular base; anteriorly; with robust apex; of equal size between then; lateral protuberance absent; with right posterior rim expanded; over next segment; without slender sensilla on each posterior rim; without posterior-dorsal process. Genital double-somite opercular pad present; broader than longer; symmetrical; development laterally; expanded posteriorly; covering partially; double gonoporal slit; located ventrally; with arthrodial membrane; inserted anteriorly; post-genital process present; single; disto-ventral tumescence absent; ventral vertical folds absent; dorsal sensilla absent. Second urosome segment without ventral fusion to anal segment; right distal process absent. Caudal rami patch of setules on outer surface present; patch of spinules on outer surface absent.

##### Appendices features

Rostrum basal process absent. **Antennules**. Symmetrical. Right antennule surpassing to genital double-segment; extending beyond caudal rami. Right antennule not exceeding the caudal setae. Right antennule ornamentation pattern equals to male left antennule; fully.

##### Fifth swimming legs

Symmetrical; Fifth swimming legs biramous. Fifth swimming legs intercoxal plate longer than wide; separated from the legs. Fifth swimming legs praecoxa with sclerite praecoxal; separated from the coxae; without ornamentation. Fifth swimming legs coxa with process; conical; at the outer rim; distally; sensilla present; stout; at apex; projecting over basal segment; no longer 2x than basal insertion; marginal extension absent; without swelling; without seta; without spinules. Fifth swimming legs basis sub-triangular; unequal size between inner and outer sides; shorter outer than inner side; with convex inner side; without proximal inner outgrowth; without groove; with distal extension; on posterior surface; with seta; outerly inserted; on anterior surface; no longer 2x than origin segment. Fifth swimming legs endopod segments 1 and 2 fused; segments 2 and 3 fused; 1-segmented; stout; separated from the basis; absent discontinuity cuticle; with spinules; as a row; single; non-oblique; sub-terminally; at anterior surface; with seta; double; one medially; on posterior surface; rectilinear; one distally; on posterior surface; rectilinear; of unequal size; distal seta longer than medial seta. Fifth swimming legs exopod segments 1 and 2 separated; segments 2 and 3 separated; 3-segmented; separated from the basis. Fifth swimming legs exopod 1 sub-cylindrical; longer than wide; longer or equal than 2 times; with unequal size between inner and outer side; shorter inner than outer side; with convex inner side; with rectilinear outer side; without swelling; without marginal extension; without posterior process; without spine; without seta. Fifth swimming legs exopod 2 sub-cylindrical; longer than broad; longer or equal than 2 times; without swelling; without marginal extension; without process; without lobe; with spine; inserted laterally; rectilinear; without ornamentation; sharp tip; equal size or larger than next segment; without seta. Fifth swimming legs exopod 3 cylindrical; longer than wide; without swelling; without process; without lobe; without spine; with seta; double; inserted terminally; unequal size between them; outer seta smaller than inner; nearly 3 times; outer seta not ornamented by setules; without ornamentation; presence of terminal claw; sclerotized; arched; externally directed; convex inner side; with ornamentation; of denticles; as a row; on surface partially; at medial region; concave outer side; with ornamentation; of denticles; as a row; on surface partially; at medial region; blunt tip; 6 times longer than origin segment.

##### Distribution records

###### VENEZUELA

**Monagas**: Orinoco River at the Barrancas (Dussart, 1984a). **Bolívar**: Orinoco River, right margin, in Bolívar City (Dussart, 1984a). GUYANA. Essequibo River, and associated places; Georgetown (Wright, 1927). BRAZIL. **Roraima:** Americanos Lake (Lavrado), Boa Vista (Geraldes-Primeiro *et al*., 2017). **Amazonas**: Janauari Lake, Negro River near to Manaus City (Brandorff, 1972; 1973b); Catalão Lake, Amazonas River/Negro River, near to Manaus (Brandorff, 1972; 1973); paraná of the Curari, Amazonas River (Brandorff, 1972); lake of the Rei, isle of the Careiro, Amazonas River, near to Manaus (Brandorff, 1972; Santos-Silva *et al*., 1989); lake and paraná of the Piranha, Amazonas River; Mata Fome Lake, Madeira River (Brandorff, 1972); Castanho, Jacaretinga, and Redondo Lakes, Amazonas River (Hardy, 1980); Grande Lake, Amazonas River, 03°22’S, 60°35’W (Carvalho, 1983); Jacaretinga Lake, Amazonas River (Matsumura-Tundisi, 1986); Calado Lake, Amazonas River (Santos-Silva *et al*., 1989; Santos-Silva, 1991); Amanã Lake, Japurá River (Santos-Silva & Robertson, 1993); Balbina reservoir, Uatumã River; lake I, isle of the Marchantaria, Amazonas River (this study); Juruazinho Lake, Mamirauá (Santos-Silva *et al*., 2015). **Pará**: Tapajós River, near to Santarém; Arari Lake, isle of Marajó; Arama River (Wright, 1927); Curuá-Una reservoir, 02°48’38“S, 54°18’55“W (Santos-Silva *et al*., 1989); Trombetas River; lago Batata, 01°30’S, 56°20’W; and lake Mussurá, 01°15’S, 56°20W (Bozelli, 1992). **Pernambuco**: BR-232, Km 131 (Brandorff, com. pessoal). **Mato Grosso do Sul**: Guaraná Lake, and Baía River, foodplain of a tributary of the Paraná River (Lima *et al*., 1996); Pato Lake, Baía River, Paraná e Cortado (Lansac-Tôha *et al*., 1997). **Rio Grande do Sul**: lagoon of the Patos (Montú & Gloeden, 1986; Bohrer & Araújo, 1999). ARGENTINA. **Santa Fé**: Madrejón Don Felipe, arroios Colastiné, and Ubajay, Rincón (Ringuelet & Martinez de Ferrato, 1967).

##### Habitat

Habitat in freshwaters: natural lakes, and reservoirs.

##### Remarks

**(1)** *D.* (s.l.) *amazonicus* was originally described from specimens collected in Arari Lake, Marajó Island, Pará, Brazil, possibly. Wright (1927) in reviewing the diaptomids of South America, mistakenly identified the taxon as *D.* (s.l.) *henseni*, resolving the misconception only in 1935 with the inclusion in the *nordestinus complex*. Although this, records indicate this as the type locality of the species (Santos-Silva *et. al.*, 2015), which received the current recombination during the creation of *Notodiaptomus* (Kiefer, 1936).

The close morphological relationship with *N. nordestinus* (Wright, 1935; Santos-Silva *et al*., 2015) was corroborated in the present effort. Outstanding attributes for *nordestinus* complex and *Notodiaptomus* could also be present for taxa, for example, male fifth right swimming leg endopod unisegmented (Wright, 1935), male right antennule with modified seta on segments 10, 11, 13, and spinous process on segments 15, and 16 (Kiefer, 1936). Among the attributes related in the amplification of genus (Kiefer, 1956) it was possible to corroborate the female fifth swimming legs with endopod unisegmented, and male fifth right swimming leg with outer spine on exopod 2 sub-distally.

Although this, significant differences between both species were identified: (1) male fifth right swimming leg with inner protuberance on basis proximally, (2) male fifth right swimming leg basis with groove not reaching the endopodal lobe, (3) female fifth swimming legs with outer seta not reaching to exopod 1 distally, and (4) female fifth swimming legs endopod absent discontinuity cuticle. All attributes present only for *N. amazonicus*.

#### Notodiaptomus anisitsi (Daday, 1905)

##### Synonymy

*Diaptomus anisitsi* Daday, 1905: 149, 151, *Biol. Geral Exper.* 29 152, pl. 9, figs. 16–22; Tollinger, 1911: 65, 270, 271, fig. Y; Pesta, 1927: 80; Wright, 1927: 73, 74, 77, 100, 102, pl. 1, figs. 4–6; 1937: 76; 1938b: 562; 1939: 647; Kiefer, 1928b: 172, figs. 2a-b; Brehm, 1935a: 12, 13; 1935b: 308. *Diaptomus* “*anisitsi*”; Kiefer, 1928b: 172. *Diaptomus inflexus* Brian, 1926: 180, figs. 4–6; Kiefer, 1928b: 170, 172; Brehm, 1958a: 166; 1965: 3, 7; Reid, 1991: 738. *Notodiaptomus anisitsi*; Kiefer, 1936a: 197; 1956: 242; Brehm, 1939: 42, figs. 2–3; 1958c: 575, 576, 578, 579; Ringuelet, 1958a: 45, 50; 1958b: 18; 1962: 87; Pesta, 1959: 148; Ringuelet & Martinez de Ferrato, 1967: 411, 416, 417, pl. 2, figs. 7–10; Brandorff, 1972: 43; 1976: 614, 616, 622, fig. 2; Paggi, 1976b: 85; Löffler, 1981: 15; Dussart & Defaye, 1983: 133, 135, 138; Dussart & Frutos, 1985: 306; Matsumura-Tundisi, 1986: 547, fig. 100; Reid, 1987: 377, tab. 1; 1991: 738; Paggi & José de Paggi, 1990: 690, tab. 2; Sendacz, 1993: 35; Battistoni, 1995: 958; Rocha *et al*., 1995: 156; Lopes *et al*., 1997: 45, 46, tab. 1c; Santos-Silva, 1998: 206; Santos-Silva *et al*., 1999: 127. *Notodiaptomus inflexus*; Brehm, 1938: 29. *“Diaptomus” bidigitatus* Brehm, 1965: 3; Brandorff, 1976: 618, fig. 3; José de Paggi, 1978: 150, 151; 1984: 141; 1985: 17. *Notodiaptomus bidigitatus*; Dussart & Defaye, 1983: 138. *Notodiaptomus anisitsi*; Rocha *et al*., 1995: 155; Santos-Silva *et al*., 2015: 29–36, figs. 15–17, identification key to male and female; Perbiche-Neves *et al*., 2015: 34–37, figs. 25–28, identification key to male and female [*error*]; Perbiche-Neves *et al*., 2020: 696-697, key to the Neotropical diaptomid, fig. 21.15 B. *Notodiaptomus (Notodiaptomus) anisitsi*; Dussart, 1985a: 201, 208.

##### Type locality

Daday (1905) did not clearly specify the type locality of this species. Mentions for Paraguay, specifically for the Permanente lagoon, in Caearapa and Villa Rica are made in the original work.

##### Type material

Not specified in the original species description. Daday’s private collection (“Collection Dadayana”) registered for the Hungarian Natural History Museum in Budapest, mentions the existence of slides of the species probably, used to describe the species (Santos-Silva *et al*. 2015). However, it was not possible to access this mentioned material.

##### Material examined

Non-type material: 2 males, and 1 female, entire in alcohol, from the Reservoir (UHE) Salto Santiago, Paraná. 17.III.1994. Detail ENSS on 9.IX.1997, Collection of the Plankton Laboratory in INPA, Brazil; 1 male (INPA-COP005, slides a-h) and 1 female (INPA-COP006, slides a-h) were selected to be dissection on eight slides each and deposited in the Zoological Collection of the INPA, Brazil. Additional material examined: 1 male, and 1 female (MZUSP 32930), entire in alcohol, from the ponds of Porto Rico city, Paraná State, Brazil, collected by G. Perbiche-Neves, 23.VII.2010, and stored in MZUSP. 2 males, entire in alcohol, from the pond in Barra de Santa Lucia, no date, collected by Thomsen, stored in collection of Kiefer (LNK 1105).

##### Diagnosis

**(1)** male actual segment 13 with modified seta surpassing to distal margin to the middle-point of the sequence segment; **(2)** male actual segment 11 with 2 (two) setae; **(3)** male fifth right swimming leg basis absence inner tumescence; **(4)** male fifth right swimming leg basis absence distal process, anterior or posterior; **(5)** male fifth right swimming leg exopod 1 presenting distal triangular process on posterior surface; **(6)** male fifth right swimming leg exopod 1 presenting distal triangular process on posterior surface not projecting over next segment; **(7)** male fifth right swimming leg exopod 2 presenting 3 (three) chitinous knob posteriorly; **(8)** male fifth right swimming leg exopod 2 with outer spine inserted medially; **(9)** female genital double-somite presenting on each side one rounded swollen with unequal size between them; **(10)** female genital double-somite without right posterior rim expanded; **(11)** female genital double-somite presenting posterior distal process on either side, single on the left and double on the right; **(12)** female genital double-somite with ventral vertical folds; **(13)** female right antennule with length exceeding the caudal setae; **(14)** female fifth swimming legs basis with outer seta longer 2x than origin segment, not reaching to exopod 1 distally; **(15)** female fifth swimming legs endopod 2-segmented.

##### Redescription

###### MALE

Body 1166 micrometers excluding caudal setae. Male body smaller and slenderer than female. Nerve axons myelinated. Prosome 6-segmented; widest at first metasome segment; without one line of setules at posterior margin; with spinules at least at one segment. Cephalosome anterior margin sub-triangular; with dorsal suture; incomplete; separate from first metasome segment. First metasome segment without sensilla. Second metasome segment with sensilla; 2 dorsally; of equal size. Third metasome segment with sensillae; 4 laterally; of unequal size; ornamented posterior margin; with spinules; as a row; double; dorsally, or laterally. Fourth metasome segment with sensillae; 2 dorsally; 4 laterally; of equal size; separated from the fifth metasome. Limit between fourth and fifth metasome segments ornamented; with spinules; as a row; on dorsal doubly; on lateral doubly; same size. Fifth metasome segment with sensilla; 4 laterally; Fifth metasome segment equal size; Fifth metasome segment without ornamentation; Fifth metasome segment without dorsal conical process; with epimeral plates. Epimeral plates symmetrical. Right epimeral plates reduced, as rounded distal corner segment limit; with sensilla; at the apex of projection; without ornamentation.

##### Urosome

5-segmented; Urosome 5 - free segments. Genital somite symmetrical in dorsal view; with single aperture; located on left side; ventrolaterally on posterior rim; with sensillae; on both sides; one; at left lateral; posteriorly; one; at right rim; posteriorly; of equal size between then. Third urosome segment without spinules; without external seta. Fourth urosome segment without spinules; without sub-conical blunt dorsal-lateral process. Anal segment presence of dorsal sensillae; one on each side; medially inserted; presence of operculum; convex; covering the anal aperture fully. Caudal rami symmetrical; separated from anal segment; longer than wide; with setules; continuous on; inner side; each ramus bearing 6 caudal setae; 5 marginals; plumose; and 1 internal dorsally; straight; not reticulated main axis; outermost seta with outer spiniform process absent.

##### Appendices features

Rostrum symmetrical; separated from dorsal cephalic shield; by complete suture; sensillae present; one pair; anteriorly inserted on surface tegument; with rostral filament; double; paired; extended; into point; with basal process; in ventral view, rounded on left side; without a smaller basal expansion on the right side.

##### Antennules

Asymmetrical. **Right antennules**. Uniramous; right antennule surpassing to genital segment; right antennule extending beyond caudal rami.

Right antennule ancestral segment I and II separated. Ancestral segment II and III fused. Ancestral segment III and IV fused. Ancestral segment IV and V separated. Ancestral segment V and VI separated. Ancestral segment VI and VII separated. Ancestral segment VII and VIII separated. Ancestral segment VIII and IX separated. Ancestral segment IX and X separated. Ancestral segment X and XI separated. Ancestral segment XI and XII separated. Ancestral segment XII and XIII separated. Ancestral segment XIII and XIV separated. Ancestral segment XIV and XV separated. Ancestral segment XV and XVI separated. Ancestral segment XVI and XVII separated. Ancestral segment XVII and XVIII separated. Ancestral segment XVIII and XIX separated. Ancestral segment XIX and XX separated. Ancestral segment XX and XXI separated. Ancestral segment XXI and XXII fused. Ancestral segment XXII and XXIII fused. Ancestral segment XXIII and XXIV separated. Ancestral segment XXIV and XXV fused. Ancestral segment XXV and XXVI separated. Ancestral segment XXVI and XXVII separated. Ancestral segment XXVII and XXVIII fused.

Right antennule actual 22-segmented; geniculated; between the segment 18 and segment 19; with swollen and modified region; formed by 5 segments; between 13 and 17 segments. Actual segment 1 with seta; one element; straight; none larger than segment; without spinules; without vestigial seta; without conical seta; without modified seta; without spinous process; with aesthetasc; one element. Actual segment 2 with seta; three elements; of unequal size; straight; none larger than segment; without spinules; with vestigial seta; one element; without conical seta; without modified seta; without spinous process; with aesthetasc; one element. Actual segment 3 with seta; one element; one larger than segment; surpassing to distal margin; beyond three sequential segments; straight; blunt apex; without spinules; with vestigial seta; one element; without conical seta; without modified seta; without spinous process; with aesthetasc. Actual segment 4 with seta; one element; one larger than segment; surpassing to distal margin; straight; not beyond three sequential segments; without spinules; without vestigial seta; without conical seta; without modified seta; without spinous process; without aesthetasc. Actual segment 5 with seta; one element; straight; one larger than segment; surpassing to distal margin; not beyond three sequential segments; without spinules; with vestigial seta; one element; without conical seta; without modified seta; without spinous process; with aesthetasc; one element. Actual segment 6 with seta; one element; none larger than segment; straight; without spinules; without vestigial seta; without conical seta; without modified seta; without spinous process; without aesthetasc. Actual segment 7 with seta; one element; straight; one larger than segment; surpassing to distal margin; beyond three sequential segments; blunt apex; without spinules; without vestigial seta; without conical seta; without modified seta; without spinous process; with aesthetasc; one element. Actual segment 8 with seta; one element; straight; none larger than segment; without spinules; without vestigial seta; with conical seta; one element; reaching to middle-point of the sequent segment; without modified seta; without spinous process; without aesthetasc. Actual segment 9 with seta; two elements; of unequal size; straight; one larger than segment; surpassing to distal margin; beyond three sequential segments; blunt apex; without spinules; without vestigial seta; without conical seta; without modified seta; without spinous process; with aesthetasc; one element. Actual segment 10 with seta; one element; straight; none larger than segment; without spinules; without vestigial seta; without conical seta; with modified seta; presenting blunt apex; slender form; surpassing to distal margin; beyond of the sequential segment; parallel to antennule direction; without spinous process; without aesthetasc. Actual segment 11 with seta; one element; straight; one larger than segment; surpassing to distal margin; not beyond three sequential segments; without spinules; without vestigial seta; without conical seta; with modified seta; slender form; presenting blunt apex; surpassing to distal margin; beyond of the sequential segment; parallel to antennule direction; shorter length than homologous of actual segment 13; without spinous process; without aesthetasc. Actual segment 12 with seta; one element; straight; one larger than segment; surpassing to distal margin; not beyond three sequential segments; without spinules; without vestigial seta; with conical seta; one element; smaller than to segment 8; without modified seta; without spinous process; with aesthetasc; one element; absent internal perpendicular fission. Actual segment 13 with seta; one element; straight; one larger than segment; surpassing to distal margin; not beyond three sequential segments; without spinules; without vestigial seta; without conical seta; with modified seta; stout form; surpassing to distal margin; to the middle-point of the sequence segment; parallel to antennule direction; presenting bifid apex; without spinous process; with aesthetasc; one element. Actual segment 14 with seta; two elements; of unequal size; straight; one larger than segment; surpassing to distal margin; beyond three sequential segments; blunt apex; without spinules; without vestigial seta; without conical seta; without modified seta; without spinous process; with aesthetasc; one element. Actual segment 15 with seta; two elements; of unequal size; straight; not bifidform; none larger than segment; without spinules; without vestigial seta; without conical seta; without modified seta; with spinous process; on outer margin; surpassing distal margin; with aesthetasc; one element. Actual segment 16 with seta; two elements; of unequal size; plumose; one larger than segment; surpassing to distal margin; not beyond three sequential segments; not bifidform; without spinules; without vestigial seta; without conical seta; without modified seta; with spinous process; on outer margin; surpassing distal margin; unequal size to process on preceding segment; with aesthetasc; one element. Actual segment 17 with seta; two elements; of unequal size; straight; none larger than segment; bifidform; without spinules; without vestigial seta; without conical seta; with modified seta; one element; stout form; surpassing to distal margin; not beyond of the sequential segment; parallel to antennule direction; without spinous process; without aesthetasc. Actual segment 18 with seta; two elements; of equal size; straight; none larger than segment; without spinules; without vestigial seta; without conical seta; with modified seta; one element; stout form; surpassing distal margin; parallel to antennule direction; without spinous process; without aesthetasc. Actual segment 19 with seta; two elements; of unequal size; plumose; none larger than segment; without spinules; without vestigial seta; without conical seta; with modified seta; two elements; stout form; at least one bifid form; surpassing distal margin; parallel to antennule direction; without spinous process; with aesthetasc; one element. Actual segment 20 with seta; four elements; of unequal size; straight; one larger than segment; surpassing to distal margin; beyond three sequential segments; without spinules; without vestigial seta; without conical seta; without modified seta; without spinous process; without aesthetasc. Actual segment 21 with seta; two elements; of equal size; plumose; one larger than segment; surpassing to distal margin; greater 3x than original segment; without spinules; without vestigial seta; without conical seta; without modified seta; without spinous process; without aesthetasc. Actual segment 22 with seta; four elements; of equal size; one larger than segment; plumose; surpassing to distal margin; greater 3x than original segment; without spinules; without vestigial seta; without conical seta; without modified seta; without spinous process; with aesthetasc; one element.

##### Left antennules

Uniramous; Left antennule surpassing to prosome; Left antennule extending beyond caudal rami. Ancestral segment I and II separated. Ancestral segment II and III fused. Ancestral segment III and IV fused. Ancestral segment IV and V separated. Ancestral segment V and VI separated. Ancestral segment VI and VII separated. Ancestral segment VII and VIII separated. Ancestral segment VIII and IX separated. Ancestral segment IX and X separated. Ancestral segment X and XI separated. Ancestral segment XI and XII separated. Ancestral segment XII and XIII separated. Ancestral segment XIII and XIV separated. Ancestral segment XIV and XV separated. Ancestral segment XV and XVI separated. Ancestral segment XVI and XVII separated. Ancestral segment XVII and XVIII separated. Ancestral segment XVIII and XIX separated. Ancestral segment XIX and XX separated. Ancestral segment XX and XXI separated. Ancestral segment XXI and XXII separated. Ancestral segment XXII and XXIII separated. Ancestral segment XXIII and XXIV separated. Ancestral segment XXIV and XXV separated. Ancestral segment XXV and XXVI separated. Ancestral segment XXVI and XXVII separated. Ancestral segment XXVII and XXVIII fused.

Left antennule actual 25-segmented; not-geniculated. Actual segment 1 with seta; one element; none larger than segment; straight; without spinules; without vestigial seta; without conical seta; without modified seta; without spinous process; with aesthetasc; one element. Actual segment 2 with seta; three elements; of equal size; none larger than segment; straight; without spinules; with vestigial seta; one element; without conical seta; without modified seta; without spinous process; with aesthetasc; one element. Actual segment 3 with seta; one element; one larger than segment; straight; surpassing to distal margin; beyond three sequential segments; without spinules; with vestigial seta; one element; without conical seta; without modified seta; without spinous process; with aesthetasc. Actual segment 4 with seta; one element; none larger than segment; straight; without spinules; without vestigial seta; without conical seta; without modified seta; without spinous process; without aesthetasc. Actual segment 5 with seta; one element; one larger than segment; straight; surpassing to distal margin; not beyond three sequential segments; without spinules; with vestigial seta; one element; without conical seta; without modified seta; without spinous process; with aesthetasc; one element. Actual segment 6 with seta; one element; none larger than segment; straight; without spinules; without vestigial seta; without conical seta; without modified seta; without spinous process; without aesthetasc. Actual segment 7 with seta; one element; one larger than segment; straight; surpassing to distal margin; beyond three sequential segments; without spinules; without vestigial seta; without conical seta; without modified seta; without spinous process; with aesthetasc; one element. Actual segment 8 with seta; one element; one larger than segment; straight; surpassing distal margin; without spinules; without vestigial seta; with conical seta; without modified seta; without spinous process; without aesthetasc. Actual segment 9 with seta; two elements; of unequal size; one larger than segment; straight; surpassing to distal margin; beyond three sequential segments; without spinules; without vestigial seta; without conical seta; without modified seta; without spinous process; with aesthetasc; one element. Actual segment 10 with seta; one element; none larger than segment; straight; without spinules; without vestigial seta; without conical seta; without modified seta; without spinous process; without aesthetasc. Actual segment 11 with seta; two elements; of unequal size; one larger than segment; straight; surpassing to distal margin; beyond three sequential segments; without spinules; without vestigial seta; without conical seta; without modified seta; without spinous process; without aesthetasc. Actual segment 12 with seta; one element; one larger than segment; straight; surpassing distal margin; without spinules; without vestigial seta; with conical seta; without modified seta; without spinous process; with aesthetasc; one element. Actual segment 13 with seta; one element; none elongated; straight; surpassing distal margin; without spinules; without vestigial seta; without conical seta; without modified seta; without spinous process; without aesthetasc. Actual segment 14 with seta; one element; elongated; straight; surpassing to distal margin; beyond three sequential segments; without spinules; without vestigial seta; without conical seta; without modified seta; without spinous process; with aesthetasc; one element. Actual segment 15 with seta; one element; larger than segment; straight; surpassing to distal margin; not beyond three sequential segments; without spinules; without vestigial seta; without conical seta; without modified seta; without spinous process; without aesthetasc. Actual segment 16 with seta; one element; larger than segment; plumose; surpassing to distal margin; not beyond three sequential segments; without spinules; without vestigial seta; without conical seta; without modified seta; without spinous process; with aesthetasc; one element. Actual segment 17 with seta; one element; not larger than segment; straight; without spinules; without vestigial seta; without conical seta; without modified seta; without spinous process; without aesthetasc. Actual segment 18 with seta; one element; larger than segment; straight; surpassing to distal margin; beyond three sequential segments; without spinules; without vestigial seta; without conical seta; without modified seta; without spinous process; without aesthetasc. Actual segment 19 with seta; one element; not larger than segment; straight; surpassing distal margin; without spinules; without vestigial seta; without conical seta; without modified seta; without spinous process; with aesthetasc; one element. Actual segment 20 with seta; one element; not larger than segment; straight; surpassing distal margin; without spinules; without vestigial seta; without conical seta; without modified seta; without spinous process; without aesthetasc. Actual segment 21 with seta; one element; larger than segment; plumose; surpassing to distal margin; beyond three sequential segments; without spinules; without vestigial seta; without conical seta; without modified seta; without spinous process; without aesthetasc. Actual segment 22 with seta; two elements; of unequal size; one of them elongated; plumose; surpassing to distal margin; without spinules; without vestigial seta; without conical seta; without modified seta; without spinous process; without aesthetasc. Actual segment 23 with seta; two elements; of unequal size; one larger than segment; plumose; surpassing to distal margin; greater 3x than original segment; without spinules; without vestigial seta; without conical seta; without modified seta; without spinous process; without aesthetasc. Actual segment 24 with seta; two elements; of equal size; one larger than segment; plumose; surpassing to distal margin; greater 3x than original segment; without spinules; without vestigial seta; without conical seta; without modified seta; without spinous process; without aesthetasc. Actual segment 25 with seta; five elements; of equal size; elongated; plumose; surpassing to distal margin; 4 times larger than segment; without spinules; without vestigial seta; without conical seta; without modified seta; without spinous process; with aesthetasc; one element.

##### Antenna

Biramous. Antenna coxa separated from the basis; bearing seta; 1; on inner surface; at distal corner; reaching to the endopod 1. Antenna basis (fusion) separated from the endopodal segment; bearing seta; 2; on inner surface; at distal corner. Endopodal ancestral segment I and II separated. Ancestral segment II and III fused. Ancestral segment III and IV fused. Ancestral segment III and IV fully. Antenna endopod actual 2-segmented. Actual segment 1 not bilobate; with seta; two; on inner margin; with spinules; as a row; obliquely; on outer surface; with pore. Actual segment 2 bilobate; without discontinuity on outer cuticle; inner lobe bearing 8 setae; distally; outer lobe bearing 7 setae; distally; with spinules; as a patch; on outer surface. Antenna exopod ancestral segment I and II separated. Ancestral segment II and III fused. Ancestral segment III and IV fused. Ancestral segment IV and V separated. Ancestral segment V and VI separated. Ancestral segment VI and VII separated. Ancestral segment VII and VIII separated. Ancestral segment VIII and IX separated. Ancestral segment IX and X fused. Antenna exopod actual 7-segmented. Actual segment 1 single; elongated (width-length, equal or larger ratio 2:1); with seta; one; at inner surface. Actual segment 2 compound; elongated (larger width-length ratio 2:1); with seta; three; at inner surface. Actual segment 3 single; not elongated (lesser width-length ratio 2:1); with seta; one; at inner surface. Actual segment 4 single; not elongated (lesser width-length ratio 2:1); with seta; one; at inner surface. Actual segment 5 single; not elongated (lesser width-length ratio 2:1); with seta; one; at inner surface. Actual segment 6 single; not elongated (lesser width-length ratio 2:1); with seta; one; at inner surface. Actual segment 7 compound; elongated (larger or equal width-length ratio 2:1); with seta; one; at inner surface; and three; at distal surface.

##### Oral features

**Mandible**. Coxal gnathobase sclerotized; with lobe; prominent; on caudal margin; presence of cutting blade; with tooth-like prominence; two, distinctly; 1 acute; on caudal margin; and 1 triangular; on sub-caudal margin; without acute projection between the prominences; with additional spinules; as a row; on dorsal surface; with seta; 1; dorsally; on apical surface; with spinules; apicalmost. Mandible palps biramous; comprising the basis; with seta; four; differently inserted; first medially; reaching to beyond the endopod 1; second distally; third distally; fourth distally; on inner margin; none with setulose ornamentation. Mandible endopod 2-segmented. Mandible endopod 1 with lobe; bearing seta; four; distally inserted; without spinules. Mandible endopod 2 without lobe; bearing setae; nine elements; distally inserted; with spinules; as a row; double. Mandible exopod 4-segmented. Mandible exopod 1 with seta; one element; distally; on inner margin. Mandible exopod 2 with seta; one element; distally; on inner side. Mandible exopod 3 with seta; one element; distally; on inner side. Mandible exopod 4 with setae; three elements; on terminal region. **Maxillule**. Birramous. Maxillule 3-segmented. Maxillule praecoxa with praecoxal arthrite; bearing spines; fifteen elements; ten marginally; plus, five sub-marginally; with spinules; as a patch; on sub-marginal surface. Maxillule coxa with coxal epipodite; with conspicuous outer lobe; bearing setae; nine elements; with coxal endite; elongated (larger or equal width-length ratio 2:1); bearing setae; four elements. Maxillule basis with basal endite; double; first proximal; elongated (larger width-length ratio 2:1; separated from basis; with setae; four elements; distally inserted; second distal; fused to basis; not elongated (lesser width-length ratio 2:1); with setae; four elements; distally inserted; with setules; as a row; on inner side; basal exite present; with setae; one element; on outer surface. Maxillule endopod 1-segmented. Endopod 1 bilobate; first proximal; with setae; three elements; second distal; with setae; five elements. Maxillule exopod 1-segmented. Exopod 1 with setae; six elements; with setules; as a row; on inner side; spinules absent. **Maxilla**. Uniramous. Maxilla 5-segmented. Maxilla praecoxa fused to coxa; incompletely; distinct externally; with praecoxal endite; double; first elongated endite (larger or equal width length ratio 2:1); proximally inserted; with seta; straight, or plumose; 1 straight; 4 plumose; with spine; single; without spinules; without setule; second elongated endite (larger or equal width length ratio 2:1); distally inserted; with seta; plumose; 3 plumose; without spine; with spinules; as a row; on distal margin; with setule; as a row; on distal margin; absence of outer seta. Maxilla coxa with coxal endite; double; first elongated endite (larger or equal width); proximally inserted; with seta; plumose; 3 plumose; without spine; without spinules; with setules; as a row; on proximal margin; second elongated endite (larger or equal width); distally inserted; with seta; plumose; 3 plumose; without spine; without spinules; with setules; as a row; on proximal margin; absence of outer seta. Maxilla basis with basal endite; single; elongated (larger or equal width-length ratio 2:1); with seta; plumose; 3 plumose; without spinules; absence of outer seta. Maxilla endopod 2-segmented. Endopod 1 with seta; 2 plumose; without spine; without spinules; without setules. Maxilla endopod 2 with seta; 2 plumose; without spine; without spinules; without setules. **Maxilliped**. Uniramous; Maxilliped 8-segmented. Maxilliped praecoxa fused to coxa; incompletely; distinct internally; with praecoxal endite; not elongated (lesser width-length ratio 2:1); distally inserted; with seta; 1 straight; with spinules; as a row; single; on basal surface; without setules. Maxilliped coxa with coxal endite; three coxal endite; first elongated (larger or equal width); proximally inserted; with seta; 2 plumose; with spinules; as a patch; single; on apical surface; without setules; second not elongated (lesser width-length ratio 2:1); medially inserted; with seta; 3 plumose; with spinules; as a row; single; on medial surface; without setules; third elongated (larger or equal width length ratio 2:1); distally inserted; with seta; 3 plumose; none reaching to beyond of the basis; with spinules; as a row; single; on basal surface; without setules; with lobe; prominence; at inner distal angle; ornamented; with spinules; continuously on margin. Maxilliped basis without basal endite; with seta; 3 plumose; with spinules; as a row; single; on medial surface; with setules; as a row; single; on inner margin. Maxilliped endopod segment 6-segmented. Endopod 1 with seta; 2 plumose; on inner surface. Endopod 2 with seta; 3 plumose; on inner surface. Endopod 3 with seta; 2 plumose; on inner surface. Endopod 4 with seta; 2 plumose; on inner surface. Endopod 5 with seta; 2 plumose; on inner surface, or on outer surface; outer seta absent. Endopod 6 with seta; 4 plumose; on inner surface, or on outer surface.

##### Swimming legs features

**First swimming legs.** Symmetrical; biramous. First swimming legs intercoxal plate without seta. First swimming legs praecoxa absent. First swimming legs coxa with seta; one; straight; distally inserted; on inner surface; surpassing to basal segment; with setules; one group; as a patch; on inner margin; without spinules; without spine. First swimming legs basis without seta; with setules; as a patch; single; on outer surface; without spinules; without spine. First swimming legs endopod 2-segmented. Endopod 1 with seta; straight; restricted; to inner surface; one element; without spine; with setules; as a row; single; continuously; on outer surface; without spinules; absence of Schmeil’s organ. Endopod 2 with seta; unrestricted; three on inner surface; one on outer surface; two on distal surface; straight; without spine; with setules; as a row; single; continuously; on outer surface; without spinules; absence of Schmeil’s organ. Endopod 3 absence. First swimming legs exopod 1 with seta; restricted; 1 on inner surface; with spine; 1; stout; smaller than original segment; serrated; on inner side; continuously; without setules. First swimming legs exopod 2 with seta; restricted; 1 on inner surface; straight; without spine; with setules; as a row; single; continuously; on inner margin, or on outer margin; without spinules. First swimming legs exopod 3 with setule; as a row; single; continuously; on outer surface; without spinules; with seta; unrestricted; 2 on inner surface; 2 on terminal surface; with spine; 2; unequal size; first no longer 2x than origin segment; stout; serrated; on inner side, or on outer side; equally; second longer 3x than origin segment; slender; serrated; on outer side; with ornamentation on non-serrated side; by setules. **Second swimming legs**. Symmetrical; Second swimming legs biramous. Second swimming legs intercoxal plate without seta. Second swimming legs praecoxa present; located laterally. Second swimming legs coxa with seta; straight; distally inserted; on inner surface; surpassing to basal segment; without setules; without spinules; without spine. Second swimming legs basis without seta; without setules; without spinules; without spine. Second swimming legs endopod 3-segmented. Endopod 1 with seta; straight; restricted; one on inner surface; without spine; with setules; as a row; single; continuously; on outer surface; without spinules; absence of Schmeil’s organ. Endopod 2 with seta; straight; unrestricted; two on inner surface; without spine; with setules; as a row; single; continuously; on outer side; without spinules; presence of Schmeil’s organ; on posterior surface. Endopod 3 with seta; plumose; unrestricted; three on inner surface; two on outer surface; two on distal surface; without spine; without setules; with spinules; as a row; double; distally inserted; at anterior surface; absence of Schmeil’s organ. Second swimming legs exopod 1 with seta; restricted; one on inner surface; with spine; 1; stout; not reaching to distal-third of the exopod 2; serrated; on inner side, or on outer side; with setules; as a row; single; continuously; on inner side; without spinules; absence of Schmeil’s organ. Exopod 2 with seta; unrestricted; one on inner surface; with spine; 1; stout; not surpassing the exopod 3; serrated; on inner side, or on outer side; with setules; as a row; single; continuously; on inner surface; without spinules; absence of Schmeil’s organ. Exopod 3 with seta; plurimarginal; three on inner surface; two on terminal surface; with spine; 2; unequal size; first no longer 2x than origin segment; stout; serrated; on inner side, or on outer side; equally; second longer 2x than origin segment; slender; serrated; on outer side; with ornamentation on non-serrated side; of setules; setules on outer surface; as a row; single; continuously; on inner surface; with spinules; as a row; single; distally inserted; at anterior surface; absence of Schmeil’s organ. **Third swimming legs**. Symmetrical; Third swimming legs biramous. Third swimming legs intercoxal plate without seta. Third swimming legs praecoxa present; not laterally located. Third swimming legs coxa with seta; straight; distally inserted; on inner surface; surpassing to basal segment; without setules; without spinules; without spine. Third swimming legs basis without seta; without setules; without spinules; without spine. Third swimming legs endopod 3-segmented. Endopod 1 with seta; restricted; one on inner surface; without spine; without setules; without spinules; absence of Schmeil’s organ. Endopod 2 with seta; restricted; two on inner surface; straight; without spine; without setules; without spinules; absence of Schmeil’s organ. Endopod 3 with seta; straight; plurimarginal; two on inner surface; two on outer surface; three on terminal surface; without spine; without setules; with spinules; as a row; distally inserted; double; at anterior surface; absence of Schmeil’s organ. Third swimming legs exopod 1 with seta; restricted; straight; one on inner surface; with spine; 1; stout; not reaching to the distal-third of the exopod 2; serrated; equally; on inner surface, or on outer surface; with setules; as a row; single; continuously; on inner surface; without spinules; absence of Schmeil’s organ. Exopod 2 with seta; straight; restricted; one on inner surface; with spine; 1; stout; not reaching out to exopod 3; serrated; on inner side, or on outer side; equally; with setules; as a row; single; continuously; on inner side; without spinules; absence of Schmeil’s organ. Exopod 3 without setules; with spinules; as a row; single; distally inserted; at anterior surface; with seta; straight; unrestricted; three on inner surface; two on terminal surface; with spine; 2; unequal size; first no longer 2x than origin segment; stout; serrated; on inner side, or on outer side; equally; second longer 2x than origin segment; slender; serrated; on outer side; with ornamentation on non-serrated side; of setules; absence of Schmeil’s organ. **Fourth swimming legs**. Symmetrical; biramous. Intercoxal plate without sensilla. Praecoxa present. Coxa with seta; distally inserted; on inner margin; reaching out to endopod 1; without spinules; setules absent. Basis with seta; one; medially inserted; on posterior surface; smaller than the original segment; without setules; without spinules; without spine. Fourth swimming legs endopod 3-segmented. Endopod 1 with seta; one; restricted; on inner surface; without spine; without setules; without spinules; absence of Schmeil’s organ. Endopod 2 with seta; restricted; two on inner side; without spine; with setules; as a row; single; continuously; on outer surface; without spinules; absence of Schmeil’s organ. Endopod 3 with seta; unrestricted; two on inner surface; two on outer surface; three on distal surface; without spine; without setules; with spinules; as a row; double; distally inserted; at anterior surface; absence of Schmeil’s organ. Fourth swimming legs exopod 1 with seta; restricted; one on inner surface; with spine; 1; stout; not reaching out to distal-third of the exopod 2; serrated; on inner side, or on outer side; equally; with setules; as a row; single; continuously; on inner surface; without spinules; absence of Schmeil’s organ. Exopod 2 with seta; restricted; one on inner surface; with spine; 1; stout; not reaching the end of exopod 3; serrated; on inner side, or on outer side; equally; with setules; as a row; single; continuously; on inner surface; without spinules; absence of Schmeil’s organ. Exopod 3 without setules; with spinules; as a row; single; distally inserted; at anterior surface; with seta; unrestricted; three on inner surface; two on distal surface; with spine; 2; unequal size; first no longer 2x than origin segment; stout; serrated; on inner side, or on outer side; equally; second longer 2x than origin segment; slender; serrated; on outer side; without ornamentation on non-serrated side; absence of Schmeil’s organ.

##### Fifth swimming legs features

Asymmetrical. Fifth swimming leg intercoxal plate with length not equal or greater than width on 1.5x; with irregular proximal margin; discontinuous to; the anterior margin of the left coxa, or the anterior margin of the right coxa; posterior sensilla on the right lateral absent. **Fifth left swimming leg**. Fifth left swimming leg biramous; leg reaching first right exopod segment; proximally. Fifth left swimming leg praecoxa present; rudimentary; separated from the coxae; without ornamentation. Fifth left swimming leg coxa concave inner side; without teeth-like structures; with process; conical; on posterior surface; outer side; distally inserted; not projecting over basis; with sensilla; stout; triangular; at apex; no longer 2x than insertion basis; without swelling; without seta; without spinules. Fifth left swimming leg basis sub-cylindrical; unequal size between inner and outer side; shorter outer than inner side; with concave inner side; rounded internal proximal expansion absent; without outgrowth; with groove; deep; obliquely; on posterior surface; not reaching the endopodal lobe; not ornamented; absence of protuberance; with seta; outerly inserted; no longer 2x than origin segment; absence of minutely granular. Fifth left swimming leg endopod segments 1 and 2 fused; segments 2 and 3 fused; 1-segmented; stout; separated from the basis; ornamented; on inner side; with spinules; more than four elements; as a row; terminally; row of setules absent; without seta. Fifth left swimming leg exopod segments 1 and 2 separated; segments 2 and 3 fused; 2-segmented; stout; separated from the basis. Fifth left swimming leg exopod 1 sub-triangular; longer than broad; equal size between inner and outer side; rectilinear inner side; convex outer side; without swelling; without marginal extension; without process; with lobe; single; semicircular; medially inserted; on inner side; covered; by setules; without outer spine; absence seta. Fifth left swimming leg exopod 2 digitiform; longer than broad; equal size between inner and outer side; disform inner side; with rectilinear outer side; setulose pad present; prominently rounded; proximally; on inner side; inflated medial region present; setulose; anteriorly; distal process present; digitiform; denticulate; not bicuspidate; without transverse row of denticles; none oblique row of 5 denticles; at anterior surface; not innerly directed; with seta; spiniform; ornamented by spinules; not surpassing the distal-point of the segment; without outer spine; terminal claw absent.

##### Fifth right swimming leg

Biramous. Fifth right swimming leg praecoxa present; separated from the coxae; without ornamentation. Fifth right swimming leg coxa convex inner side; without teeth-like structures; with process; rounded; distally inserted; on posterior surface; closest to the outer rim; projecting over basis; beyond the first third; until the medial surface; without triangular protuberance innerly; with sensilla; slender; at apex; no longer 2x than basal insertion; without marginal extension; without seta; without spinules. Fifth right swimming leg basis cylindrical; unequal size between inner and outer side; shorter outer than inner side; rectilinear inner side; tumescence absent; without protuberance; absence of distinct minutely granular; additional inner process absent; with posterior groove; deep; obliquely; not reaching the endopodal lobe; not ornamented; with seta; outerly inserted; on anterior surface; no longer 2x than origin segment; posterior protrusion present; distal process absent. Fifth right swimming leg with endopodite present; fused to basis; on anterior surface; ancestral segments 1 and 2 fused; ancestral segments 2 and 3 fused; stout; ornamented; with setules; as a row; on inner side; terminally; without seta. Fifth right swimming leg exopod segments 1 and 2 separated; segments 2 and 3 fused; 2-segmented; stout; separated from the basis. Fifth right swimming leg exopod 1 sub-cylindrical; longer than broad; nearly 1.25 times; unequal size between both sides; shorter inner than outer side; convex inner side; rectilinear outer side; with marginal extension; sub-triangular; distally inserted; at outer rim; spinules absent; with process; triangular; rectilinear; blunt tip; sclerotized; without ornamentation; distally inserted; at posterior surface; not projecting over next segment; without outer spine; without seta; internal prominence absent; lamella on posterior surface absent. Fifth right swimming leg exopod 2 cylindrical; longer than broad; nearly 2 times; equal size between both sides; disform inner side; convex outer side; with posterior proximal swelling; inner-posterior process absent; without marginal expansion; curved ridge on distal posterior surface absent; chitinous knobs present; with 3–6 posteriorly; with outer spine; inserted medially; arched; internally directed; ornamented innerly; by spinules; as a row; not ornamented outerly; sharp tip; with apparent curve; outerly directed; lesser than the length of the exopod 2; until to 2 times its size; 1.5x; sensilla absent; terminal claw present; not equal or longer 1.5 times than insertion segment; sclerotized; arched; inward; with conspicuous curve; proximally; ornamented innerly; by spinules; as a row; partially on extension; medially, or distally; ornamented outerly; sharp tip; not curved tip; without medial constriction; hyaline process absent.

##### FEMALE

Body longer and wider than male; body 1168 micrometers excluding caudal setae. Widest at first metasome segment. Distal margin of the prosomal segments without one line of setules at posterior margin. Prosome segments with spinules at least at one prosomal segment. Fourth metasome segment absence of dorsal protuberance. Fourth and fifth metasome segments fused; partially; on dorsal surface. Limit between fourth and fifth metasome segments ornamented; with spinules; as a row; double; complete; same size; partially over limit; dorsally. **Fifth metasome segment**. Fifth metasome segment with sensilla; dorsally; 2 elements; with epimeral plates. Epimeral plates asymmetrical. Right epimeral plates prominent, as projections; thinner than the left; one posterior-laterally directed; not reaching half length of the genital segment; with sensilla at the apex; dorsal-posterior sensilla present; slender; without ornamentation. Left epimeral plate without expansion.

##### Urosome

3-segmented. **Genital double-somite**. Asymmetrical in dorsal view; longer than broad; longer than other urosomites combined; dorsal suture at mid-length absent; not covered by spinules; with swelling; rounded; unequal size; greater right than left; anteriorly; with sensillae; on both sides; one; stout; with robust apex; at left lateral; not on lobular base; medially; one; stout; at right lateral; not on lobular base; anteriorly; with robust apex; of equal size between then; lateral protuberance absent; without right posterior rim expanded; without slender sensilla on each posterior rim; with posterior-dorsal process; on either side; single on the left; double on the right. Genital double-somite opercular pad present; broader than longer; symmetrical; development laterally; expanded posteriorly; covering partially; double gonoporal slit; located ventrally; with arthrodial membrane; inserted anteriorly; post-genital process absent; disto-ventral tumescence absent; ventral vertical folds present; dorsal sensilla absent. Second urosome segment with ventral fusion to anal segment; right distal process absent. Caudal rami patch of setules on outer surface absent; patch of spinules on outer surface absent.

##### Appendices features

Rostrum basal process absent. **Antennules**. Symmetrical. Right antennule surpassing to genital double-segment; extending beyond caudal rami. Right antennule exceeding the caudal setae. Right antennule ornamentation pattern equals to male left antennule; mostly. Actual segment 13 without seta; without aesthetasc. Actual segment 14 without seta; without aesthetasc. Actual segment 15 without seta; without aesthetasc. Actual segment 16 without seta; without aesthetasc. Actual segment 17 without seta. Actual segment 18 without seta.

##### Fifth swimming legs

Symmetrical; Fifth swimming legs biramous. Fifth swimming legs intercoxal plate longer than wide; separated from the legs. Fifth swimming legs praecoxa with sclerite praecoxal; separated from the coxae; without ornamentation. Fifth swimming legs coxa with process; conical; at the outer rim; distally; sensilla present; stout; at apex; projecting over basal segment; no longer 2x than basal insertion; marginal extension absent; without swelling; without seta; without spinules. Fifth swimming legs basis sub-triangular; unequal size between inner and outer sides; shorter outer than inner side; with convex inner side; without proximal inner outgrowth; without groove; with distal extension; on posterior surface; with seta; outerly inserted; on anterior surface; longer 2x than origin segment; not reaching to exopod 1 distally. Fifth swimming legs endopod segments 1 and 2 separated; segments 2 and 3 fused; 2-segmented; with complete suture; stout; separated from the basis; ornamentation on segment 2; with spinules; as a row; single; non-oblique; sub-terminally; at anterior surface; with seta; double; one medially; on posterior surface; rectilinear; one distally; on posterior surface; arched; of unequal size; distal seta longer than medial seta. Fifth swimming legs exopod segments 1 and 2 separated; segments 2 and 3 separated; 3-segmented; separated from the basis. Fifth swimming legs exopod 1 sub-cylindrical; longer than wide; longer or equal than 2 times; with unequal size between inner and outer side; shorter inner than outer side; with convex inner side; with rectilinear outer side; without swelling; without marginal extension; without posterior process; without spine; without seta. Fifth swimming legs exopod 2 sub-cylindrical; longer than broad; longer or equal than 2 times; without swelling; without marginal extension; without process; without lobe; with spine; inserted laterally; rectilinear; without ornamentation; sharp tip; equal size or larger than next segment; without seta. Fifth swimming legs exopod 3 cylindrical; longer than wide; without swelling; without process; without lobe; without spine; with seta; double; inserted terminally; unequal size between them; outer seta smaller than inner; nearly 3 times; outer seta not ornamented by setules; without ornamentation; presence of terminal claw; sclerotized; arched; externally directed; convex inner side; with ornamentation; of denticles; as a row; on surface partially; at medial region; concave outer side; with ornamentation; of denticles; as a row; on surface partially; at medial region; blunt tip; 6 times longer than origin segment.

##### Distribution records

###### BRAZIL

**Paraná**: Segredo Reservoir, Iguaçú River (Lopes *et al*., 1997); Salto Santiago Reservoir, Iguaçú River (this study). **Rio Grande do Sul** (Brandorff, 1976). PARAGUAY. Swamps in Caerapa, and Villa Rica (Daday, 1905). ARGENTINA. Middle Paraná River (Paggi & José de Paggi, 1990). **Buenos Aires**: Pergamino Stream (Ringuelet, 1958a); Hoya del Plata (Ringuelet, 1962). **Capital Federal**: artificial lake near to Buenos Aires Cricket Club, and Lake La Administración in Park 3 of Febrero, both in Palermo (Wright, 1939). **Chaco**: pond in Makallé (Ringuelet, 1958a). **Entre Rios**: Concordia, and Colón, Uruguay River (Brian, 1926). **Formosa**: lagoon Yema (Brehm, 1958a; 1965). **Santa Fé**: ponds in Crespo, Calchaquí, and Guadalupe (Ringuelet, 1958a); Calchaqui Brehm, 1958; 1965); arroio Ubajay, La Capital (Ringuelet & Martinez de Ferrato, 1967). URUGUAY. **Salto**: Salto, Uruguai River (Brian, 1926). River mount of La Plata, in Montevideo, probably (Brehm, 1938; 1939). **Plata river basin**: low Paraná River (Perbiche-Neves *et al*., 2015).

##### Habitat

Habitat in freshwaters: rivers, reservoirs, swamps, stream, and ponds.

##### Remarks

This is a nominal species presented to science through specimens from Paraguay, possibly (Daday, 1905). In the original description are related only the localities “permanent lagoon” in Caearapa and Villa Rica, without the formal designation of type material. Its current recombination was carried out in the foundation of *Notodiaptomus*, considered part of *nordestinus* only in the amplification (Wright, 1937) and revision from specimens from Argentina (USNM 92971), Uruguay (Kiefer Collection 1105) and Paraná, Brazil (Santos-Silva *et al*., 2015).

The taxonomic trajectory of the species is based on unknowns about its true identity, historically confused with *N. inflexus* Brian 1926, *N. perelegans* (Wright, 1927) and a complex named the “bidigitatus group” (Brandorff, 1976). Some recent studies maintain the issue (Perbiche-Neves *et al*., 2015), others have evidenced clarifications for its taxonomic status (Paggi, 2001; Santos-Silva *et al*., 2015). The morphological evidence presented in the *nordestinus* review was corroborated for female: (1) genital double-somite with posterior-dorsal process on either side, right double; (2) P5 endopod 2-segmented, (3) presence of dorsal-posterior sensilla on epimeral plates; and male: (1) P5R exopod 1 with inner process distally; (2) P5R exopod 2 with chitinous knob posteriorly; (3) P5R exopod 1 with inner-distal process posteriorly.

In the present effort, the present variations are for the amount of chitinous knobs on male P5R exopod 2, always present three semicircular processes and not two as evidenced in Santos-Silva *et al*. (2015). Other conditions were present for the male P5R exopod 1 inner-distal process posteriorly, such as triangular shape, rectilinear direction (different from *N. deitersi*, in arched condition), and not projecting over exopod 2. Additionally, they were present for male: A1R actual segment 8 with conical seta length reaching to middle-point of sequent segment, and fourth metasome segment with ornamented posterior-margin with double spinules row, dorsally and laterally. For females, the observation of the genital double-somite with ventral vertical folds is relevant for the distinction of the species from its congeners.

Among the attributes originally reported for the creation and amplification of *Notodiaptomus* (Kiefer, 1936; 1956), only the conditions for female P5 endopod unisegmented, and for male P5R exopod 2 with outer spine sub-distally have not been corroborated. The female endopod and the positioning of the outer spine masculine are 2-segmented, and on exopod 2 medially, respectively. Additionally, the fulfillment of the outer arrow on P5 basis highlighted by Brehm (1939) has been confirmed, although it does not reach exopod 1 distally as in *N. deitersi.* Nevertheless, all other characteristics are present and reinforce the position of the species in *Notodiaptomus*.

#### Notodiaptomus brandorffi Reid, 1987

##### Synonymy

*Notodiaptomus brandorffi* Reid, 1987: 364, 372, 377, figs. 32–59; Reid & Turner, 1988: 489, 492; Sendacz, 1993: 35; Rocha *et al*., 1995: 156; Santos-Silva, 1998: 206; Perbiche-Neves *et al*., 2020: 681-682, key to the Neotropical diaptomid, fig. 21.8 L.

##### Type locality

Lake Açu, Maranhão State, Brazil, 3°50m South, 44°55m West.

##### Type material

Holotype: female (MZUSP 7231); paratypes: 01 male (MZUSP 7232), 15 females (MZUSP 7233), and 19 males (MZUSP 7234); 20 females and 20 males (USNM 227108); 2 females, and 3 males (built in slides); from the Lake Açu (3°50m S, 44°55m W), Maranhão State, Brazil, 11.X.1984, M.S.R. Ibañez collector. Additional material: 4 males, and 3 females (USNM 227123); 1 male (built in slides); all from the Betume city, Sergipe State, Brazil (10° 19m S, 36° 35m W), 18.III.1983, E.R. dos Santos coll. The undissected material is reported as preserved entire in alcohol. All specimens without a deposit code were stored in the J. Reid collection.

##### Material examined

Holotype: male (MZUSP 7231). Paratype: female (MZUSP 7232), 4 females (MZUSP 7233). Non-type material: 5 males, and 6 females, from the Lake Açu, Maranhão States, 13.X.1983, A. Darwich Coll. Except the holotype, all individuals examined were preserved entire in alcohol; 1 male (INPA-COP007, slides a-h) and 1 female (INPA-COP008, slides a-h) were selected to be dissection on eight slides each and deposited in the Zoological Collection of the INPA, Brazil.

##### Diagnosis

**(1)** Male right and left antennules presenting length not surpassing to genital segment; **(2)** first swimming legs coxa presenting anterior spinules row distally; **(3)** P2 to P4 endopod 3 presenting double spinules row on posterior surface distally; **(4)** P2 exopods 2, and 3 absence setules on inner surface; **(5)** P3 exopod 3 absence row of spinules at posterior surface distally; **(6)** P4 coxa presenting inner distal setae with plumose composition, and surpassing to basal segment; **(7)** P1 to P4 exopod 2 absence setules on inner surface; **(8)** male fifth left swimming leg coxa presenting sensilla length no longer 2x than basal insertion; **(9)** male fifth left swimming leg basis presenting rounded internal proximal expansion; **(10)** male fifth left swimming leg basis absence groove posteriorly; **(11)** male fifth left swimming leg exopod 1 with sub-cylindrical form; **(12)** male fifth left swimming leg exopod 1 presenting unequal size between inner and outer side, concave inner side; **(13)** male fifth left swimming leg exopod 1 presenting posterior proximal process; **(14)** male fifth left swimming leg exopod 2 with spiniform seta not surpassing the distal-point of the segment; **(15)** male fifth right swimming leg basis presenting convex inner side; **(16)** male fifth right swimming leg basis absence inner intumescence proximally; **(17)** male fifth right swimming leg basis absence protuberance on inner side; **(18)** male fifth right swimming leg exopod 2 elliptical, longer than broad, nearly 2 times; **(19)** male fifth right swimming leg exopod 2 presenting inner ornamentation on outer spine; **(20)** male fifth right swimming leg exopod 2 presenting terminal claw with inner ornamentation, not curved tip; **(21)** female fifth metasome segments ornamented with dorsal spinules row, single, incomplete; **(22)** female right epimeral plate presenting dorsal posterior sensilla with slender condition; **(23)** female genital double-somite presenting post-genital process, single; **(24)** female fifth swimming legs basis presenting outer seta on posterior surface; **(25)** female fifth swimming legs endopod presenting distal seta rectilinear.

##### Redescription

###### MALE

Body 1089 micrometers excluding caudal setae. Body smaller and slenderer than female. Nerve axons myelinated. Prosome 6-segmented; widest at first metasome segment; without one line of setules at posterior margin; without spinules at segments. Cephalosome anterior margin rounded; with dorsal suture; complete; separate from first metasome segment. First metasome segment without sensilla. Second metasome segment without sensilla. Third metasome segment without sensillae; non-ornamented posterior margin. Fourth metasome segment without sensillae; separated from the fifth metasome. Limit between fourth and fifth metasome segments without ornamentation. Fifth metasome segment without sensilla; Fifth metasome segment without ornamentation; Fifth metasome segment without dorsal conical process; with epimeral plates. Epimeral plates symmetrical. Right epimeral plates reduced, as rounded distal corner segment limit; with sensilla; at the apex of projection; without ornamentation.

##### Urosome

5-segmented; Urosome 5-free segments. Genital somite asymmetrical in dorsal view; with single aperture; located on left side; ventrolaterally on posterior rim; with sensillae; on both sides; one; at left lateral; posteriorly; one; at right rim; posteriorly; of equal size between then. Third urosome segment without spinules; without external seta. Fourth urosome segment without spinules; without sub-conical blunt dorsal-lateral process. Anal segment absence of dorsal sensillae; presence of operculum; convex; covering the anal aperture fully. Caudal rami symmetrical; separated from anal segment; longer than wide; with setules; continuous on; inner side; each ramus bearing 6 caudal setae; 5 marginals; plumose; and 1 internal dorsally; straight; not reticulated main axis; outermost seta with outer spiniform process absent.

##### Appendices features

Rostrum symmetrical; separated from dorsal cephalic shield; by complete suture; sensillae present; one pair; anteriorly inserted on surface tegument; with rostral filament; double; paired; extended; into point; with basal process; in ventral view, rounded on left side; without a smaller basal expansion on the right side.

##### Antennules

Asymmetrical. **Right antennules**. Uniramous; right antennule not surpassing to genital segment.

Right antennule ancestral segment I and II separated. Ancestral segment II and III fused. Ancestral segment III and IV fused. Ancestral segment IV and V separated. Ancestral segment V and VI separated. Ancestral segment VI and VII separated. Ancestral segment VII and VIII separated. Ancestral segment VIII and IX separated. Ancestral segment IX and X separated. Ancestral segment X and XI separated. Ancestral segment XI and XII separated. Ancestral segment XII and XIII separated. Ancestral segment XIII and XIV separated. Ancestral segment XIV and XV separated. Ancestral segment XV and XVI separated. Ancestral segment XVI and XVII separated. Ancestral segment XVII and XVIII separated. Ancestral segment XVIII and XIX separated. Ancestral segment XIX and XX separated. Ancestral segment XX and XXI separated. Ancestral segment XXI and XXII fused. Ancestral segment XXII and XXIII fused. Ancestral segment XXIII and XXIV separated. Ancestral segment XXIV and XXV fused. Ancestral segment XXV and XXVI separated. Ancestral segment XXVI and XXVII separated. Ancestral segment XXVII and XXVIII fused.

Right antennule actual 22-segmented; geniculated; between the segment 18 and segment 19; with swollen and modified region; formed by 5 segments; between 13 and 17 segments. Actual segment 1 with seta; one element; straight; none larger than segment; without spinules; without vestigial seta; without conical seta; without modified seta; without spinous process; with aesthetasc; one element. Actual segment 2 with seta; three elements; of unequal size; straight; none larger than segment; without spinules; with vestigial seta; one element; without conical seta; without modified seta; without spinous process; with aesthetasc; one element. Actual segment 3 with seta; one element; one larger than segment; surpassing to distal margin; beyond three sequential segments; straight; blunt apex; without spinules; with vestigial seta; one element; without conical seta; without modified seta; without spinous process; with aesthetasc. Actual segment 4 with seta; one element; one larger than segment; surpassing to distal margin; straight; not beyond three sequential segments; without spinules; without vestigial seta; without conical seta; without modified seta; without spinous process; without aesthetasc. Actual segment 5 with seta; one element; straight; one larger than segment; surpassing to distal margin; not beyond three sequential segments; without spinules; with vestigial seta; one element; without conical seta; without modified seta; without spinous process; with aesthetasc; one element. Actual segment 6 with seta; one element; none larger than segment; straight; without spinules; without vestigial seta; without conical seta; without modified seta; without spinous process; without aesthetasc. Actual segment 7 with seta; one element; straight; one larger than segment; surpassing to distal margin; beyond three sequential segments; blunt apex; without spinules; without vestigial seta; without conical seta; without modified seta; without spinous process; with aesthetasc; one element. Actual segment 8 with seta; one element; straight; none larger than segment; without spinules; without vestigial seta; with conical seta; one element; not reaching to middle-point of the sequent segment; without modified seta; without spinous process; without aesthetasc. Actual segment 9 with seta; two elements; of unequal size; straight; one larger than segment; surpassing to distal margin; beyond three sequential segments; blunt apex; without spinules; without vestigial seta; without conical seta; without modified seta; without spinous process; with aesthetasc; one element. Actual segment 10 with seta; one element; straight; none larger than segment; without spinules; without vestigial seta; without conical seta; with modified seta; presenting blunt apex; slender form; surpassing to distal margin; beyond of the sequential segment; parallel to antennule direction; without spinous process; without aesthetasc. Actual segment 11 with seta; one element; straight; one larger than segment; surpassing to distal margin; not beyond three sequential segments; without spinules; without vestigial seta; without conical seta; with modified seta; slender form; presenting blunt apex; surpassing to distal margin; beyond of the sequential segment; parallel to antennule direction; shorter length than homologous of actual segment 13; without spinous process; without aesthetasc. Actual segment 12 with seta; one element; straight; one larger than segment; surpassing to distal margin; not beyond three sequential segments; without spinules; without vestigial seta; with conical seta; one element; not smaller than to segment 8; without modified seta; without spinous process; with aesthetasc; one element; absent internal perpendicular fission. Actual segment 13 with seta; one element; straight; one larger than segment; surpassing to distal margin; not beyond three sequential segments; without spinules; without vestigial seta; without conical seta; with modified seta; stout form; surpassing to distal margin; to the middle-point of the sequence segment; perpendicular to antennule direction; presenting bifid apex; without spinous process; with aesthetasc; one element. Actual segment 14 with seta; two elements; of unequal size; straight; one larger than segment; surpassing to distal margin; beyond three sequential segments; blunt apex; without spinules; without vestigial seta; without conical seta; without modified seta; without spinous process; with aesthetasc; one element. Actual segment 15 with seta; two elements; of unequal size; straight; not bifidform; none larger than segment; without spinules; without vestigial seta; without conical seta; without modified seta; with spinous process; on outer margin; surpassing distal margin; with aesthetasc; one element. Actual segment 16 with seta; two elements; of unequal size; plumose; one larger than segment; surpassing to distal margin; not beyond three sequential segments; not bifidform; without spinules; without vestigial seta; without conical seta; without modified seta; with spinous process; on outer margin; surpassing distal margin; unequal size to process on preceding segment; with aesthetasc; one element. Actual segment 17 with seta; two elements; of unequal size; straight; none larger than segment; bifidform; without spinules; without vestigial seta; without conical seta; with modified seta; one element; stout form; surpassing to distal margin; not beyond of the sequential segment; parallel to antennule direction; without spinous process; without aesthetasc. Actual segment 18 with seta; two elements; of equal size; straight; none larger than segment; without spinules; without vestigial seta; without conical seta; with modified seta; one element; stout form; surpassing distal margin; parallel to antennule direction; without spinous process; without aesthetasc. Actual segment 19 with seta; two elements; of unequal size; plumose; none larger than segment; without spinules; without vestigial seta; without conical seta; with modified seta; two elements; stout form; at least one bifid form; surpassing distal margin; parallel to antennule direction; without spinous process; with aesthetasc; one element. Actual segment 20 with seta; four elements; of unequal size; straight; one larger than segment; surpassing to distal margin; beyond three sequential segments; without spinules; without vestigial seta; without conical seta; without modified seta; without spinous process; without aesthetasc. Actual segment 21 with seta; two elements; of equal size; plumose; one larger than segment; surpassing to distal margin; greater 3x than original segment; without spinules; without vestigial seta; without conical seta; without modified seta; without spinous process; without aesthetasc. Actual segment 22 with seta; four elements; of equal size; one larger than segment; plumose; surpassing to distal margin; greater 3x than original segment; without spinules; without vestigial seta; without conical seta; without modified seta; without spinous process; with aesthetasc; one element.

##### Left antennules

Uniramous; Left antennule not surpassing to prosome. Ancestral segment I and II separated. Ancestral segment II and III fused. Ancestral segment III and IV fused. Ancestral segment IV and V separated. Ancestral segment V and VI separated. Ancestral segment VI and VII separated. Ancestral segment VII and VIII separated. Ancestral segment VIII and IX separated. Ancestral segment IX and X separated. Ancestral segment X and XI separated. Ancestral segment XI and XII separated. Ancestral segment XII and XIII separated. Ancestral segment XIII and XIV separated. Ancestral segment XIV and XV separated. Ancestral segment XV and XVI separated. Ancestral segment XVI and XVII separated. Ancestral segment XVII and XVIII separated. Ancestral segment XVIII and XIX separated. Ancestral segment XIX and XX separated. Ancestral segment XX and XXI separated. Ancestral segment XXI and XXII separated. Ancestral segment XXII and XXIII separated. Ancestral segment XXIII and XXIV separated. Ancestral segment XXIV and XXV separated. Ancestral segment XXV and XXVI separated. Ancestral segment XXVI and XXVII separated. Ancestral segment XXVII and XXVIII fused.

Left antennule actual 25-segmented; not-geniculated. Actual segment 1 with seta; one element; none larger than segment; straight; without spinules; without vestigial seta; without conical seta; without modified seta; without spinous process; with aesthetasc; one element. Actual segment 2 with seta; three elements; of equal size; none larger than segment; straight; without spinules; with vestigial seta; one element; without conical seta; without modified seta; without spinous process; with aesthetasc; one element. Actual segment 3 with seta; one element; one larger than segment; straight; surpassing to distal margin; beyond three sequential segments; without spinules; with vestigial seta; one element; without conical seta; without modified seta; without spinous process; with aesthetasc. Actual segment 4 with seta; one element; none larger than segment; straight; without spinules; without vestigial seta; without conical seta; without modified seta; without spinous process; without aesthetasc. Actual segment 5 with seta; one element; one larger than segment; straight; surpassing to distal margin; not beyond three sequential segments; without spinules; with vestigial seta; one element; without conical seta; without modified seta; without spinous process; with aesthetasc; one element. Actual segment 6 with seta; one element; none larger than segment; straight; without spinules; without vestigial seta; without conical seta; without modified seta; without spinous process; without aesthetasc. Actual segment 7 with seta; one element; one larger than segment; straight; surpassing to distal margin; beyond three sequential segments; without spinules; without vestigial seta; without conical seta; without modified seta; without spinous process; with aesthetasc; one element. Actual segment 8 with seta; one element; one larger than segment; straight; surpassing distal margin; without spinules; without vestigial seta; with conical seta; without modified seta; without spinous process; without aesthetasc. Actual segment 9 with seta; two elements; of unequal size; one larger than segment; straight; surpassing to distal margin; beyond three sequential segments; without spinules; without vestigial seta; without conical seta; without modified seta; without spinous process; with aesthetasc; one element. Actual segment 10 with seta; one element; none larger than segment; straight; without spinules; without vestigial seta; without conical seta; without modified seta; without spinous process; without aesthetasc. Actual segment 11 with seta; one element; one larger than segment; straight; surpassing to distal margin; beyond three sequential segments; without spinules; without vestigial seta; without conical seta; without modified seta; without spinous process; without aesthetasc. Actual segment 12 with seta; one element; one larger than segment; straight; surpassing distal margin; without spinules; without vestigial seta; with conical seta; without modified seta; without spinous process; with aesthetasc; one element. Actual segment 13 with seta; one element; none elongated; straight; surpassing distal margin; without spinules; without vestigial seta; without conical seta; without modified seta; without spinous process; without aesthetasc. Actual segment 14 with seta; one element; elongated; straight; surpassing to distal margin; beyond three sequential segments; without spinules; without vestigial seta; without conical seta; without modified seta; without spinous process; with aesthetasc; one element. Actual segment 15 with seta; one element; larger than segment; straight; surpassing to distal margin; not beyond three sequential segments; without spinules; without vestigial seta; without conical seta; without modified seta; without spinous process; without aesthetasc. Actual segment 16 with seta; one element; larger than segment; plumose; surpassing to distal margin; not beyond three sequential segments; without spinules; without vestigial seta; without conical seta; without modified seta; without spinous process; with aesthetasc; one element. Actual segment 17 with seta; one element; not larger than segment; straight; without spinules; without vestigial seta; without conical seta; without modified seta; without spinous process; without aesthetasc. Actual segment 18 with seta; one element; larger than segment; straight; surpassing to distal margin; beyond three sequential segments; without spinules; without vestigial seta; without conical seta; without modified seta; without spinous process; without aesthetasc. Actual segment 19 with seta; one element; not larger than segment; straight; surpassing distal margin; without spinules; without vestigial seta; without conical seta; without modified seta; without spinous process; with aesthetasc; one element. Actual segment 20 with seta; one element; not larger than segment; straight; surpassing distal margin; without spinules; without vestigial seta; without conical seta; without modified seta; without spinous process; without aesthetasc. Actual segment 21 with seta; one element; larger than segment; plumose; surpassing to distal margin; beyond three sequential segments; without spinules; without vestigial seta; without conical seta; without modified seta; without spinous process; without aesthetasc. Actual segment 22 with seta; two elements; of unequal size; one of them elongated; plumose; surpassing to distal margin; without spinules; without vestigial seta; without conical seta; without modified seta; without spinous process; without aesthetasc. Actual segment 23 with seta; two elements; of unequal size; one larger than segment; plumose; surpassing to distal margin; greater 3x than original segment; without spinules; without vestigial seta; without conical seta; without modified seta; without spinous process; without aesthetasc. Actual segment 24 with seta; two elements; of equal size; one larger than segment; plumose; surpassing to distal margin; greater 3x than original segment; without spinules; without vestigial seta; without conical seta; without modified seta; without spinous process; without aesthetasc. Actual segment 25 with seta; four elements; of equal size; elongated; plumose; surpassing to distal margin; 4 times larger than segment; without spinules; without vestigial seta; without conical seta; without modified seta; without spinous process; with aesthetasc; one element.

##### Antenna

Biramous. Antenna coxa separated from the basis; bearing seta; 1; on inner surface; at distal corner; reaching to the endopod 1. Antenna basis (fusion) separated from the endopodal segment; bearing seta; 2; on inner surface; at distal corner. Endopodal ancestral segment I and II separated. Ancestral segment II and III fused. Ancestral segment III and IV separated. Antenna endopod actual 3-segmented. Actual segment 1 not bilobate; with seta; two; on inner margin; with spinules; as a row; obliquely; on outer surface; without pore. Actual segment 2 not bilobate; without discontinuity on outer cuticle; inner lobe bearing 9 setae; distally; without spinules. Actual segment 3 outer lobe bearing 8 setae; distally; with spinules; as a row; not obliquely; on outer surface. Antenna exopod ancestral segment I and II separated. Ancestral segment II and III fused. Ancestral segment III and IV fused. Ancestral segment IV and V separated. Ancestral segment V and VI separated. Ancestral segment VI and VII separated. Ancestral segment VII and VIII separated. Ancestral segment VIII and IX separated. Ancestral segment IX and X fused. Antenna exopod actual 7-segmented. Actual segment 1 single; elongated (width-length, equal or larger ratio 2:1); with seta; one; at inner surface. Actual segment 2 compound; elongated (larger width-length ratio 2:1); with seta; three; at inner surface. Actual segment 3 single; not elongated (lesser width-length ratio 2:1); with seta; one; at inner surface. Actual segment 4 single; not elongated (lesser width-length ratio 2:1); with seta; one; at inner surface. Actual segment 5 single; not elongated (lesser width-length ratio 2:1); with seta; one; at inner surface. Actual segment 6 single; not elongated (lesser width-length ratio 2:1); with seta; one; at inner surface. Actual segment 7 compound; elongated (larger or equal width-length ratio 2:1); with seta; one; at inner surface; and three; at distal surface.

##### Oral features

**Mandible**. Coxal gnathobase sclerotized; with lobe; not prominent; on caudal margin; presence of cutting blade; with tooth-like prominence; two, distinctly; 1 acute; on caudal margin; and 1 triangular; on sub-caudal margin; without acute projection between the prominences; with additional spinules; as a row; on dorsal surface; with seta; 1; dorsally; on apical surface; with spinules; apicalmost. Mandible palps biramous; comprising the basis; with seta; four; differently inserted; first medially; reaching to beyond the endopod 1; second distally; third distally; fourth distally; on inner margin; none with setulose ornamentation. Mandible endopod 2-segmented. Mandible endopod 1 with lobe; bearing seta; four; distally inserted; without spinules. Mandible endopod 2 without lobe; bearing setae; nine elements; distally inserted; with spinules; as a row; double. Mandible exopod 4-segmented. Mandible exopod 1 with seta; one element; distally; on inner margin. Mandible exopod 2 with seta; one element; distally; on inner side. Mandible exopod 3 with seta; one element; distally; on inner side. Mandible exopod 4 with setae; three elements; on terminal region. **Maxillule**. Birramous. Maxillule 3-segmented. Maxillule praecoxa with praecoxal arthrite; bearing spines; fifteen elements; ten marginally; plus, five sub-marginally; with spinules; as a patch; on sub-marginal surface. Maxillule coxa with coxal epipodite; with conspicuous outer lobe; bearing setae; nine elements; with coxal endite; elongated (larger or equal width-length ratio 2:1); bearing setae; four elements. Maxillule basis with basal endite; double; first proximal; elongated (larger width-length ratio 2:1; separated from basis; with setae; four elements; distally inserted; second distal; fused to basis; not elongated (lesser width-length ratio 2:1); with setae; four elements; distally inserted; with setules; as a row; on inner side; basal exite present; with setae; one element; on outer surface. Maxillule endopod 1-segmented. Endopod 1 bilobate; first proximal; with setae; three elements; second distal; with setae; five elements. Maxillule exopod 1-segmented. Exopod 1 with setae; six elements; with setules; as a row; on inner side; spinules absent. **Maxilla**. Uniramous. Maxilla 5-segmented. Maxilla praecoxa fused to coxa; incompletely; distinct externally; with praecoxal endite; double; first elongated endite (larger or equal width length ratio 2:1); proximally inserted; with seta; straight, or plumose; 1 straight; 4 plumose; with spine; single; without spinules; without setule; second elongated endite (larger or equal width length ratio 2:1); distally inserted; with seta; plumose; 3 plumose; without spine; with spinules; as a row; on distal margin; with setule; as a row; on distal margin; absence of outer seta. Maxilla coxa with coxal endite; double; first elongated endite (larger or equal width); proximally inserted; with seta; plumose; 3 plumose; without spine; without spinules; with setules; as a row; on proximal margin; second elongated endite (larger or equal width); distally inserted; with seta; plumose; 3 plumose; without spine; without spinules; with setules; as a row; on proximal margin; absence of outer seta. Maxilla basis with basal endite; single; elongated (larger or equal width-length ratio 2:1); with seta; plumose; 3 plumose; without spinules; absence of outer seta. Maxilla endopod 2-segmented. Endopod 1 with seta; 2 plumose; without spine; without spinules; without setules. Maxilla endopod 2 with seta; 2 plumose; without spine; without spinules; without setules. **Maxilliped**. Uniramous; Maxilliped 8-segmented. Maxilliped praecoxa fused to coxa; incompletely; distinct internally; with praecoxal endite; not elongated (lesser width-length ratio 2:1); distally inserted; with seta; 1 straight; with spinules; as a row; single; on basal surface; without setules. Maxilliped coxa with coxal endite; three coxal endite; first elongated (larger or equal width); proximally inserted; with seta; 2 plumose; with spinules; as a patch; single; on apical surface; without setules; second not elongated (lesser width-length ratio 2:1); medially inserted; with seta; 3 plumose; with spinules; as a row; single; on medial surface; without setules; third elongated (larger or equal width length ratio 2:1); distally inserted; with seta; 3 plumose; none reaching to beyond of the basis; with spinules; as a row; single; on basal surface; without setules; with lobe; prominence; at inner distal angle; ornamented; with spinules; continuously on margin. Maxilliped basis without basal endite; with seta; 3 plumose; with spinules; as a row; single; on medial surface; with setules; as a row; single; on inner margin. Maxilliped endopod segment 6-segmented. Endopod 1 with seta; 2 plumose; on inner surface. Endopod 2 with seta; 3 plumose; on inner surface. Endopod 3 with seta; 2 plumose; on inner surface. Endopod 4 with seta; 2 plumose; on inner surface. Endopod 5 with seta; 2 plumose; on inner surface, or on outer surface; outer seta absent. Endopod 6 with seta; 4 plumose; on inner surface, or on outer surface.

##### Swimming legs features

**First swimming legs.** Symmetrical; biramous. First swimming legs intercoxal plate without seta. First swimming legs praecoxa absent. First swimming legs coxa with seta; one; plumose; distally inserted; on inner surface; surpassing to basal segment; with setules; one group; as a row; discontinuously; on inner margin; with spinules; as a row; single; distally inserted; at anterior surface; without spine. First swimming legs basis without seta; with setules; as a patch; single; on outer surface; without spinules; without spine. First swimming legs endopod 2-segmented. Endopod 1 with seta; straight; restricted; to inner surface; one element; without spine; with setules; as a row; single; continuously; on outer surface; without spinules; absence of Schmeil’s organ. Endopod 2 with seta; unrestricted; three on inner surface; one on outer surface; two on distal surface; plumose; without spine; with setules; as a row; single; continuously; on outer surface; without spinules; absence of Schmeil’s organ. Endopod 3 absence. First swimming legs exopod 1 with seta; restricted; 1 on inner surface; with spine; 1; stout; smaller than original segment; serrated; on inner side; continuously; without setules. First swimming legs exopod 2 with seta; restricted; 1 on inner surface; plumose; without spine; with setules; as a row; single; continuously; on outer margin; without spinules. First swimming legs exopod 3 with setule; as a row; single; continuously; on outer surface; without spinules; with seta; unrestricted; 2 on inner surface; 2 on terminal surface; with spine; 2; unequal size; first no longer 2x than origin segment; stout; serrated; on inner side, or on outer side; equally; second longer 3x than origin segment; slender; serrated; on outer side; with ornamentation on non-serrated side; by setules. **Second swimming legs**. Symmetrical; Second swimming legs biramous. Second swimming legs intercoxal plate without seta. Second swimming legs praecoxa present; located laterally. Second swimming legs coxa with seta; straight; distally inserted; on inner surface; surpassing to basal segment; without setules; without spinules; without spine. Second swimming legs basis without seta; without setules; without spinules; without spine. Second swimming legs endopod 3-segmented. Endopod 1 with seta; straight; restricted; one on inner surface; without spine; without setules; without spinules; absence of Schmeil’s organ. Endopod 2 with seta; plumose; unrestricted; two on inner surface; without spine; with setules; as a row; single; continuously; on outer side; without spinules; presence of Schmeil’s organ; on posterior surface. Endopod 3 with seta; plumose; unrestricted; three on inner surface; two on outer surface; two on distal surface; without spine; without setules; with spinules; as a row; double; distally inserted; at posterior surface; absence of Schmeil’s organ. Second swimming legs exopod 1 with seta; restricted; one on inner surface; with spine; 1; stout; not reaching to distal-third of the exopod 2; serrated; on inner side, or on outer side; with setules; as a row; single; continuously; on inner side; without spinules; absence of Schmeil’s organ. Exopod 2 with seta; unrestricted; one on inner surface; with spine; 1; stout; not surpassing the exopod 3; serrated; on inner side, or on outer side; without setules; without spinules; absence of Schmeil’s organ. Exopod 3 with seta; plurimarginal; three on inner surface; two on terminal surface; with spine; 2; unequal size; first no longer 2x than origin segment; stout; serrated; on inner side, or on outer side; equally; second longer 2x than origin segment; slender; serrated; on outer side; with ornamentation on non-serrated side; of setules; no setules on outer surface; without spinules; absence of Schmeil’s organ. **Third swimming legs**. Symmetrical; Third swimming legs biramous. Third swimming legs intercoxal plate without seta. Third swimming legs praecoxa present; not laterally located. Third swimming legs coxa with seta; plumose; distally inserted; on inner surface; surpassing to basal segment; without setules; without spinules; without spine. Third swimming legs basis without seta; without setules; without spinules; without spine. Third swimming legs endopod 3-segmented. Endopod 1 with seta; restricted; one on inner surface; without spine; without setules; without spinules; absence of Schmeil’s organ. Endopod 2 with seta; restricted; two on inner surface; plumose; without spine; without setules; with spinules; as a patch; single; proximally inserted; at anterior surface; absence of Schmeil’s organ. Endopod 3 with seta; plumose; plurimarginal; two on inner surface; two on outer surface; three on terminal surface; without spine; without setules; with spinules; as a row; distally inserted; double; at posterior surface; absence of Schmeil’s organ. Third swimming legs exopod 1 with seta; restricted; plumose; one on inner surface; with spine; 1; stout; not reaching to the distal-third of the exopod 2; serrated; equally; on inner surface, or on outer surface; with setules; as a row; single; continuously; on inner surface; without spinules; absence of Schmeil’s organ. Exopod 2 with seta; plumose; restricted; one on inner surface; with spine; 1; stout; not reaching out to exopod 3; serrated; on inner side, or on outer side; equally; without setules; without spinules; absence of Schmeil’s organ. Exopod 3 without setules; without spinules; with seta; plumose; unrestricted; three on inner surface; two on terminal surface; with spine; 2; unequal size; first no longer 2x than origin segment; stout; serrated; on inner side, or on outer side; equally; second longer 2x than origin segment; slender; serrated; on outer side; with ornamentation on non-serrated side; of setules; absence of Schmeil’s organ. **Fourth swimming legs**. Symmetrical; biramous. Intercoxal plate without sensilla. Praecoxa present. Coxa with seta; distally inserted; on inner margin; reaching out to endopod 1; without spinules; setules absent. Basis with seta; one; medially inserted; on posterior surface; smaller than the original segment; without setules; without spinules; without spine. Fourth swimming legs endopod 3-segmented. Endopod 1 with seta; one; restricted; on inner surface; without spine; without setules; without spinules; absence of Schmeil’s organ. Endopod 2 with seta; restricted; two on inner side; without spine; with setules; as a row; single; continuously; on outer surface; without spinules; absence of Schmeil’s organ. Endopod 3 with seta; unrestricted; two on inner surface; two on outer surface; three on distal surface; without spine; without setules; with spinules; as a row; double; distally inserted; at anterior surface; absence of Schmeil’s organ. Fourth swimming legs exopod 1 with seta; restricted; one on inner surface; with spine; 1; stout; not reaching out to distal-third of the exopod 2; serrated; on inner side, or on outer side; equally; with setules; as a row; single; continuously; on inner surface; without spinules; absence of Schmeil’s organ. Exopod 2 with seta; restricted; one on inner surface; with spine; 1; stout; not reaching the end of exopod 3; serrated; on inner side, or on outer side; equally; with setules; as a row; single; continuously; on inner surface; without spinules; absence of Schmeil’s organ. Exopod 3 without setules; with spinules; as a row; single; distally inserted; at anterior surface; with seta; unrestricted; three on inner surface; two on distal surface; with spine; 2; unequal size; first no longer 2x than origin segment; stout; serrated; on inner side, or on outer side; equally; second longer 2x than origin segment; slender; serrated; on outer side; without ornamentation on non-serrated side; absence of Schmeil’s organ.

##### Fifth swimming legs features

Asymmetrical. Fifth swimming leg intercoxal plate with length not equal or greater than width on 1.5x; with irregular proximal margin; discontinuous to; the anterior margin of the left coxa, or the anterior margin of the right coxa; posterior sensilla on the right lateral absent. **Fifth left swimming leg**. Fifth left swimming leg biramous; leg reaching first right exopod segment; proximally. Fifth left swimming leg praecoxa present; rudimentary; separated from the coxae; without ornamentation. Fifth left swimming leg coxa concave inner side; without teeth-like structures; with process; conical; on posterior surface; outer side; distally inserted; not projecting over basis; with sensilla; stout; triangular; at apex; no longer 2x than insertion basis; without swelling; without seta; without spinules. Fifth left swimming leg basis sub-cylindrical; unequal size between inner and outer side; shorter outer than inner side; with concave inner side; rounded internal proximal expansion present; without outgrowth; without groove; absence of protuberance; with seta; outerly inserted; no longer 2x than origin segment; absence of minutely granular. Fifth left swimming leg endopod segments 1 and 2 fused; segments 2 and 3 fused; 1-segmented; stout; separated from the basis; ornamented; on inner side; with spinules; more than four elements; as a row; terminally; row of setules absent; without seta. Fifth left swimming leg exopod segments 1 and 2 separated; segments 2 and 3 fused; 2-segmented; stout; separated from the basis. Fifth left swimming leg exopod 1 sub-cylindrical; longer than broad; unequal size between inner and outer side; shorter inner than outer side; concave inner side; convex outer side; without swelling; without marginal extension; with process; single; proximal; posteriorly; with lobe; single; circular; medially inserted; on inner side; covered; by setules; without outer spine; absence seta. Fifth left swimming leg exopod 2 sub-triangular; longer than broad; equal size between inner and outer side; disform inner side; with rectilinear outer side; setulose pad present; prominently rounded; proximally; on inner side; inflated medial region absent; distal process present; digitiform; denticulate; not bicuspidate; not innerly directed; with seta; spiniform; not ornamented by spinules; not surpassing the distal-point of the segment; without outer spine; terminal claw absent.

##### Fifth right swimming leg

Biramous. Fifth right swimming leg praecoxa present; separated from the coxae; without ornamentation. Fifth right swimming leg coxa convex inner side; without teeth-like structures; with process; rounded; distally inserted; on posterior surface; closest to the outer rim; projecting over basis; beyond the first third; until the medial surface; without triangular protuberance innerly; with sensilla; slender; at apex; no longer 2x than basal insertion; without marginal extension; without seta; without spinules. Fifth right swimming leg basis cylindrical; unequal size between inner and outer side; shorter outer than inner side; convex inner side; tumescence absent; without protuberance; absence of distinct minutely granular; additional inner process absent; with posterior groove; deep; obliquely; not reaching the endopodal lobe; ornamented; with tubercles; throughout of the outer border; with seta; outerly inserted; on anterior surface; no longer 2x than origin segment; posterior protrusion present; distal process absent. Fifth right swimming leg with endopodite present; separated from the basis; on anterior surface; ancestral segments 1 and 2 fused; ancestral segments 2 and 3 fused; 1-segmented; stout; ornamented; with spinules; as a row; on inner side; sub-terminally; without seta. Fifth right swimming leg exopod segments 1 and 2 separated; segments 2 and 3 fused; 2-segmented; stout; separated from the basis. Fifth right swimming leg exopod 1 trapezium; longer than broad; nearly 1.25 times; unequal size between both sides; shorter inner than outer side; rectilinear inner side; rectilinear outer side; with marginal extension; sub-triangular; distally inserted; at outer rim; spinules absent; without process; without outer spine; without seta; internal prominence absent; lamella on posterior surface absent. Fifth right swimming leg exopod 2 elliptical; longer than broad; nearly 2 times; equal size between both sides; disform inner side; convex outer side; with posterior proximal swelling; inner-posterior process absent; without marginal expansion; curved ridge on distal posterior surface present; chitinous knobs absent; with outer spine; inserted sub-distally; rectilinear; ornamented innerly; by spinules; as a row; not ornamented outerly; sharp tip; without apparent curve; lesser than the length of the exopod 2; beyond to 2 times its size; 3x; sensilla absent; terminal claw present; equal or longer 1.5 times than insertion segment; sclerotized; arched; inward; with conspicuous curve; proximally; ornamented innerly; by spinules; as a row; partially on extension; medially, or distally; ornamented outerly; sharp tip; not curved tip; without medial constriction; hyaline process absent.

##### FEMALE

Body longer and wider than male; Female body 1135 micrometers excluding caudal setae. Widest at first metasome segment. Distal margin of the prosomal segments without one line of setules at posterior margin. Prosome segments without spinules at prosomal segments. Fourth metasome segment absence of dorsal protuberance. Fourth and fifth metasome segments fused; partially; on dorsal surface. Limit between fourth and fifth metasome segments not ornamented. **Fifth metasome segment**. Fifth metasome segment ornamented with spinules; as a row; single; incomplete; same size; partially over limit; dorsally; without sensilla; with epimeral plates. Epimeral plates asymmetrical. Right epimeral plates prominent, as projections; thinner than the left; one posterior-dorsally directed; not reaching half length of the genital segment; with sensilla at the apex; dorsal-posterior sensilla present; slender; ornamented; with spinules; as a row; innerly; on dorsal surface. Left epimeral plate without expansion.

##### Urosome

3-segmented. **Genital double-somite**. Asymmetrical in dorsal view; longer than broad; longer than other urosomites combined; dorsal suture at mid-length absent; not covered by spinules; with swelling; rounded; unequal size; greater right than left; anteriorly; with sensillae; on both sides; one; stout; with robust apex; at left lateral; not on lobular base; anteriorly; one; stout; at right lateral; not on lobular base; anteriorly; with robust apex; of equal size between then; lateral protuberance absent; with right posterior rim expanded; over next segment; without slender sensilla on each posterior rim; without posterior-dorsal process. Genital double-somite opercular pad present; broader than longer; symmetrical; not development laterally; expanded posteriorly; covering partially; single gonoporal slit; located ventrally; with arthrodial membrane; inserted anteriorly; post-genital process present; single; disto-ventral tumescence absent; ventral vertical folds absent; dorsal sensilla absent. Second urosome segment without ventral fusion to anal segment; right distal process absent. Caudal rami patch of setules on outer surface absent.

##### Appendices features

Rostrum basal process absent. **Antennules**. Symmetrical. Right antennule surpassing to genital double-segment; not extending beyond caudal rami; ornamentation pattern equals to male left antennule; fully.

##### Fifth swimming legs

Symmetrical; Fifth swimming legs biramous. Fifth swimming legs intercoxal plate longer than wide; separated from the legs. Fifth swimming legs praecoxa with sclerite praecoxal; separated from the coxae; without ornamentation. Fifth swimming legs coxa with process; conical; at the outer rim; distally; sensilla present; stout; at apex; projecting over basal segment; no longer 2x than basal insertion; marginal extension absent; without swelling; without seta; without spinules. Fifth swimming legs basis sub-triangular; unequal size between inner and outer sides; shorter outer than inner side; with convex inner side; without proximal inner outgrowth; without groove; with distal extension; on posterior surface; with seta; outerly inserted; on posterior surface; no longer 2x than origin segment. Fifth swimming legs endopod segments 1 and 2 separated; segments 2 and 3 fused; 2-segmented; with complete suture; stout; separated from the basis; ornamentation on segment 2; with spinules; as a row; single; non-oblique; sub-terminally; at anterior surface; with seta; double; one medially; on posterior surface; rectilinear; one distally; on posterior surface; rectilinear; of unequal size; distal seta longer than medial seta. Fifth swimming legs exopod segments 1 and 2 separated; segments 2 and 3 separated; 3-segmented; separated from the basis. Fifth swimming legs exopod 1 sub-cylindrical; longer than wide; longer or equal than 2 times; with unequal size between inner and outer side; shorter inner than outer side; with convex inner side; with rectilinear outer side; without swelling; without marginal extension; without posterior process; without spine; without seta. Fifth swimming legs exopod 2 sub-cylindrical; longer than broad; longer or equal than 2 times; without swelling; without marginal extension; without process; without lobe; with spine; inserted laterally; rectilinear; without ornamentation; sharp tip; equal size or larger than next segment; without seta. Fifth swimming legs exopod 3 cylindrical; longer than wide; without swelling; without process; without lobe; without spine; with seta; double; inserted terminally; unequal size between them; outer seta smaller than inner; nearly 3 times; outer seta not ornamented by setules; without ornamentation; presence of terminal claw; sclerotized; rectilinear; with ornamentation; of denticles; as a row; on surface partially; at medial region; concave outer side; with ornamentation; of denticles; as a row; on surface partially; at medial region; blunt tip; 6 times longer than origin segment.

##### Distribution records

###### BRAZIL

**Maranhão**: Açú Lake, Mearim River, 03°50’S, 44°55’W and estuary of the Coqueiro River (Reid, 1987; Reid & Turner, 1988). **Sergipe**: Betume, near to Neápolis, in the São Francisco River Basin, 10°19’S, 36°35’W (Reid, 1987).

##### Habitat

Habitat in estuary region, and freshwaters: shallow lakes, and rivers.

##### Remarks

Reid (1987) presented the species to science from individuals collected in a lacustrine system in Maranhão, Brazil, registering for Sergipe, Brazil, in the basin system of the São Francisco River, latterly. The type-material of the species was located in the Museum of Zoology of USP, preserved in alcohol, and on slides. The holotype was located on slide and additionally were examined paratypes on slide, all stored in the same biological collection.

In the original description, the basis of the proposal in Dussart (1985) for subgenera in *Notodiaptomus* was considered and for indication of the intrageneric relationships of the species, from which specific analyses were presented. The observation in Reid (1987) for fourth metasome without spinules row on female of *N. brandorffi* was here corroborated as a distinctive attribute from the *N. spinuliferus.* For the dorsal conical process on the fourth pediger (in this effort treated as dorsal protuberance on fourth metasome segment), the sustained differentiations for the presence of the trait in *N. carteri*, and *N. isabelae* were confirmed here, but for *N. deitersi* the attribute is nonexistent, such as in *N. brandorffi*, and not useful for distinguishing both species.

Effectively, *N. brandorffi* and *N. deitersi* can be distinguished by female fifth swimming legs endopod 2-segmented for the first species, the same as that differentiates it from *N. iheringi*. In the species of Reid (1987) the male fifth right swimming leg exopod 2 with outer spine inserted medially was present as a distinction in relation to *N. jatobensis*, and *N. anisitsi*. In this approach, the relevant congener proximity was with *N. paraensis*, morphologically distinct by the male of *N. brandorffi* having: (1) fifth left swimming length not surpassing first right exopod segment, reaching to segment proximally; (2) fifth left swimming length exopod 2 with spiniform seta not surpassing to distal-point segment; (3) fifth right swimming exopod 1 longer than broad. And female: (1) second urosome segment without ventral fusion to anal segment; and (2) caudal rami with setules patch outerly.

**(1)** *N. brandorffi* can be included in *Notodiaptomus* from 14 of the 18 morphological shareings suggested in the creation and amplification of *nordestinus*, such as genus (Wright, 1935, 1936, 1937; Kiefer, 1936, 1956). Among the most frequent convergents within the genus and present in the species are: (1) male fifth left swimming leg exopod 2 in digitiform, and with spiniform seta not surpassing to distal-point segment; (2) male fifth left swimming leg endopod 1-segment; (3) male fifth right swimming leg basis with posterior protrusion; (4) male fifth right swimming leg exopod 1 longer than broad; and (5) male fifth right swimming leg exopod with outer spine inserted sub-distally.

#### Notodiaptomus cannarensis Alonso, Santos-Silva & Jaume, 2017

##### Synonymy

*Notodiaptomus cannarensis* Alonso, Santos-Silva & Jaume, 2017: 59–71, figs. 1–5.

##### Type locality

Mazar Reservoir, Cañar Province; southern Ecuador (2°35’53.08s S; 78°37m 32.16s W).

##### Type material

Holotype: male. Paratypes: 10 males, and 10 females. Both preserved in formalin vial, entire, and registered under same registration number [MECN-SI-Cal-0001], collected by Verónica Ordóñez, IV.2013, and deposited in the Museo Ecuatoriano de Ciencias Naturales of the Instituto Nacional del Biodiversidad, Quito, Ecuador [MECN).

##### Material examined

Topotype: 2 males, and 3 females, entire, in alcohol, from the zooplanktonic collection of the Plankton Laboratory, Instituto Nacional de Pesquisas da Amazônia (INPA), Brazil. 1 male (INPA-COP009, slides a-h) and 1 female (INPA-COP010, slides a-h) were selected to be dissection on eight slides each and deposited in the Zoological Collection of the INPA, Brazil.

##### Diagnosis

**(1)** male right epimeral plate with sensilla at the apex of projection and other medially; **(2)** fourth urosome segment with sub-conical blunt dorsal-lateral process; **(3)** male right antennule actual segment 6 with vestigial seta; male right antennule actual segment 13 with modified seta perpendicular to antennule direction; **(4)** male right antennule actual segment 22 with five setae; **(5)** antenna endopod actual segment 1 without spinules; **(6)** antenna exopod actual 8-segmented; **(7)** mandible endopod 2 with ten setae; **(8)** mandible exopod 5-segmented; **(9)** maxillule exopod 1 without setules; **(10)** maxilla 6-segmented; **(11)** maxilliped praecoxa separated from the coxa; **(12)** male fifth left swimming leg coxa rounded; **(13)** male fifth left swimming leg basis sub-triangular; **(14)** male fifth left swimming leg basis; **(15)** female urosome 2-segmented; **(16)** female genital double-somite with rounded expansion, like an avoid ventrolateral hypertrophy, inserted posteriorly.

##### Redescription

###### MALE

Body 1198 micrometers excluding caudal setae. Male body smaller and slenderer than female. Nerve axons myelinated. Prosome 5-segmented; widest at first metasome segment; without one line of setules at posterior margin; without spinules at segments. Cephalosome anterior margin rounded; with dorsal suture; incomplete; separate from first metasome segment. First metasome segment without sensilla. Second metasome segment without sensilla. Third metasome segment without sensillae; non-ornamented posterior margin. Fourth metasome segment without sensillae; fused to fifth metasome; fourth metasome segment totally. Limit between fourth and fifth metasome segments without ornamentation. Fifth metasome segment without sensilla; Fifth metasome segment without ornamentation; Fifth metasome segment without dorsal conical process; with epimeral plates. Epimeral plates asymmetrical. Right epimeral plates prominent, as projections; one projection; posterior-laterally directed; reaching half or surpassing the length of genital segment; with sensilla; one at the apex of projection and other medially; without ornamentation. Left epimeral plate prominent, as projection; one projection; posterior-laterally directed; reaching half length of the genital segment; with sensillae; one at the apex of projection and other medially; without ornamentation.

##### Urosome

5-segmented; Urosome 5-free segments. Genital somite asymmetrical in dorsal view; with single aperture; located on left side; ventrolaterally on posterior rim; with sensillae; on both sides; one; at left lateral; posteriorly; one; at right rim; posteriorly; of equal size between then. Third urosome segment without spinules; without external seta. Fourth urosome segment without spinules; with sub-conical blunt dorsal-lateral process; bearing sensilla on the apex. Anal segment absence of dorsal sensillae; presence of operculum; convex; not covering the anal aperture fully. Caudal rami asymmetrical; separated from anal segment; longer than wide; with setules; continuous on; inner side, or outer side; each ramus bearing 6 caudal setae; 5 marginals; plumose; and 1 internal dorsally; plumose; not reticulated main axis; outermost seta with outer spiniform process absent.

##### Appendices features

Rostrum symmetrical; separated from dorsal cephalic shield; by complete suture; sensillae present; one pair; anteriorly inserted on surface tegument; with rostral filament; double; paired; extended; into point; with basal process; in ventral view, rounded on left side; without a smaller basal expansion on the right side.

##### Antennules

Asymmetrical. **Right antennules**. Uniramous; right antennule surpassing to genital segment; right antennule not extending beyond caudal rami.

Right antennule ancestral segment I and II separated. Ancestral segment II and III fused. Ancestral segment III and IV fused. Ancestral segment IV and V separated. Ancestral segment V and VI separated. Ancestral segment VI and VII separated. Ancestral segment VII and VIII separated. Ancestral segment VIII and IX separated. Ancestral segment IX and X separated. Ancestral segment X and XI separated. Ancestral segment XI and XII separated. Ancestral segment XII and XIII separated. Ancestral segment XIII and XIV separated. Ancestral segment XIV and XV separated. Ancestral segment XV and XVI separated. Ancestral segment XVI and XVII separated. Ancestral segment XVII and XVIII separated. Ancestral segment XVIII and XIX separated. Ancestral segment XIX and XX separated. Ancestral segment XX and XXI separated. Ancestral segment XXI and XXII fused. Ancestral segment XXII and XXIII fused. Ancestral segment XXIII and XXIV separated. Ancestral segment XXIV and XXV fused. Ancestral segment XXV and XXVI separated. Ancestral segment XXVI and XXVII separated. Ancestral segment XXVII and XXVIII fused.

Right antennule actual 22-segmented; geniculated; between the segment 18 and segment 19; with swollen and modified region; formed by 5 segments; between 13 and 17 segments. Actual segment 1 with seta; one element; straight; none larger than segment; without spinules; without vestigial seta; without conical seta; without modified seta; without spinous process; with aesthetasc; one element. Actual segment 2 with seta; three elements; of unequal size; straight; none larger than segment; without spinules; with vestigial seta; one element; without conical seta; without modified seta; without spinous process; with aesthetasc; one element. Actual segment 3 with seta; one element; one larger than segment; surpassing to distal margin; beyond three sequential segments; straight; sharp apex; without spinules; with vestigial seta; one element; without conical seta; without modified seta; without spinous process; with aesthetasc. Actual segment 4 with seta; one element; one larger than segment; surpassing to distal margin; straight; not beyond three sequential segments; without spinules; without vestigial seta; without conical seta; without modified seta; without spinous process; without aesthetasc. Actual segment 5 with seta; one element; straight; one larger than segment; surpassing to distal margin; not beyond three sequential segments; without spinules; with vestigial seta; one element; without conical seta; without modified seta; without spinous process; with aesthetasc; one element. Actual segment 6 with seta; one element; none larger than segment; straight; without spinules; with vestigial seta; 1 element; without conical seta; without modified seta; without spinous process; without aesthetasc. Actual segment 7 with seta; one element; straight; one larger than segment; surpassing to distal margin; beyond three sequential segments; sharp apex; without spinules; without vestigial seta; without conical seta; without modified seta; without spinous process; with aesthetasc; one element. Actual segment 8 with seta; one element; straight; none larger than segment; without spinules; without vestigial seta; with conical seta; one element; not reaching to middle-point of the sequent segment; without modified seta; without spinous process; without aesthetasc. Actual segment 9 with seta; two elements; of unequal size; straight; one larger than segment; surpassing to distal margin; beyond three sequential segments; sharp apex; without spinules; without vestigial seta; without conical seta; without modified seta; without spinous process; with aesthetasc; one element. Actual segment 10 with seta; one element; straight; none larger than segment; without spinules; without vestigial seta; without conical seta; with modified seta; presenting blunt apex; slender form; surpassing to distal margin; beyond of the sequential segment; perpendicular to antennule direction; without spinous process; without aesthetasc. Actual segment 11 with seta; one element; straight; one larger than segment; surpassing to distal margin; not beyond three sequential segments; without spinules; without vestigial seta; without conical seta; with modified seta; slender form; presenting blunt apex; surpassing to distal margin; beyond of the sequential segment; perpendicular to antennule direction; shorter length than homologous of actual segment 13; without spinous process; without aesthetasc. Actual segment 12 with seta; one element; straight; one larger than segment; surpassing to distal margin; not beyond three sequential segments; without spinules; without vestigial seta; with conical seta; one element; not smaller than to segment 8; without modified seta; without spinous process; with aesthetasc; one element; absent internal perpendicular fission. Actual segment 13 with seta; one element; straight; one larger than segment; surpassing to distal margin; not beyond three sequential segments; without spinules; without vestigial seta; without conical seta; with modified seta; stout form; surpassing to distal margin; to the middle-point of the sequence segment; perpendicular to antennule direction; presenting bifid apex; without spinous process; with aesthetasc; one element. Actual segment 14 with seta; two elements; of unequal size; straight; one larger than segment; surpassing to distal margin; beyond three sequential segments; blunt apex; without spinules; without vestigial seta; without conical seta; without modified seta; without spinous process; with aesthetasc; one element. Actual segment 15 with seta; two elements; of unequal size; straight; not bifidform; none larger than segment; without spinules; without vestigial seta; without conical seta; without modified seta; with spinous process; on outer margin; surpassing distal margin; with aesthetasc; one element. Actual segment 16 with seta; two elements; of unequal size; plumose; one larger than segment; surpassing to distal margin; not beyond three sequential segments; not bifidform; without spinules; without vestigial seta; without conical seta; without modified seta; with spinous process; on outer margin; surpassing distal margin; unequal size to process on preceding segment; with aesthetasc; one element. Actual segment 17 with seta; one element; straight; none larger than segment; bifidform; without spinules; without vestigial seta; without conical seta; with modified seta; one element; stout form; surpassing to distal margin; not beyond of the sequential segment; parallel to antennule direction; without spinous process; without aesthetasc. Actual segment 18 with seta; one element; straight; none larger than segment; without spinules; without vestigial seta; without conical seta; with modified seta; one element; stout form; surpassing distal margin; parallel to antennule direction; without spinous process; without aesthetasc. Actual segment 19 with seta; two elements; of unequal size; plumose; none larger than segment; without spinules; without vestigial seta; without conical seta; with modified seta; two elements; stout form; at least one bifid form; surpassing distal margin; parallel to antennule direction; without spinous process; with aesthetasc; one element. Actual segment 20 with seta; four elements; of unequal size; straight; one larger than segment; surpassing to distal margin; beyond three sequential segments; without spinules; without vestigial seta; without conical seta; without modified seta; without spinous process; without aesthetasc. Actual segment 21 with seta; two elements; of equal size; plumose; one larger than segment; surpassing to distal margin; greater 3x than original segment; without spinules; without vestigial seta; without conical seta; without modified seta; without spinous process; without aesthetasc. Actual segment 22 with seta; five elements; of equal size; one larger than segment; plumose; surpassing to distal margin; greater 3x than original segment; without spinules; without vestigial seta; without conical seta; without modified seta; without spinous process; with aesthetasc; one element.

##### Left antennules

Uniramous; Left antennule surpassing to prosome; Left antennule not extending beyond caudal rami. Ancestral segment I and II separated. Ancestral segment II and III fused. Ancestral segment III and IV fused. Ancestral segment IV and V separated. Ancestral segment V and VI separated. Ancestral segment VI and VII separated. Ancestral segment VII and VIII separated. Ancestral segment VIII and IX separated. Ancestral segment IX and X separated. Ancestral segment X and XI separated. Ancestral segment XI and XII separated. Ancestral segment XII and XIII separated. Ancestral segment XIII and XIV separated. Ancestral segment XIV and XV separated. Ancestral segment XV and XVI separated. Ancestral segment XVI and XVII separated. Ancestral segment XVII and XVIII separated. Ancestral segment XVIII and XIX separated. Ancestral segment XIX and XX separated. Ancestral segment XX and XXI separated. Ancestral segment XXI and XXII separated. Ancestral segment XXII and XXIII separated. Ancestral segment XXIII and XXIV separated. Ancestral segment XXIV and XXV separated. Ancestral segment XXV and XXVI separated. Ancestral segment XXVI and XXVII separated. Ancestral segment XXVII and XXVIII fused.

Left antennule actual 25-segmented; not-geniculated. Actual segment 1 with seta; one element; none larger than segment; straight; without spinules; without vestigial seta; without conical seta; without modified seta; without spinous process; with aesthetasc; one element. Actual segment 2 with seta; three elements; of equal size; none larger than segment; straight; without spinules; with vestigial seta; one element; without conical seta; without modified seta; without spinous process; with aesthetasc; one element. Actual segment 3 with seta; one element; one larger than segment; straight; surpassing to distal margin; beyond three sequential segments; without spinules; with vestigial seta; one element; without conical seta; without modified seta; without spinous process; with aesthetasc; one element. Actual segment 4 with seta; one element; none larger than segment; straight; without spinules; without vestigial seta; without conical seta; without modified seta; without spinous process; without aesthetasc. Actual segment 5 with seta; one element; one larger than segment; straight; surpassing to distal margin; not beyond three sequential segments; without spinules; with vestigial seta; one element; without conical seta; without modified seta; without spinous process; with aesthetasc; one element. Actual segment 6 with seta; one element; none larger than segment; straight; without spinules; without vestigial seta; without conical seta; without modified seta; without spinous process; without aesthetasc. Actual segment 7 with seta; one element; one larger than segment; straight; surpassing to distal margin; beyond three sequential segments; without spinules; without vestigial seta; without conical seta; without modified seta; without spinous process; with aesthetasc; one element. Actual segment 8 with seta; one element; one larger than segment; straight; surpassing distal margin; without spinules; without vestigial seta; with conical seta; without modified seta; without spinous process; without aesthetasc. Actual segment 9 with seta; two elements; of unequal size; one larger than segment; straight; surpassing to distal margin; beyond three sequential segments; without spinules; without vestigial seta; without conical seta; without modified seta; without spinous process; with aesthetasc; one element. Actual segment 10 with seta; one element; none larger than segment; straight; without spinules; without vestigial seta; without conical seta; without modified seta; without spinous process; without aesthetasc. Actual segment 11 with seta; one element; one larger than segment; straight; surpassing to distal margin; beyond three sequential segments; without spinules; without vestigial seta; without conical seta; without modified seta; without spinous process; without aesthetasc. Actual segment 12 with seta; one element; one larger than segment; straight; surpassing distal margin; without spinules; without vestigial seta; with conical seta; without modified seta; without spinous process; with aesthetasc; one element. Actual segment 13 with seta; one element; none elongated; straight; surpassing distal margin; without spinules; without vestigial seta; without conical seta; without modified seta; without spinous process; without aesthetasc. Actual segment 14 with seta; one element; elongated; straight; surpassing to distal margin; beyond three sequential segments; without spinules; without vestigial seta; without conical seta; without modified seta; without spinous process; with aesthetasc; one element. Actual segment 15 with seta; one element; larger than segment; straight; surpassing to distal margin; not beyond three sequential segments; without spinules; without vestigial seta; without conical seta; without modified seta; without spinous process; without aesthetasc. Actual segment 16 with seta; one element; larger than segment; plumose; surpassing to distal margin; not beyond three sequential segments; without spinules; without vestigial seta; without conical seta; without modified seta; without spinous process; with aesthetasc; one element. Actual segment 17 with seta; one element; not larger than segment; straight; without spinules; without vestigial seta; without conical seta; without modified seta; without spinous process; without aesthetasc. Actual segment 18 with seta; one element; larger than segment; straight; surpassing to distal margin; beyond three sequential segments; without spinules; without vestigial seta; without conical seta; without modified seta; without spinous process; without aesthetasc. Actual segment 19 with seta; one element; not larger than segment; straight; surpassing distal margin; without spinules; without vestigial seta; without conical seta; without modified seta; without spinous process; with aesthetasc; one element. Actual segment 20 with seta; one element; not larger than segment; straight; surpassing distal margin; without spinules; without vestigial seta; without conical seta; without modified seta; without spinous process; without aesthetasc. Actual segment 21 with seta; one element; larger than segment; plumose; surpassing to distal margin; beyond three sequential segments; without spinules; without vestigial seta; without conical seta; without modified seta; without spinous process; without aesthetasc. Actual segment 22 with seta; two elements; of unequal size; one of them elongated; plumose; surpassing to distal margin; without spinules; without vestigial seta; without conical seta; without modified seta; without spinous process; without aesthetasc. Actual segment 23 with seta; two elements; of unequal size; one larger than segment; plumose; surpassing to distal margin; greater 3x than original segment; without spinules; without vestigial seta; without conical seta; without modified seta; without spinous process; without aesthetasc. Actual segment 24 with seta; two elements; of equal size; one larger than segment; plumose; surpassing to distal margin; greater 3x than original segment; without spinules; without vestigial seta; without conical seta; without modified seta; without spinous process; without aesthetasc. Actual segment 25 with seta; five elements; of equal size; elongated; plumose; surpassing to distal margin; 4 times larger than segment; without spinules; without vestigial seta; without conical seta; without modified seta; without spinous process; with aesthetasc; one element.

##### Antenna

Biramous. Antenna coxa separated from the basis; bearing seta; 1; on inner surface; at distal corner; reaching to the middle basis. Antenna basis (fusion) separated from the endopodal segment; bearing seta; 2; on inner surface; at distal corner. Endopodal ancestral segment I and II separated. Ancestral segment II and III fused. Ancestral segment III and IV fused. Ancestral segment III and IV fully. Antenna endopod actual 2-segmented. Actual segment 1 not bilobate; with seta; two; on inner margin; without spinules; without pore. Actual segment 2 bilobate; with discontinuity on outer cuticle; not developed as a suture; inner lobe bearing 7 setae; distally; outer lobe bearing 6 setae; distally; with spinules; as a row; not obliquely; on outer surface. Antenna exopod ancestral segment I and II separated. Ancestral segment II and III fused. Ancestral segment III and IV fused. Ancestral segment IV and V separated. Ancestral segment V and VI separated. Ancestral segment VI and VII separated. Ancestral segment VII and VIII separated. Ancestral segment VIII and IX fused. Ancestral segment IX and X fused. Antenna exopod actual 8-segmented. Actual segment 1 single; elongated (width-length, equal or larger ratio 2:1); with seta; one; at inner surface. Actual segment 2 compound; elongated (larger width-length ratio 2:1); with seta; three; at inner surface. Actual segment 3 single; not elongated (lesser width-length ratio 2:1); with seta; one; at inner surface. Actual segment 4 single; not elongated (lesser width-length ratio 2:1); with seta; one; at inner surface. Actual segment 5 single; not elongated (lesser width-length ratio 2:1); with seta; one; at inner surface. Actual segment 6 single; not elongated (lesser width-length ratio 2:1); with seta; one; at inner surface. Actual segment 7 compound; elongated (larger or equal width-length ratio 2:1); with seta; one; at inner surface. Actual segment 8 not elongated (lesser width-length ratio 2:1); with 3 setae; at distal surface.

##### Oral features

**Mandible**. Coxal gnathobase sclerotized; with lobe; not prominent; on caudal margin; presence of cutting blade; with tooth-like prominence; two, distinctly; 1 acute; on caudal margin; and 1 triangular; on sub-caudal margin; without acute projection between the prominences; without additional spinules; with seta; 1; dorsally; on apical surface; without spinules. Mandible palps biramous; comprising the basis; with seta; four; differently inserted; first medially; not reaching to beyond the endopod 1; second distally; third distally; fourth distally; on inner margin; all with setulose ornamentation. Mandible endopod 2-segmented. Mandible endopod 1 with lobe; bearing seta; four; distally inserted; without spinules. Mandible endopod 2 without lobe; bearing setae; ten elements; distally inserted; with spinules; as a row; double. Mandible exopod 5-segmented. Mandible exopod 1 with seta; one element; distally; on inner margin. Mandible exopod 2 with seta; one element; distally; on inner side. Mandible exopod 3 with seta; one element; distally; on inner side. Mandible exopod 4 with setae; one element; on terminal region. Mandible exopod 5 with 3 setae. **Maxillule**. Birramous. Maxillule 3-segmented. Maxillule praecoxa with praecoxal arthrite; bearing spines; fifteen elements; ten marginally; plus, five sub-marginally; with spinules; as a patch; on sub-marginal surface. Maxillule coxa with coxal epipodite; with conspicuous outer lobe; bearing setae; nine elements; with coxal endite; elongated (larger or equal width-length ratio 2:1); bearing setae; four elements. Maxillule basis with basal endite; double; first proximal; elongated (larger width-length ratio 2:1; separated from basis; with setae; four elements; distally inserted; second distal; fused to basis; not elongated (lesser width-length ratio 2:1); with setae; four elements; distally inserted; with setules; as a row; on inner side; basal exite present; with setae; one element; on outer surface. Maxillule endopod 1-segmented. Endopod 1 bilobate; first proximal; with setae; four elements; second distal; with setae; five elements. Maxillule exopod 1-segmented. Exopod 1 with setae; six elements; without setules; spinules absent. **Maxilla**. Uniramous. Maxilla 6-segmented. Maxilla praecoxa fused to coxa; completely; with praecoxal endite; triple; first elongated endite (larger or equal width length ratio 2:1); proximally inserted; with seta; straight, or plumose; 1 straight; 4 plumose; with spine; single; without spinules; without setule; second elongated endite (larger or equal width length ratio 2:1); medially inserted; with seta; plumose; 3 plumose; without spine; without spinules; without setule; third elongated endite (larger or equal width length ratio 2:1); distally inserted; with seta; 2 straight; 1 plumose; absence of outer seta. Maxilla coxa with coxal endite; triple; first elongated endite (larger or equal width); proximally inserted; with seta; straight, or plumose; 2 straight; 1 plumose; without spine; without spinules; without setules; second elongated endite (larger or equal width); medially inserted; with seta; straight, or plumose; 2 straight; 1 plumose; without spine; without spinules; without setules; absence of outer seta; third not elongated (lesser width length ratio 2:1); distally inserted; with one seta; straight. Maxilla basis with basal endite; single; not elongated (lesser width-length ratio 2:1); with seta; straight; 1 straight; without spinules; absence of outer seta. Maxilla endopod 3-segmented. Maxilla endopod 1 with seta; 1 straight; without spine; without spinules; without setules. Maxilla endopod 2 with seta; 1 straight; without spine; without spinules; without setules. Maxilla endopod 3 with 3 setae; straight. **Maxilliped**. Uniramous; Maxilliped 8-segmented. Maxilliped praecoxa separated from the coxa; with praecoxal endite; not elongated (lesser width-length ratio 2:1); distally inserted; with seta; 1 plumose; without spinules; without setules. Maxilliped coxa with coxal endite; three coxal endite; first elongated (larger or equal width); proximally inserted; with seta; 2 plumose; without spinules; without setules; second not elongated (lesser width-length ratio 2:1); medially inserted; with seta; 3 plumose; without spinules; without setules; third elongated (larger or equal width length ratio 2:1); distally inserted; with seta; 4 plumose; none reaching to beyond of the basis; without spinules; without setules; with lobe; prominence; at inner distal angle; ornamented; with spinules; continuously on margin. Maxilliped basis without basal endite; with seta; 3 plumose; with spinules; as a row; single; on medial surface; with setules; as a row; single; on inner margin. Maxilliped endopod segment 6-segmented. Endopod 1 with seta; 2 plumose; on inner surface. Endopod 2 with seta; 3 plumose; on inner surface. Endopod 3 with seta; 2 plumose; on inner surface. Endopod 4 with seta; 2 plumose; on inner surface. Endopod 5 with seta; 2 plumose; on inner surface, or on outer surface; outer seta absent. Endopod 6 with seta; 4 plumose; on inner surface, or on outer surface.

##### Swimming legs features

**First swimming legs.** Symmetrical; biramous. First swimming legs intercoxal plate without seta. First swimming legs praecoxa absent. First swimming legs coxa with seta; one; straight; distally inserted; on inner surface; surpassing to first endopodal segment; with setules; two group; as a patch; on inner margin; and as a row; double; on anterior surface; outerly; without spinules; without spine. First swimming legs basis without seta; with setules; as a patch; single; on outer surface; without spinules; without spine. First swimming legs endopod 2-segmented. Endopod 1 with seta; straight; restricted; to inner surface; one element; without spine; with setules; as a row; single; continuously; on outer surface; without spinules; absence of Schmeil’s organ. Endopod 2 with seta; unrestricted; three on inner surface; one on outer surface; two on distal surface; straight; without spine; with setules; as a row; single; continuously; on outer surface; without spinules; absence of Schmeil’s organ. Endopod 3 absence. First swimming legs exopod 1 with seta; restricted; 1 on inner surface; with spine; 1; stout; smaller than original segment; serrated; on inner side; continuously; with setules; as a row; single; as a row; innerly. First swimming legs exopod 2 with seta; restricted; 1 on inner surface; straight; without spine; with setules; as a row; single; continuously; on inner margin, or on outer margin; without spinules. First swimming legs exopod 3 with setule; as a row; single; continuously; on outer surface; without spinules; with seta; unrestricted; 2 on inner surface; 2 on terminal surface; with spine; 2; unequal size; first no longer 2x than origin segment; stout; serrated; on inner side, or on outer side; equally; second longer 3x than origin segment; slender; serrated; on outer side; with ornamentation on non-serrated side; by setules. **Second swimming legs**. Symmetrical; Second swimming legs biramous. Second swimming legs intercoxal plate without seta. Second swimming legs praecoxa present; located laterally. Second swimming legs coxa with seta; straight; distally inserted; on inner surface; surpassing to basal segment; without setules; without spinules; without spine. Second swimming legs basis without seta; without setules; without spinules; without spine. Second swimming legs endopod 3-segmented. Endopod 1 with seta; straight; restricted; one on inner surface; without spine; with setules; as a row; single; continuously; on outer surface; without spinules; absence of Schmeil’s organ. Endopod 2 with seta; straight; unrestricted; two on inner surface; without spine; with setules; as a row; single; continuously; on outer side; without spinules; presence of Schmeil’s organ; on posterior surface. Endopod 3 with seta; straight; unrestricted; three on inner surface; two on outer surface; two on distal surface; without spine; without setules; with spinules; as a row; double; distally inserted; at anterior surface; absence of Schmeil’s organ. Second swimming legs exopod 1 with seta; restricted; one on inner surface; with spine; 1; stout; not reaching to distal-third of the exopod 2; serrated; on inner side, or on outer side; with setules; as a row; single; continuously; on inner side; without spinules; absence of Schmeil’s organ. Exopod 2 with seta; unrestricted; one on inner surface; with spine; 1; stout; not surpassing the exopod 3; serrated; on inner side, or on outer side; with setules; as a row; single; continuously; on inner surface; without spinules; absence of Schmeil’s organ. Exopod 3 with seta; plurimarginal; three on inner surface; two on terminal surface; with spine; 2; unequal size; first no longer 2x than origin segment; stout; serrated; on inner side, or on outer side; equally; second longer 2x than origin segment; slender; serrated; on outer side; with ornamentation on non-serrated side; of setules; setules on outer surface; as a row; single; continuously; on inner surface; with spinules; as a row; single; distally inserted; at anterior surface; absence of Schmeil’s organ. **Third swimming legs**. Symmetrical; Third swimming legs biramous. Third swimming legs intercoxal plate without seta. Third swimming legs praecoxa present; not laterally located. Third swimming legs coxa with seta; straight; distally inserted; on inner surface; surpassing to first endopodal segment; without setules; without spinules; without spine. Third swimming legs basis without seta; without setules; without spinules; without spine. Third swimming legs endopod 3-segmented. Endopod 1 with seta; restricted; one on inner surface; without spine; without setules; without spinules; absence of Schmeil’s organ. Endopod 2 with seta; restricted; two on inner surface; straight; without spine; without setules; without spinules; absence of Schmeil’s organ. Endopod 3 with seta; straight; plurimarginal; two on inner surface; two on outer surface; three on terminal surface; without spine; without setules; with spinules; as a row; distally inserted; double; at anterior surface; absence of Schmeil’s organ. Third swimming legs exopod 1 with seta; restricted; straight; one on inner surface; with spine; 1; stout; not reaching to the distal-third of the exopod 2; serrated; equally; on inner surface, or on outer surface; with setules; as a row; single; continuously; on inner surface; without spinules; absence of Schmeil’s organ. Exopod 2 with seta; straight; restricted; one on inner surface; with spine; 1; stout; not reaching out to exopod 3; serrated; on inner side, or on outer side; equally; with setules; as a row; single; continuously; on inner side; without spinules; absence of Schmeil’s organ. Exopod 3 without setules; with spinules; as a row; single; distally inserted; at anterior surface; with seta; straight; unrestricted; three on inner surface; two on terminal surface; with spine; 2; unequal size; first no longer 2x than origin segment; stout; serrated; on inner side, or on outer side; equally; second longer 2x than origin segment; slender; serrated; on outer side; with ornamentation on non-serrated side; of setules; absence of Schmeil’s organ. **Fourth swimming legs**. Symmetrical; biramous. Intercoxal plate without sensilla. Praecoxa present. Coxa with seta; distally inserted; on inner margin; reaching out to endopod 1; without spinules; setules absent. Basis with seta; one; medially inserted; on posterior surface; smaller than the original segment; without setules; without spinules; without spine. Fourth swimming legs endopod 3-segmented. Endopod 1 with seta; one; restricted; on inner surface; without spine; without setules; without spinules; absence of Schmeil’s organ. Endopod 2 with seta; restricted; two on inner side; without spine; with setules; as a row; single; continuously; on outer surface; without spinules; absence of Schmeil’s organ. Endopod 3 with seta; unrestricted; two on inner surface; two on outer surface; three on distal surface; without spine; without setules; with spinules; as a row; double; distally inserted; at anterior surface; absence of Schmeil’s organ. Fourth swimming legs exopod 1 with seta; restricted; one on inner surface; with spine; 1; stout; not reaching out to distal-third of the exopod 2; serrated; on inner side, or on outer side; equally; with setules; as a row; single; continuously; on inner surface; without spinules; absence of Schmeil’s organ. Exopod 2 with seta; restricted; one on inner surface; with spine; 1; stout; not reaching the end of exopod 3; serrated; on inner side, or on outer side; equally; with setules; as a row; single; continuously; on inner surface; without spinules; absence of Schmeil’s organ. Exopod 3 without setules; with spinules; as a row; single; distally inserted; at anterior surface; with seta; unrestricted; three on inner surface; two on distal surface; with spine; 2; unequal size; first no longer 2x than origin segment; stout; serrated; on inner side, or on outer side; equally; second longer 2x than origin segment; slender; serrated; on outer side; without ornamentation on non-serrated side; absence of Schmeil’s organ.

##### Fifth swimming legs features

Asymmetrical. Fifth swimming leg intercoxal plate with length not equal or greater than width on 1.5x; with irregular proximal margin; continuous to; the anterior margin of the left coxa, or the anterior margin of the right coxa; posterior sensilla on the right lateral absent. **Fifth left swimming leg**. Fifth left swimming leg biramous; leg surpassing first right exopod segment. Fifth left swimming leg praecoxa present; rudimentary; fused to coxae; completely; without ornamentation. Fifth left swimming leg coxa concave inner side; without teeth-like structures; with process; triangulated; on posterior surface; outer side; distally inserted; not projecting over basis; with sensilla; stout; rounded; at apex; no longer 2x than insertion basis; without swelling; without seta; without spinules. Fifth left swimming leg basis sub-triangular; unequal size between inner and outer side; shorter inner than outer side; with convex inner side; rounded internal proximal expansion absent; without outgrowth; without groove; absence of protuberance; with seta; outerly inserted; no longer 2x than origin segment; absence of minutely granular. Fifth left swimming leg endopod segments 1 and 2 fused; segments 2 and 3 fused; 1-segmented; stout; separated from the basis; ornamented; on inner side; with spinules; more than four elements; as a row; terminally; row of setules absent; without seta. Fifth left swimming leg exopod segments 1 and 2 separated; segments 2 and 3 fused; 2-segmented; stout; separated from the basis. Fifth left swimming leg exopod 1 sub-cylindrical; longer than broad; equal size between inner and outer side; concave inner side; convex outer side; without swelling; without marginal extension; without process; with lobe; single; semicircular; medially inserted; on inner side; covered; by tubercles; without outer spine; absence seta. Fifth left swimming leg exopod 2 sub-triangular; longer than broad; equal size between inner and outer side; disform inner side; with rectilinear outer side; setulose pad present; not prominently rounded; medially; on inner side; inflated medial region absent; distal process present; acute-form; denticulate; not bicuspidate; with transverse row of denticles; plus oblique row of 5 denticles; at anterior surface; not innerly directed; with seta; spiniform; not ornamented by spinules; not surpassing the distal-point of the segment; without outer spine; terminal claw absent.

##### Fifth right swimming leg

Biramous. Fifth right swimming leg praecoxa present; fused to coxae; completely; without ornamentation. Fifth right swimming leg coxa convex inner side; without teeth-like structures; with process; conical; distally inserted; on posterior surface; closest to the outer rim; not projecting over basis; without triangular protuberance innerly; with sensilla; slender; at apex; no longer 2x than basal insertion; without marginal extension; without seta; without spinules. Fifth right swimming leg basis cylindrical; unequal size between inner and outer side; shorter outer than inner side; rectilinear inner side; tumescence absent; without protuberance; absence of distinct minutely granular; additional inner process absent; without posterior groove; with seta; outerly inserted; on anterior surface; no longer 2x than origin segment; posterior protrusion absent; distal process absent. Fifth right swimming leg with endopodite present; separated from the basis; on anterior surface; ancestral segments 1 and 2 fused; ancestral segments 2 and 3 fused; 1-segmented; stout; ornamented; with setules; as a row; on inner side; terminally; without seta. Fifth right swimming leg exopod segments 1 and 2 separated; segments 2 and 3 fused; 2-segmented; stout; separated from the basis. Fifth right swimming leg exopod 1 sub-cylindrical; broader than long; nearly 1.25 times; unequal size between both sides; shorter inner than outer side; convex inner side; rectilinear outer side; without marginal extension; spinules absent; without process; without outer spine; without seta; internal prominence present; acute; lamella on posterior surface present; trapezoidal form; surpassing to margin; outerly. Fifth right swimming leg exopod 2 elliptical; longer than broad; nearly 2 times; equal size between both sides; uniform inner side; convex outer side; without posterior proximal swelling; inner-posterior process absent; without marginal expansion; curved ridge on distal posterior surface absent; chitinous knobs absent; with outer spine; inserted premedially; arched; internally directed; ornamented innerly; by spinules; as a row; ornamented outerly; by spinules; as a row; sharp tip; with apparent curve; outerly directed; lesser than the length of the exopod 2; until to 2 times its size; 1.5x; sensilla absent; terminal claw present; equal or longer 1.5 times than insertion segment; sclerotized; arched; inward; with conspicuous curve; proximally; ornamented innerly; by spinules; as a row; partially on extension; distally; not ornamented outerly; sharp tip; not curved tip; without medial constriction; hyaline process present; on proximal third; outerly or innerly.

##### FEMALE

Body longer and wider than male; Female body 1352 micrometers excluding caudal setae. Widest at first metasome segment. Distal margin of the prosomal segments without one line of setules at posterior margin. Prosome segments without spinules at prosomal segments. Fourth metasome segment absence of dorsal protuberance. Fourth and fifth metasome segments fused; partially; on dorsal surface. Limit between fourth and fifth metasome segments without ornamentation. **Fifth metasome segment**. Fifth metasome segment without sensilla; with epimeral plates. Epimeral plates symmetrical. Right epimeral plates prominent, as projections; two posterior-dorsally directed; reaching half length of the genital segment; with sensilla at the apex; dorsal-posterior sensilla absent; without ornamentation. Left epimeral plate without expansion.

##### Urosome

2-segmented. **Genital double-somite**. Asymmetrical in dorsal view; longer than broad; longer than other urosomites combined; dorsal suture at mid-length absent; not covered by spinules; with expansion; rounded; like an ovoid ventrolateral hypertrophy; equal size; posteriorly; with sensillae; on both sides; one; stout; with robust apex; at left lateral; not on lobular base; medially; one; stout; at right lateral; not on lobular base; medially; with robust apex; of equal size between then; lateral protuberance absent; without right posterior rim expanded; without slender sensilla on each posterior rim; without posterior-dorsal process. Genital double-somite opercular pad present; broader than longer; symmetrical; development laterally; not expanded posteriorly; covering partially; double gonoporal slit; located ventrally; with arthrodial membrane; inserted anteriorly; post-genital process absent; disto-ventral tumescence absent; ventral vertical folds absent; dorsal sensilla absent. Second urosome segment without ventral fusion to anal segment; right distal process absent. Caudal rami patch of setules on outer surface present; patch of spinules on outer surface absent.

##### Appendices features

Rostrum basal process absent. **Antennules**. Symmetrical. Right antennule surpassing to genital double-segment; extending beyond caudal rami. Right antennule not exceeding the caudal setae. Right antennule ornamentation pattern equals to male left antennule; fully.

##### Fifth swimming legs

Symmetrical; Fifth swimming legs biramous. Fifth swimming legs intercoxal plate longer than wide; separated from the legs. Fifth swimming legs praecoxa with sclerite praecoxal; separated from the coxae; without ornamentation. Fifth swimming legs coxa without process; sensilla absent; marginal extension absent; without swelling; without seta; without spinules. Fifth swimming legs basis sub-triangular; unequal size between inner and outer sides; shorter outer than inner side; with convex inner side; without proximal inner outgrowth; with groove; not ornamented; with distal extension; on lateral surface; with seta; outerly inserted; on anterior surface; no longer 2x than origin segment. Fifth swimming legs endopod segments 1 and 2 fused; segments 2 and 3 fused; 1-segmented; stout; separated from the basis; absent discontinuity cuticle; with spinules; as a row; single; non-oblique; sub-terminally; at anterior surface; with seta; double; one medially; on posterior surface; rectilinear; one distally; on posterior surface; rectilinear; of unequal size; distal seta longer than medial seta. Fifth swimming legs exopod segments 1 and 2 separated; segments 2 and 3 separated; 3-segmented; separated from the basis. Fifth swimming legs exopod 1 sub-cylindrical; longer than wide; longer or equal than 2 times; with unequal size between inner and outer side; shorter inner than outer side; with convex inner side; with rectilinear outer side; without swelling; without marginal extension; without posterior process; without spine; without seta. Fifth swimming legs exopod 2 sub-cylindrical; longer than broad; longer or equal than 2 times; without swelling; without marginal extension; without process; without lobe; with spine; inserted laterally; rectilinear; without ornamentation; sharp tip; equal size or larger than next segment; without seta. Fifth swimming legs exopod 3 cylindrical; longer than wide; without swelling; without process; without lobe; without spine; with seta; double; inserted terminally; unequal size between them; outer seta smaller than inner; nearly 3 times; outer seta ornamented by setules; without ornamentation; presence of terminal claw; sclerotized; arched; internally directed; concave inner side; with ornamentation; of spinules; as a row; on surface partially; at medial region, or at distal region; rectilinear outer side; with ornamentation; of spinules; as a row; on surface partially; at medial region, or at distal region; blunt tip; 6 times longer than origin segment.

##### Distribution records

Restricted to type locality.

##### Habitat

Habitat in freshwaters: reservoir in the Andes.

##### Remarks

The taxon was presented from organisms from the Mazar Reservoir, Ecuador, 250 kilometers from the east coast of the Pacific Ocean. Until this moment it is characterized as endemic to the region, since its inauguration having not been registered its occurrence in other known records or type locality. Undoubtedly it is the most discrepant hypothesis for *Notodiaptomus* among its congeners, morphologically.

In the original description, the authors indicated the presence of processes on segments 10, 11 of the male right antennule. However, it is truly the modified setae frequent on these same segments for similar species in *Notodiaptomus.* For *N. cannarensis* both structures positioned perpendicular to the original segments and establish a variation of the type-species of the group. For the right A1 of the males examined, it was possible to observe variations to the ornamental pattern verified for the genus, such as the sharp apex of the elongated seta on segments 3, 7 and 9, variation to the pattern identified in the species of *Notodiaptomus*, which have this ornamentation with blunt apex.

Other variations are still possible to identify, for example, the presence of a vestigial seta on segment 6, treated in the original description as sensilla mistakenly. In *N. deitersi* the presence of vestigial setae are present only for segments 2, 3 and 5. Likewise, on segments 17 and 18 the presence of only 1 seta evidences new variation in the ornamental pattern of the male right antennule in *Notodiaptomus* (Santos-Silva *et al*., 2015). Similarly, the presence of a reduced seta on segment 19 and five setae in the last segment are variations when compared to the longer seta and the four setae in the last segment of the male N. *deitersi*, respectively.

For the male and female left antennule, we present on segments 2, 3 and 5 the presence of vestigial seta, absent in the original description, as well as the presence of aesthetasc on actual segment 3. As for the right A1, the last segment has five plumose setae. Both for this last described observation and for the presence of an aesthetasc on actual segment 3 represent variations to the ornamental pattern of *Notodiaptomus* as described in Santos-Silva *et al*. (1999), and Santos-Silva *et al*. (2015).

Another relevant point is for the report of the presence of dorsal-lateral protuberance on segment 3 of the male urosome. It was not possible to observe this condition in the specimens examined for this thesis, it must probably be a mistake in the original description, since the structure is present for actual segment 4 of the urosome. Additionally, from micrometric variation in the optical apparatus, we observe the “spine” at the apex of the conical protuberance with apparent internal enervation, which characterizes it as a truly sensory element (*sensu* Garm & Watling, 2013), not integumentary ornamentation as described originally. Therefore, it is correct to treat the element as a sensilla.

These and other variations identified in the reexamination of the organisms of the species, allow us to indicate a significant morphological distance with the genus *Notodiaptomus*. On the contrary, the intrinsic proximity to other Neotropical genus, such as *Tumeodiaptomus* Dussart 1979 and some species of *Rhacodiaptomus* Kiefer 1936 point to the need for a comparative analysis with these groups and their possible repositioning. Among the attributes presented in the original description and amplification of *Notodiaptomus*, the taxon has only the male, and female fifth swimming legs endopod 1-segmented, and presence of male left swimming leg exopod 2 with spiniform seta not surpassing to distal-point segment.

#### Notodiaptomus carteri (Lowndes, 1934)

##### Synonymy

*Diaptomus carteri* Lowndes, 1934: 89, 90, 91, 92, 93, 98–100, pl. 3, fig. 3a-d; Wright, 1937a: 76; 1938a: 562. *Notodiaptomus carteri* n. comb., Kiefer, 1936a: 197; 1956: 242; Ringuelet & Martínez de Ferrato, 1967: 411, 412, pl. 1, figs, 1–6; Brandorff, 1972: 43; 1976: 614, 616, fig. 2; Bowman, 1973: 199; Löffler, 1981: 15; Dussart & Defaye, 1983: 133, 136; Dussart, 1985b: 264, fig. 7C; Dussart & Frutos, 1986: 306; 1987: 244, 245, 246, pl. 1, figs. 2–9; Montú & Gloeden, 1986: 6, 82, fig. 25e-j; Reid, 1987: 377; Battistoni, 1995: 958; Rocha *et al*., 1995: 156; Santos-Silva, 1998: 206; Santos-Silva *et al*., 1999: 127; Bohrer & Araújo, 1999: 92, 94; Santos-Silva, 2008: 21–22; Perbiche-Neves *et al*., 2015: 38–42, fig. 29–30; Perbiche-Neves *et al*., 2020: 676-677, key to the Neotropical diaptomid, fig 21.6 D. *Notodiaptomus (Notodiaptomus) carteri*; Dussart, 1985a: 208.

##### Type locality

Not specified. In the original description were mentioned various places in pools, area of swamps, and flood water in the Makthlaiya, Paraguay.

##### Material examined

Non-type material: 1 male, and 1 female, entire in formalin solution (MZUSP 28390) from the Paraná River, Argentina. Perbiche-Neves coll., 29.I.2010. Additional material examined: 1 male, and 2 females, entire in alcohol, from the flood water (rice fields, 32° 04m 38s S, 52° 10m 09s W) in Rio Grande do Sul, Brazil, M. G. Bandeira coll., 15.V.2017; 1 male (INPA-COP011, slides a-h) and 1 female (INPA-COP012, slides a-h) were selected to be dissection on eight slides each and deposited in the Zoological Collection of the INPA, Brazil.

##### Diagnosis

**(1)** Male right antennule actual segment 12 with conical seta smaller than to those of the segment 8; **(2)** male fifth left swimming leg with length surpassing first right exopod segment; **(3)** male fifth left swimming leg coxa with sensilla on process no longer 2x than insertion basis; **(4)** male fifth left swimming leg basis without groove; **(5)** male fifth right swimming leg coxa with process projecting over basis not beyond the first third; **(6)** male fifth right swimming leg basis with groove latitudinally; **(7)** male fifth right swimming leg exopod 2 with inner-posterior process; **(8)** male fifth right swimming leg exopod 2 with outer spine arched outwardly; **(9)** male fifth right swimming leg exopod 2 with outer spine smaller 2x than the insertion segment; **(10)** male fifth right swimming leg exopod 2 with terminal claw not larger or equal that 1.5x than the insertion segment; **(11)** female genital double-somite with sensilla inserted on lobular base; **(12)** female fifth swimming lags endopod 2-segmented with incomplete suture.

##### Redescription

###### MALE

Body 1437 micrometers excluding caudal setae. Body smaller and slenderer than female. Nerve axons myelinated. Prosome 6-segmented; widest at first metasome segment; without one line of setules at posterior margin; without spinules at segments. Cephalosome anterior margin rounded; with dorsal suture; complete; separate from first metasome segment. First metasome segment without sensilla. Second metasome segment without sensilla. Third metasome segment without sensillae; non-ornamented posterior margin. Fourth metasome segment without sensillae; separated from the fifth metasome. Limit between fourth and fifth metasome segments without ornamentation. Fifth metasome segment with sensilla; 2 laterally; Fifth metasome segment equal size; Fifth metasome segment without ornamentation; Fifth metasome segment without dorsal conical process; with epimeral plates. Epimeral plates symmetrical. Right epimeral plates reduced, as rounded distal corner segment limit; with sensilla; at the apex of projection; without ornamentation.

##### Urosome

5-segmented; Urosome 5 - free segments. Genital somite symmetrical in dorsal view; with single aperture; located on left side; ventrolaterally on posterior rim; with sensillae; on both sides; one; at left lateral; posteriorly; one; at right rim; posteriorly; of equal size between then. Third urosome segment without spinules; without external seta. Fourth urosome segment without spinules; without sub-conical blunt dorsal-lateral process. Anal segment presence of dorsal sensillae; one on each side; medially inserted; presence of operculum; convex; covering the anal aperture fully. Caudal rami symmetrical; separated from anal segment; longer than wide; with setules; continuous on; inner side; each ramus bearing 6 caudal setae; 5 marginals; plumose; and 1 internal dorsally; straight; not reticulated main axis; outermost seta with outer spiniform process absent.

##### Appendices features

Rostrum symmetrical; separated from dorsal cephalic shield; by complete suture; sensillae present; one pair; anteriorly inserted on surface tegument; with rostral filament; double; paired; extended; into point; with basal process; in ventral view, rounded on left side; without a smaller basal expansion on the right side.

##### Antennules

Asymmetrical. **Right antennules**. Uniramous; right antennule surpassing to genital segment; right antennule not extending beyond caudal rami.

Right antennule ancestral segment I and II separated. Ancestral segment II and III fused. Ancestral segment III and IV fused. Ancestral segment IV and V separated. Ancestral segment V and VI separated. Ancestral segment VI and VII separated. Ancestral segment VII and VIII separated. Ancestral segment VIII and IX separated. Ancestral segment IX and X separated. Ancestral segment X and XI separated. Ancestral segment XI and XII separated. Ancestral segment XII and XIII separated. Ancestral segment XIII and XIV separated. Ancestral segment XIV and XV separated. Ancestral segment XV and XVI separated. Ancestral segment XVI and XVII separated. Ancestral segment XVII and XVIII separated. Ancestral segment XVIII and XIX separated. Ancestral segment XIX and XX separated. Ancestral segment XX and XXI separated. Ancestral segment XXI and XXII fused. Ancestral segment XXII and XXIII fused. Ancestral segment XXIII and XXIV separated. Ancestral segment XXIV and XXV fused. Ancestral segment XXV and XXVI separated. Ancestral segment XXVI and XXVII separated. Ancestral segment XXVII and XXVIII fused.

Right antennule actual 22-segmented; geniculated; between the segment 18 and segment 19; with swollen and modified region; formed by 5 segments; between 13 and 17 segments. Actual segment 1 with seta; one element; straight; none larger than segment; without spinules; without vestigial seta; without conical seta; without modified seta; without spinous process; with aesthetasc; one element. Actual segment 2 with seta; three elements; of unequal size; straight; none larger than segment; without spinules; with vestigial seta; one element; without conical seta; without modified seta; without spinous process; with aesthetasc; one element. Actual segment 3 with seta; one element; one larger than segment; surpassing to distal margin; beyond three sequential segments; straight; blunt apex; without spinules; with vestigial seta; one element; without conical seta; without modified seta; without spinous process; with aesthetasc. Actual segment 4 with seta; one element; one larger than segment; surpassing to distal margin; straight; not beyond three sequential segments; without spinules; without vestigial seta; without conical seta; without modified seta; without spinous process; without aesthetasc. Actual segment 5 with seta; one element; straight; one larger than segment; surpassing to distal margin; not beyond three sequential segments; without spinules; with vestigial seta; one element; without conical seta; without modified seta; without spinous process; with aesthetasc; one element. Actual segment 6 with seta; one element; none larger than segment; straight; without spinules; without vestigial seta; without conical seta; without modified seta; without spinous process; without aesthetasc. Actual segment 7 with seta; one element; straight; one larger than segment; surpassing to distal margin; beyond three sequential segments; blunt apex; without spinules; without vestigial seta; without conical seta; without modified seta; without spinous process; with aesthetasc; one element. Actual segment 8 with seta; one element; straight; none larger than segment; without spinules; without vestigial seta; with conical seta; one element; not reaching to middle-point of the sequent segment; without modified seta; without spinous process; without aesthetasc. Actual segment 9 with seta; two elements; of unequal size; straight; one larger than segment; surpassing to distal margin; beyond three sequential segments; blunt apex; without spinules; without vestigial seta; without conical seta; without modified seta; without spinous process; with aesthetasc; one element. Actual segment 10 with seta; one element; straight; none larger than segment; without spinules; without vestigial seta; without conical seta; with modified seta; presenting blunt apex; slender form; surpassing to distal margin; beyond of the sequential segment; parallel to antennule direction; without spinous process; without aesthetasc. Actual segment 11 with seta; one element; straight; one larger than segment; surpassing to distal margin; not beyond three sequential segments; without spinules; without vestigial seta; without conical seta; with modified seta; slender form; presenting blunt apex; surpassing to distal margin; beyond of the sequential segment; parallel to antennule direction; shorter length than homologous of actual segment 13; without spinous process; without aesthetasc. Actual segment 12 with seta; one element; straight; one larger than segment; surpassing to distal margin; not beyond three sequential segments; without spinules; without vestigial seta; with conical seta; one element; smaller than to segment 8; without modified seta; without spinous process; with aesthetasc; one element; absent internal perpendicular fission. Actual segment 13 with seta; one element; straight; one larger than segment; surpassing to distal margin; not beyond three sequential segments; without spinules; without vestigial seta; without conical seta; with modified seta; stout form; surpassing to distal margin; to the distal-point of the sequence segment; perpendicular to antennule direction; presenting bifid apex; without spinous process; with aesthetasc; one element. Actual segment 14 with seta; two elements; of unequal size; straight; one larger than segment; surpassing to distal margin; beyond three sequential segments; blunt apex; without spinules; without vestigial seta; without conical seta; without modified seta; without spinous process; with aesthetasc; one element. Actual segment 15 with seta; two elements; of unequal size; straight; not bifidform; none larger than segment; without spinules; without vestigial seta; without conical seta; without modified seta; with spinous process; on outer margin; surpassing distal margin; with aesthetasc; one element. Actual segment 16 with seta; two elements; of unequal size; plumose; one larger than segment; surpassing to distal margin; not beyond three sequential segments; not bifidform; without spinules; without vestigial seta; without conical seta; without modified seta; with spinous process; on outer margin; surpassing distal margin; unequal size to process on preceding segment; with aesthetasc; one element. Actual segment 17 with seta; two elements; of unequal size; straight; none larger than segment; bifidform; without spinules; without vestigial seta; without conical seta; with modified seta; one element; stout form; surpassing to distal margin; not beyond of the sequential segment; parallel to antennule direction; without spinous process; without aesthetasc. Actual segment 18 with seta; two elements; of equal size; straight; none larger than segment; without spinules; without vestigial seta; without conical seta; with modified seta; one element; stout form; surpassing distal margin; parallel to antennule direction; without spinous process; without aesthetasc. Actual segment 19 with seta; two elements; of unequal size; plumose; none larger than segment; without spinules; without vestigial seta; without conical seta; with modified seta; two elements; stout form; at least one bifid form; surpassing distal margin; parallel to antennule direction; without spinous process; with aesthetasc; one element. Actual segment 20 with seta; four elements; of unequal size; straight; one larger than segment; surpassing to distal margin; beyond three sequential segments; without spinules; without vestigial seta; without conical seta; without modified seta; with spinous process; distally; not reaching beyond of distal-point segment 21; without aesthetasc. Actual segment 21 with seta; two elements; of equal size; plumose; one larger than segment; surpassing to distal margin; greater 3x than original segment; without spinules; without vestigial seta; without conical seta; without modified seta; without spinous process; without aesthetasc. Actual segment 22 with seta; four elements; of equal size; one larger than segment; plumose; surpassing to distal margin; greater 3x than original segment; without spinules; without vestigial seta; without conical seta; without modified seta; without spinous process; with aesthetasc; one element.

##### Left antennules

Uniramous; Left antennule surpassing to prosome; Left antennule not extending beyond caudal rami. Ancestral segment I and II separated. Ancestral segment II and III fused. Ancestral segment III and IV fused. Ancestral segment IV and V separated. Ancestral segment V and VI separated. Ancestral segment VI and VII separated. Ancestral segment VII and VIII separated. Ancestral segment VIII and IX separated. Ancestral segment IX and X separated. Ancestral segment X and XI separated. Ancestral segment XI and XII separated. Ancestral segment XII and XIII separated. Ancestral segment XIII and XIV separated. Ancestral segment XIV and XV separated. Ancestral segment XV and XVI separated. Ancestral segment XVI and XVII separated. Ancestral segment XVII and XVIII separated. Ancestral segment XVIII and XIX separated. Ancestral segment XIX and XX separated. Ancestral segment XX and XXI separated. Ancestral segment XXI and XXII separated. Ancestral segment XXII and XXIII separated. Ancestral segment XXIII and XXIV separated. Ancestral segment XXIV and XXV separated. Ancestral segment XXV and XXVI separated. Ancestral segment XXVI and XXVII separated. Ancestral segment XXVII and XXVIII fused.

Left antennule actual 25-segmented; not-geniculated. Actual segment 1 with seta; one element; none larger than segment; straight; without spinules; without vestigial seta; without conical seta; without modified seta; without spinous process; with aesthetasc; one element. Actual segment 2 with seta; three elements; of equal size; none larger than segment; straight; without spinules; with vestigial seta; one element; without conical seta; without modified seta; without spinous process; with aesthetasc; one element. Actual segment 3 with seta; one element; one larger than segment; straight; surpassing to distal margin; beyond three sequential segments; without spinules; with vestigial seta; one element; without conical seta; without modified seta; without spinous process; with aesthetasc. Actual segment 4 with seta; one element; none larger than segment; straight; without spinules; without vestigial seta; without conical seta; without modified seta; without spinous process; without aesthetasc. Actual segment 5 with seta; one element; one larger than segment; straight; surpassing to distal margin; not beyond three sequential segments; without spinules; with vestigial seta; one element; without conical seta; without modified seta; without spinous process; with aesthetasc; one element. Actual segment 6 with seta; one element; none larger than segment; straight; without spinules; without vestigial seta; without conical seta; without modified seta; without spinous process; without aesthetasc. Actual segment 7 with seta; one element; one larger than segment; straight; surpassing to distal margin; beyond three sequential segments; without spinules; without vestigial seta; without conical seta; without modified seta; without spinous process; with aesthetasc; one element. Actual segment 8 with seta; one element; one larger than segment; straight; surpassing distal margin; without spinules; without vestigial seta; with conical seta; without modified seta; without spinous process; without aesthetasc. Actual segment 9 with seta; two elements; of unequal size; one larger than segment; straight; surpassing to distal margin; beyond three sequential segments; without spinules; without vestigial seta; without conical seta; without modified seta; without spinous process; with aesthetasc; one element. Actual segment 10 with seta; one element; none larger than segment; straight; without spinules; without vestigial seta; without conical seta; without modified seta; without spinous process; without aesthetasc. Actual segment 11 with seta; one element; one larger than segment; straight; surpassing to distal margin; beyond three sequential segments; without spinules; without vestigial seta; without conical seta; without modified seta; without spinous process; without aesthetasc. Actual segment 12 with seta; one element; one larger than segment; straight; surpassing distal margin; without spinules; without vestigial seta; with conical seta; without modified seta; without spinous process; with aesthetasc; one element. Actual segment 13 with seta; one element; none elongated; straight; surpassing distal margin; without spinules; without vestigial seta; without conical seta; without modified seta; without spinous process; without aesthetasc. Actual segment 14 with seta; one element; elongated; straight; surpassing to distal margin; beyond three sequential segments; without spinules; without vestigial seta; without conical seta; without modified seta; without spinous process; with aesthetasc; one element. Actual segment 15 with seta; one element; larger than segment; straight; surpassing to distal margin; not beyond three sequential segments; without spinules; without vestigial seta; without conical seta; without modified seta; without spinous process; without aesthetasc. Actual segment 16 with seta; one element; larger than segment; plumose; surpassing to distal margin; not beyond three sequential segments; without spinules; without vestigial seta; without conical seta; without modified seta; without spinous process; with aesthetasc; one element. Actual segment 17 with seta; one element; not larger than segment; straight; without spinules; without vestigial seta; without conical seta; without modified seta; without spinous process; without aesthetasc. Actual segment 18 with seta; one element; larger than segment; straight; surpassing to distal margin; beyond three sequential segments; without spinules; without vestigial seta; without conical seta; without modified seta; without spinous process; without aesthetasc. Actual segment 19 with seta; one element; not larger than segment; straight; surpassing distal margin; without spinules; without vestigial seta; without conical seta; without modified seta; without spinous process; with aesthetasc; one element. Actual segment 20 with seta; one element; not larger than segment; straight; surpassing distal margin; without spinules; without vestigial seta; without conical seta; without modified seta; without spinous process; without aesthetasc. Actual segment 21 with seta; one element; larger than segment; plumose; surpassing to distal margin; beyond three sequential segments; without spinules; without vestigial seta; without conical seta; without modified seta; without spinous process; without aesthetasc. Actual segment 22 with seta; two elements; of unequal size; one of them elongated; plumose; surpassing to distal margin; without spinules; without vestigial seta; without conical seta; without modified seta; without spinous process; without aesthetasc. Actual segment 23 with seta; two elements; of unequal size; one larger than segment; plumose; surpassing to distal margin; greater 3x than original segment; without spinules; without vestigial seta; without conical seta; without modified seta; without spinous process; without aesthetasc. Actual segment 24 with seta; two elements; of equal size; one larger than segment; plumose; surpassing to distal margin; greater 3x than original segment; without spinules; without vestigial seta; without conical seta; without modified seta; without spinous process; without aesthetasc. Actual segment 25 with seta; four elements; of equal size; elongated; plumose; surpassing to distal margin; 4 times larger than segment; without spinules; without vestigial seta; without conical seta; without modified seta; without spinous process; with aesthetasc; one element.

##### Antenna

Biramous. Antenna coxa separated from the basis; bearing seta; 1; on inner surface; at distal corner; reaching to the endopod 1. Antenna basis (fusion) separated from the endopodal segment; bearing seta; 2; on inner surface; at distal corner. Endopodal ancestral segment I and II separated. Ancestral segment II and III fused. Ancestral segment III and IV fused. Ancestral segment III and IV fully. Antenna endopod actual 2-segmented. Actual segment 1 not bilobate; with seta; two; on inner margin; with spinules; as a row; obliquely; on outer surface; with pore. Actual segment 2 bilobate; without discontinuity on outer cuticle; inner lobe bearing 8 setae; distally; outer lobe bearing 7 setae; distally; with spinules; as a patch; on outer surface. Antenna exopod ancestral segment I and II separated. Ancestral segment II and III fused. Ancestral segment III and IV fused. Ancestral segment IV and V separated. Ancestral segment V and VI separated. Ancestral segment VI and VII separated. Ancestral segment VII and VIII separated. Ancestral segment VIII and IX separated. Ancestral segment IX and X fused. Antenna exopod actual 7-segmented. Actual segment 1 single; elongated (width-length, equal or larger ratio 2:1); with seta; one; at inner surface. Actual segment 2 compound; elongated (larger width-length ratio 2:1); with seta; three; at inner surface. Actual segment 3 single; not elongated (lesser width-length ratio 2:1); with seta; one; at inner surface. Actual segment 4 single; not elongated (lesser width-length ratio 2:1); with seta; one; at inner surface. Actual segment 5 single; not elongated (lesser width-length ratio 2:1); with seta; one; at inner surface. Actual segment 6 single; not elongated (lesser width-length ratio 2:1); with seta; one; at inner surface. Actual segment 7 compound; elongated (larger or equal width-length ratio 2:1); with seta; one; at inner surface; and three; at distal surface.

##### Oral features

**Mandible**. Coxal gnathobase sclerotized; with lobe; prominent; on caudal margin; presence of cutting blade; with tooth-like prominence; two, distinctly; 1 acute; on caudal margin; and 1 triangular; on sub-caudal margin; without acute projection between the prominences; with additional spinules; as a row; on dorsal surface; with seta; 1; dorsally; on apical surface; with spinules; apicalmost. Mandible palps biramous; comprising the basis; with seta; four; differently inserted; first medially; reaching to beyond the endopod 1; second distally; third distally; fourth distally; on inner margin; none with setulose ornamentation. Mandible endopod 2-segmented. Mandible endopod 1 with lobe; bearing seta; four; distally inserted; without spinules. Mandible endopod 2 without lobe; bearing setae; nine elements; distally inserted; with spinules; as a row; double. Mandible exopod 4-segmented. Mandible exopod 1 with seta; one element; distally; on inner margin. Mandible exopod 2 with seta; one element; distally; on inner side. Mandible exopod 3 with seta; one element; distally; on inner side. Mandible exopod 4 with setae; three elements; on terminal region. **Maxillule**. Birramous. Maxillule 3-segmented. Maxillule praecoxa with praecoxal arthrite; bearing spines; fifteen elements; ten marginally; plus, five sub-marginally; with spinules; as a patch; on sub-marginal surface. Maxillule coxa with coxal epipodite; with conspicuous outer lobe; bearing setae; nine elements; with coxal endite; elongated (larger or equal width-length ratio 2:1); bearing setae; four elements. Maxillule basis with basal endite; double; first proximal; elongated (larger width-length ratio 2:1; separated from basis; with setae; four elements; distally inserted; second distal; fused to basis; not elongated (lesser width-length ratio 2:1); with setae; four elements; distally inserted; with setules; as a row; on inner side; basal exite present; with setae; one element; on outer surface. Maxillule endopod 1-segmented. Endopod 1 bilobate; first proximal; with setae; three elements; second distal; with setae; five elements. Maxillule exopod 1-segmented. Exopod 1 with setae; six elements; with setules; as a row; on inner side; spinules absent. **Maxilla**. Uniramous. Maxilla 5-segmented. Maxilla praecoxa fused to coxa; incompletely; distinct externally; with praecoxal endite; double; first elongated endite (larger or equal width length ratio 2:1); proximally inserted; with seta; straight, or plumose; 1 straight; 4 plumose; with spine; single; without spinules; without setule; second elongated endite (larger or equal width length ratio 2:1); distally inserted; with seta; plumose; 3 plumose; without spine; with spinules; as a row; on distal margin; with setule; as a row; on distal margin; absence of outer seta. Maxilla coxa with coxal endite; double; first elongated endite (larger or equal width); proximally inserted; with seta; plumose; 3 plumose; without spine; without spinules; with setules; as a row; on proximal margin; second elongated endite (larger or equal width); distally inserted; with seta; plumose; 3 plumose; without spine; without spinules; with setules; as a row; on proximal margin; absence of outer seta. Maxilla basis with basal endite; single; elongated (larger or equal width-length ratio 2:1); with seta; plumose; 3 plumose; without spinules; absence of outer seta. Maxilla endopod 2-segmented. Endopod 1 with seta; 2 plumose; without spine; without spinules; without setules. Maxilla endopod 2 with seta; 2 plumose; without spine; without spinules; without setules. **Maxilliped**. Uniramous; Maxilliped 8-segmented. Maxilliped praecoxa fused to coxa; incompletely; distinct internally; with praecoxal endite; not elongated (lesser width-length ratio 2:1); distally inserted; with seta; 1 straight; with spinules; as a row; single; on basal surface; without setules. Maxilliped coxa with coxal endite; three coxal endite; first elongated (larger or equal width); proximally inserted; with seta; 2 plumose; with spinules; as a patch; single; on apical surface; without setules; second not elongated (lesser width-length ratio 2:1); medially inserted; with seta; 3 plumose; with spinules; as a row; single; on medial surface; without setules; third elongated (larger or equal width length ratio 2:1); distally inserted; with seta; 3 plumose; none reaching to beyond of the basis; with spinules; as a row; single; on basal surface; without setules; with lobe; prominence; at inner distal angle; ornamented; with spinules; continuously on margin. Maxilliped basis without basal endite; with seta; 3 plumose; with spinules; as a row; single; on medial surface; with setules; as a row; single; on inner margin. Maxilliped endopod segment 6-segmented. Endopod 1 with seta; 2 plumose; on inner surface. Endopod 2 with seta; 3 plumose; on inner surface. Endopod 3 with seta; 2 plumose; on inner surface. Endopod 4 with seta; 2 plumose; on inner surface. Endopod 5 with seta; 2 plumose; on inner surface, or on outer surface; outer seta absent. Endopod 6 with seta; 4 plumose; on inner surface, or on outer surface.

##### Swimming legs features

**First swimming legs.** Symmetrical; biramous. First swimming legs intercoxal plate without seta. First swimming legs praecoxa absent. First swimming legs coxa with seta; one; straight; distally inserted; on inner surface; surpassing to basal segment; with setules; one group; as a patch; on inner margin; without spinules; without spine. First swimming legs basis without seta; with setules; as a patch; single; on outer surface; without spinules; without spine. First swimming legs endopod 2-segmented. Endopod 1 with seta; straight; restricted; to inner surface; one element; without spine; with setules; as a row; single; continuously; on outer surface; without spinules; absence of Schmeil’s organ. Endopod 2 with seta; unrestricted; three on inner surface; one on outer surface; two on distal surface; straight; without spine; with setules; as a row; single; continuously; on outer surface; without spinules; absence of Schmeil’s organ. Endopod 3 absence. First swimming legs exopod 1 with seta; restricted; 1 on inner surface; with spine; 1; stout; smaller than original segment; serrated; on inner side; continuously; without setules. First swimming legs exopod 2 with seta; restricted; 1 on inner surface; straight; without spine; with setules; as a row; single; continuously; on inner margin, or on outer margin; without spinules. First swimming legs exopod 3 with setule; as a row; single; continuously; on outer surface; without spinules; with seta; unrestricted; 2 on inner surface; 2 on terminal surface; with spine; 2; unequal size; first no longer 2x than origin segment; stout; serrated; on inner side, or on outer side; equally; second longer 3x than origin segment; slender; serrated; on outer side; with ornamentation on non-serrated side; by setules. **Second swimming legs**. Symmetrical; Second swimming legs biramous. Second swimming legs intercoxal plate without seta. Second swimming legs praecoxa present; located laterally. Second swimming legs coxa with seta; straight; distally inserted; on inner surface; surpassing to basal segment; without setules; without spinules; without spine. Second swimming legs basis without seta; without setules; without spinules; without spine. Second swimming legs endopod 3-segmented. Endopod 1 with seta; straight; restricted; one on inner surface; without spine; with setules; as a row; single; continuously; on outer surface; without spinules; absence of Schmeil’s organ. Endopod 2 with seta; straight; unrestricted; two on inner surface; without spine; with setules; as a row; single; continuously; on outer side; without spinules; presence of Schmeil’s organ; on posterior surface. Endopod 3 with seta; straight; unrestricted; three on inner surface; two on outer surface; two on distal surface; without spine; without setules; with spinules; as a row; double; distally inserted; at anterior surface; absence of Schmeil’s organ. Second swimming legs exopod 1 with seta; restricted; one on inner surface; with spine; 1; stout; not reaching to distal-third of the exopod 2; serrated; on inner side, or on outer side; with setules; as a row; single; continuously; on inner side; without spinules; absence of Schmeil’s organ. Exopod 2 with seta; unrestricted; one on inner surface; with spine; 1; stout; not surpassing the exopod 3; serrated; on inner side, or on outer side; with setules; as a row; single; continuously; on inner surface; without spinules; absence of Schmeil’s organ. Exopod 3 with seta; plurimarginal; three on inner surface; two on terminal surface; with spine; 2; unequal size; first no longer 2x than origin segment; stout; serrated; on inner side, or on outer side; equally; second longer 2x than origin segment; slender; serrated; on outer side; with ornamentation on non-serrated side; of setules; setules on outer surface; as a row; single; continuously; on inner surface; with spinules; as a row; single; distally inserted; at anterior surface; absence of Schmeil’s organ. **Third swimming legs**. Symmetrical; Third swimming legs biramous. Third swimming legs intercoxal plate without seta. Third swimming legs praecoxa present; not laterally located. Third swimming legs coxa with seta; straight; distally inserted; on inner surface; surpassing to first endopodal segment; without setules; without spinules; without spine. Third swimming legs basis without seta; without setules; without spinules; without spine. Third swimming legs endopod 3-segmented. Endopod 1 with seta; restricted; one on inner surface; without spine; without setules; without spinules; absence of Schmeil’s organ. Endopod 2 with seta; restricted; two on inner surface; straight; without spine; without setules; without spinules; absence of Schmeil’s organ. Endopod 3 with seta; straight; plurimarginal; two on inner surface; two on outer surface; three on terminal surface; without spine; without setules; with spinules; as a row; distally inserted; double; at anterior surface; absence of Schmeil’s organ. Third swimming legs exopod 1 with seta; restricted; straight; one on inner surface; with spine; 1; stout; not reaching to the distal-third of the exopod 2; serrated; equally; on inner surface, or on outer surface; with setules; as a row; single; continuously; on inner surface; without spinules; absence of Schmeil’s organ. Exopod 2 with seta; straight; restricted; one on inner surface; with spine; 1; stout; not reaching out to exopod 3; serrated; on inner side, or on outer side; equally; with setules; as a row; single; continuously; on inner side; without spinules; absence of Schmeil’s organ. Exopod 3 without setules; with spinules; as a row; single; distally inserted; at anterior surface; with seta; straight; unrestricted; three on inner surface; two on terminal surface; with spine; 2; unequal size; first no longer 2x than origin segment; stout; serrated; on inner side, or on outer side; equally; second longer 2x than origin segment; slender; serrated; on outer side; with ornamentation on non-serrated side; of setules; absence of Schmeil’s organ. **Fourth swimming legs**. Symmetrical; biramous. Intercoxal plate without sensilla. Praecoxa present. Coxa with seta; distally inserted; on inner margin; reaching out to endopod 1; without spinules; setules absent. Basis with seta; one; medially inserted; on posterior surface; smaller than the original segment; without setules; without spinules; without spine. Fourth swimming legs endopod 3-segmented. Endopod 1 with seta; one; restricted; on inner surface; without spine; without setules; without spinules; absence of Schmeil’s organ. Endopod 2 with seta; restricted; two on inner side; without spine; with setules; as a row; single; continuously; on outer surface; without spinules; absence of Schmeil’s organ. Endopod 3 with seta; unrestricted; two on inner surface; two on outer surface; three on distal surface; without spine; without setules; with spinules; as a row; double; distally inserted; at anterior surface; absence of Schmeil’s organ. Fourth swimming legs exopod 1 with seta; restricted; one on inner surface; with spine; 1; stout; not reaching out to distal-third of the exopod 2; serrated; on inner side, or on outer side; equally; with setules; as a row; single; continuously; on inner surface; without spinules; absence of Schmeil’s organ. Exopod 2 with seta; restricted; one on inner surface; with spine; 1; stout; not reaching the end of exopod 3; serrated; on inner side, or on outer side; equally; with setules; as a row; single; continuously; on inner surface; without spinules; absence of Schmeil’s organ. Exopod 3 without setules; with spinules; as a row; single; distally inserted; at anterior surface; with seta; unrestricted; three on inner surface; two on distal surface; with spine; 2; unequal size; first no longer 2x than origin segment; stout; serrated; on inner side, or on outer side; equally; second longer 2x than origin segment; slender; serrated; on outer side; without ornamentation on non-serrated side; absence of Schmeil’s organ.

##### Fifth swimming legs features

Asymmetrical. Fifth swimming leg intercoxal plate with length not equal or greater than width on 1.5x; with irregular proximal margin; discontinuous to; the anterior margin of the left coxa, or the anterior margin of the right coxa; posterior sensilla on the right lateral absent. **Fifth left swimming leg**. Fifth left swimming leg biramous; leg surpassing first right exopod segment. Fifth left swimming leg praecoxa present; rudimentary; separated from the coxae; without ornamentation. Fifth left swimming leg coxa concave inner side; without teeth-like structures; with process; conical; on posterior surface; outer side; distally inserted; not projecting over basis; with sensilla; stout; triangular; at apex; no longer 2x than insertion basis; without swelling; without seta; without spinules. Fifth left swimming leg basis sub-cylindrical; unequal size between inner and outer side; shorter outer than inner side; with concave inner side; rounded internal proximal expansion absent; without outgrowth; without groove; absence of protuberance; with seta; outerly inserted; no longer 2x than origin segment; presence of minutely granular; as a patch; outerly. Fifth left swimming leg endopod segments 1 and 2 fused; segments 2 and 3 fused; 1-segmented; stout; separated from the basis; ornamented; on inner side; with spinules; more than four elements; as a row; terminally, or sub-terminally; row of setules absent; without seta. Fifth left swimming leg exopod segments 1 and 2 separated; segments 2 and 3 fused; 2-segmented; stout; separated from the basis. Fifth left swimming leg exopod 1 sub-triangular; longer than broad; equal size between inner and outer side; rectilinear inner side; convex outer side; without swelling; without marginal extension; without process; with lobe; single; circular; medially inserted; on inner side; covered; by setules; without outer spine; absence seta. Fifth left swimming leg exopod 2 digitiform; longer than broad; equal size between inner and outer side; disform inner side; with convex outer side; setulose pad present; prominently rounded; proximally; on inner side; inflated medial region absent; distal process present; digitiform; non denticulate; without transverse row of denticles; none oblique row of 5 denticles; innerly directed; with seta; spiniform; not ornamented by spinules; not surpassing the distal-point of the segment; without outer spine; terminal claw absent.

##### Fifth right swimming leg

Biramous. Fifth right swimming leg praecoxa present; separated from the coxae; without ornamentation. Fifth right swimming leg coxa convex inner side; without teeth-like structures; with process; rounded; distally inserted; on posterior surface; closest to the outer rim; projecting over basis; not beyond the first third; without triangular protuberance innerly; with sensilla; slender; at apex; no longer 2x than basal insertion; without marginal extension; without seta; without spinules. Fifth right swimming leg basis trapezoidal; unequal size between inner and outer side; shorter outer than inner side; rectilinear inner side; tumescence present; not inflated; restricted on inner surface; proximally; without protuberance; absence of distinct minutely granular; additional inner process absent; with posterior groove; deep; latitudinally; not reaching the endopodal lobe; ornamented; with tubercles; throughout of the outer border; with seta; outerly inserted; on anterior surface; no longer 2x than origin segment; posterior protrusion present; distal process absent. Fifth right swimming leg with endopodite present; separated from the basis; on anterior surface; ancestral segments 1 and 2 separated; ancestral segments 2 and 3 fused; 2-segmented; stout; ornamented; with setules; as a row; on inner side; sub-terminally; without seta. Fifth right swimming leg exopod segments 1 and 2 separated; segments 2 and 3 fused; 2-segmented; stout; separated from the basis. Fifth right swimming leg exopod 1 trapezium; longer than broad; nearly 1.25 times; unequal size between both sides; shorter inner than outer side; convex inner side; rectilinear outer side; with marginal extension; sub-triangular; distally inserted; at outer rim; spinules present; without process; without outer spine; without seta; internal prominence absent; lamella on posterior surface absent. Fifth right swimming leg exopod 2 cylindrical; longer than broad; nearly 2.5 times; equal size between both sides; disform inner side; convex outer side; without posterior proximal swelling; inner-posterior process present; sub-triangular; medially; without marginal expansion; curved ridge on distal posterior surface present; chitinous knobs absent; with outer spine; inserted sub-distally; arched; externally directed; not ornamented innerly; not ornamented outerly; sharp tip; without apparent curve; lesser than the length of the exopod 2; beyond to 2 times its size; 3x; sensilla absent; terminal claw present; not equal or longer 1.5 times than insertion segment; sclerotized; arched; inward; without conspicuous curve; not ornamented innerly; not ornamented outerly; sharp tip; not curved tip; without medial constriction; hyaline process absent.

##### FEMALE

Body longer and wider than male. Widest at first metasome segment. Distal margin of the prosomal segments without one line of setules at posterior margin. Prosome segments without spinules at prosomal segments. Fourth metasome segment presence of dorsal protuberance; rounded; inserted distally; without posterior process; without anterior process; fourth metasome segment without proximal sensillae present. Fourth and fifth metasome segments fused; partially; on dorsal surface. Limit between fourth and fifth metasome segments without ornamentation. **Fifth metasome segment**. Fifth metasome segment without sensilla; with epimeral plates. Epimeral plates asymmetrical. Right epimeral plates prominent, as projections; thinner than the left; one posterior-laterally directed; not reaching half length of the genital segment; with sensilla at the apex; dorsal-posterior sensilla present; slender; without ornamentation. Left epimeral plate without expansion.

##### Urosome

3-segmented. **Genital double-somite**. Asymmetrical in dorsal view; longer than broad; longer than other urosomites combined; dorsal suture at mid-length absent; not covered by spinules; with swelling; rounded; unequal size; greater right than left; anteriorly; with sensillae; on both sides; one; stout; with robust apex; at left lateral; not on lobular base; medially; one; stout; at right lateral; on lobular base; anteriorly; with robust apex; of equal size between then; lateral protuberance absent; with right posterior rim expanded; over next segment; without slender sensilla on each posterior rim; without posterior-dorsal process. Genital double-somite opercular pad present; broader than longer; symmetrical; development laterally; expanded posteriorly; covering partially; double gonoporal slit; located ventrally; with arthrodial membrane; inserted anteriorly; post-genital process absent; disto-ventral tumescence absent; ventral vertical folds absent; dorsal sensilla absent. Second urosome segment without ventral fusion to anal segment; right distal process absent. Caudal rami patch of setules on outer surface absent; patch of spinules on outer surface absent.

##### Appendices features

Rostrum basal process absent. **Antennules**. Symmetrical. Right antennule surpassing to genital double-segment; extending beyond caudal rami. Right antennule not exceeding the caudal setae. Right antennule ornamentation pattern equals to male left antennule; fully.

##### Fifth swimming legs

Symmetrical; Fifth swimming legs biramous. Fifth swimming legs intercoxal plate longer than wide; separated from the legs. Fifth swimming legs praecoxa with sclerite praecoxal; separated from the coxae; without ornamentation. Fifth swimming legs coxa with process; conical; at the outer rim; distally; sensilla present; stout; at apex; projecting over basal segment; no longer 2x than basal insertion; marginal extension absent; without swelling; without seta; without spinules. Fifth swimming legs basis sub-triangular; unequal size between inner and outer sides; shorter outer than inner side; with convex inner side; without proximal inner outgrowth; without groove; with distal extension; on posterior surface; with seta; outerly inserted; on anterior surface; no longer 2x than origin segment. Fifth swimming legs endopod segments 1 and 2 separated; segments 2 and 3 fused; 2-segmented; with incomplete suture; stout; separated from the basis; ornamentation on segment 2; with spinules; as a row; single; non-oblique; sub-terminally; at anterior surface; with seta; double; one medially; on posterior surface; rectilinear; one distally; on posterior surface; arched; of unequal size; distal seta longer than medial seta. Fifth swimming legs exopod segments 1 and 2 separated; segments 2 and 3 separated; 3-segmented; separated from the basis. Fifth swimming legs exopod 1 sub-cylindrical; longer than wide; longer or equal than 2 times; with unequal size between inner and outer side; shorter inner than outer side; with convex inner side; with rectilinear outer side; without swelling; without marginal extension; without posterior process; without spine; without seta. Fifth swimming legs exopod 2 sub-cylindrical; longer than broad; longer or equal than 2 times; without swelling; without marginal extension; without process; without lobe; with spine; inserted laterally; rectilinear; without ornamentation; sharp tip; equal size or larger than next segment; without seta. Fifth swimming legs exopod 3 cylindrical; longer than wide; without swelling; without process; without lobe; without spine; with seta; double; inserted terminally; unequal size between them; outer seta smaller than inner; nearly 3 times; outer seta not ornamented by setules; without ornamentation; presence of terminal claw; sclerotized; arched; externally directed; convex inner side; with ornamentation; of denticles; as a row; on surface partially; at medial region; concave outer side; with ornamentation; of denticles; as a row; on surface partially; at medial region; blunt tip; 6 times longer than origin segment.

##### Distribution records

###### BRAZIL

**Rio Grande do Sul**: São Gonçalo Channel (Montú & Gloeden, 1986); Lagoa dos Patos (Bohrer & Araújo, 1999). PARAGUAY. Makthlawaiya, 23°25’S, 58°19’W (Lowndes, 1934). ARGENTINA. **Chaco**: Estero Marocho and Estero Pati (Dussart & Frutos, 1986). **Chaco**: Cangui Chico stream; Oro River; Gayacuru River; Tragadero River; Palometa River (Dussart & Frutos, 1986). **Santa Fé**: along highway N° 168, from Santa Fé to Helvecia; Laguna Los Espejos, at Sirgadero Island, in front of the city of Santa Fé; Madrejón Don Felipe, Colastiné (Ringuelet & Martínez de Ferrato, 1967). URUGUAY. **Plata River Basin**: low Paraná River (Perbiche-Neves *et al*., 2015).

##### Habitat

Habitat in freshwaters: lakes, shallow pools, and rivers.

##### Remarks

The origin of the organisms of *D.* (s.l.) *carteri* is not specified by Lowndes (1934), only indications are made for environments in swamps, shallow lakes with aquatic plants, and flooded lands in the Makthlaiya, Paraguay. The current recombination was proposed in the origin of *Notodiaptomus*, being recorded several other times to the South Neotropical region (*i.e.*, Rio Grande do Sul, Brazil; Paraguay; and Argentina). A new recombination was proposed by Dussart (1985) for *Notodiaptomus* (*Notodiaptomus*) *carteri* without sufficient foundations for validation.

Throughout the morphological trajectory of the species relevant inconsistencies are recorded. In the original description, illustrations of the female show fifth swimming legs endopod 2-segmented, and fourth and fifth metasome segments separated completely. Ringuelet & de Ferrato (1967) when adding information on specimens from Argentina, illustrated the same morphology. However, in possession of individuals also from Argentina Dussart & Frutos (1986) partially corroborated these conditions by representing the female with metasome segments 4 and 5 separated only dorsally, and endopod 1-segmented with discontinuity in the cuticle internally.

In a recent morphological review for the species, individuals from the lower stretch of the Paraná River were treated by Perbiche-Neves *et al*. (2015). In this review the female of the species was illustrated with the metasome and endopod fused completely, disagreeing not only with Lowndes, Ringuelet & de Ferrato (1967), but Dussart & Frutos (1986) for metasomal fusion. In the present effort presented for specimens from Rio Grande do Sul, Brazil, we corroborate the original description for the endopod condition and redescription of Dussart & Frutos (1986) and Perbiche-Neves *et al*. (2015) for metasome segments 4 and 5 fused laterally.

Other relevant attributes are for female genital double-somite with right sensilla on lobular base, and female fourth metasome segment with dorsal rounded protuberance distally, both noted in Perbiche-Neves *et al*. (2015) previously. Among the attributes described in the creation and amendment of *Notodiaptomus*, the only feature to which the taxon does not converge is female fifth swimming legs endopod 1-segmented. As annotation of divergence of the type species of the genus: (1) male A1R actual segment 12 with conical seta smaller than those of actual segment 8; (2) male A1R actual segment 13 with spiniform modified seta reaching to distal-point actual segment 14; and (3) male fifth right swimming leg basis with latitudinal posterior groove not reaching to endopodal lobe.

#### Notodiaptomus cearensis Wright, 1936

##### Synonymy

*Diaptomus cearensis* Wright, 1936a: 80, pl. 1, fig. 2; 1937: 73, 76; 1938a: 300; 1938b: 563; Reid, 1991: 740. *Notodiaptomus cearensis*; Kiefer, 1936a: 197; 1956: 242; Brandorff, 1972: 44; 1976: 615, 616, fig. 2; Bowman, 1973: 193, 194, figs. 1–21, 33–35; Löffler, 1981: 15; Dussart & Defaye, 1983: 137; 1995: fig. L62; Dussart, 1984a: 26, 27, 28, 34, 35, 36, 38, 39, 49, fig. 6; Reid, 1985: 589, 590; Matsumura-Tundisi, 1986: 542, 547, figs. 61–66, 100; Cicchino *et al*., 1989: 101; Reid, 1991: 740; Cicchino, 1994: 145, fig. 9; Tundisi & Matsumura-Tundisi, 1994: 27; Rocha *et al*., 1995: 156; 1998: 794, 795, tab. 1; Santos-Silva, 1998: 207; Santos-Silva *et al*., 1999: 127; Santos-Silva, 2008: 22; Santos-Silva *et al*., 2015: 53–57, figs. 30–32, identification key to male and female; Perbiche-Neves *et al*., 2015: 42–44, figs. 32–33; Perbiche-Neves *et al*., 2020: 697-698, key to the Neotropical diaptomid, fig. 21.15 C. *Notodiaptomus (Notodiaptomus) cearensis*; Dussart, 1985a: 208.

##### Type locality

Imprecisely specified in the original description. However, two localities in the State of Ceará are listed: tauapé Lake in Fortaleza, and Mecejana Lake in Mecejana. Other localities in the northeast of Ceará State are mentioned, like as pond Pilões near São João do Rio do Peixe, Paraiba State, where several bodies of water close to Caraúbas, and Assu Rio Grande do Norte are related.

##### Type material

Holotype unspecified originally, and material type probably non-existent according to Santos Silva *et al*. (2015).

##### Material examined

Non-type material: 2 males, and 3 females, 21.XII.1933, from the Piloes Pond, Ceará, Brazil. Biological material from the collection of Stillman Wright 1934–1935 (pack 4612-CN), stored in Plankton laboratory, Instituto Nacional de Pesquisas da Amazônia - INPA (under code LP19-A III); 1 male (INPA-COP013, slides a-h) and 1 female (INPA-COP014, slides a-h) were selected to be dissection on eight slides each and deposited in the Zoological Collection of the INPA, Brazil. Additional material examined: 01 male, and 2 females, entire in formalin solution (MZUSP 32933) from the upper Tiete River, at the Barra Bonita Reservoir, Brazil, collected by G. Perbiche-Neves, and stored in MZUSP.

##### Diagnosis

**(1)** Male right epimeral plates prominent, as projections directed posteriorly; **(2)** male right antennule actual segment 13 with modified seta perpendicular to antennule direction, and length reaching to the middle-point of the sequence segment; **(3)** left antennule actual segment 11 with two setae; **(4)** male fifth left swimming leg length reaching to first right exopod segment distally; **(5)** male fifth right swimming leg exopod 1 longer than broad; **(6)** female left epimeral plate with semicircular expansion on posterior surface dorsally; **(7)** female second urosome segment without ventral fusion to anal segment; **(8)** female right antennule not extending beyond caudal rami; **(9)** female fifth swimming legs basis with seta longer 2x than origin segment.

##### Redescription

###### MALE

Body 1134 micrometers excluding caudal setae. Male body smaller and slenderer than female. Nerve axons myelinated. Prosome 6-segmented; widest at first metasome segment; without one line of setules at posterior margin; without spinules at segments. Cephalosome anterior margin rounded; with dorsal suture; incomplete; separate from first metasome segment. First metasome segment without sensilla. Second metasome segment without sensilla. Third metasome segment without sensillae; non-ornamented posterior margin. Fourth metasome segment without sensillae; separated from the fifth metasome. Limit between fourth and fifth metasome segments without ornamentation. Fifth metasome segment without sensilla; Fifth metasome segment without ornamentation; Fifth metasome segment without dorsal conical process; with epimeral plates. Epimeral plates symmetrical. Right epimeral plates prominent, as projections; one projection; posterior-laterally directed; reaching half or surpassing the length of genital segment; with sensilla; at the apex of projection; without ornamentation.

##### Urosome

5-segmented; Urosome 5 - free segments. Genital somite symmetrical in dorsal view; with single aperture; located on left side; ventrolaterally on posterior rim; without sensillae. Third urosome segment without spinules; without external seta. Fourth urosome segment without spinules; without sub-conical blunt dorsal-lateral process. Anal segment absence of dorsal sensillae; presence of operculum; convex; covering the anal aperture fully. Caudal rami symmetrical; separated from anal segment; longer than wide; with setules; continuous on; inner side; each ramus bearing 6 caudal setae; 5 marginals; plumose; and 1 internal dorsally; straight; not reticulated main axis; outermost seta with outer spiniform process absent.

##### Appendices features

Rostrum symmetrical; separated from dorsal cephalic shield; by complete suture; sensillae present; one pair; anteriorly inserted on surface tegument; with rostral filament; double; paired; extended; into point; with basal process; in ventral view, rounded on left side; without a smaller basal expansion on the right side.

##### Antennules

Asymmetrical. **Right antennules**. Uniramous; right antennule surpassing to genital segment; right antennule not extending beyond caudal rami.

Right antennule ancestral segment I and II separated. Ancestral segment II and III fused. Ancestral segment III and IV fused. Ancestral segment IV and V separated. Ancestral segment V and VI separated. Ancestral segment VI and VII separated. Ancestral segment VII and VIII separated. Ancestral segment VIII and IX separated. Ancestral segment IX and X separated. Ancestral segment X and XI separated. Ancestral segment XI and XII separated. Ancestral segment XII and XIII separated. Ancestral segment XIII and XIV separated. Ancestral segment XIV and XV separated. Ancestral segment XV and XVI separated. Ancestral segment XVI and XVII separated. Ancestral segment XVII and XVIII separated. Ancestral segment XVIII and XIX separated. Ancestral segment XIX and XX separated. Ancestral segment XX and XXI separated. Ancestral segment XXI and XXII fused. Ancestral segment XXII and XXIII fused. Ancestral segment XXIII and XXIV separated. Ancestral segment XXIV and XXV fused. Ancestral segment XXV and XXVI separated. Ancestral segment XXVI and XXVII separated. Ancestral segment XXVII and XXVIII fused.

Right antennule actual 22-segmented; geniculated; between the segment 18 and segment 19; with swollen and modified region; formed by 5 segments; between 13 and 17 segments. Actual segment 1 with seta; one element; straight; none larger than segment; without spinules; without vestigial seta; without conical seta; without modified seta; without spinous process; with aesthetasc; one element. Actual segment 2 with seta; three elements; of unequal size; straight; none larger than segment; without spinules; with vestigial seta; one element; without conical seta; without modified seta; without spinous process; with aesthetasc; one element. Actual segment 3 with seta; one element; one larger than segment; surpassing to distal margin; beyond three sequential segments; straight; blunt apex; without spinules; with vestigial seta; one element; without conical seta; without modified seta; without spinous process; with aesthetasc. Actual segment 4 with seta; one element; one larger than segment; surpassing to distal margin; straight; not beyond three sequential segments; without spinules; without vestigial seta; without conical seta; without modified seta; without spinous process; without aesthetasc. Actual segment 5 with seta; one element; straight; one larger than segment; surpassing to distal margin; not beyond three sequential segments; without spinules; with vestigial seta; one element; without conical seta; without modified seta; without spinous process; with aesthetasc; one element. Actual segment 6 with seta; one element; none larger than segment; straight; without spinules; without vestigial seta; without conical seta; without modified seta; without spinous process; without aesthetasc. Actual segment 7 with seta; one element; straight; one larger than segment; surpassing to distal margin; beyond three sequential segments; blunt apex; without spinules; without vestigial seta; without conical seta; without modified seta; without spinous process; with aesthetasc; one element. Actual segment 8 with seta; one element; straight; none larger than segment; without spinules; without vestigial seta; with conical seta; one element; not reaching to middle-point of the sequent segment; without modified seta; without spinous process; without aesthetasc. Actual segment 9 with seta; two elements; of unequal size; straight; one larger than segment; surpassing to distal margin; beyond three sequential segments; blunt apex; without spinules; without vestigial seta; without conical seta; without modified seta; without spinous process; with aesthetasc; one element. Actual segment 10 with seta; one element; straight; none larger than segment; without spinules; without vestigial seta; without conical seta; with modified seta; presenting blunt apex; slender form; surpassing to distal margin; beyond of the sequential segment; parallel to antennule direction; without spinous process; without aesthetasc. Actual segment 11 with seta; one element; straight; one larger than segment; surpassing to distal margin; not beyond three sequential segments; without spinules; without vestigial seta; without conical seta; with modified seta; slender form; presenting blunt apex; surpassing to distal margin; beyond of the sequential segment; parallel to antennule direction; shorter length than homologous of actual segment 13; without spinous process; without aesthetasc. Actual segment 12 with seta; one element; straight; one larger than segment; surpassing to distal margin; not beyond three sequential segments; without spinules; without vestigial seta; with conical seta; one element; not smaller than to segment 8; without modified seta; without spinous process; with aesthetasc; one element; absent internal perpendicular fission. Actual segment 13 with seta; one element; straight; one larger than segment; surpassing to distal margin; not beyond three sequential segments; without spinules; without vestigial seta; without conical seta; with modified seta; stout form; surpassing to distal margin; to the middle-point of the sequence segment; perpendicular to antennule direction; presenting bifid apex; without spinous process; with aesthetasc; one element. Actual segment 14 with seta; two elements; of unequal size; straight; one larger than segment; surpassing to distal margin; beyond three sequential segments; blunt apex; without spinules; without vestigial seta; without conical seta; without modified seta; without spinous process; with aesthetasc; one element. Actual segment 15 with seta; two elements; of unequal size; straight; not bifidform; none larger than segment; without spinules; without vestigial seta; without conical seta; without modified seta; with spinous process; on outer margin; surpassing distal margin; with aesthetasc; one element. Actual segment 16 with seta; two elements; of unequal size; plumose; one larger than segment; surpassing to distal margin; not beyond three sequential segments; not bifidform; without spinules; without vestigial seta; without conical seta; without modified seta; with spinous process; on outer margin; surpassing distal margin; unequal size to process on preceding segment; with aesthetasc; one element. Actual segment 17 with seta; two elements; of unequal size; straight; none larger than segment; bifidform; without spinules; without vestigial seta; without conical seta; with modified seta; one element; stout form; surpassing to distal margin; not beyond of the sequential segment; parallel to antennule direction; without spinous process; without aesthetasc. Actual segment 18 with seta; two elements; of equal size; straight; none larger than segment; without spinules; without vestigial seta; without conical seta; with modified seta; one element; stout form; surpassing distal margin; parallel to antennule direction; without spinous process; without aesthetasc. Actual segment 19 with seta; two elements; of unequal size; plumose; none larger than segment; without spinules; without vestigial seta; without conical seta; with modified seta; two elements; stout form; at least one bifid form; surpassing distal margin; parallel to antennule direction; without spinous process; with aesthetasc; one element. Actual segment 20 with seta; four elements; of unequal size; straight; one larger than segment; surpassing to distal margin; beyond three sequential segments; without spinules; without vestigial seta; without conical seta; without modified seta; without spinous process; without aesthetasc. Actual segment 21 with seta; two elements; of equal size; plumose; one larger than segment; surpassing to distal margin; greater 3x than original segment; without spinules; without vestigial seta; without conical seta; without modified seta; without spinous process; without aesthetasc. Actual segment 22 with seta; four elements; of equal size; one larger than segment; plumose; surpassing to distal margin; greater 3x than original segment; without spinules; without vestigial seta; without conical seta; without modified seta; without spinous process; with aesthetasc; one element.

##### Left antennules

Uniramous; Left antennule surpassing to prosome; Left antennule not extending beyond caudal rami. Ancestral segment I and II separated. Ancestral segment II and III fused. Ancestral segment III and IV fused. Ancestral segment IV and V separated. Ancestral segment V and VI separated. Ancestral segment VI and VII separated. Ancestral segment VII and VIII separated. Ancestral segment VIII and IX separated. Ancestral segment IX and X separated. Ancestral segment X and XI separated. Ancestral segment XI and XII separated. Ancestral segment XII and XIII separated. Ancestral segment XIII and XIV separated. Ancestral segment XIV and XV separated. Ancestral segment XV and XVI separated. Ancestral segment XVI and XVII separated. Ancestral segment XVII and XVIII separated. Ancestral segment XVIII and XIX separated. Ancestral segment XIX and XX separated. Ancestral segment XX and XXI separated. Ancestral segment XXI and XXII separated. Ancestral segment XXII and XXIII separated. Ancestral segment XXIII and XXIV separated. Ancestral segment XXIV and XXV separated. Ancestral segment XXV and XXVI separated. Ancestral segment XXVI and XXVII separated. Ancestral segment XXVII and XXVIII fused.

Left antennule actual 25-segmented; not-geniculated. Actual segment 1 with seta; one element; none larger than segment; straight; without spinules; without vestigial seta; without conical seta; without modified seta; without spinous process; with aesthetasc; one element. Actual segment 2 with seta; three elements; of equal size; none larger than segment; straight; without spinules; with vestigial seta; one element; without conical seta; without modified seta; without spinous process; with aesthetasc; one element. Actual segment 3 with seta; one element; one larger than segment; straight; surpassing to distal margin; beyond three sequential segments; without spinules; with vestigial seta; one element; without conical seta; without modified seta; without spinous process; with aesthetasc. Actual segment 4 with seta; one element; none larger than segment; straight; without spinules; without vestigial seta; without conical seta; without modified seta; without spinous process; without aesthetasc. Actual segment 5 with seta; one element; one larger than segment; straight; surpassing to distal margin; not beyond three sequential segments; without spinules; with vestigial seta; one element; without conical seta; without modified seta; without spinous process; with aesthetasc; one element. Actual segment 6 with seta; one element; none larger than segment; straight; without spinules; without vestigial seta; without conical seta; without modified seta; without spinous process; without aesthetasc. Actual segment 7 with seta; one element; one larger than segment; straight; surpassing to distal margin; beyond three sequential segments; without spinules; without vestigial seta; without conical seta; without modified seta; without spinous process; with aesthetasc; one element. Actual segment 8 with seta; one element; one larger than segment; straight; surpassing distal margin; without spinules; without vestigial seta; with conical seta; without modified seta; without spinous process; without aesthetasc. Actual segment 9 with seta; two elements; of unequal size; one larger than segment; straight; surpassing to distal margin; beyond three sequential segments; without spinules; without vestigial seta; without conical seta; without modified seta; without spinous process; with aesthetasc; one element. Actual segment 10 with seta; one element; none larger than segment; straight; without spinules; without vestigial seta; without conical seta; without modified seta; without spinous process; without aesthetasc. Actual segment 11 with seta; two elements; of unequal size; one larger than segment; straight; surpassing to distal margin; beyond three sequential segments; without spinules; without vestigial seta; without conical seta; without modified seta; without spinous process; without aesthetasc. Actual segment 12 with seta; one element; one larger than segment; straight; surpassing distal margin; without spinules; without vestigial seta; with conical seta; without modified seta; without spinous process; with aesthetasc; one element. Actual segment 13 with seta; one element; none elongated; straight; surpassing distal margin; without spinules; without vestigial seta; without conical seta; without modified seta; without spinous process; without aesthetasc. Actual segment 14 with seta; one element; elongated; straight; surpassing to distal margin; beyond three sequential segments; without spinules; without vestigial seta; without conical seta; without modified seta; without spinous process; with aesthetasc; one element. Actual segment 15 with seta; one element; larger than segment; straight; surpassing to distal margin; not beyond three sequential segments; without spinules; without vestigial seta; without conical seta; without modified seta; without spinous process; without aesthetasc. Actual segment 16 with seta; one element; larger than segment; plumose; surpassing to distal margin; not beyond three sequential segments; without spinules; without vestigial seta; without conical seta; without modified seta; without spinous process; with aesthetasc; one element. Actual segment 17 with seta; one element; not larger than segment; straight; without spinules; without vestigial seta; without conical seta; without modified seta; without spinous process; without aesthetasc. Actual segment 18 with seta; one element; larger than segment; straight; surpassing to distal margin; beyond three sequential segments; without spinules; without vestigial seta; without conical seta; without modified seta; without spinous process; without aesthetasc. Actual segment 19 with seta; one element; not larger than segment; straight; surpassing distal margin; without spinules; without vestigial seta; without conical seta; without modified seta; without spinous process; with aesthetasc; one element. Actual segment 20 with seta; one element; not larger than segment; straight; surpassing distal margin; without spinules; without vestigial seta; without conical seta; without modified seta; without spinous process; without aesthetasc. Actual segment 21 with seta; one element; larger than segment; plumose; surpassing to distal margin; beyond three sequential segments; without spinules; without vestigial seta; without conical seta; without modified seta; without spinous process; without aesthetasc. Actual segment 22 with seta; two elements; of unequal size; one of them elongated; plumose; surpassing to distal margin; without spinules; without vestigial seta; without conical seta; without modified seta; without spinous process; without aesthetasc. Actual segment 23 with seta; two elements; of unequal size; one larger than segment; plumose; surpassing to distal margin; greater 3x than original segment; without spinules; without vestigial seta; without conical seta; without modified seta; without spinous process; without aesthetasc. Actual segment 24 with seta; two elements; of equal size; one larger than segment; plumose; surpassing to distal margin; greater 3x than original segment; without spinules; without vestigial seta; without conical seta; without modified seta; without spinous process; without aesthetasc. Actual segment 25 with seta; four elements; of equal size; elongated; plumose; surpassing to distal margin; 4 times larger than segment; without spinules; without vestigial seta; without conical seta; without modified seta; without spinous process; with aesthetasc; one element.

##### Antenna

Biramous. Antenna coxa separated from the basis; bearing seta; 1; on inner surface; at distal corner; reaching to the endopod 1. Antenna basis (fusion) separated from the endopodal segment; bearing seta; 2; on inner surface; at distal corner. Endopodal ancestral segment I and II separated. Ancestral segment II and III fused. Ancestral segment III and IV fused. Ancestral segment III and IV fully. Antenna endopod actual 2-segmented. Actual segment 1 not bilobate; with seta; two; on inner margin; with spinules; as a row; obliquely; on outer surface; with pore. Actual segment 2 bilobate; with discontinuity on outer cuticle; not developed as a suture; inner lobe bearing 8 setae; distally; outer lobe bearing 7 setae; distally; with spinules; as a patch; on outer surface. Antenna exopod ancestral segment I and II separated. Ancestral segment II and III fused. Ancestral segment III and IV fused. Ancestral segment IV and V separated. Ancestral segment V and VI separated. Ancestral segment VI and VII separated. Ancestral segment VII and VIII separated. Ancestral segment VIII and IX separated. Ancestral segment IX and X fused. Antenna exopod actual 7-segmented. Actual segment 1 single; elongated (width-length, equal or larger ratio 2:1); with seta; one; at inner surface. Actual segment 2 compound; elongated (larger width-length ratio 2:1); with seta; three; at inner surface. Actual segment 3 single; not elongated (lesser width-length ratio 2:1); with seta; one; at inner surface. Actual segment 4 single; not elongated (lesser width-length ratio 2:1); with seta; one; at inner surface. Actual segment 5 single; not elongated (lesser width-length ratio 2:1); with seta; one; at inner surface. Actual segment 6 single; not elongated (lesser width-length ratio 2:1); with seta; one; at inner surface. Actual segment 7 compound; elongated (larger or equal width-length ratio 2:1); with seta; one; at inner surface; and three; at distal surface.

##### Oral features

**Mandible**. Coxal gnathobase sclerotized; with lobe; prominent; on caudal margin; presence of cutting blade; with tooth-like prominence; two, distinctly; 1 acute; on caudal margin; and 1 triangular; on sub-caudal margin; without acute projection between the prominences; with additional spinules; as a row; on dorsal surface; with seta; 1; dorsally; on apical surface; with spinules; apicalmost. Mandible palps biramous; comprising the basis; with seta; four; differently inserted; first medially; reaching to beyond the endopod 1; second distally; third distally; fourth distally; on inner margin; none with setulose ornamentation. Mandible endopod 2-segmented. Mandible endopod 1 with lobe; bearing seta; four; distally inserted; without spinules. Mandible endopod 2 without lobe; bearing setae; nine elements; distally inserted; with spinules; as a row; double. Mandible exopod 4-segmented. Mandible exopod 1 with seta; one element; distally; on inner margin. Mandible exopod 2 with seta; one element; distally; on inner side. Mandible exopod 3 with seta; one element; distally; on inner side. Mandible exopod 4 with setae; three elements; on terminal region. **Maxillule**. Birramous. Maxillule 3-segmented. Maxillule praecoxa with praecoxal arthrite; bearing spines; fifteen elements; ten marginally; plus, five sub-marginally; with spinules; as a patch; on sub-marginal surface. Maxillule coxa with coxal epipodite; with conspicuous outer lobe; bearing setae; nine elements; with coxal endite; elongated (larger or equal width-length ratio 2:1); bearing setae; four elements. Maxillule basis with basal endite; double; first proximal; elongated (larger width-length ratio 2:1; separated from basis; with setae; four elements; distally inserted; second distal; fused to basis; not elongated (lesser width-length ratio 2:1); with setae; four elements; distally inserted; with setules; as a row; on inner side; basal exite present; with setae; one element; on outer surface. Maxillule endopod 1-segmented. Endopod 1 bilobate; first proximal; with setae; three elements; second distal; with setae; five elements. Maxillule exopod 1-segmented. Exopod 1 with setae; six elements; with setules; as a row; on inner side; spinules absent. **Maxilla**. Uniramous. Maxilla 5-segmented. Maxilla praecoxa fused to coxa; incompletely; distinct externally; with praecoxal endite; double; first elongated endite (larger or equal width length ratio 2:1); proximally inserted; with seta; straight, or plumose; 1 straight; 4 plumose; with spine; single; without spinules; without setule; second elongated endite (larger or equal width length ratio 2:1); distally inserted; with seta; plumose; 3 plumose; without spine; with spinules; as a row; on distal margin; with setule; as a row; on distal margin; absence of outer seta. Maxilla coxa with coxal endite; double; first elongated endite (larger or equal width); proximally inserted; with seta; plumose; 3 plumose; without spine; without spinules; with setules; as a row; on proximal margin; second elongated endite (larger or equal width); distally inserted; with seta; plumose; 3 plumose; without spine; without spinules; with setules; as a row; on proximal margin; absence of outer seta. Maxilla basis with basal endite; single; elongated (larger or equal width-length ratio 2:1); with seta; plumose; 3 plumose; without spinules; absence of outer seta. Maxilla endopod 2-segmented. Endopod 1 with seta; 2 plumose; without spine; without spinules; without setules. Maxilla endopod 2 with seta; 2 plumose; without spine; without spinules; without setules. **Maxilliped**. Uniramous; Maxilliped 8-segmented. Maxilliped praecoxa fused to coxa; incompletely; distinct internally; with praecoxal endite; not elongated (lesser width-length ratio 2:1); distally inserted; with seta; 1 straight; with spinules; as a row; single; on basal surface; without setules. Maxilliped coxa with coxal endite; three coxal endite; first elongated (larger or equal width); proximally inserted; with seta; 2 plumose; with spinules; as a patch; single; on apical surface; without setules; second not elongated (lesser width-length ratio 2:1); medially inserted; with seta; 3 plumose; with spinules; as a row; single; on medial surface; without setules; third elongated (larger or equal width length ratio 2:1); distally inserted; with seta; 3 plumose; none reaching to beyond of the basis; with spinules; as a row; single; on basal surface; without setules; with lobe; prominence; at inner distal angle; ornamented; with spinules; continuously on margin. Maxilliped basis without basal endite; with seta; 3 plumose; with spinules; as a row; single; on medial surface; with setules; as a row; single; on inner margin. Maxilliped endopod segment 6-segmented. Endopod 1 with seta; 2 plumose; on inner surface. Endopod 2 with seta; 3 plumose; on inner surface. Endopod 3 with seta; 2 plumose; on inner surface. Endopod 4 with seta; 2 plumose; on inner surface. Endopod 5 with seta; 2 plumose; on inner surface, or on outer surface; outer seta absent. Endopod 6 with seta; 4 plumose; on inner surface, or on outer surface.

##### Swimming legs features

**First swimming legs.** Symmetrical; biramous. First swimming legs intercoxal plate without seta. First swimming legs praecoxa absent. First swimming legs coxa with seta; one; straight; distally inserted; on inner surface; surpassing to first endopodal segment; with setules; two group; as a patch; on inner margin; and as a row; double; on anterior surface; outerly; without spinules; without spine. First swimming legs basis without seta; with setules; as a patch; single; on outer surface; without spinules; without spine. First swimming legs endopod 2-segmented. Endopod 1 with seta; straight; restricted; to inner surface; one element; without spine; with setules; as a row; single; continuously; on outer surface; without spinules; absence of Schmeil’s organ. Endopod 2 with seta; unrestricted; three on inner surface; one on outer surface; two on distal surface; straight; without spine; with setules; as a row; single; continuously; on outer surface; without spinules; absence of Schmeil’s organ. Endopod 3 absence. First swimming legs exopod 1 with seta; restricted; 1 on inner surface; with spine; 1; stout; smaller than original segment; serrated; on inner side; continuously; with setules; as a row; single; as a row; innerly. First swimming legs exopod 2 with seta; restricted; 1 on inner surface; straight; without spine; with setules; as a row; single; continuously; on inner margin, or on outer margin; without spinules. First swimming legs exopod 3 with setule; as a row; single; continuously; on outer surface; without spinules; with seta; unrestricted; 2 on inner surface; 2 on terminal surface; with spine; 2; unequal size; first no longer 2x than origin segment; stout; serrated; on inner side, or on outer side; equally; second longer 3x than origin segment; slender; serrated; on outer side; with ornamentation on non-serrated side; by setules. **Second swimming legs**. Symmetrical; Second swimming legs biramous. Second swimming legs intercoxal plate without seta. Second swimming legs praecoxa present; located laterally. Second swimming legs coxa with seta; straight; distally inserted; on inner surface; surpassing to basal segment; without setules; without spinules; without spine. Second swimming legs basis without seta; without setules; without spinules; without spine. Second swimming legs endopod 3-segmented. Endopod 1 with seta; straight; restricted; one on inner surface; without spine; with setules; as a row; single; continuously; on outer surface; without spinules; absence of Schmeil’s organ. Endopod 2 with seta; straight; unrestricted; two on inner surface; without spine; with setules; as a row; single; continuously; on outer side; without spinules; presence of Schmeil’s organ; on posterior surface. Endopod 3 with seta; straight; unrestricted; three on inner surface; two on outer surface; two on distal surface; without spine; without setules; with spinules; as a row; double; distally inserted; at anterior surface; absence of Schmeil’s organ. Second swimming legs exopod 1 with seta; restricted; one on inner surface; with spine; 1; stout; not reaching to distal-third of the exopod 2; serrated; on inner side, or on outer side; with setules; as a row; single; continuously; on inner side; without spinules; absence of Schmeil’s organ. Exopod 2 with seta; unrestricted; one on inner surface; with spine; 1; stout; not surpassing the exopod 3; serrated; on inner side, or on outer side; with setules; as a row; single; continuously; on inner surface; without spinules; absence of Schmeil’s organ. Exopod 3 with seta; plurimarginal; three on inner surface; two on terminal surface; with spine; 2; unequal size; first no longer 2x than origin segment; stout; serrated; on inner side, or on outer side; equally; second longer 2x than origin segment; slender; serrated; on outer side; with ornamentation on non-serrated side; of setules; setules on outer surface; as a row; single; continuously; on inner surface; with spinules; as a row; single; distally inserted; at anterior surface; absence of Schmeil’s organ. **Third swimming legs**. Symmetrical; Third swimming legs biramous. Third swimming legs intercoxal plate without seta. Third swimming legs praecoxa present; not laterally located. Third swimming legs coxa with seta; straight; distally inserted; on inner surface; surpassing to first endopodal segment; without setules; without spinules; without spine. Third swimming legs basis without seta; without setules; without spinules; without spine. Third swimming legs endopod 3-segmented. Endopod 1 with seta; restricted; one on inner surface; without spine; without setules; without spinules; absence of Schmeil’s organ. Endopod 2 with seta; restricted; two on inner surface; straight; without spine; without setules; without spinules; absence of Schmeil’s organ. Endopod 3 with seta; straight; plurimarginal; two on inner surface; two on outer surface; three on terminal surface; without spine; without setules; with spinules; as a row; distally inserted; double; at anterior surface; absence of Schmeil’s organ. Third swimming legs exopod 1 with seta; restricted; straight; one on inner surface; with spine; 1; stout; not reaching to the distal-third of the exopod 2; serrated; equally; on inner surface, or on outer surface; with setules; as a row; single; continuously; on inner surface; without spinules; absence of Schmeil’s organ. Exopod 2 with seta; straight; restricted; one on inner surface; with spine; 1; stout; not reaching out to exopod 3; serrated; on inner side, or on outer side; equally; with setules; as a row; single; continuously; on inner side; without spinules; absence of Schmeil’s organ. Exopod 3 without setules; with spinules; as a row; single; distally inserted; at anterior surface; with seta; straight; unrestricted; three on inner surface; two on terminal surface; with spine; 2; unequal size; first no longer 2x than origin segment; stout; serrated; on inner side, or on outer side; equally; second longer 2x than origin segment; slender; serrated; on outer side; with ornamentation on non-serrated side; of setules; absence of Schmeil’s organ. **Fourth swimming legs**. Symmetrical; biramous. Intercoxal plate without sensilla. Praecoxa present. Coxa with seta; distally inserted; on inner margin; reaching out to endopod 1; without spinules; setules absent. Basis with seta; one; medially inserted; on posterior surface; smaller than the original segment; without setules; without spinules; without spine. Fourth swimming legs endopod 3-segmented. Endopod 1 with seta; one; restricted; on inner surface; without spine; without setules; without spinules; absence of Schmeil’s organ. Endopod 2 with seta; restricted; two on inner side; without spine; with setules; as a row; single; continuously; on outer surface; without spinules; absence of Schmeil’s organ. Endopod 3 with seta; unrestricted; two on inner surface; two on outer surface; three on distal surface; without spine; without setules; with spinules; as a row; double; distally inserted; at anterior surface; absence of Schmeil’s organ. Fourth swimming legs exopod 1 with seta; restricted; one on inner surface; with spine; 1; stout; not reaching out to distal-third of the exopod 2; serrated; on inner side, or on outer side; equally; with setules; as a row; single; continuously; on inner surface; without spinules; absence of Schmeil’s organ. Exopod 2 with seta; restricted; one on inner surface; with spine; 1; stout; not reaching the end of exopod 3; serrated; on inner side, or on outer side; equally; with setules; as a row; single; continuously; on inner surface; without spinules; absence of Schmeil’s organ. Exopod 3 without setules; with spinules; as a row; single; distally inserted; at anterior surface; with seta; unrestricted; three on inner surface; two on distal surface; with spine; 2; unequal size; first no longer 2x than origin segment; stout; serrated; on inner side, or on outer side; equally; second longer 2x than origin segment; slender; serrated; on outer side; without ornamentation on non-serrated side; absence of Schmeil’s organ.

##### Fifth swimming legs features

Asymmetrical. Fifth swimming leg intercoxal plate with length not equal or greater than width on 1.5x; with irregular proximal margin; discontinuous to; the anterior margin of the left coxa, or the anterior margin of the right coxa; posterior sensilla on the right lateral absent. **Fifth left swimming leg**. Fifth left swimming leg biramous; leg reaching first right exopod segment; distally. Fifth left swimming leg praecoxa present; rudimentary; separated from the coxae; without ornamentation. Fifth left swimming leg coxa concave inner side; without teeth-like structures; with process; conical; on posterior surface; outer side; distally inserted; not projecting over basis; with sensilla; stout; triangular; at apex; no longer 2x than insertion basis; with swelling; on inner side; distally; without seta; without spinules. Fifth left swimming leg basis sub-cylindrical; unequal size between inner and outer side; shorter inner than outer side; with rectilinear inner side; rounded internal proximal expansion absent; without outgrowth; with groove; deep; obliquely; on posterior surface; not reaching the endopodal lobe; not ornamented; absence of protuberance; with seta; outerly inserted; no longer 2x than origin segment; absence of minutely granular. Fifth left swimming leg endopod segments 1 and 2 fused; segments 2 and 3 fused; 1-segmented; stout; separated from the basis; ornamented; on inner side; with spinules; more than four elements; as a row; terminally; row of setules absent; without seta. Fifth left swimming leg exopod segments 1 and 2 separated; segments 2 and 3 fused; 2-segmented; stout; separated from the basis. Fifth left swimming leg exopod 1 sub-triangular; longer than broad; equal size between inner and outer side; rectilinear inner side; convex outer side; without swelling; without marginal extension; without process; with lobe; single; circular; medially inserted; on inner side; covered; by setules; without outer spine; absence seta. Fifth left swimming leg exopod 2 sub-triangular; longer than broad; equal size between inner and outer side; disform inner side; with rectilinear outer side; setulose pad present; not prominently rounded; proximally; on inner side; inflated medial region present; setulose; anteriorly; distal process present; digitiform; non denticulate; without transverse row of denticles; none oblique row of 5 denticles; innerly directed; with seta; spiniform; ornamented by spinules; surpassing the distal-point of the segment; without outer spine; terminal claw absent.

##### Fifth right swimming leg

Biramous. Fifth right swimming leg praecoxa absent. Fifth right swimming leg coxa convex inner side; without teeth-like structures; with process; rounded; distally inserted; on posterior surface; closest to the outer rim; projecting over basis; beyond the first third; until the medial surface; without triangular protuberance innerly; with sensilla; slender; at apex; no longer 2x than basal insertion; without marginal extension; without seta; without spinules. Fifth right swimming leg basis cylindrical; unequal size between inner and outer side; shorter outer than inner side; rectilinear inner side; tumescence present; not inflated; restricted on inner surface; proximally; without protuberance; absence of distinct minutely granular; additional inner process absent; with posterior groove; deep; obliquely; not reaching the endopodal lobe; ornamented; with tubercles; throughout of the outer border; with seta; outerly inserted; on anterior surface; no longer 2x than origin segment; posterior protrusion present; distal process absent. Fifth right swimming leg with endopodite present; fused to basis; on anterior surface; ancestral segments 1 and 2 fused; ancestral segments 2 and 3 fused; stout; ornamented; with setules; as a row; on inner side; terminally; without seta. Fifth right swimming leg exopod segments 1 and 2 separated; segments 2 and 3 fused; 2-segmented; stout; separated from the basis. Fifth right swimming leg exopod 1 sub-cylindrical; longer than broad; nearly 1.25 times; unequal size between both sides; shorter inner than outer side; convex inner side; rectilinear outer side; with marginal extension; sub-triangular; distally inserted; at outer rim; spinules absent; with process; triangular; arched; internally directed; sharp tip; sclerotized; without ornamentation; distally inserted; at posterior surface; projecting over next segment; without outer spine; without seta; internal prominence absent; lamella on posterior surface absent. Fifth right swimming leg exopod 2 cylindrical; longer than broad; nearly 2.5 times; equal size between both sides; uniform inner side; convex outer side; without posterior proximal swelling; inner-posterior process absent; without marginal expansion; curved ridge on distal posterior surface present; chitinous knobs absent; with outer spine; inserted sub-distally; arched; internally directed; not ornamented innerly; not ornamented outerly; sharp tip; with apparent curve; innerly directed; lesser than the length of the exopod 2; beyond to 2 times its size; 3x; sensilla absent; terminal claw present; equal or longer 1.5 times than insertion segment; sclerotized; arched; inward; with conspicuous curve; proximally, or medially; ornamented innerly; by spinules; as a row; throughout extension; not ornamented outerly; sharp tip; not curved tip; with medial constriction; hyaline process absent.

##### FEMALE

Body longer and wider than male. Widest at first metasome segment. Distal margin of the prosomal segments without one line of setules at posterior margin. Prosome segments without spinules at prosomal segments. Fourth metasome segment absence of dorsal protuberance. Fourth and fifth metasome segments fused; partially; on dorsal surface. Limit between fourth and fifth metasome segments without ornamentation. **Fifth metasome segment**. Fifth metasome segment with sensilla; laterally; 2 elements; with epimeral plates. Epimeral plates asymmetrical. Right epimeral plates prominent, as projections; thinner than the left; one posterior-laterally directed; not reaching half length of the genital segment; with sensilla at the apex; dorsal-posterior sensilla present; slender; without ornamentation. Left epimeral plate with expansion; semicircular; on posterior surface; dorsally; with sensilla; at base.

##### Urosome

3-segmented. **Genital double-somite**. Asymmetrical in dorsal view; longer than broad; longer than other urosomites combined; dorsal suture at mid-length absent; not covered by spinules; with swelling; rounded; unequal size; greater right than left; anteriorly; with sensillae; on both sides; one; stout; with robust apex; at left lateral; not on lobular base; medially; one; stout; at right lateral; not on lobular base; anteriorly; with robust apex; of equal size between then; lateral protuberance absent; with right posterior rim expanded; over next segment; without slender sensilla on each posterior rim; without posterior-dorsal process. Genital double-somite opercular pad present; broader than longer; symmetrical; development laterally; expanded posteriorly; covering partially; double gonoporal slit; located ventrally; with arthrodial membrane; inserted anteriorly; post-genital process absent; disto-ventral tumescence absent; ventral vertical folds absent; dorsal sensilla absent. Second urosome segment without ventral fusion to anal segment; right distal process absent. Caudal rami patch of setules on outer surface absent; patch of spinules on outer surface absent.

##### Appendices features

Rostrum basal process absent. **Antennules**. Symmetrical. Right antennule surpassing to genital double-segment; extending beyond caudal rami; not exceeding the caudal setae; ornamentation pattern equals to male left antennule; fully.

##### Fifth swimming legs

Symmetrical; Fifth swimming legs biramous. Fifth swimming legs intercoxal plate longer than wide; separated from the legs. Fifth swimming legs praecoxa with sclerite praecoxal; separated from the coxae; without ornamentation. Fifth swimming legs coxa with process; conical; at the outer rim; distally; sensilla present; stout; at apex; projecting over basal segment; no longer 2x than basal insertion; marginal extension absent; without swelling; without seta; without spinules. Fifth swimming legs basis sub-triangular; unequal size between inner and outer sides; shorter outer than inner side; with convex inner side; without proximal inner outgrowth; without groove; with distal extension; on posterior surface; with seta; outerly inserted; on anterior surface; longer 2x than origin segment; reaching to exopod 1 distally. Fifth swimming legs endopod segments 1 and 2 fused; segments 2 and 3 fused; 1-segmented; stout; separated from the basis; present discontinuity cuticle; on inner side; with spinules; as a row; single; non-oblique; sub-terminally; at anterior surface; with seta; double; one medially; on posterior surface; rectilinear; one distally; on posterior surface; rectilinear; of unequal size; distal seta longer than medial seta. Fifth swimming legs exopod segments 1 and 2 separated; segments 2 and 3 separated; 3-segmented; separated from the basis. Fifth swimming legs exopod 1 sub-cylindrical; longer than wide; longer or equal than 2 times; with unequal size between inner and outer side; shorter inner than outer side; with convex inner side; with rectilinear outer side; without swelling; without marginal extension; without posterior process; without spine; without seta. Fifth swimming legs exopod 2 sub-cylindrical; longer than broad; longer or equal than 2 times; without swelling; without marginal extension; without process; without lobe; with spine; inserted laterally; rectilinear; without ornamentation; sharp tip; equal size or larger than next segment; without seta. Fifth swimming legs exopod 3 cylindrical; longer than wide; without swelling; without process; without lobe; without spine; with seta; double; inserted terminally; unequal size between them; outer seta smaller than inner; nearly 3 times; outer seta not ornamented by setules; without ornamentation; presence of terminal claw; sclerotized; arched; externally directed; convex inner side; with ornamentation; of denticles; as a row; on surface partially; at medial region; concave outer side; with ornamentation; of denticles; as a row; on surface partially; at medial region; blunt tip; 6 times longer than origin segment.

##### Distribution records

###### VENEZUELA

**Anzoategui**: Orinoco River, left margin, in Soledad; Charca 2, near to Unaré River, in Clarines (Dussart, 1984a). **Aragua**: artificial lagoon in Camatagua (Dussart, 1984a). **Bolivar**: Guri, artificial lagoon near to dam at the Caroni River (Dussart, 1984a). **Delta Amacuro**: Caño Manamo near to the Tucupita (Dussart, 1984a). **Guarico**: Guarico Reservoir, near to Calobozo; Caño Falcon, Portuguesa River, near to San Fernando de Apure; Los Patos Pond (natural), near to Calobozo biological station; pond (natural) near to El Sombrero (Dussart, 1984a). **Monagas**: pond between Barcelona, and Maturin, near to Urica; surroundings to Barrancas (Bowman, 1973); Orinoco River in Barrancas (Dussart, 1984a). BRAZIL. **Maranhão**: lagoons between dunes, Lençóis Maranhenses Park (Rocha *et al*., 1998). **Ceará**: lagoon Tauapé, Fortaleza, and lagoon Mecejana, Mecejana (Wright, 1936); ponds in Fortaleza City (Matsumura-Tundisi, 1986). **Rio Grande do Norte**: several bodies of water near to Caraúbas and other near to Assú (Wright, 1936). **Paraíba**: Pilões Pond, near to São João do Rio do Peixe (Wright, 1936). **Pernambuco**: “ponds” (Matsumura-Tundisi, 1986). **Minas Gerais**: Paraiba River at the Emborcação Reservoir. **São Paulo**: Barra Bonita Reservoir, Tietê River (Tundisi & Matsumura-Tundisi, 1994).

##### Habitat

Habitat in freshwaters: artificial lakes, shallow gully, and sand dune lakes.

##### Remarks

**(1)** *N. cearensis* was founded and included in the *nordestinus* group in 1936 from organisms of unspecified locality of Ceará, Brazil. In the same year Kiefer (1936) included the taxon in *Notodiaptomus*, which underwent an attempt at unaccepted recombination with the subgenus *Notodiaptomus* (*Notodiaptomus*) (Dussart, 1985). Recently, the species was redescribed in a review of the *nordestinus* complex (Santos-Silva *et al*., 2015) and diaptomids from the Prata River Basin, Southern Neotropical (Perbiche-Neves *et al*., 2015).

The species is originally presented by Wright as very close to *N. iheringi.* However, Santos-Silva *et al*. (2015) during the review of *nordestinus* presented a thorough analysis of differential morphological characteristics for both species, corroborated by Perbiche-Neves *et al*. (2015) through new illustrations and additional identification key. Throughout our research, we were able to reinforce the differences presented in both proposals, note divergences and offer new elements for the morphological establishment of the species.

During our analyses it was not possible to reproduce the observation of Bowman (1973) for male A1R actual segment 8 with absence conical seta, and actual segment 16 absence spiniform process. This corroborates Santos-Silva *et al*. (2015) and the illustration in Dussart (1984). In the redescription of Perbiche-Neves *et al*. (2015) for individuals from the Prata River Basin these characters were not presented, and it is not possible to observe new evidence of these variations found by Bowman.

In the presentation made in Perbiche-Neves *et al*. (2015), we noticed a new inconsistency for the description of the female fifth swimming legs endopod 2-segmented. Through our examinations, we could not observe this condition, all the specimens evaluated have a unisegmented structure with internal discontinuity of the cuticle. However, for the same leg, the length of the Exp3 inner seta reaching the apex of the terminal claw can be corroborated as described in Perbiche-Neves *et al*. (2015) and differs from Santos-Silva *et al*. (2015).

Another relevant divergence is for the female epimeral plates, present asymmetrically and with the left side larger than the right. This condition differs from the description presented in Perbiche-Neves *et al*. (2015) and Santos-Silva *et al*. (2015). For the first, the symmetry between the structures is sustained, for the second, although the asymmetry is verified, the right side greater than the left is described. This condition cannot be corroborated in the present effort, where we observe the left epimeral plate with dorsal posterior lobe that makes it larger than the opposite side of the structure (right).

Additionally, the examined female presented fourth and fifth metasome segments fused dorsally, with presence of suture laterally. This verification corroborates Santos-Silva *et al*. (2015) and differs from the description in Perbiche-Neves *et al*. (2015), which supports complete fusion without suture present. In addition, all other characters described for the Wright *nordestinus* complex (Wright, 1935; 1936; 1937; Santos-Silva *et al*., 2015) and *Notodiaptomus* (Kiefer, 1936; 1956) were confirmed during our examinations.

#### Notodiaptomus conifer (Sars, 1901)

##### Synonymy

*Diaptomus conifer* Sars, 1901: 13, pl. 3, figs. 1–8; Daday, 1905: 147, 151, 152, pl. 9, fig. 10; Tollinger, 1911: 68, 270, 271, fig. D; Pearse, 1921: 459; Kiefer, 1926: 24; Kiefer, 1936b: 310; Pesta, 1927: 76, 80; Wright, 1927: 73, 75, 91, 100, 102, pl. 6, figs. 10–12; 1936: 79; 1937: 66, 73, 75, 76, pl. 3, figs. 1–4; 1938a: 302; 1938b: 562; Lowndes, 1934: 89, 91, 92, 93, 96, 98, 101; Brehm, 1935b: 308; 1956: 413; 1958a: 143, 167; 1965: 3, 5, 7, 8; Brandorff, 1972: 47; Bowman, 1973: 201; Infante *et al*., 1979: 225, 230. *Notodiaptomus conifer*; Kiefer, 1954: 173; 1956: 239, 242; Ringuelet, 1958a: 45, 46, 51; Paggi & José de Paggi, 1974: tab. 1; Brandorff, 1976: 615, 616, fig. 2; Gouvêa, 1980: 1047, 1048, 1050, 1051, 1058, 1059; Löffler, 1981: 15; Sendacz & Kubo, 1982: 54, 55, 66, 71, figs. 20–24, tab. 3; Dussart, 1983: 321; 1984b: 264, fig. 7B; Dussart & Defaye, 1983: 134; Arcifa, 1984: 143, tab. 7; Dussart & Frutos, 1985: 306, 307; 1986: 246; Sendacz *et al*., 1985: 190, 193, 196, 199, 203, 207, tabs. 4, 8, 12; Matsumura-Tundisi, 1986: 542, figs. 46–50, 100; Reid, 1987: 372; Defaye & Dussart, 1988: 123; Cicchino *et al*., 1989: 98; Paggi & José de Paggi, 1990: 690, tab. 2; Sendacz, 1993: 35; 1997: 624, 625, tab. 2; Battistoni, 1995: 958; Rocha *et al*., 1995: 155, 156; Santos-Silva, 1998: 207; Santos-Silva, 2008: 23; Santos-Silva *et al*., 2015: 25–29, figs. 12–14, identification keys to male and female; Perbiche-Neves *et al*., 2015: 44–48, figs. 35–38; Perbiche-Neves *et al*., 2020: 696-697, key to the Neotropical diaptomid, fig. 21.15 D. *Notodiaptomus (Notodiaptomus) conifer*; Dussart, 1985a: 208.

##### Type locality

Not clearly specified. Recently donated material was located and identified by the author as “*Diaptomus Conifer*”, Itatiba, São Paulo, Brazil, 1901”, which possibly indicates the type locality of the species.

##### Type material

Holotype not specified, and type material not originally defined for Sars (1901). There is material donated by SARS to the London Natural History Museum (NHM) containing 3 males and 4 females entire and labels: “*Diaptomus conifer*”, Itatiba, São Paulo, Brazil, 1901.12”. Probably it is a reference to the material used to describe the species.

##### Material examined

Topotype: 2 males, and 3 females, entire, preserved in alcohol 70%, stored in Plankton laboratory’s collection as “*Diaptomus*” *conifer*. Itatiba City, São Paulo State, Brazil, no date. Loan No. CR99/60, return by 16.VII.2000; 1 male (INPA-COP015, slides a-h) and 1 female (INPA-COP016, slides a-h) were selected to be dissection on eight slides each and deposited in the Zoological Collection of the INPA, Brazil. Additional material examined: 01 male, and 1 female, entire in formalin solution (MZUSP 32932), from the upper Tiete River, at the Barra Bonita Reservoir, Brazil, collected by G. Perbiche-Neves, and stored in MZUSP.

##### Diagnosis

**(1)** Cephalosome with sub-triangular anterior margin; **(2)** male epimeral plates asymmetrical; **(3)** male right antennule actual segment 15 with spinous surpassing to distal margin; **(4)** male right antennule actual segment 22 with five setae; **(5)** male right antennule actual segment 25 with five setae; **(6)** antenna coxa with seta reaching to the endopod 1; **(7)** male fifth left swimming leg reaching first right exopod segment distally; **(8)** male fifth left swimming leg basis with outgrowth on posterior surface longitudinally **(9)** male fifth left swimming leg exopod 1 with double lobe; **(10)** male fifth right swimming leg exopod 2 with posterior proximal swelling; **(11)** female genital double-somite with rounded swelling greater left than right; **(12)** female genital double-somite without right posterior rim expanded; **(13)** female genital double-somite with ventral vertical folds; **(14)** female caudal rami with patch of setules on outer surface; **(15)** female right antennule not extending beyond caudal rami; **(16)** female fifth swimming legs exopod 3 with terminal claw not 6x longer origin segment.

##### Redescription

###### MALE

Body 1618 micrometers excluding caudal setae. Male body smaller and slenderer than female. Nerve axons myelinated. Prosome 6-segmented; widest at first metasome segment; without one line of setules at posterior margin; without spinules at segments. Cephalosome anterior margin sub-triangular; with dorsal suture; incomplete; separate from first metasome segment. First metasome segment without sensilla. Second metasome segment without sensilla. Third metasome segment without sensillae; non-ornamented posterior margin. Fourth metasome segment without sensillae; separated from the fifth metasome. Limit between fourth and fifth metasome segments without ornamentation. Fifth metasome segment with sensilla; 2 laterally; Fifth metasome segment equal size; Fifth metasome segment without ornamentation; Fifth metasome segment without dorsal conical process; with epimeral plates. Epimeral plates asymmetrical. Right epimeral plates reduced, as rounded distal corner segment limit; with sensilla; at the apex of projection; without ornamentation. Left epimeral plate prominent, as projection; one projection; posterior-dorsally directed; reaching beyond half length of the genital segment; with sensillae; at the apex of projection; without ornamentation.

##### Urosome

5-segmented; Urosome 5 - free segments. Genital somite symmetrical in dorsal view; with single aperture; located on left side; ventrolaterally on posterior rim; with sensillae; on both sides; one; at left lateral; posteriorly; one; at right rim; posteriorly; of equal size between then. Third urosome segment without spinules; without external seta. Fourth urosome segment without spinules; without sub-conical blunt dorsal-lateral process. Anal segment presence of dorsal sensillae; one on each side; medially inserted; presence of operculum; convex; not covering the anal aperture fully. Caudal rami symmetrical; separated from anal segment; longer than wide; with setules; continuous on; inner side; each ramus bearing 6 caudal setae; 5 marginals; plumose; and 1 internal dorsally; straight; not reticulated main axis; outermost seta with outer spiniform process absent.

##### Appendices features

Rostrum symmetrical; separated from dorsal cephalic shield; by complete suture; sensillae present; one pair; anteriorly inserted on surface tegument; with rostral filament; double; paired; extended; into point; with basal process; in ventral view, rounded on left side; without a smaller basal expansion on the right side.

##### Antennules

Asymmetrical. **Right antennules**. Uniramous; right antennule surpassing to genital segment; right antennule not extending beyond caudal rami.

Right antennule ancestral segment I and II separated. Ancestral segment II and III fused. Ancestral segment III and IV fused. Ancestral segment IV and V separated. Ancestral segment V and VI separated. Ancestral segment VI and VII separated. Ancestral segment VII and VIII separated. Ancestral segment VIII and IX separated. Ancestral segment IX and X separated. Ancestral segment X and XI separated. Ancestral segment XI and XII separated. Ancestral segment XII and XIII separated. Ancestral segment XIII and XIV separated. Ancestral segment XIV and XV separated. Ancestral segment XV and XVI separated. Ancestral segment XVI and XVII separated. Ancestral segment XVII and XVIII separated. Ancestral segment XVIII and XIX separated. Ancestral segment XIX and XX separated. Ancestral segment XX and XXI separated. Ancestral segment XXI and XXII fused. Ancestral segment XXII and XXIII fused. Ancestral segment XXIII and XXIV separated. Ancestral segment XXIV and XXV fused. Ancestral segment XXV and XXVI separated. Ancestral segment XXVI and XXVII separated. Ancestral segment XXVII and XXVIII fused.

Right antennule actual 22-segmented; geniculated; between the segment 18 and segment 19; with swollen and modified region; formed by 5 segments; between 13 and 17 segments. Actual segment 1 with seta; one element; straight; none larger than segment; without spinules; without vestigial seta; without conical seta; without modified seta; without spinous process; with aesthetasc; one element. Actual segment 2 with seta; three elements; of unequal size; straight; none larger than segment; without spinules; with vestigial seta; one element; without conical seta; without modified seta; without spinous process; with aesthetasc; one element. Actual segment 3 with seta; one element; one larger than segment; surpassing to distal margin; beyond three sequential segments; straight; blunt apex; without spinules; with vestigial seta; one element; without conical seta; without modified seta; without spinous process; with aesthetasc. Actual segment 4 with seta; one element; one larger than segment; surpassing to distal margin; straight; not beyond three sequential segments; without spinules; without vestigial seta; without conical seta; without modified seta; without spinous process; without aesthetasc. Actual segment 5 with seta; one element; straight; one larger than segment; surpassing to distal margin; not beyond three sequential segments; without spinules; with vestigial seta; one element; without conical seta; without modified seta; without spinous process; with aesthetasc; one element. Actual segment 6 with seta; one element; none larger than segment; straight; without spinules; without vestigial seta; without conical seta; without modified seta; without spinous process; without aesthetasc. Actual segment 7 with seta; one element; straight; one larger than segment; surpassing to distal margin; beyond three sequential segments; blunt apex; without spinules; without vestigial seta; without conical seta; without modified seta; without spinous process; with aesthetasc; one element. Actual segment 8 with seta; one element; straight; none larger than segment; without spinules; without vestigial seta; with conical seta; one element; not reaching to middle-point of the sequent segment; without modified seta; without spinous process; without aesthetasc. Actual segment 9 with seta; two elements; of unequal size; straight; one larger than segment; surpassing to distal margin; beyond three sequential segments; blunt apex; without spinules; without vestigial seta; without conical seta; without modified seta; without spinous process; with aesthetasc; one element. Actual segment 10 with seta; one element; straight; none larger than segment; without spinules; without vestigial seta; without conical seta; with modified seta; presenting blunt apex; slender form; surpassing to distal margin; beyond of the sequential segment; parallel to antennule direction; without spinous process; without aesthetasc. Actual segment 11 with seta; one element; straight; one larger than segment; surpassing to distal margin; not beyond three sequential segments; without spinules; without vestigial seta; without conical seta; with modified seta; slender form; presenting blunt apex; surpassing to distal margin; beyond of the sequential segment; parallel to antennule direction; shorter length than homologous of actual segment 13; without spinous process; without aesthetasc. Actual segment 12 with seta; one element; straight; one larger than segment; surpassing to distal margin; not beyond three sequential segments; without spinules; without vestigial seta; with conical seta; one element; not smaller than to segment 8; without modified seta; without spinous process; with aesthetasc; one element; absent internal perpendicular fission. Actual segment 13 with seta; one element; straight; one larger than segment; surpassing to distal margin; not beyond three sequential segments; without spinules; without vestigial seta; without conical seta; with modified seta; stout form; surpassing to distal margin; to the middle-point of the sequence segment; perpendicular to antennule direction; presenting bifid apex; without spinous process; with aesthetasc; one element. Actual segment 14 with seta; two elements; of unequal size; straight; one larger than segment; surpassing to distal margin; beyond three sequential segments; blunt apex; without spinules; without vestigial seta; without conical seta; without modified seta; without spinous process; with aesthetasc; one element. Actual segment 15 with seta; two elements; of unequal size; straight; not bifidform; none larger than segment; without spinules; without vestigial seta; without conical seta; without modified seta; with spinous process; on outer margin; surpassing to distal margin; with aesthetasc; one element. Actual segment 16 with seta; two elements; of unequal size; plumose; one larger than segment; surpassing to distal margin; not beyond three sequential segments; not bifidform; without spinules; without vestigial seta; without conical seta; without modified seta; with spinous process; on outer margin; surpassing distal margin; unequal size to process on preceding segment; with aesthetasc; one element. Actual segment 17 with seta; two elements; of unequal size; straight; none larger than segment; bifidform; without spinules; without vestigial seta; without conical seta; with modified seta; one element; stout form; surpassing to distal margin; not beyond of the sequential segment; parallel to antennule direction; without spinous process; without aesthetasc. Actual segment 18 with seta; two elements; of equal size; straight; none larger than segment; without spinules; without vestigial seta; without conical seta; with modified seta; one element; stout form; surpassing distal margin; parallel to antennule direction; without spinous process; without aesthetasc. Actual segment 19 with seta; two elements; of unequal size; plumose; none larger than segment; without spinules; without vestigial seta; without conical seta; with modified seta; two elements; stout form; at least one bifid form; surpassing distal margin; parallel to antennule direction; without spinous process; with aesthetasc; one element. Actual segment 20 with seta; four elements; of unequal size; straight; one larger than segment; surpassing to distal margin; beyond three sequential segments; without spinules; without vestigial seta; without conical seta; without modified seta; without spinous process; without aesthetasc. Actual segment 21 with seta; two elements; of equal size; plumose; one larger than segment; surpassing to distal margin; greater 3x than original segment; without spinules; without vestigial seta; without conical seta; without modified seta; without spinous process; without aesthetasc. Actual segment 22 with seta; five elements; of equal size; one larger than segment; plumose; surpassing to distal margin; greater 3x than original segment; without spinules; without vestigial seta; without conical seta; without modified seta; without spinous process; with aesthetasc; one element.

##### Left antennules

Uniramous; Left antennule surpassing to prosome; Left antennule not extending beyond caudal rami. Ancestral segment I and II separated. Ancestral segment II and III fused. Ancestral segment III and IV fused. Ancestral segment IV and V separated. Ancestral segment V and VI separated. Ancestral segment VI and VII separated. Ancestral segment VII and VIII separated. Ancestral segment VIII and IX separated. Ancestral segment IX and X separated. Ancestral segment X and XI separated. Ancestral segment XI and XII separated. Ancestral segment XII and XIII separated. Ancestral segment XIII and XIV separated. Ancestral segment XIV and XV separated. Ancestral segment XV and XVI separated. Ancestral segment XVI and XVII separated. Ancestral segment XVII and XVIII separated. Ancestral segment XVIII and XIX separated. Ancestral segment XIX and XX separated. Ancestral segment XX and XXI separated. Ancestral segment XXI and XXII separated. Ancestral segment XXII and XXIII separated. Ancestral segment XXIII and XXIV separated. Ancestral segment XXIV and XXV separated. Ancestral segment XXV and XXVI separated. Ancestral segment XXVI and XXVII separated. Ancestral segment XXVII and XXVIII fused.

Left antennule actual 25-segmented; not-geniculated. Actual segment 1 with seta; one element; none larger than segment; straight; without spinules; without vestigial seta; without conical seta; without modified seta; without spinous process; with aesthetasc; one element. Actual segment 2 with seta; three elements; of equal size; none larger than segment; straight; without spinules; with vestigial seta; one element; without conical seta; without modified seta; without spinous process; with aesthetasc; one element. Actual segment 3 with seta; one element; one larger than segment; straight; surpassing to distal margin; beyond three sequential segments; without spinules; with vestigial seta; one element; without conical seta; without modified seta; without spinous process; with aesthetasc. Actual segment 4 with seta; one element; none larger than segment; straight; without spinules; without vestigial seta; without conical seta; without modified seta; without spinous process; without aesthetasc. Actual segment 5 with seta; one element; one larger than segment; straight; surpassing to distal margin; not beyond three sequential segments; without spinules; with vestigial seta; one element; without conical seta; without modified seta; without spinous process; with aesthetasc; one element. Actual segment 6 with seta; one element; none larger than segment; straight; without spinules; without vestigial seta; without conical seta; without modified seta; without spinous process; without aesthetasc. Actual segment 7 with seta; one element; one larger than segment; straight; surpassing to distal margin; beyond three sequential segments; without spinules; without vestigial seta; without conical seta; without modified seta; without spinous process; with aesthetasc; one element. Actual segment 8 with seta; one element; one larger than segment; straight; surpassing distal margin; without spinules; without vestigial seta; with conical seta; without modified seta; without spinous process; without aesthetasc. Actual segment 9 with seta; two elements; of unequal size; one larger than segment; straight; surpassing to distal margin; beyond three sequential segments; without spinules; without vestigial seta; without conical seta; without modified seta; without spinous process; with aesthetasc; one element. Actual segment 10 with seta; one element; none larger than segment; straight; without spinules; without vestigial seta; without conical seta; without modified seta; without spinous process; without aesthetasc. Actual segment 11 with seta; one element; one larger than segment; straight; surpassing to distal margin; beyond three sequential segments; without spinules; without vestigial seta; without conical seta; without modified seta; without spinous process; without aesthetasc. Actual segment 12 with seta; one element; one larger than segment; straight; surpassing distal margin; without spinules; without vestigial seta; with conical seta; without modified seta; without spinous process; with aesthetasc; one element. Actual segment 13 with seta; one element; none elongated; straight; surpassing distal margin; without spinules; without vestigial seta; without conical seta; without modified seta; without spinous process; without aesthetasc. Actual segment 14 with seta; one element; elongated; straight; surpassing to distal margin; beyond three sequential segments; without spinules; without vestigial seta; without conical seta; without modified seta; without spinous process; with aesthetasc; one element. Actual segment 15 with seta; one element; larger than segment; straight; surpassing to distal margin; not beyond three sequential segments; without spinules; without vestigial seta; without conical seta; without modified seta; without spinous process; without aesthetasc. Actual segment 16 with seta; one element; larger than segment; plumose; surpassing to distal margin; not beyond three sequential segments; without spinules; without vestigial seta; without conical seta; without modified seta; without spinous process; with aesthetasc; one element. Actual segment 17 with seta; one element; not larger than segment; straight; without spinules; without vestigial seta; without conical seta; without modified seta; without spinous process; without aesthetasc. Actual segment 18 with seta; one element; larger than segment; straight; surpassing to distal margin; beyond three sequential segments; without spinules; without vestigial seta; without conical seta; without modified seta; without spinous process; without aesthetasc. Actual segment 19 with seta; one element; not larger than segment; straight; surpassing distal margin; without spinules; without vestigial seta; without conical seta; without modified seta; without spinous process; with aesthetasc; one element. Actual segment 20 with seta; one element; not larger than segment; straight; surpassing distal margin; without spinules; without vestigial seta; without conical seta; without modified seta; without spinous process; without aesthetasc. Actual segment 21 with seta; one element; larger than segment; plumose; surpassing to distal margin; beyond three sequential segments; without spinules; without vestigial seta; without conical seta; without modified seta; without spinous process; without aesthetasc. Actual segment 22 with seta; two elements; of unequal size; one of them elongated; plumose; surpassing to distal margin; without spinules; without vestigial seta; without conical seta; without modified seta; without spinous process; without aesthetasc. Actual segment 23 with seta; two elements; of unequal size; one larger than segment; plumose; surpassing to distal margin; greater 3x than original segment; without spinules; without vestigial seta; without conical seta; without modified seta; without spinous process; without aesthetasc. Actual segment 24 with seta; two elements; of equal size; one larger than segment; plumose; surpassing to distal margin; greater 3x than original segment; without spinules; without vestigial seta; without conical seta; without modified seta; without spinous process; without aesthetasc. Actual segment 25 with seta; five elements; of equal size; elongated; plumose; surpassing to distal margin; 4 times larger than segment; without spinules; without vestigial seta; without conical seta; without modified seta; without spinous process; with aesthetasc; one element.

##### Antenna

Biramous. Antenna coxa separated from the basis; bearing seta; 1; on inner surface; at distal corner; reaching to the exopod 1. Antenna basis (fusion) separated from the endopodal segment; bearing seta; 2; on inner surface; at distal corner. Endopodal ancestral segment I and II separated. Ancestral segment II and III fused. Ancestral segment III and IV fused. Ancestral segment III and IV partially. Antenna endopod actual 2-segmented. Actual segment 1 not bilobate; with seta; two; on inner margin; with spinules; as a row; obliquely; on outer surface; with pore. Actual segment 2 bilobate; with discontinuity on outer cuticle; not developed as a suture; inner lobe bearing 8 setae; distally; outer lobe bearing 7 setae; distally; with spinules; as a patch; on outer surface. Antenna exopod ancestral segment I and II separated. Ancestral segment II and III fused. Ancestral segment III and IV fused. Ancestral segment IV and V separated. Ancestral segment V and VI separated. Ancestral segment VI and VII separated. Ancestral segment VII and VIII separated. Ancestral segment VIII and IX separated. Ancestral segment IX and X fused. Antenna exopod actual 7-segmented. Actual segment 1 single; elongated (width-length, equal or larger ratio 2:1); with seta; one; at inner surface. Actual segment 2 compound; elongated (larger width-length ratio 2:1); with seta; three; at inner surface. Actual segment 3 single; not elongated (lesser width-length ratio 2:1); with seta; one; at inner surface. Actual segment 4 single; not elongated (lesser width-length ratio 2:1); with seta; one; at inner surface. Actual segment 5 single; not elongated (lesser width-length ratio 2:1); with seta; one; at inner surface. Actual segment 6 single; not elongated (lesser width-length ratio 2:1); with seta; one; at inner surface. Actual segment 7 compound; elongated (larger or equal width-length ratio 2:1); with seta; one; at inner surface; and three; at distal surface.

##### Oral features. Mandible

Coxal gnathobase sclerotized; with lobe; prominent; on caudal margin; presence of cutting blade; with tooth-like prominence; two, distinctly; 1 acute; on caudal margin; and 1 triangular; on sub-caudal margin; without acute projection between the prominences; with additional spinules; as a row; on dorsal surface; with seta; 1; dorsally; on apical surface; with spinules; apicalmost. Mandible palps biramous; comprising the basis; with seta; four; differently inserted; first medially; reaching to beyond the endopod 1; second distally; third distally; fourth distally; on inner margin; none with setulose ornamentation. Mandible endopod 2-segmented. Mandible endopod 1 with lobe; bearing seta; four; distally inserted; without spinules. Mandible endopod 2 without lobe; bearing setae; nine elements; distally inserted; with spinules; as a row; double. Mandible exopod 4-segmented. Mandible exopod 1 with seta; one element; distally; on inner margin. Mandible exopod 2 with seta; one element; distally; on inner side. Mandible exopod 3 with seta; one element; distally; on inner side. Mandible exopod 4 with setae; three elements; on terminal region. **Maxillule**. Birramous. Maxillule 3-segmented. Maxillule praecoxa with praecoxal arthrite; bearing spines; fifteen elements; ten marginally; plus five sub-marginally; with spinules; as a patch; on sub-marginal surface. Maxillule coxa with coxal epipodite; with conspicuous outer lobe; bearing setae; nine elements; with coxal endite; elongated (larger or equal width-length ratio 2:1); bearing setae; four elements. Maxillule basis with basal endite; double; first proximal; elongated (larger width-length ratio 2:1; separated from basis; with setae; four elements; distally inserted; second distal; fused to basis; not elongated (lesser width-length ratio 2:1); with setae; four elements; distally inserted; with setules; as a row; on inner side; basal exite present; with setae; one element; on outer surface. Maxillule endopod 1-segmented. Endopod 1 bilobate; first proximal; with setae; three elements; second distal; with setae; five elements. Maxillule exopod 1-segmented. Exopod 1 with setae; six elements; with setules; as a row; on inner side; spinules absent. **Maxilla**. Uniramous. Maxilla 5-segmented. Maxilla praecoxa fused to coxa; incompletely; distinct externally; with praecoxal endite; double; first elongated endite (larger or equal width length ratio 2:1); proximally inserted; with seta; straight, or plumose; 1 straight; 4 plumose; with spine; single; without spinules; without setule; second elongated endite (larger or equal width length ratio 2:1); distally inserted; with seta; plumose; 3 plumose; without spine; with spinules; as a row; on distal margin; with setule; as a row; on distal margin; absence of outer seta. Maxilla coxa with coxal endite; double; first elongated endite (larger or equal width); proximally inserted; with seta; plumose; 3 plumose; without spine; without spinules; with setules; as a row; on proximal margin; second elongated endite (larger or equal width); distally inserted; with seta; plumose; 3 plumose; without spine; without spinules; with setules; as a row; on proximal margin; absence of outer seta. Maxilla basis with basal endite; single; elongated (larger or equal width-length ratio 2:1); with seta; plumose; 3 plumose; without spinules; absence of outer seta. Maxilla endopod 2-segmented. Endopod 1 with seta; 2 plumoses; without spine; without spinules; without setules. Maxilla endopod 2 with seta; 2 plumose; without spine; without spinules; without setules. **Maxilliped**. Uniramous; Maxilliped 8-segmented. Maxilliped praecoxa fused to coxa; incompletely; distinct internally; with praecoxal endite; not elongated (lesser width-length ratio 2:1); distally inserted; with seta; 1 straight; with spinules; as a row; single; on basal surface; without setules. Maxilliped coxa with coxal endite; three coxal endite; first elongated (larger or equal width); proximally inserted; with seta; 2 plumose; with spinules; as a patch; single; on apical surface; without setules; second not elongated (lesser width-length ratio 2:1); medially inserted; with seta; 3 plumose; with spinules; as a row; single; on medial surface; without setules; third elongated (larger or equal width length ratio 2:1); distally inserted; with seta; 3 plumose; none reaching to beyond of the basis; with spinules; as a row; single; on basal surface; without setules; with lobe; prominence; at inner distal angle; ornamented; with spinules; continuously on margin. Maxilliped basis without basal endite; with seta; 3 plumose; with spinules; as a row; single; on medial surface; with setules; as a row; single; on inner margin. Maxilliped endopod segment 6-segmented. Endopod 1 with seta; 2 plumose; on inner surface. Endopod 2 with seta; 3 plumose; on inner surface. Endopod 3 with seta; 2 plumose; on inner surface. Endopod 4 with seta; 2 plumose; on inner surface. Endopod 5 with seta; 2 plumose; on inner surface, or on outer surface; outer seta absent. Endopod 6 with seta; 4 plumose; on inner surface, or on outer surface.

##### Swimming legs features

**First swimming legs.** Symmetrical; biramous. First swimming legs intercoxal plate without seta. First swimming legs praecoxa absent. First swimming legs coxa with seta; one; straight; distally inserted; on inner surface; surpassing to first endopodal segment; with setules; two group; as a patch; on inner margin; and as a row; double; on anterior surface; outerly; without spinules; without spine. First swimming legs basis without seta; with setules; as a patch; single; on outer surface; without spinules; without spine. First swimming legs endopod 2-segmented. Endopod 1 with seta; straight; restricted; to inner surface; one element; without spine; with setules; as a row; single; continuously; on outer surface; without spinules; absence of Schmeil’s organ. Endopod 2 with seta; unrestricted; three on inner surface; one on outer surface; two on distal surface; straight; without spine; with setules; as a row; single; continuously; on outer surface; without spinules; absence of Schmeil’s organ. Endopod 3 absence. First swimming legs exopod 1 with seta; restricted; 1 on inner surface; with spine; 1; stout; smaller than original segment; serrated; on inner side; continuously; with setules; as a row; single; as a row; innerly. First swimming legs exopod 2 with seta; restricted; 1 on inner surface; straight; without spine; with setules; as a row; single; continuously; on inner margin, or on outer margin; without spinules. First swimming legs exopod 3 with setule; as a row; single; continuously; on outer surface; without spinules; with seta; unrestricted; 2 on inner surface; 2 on terminal surface; with spine; 2; unequal size; first no longer 2x than origin segment; stout; serrated; on inner side, or on outer side; equally; second longer 3x than origin segment; slender; serrated; on outer side; with ornamentation on non-serrated side; by setules. **Second swimming legs**. Symmetrical; Second swimming legs biramous. Second swimming legs intercoxal plate without seta. Second swimming legs praecoxa present; located laterally. Second swimming legs coxa with seta; straight; distally inserted; on inner surface; surpassing to basal segment; without setules; without spinules; without spine. Second swimming legs basis without seta; without setules; without spinules; without spine. Second swimming legs endopod 3-segmented. Endopod 1 with seta; straight; restricted; one on inner surface; without spine; with setules; as a row; single; continuously; on outer surface; without spinules; absence of Schmeil’s organ. Endopod 2 with seta; straight; unrestricted; two on inner surface; without spine; with setules; as a row; single; continuously; on outer side; without spinules; presence of Schmeil’s organ; on posterior surface. Endopod 3 with seta; straight; unrestricted; three on inner surface; two on outer surface; two on distal surface; without spine; without setules; with spinules; as a row; double; distally inserted; at anterior surface; absence of Schmeil’s organ. Second swimming legs exopod 1 with seta; restricted; one on inner surface; with spine; 1; stout; not reaching to distal-third of the exopod 2; serrated; on inner side, or on outer side; with setules; as a row; single; continuously; on inner side; without spinules; absence of Schmeil’s organ. Exopod 2 with seta; unrestricted; one on inner surface; with spine; 1; stout; not surpassing the exopod 3; serrated; on inner side, or on outer side; with setules; as a row; single; continuously; on inner surface; without spinules; absence of Schmeil’s organ. Exopod 3 with seta; plurimarginal; three on inner surface; two on terminal surface; with spine; 2; unequal size; first no longer 2x than origin segment; stout; serrated; on inner side, or on outer side; equally; second longer 2x than origin segment; slender; serrated; on outer side; with ornamentation on non-serrated side; of setules; setules on outer surface; as a row; single; continuously; on inner surface; with spinules; as a row; single; distally inserted; at anterior surface; absence of Schmeil’s organ. **Third swimming legs**. Symmetrical; Third swimming legs biramous. Third swimming legs intercoxal plate without seta. Third swimming legs praecoxa present; not laterally located. Third swimming legs coxa with seta; straight; distally inserted; on inner surface; surpassing to first endopodal segment; without setules; without spinules; without spine. Third swimming legs basis without seta; without setules; without spinules; without spine. Third swimming legs endopod 3-segmented. Endopod 1 with seta; restricted; one on inner surface; without spine; without setules; without spinules; absence of Schmeil’s organ. Endopod 2 with seta; restricted; two on inner surface; straight; without spine; without setules; without spinules; absence of Schmeil’s organ. Endopod 3 with seta; straight; plurimarginal; two on inner surface; two on outer surface; three on terminal surface; without spine; without setules; with spinules; as a row; distally inserted; double; at anterior surface; absence of Schmeil’s organ. Third swimming legs exopod 1 with seta; restricted; straight; one on inner surface; with spine; 1; stout; not reaching to the distal-third of the exopod 2; serrated; equally; on inner surface, or on outer surface; with setules; as a row; single; continuously; on inner surface; without spinules; absence of Schmeil’s organ. Exopod 2 with seta; straight; restricted; one on inner surface; with spine; 1; stout; not reaching out to exopod 3; serrated; on inner side, or on outer side; equally; with setules; as a row; single; continuously; on inner side; without spinules; absence of Schmeil’s organ. Exopod 3 without setules; with spinules; as a row; single; distally inserted; at anterior surface; with seta; straight; unrestricted; three on inner surface; two on terminal surface; with spine; 2; unequal size; first no longer 2x than origin segment; stout; serrated; on inner side, or on outer side; equally; second longer 2x than origin segment; slender; serrated; on outer side; with ornamentation on non-serrated side; of setules; absence of Schmeil’s organ. **Fourth swimming legs**. Symmetrical; biramous. Intercoxal plate without sensilla. Praecoxa present. Coxa with seta; distally inserted; on inner margin; reaching out to endopod 1; without spinules; setules absent. Basis with seta; one; medially inserted; on posterior surface; smaller than the original segment; without setules; without spinules; without spine. Fourth swimming legs endopod 3-segmented. Endopod 1 with seta; one; restricted; on inner surface; without spine; without setules; without spinules; absence of Schmeil’s organ. Endopod 2 with seta; restricted; two on inner side; without spine; with setules; as a row; single; continuously; on outer surface; without spinules; absence of Schmeil’s organ. Endopod 3 with seta; unrestricted; two on inner surface; two on outer surface; three on distal surface; without spine; without setules; with spinules; as a row; double; distally inserted; at anterior surface; absence of Schmeil’s organ. Fourth swimming legs exopod 1 with seta; restricted; one on inner surface; with spine; 1; stout; not reaching out to distal-third of the exopod 2; serrated; on inner side, or on outer side; equally; with setules; as a row; single; continuously; on inner surface; without spinules; absence of Schmeil’s organ. Exopod 2 with seta; restricted; one on inner surface; with spine; 1; stout; not reaching the end of exopod 3; serrated; on inner side, or on outer side; equally; with setules; as a row; single; continuously; on inner surface; without spinules; absence of Schmeil’s organ. Exopod 3 without setules; with spinules; as a row; single; distally inserted; at anterior surface; with seta; unrestricted; three on inner surface; two on distal surface; with spine; 2; unequal size; first no longer 2x than origin segment; stout; serrated; on inner side, or on outer side; equally; second longer 2x than origin segment; slender; serrated; on outer side; without ornamentation on non-serrated side; absence of Schmeil’s organ.

##### Fifth swimming legs features

Asymmetrical. Fifth swimming leg intercoxal plate with length not equal or greater than width on 1.5x; with irregular proximal margin; discontinuous to; the anterior margin of the left coxa, or the anterior margin of the right coxa; posterior sensilla on the right lateral absent. **Fifth left swimming leg**. Fifth left swimming leg biramous; leg reaching first right exopod segment; distally. Fifth left swimming leg praecoxa present; rudimentary; separated from the coxae; without ornamentation. Fifth left swimming leg coxa concave inner side; without teeth-like structures; with process; conical; on posterior surface; outer side; distally inserted; not projecting over basis; with sensilla; stout; triangular; at apex; longer 2x than insertion basis; without swelling; without seta; without spinules. Fifth left swimming leg basis sub-cylindrical; unequal size between inner and outer side; shorter outer than inner side; with concave inner side; rounded internal proximal expansion absent; with outgrowth; on posterior surface; with groove; deep; longitudinally; on posterior surface; not reaching the endopodal lobe; not ornamented; absence of protuberance; with seta; outerly inserted; no longer 2x than origin segment; absence of minutely granular. Fifth left swimming leg endopod segments 1 and 2 fused; segments 2 and 3 fused; 1-segmented; stout; separated from the basis; ornamented; on inner side; with spinules; more than four elements; as a row; terminally; row of setules absent; without seta. Fifth left swimming leg exopod segments 1 and 2 separated; segments 2 and 3 fused; 2-segmented; stout; separated from the basis. Fifth left swimming leg exopod 1 sub-cylindrical; longer than broad; unequal size between inner and outer side; shorter inner than outer side; rectilinear inner side; concave outer side; without swelling; without marginal extension; without process; with lobe; double; semicircular; medially inserted; on inner side; covered; by setules; without outer spine; absence seta. Fifth left swimming leg exopod 2 digitiform; longer than broad; equal size between inner and outer side; disform inner side; with convex outer side; setulose pad present; prominently rounded; medially; on inner side; inflated medial region absent; distal process present; digitiform; non denticulate; without transverse row of denticles; none oblique row of 5 denticles; innerly directed; with seta; spiniform; not ornamented by spinules; not surpassing the distal-point of the segment; without outer spine; terminal claw absent.

##### Fifth right swimming leg

Biramous. Fifth right swimming leg praecoxa present; separated from the coxae; without ornamentation. Fifth right swimming leg coxa convex inner side; without teeth-like structures; with process; rounded; distally inserted; on posterior surface; closest to the outer rim; projecting over basis; beyond the first third; until the medial surface; without triangular protuberance innerly; with sensilla; slender; at apex; no longer 2x than basal insertion; without marginal extension; without seta; without spinules. Fifth right swimming leg basis cylindrical; unequal size between inner and outer side; shorter outer than inner side; rectilinear inner side; tumescence present; not inflated; restricted on inner surface; proximally; without protuberance; absence of distinct minutely granular; additional inner process absent; with posterior groove; deep; obliquely; not reaching the endopodal lobe; ornamented; with tubercles; throughout of the outer border; with seta; outerly inserted; on anterior surface; no longer 2x than origin segment; posterior protrusion present; distal process absent. Fifth right swimming leg with endopodite present; fused to basis; on anterior surface; ancestral segments 1 and 2 fused; ancestral segments 2 and 3 fused; stout; ornamented; with setules; as a row; on inner side; terminally; without seta. Fifth right swimming leg exopod segments 1 and 2 separated; segments 2 and 3 fused; 2-segmented; stout; separated from the basis. Fifth right swimming leg exopod 1 sub-cylindrical; longer than broad; nearly 1.25 times; unequal size between both sides; shorter inner than outer side; convex inner side; rectilinear outer side; with marginal extension; sub-triangular; distally inserted; at outer rim; spinules absent; with process; rounded; sclerotized; without ornamentation; distally inserted; at posterior surface; not projecting over next segment; without outer spine; without seta; internal prominence absent; lamella on posterior surface absent. Fifth right swimming leg exopod 2 cylindrical; longer than broad; nearly 2.5 times; equal size between both sides; disform inner side; convex outer side; with posterior proximal swelling; inner-posterior process absent; without marginal expansion; curved ridge on distal posterior surface present; chitinous knobs absent; with outer spine; inserted sub-distally; rectilinear; not ornamented innerly; not ornamented outerly; sharp tip; without apparent curve; lesser than the length of the exopod 2; beyond to 2 times its size; 3x; sensilla absent; terminal claw present; equal or longer 1.5 times than insertion segment; sclerotized; arched; inward; without conspicuous curve; ornamented innerly; by spinules; as a row; partially on extension; medially, or distally; not ornamented outerly; sharp tip; not curved tip; without medial constriction; hyaline process absent.

##### FEMALE

Body longer and wider than male; Female body 1628 micrometers excluding caudal setae. Widest at first metasome segment. Distal margin of the prosomal segments without one line of setules at posterior margin. Prosome segments without spinules at prosomal segments. Fourth metasome segment presence of dorsal protuberance; rounded; inserted distally; without posterior process; without anterior process; fourth metasome segment without proximal sensillae present. Fourth and fifth metasome segments fused; partially; on dorsal surface. Limit between fourth and fifth metasome segments without ornamentation. **Fifth metasome segment**. Fifth metasome segment without sensilla; with epimeral plates. Epimeral plates asymmetrical. Right epimeral plates prominent, as projections; thinner than the left; one posterior-laterally directed; not reaching half length of the genital segment; with sensilla at the apex; dorsal-posterior sensilla present; slender; without ornamentation. Left epimeral plate without expansion.

##### Urosome

3-segmented. **Genital double-somite**. Asymmetrical in dorsal view; longer than broad; longer than other urosomites combined; dorsal suture at mid-length absent; not covered by spinules; with swelling; rounded; unequal size; greater right than left; anteriorly; with sensillae; on both sides; one; stout; with robust apex; at left lateral; not on lobular base; anteriorly; one; stout; at right lateral; not on lobular base; anteriorly; with robust apex; of equal size between then; lateral protuberance absent; without right posterior rim expanded; without slender sensilla on each posterior rim; without posterior-dorsal process. Genital double-somite opercular pad present; broader than longer; symmetrical; development laterally; expanded posteriorly; covering partially; double gonoporal slit; located ventrally; with arthrodial membrane; inserted anteriorly; post-genital process absent; disto-ventral tumescence absent; ventral vertical folds present; dorsal sensilla absent. Second urosome segment without ventral fusion to anal segment; right distal process absent. Caudal rami patch of setules on outer surface present; patch of spinules on outer surface absent.

##### Appendices features

Rostrum basal process absent. **Antennules**. Symmetrical. Right antennule surpassing to genital double-segment; not extending beyond caudal rami; ornamentation pattern equals to male left antennule; fully.

##### Fifth swimming legs

Symmetrical; Fifth swimming legs biramous. Fifth swimming legs intercoxal plate longer than wide; separated from the legs. Fifth swimming legs praecoxa with sclerite praecoxal; separated from the coxae; without ornamentation. Fifth swimming legs coxa with process; conical; at the outer rim; distally; sensilla present; stout; at apex; projecting over basal segment; no longer 2x than basal insertion; marginal extension absent; without swelling; without seta; without spinules. Fifth swimming legs basis sub-triangular; unequal size between inner and outer sides; shorter outer than inner side; with convex inner side; without proximal inner outgrowth; without groove; with distal extension; on posterior surface; with seta; outerly inserted; on anterior surface; longer 2x than origin segment; not reaching to exopod 1 distally. Fifth swimming legs endopod segments 1 and 2 fused; segments 2 and 3 fused; 1-segmented; stout; separated from the basis; present discontinuity cuticle; on inner side; with spinules; as a row; single; non-oblique; sub-terminally; at anterior surface; with seta; double; one medially; on posterior surface; rectilinear; one distally; on posterior surface; arched; of unequal size; distal seta longer than medial seta. Fifth swimming legs exopod segments 1 and 2 separated; segments 2 and 3 separated; 3-segmented; separated from the basis. Fifth swimming legs exopod 1 sub-cylindrical; longer than wide; longer or equal than 2 times; with unequal size between inner and outer side; shorter inner than outer side; with convex inner side; with rectilinear outer side; without swelling; without marginal extension; without posterior process; without spine; without seta. Fifth swimming legs exopod 2 sub-cylindrical; longer than broad; longer or equal than 2 times; without swelling; without marginal extension; without process; without lobe; with spine; inserted laterally; rectilinear; without ornamentation; sharp tip; equal size or larger than next segment; without seta. Fifth swimming legs exopod 3 cylindrical; longer than wide; without swelling; without process; without lobe; without spine; with seta; double; inserted terminally; unequal size between them; outer seta smaller than inner; nearly 3 times; outer seta not ornamented by setules; without ornamentation; presence of terminal claw; sclerotized; arched; externally directed; convex inner side; with ornamentation; of denticles; as a row; on surface partially; at medial region; concave outer side; with ornamentation; of denticles; as a row; on surface partially; at medial region; blunt tip; not 6 times longer origin segment.

##### Distribution records

###### BRAZIL

**Mato Grosso**: Corumbá (Daday, 1905). **Bahia**: lake of Abaeté, 12°55’S, 38°22’W (Gouvêa, 1980). **São Paulo**: dry sediment from Itatiba (Sars, 1901; this study); reservoir in Sorocaba, and brick factory’s pond, near to Amparo (Wright, 1937); Batista reservoirs, Paranapanema, and Itupararanga Rivers, Tietê River (Sendacz & Kubo, 1982; Sendacz *et al*., 1985); Xavantes Reservoir, Paranapanema River (Matsumura-Tundisi, 1986); upper Paraná River (Sendacz, 1998); upper Tietê River at the Barra Bonita Reservoir (Perbiche-Neves *et al*., 2015). PARAGUAI. Aregua, floodplain of a stream that crosses the road to the lagoon Ipacaraí; pond beside the railroad; floodplain between Aregua, and Yuguari; ponds in Assunção; Campo Grande; Calle de la Cañada; ponds on an island in the Paraguay River; Gran Cahaco, Paraguay River; Laguna (Pasito); Cerro León, Bañado; Curuzu-ñu, small lake near to Marcos Romeros’ home; lake Estia Postillon; Courallhes, permanent pond; lake Ipacaraí, water blade; Lugua, pond near the train station; Pirayu, pond in street and near to brick factory; Sapucay, pond from pluviometric origin; Tebicuay, permanent swamp; floodplain, Yuguari River (Daday, 1905); Makthlawaiya, a pond from the rain in capinzal (23°25’S, 58°19W); pond in a forest em uma floresta, 5 miles northeast of Nanahua (Lowndes, 1934). ARGENTINA. Middle Paraná River, between the Santa Fé and Paraná Citys (Paggi & José de Paggi, 1974); middle Paraná River (Paggi & José de Paggi, 1990). **Buenos Aires**: laguna Totora, Laprida; laguna Videl, Chascomus and Tapalque (Brehm, 1965); La Plata (Brehm, 1965). **Cordoba**: Unguillo (Brehm, 1965). **Chaco**: Resistencia; pond in Makallé (Ringuelet, 1958); Corzuela (Brehm, 1965). **Mendoza**: La Dormids (Brehm, 1965).

##### Habitat

Habitat in freshwaters: reservoirs, shallow ponds, rivers, lotic zone of streams, and dry sediment (resting eggs).

##### Remarks

The species was presented to science from organisms hatched from dry sediment samples from South America. Santos-Silva *et al*. (2015), in the review of the *nordestinus* complex, mentioned the existence of material identified by Sars in the London Natural History Museum (NHM), identified as “*Diaptomus conifer*, Itatiba, São Paulo, Brazil, 1901” and, probably, referring to the type locality of the resting eggs received. The taxon was considered formally part of *Notodiaptomus* only in the amplification of the genus (Kiefer, 1956).

Over time, morphological variations for the species have been reported in the literature (Daday, 1905; Pearse, 1921; Wright, 1937; Brehm, 1958). Among the significant ones, Daday (1905) found important variations for the female of the species originally described: (1) antennule length reaching to caudal rami distally; (2) fifth swimming legs endopod 2-segmented; (3) fifth swimming legs endopod with spinules row oblique, plus seta distally. Of these characteristics, only the first was confirmed in Santos-Silva *et al*. (2015), and none of them mentioned in Perbiche-Neves *et al*. (2015).

In our examinations, the attributes described by Sars (1901) were corroborated, except that we identified the female antennule length reaching to caudal rami distally, and male fifth swimming legs endopod development relatively, the left reaching to exopod 2 medially, and the right reaching to exopod 1 medially. From the diagnostic attributes originally highlighted, we present the variation in the dorsal protuberance form, being rounded instead of “pyramidal”. Other defining attributes were indicated in Santos-Silva *et al*. (2015) and Perbiche-Neves *et al*. (2015), corroborated in this research and presented in the diagnosis section above.

Among the characters assigned as convergent for the species of *Notodiaptomus* originally and in the amplification, all could be present for the species, except the male A1R 20 with spinous process. Of the converging attributes noted for the type-species of the genus, those absent in *N. conifer* were mainly: male fifth left swimming leg exopod 1 with double lobe; female genital double-somite with ventral vertical folds; female caudal rami with patch of setules on outer surface; and male fifth left swimming leg basis with outgrowth on posterior surface longitudinally. There were also present differentiation for the pattern found in the male A1R of *N. deitersi*, such as: actual segment 15 with spinous surpassing to distal margin (similar to *Argyrodiaptomus furcatus* (Sars, 1901) illustrated in Dussart, 1985, p. 203: fig 1a); actual segment 22 with five setae; and actual segment 25 with five setae.

#### Notodiaptomus coniferoides (Wright, 1927)

##### Synonymy

*Diaptomus coniferoides* Wright, 1927: 75, 92, 100, 102, pl. 7, figs. 1–4; 1937a: 77; 1938b: 562; 1939: 647; Lowndes, 1934; 89, 90, 91, 92, 93, 94–96, pl. 1, figs. 1a-d; Brehm, 1938: 29; 1957: 60, figs. 72–76; 1958a: 140, 141, 142, 143, 147, 167, pl. 2, figs. 1–4; 1960: 49; 1965: 3, 7, 8; Brandorff, 1972: 4, 5, 7, 8, 9, 24, 25, 48, figs. 27–28; Reid, 1991: 737, 738. *“Diaptomus” coniferoides*; Brehm, 1958a: 147. *Notodiaptomus coniferoides* n. comb., Ringuelet, 1958a: 45, 46, 52; 1962: 87; Herbst, 1967: 96; Cicchino, 1972: 585–596; Brandorff, 1973b: 206; 1976: 616, 622, fig. 2; 1978a: 298; Paggi & José de Paggi, 1974: 109, tab. 1; 1990: 685, 686, 690, 692, tab. 2; Andrade & Brandorff, 1975: 97; José de Paggi, 1978: 150, tab. 1; 1981: 189, 199; Gouvêa, 1980: 1047, 1050, 1051, 1058; Hardy, 1980: 594, 596, 604; Löffler, 1981: 15; Carvalho, 1983: 717; Dussart & Defaye, 1983: 135; Dussart, 1984a: 34, 35, 38, 39, 54, fig. 9; 1984b: 264, fig. 7A; Robertson & Hardy, 1984: 347, tab. 3; Arcifa, 1984: 143, tab. 7; Dussart & Frutos, 1986: 306, 307, figs. 14–18; 1987: 243, 244, 245, 246, pl. 2, figs. 10–12; Reid & Moreno, 1990: 726, 729, 730–733, tabs. 2, 3; Reid, 1991: 737, 738; Santos-Silva, 1991: 33, 34, fig. 11; 1998: 207; Sendacz & Melo Costa, 1991: 466, 468, 469; Frutos, 1993: 91, 112, tab. 3; Santos-Silva & Robertson, 1993: 101; Sendacz, 1993: 35; Battistoni, 1995: 958; Rocha *et al*., 1995: 155, 156; Jersabek *et al*., 1996: 2028, 2030. *Notodiaptomus coniferoides*; Matsumura-Tundisi, 1986: 542, 547, figs. 51–54, 100; Santos-Silva, 2008: 23; Perbiche-Neves *et al*., 2015: 49–53, figs. 39–43, identification keys to male and female; Perbiche-Neves *et al*., 2020: 683-684, key to the Neotropical diaptomid. [*error*] *Notodiaptomus (Caleodiaptomus) coniferoides*; Dussart, 1985a: 201, 214.

##### Type locality

Calama, Rondônia, Brazil.

##### Type material

Holotype: male (USNM 59827). Paratype: female (USNM 59828).

##### Material examined

Topotype: 2 males, 1 female from the lake Janauarizinho, Calama. 16.xii.1993; 1 male, and 1 female from the Castanho Lake, RO, Machado River, Calama. 19.XI.1993. All specimens entire and alcohol, stored in collection of Lab Plankton, INPA; 1 male (INPA-COP017, slides a-h) and 1 female (INPA-COP018, slides a-h) were selected to be dissection on eight slides each and deposited in the Zoological Collection of the INPA, Brazil. Additional material examined: 01 male, and 1 female, entire in formalin solution (MZUSP 28389), G. Perbiche-Neves coll., stored in MZUSP.

##### Diagnosis

**(1)** male epimeral plates with sensilla at the apex of projection and other medially; **(2)** male fifth left swimming leg exopod 2 digitiform process with bicuspidate denticles; **(3)** male fifth right swimming leg coxa with distal rounded process on posterior surface; **(4)** male fifth right swimming leg exopod 1 broader than long; **(5)** female fourth metasome segment with dorsal digitiform protuberance 2x wider than long medially; **(6)** female right epimeral plate directed posterior-dorsally; **(7)** female genital double-somite without lateral protuberance on left side, and suture at mid-length dorsally; **(8)** female right antennule not extending beyond caudal rami.

##### Redescription

###### MALE

Body 1053 micrometers excluding caudal setae. Male body smaller and slenderer than female. Nerve axons myelinated. Prosome 6-segmented; widest at first metasome segment; without one line of setules at posterior margin; with spinules at least at one segment. Cephalosome anterior margin sub-triangular; with dorsal suture; complete; separate from first metasome segment. First metasome segment without sensilla. Second metasome segment without sensilla. Third metasome segment without sensillae; ornamented posterior margin; with setules; as a patch; dorsally. Fourth metasome segment with sensillae; 4 dorsally; 2 laterally; of equal size; separated from the fifth metasome. Limit between fourth and fifth metasome segments without ornamentation. Fifth metasome segment with sensilla; 2 dorsally; Fifth metasome segment equal size; Fifth metasome segment without ornamentation; Fifth metasome segment without dorsal conical process; with epimeral plates. Epimeral plates asymmetrical. Right epimeral plates prominent, as projections; one projection; posterior-dorsally directed; not reaching half length of the genital segment; with sensilla; one at the apex of projection and other medially; ornamented; with spinules; as a patch. Left epimeral plate prominent, as projection; one projection; posterior-dorsally directed; reaching half length of the genital segment; with sensillae; one at the apex of projection and other medially; without ornamentation.

##### Urosome

5-segmented; Urosome 5-free segments. Genital somite asymmetrical in dorsal view; with single aperture; located on left side; ventrolaterally on posterior rim; with sensillae; on single side; one; at right rim; posteriorly. Third urosome segment without spinules; without external seta. Fourth urosome segment without spinules; without sub-conical blunt dorsal-lateral process. Anal segment presence of dorsal sensillae; one on each side; medially inserted; presence of operculum; convex; not covering the anal aperture fully. Caudal rami symmetrical; separated from anal segment; longer than wide; with setules; continuous on; inner side; each ramus bearing 6 caudal setae; 5 marginals; plumose; and 1 internal dorsally; straight; not reticulated main axis; outermost seta with outer spiniform process absent.

##### Appendices features

Rostrum asymmetrical; separated from dorsal cephalic shield; by complete suture; sensillae present; one pair; anteriorly inserted on surface tegument; with rostral filament; double; paired; extended; into point; with basal process; in ventral view, rounded on left side; without a smaller basal expansion on the right side.

##### Antennules

Asymmetrical. **Right antennules**. Uniramous; right antennule surpassing to genital segment; right antennule not extending beyond caudal rami.

Right antennule ancestral segment I and II separated. Ancestral segment II and III fused. Ancestral segment III and IV fused. Ancestral segment IV and V separated. Ancestral segment V and VI separated. Ancestral segment VI and VII separated. Ancestral segment VII and VIII separated. Ancestral segment VIII and IX separated. Ancestral segment IX and X separated. Ancestral segment X and XI separated. Ancestral segment XI and XII separated. Ancestral segment XII and XIII separated. Ancestral segment XIII and XIV separated. Ancestral segment XIV and XV separated. Ancestral segment XV and XVI separated. Ancestral segment XVI and XVII separated. Ancestral segment XVII and XVIII separated. Ancestral segment XVIII and XIX separated. Ancestral segment XIX and XX separated. Ancestral segment XX and XXI separated. Ancestral segment XXI and XXII fused. Ancestral segment XXII and XXIII fused. Ancestral segment XXIII and XXIV separated. Ancestral segment XXIV and XXV fused. Ancestral segment XXV and XXVI separated. Ancestral segment XXVI and XXVII separated. Ancestral segment XXVII and XXVIII fused.

Right antennule actual 22-segmented; geniculated; between the segment 18 and segment 19; with swollen and modified region; formed by 5 segments; between 13 and 17 segments. Actual segment 1 with seta; one element; straight; none larger than segment; without spinules; without vestigial seta; without conical seta; without modified seta; without spinous process; with aesthetasc; one element. Actual segment 2 with seta; three elements; of equal size; straight; none larger than segment; without spinules; with vestigial seta; one element; without conical seta; without modified seta; without spinous process; with aesthetasc; one element. Actual segment 3 with seta; one element; one larger than segment; surpassing to distal margin; beyond three sequential segments; straight; blunt apex; without spinules; with vestigial seta; one element; without conical seta; without modified seta; without spinous process; with aesthetasc; one element. Actual segment 4 with seta; one element; none larger than segment; straight; without spinules; without vestigial seta; without conical seta; without modified seta; without spinous process; without aesthetasc. Actual segment 5 with seta; one element; straight; one larger than segment; surpassing to distal margin; not beyond three sequential segments; without spinules; without vestigial seta; without conical seta; without modified seta; without spinous process; with aesthetasc; one element. Actual segment 6 with seta; one element; none larger than segment; straight; without spinules; without vestigial seta; without conical seta; without modified seta; without spinous process; without aesthetasc. Actual segment 7 with seta; one element; straight; one larger than segment; surpassing to distal margin; beyond three sequential segments; blunt apex; without spinules; without vestigial seta; without conical seta; without modified seta; without spinous process; with aesthetasc; one element. Actual segment 8 with seta; one element; straight; none larger than segment; without spinules; without vestigial seta; with conical seta; one element; not reaching to middle-point of the sequent segment; without modified seta; without spinous process; without aesthetasc. Actual segment 9 with seta; two elements; of unequal size; straight; one larger than segment; surpassing to distal margin; beyond three sequential segments; blunt apex; without spinules; without vestigial seta; without conical seta; without modified seta; without spinous process; with aesthetasc; one element. Actual segment 10 with seta; one element; straight; one larger than segment; surpassing distal margin; not beyond three sequential segments; without spinules; without vestigial seta; without conical seta; with modified seta; presenting blunt apex; slender form; surpassing to distal margin; beyond of the sequential segment; parallel to antennule direction; without spinous process; without aesthetasc. Actual segment 11 with seta; one element; straight; one larger than segment; surpassing to distal margin; not beyond three sequential segments; without spinules; without vestigial seta; without conical seta; with modified seta; slender form; presenting blunt apex; surpassing to distal margin; beyond of the sequential segment; parallel to antennule direction; shorter length than homologous of actual segment 13; without spinous process; without aesthetasc. Actual segment 12 with seta; one element; straight; one larger than segment; surpassing to distal margin; not beyond three sequential segments; without spinules; without vestigial seta; with conical seta; one element; not smaller than to segment 8; without modified seta; without spinous process; with aesthetasc; one element; absent internal perpendicular fission. Actual segment 13 with seta; one element; straight; none larger than segment; without spinules; without vestigial seta; without conical seta; with modified seta; stout form; surpassing to distal margin; to the distal-point of the sequence segment; perpendicular to antennule direction; presenting bifid apex; without spinous process; with aesthetasc; one element. Actual segment 14 with seta; two elements; of unequal size; straight; one larger than segment; surpassing to distal margin; beyond three sequential segments; blunt apex; without spinules; without vestigial seta; without conical seta; without modified seta; without spinous process; with aesthetasc; one element. Actual segment 15 with seta; two elements; of unequal size; straight; bifidform; one larger than segment; surpassing to distal margin; not beyond three sequential segments; without spinules; without vestigial seta; without conical seta; without modified seta; with spinous process; on outer margin; surpassing distal margin; with aesthetasc; one element. Actual segment 16 with seta; two elements; of unequal size; straight; one larger than segment; surpassing to distal margin; not beyond three sequential segments; bifidform; without spinules; without vestigial seta; without conical seta; without modified seta; with spinous process; on outer margin; surpassing distal margin; unequal size to process on preceding segment; with aesthetasc; one element. Actual segment 17 with seta; two elements; of unequal size; straight; one larger than segment; surpassing distal margin; bifidform; without spinules; without vestigial seta; without conical seta; with modified seta; one element; stout form; surpassing to distal margin; not beyond of the sequential segment; parallel to antennule direction; without spinous process; without aesthetasc. Actual segment 18 with seta; two elements; of equal size; straight; none larger than segment; without spinules; without vestigial seta; without conical seta; with modified seta; one element; stout form; surpassing distal margin; parallel to antennule direction; without spinous process; without aesthetasc. Actual segment 19 with seta; two elements; of unequal size; plumose; none larger than segment; without spinules; without vestigial seta; without conical seta; with modified seta; two elements; stout form; at least one bifid form; surpassing distal margin; parallel to antennule direction; without spinous process; with aesthetasc; one element. Actual segment 20 with seta; four elements; of unequal size; plumose; one larger than segment; surpassing to distal margin; not beyond three sequential segments; without spinules; with vestigial seta; without conical seta; without modified seta; without spinous process; without aesthetasc. Actual segment 21 with seta; two elements; of equal size; plumose; one larger than segment; surpassing to distal margin; greater 3x than original segment; without spinules; without vestigial seta; without conical seta; without modified seta; without spinous process; without aesthetasc. Actual segment 22 with seta; four elements; of equal size; one larger than segment; plumose; surpassing to distal margin; greater 3x than original segment; without spinules; without vestigial seta; without conical seta; without modified seta; without spinous process; with aesthetasc; one element.

##### Left antennules

Uniramous; Left antennule surpassing to prosome; Left antennule not extending beyond caudal rami. Ancestral segment I and II separated. Ancestral segment II and III fused. Ancestral segment III and IV fused. Ancestral segment IV and V separated. Ancestral segment V and VI separated. Ancestral segment VI and VII separated. Ancestral segment VII and VIII separated. Ancestral segment VIII and IX separated. Ancestral segment IX and X separated. Ancestral segment X and XI separated. Ancestral segment XI and XII separated. Ancestral segment XII and XIII separated. Ancestral segment XIII and XIV separated. Ancestral segment XIV and XV separated. Ancestral segment XV and XVI separated. Ancestral segment XVI and XVII separated. Ancestral segment XVII and XVIII separated. Ancestral segment XVIII and XIX separated. Ancestral segment XIX and XX separated. Ancestral segment XX and XXI separated. Ancestral segment XXI and XXII separated. Ancestral segment XXII and XXIII separated. Ancestral segment XXIII and XXIV separated. Ancestral segment XXIV and XXV separated. Ancestral segment XXV and XXVI separated. Ancestral segment XXVI and XXVII separated. Ancestral segment XXVII and XXVIII fused.

Left antennule actual 25-segmented; not-geniculated. Actual segment 1 with seta; one element; none larger than segment; straight; without spinules; without vestigial seta; without conical seta; without modified seta; without spinous process; with aesthetasc; one element. Actual segment 2 with seta; three elements; of equal size; none larger than segment; straight; without spinules; with vestigial seta; one element; without conical seta; without modified seta; without spinous process; with aesthetasc; one element. Actual segment 3 with seta; one element; one larger than segment; straight; surpassing to distal margin; beyond three sequential segments; without spinules; with vestigial seta; one element; without conical seta; without modified seta; without spinous process; with aesthetasc; one element. Actual segment 4 with seta; one element; none larger than segment; straight; without spinules; without vestigial seta; without conical seta; without modified seta; without spinous process; without aesthetasc. Actual segment 5 with seta; one element; one larger than segment; straight; surpassing distal margin; not beyond three sequential segments; without spinules; with vestigial seta; one element; without conical seta; without modified seta; without spinous process; with aesthetasc; one element. Actual segment 6 with seta; one element; none larger than segment; straight; without spinules; without vestigial seta; without conical seta; without modified seta; without spinous process; without aesthetasc. Actual segment 7 with seta; one element; one larger than segment; straight; surpassing to distal margin; beyond three sequential segments; without spinules; without vestigial seta; without conical seta; without modified seta; without spinous process; with aesthetasc; one element. Actual segment 8 with seta; one element; one larger than segment; straight; surpassing distal margin; without spinules; without vestigial seta; with conical seta; without modified seta; without spinous process; without aesthetasc. Actual segment 9 with seta; two elements; of unequal size; one larger than segment; straight; surpassing to distal margin; beyond three sequential segments; without spinules; without vestigial seta; without conical seta; without modified seta; without spinous process; with aesthetasc; one element. Actual segment 10 with seta; one element; one larger than segment; straight; surpassing distal margin; without spinules; without vestigial seta; without conical seta; without modified seta; without spinous process; without aesthetasc. Actual segment 11 with seta; one element; one larger than segment; straight; surpassing to distal margin; beyond three sequential segments; without spinules; without vestigial seta; without conical seta; without modified seta; without spinous process; without aesthetasc. Actual segment 12 with seta; one element; one larger than segment; straight; surpassing distal margin; without spinules; without vestigial seta; with conical seta; without modified seta; without spinous process; with aesthetasc; one element. Actual segment 13 with seta; one element; none elongated; straight; surpassing distal margin; without spinules; without vestigial seta; without conical seta; without modified seta; without spinous process; without aesthetasc. Actual segment 14 with seta; one element; elongated; straight; surpassing to distal margin; beyond three sequential segments; without spinules; without vestigial seta; without conical seta; without modified seta; without spinous process; with aesthetasc; one element. Actual segment 15 with seta; one element; larger than segment; straight; surpassing to distal margin; not beyond three sequential segments; without spinules; without vestigial seta; without conical seta; without modified seta; without spinous process; without aesthetasc. Actual segment 16 with seta; one element; larger than segment; plumose; surpassing to distal margin; not beyond three sequential segments; without spinules; without vestigial seta; without conical seta; without modified seta; without spinous process; with aesthetasc; one element. Actual segment 17 with seta; one element; not larger than segment; straight; without spinules; without vestigial seta; without conical seta; without modified seta; without spinous process; without aesthetasc. Actual segment 18 with seta; one element; larger than segment; plumose; surpassing to distal margin; beyond three sequential segments; without spinules; without vestigial seta; without conical seta; without modified seta; without spinous process; without aesthetasc. Actual segment 19 with seta; one element; not larger than segment; straight; surpassing distal margin; without spinules; without vestigial seta; without conical seta; without modified seta; without spinous process; with aesthetasc; one element. Actual segment 20 with seta; one element; not larger than segment; straight; surpassing distal margin; without spinules; without vestigial seta; without conical seta; without modified seta; without spinous process; without aesthetasc. Actual segment 21 with seta; one element; larger than segment; plumose; surpassing to distal margin; beyond three sequential segments; without spinules; without vestigial seta; without conical seta; without modified seta; without spinous process; without aesthetasc. Actual segment 22 with seta; two elements; of equal size; one of them elongated; plumose; surpassing to distal margin; without spinules; without vestigial seta; without conical seta; without modified seta; without spinous process; without aesthetasc. Actual segment 23 with seta; one element; none larger than segment; straight; without spinules; without vestigial seta; without conical seta; without modified seta; without spinous process; without aesthetasc. Actual segment 24 with seta; two elements; of equal size; one larger than segment; plumose; surpassing to distal margin; greater 3x than original segment; without spinules; without vestigial seta; without conical seta; without modified seta; without spinous process; without aesthetasc. Actual segment 25 with seta; four elements; of equal size; elongated; plumose; surpassing to distal margin; 4 times larger than segment; without spinules; without vestigial seta; without conical seta; without modified seta; without spinous process; with aesthetasc; one element.

##### Antenna

Biramous. Antenna coxa separated from the basis; bearing seta; 1; on inner surface; at distal corner; reaching to the endopod 1. Antenna basis (fusion) separated from the endopodal segment; bearing seta; 2; on inner surface; at distal corner. Endopodal ancestral segment I and II separated. Ancestral segment II and III fused. Ancestral segment III and IV fused. Ancestral segment III and IV fully. Antenna endopod actual 2-segmented. Actual segment 1 not bilobate; with seta; two; on inner margin; with spinules; as a row; obliquely; on outer surface; with pore. Actual segment 2 bilobate; with discontinuity on outer cuticle; not developed as a suture; inner lobe bearing 8 setae; distally; outer lobe bearing 7 setae; distally; with spinules; as a patch; on outer surface. Antenna exopod ancestral segment I and II separated. Ancestral segment II and III fused. Ancestral segment III and IV fused. Ancestral segment IV and V separated. Ancestral segment V and VI separated. Ancestral segment VI and VII separated. Ancestral segment VII and VIII separated. Ancestral segment VIII and IX separated. Ancestral segment IX and X fused. Antenna exopod actual 7-segmented. Actual segment 1 single; elongated (width-length, equal or larger ratio 2:1); with seta; one; at inner surface. Actual segment 2 compound; elongated (larger width-length ratio 2:1); with seta; three; at inner surface. Actual segment 3 single; not elongated (lesser width-length ratio 2:1); with seta; one; at inner surface. Actual segment 4 single; not elongated (lesser width-length ratio 2:1); with seta; one; at inner surface. Actual segment 5 single; not elongated (lesser width-length ratio 2:1); with seta; one; at inner surface. Actual segment 6 single; not elongated (lesser width-length ratio 2:1); with seta; one; at inner surface. Actual segment 7 compound; elongated (larger or equal width-length ratio 2:1); with seta; one; at inner surface; and three; at distal surface.

##### Oral features

**Mandible**. Coxal gnathobase sclerotized; with lobe; prominent; on caudal margin; presence of cutting blade; with tooth-like prominence; two, distinctly; 1 acute; on caudal margin; and 1 triangular; on sub-caudal margin; without acute projection between the prominences; with additional spinules; as a row; on dorsal surface; with seta; 1; dorsally; on apical surface; with spinules; apicalmost. Mandible palps biramous; comprising the basis; with seta; four; differently inserted; first medially; reaching to beyond the endopod 1; second distally; third distally; fourth distally; on inner margin; none with setulose ornamentation. Mandible endopod 2-segmented. Mandible endopod 1 with lobe; bearing seta; four; distally inserted; without spinules. Mandible endopod 2 without lobe; bearing setae; nine elements; distally inserted; with spinules; as a row; double. Mandible exopod 4-segmented. Mandible exopod 1 with seta; one element; distally; on inner margin. Mandible exopod 2 with seta; one element; distally; on inner side. Mandible exopod 3 with seta; one element; distally; on inner side. Mandible exopod 4 with setae; three elements; on terminal region. **Maxillule**. Birramous. Maxillule 3-segmented. Maxillule praecoxa with praecoxal arthrite; bearing spines; fifteen elements; ten marginally; plus, five sub-marginally; with spinules; as a patch; on sub-marginal surface. Maxillule coxa with coxal epipodite; with conspicuous outer lobe; bearing setae; nine elements; with coxal endite; elongated (larger or equal width-length ratio 2:1); bearing setae; four elements. Maxillule basis with basal endite; double; first proximal; elongated (larger width-length ratio 2:1; separated from basis; with setae; four elements; distally inserted; second distal; fused to basis; not elongated (lesser width-length ratio 2:1); with setae; four elements; distally inserted; with setules; as a row; on inner side; basal exite present; with setae; one element; on outer surface. Maxillule endopod 1-segmented. Endopod 1 bilobate; first proximal; with setae; three elements; second distal; with setae; five elements. Maxillule exopod 1-segmented. Exopod 1 with setae; six elements; with setules; as a row; on inner side; spinules absent. **Maxilla**. Uniramous. Maxilla 5-segmented. Maxilla praecoxa fused to coxa; incompletely; distinct externally; with praecoxal endite; double; first elongated endite (larger or equal width length ratio 2:1); proximally inserted; with seta; straight, or plumose; 1 straight; 4 plumose; with spine; single; without spinules; without setule; second elongated endite (larger or equal width length ratio 2:1); distally inserted; with seta; plumose; 3 plumose; without spine; with spinules; as a row; on distal margin; with setule; as a row; on distal margin; absence of outer seta. Maxilla coxa with coxal endite; double; first elongated endite (larger or equal width); proximally inserted; with seta; plumose; 3 plumose; without spine; without spinules; with setules; as a row; on proximal margin; second elongated endite (larger or equal width); distally inserted; with seta; plumose; 3 plumose; without spine; without spinules; with setules; as a row; on proximal margin; absence of outer seta. Maxilla basis with basal endite; single; elongated (larger or equal width-length ratio 2:1); with seta; plumose; 3 plumose; without spinules; absence of outer seta. Maxilla endopod 2-segmented. Endopod 1 with seta; 2 plumose; without spine; without spinules; without setules. Maxilla endopod 2 with seta; 2 plumose; without spine; without spinules; without setules. **Maxilliped**. Uniramous; Maxilliped 8-segmented. Maxilliped praecoxa fused to coxa; incompletely; distinct internally; with praecoxal endite; not elongated (lesser width-length ratio 2:1); distally inserted; with seta; 1 straight; with spinules; as a row; single; on basal surface; without setules. Maxilliped coxa with coxal endite; three coxal endite; first elongated (larger or equal width); proximally inserted; with seta; 2 plumose; with spinules; as a patch; single; on apical surface; without setules; second not elongated (lesser width-length ratio 2:1); medially inserted; with seta; 3 plumose; with spinules; as a row; single; on medial surface; without setules; third elongated (larger or equal width length ratio 2:1); distally inserted; with seta; 3 plumose; none reaching to beyond of the basis; with spinules; as a row; single; on basal surface; without setules; with lobe; prominence; at inner distal angle; ornamented; with spinules; continuously on margin. Maxilliped basis without basal endite; with seta; 3 plumose; with spinules; as a row; single; on medial surface; with setules; as a row; single; on inner margin. Maxilliped endopod segment 6-segmented. Endopod 1 with seta; 2 plumose; on inner surface. Endopod 2 with seta; 3 plumose; on inner surface. Endopod 3 with seta; 2 plumose; on inner surface. Endopod 4 with seta; 2 plumose; on inner surface. Endopod 5 with seta; 2 plumose; on inner surface, or on outer surface; outer seta absent. Endopod 6 with seta; 4 plumose; on inner surface, or on outer surface.

##### Swimming legs features

**First swimming legs.** Symmetrical; biramous. First swimming legs intercoxal plate without seta. First swimming legs praecoxa absent. First swimming legs coxa with seta; one; straight; distally inserted; on inner surface; surpassing to first endopodal segment; with setules; two group; as a patch; on inner margin; and as a row; double; on anterior surface; outerly; without spinules; without spine. First swimming legs basis without seta; with setules; as a patch; single; on outer surface; without spinules; without spine. First swimming legs endopod 2-segmented. Endopod 1 with seta; straight; restricted; to inner surface; one element; without spine; with setules; as a row; single; continuously; on outer surface; without spinules; absence of Schmeil’s organ. Endopod 2 with seta; unrestricted; three on inner surface; one on outer surface; two on distal surface; straight; without spine; with setules; as a row; single; continuously; on outer surface; without spinules; absence of Schmeil’s organ. Endopod 3 absence. First swimming legs exopod 1 with seta; restricted; 1 on inner surface; with spine; 1; stout; smaller than original segment; serrated; on inner side; continuously; with setules; as a row; single; as a row; innerly. First swimming legs exopod 2 with seta; restricted; 1 on inner surface; straight; without spine; with setules; as a row; single; continuously; on inner margin, or on outer margin; without spinules. First swimming legs exopod 3 with setule; as a row; single; continuously; on outer surface; without spinules; with seta; unrestricted; 2 on inner surface; 2 on terminal surface; with spine; 2; unequal size; first no longer 2x than origin segment; stout; serrated; on inner side, or on outer side; equally; second longer 3x than origin segment; slender; serrated; on outer side; with ornamentation on non-serrated side; by setules. **Second swimming legs**. Symmetrical; Second swimming legs biramous. Second swimming legs intercoxal plate without seta. Second swimming legs praecoxa present; located laterally. Second swimming legs coxa with seta; straight; distally inserted; on inner surface; surpassing to basal segment; without setules; without spinules; without spine. Second swimming legs basis without seta; without setules; without spinules; without spine. Second swimming legs endopod 3-segmented. Endopod 1 with seta; straight; restricted; one on inner surface; without spine; with setules; as a row; single; continuously; on outer surface; without spinules; absence of Schmeil’s organ. Endopod 2 with seta; straight; unrestricted; two on inner surface; without spine; with setules; as a row; single; continuously; on outer side; without spinules; presence of Schmeil’s organ; on posterior surface. Endopod 3 with seta; straight; unrestricted; three on inner surface; two on outer surface; two on distal surface; without spine; without setules; with spinules; as a row; double; distally inserted; at anterior surface; absence of Schmeil’s organ. Second swimming legs exopod 1 with seta; restricted; one on inner surface; with spine; 1; stout; not reaching to distal-third of the exopod 2; serrated; on inner side, or on outer side; with setules; as a row; single; continuously; on inner side; without spinules; absence of Schmeil’s organ. Exopod 2 with seta; unrestricted; one on inner surface; with spine; 1; stout; not surpassing the exopod 3; serrated; on inner side, or on outer side; with setules; as a row; single; continuously; on inner surface; without spinules; absence of Schmeil’s organ. Exopod 3 with seta; plurimarginal; three on inner surface; two on terminal surface; with spine; 2; unequal size; first no longer 2x than origin segment; stout; serrated; on inner side, or on outer side; equally; second longer 2x than origin segment; slender; serrated; on outer side; with ornamentation on non-serrated side; of setules; setules on outer surface; as a row; single; continuously; on inner surface; with spinules; as a row; single; distally inserted; at anterior surface; absence of Schmeil’s organ. **Third swimming legs**. Symmetrical; Third swimming legs biramous. Third swimming legs intercoxal plate without seta. Third swimming legs praecoxa present; not laterally located. Third swimming legs coxa with seta; straight; distally inserted; on inner surface; surpassing to first endopodal segment; without setules; without spinules; without spine. Third swimming legs basis without seta; without setules; without spinules; without spine. Third swimming legs endopod 3-segmented. Endopod 1 with seta; restricted; one on inner surface; without spine; without setules; without spinules; absence of Schmeil’s organ. Endopod 2 with seta; restricted; two on inner surface; straight; without spine; without setules; without spinules; absence of Schmeil’s organ. Endopod 3 with seta; straight; plurimarginal; two on inner surface; two on outer surface; three on terminal surface; without spine; without setules; with spinules; as a row; distally inserted; double; at anterior surface; absence of Schmeil’s organ. Third swimming legs exopod 1 with seta; restricted; straight; one on inner surface; with spine; 1; stout; not reaching to the distal-third of the exopod 2; serrated; equally; on inner surface, or on outer surface; with setules; as a row; single; continuously; on inner surface; without spinules; absence of Schmeil’s organ. Exopod 2 with seta; straight; restricted; one on inner surface; with spine; 1; stout; not reaching out to exopod 3; serrated; on inner side, or on outer side; equally; with setules; as a row; single; continuously; on inner side; without spinules; absence of Schmeil’s organ. Exopod 3 without setules; with spinules; as a row; single; distally inserted; at anterior surface; with seta; straight; unrestricted; three on inner surface; two on terminal surface; with spine; 2; unequal size; first no longer 2x than origin segment; stout; serrated; on inner side, or on outer side; equally; second longer 2x than origin segment; slender; serrated; on outer side; with ornamentation on non-serrated side; of setules; absence of Schmeil’s organ. **Fourth swimming legs**. Symmetrical; biramous. Intercoxal plate without sensilla. Praecoxa present. Coxa with seta; distally inserted; on inner margin; reaching out to endopod 1; without spinules; setules absent. Basis with seta; one; medially inserted; on posterior surface; smaller than the original segment; without setules; without spinules; without spine. Fourth swimming legs endopod 3-segmented. Endopod 1 with seta; one; restricted; on inner surface; without spine; without setules; without spinules; absence of Schmeil’s organ. Endopod 2 with seta; restricted; two on inner side; without spine; with setules; as a row; single; continuously; on outer surface; without spinules; absence of Schmeil’s organ. Endopod 3 with seta; unrestricted; two on inner surface; two on outer surface; three on distal surface; without spine; without setules; with spinules; as a row; double; distally inserted; at anterior surface; absence of Schmeil’s organ. Fourth swimming legs exopod 1 with seta; restricted; one on inner surface; with spine; 1; stout; not reaching out to distal-third of the exopod 2; serrated; on inner side, or on outer side; equally; with setules; as a row; single; continuously; on inner surface; without spinules; absence of Schmeil’s organ. Exopod 2 with seta; restricted; one on inner surface; with spine; 1; stout; not reaching the end of exopod 3; serrated; on inner side, or on outer side; equally; with setules; as a row; single; continuously; on inner surface; without spinules; absence of Schmeil’s organ. Exopod 3 without setules; with spinules; as a row; single; distally inserted; at anterior surface; with seta; unrestricted; three on inner surface; two on distal surface; with spine; 2; unequal size; first no longer 2x than origin segment; stout; serrated; on inner side, or on outer side; equally; second longer 2x than origin segment; slender; serrated; on outer side; without ornamentation on non-serrated side; absence of Schmeil’s organ.

##### Fifth swimming legs features

Asymmetrical. Fifth swimming leg intercoxal plate with length equal or greater than width on 1.5x; with regular proximal margin; discontinuous to; the anterior margin of the left coxa; posterior sensilla on the right lateral absent. **Fifth left swimming leg**. Fifth left swimming leg biramous; leg reaching first right exopod segment; medially. Fifth left swimming leg praecoxa present; rudimentary; separated from the coxae; without ornamentation. Fifth left swimming leg coxa convex inner side; without teeth-like structures; with process; triangulated; on posterior surface; outer side; distally inserted; not projecting over basis; with sensilla; slender; triangular; at apex; longer 2x than insertion basis; without swelling; without seta; without spinules. Fifth left swimming leg basis sub-cylindrical; unequal size between inner and outer side; shorter outer than inner side; with rectilinear inner side; rounded internal proximal expansion absent; without outgrowth; with groove; deep; obliquely; on posterior surface; not reaching the endopodal lobe; not ornamented; absence of protuberance; with seta; outerly inserted; no longer 2x than origin segment; absence of minutely granular. Fifth left swimming leg endopod segments 1 and 2 fused; segments 2 and 3 fused; 1-segmented; stout; separated from the basis; ornamented; on inner side; with spinules; more than four elements; as a row; terminally; row of setules absent; without seta. Fifth left swimming leg exopod segments 1 and 2 separated; segments 2 and 3 fused; 2-segmented; stout; separated from the basis. Fifth left swimming leg exopod 1 sub-triangular; longer than broad; unequal size between inner and outer side; shorter inner than outer side; concave inner side; convex outer side; without swelling; without marginal extension; without process; with lobe; single; semicircular; medially inserted; on inner side; covered; by setules; without outer spine; absence seta. Fifth left swimming leg exopod 2 digitiform; longer than broad; equal size between inner and outer side; disform inner side; with rectilinear outer side; setulose pad present; not prominently rounded; proximally; on inner side; inflated medial region absent; distal process present; digitiform; denticulate; bicuspidate; with transverse row of denticles; at anterior surface; not innerly directed; with seta; spiniform; not ornamented by spinules; surpassing the distal-point of the segment; without outer spine; terminal claw absent.

##### Fifth right swimming leg

Biramous. Fifth right swimming leg praecoxa present; separated from the coxae; without ornamentation. Fifth right swimming leg coxa convex inner side; without teeth-like structures; with process; rounded; distally inserted; on posterior surface; closest to the outer rim; projecting over basis; not beyond the first third; without triangular protuberance innerly; with sensilla; slender; at apex; no longer 2x than basal insertion; with marginal extension; without seta; without spinules. Fifth right swimming leg basis cylindrical; unequal size between inner and outer side; shorter outer than inner side; rectilinear inner side; tumescence absent; without protuberance; absence of distinct minutely granular; additional inner process absent; without posterior groove; with seta; outerly inserted; on posterior surface; no longer 2x than origin segment; posterior protrusion present; distal process absent. Fifth right swimming leg with endopodite present; fused to basis; on anterior surface; ancestral segments 1 and 2 fused; ancestral segments 2 and 3 fused; stout; ornamented; with setules; as a row; on inner side; terminally; without seta. Fifth right swimming leg exopod segments 1 and 2 separated; segments 2 and 3 fused; 2-segmented; stout; separated from the basis. Fifth right swimming leg exopod 1 trapezium; broader than long; nearly 1.25 times; unequal size between both sides; shorter inner than outer side; rectilinear inner side; convex outer side; with marginal extension; sub-triangular; distally inserted; at outer rim; spinules absent; with process; rounded; sclerotized; without ornamentation; distally inserted; at marginal surface; projecting over next segment; without outer spine; without seta; internal prominence absent; lamella on posterior surface absent. Fifth right swimming leg exopod 2 cylindrical; longer than broad; nearly 2.5 times; equal size between both sides; uniform inner side; convex outer side; without posterior proximal swelling; inner-posterior process absent; without marginal expansion; curved ridge on distal posterior surface present; chitinous knobs present; with 1–2 posteriorly; with outer spine; inserted sub-distally; arched; internally directed; not ornamented innerly; not ornamented outerly; sharp tip; with apparent curve; innerly directed; lesser than the length of the exopod 2; beyond to 2 times its size; 6x; sensilla absent; terminal claw present; equal or longer 1.5 times than insertion segment; sclerotized; arched; inward; without conspicuous curve; ornamented innerly; by spinules; as a row; partially on extension; medially; not ornamented outerly; sharp tip; not curved tip; without medial constriction; hyaline process absent.

##### FEMALE

Body longer and wider than male; Female body 1407 micrometers excluding caudal setae. Widest at first metasome segment. Distal margin of the prosomal segments without one line of setules at posterior margin. Prosome segments with spinules at least at one prosomal segment. Fourth metasome segment presence of dorsal protuberance; digitiform; 2x wider than long; inserted medially; without posterior process; without anterior process; fourth metasome segment without proximal sensillae present. Fourth and fifth metasome segments fused; partially; on dorsal surface. Limit between fourth and fifth metasome segments without ornamentation. **Fifth metasome segment**. Fifth metasome segment without sensilla; with epimeral plates. Epimeral plates asymmetrical. Right epimeral plates prominent, as projections; thinner than the left; one posterior-dorsally directed; reaching half length of the genital segment; with sensilla at the apex; dorsal-posterior sensilla present; stout; without ornamentation. Left epimeral plate without expansion.

##### Urosome

3-segmented. **Genital double-somite**. Asymmetrical in dorsal view; longer than broad; longer than other urosomites combined; dorsal suture at mid-length absent; not covered by spinules; with swelling; rounded; equal size; anteriorly; with sensillae; on both sides; one; stout; with robust apex; at left rim; not on lobular base; anteriorly; one; stout; at right lateral; not on lobular base; anteriorly; with robust apex; of unequal size between then; greater left than right; lateral protuberance absent; with right posterior rim expanded; over next segment; with slender sensilla on each posterior rim; without posterior-dorsal process. Genital double-somite opercular pad present; broader than longer; symmetrical; development laterally; expanded posteriorly; covering partially; double gonoporal slit; located ventrally; with arthrodial membrane; inserted anteriorly; post-genital process absent; disto-ventral tumescence absent; ventral vertical folds absent; dorsal sensilla absent. Second urosome segment without ventral fusion to anal segment; right distal process absent. Caudal rami patch of setules on outer surface absent; patch of spinules on outer surface absent.

##### Appendices features

Rostrum basal process absent. **Antennules**. Symmetrical. Right antennule surpassing to genital double-segment; not extending beyond caudal rami; ornamentation pattern equals to male left antennule; fully.

##### Fifth swimming legs

Symmetrical; Fifth swimming legs biramous. Fifth swimming legs intercoxal plate longer than wide; separated from the legs. Fifth swimming legs praecoxa without sclerite praecoxal. Fifth swimming legs coxa with process; conical; at the outer rim; distally; sensilla present; stout; at apex; projecting over basal segment; no longer 2x than basal insertion; marginal extension absent; without swelling; without seta; without spinules. Fifth swimming legs basis sub-triangular; equal size between inner and outer sides; with concave inner side; without proximal inner outgrowth; without groove; with distal extension; on posterior surface; with seta; outerly inserted; on anterior surface; no longer 2x than origin segment. Fifth swimming legs endopod segments 1 and 2 separated; segments 2 and 3 fused; 2-segmented; with complete suture; stout; separated from the basis; ornamentation on segment 2; with spinules; as a row; single; non-oblique; terminally; at anterior surface; with seta; double; one medially; on posterior surface; rectilinear; one distally; on posterior surface; rectilinear; of unequal size; distal seta longer than medial seta. Fifth swimming legs exopod segments 1 and 2 separated; segments 2 and 3 separated; 3-segmented; separated from the basis. Fifth swimming legs exopod 1 sub-cylindrical; longer than wide; longer or equal than 2 times; with unequal size between inner and outer side; shorter inner than outer side; with convex inner side; with convex outer side; without swelling; without marginal extension; without posterior process; without spine; without seta. Fifth swimming legs exopod 2 sub-cylindrical; longer than broad; longer or equal than 2 times; without swelling; without marginal extension; without process; without lobe; with spine; inserted laterally; rectilinear; without ornamentation; sharp tip; lesser than next segment; without seta. Fifth swimming legs exopod 3 cylindrical; longer than wide; without swelling; without process; without lobe; without spine; with seta; double; inserted terminally; unequal size between them; outer seta smaller than inner; nearly 3 times; outer seta not ornamented by setules; without ornamentation; presence of terminal claw; sclerotized; rectilinear; with ornamentation; of denticles; as a row; on surface partially; at medial region; rectilinear outer side; with ornamentation; of denticles; as a row; on surface partially; at medial region; blunt tip; 6 times longer than origin segment.

##### Distribution records

###### BRAZIL

**Amazonas**: Joanico Lake, Careiro Island, near to Manaus (Herbst, 1967); Janauari Lake, Negro River, near to Manaus (Brandorff, 1972; 1973b); Paraná of the Curari; Lake of the Rei, Careiro Island, Amazonas River; Manacapuru River, Lake of the Piranha, and Paraná do Piranha (Brandorff, 1972); Lake Calado, Amazonas River, near to Manacapuru City (Brandorff, 1972; Santos-Silva, 1991); Lake Jacaretinga, Amazonas River (Hardy, 1980); Lake Grande, Amazonas River, 03°22’S, 60°35W (Carvalho, 1983); Lake Lua Nova, Acre River (Sendacz & Melo Costa, 1991); Amanã Lake, Japurá River (Santos-Silva & Robertson, 1993). **Pará**: vicinity of Santarém, probably in the Tapajós River (Wright, 1927); Grande Curuai Lake, in front of Fazenda Nova Itália (Brandorff, 1972; 1973b), and in front of Caraubal, Amazonas River; Tapajós River at the Santarém City; Lake Salgado, Boi, and Cabeceira do Molha Headwaters; Jurucui Lake, Tapajós River, near to the village of Alter-do-Chão; Amazonas River, near to Santarém; Paraná of the Tapará, near to Santarém (Brandorff, 1972). **Rondônia**: Calama, Madeira River (Wright, 1927); São Pedro Stream, 09°36’S, 63°37’W (Santos-Silva & Robertson, 1993). **Mato Grosso do Sul**: southern Pantanal, region of Corumbá, Paraguay River: near to Marinha Ladário, near to Port, near to Rabicho, site 2 near airport (Corumbá), Baía de Carandazal (baía 29) at Fazenda Nhumirim (18°59’S, 56°39’W), Jacadigo bay (19°01’S, 57°41’W) (Reid & Moreno, 1990). **Paraná**: Itaipu Reservoir (Matsumura-Tundisi, 1986). BOLIVIA. Beni (Brandorff, 1976). PARAGUAY. Several samples from Makthlawaiya, 23°25’S, 58°19W and Nanahua, 32°30’S, 59°30’W, regions (Lowndes, 1934); upper Paraguay River (Perbiche-Neves *et al*., 2015). ARGENTINA. Middle Paraná River between the cities of Santa Fé and Paraná (Paggi & José de Paggi, 1974); Middle Paraná River (José de Paggi, 1981; Paggi & José de Paggi, 1990); main course of the Paraná River between Santa Fe and Buenos Aires (José de Paggi, 1978). **Buenos Aires**: Delta of Río Paraná, near Tigre (Wright, 1939); Paraná Guazú, Tigre (Brehm, 1957, 1965); stream at Pergamino and canal of Santiago River at Puerto La Plata (Ringuelet, 1958a); Hoya del Plata (Ringuelet, 1962); la Plata River at the Punta Lara (Cicchino, 1972); Arroyo Pajarito, Tigre, and Terito River near to Tigre (Reid, 1991). **Chaco**: Resistencia (Ringuelet, 1958a); Barranqueras River (Brehm, 1965); Guaycurú River; La Palometa River (Dussart & Frutos, 1986). **Corrientes**: Laguna 1 (La Turbia), Isla del Cerrito, Paraná River and Laguna 2 (Los Pajaros), Isla Nueva Cerrito, Paraná River (Frutos, 1993). **Formosa**: Yema Lake (Brehm, 1957, 1965); Ingeniero Juarez (Brehm, 1965); San Hilario stream (Dussart & Frutos, 1987).

##### Habitat

Habitat in freshwaters: reservoir, swamps, floodplain lake, pools, and streams.

##### Remarks

The taxon was presented to science from organisms collected in a lake environment of the Amazon, such as *D.* (s.l.) *coniferoides*. Ringuelet (1958) proposed the recombination of the species to *Notodiaptomus* without presenting evidence or morphological set, which added heterogeneity to the genus of Kiefer (1936). Based on these intrageneric variations, Dussart (1985) proposed a new recombination for the subgenus *Notodiaptomus* (*Caleodiaptomus*), where the species would be the type and subcongenus with *N. transitans.* Later this proposal became invalid due to the weak or non-existent taxonomic foundations.

One year earlier, the species was reported to Venezuela (Dussart, 1984), erroneously. In fact it was a specimen with similar morphology, but with important variations for male fifth right swimming leg exopod 1, and female genital double-somite. The masculine structure cited as wider than long, female with genital double-somite with lateral protuberance on left side, and suture at mid-length dorsally were described as *N. simillimus* Cicchino, Santos-Silva & Robertson (2001) further on. Other important variations can be present between both species (Cicchino *et al*., 2001), among them for *N. coniferoides*: female fourth metasome segment with medial dorsal protuberance in digitiform, and 2x wider than long.

The original condition of part of these mentioned structures had already been noted in Brandorff (1972) for individuals from lake systems of the Brazilian Amazônia, opportunely described as *D.* (s.l.) *coniferoides*. Dussart & Frutos (1986) corroborated this information through comments and illustrations (fig. 10-12, “*planchet II*”) of organisms from the Southern Neotropical (Argentina). In a recent review of the species Perbiche-Neves *et al*. (2015) offered important illustrations of organisms from the Paraguay River (Southern Neotropics) and confirmed Wright (1927), and the differences between the species and *N. simillimus*.

Among the original morphological characteristics and amplification of *Notodiaptomus*, the taxon organisms are one of the least convergent, along with *N. transitans*, *N. paraensis*, *N. santaremensis*, *N. simillimus*, *N. caperatus,* and *N.* cannarensis. Nevertheless, the masculine pattern of the A1R is similar to that of the type-species (except for length of the spiniform modified seta on actual segment 13), such as male fifth right swimming leg exopod 2 with curved ridge on distal posterior surface, and endopod 2-segmented. Undoubtedly, future evidence may indicate a taxonomic repositioning of the species together with the other organisms mentioned.

#### Notodiaptomus dentatus Paggi, 2001

##### Synonymy

*Notodiaptomus anisitsi*; Ringuelet & Martínez de Ferrato 1967: 417, 7–10; Paggi, 1978: 150–151; 1980: 72; 1981: 199; 1983: 168; 1984: 141. *Notodiaptomus dentatus* Paggi, 2001: 554–556, figs. 38– 65, and 93, tab. 1; Perbiche-Neves *et al*., 2015: 53–56, figs. 44–46; Perbiche-Neves *et al*., 2020: 687, key to the Neotropical diaptomid, fig. 21.10 G.

##### Type locality

Madrejón Don Felipe, ox-bow lake near Santa Fe, Santa Fe Province, Argentina. Material examined was collected on IV.1969, V.1969; VIII.1969; X.1969, and XII.1969. No date on collector.

##### Type material

Holotype: Male (n°. 34203), entire, deposited in Museo Argentino de Ciencias Naturales ‘Bernardino Rivadavia’, Buenos Aires. Paratypes: Female (n°. 34204), entire, and deposited in Museo Argentino de Ciencias Naturales ‘Bernardino Rivadavia’, Buenos Aires. 1 female and 1 male deposited in Museo Provincial de Ciencias Naturales Dr. Florentino Ameghino. All preserved in a mixture of formaldehyde and glycerol (1/10),

##### Material examined

Non-type material: 2 male, and 3 females, entire in alcohol (MZUSP 32933), from the middle Yaciretá Dam Reservoir, located on the Parana River between Argentina and Paraguay, G. Perbiche-Neves coll., VI.2010, and stored in MZUSP. 1 male (INPA-COP019, slides a-h) and 1 female (INPA-COP020, slides a-h) were selected to be dissection on eight slides each and deposited in the Zoological Collection of the INPA, Brazil.

##### Diagnosis

**(1)** Cephalosome without dorsal suture; **(2)** male limit between fourth and fifth metasome segments with dorsal spinules row singly; **(3)** caudal rami with outer proximal spiniform process on outermost seta; **(4)** male right antennule actual segment 8 with conical seta reaching to middle-point of the sequent segment; **(5)** male right antennule actual segment 13 with modified seta parallel to antennule direction; **(6)** male fifth left swimming leg exopod 2 sub-triangular, and with distal process in acute-form inwardly; **(7)** male fifth right swimming leg coxa with posterior process projecting over basis not beyond the first third; **(8)** male fifth right swimming leg exopod 1 longer than broad 2x nearly; **(9)** male fifth right swimming leg exopod 2 in elliptical, longer than broad 2x nearly; **(10)** male fifth right swimming leg exopod 2 with 4 chitinous knobs posteriorly; **(11)** male fifth right swimming leg exopod 2 with outer spine not ornamented innerly, beyond 2x lesser than the length of the exopod 2; **(12)** female genital double-somite with lateral rounded protuberance on the right side posteriorly; **(13)** female genital double-somite with right posterior rim expanded, over next segment; **(14)** female fifth swimming legs basis with outer seta longer 2x than origin segment, reaching to exopod 1 distally; **(15)** female fifth swimming legs endopod without discontinuity cuticle innerly.

##### Redescription

###### MALE

Body 1051 micrometers excluding caudal setae. Male body smaller and slenderer than female. Nerve axons myelinated. Prosome 6-segmented; widest at first metasome segment; without one line of setules at posterior margin; with spinules at least at one segment. Cephalosome anterior margin rounded; without dorsal suture; separate from first metasome segment. First metasome segment without sensilla. Second metasome segment without sensilla. Third metasome segment without sensillae; ornamented posterior margin; with spinules; as a row; single; dorsally. Fourth metasome segment with sensillae; 2 dorsally; of equal size; separated from the fifth metasome. Limit between fourth and fifth metasome segments ornamented; with spinules; as a row; on dorsal singly; on lateral singly; same size. Fifth metasome segment with sensilla; 2 dorsally; Fifth metasome segment equal size; Fifth metasome segment without ornamentation; Fifth metasome segment without dorsal conical process; with epimeral plates. Epimeral plates symmetrical. Right epimeral plates prominent, as projections; one projection; posterior-laterally directed; not reaching half length of the genital segment; with sensilla; at the apex of projection; without ornamentation.

##### Urosome

5-segmented; Urosome 5-free segments. Genital somite asymmetrical in dorsal view; with single aperture; located on left side; ventrolaterally on posterior rim; with sensillae; on single side; one; at left lateral; posteriorly. Third urosome segment without spinules; without external seta. Fourth urosome segment without spinules; without sub-conical blunt dorsal-lateral process. Anal segment presence of dorsal sensillae; one on each side; medially inserted; presence of operculum; convex; covering the anal aperture fully. Caudal rami symmetrical; separated from anal segment; longer than wide; with setules; continuous on; inner side; each ramus bearing 6 caudal setae; 5 marginals; plumose; and 1 internal dorsally; straight; not reticulated main axis; outermost seta with outer spiniform process present.

##### Appendices features

Rostrum symmetrical; separated from dorsal cephalic shield; by complete suture; sensillae present; one pair; medially inserted on tegument; with rostral filament; double; paired; extended; into point; with basal process; in ventral view, rounded on left side; without a smaller basal expansion on the right side.

##### Antennules

Asymmetrical. **Right antennules**. Uniramous; right antennule surpassing to genital segment; right antennule extending beyond caudal rami.

Right antennule ancestral segment I and II separated. Ancestral segment II and III fused. Ancestral segment III and IV fused. Ancestral segment IV and V separated. Ancestral segment V and VI separated. Ancestral segment VI and VII separated. Ancestral segment VII and VIII separated. Ancestral segment VIII and IX separated. Ancestral segment IX and X separated. Ancestral segment X and XI separated. Ancestral segment XI and XII separated. Ancestral segment XII and XIII separated. Ancestral segment XIII and XIV separated. Ancestral segment XIV and XV separated. Ancestral segment XV and XVI separated. Ancestral segment XVI and XVII separated. Ancestral segment XVII and XVIII separated. Ancestral segment XVIII and XIX separated. Ancestral segment XIX and XX separated. Ancestral segment XX and XXI separated. Ancestral segment XXI and XXII fused. Ancestral segment XXII and XXIII fused. Ancestral segment XXIII and XXIV separated. Ancestral segment XXIV and XXV fused. Ancestral segment XXV and XXVI separated. Ancestral segment XXVI and XXVII separated. Ancestral segment XXVII and XXVIII fused.

Right antennule actual 22-segmented; geniculated; between the segment 18 and segment 19; with swollen and modified region; formed by 5 segments; between 13 and 17 segments. Actual segment 1 with seta; one element; straight; none larger than segment; without spinules; without vestigial seta; without conical seta; without modified seta; without spinous process; with aesthetasc; one element. Actual segment 2 with seta; three elements; of unequal size; straight; none larger than segment; without spinules; with vestigial seta; one element; without conical seta; without modified seta; without spinous process; with aesthetasc; one element. Actual segment 3 with seta; one element; one larger than segment; surpassing to distal margin; not beyond three sequential segments; straight; blunt apex; without spinules; without vestigial seta; without conical seta; without modified seta; without spinous process; with aesthetasc; one element. Actual segment 4 with seta; one element; one larger than segment; surpassing to distal margin; straight; not beyond three sequential segments; without spinules; without vestigial seta; without conical seta; without modified seta; without spinous process; without aesthetasc. Actual segment 5 with seta; one element; straight; one larger than segment; surpassing to distal margin; not beyond three sequential segments; without spinules; with vestigial seta; one element; without conical seta; without modified seta; without spinous process; with aesthetasc; one element. Actual segment 6 with seta; one element; one larger than segment; surpassing to distal margin; not beyond three sequential segments; straight; without spinules; without vestigial seta; without conical seta; without modified seta; without spinous process; without aesthetasc. Actual segment 7 with seta; one element; straight; one larger than segment; surpassing to distal margin; beyond three sequential segments; blunt apex; without spinules; without vestigial seta; without conical seta; without modified seta; without spinous process; with aesthetasc; one element. Actual segment 8 with seta; one element; straight; none larger than segment; without spinules; without vestigial seta; with conical seta; one element; reaching to middle-point of the sequent segment; without modified seta; without spinous process; without aesthetasc. Actual segment 9 with seta; two elements; of unequal size; straight; one larger than segment; surpassing to distal margin; beyond three sequential segments; blunt apex; without spinules; without vestigial seta; without conical seta; without modified seta; without spinous process; with aesthetasc; one element. Actual segment 10 with seta; one element; straight; none larger than segment; without spinules; without vestigial seta; without conical seta; with modified seta; presenting blunt apex; slender form; surpassing to distal margin; beyond of the sequential segment; parallel to antennule direction; without spinous process; without aesthetasc. Actual segment 11 with seta; one element; straight; one larger than segment; surpassing to distal margin; not beyond three sequential segments; without spinules; without vestigial seta; without conical seta; with modified seta; slender form; presenting blunt apex; surpassing to distal margin; beyond of the sequential segment; parallel to antennule direction; shorter length than homologous of actual segment 13; without spinous process; without aesthetasc. Actual segment 12 with seta; one element; straight; one larger than segment; surpassing to distal margin; not beyond three sequential segments; without spinules; without vestigial seta; with conical seta; one element; smaller than to segment 8; without modified seta; without spinous process; with aesthetasc; one element; absent internal perpendicular fission. Actual segment 13 with seta; one element; straight; one larger than segment; surpassing to distal margin; not beyond three sequential segments; without spinules; without vestigial seta; without conical seta; with modified seta; stout form; surpassing to distal margin; to the middle-point of the sequence segment; parallel to antennule direction; presenting bifid apex; without spinous process; without aesthetasc. Actual segment 14 with seta; one element; straight; one larger than segment; surpassing to distal margin; beyond three sequential segments; blunt apex; without spinules; without vestigial seta; without conical seta; without modified seta; without spinous process; with aesthetasc; one element. Actual segment 15 with seta; two elements; of unequal size; straight; not bifidform; none larger than segment; without spinules; without vestigial seta; without conical seta; without modified seta; with spinous process; on outer margin; surpassing distal margin; with aesthetasc; one element. Actual segment 16 with seta; two elements; of unequal size; straight; one larger than segment; surpassing to distal margin; not beyond three sequential segments; not bifidform; without spinules; without vestigial seta; without conical seta; without modified seta; with spinous process; on outer margin; surpassing distal margin; unequal size to process on preceding segment; with aesthetasc; one element. Actual segment 17 with seta; two elements; of unequal size; straight; none larger than segment; bifidform; without spinules; without vestigial seta; without conical seta; without modified seta; without spinous process; without aesthetasc. Actual segment 18 with seta; two elements; of equal size; straight; none larger than segment; without spinules; without vestigial seta; without conical seta; without modified seta; without spinous process; without aesthetasc. Actual segment 19 with seta; two elements; of unequal size; plumose; none larger than segment; without spinules; without vestigial seta; without conical seta; without modified seta; at least one bifid form; without spinous process; with aesthetasc; one element. Actual segment 20 with seta; four elements; of unequal size; straight; one larger than segment; surpassing to distal margin; beyond three sequential segments; without spinules; without vestigial seta; without conical seta; without modified seta; with spinous process; distally; not reaching beyond of distal-point segment 21; without aesthetasc. Actual segment 21 with seta; two elements; of equal size; plumose; one larger than segment; surpassing to distal margin; greater 3x than original segment; without spinules; without vestigial seta; without conical seta; without modified seta; without spinous process; without aesthetasc. Actual segment 22 with seta; four elements; of equal size; one larger than segment; plumose; surpassing to distal margin; greater 3x than original segment; without spinules; without vestigial seta; without conical seta; without modified seta; without spinous process; with aesthetasc; one element.

##### Left antennules

Uniramous; Left antennule surpassing to prosome; Left antennule extending beyond caudal rami. Ancestral segment I and II separated. Ancestral segment II and III fused. Ancestral segment III and IV fused. Ancestral segment IV and V separated. Ancestral segment V and VI separated. Ancestral segment VI and VII separated. Ancestral segment VII and VIII separated. Ancestral segment VIII and IX separated. Ancestral segment IX and X separated. Ancestral segment X and XI separated. Ancestral segment XI and XII separated. Ancestral segment XII and XIII separated. Ancestral segment XIII and XIV separated. Ancestral segment XIV and XV separated. Ancestral segment XV and XVI separated. Ancestral segment XVI and XVII separated. Ancestral segment XVII and XVIII separated. Ancestral segment XVIII and XIX separated. Ancestral segment XIX and XX separated. Ancestral segment XX and XXI separated. Ancestral segment XXI and XXII separated. Ancestral segment XXII and XXIII separated. Ancestral segment XXIII and XXIV separated. Ancestral segment XXIV and XXV separated. Ancestral segment XXV and XXVI separated. Ancestral segment XXVI and XXVII separated. Ancestral segment XXVII and XXVIII fused.

Left antennule actual 25-segmented; not-geniculated. Actual segment 1 with seta; one element; none larger than segment; straight; without spinules; without vestigial seta; without conical seta; without modified seta; without spinous process; with aesthetasc; one element. Actual segment 2 with seta; three elements; of equal size; none larger than segment; straight; without spinules; without vestigial seta; without conical seta; without modified seta; without spinous process; with aesthetasc; one element. Actual segment 3 with seta; one element; one larger than segment; straight; surpassing to distal margin; beyond three sequential segments; without spinules; without vestigial seta; without conical seta; without modified seta; without spinous process; with aesthetasc; one element. Actual segment 4 with seta; one element; none larger than segment; straight; without spinules; without vestigial seta; without conical seta; without modified seta; without spinous process; without aesthetasc. Actual segment 5 with seta; one element; one larger than segment; straight; surpassing to distal margin; not beyond three sequential segments; without spinules; without vestigial seta; without conical seta; without modified seta; without spinous process; with aesthetasc; one element. Actual segment 6 with seta; one element; none larger than segment; straight; without spinules; without vestigial seta; without conical seta; without modified seta; without spinous process; without aesthetasc. Actual segment 7 with seta; one element; one larger than segment; straight; surpassing to distal margin; beyond three sequential segments; without spinules; without vestigial seta; without conical seta; without modified seta; without spinous process; with aesthetasc; one element. Actual segment 8 with seta; one element; one larger than segment; straight; surpassing distal margin; without spinules; without vestigial seta; with conical seta; without modified seta; without spinous process; without aesthetasc. Actual segment 9 with seta; two elements; of unequal size; one larger than segment; straight; surpassing to distal margin; beyond three sequential segments; without spinules; without vestigial seta; without conical seta; without modified seta; without spinous process; with aesthetasc; one element. Actual segment 10 with seta; one element; none larger than segment; straight; without spinules; without vestigial seta; without conical seta; without modified seta; without spinous process; without aesthetasc. Actual segment 11 with seta; one element; one larger than segment; straight; surpassing to distal margin; beyond three sequential segments; without spinules; without vestigial seta; without conical seta; without modified seta; without spinous process; without aesthetasc. Actual segment 12 with seta; one element; one larger than segment; straight; surpassing distal margin; without spinules; without vestigial seta; with conical seta; without modified seta; without spinous process; with aesthetasc; one element. Actual segment 13 with seta; one element; none elongated; straight; surpassing distal margin; without spinules; without vestigial seta; without conical seta; without modified seta; without spinous process; without aesthetasc. Actual segment 14 with seta; one element; elongated; straight; surpassing to distal margin; beyond three sequential segments; without spinules; without vestigial seta; without conical seta; without modified seta; without spinous process; with aesthetasc; one element. Actual segment 15 with seta; one element; larger than segment; straight; surpassing to distal margin; not beyond three sequential segments; without spinules; without vestigial seta; without conical seta; without modified seta; without spinous process; without aesthetasc. Actual segment 16 with seta; one element; larger than segment; plumose; surpassing to distal margin; not beyond three sequential segments; without spinules; without vestigial seta; without conical seta; without modified seta; without spinous process; with aesthetasc; one element. Actual segment 17 with seta; one element; not larger than segment; straight; without spinules; without vestigial seta; without conical seta; without modified seta; without spinous process; without aesthetasc. Actual segment 18 with seta; one element; larger than segment; straight; surpassing to distal margin; beyond three sequential segments; without spinules; without vestigial seta; without conical seta; without modified seta; without spinous process; without aesthetasc. Actual segment 19 with seta; one element; not larger than segment; straight; surpassing distal margin; without spinules; without vestigial seta; without conical seta; without modified seta; without spinous process; with aesthetasc; one element. Actual segment 20 with seta; one element; not larger than segment; straight; surpassing distal margin; without spinules; without vestigial seta; without conical seta; without modified seta; without spinous process; without aesthetasc. Actual segment 21 with seta; one element; larger than segment; plumose; surpassing to distal margin; beyond three sequential segments; without spinules; without vestigial seta; without conical seta; without modified seta; without spinous process; without aesthetasc. Actual segment 22 with seta; two elements; of unequal size; one of them elongated; plumose; surpassing to distal margin; without spinules; without vestigial seta; without conical seta; without modified seta; without spinous process; without aesthetasc. Actual segment 23 with seta; two elements; of unequal size; one larger than segment; plumose; surpassing to distal margin; greater 3x than original segment; without spinules; without vestigial seta; without conical seta; without modified seta; without spinous process; without aesthetasc. Actual segment 24 with seta; two elements; of equal size; one larger than segment; plumose; surpassing to distal margin; greater 3x than original segment; without spinules; without vestigial seta; without conical seta; without modified seta; without spinous process; without aesthetasc. Actual segment 25 with seta; four elements; of equal size; elongated; plumose; surpassing to distal margin; 4 times larger than segment; without spinules; without vestigial seta; without conical seta; without modified seta; without spinous process; with aesthetasc; one element.

##### Antenna

Biramous. Antenna coxa separated from the basis; bearing seta; 1; on inner surface; at distal corner; reaching to the middle basis. Antenna basis (fusion) separated from the endopodal segment; bearing seta; 2; on inner surface; at distal corner. Endopodal ancestral segment I and II separated. Ancestral segment II and III fused. Ancestral segment III and IV fused. Ancestral segment III and IV fully. Antenna endopod actual 2-segmented. Actual segment 1 not bilobate; with seta; two; on inner margin; with spinules; as a patch; not obliquely; on outer surface; with pore. Actual segment 2 bilobate; with discontinuity on outer cuticle; not developed as a suture; inner lobe bearing 8 setae; distally; outer lobe bearing 7 setae; distally; with spinules; as a patch; on outer surface. Antenna exopod ancestral segment I and II separated. Ancestral segment II and III fused. Ancestral segment III and IV fused. Ancestral segment IV and V separated. Ancestral segment V and VI separated. Ancestral segment VI and VII separated. Ancestral segment VII and VIII separated. Ancestral segment VIII and IX separated. Ancestral segment IX and X fused. Antenna exopod actual 7-segmented. Actual segment 1 single; elongated (width-length, equal or larger ratio 2:1); with seta; one; at inner surface. Actual segment 2 compound; elongated (larger width-length ratio 2:1); with seta; three; at inner surface. Actual segment 3 single; not elongated (lesser width-length ratio 2:1); with seta; one; at inner surface. Actual segment 4 single; not elongated (lesser width-length ratio 2:1); with seta; one; at inner surface. Actual segment 5 single; not elongated (lesser width-length ratio 2:1); with seta; one; at inner surface. Actual segment 6 single; not elongated (lesser width-length ratio 2:1); with seta; one; at inner surface. Actual segment 7 compound; elongated (larger or equal width-length ratio 2:1); with seta; one; at inner surface; and three; at distal surface.

##### Oral features

**Mandible**. Coxal gnathobase sclerotized; with lobe; prominent; on sub-caudal margin; presence of cutting blade; with tooth-like prominence; two, distinctly; 1 acute; on caudal margin; and 1 triangular; on sub-caudal margin; with acute projection between the prominences; with additional spinules; as a patch; on dorsal surface; with seta; dorsally; on apical surface; without spinules. Mandible palps biramous; comprising the basis; with seta; four; differently inserted; first medially; reaching to beyond the endopod 1; second distally; third distally; fourth distally; on inner margin; none with setulose ornamentation. Mandible endopod 2-segmented. Mandible endopod 1 with lobe; bearing seta; four; distally inserted; without spinules. Mandible endopod 2 without lobe; bearing setae; nine elements; distally inserted; with spinules; as a row; double. Mandible exopod 4-segmented. Mandible exopod 1 with seta; one element; distally; on inner margin. Mandible exopod 2 with seta; one element; distally; on inner side. Mandible exopod 3 with seta; one element; distally; on inner side. Mandible exopod 4 with setae; three elements; on terminal region. **Maxillule**. Birramous. Maxillule 3-segmented. Maxillule praecoxa with praecoxal arthrite; bearing spines; fifteen elements; ten marginally; plus, five sub-marginally; with spinules; as a patch; on sub-marginal surface. Maxillule coxa with coxal epipodite; with conspicuous outer lobe; bearing setae; nine elements; with coxal endite; elongated (larger or equal width-length ratio 2:1); bearing setae; four elements. Maxillule basis with basal endite; double; first proximal; elongated (larger width-length ratio 2:1; separated from basis; with setae; four elements; distally inserted; second distal; fused to basis; not elongated (lesser width-length ratio 2:1); with setae; four elements; distally inserted; with setules; as a row; on inner side; basal exite present; with setae; one element; on outer surface. Maxillule endopod 1-segmented. Endopod 1 bilobate; first proximal; with setae; three elements; second distal; with setae; five elements. Maxillule exopod 1-segmented. Exopod 1 with setae; six elements; with setules; as a row; on inner side; spinules absent. **Maxilla**. Uniramous. Maxilla 5-segmented. Maxilla praecoxa fused to coxa; incompletely; distinct externally; with praecoxal endite; double; first elongated endite (larger or equal width length ratio 2:1); proximally inserted; with seta; straight, or plumose; 3 straight; 3 plumose; with spine; single; with spinules; as a row; on distal margin; without setule; second non-elongated endite (lesser width length ratio 2:1); distally inserted; with seta; plumose; 3 plumose; without spine; with spinules; as a row; on distal margin; without setule; absence of outer seta. Maxilla coxa with coxal endite; double; first elongated endite (larger or equal width); proximally inserted; with seta; plumose; 3 plumose; without spine; without spinules; with setules; as a row; on distal margin; second elongated endite (larger or equal width); distally inserted; with seta; plumose; 3 plumose; without spine; without spinules; with setules; as a row; on distal margin; absence of outer seta. Maxilla basis with basal endite; single; elongated (larger or equal width-length ratio 2:1); with seta; plumose; 3 plumose; with spinules; absence of outer seta. Maxilla endopod 2-segmented. Endopod 1 with seta; 2 plumose; without spine; with spinules; as a patch; on distal margin; without setules. Maxilla endopod 2 with seta; 2 plumose; without spine; with spinules; as a patch; on distal margin; without setules. **Maxilliped**. Uniramous; Maxilliped 8-segmented. Maxilliped praecoxa fused to coxa; incompletely; distinct internally; with praecoxal endite; not elongated (lesser width-length ratio 2:1); distally inserted; with seta; 1 plumose; with spinules; as a row; single; on basal surface; without setules. Maxilliped coxa with coxal endite; three coxal endite; first not elongated (lesser width-length ratio 2:1); proximally inserted; with seta; 2 plumose; with spinules; as a patch; single; on medial surface; without setules; second elongated (larger or equal width); medially inserted; with seta; 3 plumose; with spinules; as a patch; single; on medial surface; without setules; third not elongated (lesser width-length ratio 2:1); distally inserted; with seta; 3 plumose; none reaching to beyond of the basis; with spinules; as a patch; single; on basal surface; with setules; as a patch; single; with lobe; prominence; at inner distal angle; ornamented; with spinules; continuously on margin. Maxilliped basis without basal endite; with seta; 3 straights; with spinules; as a row; single; on medial surface; with setules; as a row; single; on inner margin. Maxilliped endopod segment 6-segmented. Endopod 1 with seta; 2 plumose; on inner surface. Endopod 2 with seta; 3 plumose; on inner surface. Endopod 3 with seta; 2 plumose; on inner surface. Endopod 4 with seta; 2 plumose; on inner surface. Endopod 5 with seta; 2 plumose; on inner surface; outer seta absent. Endopod 6 with seta; 4 plumose; on inner surface, or on outer surface.

##### Swimming legs features

**First swimming legs.** Symmetrical; biramous. First swimming legs intercoxal plate without seta. First swimming legs praecoxa present; not laterally located. First swimming legs coxa with seta; one; straight; distally inserted; on inner surface; surpassing to first endopodal segment; with setules; one group; as a row; discontinuously; on inner margin; without spinules; without spine. First swimming legs basis without seta; with setules; as a row; single; continuously; on outer surface; without spinules; without spine. First swimming legs endopod 2-segmented. Endopod 1 without seta; without spine; without setules; without spinules; absence of Schmeil’s organ. Endopod 2 with seta; unrestricted; three on inner surface; one on outer surface; two on distal surface; straight; without spine; without setules; without spinules; absence of Schmeil’s organ. Endopod 3 absence. First swimming legs exopod 1 without seta; with spine; 1; stout; smaller than original segment; serrated; on inner side; continuously; with setules; as a row; single; as a row; innerly. First swimming legs exopod 2 with seta; restricted; 1 on inner surface; straight; without spine; without setules; without spinules. First swimming legs exopod 3 without setule; without spinules; with seta; unrestricted; 2 on inner surface; 2 on terminal surface; with spine; 2; unequal size; first no longer 2x than origin segment; stout; serrated; on inner side, or on outer side; equally; second longer 3x than origin segment; slender; serrated; on outer side; with ornamentation on non-serrated side; by setules. **Second swimming legs**. Symmetrical; Second swimming legs biramous. Second swimming legs intercoxal plate without seta. Second swimming legs praecoxa present; located laterally. Second swimming legs coxa with seta; straight; distally inserted; on inner surface; surpassing to basal segment; without setules; without spinules; without spine. Second swimming legs basis without seta; without setules; without spinules; without spine. Second swimming legs endopod 3-segmented. Endopod 1 with seta; straight; restricted; one on inner surface; without spine; without setules; without spinules; absence of Schmeil’s organ. Endopod 2 with seta; straight; restricted; one on inner surface; without spine; with setules; as a row; single; continuously; on inner side; without spinules; presence of Schmeil’s organ; on posterior surface. Endopod 3 with seta; straight; unrestricted; two on inner surface; two on outer surface; two on distal surface; without spine; without setules; with spinules; as a row; double; distally inserted; at anterior surface; absence of Schmeil’s organ. Second swimming legs exopod 1 with seta; restricted; one on inner surface; without spine; with setules; as a row; single; continuously; on inner side; without spinules; absence of Schmeil’s organ. Exopod 2 with seta; unrestricted; one on inner surface; with spine; 1; stout; not surpassing the exopod 3; serrated; on inner side, or on outer side; with setules; as a row; single; continuously; on inner surface; without spinules; absence of Schmeil’s organ. Exopod 3 with seta; plurimarginal; three on inner surface; two on terminal surface; with spine; 2; unequal size; first no longer 2x than origin segment; stout; serrated; on inner side, or on outer side; equally; second longer 2x than origin segment; slender; serrated; on outer side; with ornamentation on non-serrated side; of setules; no setules on outer surface; with spinules; as a row; double; distally inserted; at anterior surface; absence of Schmeil’s organ. **Third swimming legs**. Symmetrical; Third swimming legs biramous. Third swimming legs intercoxal plate without seta. Third swimming legs praecoxa present; not laterally located. Third swimming legs coxa with seta; straight; distally inserted; on inner surface; surpassing to basal segment; without setules; without spinules; without spine. Third swimming legs basis without seta; without setules; without spinules; without spine. Third swimming legs endopod 3-segmented. Endopod 1 with seta; restricted; one on inner surface; without spine; without setules; without spinules; absence of Schmeil’s organ. Endopod 2 with seta; restricted; two on inner surface; straight; without spine; without setules; without spinules; absence of Schmeil’s organ. Endopod 3 with seta; straight; plurimarginal; two on inner surface; two on outer surface; three on terminal surface; without spine; without setules; with spinules; as a row; distally inserted; single; at anterior surface; absence of Schmeil’s organ. Third swimming legs exopod 1 with seta; restricted; straight; one on inner surface; with spine; 1; stout; not reaching to the distal-third of the exopod 2; serrated; equally; on inner surface, or on outer surface; with setules; as a row; single; continuously; on inner surface; without spinules; absence of Schmeil’s organ. Exopod 2 with seta; straight; restricted; one on inner surface; with spine; 1; stout; not reaching out to exopod 3; serrated; on inner side, or on outer side; equally; without setules; without spinules; absence of Schmeil’s organ. Exopod 3 without setules; with spinules; as a row; single; distally inserted; at anterior surface; with seta; straight; unrestricted; three on inner surface; two on terminal surface; with spine; 2; unequal size; first no longer 2x than origin segment; stout; serrated; on inner side, or on outer side; equally; second longer 2x than origin segment; slender; serrated; on outer side; with ornamentation on non-serrated side; of setules; absence of Schmeil’s organ. **Fourth swimming legs**. Symmetrical; biramous. Intercoxal plate without sensilla. Praecoxa present. Coxa with seta; distally inserted; on inner margin; reaching out to endopod 1; without spinules; setules absent. Basis with seta; one; medially inserted; on posterior surface; smaller than the original segment; without setules; without spinules; without spine. Fourth swimming legs endopod 3-segmented. Endopod 1 without seta; without spine; without setules; without spinules; absence of Schmeil’s organ. Endopod 2 with seta; restricted; two on inner side; without spine; with setules; as a row; single; continuously; on outer surface; without spinules; absence of Schmeil’s organ. Endopod 3 with seta; unrestricted; two on inner surface; two on outer surface; three on distal surface; without spine; without setules; without spinules; absence of Schmeil’s organ. Fourth swimming legs exopod 1 with seta; restricted; one on inner surface; with spine; 1; stout; not reaching out to distal-third of the exopod 2; serrated; on inner side, or on outer side; equally; with setules; as a row; single; continuously; on inner surface; without spinules; absence of Schmeil’s organ. Exopod 2 with seta; restricted; one on inner surface; with spine; 1; stout; not reaching the end of exopod 3; serrated; on inner side, or on outer side; equally; with setules; as a row; single; continuously; on inner surface; without spinules; absence of Schmeil’s organ. Exopod 3 without setules; without spinules; with seta; unrestricted; three on inner surface; two on distal surface; with spine; 2; unequal size; first no longer 2x than origin segment; stout; serrated; on inner side, or on outer side; equally; second longer 2x than origin segment; slender; serrated; on outer side; with ornamentation on non-serrated side; of setules; absence of Schmeil’s organ.

##### Fifth swimming legs features

Asymmetrical. Fifth swimming leg intercoxal plate with length not equal or greater than width on 1.5x; with regular proximal margin; discontinuous to; the anterior margin of the left coxa, or the anterior margin of the right coxa; posterior sensilla on the right lateral absent. **Fifth left swimming leg**. Fifth left swimming leg biramous; leg reaching first right exopod segment; proximally. Fifth left swimming leg praecoxa present; rudimentary; separated from the coxae; without ornamentation. Fifth left swimming leg coxa concave inner side; without teeth-like structures; with process; conical; on posterior surface; outer side; distally inserted; not projecting over basis; with sensilla; slender; triangular; at apex; longer 2x than insertion basis; without swelling; without seta; without spinules. Fifth left swimming leg basis sub-cylindrical; unequal size between inner and outer side; shorter outer than inner side; with convex inner side; rounded internal proximal expansion absent; without outgrowth; without groove; absence of protuberance; with seta; outerly inserted; no longer 2x than origin segment; absence of minutely granular. Fifth left swimming leg endopod segments 1 and 2 fused; segments 2 and 3 fused; 1-segmented; stout; separated from the basis; ornamented; on inner side; with spinules; more than four elements; as a patch; terminally; row of setules absent; without seta. Fifth left swimming leg exopod segments 1 and 2 separated; segments 2 and 3 fused; 2-segmented; stout; separated from the basis. Fifth left swimming leg exopod 1 sub-cylindrical; longer than broad; unequal size between inner and outer side; shorter inner than outer side; concave inner side; concave outer side; without swelling; without marginal extension; without process; with lobe; single; circular; distally inserted; on inner side; covered; by setules; without outer spine; absence seta. Fifth left swimming leg exopod 2 sub-triangular; longer than broad; equal size between inner and outer side; disform inner side; with convex outer side; setulose pad present; prominently rounded; medially; on inner side; inflated medial region present; setulose; anteriorly; distal process present; acute-form; non denticulate; without transverse row of denticles; none oblique row of 5 denticles; innerly directed; with seta; spiniform; not ornamented by spinules; not surpassing the distal-point of the segment; without outer spine; terminal claw absent.

##### Fifth right swimming leg

Biramous. Fifth right swimming leg praecoxa absent. Fifth right swimming leg coxa concave inner side; without teeth-like structures; with process; conical; distally inserted; on posterior surface; closest to the outer rim; projecting over basis; not beyond the first third; without triangular protuberance innerly; with sensilla; stout; at apex; no longer 2x than basal insertion; without marginal extension; without seta; without spinules. Fifth right swimming leg basis cylindrical; unequal size between inner and outer side; shorter outer than inner side; rectilinear inner side; tumescence absent; without protuberance; absence of distinct minutely granular; additional inner process absent; with posterior groove; deep; longitudinally; not reaching the endopodal lobe; ornamented; with tubercles; throughout of the outer border; with seta; outerly inserted; on posterior surface; no longer 2x than origin segment; posterior protrusion present; distal process absent. Fifth right swimming leg with endopodite present; separated from the basis; on anterior surface; ancestral segments 1 and 2 fused; segments 2 and 3 fused; 1-segmented; stout; ornamented; with spinules; as a row; on inner side; sub-terminally; without seta. Fifth right swimming leg exopod segments 1 and 2 separated; segments 2 and 3 fused; 2-segmented; stout; separated from the basis. Fifth right swimming leg exopod 1 sub-cylindrical; longer than broad; nearly 2 times; unequal size between both sides; shorter inner than outer side; rectilinear inner side; rectilinear outer side; with marginal extension; sub-triangular; distally inserted; at outer rim; spinules absent; with process; triangular; rectilinear; blunt tip; sclerotized; without ornamentation; distally inserted; at marginal surface; projecting over next segment; without outer spine; without seta; internal prominence absent; lamella on posterior surface absent. Fifth right swimming leg exopod 2 elliptical; longer than broad; nearly 2.5 times; equal size between both sides; disform inner side; convex outer side; with posterior proximal swelling; inner-posterior process absent; without marginal expansion; curved ridge on distal posterior surface present; chitinous knobs present; with 3–6 posteriorly; with outer spine; inserted sub-distally; arched; internally directed; not ornamented innerly; not ornamented outerly; sharp tip; with apparent curve; innerly directed; lesser than the length of the exopod 2; beyond to 2 times its size; 3x; sensilla absent; terminal claw present; equal or longer 1.5 times than insertion segment; sclerotized; arched; inward; with conspicuous curve; proximally; ornamented innerly; by spinules; as a row; throughout extension; not ornamented outerly; sharp tip; curved tip; outwards; without medial constriction; hyaline process absent.

##### FEMALE

Body smaller and slenderer than male. Widest at first metasome segment. Distal margin of the prosomal segments without one line of setules at posterior margin. Prosome segments with spinules at least at one prosomal segment. Fourth metasome segment absence of dorsal protuberance. Fourth and fifth metasome segments fused; partially; on dorsal surface. Limit between fourth and fifth metasome segments ornamented; with spinules; as a row; single; complete; same size; entirely over limit (lateral, dorsal). **Fifth metasome segment**. Fifth metasome segment without sensilla; with epimeral plates. Epimeral plates asymmetrical. Right epimeral plates prominent, as projections; thinner than the left; one posterior-laterally directed; not reaching half length of the genital segment; with sensilla at the apex; dorsal-posterior sensilla present; slender; without ornamentation. Left epimeral plate without expansion.

##### Urosome

3-segmented. **Genital double-somite**. Asymmetrical in dorsal view; longer than broad; longer than other urosomites combined; dorsal suture at mid-length absent; not covered by spinules; with swelling; rounded; unequal size; greater right than left; anteriorly; with sensillae; on both sides; one; stout; with robust apex; at left lateral; not on lobular base; anteriorly; one; stout; at right lateral; not on lobular base; anteriorly; with robust apex; of unequal size between then; left bigger than right; lateral protuberance present; rounded; on the right side; with right posterior rim expanded; over next segment; without slender sensilla on each posterior rim; without posterior-dorsal process. Genital double-somite opercular pad present; broader than longer; symmetrical; development laterally; expanded posteriorly; covering partially; double gonoporal slit; located ventrally; with arthrodial membrane; inserted anteriorly; post-genital process absent; disto-ventral tumescence absent; ventral vertical folds absent; dorsal sensilla absent. Second urosome segment without ventral fusion to anal segment; right distal process absent. Caudal rami patch of setules on outer surface absent; patch of spinules on outer surface absent.

##### Appendices features

Rostrum basal process absent. **Antennules**. Symmetrical. Right antennule surpassing to genital double-segment; extending beyond caudal rami. Right antennule not exceeding the caudal setae. Right antennule ornamentation pattern equals to male left antennule; fully.

##### Fifth swimming legs

Symmetrical; Fifth swimming legs biramous. Fifth swimming legs intercoxal plate longer than wide; separated from the legs. Fifth swimming legs praecoxa with sclerite praecoxal; separated from the coxae; without ornamentation. Fifth swimming legs coxa with process; conical; at the outer rim; distally; sensilla present; stout; at apex; projecting over basal segment; no longer 2x than basal insertion; marginal extension absent; without swelling; without seta; without spinules. Fifth swimming legs basis sub-triangular; unequal size between inner and outer sides; shorter outer than inner side; with convex inner side; without proximal inner outgrowth; without groove; without distal extension; with seta; outerly inserted; on anterior surface; longer 2x than origin segment; reaching to exopod 1 distally. Fifth swimming legs endopod segments 1 and 2 fused; segments 2 and 3 fused; 1-segmented; stout; separated from the basis; absent discontinuity cuticle; with spinules; as a row; single; oblique; sub-terminally; at anterior surface; with seta; single; one distally; on posterior surface; rectilinear. Fifth swimming legs exopod segments 1 and 2 separated; segments 2 and 3 separated; 3-segmented; separated from the basis. Fifth swimming legs exopod 1 sub-cylindrical; longer than wide; longer or equal than 2 times; with unequal size between inner and outer side; shorter inner than outer side; with rectilinear inner side; with concave outer side; without swelling; without marginal extension; without posterior process; without spine; without seta. Fifth swimming legs exopod 2 sub-cylindrical; broader than long; no longer or equal than 2 times; without swelling; without marginal extension; without process; without lobe; with spine; inserted laterally; rectilinear; without ornamentation; sharp tip; equal size or larger than next segment; without seta. Fifth swimming legs exopod 3 cylindrical; longer than wide; without swelling; without process; without lobe; without spine; with seta; double; inserted terminally; unequal size between them; outer seta smaller than inner; nearly 3 times; outer seta not ornamented by setules; without ornamentation; presence of terminal claw; sclerotized; arched; internally directed; concave inner side; with ornamentation; of denticles; as a row; on surface partially; at medial region; rectilinear outer side; with ornamentation; of denticles; as a row; on surface partially; at medial region; blunt tip; 6 times longer than origin segment.

##### Distribution records

###### ARGENTINA

**Santa Fé**: Madrejón Don felipe, ox-bow Santa Fé (Paggi, 2001); middle Paraná River (Perbiche-Neves *et al*., 2015).

##### Habitat

Habitat in freshwaters: river.

##### Remarks

The species was described from individuals recognized as *N. anisitsi* previously. Paggi (2001) when addressing intraspecific morphological variability in *N. anisitsi*, recognized a distinct set of consistent attributes for lacustrine organisms from Santa Fe, Argentina, elevating them to the level of a new species. The attributes highlighted were, mainly: (1) male fifth right swimming leg exopod 2 with sclerotized processes posteriorly (treated here as chitinous knob), and (2) caudal rami with small basal “denticle” on outer margin of external seta (for this effort treated as outermost seta with outer spiniform process). Unfortunately, we did not have access to the type-material reported for the Museo Argentino de Ciencias Naturales ‘Bernardino Rivadavia’, which made it impossible to study the variability attested by Paggi (2001).

Perbiche-Neve *et al*. (2015), in a recent review of diaptomids from the Prata River Basin, recognized *N. dentatus* from organisms in the Paraná River between Argentina and Paraguay. The diagnostic characteristics reported in the original description were confirmed in this effort, and the observation for variation about the total male body length and female was added. In the present effort, we corroborate the original observations and review on the specimens of the Prata River, adding characteristics convergent with the type-species and related to their integration into the genus: (1) male fifth right swimming leg basis with posterior groove ornamented with tubercles on outer border; (2) male fifth right swimming leg exopod 1 longer than broad; (3) male fifth right swimming leg exopod 2 with curved ridge on posterior distal innerly; (4) female second urosome without ventral fusion to anal segment; (5) female fifth swimming leg basis with outer seta reaching to exopod 1 distally.

#### Notodiaptomus dubius Dussart & Matsumura-Tundisi in Dussart, 1986

##### Synonymy

*Notodiaptomus dubius* Dussart & Matsumura-Tundisi, 1986: 250, fig. 2; Matsumura-Tundisi, 1986: 537, 552, figs. 16–21, 100; Reid, 1987: 378; Defaye & Dussart, 1988: 114; Sendacz, 1993: 35; Rocha *et al*., 1995: 156, 159; Santos-Silva, 1998: 209; Santos-Silva, 2008: 26–27; Perbiche-Neves *et al*., 2020: 672, 681, 682, key to the Neotropical diaptomid, fig. 21.8 E. *Notodiaptomus* (*Wrightius*) *dubius*; Dussart 1985a: 210, 212, 213, 214, fig. 8.

##### Type locality

Amarelo Lake: Doce River valley, Minas Gerais State, Brazil.

##### Type material

Holotype: 1 male, dissected on the slide (MZUSP 6965). Paratypes: 1 female (MZUSP 6967), and 1 male (MZUSP 6966), both dissected on the slide. Collected on 20.VII.1978, no information on the collector or mounting medium of the slides. All material was stored in the Zoology Museum of São Paulo - MZUSP.

##### Diagnosis

**(1)** male fifth right swimming leg coxa with conical process projecting on basis until distal surface posteriorly; **(2)** right swimming leg endopod with fused to basis; **(3)** male fifth right swimming leg exopod 1 broader than long; **(5)** male fifth right swimming leg exopod 2 with tip outer spine directed outerly; **(5)** male fifth right swimming legs exopod 2 with terminal claw ornamented outerly by spinules, and sharp tip; **(6)** female fourth and fifth metasome segments with fusion on lateral surface; **(7)** female fourth metasome segment with rounded dorsal protuberance distally; **(8)** female antennules not surpassing to genital double-somite.

##### Material examined

Holotype: male (MZUSP 6965). Paratype: 1 female (MZUSP 6967), and 1 male (MZUSP 6966). All specimens preserved entire, in alcohol, and stored in MZUSP.

##### Redescription

###### MALE

Body 1089 micrometers excluding caudal setae. Male body smaller and slenderer than female. Nerve axons myelinated. Prosome 6-segmented; widest at first metasome segment; without one line of setules at posterior margin; without spinules at segments. Cephalosome anterior margin rounded; with dorsal suture; incomplete; separate from first metasome segment. First metasome segment without sensilla. Second metasome segment with sensilla; 2 dorsally; of equal size. Third metasome segment with sensillae; 4 laterally; of unequal size; non-ornamented posterior margin. Fourth metasome segment with sensillae; 2 dorsally; 4 laterally; of equal size; fused to fifth metasome; fourth metasome segment partially; fourth metasome segment on lateral surface. Limit between fourth and fifth metasome segments without ornamentation. Fifth metasome segment with sensilla; 4 laterally; Fifth metasome segment equal size; Fifth metasome segment without ornamentation; Fifth metasome segment without dorsal conical process; with epimeral plates. Epimeral plates symmetrical. Right epimeral plates reduced, as rounded distal corner segment limit; with sensilla; at the apex of projection; without ornamentation.

##### Urosome

5-segmented; Urosome 5 - free segments. Genital somite symmetrical in dorsal view; with single aperture; located on left side; ventrolaterally on posterior rim; with sensillae; on both sides; one; at left lateral; posteriorly; one; at right rim; posteriorly; of equal size between then. Third urosome segment without spinules; without external seta. Fourth urosome segment without spinules; without sub-conical blunt dorsal-lateral process. Anal segment presence of dorsal sensillae; one on each side; medially inserted; presence of operculum; convex; covering the anal aperture fully. Caudal rami symmetrical; separated from anal segment; longer than wide; with setules; continuous on; inner side; each ramus bearing 6 caudal setae; 5 marginals; plumose; and 1 internal dorsally; straight; not reticulated main axis; outermost seta with outer spiniform process absent.

##### Appendices features

Rostrum symmetrical; separated from dorsal cephalic shield; by complete suture; sensillae present; one pair; anteriorly inserted on surface tegument; with rostral filament; double; paired; extended; into point; with basal process; in ventral view, rounded on left side; without a smaller basal expansion on the right side.

##### Antennules

Asymmetrical. **Right antennules**. Uniramous; right antennule not surpassing to genital segment.

Right antennule ancestral segment I and II separated. Ancestral segment II and III fused. Ancestral segment III and IV fused. Ancestral segment IV and V separated. Ancestral segment V and VI separated. Ancestral segment VI and VII separated. Ancestral segment VII and VIII separated. Ancestral segment VIII and IX separated. Ancestral segment IX and X separated. Ancestral segment X and XI separated. Ancestral segment XI and XII separated. Ancestral segment XII and XIII separated. Ancestral segment XIII and XIV separated. Ancestral segment XIV and XV separated. Ancestral segment XV and XVI separated. Ancestral segment XVI and XVII separated. Ancestral segment XVII and XVIII separated. Ancestral segment XVIII and XIX separated. Ancestral segment XIX and XX separated. Ancestral segment XX and XXI separated. Ancestral segment XXI and XXII fused. Ancestral segment XXII and XXIII fused. Ancestral segment XXIII and XXIV separated. Ancestral segment XXIV and XXV fused. Ancestral segment XXV and XXVI separated. Ancestral segment XXVI and XXVII separated. Ancestral segment XXVII and XXVIII fused.

Right antennule actual 22-segmented; geniculated; between the segment 18 and segment 19; with swollen and modified region; formed by 5 segments; between 13 and 17 segments. Actual segment 1 with seta; one element; straight; none larger than segment; without spinules; without vestigial seta; without conical seta; without modified seta; without spinous process; with aesthetasc; one element. Actual segment 2 with seta; three elements; of unequal size; straight; none larger than segment; without spinules; with vestigial seta; one element; without conical seta; without modified seta; without spinous process; with aesthetasc; one element. Actual segment 3 with seta; one element; one larger than segment; surpassing to distal margin; beyond three sequential segments; straight; blunt apex; without spinules; with vestigial seta; one element; without conical seta; without modified seta; without spinous process; with aesthetasc. Actual segment 4 with seta; one element; one larger than segment; surpassing to distal margin; straight; not beyond three sequential segments; without spinules; without vestigial seta; without conical seta; without modified seta; without spinous process; without aesthetasc. Actual segment 5 with seta; one element; straight; one larger than segment; surpassing to distal margin; not beyond three sequential segments; without spinules; with vestigial seta; one element; without conical seta; without modified seta; without spinous process; with aesthetasc; one element. Actual segment 6 with seta; one element; none larger than segment; straight; without spinules; without vestigial seta; without conical seta; without modified seta; without spinous process; without aesthetasc. Actual segment 7 with seta; one element; straight; one larger than segment; surpassing to distal margin; beyond three sequential segments; blunt apex; without spinules; without vestigial seta; without conical seta; without modified seta; without spinous process; with aesthetasc; one element. Actual segment 8 with seta; one element; straight; none larger than segment; without spinules; without vestigial seta; with conical seta; one element; not reaching to middle-point of the sequent segment; without modified seta; without spinous process; without aesthetasc. Actual segment 9 with seta; two elements; of unequal size; straight; one larger than segment; surpassing to distal margin; beyond three sequential segments; blunt apex; without spinules; without vestigial seta; without conical seta; without modified seta; without spinous process; with aesthetasc; one element. Actual segment 10 with seta; one element; straight; none larger than segment; without spinules; without vestigial seta; without conical seta; with modified seta; presenting blunt apex; slender form; surpassing distal margin; parallel to antennule direction; without spinous process; without aesthetasc. Actual segment 11 with seta; one element; straight; one larger than segment; surpassing to distal margin; not beyond three sequential segments; without spinules; without vestigial seta; without conical seta; with modified seta; slender form; presenting blunt apex; surpassing to distal margin; not beyond of the sequential segment; perpendicular to antennule direction; shorter length than homologous of actual segment 13; without spinous process; without aesthetasc. Actual segment 12 with seta; one element; straight; one larger than segment; surpassing to distal margin; not beyond three sequential segments; without spinules; without vestigial seta; with conical seta; one element; not smaller than to segment 8; without modified seta; with aesthetasc; one element; absent internal perpendicular fission. Actual segment 13 with seta; one element; straight; one larger than segment; surpassing to distal margin; not beyond three sequential segments; without spinules; without vestigial seta; without conical seta; with modified seta; stout form; surpassing to distal margin; to the distal-point of the sequence segment; perpendicular to antennule direction; presenting bifid apex; without spinous process; with aesthetasc; one element. Actual segment 14 with seta; two elements; of unequal size; straight; one larger than segment; surpassing to distal margin; beyond three sequential segments; blunt apex; without spinules; without vestigial seta; without conical seta; without modified seta; without spinous process; with aesthetasc; one element. Actual segment 15 with seta; two elements; of unequal size; straight; not bifidform; none larger than segment; without spinules; without vestigial seta; without conical seta; without modified seta; with spinous process; on outer margin; surpassing distal margin; with aesthetasc; one element. Actual segment 16 with seta; two elements; of unequal size; plumose; one larger than segment; surpassing to distal margin; not beyond three sequential segments; not bifidform; without spinules; without vestigial seta; without conical seta; without modified seta; with spinous process; on outer margin; surpassing distal margin; unequal size to process on preceding segment; with aesthetasc; one element. Actual segment 17 with seta; two elements; of unequal size; straight; none larger than segment; bifidform; without spinules; without vestigial seta; without conical seta; with modified seta; one element; stout form; surpassing to distal margin; not beyond of the sequential segment; parallel to antennule direction; without spinous process; without aesthetasc. Actual segment 18 with seta; two elements; of equal size; straight; none larger than segment; without spinules; without vestigial seta; without conical seta; with modified seta; one element; stout form; surpassing distal margin; parallel to antennule direction; without spinous process; without aesthetasc. Actual segment 19 with seta; two elements; of unequal size; plumose; none larger than segment; without spinules; without vestigial seta; without conical seta; with modified seta; two elements; stout form; at least one bifid form; surpassing distal margin; parallel to antennule direction; without spinous process; with aesthetasc; one element. Actual segment 20 with seta; four elements; of unequal size; straight; one larger than segment; surpassing to distal margin; beyond three sequential segments; without spinules; without vestigial seta; without conical seta; without modified seta; without spinous process; without aesthetasc. Actual segment 21 with seta; two elements; of equal size; plumose; one larger than segment; surpassing to distal margin; greater 3x than original segment; without spinules; without vestigial seta; without conical seta; without modified seta; without spinous process; without aesthetasc. Actual segment 22 with seta; four elements; of equal size; one larger than segment; plumose; surpassing to distal margin; greater 3x than original segment; without spinules; without vestigial seta; without conical seta; without modified seta; without spinous process; with aesthetasc; one element.

##### Left antennules

Uniramous; Left antennule not surpassing to prosome. Ancestral segment I and II separated. Ancestral segment II and III fused. Ancestral segment III and IV fused. Ancestral segment IV and V separated. Ancestral segment V and VI separated. Ancestral segment VI and VII separated. Ancestral segment VII and VIII separated. Ancestral segment VIII and IX separated. Ancestral segment IX and X separated. Ancestral segment X and XI separated. Ancestral segment XI and XII separated. Ancestral segment XII and XIII separated. Ancestral segment XIII and XIV separated. Ancestral segment XIV and XV separated. Ancestral segment XV and XVI separated. Ancestral segment XVI and XVII separated. Ancestral segment XVII and XVIII separated. Ancestral segment XVIII and XIX separated. Ancestral segment XIX and XX separated. Ancestral segment XX and XXI separated. Ancestral segment XXI and XXII separated. Ancestral segment XXII and XXIII separated. Ancestral segment XXIII and XXIV separated. Ancestral segment XXIV and XXV separated. Ancestral segment XXV and XXVI separated. Ancestral segment XXVI and XXVII separated. Ancestral segment XXVII and XXVIII fused.

Left antennule actual 25-segmented; not-geniculated. Actual segment 1 with seta; one element; none larger than segment; straight; without spinules; without vestigial seta; without conical seta; without modified seta; without spinous process; with aesthetasc; one element. Actual segment 2 with seta; three elements; of equal size; none larger than segment; straight; without spinules; with vestigial seta; one element; without conical seta; without modified seta; without spinous process; with aesthetasc; one element. Actual segment 3 with seta; one element; one larger than segment; straight; surpassing to distal margin; beyond three sequential segments; without spinules; with vestigial seta; one element; without conical seta; without modified seta; without spinous process; with aesthetasc. Actual segment 4 with seta; one element; none larger than segment; straight; without spinules; without vestigial seta; without conical seta; without modified seta; without spinous process; without aesthetasc. Actual segment 5 with seta; one element; one larger than segment; straight; surpassing to distal margin; not beyond three sequential segments; without spinules; with vestigial seta; one element; without conical seta; without modified seta; without spinous process; with aesthetasc; one element. Actual segment 6 with seta; one element; none larger than segment; straight; without spinules; without vestigial seta; without conical seta; without modified seta; without spinous process; without aesthetasc. Actual segment 7 with seta; one element; one larger than segment; straight; surpassing to distal margin; beyond three sequential segments; without spinules; without vestigial seta; without conical seta; without modified seta; without spinous process; with aesthetasc; one element. Actual segment 8 with seta; one element; one larger than segment; straight; surpassing distal margin; without spinules; without vestigial seta; with conical seta; without modified seta; without spinous process; without aesthetasc. Actual segment 9 with seta; two elements; of unequal size; one larger than segment; straight; surpassing to distal margin; beyond three sequential segments; without spinules; without vestigial seta; without conical seta; without modified seta; without spinous process; with aesthetasc; one element. Actual segment 10 with seta; one element; none larger than segment; straight; without spinules; without vestigial seta; without conical seta; without modified seta; without spinous process; without aesthetasc. Actual segment 11 with seta; one element; one larger than segment; straight; surpassing to distal margin; beyond three sequential segments; without spinules; without vestigial seta; without conical seta; without modified seta; without spinous process; without aesthetasc. Actual segment 12 with seta; one element; one larger than segment; straight; surpassing distal margin; without spinules; without vestigial seta; with conical seta; without modified seta; without spinous process; with aesthetasc; one element. Actual segment 13 with seta; one element; none elongated; straight; surpassing distal margin; without spinules; without vestigial seta; without conical seta; without modified seta; without spinous process; without aesthetasc. Actual segment 14 with seta; one element; elongated; straight; surpassing to distal margin; beyond three sequential segments; without spinules; without vestigial seta; without conical seta; without modified seta; without spinous process; with aesthetasc; one element. Actual segment 15 with seta; one element; larger than segment; straight; surpassing to distal margin; not beyond three sequential segments; without spinules; without vestigial seta; without conical seta; without modified seta; without spinous process; without aesthetasc. Actual segment 16 with seta; one element; larger than segment; plumose; surpassing to distal margin; not beyond three sequential segments; without spinules; without vestigial seta; without conical seta; without modified seta; without spinous process; with aesthetasc; one element. Actual segment 17 with seta; one element; not larger than segment; straight; without spinules; without vestigial seta; without conical seta; without modified seta; without spinous process; without aesthetasc. Actual segment 18 with seta; one element; larger than segment; straight; surpassing to distal margin; beyond three sequential segments; without spinules; without vestigial seta; without conical seta; without modified seta; without spinous process; without aesthetasc. Actual segment 19 with seta; one element; not larger than segment; straight; surpassing distal margin; without spinules; without vestigial seta; without conical seta; without modified seta; without spinous process; with aesthetasc; one element. Actual segment 20 with seta; one element; not larger than segment; straight; surpassing distal margin; without spinules; without vestigial seta; without conical seta; without modified seta; without spinous process; without aesthetasc. Actual segment 21 with seta; one element; larger than segment; plumose; surpassing to distal margin; beyond three sequential segments; without spinules; without vestigial seta; without conical seta; without modified seta; without spinous process; without aesthetasc. Actual segment 22 with seta; two elements; of unequal size; one of them elongated; plumose; surpassing to distal margin; without spinules; without vestigial seta; without conical seta; without modified seta; without spinous process; without aesthetasc. Actual segment 23 with seta; two elements; of unequal size; one larger than segment; plumose; surpassing to distal margin; greater 3x than original segment; without spinules; without vestigial seta; without conical seta; without modified seta; without spinous process; without aesthetasc. Actual segment 24 with seta; two elements; of equal size; one larger than segment; plumose; surpassing to distal margin; greater 3x than original segment; without spinules; without vestigial seta; without conical seta; without modified seta; without spinous process; without aesthetasc. Actual segment 25 with seta; four elements; of equal size; elongated; plumose; surpassing to distal margin; 4 times larger than segment; without spinules; without vestigial seta; without conical seta; without modified seta; without spinous process; with aesthetasc; one element.

##### Antenna

Biramous. Antenna coxa separated from the basis; bearing seta; 1; on inner surface; at distal corner; reaching to the endopod 1. Antenna basis (fusion) separated from the endopodal segment; bearing seta; 2; on inner surface; at distal corner. Endopodal ancestral segment I and II separated. Ancestral segment II and III fused. Ancestral segment III and IV fused. Ancestral segment III and IV fully. Antenna endopod actual 2-segmented. Actual segment 1 not bilobate; with seta; two; on inner margin; with spinules; as a row; obliquely; on outer surface; with pore. Actual segment 2 bilobate; with discontinuity on outer cuticle; not developed as a suture; inner lobe bearing 8 setae; distally; outer lobe bearing 7 setae; distally; with spinules; as a patch; on outer surface. Antenna exopod ancestral segment I and II separated. Ancestral segment II and III fused. Ancestral segment III and IV fused. Ancestral segment IV and V separated. Ancestral segment V and VI separated. Ancestral segment VI and VII separated. Ancestral segment VII and VIII separated. Ancestral segment VIII and IX separated. Ancestral segment IX and X fused. Antenna exopod actual 7-segmented. Actual segment 1 single; elongated (width-length, equal or larger ratio 2:1); with seta; one; at inner surface. Actual segment 2 compound; elongated (larger width-length ratio 2:1); with seta; three; at inner surface. Actual segment 3 single; not elongated (lesser width-length ratio 2:1); with seta; one; at inner surface. Actual segment 4 single; not elongated (lesser width-length ratio 2:1); with seta; one; at inner surface. Actual segment 5 single; not elongated (lesser width-length ratio 2:1); with seta; one; at inner surface. Actual segment 6 single; not elongated (lesser width-length ratio 2:1); with seta; one; at inner surface. Actual segment 7 compound; elongated (larger or equal width-length ratio 2:1); with seta; one; at inner surface; and three; at distal surface.

##### Oral features

**Mandible**. Coxal gnathobase sclerotized; with lobe; prominent; on caudal margin; presence of cutting blade; with tooth-like prominence; two, distinctly; 1 acute; on caudal margin; and 1 triangular; on sub-caudal margin; without acute projection between the prominences; with additional spinules; as a row; on dorsal surface; with seta; 1; dorsally; on apical surface; with spinules; apicalmost. Mandible palps biramous; comprising the basis; with seta; four; differently inserted; first medially; reaching to beyond the endopod 1; second distally; third distally; fourth distally; on inner margin; none with setulose ornamentation. Mandible endopod 2-segmented. Mandible endopod 1 with lobe; bearing seta; four; distally inserted; without spinules. Mandible endopod 2 without lobe; bearing setae; nine elements; distally inserted; with spinules; as a row; double. Mandible exopod 4-segmented. Mandible exopod 1 with seta; one element; distally; on inner margin. Mandible exopod 2 with seta; one element; distally; on inner side. Mandible exopod 3 with seta; one element; distally; on inner side. Mandible exopod 4 with setae; three elements; on terminal region. **Maxillule**. Birramous. Maxillule 3-segmented. Maxillule praecoxa with praecoxal arthrite; bearing spines; fifteen elements; ten marginally; plus five sub-marginally; with spinules; as a patch; on sub-marginal surface. Maxillule coxa with coxal epipodite; with conspicuous outer lobe; bearing setae; nine elements; with coxal endite; elongated (larger or equal width-length ratio 2:1); bearing setae; four elements. Maxillule basis with basal endite; double; first proximal; elongated (larger width-length ratio 2:1; separated from basis; with setae; four elements; distally inserted; second distal; fused to basis; not elongated (lesser width-length ratio 2:1); with setae; four elements; distally inserted; with setules; as a row; on inner side; basal exite present; with setae; one element; on outer surface. Maxillule endopod 1-segmented. Endopod 1 bilobate; first proximal; with setae; three elements; second distal; with setae; five elements. Maxillule exopod 1-segmented. Exopod 1 with setae; six elements; with setules; as a row; on inner side; spinules absent. **Maxilla**. Uniramous. Maxilla 5-segmented. Maxilla praecoxa fused to coxa; incompletely; distinct externally; with praecoxal endite; double; first elongated endite (larger or equal width length ratio 2:1); proximally inserted; with seta; straight, or plumose; 1 straight; 4 plumose; with spine; single; without spinules; without setule; second elongated endite (larger or equal width length ratio 2:1); distally inserted; with seta; plumose; 3 plumose; without spine; with spinules; as a row; on distal margin; with setule; as a row; on distal margin; absence of outer seta. Maxilla coxa with coxal endite; double; first elongated endite (larger or equal width); proximally inserted; with seta; plumose; 3 plumose; without spine; without spinules; with setules; as a row; on proximal margin; second elongated endite (larger or equal width); distally inserted; with seta; plumose; 3 plumose; without spine; without spinules; with setules; as a row; on proximal margin; absence of outer seta. Maxilla basis with basal endite; single; elongated (larger or equal width-length ratio 2:1); with seta; plumose; 3 plumose; without spinules; absence of outer seta. Maxilla endopod 2-segmented. Endopod 1 with seta; 2 plumose; without spine; without spinules; without setules. Maxilla endopod 2 with seta; 2 plumose; without spine; without spinules; without setules. **Maxilliped**. Uniramous; Maxilliped 8-segmented. Maxilliped praecoxa fused to coxa; incompletely; distinct internally; with praecoxal endite; not elongated (lesser width-length ratio 2:1); distally inserted; with seta; 1 straight; with spinules; as a row; single; on basal surface; without setules. Maxilliped coxa with coxal endite; three coxal endite; first elongated (larger or equal width); proximally inserted; with seta; 2 plumose; with spinules; as a patch; single; on apical surface; without setules; second not elongated (lesser width-length ratio 2:1); medially inserted; with seta; 3 plumose; with spinules; as a row; single; on medial surface; without setules; third elongated (larger or equal width length ratio 2:1); distally inserted; with seta; 3 plumose; none reaching to beyond of the basis; with spinules; as a row; single; on basal surface; without setules; with lobe; prominence; at inner distal angle; ornamented; with spinules; continuously on margin. Maxilliped basis without basal endite; with seta; 3 plumose; with spinules; as a row; single; on medial surface; with setules; as a row; single; on inner margin. Maxilliped endopod segment 6-segmented. Endopod 1 with seta; 2 plumose; on inner surface. Endopod 2 with seta; 3 plumose; on inner surface. Endopod 3 with seta; 2 plumose; on inner surface. Endopod 4 with seta; 2 plumose; on inner surface. Endopod 5 with seta; 2 plumose; on inner surface, or on outer surface; outer seta absent. Endopod 6 with seta; 4 plumose; on inner surface, or on outer surface.

##### Swimming legs features

**First swimming legs.** Symmetrical; biramous. First swimming legs intercoxal plate without seta. First swimming legs praecoxa absent. First swimming legs coxa with seta; one; straight; distally inserted; on inner surface; surpassing to basal segment; with setules; two group; as a patch; on inner margin; and as a row; double; on anterior surface; outerly; without spinules; without spine. First swimming legs basis without seta; with setules; as a patch; single; on outer surface; without spinules; without spine. First swimming legs endopod 2-segmented. Endopod 1 with seta; straight; restricted; to inner surface; one element; without spine; with setules; as a row; single; continuously; on outer surface; without spinules; absence of Schmeil’s organ. Endopod 2 with seta; unrestricted; three on inner surface; one on outer surface; two on distal surface; straight; without spine; with setules; as a row; single; continuously; on outer surface; without spinules; absence of Schmeil’s organ. Endopod 3 absence. First swimming legs exopod 1 with seta; restricted; 1 on inner surface; with spine; 1; stout; smaller than original segment; serrated; on inner side; continuously; without setules. First swimming legs exopod 2 with seta; restricted; 1 on inner surface; straight; without spine; with setules; as a row; single; continuously; on inner margin, or on outer margin; without spinules. First swimming legs exopod 3 with setule; as a row; single; continuously; on outer surface; without spinules; with seta; unrestricted; 2 on inner surface; 2 on terminal surface; with spine; 2; unequal size; first no longer 2x than origin segment; stout; serrated; on inner side, or on outer side; equally; second longer 3x than origin segment; slender; serrated; on outer side; with ornamentation on non-serrated side; by setules. **Second swimming legs**. Symmetrical; Second swimming legs biramous. Second swimming legs intercoxal plate without seta. Second swimming legs praecoxa present; located laterally. Second swimming legs coxa with seta; straight; distally inserted; on inner surface; surpassing to basal segment; without setules; without spinules; without spine. Second swimming legs basis without seta; without setules; without spinules; without spine. Second swimming legs endopod 3-segmented. Endopod 1 with seta; straight; restricted; one on inner surface; without spine; with setules; as a row; single; continuously; on outer surface; without spinules; absence of Schmeil’s organ. Endopod 2 with seta; straight; unrestricted; two on inner surface; without spine; with setules; as a row; single; continuously; on outer side; without spinules; presence of Schmeil’s organ; on posterior surface. Endopod 3 with seta; straight; unrestricted; three on inner surface; two on outer surface; two on distal surface; without spine; without setules; with spinules; as a row; double; distally inserted; at anterior surface; absence of Schmeil’s organ. Second swimming legs exopod 1 with seta; restricted; one on inner surface; with spine; 1; stout; not reaching to distal-third of the exopod 2; serrated; on inner side, or on outer side; with setules; as a row; single; continuously; on inner side; without spinules; absence of Schmeil’s organ. Exopod 2 with seta; unrestricted; one on inner surface; with spine; 1; stout; not surpassing the exopod 3; serrated; on inner side, or on outer side; with setules; as a row; single; continuously; on inner surface; without spinules; absence of Schmeil’s organ. Exopod 3 with seta; plurimarginal; three on inner surface; two on terminal surface; with spine; 2; unequal size; first no longer 2x than origin segment; stout; serrated; on inner side, or on outer side; equally; second longer 2x than origin segment; slender; serrated; on outer side; with ornamentation on non-serrated side; of setules; setules on outer surface; as a row; single; continuously; on inner surface; with spinules; as a row; single; distally inserted; at anterior surface; absence of Schmeil’s organ. **Third swimming legs**. Symmetrical; Third swimming legs biramous. Third swimming legs intercoxal plate without seta. Third swimming legs praecoxa present; not laterally located. Third swimming legs coxa with seta; straight; distally inserted; on inner surface; surpassing to basal segment; without setules; without spinules; without spine. Third swimming legs basis without seta; without setules; without spinules; without spine. Third swimming legs endopod 3-segmented. Endopod 1 with seta; restricted; one on inner surface; without spine; without setules; without spinules; absence of Schmeil’s organ. Endopod 2 with seta; restricted; two on inner surface; straight; without spine; without setules; without spinules; absence of Schmeil’s organ. Endopod 3 with seta; straight; plurimarginal; two on inner surface; two on outer surface; three on terminal surface; without spine; without setules; with spinules; as a row; distally inserted; double; at anterior surface; absence of Schmeil’s organ. Third swimming legs exopod 1 with seta; restricted; straight; one on inner surface; with spine; 1; stout; not reaching to the distal-third of the exopod 2; serrated; equally; on inner surface, or on outer surface; with setules; as a row; single; continuously; on inner surface; without spinules; absence of Schmeil’s organ. Exopod 2 with seta; straight; restricted; one on inner surface; with spine; 1; stout; not reaching out to exopod 3; serrated; on inner side, or on outer side; equally; with setules; as a row; single; continuously; on inner side; without spinules; absence of Schmeil’s organ. Exopod 3 without setules; with spinules; as a row; single; distally inserted; at anterior surface; with seta; straight; unrestricted; three on inner surface; two on terminal surface; with spine; 2; unequal size; first no longer 2x than origin segment; stout; serrated; on inner side, or on outer side; equally; second longer 2x than origin segment; slender; serrated; on outer side; with ornamentation on non-serrated side; of setules; absence of Schmeil’s organ. **Fourth swimming legs**. Symmetrical; biramous. Intercoxal plate without sensilla. Praecoxa present. Coxa with seta; distally inserted; on inner margin; reaching out to endopod 1; without spinules; setules absent. Basis with seta; one; medially inserted; on posterior surface; smaller than the original segment; without setules; without spinules; without spine. Fourth swimming legs endopod 3-segmented. Endopod 1 with seta; one; restricted; on inner surface; without spine; without setules; without spinules; absence of Schmeil’s organ. Endopod 2 with seta; restricted; two on inner side; without spine; with setules; as a row; single; continuously; on outer surface; without spinules; absence of Schmeil’s organ. Endopod 3 with seta; unrestricted; two on inner surface; two on outer surface; three on distal surface; without spine; without setules; with spinules; as a row; double; distally inserted; at anterior surface; absence of Schmeil’s organ. Fourth swimming legs exopod 1 with seta; restricted; one on inner surface; with spine; 1; stout; not reaching out to distal-third of the exopod 2; serrated; on inner side, or on outer side; equally; with setules; as a row; single; continuously; on inner surface; without spinules; absence of Schmeil’s organ. Exopod 2 with seta; restricted; one on inner surface; with spine; 1; stout; not reaching the end of exopod 3; serrated; on inner side, or on outer side; equally; with setules; as a row; single; continuously; on inner surface; without spinules; absence of Schmeil’s organ. Exopod 3 without setules; with spinules; as a row; single; distally inserted; at anterior surface; with seta; unrestricted; three on inner surface; two on distal surface; with spine; 2; unequal size; first no longer 2x than origin segment; stout; serrated; on inner side, or on outer side; equally; second longer 2x than origin segment; slender; serrated; on outer side; without ornamentation on non-serrated side; absence of Schmeil’s organ.

##### Fifth swimming legs features

Asymmetrical. Fifth swimming leg intercoxal plate with length not equal or greater than width on 1.5x; with irregular proximal margin; discontinuous to; the anterior margin of the left coxa, or the anterior margin of the right coxa; posterior sensilla on the right lateral absent. **Fifth left swimming leg**. Fifth left swimming leg biramous; leg reaching first right exopod segment; proximally. Fifth left swimming leg praecoxa present; rudimentary; separated from the coxae; without ornamentation. Fifth left swimming leg coxa concave inner side; without teeth-like structures; with process; conical; on posterior surface; outer side; distally inserted; not projecting over basis; with sensilla; stout; triangular; at apex; longer 2x than insertion basis; without swelling; without seta; without spinules. Fifth left swimming leg basis sub-cylindrical; unequal size between inner and outer side; shorter outer than inner side; with concave inner side; rounded internal proximal expansion absent; without outgrowth; with groove; deep; obliquely; on posterior surface; not reaching the endopodal lobe; not ornamented; absence of protuberance; with seta; outerly inserted; no longer 2x than origin segment. Fifth left swimming leg endopod segments 1 and 2 fused; segments 2 and 3 fused; 1-segmented; stout; separated from the basis; ornamented; on inner side; with spinules; more than four elements; as a row; terminally; row of setules absent; without seta. Fifth left swimming leg exopod segments 1 and 2 separated; segments 2 and 3 fused; 2-segmented; stout; separated from the basis. Fifth left swimming leg exopod 1 sub-triangular; longer than broad; equal size between inner and outer side; rectilinear inner side; convex outer side; without swelling; without marginal extension; without process; with lobe; double; semicircular; medially inserted; on inner side; covered; by setules; without outer spine; absence seta. Fifth left swimming leg exopod 2 digitiform; longer than broad; equal size between inner and outer side; disform inner side; with rectilinear outer side; setulose pad present; not prominently rounded; proximally; on inner side; inflated medial region absent; distal process present; digitiform; denticulate; not bicuspidate; without transverse row of denticles; none oblique row of 5 denticles; at anterior surface; innerly directed; with seta; spiniform; not ornamented by spinules; not surpassing the distal-point of the segment; without outer spine; terminal claw absent.

##### Fifth right swimming leg

Biramous. Fifth right swimming leg praecoxa present; separated from the coxae; without ornamentation. Fifth right swimming leg coxa convex inner side; without teeth-like structures; with process; conical; distally inserted; on posterior surface; closest to the outer rim; projecting over basis; beyond the first third; until the distal surface; without triangular protuberance innerly; with sensilla; slender; at nadir; no longer 2x than basal insertion; without marginal extension; without seta; without spinules. Fifth right swimming leg basis trapezoidal; unequal size between inner and outer side; shorter outer than inner side; concave inner side; tumescence present; inflated; not bilobed; unrestricted on inner surface; without protuberance; absence of distinct minutely granular; additional inner process absent; without posterior groove; with seta; outerly inserted; on anterior surface; no longer 2x than origin segment; posterior protrusion absent; distal process absent. Fifth right swimming leg with endopodite present; fused to basis; on anterior surface; ancestral segments 1 and 2 fused; ancestral segments 2 and 3 fused; stout; ornamented; with setules; as a row; on inner side; terminally; without seta. Fifth right swimming leg exopod segments 1 and 2 separated; segments 2 and 3 fused; 2-segmented; stout; separated from the basis. Fifth right swimming leg exopod 1 trapezium; broader than long; unequal size between both sides; shorter outer than inner side; convex inner side; convex outer side; with marginal extension; sub-triangular; distally inserted; at outer rim; spinules absent; without process; without outer spine; without seta; internal prominence absent; lamella on posterior surface absent. Fifth right swimming leg exopod 2 cylindrical; longer than broad; nearly 2 times; unequal size between both sides; disform inner side; convex outer side; without posterior proximal swelling; inner-posterior process absent; without marginal expansion; curved ridge on distal posterior surface present; chitinous knobs absent; with outer spine; inserted sub-distally; arched; internally directed; not ornamented innerly; not ornamented outerly; sharp tip; with apparent curve; outerly directed; lesser than the length of the exopod 2; until to 2 times its size; 1.5x; sensilla absent; terminal claw present; equal or longer 1.5 times than insertion segment; sclerotized; arched; inward; with conspicuous curve; medially; not ornamented innerly; ornamented outerly; sharp tip; curved tip; outwards; without medial constriction; hyaline process absent.

##### FEMALE

Body longer and wider than male; Female body 0 micrometers excluding caudal setae. Widest at first metasome segment. Distal margin of the prosomal segments without one line of setules at posterior margin. Prosome segments without spinules at prosomal segments. Fourth metasome segment presence of dorsal protuberance; rounded; inserted distally; without posterior process; without anterior process; fourth metasome segment without proximal sensillae present. Fourth and fifth metasome segments fused; partially; on lateral surface. Limit between fourth and fifth metasome segments without ornamentation. **Fifth metasome segment**. Fifth metasome segment without sensilla; with epimeral plates. Epimeral plates asymmetrical. Right epimeral plates prominent, as projections; thinner than the left; one posterior-laterally directed; not reaching half length of the genital segment; with sensilla at the apex; dorsal-posterior sensilla present; slender; without ornamentation. Left epimeral plate without expansion.

##### Urosome

3-segmented. **Genital double-somite**. Asymmetrical in dorsal view; longer than broad; longer than other urosomites combined; dorsal suture at mid-length absent; not covered by spinules; with swelling; rounded; unequal size; greater left than right; anteriorly; with sensillae; on both sides; one; stout; with robust apex; at left lateral; not on lobular base; medially; one; stout; at right lateral; not on lobular base; medially; with robust apex; of equal size between then; lateral protuberance absent; with right posterior rim expanded; over next segment; without slender sensilla on each posterior rim; without posterior-dorsal process. Genital double-somite opercular pad present; broader than longer; symmetrical; development laterally; expanded posteriorly; covering partially; double gonoporal slit; located ventrally; with arthrodial membrane; inserted anteriorly; post-genital process absent; disto-ventral tumescence absent; ventral vertical folds absent; dorsal sensilla absent. Second urosome segment without ventral fusion to anal segment; right distal process absent. Caudal rami patch of setules on outer surface absent; patch of spinules on outer surface absent.

##### Appendices features

Rostrum basal process absent. **Antennules**. Symmetrical. Right antennule not surpassing to genital double-segment; ornamentation pattern equals to male left antennule; mostly. Actual segment 13 without seta; without aesthetasc. Actual segment 14 without seta; without aesthetasc. Actual segment 15 without seta; without aesthetasc. Actual segment 16 without seta; without aesthetasc. Actual segment 17 without seta. Actual segment 18 without seta.

##### Fifth swimming legs

Symmetrical; Fifth swimming legs biramous. Fifth swimming legs intercoxal plate longer than wide; separated from the legs. Fifth swimming legs praecoxa with sclerite praecoxal; separated from the coxae; without ornamentation. Fifth swimming legs coxa with process; conical; at the outer rim; distally; sensilla present; stout; at apex; projecting over basal segment; no longer 2x than basal insertion; marginal extension absent; without swelling; without seta; without spinules. Fifth swimming legs basis sub-triangular; unequal size between inner and outer sides; shorter outer than inner side; with convex inner side; without proximal inner outgrowth; without groove; with distal extension; on posterior surface; with seta; outerly inserted; on anterior surface; no longer 2x than origin segment. Fifth swimming legs endopod segments 1 and 2 fused; segments 2 and 3 fused; 1-segmented; stout; separated from the basis; present discontinuity cuticle; on inner side; with spinules; as a row; single; non-oblique; sub-terminally; at anterior surface; with seta; double; one medially; on posterior surface; rectilinear; one distally; on posterior surface; arched; of unequal size; distal seta longer than medial seta. Fifth swimming legs exopod segments 1 and 2 separated; segments 2 and 3 separated; 3-segmented; separated from the basis. Fifth swimming legs exopod 1 sub-cylindrical; longer than wide; longer or equal than 2 times; with unequal size between inner and outer side; shorter inner than outer side; with convex inner side; with rectilinear outer side; without swelling; without marginal extension; without posterior process; without spine; without seta. Fifth swimming legs exopod 2 sub-cylindrical; longer than broad; longer or equal than 2 times; without swelling; without marginal extension; without process; without lobe; with spine; inserted laterally; rectilinear; without ornamentation; sharp tip; equal size or larger than next segment; without seta. Fifth swimming legs exopod 3 cylindrical; longer than wide; without swelling; without process; without lobe; without spine; with seta; double; inserted terminally; unequal size between them; outer seta smaller than inner; nearly 3 times; outer seta not ornamented by setules; without ornamentation; presence of terminal claw; sclerotized; arched; externally directed; convex inner side; with ornamentation; of denticles; as a row; on surface partially; at medial region; concave outer side; with ornamentation; of denticles; as a row; on surface partially; at medial region; blunt tip; 6 times longer than origin segment.

##### Distribution records

###### BRAZIL

**Minas Gerais**: Amarela Lagoon, Doce River Valley (Dussart, 1985a; Dussart & Matsumura-Tundisi, 1986; Matsumura-Tundisi, 1986).

##### Habitat

Habitat in freshwaters: lake associated to river valley.

##### Remarks

This species was described based on organisms from the environment inserted in a natural valley depression in Southeastern Brazil. In the original description the authors mention the provisional character of its inclusion in *Notodiaptomus* (“le nom de *dubius* à été choisi parce qu’il montre combine il est difficile de ranger a priori cette espèce dans le genre *Notodiaptomus*”) based on important similarities with *Pectenodiaptomus caperatus* (Bowman 1979), later redefined in Reid (1987) as *Notodiaptomus caperatus* Bowman 1979.

The holotype of the species was located in the MZUSP and its condition as *Notodiaptomus* was maintained. All the characters employed by Wright (1935; 1936; 1937) to bring together the *nordestinus* group, and those reviewed by Santos-Silva *et al*. (2015) were present in the organisms of *N. dubius*, and their inclusion in the complex would be possible from this. Among the attributes highlighted by Kiefer (1936; 1956) and Santos-Silva *et al*. (1999) in the designation of the type-species, male fifth right swimming leg basis with posterior lamella (treated in this approach as posterior protrusion) was absent, as was male fifth left swimming leg exopod 1 with single semicircular lobe, and female fifth swimming legs basis with outer seta reaching to exopod 1 distally, respectively in bilobed condition (double), and reaching to the middle exopod 1 for *N. dubius*.

The ornamental pattern of male A1R also was variable to that of *N. deitersi.* For *N. dubius* we present the perpendicular positioning for the modified seta on actual segments 10 and 11, and spiniform modified seta on actual segment 13 reaching to distal-point of segment 14. *N. dubius* can be distinguished from its congeners through combining the following attributes: male fifth right swimming leg coxa with conical process projecting on basis until distal surface posteriorly; female fourth and fifth metasome segments with fusion on lateral surface and rounded dorsal protuberance on fourth metasome segment distally.

#### Notodiaptomus gibber (Poppe in De Guerne & Richard, 1889)

##### Synonymy

*Diaptomus gibber* Poppe (in De Guerne & Richard), 1889: 95, pl. 2, figs. 2, 14, pl. 3, fig. 1, pl. 4, fig. 27; De Guerne & Richard, 1889: pl. 18, tab. 1; Poppe, 1891: 250; Herrick & Turner, 1895: 55, 63, pl. 8, fig. 1; Schmeil, 1897: 172, pl. 14, figs. 4–5; Richard, 1897a: 276, 298; Giesbrecht & Schmeil, 1898: 82; Sars, 1901: 10, 12; Mrázek, 1901: 15; Daday, 1905: 150, 152; Tollinger, 1911: 70, 272, 273, fig. F; Pesta, 1927: 80; Wright, 1927: 73, 75, 89, 100, 102, pl. 6, figs. 4–6; 1938a: 298; 1938b: 562; Brehm, 1935b: 298; 1938: 30, 31; 1958a: 167; Brandorff, 1972: 49. *Notodiaptomus gibber* n. comb., Pallares, 1963: 39, Pl. 1, figs. 1–17; Brandorff, 1976: 616, fig. 2; 1978a: 298; Löffler, 1981: 15; Dussart & Defaye, 1983: 133; 1995: 167; Dussart & Robertson, 1984: 391; Dussart & Frutos, 1986: 306; Matsumura-Tundisi, 1986: 547, fig. 100; Battistoni, 1995: 959; Rocha *et al*., 1995: 156; Santos-Silva, 1998: 209; Santos-Silva *et al*., 1999: 127; Paggi, 2006: 51, 1– 8; Santos-Silva, 2008: 27–28; Perbiche-Neves *et al*., 2020: 678, key to the Neotropical diaptomid. *Diaptomus meridionalis*; Brandorff, 1976; Dussart & Defaye, 1983: 133; Dussart & Defaye, 2002.

##### Type locality

Undefined in the original description. It is generically indicated for the Itajaí region in the Santa Catarina State, Brazil. In Santos-Silva (2008) the occurrence of organisms of this species in water bodies, ponds and artificial lakes is indicated. Possibly, this type of environment is the origin of this species for the place indicated by Popper (1889). Collector Dr. Guilielmo Müller, no date.

##### Type material

No specified in the original description, probably inexistent.

##### Diagnosis

**(1)** Third, fourth, and fifth metasome segment with sensillae dorsally; **(2)** male right antennule actual segment 14 with spinous process; **(3)** left antennule actual segment 3, 7, 9, and 14 with length seta not beyond three sequential segments; **(4)** left antennule actual segment 3 with aesthetasc; **(5)** left antennule actual segment 8 with length seta not larger than segment; **(6)** left antennule actual segment 11 with two setae; **(7)** left antennule actual segment 16 with straight seta; **(8)** male fifth left swimming leg coxa with distal conical process on outer surface posteriorly; **(9)** male fifth left swimming leg basis with rounded inner expansion; **(10)** male fifth left swimming leg exopod 2 in sub-triangular form; **(11)** male fifth left swimming leg exopod 2 with bicuspidate denticles on distal digitiform process; **(12)** male fifth right swimming leg exopod 1 with concave outer side; **(13)** male fifth right swimming leg exopod 1 with digitiform internal prominence; **(14)** male fifth right swimming leg exopod 2 in elliptical form; **(15)** male fifth right swimming leg exopod 2 longer than broad 2x nearly; **(16)** male fifth right swimming leg exopod 2 with outer spine lesser than length exopod 2 beyond to 2x its size; **(17)** female fourth metasome segment with acute-form posterior process on dorsal protuberance medially, with anterior process, and proximal sensilla; **(18)** female left epimeral plate with semicircular expansion on medial surface dorsally; **(19)** female genital double-somite with sensilla at left rim, not on lobular basis; **(20)** female genital double-somite with lateral sensillae of unequal size between then, left bigger than right; **(21)** female genital double-somite with posterior-dorsal process on the right side doubly; **(22)** female right antennule with length not exceeding the caudal setae.

##### Material examined

Non-type material: 2 males and 1 female, entire in alcohol, from the original collection of Friedrich Kiefer (n. 4077), collected in several pools in Paraguay; 1 male (INPA-COP021, slides a-h) and 1 female (INPA-COP022, slides a-h) were selected to be dissection on eight slides each and deposited in the Zoological Collection of the INPA, Brazil. Additional material examined: 1 male and 3 females, entire in alcohol, from the flood water in Rio Grande do Sul, Brazil, M. G. Bandeira coll., 30.X.2017; 2 males and 1 female, entire in alcohol, from the Yaciretá Dam Reservoir, River Parana between Argentina, and Paraguay, (27° 24m 24s S 56° 15m 9s W), Perbiche-Neves coll., 2010.

##### Redescription

###### MALE

Body 1606 micrometers excluding caudal setae. Male body smaller and slenderer than female. Nerve axons myelinated. Prosome 6-segmented; widest at first metasome segment; without one line of setules at posterior margin; without spinules at segments. Cephalosome anterior margin sub-triangular; with dorsal suture; incomplete; separate from first metasome segment. First metasome segment without sensilla. Second metasome segment without sensilla. Third metasome segment with sensillae; 2 dorsally; of equal size; non-ornamented posterior margin. Fourth metasome segment with sensillae; 4 dorsally; of equal size; separated from the fifth metasome. Limit between fourth and fifth metasome segments without ornamentation. Fifth metasome segment with sensilla; 2 dorsally; Fifth metasome segment equal size; Fifth metasome segment without ornamentation; Fifth metasome segment without dorsal conical process; with epimeral plates. Epimeral plates symmetrical. Right epimeral plates reduced, as rounded distal corner segment limit; with sensilla; one at the apex of projection and other medially; without ornamentation.

##### Urosome

5-segmented; Urosome 5 - free segments. Genital somite asymmetrical in dorsal view; with single aperture; located on left side; ventrolaterally on posterior rim; with sensillae; on both sides; one; at left lateral; posteriorly; one; at right rim; posteriorly; of equal size between then. Third urosome segment without spinules; without external seta. Fourth urosome segment without spinules; without sub-conical blunt dorsal-lateral process. Anal segment presence of dorsal sensillae; one on each side; medially inserted; presence of operculum; convex; covering the anal aperture fully. Caudal rami symmetrical; separated from anal segment; longer than wide; with setules; continuous on; inner side; each ramus bearing 6 caudal setae; 5 marginals; plumose; and 1 internal dorsally; straight; not reticulated main axis; outermost seta with outer spiniform process absent.

##### Appendices features

Rostrum asymmetrical; separated from dorsal cephalic shield; by complete suture; sensillae present; one pair; anteriorly inserted on surface tegument; with rostral filament; double; paired; extended; into blunt protrusion; with basal process; in ventral view, rounded on left side; with a smaller basal expansion on the right side.

##### Antennules

Asymmetrical. **Right antennules**. Uniramous; right antennule surpassing to genital segment; right antennule extending beyond caudal rami.

Right antennule ancestral segment I and II separated. Ancestral segment II and III fused. Ancestral segment III and IV fused. Ancestral segment IV and V separated. Ancestral segment V and VI separated. Ancestral segment VI and VII separated. Ancestral segment VII and VIII separated. Ancestral segment VIII and IX separated. Ancestral segment IX and X separated. Ancestral segment X and XI separated. Ancestral segment XI and XII separated. Ancestral segment XII and XIII separated. Ancestral segment XIII and XIV separated. Ancestral segment XIV and XV separated. Ancestral segment XV and XVI separated. Ancestral segment XVI and XVII separated. Ancestral segment XVII and XVIII separated. Ancestral segment XVIII and XIX separated. Ancestral segment XIX and XX separated. Ancestral segment XX and XXI separated. Ancestral segment XXI and XXII fused. Ancestral segment XXII and XXIII fused. Ancestral segment XXIII and XXIV separated. Ancestral segment XXIV and XXV fused. Ancestral segment XXV and XXVI separated. Ancestral segment XXVI and XXVII separated. Ancestral segment XXVII and XXVIII fused.

Right antennule actual 22-segmented; geniculated; between the segment 18 and segment 19; with swollen and modified region; formed by 5 segments; between 13 and 17 segments. Actual segment 1 with seta; one element; straight; none larger than segment; without spinules; without vestigial seta; without conical seta; without modified seta; without spinous process; with aesthetasc; one element. Actual segment 2 with seta; three elements; of unequal size; straight; none larger than segment; without spinules; with vestigial seta; one element; without conical seta; without modified seta; without spinous process; with aesthetasc; one element. Actual segment 3 with seta; one element; none larger than segment; straight; sharp apex; without spinules; with vestigial seta; one element; without conical seta; without modified seta; without spinous process; with aesthetasc. Actual segment 4 with seta; one element; one larger than segment; surpassing to distal margin; straight; not beyond three sequential segments; without spinules; without vestigial seta; without conical seta; without modified seta; without spinous process; without aesthetasc. Actual segment 5 with seta; one element; straight; one larger than segment; surpassing to distal margin; not beyond three sequential segments; without spinules; with vestigial seta; one element; without conical seta; without modified seta; without spinous process; without aesthetasc. Actual segment 6 with seta; one element; none larger than segment; straight; without spinules; without vestigial seta; without conical seta; without modified seta; without spinous process; without aesthetasc. Actual segment 7 with seta; one element; straight; none larger than segment; sharp apex; without spinules; without vestigial seta; without conical seta; without modified seta; without spinous process; with aesthetasc; one element. Actual segment 8 with seta; one element; straight; none larger than segment; without spinules; without vestigial seta; with conical seta; one element; not reaching to middle-point of the sequent segment; without modified seta; without spinous process; without aesthetasc. Actual segment 9 with seta; two elements; of unequal size; straight; none larger than segment; sharp apex; without spinules; without vestigial seta; without conical seta; without modified seta; without spinous process; with aesthetasc; one element. Actual segment 10 with seta; one element; straight; none larger than segment; without spinules; without vestigial seta; without conical seta; with modified seta; presenting blunt apex; slender form; surpassing to distal margin; not beyond of the sequential segment; parallel to antennule direction; without spinous process; without aesthetasc. Actual segment 11 with seta; one element; straight; one larger than segment; surpassing to distal margin; not beyond three sequential segments; without spinules; without vestigial seta; without conical seta; with modified seta; slender form; presenting blunt apex; surpassing to distal margin; not beyond of the sequential segment; perpendicular to antennule direction; shorter length than homologous of actual segment 13; without spinous process; without aesthetasc. Actual segment 12 with seta; one element; straight; one larger than segment; surpassing to distal margin; not beyond three sequential segments; without spinules; without vestigial seta; with conical seta; one element; not smaller than to segment 8; without modified seta; without spinous process; with aesthetasc; one element; absent internal perpendicular fission. Actual segment 13 with seta; one element; straight; one larger than segment; surpassing to distal margin; not beyond three sequential segments; without spinules; without vestigial seta; without conical seta; with modified seta; stout form; surpassing to distal margin; to the middle-point of the sequence segment; perpendicular to antennule direction; presenting rounded apex; without spinous process; with aesthetasc; one element. Actual segment 14 with seta; two elements; of unequal size; straight; one larger than segment; surpassing to distal margin; beyond three sequential segments; blunt apex; without spinules; without vestigial seta; without conical seta; without modified seta; with spinous process; without aesthetasc. Actual segment 15 with seta; two elements; of unequal size; straight; not bifidform; none larger than segment; without spinules; without vestigial seta; without conical seta; without modified seta; with spinous process; on outer margin; surpassing distal margin; with aesthetasc; one element. Actual segment 16 with seta; two elements; of unequal size; plumose; one larger than segment; surpassing to distal margin; not beyond three sequential segments; not bifidform; without spinules; without vestigial seta; without conical seta; without modified seta; with spinous process; on outer margin; surpassing distal margin; unequal size to process on preceding segment; with aesthetasc; one element. Actual segment 17 with seta; two elements; of unequal size; straight; none larger than segment; bifidform; without spinules; without vestigial seta; without conical seta; with modified seta; one element; stout form; surpassing to distal margin; not beyond of the sequential segment; parallel to antennule direction; without spinous process; without aesthetasc. Actual segment 18 with seta; two elements; of equal size; straight; none larger than segment; without spinules; without vestigial seta; without conical seta; with modified seta; one element; stout form; surpassing distal margin; parallel to antennule direction; without spinous process; without aesthetasc. Actual segment 19 with seta; two elements; of unequal size; plumose; none larger than segment; without spinules; without vestigial seta; without conical seta; with modified seta; two elements; stout form; at least one bifid form; surpassing distal margin; parallel to antennule direction; without spinous process; with aesthetasc; one element. Actual segment 20 with seta; four elements; of unequal size; straight; one larger than segment; surpassing to distal margin; beyond three sequential segments; without spinules; without vestigial seta; without conical seta; without modified seta; with spinous process; distally; not reaching beyond of distal-point segment 21; without aesthetasc. Actual segment 21 with seta; two elements; of equal size; plumose; one larger than segment; surpassing to distal margin; greater 3x than original segment; without spinules; without vestigial seta; without conical seta; without modified seta; without spinous process; without aesthetasc. Actual segment 22 with seta; four elements; of equal size; one larger than segment; plumose; surpassing to distal margin; greater 3x than original segment; without spinules; without vestigial seta; without conical seta; without modified seta; without spinous process; with aesthetasc; one element.

##### Left antennules

Uniramous; Left antennule surpassing to prosome; Left antennule extending beyond caudal rami. Ancestral segment I and II separated. Ancestral segment II and III fused. Ancestral segment III and IV fused. Ancestral segment IV and V separated. Ancestral segment V and VI separated. Ancestral segment VI and VII separated. Ancestral segment VII and VIII separated. Ancestral segment VIII and IX separated. Ancestral segment IX and X separated. Ancestral segment X and XI separated. Ancestral segment XI and XII separated. Ancestral segment XII and XIII separated. Ancestral segment XIII and XIV separated. Ancestral segment XIV and XV separated. Ancestral segment XV and XVI separated. Ancestral segment XVI and XVII separated. Ancestral segment XVII and XVIII separated. Ancestral segment XVIII and XIX separated. Ancestral segment XIX and XX separated. Ancestral segment XX and XXI separated. Ancestral segment XXI and XXII separated. Ancestral segment XXII and XXIII separated. Ancestral segment XXIII and XXIV separated. Ancestral segment XXIV and XXV separated. Ancestral segment XXV and XXVI separated. Ancestral segment XXVI and XXVII separated. Ancestral segment XXVII and XXVIII fused.

Left antennule actual 25-segmented; not-geniculated. Actual segment 1 with seta; one element; none larger than segment; straight; without spinules; without vestigial seta; without conical seta; without modified seta; without spinous process; with aesthetasc; one element. Actual segment 2 with seta; three elements; of equal size; none larger than segment; straight; without spinules; with vestigial seta; one element; without conical seta; without modified seta; without spinous process; with aesthetasc; one element. Actual segment 3 with seta; one element; one larger than segment; straight; surpassing to distal margin; not beyond three sequential segments; without spinules; with vestigial seta; one element; without conical seta; without modified seta; without spinous process; with aesthetasc; one element. Actual segment 4 with seta; one element; none larger than segment; straight; without spinules; without vestigial seta; without conical seta; without modified seta; without spinous process; without aesthetasc. Actual segment 5 with seta; one element; one larger than segment; straight; surpassing to distal margin; not beyond three sequential segments; without spinules; with vestigial seta; one element; without conical seta; without modified seta; without spinous process; with aesthetasc; one element. Actual segment 6 with seta; one element; none larger than segment; straight; without spinules; without vestigial seta; without conical seta; without modified seta; without spinous process; without aesthetasc. Actual segment 7 with seta; one element; one larger than segment; straight; surpassing to distal margin; not beyond three sequential segments; without spinules; without vestigial seta; without conical seta; without modified seta; without spinous process; with aesthetasc; one element. Actual segment 8 with seta; one element; none larger than segment; straight; surpassing distal margin; without spinules; without vestigial seta; with conical seta; without modified seta; without spinous process; without aesthetasc. Actual segment 9 with seta; two elements; of unequal size; one larger than segment; straight; surpassing to distal margin; not beyond three sequential segments; without spinules; without vestigial seta; without conical seta; without modified seta; without spinous process; with aesthetasc; one element. Actual segment 10 with seta; one element; one larger than segment; straight; surpassing to distal margin; not beyond three sequential segments; without spinules; without vestigial seta; without conical seta; without modified seta; without spinous process; without aesthetasc. Actual segment 11 with seta; two elements; of unequal size; one larger than segment; straight; surpassing to distal margin; not beyond three sequential segments; without spinules; without vestigial seta; without conical seta; without modified seta; without spinous process; without aesthetasc. Actual segment 12 with seta; one element; one larger than segment; straight; surpassing distal margin; without spinules; without vestigial seta; with conical seta; without modified seta; without spinous process; with aesthetasc; one element. Actual segment 13 with seta; one element; none elongated; straight; surpassing distal margin; without spinules; without vestigial seta; without conical seta; without modified seta; without spinous process; without aesthetasc. Actual segment 14 with seta; one element; elongated; straight; surpassing to distal margin; not beyond three sequential segments; without spinules; without vestigial seta; without conical seta; without modified seta; without spinous process; with aesthetasc; one element. Actual segment 15 with seta; one element; larger than segment; straight; surpassing to distal margin; not beyond three sequential segments; without spinules; without vestigial seta; without conical seta; without modified seta; without spinous process; without aesthetasc. Actual segment 16 with seta; one element; larger than segment; straight; surpassing to distal margin; not beyond three sequential segments; without spinules; without vestigial seta; without conical seta; without modified seta; without spinous process; with aesthetasc; one element. Actual segment 17 with seta; one element; not larger than segment; straight; without spinules; without vestigial seta; without conical seta; without modified seta; without spinous process; without aesthetasc. Actual segment 18 with seta; one element; larger than segment; straight; surpassing to distal margin; not beyond three sequential segments; without spinules; without vestigial seta; without conical seta; without modified seta; without spinous process; without aesthetasc. Actual segment 19 with seta; one element; not larger than segment; straight; surpassing distal margin; without spinules; without vestigial seta; without conical seta; without modified seta; without spinous process; with aesthetasc; one element. Actual segment 20 with seta; one element; not larger than segment; straight; surpassing distal margin; without spinules; without vestigial seta; without conical seta; without modified seta; without spinous process; without aesthetasc. Actual segment 21 with seta; one element; larger than segment; plumose; surpassing to distal margin; not beyond three sequential segments; without spinules; without vestigial seta; without conical seta; without modified seta; without spinous process; without aesthetasc. Actual segment 22 with seta; two elements; of unequal size; one of them elongated; plumose; surpassing to distal margin; without spinules; without vestigial seta; without conical seta; without modified seta; without spinous process; without aesthetasc. Actual segment 23 with seta; two elements; of unequal size; one larger than segment; plumose; surpassing to distal margin; greater 3x than original segment; without spinules; without vestigial seta; without conical seta; without modified seta; without spinous process; without aesthetasc. Actual segment 24 with seta; two elements; of equal size; one larger than segment; plumose; surpassing to distal margin; greater 3x than original segment; without spinules; without vestigial seta; without conical seta; without modified seta; without spinous process; without aesthetasc. Actual segment 25 with seta; four elements; of equal size; elongated; plumose; surpassing to distal margin; 4 times larger than segment; without spinules; without vestigial seta; without conical seta; without modified seta; without spinous process; with aesthetasc; one element.

##### Antenna

Biramous. Antenna coxa separated from the basis; bearing seta; 1; on inner surface; at distal corner; reaching to the middle basis. Antenna basis (fusion) separated from the endopodal segment; bearing seta; 2; on inner surface; at distal corner. Endopodal ancestral segment I and II separated. Ancestral segment II and III fused. Ancestral segment III and IV fused. Ancestral segment III and IV fully. Antenna endopod actual 2-segmented. Actual segment 1 not bilobate; with seta; two; on inner margin; with spinules; as a row; obliquely; on outer surface; without pore. Actual segment 2 bilobate; with discontinuity on outer cuticle; developed as a suture; incomplete; inner lobe bearing 8 setae; distally; outer lobe bearing 7 setae; distally; with spinules; as a patch; on outer surface. Antenna exopod ancestral segment I and II separated. Ancestral segment II and III fused. Ancestral segment III and IV fused. Ancestral segment IV and V separated. Ancestral segment V and VI separated. Ancestral segment VI and VII separated. Ancestral segment VII and VIII separated.

Ancestral segment VIII and IX separated. Ancestral segment IX and X fused. Antenna exopod actual 7-segmented. Actual segment 1 single; elongated (width-length, equal or larger ratio 2:1); with seta; one; at inner surface. Actual segment 2 compound; elongated (larger width-length ratio 2:1); with seta; three; at inner surface. Actual segment 3 single; not elongated (lesser width-length ratio 2:1); with seta; one; at inner surface. Actual segment 4 single; not elongated (lesser width-length ratio 2:1); with seta; one; at inner surface. Actual segment 5 single; not elongated (lesser width-length ratio 2:1); with seta; one; at inner surface. Actual segment 6 single; not elongated (lesser width-length ratio 2:1); with seta; one; at inner surface. Actual segment 7 compound; elongated (larger or equal width-length ratio 2:1); with seta; one; at inner surface; and three; at distal surface.

##### Oral features

**Mandible**. Coxal gnathobase sclerotized; with lobe; prominent; on caudal margin; presence of cutting blade; with tooth-like prominence; two, distinctly; 1 acute; on caudal margin; and 1 triangular; on sub-caudal margin; without acute projection between the prominences; without additional spinules; with seta; 1; dorsally; on apical surface; without spinules. Mandible palps biramous; comprising the basis; with seta; four; differently inserted; first medially; reaching to beyond the endopod 1; second distally; third distally; fourth distally; on inner margin; none with setulose ornamentation. Mandible endopod 2-segmented. Mandible endopod 1 with lobe; bearing seta; four; distally inserted; without spinules. Mandible endopod 2 without lobe; bearing setae; nine elements; distally inserted; with spinules; as a patch; double. Mandible exopod 4-segmented. Mandible exopod 1 with seta; one element; distally; on inner margin. Mandible exopod 2 with seta; one element; distally; on inner side. Mandible exopod 3 with seta; one element; distally; on inner side. Mandible exopod 4 with setae; three elements; on terminal region. **Maxillule**. Birramous. Maxillule 3-segmented. Maxillule praecoxa with praecoxal arthrite; bearing spines; fifteen elements; ten marginally; plus, five sub-marginally; with spinules; as a patch; on sub-marginal surface. Maxillule coxa with coxal epipodite; with conspicuous outer lobe; bearing setae; nine elements; with coxal endite; elongated (larger or equal width-length ratio 2:1); bearing setae; four elements. Maxillule basis with basal endite; double; first proximal; elongated (larger width-length ratio 2:1; separated from basis; with setae; four elements; distally inserted; second distal; fused to basis; not elongated (lesser width-length ratio 2:1); with setae; four elements; distally inserted; with setules; as a row; on inner side; basal exite present; with setae; one element; on outer surface. Maxillule endopod 1-segmented. Endopod 1 bilobate; first proximal; with setae; three elements; second distal; with setae; five elements. Maxillule exopod 1-segmented. Exopod 1 with setae; six elements; with setules; as a row; on inner side; spinules absent. **Maxilla**. Uniramous. Maxilla 5-segmented. Maxilla praecoxa fused to coxa; incompletely; distinct externally; with praecoxal endite; double; first elongated endite (larger or equal width length ratio 2:1); proximally inserted; with seta; straight, or plumose; 1 straight; 4 plumose; with spine; single; without spinules; without setule; second elongated endite (larger or equal width length ratio 2:1); distally inserted; with seta; plumose; 3 plumose; without spine; with spinules; as a row; on distal margin; with setule; as a row; on distal margin; absence of outer seta. Maxilla coxa with coxal endite; double; first elongated endite (larger or equal width); proximally inserted; with seta; plumose; 3 plumose; without spine; without spinules; with setules; as a row; on proximal margin; second elongated endite (larger or equal width); distally inserted; with seta; plumose; 3 plumose; without spine; without spinules; with setules; as a row; on proximal margin; absence of outer seta. Maxilla basis with basal endite; single; elongated (larger or equal width-length ratio 2:1); with seta; plumose; 3 plumose; without spinules; absence of outer seta. Maxilla endopod 2-segmented. Endopod 1 with seta; 2 plumose; without spine; without spinules; without setules. Maxilla endopod 2 with seta; 2 plumose; without spine; without spinules; without setules. **Maxilliped**. Uniramous; Maxilliped 8-segmented. Maxilliped praecoxa fused to coxa; completely; with praecoxal endite; not elongated (lesser width-length ratio 2:1); distally inserted; with seta; 1 straight; with spinules; as a row; single; on basal surface; without setules. Maxilliped coxa with coxal endite; three coxal endite; first elongated (larger or equal width); proximally inserted; with seta; 2 plumose; with spinules; as a patch; single; on apical surface; without setules; second not elongated (lesser width-length ratio 2:1); medially inserted; with seta; 3 plumose; with spinules; as a row; single; on medial surface; without setules; third elongated (larger or equal width length ratio 2:1); distally inserted; with seta; 3 plumose; none reaching to beyond of the basis; with spinules; as a row; single; on basal surface; without setules; with lobe; prominence; at inner distal angle; ornamented; with spinules; continuously on margin. Maxilliped basis without basal endite; with seta; 3 plumose; with spinules; as a row; single; on medial surface; with setules; as a row; single; on inner margin. Maxilliped endopod segment 6-segmented. Endopod 1 with seta; 2 plumose; on inner surface. Endopod 2 with seta; 3 plumose; on inner surface. Endopod 3 with seta; 2 plumose; on inner surface. Endopod 4 with seta; 2 plumose; on inner surface. Endopod 5 with seta; 2 plumose; on inner surface, or on outer surface; outer seta absent. Endopod 6 with seta; 4 plumose; on inner surface, or on outer surface.

##### Swimming legs features

**First swimming legs.** Symmetrical; biramous. First swimming legs intercoxal plate without seta. First swimming legs praecoxa absent. First swimming legs coxa with seta; one; straight; distally inserted; on inner surface; surpassing to first endopodal segment; with setules; two group; as a patch; on inner margin; and as a row; double; on anterior surface; outerly; without spinules; without spine. First swimming legs basis without seta; with setules; as a patch; single; on outer surface; without spinules; without spine. First swimming legs endopod 2-segmented. Endopod 1 with seta; straight; restricted; to inner surface; one element; without spine; with setules; as a row; single; continuously; on outer surface; without spinules; absence of Schmeil’s organ. Endopod 2 with seta; unrestricted; three on inner surface; one on outer surface; two on distal surface; straight; without spine; with setules; as a row; single; continuously; on outer surface; without spinules; absence of Schmeil’s organ. Endopod 3 absence. First swimming legs exopod 1 with seta; restricted; 1 on inner surface; with spine; 1; stout; smaller than original segment; serrated; on inner side; continuously; with setules; as a row; single; as a row; innerly. First swimming legs exopod 2 with seta; restricted; 1 on inner surface; straight; without spine; with setules; as a row; single; continuously; on inner margin, or on outer margin; without spinules. First swimming legs exopod 3 with setule; as a row; single; continuously; on outer surface; without spinules; with seta; unrestricted; 2 on inner surface; 2 on terminal surface; with spine; 2; unequal size; first no longer 2x than origin segment; stout; serrated; on inner side, or on outer side; equally; second longer 3x than origin segment; slender; serrated; on outer side; with ornamentation on non-serrated side; by setules. **Second swimming legs**. Symmetrical; Second swimming legs biramous. Second swimming legs intercoxal plate without seta. Second swimming legs praecoxa present; located laterally. Second swimming legs coxa with seta; straight; distally inserted; on inner surface; surpassing to basal segment; without setules; without spinules; without spine. Second swimming legs basis without seta; without setules; without spinules; without spine. Second swimming legs endopod 3-segmented. Endopod 1 with seta; straight; restricted; one on inner surface; without spine; with setules; as a row; single; continuously; on outer surface; without spinules; absence of Schmeil’s organ. Endopod 2 with seta; straight; unrestricted; two on inner surface; without spine; with setules; as a row; single; continuously; on outer side; without spinules; presence of Schmeil’s organ; on posterior surface. Endopod 3 with seta; straight; unrestricted; three on inner surface; two on outer surface; two on distal surface; without spine; without setules; with spinules; as a row; double; distally inserted; at anterior surface; absence of Schmeil’s organ. Second swimming legs exopod 1 with seta; restricted; one on inner surface; with spine; 1; stout; not reaching to distal-third of the exopod 2; serrated; on inner side, or on outer side; with setules; as a row; single; continuously; on inner side; without spinules; absence of Schmeil’s organ. Exopod 2 with seta; unrestricted; one on inner surface; with spine; 1; stout; not surpassing the exopod 3; serrated; on inner side, or on outer side; with setules; as a row; single; continuously; on inner surface; without spinules; absence of Schmeil’s organ. Exopod 3 with seta; plurimarginal; three on inner surface; two on terminal surface; with spine; 2; unequal size; first no longer 2x than origin segment; stout; serrated; on inner side, or on outer side; equally; second longer 2x than origin segment; slender; serrated; on outer side; with ornamentation on non-serrated side; of setules; setules on outer surface; as a row; single; continuously; on inner surface; with spinules; as a row; single; distally inserted; at anterior surface; absence of Schmeil’s organ. **Third swimming legs**. Symmetrical; Third swimming legs biramous. Third swimming legs intercoxal plate without seta. Third swimming legs praecoxa present; not laterally located. Third swimming legs coxa with seta; straight; distally inserted; on inner surface; surpassing to first endopodal segment; without setules; without spinules; without spine. Third swimming legs basis without seta; without setules; without spinules; without spine. Third swimming legs endopod 3-segmented. Endopod 1 with seta; restricted; one on inner surface; without spine; without setules; without spinules; absence of Schmeil’s organ. Endopod 2 with seta; restricted; two on inner surface; straight; without spine; without setules; without spinules; absence of Schmeil’s organ. Endopod 3 with seta; straight; plurimarginal; two on inner surface; two on outer surface; three on terminal surface; without spine; without setules; with spinules; as a row; distally inserted; double; at anterior surface; absence of Schmeil’s organ. Third swimming legs exopod 1 with seta; restricted; straight; one on inner surface; with spine; 1; stout; not reaching to the distal-third of the exopod 2; serrated; equally; on inner surface, or on outer surface; with setules; as a row; single; continuously; on inner surface; without spinules; absence of Schmeil’s organ. Exopod 2 with seta; straight; restricted; one on inner surface; with spine; 1; stout; not reaching out to exopod 3; serrated; on inner side, or on outer side; equally; with setules; as a row; single; continuously; on inner side; without spinules; absence of Schmeil’s organ. Exopod 3 without setules; with spinules; as a row; single; distally inserted; at anterior surface; with seta; straight; unrestricted; three on inner surface; two on terminal surface; with spine; 2; unequal size; first no longer 2x than origin segment; stout; serrated; on inner side, or on outer side; equally; second longer 2x than origin segment; slender; serrated; on outer side; with ornamentation on non-serrated side; of setules; absence of Schmeil’s organ. **Fourth swimming legs**. Symmetrical; biramous. Intercoxal plate without sensilla. Praecoxa present. Coxa with seta; distally inserted; on inner margin; reaching out to endopod 1; without spinules; setules absent. Basis with seta; one; medially inserted; on posterior surface; smaller than the original segment; without setules; without spinules; without spine. Fourth swimming legs endopod 3-segmented. Endopod 1 with seta; one; restricted; on inner surface; without spine; without setules; without spinules; absence of Schmeil’s organ. Endopod 2 with seta; restricted; two on inner side; without spine; with setules; as a row; single; continuously; on outer surface; without spinules; absence of Schmeil’s organ. Endopod 3 with seta; unrestricted; two on inner surface; two on outer surface; three on distal surface; without spine; without setules; with spinules; as a row; double; distally inserted; at anterior surface; absence of Schmeil’s organ. Fourth swimming legs exopod 1 with seta; restricted; one on inner surface; with spine; 1; stout; not reaching out to distal-third of the exopod 2; serrated; on inner side, or on outer side; equally; with setules; as a row; single; continuously; on inner surface; without spinules; absence of Schmeil’s organ. Exopod 2 with seta; restricted; one on inner surface; with spine; 1; stout; not reaching the end of exopod 3; serrated; on inner side, or on outer side; equally; with setules; as a row; single; continuously; on inner surface; without spinules; absence of Schmeil’s organ. Exopod 3 without setules; with spinules; as a row; single; distally inserted; at anterior surface; with seta; unrestricted; three on inner surface; two on distal surface; with spine; 2; unequal size; first no longer 2x than origin segment; stout; serrated; on inner side, or on outer side; equally; second longer 2x than origin segment; slender; serrated; on outer side; without ornamentation on non-serrated side; absence of Schmeil’s organ.

##### Fifth swimming legs features

Asymmetrical. Fifth swimming leg intercoxal plate with length not equal or greater than width on 1.5x; with irregular proximal margin; discontinuous to; the anterior margin of the left coxa, or the anterior margin of the right coxa; posterior sensilla on the right lateral absent. **Fifth left swimming leg**. Fifth left swimming leg biramous; leg reaching first right exopod segment; medially. Fifth left swimming leg praecoxa present; rudimentary; separated from the coxae; without ornamentation. Fifth left swimming leg coxa concave inner side; without teeth-like structures; with process; conical; on marginal surface; outer side; distally inserted; not projecting over basis; with sensilla; stout; triangular; at apex; longer 2x than insertion basis; without swelling; without seta; without spinules. Fifth left swimming leg basis sub-cylindrical; unequal size between inner and outer side; shorter outer than inner side; with concave inner side; rounded internal proximal expansion present; without outgrowth; without groove; absence of protuberance; with seta; outerly inserted; no longer 2x than origin segment; absence of minutely granular. Fifth left swimming leg endopod segments 1 and 2 separated; segments 2 and 3 fused; 2-segmented; stout; separated from the basis; ornamented; ornamented on segment 2; on inner side; with spinules; more than four elements; as a row; terminally; row of setules absent; with seta. Fifth left swimming leg exopod segments 1 and 2 separated; segments 2 and 3 fused; 2-segmented; stout; separated from the basis. Fifth left swimming leg exopod 1 sub-triangular; longer than broad; equal size between inner and outer side; rectilinear inner side; convex outer side; without swelling; without marginal extension; without process; with lobe; single; semicircular; medially inserted; on inner side; covered; by setules; without outer spine; absence seta. Fifth left swimming leg exopod 2 sub-triangular; longer than broad; equal size between inner and outer side; disform inner side; with convex outer side; setulose pad present; prominently rounded; proximally; on inner side; inflated medial region absent; distal process present; digitiform; denticulate; bicuspidate; with transverse row of denticles; none oblique row of 5 denticles; at anterior surface; innerly directed; with seta; spiniform; ornamented by spinules; surpassing the distal-point of the segment; without outer spine; terminal claw absent.

##### Fifth right swimming leg

Biramous. Fifth right swimming leg praecoxa present; separated from the coxae; without ornamentation. Fifth right swimming leg coxa convex inner side; without teeth-like structures; with process; rounded; distally inserted; on posterior surface; closest to the outer rim; projecting over basis; not beyond the first third; without triangular protuberance innerly; with sensilla; slender; at apex; no longer 2x than basal insertion; without marginal extension; without seta; without spinules. Fifth right swimming leg basis trapezoidal; unequal size between inner and outer side; shorter outer than inner side; convex inner side; tumescence present; inflated; not bilobed; unrestricted on inner surface; with protuberance; single; semicircular; on inner side; proximally; not ornamented; absence of distinct minutely granular; additional inner process present; triangular; medially; without posterior groove; with seta; outerly inserted; on anterior surface; no longer 2x than origin segment; posterior protrusion absent; distal process absent. Fifth right swimming leg with endopodite present; fused to basis; on anterior surface; ancestral segments 1 and 2 fused; ancestral segments 2 and 3 fused; slender; ornamented; with setules; as a row; on inner side; terminally; with seta. Fifth right swimming leg exopod segments 1 and 2 separated; segments 2 and 3 fused; 2-segmented; stout; separated from the basis. Fifth right swimming leg exopod 1 trapezium; longer than broad; nearly 1.25 times; unequal size between both sides; shorter inner than outer side; convex inner side; concave outer side; with marginal extension; sub-triangular; distally inserted; at outer rim; spinules absent; with process; rounded; sclerotized; without ornamentation; distally inserted; at posterior surface; projecting over next segment; without outer spine; without seta; internal prominence present; digitiform; lamella on posterior surface absent. Fifth right swimming leg exopod 2 elliptical; longer than broad; nearly 2 times; unequal size between both sides; uniform inner side; convex outer side; without posterior proximal swelling; inner-posterior process absent; without marginal expansion; curved ridge on distal posterior surface present; chitinous knobs absent; with outer spine; inserted sub-distally; rectilinear; not ornamented innerly; not ornamented outerly; sharp tip; without apparent curve; lesser than the length of the exopod 2; beyond to 2 times its size; 3x; sensilla absent; terminal claw present; equal or longer 1.5 times than insertion segment; sclerotized; arched; inward; with conspicuous curve; medially; ornamented innerly; by spinules; as a row; partially on extension; medially, or distally; not ornamented outerly; sharp tip; curved tip; inwards; with medial constriction; hyaline process absent.

##### FEMALE

Body longer and wider than male; Female body 1711 micrometers excluding caudal setae. Widest at posterior cephalosome. Distal margin of the prosomal segments without one line of setules at posterior margin. Prosome segments without spinules at prosomal segments. Fourth metasome segment presence of dorsal protuberance; digitiform; 2x longer than wide; inserted medially; with posterior process; stout-form; medially; with anterior process; acute-form; medially; fourth metasome segment with proximal sensillae present. Fourth and fifth metasome segments fused; totally. Limit between fourth and fifth metasome segments without ornamentation. **Fifth metasome segment**. Fifth metasome segment without sensilla; with epimeral plates. Epimeral plates asymmetrical. Right epimeral plates prominent, as projections; thinner than the left; two posterior-laterally directed; not reaching half length of the genital segment; with sensilla at the apex; dorsal-posterior sensilla absent; without ornamentation. Left epimeral plate with expansion; semicircular; on posterior surface; dorsally; without sensilla.

##### Urosome

3-segmented. **Genital double-somite**. Asymmetrical in dorsal view; longer than broad; longer than other urosomites combined; dorsal suture at mid-length absent; not covered by spinules; with swelling; conical; unequal size; greater right than left; anteriorly; with sensillae; on both sides; one; stout; with robust apex; at left lateral; not on lobular base; anteriorly; one; stout; at right lateral; not on lobular base; anteriorly; with robust apex; of unequal size between then; left bigger than right; lateral protuberance absent; with right posterior rim expanded; over next segment; without slender sensilla on each posterior rim; with posterior-dorsal process; in a single side; double on the right. Genital double-somite opercular pad present; broader than longer; symmetrical; development laterally; expanded posteriorly; covering partially; double gonoporal slit; located ventrally; with arthrodial membrane; inserted anteriorly; post-genital process absent; disto-ventral tumescence absent; ventral vertical folds absent; dorsal sensilla absent. Second urosome segment without ventral fusion to anal segment; right distal process absent. Caudal rami patch of setules on outer surface absent; patch of spinules on outer surface absent.

##### Appendices features

Rostrum basal process absent. **Antennules**. Symmetrical. Right antennule surpassing to genital double-segment; extending beyond caudal rami. Right antennule not exceeding the caudal setae. Right antennule ornamentation pattern equals to male left antennule; fully.

##### Fifth swimming legs

Symmetrical; Fifth swimming legs biramous. Fifth swimming legs intercoxal plate longer than wide; separated from the legs. Fifth swimming legs praecoxa with sclerite praecoxal; separated from the coxae; without ornamentation. Fifth swimming legs coxa with process; conical; at the outer rim; distally; sensilla present; stout; at apex; not projecting over basal segment; no longer 2x than basal insertion; marginal extension absent; without swelling; without seta; without spinules. Fifth swimming legs basis sub-triangular; unequal size between inner and outer sides; shorter outer than inner side; with convex inner side; without proximal inner outgrowth; without groove; with distal extension; on posterior surface; with seta; outerly inserted; on anterior surface; no longer 2x than origin segment. Fifth swimming legs endopod segments 1 and 2 separated; segments 2 and 3 fused; 2-segmented; with complete suture; stout; separated from the basis; ornamentation on segment 2; with spinules; as a row; single; non-oblique; sub-terminally; at anterior surface; with seta; double; one medially; on posterior surface; rectilinear; one distally; on posterior surface; arched; of unequal size; proximal seta longer than distal seta. Fifth swimming legs exopod segments 1 and 2 separated; segments 2 and 3 separated; 3-segmented; separated from the basis. Fifth swimming legs exopod 1 sub-cylindrical; longer than wide; longer or equal than 2 times; with equal size between inner and outer sides; with convex inner side; with rectilinear outer side; without swelling; without marginal extension; without posterior process; without spine; without seta. Fifth swimming legs exopod 2 sub-cylindrical; longer than broad; longer or equal than 2 times; without swelling; without marginal extension; without process; without lobe; with spine; inserted laterally; rectilinear; without ornamentation; sharp tip; smaller than next segment; without seta. Fifth swimming legs exopod 3 cylindrical; longer than wide; without swelling; without process; without lobe; without spine; with seta; double; inserted terminally; unequal size between them; outer seta smaller than inner; nearly 1 time; outer seta not ornamented by setules; without ornamentation; presence of terminal claw; sclerotized; arched; externally directed; convex inner side; with ornamentation; of denticles; as a row; on surface partially; at medial region; concave outer side; with ornamentation; of denticles; as a row; on surface partially; at medial region; blunt tip; 6 times longer than origin segment.

##### Distribution records

###### BRAZIL

**Santa Catarina**. **Rio Grande do Sul**: rice flood water (this study). PARAGUAY. Several pools (this study); Yaciretá Dam Reservoir, River Parana between Argentina, and Paraguay (27° 24m 24s S 56° 15m 9s W) (this study). URUGUAY: **Barras Agas**; **Montevideo:** Barra Santa Lucía. ARGENTINA. **Buenos Aires**: “Balneário Norte de Nunez”.

##### Habitat

Habitat in freshwaters: floodplains, pools, rivers, and reservoirs.

##### Remarks

The taxonomic proposal of the species was defined from organisms from the region of Itajaí in Santa Catarina and received the name of *Diaptomus* s.l. *gibber* Popper 1889. Wright (1927) in a broad review of the “*Diaptomus*” (sensu lato) of South America portrayed the morphology of the organisms of the species and indicates it as the first diaptomid of the continent but did not include it in his original proposal and expansion of the *nordestinus* complex (Wright, 1935; 1936). Kiefer (1936) in taking Wright’s proposal as a basis and elevating other “*Diaptomus*” to his creation of *Notodiaptomus*, also disregarded *D. gibber*.

In 1938, Brehm when evaluating organisms from Uruguay indicated morphological reasons that could be related to disregard of Kiefer (1936). Although the male fifth right swimming leg, and female fourth prosome segment with dorsal protuberance of *D. gibber* was similar to *Notodiaptomus isabelae* (Wright 1936), the structure of male A1R differed from the species of the *nordestinus* complex and *Notodiaptomus* (Brehm, 1938). Formally, only in 1963, Pallares transferred the species and recombined it as *Notodiaptomus gibber* (Poppe, 1889), in an effort that added amendments to the morphology of the taxon organisms and extended its distribution to Argentina.

More recently, Paggi (2006) approached the morphology of the species and proposed synonymization with *Diaptomus meridionalis* Kiefer 1933. However, while the illustrations offered are consistent evidence to accept his proposal, these are unique to the female fifth swimming legs, genital double-somite, and last prosome segment of incomplete individuals or exuvia. It is crucial that other observations are made from specimens of the type series or complete topotypes of both species.

Within *Notodiaptomus*, the male fifth right swimming leg exopod 1 with digitiform internal prominence is a attribute shared between congeners of the species, such as *N. cannarensis* and *N. santafesinus.* Another convergent attribute present in *N. gibber* is female fourth metasome segment with dorsal protuberance in digitiform, 2x longer than wide, and inserted distally. Additionally, this protuberance has an anteriorly and posteriorly process, which we evidence as an exclusive condition in *Notodiaptomus*. This additional condition differentiates the species from the *N. inflatus*, *N. anisitsi*, *N. coniferoides*, and *N. paraenses*.

For the recharacterization of the male right antennule of the species was present aesthetasc absent on actual segment 5, and setae with sharp apex on actual segments 3, 7, 9 and 14. To actual segment 13 was present spiniform modified seta with blunt apex (rounded), and on actual segment 14 the same spinous process verified in *N. anisitsi*, *N. isabelae*, and *N. santafesinus.* Additionally, only three setae on current segment 20 are present, and hyaline lamella with distal extension to the sequent segment proximally. Invariably, these are conditions divergent from the ornamental pattern of the male right antennule described in Santos-Silva *et al*. (1999) and corroborated in this thesis.

The representation of the male rostrum offered in Pallares (1963, fig. 17) insinuates the presence of ornamental elements (probably setules) present in the distal portion of the rostral filaments. In addition, in the intermediate region of the filamentous pair it is possible to observe two paired elements of similar length to the rostral filaments. Throughout our examinations it was not possible to observe any of the conditions. For the intermediate elements to the rostral filaments, the author present the ventral junction of the filaments to the integument, probably an optical effect induced by examination under the ventral plane of view only. In addition, we bring here new observations for the rostrum of individuals of this species shared by their congeners, such as the presence of a basal rounded process positioned to the left ventrally and a smaller process to the right side posteriorly. This, in fact, is a variable condition within *Notodiaptomus*.

For the fifth male leg, it was later possible to observe the right coxa with sensilla inserted on a rounded process, notably smaller than that present in other species of *Notodiaptomus*. In the same observation plane, all specimens examined had uniform integumentary expansion along the inner margin of the right base, overlapped to the sequential segment proximally. This integumentary expansion partially covers the inner digitiform expansion of exopod 1 and the rudimentary endopod positioned antero-internally in the masculine leg completely. This present condition is distinct from the other examined *Notodiaptomus* in this effort, and observation of Paggi (2006) about the structural similarity of the right endopod of *N. gibber*, and *N. isabelae* could not be corroborated. The endopod of the first species was present throughout our examinations with a discrete protrusion marked by the presence of a spinules row and seta proximally.

In the same plane of observation, it was not possible to observe the male fifth left swimming leg coxa with posterior process characteristic of the *Notodiaptomus* species, the sensilla present on apex structure is inserted to the integument postero-externally directly. For the endopod of this leg it was possible to notice a feeble suture that defines the bisegmentation as an additional characteristic to the species. In the original illustrations of Poppe (1989), reproduced in Wright (1927), the left rami is schematically presented as a process indistinct from the basis and without any segmentation. In Pallares (1963) only the fifth right leg is presented and in Paggi (2006) it is only females of the species. Another additional feature is the final portion of the second exopod, which has an internally directed curved distal process, and with denticles in a row parallel to the axis of the segment anteriorly.

Undoubtedly, *N. gibber* is a species with considerable divergences in *Notodiaptomus*. From the morphological characteristics suggested in Wright (1935) for the *nordestinus* complex is present, in the species, the absence of the condition for male fifth left swimming leg endopod unisegmented. Among the attributes of Kiefer (1936; 1956) are absent: (1) female fifth swimming legs endopod unisegmented; (2) male fifth right swimming leg basis with posterior protrusion; (3) male fifth left swimming leg exopod 2 with spiniform “short” seta (here reinterpreted as not surpassing to distal-point segment); and (4) male right antennule actual segment 14 without spinous process.

#### Notodiaptomus henseni (Dahl, 1894)

##### Synonymy

*Diaptomus henseni* Dahl, 1894: 11, 19, pl. 1, figs. 1–5, 5a; Giesbrecht & Schmeil, 1898: 78; Daday, 1905: 151, 152; Tollinger, 1911: 70; 272, 273, fig. E; Brian, 1926: 183; Pesta, 1927: 80; Wright, 1935: 214, 219, 220, 221, 222, 223, pl. 1, fig. 3; 1936a: 79; 1937: 76; 1938b: 562; Kiefer, 1956: 242; Cipólli & Carvalho, 1973: 95, 97, 98, 100, 101, tab. 2; Reid, 1991: 737. *Diaptomus henseni*; Wright, 1927 (nec Dahl, 1894): 73–75, 96, pl. 8, figs. 7–11. *Notodiaptomus henseni*; Kiefer, 1936a: 197, fig. 7; Brehm, 1958a: 168; Brandorff, 1972: 44; 1976: 616, fig. 2; Andrade & Brandorff, 1975: 97; Löffler, 1981: 15; Dussart & Defaye, 1983: 134; Dussart, 1984a: 34, 39, 43, 46, fig. 3; Robertson & Hardy, 1984: 346, tab. 3; Matsumura-Tundisi, 1986: 542, figs. 81–85; Reid & Turner, 1988: 492; Cicchino *et al*., 1989: 98–105, figs. 1a-f, 2, 3, 4, 5; Cicchino, 1994: 145, fig. 6; Zoppi de Roa, 1994: 1384–1386, tab. 1; Rocha *et al*., 1995: 156; Santos-Silva, 1998: 209; Santos-Silva *et al*., 1999: 127; Santos-Silva *et al*., 2015: 21–25, figs. 9–11, identification key to male, and female, Perbiche-Neves *et al*., 2015: 56–64, figs. 47–52, and 54; Perbiche-Neves *et al*., 2020: 696-697, key to the Neotropical diaptomid, fig. 21.15 F; Geraldes-Primeiro *et al*., 2021: 2–3. *Notodiaptomus (Notodiaptomus) henseni*; Dussart, 1985a: 208.

##### Type locality

Although not clearly defined, the type locality is indicated in the original survey as the mouth of the Tocantins River, State of Pará (see Table I and Figure 32). For further details in Dahl (1894), it is possible to observe the indication of date X.5, depth of 35m, and water temperature of 28°C (table I). Probably this is associated with the original material and collection area.

##### Type material

Holotype not specified. Santos-Silva *et al*. (2015) found that the type-material designated by Dahl (263 males and 620 females) is probably lost. In the cited research, 1 male was designated as a neotype from the Igarapé Uruazinho, Maiauatá, the mouth of the Tocantins River, VIII.27.1970, sample 40. The material was deposited at MZUSP, and additional ones were deposited at INPA, NHM, and USNM.

##### Material examined

Topotype: 2 males, and 2 females (samples 40 and 42), from the Igarapé Uruazinho, Maiuatá Village, region of the mouth of the River Tocantins, Pará State, Brazil; 1 male, and 1 female (samples 44), from Igarapé Bahia, Maiauatá Village, River Tocantins, Pará (Tocantins currently). All individuals entire in alcohol, collected on 27.VIII.1970, by M. N. Cipólli, and M. A. Juliano de Carvalho. Material stored in the zooplankton collection (copepods) from the Plankton Laboratory, INPA; 1 male (INPA-COP023, slides a-h) and 1 female (INPA-COP024, slides a-h) were selected to be dissection on eight slides each and deposited in the Zoological Collection of the INPA, Brazil. Additional-material examined: 01 male, and 1 female, entire in formalin solution (MZUSP 32934) from the Furnas Reservoir on the Grande River, Brazil, collected by G. Perbiche-Neves, and stored in MZUSP.

##### Diagnosis

**(1)** male fifth left swimming leg reaching first right exopod segment medially; **(2)** male fifth left swimming leg basis with inner protuberance doubly, ornamented with tubercles patch, granular minutely; **(3)** male fifth right swimming leg basis with inner protuberance proximally, ornamented with tubercles; **(4)** male fifth right swimming legs exopod 1 with rectilinear triangular process on distal surface posteriorly; **(5)** male fifth right swimming legs exopod 2 inner-distal sub-triangular process posteriorly; **(6)** female right epimeral plate with ornamentation of spinules patch on dorsal-ventral surface outerly; **(7)** female genital double-somite with lateral expansion greater right than left; **(8)** female genital double-somite with double post-genital process ventrally; **(9)** female second urosome segment with ventral fusion to anal segment; **(10)** female fifth swimming legs coxa with outer conical process not projecting over basal segment; **(11)** female fifth swimming legs exopod 3 with rectilinear terminal claw.

##### Redescription

###### MALE

Body 1197 micrometers excluding caudal setae. Male body smaller and slenderer than female. Nerve axons myelinated. Prosome 6-segmented; widest at first metasome segment; without one line of setules at posterior margin; without spinules at segments. Cephalosome anterior margin rounded; with dorsal suture; incomplete; separate from first metasome segment. First metasome segment without sensilla. Second metasome segment with sensilla; 2 dorsally; of equal size. Third metasome segment with sensillae; 4 laterally; of unequal size; non-ornamented posterior margin. Fourth metasome segment with sensillae; 2 dorsally; 4 laterally; of equal size; separated from the fifth metasome. Limit between fourth and fifth metasome segments without ornamentation. Fifth metasome segment with sensilla; 4 laterally; Fifth metasome segment equal size; Fifth metasome segment ornamented; Fifth metasome segment with spinules; Fifth metasome segment as a row; Fifth metasome segment double; Fifth metasome segment continuous; Fifth metasome segment dorsally; Fifth metasome segment without dorsal conical process; with epimeral plates. Epimeral plates symmetrical. Right epimeral plates reduced, as rounded distal corner segment limit; with sensilla; at the apex of projection; ornamented; with spinules; as a patch.

##### Urosome

5-segmented; Urosome 5 - free segments. Genital somite asymmetrical in dorsal view; with single aperture; located on left side; ventrolaterally on posterior rim; with sensillae; on both sides; one; at left lateral; posteriorly; one; at right rim; posteriorly; of equal size between then. Third urosome segment without spinules; without external seta. Fourth urosome segment without spinules; without sub-conical blunt dorsal-lateral process. Anal segment presence of dorsal sensillae; one on each side; medially inserted; presence of operculum; convex; covering the anal aperture fully. Caudal rami symmetrical; separated from anal segment; longer than wide; with setules; continuous on; inner side; each ramus bearing 6 caudal setae; 5 marginals; plumose; and 1 internal dorsally; straight; not reticulated main axis; outermost seta with outer spiniform process absent.

##### Appendices features

Rostrum symmetrical; separated from dorsal cephalic shield; by complete suture; sensillae present; one pair; anteriorly inserted on surface tegument; with rostral filament; double; paired; extended; into point; with basal process; in ventral view, rounded on left side; without a smaller basal expansion on the right side.

##### Antennules

Asymmetrical. **Right antennules**. Uniramous; right antennule surpassing to genital segment; right antennule extending beyond caudal rami.

Right antennule ancestral segment I and II separated. Ancestral segment II and III fused. Ancestral segment III and IV fused. Ancestral segment IV and V separated. Ancestral segment V and VI separated. Ancestral segment VI and VII separated. Ancestral segment VII and VIII separated. Ancestral segment VIII and IX separated. Ancestral segment IX and X separated. Ancestral segment X and XI separated. Ancestral segment XI and XII separated. Ancestral segment XII and XIII separated. Ancestral segment XIII and XIV separated. Ancestral segment XIV and XV separated. Ancestral segment XV and XVI separated. Ancestral segment XVI and XVII separated. Ancestral segment XVII and XVIII separated. Ancestral segment XVIII and XIX separated. Ancestral segment XIX and XX separated. Ancestral segment XX and XXI separated. Ancestral segment XXI and XXII fused. Ancestral segment XXII and XXIII fused. Ancestral segment XXIII and XXIV separated. Ancestral segment XXIV and XXV fused. Ancestral segment XXV and XXVI separated. Ancestral segment XXVI and XXVII separated. Ancestral segment XXVII and XXVIII fused.

Right antennule actual 22-segmented; geniculated; between the segment 18 and segment 19; with swollen and modified region; formed by 5 segments; between 13 and 17 segments. Actual segment 1 with seta; one element; straight; none larger than segment; without spinules; without vestigial seta; without conical seta; without modified seta; without spinous process; with aesthetasc; one element. Actual segment 2 with seta; three elements; of unequal size; straight; none larger than segment; without spinules; with vestigial seta; one element; without conical seta; without modified seta; without spinous process; with aesthetasc; one element. Actual segment 3 with seta; one element; one larger than segment; surpassing to distal margin; beyond three sequential segments; straight; blunt apex; without spinules; with vestigial seta; one element; without conical seta; without modified seta; without spinous process; with aesthetasc. Actual segment 4 with seta; one element; one larger than segment; surpassing to distal margin; straight; not beyond three sequential segments; without spinules; without vestigial seta; without conical seta; without modified seta; without spinous process; without aesthetasc. Actual segment 5 with seta; one element; straight; one larger than segment; surpassing to distal margin; not beyond three sequential segments; without spinules; with vestigial seta; one element; without conical seta; without modified seta; without spinous process; with aesthetasc; one element. Actual segment 6 with seta; one element; none larger than segment; straight; without spinules; without vestigial seta; without conical seta; without modified seta; without spinous process; without aesthetasc. Actual segment 7 with seta; one element; straight; one larger than segment; surpassing to distal margin; beyond three sequential segments; blunt apex; without spinules; without vestigial seta; without conical seta; without modified seta; without spinous process; with aesthetasc; one element. Actual segment 8 with seta; one element; straight; none larger than segment; without spinules; without vestigial seta; with conical seta; one element; not reaching to middle-point of the sequent segment; without modified seta; without spinous process; without aesthetasc. Actual segment 9 with seta; two elements; of unequal size; straight; one larger than segment; surpassing to distal margin; beyond three sequential segments; blunt apex; without spinules; without vestigial seta; without conical seta; without modified seta; without spinous process; with aesthetasc; one element. Actual segment 10 with seta; one element; straight; none larger than segment; without spinules; without vestigial seta; without conical seta; with modified seta; presenting blunt apex; slender form; surpassing to distal margin; beyond of the sequential segment; parallel to antennule direction; without spinous process; without aesthetasc. Actual segment 11 with seta; one element; straight; one larger than segment; surpassing to distal margin; not beyond three sequential segments; without spinules; without vestigial seta; without conical seta; with modified seta; slender form; presenting blunt apex; surpassing to distal margin; beyond of the sequential segment; parallel to antennule direction; shorter length than homologous of actual segment 13; without spinous process; without aesthetasc. Actual segment 12 with seta; one element; straight; one larger than segment; surpassing to distal margin; not beyond three sequential segments; without spinules; without vestigial seta; with conical seta; one element; not smaller than to segment 8; without modified seta; without spinous process; with aesthetasc; one element; absent internal perpendicular fission. Actual segment 13 with seta; one element; straight; one larger than segment; surpassing to distal margin; not beyond three sequential segments; without spinules; without vestigial seta; without conical seta; with modified seta; stout form; surpassing to distal margin; to the middle-point of the sequence segment; perpendicular to antennule direction; presenting bifid apex; without spinous process; with aesthetasc; one element. Actual segment 14 with seta; two elements; of unequal size; straight; one larger than segment; surpassing to distal margin; beyond three sequential segments; blunt apex; without spinules; without vestigial seta; without conical seta; without modified seta; without spinous process; with aesthetasc; one element. Actual segment 15 with seta; two elements; of unequal size; straight; not bifidform; none larger than segment; without spinules; without vestigial seta; without conical seta; without modified seta; with spinous process; on outer margin; surpassing distal margin; with aesthetasc; one element. Actual segment 16 with seta; two elements; of unequal size; plumose; one larger than segment; surpassing to distal margin; not beyond three sequential segments; not bifidform; without spinules; without vestigial seta; without conical seta; without modified seta; with spinous process; on outer margin; surpassing distal margin; unequal size to process on preceding segment; with aesthetasc; one element. Actual segment 17 with seta; two elements; of unequal size; straight; none larger than segment; bifidform; without spinules; without vestigial seta; without conical seta; with modified seta; one element; stout form; surpassing to distal margin; not beyond of the sequential segment; parallel to antennule direction; without spinous process; without aesthetasc. Actual segment 18 with seta; two elements; of equal size; straight; none larger than segment; without spinules; without vestigial seta; without conical seta; with modified seta; one element; stout form; surpassing distal margin; parallel to antennule direction; without spinous process; without aesthetasc. Actual segment 19 with seta; two elements; of unequal size; plumose; none larger than segment; without spinules; without vestigial seta; without conical seta; with modified seta; two elements; stout form; at least one bifid form; surpassing distal margin; parallel to antennule direction; without spinous process; with aesthetasc; one element. Actual segment 20 with seta; four elements; of unequal size; straight; one larger than segment; surpassing to distal margin; beyond three sequential segments; without spinules; without vestigial seta; without conical seta; without modified seta; without spinous process; without aesthetasc. Actual segment 21 with seta; two elements; of equal size; plumose; one larger than segment; surpassing to distal margin; greater 3x than original segment; without spinules; without vestigial seta; without conical seta; without modified seta; without spinous process; without aesthetasc. Actual segment 22 with seta; four elements; of equal size; one larger than segment; plumose; surpassing to distal margin; greater 3x than original segment; without spinules; without vestigial seta; without conical seta; without modified seta; without spinous process; with aesthetasc; one element.

##### Left antennules

Uniramous; Left antennule surpassing to prosome; Left antennule extending beyond caudal rami. Ancestral segment I and II separated. Ancestral segment II and III fused. Ancestral segment III and IV fused. Ancestral segment IV and V separated. Ancestral segment V and VI separated. Ancestral segment VI and VII separated. Ancestral segment VII and VIII separated. Ancestral segment VIII and IX separated. Ancestral segment IX and X separated. Ancestral segment X and XI separated. Ancestral segment XI and XII separated. Ancestral segment XII and XIII separated. Ancestral segment XIII and XIV separated. Ancestral segment XIV and XV separated. Ancestral segment XV and XVI separated. Ancestral segment XVI and XVII separated. Ancestral segment XVII and XVIII separated. Ancestral segment XVIII and XIX separated. Ancestral segment XIX and XX separated. Ancestral segment XX and XXI separated. Ancestral segment XXI and XXII separated. Ancestral segment XXII and XXIII separated. Ancestral segment XXIII and XXIV separated. Ancestral segment XXIV and XXV separated. Ancestral segment XXV and XXVI separated. Ancestral segment XXVI and XXVII separated. Ancestral segment XXVII and XXVIII fused.

Left antennule actual 25-segmented; not-geniculated. Actual segment 1 with seta; one element; none larger than segment; straight; without spinules; without vestigial seta; without conical seta; without modified seta; without spinous process; with aesthetasc; one element. Actual segment 2 with seta; three elements; of equal size; none larger than segment; straight; without spinules; with vestigial seta; one element; without conical seta; without modified seta; without spinous process; with aesthetasc; one element. Actual segment 3 with seta; one element; one larger than segment; straight; surpassing to distal margin; beyond three sequential segments; without spinules; with vestigial seta; one element; without conical seta; without modified seta; without spinous process; with aesthetasc. Actual segment 4 with seta; one element; none larger than segment; straight; without spinules; without vestigial seta; without conical seta; without modified seta; without spinous process; without aesthetasc. Actual segment 5 with seta; one element; one larger than segment; straight; surpassing to distal margin; not beyond three sequential segments; without spinules; with vestigial seta; one element; without conical seta; without modified seta; without spinous process; with aesthetasc; one element. Actual segment 6 with seta; one element; none larger than segment; straight; without spinules; without vestigial seta; without conical seta; without modified seta; without spinous process; without aesthetasc. Actual segment 7 with seta; one element; one larger than segment; straight; surpassing to distal margin; beyond three sequential segments; without spinules; without vestigial seta; without conical seta; without modified seta; without spinous process; with aesthetasc; one element. Actual segment 8 with seta; one element; one larger than segment; straight; surpassing distal margin; without spinules; without vestigial seta; with conical seta; without modified seta; without spinous process; without aesthetasc. Actual segment 9 with seta; two elements; of unequal size; one larger than segment; straight; surpassing to distal margin; beyond three sequential segments; without spinules; without vestigial seta; without conical seta; without modified seta; without spinous process; with aesthetasc; one element. Actual segment 10 with seta; one element; none larger than segment; straight; without spinules; without vestigial seta; without conical seta; without modified seta; without spinous process; without aesthetasc. Actual segment 11 with seta; one element; one larger than segment; straight; surpassing to distal margin; beyond three sequential segments; without spinules; without vestigial seta; without conical seta; without modified seta; without spinous process; without aesthetasc. Actual segment 12 with seta; one element; one larger than segment; straight; surpassing distal margin; without spinules; without vestigial seta; with conical seta; without modified seta; without spinous process; with aesthetasc; one element. Actual segment 13 with seta; one element; none elongated; straight; surpassing distal margin; without spinules; without vestigial seta; without conical seta; without modified seta; without spinous process; without aesthetasc. Actual segment 14 with seta; one element; elongated; straight; surpassing to distal margin; beyond three sequential segments; without spinules; without vestigial seta; without conical seta; without modified seta; without spinous process; with aesthetasc; one element. Actual segment 15 with seta; one element; larger than segment; straight; surpassing to distal margin; not beyond three sequential segments; without spinules; without vestigial seta; without conical seta; without modified seta; without spinous process; without aesthetasc. Actual segment 16 with seta; one element; larger than segment; plumose; surpassing to distal margin; not beyond three sequential segments; without spinules; without vestigial seta; without conical seta; without modified seta; without spinous process; with aesthetasc; one element. Actual segment 17 with seta; one element; not larger than segment; straight; without spinules; without vestigial seta; without conical seta; without modified seta; without spinous process; without aesthetasc. Actual segment 18 with seta; one element; larger than segment; straight; surpassing to distal margin; beyond three sequential segments; without spinules; without vestigial seta; without conical seta; without modified seta; without spinous process; without aesthetasc. Actual segment 19 with seta; one element; not larger than segment; straight; surpassing distal margin; without spinules; without vestigial seta; without conical seta; without modified seta; without spinous process; with aesthetasc; one element. Actual segment 20 with seta; one element; not larger than segment; straight; surpassing distal margin; without spinules; without vestigial seta; without conical seta; without modified seta; without spinous process; without aesthetasc. Actual segment 21 with seta; one element; larger than segment; plumose; surpassing to distal margin; beyond three sequential segments; without spinules; without vestigial seta; without conical seta; without modified seta; without spinous process; without aesthetasc. Actual segment 22 with seta; two elements; of unequal size; one of them elongated; plumose; surpassing to distal margin; without spinules; without vestigial seta; without conical seta; without modified seta; without spinous process; without aesthetasc. Actual segment 23 with seta; two elements; of unequal size; one larger than segment; plumose; surpassing to distal margin; greater 3x than original segment; without spinules; without vestigial seta; without conical seta; without modified seta; without spinous process; without aesthetasc. Actual segment 24 with seta; two elements; of equal size; one larger than segment; plumose; surpassing to distal margin; greater 3x than original segment; without spinules; without vestigial seta; without conical seta; without modified seta; without spinous process; without aesthetasc. Actual segment 25 with seta; four elements; of equal size; elongated; plumose; surpassing to distal margin; 4 times larger than segment; without spinules; without vestigial seta; without conical seta; without modified seta; without spinous process; with aesthetasc; one element.

##### Antenna

Biramous. Antenna coxa separated from the basis; bearing seta; 1; on inner surface; at distal corner; reaching to the endopod 1. Antenna basis (fusion) separated from the endopodal segment; bearing seta; 2; on inner surface; at distal corner. Endopodal ancestral segment I and II separated. Ancestral segment II and III fused. Ancestral segment III and IV fused. Ancestral segment III and IV fully. Antenna endopod actual 2-segmented. Actual segment 1 not bilobate; with seta; two; on inner margin; with spinules; as a row; obliquely; on outer surface; with pore. Actual segment 2 bilobate; with discontinuity on outer cuticle; not developed as a suture; inner lobe bearing 8 setae; distally; outer lobe bearing 7 setae; distally; with spinules; as a patch; on outer surface. Antenna exopod ancestral segment I and II separated. Ancestral segment II and III fused.

Ancestral segment III and IV fused. Ancestral segment IV and V separated. Ancestral segment V and VI separated. Ancestral segment VI and VII separated. Ancestral segment VII and VIII separated. Ancestral segment VIII and IX separated. Ancestral segment IX and X fused. Antenna exopod actual 7-segmented. Actual segment 1 single; elongated (width-length, equal or larger ratio 2:1); with seta; one; at inner surface. Actual segment 2 compound; elongated (larger width-length ratio 2:1); with seta; three; at inner surface. Actual segment 3 single; not elongated (lesser width-length ratio 2:1); with seta; one; at inner surface. Actual segment 4 single; not elongated (lesser width-length ratio 2:1); with seta; one; at inner surface. Actual segment 5 single; not elongated (lesser width-length ratio 2:1); with seta; one; at inner surface. Actual segment 6 single; not elongated (lesser width-length ratio 2:1); with seta; one; at inner surface. Actual segment 7 compound; elongated (larger or equal width-length ratio 2:1); with seta; one; at inner surface; and three; at distal surface.

##### Oral features

**Mandible**. Coxal gnathobase sclerotized; with lobe; prominent; on caudal margin; presence of cutting blade; with tooth-like prominence; two, distinctly; 1 acute; on caudal margin; and 1 triangular; on sub-caudal margin; without acute projection between the prominences; with additional spinules; as a row; on dorsal surface; with seta; 1; dorsally; on apical surface; with spinules; apicalmost. Mandible palps biramous; comprising the basis; with seta; four; differently inserted; first medially; reaching to beyond the endopod 1; second distally; third distally; fourth distally; on inner margin; none with setulose ornamentation. Mandible endopod 2-segmented. Mandible endopod 1 with lobe; bearing seta; four; distally inserted; without spinules. Mandible endopod 2 without lobe; bearing setae; nine elements; distally inserted; with spinules; as a row; double. Mandible exopod 4-segmented. Mandible exopod 1 with seta; one element; distally; on inner margin. Mandible exopod 2 with seta; one element; distally; on inner side. Mandible exopod 3 with seta; one element; distally; on inner side. Mandible exopod 4 with setae; three elements; on terminal region. **Maxillule**. Birramous. Maxillule 3-segmented. Maxillule praecoxa with praecoxal arthrite; bearing spines; fifteen elements; ten marginally; plus, five sub-marginally; with spinules; as a patch; on sub-marginal surface. Maxillule coxa with coxal epipodite; with conspicuous outer lobe; bearing setae; nine elements; with coxal endite; elongated (larger or equal width-length ratio 2:1); bearing setae; four elements. Maxillule basis with basal endite; double; first proximal; elongated (larger width-length ratio 2:1; separated from basis; with setae; four elements; distally inserted; second distal; fused to basis; not elongated (lesser width-length ratio 2:1); with setae; four elements; distally inserted; with setules; as a row; on inner side; basal exite present; with setae; one element; on outer surface. Maxillule endopod 1-segmented. Endopod 1 bilobate; first proximal; with setae; three elements; second distal; with setae; five elements. Maxillule exopod 1-segmented. Exopod 1 with setae; six elements; with setules; as a row; on inner side; spinules absent. **Maxilla**. Uniramous. Maxilla 5-segmented. Maxilla praecoxa fused to coxa; incompletely; distinct externally; with praecoxal endite; double; first elongated endite (larger or equal width length ratio 2:1); proximally inserted; with seta; straight, or plumose; 1 straight; 4 plumose; with spine; single; without spinules; without setule; second elongated endite (larger or equal width length ratio 2:1); distally inserted; with seta; plumose; 3 plumose; without spine; with spinules; as a row; on distal margin; with setule; as a row; on distal margin; absence of outer seta. Maxilla coxa with coxal endite; double; first elongated endite (larger or equal width); proximally inserted; with seta; plumose; 3 plumose; without spine; without spinules; with setules; as a row; on proximal margin; second elongated endite (larger or equal width); distally inserted; with seta; plumose; 3 plumose; without spine; without spinules; with setules; as a row; on proximal margin; absence of outer seta. Maxilla basis with basal endite; single; elongated (larger or equal width-length ratio 2:1); with seta; plumose; 3 plumose; without spinules; absence of outer seta. Maxilla endopod 2-segmented. Endopod 1 with seta; 2 plumose; without spine; without spinules; without setules. Maxilla endopod 2 with seta; 2 plumose; without spine; without spinules; without setules. **Maxilliped**. Uniramous; Maxilliped 8-segmented. Maxilliped praecoxa fused to coxa; incompletely; distinct internally; with praecoxal endite; not elongated (lesser width-length ratio 2:1); distally inserted; with seta; 1 straight; with spinules; as a row; single; on basal surface; without setules. Maxilliped coxa with coxal endite; three coxal endite; first elongated (larger or equal width); proximally inserted; with seta; 2 plumose; with spinules; as a patch; single; on apical surface; without setules; second not elongated (lesser width-length ratio 2:1); medially inserted; with seta; 3 plumose; with spinules; as a row; single; on medial surface; without setules; third elongated (larger or equal width length ratio 2:1); distally inserted; with seta; 3 plumose; none reaching to beyond of the basis; with spinules; as a row; single; on basal surface; without setules; with lobe; prominence; at inner distal angle; ornamented; with spinules; continuously on margin. Maxilliped basis without basal endite; with seta; 3 plumose; with spinules; as a row; single; on medial surface; with setules; as a row; single; on inner margin. Maxilliped endopod segment 6-segmented. Endopod 1 with seta; 2 plumose; on inner surface. Endopod 2 with seta; 3 plumose; on inner surface. Endopod 3 with seta; 2 plumose; on inner surface. Endopod 4 with seta; 2 plumose; on inner surface. Endopod 5 with seta; 2 plumose; on inner surface, or on outer surface; outer seta absent. Endopod 6 with seta; 4 plumose; on inner surface, or on outer surface.

##### Swimming legs features

**First swimming legs.** Symmetrical; biramous. First swimming legs intercoxal plate without seta. First swimming legs praecoxa absent. First swimming legs coxa with seta; one; straight; distally inserted; on inner surface; surpassing to first endopodal segment; with setules; two group; as a patch; on inner margin; and as a row; double; on anterior surface; outerly; without spinules; without spine. First swimming legs basis without seta; with setules; as a patch; single; on outer surface; without spinules; without spine. First swimming legs endopod 2-segmented. Endopod 1 with seta; straight; restricted; to inner surface; one element; without spine; with setules; as a row; single; continuously; on outer surface; without spinules; absence of Schmeil’s organ. Endopod 2 with seta; unrestricted; three on inner surface; one on outer surface; two on distal surface; straight; without spine; with setules; as a row; single; continuously; on outer surface; without spinules; absence of Schmeil’s organ. Endopod 3 absence. First swimming legs exopod 1 with seta; restricted; 1 on inner surface; with spine; 1; stout; smaller than original segment; serrated; on inner side; continuously; with setules; as a row; single; as a row; innerly. First swimming legs exopod 2 with seta; restricted; 1 on inner surface; straight; without spine; with setules; as a row; single; continuously; on inner margin, or on outer margin; without spinules. First swimming legs exopod 3 with setule; as a row; single; continuously; on outer surface; without spinules; with seta; unrestricted; 2 on inner surface; 2 on terminal surface; with spine; 2; unequal size; first no longer 2x than origin segment; stout; serrated; on inner side, or on outer side; equally; second longer 3x than origin segment; slender; serrated; on outer side; with ornamentation on non-serrated side; by setules. **Second swimming legs**. Symmetrical; Second swimming legs biramous. Second swimming legs intercoxal plate without seta. Second swimming legs praecoxa present; located laterally. Second swimming legs coxa with seta; straight; distally inserted; on inner surface; surpassing to basal segment; without setules; without spinules; without spine. Second swimming legs basis without seta; without setules; without spinules; without spine. Second swimming legs endopod 3-segmented. Endopod 1 with seta; straight; restricted; one on inner surface; without spine; with setules; as a row; single; continuously; on outer surface; without spinules; absence of Schmeil’s organ. Endopod 2 with seta; straight; unrestricted; two on inner surface; without spine; with setules; as a row; single; continuously; on outer side; without spinules; presence of Schmeil’s organ; on posterior surface. Endopod 3 with seta; straight; unrestricted; three on inner surface; two on outer surface; two on distal surface; without spine; without setules; with spinules; as a row; double; distally inserted; at anterior surface; absence of Schmeil’s organ. Second swimming legs exopod 1 with seta; restricted; one on inner surface; with spine; 1; stout; not reaching to distal-third of the exopod 2; serrated; on inner side, or on outer side; with setules; as a row; single; continuously; on inner side; without spinules; absence of Schmeil’s organ. Exopod 2 with seta; unrestricted; one on inner surface; with spine; 1; stout; not surpassing the exopod 3; serrated; on inner side, or on outer side; with setules; as a row; single; continuously; on inner surface; without spinules; absence of Schmeil’s organ. Exopod 3 with seta; plurimarginal; three on inner surface; two on terminal surface; with spine; 2; unequal size; first no longer 2x than origin segment; stout; serrated; on inner side, or on outer side; equally; second longer 2x than origin segment; slender; serrated; on outer side; with ornamentation on non-serrated side; of setules; setules on outer surface; as a row; single; continuously; on inner surface; with spinules; as a row; single; distally inserted; at anterior surface; absence of Schmeil’s organ. **Third swimming legs**. Symmetrical; Third swimming legs biramous. Third swimming legs intercoxal plate without seta. Third swimming legs praecoxa present; not laterally located. Third swimming legs coxa with seta; straight; distally inserted; on inner surface; surpassing to first endopodal segment; without setules; without spinules; without spine. Third swimming legs basis without seta; without setules; without spinules; without spine. Third swimming legs endopod 3-segmented. Endopod 1 with seta; restricted; one on inner surface; without spine; without setules; without spinules; absence of Schmeil’s organ. Endopod 2 with seta; restricted; two on inner surface; straight; without spine; without setules; without spinules; absence of Schmeil’s organ. Endopod 3 with seta; straight; plurimarginal; two on inner surface; two on outer surface; three on terminal surface; without spine; without setules; with spinules; as a row; distally inserted; double; at anterior surface; absence of Schmeil’s organ. Third swimming legs exopod 1 with seta; restricted; straight; one on inner surface; with spine; 1; stout; not reaching to the distal-third of the exopod 2; serrated; equally; on inner surface, or on outer surface; with setules; as a row; single; continuously; on inner surface; without spinules; absence of Schmeil’s organ. Exopod 2 with seta; straight; restricted; one on inner surface; with spine; 1; stout; not reaching out to exopod 3; serrated; on inner side, or on outer side; equally; with setules; as a row; single; continuously; on inner side; without spinules; absence of Schmeil’s organ. Exopod 3 without setules; with spinules; as a row; single; distally inserted; at anterior surface; with seta; straight; unrestricted; three on inner surface; two on terminal surface; with spine; 2; unequal size; first no longer 2x than origin segment; stout; serrated; on inner side, or on outer side; equally; second longer 2x than origin segment; slender; serrated; on outer side; with ornamentation on non-serrated side; of setules; absence of Schmeil’s organ. **Fourth swimming legs**. Symmetrical; biramous. Intercoxal plate without sensilla. Praecoxa present. Coxa with seta; distally inserted; on inner margin; reaching out to endopod 1; without spinules; setules absent. Basis with seta; one; medially inserted; on posterior surface; smaller than the original segment; without setules; without spinules; without spine. Fourth swimming legs endopod 3-segmented. Endopod 1 with seta; one; restricted; on inner surface; without spine; without setules; without spinules; absence of Schmeil’s organ. Endopod 2 with seta; restricted; two on inner side; without spine; with setules; as a row; single; continuously; on outer surface; without spinules; absence of Schmeil’s organ. Endopod 3 with seta; unrestricted; two on inner surface; two on outer surface; three on distal surface; without spine; without setules; with spinules; as a row; double; distally inserted; at anterior surface; absence of Schmeil’s organ. Fourth swimming legs exopod 1 with seta; restricted; one on inner surface; with spine; 1; stout; not reaching out to distal-third of the exopod 2; serrated; on inner side, or on outer side; equally; with setules; as a row; single; continuously; on inner surface; without spinules; absence of Schmeil’s organ. Exopod 2 with seta; restricted; one on inner surface; with spine; 1; stout; not reaching the end of exopod 3; serrated; on inner side, or on outer side; equally; with setules; as a row; single; continuously; on inner surface; without spinules; absence of Schmeil’s organ. Exopod 3 without setules; with spinules; as a row; single; distally inserted; at anterior surface; with seta; unrestricted; three on inner surface; two on distal surface; with spine; 2; unequal size; first no longer 2x than origin segment; stout; serrated; on inner side, or on outer side; equally; second longer 2x than origin segment; slender; serrated; on outer side; without ornamentation on non-serrated side; absence of Schmeil’s organ.

##### Fifth swimming legs features

Asymmetrical. Fifth swimming leg intercoxal plate with length not equal or greater than width on 1.5x; with irregular proximal margin; discontinuous to; the anterior margin of the left coxa, or the anterior margin of the right coxa; posterior sensilla on the right lateral absent. **Fifth left swimming leg**. Fifth left swimming leg biramous; leg reaching first right exopod segment; medially. Fifth left swimming leg praecoxa present; rudimentary; separated from the coxae; without ornamentation. Fifth left swimming leg coxa concave inner side; without teeth-like structures; with process; conical; on posterior surface; outer side; distally inserted; not projecting over basis; with sensilla; stout; triangular; at apex; no longer 2x than insertion basis; without swelling; without seta; without spinules. Fifth left swimming leg basis sub-cylindrical; unequal size between inner and outer side; shorter outer than inner side; with concave inner side; rounded internal proximal expansion absent; without outgrowth; with groove; deep; obliquely; on posterior surface; not reaching the endopodal lobe; not ornamented; presence of protuberance; double; semicircular; on inner margin; ornamented; with tubercles; as a patch; minutely granular; covering the element; until to adjacent surface; with seta; outerly inserted; no longer 2x than origin segment; absence of minutely granular. Fifth left swimming leg endopod segments 1 and 2 fused; segments 2 and 3 fused; 1-segmented; stout; separated from the basis; ornamented; on inner side; with spinules; more than four elements; as a row; terminally; row of setules absent; without seta. Fifth left swimming leg exopod segments 1 and 2 separated; segments 2 and 3 fused; 2-segmented; stout; separated from the basis. Fifth left swimming leg exopod 1 sub-triangular; longer than broad; equal size between inner and outer side; convex inner side; convex outer side; without swelling; without marginal extension; with process; double; proximal; and other distal; posteriorly; with lobe; single; semicircular; medially inserted; on inner side; covered; by setules; without outer spine; absence seta. Fifth left swimming leg exopod 2 digitiform; longer than broad; equal size between inner and outer side; disform inner side; with rectilinear outer side; setulose pad present; prominently rounded; medially; on inner side; inflated medial region absent; distal process present; digitiform; non denticulate; without transverse row of denticles; none oblique row of 5 denticles; innerly directed; with seta; spiniform; ornamented by spinules; not surpassing the distal-point of the segment; without outer spine; terminal claw absent.

##### Fifth right swimming leg

Biramous. Fifth right swimming leg praecoxa present; separated from the coxae; without ornamentation. Fifth right swimming leg coxa convex inner side; without teeth-like structures; with process; conical; distally inserted; on posterior surface; closest to the outer rim; projecting over basis; beyond the first third; until the medial surface; without triangular protuberance innerly; with sensilla; slender; at apex; no longer 2x than basal insertion; without marginal extension; without seta; without spinules. Fifth right swimming leg basis cylindrical; unequal size between inner and outer side; shorter outer than inner side; rectilinear inner side; tumescence present; not inflated; restricted on inner surface; proximally; with protuberance; single; semicircular; on inner side; proximally; ornamented; with tubercles; as a patch; minutely granular; covering the element; until to adjacent surface; absence of distinct minutely granular; additional inner process absent; with posterior groove; deep; obliquely; reaching the endopodal lobe; ornamented; with tubercles; throughout of the outer border; with seta; outerly inserted; on anterior surface; no longer 2x than origin segment; posterior protrusion present; distal process absent. Fifth right swimming leg with endopodite present; separated from the basis; on anterior surface; ancestral segments 1 and 2 fused; ancestral segments 2 and 3 fused; 1-segmented; stout; ornamented; with setules; as a row; on inner side; terminally; without seta. Fifth right swimming leg exopod segments 1 and 2 separated; segments 2 and 3 fused; 2-segmented; stout; separated from the basis. Fifth right swimming leg exopod 1 trapezium; longer than broad; nearly 1.25 times; unequal size between both sides; shorter inner than outer side; convex inner side; rectilinear outer side; with marginal extension; sub-triangular; distally inserted; at outer rim; spinules absent; with process; triangular; rectilinear; blunt tip; sclerotized; without ornamentation; distally inserted; at posterior surface; projecting over next segment; without outer spine; without seta; internal prominence absent; lamella on posterior surface absent. Fifth right swimming leg exopod 2 cylindrical; longer than broad; nearly 2 times; equal size between both sides; disform inner side; convex outer side; without posterior proximal swelling; inner-posterior process present; sub-triangular; distally; without marginal expansion; curved ridge on distal posterior surface absent; chitinous knobs absent; with outer spine; inserted sub-distally; arched; externally directed; ornamented innerly; by spinules; as a row; not ornamented outerly; sharp tip; without apparent curve; lesser than the length of the exopod 2; until to 2 times its size; 2x; sensilla absent; terminal claw present; equal or longer 1.5 times than insertion segment; sclerotized; arched; inward; with conspicuous curve; proximally; ornamented innerly; by spinules; as a row; partially on extension; medially, or distally; not ornamented outerly; sharp tip; not curved tip; without medial constriction; hyaline process absent.

##### FEMALE

Body longer and wider than male; Female body 1331 micrometers excluding caudal setae. Widest at first metasome segment. Distal margin of the prosomal segments without one line of setules at posterior margin. Prosome segments with spinules at least at one prosomal segment. Fourth metasome segment absence of dorsal protuberance. Fourth and fifth metasome segments fused; partially; on dorsal surface. Limit between fourth and fifth metasome segments without ornamentation. **Fifth metasome segment**. Fifth metasome segment with sensilla; dorsally; 2 elements; with epimeral plates. Epimeral plates symmetrical. Right epimeral plates prominent, as projections; one posterior-laterally directed; not reaching half length of the genital segment; with sensilla at the apex; dorsal-posterior sensilla present; slender; ornamented; with spinules; as a patch; outerly; on dorsal surface, or on ventral surface. Left epimeral plate without expansion.

##### Urosome

3-segmented. **Genital double-somite**. Asymmetrical in dorsal view; longer than broad; longer than other urosomites combined; dorsal suture at mid-length absent; not covered by spinules; with swelling; rounded; unequal size; greater right than left; anteriorly; with sensillae; on both sides; one; stout; with robust apex; at left lateral; not on lobular base; anteriorly; one; stout; at right lateral; not on lobular base; anteriorly; with robust apex; of equal size between then; lateral protuberance absent; with right posterior rim expanded; over next segment; without slender sensilla on each posterior rim; without posterior-dorsal process. Genital double-somite opercular pad present; broader than longer; symmetrical; development laterally; expanded posteriorly; covering partially; double gonoporal slit; located ventrally; with arthrodial membrane; inserted anteriorly; post-genital process present; double; disto-ventral tumescence absent; ventral vertical folds absent; dorsal sensilla absent. Second urosome segment with ventral fusion to anal segment; right distal process absent. Caudal rami patch of setules on outer surface absent; patch of spinules on outer surface present.

##### Oral appendices feature

Rostrum basal process absent. **Antennules**. Symmetrical. Right antennule surpassing to genital double-segment; extending beyond caudal rami. Right antennule not exceeding the caudal setae. Right antennule ornamentation pattern equals to male left antennule; fully.

##### Fifth swimming legs

Symmetrical; Fifth swimming legs biramous. Fifth swimming legs intercoxal plate longer than wide; separated from the legs. Fifth swimming legs praecoxa with sclerite praecoxal; separated from the coxae; without ornamentation. Fifth swimming legs coxa with process; conical; at the outer rim; distally; sensilla present; stout; at apex; not projecting over basal segment; no longer 2x than basal insertion; marginal extension absent; without swelling; without seta; without spinules. Fifth swimming legs basis sub-triangular; unequal size between inner and outer sides; shorter outer than inner side; with convex inner side; without proximal inner outgrowth; without groove; with distal extension; on posterior surface; with seta; outerly inserted; on anterior surface; longer 2x than origin segment; not reaching to exopod 1 distally. Fifth swimming legs endopod segments 1 and 2 fused; segments 2 and 3 fused; 1-segmented; stout; separated from the basis; present discontinuity cuticle; on inner side; with spinules; as a row; single; non-oblique; sub-terminally; at anterior surface; with seta; double; one medially; on posterior surface; rectilinear; one distally; on posterior surface; arched; of unequal size; distal seta longer than medial seta. Fifth swimming legs exopod segments 1 and 2 separated; segments 2 and 3 separated; 3-segmented; separated from the basis. Fifth swimming legs exopod 1 sub-cylindrical; longer than wide; longer or equal than 2 times; with unequal size between inner and outer side; shorter inner than outer side; with convex inner side; with rectilinear outer side; without swelling; without marginal extension; with posterior process; distally; without spine; without seta. Fifth swimming legs exopod 2 sub-cylindrical; longer than broad; longer or equal than 2 times; without swelling; without marginal extension; without process; without lobe; with spine; inserted laterally; rectilinear; without ornamentation; sharp tip; equal size or larger than next segment; without seta. Fifth swimming legs exopod 3 cylindrical; longer than wide; without swelling; without process; without lobe; without spine; with seta; double; inserted terminally; unequal size between them; outer seta smaller than inner; nearly 3 times; outer seta not ornamented by setules; without ornamentation; presence of terminal claw; sclerotized; rectilinear; with ornamentation; of denticles; as a row; on surface partially; at medial region; rectilinear outer side; with ornamentation; of denticles; as a row; on surface partially; at medial region; blunt tip; 6 times longer than origin segment.

##### Distribution records

###### COLOMBIA

Guaviare River (Cicchino *et al*., 1989). VENEZUELA. **Apure**: Mantecal wetland savanna, 07°35’N and 69°10’W (Cicchino *et al*., 1989; Zoppi de Roa, 1994). **Carabobo:** Lake Valencia (Cicchino *et al*., 1989). **Delta Amacuro**: Caño Manamo (Cicchino *et al*., 1989; Dussart, 1984a); **Caño Guara**, near to Tucupita, Orinoco Delta (Dussart, 1984a). **Guárico**: Portuguese River (Cicchino *et al*., 1989). BRAZIL. **Amazonas**: Balbina reservoir, Uatumã river (Santos-Silva *et al*., 2015). **Pará**: mouth of the Tocantins River (Dahl, 1894); all the records listed below were made by Cipólli & Carvalho (1973: tabs. 2, 4) from the Guamá, Capim and Tocantins Rivers: Marajó Bay; Ariacana, Capim River; flooded area near to Timbiras, Caranandeua Lake; Timbiras Lake, Caranandeua; Maria Preta Lake, Capim River; Jurumundeua Lake, Caranandeua; Bernardino Lake, Santana do Capim; Uruazinho stream, Maiauatá; Jacarequara stream, Abaetetuba; São Lourenço river, Panaquera borehole; Inó stream, Panaquera hole; Coelho stream, Maratapá Bay; Pindobal River, Maratapá Bay; Grilo creek, Maratapá Bay; paraná Samuuma, Maratapá Bay; Mapará stream, Paraná Samuuma; Tocantins River, Cametá; Maloca stream, Cametá; Aricurá stream, Cametá; Espírito Santo stream, Baião; Murú stream; Tocantins River, Tucuruí; marginal lake to the Tocantins River, Jatobal; lagoon, Tucuruí; Trocará Lake between Tucuruí and Baião. **Maranhão**: José Maria Lake, Mearim River (Matsumura-Tundisi, 1986); **São Paulo**: Furnas Reservoir on the Grande River (Perbiche-Neves *et al*., 2015).

##### Habitat

Habitat in freshwaters: river mounth-lakes.

##### Remarks

This was a species described from organisms from the Tocantins River nonspecifically. Dahl provided a summary morphological description and poor illustrations of relevant structures (*e.g.*, male, and female fifth swimming legs, and female genital double-somite). The species was extensively reviewed in the efforts of Santos-Silva *et al*. (2015) and Perbiche-Neves *et al*. (2015) and offered comments on the taxonomic trajectory and important morphological amendments.

In the present approach, *N. henseni* possesses all the characteristics described by Wright (1935; 1936; 1937) for the *nordestinus* complex, and Kiefer (1936; 1956) for *Notodiaptomus*, except for the male fifth right swimming leg exopod 2 with absence of angular or rounded prominence on inner margin distally (in the present effort as curved ridge on distal posterior surface). Among the characters present in the species and not converging with the type of the genus are: (1) male fifth left swimming leg reaching first right exopod segment medially; (2) male fifth left swimming leg exopod 1 with posterior double process proximally, and distally; (3) male fifth right swimming leg exopod 1 with posterior triangular process, rectilinear, and inserted distally; (4) female genital double-somite with post-genital pore process doubly; (5) female urosome segment with ventral fusion to anal segment; (6) female antennule without setae on actual segments 13, 15, 16, 17, 18, and absence aesthetasc on 14.

#### Notodiaptomus iheringi (Wright, 1935)

##### Synonymy

*Diaptomus iheringi* Wright, 1935: 214, 219, 221, 223, 226, 229, pl. 1, fig. 4, pl. 2, figs. 3, 5–11;1936a: 80, 81; 1937: 76; 1938a: 300; 1938b: 562; Brehm, 1958a: 140, 146, 168; 1960: 49; Cipólli & Carvalho, 1973: 95, 97, 98, 101, tab. 2; Reid, 1991: 738, 740. *Notodiaptomus iheringi*; Kiefer, 1936a: 197, figs. 3, 4; 1956: 242; Brandorff, 1972: 44; 1976: 616, 621, fig. 2; Löffler, 1981: 15; Sendacz & Kubo, 1982: 54, 69– 71, 85–86, figs. 25–29, tab. 3; Dussart & Defaye, 1983: 137; Arcifa, 1984: 143, tab. 7; Reid & Esteves, 1984: 310, 311, 317, 321, 322, tab. 2; Robertson & Hardy, 1984: tab. 3; Reid, 1985: 574–579, 589, figs. 1–28; 1987: 378; 1991: 738, 740; Sendacz *et al*., 1985: 190, 193, 195, 196, 201, 203, 205, 207, tabs. 4, 6, 8,10, 12; Matsumura Tundisi, 1986: 542, 547, figs. 66–72, 100; Rocha *et al*., 1990: 94, tab. 5; Lansac-Tôha *et al*., 1992: 43, 45, 47; Tomm *et al*., 1992: 57, 58, 64, 67, 69; Sendacz, 1993: 35; 1997: 624, 625, tab. 2; Reid & Pinto-Coelho, 1994: 93, 95, 99, 100, 108; Tundisi & Matsumura-Tundisi, 1994: 27; Rocha *et al*., 1995: 155, 156; Lima *et al*., 1996: 115, fig. 3; Lansac-Tôha *et al*., 1997: 140, tab. 3; Santos-Silva, 1998: 209; Carvalho & Sendacz, 1998: 1525, 1527; Santos-Silva *et al*., 1999: 127. *Notodiaptomus* (*Wrightius*) *iheringi*; Dussart, 1985a: 210. *Notodiaptonus iheringi*; Rolla *et al*., 1990: 241, tab. 6 [*error*]; Santos-Silva, 2008: 28–29; Santos-Silva *et al*., 2015: 45–50, figs. 24–26, 40, identification keys to male and female; Perbiche-Neves *et al*., 2015: 64–69, figs. 55–60, identification keys to male and female; Perbiche-Neves *et al*., 2020: 696-697, key to the Neotropical diaptomid, fig. 21.15 G; Geraldes-Primeiro *et al*., 2021: 2.

##### Type locality

Puxinana Reservoir, Puxinana ville near to Campina Grande State, Paraiba, Brazil.

##### Type material

Holotype not specified. Reid (1991) indicates that the type-material is non-existent and topotypes in alcohol collected by S. Wright are deposited in alcohol at USNM (n° 79542), origin of the specimen designated by Santos-Silva *et al*. (2015) as a neotype of the taxon, but no indicated of the biological collection or depositary code.

##### Material examined

Topotype: 2 males, and 2 females, entire, in alcohol, no date, collected and designated by S. Wright, under code USNM 79542; 2 males, 3 females, entire, in alcohol, 05.XII.1934, collected by Lenz and belonging to Kiefer’s collection (LNK), sample 1053, stored in Plankton Laboratory, INPA; 1 male (INPA-COP025, slides a-h) and 1 female (INPA-COP026, slides a-h) were selected to be dissection on eight slides each and deposited in the Zoological Collection of the INPA, Brazil. Additional material examined: 2 males, and 2 females, entire, in alcohol from the Açude Novo, Campina Grande City, Paraíba State, Brazil, 07.VI.1934, belonging to Wright’s collection, and stored in Plankton Laboratory (LP19-A22), INPA, Brazil; 4 males (MZUSP 6192), 3 females (MZUSP 6192), and (MZUSP 6193) 4 juveniles, entire, in alcohol not were examined due precarious state of preservation present.

##### Diagnosis

**(1)** Prosome with spinules at posterior margins of the segments 3, 4, and 5; **(2)** male limit between fourth and fifth metasome ornamented with double spinules row dorsally; **(3)** male epimeral plates asymmetrical; **(4)** male right antennule with length extending beyond caudal rami; **(5)** male right antennule actual segment 1 with dorsal spinules; **(6)** male right antennule actual segment 8 with conical seta reaching to middle-point of the sequential segment; **(7)** male right antennule actual segment 12 with conical seta smaller than to those segment 8; **(8)** left antennule actual segment 1 with spinules row; **(9)** male fifth left swimming leg coxa with posterior outer process projecting over basis proximally; **(10)** male fifth left swimming leg exopod 2 with denticulate distal digitiform process anteriorly; **(11)** male fifth right swimming leg basis with groove reaching the endopodal lobe; **(12)** male fifth right swimming leg exopod 1 longer than broad 2x nearly; **(13)** male fifth right swimming leg exopod 1 with distal rectilinear triangular process posteriorly; **(14)** male fifth right swimming leg exopod 2 longer than broad 2.5x nearly; **(15)** male fifth right swimming leg exopod 2 with outer spine lesser 6x than original segment; **(16)** female fourth prosome segment with double spinules row dorsally; **(17)** female fifth swimming legs endopod without discontinuity cuticle; **(18)** female fifth swimming legs endopod with single seta; **(19)** female fifth swimming legs exopod 2 with lateral spine smaller than next segment.

##### Redescription

###### MALE

Body 1087 micrometers excluding caudal setae. Male body smaller and slenderer than female. Nerve axons myelinated. Prosome 6-segmented; widest at first metasome segment; without one line of setules at posterior margin; with spinules at least at one segment. Cephalosome anterior margin rounded; with dorsal suture; incomplete; separate from first metasome segment. First metasome segment without sensilla. Second metasome segment without sensilla. Third metasome segment without sensillae; ornamented posterior margin; with spinules; as a row; single; dorsally. Fourth metasome segment without sensillae; separated from the fifth metasome. Limit between fourth and fifth metasome segments ornamented; with spinules; as a row; on dorsal doubly; on lateral singly; same size. Fifth metasome segment without sensilla; ornamented; with spinules; as a row; single; discontinuous; dorsally. Fifth metasome segment without dorsal conical process; with epimeral plates. Epimeral plates asymmetrical. Right epimeral plates reduced, as rounded distal corner segment limit; with sensilla; at the apex of projection; without ornamentation. Left epimeral plate prominent, as projection; one projection; posterior-dorsally directed; reaching half length of the genital segment; with sensillae; at the apex of projection; without ornamentation.

##### Urosome

5-segmented; Urosome 5 - free segments. Genital somite asymmetrical in dorsal view; with single aperture; located on left side; ventrolaterally on posterior rim; with sensillae; on both sides; one; at left lateral; posteriorly; one; at right rim; posteriorly; of equal size between then. Third urosome segment without spinules; without external seta. Fourth urosome segment without spinules; without sub-conical blunt dorsal-lateral process. Anal segment absence of dorsal sensillae; presence of operculum; convex; not covering the anal aperture fully. Caudal rami symmetrical; separated from anal segment; longer than wide; with setules; continuous on; inner side; each ramus bearing 6 caudal setae; 5 marginals; plumose; and 1 internal dorsally; straight; not reticulated main axis; outermost seta with outer spiniform process absent.

##### Oral appendices feature

Rostrum symmetrical; separated from dorsal cephalic shield; by complete suture; sensillae present; one pair; anteriorly inserted on surface tegument; with rostral filament; double; paired; extended; into point; with basal process; in ventral view, rounded on left side; without a smaller basal expansion on the right side.

##### Antennules

Asymmetrical. **Right antennules**. Uniramous; right antennule surpassing to genital segment; right antennule extending beyond caudal rami.

Right antennule ancestral segment I and II separated. Ancestral segment II and III fused. Ancestral segment III and IV fused. Ancestral segment IV and V separated. Ancestral segment V and VI separated. Ancestral segment VI and VII separated. Ancestral segment VII and VIII separated. Ancestral segment VIII and IX separated. Ancestral segment IX and X separated. Ancestral segment X and XI separated. Ancestral segment XI and XII separated. Ancestral segment XII and XIII separated. Ancestral segment XIII and XIV separated. Ancestral segment XIV and XV separated. Ancestral segment XV and XVI separated. Ancestral segment XVI and XVII separated. Ancestral segment XVII and XVIII separated. Ancestral segment XVIII and XIX separated. Ancestral segment XIX and XX separated. Ancestral segment XX and XXI separated. Ancestral segment XXI and XXII fused. Ancestral segment XXII and XXIII fused. Ancestral segment XXIII and XXIV separated. Ancestral segment XXIV and XXV fused. Ancestral segment XXV and XXVI separated. Ancestral segment XXVI and XXVII separated. Ancestral segment XXVII and XXVIII fused.

Right antennule actual 22-segmented; geniculated; between the segment 18 and segment 19; with swollen and modified region; formed by 5 segments; between 13 and 17 segments. Actual segment 1 with seta; one element; straight; none larger than segment; with spinules; without vestigial seta; without conical seta; without modified seta; without spinous process; with aesthetasc; one element. Actual segment 2 with seta; three elements; of unequal size; straight; none larger than segment; without spinules; with vestigial seta; one element; without conical seta; without modified seta; without spinous process; with aesthetasc; one element. Actual segment 3 with seta; one element; one larger than segment; surpassing to distal margin; beyond three sequential segments; straight; blunt apex; without spinules; with vestigial seta; one element; without conical seta; without modified seta; without spinous process; with aesthetasc. Actual segment 4 with seta; one element; one larger than segment; surpassing to distal margin; straight; not beyond three sequential segments; without spinules; without vestigial seta; without conical seta; without modified seta; without spinous process; without aesthetasc. Actual segment 5 with seta; one element; straight; one larger than segment; surpassing to distal margin; not beyond three sequential segments; without spinules; with vestigial seta; one element; without conical seta; without modified seta; without spinous process; with aesthetasc; one element. Actual segment 6 with seta; one element; none larger than segment; straight; without spinules; without vestigial seta; without conical seta; without modified seta; without spinous process; without aesthetasc. Actual segment 7 with seta; one element; straight; one larger than segment; surpassing to distal margin; beyond three sequential segments; blunt apex; without spinules; without vestigial seta; without conical seta; without modified seta; without spinous process; with aesthetasc; one element. Actual segment 8 with seta; one element; straight; none larger than segment; without spinules; without vestigial seta; with conical seta; one element; reaching to middle-point of the sequent segment; without modified seta; without spinous process; without aesthetasc. Actual segment 9 with seta; two elements; of unequal size; straight; one larger than segment; surpassing to distal margin; beyond three sequential segments; blunt apex; without spinules; without vestigial seta; without conical seta; without modified seta; without spinous process; with aesthetasc; one element. Actual segment 10 with seta; one element; straight; none larger than segment; without spinules; without vestigial seta; without conical seta; with modified seta; presenting blunt apex; slender form; surpassing to distal margin; beyond of the sequential segment; parallel to antennule direction; without spinous process; without aesthetasc. Actual segment 11 with seta; one element; straight; one larger than segment; surpassing to distal margin; not beyond three sequential segments; without spinules; without vestigial seta; without conical seta; with modified seta; slender form; presenting blunt apex; surpassing to distal margin; beyond of the sequential segment; parallel to antennule direction; shorter length than homologous of actual segment 13; without spinous process; without aesthetasc. Actual segment 12 with seta; one element; straight; one larger than segment; surpassing to distal margin; not beyond three sequential segments; without spinules; without vestigial seta; with conical seta; one element; smaller than to segment 8; without modified seta; without spinous process; with aesthetasc; one element; absent internal perpendicular fission. Actual segment 13 with seta; one element; straight; one larger than segment; surpassing to distal margin; not beyond three sequential segments; without spinules; without vestigial seta; without conical seta; with modified seta; stout form; surpassing to distal margin; to the middle-point of the sequence segment; parallel to antennule direction; presenting bifid apex; without spinous process; with aesthetasc; one element. Actual segment 14 with seta; two elements; of unequal size; straight; one larger than segment; surpassing to distal margin; beyond three sequential segments; blunt apex; without spinules; without vestigial seta; without conical seta; without modified seta; without spinous process; with aesthetasc; one element. Actual segment 15 with seta; two elements; of unequal size; straight; not bifidform; none larger than segment; without spinules; without vestigial seta; without conical seta; without modified seta; with spinous process; on outer margin; surpassing distal margin; with aesthetasc; one element. Actual segment 16 with seta; two elements; of unequal size; plumose; one larger than segment; surpassing to distal margin; not beyond three sequential segments; not bifidform; without spinules; without vestigial seta; without conical seta; without modified seta; with spinous process; on outer margin; surpassing distal margin; unequal size to process on preceding segment; with aesthetasc; one element. Actual segment 17 with seta; one element; straight; none larger than segment; bifidform; without spinules; without vestigial seta; without conical seta; with modified seta; one element; stout form; surpassing to distal margin; not beyond of the sequential segment; parallel to antennule direction; without spinous process; without aesthetasc. Actual segment 18 with seta; two elements; of equal size; straight; none larger than segment; without spinules; without vestigial seta; without conical seta; with modified seta; one element; stout form; surpassing distal margin; parallel to antennule direction; without spinous process; without aesthetasc. Actual segment 19 with seta; two elements; of unequal size; plumose; none larger than segment; without spinules; without vestigial seta; without conical seta; with modified seta; two elements; stout form; at least one bifid form; surpassing distal margin; parallel to antennule direction; without spinous process; with aesthetasc; one element. Actual segment 20 with seta; four elements; of unequal size; straight; one larger than segment; surpassing to distal margin; beyond three sequential segments; without spinules; without vestigial seta; without conical seta; without modified seta; without spinous process; without aesthetasc. Actual segment 21 with seta; two elements; of equal size; plumose; one larger than segment; surpassing to distal margin; greater 3x than original segment; without spinules; without vestigial seta; without conical seta; without modified seta; without spinous process; without aesthetasc. Actual segment 22 with seta; four elements; of equal size; one larger than segment; plumose; surpassing to distal margin; greater 3x than original segment; without spinules; without vestigial seta; without conical seta; without modified seta; without spinous process; with aesthetasc; one element.

##### Left antennules

Uniramous; Left antennule surpassing to prosome; Left antennule extending beyond caudal rami. Ancestral segment I and II separated. Ancestral segment II and III fused. Ancestral segment III and IV fused. Ancestral segment IV and V separated. Ancestral segment V and VI separated. Ancestral segment VI and VII separated. Ancestral segment VII and VIII separated. Ancestral segment VIII and IX separated. Ancestral segment IX and X separated. Ancestral segment X and XI separated. Ancestral segment XI and XII separated. Ancestral segment XII and XIII separated. Ancestral segment XIII and XIV separated. Ancestral segment XIV and XV separated. Ancestral segment XV and XVI separated. Ancestral segment XVI and XVII separated. Ancestral segment XVII and XVIII separated. Ancestral segment XVIII and XIX separated. Ancestral segment XIX and XX separated. Ancestral segment XX and XXI separated. Ancestral segment XXI and XXII separated. Ancestral segment XXII and XXIII separated. Ancestral segment XXIII and XXIV separated. Ancestral segment XXIV and XXV separated. Ancestral segment XXV and XXVI separated. Ancestral segment XXVI and XXVII separated. Ancestral segment XXVII and XXVIII fused.

Left antennule actual 25-segmented; not-geniculated. Actual segment 1 with seta; one element; none larger than segment; straight; with spinules; as a row; without vestigial seta; without conical seta; without modified seta; without spinous process; with aesthetasc; one element. Actual segment 2 with seta; three elements; of equal size; none larger than segment; straight; without spinules; with vestigial seta; one element; without conical seta; without modified seta; without spinous process; with aesthetasc; one element. Actual segment 3 with seta; one element; one larger than segment; straight; surpassing to distal margin; beyond three sequential segments; without spinules; with vestigial seta; one element; without conical seta; without modified seta; without spinous process; with aesthetasc. Actual segment 4 with seta; one element; none larger than segment; straight; without spinules; without vestigial seta; without conical seta; without modified seta; without spinous process; without aesthetasc. Actual segment 5 with seta; one element; one larger than segment; straight; surpassing to distal margin; not beyond three sequential segments; without spinules; with vestigial seta; one element; without conical seta; without modified seta; without spinous process; with aesthetasc; one element. Actual segment 6 with seta; one element; none larger than segment; straight; without spinules; without vestigial seta; without conical seta; without modified seta; without spinous process; without aesthetasc. Actual segment 7 with seta; one element; one larger than segment; straight; surpassing to distal margin; beyond three sequential segments; without spinules; without vestigial seta; without conical seta; without modified seta; without spinous process; with aesthetasc; one element. Actual segment 8 with seta; one element; one larger than segment; straight; surpassing distal margin; without spinules; without vestigial seta; with conical seta; without modified seta; without spinous process; without aesthetasc. Actual segment 9 with seta; two elements; of unequal size; one larger than segment; straight; surpassing to distal margin; beyond three sequential segments; without spinules; without vestigial seta; without conical seta; without modified seta; without spinous process; with aesthetasc; one element. Actual segment 10 with seta; one element; none larger than segment; straight; without spinules; without vestigial seta; without conical seta; without modified seta; without spinous process; without aesthetasc. Actual segment 11 with seta; one element; one larger than segment; straight; surpassing to distal margin; beyond three sequential segments; without spinules; without vestigial seta; without conical seta; without modified seta; without spinous process; without aesthetasc. Actual segment 12 with seta; one element; one larger than segment; straight; surpassing distal margin; without spinules; without vestigial seta; with conical seta; without modified seta; without spinous process; with aesthetasc; one element. Actual segment 13 with seta; one element; none elongated; straight; surpassing distal margin; without spinules; without vestigial seta; without conical seta; without modified seta; without spinous process; without aesthetasc. Actual segment 14 with seta; one element; elongated; straight; surpassing to distal margin; beyond three sequential segments; without spinules; without vestigial seta; without conical seta; without modified seta; without spinous process; with aesthetasc; one element. Actual segment 15 with seta; one element; larger than segment; straight; surpassing to distal margin; not beyond three sequential segments; without spinules; without vestigial seta; without conical seta; without modified seta; without spinous process; without aesthetasc. Actual segment 16 with seta; one element; larger than segment; plumose; surpassing to distal margin; not beyond three sequential segments; without spinules; without vestigial seta; without conical seta; without modified seta; without spinous process; with aesthetasc; one element. Actual segment 17 with seta; one element; not larger than segment; straight; without spinules; without vestigial seta; without conical seta; without modified seta; without spinous process; without aesthetasc. Actual segment 18 with seta; one element; larger than segment; straight; surpassing to distal margin; beyond three sequential segments; without spinules; without vestigial seta; without conical seta; without modified seta; without spinous process; without aesthetasc. Actual segment 19 with seta; one element; not larger than segment; straight; surpassing distal margin; without spinules; without vestigial seta; without conical seta; without modified seta; without spinous process; with aesthetasc; one element. Actual segment 20 with seta; one element; not larger than segment; straight; surpassing distal margin; without spinules; without vestigial seta; without conical seta; without modified seta; without spinous process; without aesthetasc. Actual segment 21 with seta; one element; larger than segment; plumose; surpassing to distal margin; beyond three sequential segments; without spinules; without vestigial seta; without conical seta; without modified seta; without spinous process; without aesthetasc. Actual segment 22 with seta; two elements; of unequal size; one of them elongated; plumose; surpassing to distal margin; without spinules; without vestigial seta; without conical seta; without modified seta; without spinous process; without aesthetasc. Actual segment 23 with seta; two elements; of unequal size; one larger than segment; plumose; surpassing to distal margin; greater 3x than original segment; without spinules; without vestigial seta; without conical seta; without modified seta; without spinous process; without aesthetasc. Actual segment 24 with seta; two elements; of equal size; one larger than segment; plumose; surpassing to distal margin; greater 3x than original segment; without spinules; without vestigial seta; without conical seta; without modified seta; without spinous process; without aesthetasc. Actual segment 25 with seta; four elements; of equal size; elongated; plumose; surpassing to distal margin; 4 times larger than segment; without spinules; without vestigial seta; without conical seta; without modified seta; without spinous process; with aesthetasc; one element.

##### Antenna

Biramous. Antenna coxa separated from the basis; bearing seta; 1; on inner surface; at distal corner; reaching to the endopod 1. Antenna basis (fusion) separated from the endopodal segment; bearing seta; 2; on inner surface; at distal corner. Endopodal ancestral segment I and II separated. Ancestral segment II and III fused. Ancestral segment III and IV fused. Ancestral segment III and IV fully. Antenna endopod actual 2-segmented. Actual segment 1 not bilobate; with seta; two; on inner margin; with spinules; as a row; obliquely; on outer surface; with pore. Actual segment 2 bilobate; with discontinuity on outer cuticle; not developed as a suture; inner lobe bearing 8 setae; distally; outer lobe bearing 7 setae; distally; with spinules; as a patch; on outer surface. Antenna exopod ancestral segment I and II separated. Ancestral segment II and III fused. Ancestral segment III and IV fused. Ancestral segment IV and V separated. Ancestral segment V and VI separated. Ancestral segment VI and VII separated. Ancestral segment VII and VIII separated. Ancestral segment VIII and IX separated. Ancestral segment IX and X fused. Antenna exopod actual 7-segmented. Actual segment 1 single; elongated (width-length, equal or larger ratio 2:1); with seta; one; at inner surface. Actual segment 2 compound; elongated (larger width-length ratio 2:1); with seta; three; at inner surface. Actual segment 3 single; not elongated (lesser width-length ratio 2:1); with seta; one; at inner surface. Actual segment 4 single; not elongated (lesser width-length ratio 2:1); with seta; one; at inner surface. Actual segment 5 single; not elongated (lesser width-length ratio 2:1); with seta; one; at inner surface. Actual segment 6 single; not elongated (lesser width-length ratio 2:1); with seta; one; at inner surface. Actual segment 7 compound; elongated (larger or equal width-length ratio 2:1); with seta; one; at inner surface; and three; at distal surface.

##### Oral features

**Mandible**. Coxal gnathobase sclerotized; with lobe; prominent; on caudal margin; presence of cutting blade; with tooth-like prominence; two, distinctly; 1 acute; on caudal margin; and 1 triangular; on sub-caudal margin; without acute projection between the prominences; with additional spinules; as a row; on dorsal surface; with seta; 1; dorsally; on apical surface; with spinules; apicalmost. Mandible palps biramous; comprising the basis; with seta; four; differently inserted; first medially; reaching to beyond the endopod 1; second distally; third distally; fourth distally; on inner margin; none with setulose ornamentation. Mandible endopod 2-segmented. Mandible endopod 1 with lobe; bearing seta; four; distally inserted; without spinules. Mandible endopod 2 without lobe; bearing setae; nine elements; distally inserted; with spinules; as a row; double. Mandible exopod 4-segmented. Mandible exopod 1 with seta; one element; distally; on inner margin. Mandible exopod 2 with seta; one element; distally; on inner side. Mandible exopod 3 with seta; one element; distally; on inner side. Mandible exopod 4 with setae; three elements; on terminal region. **Maxillule**. Birramous. Maxillule 3-segmented. Maxillule praecoxa with praecoxal arthrite; bearing spines; fifteen elements; ten marginally; plus five sub-marginally; with spinules; as a patch; on sub-marginal surface. Maxillule coxa with coxal epipodite; with conspicuous outer lobe; bearing setae; nine elements; with coxal endite; elongated (larger or equal width-length ratio 2:1); bearing setae; four elements. Maxillule basis with basal endite; double; first proximal; elongated (larger width-length ratio 2:1; separated from basis; with setae; four elements; distally inserted; second distal; fused to basis; not elongated (lesser width-length ratio 2:1); with setae; four elements; distally inserted; with setules; as a row; on inner side; basal exite present; with setae; one element; on outer surface. Maxillule endopod 1-segmented. Endopod 1 bilobate; first proximal; with setae; three elements; second distal; with setae; five elements. Maxillule exopod 1-segmented. Exopod 1 with setae; six elements; with setules; as a row; on inner side; spinules absent. **Maxilla**. Uniramous. Maxilla 5-segmented. Maxilla praecoxa fused to coxa; incompletely; distinct externally; with praecoxal endite; double; first elongated endite (larger or equal width length ratio 2:1); proximally inserted; with seta; straight, or plumose; 1 straight; 4 plumose; with spine; single; without spinules; without setule; second elongated endite (larger or equal width length ratio 2:1); distally inserted; with seta; plumose; 3 plumose; without spine; with spinules; as a row; on distal margin; with setule; as a row; on distal margin; absence of outer seta. Maxilla coxa with coxal endite; double; first elongated endite (larger or equal width); proximally inserted; with seta; plumose; 3 plumose; without spine; without spinules; with setules; as a row; on proximal margin; second elongated endite (larger or equal width); distally inserted; with seta; plumose; 3 plumose; without spine; without spinules; with setules; as a row; on proximal margin; absence of outer seta. Maxilla basis with basal endite; single; elongated (larger or equal width-length ratio 2:1); with seta; plumose; 3 plumose; without spinules; absence of outer seta. Maxilla endopod 2-segmented. Endopod 1 with seta; 2 plumose; without spine; without spinules; without setules. Maxilla endopod 2 with seta; 2 plumose; without spine; without spinules; without setules. **Maxilliped**. Uniramous; Maxilliped 8-segmented. Maxilliped praecoxa fused to coxa; incompletely; distinct internally; with praecoxal endite; not elongated (lesser width-length ratio 2:1); distally inserted; with seta; 1 straight; with spinules; as a row; single; on basal surface; without setules. Maxilliped coxa with coxal endite; three coxal endite; first elongated (larger or equal width); proximally inserted; with seta; 2 plumose; with spinules; as a patch; single; on apical surface; without setules; second not elongated (lesser width-length ratio 2:1); medially inserted; with seta; 3 plumose; with spinules; as a row; single; on medial surface; without setules; third elongated (larger or equal width length ratio 2:1); distally inserted; with seta; 3 plumose; none reaching to beyond of the basis; with spinules; as a row; single; on basal surface; without setules; with lobe; prominence; at inner distal angle; ornamented; with spinules; continuously on margin. Maxilliped basis without basal endite; with seta; 3 plumose; with spinules; as a row; single; on medial surface; with setules; as a row; single; on inner margin. Maxilliped endopod segment 6-segmented. Endopod 1 with seta; 2 plumose; on inner surface. Endopod 2 with seta; 3 plumose; on inner surface. Endopod 3 with seta; 2 plumose; on inner surface. Endopod 4 with seta; 2 plumose; on inner surface. Endopod 5 with seta; 2 plumose; on inner surface, or on outer surface; outer seta absent. Endopod 6 with seta; 4 plumose; on inner surface, or on outer surface.

##### Swimming legs features

**First swimming legs.** Symmetrical; biramous. First swimming legs intercoxal plate without seta. First swimming legs praecoxa absent. First swimming legs coxa with seta; one; straight; distally inserted; on inner surface; surpassing to first endopodal segment; with setules; two group; as a patch; on inner margin; and as a row; double; on anterior surface; outerly; without spinules; without spine. First swimming legs basis without seta; with setules; as a patch; single; on outer surface; without spinules; without spine. First swimming legs endopod 2-segmented. Endopod 1 with seta; straight; restricted; to inner surface; one element; without spine; with setules; as a row; single; continuously; on outer surface; without spinules; absence of Schmeil’s organ. Endopod 2 with seta; unrestricted; three on inner surface; one on outer surface; two on distal surface; straight; without spine; with setules; as a row; single; continuously; on outer surface; without spinules; absence of Schmeil’s organ. Endopod 3 absence. First swimming legs exopod 1 with seta; restricted; 1 on inner surface; with spine; 1; stout; smaller than original segment; serrated; on inner side; continuously; with setules; as a row; single; as a row; innerly. First swimming legs exopod 2 with seta; restricted; 1 on inner surface; straight; without spine; with setules; as a row; single; continuously; on inner margin, or on outer margin; without spinules. First swimming legs exopod 3 with setule; as a row; single; continuously; on outer surface; without spinules; with seta; unrestricted; 2 on inner surface; 2 on terminal surface; with spine; 2; unequal size; first no longer 2x than origin segment; stout; serrated; on inner side, or on outer side; equally; second longer 3x than origin segment; slender; serrated; on outer side; with ornamentation on non-serrated side; by setules. **Second swimming legs**. Symmetrical; Second swimming legs biramous. Second swimming legs intercoxal plate without seta. Second swimming legs praecoxa present; located laterally. Second swimming legs coxa with seta; straight; distally inserted; on inner surface; surpassing to basal segment; without setules; without spinules; without spine. Second swimming legs basis without seta; without setules; without spinules; without spine. Second swimming legs endopod 3-segmented. Endopod 1 with seta; straight; restricted; one on inner surface; without spine; with setules; as a row; single; continuously; on outer surface; without spinules; absence of Schmeil’s organ. Endopod 2 with seta; straight; unrestricted; two on inner surface; without spine; with setules; as a row; single; continuously; on outer side; without spinules; presence of Schmeil’s organ; on posterior surface. Endopod 3 with seta; straight; unrestricted; three on inner surface; two on outer surface; two on distal surface; without spine; without setules; with spinules; as a row; double; distally inserted; at anterior surface; absence of Schmeil’s organ. Second swimming legs exopod 1 with seta; restricted; one on inner surface; with spine; 1; stout; not reaching to distal-third of the exopod 2; serrated; on inner side, or on outer side; with setules; as a row; single; continuously; on inner side; without spinules; absence of Schmeil’s organ. Exopod 2 with seta; unrestricted; one on inner surface; with spine; 1; stout; not surpassing the exopod 3; serrated; on inner side, or on outer side; with setules; as a row; single; continuously; on inner surface; without spinules; absence of Schmeil’s organ. Exopod 3 with seta; plurimarginal; three on inner surface; two on terminal surface; with spine; 2; unequal size; first no longer 2x than origin segment; stout; serrated; on inner side, or on outer side; equally; second longer 2x than origin segment; slender; serrated; on outer side; with ornamentation on non-serrated side; of setules; setules on outer surface; as a row; single; continuously; on inner surface; with spinules; as a row; single; distally inserted; at anterior surface; absence of Schmeil’s organ. **Third swimming legs**. Symmetrical; Third swimming legs biramous. Third swimming legs intercoxal plate without seta. Third swimming legs praecoxa present; not laterally located. Third swimming legs coxa with seta; straight; distally inserted; on inner surface; surpassing to basal segment; without setules; without spinules; without spine. Third swimming legs basis without seta; without setules; without spinules; without spine. Third swimming legs endopod 3-segmented. Endopod 1 with seta; restricted; one on inner surface; without spine; without setules; without spinules; absence of Schmeil’s organ. Endopod 2 with seta; restricted; two on inner surface; straight; without spine; without setules; without spinules; absence of Schmeil’s organ. Endopod 3 with seta; straight; plurimarginal; two on inner surface; two on outer surface; three on terminal surface; without spine; without setules; with spinules; as a patch; proximally inserted; double; at anterior surface; absence of Schmeil’s organ. Third swimming legs exopod 1 with seta; restricted; straight; one on inner surface; with spine; 1; stout; not reaching to the distal-third of the exopod 2; serrated; equally; on inner surface, or on outer surface; with setules; as a row; single; continuously; on inner surface; without spinules; absence of Schmeil’s organ. Exopod 2 with seta; straight; restricted; one on inner surface; with spine; 1; stout; not reaching out to exopod 3; serrated; on inner side, or on outer side; equally; with setules; as a row; single; continuously; on inner side; without spinules; absence of Schmeil’s organ. Exopod 3 without setules; with spinules; as a row; single; distally inserted; at anterior surface; with seta; straight; unrestricted; three on inner surface; two on terminal surface; with spine; 2; unequal size; first no longer 2x than origin segment; stout; serrated; on inner side, or on outer side; equally; second longer 2x than origin segment; slender; serrated; on outer side; with ornamentation on non-serrated side; of setules; absence of Schmeil’s organ. **Fourth swimming legs**. Symmetrical; biramous. Intercoxal plate without sensilla. Praecoxa present. Coxa with seta; distally inserted; on inner margin; reaching out to endopod 1; without spinules; setules absent. Basis with seta; one; medially inserted; on posterior surface; smaller than the original segment; without setules; without spinules; without spine. Fourth swimming legs endopod 3-segmented. Endopod 1 with seta; one; restricted; on inner surface; without spine; without setules; without spinules; absence of Schmeil’s organ. Endopod 2 with seta; restricted; two on inner side; without spine; with setules; as a row; single; continuously; on outer surface; without spinules; absence of Schmeil’s organ. Endopod 3 with seta; unrestricted; two on inner surface; two on outer surface; three on distal surface; without spine; without setules; with spinules; as a row; double; distally inserted; at anterior surface; absence of Schmeil’s organ. Fourth swimming legs exopod 1 with seta; restricted; one on inner surface; with spine; 1; stout; not reaching out to distal-third of the exopod 2; serrated; on inner side, or on outer side; equally; with setules; as a row; single; continuously; on inner surface; without spinules; absence of Schmeil’s organ. Exopod 2 with seta; restricted; one on inner surface; with spine; 1; stout; not reaching the end of exopod 3; serrated; on inner side, or on outer side; equally; with setules; as a row; single; continuously; on inner surface; without spinules; absence of Schmeil’s organ. Exopod 3 without setules; with spinules; as a row; single; distally inserted; at anterior surface; with seta; unrestricted; three on inner surface; two on distal surface; with spine; 2; unequal size; first no longer 2x than origin segment; stout; serrated; on inner side, or on outer side; equally; second longer 2x than origin segment; slender; serrated; on outer side; without ornamentation on non-serrated side; absence of Schmeil’s organ.

##### Fifth swimming legs features

Asymmetrical. Fifth swimming leg intercoxal plate with length not equal or greater than width on 1.5x; with irregular proximal margin; discontinuous to; the anterior margin of the left coxa, or the anterior margin of the right coxa; posterior sensilla on the right lateral absent. **Fifth left swimming leg**. Fifth left swimming leg biramous; leg reaching first right exopod segment; proximally. Fifth left swimming leg praecoxa absent. **Fifth left swimming leg** coxa concave inner side; without teeth-like structures; with process; conical; on posterior surface; outer side; distally inserted; projecting over basis; reaching to the proximal surface; with sensilla; stout; triangular; at apex; no longer 2x than insertion basis; with swelling; on inner side; distally; without seta; without spinules. Fifth left swimming leg basis sub-cylindrical; unequal size between inner and outer side; shorter outer than inner side; with rectilinear inner side; rounded internal proximal expansion absent; without outgrowth; with groove; deep; obliquely; on posterior surface; not reaching the endopodal lobe; not ornamented; absence of protuberance; with seta; outerly inserted; longer 2x than origin segment; absence of minutely granular. Fifth left swimming leg endopod segments 1 and 2 fused; segments 2 and 3 fused; 1-segmented; stout; separated from the basis; ornamented; on inner side; with spinules; more than four elements; as a row; terminally; row of setules absent; without seta. Fifth left swimming leg exopod segments 1 and 2 separated; segments 2 and 3 fused; 2-segmented; stout; separated from the basis. Fifth left swimming leg exopod 1 sub-cylindrical; longer than broad; unequal size between inner and outer side; shorter inner than outer side; concave inner side; convex outer side; without swelling; without marginal extension; without process; with lobe; single; circular; medially inserted; on inner side; covered; by setules; without outer spine; absence seta. Fifth left swimming leg exopod 2 sub-triangular; longer than broad; unequal size between inner and outer side; shorter inner than outer side; uniform inner side; with convex outer side; setulose pad present; not prominently rounded; proximally; on inner side; inflated medial region present; setulose; anteriorly; distal process present; digitiform; denticulate; not bicuspidate; without transverse row of denticles; none oblique row of 5 denticles; at anterior surface; innerly directed; with seta; spiniform; ornamented by spinules; surpassing the distal-point of the segment; without outer spine; terminal claw absent.

##### Fifth right swimming leg

Biramous. Fifth right swimming leg praecoxa present; separated from the coxae; without ornamentation. Fifth right swimming leg coxa convex inner side; without teeth-like structures; with process; rounded; distally inserted; on posterior surface; closest to the outer rim; projecting over basis; beyond the first third; until the medial surface; without triangular protuberance innerly; with sensilla; slender; at apex; no longer 2x than basal insertion; without marginal extension; without seta; without spinules. Fifth right swimming leg basis cylindrical; unequal size between inner and outer side; shorter outer than inner side; convex inner side; tumescence present; not inflated; restricted on inner surface; proximally; without protuberance; absence of distinct minutely granular; additional inner process absent; with posterior groove; deep; obliquely; reaching the endopodal lobe; ornamented; with tubercles; throughout of the outer border; with seta; outerly inserted; on anterior surface; no longer 2x than origin segment; posterior protrusion present; distal process absent. Fifth right swimming leg with endopodite present; fused to basis; on anterior surface; ancestral segments 1 and 2 fused; ancestral segments 2 and 3 fused; stout; ornamented; with spinules; as a row; on inner side; terminally; without seta. Fifth right swimming leg exopod segments 1 and 2 separated; segments 2 and 3 fused; 2-segmented; stout; separated from the basis. Fifth right swimming leg exopod 1 trapezium; longer than broad; nearly 2 times; unequal size between both sides; shorter inner than outer side; rectilinear inner side; rectilinear outer side; with marginal extension; sub-triangular; distally inserted; at outer rim; spinules absent; with process; triangular; rectilinear; sharp tip; sclerotized; without ornamentation; distally inserted; at posterior surface; projecting over next segment; without outer spine; without seta; internal prominence absent; lamella on posterior surface absent. Fifth right swimming leg exopod 2 elliptical; longer than broad; nearly 2 times; unequal size between both sides; uniform inner side; convex outer side; without posterior proximal swelling; inner-posterior process absent; without marginal expansion; curved ridge on distal posterior surface absent; chitinous knobs absent; with outer spine; inserted sub-distally; arched; internally directed; ornamented innerly; by spinules; as a row; not ornamented outerly; sharp tip; with apparent curve; innerly directed; lesser than the length of the exopod 2; beyond to 2 times its size; 6x; sensilla absent; terminal claw present; equal or longer 1.5 times than insertion segment; sclerotized; arched; inward; with conspicuous curve; proximally; ornamented innerly; by spinules; as a row; throughout extension; ornamented outerly; sharp tip; not curved tip; with medial constriction; hyaline process absent.

##### FEMALE

Body longer and wider than male; Female body 1134 micrometers excluding caudal setae. Widest at first metasome segment. Distal margin of the prosomal segments without one line of setules at posterior margin. Prosome segments with spinules at least at one prosomal segment. Fourth metasome segment absence of dorsal protuberance. Fourth and fifth metasome segments separated. Limit between fourth and fifth metasome segments ornamented; with spinules; as a row; double; incomplete; same size; entirely over limit (lateral, dorsal). **Fifth metasome segment**. Fifth metasome segment with sensilla; dorsally; 2 elements; with epimeral plates. Epimeral plates asymmetrical. Right epimeral plates prominent, as projections; thinner than the left; one posterior-laterally directed; not reaching half length of the genital segment; with sensilla at the apex; dorsal-posterior sensilla present; slender; without ornamentation. Left epimeral plate without expansion.

##### Urosome

3-segmented. **Genital double-somite**. Asymmetrical in dorsal view; longer than broad; longer than other urosomites combined; dorsal suture at mid-length absent; not covered by spinules; with swelling; rounded; unequal size; greater right than left; anteriorly; with sensillae; on both sides; one; stout; with robust apex; at left lateral; not on lobular base; anteriorly; one; stout; at right lateral; not on lobular base; anteriorly; with robust apex; of equal size between then; lateral protuberance absent; with right posterior rim expanded; over next segment; without slender sensilla on each posterior rim; without posterior-dorsal process. Genital double-somite opercular pad present; broader than longer; symmetrical; development laterally; expanded posteriorly; covering partially; double gonoporal slit; located ventrally; with arthrodial membrane; inserted anteriorly; post-genital process absent; disto-ventral tumescence absent; ventral vertical folds absent; dorsal sensilla absent. Second urosome segment without ventral fusion to anal segment; right distal process absent. Caudal rami patch of setules on outer surface absent; patch of spinules on outer surface absent.

##### Oral appendices feature

Rostrum basal process absent. **Antennules**. Symmetrical. Right antennule surpassing to genital double-segment; extending beyond caudal rami. Right antennule exceeding the caudal setae. Right antennule ornamentation pattern equals to male left antennule; fully.

##### Fifth swimming legs

Symmetrical; Fifth swimming legs biramous. Fifth swimming legs intercoxal plate longer than wide; separated from the legs. Fifth swimming legs praecoxa without sclerite praecoxal. Fifth swimming legs coxa with process; conical; at the outer rim; distally; sensilla present; stout; at apex; projecting over basal segment; no longer 2x than basal insertion; marginal extension absent; without swelling; without seta; without spinules. Fifth swimming legs basis sub-triangular; unequal size between inner and outer sides; shorter outer than inner side; with convex inner side; without proximal inner outgrowth; without groove; with distal extension; on posterior surface; with seta; outerly inserted; on anterior surface; longer 2x than origin segment; reaching to exopod 1 distally. Fifth swimming legs endopod segments 1 and 2 fused; segments 2 and 3 fused; 1-segmented; stout; separated from the basis; absent discontinuity cuticle; with spinules; as a row; single; non-oblique; sub-terminally; at anterior surface; with seta; single; one distally; on posterior surface; rectilinear. Fifth swimming legs exopod segments 1 and 2 separated; segments 2 and 3 separated; 3-segmented; separated from the basis. Fifth swimming legs exopod 1 sub-cylindrical; longer than wide; longer or equal than 2 times; with unequal size between inner and outer side; shorter inner than outer side; with convex inner side; with concave outer side; without swelling; without marginal extension; without posterior process; without spine; without seta. Fifth swimming legs exopod 2 sub-cylindrical; longer than broad; longer or equal than 2 times; without swelling; without marginal extension; without process; without lobe; with spine; inserted laterally; rectilinear; without ornamentation; sharp tip; smaller than next segment; without seta. Fifth swimming legs exopod 3 cylindrical; longer than wide; without swelling; without process; without lobe; without spine; with seta; double; inserted terminally; unequal size between them; outer seta smaller than inner; nearly 3 times; outer seta not ornamented by setules; without ornamentation; presence of terminal claw; sclerotized; arched; externally directed; convex inner side; with ornamentation; of denticles; as a row; on surface partially; at medial region; concave outer side; with ornamentation; of denticles; as a row; on surface partially; at medial region; blunt tip; 6 times longer than origin segment.

##### Distribution records

###### BRAZIL

**Pará**: various places on the Guamá, Capim, and Tocantins River Basins, recorded by Cipólli & Carvalho (1973). Timbiras Lake, Caranandeua; Jurumundeua Lake, Caranandeua; furo de Panaquera: São Lourenço River, Sororoca Stream; Maratapá Bay: Grilo stream, paraná Samuuma; Mapará stream, paraná Samuuma. Cametá: Tocantins River, Maloca stream, Aricurá stream; Murú stream; Tocantins River, Tucuruí; marginal lake in Jatobal; Laguinho, Tucuruí. **Ceará**: pond in Fortaleza (Matsumura-Tundisi, 1986). **Paraíba**: ponde Puxinanã, in ville of same label, near to Campina Grande State (Wright, 1935); pond Novo, Campina Grande City (this study). **Pernambuco**: ponde in Garanhuns (Wright, 1935). **Minas Gerais**: Volta Grande Reservoir (between 19°57’52“-20°10’00“S e 48°25’-47°35’W) (Rolla *et al*., 1990). **Mato Grosso do Sul**: lagoons in Guaraná, and Baía River, tributaries of the Paraná River (Lima *et al*., 1996); Nova Andradina, floodplain of the upper Paraná River (Lansac-Tôha *et al*., 1992); Pato Lake, Pousada das Garças, and Baía, Curutuba Rivers, Ivinhema, and Paraná Rivers (Lansac-Tôha *et al*., 1997). **São Paulo**: reservoirs in Itapeva, and Funil, Paraíba do Sul River Basin (Sendacz & Kubo, 1982; Sendacz *et al*., 1985); Capivara, and Tietê Rivers (Matsumura-Tundisi, 1986); reservoirs in Barra Bonita, Tietê River (Tundisi & Matsumura-Tundisi, 1994); upper Paraná River: reservoirs in Ilha Solteira, and Jupiá, lagoons in Comprida 1 and 2, Jota Lake, Paraná River (Sendacz, 1997); Abaixo River, sand extraction area, Paraíba do Sul River Basis (Carvalho & Sendacz, 1998). **Rio de Janeiro**: Saudade Lake, 21°42’S, 41°20’W, and Campelo Lake, 21°40’S, 41°11’W (Reid & Esteves, 1984; Reid, 1985). **Paraná**: reservoir in Itaipu (Matsumura-Tundisi, 1986; Tomm *et al*., 1992); Porto Rico, floodplain of the upper Paraná River (Lansac-Tôha *et al*., 1992); reservoirs in Salto Osório, and Foz de Areia, Iguaçu River (Santos-Silva *et al*., 2015; Perbiche-Neves *et al*., 2015).

##### Habitat

Habitat in freshwaters: lakes, ponds, and reservoirs.

##### Remarks

The organisms of the species come from a reservoir area in the Brazilian Northeast and were included in the *nordestinus* complex during its foundation (Wright, 1935). From this, they were recorded for several geographic regions of Brazil, ranging from Pará (Cipólli & Carvalho) to the Iguaçu River, Paraná (Lansac-Tôha *et al*., 1992; Santos-Silva *et al*., 2015) currently. Kiefer (1936) considered the taxon for his initial proposal of *Notodiaptomus*, offering his current taxonomic recombination.

In the taxonomic trajectory of *N. iheringi* several studies approached the morphological variations of the species. Wright (1935) illustrated the male and female of the species, mainly: (1) male fifth right swimming leg exopod 1 longer than broad, around 2x; (2) male fifth right swimming leg exopod 2 with “short” outer seta; (3) male fifth left swimming leg exopod 2 directed inwardly; (4) female fifth metasome segment with dorsal elevation covering with spinules row; (5) female genital double-somite with lateral expansion proximally; (6) female fifth swimming legs endopod 1-segmented; (7) female fifth swimming legs exopod 2 with lateral spine shorter than exopod 3. Except for attribute 4, and proximal constriction condition (originally described), all other characteristics were corroborated in this effort.

Reid (1985) when expanding the geographic record of the species to the region of Rio de Janeiro offered important illustrations for morphology of the species. Variations to the description of Wright were found by the author, who on the occasion had already mentioned about the non-existence of the female fifth metasome segment without dorsal elevation. Santos-Silva *et al*. (2015) reviewing the organisms of the species from the Paraná State corroborated this variation but did not agree with the condition of the female fifth swimming leg basis with outer seta “short”, and endopod with proximal constriction and double apical setae. In the present effort we corroborate the additional conditions verified in Santos-Silva *et al*. (2015), the examined females from the Wright’s collection have fifth swimming leg basis with outer seta reaching to exopod 1 distally, and endopod without proximal constriction with single seta apically.

Perbiche-Neves *et al*. (2015) reviewed the species from organisms from the Furnas Reservoir in Rio Grande (Minas Gerais) and found additional variations. The male from this locality has third, fourth, and fifth metasome segments with spinules row posteriorly. This condition for fifth metasome segment with spinules row is partially varied, with the row apparently not continuous. Furthermore, were reported the male fifth right swimming leg exopod 2 with outer spine in rectilinear form, and female fourth and fifth metasome segments with incomplete suture dorsally. In this present approach was corroborated the observation for the ornamentation in the segments of the masculine metasoma, fifth right swimming leg exopod 2 with outer spine in curved inwardly, and female fourth and fifth metasome segments with complete suture dorsally, evidencing the separation between the segments totally.

Among the attributes used by Wright (1935; 1936; 1937) and Kiefer (1936; 1956) for the original groupings of the species, only male fifth right swimming leg basis with inner protuberance cannot be verified. Additionally some convergent attributes from the type-species were absent for *N. iheringi* and had the following variations: (1) male and female limit between fourth and fifth metasome segments with ornamentation; (2) male and female antennules reaching to beyond caudal setae; (3) male right antennule actual segment 8 with conical seta reaching to middle-point sequent segment, and conical seta on actual segment 12 smaller; (4) male fifth left swimming leg exopod 2 with denticulate digitiform process; (5) male fifth right swimming leg exopod 1 with blunt distal process posteriorly; (6) male fifth right swimming leg exopod 2 with outer spine 6x lesser than length of original segment; and (7) female fifth swimming legs endopod without discontinuity cuticle.

#### Notodiaptomus incompositus (Brian, 1926)

##### Synonymy

*Diaptomus incompositus* Brian, 1926: 182, figs. 7–9; 1927: 131; Brehm, 1933a: 284; 1935b: 298, 299, 305; 1958a: 168; 1965: 3; Wright, 1937: 76; 1938a: 298, 299, 301; 1938b: 562; 1939: 645, 647, 648; Ringuelet (in Olivier), 1955: 299; Reid, 1991: 738. *Diaptomus paranaensis* Pesta, 1927: 68, figs. 1a-d; Brehm, 1965: 7, 8, 11. *Notodiaptomus incompositus*; Kiefer, 1936a: 197; 1956: 242; Brehm, 1938: 27, 29; Ringuelet, 1958a: 45, 47, 52; 1958b: 18, 22, 23, 24, 25; 1962: 87, 92; 1968: 265; Brandorff, 1972: 44; 1976: 616, 620, 621, 622, fig. 2; Bowman, 1973: 199; Paggi & José de Paggi, 1974: tab. 1; 1990: 690, 692, tab. 2; Pezzani, 1977: 139; Dussart, 1979: 6; Löffler, 1981: 15; Dussart & Defaye, 1983: 135; Dussart & Frutos, 1985: 306, 307; 1986: 243, 244, 245, 246, 248, pl. 3, figs. 13–16; Montú & Gloeden, 1986: 6, 80, fig. 25a-d; José de Paggi & Paggi, 1988: 98; Reid & Moreno, 1990: 732; Reid, 1991: 738; Sendacz, 1993: 34, 35; Frutos, 1993: 112, tab. 3; Battistoni, 1995: 959; Rocha *et al*., 1995: 155, 156; Santos-Silva, 1998: 210; Santos-Silva *et al*., 1999: 127; Bohrer & Araújo, 1999: 92, 94, figs. 1–4; Santos-Silva, 2008: 29–30, fig. 6; Santos-Silva *et al*., 2015: 36–40, figs. 18–20, 40; Perbiche-Neves *et al*., 2015: 69–74, figs. 61–65, identification keys to male and female; Perbiche-Neves *et al*., 2020: 696-697, key to the Neotropical diaptomid, fig. 21.15 H. *Notodiaptomus* (*Notodiaptomus*) *incompositus*; Dussart, 1985a: 201, 208.

##### Type locality

Not clearly specified. In the original literature, some localities are mentioned, as Prata River, Uruguay River, and Trocadero River, all south of the Neotropical region.

##### Type material

Originally not specified. Santos-Silva *et al*., 2015 indicate that there is collection material deposited in Brazil (INPA), Germany (MNK), and the United States (USNM).

##### Material examined

Non-type material: 13 males, and 7 females, entire, in alcohol, from Lake Alalay, Cochabamba (17° 23m 43s S, 66°09m 35s W), Bolivia, no date, stored in Plankton Laboratory (n° 13474), INPA; 1 male (INPA-COP027, slides a-h) and 1 female (INPA-COP028, slides a-h) were selected to be dissection on eight slides each and deposited in the Zoological Collection of the INPA, Brazil. Additional material examined: 2 males, and 2 female (MZUSP 32936) from the River Uruguay (middle stretch) (31°38’12’’S, 60°2’21’’ W), Argentina, G. Perbiche-Neves collector, 29.I.2010, and stored in MZUSP; 2 males, and 2 females from the middle stretch River Uruguay, Argentina, G. Perbiche-Neves coll., 16.VI.2010; 2 males, River Iguaçu, at the reservoir of Foz do Areia, G. Perbiche-Neves coll., 12.II.2010; 1 male, and 1 female, River Parana (middle stretch), G. Perbiche-Neves coll., 21.I.2010.

##### Diagnosis

**(1)** Male fourth metasome segment ornamented posterior margin with spinules row minutely; **(2)** caudal rami with reticulated main axis; **(3)** male right antennule actual segment 22 with five setae; **(4)** male fifth left swimming leg exopod 1 with inner semicircular lobe doubly; **(5)** male fifth left swimming leg exopod 2 with denticulate distal digitiform process; **(6)** male fifth left swimming leg exopod 2 with spiniform seta surpassing distal-point of the segment; **(7)** male fifth right swimming leg exopod 1 broader than long; **(8)** male fifth right swimming leg exopod 2 with inner-posterior process medially; **(9)** female fourth and fifth metasome segments separated; **(10)** female fifth metasome segment ornamented with dorsal setules row medially; **(11)** female fifth swimming legs endopod 2-segmented.

##### Redescription

###### MALE

Body 1166 micrometers excluding caudal setae. Male body smaller and slenderer than female. Nerve axons myelinated. Prosome 6-segmented; widest at first metasome segment; without one line of setules at posterior margin; with spinules at least at one segment. Cephalosome anterior margin sub-triangular; with dorsal suture; incomplete; separate from first metasome segment. First metasome segment without sensilla. Second metasome segment with sensilla; 2 dorsally; of equal size. Third metasome segment with sensillae; 4 laterally; of unequal size; non-ornamented posterior margin. Fourth metasome segment with sensillae; 4 dorsally; of equal size; separated from the fifth metasome; ornamented; with spinules; as a row (minutely); on dorsally. Limit between fourth and fifth metasome segments ornamented; with spinules; as a row; on dorsal singly; on lateral singly; same size. Fifth metasome segment with sensilla; 2 dorsally; 2 laterally; Fifth metasome segment equal size; Fifth metasome segment without ornamentation; Fifth metasome segment without dorsal conical process; with epimeral plates. Epimeral plates symmetrical. Right epimeral plates reduced, as rounded distal corner segment limit; with sensilla; at the apex of projection; without ornamentation.

##### Urosome

5-segmented; Urosome 5-free segments. Genital somite asymmetrical in dorsal view; with single aperture; located on left side; ventrolaterally on posterior rim; with sensillae; on both sides; one; at left lateral; medially; one; at right rim; posteriorly; of equal size between then. Third urosome segment with spinules; as patch; without external seta. Fourth urosome segment with spinules; as patch; without sub-conical blunt dorsal-lateral process. Anal segment presence of dorsal sensillae; one on each side; medially inserted; presence of operculum; convex; covering the anal aperture fully. Caudal rami symmetrical; separated from anal segment; longer than wide; with setules; continuous on; inner side; each ramus bearing 6 caudal setae; 5 marginals; plumose; and 1 internal dorsally; straight; reticulated main axis; outermost seta with outer spiniform process absent.

##### Oral appendices feature

Rostrum symmetrical; separated from dorsal cephalic shield; by complete suture; sensillae present; one pair; anteriorly inserted on surface tegument; with rostral filament; double; paired; extended; into point; with basal process; in ventral view, rounded on left side; without a smaller basal expansion on the right side.

##### Antennules

Asymmetrical. **Right antennules**. Uniramous; right antennule surpassing to genital segment; right antennule not extending beyond caudal rami.

Right antennule ancestral segment I and II separated. Ancestral segment II and III fused. Ancestral segment III and IV fused. Ancestral segment IV and V separated. Ancestral segment V and VI separated. Ancestral segment VI and VII separated. Ancestral segment VII and VIII separated. Ancestral segment VIII and IX separated. Ancestral segment IX and X separated. Ancestral segment X and XI separated. Ancestral segment XI and XII separated. Ancestral segment XII and XIII separated. Ancestral segment XIII and XIV separated. Ancestral segment XIV and XV separated. Ancestral segment XV and XVI separated. Ancestral segment XVI and XVII separated. Ancestral segment XVII and XVIII separated. Ancestral segment XVIII and XIX separated. Ancestral segment XIX and XX separated. Ancestral segment XX and XXI separated. Ancestral segment XXI and XXII fused. Ancestral segment XXII and XXIII fused. Ancestral segment XXIII and XXIV separated. Ancestral segment XXIV and XXV fused. Ancestral segment XXV and XXVI separated. Ancestral segment XXVI and XXVII separated. Ancestral segment XXVII and XXVIII fused.

Right antennule actual 22-segmented; geniculated; between the segment 18 and segment 19; with swollen and modified region; formed by 5 segments; between 13 and 17 segments. Actual segment 1 with seta; one element; straight; none larger than segment; without spinules; without vestigial seta; without conical seta; without modified seta; without spinous process; with aesthetasc; one element. Actual segment 2 with seta; three elements; of unequal size; straight; none larger than segment; without spinules; with vestigial seta; one element; without conical seta; without modified seta; without spinous process; with aesthetasc; one element. Actual segment 3 with seta; one element; one larger than segment; surpassing to distal margin; beyond three sequential segments; straight; blunt apex; without spinules; with vestigial seta; one element; without conical seta; without modified seta; without spinous process; with aesthetasc. Actual segment 4 with seta; one element; one larger than segment; surpassing to distal margin; straight; not beyond three sequential segments; without spinules; without vestigial seta; without conical seta; without modified seta; without spinous process; without aesthetasc. Actual segment 5 with seta; one element; straight; one larger than segment; surpassing to distal margin; not beyond three sequential segments; without spinules; with vestigial seta; one element; without conical seta; without modified seta; without spinous process; with aesthetasc; one element. Actual segment 6 with seta; one element; none larger than segment; straight; without spinules; without vestigial seta; without conical seta; without modified seta; without spinous process; without aesthetasc. Actual segment 7 with seta; one element; straight; one larger than segment; surpassing to distal margin; beyond three sequential segments; blunt apex; without spinules; without vestigial seta; without conical seta; without modified seta; without spinous process; with aesthetasc; one element. Actual segment 8 with seta; one element; straight; none larger than segment; without spinules; without vestigial seta; with conical seta; one element; reaching to middle-point of the sequent segment; without modified seta; without spinous process; without aesthetasc. Actual segment 9 with seta; two elements; of unequal size; straight; one larger than segment; surpassing to distal margin; beyond three sequential segments; blunt apex; without spinules; without vestigial seta; without conical seta; without modified seta; without spinous process; with aesthetasc; one element. Actual segment 10 with seta; one element; straight; none larger than segment; without spinules; without vestigial seta; without conical seta; with modified seta; presenting blunt apex; slender form; surpassing to distal margin; beyond of the sequential segment; parallel to antennule direction; without spinous process; without aesthetasc. Actual segment 11 with seta; one element; straight; one larger than segment; surpassing to distal margin; not beyond three sequential segments; without spinules; without vestigial seta; without conical seta; with modified seta; slender form; presenting blunt apex; surpassing to distal margin; beyond of the sequential segment; parallel to antennule direction; shorter length than homologous of actual segment 13; without spinous process; without aesthetasc. Actual segment 12 with seta; one element; straight; one larger than segment; surpassing to distal margin; not beyond three sequential segments; without spinules; without vestigial seta; with conical seta; one element; smaller than those of the segment 8; without modified seta; without spinous process; with aesthetasc; one element; absent internal perpendicular fission. Actual segment 13 with seta; one element; straight; one larger than segment; surpassing to distal margin; not beyond three sequential segments; without spinules; without vestigial seta; without conical seta; with modified seta; stout form; surpassing to distal margin; to the distal-point of the sequence segment; parallel to antennule direction; presenting bifid apex; without spinous process; with aesthetasc; one element. Actual segment 14 with seta; two elements; of unequal size; straight; one larger than segment; surpassing to distal margin; beyond three sequential segments; blunt apex; without spinules; without vestigial seta; without conical seta; without modified seta; without spinous process; with aesthetasc; one element. Actual segment 15 with seta; two elements; of unequal size; straight; not bifidform; none larger than segment; without spinules; without vestigial seta; without conical seta; without modified seta; with spinous process; on outer margin; surpassing distal margin; with aesthetasc; one element. Actual segment 16 with seta; two elements; of unequal size; plumose; one larger than segment; surpassing to distal margin; not beyond three sequential segments; not bifidform; without spinules; without vestigial seta; without conical seta; without modified seta; with spinous process; on outer margin; surpassing distal margin; unequal size to process on preceding segment; with aesthetasc; one element. Actual segment 17 with seta; two elements; of unequal size; straight; none larger than segment; bifidform; without spinules; without vestigial seta; without conical seta; with modified seta; one element; stout form; surpassing to distal margin; not beyond of the sequential segment; parallel to antennule direction; without spinous process; without aesthetasc. Actual segment 18 with seta; two elements; of equal size; straight; none larger than segment; without spinules; without vestigial seta; without conical seta; with modified seta; one element; stout form; surpassing distal margin; parallel to antennule direction; without spinous process; without aesthetasc. Actual segment 19 with seta; two elements; of unequal size; plumose; none larger than segment; without spinules; without vestigial seta; without conical seta; with modified seta; two elements; stout form; at least one bifid form; surpassing distal margin; parallel to antennule direction; without spinous process; with aesthetasc; one element. Actual segment 20 with seta; four elements; of unequal size; straight; one larger than segment; surpassing to distal margin; beyond three sequential segments; without spinules; without vestigial seta; without conical seta; without modified seta; without spinous process; without aesthetasc. Actual segment 21 with seta; two elements; of equal size; plumose; one larger than segment; surpassing to distal margin; greater 3x than original segment; without spinules; without vestigial seta; without conical seta; without modified seta; without spinous process; without aesthetasc. Actual segment 22 with seta; five elements; of equal size; one larger than segment; plumose; surpassing to distal margin; greater 3x than original segment; without spinules; without vestigial seta; without conical seta; without modified seta; without spinous process; with aesthetasc; one element.

##### Left antennules

Uniramous; Left antennule surpassing to prosome; Left antennule not extending beyond caudal rami. Ancestral segment I and II separated. Ancestral segment II and III fused. Ancestral segment III and IV fused. Ancestral segment IV and V separated. Ancestral segment V and VI separated. Ancestral segment VI and VII separated. Ancestral segment VII and VIII separated. Ancestral segment VIII and IX separated. Ancestral segment IX and X separated. Ancestral segment X and XI separated. Ancestral segment XI and XII separated. Ancestral segment XII and XIII separated. Ancestral segment XIII and XIV separated. Ancestral segment XIV and XV separated. Ancestral segment XV and XVI separated. Ancestral segment XVI and XVII separated. Ancestral segment XVII and XVIII separated. Ancestral segment XVIII and XIX separated. Ancestral segment XIX and XX separated. Ancestral segment XX and XXI separated. Ancestral segment XXI and XXII separated. Ancestral segment XXII and XXIII separated. Ancestral segment XXIII and XXIV separated. Ancestral segment XXIV and XXV separated. Ancestral segment XXV and XXVI separated. Ancestral segment XXVI and XXVII separated. Ancestral segment XXVII and XXVIII fused.

Left antennule actual 25-segmented; not-geniculated. Actual segment 1 with seta; one element; none larger than segment; straight; without spinules; without vestigial seta; without conical seta; without modified seta; without spinous process; with aesthetasc; one element. Actual segment 2 with seta; three elements; of equal size; none larger than segment; straight; without spinules; with vestigial seta; one element; without conical seta; without modified seta; without spinous process; with aesthetasc; one element. Actual segment 3 with seta; one element; one larger than segment; straight; surpassing to distal margin; beyond three sequential segments; without spinules; with vestigial seta; one element; without conical seta; without modified seta; without spinous process; with aesthetasc. Actual segment 4 with seta; one element; none larger than segment; straight; without spinules; without vestigial seta; without conical seta; without modified seta; without spinous process; without aesthetasc. Actual segment 5 with seta; one element; one larger than segment; straight; surpassing to distal margin; not beyond three sequential segments; without spinules; with vestigial seta; one element; without conical seta; without modified seta; without spinous process; with aesthetasc; one element. Actual segment 6 with seta; one element; none larger than segment; straight; without spinules; without vestigial seta; without conical seta; without modified seta; without spinous process; without aesthetasc. Actual segment 7 with seta; one element; one larger than segment; straight; surpassing to distal margin; beyond three sequential segments; without spinules; without vestigial seta; without conical seta; without modified seta; without spinous process; with aesthetasc; one element. Actual segment 8 with seta; one element; one larger than segment; straight; surpassing distal margin; without spinules; without vestigial seta; with conical seta; without modified seta; without spinous process; without aesthetasc. Actual segment 9 with seta; two elements; of unequal size; one larger than segment; straight; surpassing to distal margin; beyond three sequential segments; without spinules; without vestigial seta; without conical seta; without modified seta; without spinous process; with aesthetasc; one element. Actual segment 10 with seta; one element; none larger than segment; straight; without spinules; without vestigial seta; without conical seta; without modified seta; without spinous process; without aesthetasc. Actual segment 11 with seta; one element; one larger than segment; straight; surpassing to distal margin; beyond three sequential segments; without spinules; without vestigial seta; without conical seta; without modified seta; without spinous process; without aesthetasc. Actual segment 12 with seta; one element; one larger than segment; straight; surpassing distal margin; without spinules; without vestigial seta; with conical seta; without modified seta; without spinous process; with aesthetasc; one element. Actual segment 13 with seta; one element; none elongated; straight; surpassing distal margin; without spinules; without vestigial seta; without conical seta; without modified seta; without spinous process; without aesthetasc. Actual segment 14 with seta; one element; elongated; straight; surpassing to distal margin; beyond three sequential segments; without spinules; without vestigial seta; without conical seta; without modified seta; without spinous process; with aesthetasc; one element. Actual segment 15 with seta; one element; larger than segment; straight; surpassing to distal margin; not beyond three sequential segments; without spinules; without vestigial seta; without conical seta; without modified seta; without spinous process; without aesthetasc. Actual segment 16 with seta; one element; larger than segment; plumose; surpassing to distal margin; not beyond three sequential segments; without spinules; without vestigial seta; without conical seta; without modified seta; without spinous process; with aesthetasc; one element. Actual segment 17 with seta; one element; not larger than segment; straight; without spinules; without vestigial seta; without conical seta; without modified seta; without spinous process; without aesthetasc. Actual segment 18 with seta; one element; larger than segment; straight; surpassing to distal margin; beyond three sequential segments; without spinules; without vestigial seta; without conical seta; without modified seta; without spinous process; without aesthetasc. Actual segment 19 with seta; one element; not larger than segment; straight; surpassing distal margin; without spinules; without vestigial seta; without conical seta; without modified seta; without spinous process; with aesthetasc; one element. Actual segment 20 with seta; one element; not larger than segment; straight; surpassing distal margin; without spinules; without vestigial seta; without conical seta; without modified seta; without spinous process; without aesthetasc. Actual segment 21 with seta; one element; larger than segment; plumose; surpassing to distal margin; beyond three sequential segments; without spinules; without vestigial seta; without conical seta; without modified seta; without spinous process; without aesthetasc. Actual segment 22 with seta; two elements; of unequal size; one of them elongated; plumose; surpassing to distal margin; without spinules; without vestigial seta; without conical seta; without modified seta; without spinous process; without aesthetasc. Actual segment 23 with seta; two elements; of unequal size; one larger than segment; plumose; surpassing to distal margin; greater 3x than original segment; without spinules; without vestigial seta; without conical seta; without modified seta; without spinous process; without aesthetasc. Actual segment 24 with seta; two elements; of equal size; one larger than segment; plumose; surpassing to distal margin; greater 3x than original segment; without spinules; without vestigial seta; without conical seta; without modified seta; without spinous process; without aesthetasc. Actual segment 25 with seta; four elements; of equal size; elongated; plumose; surpassing to distal margin; 4 times larger than segment; without spinules; without vestigial seta; without conical seta; without modified seta; without spinous process; with aesthetasc; one element.

##### Antenna

Biramous. Antenna coxa separated from the basis; bearing seta; 1; on inner surface; at distal corner; reaching to the endopod 1. Antenna basis (fusion) separated from the endopodal segment; bearing seta; 2; on inner surface; at distal corner. Endopodal ancestral segment I and II separated. Ancestral segment II and III fused. Ancestral segment III and IV fused. Ancestral segment III and IV fully. Antenna endopod actual 2-segmented. Actual segment 1 not bilobate; with seta; two; on inner margin; with spinules; as a row; obliquely; on outer surface; with pore. Actual segment 2 bilobate; without discontinuity on outer cuticle; inner lobe bearing 8 setae; distally; outer lobe bearing 7 setae; distally; with spinules; as a patch; on outer surface. Antenna exopod ancestral segment I and II separated. Ancestral segment II and III fused. Ancestral segment III and IV fused. Ancestral segment IV and V separated. Ancestral segment V and VI separated. Ancestral segment VI and VII separated. Ancestral segment VII and VIII separated. Ancestral segment VIII and IX separated. Ancestral segment IX and X fused. Antenna exopod actual 7-segmented. Actual segment 1 single; elongated (width-length, equal or larger ratio 2:1); with seta; one; at inner surface. Actual segment 2 compound; elongated (larger width-length ratio 2:1); with seta; three; at inner surface. Actual segment 3 single; not elongated (lesser width-length ratio 2:1); with seta; one; at inner surface. Actual segment 4 single; not elongated (lesser width-length ratio 2:1); with seta; one; at inner surface. Actual segment 5 single; not elongated (lesser width-length ratio 2:1); with seta; one; at inner surface. Actual segment 6 single; not elongated (lesser width-length ratio 2:1); with seta; one; at inner surface. Actual segment 7 compound; elongated (larger or equal width-length ratio 2:1); with seta; one; at inner surface; and three; at distal surface.

##### Oral features

**Mandible**. Coxal gnathobase sclerotized; with lobe; prominent; on caudal margin; presence of cutting blade; with tooth-like prominence; two, distinctly; 1 acute; on caudal margin; and 1 triangular; on sub-caudal margin; without acute projection between the prominences; with additional spinules; as a row; on dorsal surface; with seta; 1; dorsally; on apical surface; with spinules; apicalmost. Mandible palps biramous; comprising the basis; with seta; four; differently inserted; first medially; reaching to beyond the endopod 1; second distally; third distally; fourth distally; on inner margin; none with setulose ornamentation. Mandible endopod 2-segmented. Mandible endopod 1 with lobe; bearing seta; four; distally inserted; without spinules. Mandible endopod 2 without lobe; bearing setae; nine elements; distally inserted; with spinules; as a row; double. Mandible exopod 4-segmented. Mandible exopod 1 with seta; one element; distally; on inner margin. Mandible exopod 2 with seta; one element; distally; on inner side. Mandible exopod 3 with seta; one element; distally; on inner side. Mandible exopod 4 with setae; three elements; on terminal region. **Maxillule**. Birramous. Maxillule 3-segmented. Maxillule praecoxa with praecoxal arthrite; bearing spines; fifteen elements; ten marginally; plus, five sub-marginally; with spinules; as a patch; on sub-marginal surface. Maxillule coxa with coxal epipodite; with conspicuous outer lobe; bearing setae; nine elements; with coxal endite; elongated (larger or equal width-length ratio 2:1); bearing setae; four elements. Maxillule basis with basal endite; double; first proximal; elongated (larger width-length ratio 2:1; separated from basis; with setae; four elements; distally inserted; second distal; fused to basis; not elongated (lesser width-length ratio 2:1); with setae; four elements; distally inserted; with setules; as a row; on inner side; basal exite present; with setae; one element; on outer surface. Maxillule endopod 1-segmented. Endopod 1 bilobate; first proximal; with setae; three elements; second distal; with setae; five elements. Maxillule exopod 1-segmented. Exopod 1 with setae; six elements; with setules; as a row; on inner side; spinules absent. **Maxilla**. Uniramous. Maxilla 5-segmented. Maxilla praecoxa fused to coxa; incompletely; distinct externally; with praecoxal endite; double; first elongated endite (larger or equal width length ratio 2:1); proximally inserted; with seta; straight, or plumose; 1 straight; 4 plumose; with spine; single; without spinules; without setule; second elongated endite (larger or equal width length ratio 2:1); distally inserted; with seta; plumose; 3 plumose; without spine; with spinules; as a row; on distal margin; with setule; as a row; on distal margin; absence of outer seta. Maxilla coxa with coxal endite; double; first elongated endite (larger or equal width); proximally inserted; with seta; plumose; 3 plumose; without spine; without spinules; with setules; as a row; on proximal margin; second elongated endite (larger or equal width); distally inserted; with seta; plumose; 3 plumose; without spine; without spinules; with setules; as a row; on proximal margin; absence of outer seta. Maxilla basis with basal endite; single; elongated (larger or equal width-length ratio 2:1); with seta; plumose; 3 plumose; without spinules; absence of outer seta. Maxilla endopod 2-segmented. Endopod 1 with seta; 2 plumose; without spine; without spinules; without setules. Maxilla endopod 2 with seta; 2 plumose; without spine; without spinules; without setules. **Maxilliped**. Uniramous; Maxilliped 8-segmented. Maxilliped praecoxa fused to coxa; incompletely; distinct internally; with praecoxal endite; not elongated (lesser width-length ratio 2:1); distally inserted; with seta; 1 straight; with spinules; as a row; single; on basal surface; without setules. Maxilliped coxa with coxal endite; three coxal endite; first elongated (larger or equal width); proximally inserted; with seta; 2 plumose; with spinules; as a patch; single; on apical surface; without setules; second not elongated (lesser width-length ratio 2:1); medially inserted; with seta; 3 plumose; with spinules; as a row; single; on medial surface; without setules; third elongated (larger or equal width length ratio 2:1); distally inserted; with seta; 3 plumose; none reaching to beyond of the basis; with spinules; as a row; single; on basal surface; without setules; with lobe; prominence; at inner distal angle; ornamented; with spinules; continuously on margin. Maxilliped basis without basal endite; with seta; 3 plumose; with spinules; as a row; single; on medial surface; with setules; as a row; single; on inner margin. Maxilliped endopod segment 6-segmented. Endopod 1 with seta; 2 plumose; on inner surface. Endopod 2 with seta; 3 plumose; on inner surface. Endopod 3 with seta; 2 plumose; on inner surface. Endopod 4 with seta; 2 plumose; on inner surface. Endopod 5 with seta; 2 plumose; on inner surface, or on outer surface; outer seta absent. Endopod 6 with seta; 4 plumose; on inner surface, or on outer surface.

##### Swimming legs features

**First swimming legs.** Symmetrical; biramous. First swimming legs intercoxal plate without seta. First swimming legs praecoxa absent. First swimming legs coxa with seta; one; straight; distally inserted; on inner surface; surpassing to first endopodal segment; with setules; one group; as a patch; on inner margin; without spinules; without spine. First swimming legs basis without seta; with setules; as a patch; single; on outer surface; without spinules; without spine. First swimming legs endopod 2-segmented. Endopod 1 with seta; straight; restricted; to inner surface; one element; without spine; with setules; as a row; single; continuously; on outer surface; without spinules; absence of Schmeil’s organ. Endopod 2 with seta; unrestricted; three on inner surface; one on outer surface; two on distal surface; straight; without spine; with setules; as a row; single; continuously; on outer surface; without spinules; absence of Schmeil’s organ. Endopod 3 absence. First swimming legs exopod 1 with seta; restricted; 1 on inner surface; with spine; 1; stout; smaller than original segment; serrated; on inner side; continuously; with setules; as a row; single; as a row; innerly. First swimming legs exopod 2 with seta; restricted; 1 on inner surface; straight; without spine; with setules; as a row; single; continuously; on inner margin, or on outer margin; without spinules. First swimming legs exopod 3 with setule; as a row; single; continuously; on outer surface; without spinules; with seta; unrestricted; 2 on inner surface; 2 on terminal surface; with spine; 2; unequal size; first no longer 2x than origin segment; stout; serrated; on inner side, or on outer side; equally; second longer 3x than origin segment; slender; serrated; on outer side; with ornamentation on non-serrated side; by setules. **Second swimming legs**. Symmetrical; Second swimming legs biramous. Second swimming legs intercoxal plate without seta. Second swimming legs praecoxa present; located laterally. Second swimming legs coxa with seta; straight; distally inserted; on inner surface; surpassing to basal segment; without setules; without spinules; without spine. Second swimming legs basis without seta; without setules; without spinules; without spine. Second swimming legs endopod 3-segmented. Endopod 1 with seta; straight; restricted; one on inner surface; without spine; with setules; as a row; single; continuously; on outer surface; without spinules; absence of Schmeil’s organ. Endopod 2 with seta; straight; unrestricted; two on inner surface; without spine; with setules; as a row; single; continuously; on outer side; without spinules; presence of Schmeil’s organ; on posterior surface. Endopod 3 with seta; straight; restricted; three on inner surface; two on outer surface; two on distal surface; without spine; without setules; with spinules; as a row; double; distally inserted; at anterior surface; absence of Schmeil’s organ. Second swimming legs exopod 1 with seta; restricted; one on inner surface; with spine; 1; stout; not reaching to distal-third of the exopod 2; serrated; on inner side, or on outer side; with setules; as a row; single; continuously; on inner side; without spinules; absence of Schmeil’s organ. Exopod 2 with seta; unrestricted; one on inner surface; with spine; 1; stout; not surpassing the exopod 3; serrated; on inner side, or on outer side; with setules; as a row; single; continuously; on inner surface; without spinules; absence of Schmeil’s organ. Exopod 3 with seta; plurimarginal; three on inner surface; two on terminal surface; with spine; 2; unequal size; first no longer 2x than origin segment; stout; serrated; on inner side, or on outer side; equally; second longer 2x than origin segment; slender; serrated; on outer side; with ornamentation on non-serrated side; of setules; setules on outer surface; as a row; single; continuously; on inner surface; with spinules; as a row; single; distally inserted; at anterior surface; absence of Schmeil’s organ. **Third swimming legs**. Symmetrical; Third swimming legs biramous. Third swimming legs intercoxal plate without seta. Third swimming legs praecoxa present; not laterally located. Third swimming legs coxa with seta; straight; distally inserted; on inner surface; surpassing to first endopodal segment; without setules; without spinules; without spine. Third swimming legs basis without seta; without setules; without spinules; without spine. Third swimming legs endopod 3-segmented. Endopod 1 with seta; restricted; one on inner surface; without spine; without setules; without spinules; absence of Schmeil’s organ. Endopod 2 with seta; restricted; two on inner surface; straight; without spine; without setules; without spinules; absence of Schmeil’s organ. Endopod 3 with seta; straight; plurimarginal; two on inner surface; two on outer surface; three on terminal surface; without spine; without setules; with spinules; as a row; distally inserted; double; at anterior surface; absence of Schmeil’s organ. Third swimming legs exopod 1 with seta; restricted; straight; one on inner surface; with spine; 1; stout; not reaching to the distal-third of the exopod 2; serrated; equally; on inner surface, or on outer surface; with setules; as a row; single; continuously; on inner surface; without spinules; absence of Schmeil’s organ. Exopod 2 with seta; straight; restricted; one on inner surface; with spine; 1; stout; not reaching out to exopod 3; serrated; on inner side, or on outer side; equally; with setules; as a row; single; continuously; on inner side; without spinules; absence of Schmeil’s organ. Exopod 3 without setules; with spinules; as a row; single; distally inserted; at anterior surface; with seta; straight; unrestricted; three on inner surface; two on terminal surface; with spine; 2; unequal size; first no longer 2x than origin segment; stout; serrated; on inner side, or on outer side; equally; second longer 2x than origin segment; slender; serrated; on outer side; with ornamentation on non-serrated side; of setules; absence of Schmeil’s organ. **Fourth swimming legs**. Symmetrical; biramous. Intercoxal plate without sensilla. Praecoxa present. Coxa with seta; distally inserted; on inner margin; reaching out to endopod 1; without spinules; setules absent. Basis with seta; one; medially inserted; on posterior surface; smaller than the original segment; without setules; without spinules; without spine. Fourth swimming legs endopod 3-segmented. Endopod 1 with seta; one; restricted; on inner surface; without spine; without setules; without spinules; absence of Schmeil’s organ. Endopod 2 with seta; restricted; two on inner side; without spine; with setules; as a row; single; continuously; on outer surface; without spinules; absence of Schmeil’s organ. Endopod 3 with seta; unrestricted; two on inner surface; two on outer surface; three on distal surface; without spine; without setules; with spinules; as a row; double; distally inserted; at anterior surface; absence of Schmeil’s organ. Fourth swimming legs exopod 1 with seta; restricted; one on inner surface; with spine; 1; stout; not reaching out to distal-third of the exopod 2; serrated; on inner side, or on outer side; equally; with setules; as a row; single; continuously; on inner surface; without spinules; absence of Schmeil’s organ. Exopod 2 with seta; restricted; one on inner surface; with spine; 1; stout; not reaching the end of exopod 3; serrated; on inner side, or on outer side; equally; with setules; as a row; single; continuously; on inner surface; without spinules; absence of Schmeil’s organ. Exopod 3 without setules; with spinules; as a row; single; distally inserted; at anterior surface; with seta; unrestricted; three on inner surface; two on distal surface; with spine; 2; unequal size; first no longer 2x than origin segment; stout; serrated; on inner side, or on outer side; equally; second longer 2x than origin segment; slender; serrated; on outer side; without ornamentation on non-serrated side; absence of Schmeil’s organ.

##### Fifth swimming legs features

Asymmetrical. Fifth swimming leg intercoxal plate with length not equal or greater than width on 1.5x; with irregular proximal margin; discontinuous to; the anterior margin of the left coxa, or the anterior margin of the right coxa; posterior sensilla on the right lateral absent. **Fifth left swimming leg**. Fifth left swimming leg biramous; leg reaching first right exopod segment; proximally. Fifth left swimming leg praecoxa present; rudimentary; separated from the coxae; without ornamentation. Fifth left swimming leg coxa concave inner side; without teeth-like structures; with process; conical; on posterior surface; outer side; distally inserted; not projecting over basis; with sensilla; stout; triangular; at apex; longer 2x than insertion basis; without swelling; without seta; without spinules. Fifth left swimming leg basis sub-cylindrical; unequal size between inner and outer side; shorter outer than inner side; with concave inner side; rounded internal proximal expansion absent; without outgrowth; without groove; absence of protuberance; with seta; outerly inserted; no longer 2x than origin segment; absence of minutely granular. Fifth left swimming leg endopod segments 1 and 2 fused; segments 2 and 3 fused; 1-segmented; stout; separated from the basis; ornamented; on inner side; with spinules; more than four elements; as a row; terminally; row of setules absent; without seta. Fifth left swimming leg exopod segments 1 and 2 separated; segments 2 and 3 fused; 2-segmented; stout; separated from the basis. Fifth left swimming leg exopod 1 sub-cylindrical; longer than broad; equal size between inner and outer side; rectilinear inner side; convex outer side; without swelling; without marginal extension; without process; with lobe; double; semicircular; medially inserted; on inner side; covered; by setules; without outer spine; absence seta. Fifth left swimming leg exopod 2 digitiform; longer than broad; equal size between inner and outer side; disform inner side; with rectilinear outer side; setulose pad present; prominently rounded; proximally; on inner side; inflated medial region absent; distal process present; digitiform; denticulate; not bicuspidate; without transverse row of denticles; none oblique row of 5 denticles; at anterior surface; not innerly directed; with seta; spiniform; ornamented by spinules; surpassing the distal-point of the segment; without outer spine; terminal claw absent.

##### Fifth right swimming leg

Biramous. Fifth right swimming leg praecoxa present; separated from the coxae; without ornamentation. Fifth right swimming leg coxa convex inner side; without teeth-like structures; with process; rounded; distally inserted; on posterior surface; closest to the outer rim; projecting over basis; not beyond the first third; without triangular protuberance innerly; with sensilla; stout; at apex; no longer 2x than basal insertion; without marginal extension; without seta; without spinules. Fifth right swimming leg basis cylindrical; unequal size between inner and outer side; shorter outer than inner side; rectilinear inner side; tumescence present; not inflated; restricted on inner surface; proximally; without protuberance; absence of distinct minutely granular; additional inner process absent; with posterior groove; shallow; longitudinally; not reaching the endopodal lobe; not ornamented; with seta; outerly inserted; on anterior surface; no longer 2x than origin segment; posterior protrusion absent; distal process absent. Fifth right swimming leg with endopodite present; fused to basis; on anterior surface; ancestral segments 1 and 2 fused; ancestral segments 2 and 3 fused; stout; ornamented; with setules; as a row; on inner side; terminally; without seta. Fifth right swimming leg exopod segments 1 and 2 separated; segments 2 and 3 fused; 2-segmented; stout; separated from the basis. Fifth right swimming leg exopod 1 sub-cylindrical; longer than broad; nearly 1.25 times; unequal size between both sides; shorter inner than outer side; convex inner side; rectilinear outer side; with marginal extension; sub-triangular; distally inserted; at outer rim; spinules absent; with process; triangular; arched; internally directed; sharp tip; sclerotized; without ornamentation; distally inserted; at posterior surface; projecting over next segment; without outer spine; without seta; internal prominence present; acute; lamella on posterior surface absent. Fifth right swimming leg exopod 2 cylindrical; longer than broad; nearly 2 times; equal size between both sides; disform inner side; convex outer side; without posterior proximal swelling; inner-posterior process present; tubercular; medially; without marginal expansion; curved ridge on distal posterior surface absent; chitinous knobs absent; with outer spine; inserted sub-distally; arched; internally directed; ornamented innerly; by spinules; as a row; not ornamented outerly; sharp tip; with apparent curve; outerly directed; lesser than the length of the exopod 2; until to 2 times its size; 2x; sensilla absent; terminal claw present; equal or longer 1.5 times than insertion segment; sclerotized; arched; inward; with conspicuous curve; proximally; ornamented innerly; by spinules; as a row; partially on extension; medially, or distally; ornamented outerly; sharp tip; curved tip; outwards (slightly); without medial constriction; hyaline process absent.

##### FEMALE

Body longer and wider than male; Female body 1168 micrometers excluding caudal setae. Widest at first metasome segment. Distal margin of the prosomal segments with one line of setules at posterior margin. Prosome segments with spinules at least at one prosomal segment. Fourth metasome segment absence of dorsal protuberance. Fourth and fifth metasome segments separated. Limit between fourth and fifth metasome segments ornamented; with spinules; as a row; single; incomplete; same size; partially over limit; dorsally. **Fifth metasome segment**. Fifth metasome segment without sensilla; with epimeral plates; ornamented with dorsal setules row medially. Epimeral plates asymmetrical. Right epimeral plates prominent, as projections; thinner than the left; one posterior-laterally directed; reaching half length of the genital segment; with sensilla at the apex; dorsal-posterior sensilla present; stout; without ornamentation. Left epimeral plate with expansion; conical; on posterior surface; dorsally; with sensilla; at tip.

##### Urosome

3-segmented. **Genital double-somite**. Asymmetrical in dorsal view; longer than broad; longer than other urosomites combined; dorsal suture at mid-length absent; not covered by spinules; with swelling; rounded; unequal size; greater left than right; anteriorly; with sensillae; on both sides; one; stout; with robust apex; at left lateral; not on lobular base; medially; one; stout; at right lateral; not on lobular base; anteriorly; with robust apex; of equal size between then; lateral protuberance absent; with right posterior rim expanded; over next segment; without slender sensilla on each posterior rim; without posterior-dorsal process. Genital double-somite opercular pad present; broader than longer; symmetrical; development laterally; expanded posteriorly; covering partially; double gonoporal slit; located ventrally; with arthrodial membrane; inserted anteriorly; post-genital process absent; disto-ventral tumescence absent; ventral vertical folds absent; dorsal sensilla absent. Second urosome segment without ventral fusion to anal segment; right distal process absent. Caudal rami patch of setules on outer surface absent; patch of spinules on outer surface absent.

##### Oral appendices feature

Rostrum basal process absent. **Antennules**. Symmetrical. Right antennule surpassing to genital double-segment; extending beyond caudal rami. Right antennule not exceeding the caudal setae. Right antennule ornamentation pattern equals to male left antennule; mostly. Actual segment 13 without seta; without aesthetasc. Actual segment 14 without seta; without aesthetasc. Actual segment 15 without seta; without aesthetasc. Actual segment 16 without seta; without aesthetasc. Actual segment 17 without seta. Actual segment 18 without seta.

##### Fifth swimming legs

Symmetrical; Fifth swimming legs biramous. Fifth swimming legs intercoxal plate longer than wide; separated from the legs. Fifth swimming legs praecoxa with sclerite praecoxal; separated from the coxae; without ornamentation. Fifth swimming legs coxa with process; conical; at the outer rim; distally; sensilla present; stout; at apex; projecting over basal segment; no longer 2x than basal insertion; marginal extension absent; without swelling; without seta; without spinules. Fifth swimming legs basis sub-triangular; unequal size between inner and outer sides; shorter outer than inner side; with convex inner side; without proximal inner outgrowth; without groove; with distal extension; on posterior surface; with seta; outerly inserted; on anterior surface; longer 2x than origin segment; not reaching to exopod 1 distally. Fifth swimming legs endopod segments 1 and 2 separated; segments 2 and 3 fused; 2-segmented; with complete suture; stout; separated from the basis; ornamentation on segment 2; with spinules; as a row; single; non-oblique; sub-terminally; at anterior surface; with seta; double; one medially; on posterior surface; rectilinear; one distally; on posterior surface; arched; of unequal size; distal seta longer than medial seta. Fifth swimming legs exopod segments 1 and 2 separated; segments 2 and 3 separated; 3-segmented; separated from the basis. Fifth swimming legs exopod 1 sub-cylindrical; longer than wide; longer or equal than 2 times; with unequal size between inner and outer side; shorter inner than outer side; with convex inner side; with rectilinear outer side; without swelling; without marginal extension; without posterior process; without spine; without seta. Fifth swimming legs exopod 2 sub-cylindrical; longer than broad; longer or equal than 2 times; without swelling; without marginal extension; without process; without lobe; with spine; inserted laterally; rectilinear; without ornamentation; sharp tip; smaller than next segment; without seta. Fifth swimming legs exopod 3 cylindrical; longer than wide; without swelling; without process; without lobe; without spine; with seta; double; inserted terminally; unequal size between them; outer seta smaller than inner; nearly 3 times; outer seta not ornamented by setules; without ornamentation; presence of terminal claw; sclerotized; arched; externally directed; convex inner side; with ornamentation; of denticles; as a row; on surface partially; at medial region; concave outer side; with ornamentation; of denticles; as a row; on surface partially; at medial region; blunt tip; 6 times longer than origin segment.

##### Distribution records

###### BOLÍVIA

Alalay Lagoon, Cochabamba (Santos-Silva *et al*., 2015). BRAZIL. **Rio Grande do Sul**: Patos Lagoon (Montú & Gloeden, 1986; Bohrer & Araújo, 1999); Quadros Lagoon, in Porto Alegre, and Negra Lagoon, in Viamão City (Bohrer & Araújo, 1999); Machadinho Reservoir on the Uruguay River. ARGENTINA. Middle Paraná River between Santa Fé, and Paraná Citys (Paggi & José de Paggi, 1974); middle Paraná River (Paggi & José de Paggi, 1990). **Buenos Aires**: La Plata River, Tigre (Brian, 1926); Abra Nueva on Paraná Delta, near to Tigre (Pesta, 1927); Vivero Lake, Palermo; pond beside to road (3 kilometers to Glew South), road to San Vicent (Wright, 1938a); two places near to Dufaur; various localities near to Buenos Aires (Wright, 1939); the recordings presented below were made by Ringuelet (1958a): Olivera, between Luján, and Mercedes; pond in Maciel Island; pond near to Del Gato stream; Santiago River; on around to La Plata; pond in La Plata; pond Amichetti in Los Talas; Carpincho Lagoon, Junín; Alcollaradas de Bolívar Lagoon; Lobos Lagoon; Flores Grandes Lagoon; arroio Saladillo, in Atucha; Plaza Montero Lagoon, in Las Flores; Monte Lagoon; Las Perdices Lagoon; Vitel Lagoon; pond in Chascomús; Adela Lagoon; Del Burro Lagoon; Chis Lagoon; San Ramón Lagoon in Bragado; Tapalqué stream; Camarón Grande Lagoon, Pila; El Talita Lagoon; La Ttora Lagoon; Del Estado Lagoon; Sauce Grande Lagoon; Alsina Lagoon; Cochicó Lagoon; Del Pastero Lagoon; La Brava Lagoon; Los Padres Lagoon; mount of the Sauce Grande stream (Ringuelet, in Olivier, 1955); Chascomús Lagoon (Wright, 1938a); Hoya del Plata (Ringuelet, 1962); Monteros Lagoon, Laprida (Brehm, 1965); La Brava Lagoon, Mar del Plata (Brehm, 1965); artificial Lagoon in Balneario de Quilmes (Reid, 1991). **Capital Federal**: Riachuelo River in La Boca; Palermo (Brian, 1926); Zoological Garden, Buenos Aires (Pesta, 1927). **Chaco**: Tragadero River, Colonia Benitz (Brian, 1926); Resistencia (Brehm, 1965); Oro River (Dussart & Frutos, 1986). **Corrientes**: Lagoon 1 (La Turbia), Cerrito Lagoon, Paraná River, and lagoon 2 (Los Pajaros), Nueva Cerrito Island, Paraná River (Frutos, 1993). **Entre Rios**: Colón, and Concepción, Uruguay River (Brian, 1926). **Formosa:** Pilagá stream, and arroyo Salado (Dussart & Frutos, 1986). **Rio Negro**: Valcheta stream (Ringuelet, 1958a). **San Luis**: 25 lagoons in South of the province, major in Pedernera (Wright, 1939); Tres Lagunas (Reid, 1991). **Santa Fé**: Five Lille stream (Brehm, 1965); Resistencia Chaco (Brehm, 1965). URUGUAI. **Soriano**: Palmira, Uruguay River (Brian, 1926). **Montevideo**: pluviogenics ponds in Barra de Santa Lucia, near to Montevideo, and Paso de Arena (Wright, 1938a).

##### Habitat

Habitat in freshwaters: shallow lakes, pools, and ponds associated to streams.

##### Remarks

The species was introduced by Brian from specimens from the Silver River Basin in South Neotropical, probably. It was described for Argentina as *Diaptomus paranaensis* Pesta 1927 mistakenly, being accepted for the *nordestinus* complex only in its amplification (Wright, 1937), previously recombined to *Notodiaptomus* in Kiefer (1936). Throughout its taxonomic trajectory several studies have dealt with the species without presenting relevant variations (Pesta, 1927; Dussart & Fruits, 1986; Montú & Gloeden, 1986).

Perbiche-Neves *et al*. (2015) and Santos-Silva *et al*. (2015) offered a morphological review on the interest of the diaptomids of the Prata River Basin, and *nordestinus* complex respectively. In the first contribution cited, scanning electron microscopy evaluation was presented and brought important clarifications to the organisms of the species, evidencing the male condition to fourth metasome segment ornamented with spinules row, third and fourth urosome segments ornamented with spinules patch dorsally, female fifth metasome segment ornamented with dorsal setules row medially, and fifth metasome segment with epimeral plates bearing dorsal expansion with stout sensilla at apex.

Santos-Silva *et al*. (2015) when examining organisms from Bolivia and Argentina, collected by Wright (1936), described morphological conditions concordant to those reported in Perbiche-Neves *et al*. (2015). However, in the study of *nordestinus*, divergent characterizations of the organisms are presented in comparison to Perbiche-Neves *et al*. (2015) through the male fifth right swimming leg exopod 1 broader than long, and female fifth swimming leg endopod 2-segmented. In the present effort, we corroborate the morphological conditions described such as Santos-Silva *et al*. (2015).

Additionally, we identified interesting morphological attributes for the species:(1) male right antennule actual segment 22 with five setae; (2) male fifth left swimming leg exopod 1 with inner semicircular lobe doubly; (3) male fifth left swimming leg exopod 2 with denticulate distal digitiform process; (4) male fifth left swimming leg exopod 2 with spiniform seta surpassing distal-point of the segment. Previously other described characteristics (Perbiche-Neves *et al*., 2015; Santos-Silva *et al*., 2015) were confirmed: (1) male fifth right swimming leg exopod 2 with inner-posterior process medially; (2) female fourth and fifth metasome segments separated; and (3) caudal rami with reticulated main axis.

It is a morphological condition of *N. incompositus* and divergent from those used to group *nordestinus* (Kiefer, 1935; 1936; 1937): (1) male fifth right swimming leg basis without inner protuberance. Distinct species attributes of *Notodiaptomus* (Kiefer 1936, 1956) are also: (1) female fifth swimming legs endopod 2-segmented; (2) male fifth right swimming leg exopod 1 broader than long; and (3) male fifth right swimming leg exopod 2 without rounded hyaline protuberance on the posterior apical surface (in this thesis as curved ridge on distal posterior surface innerly). Among the divergent characteristics of the type species of the genus are: (1) male right antennule actual segment 8 with conical seta reaching to middle-point sequent segment; and (2) male fifth right swimming leg basis without ornamentation on posterior groove.

#### Notodiaptomus inflatus (Kiefer, 1933)

##### Synonymy

*Diaptomus inflatus* Kiefer, 1933: 38, pl. 1, figs. 1–7; Brandorff, 1976: 618, fig. 3. *Diaptomus inflatus*; Wright, 1936: 79; 1937: 76; 1938b: 562; Thomasson, 1953: 194; Brehm, 1958a: 166; Andrade & Brandorff, 1975: 102. *Notodiaptomus inflatus*; Kiefer, 1936a: 197; 1956: 242; Brandorff, 1972: 45; Andrade & Brandorff, 1975: 97; Löffler, 1981: 15; Dussart & Defaye, 1983: 136; Dussart & Robertson, 1984: 391; Robertson & Hardy, 1984: tab. 3; Rocha *et al*., 1995: 154, 156; Santos-Silva *et al*., 1999: 127; Santos-Silva, 2008: 30–31, fig. 6; Santos-Silva *et al*., 2015: 65–67, figs. 39A-G, 40. *Notodiaptomus* (*Wrightius*) *inflatus;* Dussart, 1985a: 210.

##### Type locality

Not clearly specified. There is mention in the original description of 2 males, 1 female, and 1 juvenile, owned by P.A. Chappuis, identified to “Manaus, 17.XI.1927”. Santos-Silva et al (2015) indicate that the material was probably collected in the Negro River.

##### Type material

Not originally specified. In Santos-Silva et al (2015) is indicated that probably the type-material is currently non-existent.

##### Material examined

Non-type material: 3 males, and 2 females from the De los Amigos River, watershed Madre de Dios, Peru. no date, collected by I. Samanez, and stored in Plankton Laboratory, INPA. 1 male (INPA-COP029, slides a-h) and 1 female (INPA-COP030, slides a-h) were selected to be dissection on eight slides each and deposited in the Zoological Collection of the INPA, Brazil.

##### Diagnosis

**(1)** Cephalosome with complete dorsal suture; **(2)** male epimeral plates asymmetrical; **(3)** male fifth left swimming leg with length reaching first right exopod segment distally; **(4)** male fifth left swimming leg exopod 2 with plus oblique row of 5 denticles on distal digitiform process anteriorly; **(5)** male fifth left swimming leg endopod 2-segmented; **(6)** male fifth right swimming leg exopod 1 broader than long 1.25x nearly; **(7)** male fifth right swimming leg exopod 1 with outer acute extension distally; **(8)** male fifth right swimming leg exopod 1 with trapezoidal lamella on posterior surface; **(9)** male fifth right swimming leg exopod 2 with inner expansion medially; **(10)** female fifth swimming legs exopod 3 with single seta terminally.

##### Redescription

###### MALE

Body 1430 micrometers excluding caudal setae. Male body smaller and slenderer than female. Nerve axons myelinated. Prosome 6-segmented; widest at first metasome segment; without one line of setules at posterior margin; without spinules at segments. Cephalosome anterior margin rounded; with dorsal suture; complete; separate from first metasome segment. First metasome segment without sensilla. Second metasome segment without sensilla. Third metasome segment without sensillae; non-ornamented posterior margin. Fourth metasome segment without sensillae; separated from the fifth metasome. Limit between fourth and fifth metasome segments without ornamentation. Fifth metasome segment without sensilla; Fifth metasome segment without ornamentation; Fifth metasome segment without dorsal conical process; with epimeral plates. Epimeral plates asymmetrical. Right epimeral plates prominent, as projections; one projection; posterior-dorsally directed; not reaching half length of the genital segment; with sensilla; at the apex of projection; without ornamentation. Left epimeral plate reduced, as rounded distal corner segment limit; with sensillae; at the apex of projection; without ornamentation.

##### Urosome

5-segmented; Urosome 5 - free segments. Genital somite symmetrical in dorsal view; with single aperture; located on left side; ventrolaterally on posterior rim; with sensillae; on both sides; one; at left lateral; posteriorly; one; at right rim; posteriorly; of equal size between then. Third urosome segment without spinules; without external seta. Fourth urosome segment without spinules; without sub-conical blunt dorsal-lateral process. Anal segment presence of dorsal sensillae; one on each side; medially inserted; presence of operculum; convex; covering the anal aperture fully. Caudal rami symmetrical; separated from anal segment; longer than wide; with setules; continuous on; inner side; each ramus bearing 6 caudal setae; 5 marginals; plumose; and 1 internal dorsally; straight; not reticulated main axis; outermost seta with outer spiniform process absent.

##### Oral appendices feature

Rostrum asymmetrical; separated from dorsal cephalic shield; by complete suture; sensillae present; one pair; anteriorly inserted on surface tegument; with rostral filament; double; paired; extended; into point; with basal process; in ventral view, rounded on left side; without a smaller basal expansion on the right side.

##### Antennules

Asymmetrical. **Right antennules**. Uniramous; right antennule surpassing to genital segment; right antennule extending beyond caudal rami.

Right antennule ancestral segment I and II separated. Ancestral segment II and III fused. Ancestral segment III and IV fused. Ancestral segment IV and V separated. Ancestral segment V and VI separated. Ancestral segment VI and VII separated. Ancestral segment VII and VIII separated. Ancestral segment VIII and IX separated. Ancestral segment IX and X separated. Ancestral segment X and XI separated. Ancestral segment XI and XII separated. Ancestral segment XII and XIII separated. Ancestral segment XIII and XIV separated. Ancestral segment XIV and XV separated. Ancestral segment XV and XVI separated. Ancestral segment XVI and XVII separated. Ancestral segment XVII and XVIII separated. Ancestral segment XVIII and XIX separated. Ancestral segment XIX and XX separated. Ancestral segment XX and XXI separated. Ancestral segment XXI and XXII fused. Ancestral segment XXII and XXIII fused. Ancestral segment XXIII and XXIV separated. Ancestral segment XXIV and XXV fused. Ancestral segment XXV and XXVI separated. Ancestral segment XXVI and XXVII separated. Ancestral segment XXVII and XXVIII fused.

Right antennule actual 22-segmented; geniculated; between the segment 18 and segment 19; with swollen and modified region; formed by 5 segments; between 13 and 17 segments. Actual segment 1 with seta; one element; straight; none larger than segment; without spinules; without vestigial seta; without conical seta; without modified seta; without spinous process; with aesthetasc; one element. Actual segment 2 with seta; three elements; of unequal size; straight; none larger than segment; without spinules; with vestigial seta; one element; without conical seta; without modified seta; without spinous process; with aesthetasc; one element. Actual segment 3 with seta; one element; one larger than segment; surpassing to distal margin; beyond three sequential segments; straight; blunt apex; without spinules; with vestigial seta; one element; without conical seta; without modified seta; without spinous process; with aesthetasc. Actual segment 4 with seta; one element; one larger than segment; surpassing to distal margin; straight; not beyond three sequential segments; without spinules; without vestigial seta; without conical seta; without modified seta; without spinous process; without aesthetasc. Actual segment 5 with seta; one element; straight; one larger than segment; surpassing to distal margin; not beyond three sequential segments; without spinules; with vestigial seta; one element; without conical seta; without modified seta; without spinous process; with aesthetasc; one element. Actual segment 6 with seta; one element; none larger than segment; straight; without spinules; without vestigial seta; without conical seta; without modified seta; without spinous process; without aesthetasc. Actual segment 7 with seta; one element; straight; one larger than segment; surpassing to distal margin; beyond three sequential segments; blunt apex; without spinules; without vestigial seta; without conical seta; without modified seta; without spinous process; with aesthetasc; one element. Actual segment 8 with seta; one element; straight; none larger than segment; without spinules; without vestigial seta; with conical seta; one element; not reaching to middle-point of the sequent segment; without modified seta; without spinous process; without aesthetasc. Actual segment 9 with seta; two elements; of unequal size; straight; one larger than segment; surpassing to distal margin; beyond three sequential segments; blunt apex; without spinules; without vestigial seta; without conical seta; without modified seta; without spinous process; with aesthetasc; one element. Actual segment 10 with seta; one element; straight; none larger than segment; without spinules; without vestigial seta; without conical seta; with modified seta; presenting blunt apex; slender form; surpassing to distal margin; beyond of the sequential segment; parallel to antennule direction; without spinous process; without aesthetasc. Actual segment 11 with seta; one element; straight; one larger than segment; surpassing to distal margin; not beyond three sequential segments; without spinules; without vestigial seta; without conical seta; with modified seta; slender form; presenting blunt apex; surpassing to distal margin; beyond of the sequential segment; parallel to antennule direction; shorter length than homologous of actual segment 13; without spinous process; without aesthetasc. Actual segment 12 with seta; one element; straight; one larger than segment; surpassing to distal margin; not beyond three sequential segments; without spinules; without vestigial seta; with conical seta; one element; not smaller than to segment 8; without modified seta; without spinous process; with aesthetasc; one element; absent internal perpendicular fission. Actual segment 13 with seta; one element; straight; one larger than segment; surpassing to distal margin; not beyond three sequential segments; without spinules; without vestigial seta; without conical seta; with modified seta; stout form; surpassing to distal margin; to the middle-point of the sequence segment; perpendicular to antennule direction; presenting bifid apex; without spinous process; with aesthetasc; one element. Actual segment 14 with seta; two elements; of unequal size; straight; one larger than segment; surpassing to distal margin; beyond three sequential segments; blunt apex; without spinules; without vestigial seta; without conical seta; without modified seta; without spinous process; with aesthetasc; one element. Actual segment 15 with seta; two elements; of unequal size; straight; not bifidform; none larger than segment; without spinules; without vestigial seta; without conical seta; without modified seta; with spinous process; on outer margin; surpassing distal margin; with aesthetasc; one element. Actual segment 16 with seta; two elements; of unequal size; plumose; one larger than segment; surpassing to distal margin; not beyond three sequential segments; not bifidform; without spinules; without vestigial seta; without conical seta; without modified seta; with spinous process; on outer margin; surpassing distal margin; unequal size to process on preceding segment; with aesthetasc; one element. Actual segment 17 with seta; two elements; of unequal size; straight; none larger than segment; bifidform; without spinules; without vestigial seta; without conical seta; with modified seta; one element; stout form; surpassing to distal margin; not beyond of the sequential segment; parallel to antennule direction; without spinous process; without aesthetasc. Actual segment 18 with seta; two elements; of equal size; straight; none larger than segment; without spinules; without vestigial seta; without conical seta; with modified seta; one element; stout form; surpassing distal margin; parallel to antennule direction; without spinous process; without aesthetasc. Actual segment 19 with seta; two elements; of unequal size; plumose; none larger than segment; without spinules; without vestigial seta; without conical seta; with modified seta; two elements; stout form; at least one bifid form; surpassing distal margin; parallel to antennule direction; without spinous process; with aesthetasc; one element. Actual segment 20 with seta; four elements; of unequal size; straight; one larger than segment; surpassing to distal margin; beyond three sequential segments; without spinules; without vestigial seta; without conical seta; without modified seta; without spinous process; without aesthetasc. Actual segment 21 with seta; two elements; of equal size; plumose; one larger than segment; surpassing to distal margin; greater 3x than original segment; without spinules; without vestigial seta; without conical seta; without modified seta; without spinous process; without aesthetasc. Actual segment 22 with seta; four elements; of equal size; one larger than segment; plumose; surpassing to distal margin; greater 3x than original segment; without spinules; without vestigial seta; without conical seta; without modified seta; without spinous process; with aesthetasc; one element.

##### Left antennules

Uniramous; Left antennule surpassing to prosome; Left antennule extending beyond caudal rami. Ancestral segment I and II separated. Ancestral segment II and III fused. Ancestral segment III and IV fused. Ancestral segment IV and V separated. Ancestral segment V and VI separated. Ancestral segment VI and VII separated. Ancestral segment VII and VIII separated. Ancestral segment VIII and IX separated. Ancestral segment IX and X separated. Ancestral segment X and XI separated. Ancestral segment XI and XII separated. Ancestral segment XII and XIII separated. Ancestral segment XIII and XIV separated.

Ancestral segment XIV and XV separated. Ancestral segment XV and XVI separated. Ancestral segment XVI and XVII separated. Ancestral segment XVII and XVIII separated. Ancestral segment XVIII and XIX separated. Ancestral segment XIX and XX separated. Ancestral segment XX and XXI separated. Ancestral segment XXI and XXII separated. Ancestral segment XXII and XXIII separated. Ancestral segment XXIII and XXIV separated. Ancestral segment XXIV and XXV separated. Ancestral segment XXV and XXVI separated. Ancestral segment XXVI and XXVII separated. Ancestral segment XXVII and XXVIII fused.

Left antennule actual 25-segmented; not-geniculated. Actual segment 1 with seta; one element; none larger than segment; straight; without spinules; without vestigial seta; without conical seta; without modified seta; without spinous process; with aesthetasc; one element. Actual segment 2 with seta; three elements; of equal size; none larger than segment; straight; without spinules; with vestigial seta; one element; without conical seta; without modified seta; without spinous process; with aesthetasc; one element. Actual segment 3 with seta; one element; one larger than segment; straight; surpassing to distal margin; beyond three sequential segments; without spinules; with vestigial seta; one element; without conical seta; without modified seta; without spinous process; with aesthetasc. Actual segment 4 with seta; one element; none larger than segment; straight; without spinules; without vestigial seta; without conical seta; without modified seta; without spinous process; without aesthetasc. Actual segment 5 with seta; one element; one larger than segment; straight; surpassing to distal margin; not beyond three sequential segments; without spinules; with vestigial seta; one element; without conical seta; without modified seta; without spinous process; with aesthetasc; one element. Actual segment 6 with seta; one element; none larger than segment; straight; without spinules; without vestigial seta; without conical seta; without modified seta; without spinous process; without aesthetasc. Actual segment 7 with seta; one element; one larger than segment; straight; surpassing to distal margin; beyond three sequential segments; without spinules; without vestigial seta; without conical seta; without modified seta; without spinous process; with aesthetasc; one element. Actual segment 8 with seta; one element; one larger than segment; straight; surpassing distal margin; without spinules; without vestigial seta; with conical seta; without modified seta; without spinous process; without aesthetasc. Actual segment 9 with seta; two elements; of unequal size; one larger than segment; straight; surpassing to distal margin; beyond three sequential segments; without spinules; without vestigial seta; without conical seta; without modified seta; without spinous process; with aesthetasc; one element. Actual segment 10 with seta; one element; none larger than segment; straight; without spinules; without vestigial seta; without conical seta; without modified seta; without spinous process; without aesthetasc. Actual segment 11 with seta; one element; one larger than segment; straight; surpassing to distal margin; beyond three sequential segments; without spinules; without vestigial seta; without conical seta; without modified seta; without spinous process; without aesthetasc. Actual segment 12 with seta; one element; one larger than segment; straight; surpassing distal margin; without spinules; without vestigial seta; with conical seta; without modified seta; without spinous process; with aesthetasc; one element. Actual segment 13 with seta; one element; none elongated; straight; surpassing distal margin; without spinules; without vestigial seta; without conical seta; without modified seta; without spinous process; without aesthetasc. Actual segment 14 with seta; one element; elongated; straight; surpassing to distal margin; beyond three sequential segments; without spinules; without vestigial seta; without conical seta; without modified seta; without spinous process; with aesthetasc; one element. Actual segment 15 with seta; one element; larger than segment; straight; surpassing to distal margin; not beyond three sequential segments; without spinules; without vestigial seta; without conical seta; without modified seta; without spinous process; without aesthetasc. Actual segment 16 with seta; one element; larger than segment; plumose; surpassing to distal margin; not beyond three sequential segments; without spinules; without vestigial seta; without conical seta; without modified seta; without spinous process; with aesthetasc; one element. Actual segment 17 with seta; one element; not larger than segment; straight; without spinules; without vestigial seta; without conical seta; without modified seta; without spinous process; without aesthetasc. Actual segment 18 with seta; one element; larger than segment; straight; surpassing to distal margin; beyond three sequential segments; without spinules; without vestigial seta; without conical seta; without modified seta; without spinous process; without aesthetasc. Actual segment 19 with seta; one element; not larger than segment; straight; surpassing distal margin; without spinules; without vestigial seta; without conical seta; without modified seta; without spinous process; with aesthetasc; one element. Actual segment 20 with seta; one element; not larger than segment; straight; surpassing distal margin; without spinules; without vestigial seta; without conical seta; without modified seta; without spinous process; without aesthetasc. Actual segment 21 with seta; one element; larger than segment; plumose; surpassing to distal margin; beyond three sequential segments; without spinules; without vestigial seta; without conical seta; without modified seta; without spinous process; without aesthetasc. Actual segment 22 with seta; two elements; of unequal size; one of them elongated; plumose; surpassing to distal margin; without spinules; without vestigial seta; without conical seta; without modified seta; without spinous process; without aesthetasc. Actual segment 23 with seta; two elements; of unequal size; one larger than segment; plumose; surpassing to distal margin; greater 3x than original segment; without spinules; without vestigial seta; without conical seta; without modified seta; without spinous process; without aesthetasc. Actual segment 24 with seta; two elements; of equal size; one larger than segment; plumose; surpassing to distal margin; greater 3x than original segment; without spinules; without vestigial seta; without conical seta; without modified seta; without spinous process; without aesthetasc. Actual segment 25 with seta; four elements; of equal size; elongated; plumose; surpassing to distal margin; 4 times larger than segment; without spinules; without vestigial seta; without conical seta; without modified seta; without spinous process; with aesthetasc; one element.

##### Antenna

Biramous. Antenna coxa separated from the basis; bearing seta; 1; on inner surface; at distal corner; reaching to the endopod 1. Antenna basis (fusion) separated from the endopodal segment; bearing seta; 2; on inner surface; at distal corner. Endopodal ancestral segment I and II separated. Ancestral segment II and III fused. Ancestral segment III and IV fused. Ancestral segment III and IV fully. Antenna endopod actual 2-segmented. Actual segment 1 not bilobate; with seta; two; on inner margin; with spinules; as a row; obliquely; on outer surface; with pore. Actual segment 2 bilobate; with discontinuity on outer cuticle; not developed as a suture; inner lobe bearing 8 setae; distally; outer lobe bearing 7 setae; distally; with spinules; as a patch; on outer surface. Antenna exopod ancestral segment I and II separated. Ancestral segment II and III fused. Ancestral segment III and IV fused. Ancestral segment IV and V separated. Ancestral segment V and VI separated. Ancestral segment VI and VII separated. Ancestral segment VII and VIII separated. Ancestral segment VIII and IX separated. Ancestral segment IX and X fused. Antenna exopod actual 7-segmented. Actual segment 1 single; elongated (width-length, equal or larger ratio 2:1); with seta; one; at inner surface. Actual segment 2 compound; elongated (larger width-length ratio 2:1); with seta; three; at inner surface. Actual segment 3 single; not elongated (lesser width-length ratio 2:1); with seta; one; at inner surface. Actual segment 4 single; not elongated (lesser width-length ratio 2:1); with seta; one; at inner surface. Actual segment 5 single; not elongated (lesser width-length ratio 2:1); with seta; one; at inner surface. Actual segment 6 single; not elongated (lesser width-length ratio 2:1); with seta; one; at inner surface. Actual segment 7 compound; elongated (larger or equal width-length ratio 2:1); with seta; one; at inner surface; and three; at distal surface.

##### Oral features

**Mandible**. Coxal gnathobase sclerotized; with lobe; prominent; on caudal margin; presence of cutting blade; with tooth-like prominence; two, distinctly; 1 acute; on caudal margin; and 1 triangular; on sub-caudal margin; without acute projection between the prominences; with additional spinules; as a row; on dorsal surface; with seta; 1; dorsally; on apical surface; with spinules; apicalmost. Mandible palps biramous; comprising the basis; with seta; four; differently inserted; first medially; reaching to beyond the endopod 1; second distally; third distally; fourth distally; on inner margin; none with setulose ornamentation. Mandible endopod 2-segmented. Mandible endopod 1 with lobe; bearing seta; four; distally inserted; without spinules. Mandible endopod 2 without lobe; bearing setae; nine elements; distally inserted; with spinules; as a row; double. Mandible exopod 4-segmented. Mandible exopod 1 with seta; one element; distally; on inner margin. Mandible exopod 2 with seta; one element; distally; on inner side. Mandible exopod 3 with seta; one element; distally; on inner side. Mandible exopod 4 with setae; three elements; on terminal region. **Maxillule**. Birramous. Maxillule 3-segmented. Maxillule praecoxa with praecoxal arthrite; bearing spines; fifteen elements; ten marginally; plus, five sub-marginally; with spinules; as a patch; on sub-marginal surface. Maxillule coxa with coxal epipodite; with conspicuous outer lobe; bearing setae; nine elements; with coxal endite; elongated (larger or equal width-length ratio 2:1); bearing setae; four elements. Maxillule basis with basal endite; double; first proximal; elongated (larger width-length ratio 2:1; separated from basis; with setae; four elements; distally inserted; second distal; fused to basis; not elongated (lesser width-length ratio 2:1); with setae; four elements; distally inserted; with setules; as a row; on inner side; basal exite present; with setae; one element; on outer surface. Maxillule endopod 1-segmented. Endopod 1 bilobate; first proximal; with setae; three elements; second distal; with setae; five elements. Maxillule exopod 1-segmented. Exopod 1 with setae; six elements; with setules; as a row; on inner side; spinules absent. **Maxilla**. Uniramous. Maxilla 5-segmented. Maxilla praecoxa fused to coxa; incompletely; distinct externally; with praecoxal endite; double; first elongated endite (larger or equal width length ratio 2:1); proximally inserted; with seta; straight, or plumose; 1 straight; 4 plumose; with spine; single; without spinules; without setule; second elongated endite (larger or equal width length ratio 2:1); distally inserted; with seta; plumose; 3 plumose; without spine; with spinules; as a row; on distal margin; with setule; as a row; on distal margin; absence of outer seta. Maxilla coxa with coxal endite; double; first elongated endite (larger or equal width); proximally inserted; with seta; plumose; 3 plumose; without spine; without spinules; with setules; as a row; on proximal margin; second elongated endite (larger or equal width); distally inserted; with seta; plumose; 3 plumose; without spine; without spinules; with setules; as a row; on proximal margin; absence of outer seta. Maxilla basis with basal endite; single; elongated (larger or equal width-length ratio 2:1); with seta; plumose; 3 plumose; without spinules; absence of outer seta. Maxilla endopod 2-segmented. Endopod 1 with seta; 2 plumose; without spine; without spinules; without setules. Maxilla endopod 2 with seta; 2 plumose; without spine; without spinules; without setules. **Maxilliped**. Uniramous; Maxilliped 8-segmented. Maxilliped praecoxa fused to coxa; incompletely; distinct internally; with praecoxal endite; not elongated (lesser width-length ratio 2:1); distally inserted; with seta; 1 straight; with spinules; as a row; single; on basal surface; without setules. Maxilliped coxa with coxal endite; three coxal endite; first elongated (larger or equal width); proximally inserted; with seta; 2 plumose; with spinules; as a patch; single; on apical surface; without setules; second not elongated (lesser width-length ratio 2:1); medially inserted; with seta; 3 plumose; with spinules; as a row; single; on medial surface; without setules; third elongated (larger or equal width length ratio 2:1); distally inserted; with seta; 3 plumose; none reaching to beyond of the basis; with spinules; as a row; single; on basal surface; without setules; with lobe; prominence; at inner distal angle; ornamented; with spinules; continuously on margin. Maxilliped basis without basal endite; with seta; 3 plumose; with spinules; as a row; single; on medial surface; with setules; as a row; single; on inner margin. Maxilliped endopod segment 6-segmented. Endopod 1 with seta; 2 plumose; on inner surface. Endopod 2 with seta; 3 plumose; on inner surface. Endopod 3 with seta; 2 plumose; on inner surface. Endopod 4 with seta; 2 plumose; on inner surface. Endopod 5 with seta; 2 plumose; on inner surface, or on outer surface; outer seta absent. Endopod 6 with seta; 4 plumose; on inner surface, or on outer surface.

##### Swimming legs features

**First swimming legs.** Symmetrical; biramous. First swimming legs intercoxal plate without seta. First swimming legs praecoxa absent. First swimming legs coxa with seta; one; straight; distally inserted; on inner surface; surpassing to first endopodal segment; with setules; two group; as a patch; on inner margin; and as a row; double; on anterior surface; outerly; without spinules; without spine. First swimming legs basis without seta; with setules; as a patch; single; on outer surface; without spinules; without spine. First swimming legs endopod 2-segmented. Endopod 1 with seta; straight; restricted; to inner surface; one element; without spine; with setules; as a row; single; continuously; on outer surface; without spinules; absence of Schmeil’s organ. Endopod 2 with seta; unrestricted; three on inner surface; one on outer surface; two on distal surface; straight; without spine; with setules; as a row; single; continuously; on outer surface; without spinules; absence of Schmeil’s organ. Endopod 3 absence. First swimming legs exopod 1 with seta; restricted; 1 on inner surface; with spine; 1; stout; smaller than original segment; serrated; on inner side; continuously; with setules; as a row; single; as a row; innerly. First swimming legs exopod 2 with seta; restricted; 1 on inner surface; straight; without spine; with setules; as a row; single; continuously; on inner margin, or on outer margin; without spinules. First swimming legs exopod 3 with setule; as a row; single; continuously; on outer surface; without spinules; with seta; unrestricted; 2 on inner surface; 2 on terminal surface; with spine; 2; unequal size; first no longer 2x than origin segment; stout; serrated; on inner side, or on outer side; equally; second longer 3x than origin segment; slender; serrated; on outer side; with ornamentation on non-serrated side; by setules. **Second swimming legs**. Symmetrical; Second swimming legs biramous. Second swimming legs intercoxal plate without seta. Second swimming legs praecoxa present; located laterally. Second swimming legs coxa with seta; straight; distally inserted; on inner surface; surpassing to basal segment; without setules; without spinules; without spine. Second swimming legs basis without seta; without setules; without spinules; without spine. Second swimming legs endopod 3-segmented. Endopod 1 with seta; straight; restricted; one on inner surface; without spine; with setules; as a row; single; continuously; on outer surface; without spinules; absence of Schmeil’s organ. Endopod 2 with seta; straight; unrestricted; two on inner surface; without spine; with setules; as a row; single; continuously; on outer side; without spinules; presence of Schmeil’s organ; on posterior surface. Endopod 3 with seta; straight; unrestricted; three on inner surface; two on outer surface; two on distal surface; without spine; without setules; with spinules; as a row; double; distally inserted; at anterior surface; absence of Schmeil’s organ. Second swimming legs exopod 1 with seta; restricted; one on inner surface; with spine; 1; stout; not reaching to distal-third of the exopod 2; serrated; on inner side, or on outer side; with setules; as a row; single; continuously; on inner side; without spinules; absence of Schmeil’s organ. Exopod 2 with seta; unrestricted; one on inner surface; with spine; 1; stout; not surpassing the exopod 3; serrated; on inner side, or on outer side; with setules; as a row; single; continuously; on inner surface; without spinules; absence of Schmeil’s organ. Exopod 3 with seta; plurimarginal; three on inner surface; two on terminal surface; with spine; 2; unequal size; first no longer 2x than origin segment; stout; serrated; on inner side, or on outer side; equally; second longer 2x than origin segment; slender; serrated; on outer side; with ornamentation on non-serrated side; of setules; setules on outer surface; as a row; single; continuously; on inner surface; with spinules; as a row; single; distally inserted; at anterior surface; absence of Schmeil’s organ. **Third swimming legs**. Symmetrical; Third swimming legs biramous. Third swimming legs intercoxal plate without seta. Third swimming legs praecoxa present; not laterally located. Third swimming legs coxa with seta; straight; distally inserted; on inner surface; surpassing to first endopodal segment; without setules; without spinules; without spine. Third swimming legs basis without seta; without setules; without spinules; without spine. Third swimming legs endopod 3-segmented. Endopod 1 with seta; restricted; one on inner surface; without spine; without setules; without spinules; absence of Schmeil’s organ. Endopod 2 with seta; restricted; two on inner surface; straight; without spine; without setules; without spinules; absence of Schmeil’s organ. Endopod 3 with seta; straight; plurimarginal; two on inner surface; two on outer surface; three on terminal surface; without spine; without setules; with spinules; as a row; distally inserted; double; at anterior surface; absence of Schmeil’s organ. Third swimming legs exopod 1 with seta; restricted; straight; one on inner surface; with spine; 1; stout; not reaching to the distal-third of the exopod 2; serrated; equally; on inner surface, or on outer surface; with setules; as a row; single; continuously; on inner surface; without spinules; absence of Schmeil’s organ. Exopod 2 with seta; straight; restricted; one on inner surface; with spine; 1; stout; not reaching out to exopod 3; serrated; on inner side, or on outer side; equally; with setules; as a row; single; continuously; on inner side; without spinules; absence of Schmeil’s organ. Exopod 3 without setules; with spinules; as a row; single; distally inserted; at anterior surface; with seta; straight; unrestricted; three on inner surface; two on terminal surface; with spine; 2; unequal size; first no longer 2x than origin segment; stout; serrated; on inner side, or on outer side; equally; second longer 2x than origin segment; slender; serrated; on outer side; with ornamentation on non-serrated side; of setules; absence of Schmeil’s organ. **Fourth swimming legs**. Symmetrical; biramous. Intercoxal plate without sensilla. Praecoxa present. Coxa with seta; distally inserted; on inner margin; reaching out to endopod 1; without spinules; setules absent. Basis with seta; one; medially inserted; on posterior surface; smaller than the original segment; without setules; without spinules; without spine. Fourth swimming legs endopod 3-segmented. Endopod 1 with seta; one; restricted; on inner surface; without spine; without setules; without spinules; absence of Schmeil’s organ. Endopod 2 with seta; restricted; two on inner side; without spine; with setules; as a row; single; continuously; on outer surface; without spinules; absence of Schmeil’s organ. Endopod 3 with seta; unrestricted; two on inner surface; two on outer surface; three on distal surface; without spine; without setules; with spinules; as a row; double; distally inserted; at anterior surface; absence of Schmeil’s organ. Fourth swimming legs exopod 1 with seta; restricted; one on inner surface; with spine; 1; stout; not reaching out to distal-third of the exopod 2; serrated; on inner side, or on outer side; equally; with setules; as a row; single; continuously; on inner surface; without spinules; absence of Schmeil’s organ. Exopod 2 with seta; restricted; one on inner surface; with spine; 1; stout; not reaching the end of exopod 3; serrated; on inner side, or on outer side; equally; with setules; as a row; single; continuously; on inner surface; without spinules; absence of Schmeil’s organ. Exopod 3 without setules; with spinules; as a row; single; distally inserted; at anterior surface; with seta; unrestricted; three on inner surface; two on distal surface; with spine; 2; unequal size; first no longer 2x than origin segment; stout; serrated; on inner side, or on outer side; equally; second longer 2x than origin segment; slender; serrated; on outer side; without ornamentation on non-serrated side; absence of Schmeil’s organ.

##### Fifth swimming legs features

Asymmetrical. Fifth swimming leg intercoxal plate with length equal or greater than width on 1.5x; with irregular proximal margin; discontinuous to; the anterior margin of the right coxa; posterior sensilla on the right lateral absent. **Fifth left swimming leg**. Fifth left swimming leg biramous; leg reaching first right exopod segment; distally. Fifth left swimming leg praecoxa present; rudimentary; separated from the coxae; without ornamentation. Fifth left swimming leg coxa concave inner side; without teeth-like structures; with process; conical; on posterior surface; outer side; distally inserted; not projecting over basis; with sensilla; stout; triangular; at apex; no longer 2x than insertion basis; without swelling; without seta; without spinules. Fifth left swimming leg basis sub-cylindrical; unequal size between inner and outer side; shorter outer than inner side; with convex inner side; rounded internal proximal expansion absent; without outgrowth; without groove; absence of protuberance; with seta; outerly inserted; no longer 2x than origin segment; absence of minutely granular. Fifth left swimming leg endopod segments 1 and 2 separated; segments 2 and 3 fused; 2-segmented; stout; separated from the basis; ornamented; ornamented on segment 2; on inner side; with spinules; more than four elements; as a row; terminally; row of setules absent; without seta. Fifth left swimming leg exopod segments 1 and 2 separated; segments 2 and 3 fused; 2-segmented; stout; separated from the basis. Fifth left swimming leg exopod 1 sub-cylindrical; longer than broad; equal size between inner and outer side; concave inner side; rectilinear outer side; without swelling; without marginal extension; without process; with lobe; single; circular; medially inserted; on inner side; covered; by setules; without outer spine; absence seta. Fifth left swimming leg exopod 2 digitiform; broader than long; unequal size between inner and outer side; shorter outer than inner side; disform inner side; with convex outer side; setulose pad present; prominently rounded; proximally; on inner side; inflated medial region absent; distal process present; digitiform; denticulate; not bicuspidate; with transverse row of denticles; plus oblique row of 5 denticles; at anterior surface; innerly directed; with seta; spiniform; ornamented by spinules; not surpassing the distal-point of the segment; without outer spine; terminal claw absent.

##### Fifth right swimming leg

Biramous. Fifth right swimming leg praecoxa present; separated from the coxae; without ornamentation. Fifth right swimming leg coxa concave inner side; without teeth-like structures; with process; conical; distally inserted; on posterior surface; closest to the outer rim; projecting over basis; beyond the first third; until the medial surface; without triangular protuberance innerly; with sensilla; stout; at apex; no longer 2x than basal insertion; without marginal extension; without seta; without spinules. Fifth right swimming leg basis cylindrical; unequal size between inner and outer side; shorter outer than inner side; concave inner side; tumescence present; not inflated; restricted on inner surface; proximally; without protuberance; absence of distinct minutely granular; additional inner process absent; without posterior groove; with seta; outerly inserted; on posterior surface; no longer 2x than origin segment; posterior protrusion absent; distal process absent. Fifth right swimming leg with endopodite present; separated from the basis; on anterior surface; ancestral segments 1 and 2 fused; ancestral segments 2 and 3 fused; 1-segmented; stout; ornamented; with setules; as a row; on inner side; terminally; with seta. Fifth right swimming leg exopod segments 1 and 2 separated; segments 2 and 3 fused; 2-segmented; stout; separated from the basis. Fifth right swimming leg exopod 1 trapezium; broader than long; nearly 1.25 times; unequal size between both sides; shorter inner than outer side; convex inner side; convex outer side; with marginal extension; acute; distally inserted; at outer rim; spinules absent; with process; rounded; sclerotized; without ornamentation; distally inserted; at posterior surface; projecting over next segment; without outer spine; without seta; internal prominence absent; lamella on posterior surface present; trapezoidal form; not surpassing to margin. Fifth right swimming leg exopod 2 cylindrical; longer than broad; nearly 2.5 times; equal size between both sides; disform inner side; convex outer side; without posterior proximal swelling; inner-posterior process absent; with marginal expansion; medial; innerly; curved ridge on distal posterior surface present; chitinous knobs absent; with outer spine; inserted sub-distally; rectilinear; not ornamented innerly; not ornamented outerly; sharp tip; without apparent curve; lesser than the length of the exopod 2; until to 2 times its size; 2x; sensilla absent; terminal claw present; equal or longer 1.5 times than insertion segment; sclerotized; arched; inward; with conspicuous curve; proximally; ornamented innerly; by spinules; as a row; partially on extension; medially, or distally; not ornamented outerly; sharp tip; not curved tip; without medial constriction; hyaline process absent.

##### FEMALE

Body longer and wider than male; Female body 1300 micrometers excluding caudal setae. Widest at posterior cephalosome. Distal margin of the prosomal segments without one line of setules at posterior margin. Prosome segments without spinules at prosomal segments. Fourth metasome segment absence of dorsal protuberance. Fourth and fifth metasome segments fused; totally. Limit between fourth and fifth metasome segments without ornamentation. **Fifth metasome segment**. Fifth metasome segment without sensilla; with epimeral plates. Epimeral plates asymmetrical. Right epimeral plates prominent, as projections; not thinner than the left; one posterior-dorsally directed; not reaching half length of the genital segment; with sensilla at the apex; dorsal-posterior sensilla present; stout; without ornamentation. Left epimeral plate without expansion.

##### Urosome

3-segmented. **Genital double-somite**. Asymmetrical in dorsal view; longer than broad; longer than other urosomites combined; dorsal suture at mid-length absent; not covered by spinules; with swelling; conical; unequal size; greater left than right; anteriorly; with sensillae; on both sides; one; stout; with robust apex; at left rim; not on lobular base; anteriorly; one; stout; at right lateral; on lobular base; anteriorly; with robust apex; of equal size between then; lateral protuberance absent; with right posterior rim expanded; over next segment; without slender sensilla on each posterior rim; without posterior-dorsal process. Genital double-somite opercular pad present; broader than longer; symmetrical; development laterally; expanded posteriorly; covering partially; double gonoporal slit; located ventrally; with arthrodial membrane; inserted anteriorly; post-genital process absent; disto-ventral tumescence absent; ventral vertical folds absent; dorsal sensilla absent. Second urosome segment without ventral fusion to anal segment; right distal process absent. Caudal rami patch of setules on outer surface absent; patch of spinules on outer surface absent.

##### Oral appendices feature

Rostrum basal process absent. **Antennules**. Symmetrical. Right antennule surpassing to genital double-segment; extending beyond caudal rami. Right antennule exceeding the caudal setae. Right antennule ornamentation pattern equals to male left antennule; fully.

##### Fifth swimming legs

Symmetrical; Fifth swimming legs biramous. Fifth swimming legs intercoxal plate longer than wide; separated from the legs. Fifth swimming legs praecoxa with sclerite praecoxal; separated from the coxae; without ornamentation. Fifth swimming legs coxa with process; conical; at the outer rim; distally; sensilla present; stout; at apex; projecting over basal segment; no longer 2x than basal insertion; marginal extension absent; without swelling; without seta; without spinules. Fifth swimming legs basis sub-triangular; unequal size between inner and outer sides; shorter outer than inner side; with convex inner side; without proximal inner outgrowth; without groove; with distal extension; on posterior surface; with seta; outerly inserted; on anterior surface; no longer 2x than origin segment. Fifth swimming legs endopod segments 1 and 2 fused; segments 2 and 3 fused; 1-segmented; stout; separated from the basis; present discontinuity cuticle; on inner side; with spinules; as a row; single; non-oblique; terminally; at anterior surface; without seta. Fifth swimming legs exopod segments 1 and 2 separated; segments 2 and 3 separated; 3-segmented; separated from the basis. Fifth swimming legs exopod 1 sub-cylindrical; longer than wide; longer or equal than 2 times; with unequal size between inner and outer side; shorter inner than outer side; with convex inner side; with rectilinear outer side; without swelling; without marginal extension; with posterior process; distally; without spine; without seta. Fifth swimming legs exopod 2 sub-cylindrical; longer than broad; longer or equal than 2 times; without swelling; without marginal extension; without process; without lobe; with spine; inserted laterally; arched; externally directed; without ornamentation; sharp tip; smaller than next segment; without seta. Fifth swimming legs exopod 3 cylindrical; longer than wide; without swelling; without process; without lobe; without spine; with seta; single; inserted terminally; outer seta not ornamented by setules; without ornamentation; presence of terminal claw; sclerotized; arched; externally directed; convex inner side; with ornamentation; of denticles; as a row; on surface partially; at medial region; concave outer side; with ornamentation; of denticles; as a row; on surface partially; at medial region; blunt tip; not 6 times longer origin segment.

##### Distribution records

###### BRAZIL

**Amazonas**: proximity to Manaus City (Kiefer, 1933). PERÚ. De los Amigos River, Madre de Dios Watershed (this study).

##### Habitat

Habitat in freshwaters: rivers.

##### Remarks

The taxon was founded from organisms from the Brazilian Amazon probably. In the original description Kiefer (1933) mentions material belonging to P. A. Chappuis, identified such as “Manaus, 17.XI.1927”. Santos-Silva *et al*. (2015) indicate that the material was collected in the Negro River probably. The species was recombined to *Notodiaptomus* at the creation of the genus and *nordestinus* complex at its amplification (Wright, 1936). It was also suggested to be transferred to the subgenus *Notodiaptomus* (*Wrightius*) (Dussart, 1985), an unfounded and never accepted proposal.

In the present study organisms were examined from the De los Amigos River, watershed Madre de Dios, Peru, collected by I. Semanez. Among the main characteristics present and constant in the diagnosis of the species are: (1) male fifth left swimming leg exopod 2 with plus oblique row of 5 denticles on distal digitiform process anteriorly; (2) male fifth right swimming leg exopod 1 broader than long 1.25x nearly; (3) male fifth right swimming leg exopod 1 with outer acute extension distally; (4) male fifth right swimming leg exopod 1 with trapezoidal lamella on posterior surface; and (5) male fifth right swimming leg exopod 2 with inner expansion medially. Among the conditions of *N. inflatus* divergent from Wright’s morphological set (1935; 1936; 1937) is the male fifth left swimming leg endopod 2-segmented. For *Notodiaptomus* originally (Kiefer, 1936) and amplification (Kiefer, 1956), the main divergences are for male fifth right swimming leg basis without posterior protrusion, and exopod 1 broader than long.

Considering that there have never been other public records for the occurrence of this taxon since its foundation and the impossibility of specifying the date of collection of the specimens examined here, it would be possible to suggest the extinction status for their organisms. Santos-Silva *et al*. (2015) drew previous attention to this fact and also highlighted that the originality of the organisms of this species is for a region of consistent sampling efforts of zooplankton, which reinforces the hypothesis of extinction of these organisms.

#### Notodiaptomus isabelae (Wright, 1936)

##### Synonymy

*Diaptomus isabelae* Wright, 1936a: 81, 82, pl.2, fig. 5; 1937: 76; 1938b: 563; Brehm, 1938: 30, 31; 1958a: 143; Brandorff, 1972: 50; Reid, 1991: 740. *Notodiaptomus isabelae*; Kiefer, 1956: 242; Bowman, 1973: 199; Brandorff, 1976: 616, fig. 2; Paggi, 1976a: 153, 154, figs. 1–25; Löffler, 1981: 15; Dussart & Defaye, 1983: 137; Dussart & Frutos, 1985: 307, figs. 3–6; Matsumura-Tundisi, 1986: 542, 547, 552, figs. 55–60, 100; Reid, 1987: 377, tab. 1; 1991: 740; José de Paggi & Paggi, 1988: 101, tab. 2; Lansac-Tôha *et al*., 1992: 43, 45, 47; 1995: 73; 1997: 140, 141, tab. 3; Sendacz, 1993: 35; 1997: 624, 625, tab. 2; Frutos, 1993: 112: tab. 3; Battistoni, 1995: 959; Rocha *et al*., 1995: 155, 156; Lima *et al*., 1996: 115, fig. 3; Bonecker *et al*., 1996: 897, fig. 3; Rocha & Matsumura-Tundisi, 1997: 293, tabs. 7, 9; Tundisi *et al*., 1997: 425, 434, tab. 11; Santos-Silva, 1998: 210; Santos-Silva, 2008: 31, fig. 6; Santos-Silva *et al*., 2015: 57–62, figs. 33– 35, 40, identification keys to male and female; Perbiche-Neves *et al*., 2015: 74–77, figs. 66–68, identification keys to male and female; Perbiche-Neves *et al*., 2020: 696-697, key to the Neotropical diaptomid, fig. 21.15 I. *Notodiaptomus* (*Notodiaptomus*) *isabelae*; Dussart, 1985a: 208.

##### Type locality

Lakes linked to São Francisco River, near to Jatobá, Pernambuco, Brazil.

##### Type material

Holotype is not designed in the original description; type material was not localized. By Santos-Silva *et al*. (2015) is probably does not existent more.

##### Material examined

Non-type material: 2 males, and 4 females, entire, in alcohol (USNM 267321), from the Dom Helvécio Lake, Doce River Valley (State Park), 03.XII.1993, J. Reid collector and designator; 1 male (INPA-COP031, slides a-h) and 1 female (INPA-COP032, slides a-h) were selected to be dissection on eight slides each and deposited in the Zoological Collection of the INPA, Brazil. Additional material examined: 2 males, and 2 females, entire, in alcohol, form the Brejo Grande, Sergipe State, São Francisco River. 21.IX.1993, Lucy M. Oliveira coll.; 4 males, and 5 females, entire, in alcohol, from the Gaspar Lake, Santa Catarina States, G. Perbiche-Neves coll., 2011.

##### Diagnosis

**(1)** male limit between fourth and fifth metasome segments ornamented with spinules row dorsally; **(2)** male fifth right swimming leg basis with inflated inner bilobed tumescence; **(3)** male fifth right swimming leg exopod 2 in elliptical form; **(4)** male fifth right swimming leg exopod 2 with inner expansion medially; **(5)** male fifth right swimming leg exopod 2 with rectilinear outer spine inserted sub-distally, and lesser 6x than length original segment; **(6)** female fourth metasome segment with dorsal rounded protuberance medially; **(7)** female limit between fourth and fifth metasome segments with spinules row on limit entirely; **(8)** female genital double-somite with dorsal suture at mid-length; **(9)** female genital double-somite with ventral vertical folds; **(10)** female right antennule with length exceeding the caudal setae.

##### Redescription

###### MALE

Body 1166 micrometers excluding caudal setae. Male body smaller and slenderer than female. Nerve axons myelinated. Prosome 6-segmented; widest at first metasome segment; without one line of setules at posterior margin; with spinules at least at one segment. Cephalosome anterior margin sub-triangular; with dorsal suture; incomplete; separate from first metasome segment. First metasome segment without sensilla. Second metasome segment with sensilla; 2 dorsally; of equal size. Third metasome segment with sensillae; 4 dorsally; of equal size; ornamented posterior margin; with spinules; as a row; double; dorsally. Fourth metasome segment with sensillae; 4 dorsally; of equal size; separated from the fifth metasome. Limit between fourth and fifth metasome segments ornamented; with spinules; as a row; on dorsal singly; on lateral singly. Fifth metasome segment with sensilla; 2 dorsally; Fifth metasome segment equal size; Fifth metasome segment without ornamentation; Fifth metasome segment without dorsal conical process; with epimeral plates. Epimeral plates asymmetrical. Right epimeral plates reduced, as rounded distal corner segment limit; with sensilla; at the apex of projection; without ornamentation. Left epimeral plate reduced, as rounded distal corner segment limit; with sensillae; at the apex of projection; without ornamentation.

##### Urosome

5-segmented; Urosome 5 - free segments. Genital somite symmetrical in dorsal view; with single aperture; located on left side; ventrolaterally on posterior rim; with sensillae; on both sides; one; at left lateral; posteriorly; one; at right rim; posteriorly; of equal size between then. Third urosome segment without spinules; without external seta. Fourth urosome segment without spinules; without sub-conical blunt dorsal-lateral process. Anal segment presence of dorsal sensillae; one on each side; medially inserted; presence of operculum; convex; covering the anal aperture fully. Caudal rami symmetrical; separated from anal segment; longer than wide; with setules; continuous on; inner side; each ramus bearing 6 caudal setae; 5 marginals; plumose; and 1 internal dorsally; straight; not reticulated main axis; outermost seta with outer spiniform process absent.

##### Oral appendices feature

Rostrum symmetrical; separated from dorsal cephalic shield; by complete suture; sensillae present; one pair; anteriorly inserted on surface tegument; with rostral filament; double; paired; extended; into point; with basal process; in ventral view, rounded on left side; without a smaller basal expansion on the right side.

##### Antennules

Asymmetrical. **Right antennules**. Uniramous; right antennule surpassing to genital segment; right antennule not extending beyond caudal rami.

Right antennule ancestral segment I and II separated. Ancestral segment II and III fused. Ancestral segment III and IV fused. Ancestral segment IV and V separated. Ancestral segment V and VI separated. Ancestral segment VI and VII separated. Ancestral segment VII and VIII separated. Ancestral segment VIII and IX separated. Ancestral segment IX and X separated. Ancestral segment X and XI separated. Ancestral segment XI and XII separated. Ancestral segment XII and XIII separated. Ancestral segment XIII and XIV separated. Ancestral segment XIV and XV separated. Ancestral segment XV and XVI separated. Ancestral segment XVI and XVII separated. Ancestral segment XVII and XVIII separated. Ancestral segment XVIII and XIX separated. Ancestral segment XIX and XX separated. Ancestral segment XX and XXI separated. Ancestral segment XXI and XXII fused. Ancestral segment XXII and XXIII fused. Ancestral segment XXIII and XXIV separated. Ancestral segment XXIV and XXV fused. Ancestral segment XXV and XXVI separated. Ancestral segment XXVI and XXVII separated. Ancestral segment XXVII and XXVIII fused.

Right antennule actual 22-segmented; geniculated; between the segment 18 and segment 19; with swollen and modified region; formed by 5 segments; between 13 and 17 segments. Actual segment 1 with seta; one element; straight; none larger than segment; without spinules; without vestigial seta; without conical seta; without modified seta; without spinous process; with aesthetasc; one element. Actual segment 2 with seta; three elements; of unequal size; straight; none larger than segment; without spinules; with vestigial seta; one element; without conical seta; without modified seta; without spinous process; with aesthetasc; one element. Actual segment 3 with seta; one element; one larger than segment; surpassing to distal margin; beyond three sequential segments; straight; blunt apex; without spinules; with vestigial seta; one element; without conical seta; without modified seta; without spinous process; with aesthetasc. Actual segment 4 with seta; one element; one larger than segment; surpassing to distal margin; straight; not beyond three sequential segments; without spinules; without vestigial seta; without conical seta; without modified seta; without spinous process; without aesthetasc. Actual segment 5 with seta; one element; straight; one larger than segment; surpassing to distal margin; not beyond three sequential segments; without spinules; with vestigial seta; one element; without conical seta; without modified seta; without spinous process; with aesthetasc; one element. Actual segment 6 with seta; one element; none larger than segment; straight; without spinules; without vestigial seta; without conical seta; without modified seta; without spinous process; without aesthetasc. Actual segment 7 with seta; one element; straight; one larger than segment; surpassing to distal margin; beyond three sequential segments; blunt apex; without spinules; without vestigial seta; without conical seta; without modified seta; without spinous process; with aesthetasc; one element. Actual segment 8 with seta; one element; straight; none larger than segment; without spinules; without vestigial seta; with conical seta; one element; reaching to middle-point of the sequent segment; without modified seta; without spinous process; without aesthetasc. Actual segment 9 with seta; two elements; of unequal size; straight; one larger than segment; surpassing to distal margin; beyond three sequential segments; blunt apex; without spinules; without vestigial seta; without conical seta; without modified seta; without spinous process; with aesthetasc; one element. Actual segment 10 with seta; one element; straight; none larger than segment; without spinules; without vestigial seta; without conical seta; with modified seta; presenting blunt apex; slender form; surpassing to distal margin; beyond of the sequential segment; parallel to antennule direction; without spinous process; without aesthetasc. Actual segment 11 with seta; one element; straight; one larger than segment; surpassing to distal margin; not beyond three sequential segments; without spinules; without vestigial seta; without conical seta; with modified seta; slender form; presenting blunt apex; surpassing to distal margin; beyond of the sequential segment; parallel to antennule direction; shorter length than homologous of actual segment 13; without spinous process; without aesthetasc. Actual segment 12 with seta; one element; straight; one larger than segment; surpassing to distal margin; not beyond three sequential segments; without spinules; without vestigial seta; with conical seta; one element; smaller than to segment 8; without modified seta; without spinous process; with aesthetasc; one element; absent internal perpendicular fission. Actual segment 13 with seta; one element; straight; one larger than segment; surpassing to distal margin; not beyond three sequential segments; without spinules; without vestigial seta; without conical seta; with modified seta; stout form; surpassing to distal margin; to the distal-point of the sequence segment; parallel to antennule direction; presenting bifid apex; without spinous process; with aesthetasc; one element. Actual segment 14 with seta; two elements; of unequal size; straight; one larger than segment; surpassing to distal margin; beyond three sequential segments; blunt apex; without spinules; without vestigial seta; without conical seta; without modified seta; without spinous process; with aesthetasc; one element. Actual segment 15 with seta; two elements; of unequal size; straight; not bifidform; none larger than segment; without spinules; without vestigial seta; without conical seta; without modified seta; with spinous process; on outer margin; surpassing distal margin; with aesthetasc; one element. Actual segment 16 with seta; two elements; of unequal size; plumose; one larger than segment; surpassing to distal margin; not beyond three sequential segments; not bifidform; without spinules; without vestigial seta; without conical seta; without modified seta; with spinous process; on outer margin; surpassing distal margin; unequal size to process on preceding segment; with aesthetasc; one element. Actual segment 17 with seta; two elements; of unequal size; straight; none larger than segment; bifidform; without spinules; without vestigial seta; without conical seta; with modified seta; one element; stout form; surpassing to distal margin; not beyond of the sequential segment; parallel to antennule direction; without spinous process; without aesthetasc. Actual segment 18 with seta; two elements; of equal size; straight; none larger than segment; without spinules; without vestigial seta; without conical seta; with modified seta; one element; stout form; surpassing distal margin; parallel to antennule direction; without spinous process; without aesthetasc. Actual segment 19 with seta; two elements; of unequal size; plumose; none larger than segment; without spinules; without vestigial seta; without conical seta; with modified seta; two elements; stout form; at least one bifid form; surpassing distal margin; parallel to antennule direction; without spinous process; with aesthetasc; one element. Actual segment 20 with seta; four elements; of unequal size; straight; one larger than segment; surpassing to distal margin; beyond three sequential segments; without spinules; without vestigial seta; without conical seta; without modified seta; without spinous process; without aesthetasc. Actual segment 21 with seta; two elements; of equal size; plumose; one larger than segment; surpassing to distal margin; greater 3x than original segment; without spinules; without vestigial seta; without conical seta; without modified seta; without spinous process; without aesthetasc. Actual segment 22 with seta; four elements; of equal size; one larger than segment; plumose; surpassing to distal margin; greater 3x than original segment; without spinules; without vestigial seta; without conical seta; without modified seta; without spinous process; with aesthetasc; one element.

##### Left antennules

Uniramous; Left antennule surpassing to prosome; Left antennule not extending beyond caudal rami. Ancestral segment I and II separated. Ancestral segment II and III fused. Ancestral segment III and IV fused. Ancestral segment IV and V separated. Ancestral segment V and VI separated. Ancestral segment VI and VII separated. Ancestral segment VII and VIII separated. Ancestral segment VIII and IX separated. Ancestral segment IX and X separated. Ancestral segment X and XI separated. Ancestral segment XI and XII separated. Ancestral segment XII and XIII separated. Ancestral segment XIII and XIV separated. Ancestral segment XIV and XV separated. Ancestral segment XV and XVI separated. Ancestral segment XVI and XVII separated. Ancestral segment XVII and XVIII separated. Ancestral segment XVIII and XIX separated. Ancestral segment XIX and XX separated. Ancestral segment XX and XXI separated. Ancestral segment XXI and XXII separated. Ancestral segment XXII and XXIII separated. Ancestral segment XXIII and XXIV separated. Ancestral segment XXIV and XXV separated. Ancestral segment XXV and XXVI separated. Ancestral segment XXVI and XXVII separated. Ancestral segment XXVII and XXVIII fused.

Left antennule actual 25-segmented; not-geniculated. Actual segment 1 with seta; one element; none larger than segment; straight; without spinules; without vestigial seta; without conical seta; without modified seta; without spinous process; with aesthetasc; one element. Actual segment 2 with seta; three elements; of equal size; none larger than segment; straight; without spinules; with vestigial seta; one element; without conical seta; without modified seta; without spinous process; with aesthetasc; one element. Actual segment 3 with seta; one element; one larger than segment; straight; surpassing to distal margin; beyond three sequential segments; without spinules; with vestigial seta; one element; without conical seta; without modified seta; without spinous process; with aesthetasc. Actual segment 4 with seta; one element; none larger than segment; straight; without spinules; without vestigial seta; without conical seta; without modified seta; without spinous process; without aesthetasc. Actual segment 5 with seta; one element; one larger than segment; straight; surpassing to distal margin; not beyond three sequential segments; without spinules; with vestigial seta; one element; without conical seta; without modified seta; without spinous process; with aesthetasc; one element. Actual segment 6 with seta; one element; none larger than segment; straight; without spinules; without vestigial seta; without conical seta; without modified seta; without spinous process; without aesthetasc. Actual segment 7 with seta; one element; one larger than segment; straight; surpassing to distal margin; beyond three sequential segments; without spinules; without vestigial seta; without conical seta; without modified seta; without spinous process; with aesthetasc; one element. Actual segment 8 with seta; one element; one larger than segment; straight; surpassing distal margin; without spinules; without vestigial seta; with conical seta; without modified seta; without spinous process; without aesthetasc. Actual segment 9 with seta; two elements; of unequal size; one larger than segment; straight; surpassing to distal margin; beyond three sequential segments; without spinules; without vestigial seta; without conical seta; without modified seta; without spinous process; with aesthetasc; one element. Actual segment 10 with seta; one element; none larger than segment; straight; without spinules; without vestigial seta; without conical seta; without modified seta; without spinous process; without aesthetasc. Actual segment 11 with seta; one element; one larger than segment; straight; surpassing to distal margin; beyond three sequential segments; without spinules; without vestigial seta; without conical seta; without modified seta; without spinous process; without aesthetasc. Actual segment 12 with seta; one element; one larger than segment; straight; surpassing distal margin; without spinules; without vestigial seta; with conical seta; without modified seta; without spinous process; with aesthetasc; one element. Actual segment 13 with seta; one element; none elongated; straight; surpassing distal margin; without spinules; without vestigial seta; without conical seta; without modified seta; without spinous process; without aesthetasc. Actual segment 14 with seta; one element; elongated; straight; surpassing to distal margin; beyond three sequential segments; without spinules; without vestigial seta; without conical seta; without modified seta; without spinous process; with aesthetasc; one element. Actual segment 15 with seta; one element; larger than segment; straight; surpassing to distal margin; not beyond three sequential segments; without spinules; without vestigial seta; without conical seta; without modified seta; without spinous process; without aesthetasc. Actual segment 16 with seta; one element; larger than segment; plumose; surpassing to distal margin; not beyond three sequential segments; without spinules; without vestigial seta; without conical seta; without modified seta; without spinous process; with aesthetasc; one element. Actual segment 17 with seta; one element; not larger than segment; straight; without spinules; without vestigial seta; without conical seta; without modified seta; without spinous process; without aesthetasc. Actual segment 18 with seta; one element; larger than segment; straight; surpassing to distal margin; beyond three sequential segments; without spinules; without vestigial seta; without conical seta; without modified seta; without spinous process; without aesthetasc. Actual segment 19 with seta; one element; not larger than segment; straight; surpassing distal margin; without spinules; without vestigial seta; without conical seta; without modified seta; without spinous process; with aesthetasc; one element. Actual segment 20 with seta; one element; not larger than segment; straight; surpassing distal margin; without spinules; without vestigial seta; without conical seta; without modified seta; without spinous process; without aesthetasc. Actual segment 21 with seta; one element; larger than segment; plumose; surpassing to distal margin; beyond three sequential segments; without spinules; without vestigial seta; without conical seta; without modified seta; without spinous process; without aesthetasc. Actual segment 22 with seta; two elements; of unequal size; one of them elongated; plumose; surpassing to distal margin; without spinules; without vestigial seta; without conical seta; without modified seta; without spinous process; without aesthetasc. Actual segment 23 with seta; two elements; of unequal size; one larger than segment; plumose; surpassing to distal margin; greater 3x than original segment; without spinules; without vestigial seta; without conical seta; without modified seta; without spinous process; without aesthetasc. Actual segment 24 with seta; two elements; of equal size; one larger than segment; plumose; surpassing to distal margin; greater 3x than original segment; without spinules; without vestigial seta; without conical seta; without modified seta; without spinous process; without aesthetasc. Actual segment 25 with seta; four elements; of equal size; elongated; plumose; surpassing to distal margin; 4 times larger than segment; without spinules; without vestigial seta; without conical seta; without modified seta; without spinous process; with aesthetasc; one element.

##### Antenna

Biramous. Antenna coxa separated from the basis; bearing seta; 1; on inner surface; at distal corner; reaching to the endopod 1. Antenna basis (fusion) separated from the endopodal segment; bearing seta; 2; on inner surface; at distal corner. Endopodal ancestral segment I and II separated. Ancestral segment II and III fused. Ancestral segment III and IV fused. Ancestral segment III and IV fully. Antenna endopod actual 2-segmented. Actual segment 1 not bilobate; with seta; two; on inner margin; with spinules; as a row; obliquely; on outer surface; with pore. Actual segment 2 bilobate; without discontinuity on outer cuticle; inner lobe bearing 8 setae; distally; outer lobe bearing 7 setae; distally; with spinules; as a patch; on outer surface. Antenna exopod ancestral segment I and II separated. Ancestral segment II and III fused. Ancestral segment III and IV fused. Ancestral segment IV and V separated. Ancestral segment V and VI separated. Ancestral segment VI and VII separated. Ancestral segment VII and VIII separated. Ancestral segment VIII and IX separated. Ancestral segment IX and X fused. Antenna exopod actual 7-segmented. Actual segment 1 single; elongated (width-length, equal or larger ratio 2:1); with seta; one; at inner surface. Actual segment 2 compound; elongated (larger width-length ratio 2:1); with seta; three; at inner surface. Actual segment 3 single; not elongated (lesser width-length ratio 2:1); with seta; one; at inner surface. Actual segment 4 single; not elongated (lesser width-length ratio 2:1); with seta; one; at inner surface. Actual segment 5 single; not elongated (lesser width-length ratio 2:1); with seta; one; at inner surface. Actual segment 6 single; not elongated (lesser width-length ratio 2:1); with seta; one; at inner surface. Actual segment 7 compound; elongated (larger or equal width-length ratio 2:1); with seta; one; at inner surface; and three; at distal surface.

##### Oral features

**Mandible**. Coxal gnathobase sclerotized; with lobe; prominent; on caudal margin; presence of cutting blade; with tooth-like prominence; two, distinctly; 1 acute; on caudal margin; and 1 triangular; on sub-caudal margin; without acute projection between the prominences; with additional spinules; as a row; on dorsal surface; with seta; 1; dorsally; on apical surface; with spinules; apicalmost. Mandible palps biramous; comprising the basis; with seta; four; differently inserted; first medially; reaching to beyond the endopod 1; second distally; third distally; fourth distally; on inner margin; none with setulose ornamentation. Mandible endopod 2-segmented. Mandible endopod 1 with lobe; bearing seta; four; distally inserted; without spinules. Mandible endopod 2 without lobe; bearing setae; nine elements; distally inserted; with spinules; as a row; double. Mandible exopod 4-segmented. Mandible exopod 1 with seta; one element; distally; on inner margin. Mandible exopod 2 with seta; one element; distally; on inner side. Mandible exopod 3 with seta; one element; distally; on inner side. Mandible exopod 4 with setae; three elements; on terminal region. **Maxillule**. Birramous. Maxillule 3-segmented. Maxillule praecoxa with praecoxal arthrite; bearing spines; fifteen elements; ten marginally; plus, five sub-marginally; with spinules; as a patch; on sub-marginal surface. Maxillule coxa with coxal epipodite; with conspicuous outer lobe; bearing setae; nine elements; with coxal endite; elongated (larger or equal width-length ratio 2:1); bearing setae; four elements. Maxillule basis with basal endite; double; first proximal; elongated (larger width-length ratio 2:1; separated from basis; with setae; four elements; distally inserted; second distal; fused to basis; not elongated (lesser width-length ratio 2:1); with setae; four elements; distally inserted; with setules; as a row; on inner side; basal exite present; with setae; one element; on outer surface. Maxillule endopod 1-segmented. Endopod 1 bilobate; first proximal; with setae; three elements; second distal; with setae; five elements. Maxillule exopod 1-segmented. Exopod 1 with setae; six elements; with setules; as a row; on inner side; spinules absent. **Maxilla**. Uniramous. Maxilla 5-segmented. Maxilla praecoxa fused to coxa; incompletely; distinct externally; with praecoxal endite; double; first elongated endite (larger or equal width length ratio 2:1); proximally inserted; with seta; straight, or plumose; 1 straight; 4 plumose; with spine; single; without spinules; without setule; second elongated endite (larger or equal width length ratio 2:1); distally inserted; with seta; plumose; 3 plumose; without spine; with spinules; as a row; on distal margin; with setule; as a row; on distal margin; absence of outer seta. Maxilla coxa with coxal endite; double; first elongated endite (larger or equal width); proximally inserted; with seta; plumose; 3 plumose; without spine; without spinules; with setules; as a row; on proximal margin; second elongated endite (larger or equal width); distally inserted; with seta; plumose; 3 plumose; without spine; without spinules; with setules; as a row; on proximal margin; absence of outer seta. Maxilla basis with basal endite; single; elongated (larger or equal width-length ratio 2:1); with seta; plumose; 3 plumose; without spinules; absence of outer seta. Maxilla endopod 2-segmented. Endopod 1 with seta; 2 plumoses; without spine; without spinules; without setules. Maxilla endopod 2 with seta; 2 plumose; without spine; without spinules; without setules. **Maxilliped**. Uniramous; Maxilliped 8-segmented. Maxilliped praecoxa fused to coxa; incompletely; distinct internally; with praecoxal endite; not elongated (lesser width-length ratio 2:1); distally inserted; with seta; 1 straight; with spinules; as a row; single; on basal surface; without setules. Maxilliped coxa with coxal endite; three coxal endite; first elongated (larger or equal width); proximally inserted; with seta; 2 plumose; with spinules; as a patch; single; on apical surface; without setules; second not elongated (lesser width-length ratio 2:1); medially inserted; with seta; 3 plumose; with spinules; as a row; single; on medial surface; without setules; third elongated (larger or equal width length ratio 2:1); distally inserted; with seta; 3 plumose; none reaching to beyond of the basis; with spinules; as a row; single; on basal surface; without setules; with lobe; prominence; at inner distal angle; ornamented; with spinules; continuously on margin. Maxilliped basis without basal endite; with seta; 3 plumose; with spinules; as a row; single; on medial surface; with setules; as a row; single; on inner margin. Maxilliped endopod segment 6-segmented. Endopod 1 with seta; 2 plumose; on inner surface. Endopod 2 with seta; 3 plumose; on inner surface. Endopod 3 with seta; 2 plumose; on inner surface. Endopod 4 with seta; 2 plumose; on inner surface. Endopod 5 with seta; 2 plumose; on inner surface, or on outer surface; outer seta absent. Endopod 6 with seta; 4 plumose; on inner surface, or on outer surface.

##### Swimming legs features

**First swimming legs.** Symmetrical; biramous. First swimming legs intercoxal plate without seta. First swimming legs praecoxa absent. First swimming legs coxa with seta; one; straight; distally inserted; on inner surface; surpassing to first endopodal segment; with setules; two group; as a patch; on inner margin; and as a row; double; on anterior surface; outerly; without spinules; without spine. First swimming legs basis without seta; with setules; as a patch; single; on outer surface; without spinules; without spine. First swimming legs endopod 2-segmented. Endopod 1 with seta; straight; restricted; to inner surface; one element; without spine; with setules; as a row; single; continuously; on outer surface; without spinules; absence of Schmeil’s organ. Endopod 2 with seta; unrestricted; three on inner surface; one on outer surface; two on distal surface; straight; without spine; with setules; as a row; single; continuously; on outer surface; without spinules; absence of Schmeil’s organ. Endopod 3 absence. First swimming legs exopod 1 with seta; restricted; 1 on inner surface; with spine; 1; stout; smaller than original segment; serrated; on inner side; continuously; with setules; as a row; single; as a row; innerly. First swimming legs exopod 2 with seta; restricted; 1 on inner surface; straight; without spine; with setules; as a row; single; continuously; on inner margin, or on outer margin; without spinules. First swimming legs exopod 3 with setule; as a row; single; continuously; on outer surface; without spinules; with seta; unrestricted; 2 on inner surface; 2 on terminal surface; with spine; 2; unequal size; first no longer 2x than origin segment; stout; serrated; on inner side, or on outer side; equally; second longer 3x than origin segment; slender; serrated; on outer side; with ornamentation on non-serrated side; by setules. **Second swimming legs**. Symmetrical; Second swimming legs biramous. Second swimming legs intercoxal plate without seta. Second swimming legs praecoxa present; located laterally. Second swimming legs coxa with seta; straight; distally inserted; on inner surface; surpassing to basal segment; without setules; without spinules; without spine. Second swimming legs basis without seta; without setules; without spinules; without spine. Second swimming legs endopod 3-segmented. Endopod 1 with seta; straight; restricted; one on inner surface; without spine; with setules; as a row; single; continuously; on outer surface; without spinules; absence of Schmeil’s organ. Endopod 2 with seta; straight; unrestricted; two on inner surface; without spine; with setules; as a row; single; continuously; on outer side; without spinules; presence of Schmeil’s organ; on posterior surface. Endopod 3 with seta; straight; unrestricted; three on inner surface; two on outer surface; two on distal surface; without spine; without setules; with spinules; as a row; double; distally inserted; at anterior surface; absence of Schmeil’s organ. Second swimming legs exopod 1 with seta; restricted; one on inner surface; with spine; 1; stout; not reaching to distal-third of the exopod 2; serrated; on inner side, or on outer side; with setules; as a row; single; continuously; on inner side; without spinules; absence of Schmeil’s organ. Exopod 2 with seta; unrestricted; one on inner surface; with spine; 1; stout; not surpassing the exopod 3; serrated; on inner side, or on outer side; with setules; as a row; single; continuously; on inner surface; without spinules; absence of Schmeil’s organ. Exopod 3 with seta; plurimarginal; three on inner surface; two on terminal surface; with spine; 2; unequal size; first no longer 2x than origin segment; stout; serrated; on inner side, or on outer side; equally; second longer 2x than origin segment; slender; serrated; on outer side; with ornamentation on non-serrated side; of setules; setules on outer surface; as a row; single; continuously; on inner surface; with spinules; as a row; single; distally inserted; at anterior surface; absence of Schmeil’s organ. **Third swimming legs**. Symmetrical; Third swimming legs biramous. Third swimming legs intercoxal plate without seta. Third swimming legs praecoxa present; not laterally located. Third swimming legs coxa with seta; straight; distally inserted; on inner surface; surpassing to first endopodal segment; without setules; without spinules; without spine. Third swimming legs basis without seta; without setules; without spinules; without spine. Third swimming legs endopod 3-segmented. Endopod 1 with seta; restricted; one on inner surface; without spine; without setules; without spinules; absence of Schmeil’s organ. Endopod 2 with seta; restricted; two on inner surface; straight; without spine; without setules; without spinules; absence of Schmeil’s organ. Endopod 3 with seta; straight; plurimarginal; two on inner surface; two on outer surface; three on terminal surface; without spine; without setules; with spinules; as a row; distally inserted; double; at anterior surface; absence of Schmeil’s organ. Third swimming legs exopod 1 with seta; restricted; straight; one on inner surface; with spine; 1; stout; not reaching to the distal-third of the exopod 2; serrated; equally; on inner surface, or on outer surface; with setules; as a row; single; continuously; on inner surface; without spinules; absence of Schmeil’s organ. Exopod 2 with seta; straight; restricted; one on inner surface; with spine; 1; stout; not reaching out to exopod 3; serrated; on inner side, or on outer side; equally; with setules; as a row; single; continuously; on inner side; without spinules; absence of Schmeil’s organ. Exopod 3 without setules; with spinules; as a row; single; distally inserted; at anterior surface; with seta; straight; unrestricted; three on inner surface; two on terminal surface; with spine; 2; unequal size; first no longer 2x than origin segment; stout; serrated; on inner side, or on outer side; equally; second longer 2x than origin segment; slender; serrated; on outer side; with ornamentation on non-serrated side; of setules; absence of Schmeil’s organ. **Fourth swimming legs**. Symmetrical; biramous. Intercoxal plate without sensilla. Praecoxa present. Coxa with seta; distally inserted; on inner margin; reaching out to endopod 1; without spinules; setules absent. Basis with seta; one; medially inserted; on posterior surface; smaller than the original segment; without setules; without spinules; without spine. Fourth swimming legs endopod 3-segmented. Endopod 1 with seta; one; restricted; on inner surface; without spine; without setules; without spinules; absence of Schmeil’s organ. Endopod 2 with seta; restricted; two on inner side; without spine; with setules; as a row; single; continuously; on outer surface; without spinules; absence of Schmeil’s organ. Endopod 3 with seta; unrestricted; two on inner surface; two on outer surface; three on distal surface; without spine; without setules; with spinules; as a row; double; distally inserted; at anterior surface; absence of Schmeil’s organ. Fourth swimming legs exopod 1 with seta; restricted; one on inner surface; with spine; 1; stout; not reaching out to distal-third of the exopod 2; serrated; on inner side, or on outer side; equally; with setules; as a row; single; continuously; on inner surface; without spinules; absence of Schmeil’s organ. Exopod 2 with seta; restricted; one on inner surface; with spine; 1; stout; not reaching the end of exopod 3; serrated; on inner side, or on outer side; equally; with setules; as a row; single; continuously; on inner surface; without spinules; absence of Schmeil’s organ. Exopod 3 without setules; with spinules; as a row; single; distally inserted; at anterior surface; with seta; unrestricted; three on inner surface; two on distal surface; with spine; 2; unequal size; first no longer 2x than origin segment; stout; serrated; on inner side, or on outer side; equally; second longer 2x than origin segment; slender; serrated; on outer side; without ornamentation on non-serrated side; absence of Schmeil’s organ.

##### Fifth swimming legs features

Asymmetrical. Fifth swimming leg intercoxal plate with length not equal or greater than width on 1.5x; with irregular proximal margin; discontinuous to; the anterior margin of the left coxa, or the anterior margin of the right coxa; posterior sensilla on the right lateral absent. **Fifth left swimming leg**. Fifth left swimming leg biramous; leg reaching first right exopod segment; proximally. Fifth left swimming leg praecoxa present; rudimentary; separated from the coxae; without ornamentation. Fifth left swimming leg coxa concave inner side; without teeth-like structures; with process; conical; on posterior surface; outer side; distally inserted; not projecting over basis; with sensilla; stout; triangular; at apex; longer 2x than insertion basis; without swelling; without seta; without spinules. Fifth left swimming leg basis sub-cylindrical; unequal size between inner and outer side; shorter outer than inner side; with concave inner side; rounded internal proximal expansion absent; without outgrowth; with groove; deep; obliquely; on posterior surface; not reaching the endopodal lobe; not ornamented; absence of protuberance; with seta; outerly inserted; no longer 2x than origin segment; absence of minutely granular. Fifth left swimming leg endopod segments 1 and 2 fused; segments 2 and 3 fused; 1-segmented; stout; separated from the basis; ornamented; on inner side; with spinules; more than four elements; as a row; terminally; row of setules absent; without seta. Fifth left swimming leg exopod segments 1 and 2 separated; segments 2 and 3 fused; 2-segmented; stout; separated from the basis. Fifth left swimming leg exopod 1 sub-triangular; longer than broad; equal size between inner and outer side; rectilinear inner side; convex outer side; without swelling; without marginal extension; without process; with lobe; single; circular; medially inserted; on inner side; covered; by setules; without outer spine; absence seta. Fifth left swimming leg exopod 2 digitiform; longer than broad; equal size between inner and outer side; disform inner side; with rectilinear outer side; setulose pad present; prominently rounded; proximally; on inner side; inflated medial region absent; distal process present; digitiform; non denticulate; without transverse row of denticles; none oblique row of 5 denticles; not innerly directed; with seta; spiniform; ornamented by spinules; not surpassing the distal-point of the segment; without outer spine; terminal claw absent.

##### Fifth right swimming leg

Biramous. Fifth right swimming leg praecoxa present; separated from the coxae; without ornamentation. Fifth right swimming leg coxa convex inner side; without teeth-like structures; with process; rounded; distally inserted; on posterior surface; closest to the outer rim; projecting over basis; not beyond the first third; without triangular protuberance innerly; with sensilla; slender; at apex; no longer 2x than basal insertion; without marginal extension; without seta; without spinules. Fifth right swimming leg basis trapezoidal; unequal size between inner and outer side; shorter outer than inner side; convex inner side; tumescence present; inflated; bilobed; unrestricted on inner surface; without protuberance; absence of distinct minutely granular; additional inner process absent; with posterior groove; shallow; obliquely; not reaching the endopodal lobe; ornamented; with tubercles; throughout of the inner border, or throughout of the outer border; with seta; outerly inserted; on anterior surface; no longer 2x than origin segment; posterior protrusion present; distal process absent. Fifth right swimming leg with endopodite present; fused to basis; on anterior surface; ancestral segments 1 and 2 fused; ancestral segments 2 and 3 fused; stout; ornamented; with spinules; as a row; on inner side; terminally; without seta. Fifth right swimming leg exopod segments 1 and 2 separated; segments 2 and 3 fused; 2-segmented; stout; separated from the basis. Fifth right swimming leg exopod 1 sub-cylindrical; longer than broad; nearly 1.25 times; unequal size between both sides; shorter inner than outer side; convex inner side; rectilinear outer side; with marginal extension; sub-triangular; distally inserted; at outer rim; spinules absent; with process; triangular; arched; internally directed; sharp tip; sclerotized; without ornamentation; distally inserted; at posterior surface; projecting over next segment; without outer spine; without seta; internal prominence absent; lamella on posterior surface absent. Fifth right swimming leg exopod 2 elliptical; longer than broad; nearly 2 times; equal size between both sides; disform inner side; convex outer side; without posterior proximal swelling; inner-posterior process absent; with marginal expansion; medial; innerly; curved ridge on distal posterior surface present; chitinous knobs absent; with outer spine; inserted sub-distally; rectilinear; ornamented innerly; by spinules; as a row; not ornamented outerly; sharp tip; without apparent curve; lesser than the length of the exopod 2; beyond to 2 times its size; 6x; sensilla absent; terminal claw present; equal or longer 1.5 times than insertion segment; sclerotized; rectilineal; with conspicuous curve; proximally; ornamented innerly; by spinules; as a row; partially on extension; medially, or distally; not ornamented outerly; sharp tip; not curved tip; without medial constriction; hyaline process absent.

##### FEMALE

Body longer and wider than male. Widest at posterior cephalosome. Distal margin of the prosomal segments without one line of setules at posterior margin. Prosome segments with spinules at least at one prosomal segment. Fourth metasome segment presence of dorsal protuberance; rounded; inserted medially; without posterior process; without anterior process; fourth metasome segment without proximal sensillae present. Fourth and fifth metasome segments fused; totally. Limit between fourth and fifth metasome segments ornamented; with spinules; as a row; single; complete; same size; entirely over limit (lateral, dorsal). **Fifth metasome segment**. Fifth metasome segment with sensilla; dorsally; 2 elements; with epimeral plates. Epimeral plates symmetrical. Right epimeral plates prominent, as projections; one posterior-laterally directed; not reaching half length of the genital segment; with sensilla at the apex; dorsal-posterior sensilla present; stout; without ornamentation. Left epimeral plate without expansion.

##### Urosome

3-segmented. **Genital double-somite**. Asymmetrical in dorsal view; longer than broad; longer than other urosomites combined; dorsal suture at mid-length present; not covered by spinules; with swelling; rounded; unequal size; greater left than right; anteriorly; with sensillae; on both sides; one; stout; with robust apex; at left lateral; not on lobular base; medially; one; stout; at right lateral; not on lobular base; anteriorly; with robust apex; of equal size between then; lateral protuberance absent; with right posterior rim expanded; over next segment; without slender sensilla on each posterior rim; without posterior-dorsal process. Genital double-somite opercular pad present; broader than longer; symmetrical; development laterally; expanded posteriorly; covering partially; double gonoporal slit; located ventrally; with arthrodial membrane; inserted anteriorly; post-genital process absent; disto-ventral tumescence absent; ventral vertical folds present; dorsal sensilla absent. Second urosome segment without ventral fusion to anal segment; right distal process absent. Caudal rami patch of setules on outer surface absent; patch of spinules on outer surface absent.

##### Oral appendices feature

Rostrum basal process absent. **Antennules**. Symmetrical. Right antennule surpassing to genital double-segment; extending beyond caudal rami. Right antennule exceeding the caudal setae. Right antennule ornamentation pattern equals to male left antennule; mostly. Actual segment 13 without seta; without aesthetasc. Actual segment 14 without seta; without aesthetasc. Actual segment 15 without seta; without aesthetasc. Actual segment 16 without seta; without aesthetasc. Actual segment 17 without seta. Actual segment 18 without seta.

##### Fifth swimming legs

Symmetrical; Fifth swimming legs biramous. Fifth swimming legs intercoxal plate longer than wide; separated from the legs. Fifth swimming legs praecoxa with sclerite praecoxal; separated from the coxae; without ornamentation. Fifth swimming legs coxa with process; conical; at the outer rim; distally; sensilla present; stout; at apex; projecting over basal segment; no longer 2x than basal insertion; marginal extension absent; without swelling; without seta; without spinules. Fifth swimming legs basis sub-triangular; unequal size between inner and outer sides; shorter outer than inner side; with convex inner side; without proximal inner outgrowth; without groove; with distal extension; on posterior surface; with seta; outerly inserted; on anterior surface; longer 2x than origin segment; not reaching to exopod 1 distally. Fifth swimming legs endopod segments 1 and 2 fused; segments 2 and 3 fused; 1-segmented; stout; separated from the basis; absent discontinuity cuticle; with spinules; as a row; single; non-oblique; sub-terminally; at anterior surface; with seta; double; one medially; on posterior surface; rectilinear; one distally; on posterior surface; arched; of unequal size; distal seta longer than medial seta. Fifth swimming legs exopod segments 1 and 2 separated; segments 2 and 3 separated; 3-segmented; separated from the basis. Fifth swimming legs exopod 1 sub-cylindrical; longer than wide; longer or equal than 2 times; with unequal size between inner and outer side; shorter inner than outer side; with convex inner side; with rectilinear outer side; without swelling; without marginal extension; without posterior process; without spine; without seta. Fifth swimming legs exopod 2 sub-cylindrical; longer than broad; longer or equal than 2 times; without swelling; without marginal extension; without process; without lobe; with spine; inserted laterally; rectilinear; without ornamentation; sharp tip; equal size or larger than next segment; without seta. Fifth swimming legs exopod 3 cylindrical; longer than wide; without swelling; without process; without lobe; without spine; with seta; double; inserted terminally; unequal size between them; outer seta smaller than inner; nearly 3 times; outer seta not ornamented by setules; without ornamentation; presence of terminal claw; sclerotized; arched; externally directed; convex inner side; with ornamentation; of denticles; as a row; on surface partially; at medial region; concave outer side; with ornamentation; of denticles; as a row; on surface partially; at medial region; blunt tip; 6 times longer than origin segment.

##### Distribution records

###### BRAZIL

**Pernambuco**: two ponds near to Jatobá, each connected to São Francisco River on flood season (Wright, 1936); “açudes” (Matsumura-Tundisi, 1986). **Minas Gerais**: Bonita Lagoon, Doce River Valley (Matsumura-Tundisi, 1986); Doce River on Belo Oriente, near to Ipatinga, upstream of its confluence with the Santo Antônio River (Bonecker *et al*., 1996); Palmeiras Lake, Almacega, Carvão, and Azeite, pond Fundo, Águas Claras, Jacaré, Ariranha, Palmeirinha, and Ferrugem, Doce River Valley (Tundisi *et al*., 1997). **Mato Grosso do Sul**: floodplain in upper Paraná River, near to Nova Andradina (Lansac-Tôha *et al*., 1992); Pousada das Garças Lake, floodplain in upper Paraná River (Lansac-Tôha *et al*., 1995); Guaraná Lake, and Baía River, floodplain of the Paraná River (Lima *et al*., 1996); Pato Lake, Guaraná, Pousada das Garças, and Fechada, Baía River, Ivinhema, and Paraná (Lansac-Tôha *et al*., 1997). **São Paulo**: Comprida Lagoom 1, and 2, and Jota Lagoon, upper Paraná River (Sendacz, 1997). **Paraná**: floodplain of the upper Paraná River, near to Porto Rico (Lansac-Tôha *et al*., 1992). ARGENTINA. **Corrientes**: Turbia Lagoon, Del Cerrito Island, Paraná River (Dussart & Frutos, 1985). **Santa Fé**: Madrejón Don Felipe; Madrejón El Negro, Carbajal Isalnd; Santa Fé River (Paggi, 1976a); Santa Fé River (José de Paggi & Paggi, 1988). PARAGUAY. Paraná River at Yaciretá Reservoir (Perbiche-Neves *et al*., 2015).

##### Habitat

Habitat in freshwaters: rivers, pools, lakes.

##### Remarks

The taxon was presented to science from organisms of lacustrine systems in the Brazilian Northeast, believed for years to be exclusive to this region. Forwards studies extended its current occurrence to the Southern Neotropics, specifically to river environments at Santa Fé in Argentina (Paggi & Paggi, 1988). *D.* (s.l.) *isabelae* was integrated into the *nordestinus* complex during its first expansion (Wright, 1936), undergoing the current recombination in the amplification of *Notodiaptomus* (Kiefer, 1956).

In the original description Wright suggested among the differential attributes of the species in *nordestinus*, exactly the morphological criterion used for its inclusion: male fifth right swimming leg exopod 1 with distal process posteriorly. Santos-Silva *et al*. (2015) when reviewing the Wright’s complex corroborated this characteristic for the species, but attributed broad convergence to other taxa that are part of the grouping, which would fatefully integrate it into the group, but would not be enough for a morphological differentiation. Among the other 4 attributes highlighted in the founding of the species are: (1) male fifth right swimming leg basis with inflated inner bilobed tumescence; (2) male fifth right swimming leg exopod 2 in elliptical form; (3) male fifth right swimming leg exopod 2 with “short” outer spine; and (4) male fifth right swimming leg exopod 2 with curved narrow terminal claw.

For the present effort only attribute 4 cannot be corroborated. The males examined from the Dom Helvécio Lake, Doce River Valley have the rectilinear terminal claw on exopod 2 of the male fifth right leg. Additionally, the subjectivity originally highlighted for “short” outer spine was here reinterpreted as lesser 6x than length original segment and, through this objective criterion, can be confirmed. Perbiche-Neves *et al*. (2015) offered revision for the species and presented illustrated the internally curved terminal claw and interpreted the inflated inner bilobed tumescence on basis as “two” small expansions present proximally on internal margin of right basis.

In the study of the diaptomids of the “La Plata” River Basin (Perbiche-Neves *et al*., 2015), the individuals examined do not have setae on male left antennule actual segment 11, and ornamentation on male fifth right swimming leg basis posteriorly was absent (*vide*, Perbiche-Neves *et al*., 2015, p. 12, dicotomic key).

In the present study, all males examined had left antennule actual segment 11 with seta, and tubercules on inner, and outer border of the fifth right swimming leg basis posteriorly. However, Perbiche-Neves *et al*. (2015) added observations corroborated in the present effort, such as: (1) male limit between fourth and fifth metasome segments ornamented with spinules row dorsally; and (2) female limit between fourth and fifth metasome segments with spinules row on limit entirely.

Additionally, we present for females of the species divergent characteristics relative to the type-species of *Notodiaptomus*: (1) female fourth metasome segment with dorsal rounded protuberance medially; (2) female genital double-somite with dorsal suture at mid-length; and (3) female genital double-somite with ventral vertical folds. Santos-Silva *et al*. (2015) had already verified this condition. In the redescription of the species in Paggi (1976), no mention was made of the attributes 2 and 3 highlighted, as well as Perbiche-Neves *et al*. (2015) recently. Among the characteristics applied to the grouping *nordestinus* (Wright, 1935; 1936; 1937) and *Notodiaptomus* (Kiefer, 1936; 1956), *N. isabelae* shares them all. However, the distinction lies in the converging attributes we identify in the type species and congeners. Of these, the main divergences are for male fifth right swimming leg: (1) basis with bilobate inner intumescence; (2) basis with posterior groove not reaching to endopod; (3) exopod 2 with outer spine 6x lesser than length original segment. For female: (1) fourth metasome segment with dorsal protuberance; (2) limit between fourth and fifth metasome segment with ornamentation; and (3) fifth swimming legs endopod without discontinuity cuticle.

#### Notodiaptomus jatobensis (Wright, 1936)

##### Synonymy

*Diaptomus jatobensis* Wright, 1936a: 82, pl. 2, fig. 4; 1937: 76; 1938b: 563; Brandorff, 1972: 50; Cipólli and Carvalho, 1973: 95, 97, 98, 101, tab. 2; Reid, 1991: 740. *Notodiaptomus jatobensis*; Kiefer, 1956: 242; Brehm, 1958a: 145; Brandorff, 1976: 616, fig.2; Löffler, 1981: 15; Dussart & Defaye, 1983: 137; Robertson & Hardy, 1984: tab. 3; Matsumura-Tundisi, 1986: 542, 547, figs. 73–77, 100; Reid, 1987: 377; 1991: 740; Sendacz, 1993: 35; 1997: 624, 625, tab. 2; Rocha *et al*., 1995: 155, 156; Santos-Silva, 1998: 211; Santos-Silva, 2008: 31–32, fig. 6; Santos-Silva *et al*., 2015: 62–65, figs. 36–38, 40; Perbiche-Neves *et al*., 2020: 696-697, key to the Neotropical diaptomid, fig. 21.15 J. *Notodiaptomus (Notodiaptomus) jatobensis*; Dussart, 1985a: 208.

##### Type locality

The original description indicates a lake at the Itaparica waterfall on the São Francisco River, near the city of Jatobá, State of Pernambuco, Brazil.

##### Type material

Holotype not specified in the original description. According to Santos-Silva *et al*. (2015) the type material no exists more probably.

##### Material examined

Topotype: 2 males, and 1 female, entire, in alcohol from the Lake at the margin of the Sao Francisco River (7 km to Iboticama), Bahia State, Brazil, 30.VI.1980, collected by L. Elmoor and specified by J. Reid (USNM 241584). 1 male (INPA-COP033, slides a-h) and 1 female (INPA-COP034, slides a-h) were selected to be dissection on eight slides each and deposited in the Zoological Collection of the INPA, Brazil.

##### Diagnosis

**(1)** male limit between fourth and fifth metasome segments with single spinules row laterally; **(2)** male right antennule actual segments 22 and 25 with five setae; **(3)** male fifth left swimming leg with length surpassing first right exopod segment; **(4)** male fifth left swimming leg exopod 1 with proximal single process posteriorly; **(5)** male fifth right swimming leg coxa with triangular protuberance on conical process innerly; **(6)** male fifth right swimming leg basis with inflated inner tumescence, not bilobed; **(7)** male fifth right swimming leg basis without posterior protrusion; **(8)** male fifth right swimming leg exopod 2 with 3 chitinous knobs posteriorly; **(9)** female limit between fourth and fifth metasome segments with double spinules row of different size; **(10)** female right epimeral plate without dorsal-posterior sensilla; **(11)** female genital double-somite with rounded expansion like a bulging dilation; **(12)** female genital double-somite with dorsal suture at mid-length; **(13)** female genital double-somite with ventral vertical folds **(14)** female fifth swimming legs basis with outer seta longer 2x than origin segment, reaching to middle-point exopod 1.

##### Redescription

###### MALE

Body 1067 micrometers excluding caudal setae. Male body smaller and slenderer than female. Nerve axons myelinated. Prosome 6-segmented; widest at first metasome segment; without one line of setules at posterior margin; with spinules at least at one segment. Cephalosome anterior margin rounded; with dorsal suture; incomplete; separate from first metasome segment. First metasome segment without sensilla. Second metasome segment without sensilla. Third metasome segment without sensillae; non-ornamented posterior margin. Fourth metasome segment without sensillae; separated from the fifth metasome. Limit between fourth and fifth metasome segments ornamented; with spinules; as a row; on lateral singly; same size. Fifth metasome segment with sensilla; 2 dorsally; Fifth metasome segment equal size; Fifth metasome segment without ornamentation; Fifth metasome segment without dorsal conical process; with epimeral plates. Epimeral plates asymmetrical. Right epimeral plates prominent, as projections; one projection; posterior-laterally directed; not reaching half length of the genital segment; with sensilla; at the apex of projection; without ornamentation. Left epimeral plate prominent, as projection; one projection; posterior-dorsally directed; reaching half length of the genital segment; with sensillae; at the apex of projection; without ornamentation.

##### Urosome

5-segmented; Urosome 5-free segments. Genital somite asymmetrical in dorsal view; with single aperture; located on left side; ventrolaterally on posterior rim; with sensillae; on both sides; one; at left lateral; posteriorly; one; at right rim; posteriorly; of unequal size between then; right bigger than left. Third urosome segment without spinules; without external seta. Fourth urosome segment without spinules; without sub-conical blunt dorsal-lateral process. Anal segment presence of dorsal sensillae; one on each side; medially inserted; presence of operculum; convex; covering the anal aperture fully. Caudal rami symmetrical; separated from anal segment; longer than wide; with setules; continuous on; inner side; each ramus bearing 6 caudal setae; 5 marginals; plumose; and 1 internal dorsally; straight; not reticulated main axis; outermost seta with outer spiniform process absent.

##### Oral appendices feature

Rostrum symmetrical; separated from dorsal cephalic shield; by complete suture; sensillae present; one pair; anteriorly inserted on surface tegument; with rostral filament; double; paired; extended; into point; with basal process; in ventral view, rounded on left side; without a smaller basal expansion on the right side.

##### Antennules

Asymmetrical. **Right antennules**. Uniramous; right antennule surpassing to genital segment; right antennule extending beyond caudal rami.

Right antennule ancestral segment I and II separated. Ancestral segment II and III fused. Ancestral segment III and IV fused. Ancestral segment IV and V separated. Ancestral segment V and VI separated. Ancestral segment VI and VII separated. Ancestral segment VII and VIII separated. Ancestral segment VIII and IX separated. Ancestral segment IX and X separated. Ancestral segment X and XI separated. Ancestral segment XI and XII separated. Ancestral segment XII and XIII separated. Ancestral segment XIII and XIV separated. Ancestral segment XIV and XV separated. Ancestral segment XV and XVI separated. Ancestral segment XVI and XVII separated. Ancestral segment XVII and XVIII separated. Ancestral segment XVIII and XIX separated. Ancestral segment XIX and XX separated. Ancestral segment XX and XXI separated. Ancestral segment XXI and XXII fused. Ancestral segment XXII and XXIII fused. Ancestral segment XXIII and XXIV separated. Ancestral segment XXIV and XXV fused. Ancestral segment XXV and XXVI separated. Ancestral segment XXVI and XXVII separated. Ancestral segment XXVII and XXVIII fused.

Right antennule actual 22-segmented; geniculated; between the segment 18 and segment 19; with swollen and modified region; formed by 5 segments; between 13 and 17 segments. Actual segment 1 with seta; one element; straight; none larger than segment; without spinules; without vestigial seta; without conical seta; without modified seta; without spinous process; with aesthetasc; one element. Actual segment 2 with seta; three elements; of unequal size; straight; none larger than segment; without spinules; with vestigial seta; one element; without conical seta; without modified seta; without spinous process; with aesthetasc; one element. Actual segment 3 with seta; one element; one larger than segment; surpassing to distal margin; beyond three sequential segments; straight; blunt apex; without spinules; with vestigial seta; one element; without conical seta; without modified seta; without spinous process; with aesthetasc. Actual segment 4 with seta; one element; one larger than segment; surpassing to distal margin; straight; not beyond three sequential segments; without spinules; without vestigial seta; without conical seta; without modified seta; without spinous process; without aesthetasc. Actual segment 5 with seta; one element; straight; one larger than segment; surpassing to distal margin; not beyond three sequential segments; without spinules; with vestigial seta; one element; without conical seta; without modified seta; without spinous process; with aesthetasc; one element. Actual segment 6 with seta; one element; none larger than segment; straight; without spinules; without vestigial seta; without conical seta; without modified seta; without spinous process; without aesthetasc. Actual segment 7 with seta; one element; straight; one larger than segment; surpassing to distal margin; beyond three sequential segments; blunt apex; without spinules; without vestigial seta; without conical seta; without modified seta; without spinous process; with aesthetasc; one element. Actual segment 8 with seta; one element; straight; none larger than segment; without spinules; without vestigial seta; with conical seta; one element; not reaching to middle-point of the sequent segment; without modified seta; without spinous process; without aesthetasc. Actual segment 9 with seta; two elements; of unequal size; straight; one larger than segment; surpassing to distal margin; beyond three sequential segments; blunt apex; without spinules; without vestigial seta; without conical seta; without modified seta; without spinous process; with aesthetasc; one element. Actual segment 10 with seta; one element; straight; none larger than segment; without spinules; without vestigial seta; without conical seta; with modified seta; presenting blunt apex; slender form; surpassing to distal margin; beyond of the sequential segment; parallel to antennule direction; without spinous process; without aesthetasc. Actual segment 11 with seta; one element; straight; one larger than segment; surpassing to distal margin; not beyond three sequential segments; without spinules; without vestigial seta; without conical seta; with modified seta; slender form; presenting blunt apex; surpassing to distal margin; beyond of the sequential segment; parallel to antennule direction; shorter length than homologous of actual segment 13; without spinous process; without aesthetasc. Actual segment 12 with seta; one element; straight; one larger than segment; surpassing to distal margin; not beyond three sequential segments; without spinules; without vestigial seta; with conical seta; one element; smaller than to segment 8; without modified seta; without spinous process; with aesthetasc; one element; absent internal perpendicular fission. Actual segment 13 with seta; one element; straight; one larger than segment; surpassing to distal margin; not beyond three sequential segments; without spinules; without vestigial seta; without conical seta; with modified seta; stout form; surpassing to distal margin; to the distal-point of the sequence segment; parallel to antennule direction; presenting bifid apex; without spinous process; with aesthetasc; one element. Actual segment 14 with seta; two elements; of unequal size; straight; one larger than segment; surpassing to distal margin; beyond three sequential segments; blunt apex; without spinules; without vestigial seta; without conical seta; without modified seta; without spinous process; with aesthetasc; one element. Actual segment 15 with seta; two elements; of unequal size; straight; not bifidform; none larger than segment; without spinules; without vestigial seta; without conical seta; without modified seta; with spinous process; on outer margin; surpassing distal margin; with aesthetasc; one element. Actual segment 16 with seta; two elements; of unequal size; plumose; one larger than segment; surpassing to distal margin; not beyond three sequential segments; not bifidform; without spinules; without vestigial seta; without conical seta; without modified seta; with spinous process; on outer margin; surpassing distal margin; unequal size to process on preceding segment; with aesthetasc; one element. Actual segment 17 with seta; two elements; of unequal size; straight; none larger than segment; bifidform; without spinules; without vestigial seta; without conical seta; with modified seta; one element; stout form; surpassing to distal margin; not beyond of the sequential segment; parallel to antennule direction; without spinous process; without aesthetasc. Actual segment 18 with seta; two elements; of equal size; straight; none larger than segment; without spinules; without vestigial seta; without conical seta; with modified seta; one element; stout form; surpassing distal margin; parallel to antennule direction; without spinous process; without aesthetasc. Actual segment 19 with seta; two elements; of unequal size; plumose; none larger than segment; without spinules; without vestigial seta; without conical seta; with modified seta; two elements; stout form; at least one bifid form; surpassing distal margin; parallel to antennule direction; without spinous process; with aesthetasc; one element. Actual segment 20 with seta; four elements; of unequal size; straight; one larger than segment; surpassing to distal margin; beyond three sequential segments; without spinules; without vestigial seta; without conical seta; without modified seta; without spinous process; without aesthetasc. Actual segment 21 with seta; two elements; of equal size; plumose; one larger than segment; surpassing to distal margin; greater 3x than original segment; without spinules; without vestigial seta; without conical seta; without modified seta; without spinous process; without aesthetasc. Actual segment 22 with seta; five elements; of equal size; one larger than segment; plumose; surpassing to distal margin; greater 3x than original segment; without spinules; without vestigial seta; without conical seta; without modified seta; without spinous process; with aesthetasc; one element.

**Left antennules**. Uniramous; Left antennule surpassing to prosome; Left antennule extending beyond caudal rami. Ancestral segment I and II separated. Ancestral segment II and III fused. Ancestral segment III and IV fused. Ancestral segment IV and V separated. Ancestral segment V and VI separated. Ancestral segment VI and VII separated. Ancestral segment VII and VIII separated. Ancestral segment VIII and IX separated. Ancestral segment IX and X separated. Ancestral segment X and XI separated. Ancestral segment XI and XII separated. Ancestral segment XII and XIII separated. Ancestral segment XIII and XIV separated. Ancestral segment XIV and XV separated. Ancestral segment XV and XVI separated. Ancestral segment XVI and XVII separated. Ancestral segment XVII and XVIII separated. Ancestral segment XVIII and XIX separated. Ancestral segment XIX and XX separated. Ancestral segment XX and XXI separated. Ancestral segment XXI and XXII separated. Ancestral segment XXII and XXIII separated. Ancestral segment XXIII and XXIV separated. Ancestral segment XXIV and XXV separated. Ancestral segment XXV and XXVI separated. Ancestral segment XXVI and XXVII separated. Ancestral segment XXVII and XXVIII fused.

Left antennule actual 25-segmented; not-geniculated. Actual segment 1 with seta; one element; none larger than segment; straight; without spinules; without vestigial seta; without conical seta; without modified seta; without spinous process; with aesthetasc; one element. Actual segment 2 with seta; three elements; of equal size; none larger than segment; straight; without spinules; with vestigial seta; one element; without conical seta; without modified seta; without spinous process; with aesthetasc; one element. Actual segment 3 with seta; one element; one larger than segment; straight; surpassing to distal margin; beyond three sequential segments; without spinules; with vestigial seta; one element; without conical seta; without modified seta; without spinous process; with aesthetasc. Actual segment 4 with seta; one element; none larger than segment; straight; without spinules; without vestigial seta; without conical seta; without modified seta; without spinous process; without aesthetasc. Actual segment 5 with seta; one element; one larger than segment; straight; surpassing to distal margin; not beyond three sequential segments; without spinules; with vestigial seta; one element; without conical seta; without modified seta; without spinous process; with aesthetasc; one element. Actual segment 6 with seta; one element; none larger than segment; straight; without spinules; without vestigial seta; without conical seta; without modified seta; without spinous process; without aesthetasc. Actual segment 7 with seta; one element; one larger than segment; straight; surpassing to distal margin; beyond three sequential segments; without spinules; without vestigial seta; without conical seta; without modified seta; without spinous process; with aesthetasc; one element. Actual segment 8 with seta; one element; one larger than segment; straight; surpassing distal margin; without spinules; without vestigial seta; with conical seta; without modified seta; without spinous process; without aesthetasc. Actual segment 9 with seta; two elements; of unequal size; one larger than segment; straight; surpassing to distal margin; beyond three sequential segments; without spinules; without vestigial seta; without conical seta; without modified seta; without spinous process; with aesthetasc; one element. Actual segment 10 with seta; one element; none larger than segment; straight; without spinules; without vestigial seta; without conical seta; without modified seta; without spinous process; without aesthetasc. Actual segment 11 with seta; one element; one larger than segment; straight; surpassing to distal margin; beyond three sequential segments; without spinules; without vestigial seta; without conical seta; without modified seta; without spinous process; without aesthetasc. Actual segment 12 with seta; one element; one larger than segment; straight; surpassing distal margin; without spinules; without vestigial seta; with conical seta; without modified seta; without spinous process; with aesthetasc; one element. Actual segment 13 with seta; one element; none elongated; straight; surpassing distal margin; without spinules; without vestigial seta; without conical seta; without modified seta; without spinous process; without aesthetasc. Actual segment 14 with seta; one element; elongated; straight; surpassing to distal margin; beyond three sequential segments; without spinules; without vestigial seta; without conical seta; without modified seta; without spinous process; with aesthetasc; one element. Actual segment 15 with seta; one element; larger than segment; straight; surpassing to distal margin; not beyond three sequential segments; without spinules; without vestigial seta; without conical seta; without modified seta; without spinous process; without aesthetasc. Actual segment 16 with seta; one element; larger than segment; plumose; surpassing to distal margin; not beyond three sequential segments; without spinules; without vestigial seta; without conical seta; without modified seta; without spinous process; with aesthetasc; one element. Actual segment 17 with seta; one element; not larger than segment; straight; without spinules; without vestigial seta; without conical seta; without modified seta; without spinous process; without aesthetasc. Actual segment 18 with seta; one element; larger than segment; straight; surpassing to distal margin; beyond three sequential segments; without spinules; without vestigial seta; without conical seta; without modified seta; without spinous process; without aesthetasc. Actual segment 19 with seta; one element; not larger than segment; straight; surpassing distal margin; without spinules; without vestigial seta; without conical seta; without modified seta; without spinous process; with aesthetasc; one element. Actual segment 20 with seta; one element; not larger than segment; straight; surpassing distal margin; without spinules; without vestigial seta; without conical seta; without modified seta; without spinous process; without aesthetasc. Actual segment 21 with seta; one element; larger than segment; plumose; surpassing to distal margin; beyond three sequential segments; without spinules; without vestigial seta; without conical seta; without modified seta; without spinous process; without aesthetasc. Actual segment 22 with seta; two elements; of unequal size; one of them elongated; plumose; surpassing to distal margin; without spinules; without vestigial seta; without conical seta; without modified seta; without spinous process; without aesthetasc. Actual segment 23 with seta; two elements; of unequal size; one larger than segment; plumose; surpassing to distal margin; greater 3x than original segment; without spinules; without vestigial seta; without conical seta; without modified seta; without spinous process; without aesthetasc. Actual segment 24 with seta; two elements; of equal size; one larger than segment; plumose; surpassing to distal margin; greater 3x than original segment; without spinules; without vestigial seta; without conical seta; without modified seta; without spinous process; without aesthetasc. Actual segment 25 with seta; five elements; of equal size; elongated; plumose; surpassing to distal margin; 4 times larger than segment; without spinules; without vestigial seta; without conical seta; without modified seta; without spinous process; with aesthetasc; one element.

##### Antenna

Biramous. Antenna coxa separated from the basis; bearing seta; 1; on inner surface; at distal corner; reaching to the endopod 1. Antenna basis (fusion) separated from the endopodal segment; bearing seta; 2; on inner surface; at distal corner. Endopodal ancestral segment I and II separated. Ancestral segment II and III fused. Ancestral segment III and IV fused. Ancestral segment III and IV fully. Antenna endopod actual 2-segmented. Actual segment 1 not bilobate; with seta; two; on inner margin; with spinules; as a row; obliquely; on outer surface; with pore. Actual segment 2 bilobate; with discontinuity on outer cuticle; not developed as a suture; inner lobe bearing 8 setae; distally; outer lobe bearing 7 setae; distally; with spinules; as a patch; on outer surface. Antenna exopod ancestral segment I and II separated. Ancestral segment II and III fused. Ancestral segment III and IV fused. Ancestral segment IV and V separated. Ancestral segment V and VI separated. Ancestral segment VI and VII separated. Ancestral segment VII and VIII separated. Ancestral segment VIII and IX separated. Ancestral segment IX and X fused. Antenna exopod actual 7-segmented. Actual segment 1 single; elongated (width-length, equal or larger ratio 2:1); with seta; one; at inner surface.

Actual segment 2 compound; elongated (larger width-length ratio 2:1); with seta; three; at inner surface. Actual segment 3 single; not elongated (lesser width-length ratio 2:1); with seta; one; at inner surface. Actual segment 4 single; not elongated (lesser width-length ratio 2:1); with seta; one; at inner surface. Actual segment 5 single; not elongated (lesser width-length ratio 2:1); with seta; one; at inner surface. Actual segment 6 single; not elongated (lesser width-length ratio 2:1); with seta; one; at inner surface. Actual segment 7 compound; elongated (larger or equal width-length ratio 2:1); with seta; one; at inner surface; and three; at distal surface.

##### Oral features

**Mandible**. Coxal gnathobase sclerotized; with lobe; prominent; on caudal margin; presence of cutting blade; with tooth-like prominence; two, distinctly; 1 acute; on caudal margin; and 1 triangular; on sub-caudal margin; without acute projection between the prominences; with additional spinules; as a row; on dorsal surface; with seta; 1; dorsally; on apical surface; with spinules; apicalmost. Mandible palps biramous; comprising the basis; with seta; four; differently inserted; first medially; reaching to beyond the endopod 1; second distally; third distally; fourth distally; on inner margin; none with setulose ornamentation. Mandible endopod 2-segmented. Mandible endopod 1 with lobe; bearing seta; four; distally inserted; without spinules. Mandible endopod 2 without lobe; bearing setae; nine elements; distally inserted; with spinules; as a row; double. Mandible exopod 4-segmented. Mandible exopod 1 with seta; one element; distally; on inner margin. Mandible exopod 2 with seta; one element; distally; on inner side. Mandible exopod 3 with seta; one element; distally; on inner side. Mandible exopod 4 with setae; three elements; on terminal region. **Maxillule**. Birramous. Maxillule 3-segmented. Maxillule praecoxa with praecoxal arthrite; bearing spines; fifteen elements; ten marginally; plus, five sub-marginally; with spinules; as a patch; on sub-marginal surface. Maxillule coxa with coxal epipodite; with conspicuous outer lobe; bearing setae; nine elements; with coxal endite; elongated (larger or equal width-length ratio 2:1); bearing setae; four elements. Maxillule basis with basal endite; double; first proximal; elongated (larger width-length ratio 2:1; separated from basis; with setae; four elements; distally inserted; second distal; fused to basis; not elongated (lesser width-length ratio 2:1); with setae; four elements; distally inserted; with setules; as a row; on inner side; basal exite present; with setae; one element; on outer surface. Maxillule endopod 1-segmented. Endopod 1 bilobate; first proximal; with setae; three elements; second distal; with setae; five elements. Maxillule exopod 1-segmented. Exopod 1 with setae; six elements; with setules; as a row; on inner side; spinules absent. **Maxilla**. Uniramous. Maxilla 5-segmented. Maxilla praecoxa fused to coxa; incompletely; distinct externally; with praecoxal endite; double; first elongated endite (larger or equal width length ratio 2:1); proximally inserted; with seta; straight, or plumose; 1 straight; 4 plumose; with spine; single; without spinules; without setule; second elongated endite (larger or equal width length ratio 2:1); distally inserted; with seta; plumose; 3 plumose; without spine; with spinules; as a row; on distal margin; with setule; as a row; on distal margin; absence of outer seta. Maxilla coxa with coxal endite; double; first elongated endite (larger or equal width); proximally inserted; with seta; plumose; 3 plumose; without spine; without spinules; with setules; as a row; on proximal margin; second elongated endite (larger or equal width); distally inserted; with seta; plumose; 3 plumose; without spine; without spinules; with setules; as a row; on proximal margin; absence of outer seta. Maxilla basis with basal endite; single; elongated (larger or equal width-length ratio 2:1); with seta; plumose; 3 plumose; without spinules; absence of outer seta. Maxilla endopod 2-segmented. Endopod 1 with seta; 2 plumose; without spine; without spinules; without setules. Maxilla endopod 2 with seta; 2 plumose; without spine; without spinules; without setules. **Maxilliped**. Uniramous; Maxilliped 8-segmented. Maxilliped praecoxa fused to coxa; incompletely; distinct internally; with praecoxal endite; not elongated (lesser width-length ratio 2:1); distally inserted; with seta; 1 straight; with spinules; as a row; single; on basal surface; without setules. Maxilliped coxa with coxal endite; three coxal endite; first elongated (larger or equal width); proximally inserted; with seta; 2 plumose; with spinules; as a patch; single; on apical surface; without setules; second not elongated (lesser width-length ratio 2:1); medially inserted; with seta; 3 plumose; with spinules; as a row; single; on medial surface; without setules; third elongated (larger or equal width length ratio 2:1); distally inserted; with seta; 3 plumose; none reaching to beyond of the basis; with spinules; as a row; single; on basal surface; without setules; with lobe; prominence; at inner distal angle; ornamented; with spinules; continuously on margin. Maxilliped basis without basal endite; with seta; 3 plumose; with spinules; as a row; single; on medial surface; with setules; as a row; single; on inner margin. Maxilliped endopod segment 6-segmented. Endopod 1 with seta; 2 plumose; on inner surface. Endopod 2 with seta; 3 plumose; on inner surface. Endopod 3 with seta; 2 plumose; on inner surface. Endopod 4 with seta; 2 plumose; on inner surface. Endopod 5 with seta; 2 plumose; on inner surface, or on outer surface; outer seta absent. Endopod 6 with seta; 4 plumose; on inner surface, or on outer surface.

##### Swimming legs features

**First swimming legs.** Symmetrical; biramous. First swimming legs intercoxal plate without seta. First swimming legs praecoxa absent. First swimming legs coxa with seta; one; straight; distally inserted; on inner surface; surpassing to first endopodal segment; with setules; two group; as a patch; on inner margin; and as a row; double; on anterior surface; outerly; without spinules; without spine. First swimming legs basis without seta; with setules; as a patch; single; on outer surface; without spinules; without spine. First swimming legs endopod 2-segmented. Endopod 1 with seta; straight; restricted; to inner surface; one element; without spine; with setules; as a row; single; continuously; on outer surface; without spinules; absence of Schmeil’s organ. Endopod 2 with seta; unrestricted; three on inner surface; one on outer surface; two on distal surface; straight; without spine; with setules; as a row; single; continuously; on outer surface; without spinules; absence of Schmeil’s organ. Endopod 3 absence. First swimming legs exopod 1 with seta; restricted; 1 on inner surface; with spine; 1; stout; smaller than original segment; serrated; on inner side; continuously; with setules; as a row; single; as a row; innerly. First swimming legs exopod 2 with seta; restricted; 1 on inner surface; straight; without spine; with setules; as a row; single; continuously; on inner margin, or on outer margin; without spinules. First swimming legs exopod 3 with setule; as a row; single; continuously; on outer surface; without spinules; with seta; unrestricted; 2 on inner surface; 2 on terminal surface; with spine; 2; unequal size; first no longer 2x than origin segment; stout; serrated; on inner side, or on outer side; equally; second longer 3x than origin segment; slender; serrated; on outer side; with ornamentation on non-serrated side; by setules. **Second swimming legs**. Symmetrical; Second swimming legs biramous. Second swimming legs intercoxal plate without seta. Second swimming legs praecoxa present; located laterally. Second swimming legs coxa with seta; straight; distally inserted; on inner surface; surpassing to basal segment; without setules; without spinules; without spine. Second swimming legs basis without seta; without setules; without spinules; without spine. Second swimming legs endopod 3-segmented. Endopod 1 with seta; straight; restricted; one on inner surface; without spine; with setules; as a row; single; continuously; on outer surface; without spinules; absence of Schmeil’s organ. Endopod 2 with seta; straight; unrestricted; two on inner surface; without spine; with setules; as a row; single; continuously; on outer side; without spinules; presence of Schmeil’s organ; on posterior surface. Endopod 3 with seta; straight; unrestricted; three on inner surface; two on outer surface; two on distal surface; without spine; without setules; with spinules; as a row; double; distally inserted; at anterior surface; absence of Schmeil’s organ. Second swimming legs exopod 1 with seta; restricted; one on inner surface; with spine; 1; stout; not reaching to distal-third of the exopod 2; serrated; on inner side, or on outer side; with setules; as a row; single; continuously; on inner side; without spinules; absence of Schmeil’s organ. Exopod 2 with seta; unrestricted; one on inner surface; with spine; 1; stout; not surpassing the exopod 3; serrated; on inner side, or on outer side; with setules; as a row; single; continuously; on inner surface; without spinules; absence of Schmeil’s organ. Exopod 3 with seta; plurimarginal; three on inner surface; two on terminal surface; with spine; 2; unequal size; first no longer 2x than origin segment; stout; serrated; on inner side, or on outer side; equally; second longer 2x than origin segment; slender; serrated; on outer side; with ornamentation on non-serrated side; of setules; setules on outer surface; as a row; single; continuously; on inner surface; with spinules; as a row; single; distally inserted; at anterior surface; absence of Schmeil’s organ. **Third swimming legs**. Symmetrical; Third swimming legs biramous. Third swimming legs intercoxal plate without seta. Third swimming legs praecoxa present; not laterally located. Third swimming legs coxa with seta; straight; distally inserted; on inner surface; surpassing to first endopodal segment; without setules; without spinules; without spine. Third swimming legs basis without seta; without setules; without spinules; without spine. Third swimming legs endopod 3-segmented. Endopod 1 with seta; restricted; one on inner surface; without spine; without setules; without spinules; absence of Schmeil’s organ. Endopod 2 with seta; restricted; two on inner surface; straight; without spine; without setules; without spinules; absence of Schmeil’s organ. Endopod 3 with seta; straight; plurimarginal; two on inner surface; two on outer surface; three on terminal surface; without spine; without setules; with spinules; as a row; distally inserted; double; at anterior surface; absence of Schmeil’s organ. Third swimming legs exopod 1 with seta; restricted; straight; one on inner surface; with spine; 1; stout; not reaching to the distal-third of the exopod 2; serrated; equally; on inner surface, or on outer surface; with setules; as a row; single; continuously; on inner surface; without spinules; absence of Schmeil’s organ. Exopod 2 with seta; straight; restricted; one on inner surface; with spine; 1; stout; not reaching out to exopod 3; serrated; on inner side, or on outer side; equally; with setules; as a row; single; continuously; on inner side; without spinules; absence of Schmeil’s organ. Exopod 3 without setules; with spinules; as a row; single; distally inserted; at anterior surface; with seta; straight; unrestricted; three on inner surface; two on terminal surface; with spine; 2; unequal size; first no longer 2x than origin segment; stout; serrated; on inner side, or on outer side; equally; second longer 2x than origin segment; slender; serrated; on outer side; with ornamentation on non-serrated side; of setules; absence of Schmeil’s organ. **Fourth swimming legs**. Symmetrical; biramous. Intercoxal plate without sensilla. Praecoxa present. Coxa with seta; distally inserted; on inner margin; reaching out to endopod 1; without spinules; setules absent. Basis with seta; one; medially inserted; on posterior surface; smaller than the original segment; without setules; without spinules; without spine. Fourth swimming legs endopod 3-segmented. Endopod 1 with seta; one; restricted; on inner surface; without spine; without setules; without spinules; absence of Schmeil’s organ. Endopod 2 with seta; restricted; two on inner side; without spine; with setules; as a row; single; continuously; on outer surface; without spinules; absence of Schmeil’s organ. Endopod 3 with seta; unrestricted; two on inner surface; two on outer surface; three on distal surface; without spine; without setules; with spinules; as a row; double; distally inserted; at anterior surface; absence of Schmeil’s organ. Fourth swimming legs exopod 1 with seta; restricted; one on inner surface; with spine; 1; stout; not reaching out to distal-third of the exopod 2; serrated; on inner side, or on outer side; equally; with setules; as a row; single; continuously; on inner surface; without spinules; absence of Schmeil’s organ. Exopod 2 with seta; restricted; one on inner surface; with spine; 1; stout; not reaching the end of exopod 3; serrated; on inner side, or on outer side; equally; with setules; as a row; single; continuously; on inner surface; without spinules; absence of Schmeil’s organ. Exopod 3 without setules; with spinules; as a row; single; distally inserted; at anterior surface; with seta; unrestricted; three on inner surface; two on distal surface; with spine; 2; unequal size; first no longer 2x than origin segment; stout; serrated; on inner side, or on outer side; equally; second longer 2x than origin segment; slender; serrated; on outer side; without ornamentation on non-serrated side; absence of Schmeil’s organ.

##### Fifth swimming legs features

Asymmetrical. Fifth swimming leg intercoxal plate with length equal or greater than width on 1.5x; with irregular proximal margin; discontinuous to; the anterior margin of the left coxa, or the anterior margin of the right coxa; posterior sensilla on the right lateral absent. **Fifth left swimming leg**. Fifth left swimming leg biramous; leg surpassing first right exopod segment. Fifth left swimming leg praecoxa present; rudimentary; separated from the coxae; without ornamentation. Fifth left swimming leg coxa concave inner side; without teeth-like structures; with process; conical; on posterior surface; outer side; distally inserted; not projecting over basis; with sensilla; stout; triangular; at apex; longer 2x than insertion basis; without swelling; without seta; without spinules. Fifth left swimming leg basis sub-cylindrical; unequal size between inner and outer side; shorter outer than inner side; with convex inner side; rounded internal proximal expansion absent; without outgrowth; without groove; absence of protuberance; with seta; outerly inserted; no longer 2x than origin segment; absence of minutely granular. Fifth left swimming leg endopod segments 1 and 2 fused; segments 2 and 3 fused; 1-segmented; stout; separated from the basis; ornamented; on inner side; with spinules; more than four elements; as a row; terminally; row of setules absent; without seta. Fifth left swimming leg exopod segments 1 and 2 separated; segments 2 and 3 fused; 2-segmented; stout; separated from the basis. Fifth left swimming leg exopod 1 sub-triangular; longer than broad; unequal size between inner and outer side; shorter inner than outer side; concave inner side; convex outer side; without swelling; without marginal extension; with process; single; proximal; posteriorly; with lobe; single; circular; distally inserted; on inner side; covered; by setules; without outer spine; absence seta. Fifth left swimming leg exopod 2 digitiform; longer than broad; equal size between inner and outer side; disform inner side; with rectilinear outer side; setulose pad present; prominently rounded; medially; on inner side; inflated medial region absent; distal process present; digitiform; denticulate; not bicuspidate; without transverse row of denticles; none oblique row of 5 denticles; at anterior surface; not innerly directed; with seta; spiniform; not ornamented by spinules; not surpassing the distal-point of the segment; without outer spine; terminal claw absent.

##### Fifth right swimming leg

Biramous. Fifth right swimming leg praecoxa present; separated from the coxae; without ornamentation. Fifth right swimming leg coxa convex inner side; without teeth-like structures; with process; rounded; distally inserted; on posterior surface; closest to the outer rim; projecting over basis; beyond the first third; until the medial surface; with triangular protuberance innerly; with sensilla; slender; at apex; no longer 2x than basal insertion; without marginal extension; without seta; without spinules. Fifth right swimming leg basis cylindrical; unequal size between inner and outer side; shorter outer than inner side; convex inner side; tumescence present; inflated; not bilobed; unrestricted on inner surface; without protuberance; absence of distinct minutely granular; additional inner process absent; without posterior groove; with seta; outerly inserted; on anterior surface; no longer 2x than origin segment; posterior protrusion absent; distal process absent. Fifth right swimming leg with endopodite present; separated from the basis; on anterior surface; ancestral segments 1 and 2 fused; ancestral segments 2 and 3 fused; 1-segmented; stout; ornamented; with setules; as a row; on inner side; terminally; without seta. Fifth right swimming leg exopod segments 1 and 2 separated; segments 2 and 3 fused; 2-segmented; stout; separated from the basis. Fifth right swimming leg exopod 1 sub-cylindrical; longer than broad; nearly 1.25 times; unequal size between both sides; shorter inner than outer side; rectilinear inner side; rectilinear outer side; with marginal extension; sub-triangular; distally inserted; at outer rim; spinules absent; with process; rounded; sclerotized; without ornamentation; distally inserted; at posterior surface; projecting over next segment; without outer spine; without seta; internal prominence absent; lamella on posterior surface absent. Fifth right swimming leg exopod 2 elliptical; longer than broad; nearly 2 times; equal size between both sides; disform inner side; convex outer side; without posterior proximal swelling; inner-posterior process absent; without marginal expansion; curved ridge on distal posterior surface present; chitinous knobs present; with 3–6 posteriorly; with outer spine; inserted medially; arched; internally directed; ornamented innerly; by spinules; as a row; not ornamented outerly; sharp tip; without apparent curve; lesser than the length of the exopod 2; until to 2 times its size; 2x; sensilla absent; terminal claw present; equal or longer 1.5 times than insertion segment; sclerotized; arched; inward; with conspicuous curve; medially; ornamented innerly; by spinules; as a row; throughout extension; not ornamented outerly; sharp tip; not curved tip; without medial constriction; hyaline process present; on proximal third; innerly.

##### FEMALE

Body longer and wider than male; Female body 1171 micrometers excluding caudal setae. Widest at first metasome segment. Distal margin of the prosomal segments without one line of setules at posterior margin. Prosome segments with spinules at least at one prosomal segment. Fourth metasome segment absence of dorsal protuberance. Fourth and fifth metasome segments fused; totally. Limit between fourth and fifth metasome segments ornamented; with spinules; as a row; double; complete; different size; entirely over limit (lateral, dorsal). **Fifth metasome segment**. Fifth metasome segment with sensilla; dorsally; 2 elements; with epimeral plates. Epimeral plates asymmetrical. Right epimeral plates prominent, as projections; thinner than the left; one posterior-laterally directed; not reaching half length of the genital segment; with sensilla at the apex; dorsal-posterior sensilla absent; without ornamentation. Left epimeral plate without expansion.

##### Urosome

3-segmented. **Genital double-somite**. Asymmetrical in dorsal view; longer than broad; longer than other urosomites combined; dorsal suture at mid-length present; not covered by spinules; with expansion; rounded; like a bulging dilation; unequal size; greater left than right; anteriorly; with sensillae; on both sides; one; stout; with robust apex; at left lateral; not on lobular base; medially; one; stout; at right lateral; not on lobular base; medially; with robust apex; of equal size between then; lateral protuberance absent; without right posterior rim expanded; without slender sensilla on each posterior rim; without posterior-dorsal process. Genital double-somite opercular pad present; broader than longer; symmetrical; development laterally; expanded posteriorly; covering partially; double gonoporal slit; located ventrally; with arthrodial membrane; inserted anteriorly; post-genital process absent; disto-ventral tumescence absent; ventral vertical folds present; dorsal sensilla absent. Second urosome segment without ventral fusion to anal segment; right distal process absent. Caudal rami patch of setules on outer surface absent; patch of spinules on outer surface absent.

##### Oral appendices feature

Rostrum basal process absent. **Antennules**. Symmetrical. Right antennule surpassing to genital double-segment; extending beyond caudal rami. Right antennule exceeding the caudal setae. Right antennule ornamentation pattern equals to male left antennule; fully.

##### Fifth swimming legs

Symmetrical; Fifth swimming legs biramous. Fifth swimming legs intercoxal plate longer than wide; separated from the legs. Fifth swimming legs praecoxa with sclerite praecoxal; separated from the coxae; without ornamentation. Fifth swimming legs coxa with process; conical; at the outer rim; distally; sensilla present; stout; at apex; projecting over basal segment; no longer 2x than basal insertion; marginal extension absent; without swelling; without seta; without spinules. Fifth swimming legs basis sub-triangular; unequal size between inner and outer sides; shorter outer than inner side; with convex inner side; without proximal inner outgrowth; without groove; with distal extension; on posterior surface; with seta; outerly inserted; on anterior surface; longer 2x than origin segment; not reaching to exopod 1 distally. Fifth swimming legs endopod segments 1 and 2 fused; segments 2 and 3 fused; 1-segmented; stout; separated from the basis; present discontinuity cuticle; on inner side; with spinules; as a row; single; non-oblique; sub-terminally; at anterior surface; with seta; double; one medially; on posterior surface; rectilinear; one distally; on posterior surface; rectilinear; of unequal size; distal seta longer than medial seta. Fifth swimming legs exopod segments 1 and 2 separated; segments 2 and 3 separated; 3-segmented; separated from the basis. Fifth swimming legs exopod 1 sub-cylindrical; longer than wide; longer or equal than 2 times; with unequal size between inner and outer side; shorter inner than outer side; with rectilinear inner side; with convex outer side; without swelling; without marginal extension; without posterior process; without spine; without seta. Fifth swimming legs exopod 2 sub-cylindrical; longer than broad; longer or equal than 2 times; without swelling; without marginal extension; without process; without lobe; with spine; inserted laterally; rectilinear; without ornamentation; sharp tip; equal size or larger than next segment; without seta. Fifth swimming legs exopod 3 cylindrical; longer than wide; without swelling; without process; without lobe; without spine; with seta; double; inserted terminally; unequal size between them; outer seta smaller than inner; nearly 3 times; outer seta not ornamented by setules; without ornamentation; presence of terminal claw; sclerotized; arched; internally directed; concave inner side; with ornamentation; of denticles; as a row; on surface partially; at medial region; concave outer side; with ornamentation; of denticles; as a row; on surface partially; at medial region; blunt tip; 6 times longer than origin segment.

##### Distribution records

###### BRAZIL

**Pará**: Urubu stream, between Tucuruí, and Baião (Cipólli & Carvalho, 1973). **Bahia**: pond in Itaparica waterfall, at baiana side of the São Francisco River, near to Jatobá (Wright, 1936); marginal lagoon in São Francisco River, around 7 Km to Iboticama (this study). **Distrito Federal**: Paranoá Lake, in Brasília (Matsumura-Tundisi, 1986). **São Paulo**: Ilha Solteira Reservoir, and Paraná River (Sendacz, 1998). **Paraná**: Itaipu Reservoir (Matsumura-Tundisi, 1986).

##### Habitat

Habitat in freshwaters: streams, ponds, rivers, lakes associated to rivers, lakes, and reservoirs.

##### Remarks

The species was presented to science from organisms collected in a lake environment in the Northeast Brazil. Over time other records had expanded the distribution of these organisms to the North (Cipólli & Carvalho, 1973), Center, Southeast and South (Matsumura-Tundisi, 1986) of Brazil. The species was immediately included in *nordestinus* after its foundation, undergoing the current recombination only in the enlargement of *Notodiaptomus* (Kiefer, 1956).

The main feature highlighted in Wright (1936) for the inclusion of the taxon in *nordestinus* was male fifth right leg exopod 1 longer than broad, forward confirmed by Kiefer (1956) for the grouping in *Notodiaptomus*. Originally the organisms of the species were differentiated within *nordestinus* by the male fifth right swimming leg exopod 2 with outer spine medially. Santos-Silva *et al*. (2015) when reviewing the grouping found that the male of *N. anisitsi* also has the same condition and that the attribute could not be differential for the Wright species within the grouping.

In the present effort, we were able to corroborate the findings of Wright (1936) and Kiefer (1956) for male fifth right swimming leg exopod 1 longer than broad as convergent for 27 species in *Notodiaptomus* currently recognized. The verification for the medial positioning of the outer spine on male fifth leg exopod 2 was corroborated and verified for other species of the genus besides *N. anisitsi*, such as *N. paraensis*, and *N. kieferi.* Additionally, when examining topotypes, we found other conditions that represent variation of the species in relation to the type-species and differentiate it within *Notodiaptomus*: (1) male limit between fourth and fifth metasome segments with single spinules row laterally; (2) male right antennule actual segments 22 and 25 with five setae; (3) male fifth right swimming leg basis with inflated inner tumescence, not bilobed; (4) female limit between fourth and fifth metasome segments with double spinules row of different size; (5) female genital double-somite with dorsal suture at mid-length; female genital double-somite with ventral vertical folds; (5) female fifth swimming legs basis with outer seta longer 2x than origin segment, reaching to middle-point exopod 1; and (6) female genital double-somite with rounded expansion like a bulging dilation.

Predominantly, these verified morphological characteristics corroborate the observations described in the broad review for *N. jatobensis* encompassing the *nordestinus* complex. In contribution of Santos-Silva *et al*. (2015) the attribute 6 was described as a double genital segment with a more “strongly” expanded anterior region. Under the rigor of taxonomic objectivity, here we reinterpret the character as asserted in description 6, in this effort configuring as part of the diagnosis of the species.

#### Notodiaptomus kieferi Brandorff, 1973

##### Synonymy

*Notodiaptomus kieferi* Brandorff, 1972: 4, 30, 50, figs. 40–48; 1973b: 205, 206, pl.1, figs. 1–6, pl. 2, figs. 1–5; 1976: 616, fig. 2; Andrade & Brandorff, 1975: 97; Löffler, 1981: 15; Dussart & Defaye, 1983: 138; Dussart, 1984a: 35, 38, 39, 49, fig. 7; Dussart & Robertson, 1984: 391; Hardy *et al*., 1984: 530; Robertson & Hardy, 1984: tab. 3; Defaye & Dussart, 1988: 113; Magalhães *et al*., 1988: 271; Cicchino, 1994: 145, fig. 8; Rocha *et al*., 1995: 156; Santos-Silva *et al*., 1989: 726, 728, figs. 116–135; Santos-Silva, 2008: 32, fig. 7; Perbiche-Neves *et al*., 2020: 685, key to the Neotropical diaptomid, fig. 21.10 B. *Notodiaptomus (Wrightius) kieferi*; Dussart, 1985a: 210. *Notodiaptomus echinatus* Defaye & Dussart, 1988: 113. *“Diaptomus echinatus”*; Defaye & Dussart, 1988: 113.

##### Type locality

Catalão Lake, Manaus, Amazonas, Brazil.

##### Type material

Originally the holotype or type series was specified for the species. Hardy *et al*. (1984) formally designated lectotype on slide “in poor condition”, (INPA CRUST-033), and “paralectotype” (which represented 1 female) on slide “in poor condition”. Both collected on 22.IX.1959, “leg. H. Siole” and deposited at INPA, Amazonas, Brazil.

##### Material examined

Topotype: 3 males, and 4 females, entire in alcohol, from Catalão Lake, Iranduba City (near Manaus), Amazonas, 22.IX.59, specified by L. Geraldes-Primeiro, and belonging to the collection of the Plankton Laboratory (n° 211), INPA. 1 male (INPA-COP035, slides a-h) and 1 female (INPA-COP036, slides a-h) were selected to be dissection on eight slides each and deposited in the Zoological Collection of the INPA, Brazil.

##### Diagnosis

**(1)** cephalosome with sub-triangular anterior margin; **(2)** cephalosome without dorsal suture; **(3)** male right antennule actual segment 8 with conical seta reaching to middle-point of the sequent segment; **(4)** male right antennule actual segment 13 with modified seta presenting rounded apex; **(5)** antenna endopod actual segment 2 with discontinuity on outer cuticle, developed as a complete suture; **(6)** male fifth left swimming leg with length reaching first right exopod segment medially; **(7)** male fifth left swimming leg exopod 2 with spiniform seta surpassing the distal-point of the segment; **(8)** male fifth right swimming leg exopod 1 with acute internal prominence; **(9)** male fifth right swimming leg exopod 2 with outer spine lesser 1.5x its length than origin segment; **(10)** female right epimeral plate reaching half length of the genital segment; **(11)** female left epimeral plate with dorsal semicircular expansion posteriorly; **(12)** female genital double-somite with lateral conical swelling, greater left than right; **(13)** female caudal rami with spinules patch on outer surface; **(14)** female fifth swimming legs basis with outer seta not 2x longer than original segment; **(15)** female fifth swimming legs exopod 3 with rectilinear terminal claw.

##### Redescription

###### MALE

Body 1157 micrometers excluding caudal setae. Male body smaller and slenderer than female. Nerve axons myelinated. Prosome 6-segmented; widest at cephalosome; posteriorly; without one line of setules at posterior margin; without spinules at segments. Cephalosome anterior margin sub-triangular; without dorsal suture; separate from first metasome segment. First metasome segment without sensilla. Second metasome segment without sensilla. Third metasome segment without sensillae; non-ornamented posterior margin. Fourth metasome segment without sensillae; separated from the fifth metasome. Limit between fourth and fifth metasome segments without ornamentation. Fifth metasome segment without sensilla; Fifth metasome segment without ornamentation; Fifth metasome segment without dorsal conical process; with epimeral plates. Epimeral plates symmetrical. Right epimeral plates prominent, as projections; one projection; posterior-dorsally directed; reaching half or surpassing the length of genital segment; with sensilla; at the apex of projection; without ornamentation.

##### Urosome

5-segmented; Urosome 5 - free segments. Genital somite asymmetrical in dorsal view; with single aperture; located on left side; ventrolaterally on posterior rim; with sensillae; on both sides; one; at left lateral; posteriorly; one; at right rim; posteriorly; of equal size between then. Third urosome segment without spinules; without external seta. Fourth urosome segment without spinules; without sub-conical blunt dorsal-lateral process. Anal segment presence of dorsal sensillae; one on each side; medially inserted; presence of operculum; convex; covering the anal aperture fully. Caudal rami symmetrical; separated from anal segment; longer than wide; with setules; continuous on; inner side; each ramus bearing 6 caudal setae; 5 marginals; plumose; and 1 internal dorsally; straight; not reticulated main axis; outermost seta with outer spiniform process absent.

##### Oral appendices feature

Rostrum symmetrical; separated from dorsal cephalic shield; by complete suture; sensillae present; one pair; anteriorly inserted on surface tegument; with rostral filament; double; paired; extended; into point; with basal process; in ventral view, rounded on left side; without a smaller basal expansion on the right side.

##### Antennules

Asymmetrical. **Right antennules**. Uniramous; right antennule surpassing to genital segment; right antennule extending beyond caudal rami.

Right antennule ancestral segment I and II separated. Ancestral segment II and III fused. Ancestral segment III and IV fused. Ancestral segment IV and V separated. Ancestral segment V and VI separated. Ancestral segment VI and VII separated. Ancestral segment VII and VIII separated. Ancestral segment VIII and IX separated. Ancestral segment IX and X separated. Ancestral segment X and XI separated. Ancestral segment XI and XII separated. Ancestral segment XII and XIII separated. Ancestral segment XIII and XIV separated. Ancestral segment XIV and XV separated. Ancestral segment XV and XVI separated. Ancestral segment XVI and XVII separated. Ancestral segment XVII and XVIII separated. Ancestral segment XVIII and XIX separated. Ancestral segment XIX and XX separated. Ancestral segment XX and XXI separated. Ancestral segment XXI and XXII fused. Ancestral segment XXII and XXIII fused. Ancestral segment XXIII and XXIV separated. Ancestral segment XXIV and XXV fused. Ancestral segment XXV and XXVI separated. Ancestral segment XXVI and XXVII separated. Ancestral segment XXVII and XXVIII fused.

Right antennule actual 22-segmented; geniculated; between the segment 18 and segment 19; with swollen and modified region; formed by 5 segments; between 13 and 17 segments. Actual segment 1 with seta; one element; straight; none larger than segment; without spinules; without vestigial seta; without conical seta; without modified seta; without spinous process; with aesthetasc; one element. Actual segment 2 with seta; three elements; of unequal size; straight; none larger than segment; without spinules; with vestigial seta; one element; without conical seta; without modified seta; without spinous process; with aesthetasc; one element. Actual segment 3 with seta; one element; one larger than segment; surpassing to distal margin; beyond three sequential segments; straight; blunt apex; without spinules; with vestigial seta; one element; without conical seta; without modified seta; without spinous process; with aesthetasc. Actual segment 4 with seta; one element; one larger than segment; surpassing to distal margin; straight; not beyond three sequential segments; without spinules; without vestigial seta; without conical seta; without modified seta; without spinous process; without aesthetasc. Actual segment 5 with seta; one element; straight; one larger than segment; surpassing to distal margin; not beyond three sequential segments; without spinules; with vestigial seta; one element; without conical seta; without modified seta; without spinous process; with aesthetasc; one element. Actual segment 6 with seta; one element; none larger than segment; straight; without spinules; without vestigial seta; without conical seta; without modified seta; without spinous process; without aesthetasc. Actual segment 7 with seta; one element; straight; one larger than segment; surpassing to distal margin; beyond three sequential segments; blunt apex; without spinules; without vestigial seta; without conical seta; without modified seta; without spinous process; with aesthetasc; one element. Actual segment 8 with seta; one element; straight; none larger than segment; without spinules; without vestigial seta; with conical seta; one element; reaching to middle-point of the sequent segment; without modified seta; without spinous process; without aesthetasc. Actual segment 9 with seta; two elements; of unequal size; straight; one larger than segment; surpassing to distal margin; beyond three sequential segments; blunt apex; without spinules; without vestigial seta; without conical seta; without modified seta; without spinous process; with aesthetasc; one element. Actual segment 10 with seta; one element; straight; none larger than segment; without spinules; without vestigial seta; without conical seta; with modified seta; presenting blunt apex; slender form; surpassing to distal margin; beyond of the sequential segment; parallel to antennule direction; without spinous process; without aesthetasc. Actual segment 11 with seta; one element; straight; one larger than segment; surpassing to distal margin; not beyond three sequential segments; without spinules; without vestigial seta; without conical seta; with modified seta; slender form; presenting blunt apex; surpassing to distal margin; beyond of the sequential segment; parallel to antennule direction; shorter length than homologous of actual segment 13; without spinous process; without aesthetasc. Actual segment 12 with seta; one element; straight; one larger than segment; surpassing to distal margin; not beyond three sequential segments; without spinules; without vestigial seta; with conical seta; one element; smaller than to segment 8; without modified seta; without spinous process; with aesthetasc; one element; absent internal perpendicular fission. Actual segment 13 with seta; one element; straight; one larger than segment; surpassing to distal margin; not beyond three sequential segments; without spinules; without vestigial seta; without conical seta; with modified seta; stout form; surpassing to distal margin; to the distal-point of the sequence segment; perpendicular to antennule direction; presenting rounded apex; without spinous process; with aesthetasc; one element. Actual segment 14 with seta; two elements; of unequal size; straight; one larger than segment; surpassing to distal margin; beyond three sequential segments; blunt apex; without spinules; without vestigial seta; without conical seta; without modified seta; without spinous process; with aesthetasc; one element. Actual segment 15 with seta; two elements; of unequal size; straight; not bifidform; none larger than segment; without spinules; without vestigial seta; without conical seta; without modified seta; with spinous process; on outer margin; surpassing distal margin; with aesthetasc; one element. Actual segment 16 with seta; two elements; of unequal size; plumose; one larger than segment; surpassing to distal margin; not beyond three sequential segments; not bifidform; without spinules; without vestigial seta; without conical seta; without modified seta; with spinous process; on outer margin; surpassing distal margin; unequal size to process on preceding segment; with aesthetasc; one element. Actual segment 17 with seta; two elements; of unequal size; straight; none larger than segment; bifidform; without spinules; without vestigial seta; without conical seta; with modified seta; one element; stout form; surpassing to distal margin; not beyond of the sequential segment; parallel to antennule direction; without spinous process; without aesthetasc. Actual segment 18 with seta; two elements; of equal size; straight; none larger than segment; without spinules; without vestigial seta; without conical seta; with modified seta; one element; stout form; surpassing distal margin; parallel to antennule direction; without spinous process; without aesthetasc. Actual segment 19 with seta; two elements; of unequal size; plumose; none larger than segment; without spinules; without vestigial seta; without conical seta; with modified seta; two elements; stout form; at least one bifid form; surpassing distal margin; parallel to antennule direction; without spinous process; with aesthetasc; one element. Actual segment 20 with seta; four elements; of unequal size; straight; one larger than segment; surpassing to distal margin; beyond three sequential segments; without spinules; without vestigial seta; without conical seta; without modified seta; with spinous process; distally; not reaching beyond of distal-point segment 21; without aesthetasc. Actual segment 21 with seta; two elements; of equal size; plumose; one larger than segment; surpassing to distal margin; greater 3x than original segment; without spinules; without vestigial seta; without conical seta; without modified seta; without spinous process; without aesthetasc. Actual segment 22 with seta; four elements; of equal size; one larger than segment; plumose; surpassing to distal margin; greater 3x than original segment; without spinules; without vestigial seta; without conical seta; without modified seta; without spinous process; with aesthetasc; one element.

##### Left antennules

Uniramous; Left antennule surpassing to prosome; Left antennule extending beyond caudal rami. Ancestral segment I and II separated. Ancestral segment II and III fused. Ancestral segment III and IV fused. Ancestral segment IV and V separated. Ancestral segment V and VI separated. Ancestral segment VI and VII separated. Ancestral segment VII and VIII separated. Ancestral segment VIII and IX separated. Ancestral segment IX and X separated. Ancestral segment X and XI separated. Ancestral segment XI and XII separated. Ancestral segment XII and XIII separated. Ancestral segment XIII and XIV separated. Ancestral segment XIV and XV separated. Ancestral segment XV and XVI separated. Ancestral segment XVI and XVII separated. Ancestral segment XVII and XVIII separated. Ancestral segment XVIII and XIX separated. Ancestral segment XIX and XX separated. Ancestral segment XX and XXI separated. Ancestral segment XXI and XXII separated. Ancestral segment XXII and XXIII separated. Ancestral segment XXIII and XXIV separated. Ancestral segment XXIV and XXV separated. Ancestral segment XXV and XXVI separated. Ancestral segment XXVI and XXVII separated. Ancestral segment XXVII and XXVIII fused.

Left antennule actual 25-segmented; not-geniculated. Actual segment 1 with seta; one element; none larger than segment; straight; without spinules; without vestigial seta; without conical seta; without modified seta; without spinous process; with aesthetasc; one element. Actual segment 2 with seta; three elements; of equal size; none larger than segment; straight; without spinules; with vestigial seta; one element; without conical seta; without modified seta; without spinous process; with aesthetasc; one element. Actual segment 3 with seta; one element; one larger than segment; straight; surpassing to distal margin; beyond three sequential segments; without spinules; with vestigial seta; one element; without conical seta; without modified seta; without spinous process; with aesthetasc. Actual segment 4 with seta; one element; none larger than segment; straight; without spinules; without vestigial seta; without conical seta; without modified seta; without spinous process; without aesthetasc. Actual segment 5 with seta; one element; one larger than segment; straight; surpassing to distal margin; not beyond three sequential segments; without spinules; with vestigial seta; one element; without conical seta; without modified seta; without spinous process; with aesthetasc; one element. Actual segment 6 with seta; one element; none larger than segment; straight; without spinules; without vestigial seta; without conical seta; without modified seta; without spinous process; without aesthetasc. Actual segment 7 with seta; one element; one larger than segment; straight; surpassing to distal margin; beyond three sequential segments; without spinules; without vestigial seta; without conical seta; without modified seta; without spinous process; with aesthetasc; one element. Actual segment 8 with seta; one element; one larger than segment; straight; surpassing distal margin; without spinules; without vestigial seta; with conical seta; without modified seta; without spinous process; without aesthetasc. Actual segment 9 with seta; two elements; of unequal size; one larger than segment; straight; surpassing to distal margin; beyond three sequential segments; without spinules; without vestigial seta; without conical seta; without modified seta; without spinous process; with aesthetasc; one element. Actual segment 10 with seta; one element; none larger than segment; straight; without spinules; without vestigial seta; without conical seta; without modified seta; without spinous process; without aesthetasc. Actual segment 11 with seta; one element; one larger than segment; straight; surpassing to distal margin; beyond three sequential segments; without spinules; without vestigial seta; without conical seta; without modified seta; without spinous process; without aesthetasc. Actual segment 12 with seta; one element; one larger than segment; straight; surpassing distal margin; without spinules; without vestigial seta; with conical seta; without modified seta; without spinous process; with aesthetasc; one element. Actual segment 13 with seta; one element; none elongated; straight; surpassing distal margin; without spinules; without vestigial seta; without conical seta; without modified seta; without spinous process; without aesthetasc. Actual segment 14 with seta; one element; elongated; straight; surpassing to distal margin; beyond three sequential segments; without spinules; without vestigial seta; without conical seta; without modified seta; without spinous process; with aesthetasc; one element. Actual segment 15 with seta; one element; larger than segment; straight; surpassing to distal margin; not beyond three sequential segments; without spinules; without vestigial seta; without conical seta; without modified seta; without spinous process; without aesthetasc. Actual segment 16 with seta; one element; larger than segment; plumose; surpassing to distal margin; not beyond three sequential segments; without spinules; without vestigial seta; without conical seta; without modified seta; without spinous process; with aesthetasc; one element. Actual segment 17 with seta; one element; not larger than segment; straight; without spinules; without vestigial seta; without conical seta; without modified seta; without spinous process; without aesthetasc. Actual segment 18 with seta; one element; larger than segment; straight; surpassing to distal margin; beyond three sequential segments; without spinules; without vestigial seta; without conical seta; without modified seta; without spinous process; without aesthetasc. Actual segment 19 with seta; one element; not larger than segment; straight; surpassing distal margin; without spinules; without vestigial seta; without conical seta; without modified seta; without spinous process; with aesthetasc; one element. Actual segment 20 with seta; one element; not larger than segment; straight; surpassing distal margin; without spinules; without vestigial seta; without conical seta; without modified seta; without spinous process; without aesthetasc. Actual segment 21 with seta; one element; larger than segment; plumose; surpassing to distal margin; beyond three sequential segments; without spinules; without vestigial seta; without conical seta; without modified seta; without spinous process; without aesthetasc. Actual segment 22 with seta; two elements; of unequal size; one of them elongated; plumose; surpassing to distal margin; without spinules; without vestigial seta; without conical seta; without modified seta; without spinous process; without aesthetasc. Actual segment 23 with seta; two elements; of unequal size; one larger than segment; plumose; surpassing to distal margin; greater 3x than original segment; without spinules; without vestigial seta; without conical seta; without modified seta; without spinous process; without aesthetasc. Actual segment 24 with seta; two elements; of equal size; one larger than segment; plumose; surpassing to distal margin; greater 3x than original segment; without spinules; without vestigial seta; without conical seta; without modified seta; without spinous process; without aesthetasc. Actual segment 25 with seta; four elements; of equal size; elongated; plumose; surpassing to distal margin; 4 times larger than segment; without spinules; without vestigial seta; without conical seta; without modified seta; without spinous process; with aesthetasc; one element.

##### Antenna

Biramous. Antenna coxa separated from the basis; bearing seta; 1; on inner surface; at distal corner; reaching to the endopod 1. Antenna basis (fusion) separated from the endopodal segment; bearing seta; 2; on inner surface; at distal corner. Endopodal ancestral segment I and II separated. Ancestral segment II and III fused. Ancestral segment III and IV fused. Ancestral segment III and IV fully. Antenna endopod actual 2-segmented. Actual segment 1 not bilobate; with seta; two; on inner margin; with spinules; as a row; obliquely; on outer surface; with pore. Actual segment 2 bilobate; with discontinuity on outer cuticle; developed as a suture; complete; inner lobe bearing 8 setae; distally; outer lobe bearing 7 setae; distally; with spinules; as a patch; on outer surface. Antenna exopod ancestral segment I and II separated. Ancestral segment II and III fused. Ancestral segment III and IV fused. Ancestral segment IV and V separated. Ancestral segment V and VI separated. Ancestral segment VI and VII separated. Ancestral segment VII and VIII separated. Ancestral segment VIII and IX separated. Ancestral segment IX and X fused. Antenna exopod actual 7-segmented. Actual segment 1 single; elongated (width-length, equal or larger ratio 2:1); with seta; one; at inner surface. Actual segment 2 compound; elongated (larger width-length ratio 2:1); with seta; three; at inner surface. Actual segment 3 single; not elongated (lesser width-length ratio 2:1); with seta; one; at inner surface. Actual segment 4 single; not elongated (lesser width-length ratio 2:1); with seta; one; at inner surface. Actual segment 5 single; not elongated (lesser width-length ratio 2:1); with seta; one; at inner surface. Actual segment 6 single; not elongated (lesser width-length ratio 2:1); with seta; one; at inner surface. Actual segment 7 compound; elongated (larger or equal width-length ratio 2:1); with seta; one; at inner surface; and three; at distal surface.

##### Oral features

**Mandible**. Coxal gnathobase sclerotized; with lobe; prominent; on caudal margin; presence of cutting blade; with tooth-like prominence; two, distinctly; 1 acute; on caudal margin; and 1 triangular; on sub-caudal margin; without acute projection between the prominences; with additional spinules; as a row; on dorsal surface; with seta; 1; dorsally; on apical surface; with spinules; apicalmost. Mandible palps biramous; comprising the basis; with seta; four; differently inserted; first medially; reaching to beyond the endopod 1; second distally; third distally; fourth distally; on inner margin; none with setulose ornamentation. Mandible endopod 2-segmented. Mandible endopod 1 with lobe; bearing seta; four; distally inserted; without spinules. Mandible endopod 2 without lobe; bearing setae; nine elements; distally inserted; with spinules; as a row; double. Mandible exopod 4-segmented. Mandible exopod 1 with seta; one element; distally; on inner margin. Mandible exopod 2 with seta; one element; distally; on inner side. Mandible exopod 3 with seta; one element; distally; on inner side. Mandible exopod 4 with setae; three elements; on terminal region. **Maxillule**. Birramous. Maxillule 3-segmented. Maxillule praecoxa with praecoxal arthrite; bearing spines; fifteen elements; ten marginally; plus, five sub-marginally; with spinules; as a patch; on sub-marginal surface. Maxillule coxa with coxal epipodite; with conspicuous outer lobe; bearing setae; nine elements; with coxal endite; elongated (larger or equal width-length ratio 2:1); bearing setae; four elements. Maxillule basis with basal endite; double; first proximal; elongated (larger width-length ratio 2:1; separated from basis; with setae; four elements; distally inserted; second distal; fused to basis; not elongated (lesser width-length ratio 2:1); with setae; four elements; distally inserted; with setules; as a row; on inner side; basal exite present; with setae; one element; on outer surface. Maxillule endopod 1-segmented. Endopod 1 bilobate; first proximal; with setae; three elements; second distal; with setae; five elements. Maxillule exopod 1-segmented. Exopod 1 with setae; six elements; with setules; as a row; on inner side; spinules absent. **Maxilla**. Uniramous. Maxilla 5-segmented. Maxilla praecoxa fused to coxa; incompletely; distinct externally; with praecoxal endite; double; first elongated endite (larger or equal width length ratio 2:1); proximally inserted; with seta; straight, or plumose; 1 straight; 4 plumose; with spine; single; without spinules; without setule; second elongated endite (larger or equal width length ratio 2:1); distally inserted; with seta; plumose; 3 plumose; without spine; with spinules; as a row; on distal margin; with setule; as a row; on distal margin; absence of outer seta. Maxilla coxa with coxal endite; double; first elongated endite (larger or equal width); proximally inserted; with seta; plumose; 3 plumose; without spine; without spinules; with setules; as a row; on proximal margin; second elongated endite (larger or equal width); distally inserted; with seta; plumose; 3 plumose; without spine; without spinules; with setules; as a row; on proximal margin; absence of outer seta. Maxilla basis with basal endite; single; elongated (larger or equal width-length ratio 2:1); with seta; plumose; 3 plumose; without spinules; absence of outer seta. Maxilla endopod 2-segmented. Endopod 1 with seta; 2 plumose; without spine; without spinules; without setules. Maxilla endopod 2 with seta; 2 plumose; without spine; without spinules; without setules. **Maxilliped**. Uniramous; Maxilliped 8-segmented. Maxilliped praecoxa fused to coxa; incompletely; distinct internally; with praecoxal endite; not elongated (lesser width-length ratio 2:1); distally inserted; with seta; 1 straight; with spinules; as a row; single; on basal surface; without setules. Maxilliped coxa with coxal endite; three coxal endite; first elongated (larger or equal width); proximally inserted; with seta; 2 plumose; with spinules; as a patch; single; on apical surface; without setules; second not elongated (lesser width-length ratio 2:1); medially inserted; with seta; 3 plumose; with spinules; as a row; single; on medial surface; without setules; third elongated (larger or equal width length ratio 2:1); distally inserted; with seta; 3 plumose; none reaching to beyond of the basis; with spinules; as a row; single; on basal surface; without setules; with lobe; prominence; at inner distal angle; ornamented; with spinules; continuously on margin. Maxilliped basis without basal endite; with seta; 3 plumose; with spinules; as a row; single; on medial surface; with setules; as a row; single; on inner margin. Maxilliped endopod segment 6-segmented. Endopod 1 with seta; 2 plumose; on inner surface. Endopod 2 with seta; 3 plumose; on inner surface. Endopod 3 with seta; 2 plumose; on inner surface. Endopod 4 with seta; 2 plumose; on inner surface. Endopod 5 with seta; 2 plumose; on inner surface, or on outer surface; outer seta absent. Endopod 6 with seta; 4 plumose; on inner surface, or on outer surface.

##### Swimming legs features

**First swimming legs.** Symmetrical; biramous. First swimming legs intercoxal plate without seta. First swimming legs praecoxa absent. First swimming legs coxa with seta; one; straight; distally inserted; on inner surface; surpassing to first endopodal segment; with setules; two group; as a patch; on inner margin; and as a row; double; on anterior surface; outerly; without spinules; without spine. First swimming legs basis without seta; with setules; as a patch; single; on outer surface; without spinules; without spine. First swimming legs endopod 2-segmented. Endopod 1 with seta; straight; restricted; to inner surface; one element; without spine; with setules; as a row; single; continuously; on outer surface; without spinules; absence of Schmeil’s organ. Endopod 2 with seta; unrestricted; three on inner surface; one on outer surface; two on distal surface; straight; without spine; with setules; as a row; single; continuously; on outer surface; without spinules; absence of Schmeil’s organ. Endopod 3 absence. First swimming legs exopod 1 with seta; restricted; 1 on inner surface; with spine; 1; stout; smaller than original segment; serrated; on inner side; continuously; with setules; as a row; single; as a row; innerly. First swimming legs exopod 2 with seta; restricted; 1 on inner surface; straight; without spine; with setules; as a row; single; continuously; on inner margin, or on outer margin; without spinules. First swimming legs exopod 3 with setule; as a row; single; continuously; on outer surface; without spinules; with seta; unrestricted; 2 on inner surface; 2 on terminal surface; with spine; 2; unequal size; first no longer 2x than origin segment; stout; serrated; on inner side, or on outer side; equally; second longer 3x than origin segment; slender; serrated; on outer side; with ornamentation on non-serrated side; by setules. **Second swimming legs**. Symmetrical; Second swimming legs biramous. Second swimming legs intercoxal plate without seta. Second swimming legs praecoxa present; located laterally. Second swimming legs coxa with seta; straight; distally inserted; on inner surface; surpassing to basal segment; without setules; without spinules; without spine. Second swimming legs basis without seta; without setules; without spinules; without spine. Second swimming legs endopod 3-segmented. Endopod 1 with seta; straight; restricted; one on inner surface; without spine; with setules; as a row; single; continuously; on outer surface; without spinules; absence of Schmeil’s organ. Endopod 2 with seta; straight; unrestricted; two on inner surface; without spine; with setules; as a row; single; continuously; on outer side; without spinules; presence of Schmeil’s organ; on posterior surface. Endopod 3 with seta; straight; unrestricted; three on inner surface; two on outer surface; two on distal surface; without spine; without setules; with spinules; as a row; double; distally inserted; at anterior surface; absence of Schmeil’s organ. Second swimming legs exopod 1 with seta; restricted; one on inner surface; with spine; 1; stout; not reaching to distal-third of the exopod 2; serrated; on inner side, or on outer side; with setules; as a row; single; continuously; on inner side; without spinules; absence of Schmeil’s organ. Exopod 2 with seta; unrestricted; one on inner surface; with spine; 1; stout; not surpassing the exopod 3; serrated; on inner side, or on outer side; with setules; as a row; single; continuously; on inner surface; without spinules; absence of Schmeil’s organ. Exopod 3 with seta; plurimarginal; three on inner surface; two on terminal surface; with spine; 2; unequal size; first no longer 2x than origin segment; stout; serrated; on inner side, or on outer side; equally; second longer 2x than origin segment; slender; serrated; on outer side; with ornamentation on non-serrated side; of setules; setules on outer surface; as a row; single; continuously; on inner surface; with spinules; as a row; single; distally inserted; at anterior surface; absence of Schmeil’s organ. **Third swimming legs**. Symmetrical; Third swimming legs biramous. Third swimming legs intercoxal plate without seta. Third swimming legs praecoxa present; not laterally located. Third swimming legs coxa with seta; straight; distally inserted; on inner surface; surpassing to first endopodal segment; without setules; without spinules; without spine. Third swimming legs basis without seta; without setules; without spinules; without spine. Third swimming legs endopod 3-segmented. Endopod 1 with seta; restricted; one on inner surface; without spine; without setules; without spinules; absence of Schmeil’s organ. Endopod 2 with seta; restricted; two on inner surface; straight; without spine; without setules; without spinules; absence of Schmeil’s organ. Endopod 3 with seta; straight; plurimarginal; two on inner surface; two on outer surface; three on terminal surface; without spine; without setules; with spinules; as a row; distally inserted; double; at anterior surface; absence of Schmeil’s organ. Third swimming legs exopod 1 with seta; restricted; straight; one on inner surface; with spine; 1; stout; not reaching to the distal-third of the exopod 2; serrated; equally; on inner surface, or on outer surface; with setules; as a row; single; continuously; on inner surface; without spinules; absence of Schmeil’s organ. Exopod 2 with seta; straight; restricted; one on inner surface; with spine; 1; stout; not reaching out to exopod 3; serrated; on inner side, or on outer side; equally; with setules; as a row; single; continuously; on inner side; without spinules; absence of Schmeil’s organ. Exopod 3 without setules; with spinules; as a row; single; distally inserted; at anterior surface; with seta; straight; unrestricted; three on inner surface; two on terminal surface; with spine; 2; unequal size; first no longer 2x than origin segment; stout; serrated; on inner side, or on outer side; equally; second longer 2x than origin segment; slender; serrated; on outer side; with ornamentation on non-serrated side; of setules; absence of Schmeil’s organ. **Fourth swimming legs**. Symmetrical; biramous. Intercoxal plate without sensilla. Praecoxa present. Coxa with seta; distally inserted; on inner margin; reaching out to endopod 1; without spinules; setules absent. Basis with seta; one; medially inserted; on posterior surface; smaller than the original segment; without setules; without spinules; without spine. Fourth swimming legs endopod 3-segmented. Endopod 1 with seta; one; restricted; on inner surface; without spine; without setules; without spinules; absence of Schmeil’s organ. Endopod 2 with seta; restricted; two on inner side; without spine; with setules; as a row; single; continuously; on outer surface; without spinules; absence of Schmeil’s organ. Endopod 3 with seta; unrestricted; two on inner surface; two on outer surface; three on distal surface; without spine; without setules; with spinules; as a row; double; distally inserted; at anterior surface; absence of Schmeil’s organ. Fourth swimming legs exopod 1 with seta; restricted; one on inner surface; with spine; 1; stout; not reaching out to distal-third of the exopod 2; serrated; on inner side, or on outer side; equally; with setules; as a row; single; continuously; on inner surface; without spinules; absence of Schmeil’s organ. Exopod 2 with seta; restricted; one on inner surface; with spine; 1; stout; not reaching the end of exopod 3; serrated; on inner side, or on outer side; equally; with setules; as a row; single; continuously; on inner surface; without spinules; absence of Schmeil’s organ. Exopod 3 without setules; with spinules; as a row; single; distally inserted; at anterior surface; with seta; unrestricted; three on inner surface; two on distal surface; with spine; 2; unequal size; first no longer 2x than origin segment; stout; serrated; on inner side, or on outer side; equally; second longer 2x than origin segment; slender; serrated; on outer side; without ornamentation on non-serrated side; absence of Schmeil’s organ.

##### Fifth swimming legs features

Asymmetrical. Fifth swimming leg intercoxal plate with length not equal or greater than width on 1.5x; with irregular proximal margin; discontinuous to; the anterior margin of the left coxa, or the anterior margin of the right coxa; posterior sensilla on the right lateral absent. **Fifth left swimming leg**. Fifth left swimming leg biramous; leg reaching first right exopod segment; medially. Fifth left swimming leg praecoxa present; rudimentary; separated from the coxae; without ornamentation. Fifth left swimming leg coxa concave inner side; without teeth-like structures; with process; conical; on posterior surface; outer side; distally inserted; not projecting over basis; with sensilla; stout; triangular; at apex; no longer 2x than insertion basis; without swelling; without seta; without spinules. Fifth left swimming leg basis sub-cylindrical; unequal size between inner and outer side; shorter outer than inner side; with concave inner side; rounded internal proximal expansion absent; without outgrowth; without groove; absence of protuberance; with seta; outerly inserted; no longer 2x than origin segment; absence of minutely granular. Fifth left swimming leg endopod segments 1 and 2 fused; segments 2 and 3 fused; 1-segmented; stout; separated from the basis; ornamented; on inner side; with spinules; more than four elements; as a row; terminally; row of setules absent; without seta. Fifth left swimming leg exopod segments 1 and 2 separated; segments 2 and 3 fused; 2-segmented; stout; separated from the basis. Fifth left swimming leg exopod 1 sub-triangular; longer than broad; equal size between inner and outer side; concave inner side; convex outer side; without swelling; without marginal extension; without process; with lobe; single; semicircular; medially inserted; on inner side; covered; by setules; without outer spine; absence seta. Fifth left swimming leg exopod 2 digitiform; longer than broad; equal size between inner and outer side; disform inner side; with rectilinear outer side; setulose pad present; prominently rounded; proximally; on inner side; inflated medial region absent; distal process present; digitiform; denticulate; not bicuspidate; without transverse row of denticles; none oblique row of 5 denticles; at anterior surface; not innerly directed; with seta; spiniform; ornamented by spinules; surpassing the distal-point of the segment; without outer spine; terminal claw absent.

##### Fifth right swimming leg

Biramous. Fifth right swimming leg praecoxa present; separated from the coxae; without ornamentation. Fifth right swimming leg coxa convex inner side; without teeth-like structures; with process; conical; distally inserted; on posterior surface; closest to the outer rim; projecting over basis; beyond the first third; until the medial surface; without triangular protuberance innerly; with sensilla; slender; at apex; no longer 2x than basal insertion; without marginal extension; without seta; without spinules. Fifth right swimming leg basis cylindrical; unequal size between inner and outer side; shorter outer than inner side; concave inner side; tumescence absent; without protuberance; absence of distinct minutely granular; additional inner process absent; without posterior groove; with seta; outerly inserted; on anterior surface; no longer 2x than origin segment; posterior protrusion present; distal process absent. Fifth right swimming leg with endopodite present; separated from the basis; on anterior surface; ancestral segments 1 and 2 fused; ancestral segments 2 and 3 fused; 1-segmented; stout; ornamented; with setules; as a row; on inner side; terminally; without seta. Fifth right swimming leg exopod segments 1 and 2 separated; segments 2 and 3 fused; 2-segmented; stout; separated from the basis. Fifth right swimming leg exopod 1 trapezium; longer than broad; nearly 1.25 times; unequal size between both sides; shorter inner than outer side; rectilinear inner side; rectilinear outer side; with marginal extension; sub-triangular; distally inserted; at outer rim; spinules absent; with process; rounded; sclerotized; without ornamentation; distally inserted; at posterior surface; projecting over next segment; without outer spine; without seta; internal prominence present; acute; lamella on posterior surface absent. Fifth right swimming leg exopod 2 elliptical; longer than broad; nearly 2 times; equal size between both sides; disform inner side; convex outer side; without posterior proximal swelling; inner-posterior process absent; without marginal expansion; curved ridge on distal posterior surface present; chitinous knobs absent; with outer spine; inserted medially; rectilinear; ornamented innerly; by spinules; as a row; not ornamented outerly; sharp tip; without apparent curve; lesser than the length of the exopod 2; until to 2 times its size; 1.5x; sensilla absent; terminal claw present; equal or longer 1.5 times than insertion segment; sclerotized; arched; inward; with conspicuous curve; proximally; ornamented innerly; by spinules; as a row; partially on extension; medially, or distally; not ornamented outerly; sharp tip; not curved tip; without medial constriction; hyaline process absent.

##### FEMALE

Body longer and wider than male; Female body 1347 micrometers excluding caudal setae. Widest at first metasome segment. Distal margin of the prosomal segments without one line of setules at posterior margin. Prosome segments without spinules at prosomal segments. Fourth metasome segment absence of dorsal protuberance. Fourth and fifth metasome segments fused; totally. Limit between fourth and fifth metasome segments without ornamentation. **Fifth metasome segment**. Fifth metasome segment without sensilla; with epimeral plates. Epimeral plates asymmetrical. Right epimeral plates prominent, as projections; not thinner than the left; one posterior-laterally directed; reaching half length of the genital segment; with sensilla at the apex; dorsal-posterior sensilla present; slender; without ornamentation. Left epimeral plate with expansion; semicircular; on posterior surface; dorsally; with sensilla; at tip.

##### Urosome

3-segmented. **Genital double-somite**. Asymmetrical in dorsal view; longer than broad; longer than other urosomites combined; dorsal suture at mid-length absent; not covered by spinules; with swelling; conical; unequal size; greater left than right; anteriorly; with sensillae; on both sides; one; stout; with robust apex; at left lateral; not on lobular base; medially; one; stout; at right lateral; not on lobular base; anteriorly; with robust apex; of equal size between then; lateral protuberance absent; with right posterior rim expanded; over next segment; without slender sensilla on each posterior rim; without posterior-dorsal process. Genital double-somite opercular pad present; broader than longer; symmetrical; development laterally; expanded posteriorly; covering partially; double gonoporal slit; located ventrally; with arthrodial membrane; inserted anteriorly; post-genital process absent; disto-ventral tumescence absent; ventral vertical folds absent; dorsal sensilla absent. Second urosome segment without ventral fusion to anal segment; right distal process absent. Caudal rami patch of setules on outer surface absent; patch of spinules on outer surface present.

##### Oral appendices feature

Rostrum basal process absent. **Antennules**. Symmetrical. Right antennule surpassing to genital double-segment; extending beyond caudal rami. Right antennule exceeding the caudal setae. Right antennule ornamentation pattern equals to male left antennule; fully.

##### Fifth swimming legs

Symmetrical; Fifth swimming legs biramous. Fifth swimming legs intercoxal plate longer than wide; separated from the legs. Fifth swimming legs praecoxa with sclerite praecoxal; separated from the coxae; without ornamentation. Fifth swimming legs coxa with process; conical; at the outer rim; distally; sensilla present; stout; at apex; projecting over basal segment; longer 2x than basal insertion; marginal extension absent; without swelling; without seta; without spinules. Fifth swimming legs basis sub-triangular; unequal size between inner and outer sides; shorter outer than inner side; with convex inner side; without proximal inner outgrowth; without groove; with distal extension; on posterior surface; with seta; outerly inserted; on anterior surface; no longer 2x than origin segment. Fifth swimming legs endopod segments 1 and 2 fused; segments 2 and 3 fused; 1-segmented; stout; separated from the basis; present discontinuity cuticle; on inner side; with spinules; as a row; single; non-oblique; sub-terminally; at anterior surface; with seta; double; one medially; on posterior surface; rectilinear; one distally; on posterior surface; arched; of unequal size; distal seta longer than medial seta. Fifth swimming legs exopod segments 1 and 2 separated; segments 2 and 3 separated; 3-segmented; separated from the basis. Fifth swimming legs exopod 1 sub-cylindrical; longer than wide; longer or equal than 2 times; with unequal size between inner and outer side; shorter inner than outer side; with convex inner side; with rectilinear outer side; without swelling; without marginal extension; without posterior process; without spine; without seta. Fifth swimming legs exopod 2 sub-cylindrical; longer than broad; longer or equal than 2 times; without swelling; without marginal extension; without process; without lobe; with spine; inserted laterally; rectilinear; without ornamentation; sharp tip; equal size or larger than next segment; without seta. Fifth swimming legs exopod 3 cylindrical; longer than wide; without swelling; without process; without lobe; without spine; with seta; double; inserted terminally; unequal size between them; outer seta smaller than inner; nearly 3 times; outer seta not ornamented by setules; without ornamentation; presence of terminal claw; sclerotized; rectilinear; with ornamentation; of denticles; as a row; on surface partially; at medial region, or at distal region; concave outer side; with ornamentation; of denticles; as a row; on surface partially; at medial region; blunt tip; 6 times longer than origin segment.

##### Distribution records

###### BRAZIL

**Amazonas**: Catalão Lake, floodplain lake and Janauari Lake, near to Manaus (Brandorff, 1972); Solimões/Amazonas River, Camaleão Lake (Marchantaria Island), Paraná do Rei; Catalão Lake and Janauari Lake, Negro River, near to Manaus (Santos-Silva *et al*., 1989). **Pará**: Curuá-Una Reservoir, 2°48’38“S, 54°18’55“W (Santos-Silva *et al*., 1989). VENEZUELA. **Amazonas**: Atabapo River (Dussart, 1984a). **Bolívar**: Guri, man-made lake near the dam on Caroni River; Orinoco River, right side, at Ciudad Bolívar (Dussart, 1984a).

##### Habitat

Habitat in freshwaters: lake; floodplain lake, rivers, and reservoir.

##### Remarks

Brandorff introduced the species to science from organisms collected in the geographic Center of the Brazilian Amazon. Since their foundation, these organisms have been recorded for other lake environments in the region and artificial environments in Pará (Santos-Silva *et al*., 1989; Santos-Silva, 2008), and Venezuelan river plains (Dussart, 1984a). The species was never included in *nordestinus* complex by Wright, and an attempt at recombination for the subgenus *Notodiaptomus* (*Wrightius*) was made in Dussart (1985) but not accepted for lack of evidence.

In 1988, Dussart & Defaye suggested the recombination of *D.* (s.l.) *echinatus* for *Notodiaptomus* from organisms from French Guiana and considered *N. kieferi* a junior synonym of this species. Santos-Silva *et al*. (1989) had already warned of the lack of scientific foundations of this proposal, which should be disregarded until further examinations in the type-material of the species. This seems even more prudent when considering the characteristics of occurrence for the taxa, which suggest hypothesis of taxonomic distinction when we verify that *D.* (s.l.) *echinatus* is recorded for the Southern Neotropics, specific to the regions of Paraguay (type locality) and Argentina (Lowndes, 1934).

Effectively, the examination of the type-material of *N. kieferi* deposited in the INPA collection was made impossible due to damage, but topotypes collected in 22.IX.59, and specified for this thesis, were accessed. Consistent and relevant differences were identified between the morphological differences examined for the topotypes and those presented by Lowndes (1934) and illustrated by Dussart & Defaye (1989): (1) female of *D. echinatus* has the fourth and fifth segment of the “non-confluent” metasome (Lowndes, 1934, fig. 4a), in *N. kieferi* this separation does not exist; (2) the female of *D. echinatus*, in lateral view, has a dorsal elevation covered by numerous spinules in the last segment of the metasoma (Lowndes, 1934, fig. 4a), in *N. kieferi* this condition also does not exist; (3) the endopod of the female fifth leg is bisegmented in *D. echinatus* (Lowndes, 1934, fig. 4b), in *N. kieferi* the endopod has a single segment; (4) (4) the fifth left leg (dorsal view) reaches only the distal portion of the base of the opposite leg in males of *D. echinatus* (Lowndes, 1934, fig. 4a), in male of *N. kieferi* the reach extends beyond the base of the leg opposite and reaches the medial portion of exopod 1; and (5) the curvature of the terminal claw of the fifth right leg of the male of *D. echinatus* has a more pronounced angulation than the male of *N. kieferi*, a difference that can also be present when comparing the tip of the terminal claw of the same leg, with externally directed curvature in the first species and without curvature in the males of the second (Lowndes, 1934, fig. 4b; and Dussart & Defaye, 1989, fig. 10).

These morphological features confirm the original description of *N. kieferi.* Santos-Silva *et al*. (1989) offered important illustrations for the species from organisms from the Rio Negro, Amazonas, in Pará, and corroborated Brandorff (1973) previously. Among the characteristics adopted by Kiefer (1936; 1956) to define *Notodiapomus*, the organisms of the species have not only male fifth left leg exopod 2 with “short and thick seta”, but male fifth left swimming leg exopod 2 with spiniform seta surpassing the distal-point of the segment. Additionally, we record here morphological attributes for the species that are divergent from the type-species of *Notodiaptomus*: (1) cephalosome without dorsal suture; (2) male right antennule actual segment 8 with conical seta reaching to middle-point of the sequent segment; (3) male right antennule actual segment 13 with modified seta presenting rounded apex; (4) female right epimeral plate reaching half length of the genital segment; (5) female left epimeral plate with dorsal semicircular expansion posteriorly; (6) female genital double-somite with lateral conical swelling, greater left than right; and (7) female caudal rami with spinules patch on outer surface.

#### Notodiaptomus maracaibensis Kiefer, 1954

##### Synonymy

*Notodiaptomus maracaibensis* Kiefer, 1954: 170–174, figs. 1–11; Fuentes-Reinés *et al*., 2021: 125–132, figs. 1–5, identification key; Perbiche-Neves *et al*., 2020: 681-682, key to the Neotropical diaptomid, fig. 21.8 K.

##### Type locality

Maracaibo Lake, Maracaibo City, Venezuela.

##### Type material

In the original description, specimens collected by “Prof. Dr. Fritz Gessner” on 13.X.1954 and used for the description of the species are generically indicated. Kiefer (1956) added information on 7 females and 4 males as redescription material. No information was provided about the depository collection or condition of the material, which remains unknown.

##### Material examined

Non-type material: 3 males, and 4 females, entire in alcohol, from the Moriaties Lagoon, Venezuela, 11.II.1981, no information about collector, store in the Plankton Laboratory, INPA. 1 male (INPA-COP037, slides a-h) and 1 female (INPA-COP038, slides a-h) were selected to be dissection on eight slides each and deposited in the Zoological Collection of the INPA, Brazil.

##### Diagnosis

**(1)** Male epimeral plates asymmetrical; **(2)** male right antennule actual segment 8 with conical seta reaching to middle-point of the sequent segment; **(3)** male right antennule actual segment 12 with conical seta smaller than to those segment 8; **(4)** male fifth left swimming leg exopod 1 with medial double semicircular lobe innerly; **(5)** male fifth left swimming leg exopod 2 with medial rounded setulose pad innerly; **(6)** male fifth right swimming leg basis without inner intumescence, and protuberance; **(7)** male fifth right swimming leg exopod 1 with acute outer distal extension; **(8)** female right epimeral plate with slender sensilla on inner medial surface, and ornamented by inner spinules patch dorsally; **(9)** female left epimeral plate with medial semicircular expansion dorsally, and sensilla at apex; **(10)** female genital double-somite with dorsal suture at mid-length; **(11)** female fifth swimming leg endopod 1-segmented; **(12)** female fifth swimming leg exopod 1 with outer spine posteriorly.

##### Redescription

###### MALE

Body 930 micrometers excluding caudal setae. Male body smaller and slenderer than female. Nerve axons myelinated. Prosome 6-segmented; widest at first metasome segment; without one line of setules at posterior margin; with spinules at least at one segment. Cephalosome anterior margin rounded; with dorsal suture; incomplete; separate from first metasome segment. First metasome segment without sensilla. Second metasome segment without sensilla. Third metasome segment without sensillae; ornamented posterior margin; with spinules; as a row; single; dorsally. Fourth metasome segment without sensillae; separated from the fifth metasome. Limit between fourth and fifth metasome segments ornamented; with spinules; as a row; on dorsally singly; on lateral singly; same size. Fifth metasome segment without sensilla; Fifth metasome segment without ornamentation; dorsal conical process absent; with epimeral plates. Epimeral plates asymmetrical. Right epimeral plates prominent, as projections; one projection; posterior-laterally directed; not reaching half length of the genital segment; with sensilla; one at the apex of projection and other medially; without ornamentation. Left epimeral plate prominent, as projection; one projection; posterior-dorsally directed; reaching half length of the genital segment; with sensillae; one at the apex of projection and other medially; without ornamentation.

##### Urosome

5-segmented; Urosome 5-free segments. Genital somite symmetrical in dorsal view; with single aperture; located on left side; ventrolaterally on posterior rim; with sensillae; on both sides; one; at left lateral; posteriorly; one; at right rim; posteriorly; of equal size between then. Third urosome segment without spinules; without external seta. Fourth urosome segment without spinules; without sub-conical blunt dorsal-lateral process. Anal segment presence of dorsal sensillae; one on each side; medially inserted; presence of operculum; convex; covering the anal aperture fully. Caudal rami symmetrical; separated from anal segment; longer than wide; with setules; continuous on; inner side; each ramus bearing 6 caudal setae; 5 marginals; plumose; and 1 internal dorsally; straight; not reticulated main axis; outermost seta with outer spiniform process absent.

##### Oral appendices feature

Rostrum symmetrical; separated from dorsal cephalic shield; by complete suture; sensillae present; one pair; anteriorly inserted on surface tegument; with rostral filament; double; paired; extended; into point; with basal process; in ventral view, rounded on left side; without a smaller basal expansion on the right side.

##### Antennules

Asymmetrical. **Right antennules**. Uniramous; right antennule surpassing to genital segment; right antennule extending beyond caudal rami.

Right antennule ancestral segment I and II separated. Ancestral segment II and III fused. Ancestral segment III and IV fused. Ancestral segment IV and V separated. Ancestral segment V and VI separated. Ancestral segment VI and VII separated. Ancestral segment VII and VIII separated. Ancestral segment VIII and IX separated. Ancestral segment IX and X separated. Ancestral segment X and XI separated. Ancestral segment XI and XII separated. Ancestral segment XII and XIII separated. Ancestral segment XIII and XIV separated. Ancestral segment XIV and XV separated. Ancestral segment XV and XVI separated. Ancestral segment XVI and XVII separated. Ancestral segment XVII and XVIII separated. Ancestral segment XVIII and XIX separated. Ancestral segment XIX and XX separated. Ancestral segment XX and XXI separated. Ancestral segment XXI and XXII fused. Ancestral segment XXII and XXIII fused. Ancestral segment XXIII and XXIV separated. Ancestral segment XXIV and XXV fused. Ancestral segment XXV and XXVI separated. Ancestral segment XXVI and XXVII separated. Ancestral segment XXVII and XXVIII fused.

Right antennule actual 22-segmented; geniculated; between the segment 18 and segment 19; with swollen and modified region; formed by 5 segments; between 13 and 17 segments. Actual segment 1 with seta; one element; straight; none larger than segment; without spinules; without vestigial seta; without conical seta; without modified seta; without spinous process; with aesthetasc; one element. Actual segment 2 with seta; three elements; of unequal size; straight; none larger than segment; without spinules; with vestigial seta; one element; without conical seta; without modified seta; without spinous process; with aesthetasc; one element. Actual segment 3 with seta; one element; one larger than segment; surpassing to distal margin; beyond three sequential segments; straight; blunt apex; without spinules; with vestigial seta; one element; without conical seta; without modified seta; without spinous process; with aesthetasc. Actual segment 4 with seta; one element; one larger than segment; surpassing to distal margin; straight; not beyond three sequential segments; without spinules; without vestigial seta; without conical seta; without modified seta; without spinous process; without aesthetasc. Actual segment 5 with seta; one element; straight; one larger than segment; surpassing to distal margin; not beyond three sequential segments; without spinules; with vestigial seta; one element; without conical seta; without modified seta; without spinous process; with aesthetasc; one element. Actual segment 6 with seta; one element; none larger than segment; straight; without spinules; without vestigial seta; without conical seta; without modified seta; without spinous process; without aesthetasc. Actual segment 7 with seta; one element; straight; one larger than segment; surpassing to distal margin; beyond three sequential segments; blunt apex; without spinules; without vestigial seta; without conical seta; without modified seta; without spinous process; with aesthetasc; one element. Actual segment 8 with seta; one element; straight; none larger than segment; without spinules; without vestigial seta; with conical seta; one element; reaching to middle-point of the sequent segment; without modified seta; without spinous process; without aesthetasc. Actual segment 9 with seta; two elements; of unequal size; straight; one larger than segment; surpassing to distal margin; beyond three sequential segments; blunt apex; without spinules; without vestigial seta; without conical seta; without modified seta; without spinous process; with aesthetasc; one element. Actual segment 10 with seta; one element; straight; none larger than segment; without spinules; without vestigial seta; without conical seta; with modified seta; presenting blunt apex; slender form; surpassing to distal margin; beyond of the sequential segment; parallel to antennule direction; without spinous process; without aesthetasc. Actual segment 11 with seta; one element; straight; one larger than segment; surpassing to distal margin; not beyond three sequential segments; without spinules; without vestigial seta; without conical seta; with modified seta; slender form; presenting blunt apex; surpassing to distal margin; beyond of the sequential segment; parallel to antennule direction; without spinous process; shorter length than homologous of actual segment 13; without aesthetasc. Actual segment 12 with seta; one element; straight; one larger than segment; surpassing to distal margin; not beyond three sequential segments; without spinules; without vestigial seta; with conical seta; one element; smaller than to segment 8; without modified seta; without spinous process; with aesthetasc; one element; absent internal perpendicular fission. Actual segment 13 with seta; one element; straight; one larger than segment; surpassing to distal margin; not beyond three sequential segments; without spinules; without vestigial seta; without conical seta; with modified seta; stout form; surpassing to distal margin; to the middle-point of the sequence segment; parallel to antennule direction; presenting bifid apex; without spinous process; with aesthetasc; one element. Actual segment 14 with seta; two elements; of unequal size; straight; one larger than segment; surpassing to distal margin; beyond three sequential segments; blunt apex; without spinules; without vestigial seta; without conical seta; without modified seta; without spinous process; with aesthetasc; one element. Actual segment 15 with seta; two elements; of unequal size; straight; not bifidform; none larger than segment; without spinules; without vestigial seta; without conical seta; without modified seta; with spinous process; on outer margin; surpassing distal margin; with aesthetasc; one element. Actual segment 16 with seta; two elements; of unequal size; plumose; one larger than segment; surpassing to distal margin; not beyond three sequential segments; not bifidform; without spinules; without vestigial seta; without conical seta; without modified seta; with spinous process; on outer margin; surpassing distal margin; unequal size to process on preceding segment; with aesthetasc; one element. Actual segment 17 with seta; two elements; of unequal size; straight; none larger than segment; bifidform; without spinules; without vestigial seta; without conical seta; with modified seta; one element; stout form; surpassing to distal margin; not beyond of the sequential segment; parallel to antennule direction; without spinous process; without aesthetasc. Actual segment 18 with seta; two elements; of equal size; straight; none larger than segment; without spinules; without vestigial seta; without conical seta; with modified seta; one element; stout form; surpassing distal margin; parallel to antennule direction; without spinous process; without aesthetasc. Actual segment 19 with seta; two elements; of unequal size; plumose; none larger than segment; without spinules; without vestigial seta; without conical seta; with modified seta; two elements; stout form; at least one bifid form; surpassing distal margin; parallel to antennule direction; without spinous process; with aesthetasc; one element. Actual segment 20 with seta; four elements; of unequal size; straight; one larger than segment; surpassing to distal margin; beyond three sequential segments; without spinules; without vestigial seta; without conical seta; without modified seta; without spinous process; without aesthetasc. Actual segment 21 with seta; two elements; of equal size; plumose; one larger than segment; surpassing to distal margin; greater 3x than original segment; without spinules; without vestigial seta; without conical seta; without modified seta; without spinous process; without aesthetasc. Actual segment 22 with seta; five elements; of equal size; one larger than segment; plumose; surpassing to distal margin; greater 3x than original segment; without spinules; without vestigial seta; without conical seta; without modified seta; without spinous process; with aesthetasc; one element.

##### Left antennules

Uniramous; Left antennule surpassing to prosome; Left antennule extending beyond caudal rami. Ancestral segment I and II separated. Ancestral segment II and III fused. Ancestral segment III and IV fused. Ancestral segment IV and V separated. Ancestral segment V and VI separated. Ancestral segment VI and VII separated. Ancestral segment VII and VIII separated. Ancestral segment VIII and IX separated. Ancestral segment IX and X separated. Ancestral segment X and XI separated. Ancestral segment XI and XII separated. Ancestral segment XII and XIII separated. Ancestral segment XIII and XIV separated. Ancestral segment XIV and XV separated. Ancestral segment XV and XVI separated. Ancestral segment XVI and XVII separated. Ancestral segment XVII and XVIII separated. Ancestral segment XVIII and XIX separated. Ancestral segment XIX and XX separated. Ancestral segment XX and XXI separated. Ancestral segment XXI and XXII separated. Ancestral segment XXII and XXIII separated. Ancestral segment XXIII and XXIV separated. Ancestral segment XXIV and XXV separated. Ancestral segment XXV and XXVI separated. Ancestral segment XXVI and XXVII separated. Ancestral segment XXVII and XXVIII fused.

Left antennule actual 25-segmented; not-geniculated. Actual segment 1 with seta; one element; none larger than segment; straight; without spinules; without vestigial seta; without conical seta; without modified seta; without spinous process; with aesthetasc; one element. Actual segment 2 with seta; three elements; of equal size; none larger than segment; straight; without spinules; with vestigial seta; one element; without conical seta; without modified seta; without spinous process; with aesthetasc; one element. Actual segment 3 with seta; one element; one larger than segment; straight; surpassing to distal margin; beyond three sequential segments; without spinules; with vestigial seta; one element; without conical seta; without modified seta; without spinous process; with aesthetasc. Actual segment 4 with seta; one element; none larger than segment; straight; without spinules; without vestigial seta; without conical seta; without modified seta; without spinous process; without aesthetasc. Actual segment 5 with seta; one element; one larger than segment; straight; surpassing to distal margin; not beyond three sequential segments; without spinules; with vestigial seta; one element; without conical seta; without modified seta; without spinous process; with aesthetasc; one element. Actual segment 6 with seta; one element; none larger than segment; straight; without spinules; without vestigial seta; without conical seta; without modified seta; without spinous process; without aesthetasc. Actual segment 7 with seta; one element; one larger than segment; straight; surpassing to distal margin; beyond three sequential segments; without spinules; without vestigial seta; without conical seta; without modified seta; without spinous process; with aesthetasc; one element. Actual segment 8 with seta; one element; one larger than segment; straight; surpassing distal margin; without spinules; without vestigial seta; with conical seta; without modified seta; without spinous process; without aesthetasc. Actual segment 9 with seta; two elements; of unequal size; one larger than segment; straight; surpassing to distal margin; beyond three sequential segments; without spinules; without vestigial seta; without conical seta; without modified seta; without spinous process; with aesthetasc; one element. Actual segment 10 with seta; one element; none larger than segment; straight; without spinules; without vestigial seta; without conical seta; without modified seta; without spinous process; without aesthetasc. Actual segment 11 with seta; one element; one larger than segment; straight; surpassing to distal margin; beyond three sequential segments; without spinules; without vestigial seta; without conical seta; without modified seta; without spinous process; without aesthetasc. Actual segment 12 with seta; one element; one larger than segment; straight; surpassing distal margin; without spinules; without vestigial seta; with conical seta; without modified seta; without spinous process; with aesthetasc; one element. Actual segment 13 with seta; one element; none elongated; straight; surpassing distal margin; without spinules; without vestigial seta; without conical seta; without modified seta; without spinous process; without aesthetasc. Actual segment 14 with seta; one element; elongated; straight; surpassing to distal margin; beyond three sequential segments; without spinules; without vestigial seta; without conical seta; without modified seta; without spinous process; with aesthetasc; one element. Actual segment 15 with seta; one element; larger than segment; straight; surpassing to distal margin; not beyond three sequential segments; without spinules; without vestigial seta; without conical seta; without modified seta; without spinous process; without aesthetasc. Actual segment 16 with seta; one element; larger than segment; plumose; surpassing to distal margin; not beyond three sequential segments; without spinules; without vestigial seta; without conical seta; without modified seta; without spinous process; with aesthetasc; one element. Actual segment 17 with seta; one element; not larger than segment; straight; without spinules; without vestigial seta; without conical seta; without modified seta; without spinous process; without aesthetasc. Actual segment 18 with seta; one element; larger than segment; straight; surpassing to distal margin; beyond three sequential segments; without spinules; without vestigial seta; without conical seta; without modified seta; without spinous process; without aesthetasc. Actual segment 19 with seta; one element; not larger than segment; straight; surpassing distal margin; without spinules; without vestigial seta; without conical seta; without modified seta; without spinous process; with aesthetasc; one element. Actual segment 20 with seta; one element; not larger than segment; straight; surpassing distal margin; without spinules; without vestigial seta; without conical seta; without modified seta; without spinous process; without aesthetasc. Actual segment 21 with seta; one element; larger than segment; plumose; surpassing to distal margin; beyond three sequential segments; without spinules; without vestigial seta; without conical seta; without modified seta; without spinous process; without aesthetasc. Actual segment 22 with seta; two elements; of unequal size; one of them elongated; plumose; surpassing to distal margin; without spinules; without vestigial seta; without conical seta; without modified seta; without spinous process; without aesthetasc. Actual segment 23 with seta; two elements; of unequal size; one larger than segment; plumose; surpassing to distal margin; greater 3x than original segment; without spinules; without vestigial seta; without conical seta; without modified seta; without spinous process; without aesthetasc. Actual segment 24 with seta; two elements; of equal size; one larger than segment; plumose; surpassing to distal margin; greater 3x than original segment; without spinules; without vestigial seta; without conical seta; without modified seta; without spinous process; without aesthetasc. Actual segment 25 with seta; four elements; of equal size; elongated; plumose; surpassing to distal margin; 4 times larger than segment; without spinules; without vestigial seta; without conical seta; without modified seta; without spinous process; with aesthetasc; one element.

##### Antenna

Biramous. Antenna coxa separated from the basis; bearing seta; 1; on inner surface; at distal corner; reaching to the endopod 1. Antenna basis (fusion) separated from the endopodal segment; bearing seta; 2; on inner surface; at distal corner. Endopodal ancestral segment I and II separated. Ancestral segment II and III fused. Ancestral segment III and IV fused. Ancestral segment III and IV fully. Antenna endopod actual 2-segmented. Actual segment 1 not bilobate; with seta; two; on inner margin; with spinules; as a row; obliquely; on outer surface; with pore. Actual segment 2 bilobate; with discontinuity on outer cuticle; not developed as a suture; inner lobe bearing 8 setae; distally; outer lobe bearing 7 setae; distally; with spinules; as a patch; on outer surface. Antenna exopod ancestral segment I and II separated. Ancestral segment II and III fused. Ancestral segment III and IV fused. Ancestral segment IV and V separated. Ancestral segment V and VI separated. Ancestral segment VI and VII separated. Ancestral segment VII and VIII separated. Ancestral segment VIII and IX separated. Ancestral segment IX and X fused. Antenna exopod actual 7-segmented. Actual segment 1 single; elongated (width-length, equal or larger ratio 2:1); with seta; one; at inner surface. Actual segment 2 compound; elongated (larger width-length ratio 2:1); with seta; three; at inner surface. Actual segment 3 single; not elongated (lesser width-length ratio 2:1); with seta; one; at inner surface. Actual segment 4 single; not elongated (lesser width-length ratio 2:1); with seta; one; at inner surface. Actual segment 5 single; not elongated (lesser width-length ratio 2:1); with seta; one; at inner surface. Actual segment 6 single; not elongated (lesser width-length ratio 2:1); with seta; one; at inner surface. Actual segment 7 compound; elongated (larger or equal width-length ratio 2:1); with seta; one; at inner surface; and three; at distal surface.

##### Oral features

**Mandible**. Coxal gnathobase sclerotized; with lobe; prominent; on caudal margin; presence of cutting blade; with tooth-like prominence; two, distinctly; 1 acute; on caudal margin; and 1 triangular; on sub-caudal margin; without acute projection between the prominences; with additional spinules; as a row; on dorsal surface; with seta; 1; dorsally; on apical surface; with spinules; apicalmost. Mandible palps biramous; comprising the basis; with seta; four; differently inserted; first medially; reaching to beyond the endopod 1; second distally; third distally; fourth distally; on inner margin; none with setulose ornamentation. Mandible endopod 2-segmented. Mandible endopod 1 with lobe; bearing seta; four; distally inserted; without spinules. Mandible endopod 2 without lobe; bearing setae; nine elements; distally inserted; with spinules; as a row; double. Mandible exopod 4-segmented. Mandible exopod 1 with seta; one element; distally; on inner margin. Mandible exopod 2 with seta; one element; distally; on inner side. Mandible exopod 3 with seta; one element; distally; on inner side. Mandible exopod 4 with setae; three elements; on terminal region. **Maxillule**. Birramous. Maxillule 3-segmented. Maxillule praecoxa with praecoxal arthrite; bearing spines; fifteen elements; ten marginally; plus five sub-marginally; with spinules; as a patch; on sub-marginal surface. Maxillule coxa with coxal epipodite; with conspicuous outer lobe; bearing setae; nine elements; with coxal endite; elongated (larger or equal width-length ratio 2:1); bearing setae; four elements. Maxillule basis with basal endite; double; first proximal; elongated (larger width-length ratio 2:1; separated from basis; with setae; four elements; distally inserted; second distal; fused to basis; not elongated (lesser width-length ratio 2:1); with setae; four elements; distally inserted; with setules; as a row; on inner side; basal exite present; with setae; one element; on outer surface. Maxillule endopod 1-segmented. Endopod 1 bilobate; first proximal; with setae; three elements; second distal; with setae; five elements. Maxillule exopod 1-segmented. Exopod 1 with setae; six elements; with setules; as a row; on inner side; spinules absent. **Maxilla**. Uniramous. Maxilla 5-segmented. Maxilla praecoxa fused to coxa; incompletely; distinct externally; with praecoxal endite; double; first elongated endite (larger or equal width length ratio 2:1); proximally inserted; with seta; straight, or plumose; 1 straight; 4 plumose; with spine; single; without spinules; without setule; second elongated endite (larger or equal width length ratio 2:1); distally inserted; with seta; plumose; 3 plumose; without spine; with spinules; as a row; on distal margin; with setule; as a row; on distal margin; absence of outer seta. Maxilla coxa with coxal endite; double; first elongated endite (larger or equal width); proximally inserted; with seta; plumose; 3 plumose; without spine; without spinules; with setules; as a row; on proximal margin; second elongated endite (larger or equal width); distally inserted; with seta; plumose; 3 plumose; without spine; without spinules; with setules; as a row; on proximal margin; absence of outer seta. Maxilla basis with basal endite; single; elongated (larger or equal width-length ratio 2:1); with seta; plumose; 3 plumose; without spinules; absence of outer seta. Maxilla endopod 2-segmented. Endopod 1 with seta; 2 plumose; without spine; without spinules; without setules. Maxilla endopod 2 with seta; 2 plumose; without spine; without spinules; without setules. **Maxilliped**. Uniramous; Maxilliped 8-segmented. Maxilliped praecoxa fused to coxa; incompletely; distinct internally; with praecoxal endite; not elongated (lesser width-length ratio 2:1); distally inserted; with seta; 1 straight; with spinules; as a row; single; on basal surface; without setules. Maxilliped coxa with coxal endite; three coxal endite; first elongated (larger or equal width); proximally inserted; with seta; 2 plumose; with spinules; as a patch; single; on apical surface; without setules; second not elongated (lesser width-length ratio 2:1); medially inserted; with seta; 3 plumose; with spinules; as a row; single; on medial surface; without setules; third elongated (larger or equal width length ratio 2:1); distally inserted; with seta; 3 plumose; none reaching to beyond of the basis; with spinules; as a row; single; on basal surface; without setules; with lobe; prominence; at inner distal angle; ornamented; with spinules; continuously on margin. Maxilliped basis without basal endite; with seta; 3 plumose; with spinules; as a row; single; on medial surface; with setules; as a row; single; on inner margin. Maxilliped endopod segment 6-segmented. Endopod 1 with seta; 2 plumose; on inner surface. Endopod 2 with seta; 3 plumose; on inner surface. Endopod 3 with seta; 2 plumose; on inner surface. Endopod 4 with seta; 2 plumose; on inner surface. Endopod 5 with seta; 2 plumose; on inner surface, or on outer surface; outer seta absent. Endopod 6 with seta; 4 plumose; on inner surface, or on outer surface.

##### Swimming legs features

**First swimming legs.** Symmetrical; biramous. First swimming legs intercoxal plate without seta. First swimming legs praecoxa absent. First swimming legs coxa with seta; one; straight; distally inserted; on inner surface; surpassing to first endopodal segment; with setules; two group; as a patch; on inner margin; and as a row; double; on anterior surface; outerly; without spinules; without spine. First swimming legs basis without seta; with setules; as a patch; single; on outer surface; without spinules; without spine. First swimming legs endopod 2-segmented. Endopod 1 with seta; straight; restricted; to inner surface; one element; without spine; with setules; as a row; single; continuously; on outer surface; without spinules; absence of Schmeil’s organ. Endopod 2 with seta; unrestricted; three on inner surface; one on outer surface; two on distal surface; straight; without spine; with setules; as a row; single; continuously; on outer surface; without spinules; absence of Schmeil’s organ. Endopod 3 absence. First swimming legs exopod 1 with seta; restricted; 1 on inner surface; with spine; 1; stout; smaller than original segment; serrated; on inner side; continuously; with setules; as a row; single; as a row; innerly. First swimming legs exopod 2 with seta; restricted; 1 on inner surface; straight; without spine; with setules; as a row; single; continuously; on inner margin, or on outer margin; without spinules. First swimming legs exopod 3 with setule; as a row; single; continuously; on outer surface; without spinules; with seta; unrestricted; 2 on inner surface; 2 on terminal surface; with spine; 2; unequal size; first no longer 2x than origin segment; stout; serrated; on inner side, or on outer side; equally; second longer 3x than origin segment; slender; serrated; on outer side; with ornamentation on non-serrated side; by setules. **Second swimming legs**. Symmetrical; Second swimming legs biramous. Second swimming legs intercoxal plate without seta. Second swimming legs praecoxa present; located laterally. Second swimming legs coxa with seta; straight; distally inserted; on inner surface; surpassing to basal segment; without setules; without spinules; without spine. Second swimming legs basis without seta; without setules; without spinules; without spine. Second swimming legs endopod 3-segmented. Endopod 1 with seta; straight; restricted; one on inner surface; without spine; with setules; as a row; single; continuously; on outer surface; without spinules; absence of Schmeil’s organ. Endopod 2 with seta; straight; unrestricted; two on inner surface; without spine; with setules; as a row; single; continuously; on outer side; without spinules; presence of Schmeil’s organ; on posterior surface. Endopod 3 with seta; straight; unrestricted; three on inner surface; two on outer surface; two on distal surface; without spine; without setules; with spinules; as a row; double; distally inserted; at anterior surface; absence of Schmeil’s organ. Second swimming legs exopod 1 with seta; restricted; one on inner surface; with spine; 1; stout; not reaching to distal-third of the exopod 2; serrated; on inner side, or on outer side; with setules; as a row; single; continuously; on inner side; without spinules; absence of Schmeil’s organ. Exopod 2 with seta; unrestricted; one on inner surface; with spine; 1; stout; not surpassing the exopod 3; serrated; on inner side, or on outer side; with setules; as a row; single; continuously; on inner surface; without spinules; absence of Schmeil’s organ. Exopod 3 with seta; plurimarginal; three on inner surface; two on terminal surface; with spine; 2; unequal size; first no longer 2x than origin segment; stout; serrated; on inner side, or on outer side; equally; second longer 2x than origin segment; slender; serrated; on outer side; with ornamentation on non-serrated side; of setules; setules on outer surface; as a row; single; continuously; on inner surface; with spinules; as a row; single; distally inserted; at anterior surface; absence of Schmeil’s organ. **Third swimming legs**. Symmetrical; Third swimming legs biramous. Third swimming legs intercoxal plate without seta. Third swimming legs praecoxa present; not laterally located. Third swimming legs coxa with seta; straight; distally inserted; on inner surface; surpassing to first endopodal segment; without setules; without spinules; without spine. Third swimming legs basis without seta; without setules; without spinules; without spine. Third swimming legs endopod 3-segmented. Endopod 1 with seta; restricted; one on inner surface; without spine; without setules; without spinules; absence of Schmeil’s organ. Endopod 2 with seta; restricted; two on inner surface; straight; without spine; without setules; without spinules; absence of Schmeil’s organ. Endopod 3 with seta; straight; plurimarginal; two on inner surface; two on outer surface; three on terminal surface; without spine; without setules; with spinules; as a row; distally inserted; double; at anterior surface; absence of Schmeil’s organ. Third swimming legs exopod 1 with seta; restricted; straight; one on inner surface; with spine; 1; stout; not reaching to the distal-third of the exopod 2; serrated; equally; on inner surface, or on outer surface; with setules; as a row; single; continuously; on inner surface; without spinules; absence of Schmeil’s organ. Exopod 2 with seta; straight; restricted; one on inner surface; with spine; 1; stout; not reaching out to exopod 3; serrated; on inner side, or on outer side; equally; with setules; as a row; single; continuously; on inner side; without spinules; absence of Schmeil’s organ. Exopod 3 without setules; with spinules; as a row; single; distally inserted; at anterior surface; with seta; straight; unrestricted; three on inner surface; two on terminal surface; with spine; 2; unequal size; first no longer 2x than origin segment; stout; serrated; on inner side, or on outer side; equally; second longer 2x than origin segment; slender; serrated; on outer side; with ornamentation on non-serrated side; of setules; absence of Schmeil’s organ. **Fourth swimming legs**. Symmetrical; biramous. Intercoxal plate without sensilla. Praecoxa present. Coxa with seta; distally inserted; on inner margin; reaching out to endopod 1; without spinules; setules absent. Basis with seta; one; medially inserted; on posterior surface; smaller than the original segment; without setules; without spinules; without spine. Fourth swimming legs endopod 3-segmented. Endopod 1 with seta; one; restricted; on inner surface; without spine; without setules; without spinules; absence of Schmeil’s organ. Endopod 2 with seta; restricted; two on inner side; without spine; with setules; as a row; single; continuously; on outer surface; without spinules; absence of Schmeil’s organ. Endopod 3 with seta; unrestricted; two on inner surface; two on outer surface; three on distal surface; without spine; without setules; with spinules; as a row; double; distally inserted; at anterior surface; absence of Schmeil’s organ. Fourth swimming legs exopod 1 with seta; restricted; one on inner surface; with spine; 1; stout; not reaching out to distal-third of the exopod 2; serrated; on inner side, or on outer side; equally; with setules; as a row; single; continuously; on inner surface; without spinules; absence of Schmeil’s organ. Exopod 2 with seta; restricted; one on inner surface; with spine; 1; stout; not reaching the end of exopod 3; serrated; on inner side, or on outer side; equally; with setules; as a row; single; continuously; on inner surface; without spinules; absence of Schmeil’s organ. Exopod 3 without setules; with spinules; as a row; single; distally inserted; at anterior surface; with seta; unrestricted; three on inner surface; two on distal surface; with spine; 2; unequal size; first no longer 2x than origin segment; stout; serrated; on inner side, or on outer side; equally; second longer 2x than origin segment; slender; serrated; on outer side; without ornamentation on non-serrated side; absence of Schmeil’s organ.

##### Fifth swimming legs features

Asymmetrical. Fifth swimming leg intercoxal plate with length not equal or greater than width on 1.5x; with irregular proximal margin; discontinuous to; the anterior margin of the left coxa, or the anterior margin of the right coxa; posterior sensilla on the right lateral absent. **Fifth left swimming leg**. Fifth left swimming leg biramous; leg surpassing first right exopod segment. Fifth left swimming leg praecoxa present; rudimentary; separated from the coxae; without ornamentation. Fifth left swimming leg coxa concave inner side; without teeth-like structures; with process; conical; on posterior surface; outer side; distally inserted; not projecting over basis; with sensilla; stout; triangular; at apex; no longer 2x than insertion basis; without swelling; without seta; without spinules. Fifth left swimming leg basis sub-cylindrical; unequal size between inner and outer side; shorter outer than inner side; with rectilinear inner side; rounded internal proximal expansion absent; without outgrowth; without groove; absence of protuberance; with seta; outerly inserted; no longer 2x than origin segment; absence of minutely granular. Fifth left swimming leg endopod segments 1 and 2 fused; segments 2 and 3 fused; 1-segmented; stout; separated from the basis; ornamented; on inner side; with spinules; more than four elements; as a row; terminally; row of setules absent; with seta. Fifth left swimming leg exopod segments 1 and 2 separated; segments 2 and 3 fused; 2-segmented; stout; separated from the basis. Fifth left swimming leg exopod 1 sub-cylindrical; longer than broad; equal size between inner and outer side; rectilinear inner side; convex outer side; without swelling; without marginal extension; without process; with lobe; double; semicircular; medially inserted; on inner side; covered; by setules; without outer spine; absence seta. Fifth left swimming leg exopod 2 sub-triangular; longer than broad; unequal size between inner and outer side; shorter outer than inner side; disform inner side; with rectilinear outer side; setulose pad present; prominently rounded; medially; on inner side; inflated medial region absent; distal process present; digitiform; denticulate; not bicuspidate; without transverse row of denticles; none oblique row of 5 denticles; at anterior surface; not innerly directed; with seta; spiniform; not ornamented by spinules; not surpassing the distal-point of the segment; without outer spine; terminal claw absent.

##### Fifth right swimming leg

Biramous. Fifth right swimming leg praecoxa present; separated from the coxae; without ornamentation. Fifth right swimming leg coxa convex inner side; without teeth-like structures; with process; conical; distally inserted; on posterior surface; closest to the outer rim; projecting over basis; beyond the first third; until the medial surface; without triangular protuberance innerly; with sensilla; slender; at apex; no longer 2x than basal insertion; without marginal extension; without seta; without spinules. Fifth right swimming leg basis trapezoidal; unequal size between inner and outer side; shorter outer than inner side; convex inner side; tumescence absent; without protuberance; absence of distinct minutely granular; additional inner process absent; with posterior groove; deep; longitudinally; not reaching the endopodal lobe; ornamented; with tubercles; throughout of the outer border; with seta; outerly inserted; on anterior surface; no longer 2x than origin segment; posterior protrusion present; distal process absent. Fifth right swimming leg with endopodite present; separated from the basis, on anterior surface; ancestral segments 1 and 2 fused; ancestral segments 2 and 3 fused; 1-segmented; stout; ornamented; with setules; as a row; on inner side; terminally; without seta. Fifth right swimming leg exopod segments 1 and 2 separated; segments 2 and 3 fused; 2-segmented; stout; separated from the basis. Fifth right swimming leg exopod 1 trapezium; longer than broad; nearly 1.25 times; unequal size between both sides; shorter inner than outer side; rectilinear inner side; concave outer side; with marginal extension; acute; distally inserted; at outer rim; spinules absent; with process; triangular; rectilinear; blunt tip; sclerotized; without ornamentation; distally inserted; at posterior surface; projecting over next segment; without outer spine; without seta; internal prominence absent; lamella on posterior surface absent. Fifth right swimming leg exopod 2 cylindrical; longer than broad; nearly 2.5 times; equal size between both sides; uniform inner side; convex outer side; without posterior proximal swelling; inner-posterior process absent; without marginal expansion; curved ridge on distal posterior surface present; chitinous knobs absent; with outer spine; inserted sub-distally; rectilinear; ornamented innerly; by spinules; as a row; not ornamented outerly; sharp tip; without apparent curve; lesser than the length of the exopod 2; until to 2 times its size; 1.5x; sensilla absent; terminal claw present; equal or longer 1.5 times than insertion segment; sclerotized; arched; inward; with conspicuous curve; distally; ornamented innerly; by spinules; as a row; partially on extension; medially, or distally; not ornamented outerly; sharp tip; not curved tip; without medial constriction; hyaline process absent.

##### FEMALE

Body longer and wider than male; Female body 1095 micrometers excluding caudal setae. Widest at first metasome segment. Distal margin of the prosomal segments without one line of setules at posterior margin. Prosome segments with spinules at least at one prosomal segment. Fourth metasome segment absence of dorsal protuberance. Fourth and fifth metasome segments fused; totally. Limit between fourth and fifth metasome segments without ornamentation. **Fifth metasome segment**. Fifth metasome segment without sensilla; with epimeral plates. Epimeral plates asymmetrical. Right epimeral plates prominent, as projections; thinner than the left; one posterior-dorsally directed; not reaching half length of the genital segment; with sensilla at the apex; dorsal-posterior sensilla present; slender; ornamented; with spinules; as a patch; innerly; on dorsal surface. Left epimeral plate with expansion; semicircular; on medial surface; dorsally; with sensilla; at tip.

##### Urosome

3-segmented. **Genital double-somite**. Asymmetrical in dorsal view; longer than broad; longer than other urosomites combined; dorsal suture at mid-length present; not covered by spinules; with swelling; rounded; unequal size; greater left than right; anteriorly; with sensillae; on both sides; one; stout; with robust apex; at left lateral; not on lobular base; anteriorly; one; stout; at right lateral; not on lobular base; anteriorly; with robust apex; of equal size between then; lateral protuberance absent; without right posterior rim expanded; without slender sensilla on each posterior rim; without posterior-dorsal process. Genital double-somite opercular pad present; broader than longer; symmetrical; development laterally; expanded posteriorly; covering partially; double gonoporal slit; located ventrally; with arthrodial membrane; inserted anteriorly; post-genital process absent; disto-ventral tumescence absent; ventral vertical folds absent; dorsal sensilla absent. Second urosome segment without ventral fusion to anal segment; right distal process absent. Caudal rami patch of setules on outer surface absent; patch of spinules on outer surface absent.

##### Oral appendices feature

Rostrum basal process absent. **Antennules**. Symmetrical. Right antennule surpassing to genital double-segment; extending beyond caudal rami. Right antennule not exceeding the caudal setae. Right antennule ornamentation pattern equals to male left antennule; fully.

##### Fifth swimming legs

Symmetrical; Fifth swimming legs biramous. Fifth swimming legs intercoxal plate longer than wide; separated from the legs. Fifth swimming legs praecoxa with sclerite praecoxal; separated from the coxae; without ornamentation. Fifth swimming legs coxa with process; conical; at the outer rim; distally; sensilla present; stout; at apex; projecting over basal segment; longer 2x than basal insertion; marginal extension absent; without swelling; without seta; without spinules. Fifth swimming legs basis sub-triangular; unequal size between inner and outer sides; shorter outer than inner side; with convex inner side; without proximal inner outgrowth; without groove; without distal extension; with seta; outerly inserted; on anterior surface; no longer 2x than origin segment. Fifth swimming legs endopod segments 1 and 2 fused; segments 2 and 3 fused; 1-segmented; stout; separated from the basis; with spinules; as a row; single; non-oblique; sub-terminally; at anterior surface; with seta; double; one medially; on posterior surface; rectilinear; one distally; on posterior surface; arched; of unequal size; distal seta longer than medial seta. Fifth swimming legs exopod segments 1 and 2 separated; segments 2 and 3 separated; 3-segmented; separated from the basis. Fifth swimming legs exopod 1 sub-cylindrical; longer than wide; longer or equal than 2 times; with unequal size between inner and outer side; shorter inner than outer side; with convex inner side; with rectilinear outer side; without swelling; without marginal extension; without posterior process; with outer spine posteriorly; without seta. Fifth swimming legs exopod 2 sub-cylindrical; longer than broad; longer or equal than 2 times; without swelling; without marginal extension; without process; without lobe; with spine; inserted laterally; rectilinear; without ornamentation; sharp tip; equal size or larger than next segment; without seta. Fifth swimming legs exopod 3 cylindrical; longer than wide; without swelling; without process; without lobe; without spine; with seta; double; inserted terminally; unequal size between them; outer seta smaller than inner; nearly 3 times; outer seta not ornamented by setules; without ornamentation; presence of terminal claw; sclerotized; arched; externally directed; convex inner side; with ornamentation; of denticles; as a row; on surface fully; concave outer side; with ornamentation; of denticles; as a row; on surface partially; at medial region; blunt tip; 6 times longer than origin segment.

##### Distribution records

###### VENEZUELA

**Maracaibo**: Maracaibo Lake (Kiefer, 1954; Kiefer, 1956); COLOMBIA. Temporary pond at Ebanal farm, northern sector of La Guajira-Colombia, (11°45’23.37” N; 72°25’10.97” W) (Fuentes-Reinés *et al*., 2021).

##### Habitat

Habitat in freshwaters: lakes, and temporary pond.

##### Remarks

The species was presented to science from organisms collected in a lake environment in Venezuela. Kiefer (1954) offered a summary description of the fifth swimming leg of the male and female, and right antennule for the first, without any mention of the place of storage of the type-material. In 1956, in the amplification of *Notodiaptomus* Kiefer indicated 7 females and 4 males in the material description of the species but did not present additional morphological information or depository collection. *N. maracaibensis* has since only been recorded again 65 years later in Colombia (Fuentes-Reinés *et al*., 2021).

In the original description a close relationship with *N. deitersi* was indicated, without the presentation of the set of morphological similarities to support this. The attributes highlighted for the female of the species were, mainly: (1) last metasome segments fused dorsally and without chitinous protuberance; (2) fifth metasome segment with rounded “left corner” and “somewhat elongated” side right, with “small and thin” spines; and (3) antennule reaching the tip of the caudal setae. For males: (1) “notable thorn” in segment 8 of the right antennule; (2) segment 14 without “spine”, 15 and 16 with “strong spine” and antepenultimate segment with narrow hyaline membrane; (3) male fifth right leg exopod 2 with outer spine sub-distally, facing out and back obliquely; (4) male fifth right leg exopod 2 with strongly curved claw terminal in the basal region, straight and curved in the final region again; (5) male fifth right and left basis without ornamentation posteriorly.

In the present effort we accessed specimens collected from the Moriaties Lagoon, Venezuela and only could not confirm the attributes for male 3 and 5. All males in this locality have (3) male fifth right swimming leg exopod 2 with outer spine rectilinear, 1.5x lesser than length original segment, and (5) male fifth right swimming leg basis with posterior groove ornamented with tubercles on outer border. Additionally, the male right antennule actual segment 20 did not have a narrow hyaline membrane, portrayed in this research as a spinous process.

In describing floodplain specimens in Colombia, it is intriguing to note that Fuentes-Reinés *et al*. (2021) did not observe any of these variations reported here. Of the original attributes, the fusion for female fourth and fifth metasomal segments, and male fifth right swimming leg exopod 2 with outer rectilinear form are not mentioned by the authors. The description and representation of the male right antennule are mistaken in relation to the recognition of the segments. Possibly, this is due to the count from the integumentary part of the joint to the cephalosome, dissected along the segment 1. Except for this, sometimes the presence of medial integumentary wrinkling on actual segment 7 can provoke the illusion of a double segment, which is also a conceivable possibility. Having said that and considering the proper count, the ornamental and sensory elements reported are corroborated in this thesis, except for the number of setae in the actual segment 22, only illustrated and with unenforceable identification.

During our examinations, we identified other remarkable attributes for the species that represent variation for the morphology of the organisms examined by Kiefer (1954): (1) female genital double-somite with dorsal suture at mid-length; (2) female fifth swimming leg endopod 1-segmented; (3) female fifth swimming leg exopod 1 with outer spine. None of these characteristics were also mentioned for Colombian organisms, but it is possible to observe illustrations in this same collaboration that identify attributes 2 and 3, at least (Fuentes-Reinés *et al*., 2021, fig. 5B and 5C). Feature 3, perhaps, is the most intriguing, as no other *Notodiaptomus* exhibits this attribute. Contrary to this, this condition is similar to the species *Diaptomus* (s.l.) *corderoi* Wright 1936, recombined as *Scolodiaptomus corderoi* (Wright 1936) in Reid (1987). This species was redescribed from individuals from Minas Gerais (Reid, 1987) and also shares characteristic 1 that groups in addition to *N. maracaibensis*, *N. jatobensis*, and *N. isabelae*.

Another similarity of *N. maracaibensis* is suggested for *N. henseni* (Fuentes-Reinés *et al*., 2021). In this study two morphological indications in *N. maracaibensis* are presented as differentials: female right epimeral plate with slender sensilla on inner medial surface, ornamented with inner spinules patch dorsally; and male right antennule actual segment 13 without “spinous process” (possibly in the case of the spiniform modified seta, due to the mistake in the segment count). During our examinations, we corroborated the first characteristic, truly the positioning of the ornamentation is divergent between the species, for *N. henseni* the elements are positioned in the fifth metasomal segment dorsally, and for *N. maracaibensis* in the projection defined as epimeral plate. The second related feature is here refuted, regardless of the segment count, both species have the same ornamentation, either spinous process or spiniform modified seta. Irrefutably, a definitive attribute that distinguishes *N. maracaibensis* is male fifth right swimming leg basis without inner intumescence, and protuberance innerly. *N. nelsoni* also presents similarities with the species, from this, *N. maracaibensis* can be differentiated through of the male epimeral plates not ornamented. However, among the characteristics of the species that are divergent from those defined for *Notodiaptomus* (Kiefer, 1936; 1956) is only female fifth swimming leg endopod 1-segmented without discontinuity cuticular. Still, future efforts may reposition the species taxonomically.

#### Notodiaptomus nelsoni Previattelli, Perbiche-Neves & Rocha, 2017

##### Synonymy

*Notodiaptomus nelsoni* Previattelli *et al*., 2017: 11–30, figs. 1–10, tab. 1–3.

##### Type locality

Xingu River Basin (3°12m 54s S 52°11m 28s W), in front of Altamira, Para State, Brazil.

##### Type material

Holotype: 1 male, entire alcohol with glycerine (MZUSP 30604), from the Arapuja Lake, 3°12’S, 52°11’W, Xingu River Basin in front of Altamira, Para State, X.21.1997, Jansen Zuanon col.. Paratypes: 10 males, and 10 females, entire in alcohol with glycerine (MZUSP 30605); 1 male, and 1 female dissected and mounted on slides in glycerine (MZUSP 30606), Arapuja Lake, Xingu River, Altamira City, Para State, 21 October 1997, Jansen Zuanon col. All material was deposited in Zoology Museum São Paulo - MZUSP, Brazil.

##### Material examined

Holotype: 1 male (MZUSP 30604). Paratypes: 5 males, and 6 females (MZUSP 30604). Non-type material: 2 males, 2 females and 5 copepodites (not sexed) from the Ressaca Lake, Tocantins River Basin, 5°11m S, 49°15m W, VI.1983, Pedro Mera collector; 1 male (INPA-COP039, slides a-h) and 1 female (INPA-COP040, slides a-h) were selected to be dissection on eight slides each and deposited in the Zoological Collection of the INPA, Brazil.

##### Diagnosis

**(1)** Male first, third, fourth, and fifth metasome segments with dorsal sensilla; **(2)** fifth metasome segment ornamented with discontinuous single spinules row dorsally; **(3)** male right epimeral plate ornamented with spinules row; **(4)** left antennule actual segment 1 with spinules patch; **(5)** male fifth right swimming leg coxa with outer conical process projecting over basis on proximal surface posteriorly; **(6)** male fifth right swimming leg basis with posterior groove obliquely, ornamented with tubercles on outer border; **(7)** male fifth right swimming leg exopod 1 with lobular lamella on posterior surface; **(8)** female fourth and fifth metasome segments fused partially on lateral surface; **(9)** female limit between fourth and fifth metasome segments with double spinules row over limit entirely; **(10)** female right epimeral plate ornamented with inner spinules patch posteriorly; **(11)** female left epimeral plate with posterior semicircular expansion dorsally; **(12)** female genital double-somite with bifid apex on left sensilla anteriorly; **(13)** female right antennule with length not extending beyond caudal rami; **(14)** female fifth swimming legs intercoxal plate fused to legs; **(15)** female fifth swimming legs endopod with inner discontinuity on cuticle; **(16)** female fifth swimming legs basis with spinules patch posteriorly.

##### Redescription

###### MALE

Body 1075 micrometers excluding caudal setae. Male body smaller and slenderer than female. Nerve axons myelinated. Prosome 6-segmented; widest at first metasome segment; without one line of setules at posterior margin; with spinules at least at one segment. Cephalosome anterior margin sub-triangular; with dorsal suture; incomplete; separate from first metasome segment. First metasome segment with sensilla; 2 laterally; of equal size. Second metasome segment with sensilla; 2 dorsally; 2 laterally; of equal size. Third metasome segment with sensillae; 4 laterally; of equal size; ornamented posterior margin; with spinules; as a row; double; dorsally, or laterally. Fourth metasome segment with sensillae; 2 dorsally; 2 laterally; of equal size; fused to fifth metasome; fourth metasome segment partially; fourth metasome segment on lateral surface. Limit between fourth and fifth metasome segments ornamented; with spinules; as a row; on dorsal doubly; on lateral singly; same size. Fifth metasome segment with sensilla; 2 laterally; Fifth metasome segment equal size; Fifth metasome segment ornamented; with spinules; as a row; single; discontinuous; dorsally; dorsal conical process absent; with epimeral plates. Epimeral plates symmetrical. Right epimeral plates reduced, as rounded distal corner segment limit; with sensilla; at the apex of projection; ornamented; with spinules; as a row.

##### Urosome

5-segmented; Urosome 5 - free segments. Genital somite symmetrical in dorsal view; with single aperture; located on left side; ventrolaterally on posterior rim; with sensillae; on both sides; one; at left lateral; posteriorly; one; at right rim; posteriorly; of equal size between then. Third urosome segment without spinules; without external seta. Fourth urosome segment without spinules; without sub-conical blunt dorsal-lateral process. Anal segment presence of dorsal sensillae; one on each side; medially inserted; presence of operculum; convex; covering the anal aperture fully. Caudal rami symmetrical; separated from anal segment; longer than wide; with setules; continuous on; inner side; each ramus bearing 6 caudal setae; 5 marginals; plumose; and 1 internal dorsally; straight; not reticulated main axis; outermost seta with outer spiniform process absent.

##### Oral appendices feature

Rostrum asymmetrical; separated from dorsal cephalic shield; by complete suture; sensillae present; one pair; anteriorly inserted on surface tegument; with rostral filament; double; paired; extended; into point; with basal process; in ventral view, rounded on left side; without a smaller basal expansion on the right side.

##### Antennules

Asymmetrical. **Right antennules**. Uniramous; right antennule surpassing to genital segment; right antennule not extending beyond caudal rami.

Right antennule ancestral segment I and II separated. Ancestral segment II and III fused. Ancestral segment III and IV fused. Ancestral segment IV and V separated. Ancestral segment V and VI separated. Ancestral segment VI and VII separated. Ancestral segment VII and VIII separated. Ancestral segment VIII and IX separated. Ancestral segment IX and X separated. Ancestral segment X and XI separated. Ancestral segment XI and XII separated. Ancestral segment XII and XIII separated. Ancestral segment XIII and XIV separated. Ancestral segment XIV and XV separated. Ancestral segment XV and XVI separated. Ancestral segment XVI and XVII separated. Ancestral segment XVII and XVIII separated. Ancestral segment XVIII and XIX separated. Ancestral segment XIX and XX separated. Ancestral segment XX and XXI separated. Ancestral segment XXI and XXII fused. Ancestral segment XXII and XXIII fused. Ancestral segment XXIII and XXIV separated. Ancestral segment XXIV and XXV fused. Ancestral segment XXV and XXVI separated. Ancestral segment XXVI and XXVII separated. Ancestral segment XXVII and XXVIII fused.

Right antennule actual 22-segmented; geniculated; between the segment 18 and segment 19; with swollen and modified region; formed by 5 segments; between 13 and 17 segments. Actual segment 1 with seta; one element; straight; none larger than segment; without spinules; without vestigial seta; without conical seta; without modified seta; without spinous process; with aesthetasc; one element. Actual segment 2 with seta; three elements; of unequal size; straight; none larger than segment; without spinules; with vestigial seta; one element; without conical seta; without modified seta; without spinous process; with aesthetasc; one element. Actual segment 3 with seta; one element; one larger than segment; surpassing to distal margin; beyond three sequential segments; straight; blunt apex; without spinules; with vestigial seta; one element; without conical seta; without modified seta; without spinous process; with aesthetasc. Actual segment 4 with seta; one element; one larger than segment; surpassing to distal margin; straight; not beyond three sequential segments; without spinules; without vestigial seta; without conical seta; without modified seta; without spinous process; without aesthetasc. Actual segment 5 with seta; one element; straight; one larger than segment; surpassing to distal margin; not beyond three sequential segments; without spinules; with vestigial seta; one element; without conical seta; without modified seta; without spinous process; with aesthetasc; one element. Actual segment 6 with seta; one element; none larger than segment; straight; without spinules; without vestigial seta; without conical seta; without modified seta; without spinous process; without aesthetasc. Actual segment 7 with seta; one element; straight; one larger than segment; surpassing to distal margin; beyond three sequential segments; blunt apex; without spinules; without vestigial seta; without conical seta; without modified seta; without spinous process; with aesthetasc; one element. Actual segment 8 with seta; one element; straight; none larger than segment; without spinules; without vestigial seta; with conical seta; one element; reaching to middle-point of the sequent segment; without modified seta; without spinous process; without aesthetasc. Actual segment 9 with seta; two elements; of unequal size; straight; one larger than segment; surpassing to distal margin; beyond three sequential segments; blunt apex; without spinules; without vestigial seta; without conical seta; without modified seta; without spinous process; with aesthetasc; one element. Actual segment 10 with seta; one element; straight; none larger than segment; without spinules; without vestigial seta; without conical seta; with modified seta; presenting blunt apex; slender form; surpassing to distal margin; beyond of the sequential segment; parallel to antennule direction; without spinous process; without aesthetasc. Actual segment 11 with seta; one element; straight; one larger than segment; surpassing to distal margin; not beyond three sequential segments; without spinules; without vestigial seta; without conical seta; with modified seta; slender form; presenting blunt apex; surpassing to distal margin; beyond of the sequential segment; parallel to antennule direction; shorter length than homologous of actual segment 13; without spinous process; without aesthetasc. Actual segment 12 with seta; one element; straight; one larger than segment; surpassing to distal margin; not beyond three sequential segments; without spinules; without vestigial seta; with conical seta; one element; smaller than to segment 8; without modified seta; without spinous process; with aesthetasc; one element; absent internal perpendicular fission. Actual segment 13 with seta; one element; straight; one larger than segment; surpassing to distal margin; not beyond three sequential segments; without spinules; without vestigial seta; without conical seta; with modified seta; stout form; surpassing to distal margin; to the distal-point of the sequence segment; parallel to antennule direction; presenting bifid apex; without spinous process; with aesthetasc; one element. Actual segment 14 with seta; two elements; of unequal size; straight; one larger than segment; surpassing to distal margin; beyond three sequential segments; blunt apex; without spinules; without vestigial seta; without conical seta; without modified seta; without spinous process; with aesthetasc; one element. Actual segment 15 with seta; two elements; of unequal size; straight; not bifidform; none larger than segment; without spinules; without vestigial seta; without conical seta; without modified seta; with spinous process; on outer margin; surpassing distal margin; with aesthetasc; one element. Actual segment 16 with seta; two elements; of unequal size; plumose; one larger than segment; surpassing to distal margin; not beyond three sequential segments; not bifidform; without spinules; without vestigial seta; without conical seta; without modified seta; with spinous process; on outer margin; surpassing distal margin; unequal size to process on preceding segment; with aesthetasc; one element. Actual segment 17 with seta; two elements; of unequal size; straight; none larger than segment; bifidform; without spinules; without vestigial seta; without conical seta; with modified seta; one element; stout form; surpassing to distal margin; not beyond of the sequential segment; parallel to antennule direction; without spinous process; without aesthetasc. Actual segment 18 with seta; two elements; of equal size; straight; none larger than segment; without spinules; without vestigial seta; without conical seta; with modified seta; one element; stout form; surpassing distal margin; parallel to antennule direction; without spinous process; without aesthetasc. Actual segment 19 with seta; two elements; of unequal size; plumose; none larger than segment; without spinules; without vestigial seta; without conical seta; with modified seta; two elements; stout form; at least one bifid form; surpassing distal margin; parallel to antennule direction; without spinous process; with aesthetasc; one element. Actual segment 20 with seta; four elements; of unequal size; straight; one larger than segment; surpassing to distal margin; beyond three sequential segments; without spinules; without vestigial seta; without conical seta; without modified seta; without spinous process; without aesthetasc. Actual segment 21 with seta; two elements; of equal size; plumose; one larger than segment; surpassing to distal margin; greater 3x than original segment; without spinules; without vestigial seta; without conical seta; without modified seta; without spinous process; without aesthetasc. Actual segment 22 with seta; four elements; of equal size; one larger than segment; plumose; surpassing to distal margin; greater 3x than original segment; without spinules; without vestigial seta; without conical seta; without modified seta; without spinous process; with aesthetasc; one element.

##### Left antennules

Uniramous; Left antennule surpassing to prosome; Left antennule not extending beyond caudal rami. Ancestral segment I and II separated. Ancestral segment II and III fused. Ancestral segment III and IV fused. Ancestral segment IV and V separated. Ancestral segment V and VI separated. Ancestral segment VI and VII separated. Ancestral segment VII and VIII separated. Ancestral segment VIII and IX separated. Ancestral segment IX and X separated. Ancestral segment X and XI separated. Ancestral segment XI and XII separated. Ancestral segment XII and XIII separated. Ancestral segment XIII and XIV separated. Ancestral segment XIV and XV separated. Ancestral segment XV and XVI separated. Ancestral segment XVI and XVII separated. Ancestral segment XVII and XVIII separated. Ancestral segment XVIII and XIX separated. Ancestral segment XIX and XX separated. Ancestral segment XX and XXI separated. Ancestral segment XXI and XXII separated. Ancestral segment XXII and XXIII separated. Ancestral segment XXIII and XXIV separated. Ancestral segment XXIV and XXV separated. Ancestral segment XXV and XXVI separated. Ancestral segment XXVI and XXVII separated. Ancestral segment XXVII and XXVIII fused.

Left antennule actual 25-segmented; not-geniculated. Actual segment 1 with seta; one element; none larger than segment; straight; with spinules; as a patch; without vestigial seta; without conical seta; without modified seta; without spinous process; with aesthetasc; one element. Actual segment 2 with seta; three elements; of equal size; none larger than segment; straight; without spinules; with vestigial seta; one element; without conical seta; without modified seta; without spinous process; with aesthetasc; one element. Actual segment 3 with seta; one element; one larger than segment; straight; surpassing to distal margin; beyond three sequential segments; without spinules; with vestigial seta; one element; without conical seta; without modified seta; without spinous process; with aesthetasc. Actual segment 4 with seta; one element; none larger than segment; straight; without spinules; without vestigial seta; without conical seta; without modified seta; without spinous process; without aesthetasc. Actual segment 5 with seta; one element; one larger than segment; straight; surpassing to distal margin; not beyond three sequential segments; without spinules; with vestigial seta; one element; without conical seta; without modified seta; without spinous process; with aesthetasc; one element. Actual segment 6 with seta; one element; none larger than segment; straight; without spinules; without vestigial seta; without conical seta; without modified seta; without spinous process; without aesthetasc. Actual segment 7 with seta; one element; one larger than segment; straight; surpassing to distal margin; beyond three sequential segments; without spinules; without vestigial seta; without conical seta; without modified seta; without spinous process; with aesthetasc; one element. Actual segment 8 with seta; one element; one larger than segment; straight; surpassing distal margin; without spinules; without vestigial seta; with conical seta; without modified seta; without spinous process; without aesthetasc. Actual segment 9 with seta; two elements; of unequal size; one larger than segment; straight; surpassing to distal margin; beyond three sequential segments; without spinules; without vestigial seta; without conical seta; without modified seta; without spinous process; with aesthetasc; one element. Actual segment 10 with seta; one element; none larger than segment; straight; without spinules; without vestigial seta; without conical seta; without modified seta; without spinous process; without aesthetasc. Actual segment 11 with seta; one element; one larger than segment; straight; surpassing to distal margin; beyond three sequential segments; without spinules; without vestigial seta; without conical seta; without modified seta; without spinous process; without aesthetasc. Actual segment 12 with seta; one element; one larger than segment; straight; surpassing distal margin; without spinules; without vestigial seta; with conical seta; without modified seta; without spinous process; with aesthetasc; one element. Actual segment 13 with seta; one element; none elongated; straight; surpassing distal margin; without spinules; without vestigial seta; without conical seta; without modified seta; without spinous process; without aesthetasc. Actual segment 14 with seta; one element; elongated; straight; surpassing to distal margin; beyond three sequential segments; without spinules; without vestigial seta; without conical seta; without modified seta; without spinous process; with aesthetasc; one element. Actual segment 15 with seta; one element; larger than segment; straight; surpassing to distal margin; not beyond three sequential segments; without spinules; without vestigial seta; without conical seta; without modified seta; without spinous process; without aesthetasc. Actual segment 16 with seta; one element; larger than segment; plumose; surpassing to distal margin; not beyond three sequential segments; without spinules; without vestigial seta; without conical seta; without modified seta; without spinous process; with aesthetasc; one element. Actual segment 17 with seta; one element; not larger than segment; straight; without spinules; without vestigial seta; without conical seta; without modified seta; without spinous process; without aesthetasc. Actual segment 18 with seta; one element; larger than segment; straight; surpassing to distal margin; beyond three sequential segments; without spinules; without vestigial seta; without conical seta; without modified seta; without spinous process; without aesthetasc. Actual segment 19 with seta; one element; not larger than segment; straight; surpassing distal margin; without spinules; without vestigial seta; without conical seta; without modified seta; without spinous process; with aesthetasc; one element. Actual segment 20 with seta; one element; not larger than segment; straight; surpassing distal margin; without spinules; without vestigial seta; without conical seta; without modified seta; without spinous process; without aesthetasc. Actual segment 21 with seta; one element; larger than segment; plumose; surpassing to distal margin; beyond three sequential segments; without spinules; without vestigial seta; without conical seta; without modified seta; without spinous process; without aesthetasc. Actual segment 22 with seta; two elements; of unequal size; one of them elongated; plumose; surpassing to distal margin; without spinules; without vestigial seta; without conical seta; without modified seta; without spinous process; without aesthetasc. Actual segment 23 with seta; two elements; of unequal size; one larger than segment; plumose; surpassing to distal margin; greater 3x than original segment; without spinules; without vestigial seta; without conical seta; without modified seta; without spinous process; without aesthetasc. Actual segment 24 with seta; two elements; of equal size; one larger than segment; plumose; surpassing to distal margin; greater 3x than original segment; without spinules; without vestigial seta; without conical seta; without modified seta; without spinous process; without aesthetasc. Actual segment 25 with seta; four elements; of equal size; elongated; plumose; surpassing to distal margin; 4 times larger than segment; without spinules; without vestigial seta; without conical seta; without modified seta; without spinous process; with aesthetasc; one element.

##### Antenna

Biramous. Antenna coxa separated from the basis; bearing seta; 1; on inner surface; at distal corner; reaching to the endopod 1. Antenna basis (fusion) separated from the endopodal segment; bearing seta; 2; on inner surface; at distal corner. Endopodal ancestral segment I and II separated. Ancestral segment II and III fused. Ancestral segment III and IV fused. Ancestral segment III and IV fully. Antenna endopod actual 2-segmented. Actual segment 1 not bilobate; with seta; two; on inner margin; with spinules; as a row; obliquely; on outer surface; with pore. Actual segment 2 bilobate; without discontinuity on outer cuticle; inner lobe bearing 8 setae; distally; outer lobe bearing 7 setae; distally; with spinules; as a patch; on outer surface. Antenna exopod ancestral segment I and II separated. Ancestral segment II and III fused. Ancestral segment III and IV fused. Ancestral segment IV and V separated. Ancestral segment V and VI separated. Ancestral segment VI and VII separated. Ancestral segment VII and VIII separated. Ancestral segment VIII and IX separated. Ancestral segment IX and X fused. Antenna exopod actual 7-segmented. Actual segment 1 single; elongated (width-length, equal or larger ratio 2:1); with seta; one; at inner surface. Actual segment 2 compound; elongated (larger width-length ratio 2:1); with seta; three; at inner surface. Actual segment 3 single; not elongated (lesser width-length ratio 2:1); with seta; one; at inner surface. Actual segment 4 single; not elongated (lesser width-length ratio 2:1); with seta; one; at inner surface. Actual segment 5 single; not elongated (lesser width-length ratio 2:1); with seta; one; at inner surface. Actual segment 6 single; not elongated (lesser width-length ratio 2:1); with seta; one; at inner surface. Actual segment 7 compound; elongated (larger or equal width-length ratio 2:1); with seta; one; at inner surface; and three; at distal surface.

##### Oral features

**Mandible**. Coxal gnathobase sclerotized; without lobe; presence of cutting blade; with tooth-like prominence; two, distinctly; 1 acute; on caudal margin; and 1 triangular; on sub-caudal margin; without acute projection between the prominences; with additional spinules; as a row; on dorsal surface; with seta; 1; dorsally; on apical surface; without spinules. Mandible palps biramous; comprising the basis; with seta; four; differently inserted; first medially; not reaching to beyond the endopod 1; second distally; third distally; fourth distally; on inner margin; all with setulose ornamentation. Mandible endopod 2-segmented. Mandible endopod 1 with lobe; bearing seta; four; distally inserted; without spinules. Mandible endopod 2 without lobe; bearing setae; nine elements; distally inserted; with spinules; as a row; double. Mandible exopod 4-segmented. Mandible exopod 1 with seta; one element; distally; on inner margin. Mandible exopod 2 with seta; one element; distally; on inner side. Mandible exopod 3 with seta; one element; distally; on inner side. Mandible exopod 4 with setae; three elements; on terminal region. **Maxillule**. Birramous. Maxillule 3-segmented. Maxillule praecoxa with praecoxal arthrite; bearing spines; fifteen elements; ten marginally; plus, five sub-marginally; with spinules; as a patch; on sub-marginal surface. Maxillule coxa with coxal epipodite; with conspicuous outer lobe; bearing setae; nine elements; with coxal endite; elongated (larger or equal width-length ratio 2:1); bearing setae; four elements. Maxillule basis with basal endite; double; first proximal; elongated (larger width-length ratio 2:1; separated from basis; with setae; four elements; distally inserted; second distal; fused to basis; not elongated (lesser width-length ratio 2:1); with setae; four elements; distally inserted; with setules; as a row; on inner side; basal exite present; with setae; one element; on outer surface. Maxillule endopod 1-segmented. Endopod 1 bilobate; first proximal; with setae; three elements; second distal; with setae; five elements. Maxillule exopod 1-segmented. Exopod 1 with setae; six elements; with setules; as a row; on inner side; spinules absent. **Maxilla**. Uniramous. Maxilla 5-segmented. Maxilla praecoxa fused to coxa; incompletely; distinct externally; with praecoxal endite; double; first elongated endite (larger or equal width length ratio 2:1); proximally inserted; with seta; straight, or plumose; 1 straight; 4 plumose; with spine; single; without spinules; without setule; second elongated endite (larger or equal width length ratio 2:1); distally inserted; with seta; plumose; 3 plumose; without spine; with spinules; as a row; on distal margin; with setule; as a row; on distal margin; absence of outer seta. Maxilla coxa with coxal endite; double; first elongated endite (larger or equal width); proximally inserted; with seta; plumose; 3 plumose; without spine; without spinules; with setules; as a row; on proximal margin; second elongated endite (larger or equal width); distally inserted; with seta; plumose; 3 plumose; without spine; without spinules; with setules; as a row; on proximal margin; absence of outer seta. Maxilla basis with basal endite; single; elongated (larger or equal width-length ratio 2:1); with seta; plumose; 3 plumose; without spinules; absence of outer seta. Maxilla endopod 2-segmented. Endopod 1 with seta; 2 plumose; without spine; without spinules; without setules. Maxilla endopod 2 with seta; 2 plumose; without spine; without spinules; without setules. **Maxilliped**. Uniramous; Maxilliped 8-segmented. Maxilliped praecoxa fused to coxa; incompletely; distinct internally; with praecoxal endite; not elongated (lesser width-length ratio 2:1); distally inserted; with seta; 1 straight; with spinules; as a row; single; on basal surface; without setules. Maxilliped coxa with coxal endite; three coxal endite; first elongated (larger or equal width); proximally inserted; with seta; 2 plumose; with spinules; as a patch; single; on apical surface; without setules; second not elongated (lesser width-length ratio 2:1); medially inserted; with seta; 3 plumose; with spinules; as a row; single; on medial surface; without setules; third elongated (larger or equal width length ratio 2:1); distally inserted; with seta; 3 plumose; none reaching to beyond of the basis; with spinules; as a row; single; on basal surface; without setules; with lobe; prominence; at inner distal angle; ornamented; with spinules; continuously on margin. Maxilliped basis without basal endite; with seta; 3 plumose; with spinules; as a row; single; on medial surface; with setules; as a row; single; on inner margin. Maxilliped endopod segment 6-segmented. Endopod 1 with seta; 2 plumose; on inner surface. Endopod 2 with seta; 3 plumose; on inner surface. Endopod 3 with seta; 2 plumose; on inner surface. Endopod 4 with seta; 2 plumose; on inner surface. Endopod 5 with seta; 2 plumose; on inner surface, or on outer surface; outer seta absent. Endopod 6 with seta; 4 plumose; on inner surface, or on outer surface.

##### Swimming legs features

**First swimming legs.** Symmetrical; biramous. First swimming legs intercoxal plate without seta. First swimming legs praecoxa absent. First swimming legs coxa with seta; one; plumose; distally inserted; on inner surface; surpassing to basal segment; with setules; two group; as a patch; on inner margin; and as a row; double; on anterior surface; outerly; with spinules; as a row; single; medially inserted; at anterior surface; without spine. First swimming legs basis without seta; with setules; as a row; single; discontinuously; on outer surface; without spinules; without spine. First swimming legs endopod 2-segmented. Endopod 1 with seta; plumose; restricted; to inner surface; one element; without spine; with setules; as a row; single; continuously; on outer surface; without spinules; absence of Schmeil’s organ. Endopod 2 with seta; unrestricted; three on inner surface; one on outer surface; two on distal surface; plumose; without spine; with setules; as a row; single; continuously; on outer surface; without spinules; absence of Schmeil’s organ. Endopod 3 absence. First swimming legs exopod 1 with seta; restricted; 1 on inner surface; with spine; 1; stout; smaller than original segment; serrated; on inner side; continuously; with setules; as a row; single; as a row; innerly. First swimming legs exopod 2 with seta; restricted; 1 on inner surface; plumose; without spine; with setules; as a row; single; continuously; on inner margin, or on outer margin; without spinules. First swimming legs exopod 3 with setule; as a row; single; continuously; on outer surface; without spinules; with seta; unrestricted; 2 on inner surface; 2 on terminal surface; with spine; 2; unequal size; first no longer 2x than origin segment; stout; serrated; on inner side, or on outer side; equally; second longer 3x than origin segment; slender; serrated; on outer side; with ornamentation on non-serrated side; by setules. **Second swimming legs**. Symmetrical; Second swimming legs biramous. Second swimming legs intercoxal plate without seta. Second swimming legs praecoxa present; located laterally. Second swimming legs coxa with seta; plumose; distally inserted; on inner surface; surpassing to basal segment; without setules; without spinules; without spine. Second swimming legs basis without seta; without setules; without spinules; without spine. Second swimming legs endopod 3-segmented. Endopod 1 with seta; plumose; restricted; one on inner surface; without spine; with setules; as a row; single; continuously; on outer surface; without spinules; absence of Schmeil’s organ. Endopod 2 with seta; plumose; unrestricted; two on inner surface; without spine; with setules; as a row; single; continuously; on outer side; without spinules; presence of Schmeil’s organ; on posterior surface. Endopod 3 with seta; plumose; unrestricted; three on inner surface; two on outer surface; two on distal surface; without spine; without setules; with spinules; as a row; double; distally inserted; at anterior surface; absence of Schmeil’s organ. Second swimming legs exopod 1 with seta; restricted; one on inner surface; with spine; 1; stout; not reaching to distal-third of the exopod 2; serrated; on inner side, or on outer side; with setules; as a row; single; continuously; on inner side; without spinules; absence of Schmeil’s organ. Exopod 2 with seta; unrestricted; one on inner surface; with spine; 1; stout; not surpassing the exopod 3; serrated; on inner side, or on outer side; with setules; as a row; single; continuously; on inner surface; without spinules; absence of Schmeil’s organ. Exopod 3 with seta; plurimarginal; three on inner surface; two on terminal surface; with spine; 2; unequal size; first no longer 2x than origin segment; stout; serrated; on inner side, or on outer side; equally; second longer 2x than origin segment; slender; serrated; on outer side; with ornamentation on non-serrated side; of setules; setules on outer surface; as a row; single; continuously; on inner surface; with spinules; as a row; single; distally inserted; at anterior surface; absence of Schmeil’s organ. **Third swimming legs**. Symmetrical; Third swimming legs biramous. Third swimming legs intercoxal plate without seta. Third swimming legs praecoxa present; not laterally located. Third swimming legs coxa with seta; plumose; distally inserted; on inner surface; surpassing to basal segment; without setules; without spinules; without spine. Third swimming legs basis without seta; without setules; without spinules; without spine. Third swimming legs endopod 3-segmented. Endopod 1 with seta; restricted; one on inner surface; without spine; without setules; without spinules; absence of Schmeil’s organ. Endopod 2 with seta; restricted; two on inner surface; plumose; without spine; without setules; without spinules; absence of Schmeil’s organ. Endopod 3 with seta; plumose; plurimarginal; two on inner surface; two on outer surface; three on terminal surface; without spine; without setules; with spinules; as a row; distally inserted; double; at anterior surface; absence of Schmeil’s organ. Third swimming legs exopod 1 with seta; restricted; plumose; one on inner surface; with spine; 1; stout; not reaching to the distal-third of the exopod 2; serrated; equally; on inner surface, or on outer surface; with setules; as a row; single; continuously; on inner surface; without spinules; absence of Schmeil’s organ. Exopod 2 with seta; plumose; restricted; one on inner surface; with spine; 1; stout; not reaching out to exopod 3; serrated; on inner side, or on outer side; equally; with setules; as a row; single; continuously; on inner side; without spinules; absence of Schmeil’s organ. Exopod 3 without setules; with spinules; as a row; single; distally inserted; at anterior surface; with seta; plumose; unrestricted; three on inner surface; two on terminal surface; with spine; 2; unequal size; first no longer 2x than origin segment; stout; serrated; on inner side, or on outer side; equally; second longer 2x than origin segment; slender; serrated; on outer side; with ornamentation on non-serrated side; of setules; absence of Schmeil’s organ. **Fourth swimming legs**. Symmetrical; biramous. Intercoxal plate without sensilla. Praecoxa present. Coxa with seta; distally inserted; on inner margin; reaching out to endopod 1; without spinules; setules absent. Basis with seta; one; medially inserted; on posterior surface; smaller than the original segment; without setules; without spinules; without spine. Fourth swimming legs endopod 3-segmented. Endopod 1 with seta; one; restricted; on inner surface; without spine; without setules; without spinules; absence of Schmeil’s organ. Endopod 2 with seta; restricted; two on inner side; without spine; with setules; as a row; single; continuously; on outer surface; without spinules; absence of Schmeil’s organ. Endopod 3 with seta; unrestricted; two on inner surface; two on outer surface; three on distal surface; without spine; without setules; with spinules; as a row; double; distally inserted; at anterior surface; absence of Schmeil’s organ. Fourth swimming legs exopod 1 with seta; restricted; one on inner surface; with spine; 1; stout; not reaching out to distal-third of the exopod 2; serrated; on inner side, or on outer side; equally; with setules; as a row; single; continuously; on inner surface; without spinules; absence of Schmeil’s organ. Exopod 2 with seta; restricted; one on inner surface; with spine; 1; stout; not reaching the end of exopod 3; serrated; on inner side, or on outer side; equally; with setules; as a row; single; continuously; on inner surface; without spinules; absence of Schmeil’s organ. Exopod 3 without setules; with spinules; as a row; single; distally inserted; at anterior surface; with seta; unrestricted; three on inner surface; two on distal surface; with spine; 2; unequal size; first no longer 2x than origin segment; stout; serrated; on inner side, or on outer side; equally; second longer 2x than origin segment; slender; serrated; on outer side; without ornamentation on non-serrated side; absence of Schmeil’s organ.

##### Fifth swimming legs features

Asymmetrical. Fifth swimming leg intercoxal plate with length not equal or greater than width on 1.5x; with irregular proximal margin; discontinuous to; the anterior margin of the left coxa, or the anterior margin of the right coxa; posterior sensilla on the right lateral absent. **Fifth left swimming leg**. Fifth left swimming leg biramous; leg reaching first right exopod segment; proximally. Fifth left swimming leg praecoxa present; rudimentary; separated from the coxae; without ornamentation. Fifth left swimming leg coxa concave inner side; without teeth-like structures; with process; conical; on posterior surface; outer side; distally inserted; projecting over basis; reaching to the proximal surface; with sensilla; stout; triangular; at apex; no longer 2x than insertion basis; without swelling; without seta; without spinules. Fifth left swimming leg basis sub-cylindrical; unequal size between inner and outer side; shorter outer than inner side; with concave inner side; rounded internal proximal expansion absent; without outgrowth; with groove; deep; obliquely; on posterior surface; not reaching the endopodal lobe; not ornamented; absence of protuberance; with seta; outerly inserted; no longer 2x than origin segment; absence of minutely granular. Fifth left swimming leg endopod segments 1 and 2 fused; segments 2 and 3 fused; 1-segmented; stout; separated from the basis; ornamented; on inner side; with spinules; more than four elements; as a row; terminally; row of setules absent; without seta. Fifth left swimming leg exopod segments 1 and 2 separated; segments 2 and 3 fused; 2-segmented; stout; separated from the basis. Fifth left swimming leg exopod 1 sub-cylindrical; longer than broad; unequal size between inner and outer side; shorter inner than outer side; concave inner side; convex outer side; without swelling; without marginal extension; without process; with lobe; double; semicircular; medially inserted; on inner side; covered; by setules; without outer spine; absence seta. Fifth left swimming leg exopod 2 digitiform; longer than broad; equal sizes between inner and outer side; disform inner side; with rectilinear outer side; setulose pad absent; inflated medial region absent; distal process present; digitiform; non denticulate; without transverse row of denticles; none oblique row of 5 denticles; innerly directed; with seta; spiniform; not ornamented by spinules; not surpassing the distal-point of the segment; without outer spine; terminal claw absent.

##### Fifth right swimming leg

Biramous. Fifth right swimming leg praecoxa present; separated from the coxae; without ornamentation. Fifth right swimming leg coxa convex inner side; without teeth-like structures; with process; rounded; distally inserted; on posterior surface; closest to the outer rim; projecting over basis; beyond the first third; until the medial surface; without triangular protuberance innerly; with sensilla; slender; at apex; no longer 2x than basal insertion; without marginal extension; without seta; without spinules. Fifth right swimming leg basis cylindrical; unequal size between inner and outer side; shorter outer than inner side; rectilinear inner side; tumescence present; not inflated; restricted on inner surface; proximally; without protuberance; absence of distinct minutely granular; additional inner process absent; with posterior groove; deep; obliquely; not reaching the endopodal lobe; ornamented; with tubercles; throughout of the outer border; with seta; outerly inserted; on anterior surface; no longer 2x than origin segment; posterior protrusion present; distal tegument expansion present; triangular; anteriorly. Fifth right swimming leg with endopodite present; separated from the basis, on anterior surface; ancestral segments 1 and 2 fused; ancestral segments 2 and 3 fused; 1-segmented; stout; ornamented; with setules; as a row; on inner side; terminally; without seta. Fifth right swimming leg exopod segments 1 and 2 separated; segments 2 and 3 fused; 2-segmented; stout; separated from the basis. Fifth right swimming leg exopod 1 trapezium; longer than broad; nearly 1.25 times; unequal size between both sides; shorter inner than outer side; convex inner side; convex outer side; with marginal extension; sub-triangular; distally inserted; at outer rim; spinules absent; with process; rounded; sclerotized; without ornamentation; distally inserted; at posterior surface; projecting over next segment; without outer spine; without seta; internal prominence absent; lamella on posterior surface present; lobular form; not surpassing to margin. Fifth right swimming leg exopod 2 cylindrical; longer than broad; nearly 2 times; equal size between both sides; disform inner side; convex outer side; without posterior proximal swelling; inner-posterior process absent; without marginal expansion; curved ridge on distal posterior surface absent; chitinous knobs absent; with outer spine; inserted sub-distally; arched; internally directed; not ornamented innerly; not ornamented outerly; sharp tip; without apparent curve; lesser than the length of the exopod 2; until to 2 times its size; 1.5x; sensilla absent; terminal claw present; not equal or longer 1.5 times than insertion segment; sclerotized; arched; inward; with conspicuous curve; proximally; not ornamented innerly; ornamented outerly; sharp tip; not curved tip; without medial constriction; hyaline process absent.

##### FEMALE

Body longer and wider than male; Female body 1670 micrometers excluding caudal setae. Widest at first metasome segment. Distal margin of the prosomal segments without one line of setules at posterior margin. Prosome segments with spinules at least at one prosomal segment. Fourth metasome segment absence of dorsal protuberance. Fourth and fifth metasome segments fused; partially; on lateral surface. Limit between fourth and fifth metasome segments ornamented; with spinules; as a row; double; complete; same size; entirely over limit (lateral, dorsal). **Fifth metasome segment**. Fifth metasome segment with sensilla; dorsally; 2 elements; with epimeral plates. Epimeral plates asymmetrical. Right epimeral plates prominent, as projections; not thinner than the left; one posterior-laterally directed; not reaching half length of the genital segment; with sensilla at the apex; dorsal-posterior sensilla present; slender; ornamented; with spinules; as a patch; innerly; on dorsal surface. Left epimeral plate with expansion; semicircular; on posterior surface; dorsally; with sensilla; at tip.

##### Urosome

3-segmented. **Genital double-somite**. Asymmetrical in dorsal view; longer than broad; longer than other urosomites combined; dorsal suture at mid-length absent; not covered by spinules; with swelling; rounded; equal size; anteriorly; with sensillae; on both sides; one; stout; with bifid apex; at left lateral; not on lobular base; anteriorly; one; stout; at right lateral; not on lobular base; anteriorly; with robust apex; of equal size between then; lateral protuberance absent; with right posterior rim expanded; over next segment; without slender sensilla on each posterior rim; without posterior-dorsal process. Genital double-somite opercular pad present; broader than longer; symmetrical; development laterally; expanded posteriorly; covering partially; double gonoporal slit; located ventrally; with arthrodial membrane; inserted anteriorly; post-genital process absent; disto-ventral tumescence absent; ventral vertical folds absent; dorsal sensilla absent. Second urosome segment without ventral fusion to anal segment; right distal process absent. Caudal rami patch of setules on outer surface absent; patch of spinules on outer surface absent.

##### Oral appendices feature

Rostrum basal process absent. **Antennules**. Symmetrical. Right antennule surpassing to genital double-segment; not extending beyond caudal rami; ornamentation pattern equals to male left antennule; mostly. Actual segment 13 without seta; without aesthetasc. Actual segment 14 without seta; without aesthetasc. Actual segment 15 without seta; without aesthetasc. Actual segment 16 without seta; without aesthetasc. Actual segment 17 without seta. Actual segment 18 without seta.

##### Fifth swimming legs

Symmetrical; Fifth swimming legs biramous. Fifth swimming legs intercoxal plate longer than wide; fused to legs. Fifth swimming legs praecoxa with sclerite praecoxal; separated from the coxae; without ornamentation. Fifth swimming legs coxa with process; conical; at the outer rim; distally; sensilla present; stout; at apex; projecting over basal segment; no longer 2x than basal insertion; marginal extension absent; without swelling; without seta; without spinules. Fifth swimming legs basis sub-triangular; unequal size between inner and outer sides; shorter outer than inner side; with convex inner side; without proximal inner outgrowth; without groove; with distal extension; on posterior surface; with seta; outerly inserted; on anterior surface; no longer 2x than origin segment; presence of spinules patch posteriorly. Fifth swimming legs endopod segments 1 and 2 fused; segments 2 and 3 fused; 1-segmented; stout; separated from the basis; present discontinuity cuticle; on inner side; with spinules; as a row; single; non-oblique; sub-terminally; at anterior surface; with seta; double; one medially; on posterior surface; rectilinear; one distally; on posterior surface; arched; of unequal size; distal seta longer than medial seta. Fifth swimming legs exopod segments 1 and 2 separated; segments 2 and 3 separated; 3-segmented; separated from the basis. Fifth swimming legs exopod 1 sub-cylindrical; longer than wide; longer or equal than 2 times; with unequal size between inner and outer side; shorter inner than outer side; with convex inner side; with rectilinear outer side; without swelling; without marginal extension; without posterior process; without spine; without seta. Fifth swimming legs exopod 2 sub-cylindrical; longer than broad; longer or equal than 2 times; without swelling; without marginal extension; without process; without lobe; with spine; inserted laterally; rectilinear; without ornamentation; sharp tip; equal size or larger than next segment; without seta. Fifth swimming legs exopod 3 cylindrical; longer than wide; without swelling; without process; without lobe; without spine; with seta; double; inserted terminally; unequal size between them; outer seta smaller than inner; nearly 3 times; outer seta not ornamented by setules; without ornamentation; presence of terminal claw; sclerotized; arched; externally directed; convex inner side; with ornamentation; of denticles; as a row; on surface partially; at medial region; concave outer side; with ornamentation; of denticles; as a row; on surface partially; at medial region; blunt tip; 6 times longer than origin segment.

##### Distribution records

###### BRAZIL

**Pará**: Arapujá Lake, 3°12’54“S, 52°11’28“W, Xingu River Basin, in front of Altamira; Arapujá Lake, Xingu River, Altamira City (Previattelli *et al*., 2017).

##### Habitat

Habitat in freshwaters: lakes, and rivers.

##### Remarks

The taxon was described from organisms of the Xingu River Basin in Northern Brazil and represents the most recent inclusion in *Notodiaptomus*. The authors defined the species from the following morphological set, mainly: (1) male first, third, fourth, and fifth metasome segments with dorsal spinules dorsal doubly; (2) male fifth right swimming leg coxa with outer conical process projecting over basis on proximal surface posteriorly; (3) female fifth swimming legs with exopod 1 with lateral seta reaching to beyond middle-point of segment; (4) male right antennule actual segments 2 and 3 without vestigial seta; (5) female limit between fourth and fifth metasome segments with double spinules row over limit entirely; (6) female right epimeral plate ornamented with inner spinules patch posteriorly; (7) female fifth swimming legs endopod with inner discontinuity on cuticle; (8) male fifth right swimming leg exopod 2 with outer spine 2/3 size of original segment; (9) male fifth right swimming leg basis without inner protuberance; and (10) female fifth swimming legs basis with spinules patch posteriorly.

From these characteristics, the authors were able to differentiate the species from relatable congeners, such as *N. paraensis* through conditions 1 and 2, *N. deitersi* through conditions 1, 4, and 8, *N. henseni* through condition 8, and “shape of lateral projections of genital segment of female, *N. amazonicus* through attributes 1, and 9, and *N. nordestinus* through characteristics 10, and 1. All these characteristics could be corroborated, except for attribute 4. In the original presentation of the species, there is a discrepancy between the description and illustrations for the male right antennule, sometimes with vestigial seta on actual segments 2 and 3, sometimes without. Through accessing the holotype informed for the MZUSP it was possible to confirm the existence of the elements for the mentioned segments of the male right antennule and, therefore, this does not represent differential characteristics of *N. deitersi*.

In the present effort, other evidence were identified for the species as differentiation of the type species of the genus: (1) male right antennule actual segment 8 with conical seta reaching to middle-point sequential segment; (2) male fifth right swimming leg basis without inner protuberance; (3) male fifth right swimming leg basis with posterior groove not reaching to endopodal lobe; (4) male fifth right swimming leg exopod 1 with distal process in rounded form; and (5) female fifth swimming legs basis with outer seta no longer 2x than origin segment. Among the attributes divergent from those considered by Kiefer for *Notodiaptomus* (1936; 1956) is only male fifth right swimming leg exopod 2 without curved ridge on distal posterior surface innerly.

#### Notodiaptomus nordestinus (Wright, 1935)

##### Synonymy

*Diaptomus nordestinus* Wright, 1935: 213, 214–221, 222, 224, 225, 226, 228, pl.1, figs. 1, 6–8, 10–14, pl. 2, figs. 1, 2, 4; 1936a: 80; 1937: 73, 76; 1938a: 300, 306; 1938b: 562; Brehm, 1960: 50; Reid, 1991: 738, 740. *Notodiaptomus nordestinus*; Kiefer, 1936a: 197, fig. 5; 1956: 242; Löffler, 1963: 208; 1981: 15; Brandorff, 1972: 45; 1976: 616, 621, fig. 2; Dussart, 1979: 6; 1984a: 46, 48, fig. 5B; Dussart & Defaye, 1983: 137; Dussart & Frutos, 1986: 246; Cicchino *et al*., 1989: 101; Reid, 1991: 738, 740; Rocha *et al*., 1995: 156; Santos-Silva, 1998; 211; Santos-Silva *et al*., 1999: 127; Santos-Silva, 2008: 32–33, fig. 6; Santos-Silva *et al*., 2015: 50–53, figs. 27–29, 40, identification keys to male and female; Perbiche-Neves *et al*., 2020: 697-698, key to the Neotropical diaptomid, fig. 21.15 K. *Notodiaptomus (Notodiaptomus) nordestinus*; Dussart, 1985a: 208.

##### Type locality

Pond Açude Simão, Campina Grande State, Paraíba City, Brazil.

##### Type material

Holotype not specified. Santos-Silva *et al*. (2015) specified 1 male as the neotype from the type-locality and in material located at the Smithsonian Institution, National Museum of Natural History, determined by S. Wright and labelled as the type locality (USNM 79543).

##### Material examined

Topotype: 2 males, and 3 females, entire in alcohol collected on 16.I.1935, by Lenz, and stored in Plankton Laboratory (n° L-831 in sample 2). Non-type material: 2 males, and 2 females from the Wright s collection (4936-CN), probably collected by S. Wright on XI.1935. This material is stored in Lab Plankton under code LP19-A35. 1 male (INPA-COP041, slides a-h) and 1 female (INPA-COP042, slides a-h) were selected to be dissection on eight slides each and deposited in the Zoological Collection of the INPA, Brazil.

##### Diagnosis

**(1)** Male fifth metasome segment without ornamentation dorsally; **(2)** male right and left antennule with length extending beyond caudal rami; **(3)** male fifth left swimming leg exopod 1 with medial double semicircular lobe innerly; **(4)** male fifth left swimming leg exopod 2 without ornamentation by spinules on spiniform seta; **(5)** male fifth right swimming leg basis with inner distinct minutely granular as a patch proximally; **(6)** male fifth right swimming leg basis with posterior groove reaching the endopodal lobe; **(7)** male fifth right swimming leg exopod 2 with curved tip outwards on terminal claw; **(8)** female fourth and fifth metasome segment fused partially on dorsal surface, without ornamentation; **(9)** female right epimeral plate not thinner than the left plate; **(10)** female genital double-somite with wrinkled lateral protuberance on the right side.

##### Redescription

###### MALE

Body 998 micrometers excluding caudal setae. Male body smaller and slenderer than female. Nerve axons myelinated. Prosome 6-segmented; widest at first metasome segment; without one line of setules at posterior margin; without spinules at segments. Cephalosome anterior margin rounded; with dorsal suture; incomplete; separate from first metasome segment. First metasome segment without sensilla. Second metasome segment without sensilla. Third metasome segment without sensillae; non-ornamented posterior margin. Fourth metasome segment without sensillae; separated from the fifth metasome. Limit between fourth and fifth metasome segments without ornamentation. Fifth metasome segment with sensilla; 2 dorsally; Fifth metasome segment equal size; Fifth metasome segment without ornamentation; Fifth metasome segment without dorsal conical process; with epimeral plates. Epimeral plates symmetrical. Right epimeral plates reduced, as rounded distal corner segment limit; with sensilla; at the apex of projection; without ornamentation.

##### Urosome

5-segmented; Urosome 5 - free segments. Genital somite symmetrical in dorsal view; with single aperture; located on left side; ventrolaterally on posterior rim; with sensillae; on both sides; one; at left lateral; posteriorly; one; at right rim; posteriorly; of equal size between then. Third urosome segment without spinules; without external seta. Fourth urosome segment without spinules; without sub-conical blunt dorsal-lateral process. Anal segment absence of dorsal sensillae; presence of operculum; convex; not covering the anal aperture fully. Caudal rami symmetrical; separated from anal segment; longer than wide; with setules; continuous on; inner side; each ramus bearing 6 caudal setae; 5 marginals; plumose; and 1 internal dorsally; straight; not reticulated main axis; outermost seta with outer spiniform process absent.

##### Oral appendices feature

Rostrum symmetrical; separated from dorsal cephalic shield; by complete suture; sensillae present; one pair; anteriorly inserted on surface tegument; with rostral filament; double; paired; extended; into point; with basal process; in ventral view, rounded on left side; without a smaller basal expansion on the right side.

##### Antennules

Asymmetrical. **Right antennules**. Uniramous; right antennule surpassing to genital segment; right antennule extending beyond caudal rami.

Right antennule ancestral segment I and II separated. Ancestral segment II and III fused. Ancestral segment III and IV fused. Ancestral segment IV and V separated. Ancestral segment V and VI separated. Ancestral segment VI and VII separated. Ancestral segment VII and VIII separated. Ancestral segment VIII and IX separated. Ancestral segment IX and X separated. Ancestral segment X and XI separated. Ancestral segment XI and XII separated. Ancestral segment XII and XIII separated. Ancestral segment XIII and XIV separated. Ancestral segment XIV and XV separated. Ancestral segment XV and XVI separated. Ancestral segment XVI and XVII separated. Ancestral segment XVII and XVIII separated. Ancestral segment XVIII and XIX separated. Ancestral segment XIX and XX separated. Ancestral segment XX and XXI separated. Ancestral segment XXI and XXII fused. Ancestral segment XXII and XXIII fused. Ancestral segment XXIII and XXIV separated. Ancestral segment XXIV and XXV fused. Ancestral segment XXV and XXVI separated. Ancestral segment XXVI and XXVII separated. Ancestral segment XXVII and XXVIII fused.

Right antennule actual 22-segmented; geniculated; between the segment 18 and segment 19; with swollen and modified region; formed by 5 segments; between 13 and 17 segments. Actual segment 1 with seta; one element; straight; none larger than segment; without spinules; without vestigial seta; without conical seta; without modified seta; without spinous process; with aesthetasc; one element. Actual segment 2 with seta; three elements; of unequal size; straight; none larger than segment; without spinules; with vestigial seta; one element; without conical seta; without modified seta; without spinous process; with aesthetasc; one element. Actual segment 3 with seta; one element; one larger than segment; surpassing to distal margin; beyond three sequential segments; straight; blunt apex; without spinules; with vestigial seta; one element; without conical seta; without modified seta; without spinous process; with aesthetasc. Actual segment 4 with seta; one element; one larger than segment; surpassing to distal margin; straight; not beyond three sequential segments; without spinules; without vestigial seta; without conical seta; without modified seta; without spinous process; without aesthetasc. Actual segment 5 with seta; one element; straight; one larger than segment; surpassing to distal margin; not beyond three sequential segments; without spinules; with vestigial seta; one element; without conical seta; without modified seta; without spinous process; with aesthetasc; one element. Actual segment 6 with seta; one element; none larger than segment; straight; without spinules; without vestigial seta; without conical seta; without modified seta; without spinous process; without aesthetasc. Actual segment 7 with seta; one element; straight; one larger than segment; surpassing to distal margin; beyond three sequential segments; blunt apex; without spinules; without vestigial seta; without conical seta; without modified seta; without spinous process; with aesthetasc; one element. Actual segment 8 with seta; one element; straight; none larger than segment; without spinules; without vestigial seta; with conical seta; one element; not reaching to middle-point of the sequent segment; without modified seta; without spinous process; without aesthetasc. Actual segment 9 with seta; two elements; of unequal size; straight; one larger than segment; surpassing to distal margin; beyond three sequential segments; blunt apex; without spinules; without vestigial seta; without conical seta; without modified seta; without spinous process; with aesthetasc; one element. Actual segment 10 with seta; one element; straight; none larger than segment; without spinules; without vestigial seta; without conical seta; with modified seta; presenting blunt apex; slender form; surpassing to distal margin; beyond of the sequential segment; parallel to antennule direction; without spinous process; without aesthetasc. Actual segment 11 with seta; one element; straight; one larger than segment; surpassing to distal margin; not beyond three sequential segments; without spinules; without vestigial seta; without conical seta; with modified seta; slender form; presenting blunt apex; surpassing to distal margin; beyond of the sequential segment; parallel to antennule direction; shorter length than homologous of actual segment 13; without spinous process; without aesthetasc. Actual segment 12 with seta; one element; straight; one larger than segment; surpassing to distal margin; not beyond three sequential segments; without spinules; without vestigial seta; with conical seta; one element; not smaller than to segment 8; without modified seta; without spinous process; with aesthetasc; one element; absent internal perpendicular fission. Actual segment 13 with seta; one element; straight; one larger than segment; surpassing to distal margin; not beyond three sequential segments; without spinules; without vestigial seta; without conical seta; with modified seta; stout form; surpassing to distal margin; to the middle-point of the sequence segment; parallel to antennule direction; presenting bifid apex; without spinous process; with aesthetasc; one element. Actual segment 14 with seta; two elements; of unequal size; straight; one larger than segment; surpassing to distal margin; beyond three sequential segments; blunt apex; without spinules; without vestigial seta; without conical seta; without modified seta; without spinous process; with aesthetasc; one element. Actual segment 15 with seta; two elements; of unequal size; straight; not bifidform; none larger than segment; without spinules; without vestigial seta; without conical seta; without modified seta; with spinous process; on outer margin; surpassing distal margin; with aesthetasc; one element. Actual segment 16 with seta; two elements; of unequal size; plumose; one larger than segment; surpassing to distal margin; not beyond three sequential segments; not bifidform; without spinules; without vestigial seta; without conical seta; without modified seta; with spinous process; on outer margin; surpassing distal margin; unequal size to process on preceding segment; with aesthetasc; one element. Actual segment 17 with seta; two elements; of unequal size; straight; none larger than segment; bifidform; without spinules; without vestigial seta; without conical seta; with modified seta; one element; stout form; surpassing to distal margin; not beyond of the sequential segment; parallel to antennule direction; without spinous process; without aesthetasc. Actual segment 18 with seta; two elements; of equal size; straight; none larger than segment; without spinules; without vestigial seta; without conical seta; with modified seta; one element; stout form; surpassing distal margin; parallel to antennule direction; without spinous process; without aesthetasc. Actual segment 19 with seta; two elements; of unequal size; plumose; none larger than segment; without spinules; without vestigial seta; without conical seta; with modified seta; two elements; stout form; at least one bifid form; surpassing distal margin; parallel to antennule direction; without spinous process; with aesthetasc; one element. Actual segment 20 with seta; four elements; of unequal size; straight; one larger than segment; surpassing to distal margin; beyond three sequential segments; without spinules; without vestigial seta; without conical seta; without modified seta; without spinous process; without aesthetasc. Actual segment 21 with seta; two elements; of equal size; plumose; one larger than segment; surpassing to distal margin; greater 3x than original segment; without spinules; without vestigial seta; without conical seta; without modified seta; without spinous process; without aesthetasc. Actual segment 22 with seta; four elements; of equal size; one larger than segment; plumose; surpassing to distal margin; greater 3x than original segment; without spinules; without vestigial seta; without conical seta; without modified seta; without spinous process; with aesthetasc; one element.

##### Left antennules

Uniramous; Left antennule surpassing to prosome; Left antennule extending beyond caudal rami. Ancestral segment I and II separated. Ancestral segment II and III fused. Ancestral segment III and IV fused. Ancestral segment IV and V separated. Ancestral segment V and VI separated. Ancestral segment VI and VII separated. Ancestral segment VII and VIII separated. Ancestral segment VIII and IX separated. Ancestral segment IX and X separated. Ancestral segment X and XI separated. Ancestral segment XI and XII separated. Ancestral segment XII and XIII separated. Ancestral segment XIII and XIV separated. Ancestral segment XIV and XV separated. Ancestral segment XV and XVI separated. Ancestral segment XVI and XVII separated. Ancestral segment XVII and XVIII separated. Ancestral segment XVIII and XIX separated. Ancestral segment XIX and XX separated. Ancestral segment XX and XXI separated. Ancestral segment XXI and XXII separated. Ancestral segment XXII and XXIII separated. Ancestral segment XXIII and XXIV separated. Ancestral segment XXIV and XXV separated. Ancestral segment XXV and XXVI separated. Ancestral segment XXVI and XXVII separated. Ancestral segment XXVII and XXVIII fused.

Left antennule actual 25-segmented; not-geniculated. Actual segment 1 with seta; one element; none larger than segment; straight; without spinules; without vestigial seta; without conical seta; without modified seta; without spinous process; with aesthetasc; one element. Actual segment 2 with seta; three elements; of equal size; none larger than segment; straight; without spinules; with vestigial seta; one element; without conical seta; without modified seta; without spinous process; with aesthetasc; one element. Actual segment 3 with seta; one element; one larger than segment; straight; surpassing to distal margin; beyond three sequential segments; without spinules; with vestigial seta; one element; without conical seta; without modified seta; without spinous process; with aesthetasc. Actual segment 4 with seta; one element; none larger than segment; straight; without spinules; without vestigial seta; without conical seta; without modified seta; without spinous process; without aesthetasc. Actual segment 5 with seta; one element; one larger than segment; straight; surpassing to distal margin; not beyond three sequential segments; without spinules; with vestigial seta; one element; without conical seta; without modified seta; without spinous process; with aesthetasc; one element. Actual segment 6 with seta; one element; none larger than segment; straight; without spinules; without vestigial seta; without conical seta; without modified seta; without spinous process; without aesthetasc. Actual segment 7 with seta; one element; one larger than segment; straight; surpassing to distal margin; beyond three sequential segments; without spinules; without vestigial seta; without conical seta; without modified seta; without spinous process; with aesthetasc; one element. Actual segment 8 with seta; one element; one larger than segment; straight; surpassing distal margin; without spinules; without vestigial seta; with conical seta; without modified seta; without spinous process; without aesthetasc. Actual segment 9 with seta; two elements; of unequal size; one larger than segment; straight; surpassing to distal margin; beyond three sequential segments; without spinules; without vestigial seta; without conical seta; without modified seta; without spinous process; with aesthetasc; one element. Actual segment 10 with seta; one element; none larger than segment; straight; without spinules; without vestigial seta; without conical seta; without modified seta; without spinous process; without aesthetasc. Actual segment 11 with seta; one element; one larger than segment; straight; surpassing to distal margin; beyond three sequential segments; without spinules; without vestigial seta; without conical seta; without modified seta; without spinous process; without aesthetasc. Actual segment 12 with seta; one element; one larger than segment; straight; surpassing distal margin; without spinules; without vestigial seta; with conical seta; without modified seta; without spinous process; with aesthetasc; one element. Actual segment 13 with seta; one element; none elongated; straight; surpassing distal margin; without spinules; without vestigial seta; without conical seta; without modified seta; without spinous process; without aesthetasc. Actual segment 14 with seta; one element; elongated; straight; surpassing to distal margin; beyond three sequential segments; without spinules; without vestigial seta; without conical seta; without modified seta; without spinous process; with aesthetasc; one element. Actual segment 15 with seta; one element; larger than segment; straight; surpassing to distal margin; not beyond three sequential segments; without spinules; without vestigial seta; without conical seta; without modified seta; without spinous process; without aesthetasc. Actual segment 16 with seta; one element; larger than segment; plumose; surpassing to distal margin; not beyond three sequential segments; without spinules; without vestigial seta; without conical seta; without modified seta; without spinous process; with aesthetasc; one element. Actual segment 17 with seta; one element; not larger than segment; straight; without spinules; without vestigial seta; without conical seta; without modified seta; without spinous process; without aesthetasc. Actual segment 18 with seta; one element; larger than segment; straight; surpassing to distal margin; beyond three sequential segments; without spinules; without vestigial seta; without conical seta; without modified seta; without spinous process; without aesthetasc. Actual segment 19 with seta; one element; not larger than segment; straight; surpassing distal margin; without spinules; without vestigial seta; without conical seta; without modified seta; without spinous process; with aesthetasc; one element. Actual segment 20 with seta; one element; not larger than segment; straight; surpassing distal margin; without spinules; without vestigial seta; without conical seta; without modified seta; without spinous process; without aesthetasc. Actual segment 21 with seta; one element; larger than segment; plumose; surpassing to distal margin; beyond three sequential segments; without spinules; without vestigial seta; without conical seta; without modified seta; without spinous process; without aesthetasc. Actual segment 22 with seta; two elements; of unequal size; one of them elongated; plumose; surpassing to distal margin; without spinules; without vestigial seta; without conical seta; without modified seta; without spinous process; without aesthetasc. Actual segment 23 with seta; two elements; of unequal size; one larger than segment; plumose; surpassing to distal margin; greater 3x than original segment; without spinules; without vestigial seta; without conical seta; without modified seta; without spinous process; without aesthetasc. Actual segment 24 with seta; two elements; of equal size; one larger than segment; plumose; surpassing to distal margin; greater 3x than original segment; without spinules; without vestigial seta; without conical seta; without modified seta; without spinous process; without aesthetasc. Actual segment 25 with seta; four elements; of equal size; elongated; plumose; surpassing to distal margin; 4 times larger than segment; without spinules; without vestigial seta; without conical seta; without modified seta; without spinous process; with aesthetasc; one element.

##### Antenna

Biramous. Antenna coxa separated from the basis; bearing seta; 1; on inner surface; at distal corner; reaching to the endopod 1. Antenna basis (fusion) separated from the endopodal segment; bearing seta; 2; on inner surface; at distal corner. Endopodal ancestral segment I and II separated. Ancestral segment II and III fused. Ancestral segment III and IV fused. Ancestral segment III and IV fully. Antenna endopod actual 2-segmented. Actual segment 1 not bilobate; with seta; two; on inner margin; with spinules; as a row; obliquely; on outer surface; with pore. Actual segment 2 bilobate; with discontinuity on outer cuticle; not developed as a suture; inner lobe bearing 8 setae; distally; outer lobe bearing 7 setae; distally; with spinules; as a patch; on outer surface. Antenna exopod ancestral segment I and II separated. Ancestral segment II and III fused. Ancestral segment III and IV fused. Ancestral segment IV and V separated. Ancestral segment V and VI separated. Ancestral segment VI and VII separated. Ancestral segment VII and VIII separated. Ancestral segment VIII and IX separated. Ancestral segment IX and X fused. Antenna exopod actual 7-segmented. Actual segment 1 single; elongated (width-length, equal or larger ratio 2:1); with seta; one; at inner surface. Actual segment 2 compound; elongated (larger width-length ratio 2:1); with seta; three; at inner surface. Actual segment 3 single; not elongated (lesser width-length ratio 2:1); with seta; one; at inner surface. Actual segment 4 single; not elongated (lesser width-length ratio 2:1); with seta; one; at inner surface. Actual segment 5 single; not elongated (lesser width-length ratio 2:1); with seta; one; at inner surface. Actual segment 6 single; not elongated (lesser width-length ratio 2:1); with seta; one; at inner surface. Actual segment 7 compound; elongated (larger or equal width-length ratio 2:1); with seta; one; at inner surface; and three; at distal surface.

##### Oral features

**Mandible**. Coxal gnathobase sclerotized; with lobe; prominent; on caudal margin; presence of cutting blade; with tooth-like prominence; two, distinctly; 1 acute; on caudal margin; and 1 triangular; on sub-caudal margin; without acute projection between the prominences; with additional spinules; as a row; on dorsal surface; with seta; 1; dorsally; on apical surface; with spinules; apicalmost. Mandible palps biramous; comprising the basis; with seta; four; differently inserted; first medially; reaching to beyond the endopod 1; second distally; third distally; fourth distally; on inner margin; none with setulose ornamentation. Mandible endopod 2-segmented. Mandible endopod 1 with lobe; bearing seta; four; distally inserted; without spinules. Mandible endopod 2 without lobe; bearing setae; nine elements; distally inserted; with spinules; as a row; double. Mandible exopod 4-segmented. Mandible exopod 1 with seta; one element; distally; on inner margin. Mandible exopod 2 with seta; one element; distally; on inner side. Mandible exopod 3 with seta; one element; distally; on inner side. Mandible exopod 4 with setae; three elements; on terminal region. **Maxillule**. Birramous. Maxillule 3-segmented. Maxillule praecoxa with praecoxal arthrite; bearing spines; fifteen elements; ten marginally; plus, five sub-marginally; with spinules; as a patch; on sub-marginal surface. Maxillule coxa with coxal epipodite; with conspicuous outer lobe; bearing setae; nine elements; with coxal endite; elongated (larger or equal width-length ratio 2:1); bearing setae; four elements. Maxillule basis with basal endite; double; first proximal; elongated (larger width-length ratio 2:1; separated from basis; with setae; four elements; distally inserted; second distal; fused to basis; not elongated (lesser width-length ratio 2:1); with setae; four elements; distally inserted; with setules; as a row; on inner side; basal exite present; with setae; one element; on outer surface. Maxillule endopod 1-segmented. Endopod 1 bilobate; first proximal; with setae; three elements; second distal; with setae; five elements. Maxillule exopod 1-segmented. Exopod 1 with setae; six elements; with setules; as a row; on inner side; spinules absent. **Maxilla**. Uniramous. Maxilla 5-segmented. Maxilla praecoxa fused to coxa; incompletely; distinct externally; with praecoxal endite; double; first elongated endite (larger or equal width length ratio 2:1); proximally inserted; with seta; straight, or plumose; 1 straight; 4 plumose; with spine; single; without spinules; without setule; second elongated endite (larger or equal width length ratio 2:1); distally inserted; with seta; plumose; 3 plumose; without spine; with spinules; as a row; on distal margin; with setule; as a row; on distal margin; absence of outer seta. Maxilla coxa with coxal endite; double; first elongated endite (larger or equal width); proximally inserted; with seta; plumose; 3 plumose; without spine; without spinules; with setules; as a row; on proximal margin; second elongated endite (larger or equal width); distally inserted; with seta; plumose; 3 plumose; without spine; without spinules; with setules; as a row; on proximal margin; absence of outer seta. Maxilla basis with basal endite; single; elongated (larger or equal width-length ratio 2:1); with seta; plumose; 3 plumose; without spinules; absence of outer seta. Maxilla endopod 2-segmented. Endopod 1 with seta; 2 plumose; without spine; without spinules; without setules. Maxilla endopod 2 with seta; 2 plumose; without spine; without spinules; without setules. **Maxilliped**. Uniramous; Maxilliped 8-segmented. Maxilliped praecoxa fused to coxa; incompletely; distinct internally; with praecoxal endite; not elongated (lesser width-length ratio 2:1); distally inserted; with seta; 1 straight; with spinules; as a row; single; on basal surface; without setules. Maxilliped coxa with coxal endite; three coxal endite; first elongated (larger or equal width); proximally inserted; with seta; 2 plumose; with spinules; as a patch; single; on apical surface; without setules; second not elongated (lesser width-length ratio 2:1); medially inserted; with seta; 3 plumose; with spinules; as a row; single; on medial surface; without setules; third elongated (larger or equal width length ratio 2:1); distally inserted; with seta; 3 plumose; none reaching to beyond of the basis; with spinules; as a row; single; on basal surface; without setules; with lobe; prominence; at inner distal angle; ornamented; with spinules; continuously on margin. Maxilliped basis without basal endite; with seta; 3 plumose; with spinules; as a row; single; on medial surface; with setules; as a row; single; on inner margin. Maxilliped endopod segment 6-segmented. Endopod 1 with seta; 2 plumose; on inner surface. Endopod 2 with seta; 3 plumose; on inner surface. Endopod 3 with seta; 2 plumose; on inner surface. Endopod 4 with seta; 2 plumose; on inner surface. Endopod 5 with seta; 2 plumose; on inner surface, or on outer surface; outer seta absent. Endopod 6 with seta; 4 plumose; on inner surface, or on outer surface.

##### Swimming legs features

**First swimming legs.** Symmetrical; biramous. First swimming legs intercoxal plate without seta. First swimming legs praecoxa absent. First swimming legs coxa with seta; one; straight; distally inserted; on inner surface; surpassing to first endopodal segment; with setules; two group; as a patch; on inner margin; and as a row; double; on anterior surface; outerly; without spinules; without spine. First swimming legs basis without seta; with setules; as a patch; single; on outer surface; without spinules; without spine. First swimming legs endopod 2-segmented. Endopod 1 with seta; straight; restricted; to inner surface; one element; without spine; with setules; as a row; single; continuously; on outer surface; without spinules; absence of Schmeil’s organ. Endopod 2 with seta; unrestricted; three on inner surface; one on outer surface; two on distal surface; straight; without spine; with setules; as a row; single; continuously; on outer surface; without spinules; absence of Schmeil’s organ. Endopod 3 absence. First swimming legs exopod 1 with seta; restricted; 1 on inner surface; with spine; 1; stout; smaller than original segment; serrated; on inner side; continuously; with setules; as a row; single; as a row; innerly. First swimming legs exopod 2 with seta; restricted; 1 on inner surface; straight; without spine; with setules; as a row; single; continuously; on inner margin, or on outer margin; without spinules. First swimming legs exopod 3 with setule; as a row; single; continuously; on outer surface; without spinules; with seta; unrestricted; 2 on inner surface; 2 on terminal surface; with spine; 2; unequal size; first no longer 2x than origin segment; stout; serrated; on inner side, or on outer side; equally; second longer 3x than origin segment; slender; serrated; on outer side; with ornamentation on non-serrated side; by setules. **Second swimming legs**. Symmetrical; Second swimming legs biramous. Second swimming legs intercoxal plate without seta. Second swimming legs praecoxa present; located laterally. Second swimming legs coxa with seta; straight; distally inserted; on inner surface; surpassing to basal segment; without setules; without spinules; without spine. Second swimming legs basis without seta; without setules; without spinules; without spine. Second swimming legs endopod 3-segmented. Endopod 1 with seta; straight; restricted; one on inner surface; without spine; with setules; as a row; single; continuously; on outer surface; without spinules; absence of Schmeil’s organ. Endopod 2 with seta; straight; unrestricted; two on inner surface; without spine; with setules; as a row; single; continuously; on outer side; without spinules; presence of Schmeil’s organ; on posterior surface. Endopod 3 with seta; straight; unrestricted; three on inner surface; two on outer surface; two on distal surface; without spine; without setules; with spinules; as a row; double; distally inserted; at anterior surface; absence of Schmeil’s organ. Second swimming legs exopod 1 with seta; restricted; one on inner surface; with spine; 1; stout; not reaching to distal-third of the exopod 2; serrated; on inner side, or on outer side; with setules; as a row; single; continuously; on inner side; without spinules; absence of Schmeil’s organ. Exopod 2 with seta; unrestricted; one on inner surface; with spine; 1; stout; not surpassing the exopod 3; serrated; on inner side, or on outer side; with setules; as a row; single; continuously; on inner surface; without spinules; absence of Schmeil’s organ. Exopod 3 with seta; plurimarginal; three on inner surface; two on terminal surface; with spine; 2; unequal size; first no longer 2x than origin segment; stout; serrated; on inner side, or on outer side; equally; second longer 2x than origin segment; slender; serrated; on outer side; with ornamentation on non-serrated side; of setules; setules on outer surface; as a row; single; continuously; on inner surface; with spinules; as a row; single; distally inserted; at anterior surface; absence of Schmeil’s organ. **Third swimming legs**. Symmetrical; Third swimming legs biramous. Third swimming legs intercoxal plate without seta. Third swimming legs praecoxa present; not laterally located. Third swimming legs coxa with seta; straight; distally inserted; on inner surface; surpassing to first endopodal segment; without setules; without spinules; without spine. Third swimming legs basis without seta; without setules; without spinules; without spine. Third swimming legs endopod 3-segmented. Endopod 1 with seta; restricted; one on inner surface; without spine; without setules; without spinules; absence of Schmeil’s organ. Endopod 2 with seta; restricted; two on inner surface; straight; without spine; without setules; without spinules; absence of Schmeil’s organ. Endopod 3 with seta; straight; plurimarginal; two on inner surface; two on outer surface; three on terminal surface; without spine; without setules; with spinules; as a row; distally inserted; double; at anterior surface; absence of Schmeil’s organ. Third swimming legs exopod 1 with seta; restricted; straight; one on inner surface; with spine; 1; stout; not reaching to the distal-third of the exopod 2; serrated; equally; on inner surface, or on outer surface; with setules; as a row; single; continuously; on inner surface; without spinules; absence of Schmeil’s organ. Exopod 2 with seta; straight; restricted; one on inner surface; with spine; 1; stout; not reaching out to exopod 3; serrated; on inner side, or on outer side; equally; with setules; as a row; single; continuously; on inner side; without spinules; absence of Schmeil’s organ. Exopod 3 without setules; with spinules; as a row; single; distally inserted; at anterior surface; with seta; straight; unrestricted; three on inner surface; two on terminal surface; with spine; 2; unequal size; first no longer 2x than origin segment; stout; serrated; on inner side, or on outer side; equally; second longer 2x than origin segment; slender; serrated; on outer side; with ornamentation on non-serrated side; of setules; absence of Schmeil’s organ. **Fourth swimming legs**. Symmetrical; biramous. Intercoxal plate without sensilla. Praecoxa present. Coxa with seta; distally inserted; on inner margin; reaching out to endopod 1; without spinules; setules absent. Basis with seta; one; medially inserted; on posterior surface; smaller than the original segment; without setules; without spinules; without spine. Fourth swimming legs endopod 3-segmented. Endopod 1 with seta; one; restricted; on inner surface; without spine; without setules; without spinules; absence of Schmeil’s organ. Endopod 2 with seta; restricted; two on inner side; without spine; with setules; as a row; single; continuously; on outer surface; without spinules; absence of Schmeil’s organ. Endopod 3 with seta; unrestricted; two on inner surface; two on outer surface; three on distal surface; without spine; without setules; with spinules; as a row; double; distally inserted; at anterior surface; absence of Schmeil’s organ. Fourth swimming legs exopod 1 with seta; restricted; one on inner surface; with spine; 1; stout; not reaching out to distal-third of the exopod 2; serrated; on inner side, or on outer side; equally; with setules; as a row; single; continuously; on inner surface; without spinules; absence of Schmeil’s organ. Exopod 2 with seta; restricted; one on inner surface; with spine; 1; stout; not reaching the end of exopod 3; serrated; on inner side, or on outer side; equally; with setules; as a row; single; continuously; on inner surface; without spinules; absence of Schmeil’s organ. Exopod 3 without setules; with spinules; as a row; single; distally inserted; at anterior surface; with seta; unrestricted; three on inner surface; two on distal surface; with spine; 2; unequal size; first no longer 2x than origin segment; stout; serrated; on inner side, or on outer side; equally; second longer 2x than origin segment; slender; serrated; on outer side; without ornamentation on non-serrated side; absence of Schmeil’s organ.

##### Fifth swimming legs features

Asymmetrical. Fifth swimming leg intercoxal plate with length not equal or greater than width on 1.5x; with irregular proximal margin; discontinuous to; the anterior margin of the left coxa, or the anterior margin of the right coxa; posterior sensilla on the right lateral absent. **Fifth left swimming leg**. Fifth left swimming leg biramous; leg surpassing first right exopod segment. Fifth left swimming leg praecoxa present; rudimentary; separated from the coxae; without ornamentation. Fifth left swimming leg coxa concave inner side; without teeth-like structures; with process; conical; on posterior surface; outer side; distally inserted; not projecting over basis; with sensilla; stout; triangular; at apex; no longer 2x than insertion basis; with swelling; on inner side; distally; without seta; without spinules. Fifth left swimming leg basis sub-cylindrical; unequal size between inner and outer side; shorter outer than inner side; with concave inner side; rounded internal proximal expansion absent; without outgrowth; with groove; deep; obliquely; on posterior surface; not reaching the endopodal lobe; not ornamented; absence of protuberance; with seta; outerly inserted; no longer 2x than origin segment; presence of minutely granular; as a patch; innerly. Fifth left swimming leg endopod segments 1 and 2 fused; segments 2 and 3 fused; 1-segmented; stout; separated from the basis; ornamented; on inner side; with spinules; more than four elements; as a row; terminally; row of setules absent; without seta. Fifth left swimming leg exopod segments 1 and 2 separated; segments 2 and 3 fused; 2-segmented; stout; separated from the basis. Fifth left swimming leg exopod 1 sub-cylindrical; longer than broad; unequal size between inner and outer side; shorter inner than outer side; concave inner side; convex outer side; without swelling; without marginal extension; without process; with lobe; double; semicircular; medially inserted; on inner side; covered; by setules; without outer spine; absence seta. Fifth left swimming leg exopod 2 digitiform; longer than broad; equal size between inner and outer side; disform inner side; with rectilinear outer side; setulose pad present; prominently rounded; proximally; on inner side; inflated medial region absent; distal process present; digitiform; non denticulate; without transverse row of denticles; none oblique row of 5 denticles; not innerly directed; with seta; spiniform; not ornamented by spinules; not surpassing the distal-point of the segment; without outer spine; terminal claw absent.

##### Fifth right swimming leg

Biramous. Fifth right swimming leg praecoxa present; separated from the coxae; without ornamentation. Fifth right swimming leg coxa convex inner side; without teeth-like structures; with process; rounded; distally inserted; on posterior surface; closest to the outer rim; projecting over basis; beyond the first third; until the medial surface; without triangular protuberance innerly; with sensilla; slender; at apex; no longer 2x than basal insertion; without marginal extension; without seta; without spinules. Fifth right swimming leg basis cylindrical; unequal size between inner and outer side; shorter outer than inner side; rectilinear inner side; tumescence present; not inflated; restricted on inner surface; proximally; without protuberance; presence of distinct minutely granular; as a patch; proximally; on inner side; additional inner process absent; with posterior groove; deep; obliquely; reaching the endopodal lobe; ornamented; with tubercles; throughout of the outer border; with seta; outerly inserted; on anterior surface; no longer 2x than origin segment; posterior protrusion present; distal process absent. Fifth right swimming leg with endopodite present; separated from the basis; on anterior surface; ancestral segments 1 and 2 fused; ancestral segments 2 and 3 fused; 1-segmented; stout; ornamented; with setules; as a row; on inner side; terminally; without seta. Fifth right swimming leg exopod segments 1 and 2 separated; segments 2 and 3 fused; 2-segmented; stout; separated from the basis. Fifth right swimming leg exopod 1 trapezium; longer than broad; nearly 1.25 times; unequal size between both sides; shorter inner than outer side; convex inner side; rectilinear outer side; with marginal extension; sub-triangular; distally inserted; at outer rim; spinules absent; with process; triangular; arched; internally directed; blunt tip; sclerotized; without ornamentation; distally inserted; at posterior surface; projecting over next segment; without outer spine; without seta; internal prominence absent; lamella on posterior surface absent. Fifth right swimming leg exopod 2 elliptical; longer than broad; nearly 2.5 times; equal size between both sides; disform inner side; convex outer side; without posterior proximal swelling; inner-posterior process absent; without marginal expansion; curved ridge on distal posterior surface present; chitinous knobs absent; with outer spine; inserted sub-distally; arched; externally directed; ornamented innerly; by spinules; as a row; not ornamented outerly; sharp tip; without apparent curve; lesser than the length of the exopod 2; until to 2 times its size; 2x; sensilla absent; terminal claw present; equal or longer 1.5 times than insertion segment; sclerotized; arched; inward; with conspicuous curve; medially; ornamented innerly; by spinules; as a row; partially on extension; medially, or distally; not ornamented outerly; sharp tip; curved tip; outwards; without medial constriction; hyaline process absent.

##### FEMALE

Body longer and wider than male; Female body 1471 micrometers excluding caudal setae. Widest at first metasome segment. Distal margin of the prosomal segments without one line of setules at posterior margin. Prosome segments without spinules at prosomal segments. Fourth metasome segment absence of dorsal protuberance. Fourth and fifth metasome segments fused; partially; on dorsal surface. Limit between fourth and fifth metasome segments without ornamentation. **Fifth metasome segment**. Fifth metasome segment without sensilla; with epimeral plates. Epimeral plates asymmetrical. Right epimeral plates prominent, as projections; not thinner than the left; one posterior-dorsally directed; not reaching half length of the genital segment; with sensilla at the apex; dorsal-posterior sensilla present; slender; without ornamentation. Left epimeral plate without expansion.

##### Urosome

3-segmented. **Genital double-somite**. Asymmetrical in dorsal view; longer than broad; longer than other urosomites combined; dorsal suture at mid-length absent; not covered by spinules; with swelling; rounded; unequal size; greater left than right; anteriorly; with sensillae; on both sides; one; stout; with robust apex; at left lateral; not on lobular base; medially; one; stout; at right lateral; not on lobular base; anteriorly; with robust apex; of equal size between then; lateral protuberance present; wrinkled; on the right side; with right posterior rim expanded; over next segment; without slender sensilla on each posterior rim; without posterior-dorsal process. Genital double-somite opercular pad present; broader than longer; symmetrical; development laterally; expanded posteriorly; covering partially; double gonoporal slit; located ventrally; with arthrodial membrane; inserted anteriorly; post-genital process present; single; disto-ventral tumescence absent; ventral vertical folds absent; dorsal sensilla absent. Second urosome segment without ventral fusion to anal segment; right distal process absent. Caudal rami patch of setules on outer surface absent; patch of spinules on outer surface absent.

##### Oral appendices feature

Rostrum basal process absent. **Antennules**. Symmetrical. Right antennule surpassing to genital double-segment; extending beyond caudal rami. Right antennule not exceeding the caudal setae. Right antennule ornamentation pattern equals to male left antennule; fully.

##### Fifth swimming legs

Symmetrical; Fifth swimming legs biramous. Fifth swimming legs intercoxal plate longer than wide; separated from the legs. Fifth swimming legs praecoxa with sclerite praecoxal; separated from the coxae; without ornamentation. Fifth swimming legs coxa with process; conical; at the outer rim; distally; sensilla present; stout; at apex; projecting over basal segment; no longer 2x than basal insertion; marginal extension absent; without swelling; without seta; without spinules. Fifth swimming legs basis sub-triangular; unequal size between inner and outer sides; shorter outer than inner side; with convex inner side; without proximal inner outgrowth; without groove; with distal extension; on posterior surface; with seta; outerly inserted; on anterior surface; longer 2x than origin segment; reaching to exopod 1 distally. Fifth swimming legs endopod segments 1 and 2 fused; segments 2 and 3 fused; 1-segmented; stout; separated from the basis; present discontinuity cuticle; on inner side; with spinules; as a row; single; non-oblique; sub-terminally; at anterior surface; with seta; double; one medially; on posterior surface; rectilinear; one distally; on posterior surface; rectilinear; of unequal size; distal seta longer than medial seta. Fifth swimming legs exopod segments 1 and 2 separated; segments 2 and 3 separated; 3-segmented; separated from the basis. Fifth swimming legs exopod 1 sub-cylindrical; longer than wide; longer or equal than 2 times; with unequal size between inner and outer side; shorter inner than outer side; with convex inner side; with rectilinear outer side; without swelling; without marginal extension; without posterior process; without spine; without seta. Fifth swimming legs exopod 2 sub-cylindrical; longer than broad; longer or equal than 2 times; without swelling; without marginal extension; without process; without lobe; with spine; inserted laterally; rectilinear; without ornamentation; sharp tip; equal size or larger than next segment; without seta. Fifth swimming legs exopod 3 cylindrical; longer than wide; without swelling; without process; without lobe; without spine; with seta; double; inserted terminally; unequal size between them; outer seta smaller than inner; nearly 3 times; outer seta not ornamented by setules; without ornamentation; presence of terminal claw; sclerotized; arched; externally directed; convex inner side; with ornamentation; of denticles; as a row; on surface partially; at medial region; concave outer side; with ornamentation; of denticles; as a row; on surface partially; at medial region; blunt tip; 6 times longer than origin segment.

##### Distribution records

###### BRAZIL

**Ceará**: five ponds of the river Jaguaribe, four ponds near to Fortaleza city, and a near to Sobral City (Wright, 1938a). **Paraíba**: Simão Pond, Campina Grande City; pond near to Campina Grande; Linda Flor Pond, Mogeiro de Baixo, and Lapa, Campina Grande; pond in the Cabedelo (Wright, 1935; Reid, 1991).

##### Habitat

Habitat in freshwaters: ponds, and floodplain associated to rivers.

##### Remarks

The species was described from floodplain organisms in Northeastern Brazil and has belonged to the *nordestinus* complex since its foundation. Kiefer (1936) in creating *Notodiaptomus* recombined the taxon which again underwent attempted recombination from the proposal for the subgenus *Notodiaptomus* (*Notodiaptomus*) (Dussart, 1984a). Santos-Silva *et al*. (2015), when reviewing the *nordestinus* complex approached the morphological inconsistencies identified in the taxonomic trajectory of the species, making comments, and contributing with illustrations that established the status of the species definitively.

For the present effort, we corroborate the observations presented in the review by Santos-Silva *et al*. (2015). Among the morphological characteristics of the species that differ from those considered in Wright (1935; 1936; 1937) and Kiefer (1936; 1956) it is only not present in the species: (1) female fifth swimming legs endopod with spinules not obliquely. Additionally, we identified other attributes that make the species unique within *Notodiaptomus* and diverge from the type-species attributes converging within the genus: (1) male fifth right swimming leg basis without inner protuberance; (2) male fifth left swimming leg exopod 1 with medial double semicircular lobe innerly; and (3) female fourth and fifth metasome segment fused partially on dorsal surface.

#### Notodiaptomus orellanai Dussart, 1979

##### Synonymy

*Notodiaptomus orellanai* Dussart, 1979: 5–6, fig. 3. *Notodiaptomus spiniger*; Perbiche-Neves *et al*., 2015: 82–85, figs. 74–78.

##### Type locality

Leconte Lake, Corrientes Province, Argentina.

##### Type material

In the original description, no holotype or any specimen of the type-series is specified. It is only specified type locality, J. de Orellana collector and dated 19.VII.1975. Material from the Province of Corrientes, Argentina was located in the National Museum of Natural History in Paris: 20 males, 20 females from Leconte Lake, 19.VII.1975. I. de Orellana coll. (MNHN-Paris 420); 19 males, 7 females from the Laguna Cabrera, Province of Corrientes. 11.VIII.1975 (MNHN-419). The first probably represents all or part of the material used to describe the species.

##### Material examined

Topotype: 2 males, 3 females, entire in alcohol, from Leconte Lake, Province Corrientes. 19.VII.1975. I. de Orellana coll. (MNHN-Paris 420). Non-type material: 2 males, 2 females, entire in alcohol, from Lagoon Cabrera, Province Corrientes, 11.VIII.1975. I. de Orellana coll. (MNHN-Paris 419). 1 male (INPA-COP043, slides a-h) and 1 female (INPA-COP044, slides a-h) were selected to be dissection on eight slides each and deposited in the Zoological Collection of the INPA, Brazil.

##### Diagnosis

**(1)** third metasome segment with double spinules row dorsally; **(2)** male limit between fourth and fifth metasome segments with spinules row doubly; **(3)** male fifth metasome segment with 4 sensillae dorsally; **(4)** male third and fourth urosome with spinules as a patch, double throughout dorsal surface; **(5)** male right antennule actual segment 13 with modified seta parallel to antennule direction; **(6)** male fifth left swimming leg coxa with outer process projecting over basis reaching to the proximal surface; **(7)** male fifth left swimming leg basis with absence inner minutely granular; **(8)** male fifth right swimming leg exopod 2 with medial conspicuous curve on terminal claw inward; **(9)** male fifth right swimming leg basis with single inner protuberance in triangular form, medially, and additional inner process present; **(10)** male fifth right swimming leg exopod 1 broader than long; **(11)** male fifth right swimming leg exopod 1 with outer extension, acute, at the rim, distally inserted; **(12)** male fifth right swimming leg exopod 1 with triangular process, rectilinear, sharp tip, inserted inner rim, projecting over next segment; **(13)** female limit between fourth and fifth metasome segments with spinules patch dorsally; **(14)** female left epimeral plate with posterior semicircular expansion dorsally, with sensilla at apex; **(15)** female genital double-somite with lateral sensilla of unequal size between then, left bigger than right; **(16)** female right antennule with length extending beyond caudal rami, exceeding the caudal setae; **(17)** female fifth swimming legs coxa with sensilla longer 2x than outer conical process; **(18)** female fifth swimming legs coxa with distal triangular marginal extension at inner rim; **(19)** female fifth swimming legs basis with distal extension posteriorly; **(20)** female fifth swimming legs exopod 2 with lateral spine longer than next segment

##### Redescription

###### MALE

Body 1475 micrometers excluding caudal setae. Male body smaller and slenderer than female. Nerve axons myelinated. Prosome 6-segmented; widest at first metasome segment; without one line of setules at posterior margin; with spinules at least at one segment. Cephalosome anterior margin sub-triangular; with dorsal suture; incomplete; separate from first metasome segment. First metasome segment without sensilla. Second metasome segment without sensilla. Third metasome segment with sensillae; 4 dorsally; of equal size; ornamented posterior margin; with spinules; as a row; double; dorsally, or laterally. Fourth metasome segment with sensillae; 4 dorsally; of equal size; separated from the fifth metasome. Limit between fourth and fifth metasome segments ornamented; with spinules; as a row; on dorsal doubly; on lateral doubly; same size. Fifth metasome segment with sensilla; 4 dorsally; Fifth metasome segment equal size; Fifth metasome segment without ornamentation; Fifth metasome segment without dorsal conical process; with epimeral plates. Epimeral plates symmetrical. Right epimeral plates prominent, as projections; one projection; posterior-dorsally directed; not reaching half length of the genital segment; with sensilla; at the apex of projection; without ornamentation.

##### Urosome

5-segmented; Urosome 5-free segments. Genital somite asymmetrical in dorsal view; with single aperture; located on left side; ventrolaterally on posterior rim; with sensillae; on both sides; one; at left lateral; posteriorly; one; at right rim; posteriorly; of equal size between then. Third urosome segment with spinules; as a patch; double throughout dorsal surface; without external seta. Fourth urosome segment with spinules; as a patch; double throughout dorsal surface; without sub-conical blunt dorsal-lateral process. Anal segment presence of dorsal sensillae; one on each side; medially inserted; presence of operculum; convex; not covering the anal aperture fully. Caudal rami symmetrical; separated from anal segment; longer than wide; with setules; continuous on; inner side; each ramus bearing 6 caudal setae; 5 marginals; plumose; and 1 internal dorsally; plumose; not reticulated main axis; outermost seta with outer spiniform process absent.

##### Oral appendices feature

Rostrum symmetrical; fused to dorsal cephalic shield; by complete suture; sensillae present; one pair; anteriorly inserted on surface tegument; with rostral filament; double; paired; extended; into blunt protrusion; with basal process; in ventral view, rounded on left side; with a smaller basal expansion on the right side.

##### Antennules

Asymmetrical. **Right antennules**. Uniramous; right antennule surpassing to genital segment; right antennule extending beyond caudal rami.

Right antennule ancestral segment I and II separated. Ancestral segment II and III fused. Ancestral segment III and IV fused. Ancestral segment IV and V separated. Ancestral segment V and VI separated. Ancestral segment VI and VII separated. Ancestral segment VII and VIII separated. Ancestral segment VIII and IX separated. Ancestral segment IX and X separated. Ancestral segment X and XI separated. Ancestral segment XI and XII separated. Ancestral segment XII and XIII separated. Ancestral segment XIII and XIV separated. Ancestral segment XIV and XV separated. Ancestral segment XV and XVI separated. Ancestral segment XVI and XVII separated. Ancestral segment XVII and XVIII separated. Ancestral segment XVIII and XIX separated. Ancestral segment XIX and XX separated. Ancestral segment XX and XXI separated. Ancestral segment XXI and XXII fused. Ancestral segment XXII and XXIII fused. Ancestral segment XXIII and XXIV separated. Ancestral segment XXIV and XXV fused. Ancestral segment XXV and XXVI separated. Ancestral segment XXVI and XXVII separated. Ancestral segment XXVII and XXVIII fused.

Right antennule actual 22-segmented; geniculated; between the segment 18 and segment 19; with swollen and modified region; formed by 5 segments; between 13 and 17 segments. Actual segment 1 with seta; one element; straight; none larger than segment; without spinules; without vestigial seta; without conical seta; without modified seta; without spinous process; with aesthetasc; one element. Actual segment 2 with seta; three elements; of unequal size; straight; none larger than segment; without spinules; with vestigial seta; one element; without conical seta; without modified seta; without spinous process; with aesthetasc; one element. Actual segment 3 with seta; one element; one larger than segment; surpassing to distal margin; beyond three sequential segments; straight; blunt apex; without spinules; with vestigial seta; one element; without conical seta; without modified seta; without spinous process; with aesthetasc. Actual segment 4 with seta; one element; one larger than segment; surpassing to distal margin; straight; not beyond three sequential segments; without spinules; without vestigial seta; without conical seta; without modified seta; without spinous process; without aesthetasc. Actual segment 5 with seta; one element; straight; one larger than segment; surpassing to distal margin; not beyond three sequential segments; without spinules; with vestigial seta; one element; without conical seta; without modified seta; without spinous process; with aesthetasc; one element. Actual segment 6 with seta; one element; none larger than segment; straight; without spinules; without vestigial seta; without conical seta; without modified seta; without spinous process; without aesthetasc. Actual segment 7 with seta; one element; straight; one larger than segment; surpassing to distal margin; beyond three sequential segments; blunt apex; without spinules; without vestigial seta; without conical seta; without modified seta; without spinous process; with aesthetasc; one element. Actual segment 8 with seta; one element; straight; none larger than segment; without spinules; without vestigial seta; with conical seta; one element; not reaching to middle-point of the sequent segment; without modified seta; without spinous process; without aesthetasc. Actual segment 9 with seta; two elements; of unequal size; straight; one larger than segment; surpassing to distal margin; beyond three sequential segments; blunt apex; without spinules; without vestigial seta; without conical seta; without modified seta; without spinous process; with aesthetasc; one element. Actual segment 10 with seta; one element; straight; none larger than segment; without spinules; without vestigial seta; without conical seta; with modified seta; presenting blunt apex; slender form; surpassing to distal margin; beyond of the sequential segment; parallel to antennule direction; without spinous process; without aesthetasc. Actual segment 11 with seta; one element; straight; one larger than segment; surpassing to distal margin; not beyond three sequential segments; without spinules; without vestigial seta; without conical seta; with modified seta; slender form; presenting blunt apex; surpassing to distal margin; beyond of the sequential segment; parallel to antennule direction; shorter length than homologous of actual segment 13; without spinous process; without aesthetasc. Actual segment 12 with seta; one element; straight; one larger than segment; surpassing to distal margin; not beyond three sequential segments; without spinules; without vestigial seta; with conical seta; one element; not smaller than to segment 8; without modified seta; without spinous process; with aesthetasc; one element; absent internal perpendicular fission. Actual segment 13 with seta; one element; straight; one larger than segment; surpassing to distal margin; not beyond three sequential segments; without spinules; without vestigial seta; without conical seta; with modified seta; stout form; surpassing to distal margin; to the middle-point of the sequence segment; parallel to antennule direction; presenting bifid apex; without spinous process; with aesthetasc; one element. Actual segment 14 with seta; two elements; of unequal size; straight; one larger than segment; surpassing to distal margin; beyond three sequential segments; blunt apex; without spinules; without vestigial seta; without conical seta; without modified seta; without spinous process; with aesthetasc; one element. Actual segment 15 with seta; two elements; of unequal size; straight; not bifidform; none larger than segment; without spinules; without vestigial seta; without conical seta; without modified seta; with spinous process; on outer margin; surpassing distal margin; with aesthetasc; one element. Actual segment 16 with seta; two elements; of unequal size; plumose; one larger than segment; surpassing to distal margin; not beyond three sequential segments; not bifidform; without spinules; without vestigial seta; without conical seta; without modified seta; with spinous process; on outer margin; surpassing distal margin; unequal size to process on preceding segment; with aesthetasc; one element. Actual segment 17 with seta; two elements; of unequal size; straight; none larger than segment; bifidform; without spinules; without vestigial seta; without conical seta; with modified seta; one element; stout form; surpassing to distal margin; not beyond of the sequential segment; parallel to antennule direction; without spinous process; without aesthetasc. Actual segment 18 with seta; two elements; of equal size; straight; none larger than segment; without spinules; without vestigial seta; without conical seta; with modified seta; one element; stout form; surpassing distal margin; parallel to antennule direction; without spinous process; without aesthetasc. Actual segment 19 with seta; two elements; of unequal size; plumose; none larger than segment; without spinules; without vestigial seta; without conical seta; with modified seta; two elements; stout form; at least one bifid form; surpassing distal margin; parallel to antennule direction; without spinous process; with aesthetasc; one element. Actual segment 20 with seta; four elements; of unequal size; straight; one larger than segment; surpassing to distal margin; beyond three sequential segments; without spinules; without vestigial seta; without conical seta; without modified seta; with spinous process; distally; not reaching beyond of distal-point segment 21; without aesthetasc. Actual segment 21 with seta; two elements; of equal size; plumose; one larger than segment; surpassing to distal margin; greater 3x than original segment; without spinules; without vestigial seta; without conical seta; without modified seta; without spinous process; without aesthetasc. Actual segment 22 with seta; four elements; of equal size; one larger than segment; plumose; surpassing to distal margin; greater 3x than original segment; without spinules; without vestigial seta; without conical seta; without modified seta; without spinous process; with aesthetasc; one element.

##### Left antennules

Uniramous; Left antennule surpassing to prosome; Left antennule extending beyond caudal rami. Ancestral segment I and II separated. Ancestral segment II and III fused. Ancestral segment III and IV fused. Ancestral segment IV and V separated. Ancestral segment V and VI separated. Ancestral segment VI and VII separated. Ancestral segment VII and VIII separated. Ancestral segment VIII and IX separated. Ancestral segment IX and X separated. Ancestral segment X and XI separated. Ancestral segment XI and XII separated. Ancestral segment XII and XIII separated. Ancestral segment XIII and XIV separated. Ancestral segment XIV and XV separated. Ancestral segment XV and XVI separated. Ancestral segment XVI and XVII separated. Ancestral segment XVII and XVIII separated. Ancestral segment XVIII and XIX separated. Ancestral segment XIX and XX separated. Ancestral segment XX and XXI separated. Ancestral segment XXI and XXII separated. Ancestral segment XXII and XXIII separated. Ancestral segment XXIII and XXIV separated. Ancestral segment XXIV and XXV separated. Ancestral segment XXV and XXVI separated. Ancestral segment XXVI and XXVII separated. Ancestral segment XXVII and XXVIII fused.

Left antennule actual 25-segmented; not-geniculated. Actual segment 1 with seta; one element; none larger than segment; straight; without spinules; without vestigial seta; without conical seta; without modified seta; without spinous process; with aesthetasc; one element. Actual segment 2 with seta; three elements; of equal size; none larger than segment; straight; without spinules; with vestigial seta; one element; without conical seta; without modified seta; without spinous process; with aesthetasc; one element. Actual segment 3 with seta; one element; one larger than segment; straight; surpassing to distal margin; beyond three sequential segments; without spinules; with vestigial seta; one element; without conical seta; without modified seta; without spinous process; with aesthetasc. Actual segment 4 with seta; one element; none larger than segment; straight; without spinules; without vestigial seta; without conical seta; without modified seta; without spinous process; without aesthetasc. Actual segment 5 with seta; one element; one larger than segment; straight; surpassing to distal margin; not beyond three sequential segments; without spinules; with vestigial seta; one element; without conical seta; without modified seta; without spinous process; with aesthetasc; one element. Actual segment 6 with seta; one element; none larger than segment; straight; without spinules; without vestigial seta; without conical seta; without modified seta; without spinous process; without aesthetasc. Actual segment 7 with seta; one element; one larger than segment; straight; surpassing to distal margin; beyond three sequential segments; without spinules; without vestigial seta; without conical seta; without modified seta; without spinous process; with aesthetasc; one element. Actual segment 8 with seta; one element; one larger than segment; straight; surpassing distal margin; without spinules; without vestigial seta; with conical seta; without modified seta; without spinous process; without aesthetasc. Actual segment 9 with seta; two elements; of unequal size; one larger than segment; straight; surpassing to distal margin; beyond three sequential segments; without spinules; without vestigial seta; without conical seta; without modified seta; without spinous process; with aesthetasc; one element. Actual segment 10 with seta; one element; none larger than segment; straight; without spinules; without vestigial seta; without conical seta; without modified seta; without spinous process; without aesthetasc. Actual segment 11 with seta; one element; one larger than segment; straight; surpassing to distal margin; beyond three sequential segments; without spinules; without vestigial seta; without conical seta; without modified seta; without spinous process; without aesthetasc. Actual segment 12 with seta; one element; one larger than segment; straight; surpassing distal margin; without spinules; without vestigial seta; with conical seta; without modified seta; without spinous process; with aesthetasc; one element. Actual segment 13 with seta; one element; none elongated; straight; surpassing distal margin; without spinules; without vestigial seta; without conical seta; without modified seta; without spinous process; without aesthetasc. Actual segment 14 with seta; one element; elongated; straight; surpassing to distal margin; beyond three sequential segments; without spinules; without vestigial seta; without conical seta; without modified seta; without spinous process; with aesthetasc; one element. Actual segment 15 with seta; one element; larger than segment; straight; surpassing to distal margin; not beyond three sequential segments; without spinules; without vestigial seta; without conical seta; without modified seta; without spinous process; without aesthetasc. Actual segment 16 with seta; one element; larger than segment; plumose; surpassing to distal margin; not beyond three sequential segments; without spinules; without vestigial seta; without conical seta; without modified seta; without spinous process; with aesthetasc; one element. Actual segment 17 with seta; one element; not larger than segment; straight; without spinules; without vestigial seta; without conical seta; without modified seta; without spinous process; without aesthetasc. Actual segment 18 with seta; one element; larger than segment; straight; surpassing to distal margin; beyond three sequential segments; without spinules; without vestigial seta; without conical seta; without modified seta; without spinous process; without aesthetasc. Actual segment 19 with seta; one element; not larger than segment; straight; surpassing distal margin; without spinules; without vestigial seta; without conical seta; without modified seta; without spinous process; with aesthetasc; one element. Actual segment 20 with seta; one element; not larger than segment; straight; surpassing distal margin; without spinules; without vestigial seta; without conical seta; without modified seta; without spinous process; without aesthetasc. Actual segment 21 with seta; one element; larger than segment; plumose; surpassing to distal margin; beyond three sequential segments; without spinules; without vestigial seta; without conical seta; without modified seta; without spinous process; without aesthetasc. Actual segment 22 with seta; two elements; of unequal size; one of them elongated; plumose; surpassing to distal margin; without spinules; without vestigial seta; without conical seta; without modified seta; without spinous process; without aesthetasc. Actual segment 23 with seta; two elements; of unequal size; one larger than segment; plumose; surpassing to distal margin; greater 3x than original segment; without spinules; without vestigial seta; without conical seta; without modified seta; without spinous process; without aesthetasc. Actual segment 24 with seta; two elements; of equal size; one larger than segment; plumose; surpassing to distal margin; greater 3x than original segment; without spinules; without vestigial seta; without conical seta; without modified seta; without spinous process; without aesthetasc. Actual segment 25 with seta; four elements; of equal size; elongated; plumose; surpassing to distal margin; 4 times larger than segment; without spinules; without vestigial seta; without conical seta; without modified seta; without spinous process; with aesthetasc; one element.

##### Antenna

Biramous. Antenna coxa separated from the basis; bearing seta; 1; on inner surface; at distal corner; reaching to the endopod 1. Antenna basis (fusion) separated from the endopodal segment; bearing seta; 2; on inner surface; at distal corner. Endopodal ancestral segment I and II separated. Ancestral segment II and III fused. Ancestral segment III and IV fused. Ancestral segment III and IV fully. Antenna endopod actual 2-segmented. Actual segment 1 not bilobate; with seta; two; on inner margin; with spinules; as a row; obliquely; on outer surface; without pore. Actual segment 2 bilobate; with discontinuity on outer cuticle; not developed as a suture; inner lobe bearing 8 setae; distally; outer lobe bearing 7 setae; distally; with spinules; as a patch; on outer surface. Antenna exopod ancestral segment I and II separated. Ancestral segment II and III fused. Ancestral segment III and IV fused. Ancestral segment IV and V separated. Ancestral segment V and VI separated. Ancestral segment VI and VII separated. Ancestral segment VII and VIII separated. Ancestral segment VIII and IX separated. Ancestral segment IX and X fused. Antenna exopod actual 7-segmented. Actual segment 1 single; elongated (width-length, equal or larger ratio 2:1); with seta; one; at inner surface. Actual segment 2 compound; elongated (larger width-length ratio 2:1); with seta; three; at inner surface. Actual segment 3 single; not elongated (lesser width-length ratio 2:1); with seta; one; at inner surface. Actual segment 4 single; not elongated (lesser width-length ratio 2:1); with seta; one; at inner surface. Actual segment 5 single; not elongated (lesser width-length ratio 2:1); with seta; one; at inner surface. Actual segment 6 single; not elongated (lesser width-length ratio 2:1); with seta; one; at inner surface. Actual segment 7 compound; elongated (larger or equal width-length ratio 2:1); with seta; one; at inner surface; and three; at distal surface.

##### Oral features

**Mandible**. Coxal gnathobase sclerotized; without lobe; presence of cutting blade; with tooth-like prominence; two, distinctly; 1 acute; on caudal margin; and 1 triangular; on sub-caudal margin; without acute projection between the prominences; without additional spinules; with seta; 1; dorsally; on apical surface; with spinules; apicalmost. Mandible palps biramous; comprising the basis; with seta; four; differently inserted; first medially; reaching to beyond the endopod 1; second distally; third distally; fourth distally; on inner margin; none with setulose ornamentation. Mandible endopod 2-segmented. Mandible endopod 1 with lobe; bearing seta; four; distally inserted; without spinules. Mandible endopod 2 without lobe; bearing setae; nine elements; distally inserted; with spinules; as a row; double. Mandible exopod 4-segmented. Mandible exopod 1 with seta; one element; distally; on inner margin. Mandible exopod 2 with seta; one element; distally; on inner side. Mandible exopod 3 with seta; one element; distally; on inner side. Mandible exopod 4 with setae; three elements; on terminal region. **Maxillule**. Birramous. Maxillule 3-segmented. Maxillule praecoxa with praecoxal arthrite; bearing spines; fifteen elements; ten marginally; plus, five sub-marginally; with spinules; as a patch; on sub-marginal surface. Maxillule coxa with coxal epipodite; with conspicuous outer lobe; bearing setae; nine elements; with coxal endite; elongated (larger or equal width-length ratio 2:1); bearing setae; four elements. Maxillule basis with basal endite; double; first proximal; elongated (larger width-length ratio 2:1; separated from basis; with setae; four elements; distally inserted; second distal; fused to basis; not elongated (lesser width-length ratio 2:1); with setae; four elements; distally inserted; with setules; as a row; on inner side; basal exite present; with setae; one element; on outer surface. Maxillule endopod 1-segmented. Endopod 1 bilobate; first proximal; with setae; three elements; second distal; with setae; five elements. Maxillule exopod 1-segmented. Exopod 1 with setae; six elements; with setules; as a row; on inner side; spinules present; on anterior surface. **Maxilla**. Uniramous. Maxilla 5-segmented. Maxilla praecoxa separated from the coxa; completely; with praecoxal endite; double; first elongated endite (larger or equal width length ratio 2:1); proximally inserted; with seta; straight, or plumose; 1 straight; 4 plumose; without spine; without spinules; without setule; second elongated endite (larger or equal width length ratio 2:1); distally inserted; with seta; plumose; 3 plumose; without spine; without spinules; without setule; absence of outer seta. Maxilla coxa with coxal endite; double; first elongated endite (larger or equal width); proximally inserted; with seta; plumose; 3 plumose; without spine; without spinules; with setules; as a row; on proximal margin; second elongated endite (larger or equal width); distally inserted; with seta; plumose; 3 plumose; without spine; without spinules; with setules; as a row; on proximal margin; absence of outer seta. Maxilla basis with basal endite; single; elongated (larger or equal width-length ratio 2:1); with seta; plumose; 3 plumose; without spinules; absence of outer seta. Maxilla endopod 2-segmented. Endopod 1 with seta; 2 plumose; without spine; without spinules; without setules. Maxilla endopod 2 with seta; 2 plumose; without spine; without spinules; without setules. **Maxilliped**. Uniramous; Maxilliped 8-segmented. Maxilliped praecoxa fused to coxa; incompletely; distinct internally; with praecoxal endite; not elongated (lesser width-length ratio 2:1); distally inserted; with seta; 1 straight; with spinules; as a row; single; on basal surface; without setules. Maxilliped coxa with coxal endite; three coxal endite; first elongated (larger or equal width); proximally inserted; with seta; 2 plumose; with spinules; as a patch; single; on apical surface; without setules; second not elongated (lesser width-length ratio 2:1); medially inserted; with seta; 3 plumose; with spinules; as a row; single; on medial surface; without setules; third elongated (larger or equal width length ratio 2:1); distally inserted; with seta; 3 plumose; none reaching to beyond of the basis; with spinules; as a row; single; on basal surface; without setules; with lobe; not prominence; at inner distal angle; ornamented; with spinules; continuously on margin. Maxilliped basis without basal endite; with seta; 3 plumose; with spinules; as a row; single; on medial surface; without setules. Endopod 5-segmented. Endopod 1 with seta; 2 plumose; on inner surface. Endopod 2 with seta; 3 plumose; on inner surface. Endopod 3 with seta; 2 plumose; on inner surface. Endopod 4 with seta; 2 plumose; on inner surface. Endopod 5 with seta; 4 plumose; on inner surface, or on outer surface; outer seta absent.

##### Swimming legs features

**First swimming legs.** Symmetrical; biramous. First swimming legs intercoxal plate without seta. First swimming legs praecoxa absent. First swimming legs coxa with seta; one; straight; distally inserted; on inner surface; surpassing to first endopodal segment; with setules; two group; as a patch; on inner margin; and as a row; double; on anterior surface; outerly; without spinules; without spine. First swimming legs basis without seta; with setules; as a patch; single; on outer surface; without spinules; without spine. First swimming legs endopod 2-segmented. Endopod 1 with seta; straight; restricted; to inner surface; one element; without spine; with setules; as a row; single; continuously; on outer surface; without spinules; absence of Schmeil’s organ. Endopod 2 with seta; unrestricted; three on inner surface; one on outer surface; two on distal surface; straight; without spine; with setules; as a row; single; continuously; on outer surface; without spinules; absence of Schmeil’s organ. Endopod 3 absence. First swimming legs exopod 1 with seta; restricted; 1 on inner surface; with spine; 1; stout; smaller than original segment; serrated; on inner side; continuously; with setules; as a row; double; as a row; innerly, or outerly. First swimming legs exopod 2 with seta; restricted; 1 on inner surface; straight; without spine; with setules; as a row; single; continuously; on inner margin, or on outer margin; without spinules. First swimming legs exopod 3 with setule; as a row; single; continuously; on outer surface; without spinules; with seta; unrestricted; 2 on inner surface; 2 on terminal surface; with spine; 2; unequal size; first no longer 2x than origin segment; stout; serrated; on inner side, or on outer side; equally; second longer 3x than origin segment; slender; serrated; on outer side; with ornamentation on non-serrated side; by setules. **Second swimming legs**. Symmetrical; Second swimming legs biramous. Second swimming legs intercoxal plate without seta. Second swimming legs praecoxa present; located laterally. Second swimming legs coxa with seta; straight; distally inserted; on inner surface; surpassing to basal segment; without setules; without spinules; without spine. Second swimming legs basis without seta; without setules; without spinules; without spine. Second swimming legs endopod 3-segmented. Endopod 1 with seta; straight; restricted; one on inner surface; without spine; with setules; as a row; single; continuously; on outer surface; without spinules; absence of Schmeil’s organ. Endopod 2 with seta; straight; unrestricted; two on inner surface; without spine; with setules; as a row; single; continuously; on outer side; without spinules; presence of Schmeil’s organ; on posterior surface. Endopod 3 with seta; straight; unrestricted; three on inner surface; two on outer surface; two on distal surface; without spine; without setules; with spinules; as a row; double; distally inserted; at anterior surface; absence of Schmeil’s organ. Second swimming legs exopod 1 with seta; restricted; one on inner surface; with spine; 1; stout; not reaching to distal-third of the exopod 2; serrated; on inner side, or on outer side; with setules; as a row; single; continuously; on inner side; without spinules; absence of Schmeil’s organ.

Exopod 2 with seta; unrestricted; one on inner surface; with spine; 1; stout; not surpassing the exopod 3; serrated; on inner side, or on outer side; with setules; as a row; single; continuously; on inner surface; without spinules; absence of Schmeil’s organ. Exopod 3 with seta; plurimarginal; three on inner surface; two on terminal surface; with spine; 2; unequal size; first no longer 2x than origin segment; stout; serrated; on inner side, or on outer side; equally; second longer 2x than origin segment; slender; serrated; on outer side; with ornamentation on non-serrated side; of setules; setules on outer surface; as a row; single; continuously; on inner surface; with spinules; as a row; single; distally inserted; at anterior surface; absence of Schmeil’s organ. **Third swimming legs**. Symmetrical; Third swimming legs biramous. Third swimming legs intercoxal plate without seta. Third swimming legs praecoxa present; not laterally located. Third swimming legs coxa with seta; straight; distally inserted; on inner surface; surpassing to first endopodal segment; without setules; without spinules; without spine. Third swimming legs basis without seta; without setules; without spinules; without spine. Third swimming legs endopod 3-segmented. Endopod 1 with seta; restricted; one on inner surface; without spine; without setules; without spinules; absence of Schmeil’s organ. Endopod 2 with seta; restricted; two on inner surface; straight; without spine; without setules; without spinules; absence of Schmeil’s organ. Endopod 3 with seta; straight; plurimarginal; two on inner surface; two on outer surface; three on terminal surface; without spine; without setules; with spinules; as a row; distally inserted; double; at anterior surface; absence of Schmeil’s organ. Third swimming legs exopod 1 with seta; restricted; straight; one on inner surface; with spine; 1; stout; not reaching to the distal-third of the exopod 2; serrated; equally; on inner surface, or on outer surface; with setules; as a row; single; continuously; on inner surface; without spinules; absence of Schmeil’s organ. Exopod 2 with seta; straight; restricted; one on inner surface; with spine; 1; stout; not reaching out to exopod 3; serrated; on inner side, or on outer side; equally; with setules; as a row; single; continuously; on inner side; without spinules; absence of Schmeil’s organ. Exopod 3 without setules; with spinules; as a row; single; distally inserted; at anterior surface; with seta; straight; unrestricted; three on inner surface; two on terminal surface; with spine; 2; unequal size; first no longer 2x than origin segment; stout; serrated; on inner side, or on outer side; equally; second longer 2x than origin segment; slender; serrated; on outer side; with ornamentation on non-serrated side; of setules; absence of Schmeil’s organ. **Fourth swimming legs**. Symmetrical; biramous. Intercoxal plate without sensilla. Praecoxa present. Coxa with seta; distally inserted; on inner margin; reaching out to endopod 1; without spinules; setules absent. Basis with seta; one; medially inserted; on posterior surface; smaller than the original segment; without setules; without spinules; without spine. Fourth swimming legs endopod 3-segmented. Endopod 1 with seta; one; restricted; on inner surface; without spine; without setules; without spinules; absence of Schmeil’s organ. Endopod 2 with seta; restricted; two on inner side; without spine; with setules; as a row; single; continuously; on outer surface; without spinules; absence of Schmeil’s organ. Endopod 3 with seta; unrestricted; two on inner surface; two on outer surface; three on distal surface; without spine; without setules; with spinules; as a row; double; distally inserted; at anterior surface; absence of Schmeil’s organ. Fourth swimming legs exopod 1 with seta; restricted; one on inner surface; with spine; 1; stout; not reaching out to distal-third of the exopod 2; serrated; on inner side, or on outer side; equally; with setules; as a row; single; continuously; on inner surface; without spinules; absence of Schmeil’s organ. Exopod 2 with seta; restricted; one on inner surface; with spine; 1; stout; not reaching the end of exopod 3; serrated; on inner side, or on outer side; equally; with setules; as a row; single; continuously; on inner surface; without spinules; absence of Schmeil’s organ. Exopod 3 without setules; with spinules; as a row; single; distally inserted; at anterior surface; with seta; unrestricted; three on inner surface; two on distal surface; with spine; 2; unequal size; first no longer 2x than origin segment; stout; serrated; on inner side, or on outer side; equally; second longer 2x than origin segment; slender; serrated; on outer side; without ornamentation on non-serrated side; absence of Schmeil’s organ.

##### Fifth swimming legs features

Asymmetrical. Fifth swimming leg intercoxal plate with length equal or greater than width on 1.5x; with regular proximal margin; continuous to; the anterior margin of the left coxa; posterior sensilla on the right lateral absent. **Fifth left swimming leg**. Fifth left swimming leg biramous; leg surpassing first right exopod segment. Fifth left swimming leg praecoxa present; developed; separated from the coxae; without ornamentation. Fifth left swimming leg coxa concave inner side; without teeth-like structures; with process; conical; on posterior surface; outer side; distally inserted; projecting over basis; reaching to the proximal surface; with sensilla; stout; triangular; at apex; longer 2x than insertion basis; without swelling; without seta; without spinules. Fifth left swimming leg basis sub-cylindrical; unequal size between inner and outer side; shorter outer than inner side; with convex inner side; rounded internal proximal expansion absent; without outgrowth; without groove; absence of protuberance; with seta; outerly inserted; no longer 2x than origin segment; absence of minutely granular. Fifth left swimming leg endopod segments 1 and 2 fused; segments 2 and 3 fused; 1-segmented; stout; separated from the basis; ornamented; on inner side; with spinules; more than four elements; as a row; terminally; row of setules absent; without seta. Fifth left swimming leg exopod segments 1 and 2 separated; segments 2 and 3 fused; 2-segmented; stout; separated from the basis. Fifth left swimming leg exopod 1 sub-triangular; longer than broad; equal size between inner and outer side; rectilinear inner side; convex outer side; without swelling; without marginal extension; without process; with lobe; single; semicircular; medially inserted; on inner side; covered; by setules; without outer spine; absence seta. Fifth left swimming leg exopod 2 digitiform; longer than broad; equal size between inner and outer side; disform inner side; with rectilinear outer side; setulose pad present; not prominently rounded; medially; on inner side; inflated medial region absent; distal process present; digitiform; denticulate; not bicuspidate; with transverse row of denticles; none oblique row of 5 denticles; at anterior surface; not innerly directed; with seta; spiniform; not ornamented by spinules; not surpassing the distal-point of the segment; without outer spine; terminal claw absent.

##### Fifth right swimming leg

Biramous. Fifth right swimming leg praecoxa absent. Fifth right swimming leg coxa convex inner side; without teeth-like structures; with process; conical; distally inserted; on posterior surface; closest to the outer rim; projecting over basis; beyond the first third; until the medial surface; without triangular protuberance innerly; with sensilla; stout; at nadir; no longer 2x than basal insertion; without marginal extension; without seta; without spinules. Fifth right swimming leg basis trapezoidal; unequal size between inner and outer side; shorter outer than inner side; concave inner side; tumescence absent; with protuberance; single; triangular; on inner side; medially; not ornamented; absence of distinct minutely granular; additional inner process present; triangular; medially; without posterior groove; with seta; outerly inserted; on anterior surface; no longer 2x than origin segment; posterior protrusion present; distal process absent. Fifth right swimming leg with endopodite present; separated from the basis; on anterior surface; ancestral segments 1 and 2 fused; ancestral segments 2 and 3 fused; 1-segmented; stout; ornamented; with spinules; as a row; on inner side; terminally, or sub-terminally; without seta. Fifth right swimming leg exopod segments 1 and 2 separated; segments 2 and 3 fused; 2-segmented; stout; separated from the basis. Fifth right swimming leg exopod 1 trapezium; broader than long; nearly 1.25 times; unequal size between both sides; shorter inner than outer side; convex inner side; concave outer side; with marginal extension; acute; distally inserted; at outer rim; spinules absent; with process; triangular; rectilinear; sharp tip; sclerotized; without ornamentation; distally inserted; at marginal surface; projecting over next segment; without outer spine; without seta; internal prominence absent; lamella on posterior surface absent. Fifth right swimming leg exopod 2 cylindrical; longer than broad; nearly 2 times; equal size between both sides; disform inner side; concave outer side; without posterior proximal swelling; inner-posterior process absent; without marginal expansion; curved ridge on distal posterior surface present; chitinous knobs present; with 1–2 posteriorly; with outer spine; inserted sub-distally; rectilinear; ornamented innerly; by spinules; as a row; not ornamented outerly; sharp tip; without apparent curve; lesser than the length of the exopod 2; until to 2 times its size; 1.5x; sensilla absent; terminal claw present; equal or longer 1.5 times than insertion segment; sclerotized; arched; inward; with conspicuous curve; medially; ornamented innerly; by spinules; as a row; partially on extension; medially, or distally; not ornamented outerly; sharp tip; curved tip; outwards; without medial constriction; hyaline process absent.

##### FEMALE

Body longer and wider than male; Female body 1693 micrometers excluding caudal setae. Widest at first metasome segment. Distal margin of the prosomal segments without one line of setules at posterior margin. Prosome segments with spinules at least at one prosomal segment. Fourth metasome segment presence of dorsal protuberance; rounded; inserted distally; without posterior process; without anterior process; fourth metasome segment without proximal sensillae present. Fourth and fifth metasome segments fused; partially; on dorsal surface. Limit between fourth and fifth metasome segments ornamented; with spinules; as a patch; same size; partially over limit; dorsally. **Fifth metasome segment**. Fifth metasome segment without sensilla; with epimeral plates. Epimeral plates asymmetrical. Right epimeral plates prominent, as projections; thinner than the left; one posterior-dorsally directed; not reaching half length of the genital segment; with sensilla at the apex; dorsal-posterior sensilla present; slender; without ornamentation. Left epimeral plate with sensilla on the apex; with expansion; semicircular; on posterior surface; dorsally; with sensilla; at tip.

##### Urosome

3-segmented. **Genital double-somite**. Asymmetrical in dorsal view; longer than broad; longer than other urosomites combined; dorsal suture at mid-length absent; not covered by spinules; with swelling; rounded; unequal size; greater left than right; anteriorly; with sensillae; on both sides; one; stout; with robust apex; at left lateral; not on lobular base; anteriorly; one; stout; at right rim; not on lobular base; anteriorly; with robust apex; of unequal size between then; left bigger than right; lateral protuberance absent; with right posterior rim expanded; over next segment; without slender sensilla on each posterior rim; without posterior-dorsal process. Genital double-somite opercular pad present; broader than longer; symmetrical; development laterally; expanded posteriorly; covering partially; double gonoporal slit; located ventrally; with arthrodial membrane; inserted anteriorly; post-genital process absent; disto-ventral tumescence absent; ventral vertical folds absent; dorsal sensilla absent. Second urosome segment without ventral fusion to anal segment; right distal process absent. Caudal rami patch of setules on outer surface absent; patch of spinules on outer surface absent.

##### Oral appendices feature

Rostrum basal process absent. **Antennules**. Symmetrical. Right antennule surpassing to genital double-segment; extending beyond caudal rami. Right antennule exceeding the caudal setae. Right antennule ornamentation pattern equals to male left antennule; fully.

##### Fifth swimming legs

Symmetrical; Fifth swimming legs biramous. Fifth swimming legs intercoxal plate longer than wide; separated from the legs. Fifth swimming legs praecoxa without sclerite praecoxal. Fifth swimming legs coxa with process; conical; at the outer rim; distally; sensilla present; stout; at apex; projecting over basal segment; longer 2x than basal insertion; marginal extension present; triangular; at inner rim; distally inserted; without swelling; without seta; without spinules. Fifth swimming legs basis sub-triangular; unequal size between inner and outer sides; shorter inner than outer side; with convex inner side; without proximal inner outgrowth; without groove; without distal extension; with seta; outerly inserted; on anterior surface; no longer 2x than origin segment. Fifth swimming legs endopod segments 1 and 2 fused; segments 2 and 3 fused; 1-segmented; stout; separated from the basis; absent discontinuity cuticle; with spinules; as a row; single; oblique; sub-terminally; at anterior surface; with seta; double; one medially; on posterior surface; rectilinear; one distally; on posterior surface; rectilinear; of unequal size; distal seta longer than medial seta. Fifth swimming legs exopod segments 1 and 2 separated; segments 2 and 3 separated; 3-segmented; separated from the basis. Fifth swimming legs exopod 1 sub-cylindrical; longer than wide; longer or equal than 2 times; with unequal size between inner and outer side; shorter inner than outer side; with convex inner side; with rectilinear outer side; without swelling; without marginal extension; without posterior process; without spine; without seta. Fifth swimming legs exopod 2 sub-cylindrical; longer than broad; longer or equal than 2 times; without swelling; without marginal extension; without process; without lobe; with spine; inserted laterally; rectilinear; without ornamentation; sharp tip; equal size or larger than next segment; without seta. Fifth swimming legs exopod 3 cylindrical; longer than wide; without swelling; without process; without lobe; without spine; with seta; double; inserted terminally; unequal size between them; outer seta smaller than inner; nearly 3 times; outer seta not ornamented by setules; without ornamentation; presence of terminal claw; sclerotized; arched; internally directed; concave inner side; with ornamentation; of denticles; as a row; on surface partially; at medial region; rectilinear outer side; with ornamentation; of denticles; as a row; on surface partially; at medial region; blunt tip; 6 times longer than origin segment.

##### Distribution records

###### ARGENTINA

**Corrientes**: Leconte Lake (Dussart, 1979).

##### Habitat

Habitat in freshwaters: lake.

##### Remarks

The hypothesis of Dussart (1979) was presented from lacustrine organisms of Argentina, without the specification of type-material or biological collection. The organisms were summarily described and illustrated through male fifth swimming legs posteriorly, right antennule partially, and female last metasome segments. In this thesis we examine topotypes collected by I. de Orellana and located in the MNHN, from which morphological characteristics are added to the specimens.

In the original description is indicated the fusion between the female fourth and fifth metasome segments laterally, with the presence of multiple spinules row on the suture remaining dorsally. A female fifth swimming legs endopod 1-segmented, and basis with outer seta reaching to middle-point exopod 1 was evidenced also. Throughout our examinations we were able to corroborate these conditions, except for the arrangement of the spinules on the remaining dorsal suture, apparently arranged as multiple patches. Additionally, we verified the existence of dorsal sensillae on fifth metasome segment, and medial-posterior sensilla on each side of the segment.

Perbiche-Neves *et al*. (2015) when considering the junior synonym species of *Notodiaptomus spiniger* (previously discussed here as *D.* (s.l.) *Spiniger* truly), did not agree with this diagnosis. In the redescription of the Prata River Basin organisms, the female limit between fourth and fifth metasome segment was separated completely and ornamented with multiple spinules row discontinuity (Perbiche-Neves *et al*., 2015). In addition, any mention of the presence of dorsal sensillae on fifth metasome segment was noted by authors. In contrast, Perbiche-Neves *et al*. (2015) found the same condition for the female fifth swimming legs completely to what we verified.

New divergence for the hypothesis of synonymization between *N. orellanai* and *D.* (s.l.) *spiniger* have also been present here for conditions of the male fifth swimming legs. We corroborate Dussart (1979) for male fifth left swimming leg exopod 1 with spiniform seta not surpassing to distal-point original segment, and fifth right leg coxa with conical process projecting over basis beyond first third until medial surface posteriorly. Although the spiniform seta illustrated in Perbiche-Neves *et al*. (2015) also has the same condition, the structure is notably smaller than that present during our examinations and that noted by Dussart (1979, fig. 3), likewise for the structure of the male fifth right leg thigh.

However, we also found original inconsistencies between the organisms examined here and Dussart (1979). Unfortunately, the condition mentioned for a male fifth left swimming leg exopod 1 and 2 fused partially is unobservable. During our examinations it was possible to notice the integral separation of these segments, which, as well as for the organisms of Perbiche-Neves *et al*. (2015), have exopod 2 with denticulate digitiform distal process. All other morphological conditions for the male fifth right swimming leg were present and are identical to those recorded in Perbiche-Neves *et al*. (2015).

For the right antennule of the male of the species is described actual segments 10, 11, 13, 15, and 16 with the presence of spiniform extension, treated here as modified seta for the first three, and spinous process for the last segments. It is interesting to note that although *N. orellanai* and *D.* (s.l.) *spiniger* are treated as synonymous groupings, for the former the presence of the spinous process of segment 16 is originally indicated. This is distinct from the original description of *D.* (s.l.) *spiniger* (Brian, 1926) and its redescription (Perbiche-Neves *et al*., 2015), where the condition is not clearly recorded.

Among the diagnostic attributes highlighted in Dussart (1979) the ornamentation on the male fifth metasoma segment diverges from the specimens of *D.* (s.l.) *spiniger* evidently. However, this seems to be the only distinct morphological condition among the species, because when we observe in the laboratory the fulfillment of the antennulae of *D.* (s.l.) *spiniger*, we find that this does not differ from the one originally described for *N. orellanai.* Unfortunately, the last differential attribute of Dussart (1979) also does not represent a consistent and valid characteristic, since the presence or absence of antennule actual segment 20 with spinous process (“dentiform extension” originally) is intraspecific variation recorded among various organisms in *Notodiaptomus*.

Finally, we consider the morphology indicated by Dussart to distinguish *N. orellanai* inconclusive for the maintenance of the validity of his hypothesis. Although the ornamentation for female limit fifth and fourth metasome segments is divergent, the other morphological characteristics verified for the compared species are completely similar or equal, namely as in: *D.* (s.l.) *spiniger* (Brian, 1926; Perbiche-Neves *et al*., 2015), *Diaptomus birabeni* (Brehm, 1957; Ringuelet, 1958), *N. incompositus* (Brian, 1925; Santos-Silva *et al*., 2015; Perbiche-Neves *et al*., 2015), *N. nordestinus* (Wright, 1935; Santos-Silva *et al*., 2015), and *N. maracaibensis* (Kiefer, 1954; Fuentes-Reinés *et al*., 2021). Based on the attributes defined in Kiefer (1936; 1956) for *Notodiaptomus*, the following conditions diverged in *N. orellanai*: (1) male fifth right swimming leg exopod 1 broader than long; and (2) male left swimming leg exopod 2 with spiniform seta reaching to distal-point of the original segment. Variable for the type-species, the conditions present were mainly: (1) male limit between fourth and fifth metasome segments fused partially; (2) male fifth left swimming leg exopod 1 with double inner lobe; (3) male fifth right swimming leg basis without posterior groove, and ornamentation; (4) female fifth swimming legs endopod without cuticle discontinuity; and (5) female fifth swimming legs basis with outer seta reaching middle-point exopod 1.

#### Notodiaptomus paraensis Dussart & Robertson, 1984

##### Synonymy

*Notodiaptomus paraensis* Dussart & Robertson, 1984: 389–394, figs. 1–3; Reid, 1987: 378; Magalhães *et al*., 1988: 271; Santos-Silva *et al*., 1989: 726, 728, figs. 69–93; Rocha *et al*., 1995:156; Santos-Silva, 1998: 211; Santos-Silva, 2008: 33, figs. 7; Perbiche-Neves *et al*., 2020: 685-686, key to the Neotropical diaptomid, fig. 21.10 A. *Notodiaptomus* (*Wrightius*) *paraensis*; Dussart, 1985a: 210, fig. 7.

##### Type locality

Brazil: Pará State, Curuá-Una Reservoir (02°48’38“S, 54°18’55“W) (“Santarém South Station”).

##### Type material

Holotype: 1 male, entire in alcohol, existent in INPA. Paratype: 1 male, and 1 female, entire in alcohol, reported for the Museum National d’Historie Naturalle collection in Paris, France, code not specified.

##### Material examined

Non-type material: 03 males, and 02 females, entire in alcohol, from the Curuá-Una Reservoir, Pará, no collector, VIII.1978, stored in Collection of zooplankton sample of the Plankton Laboratory, INPA, Brazil. 1 male (INPA-COP045, slides a-h) and 1 female (INPA-COP046, slides a-h) were selected to be dissection on eight slides each and deposited in the Zoological Collection of the INPA, Brazil. Material indicated on original description is in poor-condition, its examination was not feasible.

##### Diagnosis

**(1)** male fifth right swimming leg coxa with slender sensilla no longer 2x than distal conical process on posterior surface; **(2)** male fifth right swimming leg basis without posterior groove; **(3)** male fifth right swimming leg exopod 1 broader than long 1.25x nearly; **(4)** male fifth right swimming leg exopod 1 with acute outer distal extension; **(5)** male fifth right swimming leg basis with double denticule protuberance on inner surface distally; **(6)** male fifth right swimming leg exopod 2 with outer spine inserted medially, equal length than origin segment; **(7)** female fourth and fifth metasome segments fused partially on lateral surface; **(8)** female left epimeral plate with posterior semicircular expansion dorsally, with sensilla at tip; **(9)** female second urosome segment with ventral fusion to anal segment; **(10)** female second urosome segment with elongated spiniform process on right side distally; **(11)** female fifth swimming legs endopod 1-segmented; **(12)** female fifth swimming legs exopod 2 with lateral seta smaller than next segment; **(13)** female caudal rami with setules patch on outer surface.

##### Redescription

###### MALE

Body 1105 micrometers excluding caudal setae. Male body smaller and slenderer than female. Nerve axons myelinated. Prosome 6-segmented; widest at first metasome segment; with one line of setules at posterior margin; without spinules at segments. Cephalosome anterior margin sub-triangular; without dorsal suture; separate from first metasome segment. First metasome segment without sensilla. Second metasome segment without sensilla. Third metasome segment without sensillae; non-ornamented posterior margin. Fourth metasome segment without sensillae; separated from the fifth metasome. Limit between fourth and fifth metasome segments without ornamentation. Fifth metasome segment with sensilla; 2 dorsally; Fifth metasome segment equal size; Fifth metasome segment without ornamentation; Fifth metasome segment without dorsal conical process; with epimeral plates. Epimeral plates symmetrical. Right epimeral plates reduced, as rounded distal corner segment limit; with sensilla; one at the apex of projection and other medially; without ornamentation.

##### Urosome

5-segmented; Urosome 5 - free segments. Genital somite asymmetrical in dorsal view; with single aperture; located on left side; ventrolaterally on posterior rim; with sensillae; on both sides; one; at left lateral; posteriorly; one; at right rim; posteriorly; of equal size between then. Third urosome segment without spinules; without external seta. Fourth urosome segment without spinules; without sub-conical blunt dorsal-lateral process. Anal segment absence of dorsal sensillae; presence of operculum; convex; covering the anal aperture fully. Caudal rami symmetrical; separated from anal segment; longer than wide; with setules; continuous on; inner side; each ramus bearing 6 caudal setae; 5 marginals; plumose; and 1 internal dorsally; straight; not reticulated main axis; outermost seta with outer spiniform process absent.

##### Oral appendices feature

Rostrum asymmetrical; separated from dorsal cephalic shield; by complete suture; sensillae present; one pair; anteriorly inserted on surface tegument; with rostral filament; double; paired; extended; into point; with basal process; in ventral view, rounded on left side; without a smaller basal expansion on the right side.

##### Antennules

Asymmetrical. **Right antennules**. Uniramous; right antennule surpassing to genital segment; right antennule not extending beyond caudal rami.

Right antennule ancestral segment I and II separated. Ancestral segment II and III fused. Ancestral segment III and IV fused. Ancestral segment IV and V separated. Ancestral segment V and VI separated. Ancestral segment VI and VII separated. Ancestral segment VII and VIII separated. Ancestral segment VIII and IX separated. Ancestral segment IX and X separated. Ancestral segment X and XI separated. Ancestral segment XI and XII separated. Ancestral segment XII and XIII separated. Ancestral segment XIII and XIV separated. Ancestral segment XIV and XV separated. Ancestral segment XV and XVI separated. Ancestral segment XVI and XVII separated. Ancestral segment XVII and XVIII separated. Ancestral segment XVIII and XIX separated. Ancestral segment XIX and XX separated. Ancestral segment XX and XXI separated. Ancestral segment XXI and XXII fused. Ancestral segment XXII and XXIII fused. Ancestral segment XXIII and XXIV separated. Ancestral segment XXIV and XXV fused. Ancestral segment XXV and XXVI separated. Ancestral segment XXVI and XXVII separated. Ancestral segment XXVII and XXVIII fused.

Right antennule actual 22-segmented; geniculated; between the segment 18 and segment 19; with swollen and modified region; formed by 5 segments; between 13 and 17 segments. Actual segment 1 with seta; one element; straight; none larger than segment; without spinules; without vestigial seta; without conical seta; without modified seta; without spinous process; with aesthetasc; one element. Actual segment 2 with seta; three elements; of unequal size; straight; none larger than segment; without spinules; with vestigial seta; one element; without conical seta; without modified seta; without spinous process; with aesthetasc; one element. Actual segment 3 with seta; one element; one larger than segment; surpassing to distal margin; beyond three sequential segments; straight; blunt apex; without spinules; with vestigial seta; one element; without conical seta; without modified seta; without spinous process; with aesthetasc. Actual segment 4 with seta; one element; one larger than segment; surpassing to distal margin; straight; not beyond three sequential segments; without spinules; without vestigial seta; without conical seta; without modified seta; without spinous process; without aesthetasc. Actual segment 5 with seta; one element; straight; one larger than segment; surpassing to distal margin; not beyond three sequential segments; without spinules; with vestigial seta; one element; without conical seta; without modified seta; without spinous process; with aesthetasc; one element. Actual segment 6 with seta; one element; none larger than segment; straight; without spinules; without vestigial seta; without conical seta; without modified seta; without spinous process; without aesthetasc. Actual segment 7 with seta; one element; straight; one larger than segment; surpassing to distal margin; beyond three sequential segments; blunt apex; without spinules; without vestigial seta; without conical seta; without modified seta; without spinous process; with aesthetasc; one element. Actual segment 8 with seta; one element; straight; none larger than segment; without spinules; without vestigial seta; with conical seta; one element; not reaching to middle-point of the sequent segment; without modified seta; without spinous process; without aesthetasc. Actual segment 9 with seta; two elements; of unequal size; straight; one larger than segment; surpassing to distal margin; beyond three sequential segments; blunt apex; without spinules; without vestigial seta; without conical seta; without modified seta; without spinous process; with aesthetasc; one element. Actual segment 10 with seta; one element; straight; none larger than segment; without spinules; without vestigial seta; without conical seta; with modified seta; presenting blunt apex; slender form; surpassing to distal margin; beyond of the sequential segment; parallel to antennule direction; without spinous process; without aesthetasc. Actual segment 11 with seta; one element; straight; one larger than segment; surpassing to distal margin; not beyond three sequential segments; without spinules; without vestigial seta; without conical seta; with modified seta; slender form; presenting blunt apex; surpassing to distal margin; beyond of the sequential segment; parallel to antennule direction; shorter length than homologous of actual segment 13; without spinous process; without aesthetasc. Actual segment 12 with seta; one element; straight; one larger than segment; surpassing to distal margin; not beyond three sequential segments; without spinules; without vestigial seta; with conical seta; one element; not smaller than to segment 8; without modified seta; without spinous process; with aesthetasc; one element; absent internal perpendicular fission. Actual segment 13 with seta; one element; straight; one larger than segment; surpassing to distal margin; not beyond three sequential segments; without spinules; without vestigial seta; without conical seta; with modified seta; stout form; surpassing to distal margin; to the middle-point of the sequence segment; perpendicular to antennule direction; presenting bifid apex; without spinous process; with aesthetasc; one element. Actual segment 14 with seta; two elements; of unequal size; straight; one larger than segment; surpassing to distal margin; beyond three sequential segments; blunt apex; without spinules; without vestigial seta; without conical seta; without modified seta; without spinous process; with aesthetasc; one element. Actual segment 15 with seta; two elements; of unequal size; straight; not bifidform; none larger than segment; without spinules; without vestigial seta; without conical seta; without modified seta; with spinous process; on outer margin; surpassing distal margin; with aesthetasc; one element. Actual segment 16 with seta; two elements; of unequal size; plumose; one larger than segment; surpassing distal margin; not beyond three sequential segments; not bifidform; without spinules; without vestigial seta; without conical seta; without modified seta; with spinous process; on outer margin; surpassing distal margin; unequal size to process on preceding segment; with aesthetasc; one element. Actual segment 17 with seta; two elements; of unequal size; straight; none larger than segment; bifidform; without spinules; without vestigial seta; without conical seta; with modified seta; one element; stout form; surpassing to distal margin; not beyond of the sequential segment; parallel to antennule direction; without spinous process; without aesthetasc. Actual segment 18 with seta; two elements; of equal size; straight; none larger than segment; without spinules; without vestigial seta; without conical seta; with modified seta; one element; stout form; surpassing distal margin; parallel to antennule direction; without spinous process; without aesthetasc. Actual segment 19 with seta; two elements; of unequal size; plumose; none larger than segment; without spinules; without vestigial seta; without conical seta; with modified seta; two elements; stout form; at least one bifid form; surpassing distal margin; parallel to antennule direction; without spinous process; with aesthetasc; one element. Actual segment 20 with seta; four elements; of unequal size; straight; one larger than segment; surpassing to distal margin; beyond three sequential segments; without spinules; without vestigial seta; without conical seta; without modified seta; with spinous process; distally; reaching beyond of distal-point segment 21; without aesthetasc. Actual segment 21 with seta; two elements; of equal size; plumose; one larger than segment; surpassing to distal margin; greater 3x than original segment; without spinules; without vestigial seta; without conical seta; without modified seta; without spinous process; without aesthetasc. Actual segment 22 with seta; four elements; of equal size; one larger than segment; plumose; surpassing to distal margin; greater 3x than original segment; without spinules; without vestigial seta; without conical seta; without modified seta; without spinous process; with aesthetasc; one element.

##### Left antennules

Uniramous; Left antennule surpassing to prosome; Left antennule not extending beyond caudal rami. Ancestral segment I and II separated. Ancestral segment II and III fused. Ancestral segment III and IV fused. Ancestral segment IV and V separated. Ancestral segment V and VI separated. Ancestral segment VI and VII separated. Ancestral segment VII and VIII separated. Ancestral segment VIII and IX separated. Ancestral segment IX and X separated. Ancestral segment X and XI separated. Ancestral segment XI and XII separated. Ancestral segment XII and XIII separated. Ancestral segment XIII and XIV separated. Ancestral segment XIV and XV separated. Ancestral segment XV and XVI separated. Ancestral segment XVI and XVII separated. Ancestral segment XVII and XVIII separated. Ancestral segment XVIII and XIX separated. Ancestral segment XIX and XX separated. Ancestral segment XX and XXI separated. Ancestral segment XXI and XXII separated. Ancestral segment XXII and XXIII separated. Ancestral segment XXIII and XXIV separated. Ancestral segment XXIV and XXV separated. Ancestral segment XXV and XXVI separated. Ancestral segment XXVI and XXVII separated. Ancestral segment XXVII and XXVIII fused.

Left antennule actual 25-segmented; not-geniculated. Actual segment 1 with seta; one element; none larger than segment; straight; without vestigial seta; without conical seta; without modified seta; without spinous process; with aesthetasc; one element. Actual segment 2 with seta; three elements; of equal size; none larger than segment; straight; without spinules; with vestigial seta; one element; without conical seta; without modified seta; without spinous process; with aesthetasc; one element. Actual segment 3 with seta; one element; one larger than segment; straight; surpassing to distal margin; beyond three sequential segments; without spinules; with vestigial seta; one element; without conical seta; without modified seta; without spinous process; with aesthetasc. Actual segment 4 with seta; one element; none larger than segment; straight; without spinules; without vestigial seta; without conical seta; without modified seta; without spinous process; without aesthetasc. Actual segment 5 with seta; one element; one larger than segment; straight; surpassing to distal margin; not beyond three sequential segments; without spinules; with vestigial seta; one element; without conical seta; without modified seta; without spinous process; with aesthetasc; one element. Actual segment 6 with seta; one element; none larger than segment; straight; without spinules; without vestigial seta; without conical seta; without modified seta; without spinous process; without aesthetasc. Actual segment 7 with seta; one element; one larger than segment; straight; surpassing to distal margin; beyond three sequential segments; without spinules; without vestigial seta; without conical seta; without modified seta; without spinous process; with aesthetasc; one element. Actual segment 8 with seta; one element; one larger than segment; straight; surpassing distal margin; without spinules; without vestigial seta; with conical seta; without modified seta; without spinous process; without aesthetasc. Actual segment 9 with seta; two elements; of unequal size; one larger than segment; straight; surpassing to distal margin; beyond three sequential segments; without spinules; without vestigial seta; without conical seta; without modified seta; without spinous process; with aesthetasc; one element. Actual segment 10 with seta; one element; none larger than segment; straight; without spinules; without vestigial seta; without conical seta; without modified seta; without spinous process; without aesthetasc. Actual segment 11 with seta; one element; one larger than segment; straight; surpassing to distal margin; beyond three sequential segments; without spinules; without vestigial seta; without conical seta; without modified seta; without spinous process; without aesthetasc. Actual segment 12 with seta; one element; one larger than segment; straight; surpassing distal margin; without spinules; without vestigial seta; with conical seta; without modified seta; without spinous process; with aesthetasc; one element. Actual segment 13 with seta; one element; none elongated; straight; surpassing distal margin; without spinules; without vestigial seta; without conical seta; without modified seta; without spinous process; without aesthetasc. Actual segment 14 with seta; one element; elongated; straight; surpassing to distal margin; beyond three sequential segments; without spinules; without vestigial seta; without conical seta; without modified seta; without spinous process; with aesthetasc; one element. Actual segment 15 with seta; one element; larger than segment; straight; surpassing to distal margin; not beyond three sequential segments; without spinules; without vestigial seta; without conical seta; without modified seta; without spinous process; without aesthetasc. Actual segment 16 with seta; one element; larger than segment; plumose; surpassing to distal margin; not beyond three sequential segments; without spinules; without vestigial seta; without conical seta; without modified seta; without spinous process; with aesthetasc; one element. Actual segment 17 with seta; one element; not larger than segment; straight; without spinules; without vestigial seta; without conical seta; without modified seta; without spinous process; without aesthetasc. Actual segment 18 with seta; one element; larger than segment; straight; surpassing to distal margin; beyond three sequential segments; without spinules; without vestigial seta; without conical seta; without modified seta; without spinous process; without aesthetasc. Actual segment 19 with seta; one element; not larger than segment; straight; surpassing distal margin; without spinules; without vestigial seta; without conical seta; without modified seta; without spinous process; with aesthetasc; one element. Actual segment 20 with seta; one element; not larger than segment; straight; surpassing distal margin; without spinules; without vestigial seta; without conical seta; without modified seta; without spinous process; without aesthetasc. Actual segment 21 with seta; one element; larger than segment; plumose; surpassing to distal margin; beyond three sequential segments; without spinules; without vestigial seta; without conical seta; without modified seta; without spinous process; without aesthetasc. Actual segment 22 with seta; two elements; of unequal size; one of them elongated; plumose; surpassing to distal margin; without spinules; without vestigial seta; without conical seta; without modified seta; without spinous process; without aesthetasc. Actual segment 23 with seta; two elements; of unequal size; one larger than segment; plumose; surpassing to distal margin; greater 3x than original segment; without spinules; without vestigial seta; without conical seta; without modified seta; without spinous process; without aesthetasc. Actual segment 24 with seta; two elements; of equal size; one larger than segment; plumose; surpassing to distal margin; greater 3x than original segment; without spinules; without vestigial seta; without conical seta; without modified seta; without spinous process; without aesthetasc. Actual segment 25 with seta; four elements; of equal size; elongated; plumose; surpassing to distal margin; 4 times larger than segment; without spinules; without vestigial seta; without conical seta; without modified seta; without spinous process; with aesthetasc; one element.

##### Antenna

Biramous. Antenna coxa separated from the basis; bearing seta; 1; on inner surface; at distal corner; reaching to the endopod 1. Antenna basis (fusion) separated from the endopodal segment; bearing seta; 2; on inner surface; at distal corner. Endopodal ancestral segment I and II separated. Ancestral segment II and III fused. Ancestral segment III and IV separated. Antenna endopod actual 3-segmented. Actual segment 1 not bilobate; with seta; two; on inner margin; with spinules; as a row; obliquely; on outer surface; without pore. Actual segment 2 not bilobate; without discontinuity on outer cuticle; inner lobe bearing 9 setae; distally; without spinules. Actual segment 3 outer lobe bearing 8 setae; distally; without spinules. Antenna exopod ancestral segment I and II separated. Ancestral segment II and III fused. Ancestral segment III and IV fused. Ancestral segment IV and V separated. Ancestral segment V and VI separated. Ancestral segment VI and VII separated. Ancestral segment VII and VIII separated. Ancestral segment VIII and IX separated. Ancestral segment IX and X fused. Antenna exopod actual 7-segmented. Actual segment 1 single; elongated (width-length, equal or larger ratio 2:1); with seta; one; at inner surface. Actual segment 2 compound; elongated (larger width-length ratio 2:1); with seta; three; at inner surface. Actual segment 3 single; not elongated (lesser width-length ratio 2:1); with seta; one; at inner surface. Actual segment 4 single; not elongated (lesser width-length ratio 2:1); with seta; one; at inner surface. Actual segment 5 single; not elongated (lesser width-length ratio 2:1); with seta; one; at inner surface. Actual segment 6 single; not elongated (lesser width-length ratio 2:1); with seta; one; at inner surface. Actual segment 7 compound; elongated (larger or equal width-length ratio 2:1); with seta; one; at inner surface; and three; at distal surface.

##### Oral features

**Mandible**. Coxal gnathobase sclerotized; without lobe; presence of cutting blade; with tooth-like prominence; two, distinctly; 1 acute; on caudal margin; and 1 triangular; on sub-caudal margin; without acute projection between the prominences; with additional spinules; as a row; on dorsal surface; with seta; 1; dorsally; on apical surface; without spinules. Mandible palps biramous; comprising the basis; with seta; four; differently inserted; first medially; reaching to beyond the endopod 1; second distally; third distally; fourth distally; on inner margin; none with setulose ornamentation. Mandible endopod 2-segmented. Mandible endopod 1 with lobe; bearing seta; four; distally inserted; without spinules. Mandible endopod 2 without lobe; bearing setae; nine elements; distally inserted; with spinules; as a row; double. Mandible exopod 4-segmented. Mandible exopod 1 with seta; one element; distally; on inner margin. Mandible exopod 2 with seta; one element; distally; on inner side. Mandible exopod 3 with seta; one element; distally; on inner side. Mandible exopod 4 with setae; three elements; on terminal region. **Maxillule**. Birramous. Maxillule 3-segmented. Maxillule praecoxa with praecoxal arthrite; bearing spines; fifteen elements; ten marginally; plus, five sub-marginally; with spinules; as a patch; on sub-marginal surface. Maxillule coxa with coxal epipodite; with conspicuous outer lobe; bearing setae; nine elements; with coxal endite; elongated (larger or equal width-length ratio 2:1); bearing setae; four elements. Maxillule basis with basal endite; double; first proximal; elongated (larger width-length ratio 2:1; separated from basis; with setae; four elements; distally inserted; second distal; fused to basis; not elongated (lesser width-length ratio 2:1); with setae; four elements; distally inserted; with setules; as a row; on inner side; basal exite present; with setae; one element; on outer surface. Maxillule endopod 1-segmented. Endopod 1 bilobate; first proximal; with setae; three elements; second distal; with setae; five elements. Maxillule exopod 1-segmented. Exopod 1 with setae; six elements; with setules; as a row; on inner side; spinules absent. **Maxilla**. Uniramous. Maxilla 5-segmented. Maxilla praecoxa fused to coxa; incompletely; distinct externally; with praecoxal endite; double; first elongated endite (larger or equal width length ratio 2:1); proximally inserted; with seta; straight, or plumose; 1 straight; 4 plumose; with spine; single; without spinules; without setule; second elongated endite (larger or equal width length ratio 2:1); distally inserted; with seta; plumose; 3 plumose; without spine; with spinules; as a row; on distal margin; with setule; as a row; on distal margin; absence of outer seta. Maxilla coxa with coxal endite; double; first elongated endite (larger or equal width); proximally inserted; with seta; plumose; 3 plumose; without spine; without spinules; with setules; as a row; on proximal margin; second elongated endite (larger or equal width); distally inserted; with seta; plumose; 3 plumose; without spine; without spinules; with setules; as a row; on proximal margin; absence of outer seta. Maxilla basis with basal endite; single; elongated (larger or equal width-length ratio 2:1); with seta; plumose; 3 plumose; without spinules; absence of outer seta. Maxilla endopod 2-segmented. Endopod 1 with seta; 2 plumose; without spine; without spinules; without setules. Maxilla endopod 2 with seta; 2 plumose; without spine; without spinules; without setules. **Maxilliped**. Uniramous; Maxilliped 8-segmented. Maxilliped praecoxa fused to coxa; incompletely; distinct internally; with praecoxal endite; not elongated (lesser width-length ratio 2:1); distally inserted; with seta; 1 straight; without spinules; without setules. Maxilliped coxa with coxal endite; three coxal endite; first elongated (larger or equal width); proximally inserted; with seta; 2 plumose; with spinules; as a patch; single; on apical surface; without setules; second not elongated (lesser width-length ratio 2:1); medially inserted; with seta; 3 plumose; with spinules; as a row; single; on medial surface; without setules; third elongated (larger or equal width length ratio 2:1); distally inserted; with seta; 3 plumose; none reaching to beyond of the basis; with spinules; as a row; single; on basal surface; without setules; with lobe; prominence; at inner distal angle; ornamented; with spinules; continuously on margin. Maxilliped basis without basal endite; with seta; 3 plumose; with spinules; as a row; single; on medial surface; with setules; as a row; single; on inner margin. Maxilliped endopod segment 6-segmented. Endopod 1 with seta; 2 plumose; on inner surface. Endopod 2 with seta; 3 plumose; on inner surface. Endopod 3 with seta; 2 plumose; on inner surface. Endopod 4 with seta; 2 plumose; on inner surface. Endopod 5 with seta; 2 plumose; on inner surface, or on outer surface; outer seta absent. Endopod 6 with seta; 4 plumose; on inner surface, or on outer surface.

##### Swimming legs features

**First swimming legs.** Symmetrical; biramous. First swimming legs intercoxal plate without seta. First swimming legs praecoxa absent. First swimming legs coxa with seta; one; plumose; distally inserted; on inner surface; surpassing to basal segment; without setules; without spinules; without spine. First swimming legs basis without seta; with setules; as a patch; single; on outer surface; without spinules; without spine. First swimming legs endopod 2-segmented. Endopod 1 with seta; plumose; restricted; to inner surface; one element; without spine; with setules; as a row; single; continuously; on outer surface; without spinules; absence of Schmeil’s organ. Endopod 2 with seta; unrestricted; three on inner surface; one on outer surface; two on distal surface; straight; without spine; with setules; as a row; single; continuously; on outer surface; without spinules; absence of Schmeil’s organ. Endopod 3 absence. First swimming legs exopod 1 with seta; restricted; 1 on inner surface; with spine; 1; stout; smaller than original segment; serrated; on inner side; continuously; with setules; as a row; single; as a row; innerly. First swimming legs exopod 2 with seta; restricted; 1 on inner surface; straight; without spine; with setules; as a row; single; continuously; on inner margin, or on outer margin; without spinules. First swimming legs exopod 3 with setule; as a row; single; continuously; on outer surface; without spinules; with seta; unrestricted; 2 on inner surface; 2 on terminal surface; with spine; 2; unequal size; first no longer 2x than origin segment; stout; serrated; on inner side, or on outer side; equally; second longer 3x than origin segment; slender; serrated; on outer side; with ornamentation on non-serrated side; by setules. **Second swimming legs**. Symmetrical; Second swimming legs biramous. Second swimming legs intercoxal plate without seta. Second swimming legs praecoxa present; located laterally. Second swimming legs coxa with seta; plumose; distally inserted; on inner surface; surpassing to basal segment; without setules; without spinules; without spine. Second swimming legs basis without seta; without setules; without spinules; without spine. Second swimming legs endopod 3-segmented. Endopod 1 with seta; plumose; restricted; one on inner surface; without spine; with setules; as a row; single; continuously; on outer surface; without spinules; absence of Schmeil’s organ. Endopod 2 with seta; plumose; unrestricted; two on inner surface; without spine; with setules; as a row; single; continuously; on outer side; without spinules; presence of Schmeil’s organ; on posterior surface. Endopod 3 with seta; plumose; unrestricted; three on inner surface; two on outer surface; two on distal surface; without spine; without setules; with spinules; as a row; double; distally inserted; at anterior surface; absence of Schmeil’s organ. Second swimming legs exopod 1 with seta; restricted; one on inner surface; with spine; 1; stout; not reaching to distal-third of the exopod 2; serrated; on inner side, or on outer side; with setules; as a row; single; continuously; on inner side; without spinules; absence of Schmeil’s organ. Exopod 2 with seta; unrestricted; one on inner surface; with spine; 1; stout; not surpassing the exopod 3; serrated; on inner side, or on outer side; with setules; as a row; single; continuously; on inner surface; without spinules; absence of Schmeil’s organ. Exopod 3 with seta; plurimarginal; three on inner surface; two on terminal surface; with spine; 2; unequal size; first no longer 2x than origin segment; stout; serrated; on inner side, or on outer side; equally; second longer 2x than origin segment; slender; serrated; on outer side; with ornamentation on non-serrated side; of setules; setules on outer surface; as a row; single; continuously; on inner surface; with spinules; as a row; single; distally inserted; at anterior surface; absence of Schmeil’s organ. **Third swimming legs**. Symmetrical; Third swimming legs biramous. Third swimming legs intercoxal plate without seta. Third swimming legs praecoxa present; not laterally located. Third swimming legs coxa with seta; straight; distally inserted; on inner surface; surpassing to basal segment; without setules; without spinules; without spine. Third swimming legs basis without seta; without setules; without spinules; without spine. Third swimming legs endopod 3-segmented. Endopod 1 with seta; restricted; one on inner surface; without spine; without setules; without spinules; absence of Schmeil’s organ. Endopod 2 with seta; restricted; two on inner surface; plumose; without spine; without setules; without spinules; absence of Schmeil’s organ. Endopod 3 with seta; straight; plurimarginal; two on inner surface; two on outer surface; three on terminal surface; without spine; without setules; with spinules, or without spinules; as a row; distally inserted; double; at anterior surface; absence of Schmeil’s organ. Third swimming legs exopod 1 with seta; restricted; straight; one on inner surface; with spine; 1; stout; not reaching to the distal-third of the exopod 2; serrated; equally; on inner surface, or on outer surface; with setules; as a row; single; continuously; on inner surface; without spinules; absence of Schmeil’s organ. Exopod 2 with seta; straight; restricted; one on inner surface; with spine; 1; stout; not reaching out to exopod 3; serrated; on inner side, or on outer side; equally; with setules; as a row; single; continuously; on inner side; without spinules; absence of Schmeil’s organ. Exopod 3 without setules; with spinules; as a row; single; distally inserted; at anterior surface; with seta; straight; unrestricted; three on inner surface; two on terminal surface; with spine; 2; unequal size; first no longer 2x than origin segment; stout; serrated; on inner side, or on outer side; equally; second longer 2x than origin segment; slender; serrated; on outer side; with ornamentation on non-serrated side; of setules; absence of Schmeil’s organ. **Fourth swimming legs**. Symmetrical; biramous. Intercoxal plate without sensilla. Praecoxa present. Coxa with seta; distally inserted; on inner margin; reaching out to endopod 1; without spinules; setules absent. Basis with seta; one; medially inserted; on posterior surface; smaller than the original segment; without setules; without spinules; without spine. Fourth swimming legs endopod 3-segmented. Endopod 1 with seta; one; restricted; on inner surface; without spine; without setules; without spinules; absence of Schmeil’s organ. Endopod 2 with seta; restricted; two on inner side; without spine; with setules; as a row; single; continuously; on outer surface; without spinules; absence of Schmeil’s organ. Endopod 3 with seta; unrestricted; two on inner surface; two on outer surface; three on distal surface; without spine; without setules; with spinules; as a row; double; distally inserted; at anterior surface; absence of Schmeil’s organ. Fourth swimming legs exopod 1 with seta; restricted; one on inner surface; with spine; 1; stout; not reaching out to distal-third of the exopod 2; serrated; on inner side, or on outer side; equally; with setules; as a row; single; continuously; on inner surface; without spinules; absence of Schmeil’s organ. Exopod 2 with seta; restricted; one on inner surface; with spine; 1; stout; not reaching the end of exopod 3; serrated; on inner side, or on outer side; equally; with setules; as a row; single; continuously; on inner surface; without spinules; absence of Schmeil’s organ. Exopod 3 without setules; with spinules; as a row; single; distally inserted; at anterior surface; with seta; unrestricted; three on inner surface; two on distal surface; with spine; 2; unequal size; first no longer 2x than origin segment; stout; serrated; on inner side, or on outer side; equally; second longer 2x than origin segment; slender; serrated; on outer side; without ornamentation on non-serrated side; absence of Schmeil’s organ.

##### Fifth swimming legs features

Asymmetrical. Fifth swimming leg intercoxal plate with length not equal or greater than width on 1.5x; with irregular proximal margin; discontinuous to; the anterior margin of the left coxa, or the anterior margin of the right coxa; posterior sensilla on the right lateral absent. **Fifth left swimming leg**. Fifth left swimming leg biramous; leg surpassing first right exopod segment. Fifth left swimming leg praecoxa present; rudimentary; separated from the coxae; without ornamentation. Fifth left swimming leg coxa concave inner side; without teeth-like structures; with process; conical; on posterior surface; outer side; distally inserted; not projecting over basis; with sensilla; stout; triangular; at apex; longer 2x than insertion basis; without swelling; without seta; without spinules. Fifth left swimming leg basis sub-cylindrical; unequal size between inner and outer side; shorter outer than inner side; with concave inner side; rounded internal proximal expansion absent; without outgrowth; with groove; deep; obliquely; on posterior surface; not reaching the endopodal lobe; not ornamented; absence of protuberance; with seta; outerly inserted; no longer 2x than origin segment; absence of minutely granular. Fifth left swimming leg endopod segments 1 and 2 fused; segments 2 and 3 fused; 1-segmented; stout; separated from the basis; ornamented; on inner side; with spinules; more than four elements; as a row; terminally; row of setules absent; without seta. Fifth left swimming leg exopod segments 1 and 2 separated; segments 2 and 3 fused; 2-segmented; stout; separated from the basis. Fifth left swimming leg exopod 1 sub-triangular; longer than broad; equal size between inner and outer side; rectilinear inner side; convex outer side; without swelling; without marginal extension; without process; with lobe; single; semicircular; medially inserted; on inner side; covered; by setules; without outer spine; absence seta. Fifth left swimming leg exopod 2 digitiform; longer than broad; unequal size between inner and outer side; shorter inner than outer side; disform inner side; with convex outer side; setulose pad present; not prominently rounded; proximally; on inner side; inflated medial region absent; distal process present; digitiform; denticulate; not bicuspidate; without transverse row of denticles; none oblique row of 5 denticles; at anterior surface; not innerly directed; with seta; spiniform; not ornamented by spinules; surpassing the distal-point of the segment; without outer spine; terminal claw absent.

##### Fifth right swimming leg

Biramous. Fifth right swimming leg praecoxa present; separated from the coxae; without ornamentation. Fifth right swimming leg coxa convex inner side; without teeth-like structures; with process; conical; distally inserted; on posterior surface; closest to the outer rim; projecting over basis; beyond the first third; until the medial surface; without triangular protuberance innerly; with sensilla; slender; at apex; longer 2x than basal insertion; without marginal extension; without seta; without spinules. Fifth right swimming leg basis trapezoidal; unequal size between inner and outer side; shorter outer than inner side; rectilinear inner side; tumescence present; not inflated; restricted on inner surface; proximally; with protuberance; single; triangular; on inner side; distally; not ornamented; not minutely granular; not covering the element; absence of distinct minutely granular; additional inner process present; triangular; distally; without posterior groove; with seta; outerly inserted; on anterior surface; no longer 2x than origin segment; posterior protrusion present; distal process absent. Fifth right swimming leg with endopodite present; separated from the basis; on anterior surface; ancestral segments 1 and 2 fused; ancestral segments 2 and 3 fused; 1-segmented; stout ornamented; with setules; as a row; on inner side; terminally; without seta. Fifth right swimming leg exopod segments 1 and 2 separated; segments 2 and 3 fused; 2-segmented; stout; separated from the basis. Fifth right swimming leg exopod 1 trapezium; broader than long; nearly 1.25 times; unequal size between both sides; shorter inner than outer side; rectilinear inner side; rectilinear outer side; with marginal extension; acute; distally inserted; at outer rim; spinules absent; with process; triangular; rectilinear; sharp tip; sclerotized; without ornamentation; distally inserted; at marginal surface; projecting over next segment; without outer spine; without seta; internal prominence absent; lamella on posterior surface absent; trapezoidal form; not surpassing to margin. Fifth right swimming leg exopod 2 elliptical; longer than broad; nearly 2.5 times; equal size between both sides; disform inner side; convex outer side; without posterior proximal swelling; inner-posterior process absent; without marginal expansion; curved ridge on distal posterior surface present; chitinous knobs absent; with outer spine; inserted medially; rectilinear; not ornamented innerly; not ornamented outerly; sharp tip; without apparent curve; larger than or equal to the length of the exopod 2; sensilla absent; terminal claw present; equal or longer 1.5 times than insertion segment; sclerotized; arched; inward; without conspicuous curve; ornamented innerly; by denticles; as a row; partially on extension; medially, or distally; not ornamented outerly; sharp tip; curved tip; outwards; without medial constriction; hyaline process absent.

##### FEMALE

Body longer and wider than male; Female body 1305 micrometers excluding caudal setae. Widest at first metasome segment. Distal margin of the prosomal segments without one line of setules at posterior margin. Prosome segments without spinules at prosomal segments. Fourth metasome segment absence of dorsal protuberance. Fourth and fifth metasome segments fused; partially; on dorsal surface. Limit between fourth and fifth metasome segments without ornamentation. **Fifth metasome segment**. Fifth metasome segment without sensilla; with epimeral plates. Epimeral plates asymmetrical. Right epimeral plates prominent, as projections; thinner than the left; one posterior-dorsally directed; not reaching half length of the genital segment; with sensilla at the apex; dorsal-posterior sensilla present; stout; without ornamentation. Left epimeral plate with expansion; semicircular; on posterior surface; dorsally; with sensilla; at tip.

##### Urosome

3-segmented. **Genital double-somite**. Asymmetrical in dorsal view; longer than broad; longer than other urosomites combined; dorsal suture at mid-length absent; not covered by spinules; with swelling; rounded; equal size; anteriorly; with sensillae; on both sides; one; stout; with robust apex; at left lateral; not on lobular base; medially; one; stout; at right lateral; not on lobular base; anteriorly; with robust apex; of equal size between then; lateral protuberance absent; with right posterior rim expanded; over next segment; without slender sensilla on each posterior rim; without posterior-dorsal process. Genital double-somite opercular pad present; broader than longer; symmetrical; development laterally; expanded posteriorly; covering partially; double gonoporal slit; located ventrally; with arthrodial membrane; inserted anteriorly; post-genital process absent; disto-ventral tumescence absent; ventral vertical folds absent; dorsal sensilla absent. Second urosome segment with ventral fusion to anal segment; right distal process present; spiniform; elongated. Caudal rami patch of setules on outer surface present; patch of spinules on outer surface absent.

##### Oral appendices feature

Rostrum basal process absent. **Antennules**. Symmetrical. Right antennule surpassing to genital double-segment; not extending beyond caudal rami; ornamentation pattern equals to male left antennule; fully.

##### Fifth swimming legs

Symmetrical; Fifth swimming legs biramous. Fifth swimming legs intercoxal plate longer than wide; fused to legs. Fifth swimming legs praecoxa with sclerite praecoxal; separated from the coxae; without ornamentation. Fifth swimming legs coxa with process; conical; at the outer rim; distally; sensilla present; stout; at apex; projecting over basal segment; no longer 2x than basal insertion; marginal extension absent; without swelling; without seta; without spinules. Fifth swimming legs basis sub-triangular; unequal size between inner and outer sides; shorter outer than inner side; with convex inner side; with proximal inner outgrowth; without groove; with distal extension; on posterior surface; with seta; outerly inserted; on anterior surface; no longer 2x than origin segment. Fifth swimming legs endopod segments 1 and 2 fused; segments 2 and 3 fused; 1-segmented; stout; separated from the basis; absent discontinuity cuticle; with spinules; as a row; single; non-oblique; sub-terminally; at anterior surface; with seta; double; one medially; on posterior surface; rectilinear; one distally; on posterior surface; arched; of unequal size; distal seta longer than medial seta. Fifth swimming legs exopod segments 1 and 2 separated; segments 2 and 3 separated; 3-segmented; separated from the basis. Fifth swimming legs exopod 1 sub-cylindrical; longer than wide; longer or equal than 2 times; with unequal size between inner and outer side; shorter inner than outer side; with convex inner side; with rectilinear outer side; without swelling; without marginal extension; without posterior process; without spine; without seta. Fifth swimming legs exopod 2 sub-cylindrical; longer than broad; longer or equal than 2 times; without swelling; without marginal extension; without process; without lobe; with spine; inserted laterally; rectilinear; without ornamentation; sharp tip; smaller than next segment; without seta. Fifth swimming legs exopod 3 cylindrical; longer than wide; without swelling; without process; without lobe; without spine; with seta; double; inserted terminally; unequal size between them; outer seta smaller than inner; nearly 1 time; outer seta not ornamented by setules; without ornamentation; presence of terminal claw; sclerotized; arched; externally directed; convex inner side; with ornamentation; of denticles; as a row; on surface partially; at medial region, or at distal region; concave outer side; with ornamentation; of denticles; as a row; on surface partially; at medial region; blunt tip; 6 times longer than origin segment.

##### Distribution records

###### BRAZIL

**Pará**: Lake near Santarém (Wright, 1927); Marajó Island (Wright, 1938b); Curuá-Una Reservoir, 02°48’38“S, 54°18’55“W (Santos-Silva *et al*., 1989).

##### Habitat

Habitat in freshwaters: lake, and reservoir.

##### Remarks

The organisms of this species came from an artificial environment in Northern Brazil. The holotype reported to INPA was located damaged and paratypes were reported to MNHN. The species was attempted to be combined in Dussart (1985) for the subgenus *Notodiaptomus* (*Notodiaptomus*), which would also include *N. gibber*, *N. inflatus*, *N. anceps*, *N. lobifer*, *N. kieferi*, *N. orellanai*, and *N. dilatatus*. This proposal is considered unfounded due critical deficiencies in the delimitation of diagnostic characters and has never been accepted.

In this approach we deal with topotype specimens of the species, duly identified from the specifications originally reported for male: (1) fifth right swimming leg thigh with distal process projecting over basis to beyond the first third until medial surface posteriorly; (2), fifth right swimming leg basis with double triangular protuberance on internal surface distally; (3) fifth right swimming leg exopod 1 with acute marginal extension outerly; (4) female fourth and fifth metasome segments fused completely; (5) female genital second urosome elongated spiniform process; and (6) female caudal rami with patch of setules on outer surface. All the highlighted attributes were corroborated during our analyses, except for the condition of the female last metasome segment fused (4), in which we verified partial fusion only on the dorsal region of the female, with the sides separated.

In the original description, the oral appendages, legs are not described, and some varieties are noted in this effort. The main attribute to be highlighted is for antenna with the presence of 3 endopodal segments for the organisms of the species. Additionally, it was possible to observe for the maxilliped the absence of a spinules row in the praecoxa. Both features represent a contrast to the *Notodiaptomus* pattern redescribed in Santos-Silva *et al*. (1999), for *nordestinus* group of Wright (1935) and revised by Santos-Silva *et al*. (2015) – taxonomic basis for the genus proposal.

Santos-Silva *et al*. (1989) offered detailed taxonomic illustrations of these structures and, here, we also corroborate the annotations. As noted in the cited collaboration, also we verified additional morphological characteristics to the taxon: (1) male fifth right swimming leg exopod 2 with outward curved tip on terminal claw; (2) male fifth right swimming leg exopod 2 with outer spine 1.5x length of the original segment; (3) female caudal rami with setules patch on outer surface; and (4) female fifth swimming legs exopod 2 with lateral seta smaller than next segment. All these are divergent attributes for the type-species of *Notodiaptomus*, which we identified characteristics convergent to the congeners mostly and has, in *N. paraensis*, constant variations: (1) antennule length not extending beyond caudal rami; (2) male fifth left swimming leg exopod 2 with denticles on anterior surface of the digitiform process; (3) male fifth right swimming leg exopod 2 with outer spine medially; (4) female fifth swimming legs basis with outer seta not reaching to distal-point exopod 1; and (5) female fifth swimming legs endopod without cuticle discontinuity. Among the characteristics of Wright (1935) and Kiefer (1936; 1956) *N. paraensis* only has divergence through its condition for male fifth right swimming leg exopod broader than long.

#### Notodiaptomus pseudodubius Defaye & Dussart, 1988

##### Synonymy

*Notodiaptomus pseudodubius* Defaye and Dussart, 1988: 113–114, figs. 16–20; Sendacz, 1993: 31; Rocha *et al*., 1995: 220; Perbiche-Neves *et al*., 2020: 672, 681-682, key to the Neotropical diaptomid, fig. 21.8 J.

##### Type locality

Rorota Lake, Rorota City, route “2” near to Cayenne, French Guiana.

##### Type material

Holotype: 1 male, dissected in glycerin, 21.X.1985, B. Dussart and R. Rojas-Beltran collectors, deposited in the private collection of B.H. Dussart (B.H.D. 1281), indicated as transferred to the National Museum of Natural History in Paris, posteriorly. Paratype: 1 female and 1 male, dissected (B.H.D. 1282); 5 males and 5 females in liquid medium (5% formaldehyde), and deposited in Dussart’s collection (B.H.D. 807); 5 males and 5 females in liquid medium (5% formaldehyde). All paratypes also indicated as stored in NMNH, posteriorly.

##### Material examined

Paratype: 2 males and 2 females (MNHN - PAP-11), entire in in liquid medium (5% formaldehyde), from Rorota 2 Lake, Rorota, French Guiana, from the Dussart collection n° 807. 1 male (INPA-COP047, slides a-h) and 1 female (INPA-COP048, slides a-h) were selected to be dissection on eight slides each and deposited in the Zoological Collection of the INPA, Brazil.

##### Diagnosis

**(1)** male right epimeral plate with sensilla at the apex of projection and other medially; **(2)** male fifth left swimming leg with length reaching first right exopod segment medially; **(3)** male fifth left swimming leg exopod 2 with spiniform seta surpassing distal-point of the origin segment; **(4)** male fifth right swimming leg coxa with distal conical process projecting over basis not beyond the first third posteriorly; **(5)** male fifth right swimming leg basis with medial double triangular protuberance innerly; **(6)** male fifth right swimming leg exopod 1 longer than broad 2x nearly; **(7)** male fifth right swimming leg exopod 1 with distal rounded process posteriorly; **(8)** male fifth right swimming leg exopod 2 in elliptical form; **(9)** female fourth metasome segments with dorsal protuberance distally; **(10)** female genital double-somite without right posterior rim expanded.

##### Redescription

###### MALE

Body 1240 micrometers excluding caudal setae. Male body smaller and slenderer than female. Nerve axons myelinated. Prosome 6-segmented; widest at first metasome segment; without one line of setules at posterior margin; without spinules at segments. Cephalosome anterior margin rounded; with dorsal suture; complete; separate from first metasome segment. First metasome segment without sensilla. Second metasome segment with sensilla; 2 dorsally; of equal size. Third metasome segment with sensillae; 4 laterally; of unequal size; non-ornamented posterior margin. Fourth metasome segment with sensillae; 2 dorsally; 4 laterally; of equal size; fused to fifth metasome; fourth metasome segment partially; fourth metasome segment on lateral surface. Limit between fourth and fifth metasome segments without ornamentation. Fifth metasome segment with sensilla; 4 laterally; Fifth metasome segment equal size; Fifth metasome segment without ornamentation; Fifth metasome segment without dorsal conical process; with epimeral plates. Epimeral plates asymmetrical. Right epimeral plates reduced, as rounded distal corner segment limit; with sensilla; one at the apex of projection and other medially; without ornamentation. Left epimeral plate prominent, as projection; one projection; posterior-dorsally directed; reaching beyond half length of the genital segment; with sensillae; one at the apex of projection and other medially; without ornamentation.

##### Urosome

5-segmented; Urosome 5-free segments. Genital somite asymmetrical in dorsal view; with single aperture; located on left side; ventrolaterally on posterior rim; with sensillae; on single side; one; at right rim; posteriorly. Third urosome segment without spinules; without external seta. Fourth urosome segment without spinules; without sub-conical blunt dorsal-lateral process. Anal segment presence of dorsal sensillae; one on each side; medially inserted; presence of operculum; convex; covering the anal aperture fully. Caudal rami symmetrical; separated from anal segment; longer than wide; with setules; continuous on; inner side; each ramus bearing 6 caudal setae; 5 marginals; plumose; and 1 internal dorsally; straight; not reticulated main axis; outermost seta with outer spiniform process absent.

##### Oral appendices feature

Rostrum symmetrical; separated from dorsal cephalic shield; by complete suture; sensillae present; one pair; anteriorly inserted on surface tegument; with rostral filament; double; paired; extended; into point; with basal process; in ventral view, rounded on left side; without a smaller basal expansion on the right side.

##### Antennules

Asymmetrical. **Right antennules**. Uniramous; right antennule not surpassing to genital segment.

Right antennule ancestral segment I and II separated. Ancestral segment II and III fused. Ancestral segment III and IV fused. Ancestral segment IV and V separated. Ancestral segment V and VI separated. Ancestral segment VI and VII separated. Ancestral segment VII and VIII separated. Ancestral segment VIII and IX separated. Ancestral segment IX and X separated. Ancestral segment X and XI separated. Ancestral segment XI and XII separated. Ancestral segment XII and XIII separated. Ancestral segment XIII and XIV separated. Ancestral segment XIV and XV separated. Ancestral segment XV and XVI separated. Ancestral segment XVI and XVII separated. Ancestral segment XVII and XVIII separated. Ancestral segment XVIII and XIX separated. Ancestral segment XIX and XX separated. Ancestral segment XX and XXI separated. Ancestral segment XXI and XXII fused. Ancestral segment XXII and XXIII fused. Ancestral segment XXIII and XXIV separated. Ancestral segment XXIV and XXV fused. Ancestral segment XXV and XXVI separated. Ancestral segment XXVI and XXVII separated. Ancestral segment XXVII and XXVIII fused.

Right antennule actual 22-segmented; geniculated; between the segment 18 and segment 19; with swollen and modified region; formed by 5 segments; between 13 and 17 segments. Actual segment 1 with seta; one element; straight; none larger than segment; without spinules; without vestigial seta; without conical seta; without modified seta; without spinous process; with aesthetasc; one element. Actual segment 2 with seta; three elements; of unequal size; straight; none larger than segment; without spinules; with vestigial seta; one element; without conical seta; without modified seta; without spinous process; with aesthetasc; one element. Actual segment 3 with seta; one element; one larger than segment; surpassing to distal margin; beyond three sequential segments; straight; blunt apex; without spinules; with vestigial seta; one element; without conical seta; without modified seta; without spinous process; with aesthetasc. Actual segment 4 with seta; one element; one larger than segment; surpassing to distal margin; straight; not beyond three sequential segments; without spinules; without vestigial seta; without conical seta; without modified seta; without spinous process; without aesthetasc. Actual segment 5 with seta; one element; straight; one larger than segment; surpassing to distal margin; not beyond three sequential segments; without spinules; with vestigial seta; one element; without conical seta; without modified seta; without spinous process; with aesthetasc; one element. Actual segment 6 with seta; one element; none larger than segment; straight; without spinules; without vestigial seta; without conical seta; without modified seta; without spinous process; without aesthetasc. Actual segment 7 with seta; one element; straight; one larger than segment; surpassing to distal margin; beyond three sequential segments; blunt apex; without spinules; without vestigial seta; without conical seta; without modified seta; without spinous process; with aesthetasc; one element. Actual segment 8 with seta; one element; straight; none larger than segment; without spinules; without vestigial seta; with conical seta; one element; not reaching to middle-point of the sequent segment; without modified seta; without spinous process; without aesthetasc. Actual segment 9 with seta; two elements; of unequal size; straight; one larger than segment; surpassing to distal margin; beyond three sequential segments; blunt apex; without spinules; without vestigial seta; without conical seta; without modified seta; without spinous process; with aesthetasc; one element. Actual segment 10 with seta; one element; straight; none larger than segment; without spinules; without vestigial seta; without conical seta; with modified seta; presenting blunt apex; slender form; surpassing distal margin; parallel to antennule direction; without spinous process; without aesthetasc. Actual segment 11 with seta; one element; straight; one larger than segment; surpassing to distal margin; not beyond three sequential segments; without spinules; without vestigial seta; without conical seta; with modified seta; slender form; presenting blunt apex; surpassing to distal margin; not beyond of the sequential segment; parallel to antennule direction; shorter length than homologous of actual segment 13; without spinous process; without aesthetasc. Actual segment 12 with seta; one element; straight; one larger than segment; surpassing to distal margin; not beyond three sequential segments; without spinules; without vestigial seta; with conical seta; one element; not smaller than to segment 8; without modified seta; with aesthetasc; one element; absent internal perpendicular fission. Actual segment 13 with seta; one element; straight; one larger than segment; surpassing to distal margin; not beyond three sequential segments; without spinules; without vestigial seta; without conical seta; with modified seta; stout form; surpassing to distal margin; to the distal-point of the sequence segment; perpendicular to antennule direction; presenting bifid apex; without spinous process; with aesthetasc; one element. Actual segment 14 with seta; two elements; of unequal size; straight; one larger than segment; surpassing to distal margin; beyond three sequential segments; blunt apex; without spinules; without vestigial seta; without conical seta; without modified seta; without spinous process; with aesthetasc; one element. Actual segment 15 with seta; two elements; of unequal size; straight; not bifidform; none larger than segment; without spinules; without vestigial seta; without conical seta; without modified seta; with spinous process; on outer margin; surpassing distal margin; with aesthetasc; one element. Actual segment 16 with seta; two elements; of unequal size; plumose; one larger than segment; surpassing to distal margin; not beyond three sequential segments; not bifidform; without spinules; without vestigial seta; without conical seta; without modified seta; with spinous process; on outer margin; surpassing distal margin; unequal size to process on preceding segment; with aesthetasc; one element. Actual segment 17 with seta; two elements; of unequal size; straight; none larger than segment; bifidform; without spinules; without vestigial seta; without conical seta; with modified seta; one element; stout form; surpassing to distal margin; not beyond of the sequential segment; parallel to antennule direction; without spinous process; without aesthetasc. Actual segment 18 with seta; two elements; of equal size; straight; none larger than segment; without spinules; without vestigial seta; without conical seta; with modified seta; one element; stout form; surpassing distal margin; parallel to antennule direction; without spinous process; without aesthetasc. Actual segment 19 with seta; two elements; of unequal size; plumose; none larger than segment; without spinules; without vestigial seta; without conical seta; with modified seta; two elements; stout form; at least one bifid form; surpassing distal margin; parallel to antennule direction; without spinous process; with aesthetasc; one element. Actual segment 20 with seta; four elements; of unequal size; straight; one larger than segment; surpassing to distal margin; beyond three sequential segments; without spinules; without vestigial seta; without conical seta; without modified seta; without spinous process; without aesthetasc. Actual segment 21 with seta; two elements; of equal size; plumose; one larger than segment; surpassing to distal margin; greater 3x than original segment; without spinules; without vestigial seta; without conical seta; without modified seta; without spinous process; without aesthetasc. Actual segment 22 with seta; four elements; of equal size; one larger than segment; plumose; surpassing to distal margin; greater 3x than original segment; without spinules; without vestigial seta; without conical seta; without modified seta; without spinous process; with aesthetasc; one element.

##### Left antennules

Uniramous; Left antennule not surpassing to prosome. Ancestral segment I and II separated. Ancestral segment II and III fused. Ancestral segment III and IV fused. Ancestral segment IV and V separated. Ancestral segment V and VI separated. Ancestral segment VI and VII separated. Ancestral segment VII and VIII separated. Ancestral segment VIII and IX separated. Ancestral segment IX and X separated. Ancestral segment X and XI separated. Ancestral segment XI and XII separated. Ancestral segment XII and XIII separated. Ancestral segment XIII and XIV separated. Ancestral segment XIV and XV separated. Ancestral segment XV and XVI separated. Ancestral segment XVI and XVII separated. Ancestral segment XVII and XVIII separated. Ancestral segment XVIII and XIX separated. Ancestral segment XIX and XX separated. Ancestral segment XX and XXI separated. Ancestral segment XXI and XXII separated. Ancestral segment XXII and XXIII separated. Ancestral segment XXIII and XXIV separated. Ancestral segment XXIV and XXV separated. Ancestral segment XXV and XXVI separated. Ancestral segment XXVI and XXVII separated. Ancestral segment XXVII and XXVIII fused.

Left antennule actual 25-segmented; not-geniculated. Actual segment 1 with seta; one element; none larger than segment; straight; without spinules; without vestigial seta; without conical seta; without modified seta; without spinous process; with aesthetasc; one element. Actual segment 2 with seta; three elements; of equal size; none larger than segment; straight; without spinules; with vestigial seta; one element; without conical seta; without modified seta; without spinous process; with aesthetasc; one element. Actual segment 3 with seta; one element; one larger than segment; straight; surpassing to distal margin; beyond three sequential segments; without spinules; with vestigial seta; one element; without conical seta; without modified seta; without spinous process; with aesthetasc. Actual segment 4 with seta; one element; none larger than segment; straight; without spinules; without vestigial seta; without conical seta; without modified seta; without spinous process; without aesthetasc. Actual segment 5 with seta; one element; one larger than segment; straight; surpassing to distal margin; not beyond three sequential segments; without spinules; with vestigial seta; one element; without conical seta; without modified seta; without spinous process; with aesthetasc; one element. Actual segment 6 with seta; one element; none larger than segment; straight; without spinules; without vestigial seta; without conical seta; without modified seta; without spinous process; without aesthetasc. Actual segment 7 with seta; one element; one larger than segment; straight; surpassing to distal margin; beyond three sequential segments; without spinules; without vestigial seta; without conical seta; without modified seta; without spinous process; with aesthetasc; one element. Actual segment 8 with seta; one element; one larger than segment; straight; surpassing distal margin; without spinules; without vestigial seta; with conical seta; without modified seta; without spinous process; without aesthetasc. Actual segment 9 with seta; two elements; of unequal size; one larger than segment; straight; surpassing to distal margin; beyond three sequential segments; without spinules; without vestigial seta; without conical seta; without modified seta; without spinous process; with aesthetasc; one element. Actual segment 10 with seta; one element; none larger than segment; straight; without spinules; without vestigial seta; without conical seta; without modified seta; without spinous process; without aesthetasc. Actual segment 11 with seta; one element; one larger than segment; straight; surpassing to distal margin; beyond three sequential segments; without spinules; without vestigial seta; without conical seta; without modified seta; without spinous process; without aesthetasc. Actual segment 12 with seta; one element; one larger than segment; straight; surpassing distal margin; without spinules; without vestigial seta; with conical seta; without modified seta; without spinous process; with aesthetasc; one element. Actual segment 13 with seta; one element; none elongated; straight; surpassing distal margin; without spinules; without vestigial seta; without conical seta; without modified seta; without spinous process; without aesthetasc. Actual segment 14 with seta; one element; elongated; straight; surpassing to distal margin; beyond three sequential segments; without spinules; without vestigial seta; without conical seta; without modified seta; without spinous process; with aesthetasc; one element. Actual segment 15 with seta; one element; larger than segment; straight; surpassing to distal margin; not beyond three sequential segments; without spinules; without vestigial seta; without conical seta; without modified seta; without spinous process; without aesthetasc. Actual segment 16 with seta; one element; larger than segment; plumose; surpassing to distal margin; not beyond three sequential segments; without spinules; without vestigial seta; without conical seta; without modified seta; without spinous process; with aesthetasc; one element. Actual segment 17 with seta; one element; not larger than segment; straight; without spinules; without vestigial seta; without conical seta; without modified seta; without spinous process; without aesthetasc. Actual segment 18 with seta; one element; larger than segment; straight; surpassing to distal margin; beyond three sequential segments; without spinules; without vestigial seta; without conical seta; without modified seta; without spinous process; without aesthetasc. Actual segment 19 with seta; one element; not larger than segment; straight; surpassing distal margin; without spinules; without vestigial seta; without conical seta; without modified seta; without spinous process; with aesthetasc; one element. Actual segment 20 with seta; one element; not larger than segment; straight; surpassing distal margin; without spinules; without vestigial seta; without conical seta; without modified seta; without spinous process; without aesthetasc. Actual segment 21 with seta; one element; larger than segment; plumose; surpassing to distal margin; beyond three sequential segments; without spinules; without vestigial seta; without conical seta; without modified seta; without spinous process; without aesthetasc. Actual segment 22 with seta; two elements; of unequal size; one of them elongated; plumose; surpassing to distal margin; without spinules; without vestigial seta; without conical seta; without modified seta; without spinous process; without aesthetasc. Actual segment 23 with seta; two elements; of unequal size; one larger than segment; plumose; surpassing to distal margin; greater 3x than original segment; without spinules; without vestigial seta; without conical seta; without modified seta; without spinous process; without aesthetasc. Actual segment 24 with seta; two elements; of equal size; one larger than segment; plumose; surpassing to distal margin; greater 3x than original segment; without spinules; without vestigial seta; without conical seta; without modified seta; without spinous process; without aesthetasc. Actual segment 25 with seta; four elements; of equal size; elongated; plumose; surpassing to distal margin; 4 times larger than segment; without spinules; without vestigial seta; without conical seta; without modified seta; without spinous process; with aesthetasc; one element.

##### Antenna

Biramous. Antenna coxa separated from the basis; bearing seta; 1; on inner surface; at distal corner; reaching to the endopod 1. Antenna basis (fusion) separated from the endopodal segment; bearing seta; 2; on inner surface; at distal corner. Endopodal ancestral segment I and II separated. Ancestral segment II and III fused. Ancestral segment III and IV fused. Ancestral segment III and IV fully. Antenna endopod actual 2-segmented. Actual segment 1 not bilobate; with seta; two; on inner margin; with spinules; as a row; obliquely; on outer surface; with pore. Actual segment 2 bilobate; with discontinuity on outer cuticle; not developed as a suture; inner lobe bearing 8 setae; distally; outer lobe bearing 7 setae; distally; with spinules; as a patch; on outer surface. Antenna exopod ancestral segment I and II separated. Ancestral segment II and III fused. Ancestral segment III and IV fused. Ancestral segment IV and V separated. Ancestral segment V and VI separated. Ancestral segment VI and VII separated. Ancestral segment VII and VIII separated. Ancestral segment VIII and IX separated. Ancestral segment IX and X fused. Antenna exopod actual 7-segmented. Actual segment 1 single; elongated (width-length, equal or larger ratio 2:1); with seta; one; at inner surface. Actual segment 2 compound; elongated (larger width-length ratio 2:1); with seta; three; at inner surface. Actual segment 3 single; not elongated (lesser width-length ratio 2:1); with seta; one; at inner surface. Actual segment 4 single; not elongated (lesser width-length ratio 2:1); with seta; one; at inner surface. Actual segment 5 single; not elongated (lesser width-length ratio 2:1); with seta; one; at inner surface. Actual segment 6 single; not elongated (lesser width-length ratio 2:1); with seta; one; at inner surface. Actual segment 7 compound; elongated (larger or equal width-length ratio 2:1); with seta; one; at inner surface; and three; at distal surface.

##### Oral features

**Mandible**. Coxal gnathobase sclerotized; with lobe; prominent; on caudal margin; presence of cutting blade; with tooth-like prominence; two, distinctly; 1 acute; on caudal margin; and 1 triangular; on sub-caudal margin; without acute projection between the prominences; with additional spinules; as a row; on dorsal surface; with seta; 1; dorsally; on apical surface; with spinules; apicalmost. Mandible palps biramous; comprising the basis; with seta; four; differently inserted; first medially; reaching to beyond the endopod 1; second distally; third distally; fourth distally; on inner margin; none with setulose ornamentation. Mandible endopod 2-segmented. Mandible endopod 1 with lobe; bearing seta; four; distally inserted; without spinules. Mandible endopod 2 without lobe; bearing setae; nine elements; distally inserted; with spinules; as a row; double. Mandible exopod 4-segmented. Mandible exopod 1 with seta; one element; distally; on inner margin. Mandible exopod 2 with seta; one element; distally; on inner side. Mandible exopod 3 with seta; one element; distally; on inner side. Mandible exopod 4 with setae; three elements; on terminal region. **Maxillule**. Birramous. Maxillule 3-segmented. Maxillule praecoxa with praecoxal arthrite; bearing spines; fifteen elements; ten marginally; plus, five sub-marginally; with spinules; as a patch; on sub-marginal surface. Maxillule coxa with coxal epipodite; with conspicuous outer lobe; bearing setae; nine elements; with coxal endite; elongated (larger or equal width-length ratio 2:1); bearing setae; four elements. Maxillule basis with basal endite; double; first proximal; elongated (larger width-length ratio 2:1; separated from basis; with setae; four elements; distally inserted; second distal; fused to basis; not elongated (lesser width-length ratio 2:1); with setae; four elements; distally inserted; with setules; as a row; on inner side; basal exite present; with setae; one element; on outer surface. Maxillule endopod 1-segmented. Endopod 1 bilobate; first proximal; with setae; three elements; second distal; with setae; five elements. Maxillule exopod 1-segmented. Exopod 1 with setae; six elements; with setules; as a row; on inner side; spinules absent. **Maxilla**. Uniramous. Maxilla 5-segmented. Maxilla praecoxa fused to coxa; incompletely; distinct externally; with praecoxal endite; double; first elongated endite (larger or equal width length ratio 2:1); proximally inserted; with seta; straight, or plumose; 1 straight; 4 plumose; with spine; single; without spinules; without setule; second elongated endite (larger or equal width length ratio 2:1); distally inserted; with seta; plumose; 3 plumose; without spine; with spinules; as a row; on distal margin; with setule; as a row; on distal margin; absence of outer seta. Maxilla coxa with coxal endite; double; first elongated endite (larger or equal width); proximally inserted; with seta; plumose; 3 plumose; without spine; without spinules; with setules; as a row; on proximal margin; second elongated endite (larger or equal width); distally inserted; with seta; plumose; 3 plumose; without spine; without spinules; with setules; as a row; on proximal margin; absence of outer seta. Maxilla basis with basal endite; single; elongated (larger or equal width-length ratio 2:1); with seta; plumose; 3 plumose; without spinules; absence of outer seta. Maxilla endopod 2-segmented. Endopod 1 with seta; 2 plumose; without spine; without spinules; without setules. Maxilla endopod 2 with seta; 2 plumose; without spine; without spinules; without setules. **Maxilliped**. Uniramous; Maxilliped 8-segmented. Maxilliped praecoxa fused to coxa; incompletely; distinct internally; with praecoxal endite; not elongated (lesser width-length ratio 2:1); distally inserted; with seta; 1 straight; with spinules; as a row; single; on basal surface; without setules. Maxilliped coxa with coxal endite; three coxal endite; first elongated (larger or equal width); proximally inserted; with seta; 2 plumose; with spinules; as a patch; single; on apical surface; without setules; second not elongated (lesser width-length ratio 2:1); medially inserted; with seta; 3 plumose; with spinules; as a row; single; on medial surface; without setules; third elongated (larger or equal width length ratio 2:1); distally inserted; with seta; 3 plumose; none reaching to beyond of the basis; with spinules; as a row; single; on basal surface; without setules; with lobe; prominence; at inner distal angle; ornamented; with spinules; continuously on margin. Maxilliped basis without basal endite; with seta; 3 plumose; with spinules; as a row; single; on medial surface; with setules; as a row; single; on inner margin. Maxilliped endopod segment 6-segmented. Endopod 1 with seta; 2 plumose; on inner surface. Endopod 2 with seta; 3 plumose; on inner surface. Endopod 3 with seta; 2 plumose; on inner surface. Endopod 4 with seta; 2 plumose; on inner surface. Endopod 5 with seta; 2 plumose; on inner surface, or on outer surface; outer seta absent. Endopod 6 with seta; 4 plumose; on inner surface, or on outer surface.

##### Swimming legs features

**First swimming legs.** Symmetrical; biramous. First swimming legs intercoxal plate without seta. First swimming legs praecoxa absent. First swimming legs coxa with seta; one; straight; distally inserted; on inner surface; surpassing to basal segment; with setules; two group; as a patch; on inner margin; and as a row; double; on anterior surface; outerly; without spinules; without spine. First swimming legs basis without seta; with setules; as a patch; single; on outer surface; without spinules; without spine. First swimming legs endopod 2-segmented. Endopod 1 with seta; straight; restricted; to inner surface; one element; without spine; with setules; as a row; single; continuously; on outer surface; without spinules; absence of Schmeil’s organ. Endopod 2 with seta; unrestricted; three on inner surface; one on outer surface; two on distal surface; straight; without spine; with setules; as a row; single; continuously; on outer surface; without spinules; absence of Schmeil’s organ. Endopod 3 absence. First swimming legs exopod 1 with seta; restricted; 1 on inner surface; with spine; 1; stout; smaller than original segment; serrated; on inner side; continuously; without setules. First swimming legs exopod 2 with seta; restricted; 1 on inner surface; straight; without spine; with setules; as a row; single; continuously; on inner margin, or on outer margin; without spinules. First swimming legs exopod 3 with setule; as a row; single; continuously; on outer surface; without spinules; with seta; unrestricted; 2 on inner surface; 2 on terminal surface; with spine; 2; unequal size; first no longer 2x than origin segment; stout; serrated; on inner side, or on outer side; equally; second longer 3x than origin segment; slender; serrated; on outer side; with ornamentation on non-serrated side; by setules. **Second swimming legs**. Symmetrical; Second swimming legs biramous. Second swimming legs intercoxal plate without seta. Second swimming legs praecoxa present; located laterally. Second swimming legs coxa with seta; straight; distally inserted; on inner surface; surpassing to basal segment; without setules; without spinules; without spine. Second swimming legs basis without seta; without setules; without spinules; without spine. Second swimming legs endopod 3-segmented. Endopod 1 with seta; straight; restricted; one on inner surface; without spine; with setules; as a row; single; continuously; on outer surface; without spinules; absence of Schmeil’s organ. Endopod 2 with seta; straight; unrestricted; two on inner surface; without spine; with setules; as a row; single; continuously; on outer side; without spinules; presence of Schmeil’s organ; on posterior surface. Endopod 3 with seta; straight; unrestricted; three on inner surface; two on outer surface; two on distal surface; without spine; without setules; with spinules; as a row; double; distally inserted; at anterior surface; absence of Schmeil’s organ. Second swimming legs exopod 1 with seta; restricted; one on inner surface; with spine; 1; stout; not reaching to distal-third of the exopod 2; serrated; on inner side, or on outer side; with setules; as a row; single; continuously; on inner side; without spinules; absence of Schmeil’s organ. Exopod 2 with seta; unrestricted; one on inner surface; with spine; 1; stout; not surpassing the exopod 3; serrated; on inner side, or on outer side; with setules; as a row; single; continuously; on inner surface; without spinules; absence of Schmeil’s organ. Exopod 3 with seta; plurimarginal; three on inner surface; two on terminal surface; with spine; 2; unequal size; first no longer 2x than origin segment; stout; serrated; on inner side, or on outer side; equally; second longer 2x than origin segment; slender; serrated; on outer side; with ornamentation on non-serrated side; of setules; setules on outer surface; as a row; single; continuously; on inner surface; with spinules; as a row; single; distally inserted; at anterior surface; absence of Schmeil’s organ. **Third swimming legs**. Symmetrical; Third swimming legs biramous. Third swimming legs intercoxal plate without seta. Third swimming legs praecoxa present; not laterally located. Third swimming legs coxa with seta; straight; distally inserted; on inner surface; surpassing to basal segment; without setules; without spinules; without spine. Third swimming legs basis without seta; without setules; without spinules; without spine. Third swimming legs endopod 3-segmented. Endopod 1 with seta; restricted; one on inner surface; without spine; without setules; without spinules; absence of Schmeil’s organ. Endopod 2 with seta; restricted; two on inner surface; straight; without spine; without setules; without spinules; absence of Schmeil’s organ. Endopod 3 with seta; straight; plurimarginal; two on inner surface; two on outer surface; three on terminal surface; without spine; without setules; with spinules; as a row; distally inserted; double; at anterior surface; absence of Schmeil’s organ. Third swimming legs exopod 1 with seta; restricted; straight; one on inner surface; with spine; 1; stout; not reaching to the distal-third of the exopod 2; serrated; equally; on inner surface, or on outer surface; with setules; as a row; single; continuously; on inner surface; without spinules; absence of Schmeil’s organ. Exopod 2 with seta; straight; restricted; one on inner surface; with spine; 1; stout; not reaching out to exopod 3; serrated; on inner side, or on outer side; equally; with setules; as a row; single; continuously; on inner side; without spinules; absence of Schmeil’s organ. Exopod 3 without setules; with spinules; as a row; single; distally inserted; at anterior surface; with seta; straight; unrestricted; three on inner surface; two on terminal surface; with spine; 2; unequal size; first no longer 2x than origin segment; stout; serrated; on inner side, or on outer side; equally; second longer 2x than origin segment; slender; serrated; on outer side; with ornamentation on non-serrated side; of setules; absence of Schmeil’s organ. **Fourth swimming legs**. Symmetrical; biramous. Intercoxal plate without sensilla. Praecoxa present. Coxa with seta; distally inserted; on inner margin; reaching out to endopod 1; without spinules; setules absent. Basis with seta; one; medially inserted; on posterior surface; smaller than the original segment; without setules; without spinules; without spine. Fourth swimming legs endopod 3-segmented. Endopod 1 with seta; one; restricted; on inner surface; without spine; without setules; without spinules; absence of Schmeil’s organ. Endopod 2 with seta; restricted; two on inner side; without spine; with setules; as a row; single; continuously; on outer surface; without spinules; absence of Schmeil’s organ. Endopod 3 with seta; unrestricted; two on inner surface; two on outer surface; three on distal surface; without spine; without setules; with spinules; as a row; double; distally inserted; at anterior surface; absence of Schmeil’s organ. Fourth swimming legs exopod 1 with seta; restricted; one on inner surface; with spine; 1; stout; not reaching out to distal-third of the exopod 2; serrated; on inner side, or on outer side; equally; with setules; as a row; single; continuously; on inner surface; without spinules; absence of Schmeil’s organ. Exopod 2 with seta; restricted; one on inner surface; with spine; 1; stout; not reaching the end of exopod 3; serrated; on inner side, or on outer side; equally; with setules; as a row; single; continuously; on inner surface; without spinules; absence of Schmeil’s organ. Exopod 3 without setules; with spinules; as a row; single; distally inserted; at anterior surface; with seta; unrestricted; three on inner surface; two on distal surface; with spine; 2; unequal size; first no longer 2x than origin segment; stout; serrated; on inner side, or on outer side; equally; second longer 2x than origin segment; slender; serrated; on outer side; without ornamentation on non-serrated side; absence of Schmeil’s organ.

##### Fifth swimming legs features

Asymmetrical. Fifth swimming leg intercoxal plate with length not equal or greater than width on 1.5x; with irregular proximal margin; discontinuous to; the anterior margin of the left coxa, or the anterior margin of the right coxa; posterior sensilla on the right lateral absent. **Fifth left swimming leg**. Fifth left swimming leg biramous; leg reaching first right exopod segment; medially. Fifth left swimming leg praecoxa present; rudimentary; separated from the coxae; without ornamentation. Fifth left swimming leg coxa concave inner side; without teeth-like structures; with process; conical; on posterior surface; outer side; distally inserted; not projecting over basis; with sensilla; stout; triangular; at apex; longer 2x than insertion basis; without swelling; without seta; without spinules. Fifth left swimming leg basis sub-cylindrical; unequal size between inner and outer side; shorter outer than inner side; with concave inner side; rounded internal proximal expansion absent; without outgrowth; with groove; deep; obliquely; on posterior surface; not reaching the endopodal lobe; not ornamented; absence of protuberance; with seta; outerly inserted; no longer 2x than origin segment; absence of minutely granular. Fifth left swimming leg endopod segments 1 and 2 fused; segments 2 and 3 fused; 1-segmented; stout; separated from the basis; ornamented; on inner side; with spinules; more than four elements; as a row; terminally; row of setules absent; without seta. Fifth left swimming leg exopod segments 1 and 2 separated; segments 2 and 3 fused; 2-segmented; stout; separated from the basis. Fifth left swimming leg exopod 1 sub-triangular; longer than broad; equal size between inner and outer side; rectilinear inner side; convex outer side; without swelling; without marginal extension; without process; with lobe; single; semicircular; medially inserted; on inner side; covered; by setules; without outer spine; absence seta. Fifth left swimming leg exopod 2 digitiform; longer than broad; equal size between inner and outer side; disform inner side; with rectilinear outer side; setulose pad present; not prominently rounded; proximally; on inner side; inflated medial region absent; distal process present; digitiform; denticulate; not bicuspidate; without transverse row of denticles; none oblique row of 5 denticles; at anterior surface; not innerly directed; with seta; spiniform; not ornamented by spinules; surpassing the distal-point of the segment; without outer spine; terminal claw absent.

##### Fifth right swimming leg

Biramous. Fifth right swimming leg praecoxa present; separated from the coxae; without ornamentation. Fifth right swimming leg coxa convex inner side; without teeth-like structures; with process; conical; distally inserted; on posterior surface; closest to the outer rim; projecting over basis; not beyond the first third; without triangular protuberance innerly; with sensilla; slender; at apex; no longer 2x than basal insertion; without marginal extension; without seta; without spinules. Fifth right swimming leg basis trapezoidal; unequal size between inner and outer side; shorter outer than inner side; concave inner side; tumescence present; inflated; not bilobed; unrestricted on inner surface; with protuberance; double; triangular; on inner side; medially; ornamented; with denticles; not minutely granular; absence of distinct minutely granular; additional inner process absent; without posterior groove; with seta; outerly inserted; on anterior surface; no longer 2x than origin segment; posterior protrusion absent; distal process absent. Fifth right swimming leg with endopodite present; fused to basis; on anterior surface; ancestral segments 1 and 2 fused; ancestral segments 2 and 3 fused; stout ornamented; with setules; as a row; on inner side; terminally; without seta. Fifth right swimming leg exopod segments 1 and 2 separated; segments 2 and 3 fused; 2-segmented; stout; separated from the basis. Fifth right swimming leg exopod 1 trapezium; longer than broad; nearly 2 times; unequal size between both sides; shorter outer than inner side; rectilinear inner side; convex outer side; with marginal extension; sub-triangular; distally inserted; at outer rim; spinules absent; with process; rounded; sclerotized; without ornamentation; distally inserted; at posterior surface; projecting over next segment; without outer spine; without seta; internal prominence absent; lamella on posterior surface absent. Fifth right swimming leg exopod 2 elliptical; longer than broad; nearly 2 times; unequal size between both sides; disform inner side; convex outer side; without posterior proximal swelling; inner-posterior process absent; without marginal expansion; curved ridge on distal posterior surface present; chitinous knobs absent; with outer spine; inserted sub-distally; rectilinear; not ornamented innerly; not ornamented outerly; sharp tip; without apparent curve; lesser than the length of the exopod 2; until to 2 times its size; 1.5x; sensilla absent; terminal claw present; equal or longer 1.5 times than insertion segment; sclerotized; arched; inward; with conspicuous curve; medially; not ornamented innerly; not ornamented outerly; sharp tip; not curved tip; without medial constriction; hyaline process absent.

##### FEMALE

Body longer and wider than male; Female body 1310 micrometers excluding caudal setae. Widest at first metasome segment. Distal margin of the prosomal segments without one line of setules at posterior margin. Prosome segments without spinules at prosomal segments. Fourth metasome segment presence of dorsal protuberance; rounded; inserted distally; without posterior process; without anterior process; fourth metasome segment without proximal sensillae present. Fourth and fifth metasome segments fused; partially; on dorsal surface. Limit between fourth and fifth metasome segments without ornamentation. **Fifth metasome segment**. Fifth metasome segment without sensilla; with epimeral plates. Epimeral plates asymmetrical. Right epimeral plates prominent, as projections; not thinner than the left; one posterior-laterally directed; not reaching half length of the genital segment; with sensilla at the apex; dorsal-posterior sensilla present; stout; without ornamentation. Left epimeral plate without expansion.

##### Urosome

3-segmented. **Genital double-somite**. Asymmetrical in dorsal view; longer than broad; longer than other urosomites combined; dorsal suture at mid-length absent; not covered by spinules; with swelling; rounded; unequal size; greater left than right; anteriorly; with sensillae; on both sides; one; stout; with robust apex; at left lateral; not on lobular base; medially; one; stout; at right lateral; not on lobular base; medially; with robust apex; of equal size between then; lateral protuberance absent; without right posterior rim expanded; without slender sensilla on each posterior rim; without posterior-dorsal process. Genital double-somite opercular pad present; broader than longer; symmetrical; development laterally; expanded posteriorly; covering partially; double gonoporal slit; located ventrally; with arthrodial membrane; inserted anteriorly; post-genital process absent; disto-ventral tumescence absent; ventral vertical folds absent; dorsal sensilla absent. Second urosome segment without ventral fusion to anal segment; right distal process absent. Caudal rami patch of setules on outer surface absent; patch of spinules on outer surface absent.

##### Oral appendices feature

Rostrum basal process absent. **Antennules**. Symmetrical. Right antennule not surpassing to genital double-segment; ornamentation pattern equals to male left antennule; mostly. Actual segment 13 without seta; without aesthetasc. Actual segment 14 without seta; without aesthetasc. Actual segment 15 without seta; without aesthetasc. Actual segment 16 without seta; without aesthetasc. Actual segment 17 without seta. Actual segment 18 without seta.

##### Fifth swimming legs

Symmetrical; Fifth swimming legs biramous. Fifth swimming legs intercoxal plate longer than wide; separated from the legs. Fifth swimming legs praecoxa with sclerite praecoxal; separated from the coxae; without ornamentation. Fifth swimming legs coxa with process; conical; at the outer rim; distally; sensilla present; stout; at apex; projecting over basal segment; no longer 2x than basal insertion; marginal extension absent; without swelling; without seta; without spinules. Fifth swimming legs basis sub-triangular; unequal size between inner and outer sides; shorter outer than inner side; with convex inner side; without proximal inner outgrowth; without groove; with distal extension; on posterior surface; with seta; outerly inserted; on anterior surface; no longer 2x than origin segment. Fifth swimming legs endopod segments 1 and 2 fused; segments 2 and 3 fused; 1-segmented; stout; separated from the basis; present discontinuity cuticle; on inner side; with spinules; as a row; single; non-oblique; sub-terminally; at anterior surface; with seta; double; one medially; on posterior surface; rectilinear; one distally; on posterior surface; arched; of unequal size; distal seta longer than medial seta. Fifth swimming legs exopod segments 1 and 2 separated; segments 2 and 3 separated; 3-segmented; separated from the basis. Fifth swimming legs exopod 1 sub-cylindrical; longer than wide; longer or equal than 2 times; with unequal size between inner and outer side; shorter inner than outer side; with convex inner side; with rectilinear outer side; without swelling; without marginal extension; without posterior process; without spine; without seta. Fifth swimming legs exopod 2 sub-cylindrical; longer than broad; longer or equal than 2 times; without swelling; without marginal extension; without process; without lobe; with spine; inserted laterally; rectilinear; without ornamentation; sharp tip; equal size or larger than next segment; without seta. Fifth swimming legs exopod 3 cylindrical; longer than wide; without swelling; without process; without lobe; without spine; with seta; double; inserted terminally; unequal size between them; outer seta smaller than inner; nearly 3 times; outer seta not ornamented by setules; without ornamentation; presence of terminal claw; sclerotized; arched; externally directed; convex inner side; with ornamentation; of denticles; as a row; on surface partially; at medial region; concave outer side; with ornamentation; of denticles; as a row; on surface partially; at medial region; blunt tip; 6 times longer than origin segment.

##### Distribution records

###### FRENCH GUIANA

Rorota 2 Lake, Rorota (Defaye and Dussart, 1988).

##### Habitat

Habitat in freshwaters: lake.

##### Remarks

The hypothesis of Defaye &Dussart (1988) was presented from organisms from French Guiana, in Northern South America. Originally individuals were specified for the NMNH in Paris and paratypes were examined for this thesis. In the original description, Defaye & Dussart (1988) describe the species as similar to *N. dubius*, distinct from it by the female: (1) genital double-somite without right distal expansion dorsally; (2) fifth swimming legs with endopod “shorter”; and (3) exopod 2 “shorter” especially. For male: (1) fifth right swimming leg basis; (2) fifth right swimming leg endopod “shorter” especially; (3) fifth right leg exopod 2 with outer spine rectilinear and “strong“; and (4) fifth right leg exopod 2 with terminal claw with “bulbiform basis”.

During our examinations, all the attributes highlighted originally to distinguish the taxon were corroborated. The size difference between the fifth swimming legs endopods of both is unclear, but it is possible to verify that in *N. pseudodubius* it is slightly smaller. Another attribute that seems relevant to us to comment on is male fifth right swimming leg basis, which originally no further details are given about the differential condition between both males. However, through our conclusions, it is possible to verify variation in general form of the segment and inner expansion. In *N. pseudodubius* the male fifth right swimming leg basis is longer and has irregular inner protuberance double, while in *N. dubius* it is shorter and without any expansion.

Other differential characteristics were identified among the species in this effort. It is surprising to note that the authors of the taxon did not highlight the conspicuous difference for the size of the distal conical process posteriorly on male fifth right coxa. For *N. pseudodubius* the condition of this element is shorter considerably, while in *N. dubius* it is projecting over basis until distal surface posteriorly. Additionally, two other characteristics are present only for *N. pseudodubius*: male fifth right swimming leg exopod 1 longer than broad with distal rounded process posteriorly, and exopod 2 in elliptical form.

Defaye & Dussart (1988) described the male fifth metasome segment with double sensillae on left epimeral plate, and one sensilla on right epimeral plate. We have identified that the right sensilla is existing for all males examined here. Furthermore, an objectification must be made about male right antennule, the authors describe the presence of process on actual segments 8, 10 and 11, truly it is a conical seta for the first, and modified seta for the others, respectively. However, *N. pseudodubius* presents morphological divergences to the characteristics highlighted in Kiefer (1936; 1956) for *Notodiaptomus*: (1) female fifth swimming legs endopod with non-oblique spinules row; (2) male fifth right swimming leg basis without posterior protrusion; and (3) male fifth left swimming leg exopod 2 with spiniform seta surpassing to distal-point of segment inserted. Among the converging characteristics within the genus from the type-species, the taxon still has other divergent attributes: (1) male right antennule not surpassing to genital segment; (2) male right antennule actual segment 13 with spiniform modified seta reaching to distal-point of the sequence segment; (3) male fifth left swimming leg length reaching first right exopod segment medially; (4) male fifth right swimming leg basis with inner protuberance in triangular form doubly; (5) female fourth metasome segments with dorsal protuberance; (6) female fifth swimming legs basis with outer seta reaching to exopod 1 medially.

#### Notodiaptomus santafesinus (Ringuelet & Ferrato, 1967)

##### Synonymy

*Diaptomus santafesinus* Ringuelet and Martínez de Ferrato, 1967: 415–416, Pl. II (1–6); Paggi & José de Paggi, 1974: 99, table 1; Brandorff, 1976; Paggi, 1967: 157-159, figs. 26-36; Dussart & Defaye, 1983. *Notodiaptomus santafesinus* n. comb. Dussart & Frutos, 1985: 307, figs. 1-2; Perbiche-Neves *et al*., 2015: 77–81, figs. 69–73; Perbiche-Neves *et al*., 2020: 677-678, key to the Neotropical diaptomid, fig. 21.6 I.

##### Type locality

Not specified clearly. Originally were mentioned individuals found in samples of lotic floodplains from the streams of Sirgadero Island (25.IV.1962), of Ubajay (25.4.1964), Coronda River (19.III.1965), Colorado Stream (12.III.1965), and Madrejón Don Felipe (9.IV.1964). All environments in La Capital, Santa Fé, Argentina.

##### Type material

In the original description, 2 individuals are specified, entire, and several dissected. This study treats this material as “cotypes” (probably paratypes), and deposited at the Instituto Nacional de Limnologia, Santa Fé, Argentina. No depositary code or collection details are provided.

##### Material examined

Non-type material: 2 males, and 1 female, entire in alcohol, from the lagoon of the Naranjos, near to Santa Fé, Argentina, 6.VI.67, store in the collection of the Lab of Plankton (n°. 2); 2 males, and 3 females, entire, in alcohol, from the Reservoir (UHE) Salto Santiago, Paraná, 17.III.1994, 9.IX1997, stored in collection of the Plankton Laboratory, INPA, no code; 1 male (INPA-COP049, slides a-h) and 1 female (INPA-COP050, slides a-h) were selected to be dissection on eight slides each and deposited in the Zoological Collection of the INPA, Brazil. Additional material examined: 01 male, and 1 female, entire in formalin solution (MZUSP 28391) from the middle and lower stretches of the Paraná River, Argentina, stored in MZUSP.

##### Diagnosis

**(1)** Male third and fourth metasome segments with setules patch on lateral and dorsal surface; **(2)** male right antennule actual segment 1 with spinules; **(3)** male fifth left swimming leg reaching first right exopod segment medially; **(4)** male fifth left swimming leg with spiniform seta surpassing distal-point of the origin segment; **(5)** male fifth right swimming leg basis without posterior groove; **(6)** male fifth right swimming leg endopod separated from the basis; **(7)** male fifth right swimming leg exopod 1 with acute internal prominence; **(8)** male fifth right swimming leg exopod 2 with curved ridge on distal posterior surface; **(9)** male fifth right swimming leg exopod 2 with outer spine arched outwards; **(10)** male fifth right swimming leg exopod 2 with outer spine 1.5x length of origin segment; **(11)** male fifth right swimming leg exopod 2 with outwards curved tip on terminal claw; **(12)** female fourth metasome segment with digitiform dorsal protuberance 2x wider than long inserted medially; **(13)** female fourth and fifth metasome segments fused totally; **(14)** female fourth and fifth metasome segments not ornamented; **(15)** female fifth swimming legs coxa with sensilla longer 2x than outer process; **(16)** female fifth swimming legs exopod 2 with lateral spine smaller than next segment.

##### Redescription

###### MALE

Body 998 micrometers excluding caudal setae. Male body smaller and slenderer than female. Nerve axons myelinated. Prosome 6-segmented; widest at first metasome segment; without one line of setules at posterior margin; without spinules at least at one segment. Cephalosome anterior margin sub-triangular; with dorsal suture; incomplete; separate from first metasome segment. First metasome segment without sensilla. Second metasome segment with sensilla; 2 dorsally; of equal size. Third metasome segment with sensillae; 4 laterally; of unequal size; ornamented posterior margin; with setules; as a patch; dorsally, or laterally. Fourth metasome segment with sensillae; 2 dorsally; 4 laterally; of equal size; separated from the fifth metasome. Limit between fourth and fifth metasome segments ornamented; with setules. Fifth metasome segment with sensilla; 4 laterally; Fifth metasome segment equal size; Fifth metasome segment without ornamentation; Fifth metasome segment without dorsal conical process; with epimeral plates. Epimeral plates symmetrical. Right epimeral plates reduced, as rounded distal corner segment limit; with sensilla; at the apex of projection; without ornamentation.

##### Urosome

5-segmented; Urosome 5 - free segments. Genital somite symmetrical in dorsal view; with single aperture; located on left side; ventrolaterally on posterior rim; with sensillae; on both sides; one; at left lateral; posteriorly; one; at right rim; posteriorly; of equal size between then. Third urosome segment without spinules; without external seta. Fourth urosome segment without spinules; without sub-conical blunt dorsal-lateral process. Anal segment presence of dorsal sensillae; one on each side; medially inserted; presence of operculum; convex; covering the anal aperture fully. Caudal rami symmetrical; separated from anal segment; longer than wide; with setules; continuous on; inner side; each ramus bearing 6 caudal setae; 5 marginals; plumose; and 1 internal dorsally; straight; not reticulated main axis; outermost seta with outer spiniform process absent.

##### Oral appendices feature

Rostrum symmetrical; separated from dorsal cephalic shield; by complete suture; sensillae present; one pair; anteriorly inserted on surface tegument; with rostral filament; double; paired; extended; into point; with basal process; in ventral view, rounded on left side; without a smaller basal expansion on the right side.

##### Antennules

Asymmetrical. **Right antennules**. Uniramous; right antennule surpassing to genital segment; right antennule extending beyond caudal rami.

Right antennule ancestral segment I and II separated. Ancestral segment II and III fused. Ancestral segment III and IV fused. Ancestral segment IV and V separated. Ancestral segment V and VI separated. Ancestral segment VI and VII separated. Ancestral segment VII and VIII separated. Ancestral segment VIII and IX separated. Ancestral segment IX and X separated. Ancestral segment X and XI separated. Ancestral segment XI and XII separated. Ancestral segment XII and XIII separated. Ancestral segment XIII and XIV separated. Ancestral segment XIV and XV separated. Ancestral segment XV and XVI separated. Ancestral segment XVI and XVII separated. Ancestral segment XVII and XVIII separated. Ancestral segment XVIII and XIX separated. Ancestral segment XIX and XX separated. Ancestral segment XX and XXI separated. Ancestral segment XXI and XXII fused. Ancestral segment XXII and XXIII fused. Ancestral segment XXIII and XXIV separated. Ancestral segment XXIV and XXV fused. Ancestral segment XXV and XXVI separated. Ancestral segment XXVI and XXVII separated. Ancestral segment XXVII and XXVIII fused.

Right antennule actual 22-segmented; geniculated; between the segment 18 and segment 19; with swollen and modified region; formed by 5 segments; between 13 and 17 segments. Actual segment 1 with seta; one element; straight; none larger than segment; with spinules; without vestigial seta; without conical seta; without modified seta; without spinous process; with aesthetasc; one element. Actual segment 2 with seta; three elements; of unequal size; straight; none larger than segment; without spinules; with vestigial seta; one element; without conical seta; without modified seta; without spinous process; with aesthetasc; one element. Actual segment 3 with seta; one element; one larger than segment; surpassing to distal margin; not beyond three sequential segments; straight; blunt apex; without spinules; with vestigial seta; one element; without conical seta; without modified seta; without spinous process; with aesthetasc. Actual segment 4 with seta; one element; one larger than segment; surpassing to distal margin; straight; not beyond three sequential segments; without spinules; without vestigial seta; without conical seta; without modified seta; without spinous process; without aesthetasc. Actual segment 5 with seta; one element; straight; one larger than segment; surpassing to distal margin; not beyond three sequential segments; without spinules; with vestigial seta; one element; without conical seta; without modified seta; without spinous process; with aesthetasc; one element. Actual segment 6 with seta; one element; none larger than segment; straight; without spinules; without vestigial seta; without conical seta; without modified seta; without spinous process; without aesthetasc. Actual segment 7 with seta; one element; straight; one larger than segment; surpassing to distal margin; beyond three sequential segments; blunt apex; without spinules; without vestigial seta; without conical seta; without modified seta; without spinous process; with aesthetasc; one element. Actual segment 8 with seta; one element; straight; none larger than segment; without spinules; without vestigial seta; with conical seta; one element; reaching to middle-point of the sequent segment; without modified seta; without spinous process; without aesthetasc. Actual segment 9 with seta; two elements; of unequal size; straight; one larger than segment; surpassing to distal margin; beyond three sequential segments; blunt apex; without spinules; without vestigial seta; without conical seta; without modified seta; without spinous process; with aesthetasc; one element. Actual segment 10 with seta; one element; straight; none larger than segment; without spinules; without vestigial seta; without conical seta; with modified seta; presenting blunt apex; slender form; surpassing to distal margin; beyond of the sequential segment; parallel to antennule direction; without spinous process; without aesthetasc. Actual segment 11 with seta; one element; straight; one larger than segment; surpassing to distal margin; not beyond three sequential segments; without spinules; without vestigial seta; without conical seta; with modified seta; slender form; presenting blunt apex; surpassing to distal margin; beyond of the sequential segment; parallel to antennule direction; shorter length than homologous of actual segment 13; without spinous process; without aesthetasc. Actual segment 12 with seta; one element; straight; one larger than segment; surpassing to distal margin; not beyond three sequential segments; without spinules; without vestigial seta; with conical seta; one element; smaller than to segment 8; without modified seta; without spinous process; with aesthetasc; one element; absent internal perpendicular fission. Actual segment 13 with seta; one element; straight; one larger than segment; surpassing to distal margin; not beyond three sequential segments; without spinules; without vestigial seta; without conical seta; with modified seta; stout form; surpassing to distal margin; to the distal-point of the sequence segment; perpendicular to antennule direction; presenting bifid apex; without spinous process; with aesthetasc; one element. Actual segment 14 with seta; two elements; of unequal size; straight; one larger than segment; surpassing to distal margin; beyond three sequential segments; blunt apex; without spinules; without vestigial seta; without conical seta; without modified seta; without spinous process; with aesthetasc; one element. Actual segment 15 with seta; two elements; of unequal size; straight; not bifidform; none larger than segment; without spinules; without vestigial seta; without conical seta; without modified seta; with spinous process; on outer margin; surpassing distal margin; with aesthetasc; one element. Actual segment 16 with seta; two elements; of unequal size; plumose; one larger than segment; surpassing to distal margin; not beyond three sequential segments; not bifidform; without spinules; without vestigial seta; without conical seta; without modified seta; with spinous process; on outer margin; surpassing distal margin; unequal size to process on preceding segment; with aesthetasc; one element. Actual segment 17 with seta; two elements; of unequal size; straight; none larger than segment; bifidform; without spinules; without vestigial seta; without conical seta; with modified seta; one element; stout form; surpassing to distal margin; not beyond of the sequential segment; parallel to antennule direction; without spinous process; without aesthetasc. Actual segment 18 with seta; two elements; of equal size; straight; none larger than segment; without spinules; without vestigial seta; without conical seta; with modified seta; one element; stout form; surpassing distal margin; parallel to antennule direction; without spinous process; without aesthetasc. Actual segment 19 with seta; two elements; of unequal size; plumose; none larger than segment; without spinules; without vestigial seta; without conical seta; with modified seta; two elements; stout form; at least one bifid form; surpassing distal margin; parallel to antennule direction; without spinous process; with aesthetasc; one element. Actual segment 20 with seta; four elements; of unequal size; straight; one larger than segment; surpassing to distal margin; beyond three sequential segments; without spinules; without vestigial seta; without conical seta; without modified seta; with spinous process; distally; not reaching beyond of distal-point segment 21; without aesthetasc. Actual segment 21 with seta; two elements; of equal size; plumose; one larger than segment; surpassing to distal margin; greater 3x than original segment; without spinules; without vestigial seta; without conical seta; without modified seta; without spinous process; without aesthetasc. Actual segment 22 with seta; four elements; of equal size; one larger than segment; plumose; surpassing to distal margin; greater 3x than original segment; without spinules; without vestigial seta; without conical seta; without modified seta; without spinous process; with aesthetasc; one element.

##### Left antennules

Uniramous; Left antennule surpassing to prosome; Left antennule extending beyond caudal rami. Ancestral segment I and II separated. Ancestral segment II and III fused. Ancestral segment III and IV fused. Ancestral segment IV and V separated. Ancestral segment V and VI separated. Ancestral segment VI and VII separated. Ancestral segment VII and VIII separated. Ancestral segment VIII and IX separated. Ancestral segment IX and X separated. Ancestral segment X and XI separated. Ancestral segment XI and XII separated. Ancestral segment XII and XIII separated. Ancestral segment XIII and XIV separated. Ancestral segment XIV and XV separated. Ancestral segment XV and XVI separated. Ancestral segment XVI and XVII separated. Ancestral segment XVII and XVIII separated. Ancestral segment XVIII and XIX separated. Ancestral segment XIX and XX separated. Ancestral segment XX and XXI separated. Ancestral segment XXI and XXII separated. Ancestral segment XXII and XXIII separated. Ancestral segment XXIII and XXIV separated. Ancestral segment XXIV and XXV separated. Ancestral segment XXV and XXVI separated. Ancestral segment XXVI and XXVII separated. Ancestral segment XXVII and XXVIII fused.

Left antennule actual 25-segmented; not-geniculated. Actual segment 1 with seta; one element; none larger than segment; straight; without spinules; without vestigial seta; without conical seta; without modified seta; without spinous process; with aesthetasc; one element. Actual segment 2 with seta; three elements; of equal size; none larger than segment; straight; without spinules; with vestigial seta; one element; without conical seta; without modified seta; without spinous process; with aesthetasc; one element. Actual segment 3 with seta; one element; one larger than segment; straight; surpassing to distal margin; beyond three sequential segments; without spinules; with vestigial seta; one element; without conical seta; without modified seta; without spinous process; with aesthetasc. Actual segment 4 with seta; one element; none larger than segment; straight; without spinules; without vestigial seta; without conical seta; without modified seta; without spinous process; without aesthetasc. Actual segment 5 with seta; one element; one larger than segment; straight; surpassing to distal margin; not beyond three sequential segments; without spinules; with vestigial seta; one element; without conical seta; without modified seta; without spinous process; with aesthetasc; one element. Actual segment 6 with seta; one element; none larger than segment; straight; without spinules; without vestigial seta; without conical seta; without modified seta; without spinous process; without aesthetasc. Actual segment 7 with seta; one element; one larger than segment; straight; surpassing to distal margin; beyond three sequential segments; without spinules; without vestigial seta; without conical seta; without modified seta; without spinous process; with aesthetasc; one element. Actual segment 8 with seta; one element; one larger than segment; straight; surpassing distal margin; without spinules; without vestigial seta; with conical seta; without modified seta; without spinous process; without aesthetasc. Actual segment 9 with seta; two elements; of unequal size; one larger than segment; straight; surpassing to distal margin; beyond three sequential segments; without spinules; without vestigial seta; without conical seta; without modified seta; without spinous process; with aesthetasc; one element. Actual segment 10 with seta; one element; none larger than segment; straight; without spinules; without vestigial seta; without conical seta; without modified seta; without spinous process; without aesthetasc. Actual segment 11 with seta; one element; one larger than segment; straight; surpassing to distal margin; beyond three sequential segments; without spinules; without vestigial seta; without conical seta; without modified seta; without spinous process; without aesthetasc. Actual segment 12 with seta; one element; one larger than segment; straight; surpassing distal margin; without spinules; without vestigial seta; with conical seta; without modified seta; without spinous process; with aesthetasc; one element. Actual segment 13 with seta; one element; none elongated; straight; surpassing distal margin; without spinules; without vestigial seta; without conical seta; without modified seta; without spinous process; without aesthetasc. Actual segment 14 with seta; one element; elongated; straight; surpassing to distal margin; beyond three sequential segments; without spinules; without vestigial seta; without conical seta; without modified seta; without spinous process; with aesthetasc; one element. Actual segment 15 with seta; one element; larger than segment; straight; surpassing to distal margin; not beyond three sequential segments; without spinules; without vestigial seta; without conical seta; without modified seta; without spinous process; without aesthetasc. Actual segment 16 with seta; one element; larger than segment; plumose; surpassing to distal margin; not beyond three sequential segments; without spinules; without vestigial seta; without conical seta; without modified seta; without spinous process; with aesthetasc; one element. Actual segment 17 with seta; one element; not larger than segment; straight; without spinules; without vestigial seta; without conical seta; without modified seta; without spinous process; without aesthetasc. Actual segment 18 with seta; one element; larger than segment; straight; surpassing to distal margin; beyond three sequential segments; without spinules; without vestigial seta; without conical seta; without modified seta; without spinous process; without aesthetasc. Actual segment 19 with seta; one element; not larger than segment; straight; surpassing distal margin; without spinules; without vestigial seta; without conical seta; without modified seta; without spinous process; with aesthetasc; one element. Actual segment 20 with seta; one element; not larger than segment; straight; surpassing distal margin; without spinules; without vestigial seta; without conical seta; without modified seta; without spinous process; without aesthetasc. Actual segment 21 with seta; one element; larger than segment; plumose; surpassing to distal margin; beyond three sequential segments; without spinules; without vestigial seta; without conical seta; without modified seta; without spinous process; without aesthetasc. Actual segment 22 with seta; two elements; of unequal size; one of them elongated; plumose; surpassing to distal margin; without spinules; without vestigial seta; without conical seta; without modified seta; without spinous process; without aesthetasc. Actual segment 23 with seta; two elements; of unequal size; one larger than segment; plumose; surpassing to distal margin; greater 3x than original segment; without spinules; without vestigial seta; without conical seta; without modified seta; without spinous process; without aesthetasc. Actual segment 24 with seta; two elements; of equal size; one larger than segment; plumose; surpassing to distal margin; greater 3x than original segment; without spinules; without vestigial seta; without conical seta; without modified seta; without spinous process; without aesthetasc. Actual segment 25 with seta; five elements; of equal size; elongated; plumose; surpassing to distal margin; 4 times larger than segment; without spinules; without vestigial seta; without conical seta; without modified seta; without spinous process; with aesthetasc; one element.

##### Antenna

Biramous. Antenna coxa separated from the basis; bearing seta; 1; on inner surface; at distal corner; reaching to the middle base. Antenna basis (fusion) separated from the endopodal segment; bearing seta; 2; on inner surface; at distal corner. Endopodal ancestral segment I and II separated. Ancestral segment II and III fused. Ancestral segment III and IV fused. Ancestral segment III and IV fully. Antenna endopod actual 2-segmented. Actual segment 1 not bilobate; with seta; two; on inner margin; with spinules; as a row; obliquely; on outer surface; with pore. Actual segment 2 bilobate; without discontinuity on outer cuticle; inner lobe bearing 8 setae; distally; outer lobe bearing 7 setae; distally; with spinules; as a patch; on outer surface. Antenna exopod ancestral segment I and II separated. Ancestral segment II and III fused. Ancestral segment III and IV fused. Ancestral segment IV and V separated. Ancestral segment V and VI separated. Ancestral segment VI and VII separated. Ancestral segment VII and VIII separated. Ancestral segment VIII and IX separated. Ancestral segment IX and X fused. Antenna exopod actual 7-segmented. Actual segment 1 single; elongated (width-length, equal or larger ratio 2:1); with seta; one; at inner surface. Actual segment 2 compound; elongated (larger width-length ratio 2:1); with seta; three; at inner surface. Actual segment 3 single; not elongated (lesser width-length ratio 2:1); with seta; one; at inner surface. Actual segment 4 single; not elongated (lesser width-length ratio 2:1); with seta; one; at inner surface. Actual segment 5 single; not elongated (lesser width-length ratio 2:1); with seta; one; at inner surface. Actual segment 6 single; not elongated (lesser width-length ratio 2:1); with seta; one; at inner surface. Actual segment 7 compound; elongated (larger or equal width-length ratio 2:1); with seta; one; at inner surface; and three; at distal surface.

##### Oral features

**Mandible**. Coxal gnathobase sclerotized; with lobe; prominent; on caudal margin; presence of cutting blade; with tooth-like prominence; two, distinctly; 1 acute; on caudal margin; and 1 triangular; on sub-caudal margin; without acute projection between the prominences; with additional spinules; as a row; on dorsal surface; with seta; 1; dorsally; on apical surface; with spinules; apicalmost. Mandible palps biramous; comprising the basis; with seta; four; differently inserted; first medially; reaching to beyond the endopod 1; second distally; third distally; fourth distally; on inner margin; none with setulose ornamentation. Mandible endopod 2-segmented. Mandible endopod 1 with lobe; bearing seta; four; distally inserted; without spinules. Mandible endopod 2 without lobe; bearing setae; nine elements; distally inserted; with spinules; as a row; double. Mandible exopod 4-segmented. Mandible exopod 1 with seta; one element; distally; on inner margin. Mandible exopod 2 with seta; one element; distally; on inner side. Mandible exopod 3 with seta; one element; distally; on inner side. Mandible exopod 4 with setae; three elements; on terminal region. **Maxillule**. Birramous. Maxillule 3-segmented. Maxillule praecoxa with praecoxal arthrite; bearing spines; fifteen elements; ten marginally; plus, five sub-marginally; with spinules; as a patch; on sub-marginal surface. Maxillule coxa with coxal epipodite; with conspicuous outer lobe; bearing setae; nine elements; with coxal endite; elongated (larger or equal width-length ratio 2:1); bearing setae; four elements. Maxillule basis with basal endite; double; first proximal; elongated (larger width-length ratio 2:1; separated from basis; with setae; four elements; distally inserted; second distal; fused to basis; not elongated (lesser width-length ratio 2:1); with setae; four elements; distally inserted; with setules; as a row; on inner side; basal exite present; with setae; one element; on outer surface. Maxillule endopod 1-segmented. Endopod 1 bilobate; first proximal; with setae; three elements; second distal; with setae; five elements. Maxillule exopod 1-segmented. Exopod 1 with setae; six elements; with setules; as a row; on inner side; spinules absent. **Maxilla**. Uniramous. Maxilla 5-segmented. Maxilla praecoxa fused to coxa; incompletely; distinct externally; with praecoxal endite; double; first elongated endite (larger or equal width length ratio 2:1); proximally inserted; with seta; straight, or plumose; 1 straight; 4 plumose; with spine; single; without spinules; without setule; second elongated endite (larger or equal width length ratio 2:1); distally inserted; with seta; plumose; 3 plumose; without spine; with spinules; as a row; on distal margin; with setule; as a row; on distal margin; absence of outer seta. Maxilla coxa with coxal endite; double; first elongated endite (larger or equal width); proximally inserted; with seta; plumose; 3 plumose; without spine; without spinules; with setules; as a row; on proximal margin; second elongated endite (larger or equal width); distally inserted; with seta; plumose; 3 plumose; without spine; without spinules; with setules; as a row; on proximal margin; absence of outer seta. Maxilla basis with basal endite; single; elongated (larger or equal width-length ratio 2:1); with seta; plumose; 3 plumose; without spinules; absence of outer seta. Maxilla endopod 2-segmented. Endopod 1 with seta; 2 plumose; without spine; without spinules; without setules. Maxilla endopod 2 with seta; 2 plumose; without spine; without spinules; without setules. **Maxilliped**. Uniramous; Maxilliped 8-segmented. Maxilliped praecoxa fused to coxa; incompletely; distinct internally; with praecoxal endite; not elongated (lesser width-length ratio 2:1); distally inserted; with seta; 1 straight; with spinules; as a row; single; on basal surface; without setules. Maxilliped coxa with coxal endite; three coxal endite; first elongated (larger or equal width); proximally inserted; with seta; 2 plumose; with spinules; as a patch; single; on apical surface; without setules; second not elongated (lesser width-length ratio 2:1); medially inserted; with seta; 3 plumose; with spinules; as a row; single; on medial surface; without setules; third elongated (larger or equal width length ratio 2:1); distally inserted; with seta; 3 plumose; none reaching to beyond of the basis; with spinules; as a row; single; on basal surface; without setules; with lobe; prominence; at inner distal angle; ornamented; with spinules; continuously on margin. Maxilliped basis without basal endite; with seta; 3 plumose; with spinules; as a row; single; on medial surface; with setules; as a row; single; on inner margin. Maxilliped endopod segment 6-segmented. Endopod 1 with seta; 2 plumose; on inner surface. Endopod 2 with seta; 3 plumose; on inner surface. Endopod 3 with seta; 2 plumose; on inner surface. Endopod 4 with seta; 2 plumose; on inner surface. Endopod 5 with seta; 2 plumose; on inner surface, or on outer surface; outer seta absent. Endopod 6 with seta; 4 plumose; on inner surface, or on outer surface.

##### Swimming legs features

**First swimming legs.** Symmetrical; biramous. First swimming legs intercoxal plate without seta. First swimming legs praecoxa absent. First swimming legs coxa with seta; one; straight; distally inserted; on inner surface; surpassing to basal segment; with setules; one group; as a patch; on inner margin; without spinules; without spine. First swimming legs basis without seta; with setules; as a patch; single; on outer surface; without spinules; without spine. First swimming legs endopod 2-segmented. Endopod 1 with seta; straight; restricted; to inner surface; one element; without spine; with setules; as a row; single; continuously; on outer surface; without spinules; absence of Schmeil’s organ. Endopod 2 with seta; unrestricted; three on inner surface; one on outer surface; two on distal surface; straight; without spine; with setules; as a row; single; continuously; on outer surface; without spinules; absence of Schmeil’s organ. Endopod 3 absence. First swimming legs exopod 1 with seta; restricted; 1 on inner surface; with spine; 1; stout; smaller than original segment; serrated; on inner side; continuously; without setules. First swimming legs exopod 2 with seta; restricted; 1 on inner surface; straight; without spine; with setules; as a row; single; continuously; on inner margin, or on outer margin; without spinules. First swimming legs exopod 3 with setule; as a row; single; continuously; on outer surface; without spinules; with seta; unrestricted; 2 on inner surface; 2 on terminal surface; with spine; 2; unequal size; first no longer 2x than origin segment; stout; serrated; on inner side, or on outer side; equally; second longer 3x than origin segment; slender; serrated; on outer side; with ornamentation on non-serrated side; by setules. **Second swimming legs**. Symmetrical; Second swimming legs biramous. Second swimming legs intercoxal plate without seta. Second swimming legs praecoxa present; located laterally. Second swimming legs coxa with seta; straight; distally inserted; on inner surface; surpassing to basal segment; without setules; without spinules; without spine. Second swimming legs basis without seta; without setules; without spinules; without spine. Second swimming legs endopod 3-segmented. Endopod 1 with seta; straight; restricted; one on inner surface; without spine; with setules; as a row; single; continuously; on outer surface; without spinules; absence of Schmeil’s organ. Endopod 2 with seta; straight; unrestricted; two on inner surface; without spine; with setules; as a row; single; continuously; on outer side; without spinules; presence of Schmeil’s organ; on posterior surface. Endopod 3 with seta; plumose; unrestricted; three on inner surface; two on outer surface; two on distal surface; without spine; without setules; with spinules; as a row; double; distally inserted; at anterior surface; absence of Schmeil’s organ. Second swimming legs exopod 1 with seta; restricted; one on inner surface; with spine; 1; stout; not reaching to distal-third of the exopod 2; serrated; on inner side, or on outer side; with setules; as a row; single; continuously; on inner side; without spinules; absence of Schmeil’s organ. Exopod 2 with seta; unrestricted; one on inner surface; with spine; 1; stout; not surpassing the exopod 3; serrated; on inner side, or on outer side; with setules; as a row; single; continuously; on inner surface; without spinules; absence of Schmeil’s organ. Exopod 3 with seta; plurimarginal; three on inner surface; two on terminal surface; with spine; 2; unequal size; first no longer 2x than origin segment; stout; serrated; on inner side, or on outer side; equally; second longer 2x than origin segment; slender; serrated; on outer side; with ornamentation on non-serrated side; of setules; setules on outer surface; as a row; single; continuously; on inner surface; with spinules; as a row; single; distally inserted; at anterior surface; absence of Schmeil’s organ. **Third swimming legs**. Symmetrical; Third swimming legs biramous. Third swimming legs intercoxal plate without seta. Third swimming legs praecoxa present; not laterally located. Third swimming legs coxa with seta; straight; distally inserted; on inner surface; surpassing to basal segment; without setules; without spinules; without spine. Third swimming legs basis without seta; without setules; without spinules; without spine. Third swimming legs endopod 3-segmented. Endopod 1 with seta; restricted; one on inner surface; without spine; without setules; without spinules; absence of Schmeil’s organ. Endopod 2 with seta; restricted; two on inner surface; straight; without spine; without setules; without spinules; absence of Schmeil’s organ. Endopod 3 with seta; straight; plurimarginal; two on inner surface; two on outer surface; three on terminal surface; without spine; without setules; with spinules; as a row; distally inserted; double; at anterior surface; absence of Schmeil’s organ. Third swimming legs exopod 1 with seta; restricted; straight; one on inner surface; with spine; 1; stout; not reaching to the distal-third of the exopod 2; serrated; equally; on inner surface, or on outer surface; with setules; as a row; single; continuously; on inner surface; without spinules; absence of Schmeil’s organ. Exopod 2 with seta; straight; restricted; one on inner surface; with spine; 1; stout; not reaching out to exopod 3; serrated; on inner side, or on outer side; equally; with setules; as a row; single; continuously; on inner side; without spinules; absence of Schmeil’s organ. Exopod 3 without setules; with spinules; as a row; single; distally inserted; at anterior surface; with seta; straight; unrestricted; three on inner surface; two on terminal surface; with spine; 2; unequal size; first no longer 2x than origin segment; stout; serrated; on inner side, or on outer side; equally; second longer 2x than origin segment; slender; serrated; on outer side; with ornamentation on non-serrated side; of setules; absence of Schmeil’s organ. **Fourth swimming legs**. Symmetrical; biramous. Intercoxal plate without sensilla. Praecoxa present. Coxa with seta; distally inserted; on inner margin; reaching out to endopod 1; without spinules; setules absent. Basis with seta; one; medially inserted; on posterior surface; smaller than the original segment; without setules; without spinules; without spine. Fourth swimming legs endopod 3-segmented. Endopod 1 with seta; one; restricted; on inner surface; without spine; without setules; without spinules; absence of Schmeil’s organ. Endopod 2 with seta; restricted; two on inner side; without spine; with setules; as a row; single; continuously; on outer surface; without spinules; absence of Schmeil’s organ. Endopod 3 with seta; unrestricted; two on inner surface; two on outer surface; three on distal surface; without spine; without setules; with spinules; as a row; double; distally inserted; at anterior surface; absence of Schmeil’s organ. Fourth swimming legs exopod 1 with seta; restricted; one on inner surface; with spine; 1; stout; not reaching out to distal-third of the exopod 2; serrated; on inner side, or on outer side; equally; with setules; as a row; single; continuously; on inner surface; without spinules; absence of Schmeil’s organ. Exopod 2 with seta; restricted; one on inner surface; with spine; 1; stout; not reaching the end of exopod 3; serrated; on inner side, or on outer side; equally; with setules; as a row; single; continuously; on inner surface; without spinules; absence of Schmeil’s organ. Exopod 3 without setules; with spinules; as a row; single; distally inserted; at anterior surface; with seta; unrestricted; three on inner surface; two on distal surface; with spine; 2; unequal size; first no longer 2x than origin segment; stout; serrated; on inner side, or on outer side; equally; second longer 2x than origin segment; slender; serrated; on outer side; without ornamentation on non-serrated side; absence of Schmeil’s organ.

##### Fifth swimming legs features

Asymmetrical. Fifth swimming leg intercoxal plate with length not equal or greater than width on 1.5x; with irregular proximal margin; discontinuous to; the anterior margin of the left coxa, or the anterior margin of the right coxa; posterior sensilla on the right lateral absent. **Fifth left swimming leg**. Fifth left swimming leg biramous; leg reaching first right exopod segment; medially. Fifth left swimming leg praecoxa present; rudimentary; separated from the coxae; without ornamentation. Fifth left swimming leg coxa concave inner side; without teeth-like structures; with process; conical; on posterior surface; outer side; distally inserted; not projecting over basis; with sensilla; stout; triangular; at apex; no longer 2x than insertion basis; without swelling; without seta; without spinules. Fifth left swimming leg basis sub-cylindrical; unequal size between inner and outer side; shorter outer than inner side; with concave inner side; rounded internal proximal expansion absent; without outgrowth; with groove; deep; obliquely; on posterior surface; not reaching the endopodal lobe; not ornamented; absence of protuberance; with seta; outerly inserted; no longer 2x than origin segment; absence of minutely granular. Fifth left swimming leg endopod segments 1 and 2 fused; segments 2 and 3 fused; 1-segmented; stout; separated from the basis; ornamented; on inner side; with spinules; more than four elements; as a row; terminally; row of setules absent; without seta. Fifth left swimming leg exopod segments 1 and 2 separated; segments 2 and 3 fused; 2-segmented; stout; separated from the basis. Fifth left swimming leg exopod 1 sub-triangular; longer than broad; equal size between inner and outer side; rectilinear inner side; convex outer side; without swelling; without marginal extension; without process; with lobe; single; semicircular; medially inserted; on inner side; covered; by setules; absence seta. Fifth left swimming leg exopod 2 digitiform; longer than broad; equal size between inner and outer side; disform inner side; with rectilinear outer side; setulose pad present; prominently rounded; proximally; on inner side; inflated medial region present; setulose; anteriorly; distal process present; digitiform; denticulate; not bicuspidate; without transverse row of denticles; none oblique row of 5 denticles; at anterior surface; not innerly directed; with seta; spiniform; ornamented by spinules; surpassing the distal-point of the segment; without outer spine; terminal claw absent.

##### Fifth right swimming leg

Biramous. Fifth right swimming leg praecoxa present; separated from the coxae; without ornamentation. Fifth right swimming leg coxa convex inner side; without teeth-like structures; with process; rounded; distally inserted; on posterior surface; closest to the outer rim; projecting over basis; beyond the first third; until the medial surface; without triangular protuberance innerly; with sensilla; slender; at nadir; longer 2x than basal insertion; without marginal extension; without seta; without spinules. Fifth right swimming leg basis cylindrical; unequal size between inner and outer side; shorter outer than inner side; rectilinear inner side; tumescence present; not inflated; restricted on inner surface; proximally; without protuberance; absence of distinct minutely granular; additional inner process absent; without posterior groove; with seta; outerly inserted; on anterior surface; no longer 2x than origin segment; posterior protrusion present; distal tegument expansion present; triangular; anteriorly. Fifth right swimming leg with endopodite present; separated from the basis; on anterior surface; ancestral segments 1 and 2 fused; ancestral segments 2 and 3 fused; 1-segmented; stout; ornamented; with setules; as a row; on inner side; sub-terminally; without seta. Fifth right swimming leg exopod segments 1 and 2 separated; segments 2 and 3 fused; 2-segmented; stout; separated from the basis. Fifth right swimming leg exopod 1 sub-cylindrical; longer than broad; nearly 2 times; unequal size between both sides; shorter inner than outer side; convex inner side; rectilinear outer side; with marginal extension; sub-triangular; distally inserted; at outer rim; spinules absent; with process; rounded; sclerotized; without ornamentation; distally inserted; at posterior surface; projecting over next segment; without outer spine; without seta; internal prominence present; acute; lamella on posterior surface absent. Fifth right swimming leg exopod 2 cylindrical; longer than broad; nearly 2 times; equal size between both sides; disform inner side; convex outer side; without posterior proximal swelling; inner-posterior process absent; without marginal expansion; curved ridge on distal posterior surface present; chitinous knobs present; with 1–2 posteriorly; with outer spine; inserted sub-distally; arched; externally directed; ornamented innerly; by spinules; as a row; not ornamented outerly; sharp tip; with apparent curve; outerly directed; lesser than the length of the exopod 2; until to 2 times its size; 2x; sensilla absent; terminal claw present; equal or longer 1.5 times than insertion segment; sclerotized; arched; inward; with conspicuous curve; proximally; ornamented innerly; by spinules; as a row; partially on extension; medially, or distally; not ornamented outerly; sharp tip; curved tip; outwards; without medial constriction; hyaline process absent.

###### FEMALE

Body longer and wider than male; Female body 1121 micrometers excluding caudal setae. Widest at first metasome segment. Distal margin of the prosomal segments without one line of setules at posterior margin. Prosome segments with spinules at least at one prosomal segment. Fourth metasome segment presence of dorsal protuberance; digitiform; 2x wider than long; inserted medially; without posterior process; without anterior process; fourth metasome segment without proximal sensillae present. Fourth and fifth metasome segments fused; totally. Limit between fourth and fifth metasome segments without ornamentation. **Fifth metasome segment**. Fifth metasome segment without sensilla; with epimeral plates. Epimeral plates asymmetrical. Right epimeral plates prominent, as projections; thinner than the left; one posterior-laterally directed; not reaching half length of the genital segment; with sensilla at the apex; dorsal-posterior sensilla present; slender; without ornamentation. Left epimeral plate without expansion.

##### Urosome

3-segmented. **Genital double-somite**. Asymmetrical in dorsal view; longer than broad; longer than other urosomites combined; dorsal suture at mid-length absent; not covered by spinules; with swelling; rounded; equal size; anteriorly; with sensillae; on both sides; one; stout; with robust apex; at left lateral; not on lobular base; medially; one; stout; at right lateral; not on lobular base; anteriorly; with robust apex; of equal size between then; lateral protuberance absent; with right posterior rim expanded; over next segment; without slender sensilla on each posterior rim; without posterior-dorsal process. Genital double-somite opercular pad present; broader than longer; symmetrical; development laterally; expanded posteriorly; covering partially; double gonoporal slit; located ventrally; with arthrodial membrane; inserted anteriorly; post-genital process absent; disto-ventral tumescence absent; ventral vertical folds absent; dorsal sensilla absent. Second urosome segment with ventral fusion to anal segment; right distal process absent. Caudal rami patch of setules on outer surface absent; patch of spinules on outer surface absent.

##### Oral appendices feature

Rostrum basal process absent. **Antennules**. Symmetrical. Right antennule surpassing to genital double-segment; extending beyond caudal rami. Right antennule not exceeding the caudal setae. Right antennule ornamentation pattern equals to male left antennule; mostly. Actual segment 13 without seta; without aesthetasc. Actual segment 14 without seta; without aesthetasc. Actual segment 15 without seta; without aesthetasc. Actual segment 16 without seta; without aesthetasc. Actual segment 17 without seta. Actual segment 18 without seta.

##### Fifth swimming legs

Symmetrical; Fifth swimming legs biramous. Fifth swimming legs intercoxal plate longer than wide; separated from the legs. Fifth swimming legs praecoxa with sclerite praecoxal; separated from the coxae; without ornamentation. Fifth swimming legs coxa with process; conical; at the outer rim; distally; sensilla present; stout; at apex; projecting over basal segment; longer 2x than basal insertion; marginal extension absent; without swelling; without seta; without spinules. Fifth swimming legs basis sub-triangular; unequal size between inner and outer sides; shorter outer than inner side; with convex inner side; with proximal inner outgrowth; without groove; with distal extension; on posterior surface; with seta; outerly inserted; on anterior surface; no longer 2x than origin segment. Fifth swimming legs endopod segments 1 and 2 fused; segments 2 and 3 fused; 1-segmented; stout; separated from the basis; absent discontinuity cuticle; with spinules; as a row; single; non-oblique; sub-terminally; at anterior surface; with seta; double; one medially; on posterior surface; rectilinear; one distally; on posterior surface; rectilinear; of unequal size; distal seta longer than medial seta. Fifth swimming legs exopod segments 1 and 2 separated; segments 2 and 3 separated; 3-segmented; separated from the basis. Fifth swimming legs exopod 1 sub-cylindrical; longer than wide; longer or equal than 2 times; with unequal size between inner and outer side; shorter inner than outer side; with convex inner side; with rectilinear outer side; without swelling; without marginal extension; without posterior process; without spine; without seta. Fifth swimming legs exopod 2 sub-cylindrical; longer than broad; longer or equal than 2 times; without swelling; without marginal extension; without process; without lobe; with spine; inserted laterally; rectilinear; without ornamentation; sharp tip; smaller than next segment; without seta. Fifth swimming legs exopod 3 cylindrical; longer than wide; without swelling; without process; without lobe; without spine; with seta; double; inserted terminally; unequal size between them; outer seta smaller than inner; nearly 3 times; outer seta not ornamented by setules; without ornamentation; presence of terminal claw; sclerotized; arched; externally directed; convex inner side; with ornamentation; of denticles; as a row; on surface partially; at medial region; concave outer side; with ornamentation; of denticles; as a row; on surface partially; at medial region; blunt tip; 6 times longer than origin segment.

##### Distribution records

###### ARGENTINA

**Santa Fé**: La Capital, Madrejón Don Felipe; Arroyo Colorado; Río Coronda; Arroyo Ubajay; Sirgadero (Ringuelet and Ferrato, 1967); lagoon of the Naranjos (this study). Middle and lower stretch of the Paraguay River and Paraná River (Perbiche-Neves *et al*., 2015).

##### Habitat

Habitat in freshwaters: streams, ponds associated to rivers, and rivers.

##### Remarks

The taxonomic hypothesis was supported from organisms collected in North-Central Argentina, in lotic freshwater environments associated with rivers and streams. The morphology of the taxon was based on male attributes only, and exclusive illustrations were offered for fifth swimming legs posteriorly, fifth left swimming leg anteriorly, and right antennule actual segments between 7 to 16 (detail for segment 20 and 21). The morphology of the female was known only through the consistent effort of Paggi (1976) for specimens of the middle Paraná River, adding illustrations for the habit, fifth metasome segment and genital double-somite, fifth swimming legs, and caudal rami.

For Paggi (1976) the characteristic for the female fifth swimming legs endopod would be a definitive attribute for the inclusion of the species in *Notodiaptomus*, since it was the missing convergence between those identified for the male originally. Dussart & Frutos (1985) also contributed to morphological understandings of the species, adding illustrations of male fifth swimming legs posteriorly (detail for right leg), and male right antennule. In this effort, the authors illustrated male fifth right swimming leg coxa with a less prolonged outer conical process, and basis with posterior protrusion, both attributes not Illustrated by Ringuelet & Ferrato (1967).

Perbiche-Neves *et al*. (2015) when approaching specimens of the Prata River Basin, corroborated the observations of Dussart & Frutos (1985) and evidenced these characteristics in electronic microscopic photos. Additionally, the authors illustrated posterior protrusion on male fifth right swimming leg basis, a convergent characteristic in *Notodiaptomus* (Kiefer, 1936; 1936), and for type-species (Santos-Silva *et al*., 1999).

In this present effort we do not corroborate the original observation for male fifth right swimming leg basis with granulation on the base of the outer seta. In addition, we could not insert in our diagnosis of the male the fifth left swimming leg exopod 2 with “double digitiform process” such as described in Ringuelet & Ferrato (1967). Possibly the authors were referring to the distal digitiform process and the spiniform seta inserted more internally, both in the same segment. Furthermore, it is important to mention that in the work cited the equal size between both characters cannot be corroborated, having seen the spiniform seta being larger than the digitiform process, and surpassing to the distal-point of segment for the males examined.

For the male right antennule, the “spinous process” on the actual segments 8, 10, 11, and 13 of the geniculate antennule of the original male are, respectively, one conical seta for the first, and modified seta for the other segments, being for the 13th one robust modified seta with bifid apex. For the “small bump” indicated on actual segment 15 this is certainly a spinous process, also present on actual segment 16 of the same male antennule.

Finally, the Ringuelet & Ferrato hypothesis with female fourth metasome segment with dorsal protuberance in digitiform in *Notodiaptomus* is also suitable for organisms of *N. coniferoides*, *N. gibber*, and *N. transitans*, but can be distinguished through the fifth swimming legs endopod 1-segmented. For males, the size and form of the outer spine on fifth right swimming leg exopod is similar in the *N. dilatatus*, *N. dubius*, *N. kieferi*, *N. maracaibensis*, *N. nelsoni*, *N. orellanai*, *N. pseudodubius*, but can be distinguished through the male right antennule actual segment 1 with spinules, actual segment 8 with conical seta reaching to middle-point of sequent segment, and fifth right swimming leg exopod 1 longer than broad with internal acute prominence.

#### Notodiaptomus santaremensis (Wright, 1927)

##### Synonymy

*Diaptomus santaremensis* Wright, 1927: 75, 82, 100, 102, pl. 2, figs. 6–9; 1937a: 76; 1938b: 562. *Notodiaptomus santaremensis* n. comb., Kiefer, 1936a: 197; 1956: 242; Brehm, 1958a: 147; Brandorff, 1972: 45; Andrade & Brandorff, 1975: 97; Dussart & Defaye, 1983: 136; Robertson & Hardy, 1984: tab. 3; Santos-Silva *et al*., 1989: 726, 728, figs. 94–115; Rocha *et al*., 1995: 156; Santos-Silva, 1998: 211; Santos-Silva *et al*., 1999: 127; Santos-Silva *et al*., 2008: 33, fig. 7; Perbiche-Neves *et al*., 2020: 677-678, key to the Neotropical diaptomid, fig 21.6 H. “*Diaptomus*” *santaremensis*; Brandorff, 1976: 618, fig. 3; Löffler, 1981: 15; Reid, 1991: 737. *Notodiaptomus* (*Notodiaptomus*) *santaremensis*; Dussart, 1985a: 208.

##### Type locality

Lake near Santarem, Grão Pará, Brazil.

##### Type material

Holotype: female, entire in alcohol, stored in USNM under code 59514. No additional information is provided.

##### Material examined

Non-type material: 02 males, and 03 female, entire in alcohol, from the Curuá-Una Reservoir, Pará, no collector, VIII.1978, stored in Collection of zooplankton sample of the Plankton Laboratory, INPA, Brazil. 1 male (INPA-COP051, slides a-h) and 1 female (INPA-COP052, slides a-h) were selected to be dissection on eight slides each and deposited in the Zoological Collection of the INPA, Brazil.

##### Diagnosis

**(1)** Prosome widest at first metasome segment posteriorly; **(2)** cephalosome without dorsal suture; **(3)** male right antennule actual segment 15 and 16 without spinous process; **(4)** male fifth left swimming leg with length reaching first right exopod segment distally; **(5)** male fifth left swimming leg exopod 2 with spiniform seta not ornamented with spinules; **(6)** male fifth right swimming leg basis without posterior groove; **(8)** male fifth right swimming leg basis without posterior protrusion; **(9)** male fifth right swimming leg exopod 1 longer than broad 2x nearly; **(10)** male fifth right swimming leg exopod 2 in elliptical form; **(11)** female fourth metasome segment with digitiform dorsal protuberance 2x wider than long medially; **(12)** female left epimeral plate directed posterior-dorsally, reaching half length of the genital segment; **(13)** female right epimeral plate with sensilla on dorsal-posterior swelling, at the apex; **(14)** female second urosome with right distal process in lobular form posteriorly; **(15)** female caudal rami with setules patch on outer surface; **(16)** female right antennule with length not extending beyond caudal rami; **(17)** female right swimming legs exopod 3 with outer seta smaller than inner 1x nearly.

##### Redescription

###### MALE

Body 1096 micrometers excluding caudal setae. Male body smaller and slenderer than female. Nerve axons myelinated. Prosome 6-segmented; widest at first metasome segment; posteriorly; without one line of setules at posterior margin; without spinules at segments. Cephalosome anterior margin rounded; without dorsal suture; separate from first metasome segment. First metasome segment without sensilla. Second metasome segment with sensilla; 2 dorsally; of equal size. Third metasome segment with sensillae; 4 laterally; of unequal size; non-ornamented posterior margin. Fourth metasome segment with sensillae; 2 dorsally; 4 laterally; of equal size; separated from the fifth metasome. Limit between fourth and fifth metasome segments without ornamentation. Fifth metasome segment with sensilla; 4 laterally; Fifth metasome segment equal size; Fifth metasome segment without ornamentation; Fifth metasome segment without dorsal conical process; with epimeral plates. Epimeral plates symmetrical. Right epimeral plates reduced, as rounded distal corner segment limit; with sensilla; at the apex of projection; without ornamentation.

##### Urosome

5-segmented; Urosome 5 - free segments. Genital somite symmetrical in dorsal view; with single aperture; located on left side; ventrolaterally on posterior rim; with sensillae; on both sides; one; at left lateral; posteriorly; one; at right rim; posteriorly; of equal size between then. Third urosome segment without spinules; without external seta. Fourth urosome segment without spinules; without sub-conical blunt dorsal-lateral process. Anal segment presence of dorsal sensillae; one on each side; medially inserted; presence of operculum; convex; covering the anal aperture fully. Caudal rami symmetrical; separated from anal segment; longer than wide; with setules; continuous on; inner side; each ramus bearing 6 caudal setae; 5 marginals; plumose; and 1 internal dorsally; straight; not reticulated main axis; outermost seta with outer spiniform process absent.

##### Oral appendices feature

Rostrum symmetrical; separated from dorsal cephalic shield; by complete suture; sensillae present; one pair; anteriorly inserted on surface tegument; with rostral filament; double; paired; extended; into point; with basal process; in ventral view, rounded on left side; without a smaller basal expansion on the right side.

##### Antennules

Asymmetrical. **Right antennules**. Uniramous; right antennule surpassing to genital segment; right antennule not extending beyond caudal rami.

Right antennule ancestral segment I and II separated. Ancestral segment II and III fused. Ancestral segment III and IV fused. Ancestral segment IV and V separated. Ancestral segment V and VI separated. Ancestral segment VI and VII separated. Ancestral segment VII and VIII separated. Ancestral segment VIII and IX separated. Ancestral segment IX and X separated. Ancestral segment X and XI separated. Ancestral segment XI and XII separated. Ancestral segment XII and XIII separated. Ancestral segment XIII and XIV separated. Ancestral segment XIV and XV separated. Ancestral segment XV and XVI separated. Ancestral segment XVI and XVII separated. Ancestral segment XVII and XVIII separated. Ancestral segment XVIII and XIX separated. Ancestral segment XIX and XX separated. Ancestral segment XX and XXI separated. Ancestral segment XXI and XXII fused. Ancestral segment XXII and XXIII fused. Ancestral segment XXIII and XXIV separated. Ancestral segment XXIV and XXV fused. Ancestral segment XXV and XXVI separated. Ancestral segment XXVI and XXVII separated. Ancestral segment XXVII and XXVIII fused.

Right antennule actual 22-segmented; geniculated; between the segment 18 and segment 19; with swollen and modified region; formed by 5 segments; between 13 and 17 segments. Actual segment 1 with seta; one element; straight; none larger than segment; without spinules; without vestigial seta; without conical seta; without modified seta; without spinous process; with aesthetasc; one element. Actual segment 2 with seta; three elements; of unequal size; straight; none larger than segment; without spinules; with vestigial seta; one element; without conical seta; without modified seta; without spinous process; with aesthetasc; one element. Actual segment 3 with seta; one element; one larger than segment; surpassing to distal margin; beyond three sequential segments; straight; blunt apex; without spinules; with vestigial seta; one element; without conical seta; without modified seta; without spinous process; with aesthetasc. Actual segment 4 with seta; one element; one larger than segment; surpassing to distal margin; straight; not beyond three sequential segments; without spinules; without vestigial seta; without conical seta; without modified seta; without spinous process; without aesthetasc. Actual segment 5 with seta; one element; straight; one larger than segment; surpassing to distal margin; not beyond three sequential segments; without spinules; with vestigial seta; one element; without conical seta; without modified seta; without spinous process; with aesthetasc; one element. Actual segment 6 with seta; one element; none larger than segment; straight; without spinules; without vestigial seta; without conical seta; without modified seta; without spinous process; without aesthetasc. Actual segment 7 with seta; one element; straight; one larger than segment; surpassing to distal margin; beyond three sequential segments; blunt apex; without spinules; without vestigial seta; without conical seta; without modified seta; without spinous process; with aesthetasc; one element. Actual segment 8 with seta; one element; straight; none larger than segment; without spinules; without vestigial seta; with conical seta; one element; reaching to middle-point of the sequent segment; without modified seta; without spinous process; without aesthetasc. Actual segment 9 with seta; two elements; of unequal size; straight; one larger than segment; surpassing to distal margin; beyond three sequential segments; blunt apex; without spinules; without vestigial seta; without conical seta; without modified seta; without spinous process; with aesthetasc; one element. Actual segment 10 with seta; one element; straight; none larger than segment; without spinules; without vestigial seta; without conical seta; with modified seta; presenting blunt apex; slender form; surpassing to distal margin; beyond of the sequential segment; parallel to antennule direction; without spinous process; without aesthetasc. Actual segment 11 with seta; one element; straight; one larger than segment; surpassing to distal margin; not beyond three sequential segments; without spinules; without vestigial seta; without conical seta; with modified seta; slender form; presenting blunt apex; surpassing to distal margin; beyond of the sequential segment; parallel to antennule direction; shorter length than homologous of actual segment 13; without spinous process; without aesthetasc. Actual segment 12 with seta; one element; straight; one larger than segment; surpassing to distal margin; not beyond three sequential segments; without spinules; without vestigial seta; with conical seta; one element; smaller than to segment 8; without modified seta; without spinous process; with aesthetasc; one element; absent internal perpendicular fission. Actual segment 13 with seta; one element; straight; one larger than segment; surpassing to distal margin; not beyond three sequential segments; without spinules; without vestigial seta; without conical seta; with modified seta; stout form; surpassing to distal margin; to the distal-point of the sequence segment; perpendicular to antennule direction; presenting bifid apex; without spinous process; with aesthetasc; one element. Actual segment 14 with seta; two elements; of unequal size; straight; one larger than segment; surpassing to distal margin; beyond three sequential segments; blunt apex; without spinules; without vestigial seta; without conical seta; without modified seta; without spinous process; with aesthetasc; one element. Actual segment 15 with seta; two elements; of unequal size; straight; not bifidform; none larger than segment; without spinules; without vestigial seta; without conical seta; without modified seta; without spinous process; with aesthetasc; one element. Actual segment 16 with seta; two elements; of unequal size; plumose; one larger than segment; surpassing to distal margin; not beyond three sequential segments; not bifidform; without spinules; without vestigial seta; without conical seta; without modified seta; without spinous process; with aesthetasc; one element. Actual segment 17 with seta; two elements; of unequal size; straight; none larger than segment; bifidform; without spinules; without vestigial seta; without conical seta; with modified seta; one element; stout form; surpassing to distal margin; not beyond of the sequential segment; parallel to antennule direction; without spinous process; without aesthetasc. Actual segment 18 with seta; two elements; of equal size; straight; none larger than segment; without spinules; without vestigial seta; without conical seta; with modified seta; one element; stout form; surpassing distal margin; parallel to antennule direction; without spinous process; without aesthetasc. Actual segment 19 with seta; two elements; of unequal size; plumose; none larger than segment; without spinules; without vestigial seta; without conical seta; with modified seta; two elements; stout form; at least one bifid form; surpassing distal margin; parallel to antennule direction; without spinous process; with aesthetasc; one element. Actual segment 20 with seta; four elements; of unequal size; straight; one larger than segment; surpassing to distal margin; beyond three sequential segments; without spinules; without vestigial seta; without conical seta; without modified seta; with spinous process; medially; not reaching beyond of distal-point segment 21; without aesthetasc. Actual segment 21 with seta; two elements; of equal size; plumose; one larger than segment; surpassing to distal margin; greater 3x than original segment; without spinules; without vestigial seta; without conical seta; without modified seta; without spinous process; without aesthetasc. Actual segment 22 with seta; four elements; of equal size; one larger than segment; plumose; surpassing to distal margin; greater 3x than original segment; without spinules; without vestigial seta; without conical seta; without modified seta; without spinous process; with aesthetasc; one element.

##### Left antennules

Uniramous; Left antennule surpassing to prosome; Left antennule not extending beyond caudal rami. Ancestral segment I and II separated. Ancestral segment II and III fused. Ancestral segment III and IV fused. Ancestral segment IV and V separated. Ancestral segment V and VI separated. Ancestral segment VI and VII separated. Ancestral segment VII and VIII separated. Ancestral segment VIII and IX separated. Ancestral segment IX and X separated. Ancestral segment X and XI separated. Ancestral segment XI and XII separated. Ancestral segment XII and XIII separated. Ancestral segment XIII and XIV separated. Ancestral segment XIV and XV separated. Ancestral segment XV and XVI separated. Ancestral segment XVI and XVII separated. Ancestral segment XVII and XVIII separated. Ancestral segment XVIII and XIX separated. Ancestral segment XIX and XX separated. Ancestral segment XX and XXI separated. Ancestral segment XXI and XXII separated. Ancestral segment XXII and XXIII separated. Ancestral segment XXIII and XXIV separated. Ancestral segment XXIV and XXV separated. Ancestral segment XXV and XXVI separated. Ancestral segment XXVI and XXVII separated. Ancestral segment XXVII and XXVIII fused.

Left antennule actual 25-segmented; not-geniculated. Actual segment 1 with seta; one element; none larger than segment; straight; without spinules; without vestigial seta; without conical seta; without modified seta; without spinous process; with aesthetasc; one element. Actual segment 2 with seta; three elements; of equal size; none larger than segment; straight; without spinules; with vestigial seta; one element; without conical seta; without modified seta; without spinous process; with aesthetasc; one element. Actual segment 3 with seta; one element; one larger than segment; straight; surpassing to distal margin; beyond three sequential segments; without spinules; with vestigial seta; one element; without conical seta; without modified seta; without spinous process; with aesthetasc. Actual segment 4 with seta; one element; none larger than segment; straight; without spinules; without vestigial seta; without conical seta; without modified seta; without spinous process; without aesthetasc. Actual segment 5 with seta; one element; one larger than segment; straight; surpassing to distal margin; not beyond three sequential segments; without spinules; with vestigial seta; one element; without conical seta; without modified seta; without spinous process; with aesthetasc; one element. Actual segment 6 with seta; one element; none larger than segment; straight; without spinules; without vestigial seta; without conical seta; without modified seta; without spinous process; without aesthetasc. Actual segment 7 with seta; one element; one larger than segment; straight; surpassing to distal margin; beyond three sequential segments; without spinules; without vestigial seta; without conical seta; without modified seta; without spinous process; with aesthetasc; one element. Actual segment 8 with seta; one element; one larger than segment; straight; surpassing distal margin; without spinules; without vestigial seta; with conical seta; without modified seta; without spinous process; without aesthetasc. Actual segment 9 with seta; two elements; of unequal size; one larger than segment; straight; surpassing to distal margin; beyond three sequential segments; without spinules; without vestigial seta; without conical seta; without modified seta; without spinous process; with aesthetasc; one element. Actual segment 10 with seta; one element; none larger than segment; straight; without spinules; without vestigial seta; without conical seta; without modified seta; without spinous process; without aesthetasc. Actual segment 11 with seta; one element; one larger than segment; straight; surpassing to distal margin; beyond three sequential segments; without spinules; without vestigial seta; without conical seta; without modified seta; without spinous process; without aesthetasc. Actual segment 12 with seta; one element; one larger than segment; straight; surpassing distal margin; without spinules; without vestigial seta; with conical seta; without modified seta; without spinous process; with aesthetasc; one element. Actual segment 13 with seta; one element; none elongated; straight; surpassing distal margin; without spinules; without vestigial seta; without conical seta; without modified seta; without spinous process; without aesthetasc. Actual segment 14 with seta; one element; elongated; straight; surpassing to distal margin; beyond three sequential segments; without spinules; without vestigial seta; without conical seta; without modified seta; without spinous process; with aesthetasc; one element. Actual segment 15 with seta; one element; larger than segment; straight; surpassing to distal margin; not beyond three sequential segments; without spinules; without vestigial seta; without conical seta; without modified seta; without spinous process; without aesthetasc. Actual segment 16 with seta; one element; larger than segment; plumose; surpassing to distal margin; not beyond three sequential segments; without spinules; without vestigial seta; without conical seta; without modified seta; without spinous process; with aesthetasc; one element. Actual segment 17 with seta; one element; not larger than segment; straight; without spinules; without vestigial seta; without conical seta; without modified seta; without spinous process; without aesthetasc. Actual segment 18 with seta; one element; larger than segment; straight; surpassing to distal margin; beyond three sequential segments; without spinules; without vestigial seta; without conical seta; without modified seta; without spinous process; without aesthetasc. Actual segment 19 with seta; one element; not larger than segment; straight; surpassing distal margin; without spinules; without vestigial seta; without conical seta; without modified seta; without spinous process; with aesthetasc; one element. Actual segment 20 with seta; one element; not larger than segment; straight; surpassing distal margin; without spinules; without vestigial seta; without conical seta; without modified seta; without spinous process; without aesthetasc. Actual segment 21 with seta; one element; larger than segment; plumose; surpassing to distal margin; beyond three sequential segments; without spinules; without vestigial seta; without conical seta; without modified seta; without spinous process; without aesthetasc. Actual segment 22 with seta; two elements; of unequal size; one of them elongated; plumose; surpassing to distal margin; without spinules; without vestigial seta; without conical seta; without modified seta; without spinous process; without aesthetasc. Actual segment 23 with seta; two elements; of unequal size; one larger than segment; plumose; surpassing to distal margin; greater 3x than original segment; without spinules; without vestigial seta; without conical seta; without modified seta; without spinous process; without aesthetasc. Actual segment 24 with seta; two elements; of equal size; one larger than segment; plumose; surpassing to distal margin; greater 3x than original segment; without spinules; without vestigial seta; without conical seta; without modified seta; without spinous process; without aesthetasc. Actual segment 25 with seta; four elements; of equal size; elongated; plumose; surpassing to distal margin; 4 times larger than segment; without spinules; without vestigial seta; without conical seta; without modified seta; without spinous process; with aesthetasc; one element.

##### Antenna

Biramous. Antenna coxa separated from the basis; bearing seta; 1; on inner surface; at distal corner; reaching to the endopod 1. Antenna basis (fusion) separated from the endopodal segment; bearing seta; 2; on inner surface; at distal corner. Endopodal ancestral segment I and II separated. Ancestral segment II and III fused. Ancestral segment III and IV fused. Ancestral segment III and IV fully. Antenna endopod actual 2-segmented. Actual segment 1 not bilobate; with seta; two; on inner margin; with spinules; as a row; obliquely; on outer surface; with pore. Actual segment 2 bilobate; with discontinuity on outer cuticle; not developed as a suture; inner lobe bearing 8 setae; distally; outer lobe bearing 7 setae; distally; with spinules; as a patch; on outer surface. Antenna exopod ancestral segment I and II separated. Ancestral segment II and III fused. Ancestral segment III and IV fused. Ancestral segment IV and V separated. Ancestral segment V and VI separated. Ancestral segment VI and VII separated. Ancestral segment VII and VIII separated. Ancestral segment VIII and IX separated. Ancestral segment IX and X fused. Antenna exopod actual 7-segmented. Actual segment 1 single; elongated (width-length, equal or larger ratio 2:1); with seta; one; at inner surface. Actual segment 2 compound; elongated (larger width-length ratio 2:1); with seta; three; at inner surface. Actual segment 3 single; not elongated (lesser width-length ratio 2:1); with seta; one; at inner surface. Actual segment 4 single; not elongated (lesser width-length ratio 2:1); with seta; one; at inner surface. Actual segment 5 single; not elongated (lesser width-length ratio 2:1); with seta; one; at inner surface. Actual segment 6 single; not elongated (lesser width-length ratio 2:1); with seta; one; at inner surface. Actual segment 7 compound; elongated (larger or equal width-length ratio 2:1); with seta; one; at inner surface; and three; at distal surface.

##### Oral features

**Mandible**. Coxal gnathobase sclerotized; with lobe; prominent; on caudal margin; presence of cutting blade; with tooth-like prominence; two, distinctly; 1 acute; on caudal margin; and 1 triangular; on sub-caudal margin; without acute projection between the prominences; with additional spinules; as a row; on dorsal surface; with seta; 1; dorsally; on apical surface; with spinules; apicalmost. Mandible palps biramous; comprising the basis; with seta; four; differently inserted; first medially; reaching to beyond the endopod 1; second distally; third distally; fourth distally; on inner margin; none with setulose ornamentation. Mandible endopod 2-segmented. Mandible endopod 1 with lobe; bearing seta; four; distally inserted; without spinules. Mandible endopod 2 without lobe; bearing setae; nine elements; distally inserted; with spinules; as a row; double. Mandible exopod 4-segmented. Mandible exopod 1 with seta; one element; distally; on inner margin. Mandible exopod 2 with seta; one element; distally; on inner side. Mandible exopod 3 with seta; one element; distally; on inner side. Mandible exopod 4 with setae; three elements; on terminal region. **Maxillule**. Birramous. Maxillule 3-segmented. Maxillule praecoxa with praecoxal arthrite; bearing spines; fifteen elements; ten marginally; plus, five sub-marginally; with spinules; as a patch; on sub-marginal surface. Maxillule coxa with coxal epipodite; with conspicuous outer lobe; bearing setae; nine elements; with coxal endite; elongated (larger or equal width-length ratio 2:1); bearing setae; four elements. Maxillule basis with basal endite; double; first proximal; elongated (larger width-length ratio 2:1; separated from basis; with setae; four elements; distally inserted; second distal; fused to basis; not elongated (lesser width-length ratio 2:1); with setae; four elements; distally inserted; with setules; as a row; on inner side; basal exite present; with setae; one element; on outer surface. Maxillule endopod 1-segmented. Endopod 1 bilobate; first proximal; with setae; three elements; second distal; with setae; five elements. Maxillule exopod 1-segmented. Exopod 1 with setae; six elements; with setules; as a row; on inner side; spinules absent. **Maxilla**. Uniramous. Maxilla 5-segmented. Maxilla praecoxa fused to coxa; incompletely; distinct externally; with praecoxal endite; double; first elongated endite (larger or equal width length ratio 2:1); proximally inserted; with seta; straight, or plumose; 1 straight; 4 plumose; with spine; single; without spinules; without setule; second elongated endite (larger or equal width length ratio 2:1); distally inserted; with seta; plumose; 3 plumose; without spine; with spinules; as a row; on distal margin; with setule; as a row; on distal margin; absence of outer seta. Maxilla coxa with coxal endite; double; first elongated endite (larger or equal width); proximally inserted; with seta; plumose; 3 plumose; without spine; without spinules; with setules; as a row; on proximal margin; second elongated endite (larger or equal width); distally inserted; with seta; plumose; 3 plumose; without spine; without spinules; with setules; as a row; on proximal margin; absence of outer seta. Maxilla basis with basal endite; single; elongated (larger or equal width-length ratio 2:1); with seta; plumose; 3 plumose; without spinules; absence of outer seta. Maxilla endopod 2-segmented. Endopod 1 with seta; 2 plumose; without spine; without spinules; without setules. Maxilla endopod 2 with seta; 2 plumose; without spine; without spinules; without setules. **Maxilliped**. Uniramous; Maxilliped 8-segmented. Maxilliped praecoxa fused to coxa; incompletely; distinct internally; with praecoxal endite; not elongated (lesser width-length ratio 2:1); distally inserted; with seta; 1 straight; with spinules; as a row; single; on basal surface; without setules. Maxilliped coxa with coxal endite; three coxal endite; first elongated (larger or equal width); proximally inserted; with seta; 2 plumose; with spinules; as a patch; single; on apical surface; without setules; second not elongated (lesser width-length ratio 2:1); medially inserted; with seta; 3 plumose; with spinules; as a row; single; on medial surface; without setules; third elongated (larger or equal width length ratio 2:1); distally inserted; with seta; 3 plumose; none reaching to beyond of the basis; with spinules; as a row; single; on basal surface; without setules; with lobe; prominence; at inner distal angle; ornamented; with spinules; continuously on margin. Maxilliped basis without basal endite; with seta; 3 plumose; with spinules; as a row; single; on medial surface; with setules; as a row; single; on inner margin. Maxilliped endopod segment 6-segmented. Endopod 1 with seta; 2 plumose; on inner surface. Endopod 2 with seta; 3 plumose; on inner surface. Endopod 3 with seta; 2 plumose; on inner surface. Endopod 4 with seta; 2 plumose; on inner surface. Endopod 5 with seta; 2 plumose; on inner surface, or on outer surface; outer seta absent. Endopod 6 with seta; 4 plumose; on inner surface, or on outer surface.

##### Swimming legs features

**First swimming legs.** Symmetrical; biramous. First swimming legs intercoxal plate without seta. First swimming legs praecoxa absent. First swimming legs coxa with seta; one; straight; distally inserted; on inner surface; surpassing to first endopodal segment; with setules; two group; as a patch; on inner margin; and as a row; double; on anterior surface; outerly; without spinules; without spine. First swimming legs basis without seta; with setules; as a patch; single; on outer surface; without spinules; without spine. First swimming legs endopod 2-segmented. Endopod 1 with seta; straight; restricted; to inner surface; one element; without spine; with setules; as a row; single; continuously; on outer surface; without spinules; absence of Schmeil’s organ. Endopod 2 with seta; unrestricted; three on inner surface; one on outer surface; two on distal surface; straight; without spine; with setules; as a row; single; continuously; on outer surface; without spinules; absence of Schmeil’s organ. Endopod 3 absence. First swimming legs exopod 1 with seta; restricted; 1 on inner surface; with spine; 1; stout; smaller than original segment; serrated; on inner side; continuously; with setules; as a row; single; as a row; innerly. First swimming legs exopod 2 with seta; restricted; 1 on inner surface; straight; without spine; with setules; as a row; single; continuously; on inner margin, or on outer margin; without spinules. First swimming legs exopod 3 with setule; as a row; single; continuously; on outer surface; without spinules; with seta; unrestricted; 2 on inner surface; 2 on terminal surface; with spine; 2; unequal size; first no longer 2x than origin segment; stout; serrated; on inner side, or on outer side; equally; second longer 3x than origin segment; slender; serrated; on outer side; with ornamentation on non-serrated side; by setules. **Second swimming legs**. Symmetrical; Second swimming legs biramous. Second swimming legs intercoxal plate without seta. Second swimming legs praecoxa present; located laterally. Second swimming legs coxa with seta; straight; distally inserted; on inner surface; surpassing to basal segment; without setules; without spinules; without spine. Second swimming legs basis without seta; without setules; without spinules; without spine. Second swimming legs endopod 3-segmented. Endopod 1 with seta; straight; restricted; one on inner surface; without spine; with setules; as a row; single; continuously; on outer surface; without spinules; absence of Schmeil’s organ. Endopod 2 with seta; straight; unrestricted; two on inner surface; without spine; with setules; as a row; single; continuously; on outer side; without spinules; presence of Schmeil’s organ; on posterior surface. Endopod 3 with seta; straight; unrestricted; three on inner surface; two on outer surface; two on distal surface; without spine; without setules; with spinules; as a row; double; distally inserted; at anterior surface; absence of Schmeil’s organ. Second swimming legs exopod 1 with seta; restricted; one on inner surface; with spine; 1; stout; not reaching to distal-third of the exopod 2; serrated; on inner side, or on outer side; with setules; as a row; single; continuously; on inner side; without spinules; absence of Schmeil’s organ. Exopod 2 with seta; unrestricted; one on inner surface; with spine; 1; stout; not surpassing the exopod 3; serrated; on inner side, or on outer side; with setules; as a row; single; continuously; on inner surface; without spinules; absence of Schmeil’s organ. Exopod 3 with seta; plurimarginal; three on inner surface; two on terminal surface; with spine; 2; unequal size; first no longer 2x than origin segment; stout; serrated; on inner side, or on outer side; equally; second longer 2x than origin segment; slender; serrated; on outer side; with ornamentation on non-serrated side; of setules; setules on outer surface; as a row; single; continuously; on inner surface; with spinules; as a row; single; distally inserted; at anterior surface; absence of Schmeil’s organ. **Third swimming legs**. Symmetrical; Third swimming legs biramous. Third swimming legs intercoxal plate without seta. Third swimming legs praecoxa present; not laterally located. Third swimming legs coxa with seta; straight; distally inserted; on inner surface; surpassing to first endopodal segment; without setules; without spinules; without spine. Third swimming legs basis without seta; without setules; without spinules; without spine. Third swimming legs endopod 3-segmented. Endopod 1 with seta; restricted; one on inner surface; without spine; without setules; without spinules; absence of Schmeil’s organ. Endopod 2 with seta; restricted; two on inner surface; straight; without spine; without setules; without spinules; absence of Schmeil’s organ. Endopod 3 with seta; straight; plurimarginal; two on inner surface; two on outer surface; three on terminal surface; without spine; without setules; with spinules; as a row; distally inserted; double; at anterior surface; absence of Schmeil’s organ. Third swimming legs exopod 1 with seta; restricted; straight; one on inner surface; with spine; 1; stout; not reaching to the distal-third of the exopod 2; serrated; equally; on inner surface, or on outer surface; with setules; as a row; single; continuously; on inner surface; without spinules; absence of Schmeil’s organ. Exopod 2 with seta; straight; restricted; one on inner surface; with spine; 1; stout; not reaching out to exopod 3; serrated; on inner side, or on outer side; equally; with setules; as a row; single; continuously; on inner side; without spinules; absence of Schmeil’s organ. Exopod 3 without setules; with spinules; as a row; single; distally inserted; at anterior surface; with seta; straight; unrestricted; three on inner surface; two on terminal surface; with spine; 2; unequal size; first no longer 2x than origin segment; stout; serrated; on inner side, or on outer side; equally; second longer 2x than origin segment; slender; serrated; on outer side; with ornamentation on non-serrated side; of setules; absence of Schmeil’s organ. **Fourth swimming legs**. Symmetrical; biramous. Intercoxal plate without sensilla. Praecoxa present. Coxa with seta; distally inserted; on inner margin; reaching out to endopod 1; without spinules; setules absent. Basis with seta; one; medially inserted; on posterior surface; smaller than the original segment; without setules; without spinules; without spine. Fourth swimming legs endopod 3-segmented. Endopod 1 with seta; one; restricted; on inner surface; without spine; without setules; without spinules; absence of Schmeil’s organ. Endopod 2 with seta; restricted; two on inner side; without spine; with setules; as a row; single; continuously; on outer surface; without spinules; absence of Schmeil’s organ. Endopod 3 with seta; unrestricted; two on inner surface; two on outer surface; three on distal surface; without spine; without setules; with spinules; as a row; double; distally inserted; at anterior surface; absence of Schmeil’s organ. Fourth swimming legs exopod 1 with seta; restricted; one on inner surface; with spine; 1; stout; not reaching out to distal-third of the exopod 2; serrated; on inner side, or on outer side; equally; with setules; as a row; single; continuously; on inner surface; without spinules; absence of Schmeil’s organ. Exopod 2 with seta; restricted; one on inner surface; with spine; 1; stout; not reaching the end of exopod 3; serrated; on inner side, or on outer side; equally; with setules; as a row; single; continuously; on inner surface; without spinules; absence of Schmeil’s organ. Exopod 3 without setules; with spinules; as a row; single; distally inserted; at anterior surface; with seta; unrestricted; three on inner surface; two on distal surface; with spine; 2; unequal size; first no longer 2x than origin segment; stout; serrated; on inner side, or on outer side; equally; second longer 2x than origin segment; slender; serrated; on outer side; without ornamentation on non-serrated side; absence of Schmeil’s organ.

##### Fifth swimming legs features

Asymmetrical. Fifth swimming leg intercoxal plate with length not equal or greater than width on 1.5x; with irregular proximal margin; discontinuous to; the anterior margin of the left coxa, or the anterior margin of the right coxa; posterior sensilla on the right lateral absent. **Fifth left swimming leg**. Fifth left swimming leg biramous; leg reaching first right exopod segment; distally. Fifth left swimming leg praecoxa present; rudimentary; separated from the coxae; without ornamentation. Fifth left swimming leg coxa concave inner side; without teeth-like structures; with process; conical; on posterior surface; outer side; distally inserted; not projecting over basis; with sensilla; stout; triangular; at apex; no longer 2x than insertion basis; without swelling; without seta; without spinules. Fifth left swimming leg basis sub-cylindrical; unequal size between inner and outer side; shorter outer than inner side; with concave inner side; rounded internal proximal expansion absent; without outgrowth; without groove; absence of protuberance; with seta; outerly inserted; no longer 2x than origin segment; absence of minutely granular. Fifth left swimming leg endopod segments 1 and 2 fused; segments 2 and 3 fused; 1-segmented; stout; separated from the basis; ornamented; on inner side; with spinules; more than four elements; as a row; terminally; row of setules absent; without seta. Fifth left swimming leg exopod segments 1 and 2 separated; segments 2 and 3 fused; 2-segmented; stout; separated from the basis. Fifth left swimming leg exopod 1 sub-triangular; longer than broad; equal size between inner and outer side; rectilinear inner side; convex outer side; without swelling; without marginal extension; without process; with lobe; single; semicircular; medially inserted; on inner side; covered; by setules; without outer spine; absence seta. Fifth left swimming leg exopod 2 digitiform; longer than broad; equal size between inner and outer side; disform inner side; with rectilinear outer side; setulose pad present; prominently rounded; medially; on inner side; inflated medial region absent; distal process present; digitiform; non denticulate; without transverse row of denticles; none oblique row of 5 denticles; not innerly directed; with seta; spiniform; not ornamented by spinules; not surpassing the distal-point of the segment; without outer spine; terminal claw absent.

##### Fifth right swimming leg

Biramous. Fifth right swimming leg praecoxa present; separated from the coxae; without ornamentation. Fifth right swimming leg coxa convex inner side; without teeth-like structures; with process; rounded; distally inserted; on posterior surface; closest to the outer rim; projecting over basis; not beyond the first third; without triangular protuberance innerly; with sensilla; slender; at nadir; no longer 2x than basal insertion; without marginal extension; without seta; without spinules. Fifth right swimming leg basis cylindrical; unequal size between inner and outer side; shorter outer than inner side; rectilinear inner side; tumescence present; not inflated; restricted on inner surface; proximally; without protuberance; absence of distinct minutely granular; additional inner process absent; without posterior groove; with seta; outerly inserted; on posterior surface; no longer 2x than origin segment; posterior protrusion absent; distal tegument expansion present; rounded; posteriorly. Fifth right swimming leg with endopodite present; separated from the basis; on anterior surface; ancestral segments 1 and 2 fused; ancestral segments 2 and 3 fused; 1-segmented; stout; ornamented; with setules; as a row; on inner side; terminally; without seta. Fifth right swimming leg exopod segments 1 and 2 separated; segments 2 and 3 fused; 2-segmented; stout; separated from the basis. Fifth right swimming leg exopod 1 sub-cylindrical; longer than broad; nearly 2 times; unequal size between both sides; shorter inner than outer side; convex inner side; rectilinear outer side; with marginal extension; sub-triangular; distally inserted; at outer rim; spinules absent; with process; rounded; sclerotized; without ornamentation; distally inserted; at posterior surface; projecting over next segment; without outer spine; without seta; internal prominence absent; lamella on posterior surface absent. Fifth right swimming leg exopod 2 elliptical; longer than broad; nearly 2 times; equal size between both sides; disform inner side; convex outer side; without posterior proximal swelling; inner-posterior process absent; without marginal expansion; curved ridge on distal posterior surface absent; chitinous knobs present; with 1–2 posteriorly; with outer spine; inserted sub-distally; arched; internally directed; ornamented innerly; by spinules; as a row; not ornamented outerly; sharp tip; without apparent curve; lesser than the length of the exopod 2; until to 2 times its size; 2x; sensilla absent; terminal claw present; not equal or longer 1.5 times than insertion segment; sclerotized; arched; inward; without conspicuous curve; ornamented innerly; by spinules; as a row; partially on extension; medially, or distally; not ornamented outerly; sharp tip; not curved tip; without medial constriction; hyaline process absent.

##### FEMALE

Body longer and wider than male; Female body 1139 micrometers excluding caudal setae. Widest at first metasome segment. Distal margin of the prosomal segments without one line of setules at posterior margin. Prosome segments without spinules at prosomal segments. Fourth metasome segment presence of dorsal protuberance; digitiform; 2x wider than long; inserted medially; without posterior process; without anterior process; fourth metasome segment without proximal sensillae present. Fourth and fifth metasome segments fused; totally. Limit between fourth and fifth metasome segments without ornamentation. **Fifth metasome segment**. Fifth metasome segment without sensilla; with epimeral plates. Epimeral plates asymmetrical. Right epimeral plates prominent, as projections; thinner than the left; one posterior-dorsally directed; reaching half length of the genital segment; with sensilla at the apex; dorsal-posterior sensilla present; stout; without ornamentation. Left epimeral plate without expansion.

##### Urosome

3-segmented. **Genital double-somite**. Asymmetrical in dorsal view; longer than broad; longer than other urosomites combined; dorsal suture at mid-length absent; not covered by spinules; with swelling; rounded; unequal size; greater left than right; anteriorly; with sensillae; on both sides; one; stout; with robust apex; at left lateral; not on lobular base; anteriorly; one; stout; at right lateral; not on lobular base; anteriorly; with robust apex; of equal size between then; lateral protuberance absent; without right posterior rim expanded; without slender sensilla on each posterior rim; without posterior-dorsal process. Genital double-somite opercular pad present; broader than longer; symmetrical; development laterally; expanded posteriorly; covering partially; double gonoporal slit; located ventrally; with arthrodial membrane; inserted anteriorly; post-genital process absent; disto-ventral tumescence absent; ventral vertical folds absent. Second urosome segment without ventral fusion to anal segment; right distal process present; lobed form; not elongated. Caudal rami patch of setules on outer surface present; patch of spinules on outer surface absent.

##### Oral appendices feature

Rostrum basal process absent. **Antennules**. Symmetrical. Right antennule surpassing to genital double-segment; not extending beyond caudal rami; ornamentation pattern equals to male left antennule; fully.

##### Fifth swimming legs

Symmetrical; Fifth swimming legs biramous. Fifth swimming legs intercoxal plate longer than wide; separated from the legs. Fifth swimming legs praecoxa with sclerite praecoxal; separated from the coxae; without ornamentation. Fifth swimming legs coxa with process; conical; at the outer rim; distally; sensilla present; stout; at apex; projecting over basal segment; no longer 2x than basal insertion; marginal extension absent; without swelling; without seta; without spinules. Fifth swimming legs basis sub-triangular; unequal size between inner and outer sides; shorter outer than inner side; with convex inner side; without proximal inner outgrowth; without groove; with distal extension; on posterior surface; with seta; outerly inserted; on anterior surface; no longer 2x than origin segment. Fifth swimming legs endopod segments 1 and 2 fused; segments 2 and 3 fused; 1-segmented; stout; separated from the basis; present discontinuity cuticle; on inner side; with spinules; as a row; single; non-oblique; sub-terminally; at anterior surface; with seta; double; one medially; on posterior surface; rectilinear; one distally; on posterior surface; arched; of unequal size; distal seta longer than medial seta. Fifth swimming legs exopod segments 1 and 2 separated; segments 2 and 3 separated; 3-segmented; separated from the basis. Fifth swimming legs exopod 1 sub-cylindrical; longer than wide; longer or equal than 2 times; with unequal size between inner and outer side; shorter inner than outer side; with convex inner side; with rectilinear outer side; without swelling; without marginal extension; without posterior process; without spine; without seta. Fifth swimming legs exopod 2 sub-cylindrical; longer than broad; longer or equal than 2 times; without swelling; without marginal extension; without process; without lobe; with spine; inserted laterally; rectilinear; without ornamentation; sharp tip; smaller than next segment; without seta. Fifth swimming legs exopod 3 cylindrical; longer than wide; without swelling; without process; without lobe; without spine; with seta; double; inserted terminally; unequal size between them; outer seta smaller than inner; nearly 1 time; outer seta not ornamented by setules; without ornamentation; presence of terminal claw; sclerotized; arched; externally directed; convex inner side; with ornamentation; of denticles; as a row; on surface partially; at medial region; concave outer side; with ornamentation; of denticles; as a row; on surface partially; at medial region; blunt tip; 6 times longer than origin segment.

##### Distribution records

###### BRAZIL

**Pará:** Lake near Santarém, Grão Pará (Wright, 1927); Curuá-Una Reservoir (Santos-Silva *et al*., 1989).

##### Habitat

Habitat in freshwaters: lakes, and reservoirs: lake, and reservoir.

##### Remarks

The species is originally described as “*Diaptomus santaremensis“* and has a controversial history. Wright (1935) in founding the *nordestinus* complex did not consider the taxon as an integral part. Kiefer (1936) in creating *Notodiaptomus* took as a basis the Wright’s complex and suggested this as a true *Notodiaptomus*. Wright (1937) in extending his species complex with two species proposed by Kiefer in *Notodiaptomus* was against including *D. santaremensis* as part of his complex. This seems to have influenced Kiefer’s initial position during the amplification of the genus in 1956 (Kiefer, 1956), as it failed to list *D. santaremensis* among the *Notodiaptomus*, and only suggested that it could be included in the group forwardly. Despite this, other works have come to mention *D. santaremensis* as “*Notodiaptomus santaremensis“* (*e.g.*, Brehm, 1958; Brandorff, 1972; Andrade and Brandorff, 1974; Brandorff, 1976; Santos-Silva *et al*., 1989; Santos-Silva, 2008) and, due to the initial relation to the genus (Kiefer, 1936), we consider it a *Notodiaptomus* for this effort.

Wright (1927) offered an identification key based on males and indicated as a differential attribute of the species male fifth right swimming leg exopod 1 longer than broad 2x. Additionally, in the course of the description other diagnostic features are presented for females: (1) fourth metasome segment with dorsal conical protuberance; (2) fifth metasome segment with epimeral plates asymmetrical, left side more prolonged as “an irregular cone” plus “short spine” on the apex and “spine” innerly of same size; (3) antennule reaching to the end of the second urosome segment; (4) genital double-somite broader than long; (5) second urosome segment with right distal process lobed form. For male mainly: (1) prosome wide between first metasome segment posteriorly; (2) right antennule actual segment 20 with hyaline “plate“; (3) right antennule actual segment 15 with “slender” process reaching to distal-point of actual segment 16; (4) fifth right swimming leg basis with irregular process “which extends downward“; (5) fifth right leg exopod 1 with “lateral border distinctly expanded“; (6) fifth right leg exopod 2 with outer spine inserted medially; (7) fifth left swimming leg coxa with a “long arm-like projection which is bifurcated at the end“; (8) fifth left swimming leg exopod 2 in digital form with spiniform seta; (9) Fifth Left Swimming Leg Endopod 2-segmented.

During our examinations we were able to partially corroborate the highlighted attributes, except for the female the presence of the spines on epimeral plates, that truly are sensillae. For the male attribute 3 cannot be corroborated, because the spinous process on actual segment 15 of the right antennule surpassing distal margin of original segment. Another important variation verified is for male fifth right swimming leg exopod 2 with outer spine sub-distally, not medially such as original description for attribute 6. Objectively, this outer spine is not close to the terminal claw as in the type-species of *Notodiaptomus* and other congeners but cannot be recognized as inserted medially on external margin of exopod 2. One last discrepancy is for characteristic 9, definitely identified for this thesis as fifth left swimming leg endopod 1-segmented. In the original attribute 4 for male, it is a rounded distal tegument expansion posteriorly, recognized in other congeners such as *N. nelsoni*, and *N. santafesinus*, for the first taxon anteriorly, for second in triangular form. This attribute is also identifiable for other taxonomic genera such as *Argyrodiaptomus bergi* (Richard 1897), albeit discreetly.

Santos-Silva *et al*. (1989) added details to the morphology of the species through illustrations of the habit, oral appendages, swimming legs, and male and female antennulae. It is interesting to note that some of the original attributes (*i.e*., for female 5; for male 3, 7, and 9) were not represented in this collaboration and we understand them as not corroborated by the authors. As during our examinations, Santos-Silva *et al*. (1989) did not record the condition for male 7, and 9, the fifth left swimming leg coxa does not have projection or element bifurcated on the apex.

We can reaffirm *N. santaremensis* as a legitimate member of *Notodiaptomus*. Its present characteristics are convergent to the morphology highlighted in the formation of the *nordestinus* complex (Wright, 1935; 1936; 1937). For *Notodiaptomus* (Kiefer, 1936; 1956) has some divergences, namely: (1) female fifth swimming legs endopod with spinules row not oblique; (2) male fifth right swimming leg base without posterior protrusion; and (3) male fifth right swimming leg exopod 2 without distal curved ridge innerly on posterior surface.

#### Notodiaptomus simillimus Cicchino, Santos-Silva & Robertson, 2001

##### Synonymy

*Notodiaptomus simillimus* Ciccino, Santos-Silva, and Robertson, 2001: 539–548, figs. 1–4, tab. 1–3; Perbiche-Neves *et al*., 2020: 683-684, key to the Neotropical diaptomid. *Rhacodiaptomus calatus*, partim (Brandorff 1973: 345, pl. 1, Fig. 4d, p1. 4, Fig. 2d, e, pl. 5, Fig. 1a, b, e). *Notodiaptomus coniferoides* (Dussart & Frutos, 1986: 243, 245, 246; 248: Figs. 10-12).

##### Type locality

Originally specified as coming from plankton samples from Atabapo River, near San Fernando de Atabapo, Amazonas State, Venezuela.

##### Type material

Holotype: female, dissected on slide, 24.II.1974, G. Colomine coll. Paratypes: 1 male, dissected on slide (UPEL), 10 female, and 14 male, entire in ethanol. All material stored in Universidad Pedagógica Libertador, UPEL.

##### Material examined

Non-type material: 2 females, and 2 males, entire in alcohol, from Calado Lake, Amazonas State, Brazil, 13.VIII.1941, H. Sioli coll.; 2 males, and 2 female, entire in alcohol, from Calado Lake, Amazonas State, Brazil, collected through IX.1983 to XII.1984. All examined material is stored in the Plankton Laboratory’s collection (code n°. 0004, and n°. 0015d). 1 male (INPA-COP053, slides a-h) and 1 female (INPA-COP054, slides a-h) were selected to be dissection on eight slides each and deposited in the Zoological Collection of the INPA, Brazil.

##### Diagnosis

**(1)** Male fifth left swimming leg endopod 2-segmented; **(2)** male fifth right swimming leg exopod 2 without curved ridge on distal posterior surface; **(3)** female genital double-somite with conical swollen laterally; **(4)** female genital double-somite with lateral rounded protuberance on the left side; **(5)** female genital double-somite without right posterior rim expanded; **(6)** female fifth swimming legs exopod 2 with lateral spine equal size than next segment.

##### Redescription

**MALE.** Body 1259 micrometers excluding caudal setae. Male body smaller and slenderer than female. Nerve axons myelinated. Prosome 6-segmented; widest at first metasome segment; without one line of setules at posterior margin; without spinules at segments. Cephalosome anterior margin sub-triangular; with dorsal suture; incomplete; separate from first metasome segment. First metasome segment without sensilla. Second metasome segment without sensilla. Third metasome segment without sensillae; non-ornamented posterior margin. Fourth metasome segment without sensillae; separated from the fifth metasome. Limit between fourth and fifth metasome segments without ornamentation. Fifth metasome segment with sensilla; 2 dorsally; Fifth metasome segment equal size; Fifth metasome segment without ornamentation; Fifth metasome segment without dorsal conical process; with epimeral plates. Epimeral plates asymmetrical. Right epimeral plates prominent, as projections; one projection; posterior-dorsally directed; not reaching half length of the genital segment; with sensilla; at the apex of projection; without ornamentation. Left epimeral plate prominent, as projection; one projection; posterior-dorsally directed; reaching half length of the genital segment; with sensillae; at the apex of projection; without ornamentation.

##### Urosome

5-segmented; Urosome 5 - free segments. Genital somite asymmetrical in dorsal view; with single aperture; located on left side; ventrolaterally on posterior rim; with sensillae; on single side; one; at right rim; posteriorly. Third urosome segment without spinules; without external seta. Fourth urosome segment without spinules; without sub-conical blunt dorsal-lateral process. Anal segment presence of dorsal sensillae; one on each side; medially inserted; presence of operculum; convex; not covering the anal aperture fully. Caudal rami symmetrical; separated from anal segment; longer than wide; with setules; continuous on; inner side; each ramus bearing 6 caudal setae; 5 marginals; plumose; and 1 internal dorsally; straight; not reticulated main axis; outermost seta with outer spiniform process absent.

##### Oral appendices feature

Rostrum asymmetrical; separated from dorsal cephalic shield; by complete suture; sensillae present; one pair; anteriorly inserted on surface tegument; with rostral filament; double; paired; extended; into point; with basal process; in ventral view, rounded on left side; without a smaller basal expansion on the right side.

##### Antennules

Asymmetrical. **Right antennules**. Uniramous; right antennule surpassing to genital segment; right antennule not extending beyond caudal rami.

Right antennule ancestral segment I and II separated. Ancestral segment II and III fused. Ancestral segment III and IV fused. Ancestral segment IV and V separated. Ancestral segment V and VI separated. Ancestral segment VI and VII separated. Ancestral segment VII and VIII separated. Ancestral segment VIII and IX separated. Ancestral segment IX and X separated. Ancestral segment X and XI separated. Ancestral segment XI and XII separated. Ancestral segment XII and XIII separated. Ancestral segment XIII and XIV separated. Ancestral segment XIV and XV separated. Ancestral segment XV and XVI separated. Ancestral segment XVI and XVII separated. Ancestral segment XVII and XVIII separated. Ancestral segment XVIII and XIX separated. Ancestral segment XIX and XX separated. Ancestral segment XX and XXI separated. Ancestral segment XXI and XXII fused. Ancestral segment XXII and XXIII fused. Ancestral segment XXIII and XXIV separated. Ancestral segment XXIV and XXV fused. Ancestral segment XXV and XXVI separated. Ancestral segment XXVI and XXVII separated. Ancestral segment XXVII and XXVIII fused.

Right antennule actual 22-segmented; geniculated; between the segment 18 and segment 19; with swollen and modified region; formed by 5 segments; between 13 and 17 segments. Actual segment 1 with seta; one element; straight; none larger than segment; without spinules; without vestigial seta; without conical seta; without modified seta; without spinous process; with aesthetasc; one element. Actual segment 2 with seta; three elements; of equal size; straight; none larger than segment; without spinules; with vestigial seta; one element; without conical seta; without modified seta; without spinous process; with aesthetasc; one element. Actual segment 3 with seta; one element; one larger than segment; surpassing to distal margin; beyond three sequential segments; straight; blunt apex; without spinules; with vestigial seta; one element; without conical seta; without modified seta; without spinous process; with aesthetasc; one element. Actual segment 4 with seta; one element; none larger than segment; straight; without spinules; without vestigial seta; without conical seta; without modified seta; without spinous process; without aesthetasc. Actual segment 5 with seta; one element; straight; one larger than segment; surpassing to distal margin; not beyond three sequential segments; without spinules; without vestigial seta; without conical seta; without modified seta; without spinous process; with aesthetasc; one element. Actual segment 6 with seta; one element; none larger than segment; straight; without spinules; without vestigial seta; without conical seta; without modified seta; without spinous process; without aesthetasc. Actual segment 7 with seta; one element; straight; one larger than segment; surpassing to distal margin; beyond three sequential segments; blunt apex; without spinules; without vestigial seta; without conical seta; without modified seta; without spinous process; with aesthetasc; one element. Actual segment 8 with seta; one element; straight; none larger than segment; without spinules; without vestigial seta; with conical seta; one element; not reaching to middle-point of the sequent segment; without modified seta; without spinous process; without aesthetasc. Actual segment 9 with seta; two elements; of unequal size; straight; one larger than segment; surpassing to distal margin; beyond three sequential segments; blunt apex; without spinules; without vestigial seta; without conical seta; without modified seta; without spinous process; with aesthetasc; one element. Actual segment 10 with seta; one element; straight; one larger than segment; surpassing distal margin; not beyond three sequential segments; without spinules; without vestigial seta; without conical seta; with modified seta; presenting blunt apex; slender form; surpassing to distal margin; beyond of the sequential segment; parallel to antennule direction; without spinous process; without aesthetasc. Actual segment 11 with seta; one element; straight; one larger than segment; surpassing to distal margin; not beyond three sequential segments; without spinules; without vestigial seta; without conical seta; with modified seta; slender form; presenting blunt apex; surpassing to distal margin; beyond of the sequential segment; parallel to antennule direction; shorter length than homologous of actual segment 13; without spinous process; without aesthetasc. Actual segment 12 with seta; one element; straight; one larger than segment; surpassing to distal margin; not beyond three sequential segments; without spinules; without vestigial seta; with conical seta; one element; not smaller than to segment 8; without modified seta; without spinous process; with aesthetasc; one element; absent internal perpendicular fission. Actual segment 13 with seta; one element; straight; none larger than segment; without spinules; without vestigial seta; without conical seta; with modified seta; stout form; surpassing to distal margin; to the distal point of the sequence segment; perpendicular to antennule direction; presenting bifid apex; without spinous process; with aesthetasc; one element. Actual segment 14 with seta; two elements; of unequal size; straight; one larger than segment; surpassing to distal margin; beyond three sequential segments; blunt apex; without spinules; without vestigial seta; without conical seta; without modified seta; without spinous process; with aesthetasc; one element. Actual segment 15 with seta; two elements; of unequal size; straight; bifidform; one larger than segment; surpassing to distal margin; not beyond three sequential segments; without spinules; without vestigial seta; without conical seta; without modified seta; with spinous process; on outer margin; surpassing distal margin; with aesthetasc; one element. Actual segment 16 with seta; two elements; of unequal size; straight; one larger than segment; surpassing to distal margin; not beyond three sequential segments; bifidform; without spinules; without vestigial seta; without conical seta; without modified seta; with spinous process; on outer margin; surpassing distal margin; unequal size to process on preceding segment; with aesthetasc; one element. Actual segment 17 with seta; two elements; of unequal size; straight; one larger than segment; surpassing distal margin; bifidform; without spinules; without vestigial seta; without conical seta; with modified seta; one element; stout form; surpassing to distal margin; not beyond of the sequential segment; parallel to antennule direction; without spinous process; without aesthetasc. Actual segment 18 with seta; two elements; of equal size; straight; none larger than segment; without spinules; without vestigial seta; without conical seta; with modified seta; one element; stout form; surpassing distal margin; parallel to antennule direction; without spinous process; without aesthetasc. Actual segment 19 with seta; two elements; of unequal size; plumose; none larger than segment; without spinules; without vestigial seta; without conical seta; with modified seta; two elements; stout form; at least one bifid form; surpassing distal margin; parallel to antennule direction; without spinous process; with aesthetasc; one element. Actual segment 20 with seta; four elements; of unequal size; plumose; one larger than segment; surpassing to distal margin; not beyond three sequential segments; without spinules; with vestigial seta; without conical seta; without modified seta; without spinous process; without aesthetasc. Actual segment 21 with seta; two elements; of equal size; plumose; one larger than segment; surpassing to distal margin; greater 3x than original segment; without spinules; without vestigial seta; without conical seta; without modified seta; without spinous process; without aesthetasc. Actual segment 22 with seta; four elements; of equal size; one larger than segment; plumose; surpassing to distal margin; greater 3x than original segment; without spinules; without vestigial seta; without conical seta; without modified seta; without spinous process; with aesthetasc; one element.

##### Left antennules

Uniramous; Left antennule surpassing to prosome; Left antennule not extending beyond caudal rami. Ancestral segment I and II separated. Ancestral segment II and III fused. Ancestral segment III and IV fused. Ancestral segment IV and V separated. Ancestral segment V and VI separated. Ancestral segment VI and VII separated. Ancestral segment VII and VIII separated. Ancestral segment VIII and IX separated. Ancestral segment IX and X separated. Ancestral segment X and XI separated. Ancestral segment XI and XII separated. Ancestral segment XII and XIII separated. Ancestral segment XIII and XIV separated. Ancestral segment XIV and XV separated. Ancestral segment XV and XVI separated. Ancestral segment XVI and XVII separated. Ancestral segment XVII and XVIII separated. Ancestral segment XVIII and XIX separated. Ancestral segment XIX and XX separated. Ancestral segment XX and XXI separated. Ancestral segment XXI and XXII separated. Ancestral segment XXII and XXIII separated. Ancestral segment XXIII and XXIV separated. Ancestral segment XXIV and XXV separated. Ancestral segment XXV and XXVI separated. Ancestral segment XXVI and XXVII separated. Ancestral segment XXVII and XXVIII fused.

Left antennule actual 25-segmented; not-geniculated. Actual segment 1 with seta; one element; none larger than segment; straight; without spinules; without vestigial seta; without conical seta; without modified seta; without spinous process; with aesthetasc; one element. Actual segment 2 with seta; three elements; of equal size; none larger than segment; straight; without spinules; with vestigial seta; one element; without conical seta; without modified seta; without spinous process; with aesthetasc; one element. Actual segment 3 with seta; one element; one larger than segment; straight; surpassing to distal margin; beyond three sequential segments; without spinules; with vestigial seta; one element; without conical seta; without modified seta; without spinous process; with aesthetasc; one element. Actual segment 4 with seta; one element; none larger than segment; straight; without spinules; without vestigial seta; without conical seta; without modified seta; without spinous process; without aesthetasc. Actual segment 5 with seta; one element; one larger than segment; straight; surpassing distal margin; not beyond three sequential segments; without spinules; with vestigial seta; one element; without conical seta; without modified seta; without spinous process; with aesthetasc; one element. Actual segment 6 with seta; one element; none larger than segment; straight; without spinules; without vestigial seta; without conical seta; without modified seta; without spinous process; without aesthetasc. Actual segment 7 with seta; one element; one larger than segment; straight; surpassing to distal margin; beyond three sequential segments; without spinules; without vestigial seta; without conical seta; without modified seta; without spinous process; with aesthetasc; one element. Actual segment 8 with seta; one element; one larger than segment; straight; surpassing distal margin; without spinules; without vestigial seta; with conical seta; without modified seta; without spinous process; without aesthetasc. Actual segment 9 with seta; two elements; of unequal size; one larger than segment; straight; surpassing to distal margin; beyond three sequential segments; without spinules; without vestigial seta; without conical seta; without modified seta; without spinous process; with aesthetasc; one element. Actual segment 10 with seta; one element; one larger than segment; straight; surpassing distal margin; without spinules; without vestigial seta; without conical seta; without modified seta; without spinous process; without aesthetasc. Actual segment 11 with seta; one element; one larger than segment; straight; surpassing to distal margin; beyond three sequential segments; without spinules; without vestigial seta; without conical seta; without modified seta; without spinous process; without aesthetasc. Actual segment 12 with seta; one element; one larger than segment; straight; surpassing distal margin; without spinules; without vestigial seta; with conical seta; without modified seta; without spinous process; with aesthetasc; one element. Actual segment 13 with seta; one element; none elongated; straight; surpassing distal margin; without spinules; without vestigial seta; without conical seta; without modified seta; without spinous process; without aesthetasc. Actual segment 14 with seta; one element; elongated; straight; surpassing to distal margin; beyond three sequential segments; without spinules; without vestigial seta; without conical seta; without modified seta; without spinous process; with aesthetasc; one element. Actual segment 15 with seta; one element; larger than segment; straight; surpassing to distal margin; not beyond three sequential segments; without spinules; without vestigial seta; without conical seta; without modified seta; without spinous process; without aesthetasc. Actual segment 16 with seta; one element; larger than segment; plumose; surpassing to distal margin; not beyond three sequential segments; without spinules; without vestigial seta; without conical seta; without modified seta; without spinous process; with aesthetasc; one element. Actual segment 17 with seta; one element; not larger than segment; straight; without spinules; without vestigial seta; without conical seta; without modified seta; without spinous process; without aesthetasc. Actual segment 18 with seta; one element; larger than segment; plumose; surpassing to distal margin; beyond three sequential segments; without spinules; without vestigial seta; without conical seta; without modified seta; without spinous process; without aesthetasc. Actual segment 19 with seta; one element; not larger than segment; straight; surpassing distal margin; without spinules; without vestigial seta; without conical seta; without modified seta; without spinous process; with aesthetasc; one element. Actual segment 20 with seta; one element; not larger than segment; straight; surpassing distal margin; without spinules; without vestigial seta; without conical seta; without modified seta; without spinous process; without aesthetasc. Actual segment 21 with seta; one element; larger than segment; plumose; surpassing to distal margin; beyond three sequential segments; without spinules; without vestigial seta; without conical seta; without modified seta; without spinous process; without aesthetasc. Actual segment 22 with seta; two elements; of equal size; one of them elongated; plumose; surpassing to distal margin; without spinules; without vestigial seta; without conical seta; without modified seta; without spinous process; without aesthetasc. Actual segment 23 with seta; one element; none larger than segment; straight; without spinules; without vestigial seta; without conical seta; without modified seta; without spinous process; without aesthetasc. Actual segment 24 with seta; two elements; of equal size; one larger than segment; plumose; surpassing to distal margin; greater 3x than original segment; without spinules; without vestigial seta; without conical seta; without modified seta; without spinous process; without aesthetasc. Actual segment 25 with seta; four elements; of equal size; elongated; plumose; surpassing to distal margin; 4 times larger than segment; without spinules; without vestigial seta; without conical seta; without modified seta; without spinous process; with aesthetasc; one element.

##### Antenna

Biramous. Antenna coxa separated from the basis; bearing seta; 1; on inner surface; at distal corner; reaching to the endopod 1. Antenna basis (fusion) separated from the endopodal segment; bearing seta; 2; on inner surface; at distal corner. Endopodal ancestral segment I and II separated. Ancestral segment II and III fused. Ancestral segment III and IV fused. Ancestral segment III and IV fully. Antenna endopod actual 2-segmented. Actual segment 1 not bilobate; with seta; two; on inner margin; with spinules; as a row; obliquely; on outer surface; with pore. Actual segment 2 bilobate; with discontinuity on outer cuticle; not developed as a suture; inner lobe bearing 8 setae; distally; outer lobe bearing 7 setae; distally; with spinules; as a patch; on outer surface. Antenna exopod ancestral segment I and II separated. Ancestral segment II and III fused. Ancestral segment III and IV fused. Ancestral segment IV and V separated. Ancestral segment V and VI separated. Ancestral segment VI and VII separated. Ancestral segment VII and VIII separated. Ancestral segment VIII and IX separated. Ancestral segment IX and X fused. Antenna exopod actual 7-segmented. Actual segment 1 single; elongated (width-length, equal or larger ratio 2:1); with seta; one; at inner surface. Actual segment 2 compound; elongated (larger width-length ratio 2:1); with seta; three; at inner surface. Actual segment 3 single; not elongated (lesser width-length ratio 2:1); with seta; one; at inner surface. Actual segment 4 single; not elongated (lesser width-length ratio 2:1); with seta; one; at inner surface. Actual segment 5 single; not elongated (lesser width-length ratio 2:1); with seta; one; at inner surface. Actual segment 6 single; not elongated (lesser width-length ratio 2:1); with seta; one; at inner surface. Actual segment 7 compound; elongated (larger or equal width-length ratio 2:1); with seta; one; at inner surface; and three; at distal surface.

##### Oral features

**Mandible**. Coxal gnathobase sclerotized; with lobe; prominent; on caudal margin; presence of cutting blade; with tooth-like prominence; two, distinctly; 1 acute; on caudal margin; and 1 triangular; on sub-caudal margin; without acute projection between the prominences; with additional spinules; as a row; on dorsal surface; with seta; 1; dorsally; on apical surface; with spinules; apicalmost. Mandible palps biramous; comprising the basis; with seta; four; differently inserted; first medially; reaching to beyond the endopod 1; second distally; third distally; fourth distally; on inner margin; none with setulose ornamentation. Mandible endopod 2-segmented. Mandible endopod 1 with lobe; bearing seta; four; distally inserted; without spinules. Mandible endopod 2 without lobe; bearing setae; nine elements; distally inserted; with spinules; as a row; double. Mandible exopod 4-segmented. Mandible exopod 1 with seta; one element; distally; on inner margin. Mandible exopod 2 with seta; one element; distally; on inner side. Mandible exopod 3 with seta; one element; distally; on inner side. Mandible exopod 4 with setae; three elements; on terminal region. **Maxillule**. Birramous. Maxillule 3-segmented. Maxillule praecoxa with praecoxal arthrite; bearing spines; fifteen elements; ten marginally; plus, five sub-marginally; with spinules; as a patch; on sub-marginal surface. Maxillule coxa with coxal epipodite; with conspicuous outer lobe; bearing setae; nine elements; with coxal endite; elongated (larger or equal width-length ratio 2:1); bearing setae; four elements. Maxillule basis with basal endite; double; first proximal; elongated (larger width-length ratio 2:1; separated from basis; with setae; four elements; distally inserted; second distal; fused to basis; not elongated (lesser width-length ratio 2:1); with setae; four elements; distally inserted; with setules; as a row; on inner side; basal exite present; with setae; one element; on outer surface. Maxillule endopod 1-segmented. Endopod 1 bilobate; first proximal; with setae; three elements; second distal; with setae; five elements. Maxillule exopod 1-segmented. Exopod 1 with setae; six elements; with setules; as a row; on inner side; spinules absent. **Maxilla**. Uniramous. Maxilla 5-segmented. Maxilla praecoxa fused to coxa; incompletely; distinct externally; with praecoxal endite; double; first elongated endite (larger or equal width length ratio 2:1); proximally inserted; with seta; straight, or plumose; 1 straight; 4 plumose; with spine; single; without spinules; without setule; second elongated endite (larger or equal width length ratio 2:1); distally inserted; with seta; plumose; 3 plumose; without spine; with spinules; as a row; on distal margin; with setule; as a row; on distal margin; absence of outer seta. Maxilla coxa with coxal endite; double; first elongated endite (larger or equal width); proximally inserted; with seta; plumose; 3 plumose; without spine; without spinules; with setules; as a row; on proximal margin; second elongated endite (larger or equal width); distally inserted; with seta; plumose; 3 plumose; without spine; without spinules; with setules; as a row; on proximal margin; absence of outer seta. Maxilla basis with basal endite; single; elongated (larger or equal width-length ratio 2:1); with seta; plumose; 3 plumose; without spinules; absence of outer seta. Maxilla endopod 2-segmented. Endopod 1 with seta; 2 plumose; without spine; without spinules; without setules. Maxilla endopod 2 with seta; 2 plumose; without spine; without spinules; without setules. **Maxilliped**. Uniramous; Maxilliped 8-segmented. Maxilliped praecoxa fused to coxa; incompletely; distinct internally; with praecoxal endite; not elongated (lesser width-length ratio 2:1); distally inserted; with seta; 1 straight; with spinules; as a row; single; on basal surface; without setules. Maxilliped coxa with coxal endite; three coxal endite; first elongated (larger or equal width); proximally inserted; with seta; 2 plumose; with spinules; as a patch; single; on apical surface; without setules; second not elongated (lesser width-length ratio 2:1); medially inserted; with seta; 3 plumose; with spinules; as a row; single; on medial surface; without setules; third elongated (larger or equal width length ratio 2:1); distally inserted; with seta; 3 plumose; none reaching to beyond of the basis; with spinules; as a row; single; on basal surface; without setules; with lobe; prominence; at inner distal angle; ornamented; with spinules; continuously on margin. Maxilliped basis without basal endite; with seta; 3 plumose; with spinules; as a row; single; on medial surface; with setules; as a row; single; on inner margin. Maxilliped endopod segment 6-segmented. Endopod 1 with seta; 2 plumose; on inner surface. Endopod 2 with seta; 3 plumose; on inner surface. Endopod 3 with seta; 2 plumose; on inner surface. Endopod 4 with seta; 2 plumose; on inner surface. Endopod 5 with seta; 2 plumose; on inner surface, or on outer surface; outer seta absent. Endopod 6 with seta; 4 plumose; on inner surface, or on outer surface.

##### Swimming legs features

**First swimming legs.** Symmetrical; biramous. First swimming legs intercoxal plate without seta. First swimming legs praecoxa absent. First swimming legs coxa with seta; one; straight; distally inserted; on inner surface; surpassing to first endopodal segment; with setules; two group; as a patch; on inner margin; and as a row; double; on anterior surface; outerly; without spinules; without spine. First swimming legs basis without seta; with setules; as a patch; single; on outer surface; without spinules; without spine. First swimming legs endopod 2-segmented. Endopod 1 with seta; straight; restricted; to inner surface; one element; without spine; with setules; as a row; single; continuously; on outer surface; without spinules; absence of Schmeil’s organ. Endopod 2 with seta; unrestricted; three on inner surface; one on outer surface; two on distal surface; straight; without spine; with setules; as a row; single; continuously; on outer surface; without spinules; absence of Schmeil’s organ. Endopod 3 absence. First swimming legs exopod 1 with seta; restricted; 1 on inner surface; with spine; 1; stout; smaller than original segment; serrated; on inner side; continuously; with setules; as a row; single; as a row; innerly. First swimming legs exopod 2 with seta; restricted; 1 on inner surface; straight; without spine; with setules; as a row; single; continuously; on inner margin, or on outer margin; without spinules. First swimming legs exopod 3 with setule; as a row; single; continuously; on outer surface; without spinules; with seta; unrestricted; 2 on inner surface; 2 on terminal surface; with spine; 2; unequal size; first no longer 2x than origin segment; stout; serrated; on inner side, or on outer side; equally; second longer 3x than origin segment; slender; serrated; on outer side; with ornamentation on non-serrated side; by setules. **Second swimming legs**. Symmetrical; Second swimming legs biramous. Second swimming legs intercoxal plate without seta. Second swimming legs praecoxa present; located laterally. Second swimming legs coxa with seta; straight; distally inserted; on inner surface; surpassing to basal segment; without setules; without spinules; without spine. Second swimming legs basis without seta; without setules; without spinules; without spine. Second swimming legs endopod 3-segmented. Endopod 1 with seta; straight; restricted; one on inner surface; without spine; with setules; as a row; single; continuously; on outer surface; without spinules; absence of Schmeil’s organ. Endopod 2 with seta; straight; unrestricted; two on inner surface; without spine; with setules; as a row; single; continuously; on outer side; without spinules; presence of Schmeil’s organ; on posterior surface. Endopod 3 with seta; straight; unrestricted; three on inner surface; two on outer surface; two on distal surface; without spine; without setules; with spinules; as a row; double; distally inserted; at anterior surface; absence of Schmeil’s organ. Second swimming legs exopod 1 with seta; restricted; one on inner surface; with spine; 1; stout; not reaching to distal-third of the exopod 2; serrated; on inner side, or on outer side; with setules; as a row; single; continuously; on inner side; without spinules; absence of Schmeil’s organ. Exopod 2 with seta; unrestricted; one on inner surface; with spine; 1; stout; not surpassing the exopod 3; serrated; on inner side, or on outer side; with setules; as a row; single; continuously; on inner surface; without spinules; absence of Schmeil’s organ. Exopod 3 with seta; plurimarginal; three on inner surface; two on terminal surface; with spine; 2; unequal size; first no longer 2x than origin segment; stout; serrated; on inner side, or on outer side; equally; second longer 2x than origin segment; slender; serrated; on outer side; with ornamentation on non-serrated side; of setules; setules on outer surface; as a row; single; continuously; on inner surface; with spinules; as a row; single; distally inserted; at anterior surface; absence of Schmeil’s organ. **Third swimming legs**. Symmetrical; Third swimming legs biramous. Third swimming legs intercoxal plate without seta. Third swimming legs praecoxa present; not laterally located. Third swimming legs coxa with seta; straight; distally inserted; on inner surface; surpassing to first endopodal segment; without setules; without spinules; without spine. Third swimming legs basis without seta; without setules; without spinules; without spine. Third swimming legs endopod 3-segmented. Endopod 1 with seta; restricted; one on inner surface; without spine; without setules; without spinules; absence of Schmeil’s organ. Endopod 2 with seta; restricted; two on inner surface; straight; without spine; without setules; without spinules; absence of Schmeil’s organ. Endopod 3 with seta; straight; plurimarginal; two on inner surface; two on outer surface; three on terminal surface; without spine; without setules; with spinules; as a row; distally inserted; double; at anterior surface; absence of Schmeil’s organ. Third swimming legs exopod 1 with seta; restricted; straight; one on inner surface; with spine; 1; stout; not reaching to the distal-third of the exopod 2; serrated; equally; on inner surface, or on outer surface; with setules; as a row; single; continuously; on inner surface; without spinules; absence of Schmeil’s organ. Exopod 2 with seta; straight; restricted; one on inner surface; with spine; 1; stout; not reaching out to exopod 3; serrated; on inner side, or on outer side; equally; with setules; as a row; single; continuously; on inner side; without spinules; absence of Schmeil’s organ. Exopod 3 without setules; with spinules; as a row; single; distally inserted; at anterior surface; with seta; straight; unrestricted; three on inner surface; two on terminal surface; with spine; 2; unequal size; first no longer 2x than origin segment; stout; serrated; on inner side, or on outer side; equally; second longer 2x than origin segment; slender; serrated; on outer side; with ornamentation on non-serrated side; of setules; absence of Schmeil’s organ. **Fourth swimming legs**. Symmetrical; biramous. Intercoxal plate without sensilla. Praecoxa present. Coxa with seta; distally inserted; on inner margin; reaching out to endopod 1; without spinules; setules absent. Basis with seta; one; medially inserted; on posterior surface; smaller than the original segment; without setules; without spinules; without spine. Fourth swimming legs endopod 3-segmented. Endopod 1 with seta; one; restricted; on inner surface; without spine; without setules; without spinules; absence of Schmeil’s organ. Endopod 2 with seta; restricted; two on inner side; without spine; with setules; as a row; single; continuously; on outer surface; without spinules; absence of Schmeil’s organ. Endopod 3 with seta; unrestricted; two on inner surface; two on outer surface; three on distal surface; without spine; without setules; with spinules; as a row; double; distally inserted; at anterior surface; absence of Schmeil’s organ. Fourth swimming legs exopod 1 with seta; restricted; one on inner surface; with spine; 1; stout; not reaching out to distal-third of the exopod 2; serrated; on inner side, or on outer side; equally; with setules; as a row; single; continuously; on inner surface; without spinules; absence of Schmeil’s organ. Exopod 2 with seta; restricted; one on inner surface; with spine; 1; stout; not reaching the end of exopod 3; serrated; on inner side, or on outer side; equally; with setules; as a row; single; continuously; on inner surface; without spinules; absence of Schmeil’s organ. Exopod 3 without setules; with spinules; as a row; single; distally inserted; at anterior surface; with seta; unrestricted; three on inner surface; two on distal surface; with spine; 2; unequal size; first no longer 2x than origin segment; stout; serrated; on inner side, or on outer side; equally; second longer 2x than origin segment; slender; serrated; on outer side; without ornamentation on non-serrated side; absence of Schmeil’s organ.

##### Fifth swimming legs features

Asymmetrical. Fifth swimming leg intercoxal plate with length equal or greater than width on 1.5x; with regular proximal margin; discontinuous to; the anterior margin of the left coxa; posterior sensilla on the right lateral absent. **Fifth left swimming leg**. Fifth left swimming leg biramous; leg reaching first right exopod segment; medially. Fifth left swimming leg praecoxa present; rudimentary; separated from the coxae; without ornamentation. Fifth left swimming leg coxa convex inner side; without teeth-like structures; with process; triangulated; on posterior surface; outer side; distally inserted; not projecting over basis; with sensilla; slender; triangular; at apex; longer 2x than insertion basis; without swelling; without seta; without spinules. Fifth left swimming leg basis sub-cylindrical; unequal size between inner and outer side; shorter outer than inner side; with rectilinear inner side; rounded internal proximal expansion absent; without outgrowth; with groove; deep; obliquely; on posterior surface; not reaching the endopodal lobe; not ornamented; absence of protuberance; with seta; outerly inserted; no longer 2x than origin segment; absence of minutely granular. Fifth left swimming leg endopod segments 1 and 2 fused; segments 2 and 3 fused; 2-segmented; stout; separated from the basis; ornamented; ornamented on segment 2; on inner side; with spinules; more than four elements; as a row; terminally; row of setules absent; without seta. Fifth left swimming leg exopod segments 1 and 2 separated; segments 2 and 3 fused; 2-segmented; stout; separated from the basis. Fifth left swimming leg exopod 1 sub-triangular; longer than broad; unequal size between inner and outer side; shorter inner than outer side; concave inner side; convex outer side; without swelling; without marginal extension; without process; with lobe; single; semicircular; medially inserted; on inner side; covered; by setules; without outer spine; absence seta. Fifth left swimming leg exopod 2 digitiform; longer than broad; equal size between inner and outer side; disform inner side; with rectilinear outer side; setulose pad present; prominently rounded; proximally; on inner side; inflated medial region present; setulose; anteriorly; distal process present; digitiform; denticulate; not bicuspidate; with transverse row of denticles; none oblique row of 5 denticles; at anterior surface; not innerly directed; with seta; spiniform; ornamented by spinules; surpassing the distal-point of the segment; without outer spine; terminal claw absent.

##### Fifth right swimming leg

Biramous. Fifth right swimming leg praecoxa present; separated from the coxae; without ornamentation. Fifth right swimming leg coxa convex inner side; without teeth-like structures; with process; conical; distally inserted; on posterior surface; closest to the outer rim; not projecting over basis; without triangular protuberance innerly; with sensilla; slender; at apex; no longer 2x than basal insertion; with marginal extension; without seta; without spinules. Fifth right swimming leg basis cylindrical; unequal size between inner and outer side; shorter outer than inner side; rectilinear inner side; tumescence absent; without protuberance; absence of distinct minutely granular; additional inner process absent; with posterior groove; shallow; longitudinally; reaching the endopodal lobe; not ornamented; with seta; outerly inserted; on posterior surface; no longer 2x than origin segment; posterior protrusion present; distal process absent. Fifth right swimming leg with endopodite present; separated from the basis; on anterior surface; ancestral segments 1 and 2 fused; ancestral segments 2 and 3 fused; 1-segmented; stout; ornamented; with setules; as a row; on inner side; terminally; without seta. Fifth right swimming leg exopod segments 1 and 2 separated; segments 2 and 3 fused; 2-segmented; stout; separated from the basis. Fifth right swimming leg exopod 1 trapezium; longer than broad; nearly 1.25 times; unequal size between both sides; shorter inner than outer side; rectilinear inner side; convex outer side; with marginal extension; sub-triangular; distally inserted; at outer rim; spinules absent; with process; rounded; sclerotized; without ornamentation; distally inserted; at marginal surface; projecting over next segment; without outer spine; without seta; internal prominence absent; lamella on posterior surface absent. Fifth right swimming leg exopod 2 cylindrical; longer than broad; nearly 2.5 times; equal size between both sides; uniform inner side; convex outer side; without posterior proximal swelling; inner-posterior process absent; without marginal expansion; curved ridge on distal posterior surface absent; chitinous knobs absent; with outer spine; inserted sub-distally; arched; internally directed; not ornamented innerly; not ornamented outerly; sharp tip; with apparent curve; innerly directed; lesser than the length of the exopod 2; beyond to 2 times its size; 6x; sensilla absent; terminal claw present; equal or longer 1.5 times than insertion segment; sclerotized; arched; inward; without conspicuous curve; ornamented innerly; by spinules; as a row; partially on extension; medially; not ornamented outerly; sharp tip; curved tip; outwards; without medial constriction; hyaline process absent.

##### FEMALE

Body longer and wider than male; Female body 1655 micrometers excluding caudal setae. Widest at first metasome segment. Distal margin of the prosomal segments without one line of setules at posterior margin. Prosome segments with spinules at least at one prosomal segment. Fourth metasome segment absence of dorsal protuberance. Fourth and fifth metasome segments fused; partially; on dorsal surface. Limit between fourth and fifth metasome segments without ornamentation. **Fifth metasome segment**. Fifth metasome segment without sensilla; with epimeral plates. Epimeral plates asymmetrical. Right epimeral plates prominent, as projections; thinner than the left; one posterior-laterally directed; reaching half length of the genital segment; with sensilla at the apex; dorsal-posterior sensilla present; stout; without ornamentation. Left epimeral plate without expansion.

##### Urosome

3-segmented. **Genital double-somite**. Asymmetrical in dorsal view; longer than broad; longer than other urosomites combined; dorsal suture at mid-length absent; not covered by spinules; with swelling; conical; equal size; anteriorly; with sensillae; on both sides; one; stout; with robust apex; at left rim; on lobular base; anteriorly; one; stout; at right lateral; on lobular base; anteriorly; with robust apex; of unequal size between then; greater left than right; lateral protuberance present; rounded; on the left side; without right posterior rim expanded; with slender sensilla on each posterior rim; without posterior-dorsal process. Genital double-somite opercular pad present; broader than longer; symmetrical; development laterally; expanded posteriorly; covering partially; double gonoporal slit; located ventrally; with arthrodial membrane; inserted anteriorly; post-genital process absent; disto-ventral tumescence absent; ventral vertical folds absent; dorsal sensilla absent. Second urosome segment without ventral fusion to anal segment; right distal process absent. Caudal rami patch of setules on outer surface absent; patch of spinules on outer surface absent.

##### Oral appendices feature

Rostrum basal process absent. **Antennules**. Symmetrical. Right antennule surpassing to genital double-segment; extending beyond caudal rami. Right antennule not exceeding the caudal setae. Right antennule ornamentation pattern equals to male left antennule; fully.

##### Fifth swimming legs

Symmetrical; Fifth swimming legs biramous. Fifth swimming legs intercoxal plate longer than wide; separated from the legs. Fifth swimming legs praecoxa without sclerite praecoxal. Fifth swimming legs coxa with process; conical; at the outer rim; distally; sensilla present; stout; at apex; projecting over basal segment; no longer 2x than basal insertion; marginal extension absent; without swelling; without seta; without spinules. Fifth swimming legs basis sub-triangular; equal size between inner and outer sides; with concave inner side; without proximal inner outgrowth; without groove; with distal extension; on posterior surface; with seta; outerly inserted; on anterior surface; no longer 2x than origin segment. Fifth swimming legs endopod segments 1 and 2 separated; segments 2 and 3 fused; 2-segmented; with complete suture; stout; separated from the basis; ornamentation on segment 2; with spinules; as a row; single; non-oblique; terminally; at anterior surface; with seta; double; one medially; on posterior surface; rectilinear; one distally; on posterior surface; rectilinear; of unequal size; distal seta longer than medial seta. Fifth swimming legs exopod segments 1 and 2 separated; segments 2 and 3 separated; 3-segmented; separated from the basis. Fifth swimming legs exopod 1 sub-cylindrical; longer than wide; longer or equal than 2 times; with unequal size between inner and outer side; shorter inner than outer side; with convex inner side; with convex outer side; without swelling; without marginal extension; without posterior process; without spine; without seta. Fifth swimming legs exopod 2 sub-cylindrical; longer than broad; longer or equal than 2 times; without swelling; without marginal extension; without process; without lobe; with spine; inserted laterally; rectilinear; without ornamentation; sharp tip; equal size or larger than next segment; without seta. Fifth swimming legs exopod 3 cylindrical; longer than wide; without swelling; without process; without lobe; without spine; with seta; double; inserted terminally; unequal size between them; outer seta smaller than inner; nearly 1 time; outer seta not ornamented by setules; without ornamentation; presence of terminal claw; sclerotized; rectilinear; with ornamentation; of denticles; as a row; on surface partially; at medial region; rectilinear outer side; with ornamentation; of denticles; as a row; on surface partially; at medial region; blunt tip; 6 times longer than origin segment.

##### Distribution records

###### BRAZIL

**Amazonas**: Calado Lake (Ciccino, Santos-Silva and Robertson, 2001); VENEZUELA. **Amazonas**: San Fernando de Atabapo, and Guaviare River Basins.

##### Habitat

Habitat in freshwaters: lake, and rivers.

##### Remarks

The hypothesis of Cicchino *et al*. (2001) was presented for science from lacustrine organisms of the Brazilian Amazonia, and rivers of Venezuela and Colombia. Originally the female of *N. simillimus* was presented as conspecific in the species *Rhacodiaptomus calatus* Brandorff 1973 mistakenly, and correctly related to the male in plankton samples from Atabapo (Venezuela), and Guaviare Rivers (Colombia) for its redescription as *Notodiaptomus* (Cicchino *et al*., 2001). At its foundation it was highlighted as a diagnosis of the species a female fourth metasome segment without dorsal protuberance, and genital double-somite with left rounded protuberance below to conical swelling with sensilla on the apex. Additionally male has a fifth right swimming leg exopod 1 longer than broad, and fifth left swimming leg exopod 2 with spiniform seta surpassing to distal-point of original segment. These characteristics also were indicated as differential attributes between the species and *N. coniferoides*, the most related taxon closely.

In the present effort we have added new differential identifications between species. The male of *N. simillimus* has fifth left swimming leg endopod 2-segmented, while the condition identified here for the same structure of the male of N. coniferoides is endopod 1-segmented. For females the epimeral plates are prominences but for *N. simillimus* a left epimeral plate is projected dorsal-posteriorly and reaches half of the female genital segment. For the female of *N. coniferoides* the same structure is projected dorsal-laterally and does not reach the medial region of the genital double segment. Another interesting observation is the original record of difference between species through the female of *N. simillimus* with fifth swimming legs exopod 2 with lateral spine to the detriment of the absence of the element in the homologous structure in *N. coniferoides.* Throughout our examinations all individuals present for both taxa possessed the ornamental element, being in *N. coniferoides* smaller considerably.

**(1)** *N. simillimus* is one of the taxa that has the most divergent organisms of the morphology considered for the creation of *Notodiaptomus*. Among the attributes highlighted for the *nordestinus* complex (Wright, 1935; 1936; 1937) and *Notodiaptomus* (Kiefer, 1936; 1937), the species presents the following differences: (1) male fifth left swimming leg basis with proximal tumescence innerly; (2) male fifth right swimming leg exopod 2 without inner curved ridge posteriorly; (3) male fifth left swimming leg endopod 2-segmented; (4) female fifth swimming leg endopod 2-segmented; (5) female fifth swimming leg endopod with spinules row non-oblique; (6) male fifth left swimming leg exopod 2 with spiniform seta surpassing to distal-point original segment. Divergent from the type-species, we highlight: (1) female antennule not extending beyond caudal rami; (2) male right antennule actual segment 13 with spiniform modified seta reaching to the distal-point of the sequence segment; (3) male fifth left swimming leg reaching to first right exopod segment; and (4) female fifth swimming legs basis with outer seta not reaching to distal exopod 1.

#### Notodiaptomus spinuliferus Dussart & Matsumura-Tundisi in Dussart, 1985

##### Synonymy

*Notodiaptomus spinuliferus*; Dussart, 1985a: 208, fig. 6; Dussart & Frutos, 1986: 307, 308; Dussart & Matsumura-Tundisi, 1986: 250, fig. 1; Matsumura-Tundisi, 1986: 537, figs. 34–37, 100; Reid, 1987: 377; José de Paggi & Paggi, 1988: 101, tab. 2; Sendacz, 1993: 35; 1997: 624, 625, tab. 2; Frutos, 1993: 112, tab. 3; Battistoni, 1995: 959; Rocha *et al*., 1995: 156; Lansac-Tôha *et al*., 1997: 140, tab. 3; Santos-Silva, 1998: 212; Santos-Silva, 2008: 33–34, fig. 7; Geraldes-Primeiro *et al*., 2021: 3, fig. 1. *Notodiaptomus* cf. *spinuliferus*; Reid & Moreno, 1990: 726, 729, 730, tabs. 2, 3; Perbiche-Neves *et al*., 2015: 102 and 105, fig. 93; Perbiche-Neves *et al*., 2020: 684-685, key to the Neotropical diaptomid, fig. 21.9 K. *Notodiaptomus (Notodiaptomus) spinuliferus*; Dussart, 1985a: 208.

##### Type locality

Ilha Solteira Reservoir, Sao Paulo State, Brazil.

##### Type material

Holotype: 1 male, dissected on the slide (MZUSP 6969). Paratype: 1 male, dissected on the slide (MZUSP 6970); 5 males, and 5 females, entire in formalin 4% (MZUSP 6971). Material deposited in the Zoology Museum of São Paulo - MZUSP. Additional material: females and males deposited in the laboratory of Limnology at the Federal University of São Carlos.

##### Material examined

Holotype: 1 male, entire in alcohol (MZUSP 6969). Paratype: 1 male, entire in alcohol (MZUSP 6970). 4 males, and 5 females in humid medium (formalin solution) (MZUSP 6971), 1 male (INPA-COP055, slides a-h) and 1 female (INPA-COP056, slides a-h) were selected to be dissection on eight slides each and deposited in the Zoological Collection of the INPA, Brazil.

##### Diagnosis

**(1)** male right antennule with length not extending beyond caudal rami; **(2)** left antennule actual segment 11 with two setae of unequal size, none beyond three sequential segments; **(3)** antenna endopod actual segment 2 bilobate without discontinuity on outer cuticle; **(4)** male fifth left swimming leg with length reaching to first right exopod segment distally; **(5)** male fifth right swimming leg coxa with posterior rounded process distally, and projecting over basis until to medial surface; **(6)** male fifth right swimming leg basis with posterior shallow groove obliquely, not ornamented; **(7)** male fifth right swimming leg basis without posterior protrusion; **(8)** male fifth right swimming leg exopod 2 with rectilinear outer spine, 2x lesser than original segment; **(9)** female limit between fourth and fifth metasome segments separated and with single spinules row laterally; **(10)** female right epimeral plate without dorsal-posterior sensilla, ornamented with dorsal spinules patch innerly; **(11)** female antennule surpassing to caudal setae; **(12)** female fifth swimming legs exopod 3 with outer seta smaller than inner 1x nearly; **(13)** female fifth swimming legs endopod 2-segmented with incomplete suture.

##### Redescription

###### MALE

Body 1079.8 micrometers excluding caudal setae. Male body smaller and slenderer than female. Nerve axons myelinated. Prosome 6-segmented; widest at first metasome segment; without one line of setules at posterior margin; with spinules at segments. Cephalosome anterior margin rounded; with dorsal suture; incomplete; separate from first metasome segment. First metasome segment without sensilla. Second metasome segment without sensilla. Third metasome segment without sensillae; ornamented posterior margin; with spinules; as a patch; laterally. Fourth metasome segment with sensillae; 4 dorsally; 2 laterally; of unequal size; separated from the fifth metasome. Limit between fourth and fifth metasome segments ornamented; with spinules; as a row; on dorsal doubly; on lateral singly; same size. Fifth metasome segment with sensilla; 2 dorsally; Fifth metasome segment equal size; Fifth metasome segment without ornamentation; Fifth metasome segment without dorsal conical process; with epimeral plates. Epimeral plates symmetrical. Right epimeral plates prominent, as projections; one projection; posterior-laterally directed; not reaching half length of the genital segment; with sensilla; at the apex of projection; without ornamentation.

##### Urosome

5-segmented; Urosome 5 - free segments. Genital somite asymmetrical in dorsal view; with single aperture; located on left side; ventrolaterally on posterior rim; with sensillae; on both sides; one; at right lateral; posteriorly; of equal size between then. Third urosome segment without spinules; without external seta. Fourth urosome segment without spinules; without sub-conical blunt dorsal-lateral process. Anal segment presence of dorsal sensillae; one on each side; medially inserted; presence of operculum; convex; covering the anal aperture fully. Caudal rami symmetrical; separated from anal segment; longer than wide; with setules; continuous on; inner side; each ramus bearing 6 caudal setae; 5 marginals; plumose; and 1 internal dorsally; straight; not reticulated main axis; outermost seta with outer spiniform process absent.

##### Oral appendices feature

Rostrum symmetrical; separated from dorsal cephalic shield; by complete suture; sensillae present; one pair; medially inserted on tegument; with rostral filament; double; paired; extended; into point; basal process absent; without a smaller basal expansion on the right side.

##### Antennules

Asymmetrical. **Right antennules**. Uniramous; right antennule surpassing to genital segment; right antennule not extending beyond caudal rami.

Right antennule ancestral segment I and II separated. Ancestral segment II and III fused. Ancestral segment III and IV fused. Ancestral segment IV and V separated. Ancestral segment V and VI separated. Ancestral segment VI and VII separated. Ancestral segment VII and VIII separated. Ancestral segment VIII and IX separated. Ancestral segment IX and X separated. Ancestral segment X and XI separated. Ancestral segment XI and XII separated. Ancestral segment XII and XIII separated. Ancestral segment XIII and XIV separated. Ancestral segment XIV and XV separated. Ancestral segment XV and XVI separated. Ancestral segment XVI and XVII separated. Ancestral segment XVII and XVIII separated. Ancestral segment XVIII and XIX separated. Ancestral segment XIX and XX separated. Ancestral segment XX and XXI separated. Ancestral segment XXI and XXII fused. Ancestral segment XXII and XXIII fused. Ancestral segment XXIII and XXIV separated. Ancestral segment XXIV and XXV fused. Ancestral segment XXV and XXVI separated. Ancestral segment XXVI and XXVII separated. Ancestral segment XXVII and XXVIII fused.

Right antennule actual 22-segmented; geniculated; between the segment 18 and segment 19; with swollen and modified region; formed by 5 segments; between 13 and 17 segments. Actual segment 1 with seta; one element; straight; none larger than segment; without spinules; without vestigial seta; without conical seta; without modified seta; without spinous process; with aesthetasc; one element. Actual segment 2 with seta; three elements; of unequal size; straight; none larger than segment; without spinules; with vestigial seta; one element; without conical seta; without modified seta; without spinous process; with aesthetasc; one element. Actual segment 3 with seta; one element; one larger than segment; surpassing to distal margin; beyond three sequential segments; straight; blunt apex; without spinules; without vestigial seta; without conical seta; without modified seta; without spinous process; with aesthetasc; one element. Actual segment 4 with seta; one element; one larger than segment; surpassing to distal margin; straight; not beyond three sequential segments; without spinules; without vestigial seta; without conical seta; without modified seta; without spinous process; without aesthetasc. Actual segment 5 with seta; one element; straight; one larger than segment; surpassing to distal margin; not beyond three sequential segments; without spinules; with vestigial seta; one element; without conical seta; without modified seta; without spinous process; with aesthetasc; one element. Actual segment 6 with seta; one element; one larger than segment; surpassing to distal margin; not beyond three sequential segments; straight; without spinules; without vestigial seta; without conical seta; without modified seta; without spinous process; without aesthetasc. Actual segment 7 with seta; one element; straight; one larger than segment; surpassing to distal margin; beyond three sequential segments; blunt apex; without spinules; without vestigial seta; without conical seta; without modified seta; without spinous process; with aesthetasc; one element. Actual segment 8 with seta; one element; straight; none larger than segment; without spinules; without vestigial seta; with conical seta; one element; not reaching to middle-point of the sequent segment; without modified seta; without spinous process; without aesthetasc. Actual segment 9 with seta; two elements; of unequal size; straight; one larger than segment; surpassing to distal margin; beyond three sequential segments; blunt apex; without spinules; without vestigial seta; without conical seta; without modified seta; without spinous process; with aesthetasc; one element. Actual segment 10 with seta; one element; straight; none larger than segment; without spinules; without vestigial seta; without conical seta; with modified seta; presenting blunt apex; slender form; surpassing to distal margin; beyond of the sequential segment; parallel to antennule direction; without spinous process; without aesthetasc. Actual segment 11 with seta; one element; straight; one larger than segment; surpassing to distal margin; not beyond three sequential segments; without spinules; without vestigial seta; without conical seta; with modified seta; slender form; presenting blunt apex; surpassing to distal margin; beyond of the sequential segment; parallel to antennule direction; shorter length than homologous of actual segment 13; without spinous process; without aesthetasc. Actual segment 12 with seta; one element; straight; one larger than segment; surpassing to distal margin; not beyond three sequential segments; without spinules; without vestigial seta; with conical seta; one element; smaller than to segment 8; without modified seta; without spinous process; with aesthetasc; one element; absent internal perpendicular fission. Actual segment 13 with seta; one element; straight; one larger than segment; surpassing to distal margin; not beyond three sequential segments; without spinules; without vestigial seta; without conical seta; with modified seta; stout form; surpassing to distal margin; to the middle-point of the sequence segment; perpendicular to antennule direction; presenting bifid apex; without spinous process; without aesthetasc. Actual segment 14 with seta; one element; straight; one larger than segment; surpassing to distal margin; beyond three sequential segments; blunt apex; without spinules; without vestigial seta; without conical seta; without modified seta; without spinous process; with aesthetasc; one element. Actual segment 15 with seta; two elements; of unequal size; straight; not bifidform; none larger than segment; without spinules; without vestigial seta; without conical seta; without modified seta; with spinous process; on outer margin; surpassing distal margin; with aesthetasc; one element. Actual segment 16 with seta; two elements; of unequal size; straight; one larger than segment; surpassing to distal margin; not beyond three sequential segments; not bifidform; without spinules; without vestigial seta; without conical seta; without modified seta; with spinous process; on outer margin; surpassing distal margin; unequal size to process on preceding segment; with aesthetasc; one element. Actual segment 17 with seta; two elements; of unequal size; straight; none larger than segment; bifidform; without spinules; without vestigial seta; without conical seta; without modified seta; without spinous process; without aesthetasc. Actual segment 18 with seta; two elements; of equal size; straight; none larger than segment; without spinules; without vestigial seta; without conical seta; without modified seta; without spinous process; without aesthetasc. Actual segment 19 with seta; two elements; of unequal size; plumose; none larger than segment; without spinules; without vestigial seta; without conical seta; without modified seta; at least one bifid form; without spinous process; with aesthetasc; one element. Actual segment 20 with seta; four elements; of unequal size; straight; one larger than segment; surpassing to distal margin; beyond three sequential segments; without spinules; without vestigial seta; without conical seta; without modified seta; with spinous process; distally; reaching beyond of distal-point segment 21; without aesthetasc. Actual segment 21 with seta; two elements; of equal size; plumose; one larger than segment; surpassing to distal margin; greater 3x than original segment; without spinules; without vestigial seta; without conical seta; without modified seta; without spinous process; without aesthetasc. Actual segment 22 with seta; four elements; of equal size; one larger than segment; plumose; surpassing to distal margin; greater 3x than original segment; without spinules; without vestigial seta; without conical seta; without modified seta; without spinous process; with aesthetasc; one element.

##### Left antennules

Uniramous; Left antennule surpassing to prosome; Left antennule not extending beyond caudal rami. Ancestral segment I and II separated. Ancestral segment II and III fused. Ancestral segment III and IV fused. Ancestral segment IV and V separated. Ancestral segment V and VI separated. Ancestral segment VI and VII separated. Ancestral segment VII and VIII separated. Ancestral segment VIII and IX separated. Ancestral segment IX and X separated. Ancestral segment X and XI separated. Ancestral segment XI and XII separated. Ancestral segment XII and XIII separated. Ancestral segment XIII and XIV separated. Ancestral segment XIV and XV separated. Ancestral segment XV and XVI separated. Ancestral segment XVI and XVII separated. Ancestral segment XVII and XVIII separated. Ancestral segment XVIII and XIX separated. Ancestral segment XIX and XX separated. Ancestral segment XX and XXI separated. Ancestral segment XXI and XXII separated. Ancestral segment XXII and XXIII separated. Ancestral segment XXIII and XXIV separated. Ancestral segment XXIV and XXV separated. Ancestral segment XXV and XXVI separated. Ancestral segment XXVI and XXVII separated. Ancestral segment XXVII and XXVIII fused.

Left antennule actual 25-segmented; not-geniculated. Actual segment 1 with seta; one element; none larger than segment; straight; without spinules; without vestigial seta; without conical seta; without modified seta; without spinous process; with aesthetasc; one element. Actual segment 2 with seta; three elements; of equal size; none larger than segment; straight; without spinules; without vestigial seta; without conical seta; without modified seta; without spinous process; with aesthetasc; one element. Actual segment 3 with seta; one element; one larger than segment; straight; surpassing to distal margin; beyond three sequential segments; without spinules; without vestigial seta; without conical seta; without modified seta; without spinous process; with aesthetasc; one element. Actual segment 4 with seta; one element; none larger than segment; straight; without spinules; without vestigial seta; without conical seta; without modified seta; without spinous process; without aesthetasc. Actual segment 5 with seta; one element; one larger than segment; straight; surpassing to distal margin; not beyond three sequential segments; without spinules; without vestigial seta; without conical seta; without modified seta; without spinous process; with aesthetasc; one element. Actual segment 6 with seta; one element; none larger than segment; straight; without spinules; without vestigial seta; without conical seta; without modified seta; without spinous process; without aesthetasc. Actual segment 7 with seta; one element; one larger than segment; straight; surpassing to distal margin; beyond three sequential segments; without spinules; without vestigial seta; without conical seta; without modified seta; without spinous process; with aesthetasc; one element. Actual segment 8 with seta; one element; one larger than segment; straight; surpassing distal margin; without spinules; without vestigial seta; with conical seta; without modified seta; without spinous process; without aesthetasc. Actual segment 9 with seta; two elements; of unequal size; one larger than segment; straight; surpassing to distal margin; beyond three sequential segments; without spinules; without vestigial seta; without conical seta; without modified seta; without spinous process; with aesthetasc; one element. Actual segment 10 with seta; one element; none larger than segment; straight; without spinules; without vestigial seta; without conical seta; without modified seta; without spinous process; without aesthetasc. Actual segment 11 with seta; two elements; of unequal size; one larger than segment; straight; surpassing to distal margin; not beyond three sequential segments; without spinules; without vestigial seta; without conical seta; without modified seta; without spinous process; without aesthetasc. Actual segment 12 with seta; one element; one larger than segment; straight; surpassing distal margin; without spinules; without vestigial seta; with conical seta; without modified seta; without spinous process; with aesthetasc; one element. Actual segment 13 with seta; one element; none elongated; straight; surpassing distal margin; without spinules; without vestigial seta; without conical seta; without modified seta; without spinous process; without aesthetasc. Actual segment 14 with seta; one element; elongated; straight; surpassing to distal margin; beyond three sequential segments; without spinules; without vestigial seta; without conical seta; without modified seta; without spinous process; with aesthetasc; one element. Actual segment 15 with seta; one element; larger than segment; straight; surpassing to distal margin; not beyond three sequential segments; without spinules; without vestigial seta; without conical seta; without modified seta; without spinous process; without aesthetasc. Actual segment 16 with seta; one element; larger than segment; plumose; surpassing to distal margin; not beyond three sequential segments; without spinules; without vestigial seta; without conical seta; without modified seta; without spinous process; with aesthetasc; one element. Actual segment 17 with seta; one element; not larger than segment; straight; without spinules; without vestigial seta; without conical seta; without modified seta; without spinous process; without aesthetasc. Actual segment 18 with seta; one element; larger than segment; straight; surpassing to distal margin; beyond three sequential segments; without spinules; without vestigial seta; without conical seta; without modified seta; without spinous process; without aesthetasc. Actual segment 19 with seta; one element; not larger than segment; straight; surpassing distal margin; without spinules; without vestigial seta; without conical seta; without modified seta; without spinous process; with aesthetasc; one element. Actual segment 20 with seta; one element; not larger than segment; straight; surpassing distal margin; without spinules; without vestigial seta; without conical seta; without modified seta; without spinous process; without aesthetasc. Actual segment 21 with seta; one element; larger than segment; plumose; surpassing to distal margin; beyond three sequential segments; without spinules; without vestigial seta; without conical seta; without modified seta; without spinous process; without aesthetasc. Actual segment 22 with seta; two elements; of unequal size; one of them elongated; plumose; surpassing to distal margin; without spinules; without vestigial seta; without conical seta; without modified seta; without spinous process; without aesthetasc. Actual segment 23 with seta; two elements; of unequal size; one larger than segment; plumose; surpassing to distal margin; greater 3x than original segment; without spinules; without vestigial seta; without conical seta; without modified seta; without spinous process; without aesthetasc. Actual segment 24 with seta; two elements; of equal size; one larger than segment; plumose; surpassing to distal margin; greater 3x than original segment; without spinules; without vestigial seta; without conical seta; without modified seta; without spinous process; without aesthetasc. Actual segment 25 with seta; four elements; of equal size; elongated; plumose; surpassing to distal margin; 4 times larger than segment; without spinules; without vestigial seta; without conical seta; without modified seta; without spinous process; with aesthetasc; one element.

##### Antenna

Biramous. Antenna coxa separated from the basis; bearing seta; 1; on inner surface; at distal corner; reaching to the middle basis. Antenna basis (fusion) separated from the endopodal segment; bearing seta; 2; on inner surface; at distal corner. Endopodal ancestral segment I and II separated. Ancestral segment II and III fused. Ancestral segment III and IV fused. Ancestral segment III and IV fully. Antenna endopod actual 2-segmented. Actual segment 1 not bilobate; with seta; two; on inner margin; with spinules; as a patch; not obliquely; on outer surface; with pore. Actual segment 2 bilobate; without discontinuity on outer cuticle; inner lobe bearing 8 setae; distally; outer lobe bearing 7 setae; distally; with spinules; as a patch; on outer surface. Antenna exopod ancestral segment I and II separated. Ancestral segment II and III fused. Ancestral segment III and IV fused. Ancestral segment IV and V separated. Ancestral segment V and VI separated. Ancestral segment VI and VII separated. Ancestral segment VII and VIII separated. Ancestral segment VIII and IX separated. Ancestral segment IX and X fused. Antenna exopod actual 7-segmented. Actual segment 1 single; elongated (width-length, equal or larger ratio 2:1); with seta; one; at inner surface. Actual segment 2 compound; elongated (larger width-length ratio 2:1); with seta; three; at inner surface. Actual segment 3 single; not elongated (lesser width-length ratio 2:1); with seta; one; at inner surface. Actual segment 4 single; not elongated (lesser width-length ratio 2:1); with seta; one; at inner surface. Actual segment 5 single; not elongated (lesser width-length ratio 2:1); with seta; one; at inner surface. Actual segment 6 single; not elongated (lesser width-length ratio 2:1); with seta; one; at inner surface. Actual segment 7 compound; elongated (larger or equal width-length ratio 2:1); with seta; one; at inner surface; and three; at distal surface.

##### Oral features

**Mandible**. Coxal gnathobase sclerotized; with lobe; prominent; on sub-caudal margin; presence of cutting blade; with tooth-like prominence; two, distinctly; 1 acute; on caudal margin; and 1 triangular; on sub-caudal margin; with acute projection between the prominences; with additional spinules; as a patch; on dorsal surface; with seta; dorsally; on apical surface; without spinules. Mandible palps biramous; comprising the basis; with seta; four; differently inserted; first medially; not reaching to beyond the endopod 1; second distally; third distally; fourth distally; on inner margin; none with setulose ornamentation. Mandible endopod 2-segmented. Mandible endopod 1 with lobe; bearing seta; four; distally inserted; without spinules. Mandible endopod 2 without lobe; bearing setae; nine elements; distally inserted; with spinules; as a row; double. Mandible exopod 4-segmented. Mandible exopod 1 with seta; one element; distally; on inner margin. Mandible exopod 2 with seta; one element; distally; on inner side. Mandible exopod 3 with seta; one element; distally; on inner side. Mandible exopod 4 with setae; three elements; on terminal region. **Maxillule**. Birramous. Maxillule 3-segmented. Maxillule praecoxa with praecoxal arthrite; bearing spines; fifteen elements; ten marginally; plus, five sub-marginally; with spinules; as a patch; on sub-marginal surface. Maxillule coxa with coxal epipodite; with conspicuous outer lobe; bearing setae; nine elements; with coxal endite; elongated (larger or equal width-length ratio 2:1); bearing setae; four elements. Maxillule basis with basal endite; double; first proximal; elongated (larger width-length ratio 2:1; separated from basis; with setae; four elements; distally inserted; second distal; fused to basis; not elongated (lesser width-length ratio 2:1); with setae; four elements; distally inserted; with setules; as a row; on inner side; basal exite present; with setae; one element; on outer surface. Maxillule endopod 1-segmented. Endopod 1 bilobate; first proximal; with setae; three elements; second distal; with setae; five elements. Maxillule exopod 1-segmented. Exopod 1 with setae; six elements; with setules; as a row; on inner side; spinules absent. **Maxilla**. Uniramous. Maxilla 5-segmented. Maxilla praecoxa fused to coxa; incompletely; distinct externally; with praecoxal endite; double; first elongated endite (larger or equal width length ratio 2:1); proximally inserted; with seta; straight, or plumose; 3 straight; 3 plumose; with spine; single; with spinules; as a row; on distal margin; without setule; second non-elongated endite (lesser width length ratio 2:1); distally inserted; with seta; plumose; 3 plumose; without spine; with spinules; as a row; on distal margin; without setule; absence of outer seta. Maxilla coxa with coxal endite; double; first elongated endite (larger or equal width); proximally inserted; with seta; plumose; 3 plumose; without spine; without spinules; with setules; as a row; on distal margin; second elongated endite (larger or equal width); distally inserted; with seta; plumose; 3 plumose; without spine; without spinules; with setules; as a row; on distal margin; absence of outer seta. Maxilla basis with basal endite; single; elongated (larger or equal width-length ratio 2:1); with seta; plumose; 3 plumose; with spinules; absence of outer seta. Maxilla endopod 2-segmented. Endopod 1 with seta; 2 plumose; without spine; with spinules; as a patch; on distal margin; without setules. Maxilla endopod 2 with seta; 2 plumose; without spine; with spinules; as a patch; on distal margin; without setules. **Maxilliped**. Uniramous; Maxilliped 8-segmented. Maxilliped praecoxa fused to coxa; incompletely; distinct internally; with praecoxal endite; not elongated (lesser width-length ratio 2:1); distally inserted; with seta; 1 plumose; with spinules; as a row; single; on basal surface; without setules. Maxilliped coxa with coxal endite; three coxal endite; first not elongated (lesser width-length ratio 2:1); proximally inserted; with seta; 2 plumose; with spinules; as a patch; single; on medial surface; without setules; second elongated (larger or equal width); medially inserted; with seta; 3 plumose; with spinules; as a patch; single; on medial surface; without setules; third not elongated (lesser width-length ratio 2:1); distally inserted; with seta; 3 plumose; none reaching to beyond of the basis; with spinules; as a patch; single; on basal surface; with setules; as a patch; single; with lobe; prominence; at inner distal angle; ornamented; with spinules; continuously on margin. Maxilliped basis without basal endite; with seta; 3 straights; with spinules; as a row; single; on medial surface; with setules; as a row; single; on inner margin. Maxilliped endopod segment 6-segmented. Endopod 1 with seta; 2 plumose; on inner surface. Endopod 2 with seta; 3 plumose; on inner surface. Endopod 3 with seta; 2 plumose; on inner surface. Endopod 4 with seta; 2 plumose; on inner surface. Endopod 5 with seta; 2 plumose; on inner surface; outer seta absent. Endopod 6 with seta; 4 plumose; on inner surface, or on outer surface.

##### Swimming legs features

**First swimming legs.** Symmetrical; biramous. First swimming legs intercoxal plate without seta. First swimming legs praecoxa present; not laterally located. First swimming legs coxa with seta; one; straight; distally inserted; on inner surface; surpassing to first endopodal segment; with setules; one group; as a row; discontinuously; on inner margin; without spinules; without spine. First swimming legs basis without seta; with setules; as a row; single; continuously; on outer surface; without spinules; without spine. First swimming legs endopod 2-segmented. Endopod 1 without seta; without spine; without setules; without spinules; absence of Schmeil’s organ. Endopod 2 with seta; unrestricted; three on inner surface; one on outer surface; two on distal surface; straight; without spine; without setules; without spinules; absence of Schmeil’s organ. Endopod 3 absence. First swimming legs exopod 1 without seta; with spine; 1; stout; smaller than original segment; serrated; on inner side; continuously; with setules; as a row; single; as a row; innerly. First swimming legs exopod 2 with seta; restricted; 1 on inner surface; straight; without spine; without setules; without spinules. First swimming legs exopod 3 without setule; without spinules; with seta; unrestricted; 2 on inner surface; 2 on terminal surface; with spine; 2; unequal size; first no longer 2x than origin segment; stout; serrated; on inner side, or on outer side; equally; second longer 3x than origin segment; slender; serrated; on outer side; with ornamentation on non-serrated side; by setules. **Second swimming legs**. Symmetrical; Second swimming legs biramous. Second swimming legs intercoxal plate without seta. Second swimming legs praecoxa present; located laterally. Second swimming legs coxa with seta; straight; distally inserted; on inner surface; surpassing to basal segment; without setules; without spinules; without spine. Second swimming legs basis without seta; without setules; without spinules; without spine. Second swimming legs endopod 3-segmented. Endopod 1 with seta; straight; restricted; one on inner surface; without spine; without setules; without spinules; absence of Schmeil’s organ. Endopod 2 with seta; straight; restricted; one on inner surface; without spine; with setules; as a row; single; continuously; on inner side; without spinules; presence of Schmeil’s organ; on posterior surface. Endopod 3 with seta; straight; unrestricted; two on inner surface; two on outer surface; two on distal surface; without spine; without setules; with spinules; as a row; double; distally inserted; at anterior surface; absence of Schmeil’s organ. Second swimming legs exopod 1 with seta; restricted; one on inner surface; without spine; with setules; as a row; single; continuously; on inner side; without spinules; absence of Schmeil’s organ. Exopod 2 with seta; unrestricted; one on inner surface; with spine; 1; stout; not surpassing the exopod 3; serrated; on inner side, or on outer side; with setules; as a row; single; continuously; on inner surface; without spinules; absence of Schmeil’s organ. Exopod 3 with seta; plurimarginal; three on inner surface; two on terminal surface; with spine; 2; unequal size; first no longer 2x than origin segment; stout; serrated; on inner side, or on outer side; equally; second longer 2x than origin segment; slender; serrated; on outer side; with ornamentation on non-serrated side; of setules; no setules on outer surface; with spinules; as a row; double; distally inserted; at anterior surface; absence of Schmeil’s organ. **Third swimming legs**. Symmetrical; Third swimming legs biramous. Third swimming legs intercoxal plate without seta. Third swimming legs praecoxa present; not laterally located. Third swimming legs coxa with seta; straight; distally inserted; on inner surface; surpassing to basal segment; without setules; without spinules; without spine. Third swimming legs basis without seta; without setules; without spinules; without spine. Third swimming legs endopod 3-segmented. Endopod 1 with seta; restricted; one on inner surface; without spine; without setules; without spinules; absence of Schmeil’s organ. Endopod 2 with seta; restricted; two on inner surface; straight; without spine; without setules; without spinules; absence of Schmeil’s organ. Endopod 3 with seta; straight; plurimarginal; two on inner surface; two on outer surface; three on terminal surface; without spine; without setules; with spinules; as a row; distally inserted; single; at anterior surface; absence of Schmeil’s organ. Third swimming legs exopod 1 with seta; restricted; straight; one on inner surface; with spine; 1; stout; not reaching to the distal-third of the exopod 2; serrated; equally; on inner surface, or on outer surface; with setules; as a row; single; continuously; on inner surface; without spinules; absence of Schmeil’s organ. Exopod 2 with seta; straight; restricted; one on inner surface; with spine; 1; stout; not reaching out to exopod 3; serrated; on inner side, or on outer side; equally; without setules; without spinules; absence of Schmeil’s organ. Exopod 3 without setules; with spinules; as a row; single; distally inserted; at anterior surface; with seta; straight; unrestricted; three on inner surface; two on terminal surface; with spine; 2; unequal size; first no longer 2x than origin segment; stout; serrated; on inner side, or on outer side; equally; second longer 2x than origin segment; slender; serrated; on outer side; with ornamentation on non-serrated side; of setules; absence of Schmeil’s organ. **Fourth swimming legs**. Symmetrical; biramous. Intercoxal plate without sensilla. Praecoxa present. Coxa with seta; distally inserted; on inner margin; reaching out to endopod 1; without spinules; setules absent. Basis with seta; one; medially inserted; on posterior surface; smaller than the original segment; without setules; without spinules; without spine. Fourth swimming legs endopod 3-segmented. Endopod 1 without seta; without spine; without setules; without spinules; absence of Schmeil’s organ. Endopod 2 with seta; restricted; two on inner side; without spine; with setules; as a row; single; continuously; on outer surface; without spinules; absence of Schmeil’s organ. Endopod 3 with seta; unrestricted; two on inner surface; two on outer surface; three on distal surface; without spine; without setules; without spinules; absence of Schmeil’s organ. Fourth swimming legs exopod 1 with seta; restricted; one on inner surface; with spine; 1; stout; not reaching out to distal-third of the exopod 2; serrated; on inner side, or on outer side; equally; with setules; as a row; single; continuously; on inner surface; without spinules; absence of Schmeil’s organ. Exopod 2 with seta; restricted; one on inner surface; with spine; 1; stout; not reaching the end of exopod 3; serrated; on inner side, or on outer side; equally; with setules; as a row; single; continuously; on inner surface; without spinules; absence of Schmeil’s organ. Exopod 3 without setules; without spinules; with seta; unrestricted; three on inner surface; two on distal surface; with spine; 2; unequal size; first no longer 2x than origin segment; stout; serrated; on inner side, or on outer side; equally; second longer 2x than origin segment; slender; serrated; on outer side; with ornamentation on non-serrated side; of setules; absence of Schmeil’s organ.

##### Fifth swimming legs features

Asymmetrical. Fifth swimming leg intercoxal plate with length not equal or greater than width on 1.5x; with regular proximal margin; discontinuous to; the anterior margin of the left coxa, or the anterior margin of the right coxa; posterior sensilla on the right lateral absent. **Fifth left swimming leg**. Fifth left swimming leg biramous; leg reaching first right exopod segment; distally. Fifth left swimming leg praecoxa present; developed; separated from the coxae; without ornamentation. Fifth left swimming leg coxa concave inner side; without teeth-like structures; with process; conical; on posterior surface; outer side; distally inserted; not projecting over basis; with sensilla; slender; triangular; at apex; longer 2x than insertion basis; without swelling; without seta; without spinules. Fifth left swimming leg basis sub-cylindrical; unequal size between inner and outer side; shorter outer than inner side; with rectilinear inner side; rounded internal proximal expansion absent; without outgrowth; with groove; shallow; obliquely; on posterior surface; not reaching the endopodal lobe; not ornamented; absence of protuberance; with seta; outerly inserted; no longer 2x than origin segment; absence of minutely granular. Fifth left swimming leg endopod segments 1 and 2 fused; segments 2 and 3 fused; 1-segmented; stout; separated from the basis; ornamented; on inner side; with spinules; more than four elements; as a patch; terminally; row of setules absent; without seta. Fifth left swimming leg exopod segments 1 and 2 separated; segments 2 and 3 fused; 2-segmented; stout; separated from the basis. Fifth left swimming leg exopod 1 sub-cylindrical; longer than broad; unequal size between inner and outer side; shorter inner than outer side; concave inner side; concave outer side; without swelling; without marginal extension; without process; with lobe; double; circular; medially inserted; on inner side; covered; by setules; without outer spine; absence seta. Fifth left swimming leg exopod 2 digitiform; longer than broad; equal size between inner and outer side; disform inner side; with convex outer side; setulose pad present; not prominently rounded; medially; on inner side; inflated medial region absent; distal process present; digitiform; non denticulate; without transverse row of denticles; none oblique row of 5 denticles; not innerly directed; with seta; spiniform; not ornamented by spinules; not surpassing the distal-point of the segment; without outer spine; terminal claw absent.

##### Fifth right swimming leg

Biramous. Fifth right swimming leg praecoxa present; separated from the coxae; without ornamentation. Fifth right swimming leg coxa convex inner side; without teeth-like structures; with process; rounded; distally inserted; on posterior surface; closest to the outer rim; projecting over basis; beyond the first third; until the medial surface; without triangular protuberance innerly; with sensilla; slender; at apex; no longer 2x than basal insertion; without marginal extension; without seta; without spinules. Fifth right swimming leg basis cylindrical; unequal size between inner and outer side; shorter inner than outer side; rectilinear inner side; tumescence absent; without protuberance; absence of distinct minutely granular; additional inner process absent; with posterior groove; shallow; obliquely; not reaching the endopodal lobe; not ornamented; with seta; outerly inserted; on posterior surface; no longer 2x than origin segment; posterior protrusion absent; distal process absent. Fifth right swimming leg with endopodite present; separated from the basis; on anterior surface; ancestral segments 1 and 2 fused; ancestral segments 2 and 3 fused; 1-segmented; stout; ornamented; with spinules; as a row; on inner side; sub-terminally; without seta. Fifth right swimming leg exopod segments 1 and 2 separated; segments 2 and 3 fused; 2-segmented; stout; separated from the basis. Fifth right swimming leg exopod 1 sub-cylindrical; longer than broad; nearly 1.25 times; unequal size between both sides; shorter inner than outer side; rectilinear inner side; rectilinear outer side; with marginal extension; sub-triangular; distally inserted; at outer rim; spinules absent; with process; rounded; sclerotized; without ornamentation; distally inserted; at marginal surface; projecting over next segment; without outer spine; without seta; internal prominence absent; lamella on posterior surface absent. Fifth right swimming leg exopod 2 elliptical; longer than broad; nearly 2 times; equal size between both sides; disform inner side; convex outer side; with posterior proximal swelling; inner-posterior process absent; without marginal expansion; curved ridge on distal posterior surface absent; chitinous knobs present; with 1–2 posteriorly; with outer spine; inserted sub-distally; rectilinear; ornamented innerly; by spinules; as a row; not ornamented outerly; sharp tip; with apparent curve; innerly directed; lesser than the length of the exopod 2; until to 2 times its size; 2x; sensilla absent; terminal claw present; equal or longer 1.5 times than insertion segment; sclerotized; arched; inward; with conspicuous curve; proximally; not ornamented innerly; ornamented outerly; sharp tip; curved tip; outwards; without medial constriction; hyaline process absent.

##### FEMALE

Body smaller and slenderer than male; Female body 1080 micrometers excluding caudal setae. Widest at first metasome segment. Distal margin of the prosomal segments without one line of setules at posterior margin. Prosome segments with spinules at least at one prosomal segment. Fourth metasome segment absence of dorsal protuberance. Fourth and fifth metasome segments separated. Limit between fourth and fifth metasome segments ornamented; with spinules; as a row; single; incomplete; same size; partially over limit; laterally. **Fifth metasome segment**. Fifth metasome segment without sensilla; with epimeral plates. Epimeral plates asymmetrical. Right epimeral plates prominent, as projections; thinner than the left; one posterior-laterally directed; not reaching half length of the genital segment; with sensilla at the apex; dorsal-posterior sensilla absent; ornamented; with spinules; as a patch; innerly; on dorsal surface. Left epimeral plate without expansion.

##### Urosome

3-segmented. **Genital double-somite**. Asymmetrical in dorsal view; longer than broad; longer than other urosomites combined; dorsal suture at mid-length absent; not covered by spinules; with swelling; rounded; unequal size; greater left than right; anteriorly; with sensillae; on both sides; one; stout; with robust apex; at left lateral; not on lobular base; anteriorly; one; stout; at right lateral; not on lobular base; anteriorly; with robust apex; of equal size between then; lateral protuberance absent; with right posterior rim expanded; over next segment; without slender sensilla on each posterior rim; without posterior-dorsal process. Genital double-somite opercular pad present; broader than longer; symmetrical; development laterally; expanded posteriorly; covering partially; double gonoporal slit; located ventrally; with arthrodial membrane; inserted anteriorly; post-genital process absent; disto-ventral tumescence absent; ventral vertical folds absent; dorsal sensilla absent. Second urosome segment without ventral fusion to anal segment; right distal process absent. Caudal rami patch of setules on outer surface absent; patch of spinules on outer surface absent.

##### Oral appendices feature

Rostrum basal process absent. **Antennules**. Symmetrical. Right antennule surpassing to genital double-segment; extending beyond caudal rami. Right antennule exceeding the caudal setae. Right antennule ornamentation pattern equals to male left antennule; fully.

##### Fifth swimming legs

Symmetrical; Fifth swimming legs biramous. Fifth swimming legs intercoxal plate longer than wide; separated from the legs. Fifth swimming legs praecoxa with sclerite praecoxal; separated from the coxae; without ornamentation. Fifth swimming legs coxa with process; conical; at the outer rim; distally; sensilla present; stout; at apex; projecting over basal segment; no longer 2x than basal insertion; marginal extension present; sub-triangular; at inner rim; distally inserted; without swelling; without seta; without spinules. Fifth swimming legs basis sub-triangular; unequal size between inner and outer sides; shorter outer than inner side; with convex inner side; without proximal inner outgrowth; without groove; with distal extension; on posterior surface; with seta; outerly inserted; on anterior surface; no longer 2x than origin segment. Fifth swimming legs endopod segments 1 and 2 fused; segments 2 and 3 fused; 2-segmented; with incomplete suture; stout; separated from the basis; with spinules; as a row; single; oblique; sub-terminally; at anterior surface; with seta; single; one distally; on posterior surface; rectilinear. Fifth swimming legs exopod segments 1 and 2 separated; segments 2 and 3 separated; 3-segmented; separated from the basis. Fifth swimming legs exopod 1 sub-cylindrical; longer than wide; longer or equal than 2 times; with unequal size between inner and outer side; shorter inner than outer side; with rectilinear inner side; with concave outer side; without swelling; without marginal extension; without posterior process; without spine; without seta. Fifth swimming legs exopod 2 sub-cylindrical; broader than long; no longer or equal than 2 times; without swelling; without marginal extension; without process; without lobe; with spine; inserted laterally; rectilinear; without ornamentation; sharp tip; equal size or larger than next segment; without seta. Fifth swimming legs exopod 3 cylindrical; longer than wide; without swelling; without process; without lobe; without spine; with seta; double; inserted terminally; unequal size between them; outer seta smaller than inner; nearly 1 time; outer seta not ornamented by setules; without ornamentation; presence of terminal claw; sclerotized; arched; internally directed; concave inner side; with ornamentation; of denticles; as a row; on surface partially; at medial region; rectilinear outer side; with ornamentation; of denticles; as a row; on surface partially; at medial region; blunt tip; 6 times longer than origin segment.

##### Distribution records

###### BRAZIL

**Mato Grosso do Sul**: Southern Pantanal, Corumbá Region, Paraguay River: near to Marinha Ladário (19°02’S, 57°34’W), near Port, near Corumbá’s entrance, 2nd access, Corumbá (19°00’S, 57°40’W); Capivari River: Berenice Farm (Reid & Moreno, 1990); Guaraná and Paraná River Basins (Lansac-Tôha *et al*., 1997). **São Paulo**: Ilha Solteira Reservoir (Dussart & Matsumura-Tundisi, 1986; Matsumura-Tundisi, 1986; Sendacz, 1997). **Paraná**: Itaipu Reservoir (Matsumura-Tundisi, 1986); Paraná River and Jupiá Reservoir (Sendacz, 1997). ARGENTINA. **Corrientes**: (Dussart & Frutos, 1986); Laguna 1 (La Turbia), Isla del Cerrito, Paraná River and Laguna 2 (Los Pajaros), Nueva Cerrito Island, Paraná River (Frutos, 1993). **Santa Fé**: Salado River, Juan de Garay Lagoon, near to Santo Tomé (José de Paggi & Paggi, 1988).

##### Habitat

Habitat in freshwaters: rivers, reservoirs, and lagoon.

##### Remarks

Morphological attributes of organisms collected in Southeastern Brazil based the foundation of this species, which presents inconsistencies along its trajectory. *N. spinuliferus* was founded in 1985 by Dussart and has authority attributed to Dussart & Matsumura-Tundisi in this work. However, even if the specified assignment is as “*cf. Dussart & Matsumura-Tundisi, sous presse”,* the description offered by Dussart (1985) satisfies the criteria of availability (article 11) and publication (article 8) of the CINZ (1999), therefore, represents the legitimate nomenclatural act for the taxon. The type-material reported for the MZUSP (Dussart & Matsumura-Tundisi, 1986) was located with the compromised holotype, a female was selected from the paratypes and added as a neotype formally (Geraldes-Primeiro *et al*., 2021).

The taxonomic instability attributed to *N. spinuliferus* is the result of the latest studies focused on the species. A year after its foundation Dussart & Matsumura-Tundisi (1986) offered images that did not corroborate the reports presented. Matsumura-Tundisi (2008) in response to this pointed out these misconceptions and proposed rectification to the original description, but with illustrations and photomicrographs that added new inconsistencies and contributed to the instability of the taxon. Comparison of the images (Matsumura-Tundisi, 2008, Matsumura-Tundisi, 1986) reveals noticeable differences, especially between them, Dussart (1985), Dussart & Matsumura-Tundisi (1986), and Paggi (2001).

In the review offered in Paggi (2001) the female of the species was identified with (1) fifth swimming legs endopod with segmentation partially; (2) fifth metasome segment with epimeral plates ornamented with spinules patch innerly; (3) antennule surpassing to caudal setae, and actual segment 11 with 2 setae; (4) second urosome segment without ventral fusion and similar size as anal segment; and male (5) fifth right swimming leg coxa with distal rounded process projecting over basis until distal surface posteriorly, exopod 2 with 1 chitinous knob posteriorly, and outer spine innerly directed; and (6) fifth left swimming leg basis with outer seta reaching to distal exopod 1. The observations recorded in this work represent an absolute discrepancy in relation to the original description (Dussart, 1985), later description (Dussart & Matsumura-Tundisi, 1986) and “rectification” offered for the species (Matsumura-Tundisi, 2008). Additionally, these attributes were recorded as differential morphology to the taxon most closely related in the work, *N. dentatus*.

In this present effort we corroborate all the observations recorded by Paggi (2001), except for the need to elucidate the condition for the female fifth swimming legs endopod registered with partial segmentation. During our examinations, we recognized the 2-segmented female structure with incomplete suture medially. In other morphological analyses in *Notodiaptomus* for this approach it is possible to recognize the female fifth swimming legs endopod 1-segmented with inner discontinuity on cuticle. This morphological clarification is important for the objectivity and reproduction of future examinations on the condition of this structure, in no way a mistake in Paggi (2001).

In contrast, analyzing the redescription offered in Paggi (2001) for male right antennule we find an involuntary misunderstanding of the definition of spiniform modified seta on actual segment 13, treated as spinous process (*error* “spine-like process”). We assume that the process should be a cuticular expansion devoid of basal suture under the junction or joint with its segment of origin (*sensu* Santos-Silva *et al*., 2015), we recognize the structure indicated by the authors as cuspidate element, bifid apex, provided with junction suture to actual segment 13, therefore, seta truly (*sensu* Garm & Watling, 2013). Notwithstanding, the work represents to date the most complete redescription of *N. spinuliferus*.

Finally, the species of Dussart & Matsumura-Tundisi can be recognized as one of the currently accepted members in *Notodiaptomus* most divergent from the characteristics defined for the creation of the genus. When considering the original attributes set (Kiefer, 1936) and amplification (Kiefer, 1956), the organisms of the species diverge with the following conditions: (1) male fifth right swimming leg basis without inner tumescence proximally; (2) male fifth right swimming leg exopod 2 without distal curved ridge innerly; (3) female fifth swimming legs endopod 2-segmented; and (4) male fifth right swimming leg basis without posterior protrusion. From the attributes of the type-species converging within *Notodiaptomus,* the taxon was divergent mainly: (1) male fifth left swimming leg reaching to first right exopod segment distally; (2) male fifth right swimming leg reaching to first right basis with posterior groove not reaching to endopodal lobe; (3) female limit between fourth and fifth metasome segments with ornamentation; (4) female fifth metasome segment with epimeral plates ornamented; and (5) female fifth swimming leg basis with outer seta, not reaching to exopod 1 distal.

#### Notodiaptomus transitans (Kiefer, 1929)

##### Synonymy

*Diaptomus transitans* Kiefer, 1929: 307, figs. 4a-d; Wright, 1938b: 562; 1939: 648; Brehm, 1958a: 167; Brandorff, 1972: 52; Brandorff, 1978a: 298; Dussart, 1984b: 255; Dussart, 1985a: 201; Forró, 1986: 560, tab. 1; Reid, 1991: 738. *Diaptomus pygmaeus* (Pearse, 1906) Brehm, 1956b: 543–545, figs. (Abb.) 4–7; 1960: 52; Dussart & Defaye, 1983: 64. *Diaptomus* s.l. *mildredae* Brehm, 1960: 52–54, figs. 114– 116; Dussart & Defaye, 1983: 64; Brandorff, 1972: 51; 1976: 618, fig. 3; Dussart, 1984b: 255; 1985a: 201. *Notodiaptomus transitans* n. comb., Ringuelet, 1958a: 45, 46, 54; Brandorff, 1976: 616, fig. 2; Löffler, 1981: 15; Dussart & Defaye, 1983: 136; Dussart & Frutos, 1986: 306, 307; 1987: 244, 245, 246; Matsumura-Tundisi, 1986: 537, 542, figs. 38–42, 100; Reid, 1991: 738; Battistoni, 1995: 959; Rocha *et al*., 1995: 156; Santos-Silva, 2008: 34, fig. 7; Perbiche-Neves *et al*., 2020: 676, key to the Neotropical diaptomid. *Notodiaptomus* (*Caleodiaptomus*) *transitans*; Dussart, 1985a: 214.

##### Type locality

San Roque Reservoir, Primeiro River, Argentina.

##### Type material

No specified in the original description, probably inexistent, collected on 21.XII.1926.

##### Material examined

Non-type material: 2 males and 1 female, entire in alcohol, from the original collection of Friedrich Kiefer (n. 4077) collected in various pools in Paraguay. 1 male (INPA-COP059, slides a-h) and 1 female (INPA-COP060, slides a-h) were selected to be dissection on eight slides each and deposited in the Zoological Collection of the INPA, Brazil. Additional material examined: 1 male and 3 females entire in alcohol from the flood water in Rio Grande do Sul, Brazil, M. G. Bandeira coll., 30.x.2017; 2 males and 1 female entire in alcohol from the Yaciretá Dam Reservoir, Parana River Basin between Argentina, and Paraguay, (27°24“24’ S 56°15“9’ W), Perbiche-Neves coll., 2010.

##### Diagnosis

**(1)** male right antennule surpassing to genital segment, not extending beyond caudal rami; **(2)** male right antennule actual segment 11 with two setae; **(3)** male fifth left swimming leg endopod 1-segmented; **(4)** male fifth left swimming leg exopod 2 with spiniform seta surpassing to distal-point of original segment; **(5)** male fifth right swimming leg exopod 2 with posterior proximal swelling, and triangular inner process posteriorly; **(6)** male fifth right swimming leg exopod 2 with inward arched terminal claw, without conspicuous curve; **(7)** female fourth and fifth metasome segments fused partially on dorsal surface, lateral suture present; **(8)** female fourth metasome segment with dorsal protuberance in digitiform; **(9)** female antennule length surpassing to the caudal setae; **(10)** female fifth swimming legs endopod 2-segmented.

##### Redescription

###### MALE

Body 1470 micrometers excluding caudal setae. Male body smaller and slenderer than female. Nerve axons myelinated. Prosome 6-segmented; widest at first metasome segment; without one line of setules at posterior margin; without spinules at segments. Cephalosome anterior margin rounded; with dorsal suture; incomplete; separate from first metasome segment. First metasome segment without sensilla. Second metasome segment without sensilla. Third metasome segment without sensillae; non-ornamented posterior margin. Fourth metasome segment without sensillae; separated from the fifth metasome. Limit between fourth and fifth metasome segments without ornamentation. Fifth metasome segment without sensilla; Fifth metasome segment without ornamentation; Fifth metasome segment without dorsal conical process; with epimeral plates. Epimeral plates symmetrical. Right epimeral plates reduced, as rounded distal corner segment limit; with sensilla; one at the apex of projection and other medially; without ornamentation.

##### Urosome

5-segmented; Urosome 5 - free segments. Genital somite symmetrical in dorsal view; with single aperture; located on left side; ventrolaterally on posterior rim; with sensillae; on both sides; one; at left lateral; posteriorly; one; at right rim; posteriorly; of equal size between then. Third urosome segment without spinules; without external seta. Fourth urosome segment without spinules; without sub-conical blunt dorsal-lateral process. Anal segment presence of dorsal sensillae; one on each side; medially inserted; presence of operculum; convex; covering the anal aperture fully. Caudal rami symmetrical; separated from anal segment; longer than wide; with setules; continuous on; inner side; each ramus bearing 6 caudal setae; 5 marginals; plumose; and 1 internal dorsally; straight; not reticulated main axis; outermost seta with outer spiniform process absent.

##### Oral appendices feature

Rostrum asymmetrical; separated from dorsal cephalic shield; by complete suture; sensillae present; one pair; anteriorly inserted on surface tegument; with rostral filament; double; paired; extended; into blunt protrusion; with basal process; in ventral view, rounded on left side; with a smaller basal expansion on the right side.

##### Antennules

Asymmetrical. **Right antennules**. Uniramous; right antennule surpassing to genital segment; right antennule not extending beyond caudal rami.

Right antennule ancestral segment I and II separated. Ancestral segment II and III fused. Ancestral segment III and IV fused. Ancestral segment IV and V separated. Ancestral segment V and VI separated. Ancestral segment VI and VII separated. Ancestral segment VII and VIII separated. Ancestral segment VIII and IX separated. Ancestral segment IX and X separated. Ancestral segment X and XI separated. Ancestral segment XI and XII separated. Ancestral segment XII and XIII separated. Ancestral segment XIII and XIV separated. Ancestral segment XIV and XV separated. Ancestral segment XV and XVI separated. Ancestral segment XVI and XVII separated. Ancestral segment XVII and XVIII separated. Ancestral segment XVIII and XIX separated. Ancestral segment XIX and XX separated. Ancestral segment XX and XXI separated. Ancestral segment XXI and XXII fused. Ancestral segment XXII and XXIII fused. Ancestral segment XXIII and XXIV separated. Ancestral segment XXIV and XXV fused. Ancestral segment XXV and XXVI separated. Ancestral segment XXVI and XXVII separated. Ancestral segment XXVII and XXVIII fused.

Right antennule actual 22-segmented; geniculated; between the segment 18 and segment 19; with swollen and modified region; formed by 5 segments; between 13 and 17 segments. Actual segment 1 with seta; one element; straight; none larger than segment; without spinules; without vestigial seta; without conical seta; without modified seta; without spinous process; with aesthetasc; one element. Actual segment 2 with seta; three elements; of unequal size; straight; none larger than segment; without spinules; with vestigial seta; one element; without conical seta; without modified seta; without spinous process; with aesthetasc; one element. Actual segment 3 with seta; one element; none larger than segment; straight; sharp apex; without spinules; with vestigial seta; one element; without conical seta; without modified seta; without spinous process; with aesthetasc. Actual segment 4 with seta; one element; one larger than segment; surpassing to distal margin; straight; not beyond three sequential segments; without spinules; without vestigial seta; without conical seta; without modified seta; without spinous process; without aesthetasc. Actual segment 5 with seta; one element; straight; one larger than segment; surpassing to distal margin; not beyond three sequential segments; without spinules; with vestigial seta; one element; without conical seta; without modified seta; without spinous process; without aesthetasc. Actual segment 6 with seta; one element; none larger than segment; straight; without spinules; without vestigial seta; without conical seta; without modified seta; without spinous process; without aesthetasc. Actual segment 7 with seta; one element; straight; none larger than segment; sharp apex; without spinules; without vestigial seta; without conical seta; without modified seta; without spinous process; with aesthetasc; one element. Actual segment 8 with seta; one element; straight; none larger than segment; without spinules; without vestigial seta; with conical seta; one element; not reaching to middle-point of the sequent segment; without modified seta; without spinous process; without aesthetasc. Actual segment 9 with seta; two elements; of unequal size; straight; none larger than segment; sharp apex; without spinules; without vestigial seta; without conical seta; without modified seta; without spinous process; with aesthetasc; one element. Actual segment 10 with seta; one element; straight; none larger than segment; without spinules; without vestigial seta; without conical seta; with modified seta; presenting blunt apex; slender form; surpassing to distal margin; not beyond of the sequential segment; parallel to antennule direction; without spinous process; without aesthetasc. Actual segment 11 with seta; two elements; of equal size; straight; one larger than segment; surpassing to distal margin; not beyond three sequential segments; without spinules; without vestigial seta; without conical seta; with modified seta; slender form; presenting blunt apex; surpassing to distal margin; not beyond of the sequential segment; perpendicular to antennule direction; shorter length than homologous of actual segment 13; without spinous process; without aesthetasc. Actual segment 12 with seta; one element; straight; one larger than segment; surpassing to distal margin; not beyond three sequential segments; without spinules; without vestigial seta; with conical seta; one element; not smaller than to segment 8; without modified seta; without spinous process; with aesthetasc; one element; absent internal perpendicular fission. Actual segment 13 with seta; one element; straight; one larger than segment; surpassing to distal margin; not beyond three sequential segments; without spinules; without vestigial seta; without conical seta; with modified seta; stout form; surpassing to distal margin; to the middle-point of the sequence segment; perpendicular to antennule direction; presenting rounded apex; without spinous process; with aesthetasc; one element. Actual segment 14 with seta; two elements; of unequal size; straight; one larger than segment; surpassing to distal margin; beyond three sequential segments; blunt apex; without spinules; without vestigial seta; without conical seta; without modified seta; without spinous process; without aesthetasc. Actual segment 15 with seta; two elements; of unequal size; straight; not bifidform; none larger than segment; without spinules; without vestigial seta; without conical seta; without modified seta; with spinous process; on outer margin; surpassing distal margin; with aesthetasc; one element. Actual segment 16 with seta; two elements; of unequal size; plumose; one larger than segment; surpassing to distal margin; not beyond three sequential segments; not bifidform; without spinules; without vestigial seta; without conical seta; without modified seta; with spinous process; on outer margin; surpassing distal margin; unequal size to process on preceding segment; with aesthetasc; one element. Actual segment 17 with seta; two elements; of unequal size; straight; none larger than segment; bifidform; without spinules; without vestigial seta; without conical seta; with modified seta; one element; stout form; surpassing to distal margin; not beyond of the sequential segment; parallel to antennule direction; without spinous process; without aesthetasc. Actual segment 18 with seta; two elements; of equal size; straight; none larger than segment; without spinules; without vestigial seta; without conical seta; with modified seta; one element; stout form; surpassing distal margin; parallel to antennule direction; without spinous process; without aesthetasc. Actual segment 19 with seta; two elements; of unequal size; plumose; none larger than segment; without spinules; without vestigial seta; without conical seta; with modified seta; two elements; stout form; at least one bifid form; surpassing distal margin; parallel to antennule direction; without spinous process; with aesthetasc; one element. Actual segment 20 with seta; four elements; of unequal size; straight; one larger than segment; surpassing to distal margin; beyond three sequential segments; without spinules; without vestigial seta; without conical seta; without modified seta; with spinous process; distally; not reaching beyond of distal-point segment 21; without aesthetasc. Actual segment 21 with seta; two elements; of equal size; plumose; one larger than segment; surpassing to distal margin; greater 3x than original segment; without spinules; without vestigial seta; without conical seta; without modified seta; without spinous process; without aesthetasc. Actual segment 22 with seta; four elements; of equal size; one larger than segment; plumose; surpassing to distal margin; greater 3x than original segment; without spinules; without vestigial seta; without conical seta; without modified seta; without spinous process; with aesthetasc; one element.

##### Left antennules

Uniramous; Left antennule surpassing to prosome; Left antennule not extending beyond caudal rami. Ancestral segment I and II separated. Ancestral segment II and III fused. Ancestral segment III and IV fused. Ancestral segment IV and V separated. Ancestral segment V and VI separated. Ancestral segment VI and VII separated. Ancestral segment VII and VIII separated. Ancestral segment VIII and IX separated. Ancestral segment IX and X separated. Ancestral segment X and XI separated. Ancestral segment XI and XII separated. Ancestral segment XII and XIII separated. Ancestral segment XIII and XIV separated. Ancestral segment XIV and XV separated. Ancestral segment XV and XVI separated. Ancestral segment XVI and XVII separated. Ancestral segment XVII and XVIII separated. Ancestral segment XVIII and XIX separated. Ancestral segment XIX and XX separated. Ancestral segment XX and XXI separated. Ancestral segment XXI and XXII separated. Ancestral segment XXII and XXIII separated. Ancestral segment XXIII and XXIV separated. Ancestral segment XXIV and XXV separated. Ancestral segment XXV and XXVI separated. Ancestral segment XXVI and XXVII separated. Ancestral segment XXVII and XXVIII fused.

Left antennule actual 25-segmented; not-geniculated. Actual segment 1 with seta; one element; none larger than segment; straight; without spinules; without vestigial seta; without conical seta; without modified seta; without spinous process; with aesthetasc; one element. Actual segment 2 with seta; three elements; of equal size; none larger than segment; straight; without spinules; with vestigial seta; one element; without conical seta; without modified seta; without spinous process; with aesthetasc; one element. Actual segment 3 with seta; one element; one larger than segment; straight; surpassing to distal margin; beyond three sequential segments; without spinules; with vestigial seta; one element; without conical seta; without modified seta; without spinous process; with aesthetasc. Actual segment 4 with seta; one element; none larger than segment; straight; without spinules; without vestigial seta; without conical seta; without modified seta; without spinous process; without aesthetasc. Actual segment 5 with seta; one element; one larger than segment; straight; surpassing to distal margin; not beyond three sequential segments; without spinules; with vestigial seta; one element; without conical seta; without modified seta; without spinous process; with aesthetasc; one element. Actual segment 6 with seta; one element; none larger than segment; straight; without spinules; without vestigial seta; without conical seta; without modified seta; without spinous process; without aesthetasc. Actual segment 7 with seta; one element; one larger than segment; straight; surpassing to distal margin; beyond three sequential segments; without spinules; without vestigial seta; without conical seta; without modified seta; without spinous process; with aesthetasc; one element. Actual segment 8 with seta; one element; one larger than segment; straight; surpassing distal margin; without spinules; without vestigial seta; with conical seta; without modified seta; without spinous process; without aesthetasc. Actual segment 9 with seta; two elements; of unequal size; one larger than segment; straight; surpassing to distal margin; beyond three sequential segments; without spinules; without vestigial seta; without conical seta; without modified seta; without spinous process; with aesthetasc; one element. Actual segment 10 with seta; one element; none larger than segment; straight; without spinules; without vestigial seta; without conical seta; without modified seta; without spinous process; without aesthetasc. Actual segment 11 with seta; one element; one larger than segment; straight; surpassing to distal margin; beyond three sequential segments; without spinules; without vestigial seta; without conical seta; without modified seta; without spinous process; without aesthetasc. Actual segment 12 with seta; one element; one larger than segment; straight; surpassing distal margin; without spinules; without vestigial seta; with conical seta; without modified seta; without spinous process; with aesthetasc; one element. Actual segment 13 with seta; one element; none elongated; straight; surpassing distal margin; without spinules; without vestigial seta; without conical seta; without modified seta; without spinous process; without aesthetasc. Actual segment 14 with seta; one element; elongated; straight; surpassing to distal margin; beyond three sequential segments; without spinules; without vestigial seta; without conical seta; without modified seta; without spinous process; with aesthetasc; one element. Actual segment 15 with seta; one element; larger than segment; straight; surpassing to distal margin; not beyond three sequential segments; without spinules; without vestigial seta; without conical seta; without modified seta; without spinous process; without aesthetasc. Actual segment 16 with seta; one element; larger than segment; plumose; surpassing to distal margin; not beyond three sequential segments; without spinules; without vestigial seta; without conical seta; without modified seta; without spinous process; with aesthetasc; one element. Actual segment 17 with seta; one element; not larger than segment; straight; without spinules; without vestigial seta; without conical seta; without modified seta; without spinous process; without aesthetasc. Actual segment 18 with seta; one element; larger than segment; straight; surpassing to distal margin; beyond three sequential segments; without spinules; without vestigial seta; without conical seta; without modified seta; without spinous process; without aesthetasc. Actual segment 19 with seta; one element; not larger than segment; straight; surpassing distal margin; without spinules; without vestigial seta; without conical seta; without modified seta; without spinous process; with aesthetasc; one element. Actual segment 20 with seta; one element; not larger than segment; straight; surpassing distal margin; without spinules; without vestigial seta; without conical seta; without modified seta; without spinous process; without aesthetasc. Actual segment 21 with seta; one element; larger than segment; plumose; surpassing to distal margin; beyond three sequential segments; without spinules; without vestigial seta; without conical seta; without modified seta; without spinous process; without aesthetasc. Actual segment 22 with seta; two elements; of unequal size; one of them elongated; plumose; surpassing to distal margin; without spinules; without vestigial seta; without conical seta; without modified seta; without spinous process; without aesthetasc. Actual segment 23 with seta; two elements; of unequal size; one larger than segment; plumose; surpassing to distal margin; greater 3x than original segment; without spinules; without vestigial seta; without conical seta; without modified seta; without spinous process; without aesthetasc. Actual segment 24 with seta; two elements; of equal size; one larger than segment; plumose; surpassing to distal margin; greater 3x than original segment; without spinules; without vestigial seta; without conical seta; without modified seta; without spinous process; without aesthetasc. Actual segment 25 with seta; four elements; of equal size; elongated; plumose; surpassing to distal margin; 4 times larger than segment; without spinules; without vestigial seta; without conical seta; without modified seta; without spinous process; with aesthetasc; one element.

##### Antenna

Biramous. Antenna coxa separated from the basis; bearing seta; 1; on inner surface; at distal corner; reaching to the endopod 1. Antenna basis (fusion) separated from the endopodal segment; bearing seta; 2; on inner surface; at distal corner. Endopodal ancestral segment I and II separated. Ancestral segment II and III fused. Ancestral segment III and IV fused. Ancestral segment III and IV fully. Antenna endopod actual 2-segmented. Actual segment 1 not bilobate; with seta; two; on inner margin; with spinules; as a row; obliquely; on outer surface; without pore. Actual segment 2 bilobate; with discontinuity on outer cuticle; developed as a suture; incomplete; inner lobe bearing 8 setae; distally; outer lobe bearing 7 setae; distally; with spinules; as a patch; on outer surface. Antenna exopod ancestral segment I and II separated. Ancestral segment II and III fused. Ancestral segment III and IV fused. Ancestral segment IV and V separated. Ancestral segment V and VI separated. Ancestral segment VI and VII separated. Ancestral segment VII and VIII separated. Ancestral segment VIII and IX separated. Ancestral segment IX and X fused. Antenna exopod actual 7-segmented. Actual segment 1 single; elongated (width-length, equal or larger ratio 2:1); with seta; one; at inner surface. Actual segment 2 compound; elongated (larger width-length ratio 2:1); with seta; three; at inner surface. Actual segment 3 single; not elongated (lesser width-length ratio 2:1); with seta; one; at inner surface. Actual segment 4 single; not elongated (lesser width-length ratio 2:1); with seta; one; at inner surface. Actual segment 5 single; not elongated (lesser width-length ratio 2:1); with seta; one; at inner surface. Actual segment 6 single; not elongated (lesser width-length ratio 2:1); with seta; one; at inner surface. Actual segment 7 compound; elongated (larger or equal width-length ratio 2:1); with seta; one; at inner surface; and three; at distal surface.

##### Oral features

**Mandible**. Coxal gnathobase sclerotized; with lobe; prominent; on caudal margin; presence of cutting blade; with tooth-like prominence; two, distinctly; 1 acute; on caudal margin; and 1 triangular; on sub-caudal margin; without acute projection between the prominences; without additional spinules; with seta; 1; dorsally; on apical surface; without spinules. Mandible palps biramous; comprising the basis; with seta; four; differently inserted; first medially; reaching to beyond the endopod 1; second distally; third distally; fourth distally; on inner margin; none with setulose ornamentation. Mandible endopod 2-segmented. Mandible endopod 1 with lobe; bearing seta; four; distally inserted; without spinules. Mandible endopod 2 without lobe; bearing setae; nine elements; distally inserted; with spinules; as a patch; double. Mandible exopod 4-segmented. Mandible exopod 1 with seta; one element; distally; on inner margin. Mandible exopod 2 with seta; one element; distally; on inner side. Mandible exopod 3 with seta; one element; distally; on inner side. Mandible exopod 4 with setae; three elements; on terminal region. **Maxillule**. Birramous. Maxillule 3-segmented. Maxillule praecoxa with praecoxal arthrite; bearing spines; fifteen elements; ten marginally; plus, five sub-marginally; with spinules; as a patch; on sub-marginal surface. Maxillule coxa with coxal epipodite; with conspicuous outer lobe; bearing setae; nine elements; with coxal endite; elongated (larger or equal width-length ratio 2:1); bearing setae; four elements. Maxillule basis with basal endite; double; first proximal; elongated (larger width-length ratio 2:1; separated from basis; with setae; four elements; distally inserted; second distal; fused to basis; not elongated (lesser width-length ratio 2:1); with setae; four elements; distally inserted; with setules; as a row; on inner side; basal exite present; with setae; one element; on outer surface. Maxillule endopod 1-segmented. Endopod 1 bilobate; first proximal; with setae; three elements; second distal; with setae; five elements. Maxillule exopod 1-segmented. Exopod 1 with setae; six elements; with setules; as a row; on inner side; spinules absent. **Maxilla**. Uniramous. Maxilla 5-segmented. Maxilla praecoxa fused to coxa; incompletely; distinct externally; with praecoxal endite; double; first elongated endite (larger or equal width length ratio 2:1); proximally inserted; with seta; straight, or plumose; 1 straight; 4 plumose; with spine; single; without spinules; without setule; second elongated endite (larger or equal width length ratio 2:1); distally inserted; with seta; plumose; 3 plumose; without spine; with spinules; as a row; on distal margin; with setule; as a row; on distal margin; absence of outer seta. Maxilla coxa with coxal endite; double; first elongated endite (larger or equal width); proximally inserted; with seta; plumose; 3 plumose; without spine; without spinules; with setules; as a row; on proximal margin; second elongated endite (larger or equal width); distally inserted; with seta; plumose; 3 plumose; without spine; without spinules; with setules; as a row; on proximal margin; absence of outer seta. Maxilla basis with basal endite; single; elongated (larger or equal width-length ratio 2:1); with seta; plumose; 3 plumose; without spinules; absence of outer seta. Maxilla endopod 2-segmented. Endopod 1 with seta; 2 plumose; without spine; without spinules; without setules. Maxilla endopod 2 with seta; 2 plumose; without spine; without spinules; without setules. **Maxilliped**. Uniramous; Maxilliped 8-segmented. Maxilliped praecoxa fused to coxa; completely; with praecoxal endite; not elongated (lesser width-length ratio 2:1); distally inserted; with seta; 1 straight; with spinules; as a row; single; on basal surface; without setules. Maxilliped coxa with coxal endite; three coxal endite; first elongated (larger or equal width); proximally inserted; with seta; 2 plumose; with spinules; as a patch; single; on apical surface; without setules; second not elongated (lesser width-length ratio 2:1); medially inserted; with seta; 3 plumose; with spinules; as a row; single; on medial surface; without setules; third elongated (larger or equal width length ratio 2:1); distally inserted; with seta; 3 plumose; none reaching to beyond of the basis; with spinules; as a row; single; on basal surface; without setules; with lobe; prominence; at inner distal angle; ornamented; with spinules; continuously on margin. Maxilliped basis without basal endite; with seta; 3 plumose; with spinules; as a row; single; on medial surface; with setules; as a row; single; on inner margin. Maxilliped endopod segment 6-segmented. Endopod 1 with seta; 2 plumose; on inner surface. Endopod 2 with seta; 3 plumose; on inner surface. Endopod 3 with seta; 2 plumose; on inner surface. Endopod 4 with seta; 2 plumose; on inner surface. Endopod 5 with seta; 2 plumose; on inner surface, or on outer surface; outer seta absent. Endopod 6 with seta; 4 plumose; on inner surface, or on outer surface.

##### Swimming legs features

**First swimming legs.** Symmetrical; biramous. First swimming legs intercoxal plate without seta. First swimming legs praecoxa absent. First swimming legs coxa with seta; one; straight; distally inserted; on inner surface; surpassing to first endopodal segment; with setules; two group; as a patch; on inner margin; and as a row; double; on anterior surface; outerly; without spinules; without spine. First swimming legs basis without seta; with setules; as a patch; single; on outer surface; without spinules; without spine. First swimming legs endopod 2-segmented. Endopod 1 with seta; straight; restricted; to inner surface; one element; without spine; with setules; as a row; single; continuously; on outer surface; without spinules; absence of Schmeil’s organ. Endopod 2 with seta; unrestricted; three on inner surface; one on outer surface; two on distal surface; straight; without spine; with setules; as a row; single; continuously; on outer surface; without spinules; absence of Schmeil’s organ. Endopod 3 absence. First swimming legs exopod 1 with seta; restricted; 1 on inner surface; with spine; 1; stout; smaller than original segment; serrated; on inner side; continuously; with setules; as a row; single; as a row; innerly. First swimming legs exopod 2 with seta; restricted; 1 on inner surface; straight; without spine; with setules; as a row; single; continuously; on inner margin, or on outer margin; without spinules. First swimming legs exopod 3 with setule; as a row; single; continuously; on outer surface; without spinules; with seta; unrestricted; 2 on inner surface; 2 on terminal surface; with spine; 2; unequal size; first no longer 2x than origin segment; stout; serrated; on inner side, or on outer side; equally; second longer 3x than origin segment; slender; serrated; on outer side; with ornamentation on non-serrated side; by setules. **Second swimming legs**. Symmetrical; Second swimming legs biramous. Second swimming legs intercoxal plate without seta. Second swimming legs praecoxa present; located laterally. Second swimming legs coxa with seta; straight; distally inserted; on inner surface; surpassing to basal segment; without setules; without spinules; without spine. Second swimming legs basis without seta; without setules; without spinules; without spine. Second swimming legs endopod 3-segmented. Endopod 1 with seta; straight; restricted; one on inner surface; without spine; with setules; as a row; single; continuously; on outer surface; without spinules; absence of Schmeil’s organ. Endopod 2 with seta; straight; unrestricted; two on inner surface; without spine; with setules; as a row; single; continuously; on outer side; without spinules; presence of Schmeil’s organ; on posterior surface. Endopod 3 with seta; straight; unrestricted; three on inner surface; two on outer surface; two on distal surface; without spine; without setules; with spinules; as a row; double; distally inserted; at anterior surface; absence of Schmeil’s organ. Second swimming legs exopod 1 with seta; restricted; one on inner surface; with spine; 1; stout; not reaching to distal-third of the exopod 2; serrated; on inner side, or on outer side; with setules; as a row; single; continuously; on inner side; without spinules; absence of Schmeil’s organ. Exopod 2 with seta; unrestricted; one on inner surface; with spine; 1; stout; not surpassing the exopod 3; serrated; on inner side, or on outer side; with setules; as a row; single; continuously; on inner surface; without spinules; absence of Schmeil’s organ. Exopod 3 with seta; plurimarginal; three on inner surface; two on terminal surface; with spine; 2; unequal size; first no longer 2x than origin segment; stout; serrated; on inner side, or on outer side; equally; second longer 2x than origin segment; slender; serrated; on outer side; with ornamentation on non-serrated side; of setules; setules on outer surface; as a row; single; continuously; on inner surface; with spinules; as a row; single; distally inserted; at anterior surface; absence of Schmeil’s organ. **Third swimming legs**. Symmetrical; Third swimming legs biramous. Third swimming legs intercoxal plate without seta. Third swimming legs praecoxa present; not laterally located. Third swimming legs coxa with seta; straight; distally inserted; on inner surface; surpassing to first endopodal segment; without setules; without spinules; without spine. Third swimming legs basis without seta; without setules; without spinules; without spine. Third swimming legs endopod 3-segmented. Endopod 1 with seta; restricted; one on inner surface; without spine; without setules; without spinules; absence of Schmeil’s organ. Endopod 2 with seta; restricted; two on inner surface; straight; without spine; without setules; without spinules; absence of Schmeil’s organ. Endopod 3 with seta; straight; plurimarginal; two on inner surface; two on outer surface; three on terminal surface; without spine; without setules; with spinules; as a row; distally inserted; double; at anterior surface; absence of Schmeil’s organ. Third swimming legs exopod 1 with seta; restricted; straight; one on inner surface; with spine; 1; stout; not reaching to the distal-third of the exopod 2; serrated; equally; on inner surface, or on outer surface; with setules; as a row; single; continuously; on inner surface; without spinules; absence of Schmeil’s organ. Exopod 2 with seta; straight; restricted; one on inner surface; with spine; 1; stout; not reaching out to exopod 3; serrated; on inner side, or on outer side; equally; with setules; as a row; single; continuously; on inner side; without spinules; absence of Schmeil’s organ. Exopod 3 without setules; with spinules; as a row; single; distally inserted; at anterior surface; with seta; straight; unrestricted; three on inner surface; two on terminal surface; with spine; 2; unequal size; first no longer 2x than origin segment; stout; serrated; on inner side, or on outer side; equally; second longer 2x than origin segment; slender; serrated; on outer side; with ornamentation on non-serrated side; of setules; absence of Schmeil’s organ. **Fourth swimming legs**. Symmetrical; biramous. Intercoxal plate without sensilla. Praecoxa present. Coxa with seta; distally inserted; on inner margin; reaching out to endopod 1; without spinules; setules absent. Basis with seta; one; medially inserted; on posterior surface; smaller than the original segment; without setules; without spinules; without spine. Fourth swimming legs endopod 3-segmented. Endopod 1 with seta; one; restricted; on inner surface; without spine; without setules; without spinules; absence of Schmeil’s organ. Endopod 2 with seta; restricted; two on inner side; without spine; with setules; as a row; single; continuously; on outer surface; without spinules; absence of Schmeil’s organ. Endopod 3 with seta; unrestricted; two on inner surface; two on outer surface; three on distal surface; without spine; without setules; with spinules; as a row; double; distally inserted; at anterior surface; absence of Schmeil’s organ. Fourth swimming legs exopod 1 with seta; restricted; one on inner surface; with spine; 1; stout; not reaching out to distal-third of the exopod 2; serrated; on inner side, or on outer side; equally; with setules; as a row; single; continuously; on inner surface; without spinules; absence of Schmeil’s organ. Exopod 2 with seta; restricted; one on inner surface; with spine; 1; stout; not reaching the end of exopod 3; serrated; on inner side, or on outer side; equally; with setules; as a row; single; continuously; on inner surface; without spinules; absence of Schmeil’s organ. Exopod 3 without setules; with spinules; as a row; single; distally inserted; at anterior surface; with seta; unrestricted; three on inner surface; two on distal surface; with spine; 2; unequal size; first no longer 2x than origin segment; stout; serrated; on inner side, or on outer side; equally; second longer 2x than origin segment; slender; serrated; on outer side; without ornamentation on non-serrated side; absence of Schmeil’s organ.

##### Fifth swimming legs features

Asymmetrical. Fifth swimming leg intercoxal plate with length not equal or greater than width on 1.5x; with irregular proximal margin; discontinuous to; the anterior margin of the left coxa, or the anterior margin of the right coxa; posterior sensilla on the right lateral absent. **Fifth left swimming leg**. Fifth left swimming leg biramous; leg reaching first right exopod segment; medially. Fifth left swimming leg praecoxa present; rudimentary; separated from the coxae; without ornamentation. Fifth left swimming leg coxa concave inner side; without teeth-like structures; with process; conical; on posterior surface; outer side; distally inserted; not projecting over basis; with sensilla; stout; triangular; at apex; longer 2x than insertion basis; without marginal extension; without swelling; without seta; without spinules. Fifth left swimming leg basis sub-cylindrical; unequal size between inner and outer side; shorter outer than inner side; with concave inner side; rounded internal proximal expansion absent; without outgrowth; without groove; absence of protuberance; with seta; outerly inserted; no longer 2x than origin segment; absence of minutely granular. Fifth left swimming leg endopod segments 1 and 2 fused; segments 2 and 3 fused; 1-segmented; stout; separated from the basis; ornamented; on inner side; with spinules; more than four elements; as a row; terminally; row of setules absent; with seta. Fifth left swimming leg exopod segments 1 and 2 separated; segments 2 and 3 fused; 2-segmented; stout; separated from the basis. Fifth left swimming leg exopod 1 sub-triangular; longer than broad; equal size between inner and outer side; rectilinear inner side; convex outer side; without swelling; without marginal extension; without process; with lobe; single; semicircular; medially inserted; on inner side; covered; by setules; without outer spine; absence seta. Fifth left swimming leg exopod 2 digitiform; longer than broad; equal size between inner and outer side; disform inner side; with convex outer side; setulose pad present; prominently rounded; proximally; on inner side; inflated medial region absent; distal process present; digitiform; denticulate; not bicuspidate; with transverse row of denticles; none oblique row of 5 denticles; at anterior surface; not innerly directed; with seta; spiniform; ornamented by spinules; surpassing the distal-point of the segment; without outer spine; terminal claw absent.

##### Fifth right swimming leg

Biramous. Fifth right swimming leg praecoxa present; separated from the coxae; without ornamentation. Fifth right swimming leg coxa convex inner side; without teeth-like structures; with process; rounded; distally inserted; on posterior surface; closest to the outer rim; projecting over basis; not beyond the first third; without triangular protuberance innerly; with sensilla; slender; at apex; no longer 2x than basal insertion; without marginal extension; without seta; without spinules. Fifth right swimming leg basis trapezoidal; unequal size between inner and outer side; shorter outer than inner side; convex inner side; tumescence present; inflated; not bilobed; unrestricted on inner surface; without protuberance; absence of distinct minutely granular; additional inner process absent; without posterior groove; with seta; outerly inserted; on anterior surface; no longer 2x than origin segment; posterior protrusion absent; distal process absent. Fifth right swimming leg with endopodite present; separated from the basis; on anterior surface; ancestral segments 1 and 2 fused; ancestral segments 2 and 3 fused; 1-segmented; slender; ornamented; with setules; as a row; on inner side; terminally; with seta. Fifth right swimming leg exopod segments 1 and 2 separated; segments 2 and 3 fused; 2-segmented; stout; separated from the basis. Fifth right swimming leg exopod 1 trapezium; longer than broad; nearly 1.25 times; unequal size between both sides; shorter inner than outer side; convex inner side; rectilinear outer side; with marginal extension; sub-triangular; distally inserted; at outer rim; spinules absent; with process; rounded; sclerotized; without ornamentation; distally inserted; at posterior surface; projecting over next segment; without outer spine; without seta; internal prominence present; acute; lamella on posterior surface absent. Fifth right swimming leg exopod 2 cylindrical; longer than broad; nearly 2.5 times; unequal size between both sides; uniform inner side; convex outer side; with posterior proximal swelling; inner-posterior process present; sub-triangular; medially; without marginal expansion; curved ridge on distal posterior surface present; chitinous knobs absent; with outer spine; inserted sub-distally; rectilinear; not ornamented innerly; not ornamented outerly; sharp tip; without apparent curve; lesser than the length of the exopod 2; until to 2 times its size; 2x; sensilla absent; terminal claw present; equal or longer 1.5 times than insertion segment; sclerotized; arched; inward; without conspicuous curve; ornamented innerly; by spinules; as a row; partially on extension; medially, or distally; not ornamented outerly; sharp tip; curved tip; inwards; with medial constriction; hyaline process absent.

##### FEMALE

Body longer and wider than male; Female body 1647 micrometers excluding caudal setae. Widest at posterior cephalosome. Distal margin of the prosomal segments without one line of setules at posterior margin. Prosome segments without spinules at prosomal segments. Fourth metasome segment presence of dorsal protuberance; digitiform; 2x longer than wide; inserted medially; without posterior process; without anterior process; fourth metasome segment without proximal sensillae present. Fourth and fifth metasome segments fused; partially; on dorsal surface. Limit between fourth and fifth metasome segments without ornamentation. **Fifth metasome segment**. Fifth metasome segment without sensilla; with epimeral plates. Epimeral plates asymmetrical. Right epimeral plates prominent, as projections; thinner than the left; one posterior-laterally directed; not reaching half length of the genital segment; with sensilla at the apex; dorsal-posterior sensilla present; without ornamentation. Left epimeral plate with expansion; conical; on medial surface; dorsally.

##### Urosome

3-segmented. **Genital double-somite**. Asymmetrical in dorsal view; longer than broad; longer than other urosomites combined; dorsal suture at mid-length absent; not covered by spinules; with swelling; rounded; unequal size; greater right than left; anteriorly; with sensillae; on both sides; one; stout; with robust apex; at left rim; not on lobular base; anteriorly; one; stout; at right lateral; not on lobular base; anteriorly; with robust apex; of equal size between then; lateral protuberance absent; with right posterior rim expanded; over next segment; without slender sensilla on each posterior rim; without posterior-dorsal process. Genital double-somite opercular pad present; broader than longer; symmetrical; development laterally; expanded posteriorly; covering partially; double gonoporal slit; located ventrally; with arthrodial membrane; inserted anteriorly; post-genital process absent; disto-ventral tumescence absent; ventral vertical folds absent; dorsal sensilla absent. Second urosome segment without ventral fusion to anal segment; right distal process absent. Caudal rami patch of setules on outer surface absent; patch of spinules on outer surface absent.

##### Oral appendices feature

Rostrum basal process absent. **Antennules**. Symmetrical. Right antennule surpassing to genital double-segment; extending beyond caudal rami. Right antennule exceeding the caudal setae. Right antennule ornamentation pattern equals to male left antennule; fully.

##### Fifth swimming legs

Symmetrical; Fifth swimming legs biramous. Fifth swimming legs intercoxal plate longer than wide; separated from the legs. Fifth swimming legs praecoxa with sclerite praecoxal; separated from the coxae; without ornamentation. Fifth swimming legs coxa with process; conical; at the outer rim; distally; sensilla present; stout; at apex; not projecting over basal segment; no longer 2x than basal insertion; marginal extension absent; without swelling; without seta; without spinules. Fifth swimming legs basis sub-triangular; unequal size between inner and outer sides; shorter outer than inner side; with convex inner side; without proximal inner outgrowth; without groove; with distal extension; on posterior surface; with seta; outerly inserted; on anterior surface; no longer 2x than origin segment. Fifth swimming legs endopod segments 1 and 2 separated; segments 2 and 3 fused; 2-segmented; with complete suture; stout; separated from the basis; ornamentation on segment 2; with spinules; as a row; single; non-oblique; sub-terminally; at anterior surface; with seta; double; one medially; on posterior surface; rectilinear; one distally; on posterior surface; arched; of unequal size; proximal seta longer than distal seta. Fifth swimming legs exopod segments 1 and 2 separated; segments 2 and 3 separated; 3-segmented; separated from the basis. Fifth swimming legs exopod 1 sub-cylindrical; longer than wide; longer or equal than 2 times; with equal size between inner and outer sides; with convex inner side; with rectilinear outer side; without swelling; without marginal extension; without posterior process; without spine; without seta. Fifth swimming legs exopod 2 sub-cylindrical; longer than broad; longer or equal than 2 times; without swelling; without marginal extension; without process; without lobe; with spine; inserted laterally; rectilinear; without ornamentation; sharp tip; smaller than next segment; without seta. Fifth swimming legs exopod 3 cylindrical; longer than wide; without swelling; without process; without lobe; without spine; with seta; double; inserted terminally; unequal size between them; outer seta smaller than inner; nearly 3 times; outer seta not ornamented by setules; without ornamentation; presence of terminal claw; sclerotized; rectilinear; with ornamentation; of denticles; as a row; on surface partially; at medial region; concave outer side; with ornamentation; of denticles; as a row; on surface partially; at medial region; blunt tip; 6 times longer than origin segment.

##### Distribution records

###### BRAZIL

**São Paulo**: Capivara Reservoir, Paranapanema River basin (Matsumura-Tundisi, 1986). **Paraná**: Itaipu Reservoir (Matsumura-Tundisi, 1986). PARAGUAY. (Kiefer, 1929). ARGENTINA. **Chaco**: Oro River (Dussart & Frutos, 1987). **Córdoba**: (Wright, 1938b); pond connected Terceiro River; pond of San Roque; lake at Sarmiento Park; Córdoba City (Wright, 1939); San Marcos (Brehm, 1956b); San Roque Pond, Primero River (Reid, 1991). **Yaciretá:** (Perbiche-Neves *et al*., 2015), Dam Reservoir, Parana River between Argentina and Paraguay. PARAGUAY. (Kiefer’s collection, Kiefer, 1929) **URUGUAY**. **Barras Agas. Montevideo:** Barra Santa Lucía. Argentina: Buenos Aires, “Balneário Norte de Nunez”.

##### Habitat

Habitat in freshwaters: reservoir, rivers, ponds, and floodplain associated to river.

##### Remarks

Kiefer’s species was briefly described as *Diaptomus* (s.l.) *transitans* from organisms from the Argentine river environment of the South Neotropical. Illustrations were originally offered of the male fifth swimming leg, right antennule partially, and female last metasome segments plus urosome dorsally (detail for female fourth metasome segment laterally). It is intriguing to note that the taxon was never considered by Kiefer for *Notodiaptomus* (Kiefer, 1936; 1956), only being recombined in 1958 in Ringuelet when approaching the freshwater copepods of Argentina, and corroborating Kiefer (1929) through of: (1) male fifth right swimming leg exopod 2 with outer spine “longer” with one third of the length terminal claw; (2) male right antennule actual segment 13 with spiniform modified seta reaching to the middle-point of sequent segment; (3) male fifth right swimming leg exopod 2 with inner “tubercle“; (4) female fifth metasome segment with epimeral plates directed laterally.

Matsumura-Tundisi (1986) while studying calanoids from the freshwater aquatic systems of Brazil added new morphological characteristics to the species: (1) female fifth swimming legs endopod 2-segmented; and (2) female limit between fourth and fifth metasome segments fused dorsally, with lateral suture evidently. The attribute 2 highlighted by author represented a variation to the original description of the female of the species, which was recorded with fusion completely throughout the boundary between the segments. Additionally, a dorsal protuberance on female fourth metasome segment was presented in Matsumura-Tundisi (1986) on the limit between third and fourth metasome segments, and female antennule length surpassing to the caudal setae.

In the present effort we corroborate the observations recorded in Kiefer (1929) and Ringuelet (1958) for the taxon, except for the condition of the female limit between fourth and fifth metasome segments present as in Matsumura-Tundisi (1986). In contrast, we record here the positioning of the dorsal protuberance on female last metasome segments as originally described by Kiefer. The conditions highlighted for female fifth swimming legs endopod 2-segmented, and antennule length also were present in the specimens examined in this approach. It is important to highlight the male fifth right swimming leg exopod 1 with rounded process distally inserted on posterior surface and projecting over next segment, this was not originally present for the species, which still possessed the internal prominence in acute form on exopod 1.

Santos-Silva (2008) in reviewing the taxonomy and geography of Diaptomidae highlighted the confusing trajectory of *N. transitans.* Truly, Dussart (1984a) suggested the synonymization of the species to *D.* (s.l.) *mildredae* (Brehm 1956), and subsequently proposed its recombination to *Notodiaptomus* (*Caleodiaptomus*). These propositions were never accepted due to the lack of morphological foundations (Reid, 1987) and, subsequently, because it was discovered that the organisms treated as N. *coniferoides* were *N. simillimus* truly (Cicchino *et al*., 2001). Additionally, *N. transitans* and *N. simillimus* are species with evident morphological discrepancy between their organisms, for first species: (1) male fifth left swimming leg endopod 2-segmented; and (2) female fourth metasome segment with dorsal protuberance.

However, *N. transitans* does not present morphological divergences in relation to the characteristics of Wright (1935; 1936; 1937) for *nordestinus*, and Kiefer (1936) for the creation of *Notodiaptomus.* Divergences are evident among the attributes added in Kiefer (1956) for the amplification of the genus, being them for the species: (1) female fifth swimming legs endopod not unisegmented; (2) female fifth swimming legs endopod with spinules row non-oblique; (3) male fifth right swimming leg base without posterior protrusion; and (4) male fifth left swimming leg exopod 2 with spiniform seta not short, *i.e.,* surpassing to the distal-point of original segment. Among the characteristics identified as converging within *Notodiaptomus* from the type-species, *N. transitans* diverges into: (1) male fifth left swimming leg length reaching to first right exopod segment medially; (2) male fifth left swimming leg exopod 2 with digitiform process ornamented with denticles on anterior surface; (3) male fifth right swimming leg basis with inflated tumescence on proximal surface innerly; (4) male fifth right swimming leg exopod 1 with acute process innerly; (5) female fourth metasome segment with dorsal protuberance; and (6) female fifth swimming legs basis with outer seta reaching to middle-point of exopod 1.

### Taxonomic analysis

Through the present effort, the expansion of the morphological description of *Notodiaptomus* receives an amendment from the 2.627 new characters present in consideration of males and females of each species, resulting in 38 diagnostic characters to genus. Originally, and taking benefit of the effort *nordestinus* complex (Wright 1935), Kiefer (1936) presented *Notodiaptomus* to Science as a proposal for the division of *Diaptomus* (sensu lato) from South America. The initial grouping of 11 species was defined from 3 male characters (table 2, B), one unrecognizable due to subjectivity (1b), another with broad morphological plasticity among individuals of the same species (char 3b) and a third with a high frequency of sharing among the accepted taxa for the genus currently (char 2b), same present in relatable taxonomic groups, such as, for example, *Argyrodiaptomus* Brehm 1933 and *Rhacodiaptomus* Kiefer 1936, both Neotropical diaptomids.

In a later period, Kiefer (1956) extended the defining characteristics of *Notodiaptomus* through 11 additional morphological observations, counting on this occasion with females. The new effort on South American diaptomids considered 15 species for the genus considering 12 attributes supposedly (table 2, C). From this new morphological set, it is possible to recognize, among the characteristics most shared for males, the presence of “hyaline” spine at the rounded lobe on fifth right leg (3c = 100%); fifth right leg exopod 1 longer than broad, mostly (5c = 84.37%); fifth right leg exopod 2 with outer spine “generally inserted over the base of the terminal claw” (7c = 84.37%); and, fifth left leg exopod 2 with digitiform distal portion (8c = 84.37%). For females, both described characteristics are present with discrete frequency among the organisms of the species currently included in the genus (char 1c = 62.5% and char 2c = 12.5%). Among males, the observation of rounded and wide hyaline protuberance on the “internal apical surface of the caudal surface” of exopod 2 of the fifth right leg (fig. 21, #2388) seems to be evidence exclusive to the genus, although variable between organisms of species, actually (char 6c = 62.5%). However, no other offered attribute by Kiefer (1936; 1956) is unique to *Notodiaptomus*, significant in clearly defining the boundaries of the genus and differentiating it from other relatable taxonomic groupings.

This panorama seems to do justice to the observations of Wright (1937), openly opposed Kiefer’s proposal to divide the *Diaptomus* (s.l.) of South America. For the author, the insufficiency of data for the effort and the contingency of new organisms described and/or to be described at the time characterized an inappropriate decision at an inopportune moment. Santos-Silva *et al*. (2015) when approaching the *nordestinus* complex and, naturally, Wright’s arguments, ratified this perception. For the authors, although Kiefer’s contribution (1936) was the only system available at that time, it represented a “trigger” for several problems regarding taxonomic accuracy, precisely because of the lack of clear morphological definitions. Apparently, these arguments were confirmed over the timeline through the presentation of other proposals for Notodiaptomus, with inaccuracies that accentuated the morphological heterogeneity and inflicted on its taxonomic identity, without any accurate attempt to supply these complicating factors.

A first amplified description for *Notodiaptomus* was provided in Santos-Silva *et al*. (1999), when was designated *N. deitersi* as the type-species of the genus, formally. In this redescription, it is possible to consider 72 characters (table 2, D) in a diagnosis that redefined the basis for description and redescription of species to genus. Nevertheless, Santos-Silva *et al*. (2015), in a new act to resolve the taxonomic deficiencies of the group, returned to *nordestinus* complex as the “cradle” of the first *Notodiaptomus*, with the same observations. In the contribution it is possible to observe news evidence that refutes the set of differential characters in Santos-Silva *et al*. (1999) as a diagnosis in the strict sense (see definition in Mayr, 1969; Dubois, 2017). Indeed, in Santos-Silva *et al*. (1999), this is an expanded description with characters shared by other South American diaptomids. In the contribution on the *nordestinus* complex, although the improvements implemented are irrefutable, only a few comments allow inferring differential characteristics for the grouping.

In the analysis of the characteristics present in the studies mentioned previously, a morphological pattern was not clear among the organisms of all the species considered. Among the characteristics evaluated chronologically, only 12.5% (1) in Wright (1935), 0% in Kiefer (1936), 9% (1) in Kiefer (1956), and 54% (39) in Santos-Silva *et al*. (1999) are shared among all examined organisms but present in other diaptomineans described in the literature and confirmed in this study. On the other hand, an identity of *Notodiaptomus* seems appropriate when reconsidering characteristics predominantly shared among organisms of the current species to genus.

In Wright 1935 the observation of the fifth left leg (P5L) of males with narrowly segmented exopods and setules pad (4a) remains for all current *Notodiaptomus* organisms. However, it can also be found in organisms of the genus *Rhacodiaptomus* (*e.g.*, Santos-Silva & Robertson, 1993), *Aspinus* (*e.g.*, Brandorff, 1973), and *Argyrodiaptomus* (*e.g.*, Perbiche-Neves *et al*., 2011) and does not represent a defining attribute, unfortunately. Despite this, another male attribute with less frequency of sharing seem to contribute to gender identity: (3a) angular or rounded inner prominence on exopod 2 of the right fifth leg, treated here as an internal-distal ridge posterior on exopod 2 of the right fifth leg (P5R) (fig. 21, #2388). This attribute appears to be exclusive to the genus and is present in 65.6% (21) of the organisms in *Notodiaptomus*, excluding *N. henseni*, *N. iheringi*, *N. anisitsi*, *N. incompositus*, *N. santaremensis*, *N. caperatus*, *N. spinuliferus*, *N. simillimu*s, *N. cannarensis*, and *N. nelsoni*.

Another interesting characteristic appears in 28 evaluated females: (8a) posterior expansion on distal-internal rim of the basis of the fifth leg (P5) (fig. 29, #2602). Although it also appears in *Rhacodiaptomus* (*e.g.*, Kiefer, 1936), the character is in 87.5% of organisms assigned to *Notodiaptomus*, absent in *N. dentatus*, *N. maracaibensis*, and *N. orellanai*. We present that the only species with male and female excluded from the mentioned characteristics are *N. dentatus*, *N. cannarensis*, and *N. orellanai*, the current *Notodiaptomus* taxa that share less the characteristics suggested in Wright (1935). Among the species gathered by whole attributes set are *N. amazonicus*, *N. nordestinus*, *N. deitersi*, *N. cearensis*, *N. isabelae*, *N. jatobensis*, *N. conifer*, *N. carteri*, *N. transitans*, *N. santafesinus*, *N. paraensis*, *N. dubius*, *N. pseudodubius*. Of the originals of the *nordestinus* group, the analyzed of *N. henseni*, *N. iheringi* and *N. incompositus* are excluded, due to the absence of a distal-internal ridge on exopod 2 of the male P5R (3a), and *N. anisitsi*, which in addition to the previous condition, it does not have rounded “shoulder” on inner proximal rim of the basis of the male P5R (table 2, 1a), retracted in the present study as inner proximal tumescence on basis.

The presumption that the *nordestinus* complex was the basis for the foundation of *Notodiaptomus*, allows us to consider that Kiefer (1936) added three other characteristics to the genus. However, the presence of a conspicuous protuberance in “spine” form on segments 10, 11, 13, 15, and 16 of the male right antennule (A1R (2b) seems to us the only clear, objective and invariable approach, retracted in the present study as modified seta on segments 10, 11 and 13, and spinous process on segments 15 and 16. The attribute was shared for 31 (96.87%) males examined, anyone process was present on segment 15 in *N. santaremensis*. For the attribute that Kiefer (1936) considered as the most significant (1b), we judge that “nature” of the exopod 2 on male P5L resorts to variable understandings from its shape, size, and position until its possessed ornamental and sensorial elements, to depending on the observation plan, especially. This makes indication in Kiefer (1936) unfeasible to reproduce due to the verified subjectivity. This does not occur for character 3b (table 2), insecure due to unknown variability in the same population, as present in this effort throughout the species redescriptions, and discussed in Wright (1935) and Perbiche-Neves *et al*. (2015).

In amplification of *Notodiaptomus* in Kiefer (1956), a single characteristic appears exclusive, although variable among organisms of current species: (6c = 65.6%) internal-distal ridge posteriorly on exopod 2 of male P5R. The presence of this attribute was previously reported in Wright (1935) for *Nordestinus* and confirmed in this amplification, as was the absence of a spiniform process on segment 14 of male A1R (table 2, 10c = 2b). Although these more inclusive characteristics of Kiefer (1956) were not unprecedented, the addition of others less comprehensive to current species compose an interesting morphological set for *Notodiaptomus*: (1c) female P5 endopod 1-segmented, (4c) posterior protrusion on basis of male P5R, and (5c) exopod 1 longer than wide on male P5R.

The simultaneous observation of these attributes, isolated among groups close to the genus (*e.g.*, Walter, 1994; Santos-Silva & Robertson, 1993; Perbiche-Neves *et al*., 2011; Previattelli *et al*., 2015), evidence organisms in *Notodiaptomus* that converge these attributes. They are from the species: *N. amazonicus*, *N. nordestinus*, *N. deitersi*, *N. cearensis*, *N. isabelae*, *N. conifer*, *N. maracaibensis*, *N. santafesinus*, *N. kieferi*, and *N. dentatus*. Organisms of the species *N. henseni*, and *N. iheringi* do not have char 6c, and *N. jatobensis* does not have char 4c (table 2, C), but could belong to this grouping due to the morphological proximity verified. When considering the original proposal (Kiefer, 1936) discussed above, *N. amazonicus*, *N. nordestinus*, *N. deitersi*, *N. cearensis*, *N. isabelae*, *N. conifer*, and *N. santafesinus* remain.

In Santos-Silva *et al*. (1999), and Santos-Silva *et al*. (2015) the morphological set focused on the type species of the genus and members of the *nordestinus* complex, respectively, seems to have 18 defining attributes for the 32 species currently accepted in *Notodiaptomus* (table 2, D). Among the most inclusive, we highlight four characteristics shared for all organisms (table 2, D: 31d, 35d, 40d and 69d), among which one appears shared only with the genus *Argyrodiaptomus* (*e.g.*, *A. bergi* in Perbiche-Neves *et al*., 2011), which reflects a sign of proximity between the clusters: presence of vestigial seta on segments 2, 3 and 5 of the male A1D (4d). All other attributes are shared with two or more relatable taxonomic groups, isolated or grouped distinctly.

On the other hand, other attributes not shared among all *Notodiaptomus* appear unique to the genus and contribute to its diagnosis (table 2, D): antennules with elongated seta on segments 3, 7, 9 and 14, and blunt apex (5d = 90,60%); first segment with spinules row, and pore (8d = 81.25%); male cephalosome with incomplete dorsal suture (18d = 71.87%); male P5R basis with posterior protrusion (posterior medial growth) (43d = 68.75%); and male P5R exopod 2 with distal-internal ridge posteriorly (51d = 65.62%). Two of these characteristics are unprecedented observations and highlight the importance of morphological fragments under-discussed in the literature: antennae (8d) and cephalosome (18d). For male and female antennae, the spinules row is shared for all *Notodiaptomus* (except *N. cannarensis*) and *A. bergi*, *Rhacodiaptomus calamensis* (Wright 1927) and *Boeckella bergi*, but with the absence of the outer pore for *N. orellanai*, *N. brandorffi*, *N. paraensis*, *N. gibber*, and *N. transitans*. On the male cephalosome, the dorsal suture is absent in *N. kieferi*, *N. dentatus*, *N. paraensis* and *N. santaremensis* and complete in *N. carteri*, *N. brandorffi*, *N. pseudodubius*, N. inflatus, and *N. coniferoides*, as *in R. calamensis* and *Diaptomus castor*. In *Notodiaptomus*, this isolates those mentioned in the grouping.

It is interesting to note that two of these attributes corroborate observations in Wright (1935) during the founding of *nordestinus* (43d and 51d), which demonstrates its logical consistency over the timeline. It would be natural to consider that, over time, these arguments would be refuted as new members were included in the group and methodological advances were made. Perhaps this reinforces the thesis that really the division of diaptomids in that period and conditions was inopportune and would bring risks to the desired systematic accuracy, which somehow ended up being confirmed for *Notodiaptomus*.

However, for the highlighted attributes they would be grouped in *Notodiatomus*: *N. deitersi*, *N. cearensis*, *N. isabelae*, *N. amazonicus*, *N. nordestinus N. conifer*, *N. henseni*, *N. iheringi* and *N. maracaibensis*. For male P5R, the absence of oblique deep groove on basis (44d), and exopod 2 with distal-internal ridge posteriorly (51d) are conditions that isolate in the genus *N. santafesinus* and *N. jatobensis*, and *N. anisisti* and *N. nelsoni*, respectively. Of these attributes, the organisms that share less of them are *N. cannarensis*, *N. transitans*, and *N. caperatus*. Finally, through the evaluation of the historical series of attributes applied to define *Notodiaptomus*, the remaining organisms are: **(1)** *N. deitersi*, **(2)** *N. cearensis*, **(3)** *N. isabelae*, **(4)** *N. amazonicus*, **(5)** *N. nordestinus*, and **(6)** *N. conifer*.

In consideration of this, only 19% or 6 out of 31 taxa examined are grouped in the highlighted morphological pattern chronologically. It is obvious that other non-exclusive and variable characteristics among the current valid species support the diagnosis of the genus, but it is interesting to note how over timeline the inclusion of new species without clear delimitation has increased the heterogeneity in the morphological basis of *Notodiaptomus*. This reduced the number of common attributes and species that fully share them significantly, which constituted a complication for the support of all currently accepted taxa and the distinction of the genus within Diaptominae.

From the morphological set recognized in Santos-Silva *et al*. (1999) it is possible to recognize a common baseline for the genus and closely related groups in the literature (*e.g.*, Cicchino *et al*., 2001; Bradford-Grieve *et al*., 2010). The naturalness of this assessment occurs as other diagnostic characteristics of the genus compose a set of attributes that evidence its common morphological origin and, also, distinct as a true taxonomic group. This is exactly what we were able to verify when evaluating the previously discussed attributes (table 3), in which common, partially distinct, and distinctly stable characteristics make up the diagnosis identified over time for *Notodiaptomus* (*sensu* Blackwelder, 1967, p. 285).

### Accreting morphological deductions

For this research, we were able to observe 36 others unique characters for the genus (table 3), derived or not from the characteristics presented in the timeline. These allowed us to recognize an even more accurate morphological pattern for analyzed organisms, through them deducing shared morphological fractions mainly. 03 attributes could be admitted as amendments to *Notodiaptomus* diagnosis, added to those recognized historically.

The analysis of 11 morphological fractions between males and females (table 4) allowed us to sustain (5) antenna, and (7) male fifth swimming leg the most inclusive unique attributes in *Notodiaptomus* (table 3). Each of these fractions has a respective defining attribute, as follows: (5a = 81.2%) antenna coxa with inner seta at distal corner reaching to the endopod 1 (fig. 6, #980), and (7m = 65.6%) P5 right exopod 2 with curved ridge on distal posterior surface (fig. 21, #2388). The organisms gathered by this morphological group correspond to 19 species: *N. cearensis, N. coniferoides, N. isabelae, N. amazonicus, N. deitersi, N. maracaibensis, N. carteri, N. brandorffi, N. pseudodubius, N. dubius, N. nordestinus, N. paraensis, N. kieferi, N. inflatus, N. jatobensis, N. transitans, N. dentatus, N. gibber*, and *N. orellanai*.

**TABLE 4.** Increment of morphological attributes unique to organisms in *Notodiaptomus*. Analysis through morphological fraction and relative quantity of species grouped by attribute (% grouping), respectively.

| 1. ♂ PROSOME SEGMENTS (76 attributes examined) |  | % grouping |
| --- | --- | --- |
| 1a. | Limit between fourth and fifth metasome with double spinules row <sup>(N)</sup> | 15,6% |
| 1b. | Fifth metasome segment with double spinules row transversely | 6,25% |
| 1c. | Male right epimeral plate prominent, and not reaching half length of the genital segment | 28,1% |
| 1d. | Male left epimeral plate prominent, and reaching half length of the genital segment | 6,25% |
| 1e. | Male left epimeral plate with sensilla at the apex of projection and other medially | 15,6% |
| 2. ♂ UROSOME (44 attributes examined) |  | % grouping |
| 2a. | Fourth urosome segment with sub-conical blunt dorsal-lateral process with sensillae at apex | 3,1% |
| 3. ♂ RIGHT ANTENNULE (A1R) (416 attributes examined) |  | % grouping |
| 3a. | A1R actual segment 8 with conical seta length reaching to middle-point of sequential segment | 31,2% |
| 3b. | A1R actual segment 12 with conical seta length smaller than those of segment 8 <sup>(N)</sup> | 37,5% |
| 3c. | A1R actual segment 13 with modified seta reaching to distal-point of sequential segment <sup>(N)</sup> | 40,6% |

| **4. LEFT ANTENNULE (A1L) (421 attributes examined)** |  | **% grouping** |
| --- | --- | --- |
| **4a.** | A1L actual segment 13 e 15 without aesthetasc <sup>(N)</sup> <sup>(D)</sup> | 100% |
| **5. ANTENNA, and ORAL APPENDICES FEATURES (465 attributes examined)** |  | **% grouping** |
| **5a.** | Antenna coxa with inner seta at distal corner reaching to the endopod 1 <sup>(D)</sup> | 81,2% |
| **5b.** | Mandible coxal gnathobase with lobe not prominent | 6,25% |
| **5c.** | Mandible coxal gnathobase with acute projection between prominences | 6,25% |
| **6. FIRST TO FOURTH SWIMMING LEGS (659 attributes examined)** |  | **% grouping** |
| **6a.** | First swimming legs coxa with setules arrangement as a row anteriorly <sup>(N)</sup> | 9,3% |
| **7. ♂ FIFTH SWIMMING LEGS (P5) (338 attributes examined)** |  | **% grouping** |
| **7a.** | P5 left length reaching first right exopod segment medially or distally <sup>(N)</sup> | 40,6% |
| **7b.** | P5 left basis with posterior deep groove obliquely <sup>(D)</sup> | 53,1% |
| **7c.** | P5 left exopod 1 with posterior process, single or double, proximal and/or distal | 9,3% |
| **7d.** | P5 left exopod 1 with lobe in circular form on inner margin <sup>(N)</sup> | 34,4% |
| **7e.** | P5 left exopod 2 with non-denticulated distal process <sup>(N)</sup> | 40,6% |
| **7f.** | P5 right basis with inflated inner intumescence, bilobed or not, proximally or medially | 18,7% |
| **7g.** | P5 right basis with inner protuberance medially | 18,7% |
| **7h.** | P5 right basis with posterior groove not reaching the endopodal lobe <sup>(N)</sup> | 43,7% |
| **7i.** | P5 right basis without ornamentation on posterior groove <sup>(N)</sup> | 15,6% |
| **7j.** | P5 right exopod 1 with arched triangular process, outward or inward, sharp or blunt tip <sup>(N)</sup> | 21,8% |
| **7l.** | P5 right exopod 1 with posterior lamella in trapezoidal form | 9,3% |
| **7m.** | P5 right exopod 2 with curved ridge on distal posterior surface <sup>(N)</sup> <sup>(D)</sup> | 65,6% |
| **7n.** | P5 right exopod 2 with outer spine inserted medially <sup>(N)</sup> | 12,5% |
| **7o.** | P5 right exopod 2 with outer spine 6x lesser than the length of the exopod 2 | 12,5% |
| **8. ♀ METASOMAL SEGMENTS (35 attributes examined)** |  | **% grouping** |
| **8a.** | Fourth metasome segment with dorsal protuberance, digitiform or rounded, medially or distally | 34,4% |
| **8b.** | Limit between fourth and fifth metasome segments with double spinules row <sup>(N)</sup> | 12,5% |
| **8c.** | Right epimeral plate with inner spinules, ornamented as a row or patch <sup>(N)</sup> | 12,5% |
| **9. ♀ GENITAL DOUBLE-SOMITE, and UROSOME (60 attributes examined)** |  | **% grouping** |
| **9a.** | Genital double-somite with post-genital process ventrally <sup>(N)</sup> | 12,5% |
| **9b.** | Second urosome segment with ventral fusion to anal segment <sup>(N)</sup> | 15,6% |
| **10. ♀ ANTENNULES (A1) (25 attributes examined)** |  | **% grouping** |
| **10a.** | Actual segments 13, 15, 16, 17, and 18 without seta | 25% |
| **10b.** | Actual segment 14 without aesthetasc | 25% |
| **11. ♀ FIFTH SWIMMING LEGS (P5) (131 attributes examined)** |  | **% grouping** |
| **11a.** | P5 basis with outer seta longer 2x than original segment, reaching or not to exopod 1 distally <sup>(N)</sup> | 37,5% |
| **11b.** | P5 endopod with discontinuity cuticle on inner side <sup>(N)</sup> | 40,6% |

<sup>(N)</sup> Morphological attribute added to the *nordestinus* complex previously (in Santos-Silva *et al.*, 2015); <sup>(D)</sup> Diagnostic attributes identified to genus *Notodiaptomus*.

The fraction (11) fifth swimming legs had 131 attributes shared among the females examined. Of these, 2 attributes were exclusive to the genus, (11b = 40.6%) P5 endopod with discontinuity cuticle on inner side (fig. 29, #2617) represented the largest inclusion with 13 species. The addition of this new convergent criterion includes among the species mentioned for *Notodiaptomus* previously, *N. kieferi, N. jatobensis, N. pseudodubius, N. dubius, N. cearensis, N. deitersi, N. inflatus,* and *N. nordestinus*. In this specific morphological trait, *N. nelsoni, N. santaremensis, N. conifer,* and *N. henseni* are still included.

For the fraction (3) male right antennule were recognized three unique attributes involving 26 species, distinctly. The most inclusive attribute (3c = 40.6%) A1R actual segment 13 with modified seta reaching to distal-point of sequential segment (fig. 5, #388) gathers 13 taxa, of which are remnants of both morphological sets discussed so far: *N. maracaibensis, N. jatobensis, N. pseudodubius, N. dubius, and N. kieferi.* Additionally, *N. santaremensis, N. isabelae, N. nelsoni, N. santafesinus, N. incompositus, N. carteri, N. coniferoides,* and *N. simillimus* belong to the scope of this characteristic.

The consideration of this last exclusive attribute in *Notodiaptomus* seems inappropriate as a diagnostic characteristic of the genus, since it not only excludes the original inclusions of Kiefer (1936), but also deprecates the designated type species (Santos-Silva *et al*., 1999). Although this attribute of the male antennule (3c) and female P5 (11b) represent 2 of the 4 most inclusive attributes among those analyzed as unique to *Notodiaptomus*, their considerations as differential criteria seem ineffective for group identity, to the detriment of other closely related taxonomic groupings, such as *Argyrodiaptomus* and *Rhacodiaptomus*. Nevertheless, they are morphological traits that add value of intrageneric recognition and elements for a forward understanding of morphological derivations and their evolutionary paths.

Effectively, when considering *N. deitersi* as an accepted hypothesis for the organisms fixing the morphological pattern of the genus, 11 attributes are evidence of convergence of this type-species and its congeners (table 3., attributes 5a, 7m, 11b, 7e, 11a, 7d, 10a, 10b, 7j, and 7g). Among these, the most inclusive attribute is (5a) antenna coxa with inner seta at distal corner reaching to the endopod 1, which brings together 25 accepted species currently. This criterion seems to indicate an intimate relationship between these organisms and groups in *Notodiaptomus*, hypothetically: *N. deitersi, N. cearensis, N. conifer, N. isabelae, N. amazonicus, N. maracaibensis, N. nordestinus, N. carteri, N. brandorffi, N. pseudodubius, N. dubius, N. paraensis, N. simillimus, N. coniferoides, N. kieferi, N. inflatus, N. jatobensis, N. incompositus, N. henseni, N. iheringi, N. anisitsi, N. transitans, N. orellanai, N. nelsoni,* and *N. santaremensis*.

This morphological foundation evidences a set of species that reinforces the hypothesis complex *nordestinus* and its basis as the foundation of *Notodiaptomus*. Except *N. dahli* (which cannot be examined), all the initial considerations from the Wright (1935) and broadening (Wright, 1936; Wright, 1937) are included in the present proposal for convergent species. The original hypothesis of Kiefer (1936) and broadening (Kiefer, 1956) also are accepted in the present effort, in which the criterion would extend *Notodiaptomus* with ten other taxa – including *N. santaremensis*, and *N. carteri* not accepted by Wright (1937) historically.

In order to maintain the grouping criterion based on the type-species and consider the informative condition of the male geniculated antennule, we verified two variations in the ornamental pattern between AS (actual segment) 7 and 22 (Fig. 32, N, A, and B). The characterization of *N. deitersi* (Santos-Silva *et al*., 1999) was corroborated and converged for 10 taxa within the highlighted grouping previously, being (**N**): *N. cearensis, N. amazonicus, N. nordestinus, N. henseni, N. inflatus, N. brandorffi, N. paraensis, N. transitans,* and *N. orellanai.* The first variation of this pattern is characterized through of conical seta on AS12 smaller than that one on AS8, and conical seta on AS8 and spiniform modified seta on AS13 reaching the medial and distal portion of the sequent segments respectively. This variation is significant for 11 other taxa, these being (**A**): *N. anisitsi, N. iheringi, N. isabelae, N. santaremensis, N. kieferi, N. nelsoni, N. carteri, N. coniferoides, N. simillimus, N. dubius,* and *N. pseudubius*. Although the spiniform modified seta has a condition common to the taxa included in pattern A, the last three and two species have conical seta on AS8 and AS12 as in the type-species (N), respectively. In *N. iheringi*, and *N. anisitsi* the convergence to the variable pattern is the condition of the conical setae, being peculiar 2 setae on AS17 for the first species.

A second variation to the ornamental pattern of the male right antennule is based on the presence of 5 setae on AS22. Among the species gathered by this condition are (**B**): *N. incompositus, N. jatobensis, N. maracaibensis,* and *N. conifer*. The characterization for AS 8, 12, and 13 is convergent to that present in pattern A for the first three species, except for the conical seta on AS8 in *N. jatobensis*, and spiniform modified seta on AS13 in *N. maracaibensis* that have the same pattern for type-species of the genus (N). In *N. conifer* the spinous process surpasses the original segment (AS15) distally, evidencing a condition variable for male A1R and peculiar among *Notodiaptomus*. The other highlighted attributes are the same as the pattern N presented here.

Among all the distinct morphological criteria so far, 6 taxa stand out as the most divergent of the type-species of the genus. They are *N. cannarensis, N. caperatus, N. gibber, N. santafesinus, N. dentatus,* and *N. spinuliferus*, with organisms exclusive or occurring for the Southeast and South Neotropical region (except the first two from the Andes and central Atlantic Ocean’s Island). Based on the unique attributes recognized among the morphological fractions analyzed, the taxa with the lowest convergence of characteristics are *N. cannarensis* and *N*. *caperatus*, with 1 and 3 shared attributes respectively. Both species are included through (1e) male left epimeral plate with sensilla at the apex of projection and other medially (into 15.6% of the species examined), and *N. caperatus* by 2 characteristics other: 7b, and 7h (table 4).

### Phylogenetic analysis

The heuristic search retrieved 1 most parsimonious tree of length = 2540 steps, consistency index = 0.42, retention index = 0.41 and rescaled consistency index (RC) = 0.17. The evidence that the genus ***Notodiaptomus*** represents a monophyletic group was found (Fig. 33) to be supported by a synapomorphy composed of 80 of these characters (Table 5) among 627 informative.

**TABLE 5.** Morphological characters and states than compose to the synapomorphy of *Notodiaptomus* Kiefer 1936, consistency index (CI).

| CHARACTER (TRACED STATE) | CI |
| --- | --- |
| 1. Male fourth metasome segment – fusion or separation indicated by suture (**separated from the fifth metasome**, fused to fifth metasome) | 0.20 |
| 2. Male limit between fourth and fifth metasome segments – composition dorsal spinules row (on dorsal singly, **on dorsal doubly**) * | 0.25 |
| 3. Male fifth metasome segment – arrangement of ornamentation on posterior margin (continuous, **discontinuous**) * | 0.50 |
| 4. Male left epimeral plate – position of sensillae (at the apex of projection, **one at the apex of projection and other medially**) | 0.20 |
| 5. Male genital somite – location of sensillae on left side (at left rim, **at left lateral**) * | 1.00 |
| 6. Male third urosome segment – arrangement of ornamentation (as a row, **as a patch**) | 1.00 |
| 7. Male third urosome segment – number of row (single throughout dorsal surface, **double throughout dorsal surface**) | 1.00 |
| 8. Male fourth urosome segment – arrangement of ornamentation (as a row, **as a patch**) | 1.00 |
| 9. Male fourth urosome segment – number of row (single throughout dorsal surface, **double throughout dorsal surface**) | 1.00 |
| 10. Male right antennule – relative length (**extending beyond caudal rami**, not extending beyond caudal rami) * | 0.09 |
| 11. Male right antennule actual segment 3 – apex of the seta (**blunt apex**, sharp apex) * | 0.33 |
| 12. Male right antennule actual segment 3 – presence of aesthetasc (**without aesthetasc**, with aesthetasc) | 0.25 |
| 13. Male right antennule actual segment 4 – length of seta (none larger than segment, **one larger than segment**) | 0.33 |
| 14. Male right antennule actual segment 6 – range of the seta (beyond three sequential segments, **not beyond three sequential segments**) * | 1.00 |
| 15. Male right antennule actual segment 7 – apex of the seta (**blunt apex**, sharp apex) * | 0.33 |
| 16. Male right antennule actual segment 9 – apex of the seta (**blunt apex**, sharp apex) * | 0.33 |
| 17. Male right antennule actual segment 10 – modified seta range (**beyond of the sequential segment**, not beyond of the next segment) | 0.33 |
| 18. Male right antennule actual segment 11 – length of seta (none larger than segment, **one larger than segment**) | 0.50 |
| 19. Male right antennule actual segment 12 – length of seta (none larger than segment, **one larger than segment**) | 1.00 |
| 20. Male right antennule actual segment 14 – apex of the seta (**blunt apex**, sharp apex) | 0.50 |
| 21. Male right antennule actual segment 15 – size comparison of the setae (of equal size, **of unequal size**) * | 1.00 |
| 22. Male right antennule actual segment 16 – composition of seta (straight, **plumose**) * | 0.33 |
| 23. Male right antennule actual segment 17 – range of the seta (surpassing to distal margin, **surpassing distal margin**) * | 1.00 |
| 24. Male right antennule actual segment 18 – number of setae (one element, **two elements**) | 0.33 |
| 25. Male right antennule actual segment 19 – composition of the setae (straight, **plumose**) * | 1.00 |
| 26. Male right antennule actual segment 20 – reach of spinous process (reaching beyond of distal-point segment 21, **not reaching beyond**) | 0.20 |
| 27. Left antennule – relative length (**extending beyond caudal setae**, not extending beyond caudal setae) * | 0.09 |
| 28. Left antennule actual segment 2 – presence of vestigial seta (without vestigial seta, **with vestigial seta**) | 0.33 |
| 29. Left antennule actual segment 3 – presence of vestigial seta (without vestigial seta, **with vestigial seta**) * | 1.00 |
| 30. Left antennule actual segment 5 - presence of vestigial seta (without vestigial seta, **with vestigial seta**) | 0.33 |
| 31. Left antennule actual segment 10 – range of the seta (surpassing to distal margin, **surpassing distal margin**) * | 1.00 |
| 32. Left antennule actual segment 11 – size comparison of the setae (of equal size, **of unequal size**) * | 1.00 |
| 33. Left antennule actual segment 12 – length of seta (**one larger than segment**, none larger than segment) | 0.50 |
| 34. Left antennule actual segment 13 – range of the seta (surpassing to distal margin, **surpassing distal margin**) * | 1.00 |
| 35. Left antennule actual segment 16 – composition of seta (straight, **plumose**) | 0.50 |
| 36. Left antennule actual segment 21 – composition of seta (straight, **plumose**) * | 1.00 |
| 37. Left antennule actual segment 22 – composition of seta (straight, **plumose**) * | 1.00 |
| 38. Left antennule actual segment 23 – number of the setae (one element, **two elements**) | 0.33 |
| 39. Left antennule actual segment 24 – composition of seta (straight, **plumose**) | 0.50 |
| 40. Left antennule actual segment 25 – composition of the seta (straight, **plumose**) | 0.50 |
| 41. Antenna coxa – reach of the seta (reaching to the middle base, **reaching to the endopod 1**) * | 0.16 |
| 42. Antenna endopod actual segment 2 – condition of the discontinuity present (developed as a suture, **not developed as a suture**) * | 0.50 |
| 43. Antenna endopod actual segment 2 – arrangement of spinules (as a row, **as a patch**) | 0.50 |
| 44. Antenna endopod actual segment 3 – number of setae on outer lobe (outer lobe bearing 7 setae, **outer lobe bearing 8 setae**) * | 1.00 |
| 45. Maxilla coxa – size of coxal endite (third elongated (larger width length ratio 2:1), **third not elongated (lesser width length ratio 2:1)**) * | 1.00 |
| 46. Maxilla coxa – composition of the seta on the third coxal endite (plumose, **straight**) * | 1.00 |
| 47. Maxilla endopod 3 – composition of the seta (plumose, **straight**) * | 1.00 |
| 48. First swimming legs exopod 2 – location of setules (on inner margin, **on outer margin**) | 1.00 |
| 49. First swimming legs exopod 3 – presence of spinules (with spinules, **without spinules**) | 0.50 |
| 50. Third swimming legs endopod 2 – arrangement of spinules (as a row, **as a patch**) * | 1.00 |
| 51. Fourth swimming legs exopod 3 – type of ornamentation on non-serrated side (of spinules, **of setules**) * | 1.00 |
| 52. Male fifth left swimming leg basis – quantify of protuberances (single, **double**) * | 1.00 |
| 53. Male fifth left swimming leg exopod 2 – form of the inner side (**disform inner side**, uniform inner side) | 0.33 |
| 54. Male fifth left swimming leg exopod 2 – form of the outer side (**with rectilinear outer side**, with convex outer side) | 0.11 |
| 55. Male fifth left swimming leg exopod 2 – condition of the setulose pad (rounded prominently, **not rounded prominently**) | 0.11 |
| 56. Male fifth left swimming leg exopod 2 – position of the prominently pad (proximally, **medially**) | 0.16 |
| 57. Male fifth left swimming leg exopod 2 – reach seta (**not surpassing the distal-point segment**, surpassing the distal-point segment) * <sup>W, K</sup> | 0.12 |
| 58. Male fifth right swimming leg coxa – form of the inner side (concave inner side, **convex inner side**) | 0.25 |
| 59. Male fifth right swimming leg basis – form of the additional internal process present (digitiform, **triangular**) * | 1.00 |
| 60. Male fifth right swimming leg basis – reach of groove (reaching the endopodal lobe, **not reaching the endopodal lobe**) * | 0.20 |
| 61. Male fifth right swimming leg basis – posterior protrusion (posterior protrusion absent, **posterior protrusion present**) * <sup>K</sup> | 0.11 |
| 62. Male fifth right swimming leg exopod 2 – size proportion of the segment longer than broad (**nearly 2 times**, nearly 2.5 times) * | 0.10 |
| 63. Male fifth right swimming legs exopod 2 – size comparison between sides (**equal size between both sides**, unequal size) | 0.16 |
| 64. Male fifth right swimming legs exopod 2 – curved ridge (absent, **curved ridge on distal posterior surface present**) * <sup>W, K</sup> | 0.09 |
| 65. Male fifth right swimming legs exopod 2 – form of the tip of the outer spine (**without apparent curve**, with apparent curve) | 0.11 |
| 66. Female limit between fourth and fifth metasome segments – spinule row arrangement (entirely over limit, **partially over limit**) * | 0.20 |
| 67. Female right antennule actual segment 13 – presence of seta (with seta, **without seta**) * | 1.00 |
| 68. Female right antennule actual segment 13 – presence of aesthetasc (with aesthetasc, **without aesthetasc**) * | 1.00 |
| 69. Female right antennule actual segment 14 – presence of seta (with seta, **without seta**) * | 1.00 |
| 70. Female right antennule actual segment 14 – presence of aesthetasc (with aesthetasc, **without aesthetasc**) * | 1.00 |
| 71. Female right antennule actual segment 15 – presence of seta (with seta, **without seta**) * | 1.00 |
| 72. Female right antennule actual segment 16 – presence of seta (with seta, **without seta**) * | 1.00 |
| 73. Female right antennule actual segment 17 – presence of seta (with seta, **without seta**) * | 1.00 |
| 74. Female right antennule actual segment 18 – presence of seta (with seta, **without seta**) * | 1.00 |
| 75. Female fifth swimming legs coxa – projection of the process (not projecting over basal segment, **projecting over basal segment**) | 0.25 |
| 76. Female fifth swimming legs coxa – relative length of the sensilla (not longer 2x than basal insertion, **longer 2x than basal insertion**) | 0.20 |
| 77. Female fifth swimming legs endopod – fusion between ancestral segments 1 and 2 (segments 1 and 2 separated, **segments 1 and 2 fused**) | 0.11 |
| 78. Female fifth swimming legs endopod – insertion of ornamentation (terminally, **sub-terminally**) | 0.33 |
| 79. Female fifth swimming legs endopod – location of setae 1 (on anterior surface, **posterior surface**) | 0.50 |
| 80. Female fifth swimming legs exopod 2 – relative size of the spine to next segment (**equal size or larger than next segment**, smaller than) | 0.09 |

\* Exclusive condition in *Notodiaptomus*; <sup>W</sup> character condition originally noted in Wright (1935); <sup>K</sup> character condition originally noted in Kiefer (1956).

The phylogeny of *Notodiaptomus* allows us to estimate the condition of some uncertain or previously unknown character states, their distribution hypotheses, and the confirmation of the homoplasy of many characters (homoplasy index (HI) = 0.5709) present in the characterized hypothetical common and exclusive ancestor (table 5). The main trends in character state changes in the highlighted synapomorphy were traced to male prosome (cephalosome and metasome), male urosome, male right antennule, left antennule, antennae, maxilla, first, third, and fourth swimmings legs, male fifth swimmings legs, female body (habitus, metasome, genital double-somite), and female fifth swimming legs. Among these characters, 43 are unique conditions for *Notodiaptomus* (table 5, indication *) and represent autopomorphies at the level most comprehensive to the outer group.

The external group was represented by 6 members in all. The result indicates that *Diaptomus* (s.l.) *susanae* cannot be maintained in its original taxonomic condition, since the taxon is nested within *Notodiaptomus* (Fig. 33, blue line). In the present representation*, D. susanae* belongs to the cladogenic composition innermost to *N. orellanai*, *N. cannarensis,* and *N. maracaibensis*, and distinguished from node d mainly through the unambiguous transitions of the male right antennule actual segment 17 with aestetasc (Char 466: 1► 2, IC = 1.0), male fifth right swimming leg basis with outer seta longer 2x than original segment (Char 2317: 1►2, CI = 1.0), female urosome 2-segmented (Char 2483: 2►1, CI = 0.66), and female genital double-somite symmetrical in dorsal view (Char 2484: 1►2, CI = 1.0). The phylogenetic positioning reconstructed for the male and female of this species corroborates the indication of Paggi (1976) about its morphological relationship with *Notodiaptomus*, and may substantiate the taxonomic transfer from a nomenclatural act in due course.

*Argyrodiaptomus bergi* was the closest paraphyletic taxon to the common and exclusive ancestor reconstructed for *Notodiaptomus*, followed by *Rhacodiaptomus calamensis* externally. Both genera represent taxonomic attempts to group South American diaptomids, the first in Brehm (1933) with the current second largest Neotropical richness, the second founded in Kiefer (1936) along with *Notodiaptomus* and a supposedly distinct group. In the phylogeny presented, *A. bergi* is unequivocally distinguished from the male fifth left swimming legs endopod segment 1 and 2 with incomplete suture line (Char 2171: 1►2, CI = 1.0), and female fifth swimming legs exopod 2 with marginal extension (Char 2658: 1►2, CI = 1.0). *R. calamensis* as the type species of an Amazonian genus was phylogenetically distinct to the cladogram of *Argyrodiaptomus* and *Notodiaptomus* especially through male fourth and fifth fused partially on dorsal surface (Char 40: 1►2, CI = 1.0), and female fifth swimming legs exopod 2 without lateral spine present (Char 2661: 2►1, CI = 1.0).

*Pseudodiaptomus* sp., and *Boeckella bergi* were other members of the external group and paraphyletic to the diaptomids previously mentioned in the phylogeny achieved. Both taxa are representative of Pseudodiaptomidae Sars 1902 and Centropagidae Giesbrecht 1893, and indicated in the phylogeny of Bradford-Grieve *et al*. (2010) as a sister group of Diaptomidae. The evidence on the phylogenetic relationship between the families corroborates the morphological relationship indicated in Sars (1902; 1903) when including them among the Heterarthrandria (Giesbrecht, 1893), that is, the calanoids with asymmetry of the male antennulae. In the present phylogeny obtained, the clade composed of *Pseudodiaptomus* sp., and *B. bergi* is distinguished by the male right antennule actual segment 10 without arrow (Char 312: 2►1, IC = 1.0), male right antennule actual segment 12 without aesthetasc (Char 371: 2►1, IC = 1.0), male fifth right swimming leg exopod 1 with outer spine (Char 2363: 1►2, IC = 1.0), and female fifth swimming legs exopod 3 with spine (Char 2674: 1►2, CI = 1.0).

Finally, *D. castor* as the type of *Diaptomus* (sensu stricto) was the most basal taxon traced in the phylogeny presented here. Although it is common knowledge that the previous understandings of the European *Diaptomus* of Westwood (1836) were references for the first efforts on the Neotropical calanoids, it is curious to note that the cladogenic positioning of the species indicates an evolutionary hypothesis absolutely distinct from the *Notodiaptomus* and other South American diaptomids considered. For the recognition of the taxon as a member of the external group of this phylogeny were unambiguous phylogenetic hypotheses, especially the male right antennule: actual segment 1 with two aesthetascs (Char 186: 1►2, IC = 1.0), actual segment 11 with modified seta presenting bifid apex (Char 346: 1►2, IC = 1.0), actual segment 16 without seta (Char 429: 1►2, IC = 1.0), current segment 18 with two modified setae (Char 346: 1►2, IC = 1.0), actual segment 19 with one seta (Char 489: 2►1, IC = 1.0), additionally the male fifth left swimming leg exopod 1 with presence seta (Char 2222: 1►2, IC = 1.0).

The evidence of the distinct cladogenic positioning between *D. susanae* and *Diaptomus castor* exposes several other hypotheses of *Diaptomus* from South America, still provisionally maintained. Perhaps the results presented here will reinforce future investigations on the organisms of this sensu lato grouping, in order to remedy their systematic relationships and clarify obscure morphological limits in other taxonomic groups of South America. Undoubtedly, it seems prudent for the systematics of Neotropical diaptomids that new taxonomies be presented exclusively from the sensu stricto of existing groups or new nomenclatural acts evidenced.

#### Synapomorphies exclusive to *Notodiaptomus*

It is relevant to consider the remarkable mix of clades and degrees of relationships within the monophyletic group *Notodiaptomus.* The general arrangements of the phylogenetic hypotheses within the grouping preclude, at this time, the formal naming of the phylogenetic ancestors represented in each internal node within the monophyletic condition obtained. In addition to *Notodiaptomus*, we recognize 11 additional clades (supplementary material – S3) indicated by the letters a through k (fig. 33), among which the synapomorphic character conditions unique to monophyletic *Notodiaptomus* occur and will be discussed below.

##### Attributes for male prosome and urosome

Through the prosomal region of the male *Notodiaptomus* were traced 2 phylogenetic hypotheses unique among the coogeners studied. O male limit between fourth and fifth metasome segments with spinules row on dorsal doubly (table 5 = #2; S3 = Char 44: 1►2) is an apomorphy to the condition present in *Pseudodiaptomus* sp., and convergent evidence in node i for *N. iheringi* and *N. spinuliferus*, node k for *N. anisitsi* and *N. nelsoni*, and in *N. orellanai*. The condition of the male fifth metasome segment with arrangement of ornamentation discontinuous on posterior margin (table 5 = #3; S3 = Char 56: 1►2) can be equally interpreted and is present in *N. iheringi* (node i) and *N. nelson* (node k). Although unique attributes, both phylogenetic hypotheses do not seem to consistently contribute to the topology of the represented cladogram, since they are confined to the terminals mentioned exclusively.

The male urosome of *Notodiaptomus* possessed the genital somite with sensilla at the left lateral (table 5 = #5; S3 = Char 88: 1►2) as the only morphological feature belonging to the common ancestor and exclusive recovered. Although the applicability of the attribute is dependent on the presence of the sensory element, the reconstructed characteristic contributes to consistency in the topology achieved. Undoubtedly, the phylogenetic hypothesis presented here depends on the polarization of the character states provided by the organisms associated with the defined members of the external group. New observations and bases of polarization can contribute to a better resolution of the relationships and corroboration of this as an inherent attribute of *Notodiaptomus*.

It is also interesting to note that although characters 6, 7, 8, and 9 (table 5, CI = 1.0) present a high consistency index for the topology obtained, these are plesiomorphies shared with *Pseudodiaptomus* sp. and *Boeckella bergi* and represent simplesiomorphies for *N. orellanai* (basal taxon), and *N. incompositus* (node k). The position of *N. orellanai* as the most primitive taxon in the genus suggests that the ancestral *Notodiaptomus* probably had ornamentations in the dorsal urosomal segments and lost them in its evolutionary trajectory. The sharing of these attributes with the representatives of Pseudodiaptomidae and Centropagidae seems to reinforce the relationship with Diaptomidae evidenced in the phylogeny of Bradford-Grieve *et al*. (2010).

##### Attributes for male antennules

Since Sars (1903) the antenulas have morphological attributes used as criteria for groupings within the calanoids. In the reconstruction of the common and exclusive ancestor of *Notodiaptomus* more than one-third of the characters were traced from these appendages for males, 17 chars for the right antennule, and 14 chars for left antennule. In right antennule, nine characters are unique evolutionary hypotheses, among which are male right antennule extending beyond caudal rami (table 5 = #10; S3 = Char 140: 2►1) is ambiguous and highly homoplastic (HI = 0.9) between node j (N. *deitersi, N.* anisitsi*, N. henseni*, and *N. santafesinus*), node g (*N. caperatus*, *N. gibber*, *N. kieferi*), and node i (*N. dentatus, N. iheringi*, *N. jatobensis*, and *N. nordestinus*). The occurrence of this condition in the most basal nodes within *Notodiaptomus* (*i.e*., node b, and *N. orellanai*) suggests that, although achieved by different pathways, the character probably represents an important evolutionary signal within the genus. Forwards investigations should characterize new evidence regarding this attribute in Diaptomidae.

Other attributes first traced in the male right antennule are for the current segments 3, 7, and 9 with blunt apex of the seta (table 5 = #11, 15, and 16; S3 = Chars 209, 270, and 303: 2►1, respectively). These are apomorphies present in all *Notodiaptomus* examined, except for the exclusive ancestor of *N. gibber* and *N. transitans* with present reversals. Another unique feature reconstructed for the hypothetical ancestor of *Notodiaptomus* is for the male right antennule actual segment 6 with range of the seta not beyond three sequential segments (table 5 = #14; S3 = Char 253: 1►2). The prolongation of this ornamental structure represents a loss among the *Notodiaptomus* and is confined to *N. dentatus* and *N. spinuliferus* (inserted in the node g). Among these observations, the condition for the apex of the current septal elements of segments 3, 7, and 9 seems to be the most significant for the *Notodiaptomus* hypothesis and perhaps indicates a mechanoreceptor evolutionary relationship (Garm & Watling, 2013). The recoveries of these morphological conditions corroborate the observations of Santos-Silva *et al*. (1999) and Santos-Silva *et al*. (2015), who present the same characteristics for the designation of the type of the genus and redescription of the *nordestinus* complex, respectively.

In addition, other features of relative size, range, and structural compositions for the male right antennule make up the unique hypotheses of *Notodiaptomus.* The actual segment 15 with setae of unequal size (table 5 = #21; S3 = Char 413: 1►2), and actual segment 19 with plumose seta (table 5 = #25; S3 = Char 491: 1►2) are consistent apomorphies for the phylogenetic hypothesis of the common ancestor and exclusive of the genus. The actual segment 16 with plumose seta (table 5 = #22; S3 = Char 432: 1►2) is a condition that undergoes reversal in the exclusive ancestor of *N. dentatus* and *N. spinuliferus* (node i), and in the exclusive ancestor of *N. coniferoides*, and *N. simillimus* (node e). The actual segment 17 with seta surpassing distal margin (table 5 = #23; S3 = Char 452: 1►2) is a condition lost within the cladogram of *Notodiaptomus* and confined to *N. coniferoides*, and *N. simillimus* (node e). The tracing of this last attribute seems to represent inconsistency in the reconstruction of the ancestor of *Notodiaptomus*. Although the high value of the consistency index belies this statement, its value is clearly influenced by a strictly convergent homoplasy in the topology presented.

For the male left antennule, seven other features were traced to the common and exclusive ancestor of *Notodiaptomus*. Among the most significant, the left antennule extending beyond caudal setae (table 5 = #27; S3 = Char 555: 2►1) is an unambiguous transition from the unique ancestor of *N. nelsoni*, *N. santafesinus*, and *N. anisitsi* to the unique ancestor of *N. santafesinus*, and *N. anisitsi*, with reversal of node j to node k and convergences in *N. amazonicus, N. cearensis*, *N. spinuliferus*, *N. transitans*, and *N. cannarensis.* The condition for actual segment 3 with vestigial seta (table 5 = #29; S3 = Char 623: 1►2) is unambiguous and reconstructed for the exclusive ancestor of *Notodiaptomus* consistently. Santos-Silva *et al*. (2015) mentions this characteristic as important for the morphological delimitation of the genus, which is corroborated in our results as an apomorphic condition identified with wide distribution along the topology, high retention index, and rescaled consistency (RC) (RI = 1.0; RC = 1.0).

New consistencies are identified for char 34 (table 5; S3 = Char 772: 1►2) and char 36 (table 5; S3 = Char 909: 1►2), which represent conditions derived from the state in all members of the outer group, except *D. castor.* The actual segment 10 with seta length surpassing distal margin (table 5 = #31; S3 = Char 720: 1►2), is inconsistent between the internal hypotheses of *Notodiaptomus* as it converges exclusively on *N. coniferoides*, and *N. simillimus* (node e). Different from this is the actual segment 22 with plumose seta (table 5 = #37; S3 = Char 936: 1►2), with reversal only for the mentioned members of clade e.

Finally, the character 32 (table 5) for the actual segment 11 with setae of unequal size (table 5; S3 = Char 736: 1►2) represents an extension to the contributions in Santos-Silva *et al*. (2015), who when analyzing the morphology of the organisms of eleven *Notodiaptomus*, indicated the loss of the proximal seta in this segment as a possible apomorphy for the genus. Although the character traced to the exclusive ancestor of *Notodiaptomus* is not exactly about the presence and absence of the element, the condition relative to comparative size is only applied by the presence of two setae in this segment. Truly, the presence of two setae in segment 11 traced in *Pseudodiaptomus* sp. and *Boeckella bergi* confirms the indication of Santos-Silva *et al*. (2015), being a simplesiomorphy in *N. amazonicus*, *N. cearensis,* and *N. spinuliferus* (node i), *N. anisitsi* (node k), and *N. gibber* (node g). On the other hand, the condition traced to the ancestor of *Notodiaptomus* (#32) are convergences to the same terminals mentioned.

##### Attributes for antennae, oral appendages, and swimmings legs

The attributes related to antennae, oral appendages, and swimming legs 1 to 4 are commonly overlooked for groupings among diaptomids. Historically, in any attempt to divide these organisms in South America, these morphological factors have been highlighted as defining characteristics. In *Notodiaptomus* this does not change. However, our results show 11 phylogenetic hypotheses traced for the reconstruction of the common and exclusive ancestor of the genus. Among them, 3 plesiomorphies and 8 exclusive apomorphies belong to the characterized synapomorphy.

Among them, the exclusive condition for antenna coxa with arrow reaching to the endopod 1 (table 5 = #41; S3 = Char 980: 1►2) is convergent to all members of the *nordestinus* complex (basis for the foundation of *Notodiaptomus* already discussed) and to the other terminals, except for the reversal present in the exclusive ancestor of *N. dentatus*, and *N. spinuliferus* (node i), *N. santafesinus*, *N. gibber, N. caperatus*, and *N. cannarensis*. Although highly homoplastic (HI = 0.8), the character has a retention index (RI = 0.5) above the average obtained for the set of hypotheses represented (Fig. 33), which evidences a number of significant and important emergencies for the definition of the common and exclusive ancestor of *Notodiaptomus.* Santos-Silva *et al*. (2015), when reviewing the morphology of the organisms of eleven species of the genus, including the type, recorded the same morphological condition. The analysis performed here reinforces the consistency of this evidence for *Notodiaptomus* definitively.

In a similar analysis, another relevant synapomorphic character can be identified for the antenna endopod actual segment 2 with discontinuity on outer cuticle not developed as a suture (table 5 = #42; S3 = Char 1006: 1►2), morphological condition derived from the condition of *D. castor, B. bergi*, and *A. bergi*, and simpleiomorphic in *N. gibber, N. kieferi*, and *N. transitans* (node f ► node g). Cuticular discontinuity on the antenna endopod 2 was a condition present for some organisms during the morphological examinations performed for the present study. From this attribute we present the variation for the presence or absence of transverse suture as a hypothesis for the development of the previously present cuticular discontinuity. The absence of this condition is characterized as a convergent apomorphy in *Notodiaptomus*, which suggests that the common and exclusive ancestor of organisms associated with congeners species possessed this attribute.

Inverse situation is identified for synapomorphic characters 44, 45, 46, and 47. The antenna endopod actual segment 3 with outer lobe bearing 8 setae (table 5 = #44; S3 = Char 1016: 1►2) represents an apomorphy convergent to *N. brandorffi,* and *N. paraensis* (node g) exclusively, which justifies the high retention rate for the character (RI = 1.0). The characters referring to maxilla (table 5 = #45, #46, and #47; S3 = Chars 1276, 1279, and 1308: 1►2) are traced autapomorphies of *N. cannarensis* (node a). Although the consistencies verified for the set of phylogenetic hypotheses of *Notodiaptomus* (CI = 1.0), these attributes presented do not represent informative characters for the reconstruction of the common and exclusive ancestor of the genus.

For swimming legs, two evolutionary hypotheses were traced for *Notodiaptomus*, third swimming legs endopod 2 ornamented with spinules as a patch posteriorly (table 5 = #50; S3 = Char 1845: 1►2), and fourth swimming legs exopod 3 longer spine with non-serrated side ornamented by setules (table 5 = #51; S3 = Char 2096: 1►2). These two conditions represent autapomorphy for *N. brandorffi*, and contingency for *N. dentatus* and *N. spinuliferus* (node i), respectively. Bradford-Grieve *et al*. (2010) present other hypotheses for both swimming legs in the common and exclusive ancestor of Calanoida, and Santos-Silva *et al*. (1999) for the description of the type species of the genus and redescription of the *nordestinus* complex (Santos-Silva *et al*., 2015) did not record observation for these attributes. The confinement evidenced for both evolutionary hypotheses suggests inconsistency about their consideration for the rescue of the common and exclusive ancestor of *Notodiaptomus*.

##### Attributes for male fifth swimming legs

Historically, male fifth swimming legs are criteria adopted for inter- and intraspecific characterization within the systematics of Diaptomidae (*e.g.*, Brehm, 1933; Wrigth, 1935; Kiefer, 1936). The results obtained through the phylogenetic analysis presented indicate 7 unique hypotheses traced for these male factors, 2 from the fifth left swimming leg, and 5 from the fifth right swimming leg. Among them, 1 exclusive hypothesis for the fith left swimming leg, and 2 exclusive hypotheses for the right swimming leg are remnants of the original proposals of Kiefer (1956) for the expansion of *Notodiaptomus*, and Wright (1935) for the *nordestinus* complex.

For the fifth left swimming leg, the basis with double protuberances innerly (table 5 = #21; S3 = Char 2154: 1►2) represents an autapomorphic condition for *N. henseni*, and apparently inconsistent in rescuing the ancestral features of *Notodiaptomus.* The fifth left swimming leg exopod 2 with spiniform seta not surpassing the distal-point segment (table 5 = #57; S3 = Char 2247: 2►1) has several emergencies in different evolutionary directions that characterize the present elevated homoplasy (HI = 0.8). Specifically, reversals are present from the node f to the node g, for the exclusive ancestor *N. coniferoides* and *N. simillimus*, and exclusive ancestor of *N. cearensis* and *N. iheringi*. It is possible to identify simplesiomorphic conditions for *N. santafesinus*, *N. incompositus*, and *N. pseudodubius*.

The condition of the spiniform seta not exceeding exopod 2 distally is here interpreted as the morphological attribute originally adopted for *nordestinus* complex of Wright (1935; 1936) (table 2, 5a). Through our results we corroborate this Wright decision, except for the simplesiomorphic condition present for *N. cearensis, N. iheringi*, and *N. incompositus.* Kiefer (1956) during the expansion of *Notodiaptomus* identified the same attribute as a criterion for the assembly of organisms associated with the species of the genus (table 2, 9c) and can have here his position reaffirmed, except for the same species mentioned earlier. However, although highly homoplastic, phylogenetic retention for character (RI = 0.5) is above the average present for the set of hypotheses (Fig. 33) and the phylogenetic analysis presented here suggests that this trait belongs to the common and exclusive ancestor of *Notodiaptomus*.

For a fifth right swimming leg, the basis with additional internal process in triangular form (table 5 = #59; S3 = Char 2305: 1►2) is state derived from the representative of Centropagidae defined for polarization. However, the elevated consistency index does not represent a robustness of the hypothesis for the common and exclusive ancestor of *Notodiaptomus*, since these are convergences confined to *N. gibber*, and *N. paraensis.* The disappearance of this character after the common and exclusive ancestor of *Notodiaptomus* and its confinement among the evolutionary hypotheses represented does not allow us to accurately trace its evolutionary history among the relationships presented. It is possible that difficulties in assigning homologies based on varying sizes, position, and additional observations have represented noise for accurate interpretations. However, it is expected that this character may be useful in future analyses, once new morphological groups and other specimens are examined and homologies better understood.

The fifth right swimming leg basis hypothesis with posterior groove not reaching the endopodal lobe (table 5 = #60; S3 = Char 2310: 1►2) represents an uninformative synapomorphy (IR = 0.2) for the actual characterization of the common and exclusive ancestor of *Notodiaptomus*. The high homoplasy index (HI = 0.8) reinforces the ambiguity of the character state between the evolutionary relationships represented, with reversals present from the node d to node e (*N. coniferoides*, *N. simillimus*, and *N. inflatus*), in the exclusive ancestor of *N. deitersi*, and *N. henseni*, and simplesiomorphic to *N. nordestinus*, and *N. iheringi*. The primitive condition of this character was abducted from *A. bergi* as a member of the outgroup and is present for four original members of the *nordestinus* complex, and *Notodiaptomus*: *N. deitersi*, *N. henseni*, *N. iheringi*, and *N. nordestinus*. Although an obvious inconsistency for this phylogenetic analysis, the state of the character will be useful for future investigations about intra- and inter-genus evolutionary relationships.

The character fifth right swimming leg basis with posterior protrusion (table 5 = #61; S3 = Char 2318: 1►2) is an ambiguous condition between the evolutionary relationships represented. The present high homoplasy index (HI = 0.88) confirms the number of emergencies along the set of hypotheses represented, with unequivocal reversals to the exclusive ancestor of *N. dubius*, and *N. pseudodubius* (node k), exclusive ancestor of *N. spinuliferus*, and *N. jatobensis* (node i), and exclusive ancestor of *N. gibber*, and *N. transitans* (node g). Synapapomorphic events are present for *N. incompositus*, *N. santaremensis, N. inflatus*, and *N. cannarensis.* The tracing of this hypothesis to the exclusive ancestor of *Notodiaptomus* corroborates Kiefer (1956) by observing the attribute as a grouping criterion for organisms of the genus (table 2, 4c). Evidently, for some members of the proposal to expand the grouping (Kiefer, 1956), the occurrence of the primitive state of character refutes some of its propositions, such as *N. inflatus*, *N. santaremensis*, *N. incompositus*, and *N. jatobensis.* Despite this, there is no doubt about the importance of the attribute as an evolutionary sign for *Notodiaptomus*.

Another attribute originally present by Kiefer (1956) was traced to the fifth right swimming leg (table 2, 6c) of the common and exclusive ancestor for genus. The condition of the exopod 2 with curved ridge on distal posterior surface (table 5 = #64; S3 = Char 2388: 1►2) represents a state with marked ambiguity between the evolutionary relationships represented. The high homoplastic index (HI = 0.90) and retention index (0.28) evidence the numerous emergencies along the set of hypotheses achieved, with reversals to the exclusive ancestors of *N. santaremensis*, *N. isabelae*, *N. incompositus, N. nelsoni, N. santafesinus*, and *N. anisitsi* (node k). The character state is simplesiomorphic for *N. susanae*, *N. henseni*, *N. iheringi*, *N. spinuliferus*, *N. caperatus*, *N. simillimus*, and *N. cannarensis.* Wright (1935) also present this characteristic for the assembling of the organisms of the *nordestinus* complex (table 2, 3a). Both Wright (1935) and Kiefer (1956) can be corroborated through of the present analysis, except for *N. henseni, N. iheringi*, and *N. santaremensis.* Considering the exclusivity identified for the attribute, we point out its relevance for the reconstruction of the common and exclusive ancestor of *Notodiaptomus*.

Finally, the fifth right swimming leg exopod 2 longer than broad nearly 2 times (table 5 = #62; S3 = Char 2377: 2►1) also represents a condition with marked ambiguity (HI = 0.9) between the evolutionary relationships represented. The character state is reversed from the node d to node f, and to the exclusive ancestors of *N. amazonicus, N. nordestinus, N. cearensis*, and *N. iheringi* (node i), and *N. brandorffi*, *N. paraensis*, *N. caperatus*, *N. gibber*, and *N. transitans* (node g). Character simplesiomorphy is identified for *N. dentatus.* New apomorphic character event is present for the exclusive ancestor of *N. carteri*, and *N. conifer.* Although the retention index (RI = 0.42) indicates the grouping of the character greater than the set of hypotheses represented (Fig. 33), the evidence of the status of this character as belonging to the exclusive ancestor of *Notodiaptomus* is inconsistent (CI = 0.10) and not recommended.

##### Attributes for female fourth and fifth metasome, and antennule

From the females examined it was possible to trace 9 evolutionary hypotheses unique to the common and exclusive ancestor of *Notodiaptomus*. Mostly, the antennulae are the ones that offer the apomorphic states to be highlighted, being possible to observe 6 other plesiomorphic hypotheses traced to the fifth swimming legs. For decades the systematics of South American diaptomids from females was scarce or non-existent, given arguments of strict similarity. The evidence presented below reinforces advances in the comparative morphology of these organisms and highlights the relevance of females for the abduction of informative character states in phylogenetic analyses.

The first hypothesis traced for female prosome deduces the limit between fourth and fifth metasome segments ornamented with spinule row on partially over limit (table 5 = #66; S3 = Char 2456: 1►2) through numerous events of ambiguous emergencies (HI = 0.80) throughout the evolutionary relationships presented. The character undergoes reversal from the node f to node h and has apomorphic emergence to the exclusive ancestor of *N. anisitsi*, and *N. santafesinus.* Convergence events are evidenced for *N. incompositus*, and *N. spinuliferus.* Undoubtedly, the low consistency index suggests a reconsideration of the importance of the attribute for the exclusive ancestor of *Notodiaptomus*, but the evolutionary signal is unequivocal and should be considered for future efforts on the evolutionary relationships on the organisms related to the genus.

New evidence of consistency for the ancestral female of *Notodiaptomus* is present through eight evolutionary hypotheses traced for the antennulae. The losses of the element setal (table 5 = #67, #69, #71, #72, #73, and #74; S3 = Chars 2552, 2555, 2559, 2561, 2565, and 2567: 1►2) and/or aesthetasc (table 5 = #68, and #70; S3 = Chars 2554, and 2557: 1►2) represented an evolutionary signal with a high consistency index in the k, and j nodes, mostly with convergences for *N. deitersi*, *N. anisitsi*, *N. incompositus*, *N. isabelae*, *N. dubius*, *N. nelsoni*, *N. pseudodubius*, and *N. santafesinus*. For the first four species mentioned, the phylogenetic analysis of these characters corroborates the observation recorded in Santos-Silva *et al*. (1999) in offering the diagnosis of *Notodiaptomus* after the designation of *N. deitersi* as a type of the genus. For all organisms represented through the terminals in node j, only *N. henseni*, *N. santaremensis*, and *N. carteri* were simplesiomorphic to the characters, which suggests relevant consistency for the reconstruction of the exclusive ancestor presented and a distinct intrageneric grouping.

#### Morphological basis of the creation for *Notodiaptomus*

The taxonomic establishment of *Notodiaptomus* considered three morphological characters for its foundation (Kiefer, 1936) and eleven for its expansion (Kiefer, 1956) (table 2). As discussed earlier, prior to this, the set of organisms called the *nordestinus* complex was the basis for the genus and had eight characters mainly (table 2). Among the evolutionary hypotheses based on these attributes 3 represent synapomorphies unique to the common and exclusive ancestor of the genus, the others are primitive conditions, widely conserved among the other calanoids of the outer group, and of little significance (table 6).

**TABLE 6.** Consistency (CI) of the evolutionary hypotheses abducted from the fundamental characters of *Notodiaptomus*.

| Original proposal to *nordestinus* complex (Wright, 1935) |  | CI |
| --- | --- | --- |
| #2354 | Male. 5th right leg exopod with distal posterior process in triangular form (“pointed”) | 0.14 |
| #2284 | Male. 5th right leg basis with conspicuous proximal inner angle, rounded “shoulder”, with or without small prominence | 0.12 |
| - | Male. 5th right leg “small” and conical endopod | Subjective |
| #2388 | Male. 5th right leg exopod 2 with angular or rounded prominence on inner margin | 0.09 |
| #2214 | Male. 5th left leg with exopods fused “narrowly”, with presence of “hair” pad | 1.00 |
| #2247 | Male. 5th left leg exopod distal segment with “little spine” and ending in a finger-like process | 0.12 |
| #2172 | Male. 5th left leg with endopod unisegmented | 0.16 |
| #2602 | Female. 5th swimming legs basis with expanded distal inner rim posteriorly | 0.25 |
| Original proposal of the genus *Notodiaptomus* (Kiefer, 1936) |  | CI |
| - | Male. “The nature of the final segment of the left exopod of the male fifth leg” | subjective |
| #424 | Male. Right antennule with conspicuous spines on segments 10, 11, 13, 15 and 16 | 0.50 |
| - | Male. Right antennule with hyaline lamella on 20 segments (“at most one”) | high variation |
| Proposal for expansion to the genus *Notodiaptomus* (Kiefer, 1956) |  | CI |
| #2612 | Female. 5th swimming legs with endopod unisegmented | 0.10 |
| #2623 | Female. 5th swimming legs endopod with row of oblique spinules + 1-2 setae | 0.25 |
| #2263 | Male. 5th swimming legs coxa with rounded lobule and “hyaline” spine | 1.00 |
| #2318 | Male. 5th right swimming leg basis with structure in varied shape and size, lamella posteriorly | 0.11 |
| #2342 | Male. 5th right swimming leg exopod 1 with length greater than width, mostly | 0.20 |
| #2388 | Male. 5th right leg exopod 2 with rounded-broad hyaline protuberance on inner “caudal surface” | 0.09 |
| #2392 | Male. 5th right swimming leg exopod 2 with outer spine on base of the terminal claw generally | 0.33 |
| #2337 | Male. 5th left swimming leg exopod 2 with distal portion digitiform | 1.00 |
| #2247 | Male. 5th left swimming leg exopod 2 with short broad seta, directed inwards, on distal portion | 0.12 |
| #408 | Male. Right antennule without “spineform protuberance” on segment 14 | 1.00 |
| - | Male. Right antennule with narrow hyaline membrane on segment 20 generally | high variation |

"#" character number corresponding to the applied character list (S1); " – " attribute not used for evolutionary analysis.

Among the attributes defined by Wright to group the original organisms of the *nordestimus* basis – later adopted for *Notodiaptomus*, the “sclerotized process” on the exopod 1 of the male fifth right swimming leg was highlighted as a common feature. Evidently, the phylogenetic hypothesis did not consistently contribute (Char #2354; CI = 0.14), being strongly homoplastic (HI = 0.85) in the topology represented (Fig. 33). For the species originally included in the *nordestinus* group, *N. amazonicus*, *N. nordestinus*, *N. henseni*, and *N. iheringi* represented simplesiomorphic terminals for the condition and corroborate Wright (1935). Santos-Silva *et al*. (2015) when observing the original non-inclusion of *N. deitersi* through illustrations that did not represent the structure adequately, evidenced and included the species among the original members of the complex, which was also confirmed here. The character state is consistent in node i, but is also occasionally found in other nodes, such as for the unique ancestor of *N. isabelae* (node k), and the exclusive ancestor of *N. brandorffi*, *N. caperatus*, and *N. paraensis* (node g). Other *Notodiaptomus* not considered by Wright showed simplesiomorphy for the attribute, such as *N. dentatus*, *N. maracaibensis*, *N. orellanai*, and *N. paraensis*.

Of the other seven character states on which Wright (1935) based his creation for the *nordestinus* group, all represent plesiomorphic attributes in some degree of relationship with members selected for the phylogenetic polarization of this evaluation. For the cladogram formed from the common and exclusive ancestor of *Notodiaptomus*, the character 2284 (1►2; HI = 0.87) has unambiguous emergence from node b to node c, and losses in the exclusive ancestor of *N. coniferoides* and *N. simillimus*, and exclusive ancestral of *N. dentatus* and *N. spinuliferus*. Among the thirteen members currently recognized, only *N. anisitsi* did not present the characteristic. Unfortunately, *N. dahli* cannot be evaluated for the present study.

Among the other phylogenetic hypotheses abducted from of the morphology of Wright (1935; 1936), some could not be corroborated for the members of the *nordestinus* complex. The character 2388 (1►2; HI = 0.90) was largely inconsistent and inapplied for *N. henseni*, *N. incompositus*, and *N. anisitsi* (node j). The character 2247 (2►1; HI = 0.87), plesiomorphic in consideration of the *D. susanae* in the outgroup, had present change for *N. cearensis*, *N. incompositus*, and *N. iheringi*. The character 2172 (2►1; HI = 0.83) arises in several directions in the achieved topology, a hypothesis being refuted only for *N. inflatus.* Finally, the character 2602 (1►2; HI = 0.75) appears as an unequivocal condition from the node c to node d, and conserved for all terminals corresponding to the members of *nordestinus*.

All these morphological observations were recorded in the review study of the *nordestinus* complex in Santos-Silva *et al*. (2015). Additionally, some morphological enlargements were part of the new observations offered to group of Wright and were also present throughout our examinations. Among them, the condition for the male fifth right swimming leg endopod fused to basis (char 2323; 1►2; CI = 0.10; HI = 0.90) was abducted as a phylogenetic hypothesis and refuted for the *nordestinus* group. Truly, the endopod of the male fifth right swimming leg was present condition fused to basis for some *Notodiaptomus*, which precludes its recognition to 1-segmented (state applied in #2327). The condition of the rami with segmentations was applied when separation of the endopod from the basis was present, recognized in this effort for *N. henseni*, *N. inflatus, N. nordestinus*, and *N. jatobensis.* Perhaps this is an observation that requires a change in our notion of evolutionary trends within *Notodiaptomus*, and in the future requires further efforts to trace proper homologies.

Among the four main nodes identified for the set of phylogenetic hypotheses achieved here (*i.e.,* node j, node i, node g, and node e), predominantly the members of the *nordestinus* complex were distributed among the node j (*N. deitersi*, *N. henseni*, *N. conifer*, *N. isabelae*, *N. incompositus*, and *N. anisitsi*) and node i (*N. amazonicus*, *N. iheringi*, *N. nordestinus*, *N. cearensis*, and *N. jatobensis*). *N. inflatus* was present exception, forming topology with *N. coniferoides*, and *N. simillimus* into the node e. Both nodes have unambiguous changes from the unique ancestor in node h. From node h to node j conditions left antennule actual segment 1 with spinules as a patch (char 2172; 1►2; CI = 1.00), male fifth left swimming leg exopod 1 with double process present (char 2210; 1►2; CI = 1.00), and female genital double-somite with double posterior-ventral process in lateral view (char 2534; 1►2; CI = 1.00) are inconstant but confined between terminals. From node h to node i, are inequivocal transition of the male fifth right swimming leg basis with ornamentation on protuberance not until to adjacent surface (char 2299; 1►2; CI = 1.00), and female limit between fourth and fifth metasome segments with spinules laterally (char 2257; 1►2; CI = 1.00).

These were some of the characteristics adopted by Kiefer (1936; 1956) for *Notodiaptomus*. With the last study of expansion of the genus, fourteen characteristics were presented for the grouping of organisms associated with the taxa presented. Among them, eleven attributes were abducted as evolutionary hypotheses in the present effort (table 6), with only two synapomorphies of the common and exclusive ancestor of *Notodiaptomus* (table 5, char 61, char 64), discussed earlier.

For the condition of male right antennule with conspicuous spines on segments 10, 11, 13, 15 and 16, all elements on the mentioned segments have present emergence among the terminal *Notodiaptomus*. The morphological condition is compounded and was present through five characters (or phylogenetic hypotheses) throughout our examinations. The larger variation between emergence and loss was present for the “conspicuous spine” on actual segments 15 (and 16), reinterpreted here as spinous process (table 6, character 424). Our results suggest that the evolutionary condition is highly preserved among the *Notodiaptomus* presented by Kiefer (1956), with retention index (IR = 0.66) indicating the character with few changes within the cladogram presented (Fig. 33).

For the female fifth swimming legs endopod unisegmented (char 2612; 2►1; CI = 0.10; HI = 0.90) homoplastic events were present at levels broader to the set of hypotheses reached (Fig. 33). Among the species indicated by Kiefer for *Notodiaptomus* are loss of the character condition *N. incompositus*, and *N. anisitsi* (node k). Furthermore, in several other regions of the topology the character is homoplastic in this sense, such as *N. carteri* (node k), *N. spiniliferus* (node i), and the exclusive ancestral of *N. transitans, N. gibber*, *N. caperatus, N. brandorffi*, *N. paraensis* (node g). For the latter terminal a new reversal to character is present.

Except for synapomorphic conditions #2318, and #2388, none of the other attributes offered in Kiefer (1936; 1956) is sufficiently consistent in the composition of the inner nodes in *Notodiaptomus*. Among the indicated rate grouping (Kiefer, 1956), the character 2318 is hypothesis refuted in the terminals *N. incompositus* (node k), *N. inflatus* (node e), and *N. jatobensis* (node i), with several independent emergencies (HI = 0.88) along the set of hypotheses represented (Fig. 33). The character 2342 has been corroborated for all taxa indicated in Kiefer (1956), but it is considerably inconsistent for the grouping of these organisms in the topology achieved. The character condition has unequivocal emergence from node a to node b, and loss in *N. paraensis.* Other terminals have loss events for this condition, they are *N. orellanai, N. coniferoides*, and *N. cannarensis*.

For the subdistal position of the outer spine on male fifth right swimming leg exopod 2 (char 2392; 1/2►3; CI = 0.33; HI = 0.66), *N. paraensis*, *N. kieferi, N. anisitsi*, and *N. jatobensis* have the evolutionary condition not conserved in the achieved topology. The latter two species have organisms that were considered for *Notodiaptomus* in Kiefer (1956), and here represents refutation to this hypothesis as they possess outer spine medially inserted on exopod 2 of the male fifth right leg. *N. cannarensis* (node a) is one of the most basal taxa among the *Notodiaptomus* and represented the only independent emergence of the condition outer spine premedially inserted, such as *D. castor* among the outgroup. Finally, the absence of spinous process on actual segment 14 of the male antennulae right (char 408; 2►1; CI = 1.00) represents an autapomorphy in *N. gibber*.

It seems that the characters presented in Kiefer (1936; 1956) present some inconsistencies and are of little use for grouping within the set of hypotheses obtained in this effort. Among the original and enlargement members offered, most are grouped in node i, and node j, but without adjusting abducted hypotheses of the originals of the *Notodiaptomus* proposal. We can also assess that *N. maracaibensis* (node b), and *N. inflatus* (node e) represent basal groups within the genus, and are evolutionarily distant from the other original members of the group. Despite all this, it is important to consider that future efforts in the identification and optimization of homologies between current and new members of *Notodiaptomus* may deduce new notions about these characteristics analyzed here.

#### Morphological relationships in *Notodiaptomus*

Throughout the taxonomic trajectory of *Notodiaptomus* hypotheses for possible intraspecific relatedness relations were evidenced through morphological similarities. Never before have any of these been phylogenetically tested, and here we have the opportunity to observe these results through the topology obtained. Most of the hypotheses were supported in the present cladistic analysis, others are refuting phylogenetic inferences and evidence evolutionary paths in divergent directions.

Paggi (2001) when addressing the set of species defined as “*bidigitatus* group” (Brehm, 1958) reported the morphological proximity and indicated an evolutionary relationship between *N. spinuliferus*, and *N. dentatus*. Evidently the species are terminals derived from a common and exclusive ancestor in node i, and this corroborates the indication of Paggi (2001). The confirmed close evolutionary relationship is supported through the reconstruction of this ancestor with the female fifth swimming legs endopod with single seta (char 2627; 2►1; CI = 0.50; RI = 0.50), female limit between fourth and fifth metasome segments with spinules row singly (char 2453; 2►1; CI = 0.33; RI = 0.33), and first swimming legs coxa with one group of setules on inner margin as a patch (char 1454; 1►2; CI = 0.50; IR = 0.50). For *N. dentatus* the emergence of the caudal rami with outermost seta with outer spiniform process present on outermost seta (char 122; 1►2; CI = 1.00) represents unambiguous evolutionary transition and separates from *N. spinuliferus*.

Reid (1987) in presenting the hypothesis of *N. brandorffi* related the taxon organisms to those of *N. spinuliferus*, *N. carteri, N. deitersi*, *N. iheringi, N. jatobensis*, *N. anisitsi,* and *N. isabelae*. Our phylogenetic results suggest that the morphological proximity hypothesis does not represent a sustained evolutionary relationship between the organisms of these species. All the morphological evidence for the relationships indicated in Reid (1987) could be corroborated through the morphological analyses performed here, except for *N. deitersi* for the condition of female fifth swimming legs endopod 2-segmented not confirmed in the present effort. Truly, the species of Reid (1987) has a closer evolutionary relationship to *N. paraenses* (node g) through the common and exclusive ancestor reconstructed with antenna endopod actual 3-segmented (char 990; 1►2; CI = 0.50; RI = 0.66), mandible coxal gnathobase with protuberance not proeminente on caudal margin (char 1074; 1►2; CI = 0.50), male fifth right swimming leg basis with inner protuberance distally (char 2293; 1/2►3; CI = 0.50; RI = 0.50), male fifth right swimming leg exopod 2 with outer spine lesser than the length of the exopod 2 beyond to 2 times its size (char 2405; 1►2; CI = 0.12; RI = 0.30), and female right antennule not extending beyond caudal rami (char 2548; 1►2; CI = 0.12; RI = 0.12).

Previattelli *et al*. (2017) when reporting the organisms of *N. nelsoni* related close to those of *N. paraensis* through similarities of male metasome segments, male fifth right leg coxa, and female fifth swimming legs. Our results suggest refutation of this hypothesis and indicate that *N. nelsoni* organisms have a closer evolutionary relationship to *N. anisitsi* and *N. santafesinus* (node k) through the exclusive ancestor retrieved with first swimming legs coxa with inner distal seta with length surpassing to basis (char 1451; 2►1; CI = 0.16; RI = 0.50), male fifth right swimming leg coxa with distal process projecting over basis beyond the first third posteriorly (char 2269; 1►2; CI = 0.11; RI = 0.20), male fifth right swimming leg exopod 2 with outer spine lesser than the length of the exopod 2 1.5x its size (char 2407; 2►1; CI = 0.14; RI = 0.40), and female fourth and fifth metasome segments fused partially (char 2448; 1►2; CI = 0.11). *N. nelsoni* is evolutionarily unique through the unequivocal emergence of the female genital double-somite with bifid apex sensilla on left side (char 2500; 1►2; CI = 1.00), and the homoplastic characters left antennule actual segment 1 with spinules (char 592 ; 1►2; CI = 0.33), and male fifth right swimming legs exopod 1 with posterior lamella present (char 2371; 1►2; CI = 0.20), mainly.

Among the four subgenus of *Notodiaptomus* proposed in Dussart (1985), *Notodiaptomus (Wrightius)* was a hypothesis of morphological relationship for *N. paraensis*, *N. gibber, N. inflatus, N. anceps, N. lobifer*, *N. kieferi, N. orellanai*, and *N. dilatatus*. Although the hypothesis of Dussart (1985) is admittedly unfounded and has consistent reason for refutation, our results corroborated the evolutionary relationship of *N. paraensis, N. gibber*, and *N. kieferi* (node g) – *N. anceps*, *N. lobifer*, and *N. dilatatus* were not included in the present effort. Among these species, *N. paraensis* is discussed above closest to *N. brandorffi*, where its unique ancestor is characterized. *N. paraensis,* in turn, represents a distinct evolutionary hypothesis through the unequivocal emergence of the first swimming legs coxa without setules on inner margin (char 1452; 1►2; CI = 1.00), and homoplastic characters female fifth swimming legs basis with proximal inner outgrowth (char 2599; 1►2; CI = 0.50), male fifth right swimming leg exopod 2 with outer spine larger than or equal to the length of the exopod 2 (char 2404; 1►2; CI = 0.50).

The hypothesis raised in Defaye & Dussart (1988) for the proximity between *N. pseudodubius* and *N. dubius* organisms is corroborated for node k in the topology presented here. Common and exclusive ancestry for both taxa is supported through the unambiguous appearance of the male fifth right swimming leg coxa with distal process projecting over basis until the distal surface posteriorly (char 2270; 1►2; CI = 1.00). From this hypothetical ancestor, *N. pseudodubius* has distinct evolutionary lineage by the homoplastic appearance of the male right epimeral plate with one sensilla at the apex of projection and other medially (char 66; 1►2; CI = 0.12; RI = 0.30), female right epimeral plate not thinner than the left (char 2464; 1►2; CI = 0.14; RI = 0.25), and female genital double-somite without right posterior rim expanded (char 2515; 2►1; CI = 0.09; RI = 0.16). *N. dubius* has evolutionary distinction supported through of the unequivocal emergence male fifth right swimming leg exopod 2 wider than long (char 2376; 1►2; CI = 1.00).

The formal presentation of *N. simillimus* from the erroneous recognition of organisms of the species as *N. coniferoides* (Cicchino *et al*., 2001) suggests the close relationship between these organisms. In our topology achieved this hypothesis is confirmed, and both taxa have a common and unique ancestor in the node e, that is still composed of *N. inflatus.* The hypothetical ancestor of these early organisms is reconstructed from the unambiguous emergence of the male right antennule actual segment 17 with two modified setae (char 460; 1►2; CI = 1.00), and homoplastic characters male fifth right swimming leg basis with tumescence absent (char 2284; 2►1; CI = 0.12; HI = 0.41), and female fifth swimming legs endopod 2-segmented (char 2612; 2►1; CI = 0.10; IR = 0.30). *N. simillimus* represented a distinct evolutionary hypothesis of *N. coniferoides* through the unambiguous appearance in *Notodiaptomus* of the female genital double-somite with sensilla on lobular base on left side (char 2502; 1►2; CI = 0.50), and the homoplastic loss of the curved ridge on distal posterior surface of the male fifth right swimming leg exopod 2 (char 2388; 2►1; CI = 0.09; IR = 0.28).

However, the phylogenetic analysis of *Notodiaptomus* organisms recovers several evolutionary lineages and widely confirms several relationships indicated in the literature. However, none or almost none of these confirmed relationships can be supported through the morphological hypotheses offered in the originals of each species. It is probably that the tracing of new evolutionary series for the male and female metasome segments, epimeral plates, urosome, legs 1 to 4, and oral appendices will sustain this trend and offer from here new notions about the evolution of character forms beyond antennules, and fifth swimming legs.

### Final considerations

The morphology of *Notodiaptomus* was extensively reviewed and expanded in the present study. Some of the structures are again described here from the recent and historical evaluations of group members, such as antennules, fifth swimming legs, and female genital double-somite (Wright, 1935; Wright 1936; Kiefer, 1936, Kiefer, 1956; Santos-Silva *et al*., 1999; Santos-Silva *et al*., 2015). Others were present in more detail, such as fifth metasome segments, epimeral plates, oral appendages, and dimensional spectra of various morphological fractions (length, width, and relative size).

Historically, the identification of *Notodiaptomus* organisms has been imputed through of the importance of the presence of male fifth right swimming leg exopod 1 with distal process posteriorly (Wright, 1935; Santos-Silva *et al*., 2015), and “the nature of the final segment of the left exopod of the male fifth leg” (Kiefer, 1936, p. 23). Unfortunately, for the characteristic indicated in Kiefer (1936) subjectivity prevented an objective evaluation of this attribute, but when we consider the structure referred to exopod 2 probably, we observe the common presence of the distal portion in digitiform for all members originally considered, and the spiniform seta not surpassing distal-point of the segment for 27 (87.5%) of all species currently valid and approached in this present effort. To the highlight of Wright (1935) and Santos-Silva *et al*. (2015), both the original members of the *nordestinus* group and those of Kiefer’s efforts possessed the attribute (except *N. carteri*). In addition, among the *Notodiaptomus* we addressed were the exception *N. caperatus*, *N. cannarensis*, *N. dubius*, and *N. brandorffi*.

Through the phylogenetic analysis presented, only the morphological reinterpretation for indication of Kiefer (1936; 1956) to respect of the spineform seta not surpassing to distal-point of the exopod 2 of the male fifth swimming leg possessed consistency for the reconstruction of the exclusive and common ancestor of *Notodiaptomus*. However, other characteristics highlighted in the review of the *nordestinus* complex as a basis for the genus (Santos-Silva *et al*., 2015) were corroborated and consistent with an evolutionary trend within the group, such as the setae on the male antennules, male endopod of the fifth swimming legs, and other important elements of the male and female fifth swimming leg, previously evaluated. It is likely that future studies will indicate new evolutionary trends for the grouping as the various *Diaptomus* (sensu lato) hypotheses from South America are reassessed and other neotropical taxonomic groupings follow the same path. Furthermore, less inclusive homologies (*i.e.*, character states) traced on the female genital double-somite, and positioning of sensillae on prosome dorsally may add new intrageneric notions. It is possible that biological morphometry could be an important application in this regard.

Although there are arguments for the formal recognition of some Notodiaptomus partially, the recovery of all evolutionary lineages (terminals) allows the tracing of unique phylogenetic conditions for the reconstruction of the common and exclusive ancestor of the genus (*e.g.*, table 5, chars 2, 3, 10, 11, 15, 16, 20, 41, 57, 66). Evidently in the topology achieved, all *Notodiaptomus* of the *nordestinus* complex of Wright (1935; 1936), and the foundation and amplification of Kiefer (1936; 1956) are mostly retained in node j, and node k. Among the new specific hypotheses inserted in the genus, seven species are evolutionarily related to this group, being *N. nelsoni*, *N. santafesinus*, *N. dubius*, *N. pseudodubius* (node j), and *N. dentatus* and *N. spinuliferus* (node i).

## Acknowledgments

We thank the Instituto Nacional de Pesquisas da Amazônia (INPA), through the Graduate Program Biologia de Água Doce e Pesca Interior (PPG BADPI) for the possibility of conducting this study. We gratefully acknowledge CAPES (Coordenação de Aperfeiçoamento de Pessoal de Nível Superior) for granting the funding for PHD project of the first author (Finance Code 001). We also thank Dr. Carlos Eduardo Falavigna da Rocha (USP) for the directions for this research and logistical support. To Dr. Marcos Domingos Siqueira Tavares for authorizing access to the carcinological collection of the Museum of Zoology of the University of São Paulo – MZUSP. We extend our thanks to Dr. Gilmar Perbiche-Neves (UFSCar) for the grant of a significant part of the biological material considered for this research.

