## Supplemental Data 1 for "Biological Systematic of the genus *Notodiaptomus* Kiefer 1936 (Copepoda: Calanoida)"

**Supplementary material S1** – Applied character list.

#1. Habitat/
1. in brackish waters/
2. in freshwaters/

#2. nerve axons <myelination>/
1. not myelinated/
2. myelinated/

#3. Male body <size compared to female in dorsal view>/
1. smaller and slenderer than female/
2. smaller and wider than female/

#4. Prosome <number of distinct segments dorsally>/
1. 5-segmented/
2. 6-segmented/

#5. Prosome <Male, widest region, in dorsal view>/
1. widest at first metasome segment/
2. widest at cephalosome/

#6. Prosome <Male, widest region of cephalosome, in dorsal view>/
1. medially/
2. posteriorly/

#7. Prosome <Male, presence of setules at posterior margins of the prosomal segments>/
1. without one line of setules at posterior margin/
2. with one line of setules at posterior margin/

#8. Prosome <Male, presence of spinules at posterior margins of the segments>/
1. without spinules at segments/
2. with spinules at least at one segment/

#9. Cephalosome <shape of the anterior margin in dorsal view>/
1. anterior margin rounded/
2. anterior margin sub-triangular/

#10. Cephalosome <presence of suture>/
1. without dorsal suture/
2. with dorsal suture/

#11. Cephalosome <suture completeness>/
1. incomplete/
2. complete/

#12. Cephalosome <fusion>/
1. separate from first metasome segment/

2. fused to first metasome segment/

#13. First metasome segment <presence of sensilla>/
1. without sensilla/
2. with sensilla/

#14. First metasome segment <sensillae location>/
1. <dorsally>/
2. <laterally>/

#15. First metasome segment <total of dorsal sensillae>/
1. 2 dorsally/
2. 4 dorsally/

#16. First metasome segment <number of lateral sensillae>/
1. 2 laterally/
2. 4 laterally/

#17. First metasome segment <size comparison between sensillae>/
1. of equal size/
2. of unequal size/

#18. Second metasome segment <presence of sensilla>/
1. without sensilla/
2. with sensilla/

#19. Second metasome segment <insertion of the sensillae>/
1. <dorsally>/
2. <laterally>/

#20. Second metasome segment <number of dorsal sensillae>/
1. 2 dorsally/
2. 4 dorsally/

#21. Second metasome segment <number of lateral sensillae>/
1. 2 laterally/
2. 4 laterally/

#22. Second metasome segment <size comparison between sensillae>/
1. of equal size/
2. of unequal size/

#23. Third metasome segment <presence of sensillae>/
1. without sensillae/
2. with sensillae/

#24. Third metasome segment <dorsal or lateral>/
1. <dorsally>/
2. <laterally>/

#25. Third metasome segment <number of dorsal sensillae>/
1. 2 dorsally/
2. 4 dorsaly/

#26. Third metasome segment <number of lateral sensillae>/
1. 2 laterally/
2. 4 laterally/

#27. Third metasome segment <size comparison to each other>/
1. of equal size/
2. of unequal size/

#28. Third metasome segment <ornamentation of posterior margin>/
1. non-ornamented posterior margin/
2. ornamented posterior margin/

#29. Third metasome segment <type of ornamentation on posterior margin>/
1. with setules/
2. with spinules/

#30. Third metasome segment <arrangement of the ornamentation on posterior margin>/
1. as a row/
2. as a patch/

#31. Third metasome segment <number of row on posterior margin>/
1. single/
2. double/

#32. Third metasome segment <position of ornamentation on posterior margin>/
1. dorsally/
2. laterally/

#33. Fourth metasome segment <presence of sensillae>/
1. without sensillae/
2. with sensillae/

#34. Fourth metasome segment <dorsal or lateral>/
1. <dorsally>/
2. <laterally>/

#35. Fourth metasome segment <number of dorsal sensillae>/
1. 2 dorsally/
2. 4 dorsally/

#36. Fourth metasome segment <number of lateral sensillae>/
1. 2 laterally/
2. 4 laterally/

#37. Fourth metasome segment <size comparison to each other>/
1. of equal size/
2. of unequal size/

#38. <male> Fourth metasome segment <fusion or separation indicated by suture presence, absence or partially presence>/
1. separated from the fifth metasome/
2. fused to fifth metasome/

#39. <male> fourth metasome segment <state of the fusion to fifth metasome>/
1. totally/
2. partially/

#40. <male> fourth metasome segment <location of the fusion to fifth metasome>/
1. on lateral surface/
2. on dorsal surface/

#41. <male> limit between fourth and fifth metasome segments <presence of ornamentation>/
1. without ornamentation/
2. ornamented/

#42. <male> limit between fourth and fifth metasome segments <type of ornamentation>/
1. with spinules/
2. with setules/

#43. <male> limit between fourth and fifth metasome segments <arrangement of the spinules>/
1. as a row/
2. as a patch/

#44. <male> limit between fourth and fifth metasome segments <composition dorsal spinules row>/
1. on dorsal singly/
2. on dorsal doubly/

#45. <male> limit between fourth and fifth metasome segments <composition lateral spinules row>/
1. on lateral singly/
2. on lateral doubly/

#46. <male> limit between fourth and fifth metasome segments <size comparison of spinules>/
1. same size/
2. different size/

#47. <male> fifth metasome segment <presence of sensilla>/
1. without sensilla/

2. with sensilla/

#48. <male> fifth metasome segment <restrict area insertion of the sensilla>/
1. <dorsally>/
2. <laterally>/

#49. <male> fifth metasome segment <number of dorsal sensillae>/
1. 2 dorsally/
2. 4 dorsally/

#50. <male> fifth metasome segment <number of lateral sensillae>/
1. 2 laterally/
2. 4 laterally/

#51. Fifth metasome segment <size comparison between sensillae>/
1. equal size/
2. unequal size/

#52. Fifth metasome segment <ornamentation>/
1. without ornamentation/
2. ornamented/

#53. Fifth metasome segment <type of ornamentation>/
1. with spinules/
2. with setules/

#54. Fifth metasome segment <arrangement of ornamentation>/
1. as a row/
2. as a patch/

#55. Fifth metasome segment <number of row>/
1. single/
2. double/

#56. Fifth metasome segment <arrangement of ornamentation on posterior margin>/
1. continuous/
2. discontinuous/

#57. Fifth metasome segment <patch position of ornamentation>/
1. dorsally/
2. medially/

#58. <Male> Fifth metasome segment <observation of dorsal hump>/
1. without dorsal conical process/
2. with dorsal conical process present/

#59. <male> fifth metasome segment <presence of epimeral plates>/
1. without epimeral plates/
2. with epimeral plates/

#60. <male> Epimeral plates <symmetry>/
1. symmetrical/
2. asymmetrical/

#61. <male> Right epimeral plates <form>/
1. reduced, as rounded distal corner segment limit/
2. prominent, as projections/

#62. <male> Right epimeral plate <number of projections>/
1. one projection/
2. two projections/

#63. <male> Right epimeral plate <direction>/
1. posterior-laterally directed/
2. posterior-dorsally directed/

#64. <male> Right epimeral plate <size of projection compared to genital segment>/
1. not reaching half length of the genital segment/
2. reaching half or surpassing the length of genital segment/

#65. <male> Right epimeral plate <presence of sensillae>/
1. without sensilla/
2. with sensilla/

#66. <male> Right epimeral plate <position of sensillae>/
1. at the apex of projection/
2. one at the apex of projection and other medially/

#67. <male> Right epimeral plate <presence of ornamentation>/
1. without ornamentation/
2. ornamented/

#68. <male> Right epimeral plate <type of ornamentation>/
1. with spinules/
2. with setules/

#69. <male> Right epimeral plate <arrangement of ornamentation>/
1. as a row/
2. as a patch/

#70. <male> Left epimeral plate <form>/
1. reduced, as rounded distal corner segment limit/
2. prominent, as projection/

#71. <male> Left epimeral plate <number of projections>/
1. one projection/
2. two projections/

#72. <male> Left epimeral plate <direction>/
1. posterior-laterally directed/
2. posterior-dorsally directed/

#73. <male> Left epimeral plate <size of projection in relation to genital segment>/
1. reaching half length of the genital segment/
2. reaching beyond half length of the genital segment/

#74. <male> Left epimeral plate <presence of sensillae>/
1. with sensillae/
2. without sensillae/

#75. <male> Left epimeral plate <position of sensillae>/
1. at the apex of projection/
2. one at the apex of projection and other medially/

#76. <male> Left epimeral plate <presence of ornamentation>/
1. without ornamentation/
2. ornamented/

#77. <male> Left epimeral plate <type of ornamentation>/
1. with spines/
2. with spinules/
3. with setules/

#78. <male> Left epimeral plate <arrangement of ornamentation>/
1. as a row/
2. as a patch/

#79. <male> Urosome <number of segments>/
1. 5-segmented/
2. 4-segmented/

#80. <male> genital somite <symmetry in dorsal view>/
1. symmetrical in dorsal view/
2. asymmetrical in dorsal view/

#81. <male> genital somite <number of genital aperture>/
1. with single aperture/
2. with two apertures/

#82. <male> genital somite <location of genital aperture>/
1. located on left side/
2. located on right side/

#83. <male> genital somite <position of genital aperture>/
1. ventrolaterally on posterior rim/

2. ventrolaterally on anterior rim/

#84. <male> genital somite <presence of sensillae>/
1. without sensillae/
2. with sensillae/

#85. <male> genital somite <location of the sensillae>/
1. on single side/
2. on both sides/

#86. <male> genital somite <location of the sensillae in dorsal view>/
1. <present on left side>/
2. <present on right side>/

#87. <male> genital somite <number of sensillae on left side>/
1. one/
2. two/

#88. <male> genital somite <location of sensillae on left side>/
1. at left rim/
2. at left lateral/

#89. <male> genital somite <insertion of sensillae on left side>/
1. medially/
2. posteriorly/

#90. <male> genital somite <number of sensillae on right side>/
1. one/
2. two/

#91. <male> genital somite <location of sensillae on right side>/
1. at right rim/
2. at right lateral/

#92. <male> genital somite <insertion of sensillae on right side>/
1. medially/
2. posteriorly/

#93. <male> genital somite <size comparison between sensillae>/
1. of equal size between then/
2. of unequal size between then/

#94. <male> genital somite <size comparison between sensillae>/
1. left bigger than right/
2. right bigger than left/

#95. Third urosome segment <presence of spinules>/
1. without spinules/

2. with spinules/

#96. Third urosome segment <arrangement of ornamentation>/
1. as a row/
2. as a patch/

#97. Third urosome segment <number of row>/
1. single throughout dorsal surface/
2. double throughout dorsal surface/

#98. Third urosome segment <seta present>/
1. without external seta/
2. with external seta/

#99. Fourth urosome segment <presence of spinules>/
1. without spinules/
2. with spinules/

#100. Fourth urosome segment <arrangement of ornamentation>/
1. as a row/
2. as a patch/

#101. Fourth urosome segment <number of row>/
1. single throughout dorsal surface/
2. double throughout dorsal surface/

#102. Fourth urosome segment <observation of sub-conical blunt dorsal-lateral process>/
1. without sub-conical blunt dorsal-lateral process/
2. with sub-conical blunt dorsal-lateral process/

#103. Fourth urosome segment <presence of sensilla on sub-conical blunt dorsal-lateral process>/
1. not bearing sensilla on the apex/
2. bearing sensilla on the apex/

#104. Anal segment <observation of sensillae>/
1. absence of dorsal sensillae/
2. presence of dorsal sensillae/

#105. Anal segment <number of sensillae>/
1. one on each side/
2. two on each side/

#106. Anal segment <position sensillae>/
1. medially inserted/
2. distally inserted/

#107. Anal segment <observation of operculum>/

1. absence of operculum/
2. presence of operculum/

#108. Anal segment <shape of operculum>/
1. convex/
2. concave/

#109. Anal segment <development of operculum>/
1. not covering the anal aperture fully/
2. covering the anal aperture fully/

#110. Caudal rami <symmetry>/
1. symmetrical/
2. asymmetrical/

#111. Caudal rami <fusion to anal segment>/
1. fused to anal segment/
2. separated from anal segment/

#112. Caudal rami <relation of size to symmetry ramus>/
1. longer than wide/
2. wider than long/

#113. Caudal rami <presence of setules>/
1. without setules/
2. with setules/

#114. Caudal rami <position of ornamentation>/
1. continuous on/
2. discontinuous on/

#115. Caudal rami <insertion of ornamentation>/
1. inner side/
2. outer side/

#116. Caudal rami <number of setae>/
1. each ramus bearing 6 caudal setae/
2. each ramus bearing 5 caudal setae/

#117. Caudal rami <number of marginal setae>/
1. 5 marginal/
2. 4 marginal/

#118. Caudal rami <composition of marginal setae>/
1. plumose/
2. straight/

#119. Caudal rami <number of dorsal setae>/
1. and 1 internal dorsally/

2. and 1 external dorsally/

#120. Caudal rami <composition of dorsal setae>/
1. plumose/
2. straight/

#121. Caudal rami <condition of the main axis of the caudal setae>/
1. not reticulated main axis/
2. reticulated main axis/

#122. Caudal rami <presence of outer proximal spiniform process on outermost seta>/
1. outermost seta with outer spiniform process absent/
2. outermost seta with outer spiniform process present/

#123. <male> Rostrum <symmetry>/
1. symmetrical/
2. asymmetrical/

#124. Rostrum <fusion to dorsal cephalic shield>/
1. fused to dorsal cephalic shield/
2. separated from dorsal cephalic shield/

#125. Rostrum <state of the suture>/
1. by complete suture/
2. by incomplete suture/

#126. Rostrum <observation of sensillae>/
1. sensillae absent/
2. sensillae present/

#127. Rostrum <number os sensillae>/
1. one pair/
2. two pair/

#128. Rostrum <position sensillae>/
1. anteriorly inserted on surface tegument/
2. medially inserted on tegument/

#129. Rostrum <presence of rostral filaments>/
1. without rostral filament/
2. with rostral filament/

#130. Rostrum <number of rostral filaments>/
1. single/
2. double/

#131. Rostrum <pairing of the rostral filaments>/
1. paired/

2. not paired/

#132. Rostrum <rostral filament extended or not extended>/
1. extended/
2. not extended/

#133. Rostrum <form of the distal margin of the rostral filaments>/
1. into point/
2. into blunt protrusion/

#134. <male> Rostrum <presence of basal process>/
1. withou basal process/
2. basal process present/

#135. <male> Rostrum <shape of the basal process>/
1. in ventral view, rounded on left side/
2. in ventral view, triangular on left side/

#136. <male> Rostrum <presence of the basal process>/
1. without a smaller basal expansion on the right side/
2. with a smaller basal expansion on the right side/

#137. <male> antennules <symmetry>/
1. symmetrical/
2. asymmetrical/

#138. <male> right antennule <number of rami, dorsal view>/
1. uniramous/
2. biramous/

#139. <male> right antennule <relative length>/
1. surpassing to genital segment/
2. not surpassing to genital segment/

#140. <male> right antennule <relative length>/
1. extending beyond caudal rami/
2. not extending beyond caudal rami/

#141. <male> right antennule ancestral segment I and II <fusion>/
1. separated/
2. fused/

#142. <male> right antennule ancestral segment II and III <fusion>/
1. separated/
2. fused/

#143. <male> right antennule ancestral segment III and IV <fusion>/
1. separated/
2. fused/

#144. <male> right antennule ancestral segment IV and V <fusion>/
1. separated/
2. fused/

#145. <male> right antennule ancestral segment V and VI <fusion>/
1. separated/
2. fusion/

#146. <male> right antennule ancestral segment VI and VII <fusion>/
1. separated/
2. fused/

#147. <male> right antennule ancestral segment VII and VIII <fusion>/
1. separated/
2. fused/

#148. <male> right antennule ancestral segment VIII and IX <fusion>/
1. separated/
2. fused/

#149. <male> right antennule ancestral segment IX and X <fusion>/
1. separated/
2. fused/

#150. <male> right antennule ancestral segment X and XI <fusion>/
1. separated/
2. fused/

#151. <male> right antennule ancestral segment XI and XII <fusion>/
1. separated/
2. fused/

#152. <male> right antennule ancestral segment XII and XIII <fusion>/
1. separated/
2. fused/

#153. <male> right antennule ancestral segment XIII and XIV <fusion>/
1. separated/
2. fused/

#154. <male> right antennule ancestral segment XIV and XV <fusion>/
1. separated/
2. fused/

#155. <male> right antennule ancestral segment XV and XVI <fusion>/
1. separated/
2. fused/

#156. <male> right antennule ancestral segment XVI and XVII <fusion>/
1. separated/
2. fused/

#157. <male> right antennule ancestral segment XVII and XVIII <fusion>/
1. separated/
2. fused/

#158. <male> right antennule ancestral segment XVIII and XIX <fusion>/
1. separated/
2. fused/

#159. <male> right antennule ancestral segment XIX and XX <fusion>/
1. separated/
2. fused/

#160. <male> right antennule ancestral segment XX and XXI <fusion>/
1. separated/
2. fused/

#161. <male> right antennule ancestral segment XXI and XXII <fusion>/
1. separated/
2. fused/

#162. <male> right antennule ancestral segment XXII and XXIII <fusion>/
1. separated/
2. fused/

#163. <male> right antennule ancestral segment XXIII and XXIV <fusion>/
1. separated/
2. fused/

#164. <male> right antennule ancestral segment XXIV and XXV <fusion>/
1. separated/
2. fused/

#165. <male> right antennule ancestral segment XXV and XXVI <fusion>/
1. separated/
2. fused/

#166. <male> right antennule ancestral segment XXVI and XXVII <fusion>/
1. separated/
2. fused/

#167. <male> right antennule ancestral segment XXVII and XXVIII <fusion>/
1. separated/
2. fused/

#168. <male> right antennule actual segment <number of segments>/

1. 21-segmented/
2. 22-segmented/

#169. <male> right antennule actual segment <geniculation>/
1. not-geniculated/
2. geniculated/

#170. <male> right antennule actual segment <position of the geniculation>/
1. between the segment 17 and segment 18/
2. between the segment 18 and segment 19/

#171. <male> right antennule actual segment <presence of swollen and modified region>/
1. with swollen and modified region/
2. without swollen and modified region/

#172. <male> right antennule actual segment <number of segments of the swollen and modified region>/
1. formed by 5 segments/
2. formed by 4 segments/

#173. <male> right antennule actual segment 1 <presence of seta>/
1. without seta/
2. with seta/

#174. <male> right antennule actual segment 1 <number of setae>/
1. one element/
2. two elements/

#175. <male> right antennule actual segment 1 <composition of seta>/
1. straight/
2. plumose/

#176. <male> right antennule actual segment 1 <size comparison of setae>/
1. of equal size/
2. of unequal size/

#177. <male> right antennule actual segment 1 <length of seta>/
1. none larger than segment/
2. one larger than segment/

#178. <male> right antennule actual segment 1 <range of the seta>/
1. not surpassing distal margin/
2. surpassing to distal margin/

#179. <male> right antennule actual segment 1 <range of the seta>/
1. beyond three sequential segments/
2. not beyond three sequential segments/

#180. <male> right antennule actual segment 1 <presence of spinules>/

1. without spinules/
2. with spinules/

#181. <male> right antennule actual segment 1 <presence of vestigial seta>/
1. without vestigial seta/
2. with vestigial seta/

#182. <male> right antennule actual segment 1 <presence of conical seta>/
1. without conical seta/
2. with conical seta/

#183. <male> right antennule actual segment 1 <presence of modified seta>/
1. without modified seta/
2. with modified seta/

#184. <male> right antennule actual segment 1 <presence of spinous process>/
1. without spinous process/
2. with spinous process/

#185. <male> right antennule actual segment 1 <presence of aesthetasc>/
1. without aesthetasc/
2. with aesthetasc/

#186. <male> right antennule actual segment 1 <number of aesthetasc>/
1. one element/
2. two elements/

#187. <male> right antennule actual segment 2 <presence of seta>/
1. with seta/
2. without seta/

#188. <male> right antennule actual segment 2 <number of setae>/
1. two elements/
2. three elements/

#189. <male> right antennule actual segment 2 <size comparison of seta>/
1. of equal size/
2. of unequal size/

#190. <male> right antennule actual segment 2 <composition of seta>/
1. straight/
2. plumose/

#191. <male> right antennule actual segment 2 <length of seta>/
1. none larger than segment/
2. one larger than segment/

#192. <male> right antennule actual segment 2 <range of the seta>/
1. surpassing distal margin/

2. surpassing to distal margin/

#193. <male> right antennule actual segment 2 <range of the seta>/
1. beyond three sequential segments/
2. not beyond three sequential segments/

#194. <male> right antennule actual segment 2 <presence of spinules>/
1. without spinules/
2. with spinules/

#195. <male> right antennule actual segment 2 <presence of vestigial seta>/
1. without vestigial seta/
2. with vestigial seta/

#196. <male> right antennule actual segment 2 <number of vestigial seta>/
1. one element/
2. three elements/

#197. <male> right antennule actual segment 2 <presence of conical seta>/
1. without conical seta/
2. with conical seta/

#198. <male> right antennule actual segment 2 <presence of modified seta>/
1. without modified seta/
2. with modified seta/

#199. <male> right antennule actual segment 2 <presence of spinous process>/
1. without spinous process/
2. with spinous process/

#200. <male> right antennule actual segment 2 <presence of aesthetasc>/
1. without aesthetasc/
2. with aesthetasc/

#201. <male> right antennule actual segment 2 <number of aesthetasc>/
1. one element/
2. two elements/

#202. <male> right antennule actual segment 3 <presence of seta>/
1. without seta/
2. with seta/

#203. <male> right antennule actual segment 3 <number of setae>/
1. one element/
2. three elements/

#204. <male> right antennule actual segment 3 <size comparison of seta>/
1. of equal size/
2. of unequal size/

#205. <male> right antennule actual segment 3 <length of seta>/
1. one larger than segment/
2. none larger than segment/

#206. <male> right antennule actual segment 3 <range of the seta>/
1. surpassing distal margin/
2. surpassing to distal margin/

#207. <male> right antennule actual segment 3 <range of the seta>/
1. not beyond three sequential segments/
2. beyond three sequential segments/

#208. <male> right antennule actual segment 3 <composition of seta>/
1. straight/
2. plumose/

#209. <male> right antennule actual segment 3 <apex of the seta>/
1. blunt apex/
2. sharp apex/

#210. <male> right antennule actual segment 3 <presence of spinules>/
1. without spinules/
2. with spinules/

#211. <male> right antennule actual segment 3 <presence of vestigial seta>/
1. without vestigial seta/
2. with vestigial seta/

#212. <male> right antennule actual segment 3 <number of vestigial seta>/
1. one element/
2. two elements/

#213. <male> right antennule actual segment 3 <presence of conical seta>/
1. without conical seta/
2. with conical seta/

#214. <male> right antennule actual segment 3 <presence of modified seta>/
1. without modified seta/
2. with modified seta/

#215. <male> right antennule actual segment 3 <presence of spinous process>/
1. without spinous process/
2. with spinous process/

#216. <male> right antennule actual segment 3 <presence of aesthetasc>/
1. without aesthetasc/
2. with aesthetasc/

#217. <male> right antennule actual segment 3 <number of aesthetasc>/
1. one element/
2. two elements/

#218. <male> right antennule actual segment 4 <presence of seta>/
1. without seta/
2. with seta/

#219. <male> right antennule actual segment 4 <number of setae>/
1. one element/
2. two elements/

#220. <male> right antennule actual segment 4 <size comparison of seta>/
1. of equal size/
2. of unequal size/

#221. <male> right antennule actual segment 4 <length of seta>/
1. none larger than segment/
2. one larger than segment/

#222. <male> right antennule actual segment 4 <range of the seta>/
1. surpassing to distal margin/
2. surpassing distal margin/

#223. <male> right antennule actual segment 4 <composition of seta>/
1. straight/
2. plumose/

#224. <male> right antennule actual segment 4 <range of the seta>/
1. beyond three sequential segments/
2. not beyond three sequential segments/

#225. <male> right antennule actual segment 4 <presence of spinules>/
1. without spinules/
2. with spinules/

#226. <male> right antennule actual segment 4 <presence of vestigial seta>/
1. without vestigial seta/
2. with vestigial seta/

#227. <male> right antennule actual segment 4 <quantity of vestigial seta>/
1. 1 element/
2. 2 elements/

#228. <male> right antennule actual segment 4 <presence of conical seta>/
1. without conical seta/
2. with conical seta/

#229. <male> right antennule actual segment 4 <presence of modified seta>/

1. without modified seta/
2. with modified seta/

#230. <male> right antennule actual segment 4 <presence of spinous process>/
1. without spinous process/
2. with spinous process/

#231. <male> right antennule actual segment 4 <presence of aesthetasc>/
1. without aesthetasc/
2. with aesthetasc/

#232. <male> right antennule actual segment 4 <number of aesthetasc>/
1. one element/
2. two elements/

#233. <male> right antennule actual segment 5 <presence of seta>/
1. without seta/
2. with seta/

#234. <male> right antennule actual segment 5 <number of setae>/
1. one element/
2. two elements/

#235. <male> right antennule actual segment 5 <composition of seta>/
1. straight/
2. plumose/

#236. <male> right antennule actual segment 5 <size comparison of seta>/
1. of equal size/
2. of unequal size/

#237. <male> right antennule actual segment 5 <length of seta>/
1. none larger than segment/
2. one larger than segment/

#238. <male> right antennule actual segment 5 <range of the seta>/
1. surpassing to distal margin/
2. surpassing distal margin/

#239. <male> right antennule actual segment 5 <range of the seta>/
1. beyond three sequential segments/
2. not beyond three sequential segments/

#240. <male> right antennule actual segment 5 <presence of spinules>/
1. without spinules/
2. with spinules/

#241. <male> right antennule actual segment 5 <presence of vestigial seta>/
1. without vestigial seta/

2. with vestigial seta/

#242. <male> right antennule actual segment 5 <number of vestigial seta>/
1. one element/
2. two elements/

#243. <male> right antennule actual segment 5 <presence of conical seta>/
1. without conical seta/
2. with conical seta/

#244. <male> right antennule actual segment 5 <presence of modified seta>/
1. without modified seta/
2. with modified seta/

#245. <male> right antennule actual segment 5 <presence of spinous process>/
1. without spinous process/
2. with spinous process/

#246. <male> right antennule actual segment 5 <presence of aesthetasc>/
1. without aesthetasc/
2. with aesthetasc/

#247. <male> right antennule actual segment 5 <number of aesthetasc>/
1. one element/
2. two elements/

#248. <male> right antennule actual segment 6 <presence of seta>/
1. with seta/
2. without seta/

#249. <male> right antennule actual segment 6 <number of setae>/
1. one element/
2. two elements/

#250. <male> right antennule actual segment 6 <size comparison of seta>/
1. of equal size/
2. of unequal size/

#251. <male> right antennule actual segment 6 <length of seta>/
1. none larger than segment/
2. one larger than segment/

#252. <male> right antennule actual segment 6 <range of the seta>/
1. surpassing to distal margin/
2. surpassing distal margin/

#253. <male> right antennule actual segment 6 <range of the seta>/
1. beyond three sequential segments/
2. not beyond three sequential segments/

#254. <male> right antennule actual segment 6 <composition of seta>/
1. straight/
2. plumose/

#255. <male> right antennule actual segment 6 <presence of spinules>/
1. without spinules/
2. with spinules/

#256. <male> right antennule actual segment 6 <presence of vestigial seta>/
1. without vestigial seta/
2. with vestigial seta/

#257. <male> right antennule actual segment 6 <quantity of vestigial seta>/
1. 1 element/
2. 2 elements/

#258. <male> right antennule actual segment 6 <presence of conical seta>/
1. without conical seta/
2. with conical seta/

#259. <male> right antennule actual segment 6 <presence of modified seta>/
1. without modified seta/
2. with modified seta/

#260. <male> right antennule actual segment 6 <presence of spinous process>/
1. without spinous process/
2. with spinous process/

#261. <male> right antennule actual segment 6 <presence of aesthetasc>/
1. without aesthetasc/
2. with aesthetasc/

#262. <male> right antennule actual segment 6 <number of aesthetasc>/
1. one element/
2. two elements/

#263. <male> right antennule actual segment 7 <presence of seta>/
1. without seta/
2. with seta/

#264. <male> right antennule actual segment 7 <number of setae>/
1. one element/
2. two elements/

#265. <male> right antennule actual segment 7 <size comparison of seta>/
1. of equal size/
2. of unequal size/

#266. <male> right antennule actual segment 7 <composition of seta>/
1. straight/
2. plumose/

#267. <male> right antennule actual segment 7 <length of seta>/
1. none larger than segment/
2. one larger than segment/

#268. <male> right antennule actual segment 7 <range of the seta>/
1. surpassing distal margin/
2. surpassing to distal margin/

#269. <male> right antennule actual segment 7 <range of the seta>/
1. not beyond three sequential segments/
2. beyond three sequential segments/

#270. <male> right antennule actual segment 7 <apex of the seta>/
1. blunt apex/
2. sharp apex/

#271. <male> right antennule actual segment 7 <presence of spinules>/
1. without spinules/
2. with spinules/

#272. <male> right antennule actual segment 7 <presence of vestigial seta>/
1. without vestigial seta/
2. with vestigial seta/

#273. <male> right antennule actual segment 7 <quantity of vestigial seta>/
1. 1 element/
2. 2 elements/

#274. <male> right antennule actual segment 7 <presence of conical seta>/
1. without conical seta/
2. with conical seta/

#275. <male> right antennule actual segment 7 <presence of modified seta>/
1. without modified seta/
2. with modified seta/

#276. <male> right antennule actual segment 7 <presence of spinous process>/
1. without spinous process/
2. with spinous process/

#277. <male> right antennule actual segment 7 <presence of aesthetasc>/
1. without aesthetasc/
2. with aesthetasc/

#278. <male> right antennule actual segment 7 <number of aesthetasc>/

1. one element/
2. two elements/

#279. <male> right antennule actual segment 8 <presence of seta>/
1. without seta/
2. with seta/

#280. <male> right antennule actual segment 8 <number of setae>/
1. one element/
2. two elements/

#281. <male> right antennule actual segment 8 <size comparison of seta>/
1. of equal size/
2. of unequal size/

#282. <male> right antennule actual segment 8 <composition of seta>/
1. straight/
2. plumose/

#283. <male> right antennule actual segment 8 <length of seta>/
1. one larger than segment/
2. none larger than segment/

#284. <male> right antennule actual segment 8 <range of the seta>/
1. surpassing to distal margin/
2. surpassing distal margin/

#285. <male> right antennule actual segment 8 <range of the seta>/
1. beyond three sequential segments/
2. not beyond three sequential segments/

#286. <male> right antennule actual segment 8 <presence of spinules>/
1. without spinules/
2. with spinules/

#287. <male> right antennule actual segment 8 <presence of vestigial seta>/
1. without vestigial seta/
2. with vestigial seta/

#288. <male> right antennule actual segment 8 <presence of conical seta>/
1. without conical seta/
2. with conical seta/

#289. <male> right antennule actual segment 8 <number of conical setae>/
1. one element/
2. two elements/

#290. <male> right antennule actual segment 8 <reaching of the conical seta>/
1. not reaching to middle-point of the sequent segment/

2. reaching to middle-point of the sequent segment/

#291. <male> right antennule actual segment 8 <presence of modified seta>/
1. without modified seta/
2. with modified seta/

#292. <male> right antennule actual segment 8 <presence of spinous process>/
1. without spinous process/
2. with spinous process/

#293. <male> right antennule actual segment 8 <relative size of the spinous process>/
1. not longer than the original segment/
2. longer than the original segment/

#294. <male> right antennule actual segment 8 <presence of aesthetasc>/
1. without aesthetasc/
2. with aesthetasc/

#295. <male> right antennule actual segment 8 <number of aesthetasc>/
1. one element/
2. two elements/

#296. <male> right antennule actual segment 9 <presence of seta>/
1. with seta/
2. without seta/

#297. <male> right antennule actual segment 9 <number of setae>/
1. one element/
2. two elements/

#298. <male> right antennule actual segment 9 <size comparison of seta>/
1. of equal size/
2. of unequal size/

#299. <male> right antennule actual segment 9 <composition of seta>/
1. straight/
2. plumose/

#300. <male> right antennule actual segment 9 <length of seta>/
1. one larger than segment/
2. none larger than segment/

#301. <male> right antennule actual segment 9 <range of the seta>/
1. surpassing to distal margin/
2. surpassing distal margin/

#302. <male> right antennule actual segment 9 <range of the seta>/
1. beyond three sequential segments/

2. not beyond three sequential segments/

#303. <male> right antennule actual segment 9 <apex of the seta>/
1. blunt apex/
2. sharp apex/

#304. <male> right antennule actual segment 9 <presence of spinules>/
1. without spinules/
2. with spinules/

#305. <male> right antennule actual segment 9 <presence of vestigial seta>/
1. without vestigial seta/
2. with vestigial seta/

#306. <male> right antennule actual segment 9 <quantity of vestigial seta>/
1. 1 element/
2. 2 elements/

#307. <male> right antennule actual segment 9 <presence of conical seta>/
1. without conical seta/
2. with conical seta/

#308. <male> right antennule actual segment 9 <presence of modified seta>/
1. without modified seta/
2. with modified seta/

#309. <male> right antennule actual segment 9 <presence of spinous process>/
1. without spinous process/
2. with spinous process/

#310. <male> right antennule actual segment 9 <presence of aesthetasc>/
1. without aesthetasc/
2. with aesthetasc/

#311. <male> right antennule actual segment 9 <number of aesthetasc>/
1. one element/
2. two elements/

#312. <male> right antennule actual segment 10 <presence of seta>/
1. without seta/
2. with seta/

#313. <male> right antennule actual segment 10 <number of setae>/
1. one element/
2. two elements/

#314. <male> right antennule actual segment 10 <size comparison of seta>/
1. of equal size/
2. of unequal size/

#315. <male> right antennule actual segment 10 <composition of seta>/
1. straight/
2. plumose/

#316. <male> right antennule actual segment 10 <length of seta>/
1. one larger than segment/
2. none larger than segment/

#317. <male> right antennule actual segment 10 <range of the seta>/
1. surpassing to distal margin/
2. surpassing distal margin/

#318. <male> right antennule actual segment 10 <range of the seta>/
1. beyond three sequential segments/
2. not beyond three sequential segments/

#319. <male> right antennule actual segment 10 <presence of spinules>/
1. without spinules/
2. with spinules/

#320. <male> right antennule actual segment 10 <presence of vestigial seta>/
1. without vestigial seta/
2. with vestigial seta/

#321. <male> right antennule actual segment 10 <quantity of vestigial seta>/
1. 1 element/
2. 2 elements/

#322. <male> right antennule actual segment 10 <presence of conical seta>/
1. without conical seta/
2. with conical seta/

#323. <male> right antennule actual segment 10 <presence of modified seta>/
1. without modified seta/
2. with modified seta/

#324. <male> right antennule actual segment 10 <form of the apex of the modified seta>/
1. presenting blunt apex/
2. presenting bifid apex/

#325. <male> right antennule actual segment 10 <modified seta form>/
1. slender form/
2. stout form/

#326. <male> right antennule actual segment 10 <range of the modified seta>/
1. surpassing to distal margin/
2. surpassing distal margin/

#327. <male> right antennule actual segment 10 <range of the modified seta>/
1. beyond of the sequential segment/
2. not beyond of the sequential segment/

#328. <male> right antennule actual segment 10 <position of the modified seta>/
1. parallel to antennule direction/
2. perpendicular to antennule direction/

#329. <male> right antennule actual segment 10 <presence of spinous process>/
1. without spinous process/
2. with spinous process/

#330. <male> right antennule actual segment 10 <relative size of the spinous process>/
1. not longer than the original segment/
2. longer than the original segment/

#331. <male> right antennule actual segment 10 <presence of aesthetasc>/
1. without aesthetasc/
2. with aesthetasc/

#332. <male> right antennule actual segment 10 <number of aesthetasc>/
1. one element/
2. two elements/

#333. <male> right antennule actual segment 11 <presence of seta>/
1. with seta/
2. without seta/

#334. <male> right antennule actual segment 11 <number of setae>/
1. one element/
2. two elements/

#335. <male> right antennule actual segment 11 <size comparison of seta>/
1. of equal size/
2. of unequal size/

#336. <male> right antennule actual segment 11 <composition of seta>/
1. straight/
2. plumose/

#337. <male> right antennule actual segment 11 <length of seta>/
1. none larger than segment/
2. one larger than segment/

#338. <male> right antennule actual segment 11 <range of the seta>/
1. surpassing to distal margin/
2. surpassing distal margin/

#339. <male> right antennule actual segment 11 <range of the seta>/
1. beyond three sequential segments/
2. not beyond three sequential segments/

#340. <male> right antennule actual segment 11 <presence of spinules>/
1. without spinules/
2. with spinules/

#341. <male> right antennule actual segment 11 <presence of vestigial seta>/
1. without vestigial seta/
2. with vestigial seta/

#342. <male> right antennule actual segment 11 <quantity of vestigial seta>/
1. 1 element/
2. 2 elements/

#343. <male> right antennule actual segment 11 <presence of conical seta>/
1. without conical seta/
2. with conical seta/

#344. <male> right antennule actual segment 11 <presence of modified seta>/
1. without modified seta/
2. with modified seta/

#345. <male> right antennule actual segment 11 <modified seta form>/
1. slender form/
2. stout form/

#346. <male> right antennule actual segment 11 <form of the apex of the modified seta>/
1. presenting blunt apex/
2. presenting bifid apex/

#347. <male> right antennule actual segment 11 <range of the modified seta>/
1. surpassing distal margin/
2. surpassing to distal margin/

#348. <male> right antennule actual segment 11 <range of the modified seta>/
1. beyond of the sequential segment/
2. not beyond of the sequential segment/

#349. <male> right antennule actual segment 11 <position of the modified seta>/
1. parallel to antennule direction/
2. perpendicular to antennule direction/

#350. <male> right antennule actual segment 11 <relative length of the modified seta>/
1. shorter length than homologous of segment 13/

2. greater length or equal than homologous of segment 13/

#351. <male> right antennule actual segment 11 <presence of spinous process>/
1. without spinous process/
2. with spinous process/

#352. <male> right antennule actual segment 11 <relative size of the spinous process>/
1. not longer than the original segment/
2. longer than the original segment/

#353. <male> right antennule actual segment 11 <presence of aesthetasc>/
1. without aesthetasc/
2. with aesthetasc/

#354. <male> right antennule actual segment 11 <number of aesthetasc>/
1. one element/
2. two elements/

#355. <male> right antennule actual segment 12 <presence of seta>/
1. without seta/
2. with seta/

#356. <male> right antennule actual segment 12 <number of setae>/
1. one element/
2. two elements/

#357. <male> right antennule actual segment 12 <size comparison of seta>/
1. of equal size/
2. of unequal size/

#358. <male> right antennule actual segment 12 <composition of seta>/
1. straight/
2. plumose/

#359. <male> right antennule actual segment 12 <length of seta>/
1. none larger than segment/
2. one larger than segment/

#360. <male> right antennule actual segment 12 <range of the seta>/
1. surpassing to distal margin/
2. surpassing distal margin/

#361. <male> right antennule actual segment 12 <range of the seta>/
1. beyond three sequential segments/
2. not beyond three sequential segments/

#362. <male> right antennule actual segment 12 <presence of spinules>/
1. without spinules/

2. with spinules/

#363. <male> right antennule actual segment 12 <presence of vestigial seta>/
1. without vestigial seta/
2. with vestigial seta/

#364. <male> right antennule actual segment 12 <quantity of vestigial seta>/
1. 1 element/
2. 2 elements/

#365. <male> right antennule actual segment 12 <presence of conical seta>/
1. without conical seta/
2. with conical seta/

#366. <male> right antennule actual segment 12 <number of conical setae>/
1. one element/
2. two elements/

#367. <male> right antennule actual segment 12 <size comparison of the conical seta to segment 8>/
1. smaller than to segment 8/
2. not smaller than to segment 8/

#368. <male> right antennule actual segment 12 <presence of modified seta>/
1. without modified seta/
2. with modified seta/

#369. <male> right antennule actual segment 12 <presence of spinous process>/
1. without spinous process/
2. with spinous process/

#370. <male> right antennule actual segment 12 <relative size of the spinous process>/
1. not longer than the original segment/
2. longer than the original segment/

#371. <male> right antennule actual segment 12 <presence of aesthetasc>/
1. without aesthetasc/
2. with aesthetasc/

#372. <male> right antennule actual segment 12 <number of aesthetasc>/
1. one element/
2. two elements/

#373. <male> right antennule actual segment 12 <observation of internal perpendicular fission>/
1. absent internal perpendicular fission/
2. present internal perpendicular fission/

#374. <male> right antennule actual segment 13 <presence of seta>/
1. without seta/
2. with seta/

#375. <male> right antennule actual segment 13 <number of setae>/
1. one element/
2. two elements/

#376. <male> right antennule actual segment 13 <size comparison of seta>/
1. of equal size/
2. of unequal size/

#377. <male> right antennule actual segment 13 <composition of seta>/
1. straight/
2. plumose/

#378. <male> right antennule actual segment 13 <length of seta>/
1. none larger than segment/
2. one larger than segment/

#379. <male> right antennule actual segment 13 <range of the seta>/
1. surpassing to distal margin/
2. surpassing distal margin/

#380. <male> right antennule actual segment 13 <range of the seta>/
1. beyond three sequential segments/
2. not beyond three sequential segments/

#381. <male> right antennule actual segment 13 <presence of spinules>/
1. without spinules/
2. with spinules/

#382. <male> right antennule actual segment 13 <presence of vestigial seta>/
1. without vestigial seta/
2. with vestigial seta/

#383. <male> right antennule actual segment 13 <quantity of vestigial seta>/
1. 1 element/
2. 2 elements/

#384. <male> right antennule actual segment 13 <presence of conical seta>/
1. without conical seta/
2. with conical seta/

#385. <male> right antennule actual segment 13 <presence of modified seta>/
1. without modified seta/
2. with modified seta/

#386. <male> right antennule actual segment 13 <modified seta form>/

1. slender form/
2. stout form/

#387. <male> right antennule actual segment 13 <range of the modified seta>/
1. surpassing to distal margin/
2. surpassing distal margin/

#388. <male> right antennule actual segment 13 <range of the modified seta>/
1. to the middle-point of the sequence segment/
2. to the distal-point of the sequence segment/

#389. <male> right antennule actual segment 13 <position of the modified seta>/
1. parallel to antennule direction/
2. perpendicular to antennule direction/

#390. <male> right antennule actual segment 13 <form of the apex of the modified seta>/
1. presenting rounded apex/
2. presenting bifid apex/

#391. <male> right antennule actual segment 13 <presence of spinous process>/
1. without spinous process/
2. with spinous process/

#392. <male> right antennule actual segment 13 <relative size of the spinous process>/
1. not longer than the original segment/
2. longer than the original segment/

#393. <male> right antennule actual segment 13 <presence of aesthetasc>/
1. without aesthetasc/
2. with aesthetasc/

#394. <male> right antennule actual segment 13 <number of aesthetasc>/
1. one element/
2. two elements/

#395. <male> right antennule actual segment 14 <presence of seta>/
1. with seta/
2. without seta/

#396. <male> right antennule actual segment 14 <number of setae>/
1. one element/
2. two elements/

#397. <male> right antennule actual segment 14 <size comparison of seta>/
1. of equal size/
2. of unequal size/

#398. <male> right antennule actual segment 14 <composition of seta>/
1. straight/
2. plumose/

#399. <male> right antennule actual segment 14 <length of seta>/
1. none larger than segment/
2. one larger than segment/

#400. <male> right antennule actual segment 14 <range of the seta>/
1. surpassing to distal margin/
2. surpassing distal margin/

#401. <male> right antennule actual segment 14 <range of the seta>/
1. beyond three sequential segments/
2. not beyond three sequential segments/

#402. <male> right antennule actual segment 14 <apex of the seta>/
1. blunt apex/
2. sharp apex/

#403. <male> right antennule actual segment 14 <presence of spinules>/
1. without spinules/
2. with spinules/

#404. <male> right antennule actual segment 14 <presence of vestigial seta>/
1. without vestigial seta/
2. with vestigial seta/

#405. <male> right antennule actual segment 14 <quantity of vestigial seta>/
1. 1 element/
2. 2 elements/

#406. <male> right antennule actual segment 14 <presence of conical seta>/
1. without conical seta/
2. with conical seta/

#407. <male> right antennule actual segment 14 <presence of modified seta>/
1. without modified seta/
2. with modified seta/

#408. <male> right antennule actual segment 14 <presence of spinous process>/
1. without spinous process/
2. with spinous process/

#409. <male> right antennule actual segment 14 <presence of aesthetasc>/
1. without aesthetasc/
2. with aesthetasc/

#410. <male> right antennule actual segment 14 <number of aesthetasc>/

1. one element/
2. two elements/

#411. <male> right antennule actual segment 15 <presence of seta>/
1. without seta/
2. with seta/

#412. <male> right antennule actual segment 15 <number of setae>/
1. one element/
2. two elements/

#413. <male> right antennule actual segment 15 <size comparison of seta>/
1. of equal size/
2. of unequal size/

#414. <male> right antennule actual segment 15 <composition of seta>/
1. straight/
2. plumose/

#415. <male> right antennule actual segment 15 <form of the seta>/
1. bifidform/
2. not bifidform/

#416. <male> right antennule actual segment 15 <length of seta>/
1. none larger than segment/
2. one larger than segment/

#417. <male> right antennule actual segment 15 <range of the seta>/
1. surpassing to distal margin/
2. surpassing distal margin/

#418. <male> right antennule actual segment 15 <range of the seta>/
1. beyond three sequential segments/
2. not beyond three sequential segments/

#419. <male> right antennule actual segment 15 <presence of spinules>/
1. without spinules/
2. with spinules/

#420. <male> right antennule actual segment 15 <presence of vestigial seta>/
1. without vestigial seta/
2. with vestigial seta/

#421. <male> right antennule actual segment 15 <quantity of vestigial seta>/
1. 1 element/
2. 2 elements/

#422. <male> right antennule actual segment 15 <presence of conical seta>/
1. without conical seta/

2. with conical seta/

#423. <male> right antennule actual segment 15 <presence of modified seta>/
1. without modified seta/
2. with modified seta/

#424. <male> right antennule actual segment 15 <presence of spinous process>/
1. without spinous process/
2. with spinous process/

#425. <male> right antennule actual segment 15 <position of spinous process>/
1. on inner margin/
2. on outer margin/

#426. <male> right antennule actual segment 15 <size comparison with preceding segments>/
1. surpassing distal margin/
2. surpassing to distal margin/

#427. <male> right antennule actual segment 15 <presence of aesthetasc>/
1. without aesthetasc/
2. with aesthetasc/

#428. <male> right antennule actual segment 15 <number of aesthetasc>/
1. one element/
2. two elements/

#429. <male> right antennule actual segment 16 <presence of seta>/
1. without seta/
2. with seta/

#430. <male> right antennule actual segment 16 <number of setae>/
1. one element/
2. two elements/

#431. <male> right antennule actual segment 16 <size comparison of seta>/
1. of equal size/
2. of unequal size/

#432. <male> right antennule actual segment 16 <composition of seta>/
1. straight/
2. plumose/

#433. <male> right antennule actual segment 16 <length of seta>/
1. none larger than segment/
2. one larger than segment/

#434. <male> right antennule actual segment 16 <range of the seta>/
1. surpassing to distal margin/

2. surpassing distal margin/

#435. <male> right antennule actual segment 16 <range of the seta>/
1. beyond three sequential segments/
2. not beyond three sequential segments/

#436. <male> right antennule actual segment 16 <form of the seta>/
1. bifidform/
2. not bifidform/

#437. <male> right antennule actual segment 16 <presence of spinules>/
1. without spinules/
2. with spinules/

#438. <male> right antennule actual segment 16 <presence of vestigial seta>/
1. without vestigial seta/
2. with vestigial seta/

#439. <male> right antennule actual segment 16 <quantity of vestigial seta>/
1. 1 element/
2. 2 elements/

#440. <male> right antennule actual segment 16 <presence of conical seta>/
1. without conical seta/
2. with conical seta/

#441. <male> right antennule actual segment 16 <presence of modified seta>/
1. without modified seta/
2. with modified seta/

#442. <male> right antennule actual segment 16 <presence of spinous process>/
1. without spinous process/
2. with spinous process/

#443. <male> right antennule actual segment 16 <position of spinous process>/
1. on inner margin/
2. on outer margin/

#444. <male> right antennule actual segment 16 <size comparison with preceding segments>/
1. surpassing to distal margin/
2. surpassing distal margin/

#445. <male> right antennule actual segment 16 <presence of aesthetasc>/
1. without aesthetasc/
2. with aesthetasc/

#446. <male> right antennule actual segment 16 <number of aesthetasc>/
1. one element/

2. two elements/

#447. <male> right antennule actual segment 17 <presence of seta>/
1. with seta/
2. without seta/

#448. <male> right antennule actual segment 17 <number of setae>/
1. one element/
2. two elements/

#449. <male> right antennule actual segment 17 <size comparison of seta>/
1. of equal size/
2. of unequal size/

#450. <male> right antennule actual segment 17 <composition of seta>/
1. straight/
2. plumose/

#451. <male> right antennule actual segment 17 <length of seta>/
1. one larger than segment/
2. none larger than segment/

#452. <male> right antennule actual segment 17 <range of the seta>/
1. surpassing to distal margin/
2. surpassing distal margin/

#453. <male> right antennule actual segment 17 <range of the seta>/
1. beyond three sequential segments/
2. not beyond three sequential segments/

#454. <male> right antennule actual segment 17 <form of the seta>/
1. bifidform/
2. not bifidform/

#455. <male> right antennule actual segment 17 <presence of spinules>/
1. without spinules/
2. with spinules/

#456. <male> right antennule actual segment 17 <presence of vestigial seta>/
1. without vestigial seta/
2. with vestigial seta/

#457. <male> right antennule actual segment 17 <quantity of vestigial seta>/
1. 1 element/
2. 2 elements/

#458. <male> right antennule actual segment 17 <presence of conical seta>/
1. without conical seta/
2. with conical seta/

#459. <male> right antennule actual segment 17 <presence of modified seta>/
1. without modified seta/
2. with modified seta/

#460. <male> right antennule actual segment 17 <number of modified seta>/
1. one element/
2. two elements/

#461. <male> right antennule actual segment 17 <form of the modified seta>/
1. stout form/
2. slender form/

#462. <male> right antennule actual segment 17 <range of the modified seta>/
1. surpassing to distal margin/
2. surpassing distal margin/

#463. <male> right antennule actual segment 17 <range of the modified seta>/
1. beyond of the sequential segment/
2. not beyond of the sequential segment/

#464. <male> right antennule actual segment 17 <position of the modified seta>/
1. parallel to antennule direction/
2. perpendicular to antennule direction/

#465. <male> right antennule actual segment 17 <presence of spinous process>/
1. without spinous process/
2. with spinous process/

#466. <male> right antennule actual segment 17 <presence of aesthetasc>/
1. without aesthetasc/
2. with aesthetasc/

#467. <male> right antennule actual segment 17 <number of aesthetasc>/
1. one element/
2. two elements/

#468. <male> right antennule actual segment 18 <presence of seta>/
1. with seta/
2. without seta/

#469. <male> right antennule actual segment 18 <number of setae>/
1. one element/
2. two elements/

#470. <male> right antennule actual segment 18 <size comparison of seta>/
1. of equal size/
2. of unequal size/

#471. <male> right antennule actual segment 18 <composition of seta>/
1. straight/
2. plumose/

#472. <male> right antennule actual segment 18 <length of seta>/
1. none larger than segment/
2. one larger than segment/

#473. <male> right antennule actual segment 18 <range of the seta>/
1. surpassing to distal margin/
2. surpassing distal margin/

#474. <male> right antennule actual segment 18 <range of the seta>/
1. beyond three sequential segments/
2. not beyond three sequential segments/

#475. <male> right antennule actual segment 18 <presence of spinules>/
1. without spinules/
2. with spinules/

#476. <male> right antennule actual segment 18 <presence of vestigial seta>/
1. without vestigial seta/
2. with vestigial seta/

#477. <male> right antennule actual segment 18 <quantity of vestigial seta>/
1. 1 element/
2. 2 elements/

#478. <male> right antennule actual segment 18 <presence of conical seta>/
1. without conical seta/
2. with conical seta/

#479. <male> right antennule actual segment 18 <presence of modified seta>/
1. without modified seta/
2. with modified seta/

#480. <male> right antennule actual segment 18 <number of modified seta>/
1. one element/
2. two elements/

#481. <male> right antennule actual segment 18 <form of the modified seta>/
1. stout form/
2. slender form/

#482. <male> right antennule actual segment 18 <range of the modified seta>/
1. surpassing to distal margin/
2. surpassing distal margin/

#483. <male> right antennule actual segment 18 <range of the modified seta>/

1. beyond of the sequential segment/
2. not beyond of the sequential segment/

#484. <male> right antennule actual segment 18 <position of the modified seta>/
1. parallel to antennule direction/
2. perpendicular to antennule direction/

#485. <male> right antennule actual segment 18 <presence of spinous process>/
1. without spinous process/
2. with spinous process/

#486. <male> right antennule actual segment 18 <presence of aesthetasc>/
1. without aesthetasc/
2. with aesthetasc/

#487. <male> right antennule actual segment 18 <number of aesthetasc>/
1. one element/
2. two elements/

#488. <male> right antennule actual segment 19 <presence of seta>/
1. without seta/
2. with seta/

#489. <male> right antennule actual segment 19 <number of setae>/
1. one element/
2. two elements/

#490. <male> right antennule actual segment 19 <size comparison of seta>/
1. of equal size/
2. of unequal size/

#491. <male> right antennule actual segment 19 <composition of seta>/
1. straight/
2. plumose/

#492. <male> right antennule actual segment 19 <length of seta>/
1. none larger than segment/
2. one larger than segment/

#493. <male> right antennule actual segment 19 <range of the seta>/
1. surpassing to distal margin/
2. surpassing distal margin/

#494. <male> right antennule actual segment 19 <range of the seta>/
1. beyond three sequential segments/
2. not beyond three sequential segments/

#495. <male> right antennule actual segment 19 <presence of spinules>/
1. without spinules/

2. with spinules/

#496. <male> right antennule actual segment 19 <presence of vestigial seta>/
1. without vestigial seta/
2. with vestigial seta/

#497. <male> right antennule actual segment 19 <quantity of vestigial seta>/
1. 1 element/
2. 2 elements/

#498. <male> right antennule actual segment 19 <presence of conical seta>/
1. without conical seta/
2. with conical seta/

#499. <male> right antennule actual segment 19 <presence of modified seta>/
1. without modified seta/
2. with modified seta/

#500. <male> right antennule actual segment 19 <number of modified seta>/
1. one element/
2. two elements/

#501. <male> right antennule actual segment 19 <form of the modified seta>/
1. stout form/
2. slender form/

#502. <male> right antennule actual segment 19 <type of the modified seta>/
1. at least one bifid form/
2. none in bifid form/

#503. <male> right antennule actual segment 19 <range of the modified seta>/
1. surpassing distal margin/
2. surpassing to distal margin/

#504. <male> right antennule actual segment 19 <range of the modified seta>/
1. not beyond of the sequential segment/
2. beyond of the sequential segment/

#505. <male> right antennule actual segment 19 <position of the modified seta>/
1. parallel to antennule direction/
2. perpendicular to antennule direction/

#506. <male> right antennule actual segment 19 <presence of spinous process>/
1. without spinous process/
2. with spinous process/

#507. <male> right antennule actual segment 19 <presence of aesthetasc>/
1. without aesthetasc/
2. with aesthetasc/

#508. <male> right antennule actual segment 19 <number of aesthetasc>/
1. one element/
2. two elements/

#509. <male> right antennule actual segment 20 <presence of seta>/
1. with seta/
2. without seta/

#510. <male> right antennule actual segment 20 <number of setae>/
1. two elements/
2. four elements/

#511. <male> right antennule actual segment 20 <size comparison of seta>/
1. of equal size/
2. of unequal size/

#512. <male> right antennule actual segment 20 <composition of seta>/
1. straight/
2. plumose/

#513. <male> right antennule actual segment 20 <length of seta>/
1. one larger than segment/
2. none larger than segment/

#514. <male> right antennule actual segment 20 <range of the seta>/
1. surpassing to distal margin/
2. surpassing distal margin/

#515. <male> right antennule actual segment 20 <range of the seta>/
1. beyond three sequential segments/
2. not beyond three sequential segments/

#516. <male> right antennule actual segment 20 <presence of spinules>/
1. without spinules/
2. with spinules/

#517. <male> right antennule actual segment 20 <presence of vestigial seta>/
1. without vestigial seta/
2. with vestigial seta/

#518. <male> right antennule actual segment 20 <presence of conical seta>/
1. without conical seta/
2. with conical seta/

#519. <male> right antennule actual segment 20 <presence of modified seta>/
1. without modified seta/
2. with modified seta/

#520. <male> right antennule actual segment 20 <presence of spinous process>/
1. without spinous process/
2. with spinous process/

#521. <male> right antennule actual segment 20 <location of the spinous process>/
1. medially/
2. distally/

#522. <male> right antennule actual segment 20 <reach of the spinous process>/
1. reaching beyond of distal-point segment 21/
2. not reaching beyond of distal-point segment 21/

#523. <male> right antennule actual segment 20 <presence of aesthetasc>/
1. without aesthetasc/
2. with aesthetasc/

#524. <male> right antennule actual segment 20 <number of aesthetasc>/
1. one element/
2. two elements/

#525. <male> right antennule actual segment 21 <presence of seta>/
1. with seta/
2. without seta/

#526. <male> right antennule actual segment 21 <number of setae>/
1. two elements/
2. four elements/

#527. <male> right antennule actual segment 21 <size comparison of seta>/
1. of equal size/
2. of unequal size/

#528. <male> right antennule actual segment 21 <composition of seta>/
1. straight/
2. plumose/

#529. <male> right antennule actual segment 21 <length of seta>/
1. one larger than segment/
2. none larger than segment/

#530. <male> right antennule actual segment 21 <range of the seta>/
1. surpassing to distal margin/
2. surpassing distal margin/

#531. <male> right antennule actual segment 21 <size of the seta>/
1. greater 3x than original segment/
2. not greater 3x than original segment/

#532. <male> right antennule actual segment 21 <presence of spinules>/
1. without spinules/
2. with spinules/

#533. <male> right antennule actual segment 21 <presence of vestigial seta>/
1. without vestigial seta/
2. with vestigial seta/

#534. <male> right antennule actual segment 21 <presence of conical seta>/
1. without conical seta/
2. with conical seta/

#535. <male> right antennule actual segment 21 <presence of modified seta>/
1. without modified seta/
2. with modified seta/

#536. <male> right antennule actual segment 21 <presence of spinous process>/
1. without spinous process/
2. with spinous process/

#537. <male> right antennule actual segment 21 <presence of aesthetasc>/
1. without aesthetasc/
2. with aesthetasc/

#538. <male> right antennule actual segment 21 <number of aesthetasc>/
1. one element/
2. two elements/

#539. <male> right antennule actual segment 22 <presence of seta>/
1. with seta/
2. without seta/

#540. <male> right antennule actual segment 22 <number of setae>/
1. four elements/
2. five elements/

#541. <male> right antennule actual segment 22 <size comparison of seta>/
1. of equal size/
2. of unequal size/

#542. <male> right antennule actual segment 22 <length of seta>/
1. one larger than segment/
2. none larger than segment/

#543. <male> right antennule actual segment 22 <composition of seta>/
1. straight/
2. plumose/

#544. <male> right antennule actual segment 22 <range of the seta>/

1. surpassing to distal margin/
2. surpassing distal margin/

#545. <male> right antennule actual segment 22 <size of the seta>/
1. greater 3x than original segment/
2. not greater 3x than original segment/

#546. <male> right antennule actual segment 22 <presence of spinules>/
1. without spinules/
2. with spinules/

#547. <male> right antennule actual segment 22 <presence of vestigial seta>/
1. without vestigial seta/
2. with vestigial seta/

#548. <male> right antennule actual segment 22 <presence of conical seta>/
1. without conical seta/
2. with conical seta/

#549. <male> right antennule actual segment 22 <presence of modified seta>/
1. without modified seta/
2. with modified seta/

#550. <male> right antennule actual segment 22 <presence of spinous process>/
1. without spinous process/
2. with spinous process/

#551. <male> right antennule actual segment 22 <presence of aesthetasc>/
1. without aesthetasc/
2. with aesthetasc/

#552. <male> right antennule actual segment 22 <number of aesthetasc>/
1. one element/
2. two elements/

#553. Left antennule <number of rami>/
1. uniramous/
2. biramous/

#554. Left antennule <relative length>/
1. surpassing to prosoma/
2. not surpassing to prosoma/

#555. Left antennule <relative length>/
1. extending beyond caudal setae/
2. not extending beyond caudal setae/

#556. Left antennule ancestral segment I and II <fusion>/
1. separated/

2. fused/

#557. Left antennule ancestral segment II and III <fusion>/
1. separated/
2. fused/

#558. Left antennule ancestral segment III and IV <fusion>/
1. separated/
2. fused/

#559. Left antennule ancestral segment IV and V <fusion>/
1. separated/
2. fused/

#560. Left antennule ancestral segment V and VI <fusion>/
1. separated/
2. fusion/

#561. Left antennule ancestral segment VI and VII <fusion>/
1. separated/
2. fusion/

#562. Left antennule ancestral segment VII and VIII <fusion>/
1. separated/
2. fused/

#563. Left antennule ancestral segment VIII and IX <fusion>/
1. separated/
2. fused/

#564. Left antennule ancestral segment IX and X <fusion>/
1. separated/
2. fused/

#565. Left antennule ancestral segment X and XI <fusion>/
1. separated/
2. fused/

#566. Left antennule ancestral segment XI and XII <fusion>/
1. separated/
2. fused/

#567. Left antennule ancestral segment XII and XIII <fusion>/
1. separated/
2. fused/

#568. Left antennule ancestral segment XIII and XIV <fusion>/
1. separated/
2. fused/

#569. Left antennule ancestral segment XIV and XV <fusion>/
1. separated/
2. fused/

#570. Left antennule ancestral segment XV and XVI <fusion>/
1. separated/
2. fused/

#571. Left antennule ancestral segment XVI and XVII <fusion>/
1. separated/
2. fused/

#572. Left antennule ancestral segment XVII and XVIII <fusion>/
1. separated/
2. fused/

#573. Left antennule ancestral segment XVIII and XIX <fusion>/
1. separated/
2. fused/

#574. Left antennule ancestral segment XIX and XX <fusion>/
1. separated/
2. fused/

#575. Left antennule ancestral segment XX and XXI <fusion>/
1. separated/
2. fused/

#576. Left antennule ancestral segment XXI and XXII <fusion>/
1. separated/
2. fused/

#577. Left antennule ancestral segment XXII and XXIII <fusion>/
1. separated/
2. fused/

#578. Left antennule ancestral segment XXIII and XXIV <fusion>/
1. separated/
2. fused/

#579. Left antennule ancestral segment XXIV and XXV <fusion>/
1. separated/
2. fused/

#580. Left antennule ancestral segment XXV and XXVI <fusion>/
1. separated/
2. fused/

#581. Left antennule ancestral segment XXVI and XXVII <fusion>/
1. separated/
2. fused/

#582. Left antennule ancestral segment XXVII and XXVIII <fusion>/
1. separated/
2. fused/

#583. Left antennule actual segment <number of segments>/
1. 22-segmented/
2. 25-segmented/

#584. Left antennule actual segment <geniculation>/
1. geniculated/
2. not-geniculated/

#585. Left antennule actual segment 1 <presence of seta>/
1. with seta/
2. without seta/

#586. Left antennule actual segment 1 <number of setae>/
1. one element/
2. two elements/

#587. Left antennule actual segment 1 <size comparison of seta>/
1. of equal size/
2. of unequal size/

#588. Left antennule actual segment 1 <length of seta>/
1. one larger than segment/
2. none larger than segment/

#589. Left antennule actual segment 1 <range of the seta>/
1. surpassing to distal margin/
2. surpassing distal margin/

#590. Left antennule actual segment 1 <composition of seta>/
1. straight/
2. plumose/

#591. Left antennule actual segment 1 <range of the seta>/
1. beyond three sequential segments/
2. not beyond three sequential segments/

#592. Left antennule actual segment 1 <presence of spinules>/
1. without spinules/
2. with spinules/

#593. Left antennule actual segment 1 <arrangement of spinules>/

1. as a row/
2. as a patch/

#594. Left antennule actual segment 1 <presence of vestigial seta>/
1. without vestigial seta/
2. with vestigial seta/

#595. Left antennule actual segment 1 <presence of conical seta>/
1. without conical seta/
2. with conical seta/

#596. Left antennule actual segment 1 <presence of modified seta>/
1. without modified seta/
2. with modified seta/

#597. Left antennule actual segment 1 <presence of spinous process>/
1. without spinous process/
2. with spinous process/

#598. Left antennule actual segment 1 <presence of aesthetasc>/
1. without aesthetasc/
2. with aesthetasc/

#599. Left antennule actual segment 1 <number of aesthetasc>/
1. one element/
2. two elements/

#600. Left antennule actual segment 2 <presence of seta>/
1. without seta/
2. with seta/

#601. Left antennule actual segment 2 <number of setae>/
1. three elements/
2. four elements/

#602. Left antennule actual segment 2 <size comparison of seta>/
1. of equal size/
2. of unequal size/

#603. Left antennule actual segment 2 <length of seta>/
1. one larger than segment/
2. none larger than segment/

#604. Left antennule actual segment 2 <composition of seta>/
1. straight/
2. plumose/

#605. Left antennule actual segment 2 <range of the seta>/
1. surpassing to distal margin/

2. surpassing distal margin/

#606. Left antennule actual segment 2 <range of the seta>/
1. beyond three sequential segments/
2. not beyond three sequential segments/

#607. Left antennule actual segment 2 <presence of spinules>/
1. without spinules/
2. with spinules/

#608. Left antennule actual segment 2 <presence of vestigial seta>/
1. without vestigial seta/
2. with vestigial seta/

#609. Left antennule actual segment 2 <number of vestigial seta>/
1. one element/
2. two elements/

#610. Left antennule actual segment 2 <presence of conical seta>/
1. without conical seta/
2. with conical seta/

#611. Left antennule actual segment 2 <presence of modified seta>/
1. without modified seta/
2. with modified seta/

#612. Left antennule actual segment 2 <presence of spinous process>/
1. without spinous process/
2. with spinous process/

#613. Left antennule actual segment 2 <presence of aesthetasc>/
1. without aesthetasc/
2. with aesthetasc/

#614. Left antennule actual segment 2 <number of aesthetasc>/
1. one element/
2. two elements/

#615. Left antennule actual segment 3 <presence of seta>/
1. with seta/
2. without seta/

#616. Left antennule actual segment 3 <number of setae>/
1. one element/
2. two elements/

#617. Left antennule actual segment 3 <size comparison of seta>/
1. of equal size/
2. of unequal size/

#618. Left antennule actual segment 3 <length of seta>/
1. one larger than segment/
2. none larger than segment/

#619. Left antennule actual segment 3 <composition of seta>/
1. straight/
2. plumose/

#620. Left antennule actual segment 3 <range of the seta>/
1. surpassing to distal margin/
2. surpassing distal margin/

#621. Left antennule actual segment 3 <range of the seta>/
1. beyond three sequential segments/
2. not beyond three sequential segments/

#622. Left antennule actual segment 3 <presence of spinules>/
1. without spinules/
2. with spinules/

#623. Left antennule actual segment 3 <presence of vestigial seta>/
1. without vestigial seta/
2. with vestigial seta/

#624. Left antennule actual segment 3 <number of vestigial seta>/
1. one element/
2. two elements/

#625. Left antennule actual segment 3 <presence of conical seta>/
1. without conical seta/
2. with conical seta/

#626. Left antennule actual segment 3 <presence of modified seta>/
1. without modified seta/
2. with modified seta/

#627. Left antennule actual segment 3 <presence of spinous process>/
1. without spinous process/
2. with spinous process/

#628. Left antennule actual segment 3 <presence of aesthetasc>/
1. without aesthetasc/
2. with aesthetasc/

#629. Left antennule actual segment 3 <number of aesthetasc>/
1. one element/
2. two elements/

#630. Left antennule actual segment 4 <presence of seta>/
1. with seta/
2. without seta/

#631. Left antennule actual segment 4 <number of setae>/
1. one element/
2. two elements/

#632. Left antennule actual segment 4 <size comparison of seta>/
1. of equal size/
2. of unequal size/

#633. Left antennule actual segment 4 <length of seta>/
1. none larger than segment/
2. one larger than segment/

#634. Left antennule actual segment 4 <composition of seta>/
1. straight/
2. plumose/

#635. Left antennule actual segment 4 <range of the seta>/
1. surpassing to distal margin/
2. surpassing distal margin/

#636. Left antennule actual segment 4 <range of the seta>/
1. beyond three sequential segments/
2. not beyond three sequential segments/

#637. Left antennule actual segment 4 <presence of spinules>/
1. without spinules/
2. with spinules/

#638. Left antennule actual segment 4 <presence of vestigial seta>/
1. without vestigial seta/
2. with vestigial seta/

#639. Left antennule actual segment 4 <presence of conical seta>/
1. without conical seta/
2. with conical seta/

#640. Left antennule actual segment 4 <presence of modified seta>/
1. without modified seta/
2. with modified seta/

#641. Left antennule actual segment 4 <presence of spinous process>/
1. without spinous process/
2. with spinous process/

#642. Left antennule actual segment 4 <presence of aesthetasc>/

1. without aesthetasc/
2. with aesthetasc/

#643. Left antennule actual segment 4 <number of aesthetasc>/
1. one element/
2. two elements/

#644. Left antennule actual segment 5 <presence of seta>/
1. with seta/
2. without seta/

#645. Left antennule actual segment 5 <number of setae>/
1. one element/
2. two elements/

#646. Left antennule actual segment 5 <size comparison of seta>/
1. of equal size/
2. of unequal size/

#647. Left antennule actual segment 5 <length of seta>/
1. one larger than segment/
2. none larger than segment/

#648. Left antennule actual segment 5 <composition of seta>/
1. straight/
2. plumose/

#649. Left antennule actual segment 5 <range of the seta>/
1. surpassing to distal margin/
2. surpassing distal margin/

#650. Left antennule actual segment 5 <range of the seta>/
1. beyond three sequential segments/
2. not beyond three sequential segments/

#651. Left antennule actual segment 5 <presence of spinules>/
1. without spinules/
2. with spinules/

#652. Left antennule actual segment 5 <presence of vestigial seta>/
1. without vestigial seta/
2. with vestigial seta/

#653. Left antennule actual segment 5 <number of vestigial seta>/
1. one element/
2. two elements/

#654. Left antennule actual segment 5 <presence of conical seta>/
1. without conical seta/

2. with conical seta/

#655. Left antennule actual segment 5 <presence of modified seta>/
1. without modified seta/
2. with modified seta/

#656. Left antennule actual segment 5 <presence of spinous process>/
1. without spinous process/
2. with spinous process/

#657. Left antennule actual segment 5 <presence of aesthetasc>/
1. without aesthetasc/
2. with aesthetasc/

#658. Left antennule actual segment 5 <number of aesthetasc>/
1. one element/
2. two elements/

#659. Left antennule actual segment 6 <presence of seta>/
1. with seta/
2. without seta/

#660. Left antennule actual segment 6 <number of setae>/
1. one element/
2. two elements/

#661. Left antennule actual segment 6 <size comparison of seta>/
1. of equal size/
2. of unequal size/

#662. Left antennule actual segment 6 <length of seta>/
1. one larger than segment/
2. none larger than segment/

#663. Left antennule actual segment 6 <composition of seta>/
1. straight/
2. plumose/

#664. Left antennule actual segment 6 <range of the seta>/
1. surpassing to distal margin/
2. surpassing distal margin/

#665. Left antennule actual segment 6 <range of the seta>/
1. beyond three sequential segments/
2. not beyond three sequential segments/

#666. Left antennule actual segment 6 <presence of spinules>/
1. without spinules/
2. with spinules/

#667. Left antennule actual segment 6 <presence of vestigial seta>/
    1. without vestigial seta/
    2. with vestigial seta/

#668. Left antennule actual segment 6 <presence of conical seta>/
    1. without conical seta/
    2. with conical seta/

#669. Left antennule actual segment 6 <presence of modified seta>/
    1. without modified seta/
    2. with modified seta/

#670. Left antennule actual segment 6 <presence of spinous process>/
    1. without spinous process/
    2. with spinous process/

#671. Left antennule actual segment 6 <presence of aesthetasc>/
    1. without aesthetasc/
    2. with aesthetasc/

#672. Left antennule actual segment 6 <number of aesthetasc>/
    1. one element/
    2. two elements/

#673. Left antennule actual segment 7 <presence of seta>/
    1. without seta/
    2. with seta/

#674. Left antennule actual segment 7 <number of setae>/
    1. one element/
    2. two elements/

#675. Left antennule actual segment 7 <size comparison of seta>/
    1. of equal size/
    2. of unequal size/

#676. Left antennule actual segment 7 <length of seta>/
    1. one larger than segment/
    2. none larger than segment/

#677. Left antennule actual segment 7 <composition of seta>/
    1. straight/
    2. plumose/

#678. Left antennule actual segment 7 <range of the seta>/
    1. surpassing to distal margin/
    2. surpassing distal margin/

#679. Left antennule actual segment 7 <range of the seta>/
1. beyond three sequential segments/
2. not beyond three sequential segments/

#680. Left antennule actual segment 7 <presence of spinules>/
1. without spinules/
2. with spinules/

#681. Left antennule actual segment 7 <presence of vestigial seta>/
1. without vestigial seta/
2. with vestigial seta/

#682. Left antennule actual segment 7 <presence of conical seta>/
1. without conical seta/
2. with conical seta/

#683. Left antennule actual segment 7 <presence of modified seta>/
1. without modified seta/
2. with modified seta/

#684. Left antennule actual segment 7 <presence of spinous process>/
1. without spinous process/
2. with spinous process/

#685. Left antennule actual segment 7 <presence of aesthetasc>/
1. without aesthetasc/
2. with aesthetasc/

#686. Left antennule actual segment 7 <number of aesthetasc>/
1. one element/
2. two elements/

#687. Left antennule actual segment 8 <presence of seta>/
1. with seta/
2. without seta/

#688. Left antennule actual segment 8 <number of setae>/
1. one element/
2. two elements/

#689. Left antennule actual segment 8 <size comparison of seta>/
1. of equal size/
2. of unequal size/

#690. Left antennule actual segment 8 <length of seta>/
1. one larger than segment/
2. none larger than segment/

#691. Left antennule actual segment 8 <composition of seta>/

1. straight/
2. plumose/

#692. Left antennule actual segment 8 <range of the seta>/
1. surpassing to distal margin/
2. surpassing distal margin/

#693. Left antennule actual segment 8 <range of the seta>/
1. beyond three sequential segments/
2. not beyond three sequential segments/

#694. Left antennule actual segment 8 <presence of spinules>/
1. without spinules/
2. with spinules/

#695. Left antennule actual segment 8 <presence of vestigial seta>/
1. without vestigial seta/
2. with vestigial seta/

#696. Left antennule actual segment 8 <presence of conical seta>/
1. without conical seta/
2. with conical seta/

#697. Left antennule actual segment 8 <presence of modified seta>/
1. without modified seta/
2. with modified seta/

#698. Left antennule actual segment 8 <presence of spinous process>/
1. without spinous process/
2. with spinous process/

#699. Left antennule actual segment 8 <presence of aesthetasc>/
1. without aesthetasc/
2. with aesthetasc/

#700. Left antennule actual segment 8 <number of aesthetasc>/
1. one element/
2. two elements/

#701. Left antennule actual segment 9 <presence of seta>/
1. with seta/
2. without seta/

#702. Left antennule actual segment 9 <number of setae>/
1. one element/
2. two elements/

#703. Left antennule actual segment 9 <size comparison of seta>/
1. of equal size/

2. of unequal size/

#704. Left antennule actual segment 9 <length of seta>/
1. one larger than segment/
2. none larger than segment/

#705. Left antennule actual segment 9 <composition of seta>/
1. straight/
2. plumose/

#706. Left antennule actual segment 9 <range of the seta>/
1. surpassing to distal margin/
2. surpassing distal margin/

#707. Left antennule actual segment 9 <range of the seta>/
1. beyond three sequential segments/
2. not beyond three sequential segments/

#708. Left antennule actual segment 9 <presence of spinules>/
1. without spinules/
2. with spinules/

#709. Left antennule actual segment 9 <presence of vestigial seta>/
1. without vestigial seta/
2. with vestigial seta/

#710. Left antennule actual segment 9 <presence of conical seta>/
1. without conical seta/
2. with conical seta/

#711. Left antennule actual segment 9 <presence of modified seta>/
1. without modified seta/
2. with modified seta/

#712. Left antennule actual segment 9 <presence of spinous process>/
1. without spinous process/
2. with spinous process/

#713. Left antennule actual segment 9 <presence of aesthetasc>/
1. without aesthetasc/
2. with aesthetasc/

#714. Left antennule actual segment 9 <number of aesthetasc>/
1. one element/
2. two elements/

#715. Left antennule actual segment 10 <presence of seta>/
1. with seta/
2. without seta/

#716. Left antennule actual segment 10 <number of setae>/
1. one element/
2. two elements/

#717. Left antennule actual segment 10 <size comparison of seta>/
1. of equal size/
2. of unequal size/

#718. Left antennule actual segment 10 <length of seta>/
1. one larger than segment/
2. none larger than segment/

#719. Left antennule actual segment 10 <composition of seta>/
1. straight/
2. plumose/

#720. Left antennule actual segment 10 <range of the seta>/
1. surpassing to distal margin/
2. surpassing distal margin/

#721. Left antennule actual segment 10 <range of the seta>/
1. beyond three sequential segments/
2. not beyond three sequential segments/

#722. Left antennule actual segment 10 <presence of spinules>/
1. without spinules/
2. with spinules/

#723. Left antennule actual segment 10 <presence of vestigial seta>/
1. without vestigial seta/
2. with vestigial seta/

#724. Left antennule actual segment 10 <presence of conical seta>/
1. without conical seta/
2. with conical seta/

#725. Left antennule actual segment 10 <presence of modified seta>/
1. without modified seta/
2. with modified seta/

#726. Left antennule actual segment 10 <form of the apex of the modified seta>/
1. presenting rounded apex/
2. presenting bifid apex/

#727. Left antennule actual segment 10 <number of modified seta>/
1. one element/
2. two elements/

#728. Left antennule actual segment 10 <range of the modified seta>/
1. surpassing to distal margin/
2. surpassing distal margin/

#729. Left antennule actual segment 10 <range of the modified seta>/
1. beyond of the sequential segment/
2. not beyond of the sequential segment/

#730. Left antennule actual segment 10 <position of the modified seta>/
1. parallel to antennule direction/
2. perpendicular to antennule direction/

#731. Left antennule actual segment 10 <presence of spinous process>/
1. without spinous process/
2. with spinous process/

#732. Left antennule actual segment 10 <presence of aesthetasc>/
1. without aesthetasc/
2. with aesthetasc/

#733. Left antennule actual segment 10 <number of aesthetasc>/
1. one element/
2. two elements/

#734. Left antennule actual segment 11 <presence of seta>/
1. with seta/
2. without seta/

#735. Left antennule actual segment 11 <number of setae>/
1. one element/
2. two elements/

#736. Left antennule actual segment 11 <size comparison of seta>/
1. of equal size/
2. of unequal size/

#737. Left antennule actual segment 11 <length of seta>/
1. one larger than segment/
2. none larger than segment/

#738. Left antennule actual segment 11 <composition of seta>/
1. straight/
2. plumose/

#739. Left antennule actual segment 11 <range of the seta>/
1. surpassing to distal margin/
2. surpassing distal margin/

#740. Left antennule actual segment 11 <range of the seta>/

1. beyond three sequential segments/
2. not beyond three sequential segments/

#741. Left antennule actual segment 11 <presence of spinules>/
1. without spinules/
2. with spinules/

#742. Left antennule actual segment 11 <presence of vestigial seta>/
1. without vestigial seta/
2. with vestigial seta/

#743. Left antennule actual segment 11 <presence of conical seta>/
1. without conical seta/
2. with conical seta/

#744. Left antennule actual segment 11 <presence of modified seta>/
1. without modified seta/
2. with modified seta/

#745. Left antennule actual segment 11 <form of the apex of the modified seta>/
1. presenting rounded apex/
2. presenting bifid apex/

#746. Left antennule actual segment 11 <number of modified seta>/
1. one element/
2. two elements/

#747. Left antennule actual segment 11 <range of the modified seta>/
1. surpassing to distal margin/
2. surpassing distal margin/

#748. Left antennule actual segment 11 <range of the modified seta>/
1. beyond of the sequential segment/
2. not beyond of the sequential segment/

#749. Left antennule actual segment 11 <position of the modified seta>/
1. parallel to antennule direction/
2. perpendicular to antennule direction/

#750. Left antennule actual segment 11 <presence of spinous process>/
1. without spinous process/
2. with spinous process/

#751. Left antennule actual segment 11 <presence of aesthetasc>/
1. without aesthetasc/
2. with aesthetasc/

#752. Left antennule actual segment 11 <number of aesthetasc>/
1. one element/

2. two elements/

#753. Left antennule actual segment 12 <presence of seta>/
1. with seta/
2. without seta/

#754. Left antennule actual segment 12 <number of setae>/
1. one element/
2. two elements/

#755. Left antennule actual segment 12 <size comparison of seta>/
1. of equal size/
2. of unequal size/

#756. Left antennule actual segment 12 <length of seta>/
1. one larger than segment/
2. none larger than segment/

#757. Left antennule actual segment 12 <composition of seta>/
1. straight/
2. plumose/

#758. Left antennule actual segment 12 <range of the seta>/
1. surpassing to distal margin/
2. surpassing distal margin/

#759. Left antennule actual segment 12 <range of the seta>/
1. beyond three sequential segments/
2. not beyond three sequential segments/

#760. Left antennule actual segment 12 <presence of spinules>/
1. without spinules/
2. with spinules/

#761. Left antennule actual segment 12 <presence of vestigial seta>/
1. without vestigial seta/
2. with vestigial seta/

#762. Left antennule actual segment 12 <presence of conical seta>/
1. without conical seta/
2. with conical seta/

#763. Left antennule actual segment 12 <presence of modified seta>/
1. without modified seta/
2. with modified seta/

#764. Left antennule actual segment 12 <presence of spinous process>/
1. without spinous process/
2. with spinous process/

#765. Left antennule actual segment 12 <presence of aesthetasc>/
1. without aesthetasc/
2. with aesthetasc/

#766. Left antennule actual segment 12 <number of aesthetasc>/
1. one element/
2. two elements/

#767. Left antennule actual segment 13 <presence of seta>/
1. with seta/
2. without seta/

#768. Left antennule actual segment 13 <number of setae>/
1. one element/
2. two elements/

#769. Left antennule actual segment 13 <size comparison of seta>/
1. of equal size/
2. of unequal size/

#770. Left antennule actual segment 13 <length of seta>/
1. one of them elongated/
2. none elongated/

#771. Left antennule actual segment 13 <composition of seta>/
1. straight/
2. plumose/

#772. Left antennule actual segment 13 <range of the seta>/
1. surpassing to distal margin/
2. surpassing distal margin/

#773. Left antennule actual segment 13 <range of the seta>/
1. beyond three sequential segments/
2. not beyond three sequential segments/

#774. Left antennule actual segment 13 <presence of spinules>/
1. without spinules/
2. with spinules/

#775. Left antennule actual segment 13 <presence of vestigial seta>/
1. without vestigial seta/
2. with vestigial seta/

#776. Left antennule actual segment 13 <presence of conical seta>/
1. without conical seta/
2. with conical seta/

#777. Left antennule actual segment 13 <presence of modified seta>/

1. without modified seta/
2. with modified seta/

#778. Left antennule actual segment 13 <form of the apex of the modified seta>/

1. presenting rounded apex/
2. presenting bifid apex/

#779. Left antennule actual segment 13 <number of modified seta>/

1. one element/
2. two elements/

#780. Left antennule actual segment 13 <range of the modified seta>/

1. surpassing to distal margin/
2. surpassing distal margin/

#781. Left antennule actual segment 13 <range of the modified seta>/

1. beyond of the sequential segment/
2. not beyond of the sequential segment/

#782. Left antennule actual segment 13 <position of the modified seta>/

1. parallel to antennule direction/
2. perpendicular to antennule direction/

#783. Left antennule actual segment 13 <presence of spinous process>/

1. without spinous process/
2. with spinous process/

#784. Left antennule actual segment 13 <presence of aesthetasc>/

1. without aesthetasc/
2. with aesthetasc/

#785. Left antennule actual segment 13 <number of aesthetasc>/

1. one element/
2. two elements/

#786. Left antennule actual segment 14 <presence of seta>/

1. with seta/
2. without seta/

#787. Left antennule actual segment 14 <number of setae>/

1. one element/
2. two elements/

#788. Left antennule actual segment 14 <size comparison of seta>/

1. of equal size/
2. of unequal size/

#789. Left antennule actual segment 14 <length of seta>/

1. elongated/
2. not elongated/

#790. Left antennule actual segment 14 <composition of seta>/
1. straight/
2. plumose/

#791. Left antennule actual segment 14 <range of the seta>/
1. surpassing to distal margin/
2. surpassing distal margin/

#792. Left antennule actual segment 14 <range of the seta>/
1. beyond three sequential segments/
2. not beyond three sequential segments/

#793. Left antennule actual segment 14 <presence of spinules>/
1. without spinules/
2. with spinules/

#794. Left antennule actual segment 14 <presence of vestigial seta>/
1. without vestigial seta/
2. with vestigial seta/

#795. Left antennule actual segment 14 <presence of conical seta>/
1. without conical seta/
2. with conical seta/

#796. Left antennule actual segment 14 <presence of modified seta>/
1. without modified seta/
2. with modified seta/

#797. Left antennule actual segment 14 <presence of spinous process>/
1. without spinous process/
2. with spinous process/

#798. Left antennule actual segment 14 <presence of aesthetasc>/
1. without aesthetasc/
2. with aesthetasc/

#799. Left antennule actual segment 14 <number of aesthetasc>/
1. one element/
2. two elements/

#800. Left antennule actual segment 15 <presence of seta>/
1. with seta/
2. without seta/

#801. Left antennule actual segment 15 <number of setae>/
1. one element/

2. two element/

#802. Left antennule actual segment 15 <size comparison of seta>/
1. of equal size/
2. of unequal size/

#803. Left antennule actual segment 15 <length of seta>/
1. larger than segment/
2. not larger than segment/

#804. Left antennule actual segment 15 <composition of seta>/
1. straight/
2. plumose/

#805. Left antennule actual segment 15 <range of the seta>/
1. surpassing to distal margin/
2. surpassing distal margin/

#806. Left antennule actual segment 15 <range of the seta>/
1. beyond three sequential segments/
2. not beyond three sequential segments/

#807. Left antennule actual segment 15 <presence of spinules>/
1. without spinules/
2. with spinules/

#808. Left antennule actual segment 15 <presence of vestigial seta>/
1. without vestigial seta/
2. with vestigial seta/

#809. Left antennule actual segment 15 <presence of conical seta>/
1. without conical seta/
2. with conical seta/

#810. Left antennule actual segment 15 <presence of modified seta>/
1. without modified seta/
2. with modified seta/

#811. Left antennule actual segment 15 <presence of spinous process>/
1. without spinous process/
2. with spinous process/

#812. Left antennule actual segment 15 <position of spinous process>/
1. on inner margin/
2. on outer margin/

#813. Left antennule actual segment 15 <reach of the spinous process>/
1. surpassing to distal margin/
2. surpassing distal margin/

#814. Left antennule actual segment 15 <presence of aesthetasc>/
1. without aesthetasc/
2. with aesthetasc/

#815. Left antennule actual segment 15 <number of aesthetasc>/
1. one element/
2. two elements/

#816. Left antennule actual segment 16 <presence of seta>/
1. with seta/
2. without seta/

#817. Left antennule actual segment 16 <number of setae>/
1. one element/
2. two elements/

#818. Left antennule actual segment 16 <size comparison of seta>/
1. of equal size/
2. of unequal size/

#819. Left antennule actual segment 16 <length of seta>/
1. larger than segment/
2. not larger than segment/

#820. Left antennule actual segment 16 <composition of seta>/
1. straight/
2. plumose/

#821. Left antennule actual segment 16 <range of the seta>/
1. surpassing to distal margin/
2. surpassing distal margin/

#822. Left antennule actual segment 16 <range of the seta>/
1. beyond three sequential segments/
2. not beyond three sequential segments/

#823. Left antennule actual segment 16 <presence of spinules>/
1. without spinules/
2. with spinules/

#824. Left antennule actual segment 16 <presence of vestigial seta>/
1. without vestigial seta/
2. with vestigial seta/

#825. Left antennule actual segment 16 <presence of conical seta>/
1. without conical seta/
2. with conical seta/

#826. Left antennule actual segment 16 <presence of modified seta>/
1. without modified seta/
2. with modified seta/

#827. Left antennule actual segment 16 <presence of spinous process>/
1. without spinous process/
2. with spinous process/

#828. Left antennule actual segment 16 <position of spinous process>/
1. on inner margin/
2. on outer margin/

#829. Left antennule actual segment 16 <size comparison with preceding segments>/
1. surpassing to distal margin/
2. surpassing distal margin/

#830. Left antennule actual segment 16 <size comparison with preceding segments>/
1. equal size to process on preceding segment/
2. unequal size to process on preceding segment/

#831. Left antennule actual segment 16 <size comparison with preceding segments>/
1. larger/
2. smaller/

#832. Left antennule actual segment 16 <presence of aesthetasc>/
1. without aesthetasc/
2. with aesthetasc/

#833. Left antennule actual segment 16 <number of aesthetasc>/
1. one element/
2. two elements/

#834. Left antennule actual segment 17 <presence of seta>/
1. with seta/
2. without seta/

#835. Left antennule actual segment 17 <number of setae>/
1. one element/
2. two element/

#836. Left antennule actual segment 17 <size comparison of seta>/
1. of equal size/
2. of unequal size/

#837. Left antennule actual segment 17 <length of seta>/
1. not larger than segment/

2. larger than segment/

#838. Left antennule actual segment 17 <composition of seta>/
1. straight/
2. plumose/

#839. Left antennule actual segment 17 <range of the seta>/
1. surpassing to distal margin/
2. surpassing distal margin/

#840. Left antennule actual segment 17 <range of the seta>/
1. beyond three sequential segments/
2. not beyond three sequential segments/

#841. Left antennule actual segment 17 <presence of spinules>/
1. without spinules/
2. with spinules/

#842. Left antennule actual segment 17 <presence of vestigial seta>/
1. with vestigial seta/
2. without vestigial seta/

#843. Left antennule actual segment 17 <presence of conical seta>/
1. with conical seta/
2. without conical seta/

#844. Left antennule actual segment 17 <presence of modified seta>/
1. without modified seta/
2. with modified seta/

#845. Left antennule actual segment 17 <form of the apex of the modified seta>/
1. presenting rounded apex/
2. presenting bifid apex/

#846. Left antennule actual segment 17 <number of modified seta>/
1. one element/
2. two elements/

#847. Left antennule actual segment 17 <range of the modified seta>/
1. surpassing to distal margin/
2. surpassing distal margin/

#848. Left antennule actual segment 17 <range of the modified seta>/
1. beyond of the sequential segment/
2. not beyond of the sequential segment/

#849. Left antennule actual segment 17 <position of the modified seta>/
1. parallel to antennule direction/
2. perpendicular to antennule direction/

#850. Left antennule actual segment 17 <presence of spinous process>/
  1. without spinous process/
  2. with spinous process/

#851. Left antennule actual segment 17 <presence of aesthetasc>/
  1. without aesthetasc/
  2. with aesthetasc/

#852. Left antennule actual segment 17 <number of aesthetasc>/
  1. one element/
  2. two elements/

#853. Left antennule actual segment 18 <presence of seta>/
  1. with seta/
  2. without seta/

#854. Left antennule actual segment 18 <number of setae>/
  1. one element/
  2. two elements/

#855. Left antennule actual segment 18 <size comparison of seta>/
  1. of equal size/
  2. of unequal size/

#856. Left antennule actual segment 18 <length of seta>/
  1. larger than segment/
  2. not larger than segment/

#857. Left antennule actual segment 18 <composition of seta>/
  1. straight/
  2. plumose/

#858. Left antennule actual segment 18 <range of the seta>/
  1. surpassing to distal margin/
  2. surpassing distal margin/

#859. Left antennule actual segment 18 <range of the seta>/
  1. beyond three sequential segments/
  2. not beyond three sequential segments/

#860. Left antennule actual segment 18 <presence of spinules>/
  1. without spinules/
  2. with spinules/

#861. Left antennule actual segment 18 <presence of vestigial seta>/
  1. without vestigial seta/
  2. with vestigial seta/

#862. Left antennule actual segment 18 <presence of conical seta>/
1. without conical seta/
2. with conical seta/

#863. Left antennule actual segment 18 <presence of modified seta>/
1. without modified seta/
2. with modified seta/

#864. Left antennule actual segment 18 <form of the apex of the modified seta>/
1. presenting rounded apex/
2. presenting bifid apex/

#865. Left antennule actual segment 18 <number of modified seta>/
1. one element/
2. two elements/

#866. Left antennule actual segment 18 <range of the modified seta>/
1. surpassing to distal margin/
2. surpassing distal margin/

#867. Left antennule actual segment 18 <range of the modified seta>/
1. beyond of the sequential segment/
2. not beyond of the sequential segment/

#868. Left antennule actual segment 18 <position of the modified seta>/
1. parallel to antennule direction/
2. perpendicular to antennule direction/

#869. Left antennule actual segment 18 <presence of spinous process>/
1. without spinous process/
2. with spinous process/

#870. Left antennule actual segment 18 <presence of aesthetasc>/
1. without aesthetasc/
2. with aesthetasc/

#871. Left antennule actual segment 18 <number of aesthetasc>/
1. one element/
2. two elements/

#872. Left antennule actual segment 19 <presence of seta>/
1. with seta/
2. without seta/

#873. Left antennule actual segment 19 <number of setae>/
1. one element/
2. two elements/

#874. Left antennule actual segment 19 <size comparison of seta>/

1. of equal size/
2. of unequal size/

#875. Left antennule actual segment 19 <length of seta>/
1. larger than segment/
2. not larger than segment/

#876. Left antennule actual segment 19 <composition of seta>/
1. straight/
2. plumose/

#877. Left antennule actual segment 19 <range of the seta>/
1. surpassing to distal margin/
2. surpassing distal margin/

#878. Left antennule actual segment 19 <range of the seta>/
1. beyond three sequential segments/
2. not beyond three sequential segments/

#879. Left antennule actual segment 19 <presence of spinules>/
1. without spinules/
2. with spinules/

#880. Left antennule actual segment 19 <presence of vestigial seta>/
1. without vestigial seta/
2. with vestigial seta/

#881. Left antennule actual segment 19 <presence of conical seta>/
1. without conical seta/
2. with conical seta/

#882. Left antennule actual segment 19 <presence of modified seta>/
1. without modified seta/
2. with modified seta/

#883. Left antennule actual segment 19 <form of the apex of the modified seta>/
1. presenting rounded apex/
2. presenting bifid apex/

#884. Left antennule actual segment 19 <number of modified seta>/
1. one element/
2. two elements/

#885. Left antennule actual segment 19 <range of the modified seta>/
1. surpassing to distal margin/
2. surpassing distal margin/

#886. Left antennule actual segment 19 <range of the modified seta>/
1. beyond of the sequential segment/

2. not beyond of the sequential segment/

#887. Left antennule actual segment 19 <position of the modified seta>/
1. parallel to antennule direction/
2. perpendicular to antennule direction/

#888. Left antennule actual segment 19 <presence of spinous process>/
1. without spinous process/
2. with spinous process/

#889. Left antennule actual segment 19 <presence of aesthetasc>/
1. without aesthetasc/
2. with aesthetasc/

#890. Left antennule actual segment 19 <number of aesthetasc>/
1. one element/
2. two elements/

#891. Left antennule actual segment 20 <presence of seta>/
1. with seta/
2. without seta/

#892. Left antennule actual segment 20 <number of setae>/
1. one element/
2. two elements/

#893. Left antennule actual segment 20 <size comparison of seta>/
1. of equal size/
2. of unequal size/

#894. Left antennule actual segment 20 <length of seta>/
1. larger than segment/
2. not larger than segment/

#895. Left antennule actual segment 20 <composition of seta>/
1. straight/
2. plumose/

#896. Left antennule actual segment 20 <range of the seta>/
1. surpassing to distal margin/
2. surpassing distal margin/

#897. Left antennule actual segment 20 <range of the seta>/
1. beyond three sequential segments/
2. not beyond three sequential segments/

#898. Left antennule actual segment 20 <presence of spinules>/
1. without spinules/
2. with spinules/

#899. Left antennule actual segment 20 <presence of vestigial seta>/
1. without vestigial seta/
2. with vestigial seta/

#900. Left antennule actual segment 20 <presence of conical seta>/
1. without conical seta/
2. with conical seta/

#901. Left antennule actual segment 20 <presence of modified seta>/
1. without modified seta/
2. with modified seta/

#902. Left antennule actual segment 20 <presence of spinous process>/
1. without spinous process/
2. with spinous process/

#903. Left antennule actual segment 20 <presence of aesthetasc>/
1. without aesthetasc/
2. with aesthetasc/

#904. Left antennule actual segment 20 <number of aesthetasc>/
1. one element/
2. two elements/

#905. Left antennule actual segment 21 <presence of seta>/
1. with seta/
2. without seta/

#906. Left antennule actual segment 21 <number of setae>/
1. one element/
2. two elements/

#907. Left antennule actual segment 21 <size comparison of seta>/
1. of equal size/
2. of unequal size/

#908. Left antennule actual segment 21 <length of seta>/
1. larger than segment/
2. not larger than segment/

#909. Left antennule actual segment 21 <composition of seta>/
1. straight/
2. plumose/

#910. Left antennule actual segment 21 <range of the seta>/
1. surpassing to distal margin/
2. surpassing distal margin/

#911. Left antennule actual segment 21 <range of the seta>/
1. beyond three sequential segments/
2. not beyond three sequential segments/

#912. Left antennule actual segment 21 <presence of spinules>/
1. without spinules/
2. with spinules/

#913. Left antennule actual segment 21 <presence of vestigial seta>/
1. without vestigial seta/
2. with vestigial seta/

#914. Left antennule actual segment 21 <presence of conical seta>/
1. without conical seta/
2. with conical seta/

#915. Left antennule actual segment 21 <presence of modified seta>/
1. without modified seta/
2. with modified seta/

#916. Left antennule actual segment 21 <presence of spinous process>/
1. without spinous process/
2. with spinous process/

#917. Left antennule actual segment 21 <presence of aesthetasc>/
1. without aesthetasc/
2. with aesthetasc/

#918. Left antennule actual segment 21 <number of aesthetasc>/
1. one element/
2. two elements/

#919. Left antennule actual segment 22 <presence of seta>/
1. with seta/
2. without seta/

#920. Left antennule actual segment 22 <number of setae>/
1. two elements/
2. six elements/

#921. Left antennule actual segment 22 <size comparison of seta>/
1. of equal size/
2. of unequal size/

#922. Left antennule actual segment 22 <length of seta>/
1. one of them elongated/
2. no elongated/

#923. Left antennule actual segment 22 <composition of seta>/

1. straight/
2. plumose/

#924. Left antennule actual segment 22 <range of the seta>/
1. surpassing to distal margin/
2. surpassing distal margin/

#925. Left antennule actual segment 22 <presence of spinules>/
1. without spinules/
2. with spinules/

#926. Left antennule actual segment 22 <presence of vestigial seta>/
1. without vestigial seta/
2. with vestigial seta/

#927. Left antennule actual segment 22 <presence of conical seta>/
1. without conical seta/
2. with conical seta/

#928. Left antennule actual segment 22 <presence of modified seta>/
1. without modified seta/
2. with modified seta/

#929. Left antennule actual segment 22 <presence of spinous process>/
1. without spinous process/
2. with spinous process/

#930. Left antennule actual segment 22 <presence of aesthetasc>/
1. without aesthetasc/
2. with aesthetasc/

#931. Left antennule actual segment 22 <number of aesthetasc>/
1. one element/
2. two elements/

#932. Left antennule actual segment 23 <presence of seta>/
1. with seta/
2. without seta/

#933. Left antennule actual segment 23 <number of setae>/
1. one element/
2. two elements/

#934. Left antennule actual segment 23 <size comparison of seta>/
1. of equal size/
2. of unequal size/

#935. Left antennule actual segment 23 <length of seta>/
1. one larger than segment/

2. none larger than segment/

#936. Left antennule actual segment 23 <composition of seta>/
1. straight/
2. plumose/

#937. Left antennule actual segment 23 <range of the seta>/
1. surpassing to distal margin/
2. surpassing distal margin/

#938. Left antennule actual segment 23 <range of the seta>/
1. greater 3x than original segment/
2. not greater 3x than original segment/

#939. Left antennule actual segment 23 <presence of spinules>/
1. with spinules/
2. without spinules/

#940. Left antennule actual segment 23 <presence of vestigial seta>/
1. with vestigial seta/
2. without vestigial seta/

#941. Left antennule actual segment 23 <presence of conical seta>/
1. with conical seta/
2. without conical seta/

#942. Left antennule actual segment 23 <presence of modified seta>/
1. with modified seta/
2. without modified seta/

#943. Left antennule actual segment 23 <presence of spinous process>/
1. with spinous process/
2. without spinous process/

#944. Left antennule actual segment 23 <presence of aesthetasc>/
1. without aesthetasc/
2. with aesthetasc/

#945. Left antennule actual segment 23 <number of aesthetasc>/
1. one element/
2. two elements/

#946. Left antennule actual segment 24 <presence of seta>/
1. with seta/
2. without seta/

#947. Left antennule actual segment 24 <number of setae>/
1. one element/
2. two elements/

#948. Left antennule actual segment 24 <size comparison of seta>/
1. of equal size/
2. of unequal size/

#949. Left antennule actual segment 24 <length of seta>/
1. one larger than segment/
2. none larger than segment/

#950. Left antennule actual segment 24 <composition of seta>/
1. straight/
2. plumose/

#951. Left antennule actual segment 24 <range of the seta>/
1. surpassing to distal margin/
2. surpassing distal margin/

#952. Left antennule actual segment 24 <range of the seta>/
1. greater 3x than original segment/
2. not greater 3x than original segment/

#953. Left antennule actual segment 24 <presence of spinules>/
1. with spinules/
2. without spinules/

#954. Left antennule actual segment 24 <presence of vestigial seta>/
1. with vestigial seta/
2. without vestigial seta/

#955. Left antennule actual segment 24 <presence of conical seta>/
1. with conical seta/
2. without conical seta/

#956. Left antennule actual segment 24 <presence of modified seta>/
1. with modified seta/
2. without modified seta/

#957. Left antennule actual segment 24 <presence of spinous process>/
1. with spinous process/
2. without spinous process/

#958. Left antennule actual segment 24 <presence of aesthetasc>/
1. without aesthetasc/
2. with aesthetasc/

#959. Left antennule actual segment 24 <number of aesthetasc>/
1. one element/
2. two elements/

#960. Left antennule actual segment 25 <presence of seta>/
1. with seta/
2. without seta/

#961. Left antennule actual segment 25 <number of setae>/
1. four elements/
2. five elements/

#962. Left antennule actual segment 25 <size comparison of seta>/
1. of equal size/
2. of unequal size/

#963. Left antennule actual segment 25 <length of seta>/
1. elongated/
2. not elongated/

#964. Left antennule actual segment 25 <composition of seta>/
1. straight/
2. plumose/

#965. Left antennule actual segment 25 <range of the seta>/
1. surpassing to distal margin/
2. surpassing distal margin/

#966. Left antennule actual segment 25 <range of the seta>/
1. 4 times larger than segment/
2. not 4 times larger than segment/

#967. Left antennule actual segment 25 <presence of spinules>/
1. with spinules/
2. without spinules/

#968. Left antennule actual segment 25 <presence of vestigial seta>/
1. with vestigial seta/
2. without vestigial seta/

#969. Left antennule actual segment 25 <presence of conical seta>/
1. with conical seta/
2. without conical seta/

#970. Left antennule actual segment 25 <presence of modified seta>/
1. with modified seta/
2. without modified seta/

#971. Left antennule actual segment 25 <presence of spinous process>/
1. with spinous process/
2. without spinous process/

#972. Left antennule actual segment 25 <presence of aesthetasc>/

1. without aesthetasc/
2. with aesthetasc/

#973. Left antennule actual segment 25 <number of aesthetasc>/
1. one element/
2. two elements/

#974. Antenna <number of rami>/
1. uniramous/
2. biramous/

#975. Antenna coxa <fusion>/
1. separated from the basis/
2. fused to basis/

#976. Antenna coxa <presence of seta>/
1. bearing seta/
2. not bearing seta/

#977. Antenna coxa <number of setae>/
1. 1/
2. 2/

#978. Antenna coxa <location of setae>/
1. on inner surface/
2. on outer surface/

#979. Antenna coxa <positioning of setae>/
1. at medial corner/
2. at distal corner/

#980. Antenna coxa <reach of the seta>/
1. reaching to the middle base/
2. reaching to the endopod 1/

#981. Antenna basis (fusion)/
1. separated from the endopodal segment/
2. fused to endopodal segment/

#982. Antenna basis <presence of seta>/
1. bearing seta/
2. not bearing seta/

#983. Antenna basis <number of setae>/
1. 1/
2. 2/

#984. Antenna basis <location of setae>/
1. on inner surface/

2. on outer surface/

#985. Antenna basis <positioning of seta>/
1. at medial corner/
2. at distal corner/

#986. Antenna endopod ancestral segment I and II <fusion>/
1. separated/
2. fused/

#987. Antenna endopod ancestral segment II and III <fusion>/
1. separated/
2. fused/

#988. Antenna endopod ancestral segment III and IV <fusion>/
1. separated/
2. fused/

#989. Antenna endopod ancestral segment III and IV <status of the fusion>/
1. partially/
2. fully/

#990. Antenna endopod actual <number of actual segment>/
1. 2-segmented/
2. 3-segmented/

#991. <Antenna endopod> actual segment 1 <formation>/
1. not bilobate/
2. bilobate/

#992. <Antenna endopod> actual segment 1 <number of setae on inner lobe of the segment bilobate>/
1. with inner lobe bearing more than 8 setae/
2. with inner lobe bearing less than 8 setae/

#993. <Antenna endopod> actual segment 1 <location of the seta on inner lobe of the segment bilobate>/
1. distally/
2. proximally/

#994. <Antenna endopod> actual segment 1 <number of setae on outer lobe of the segment bilobate>/
1. outer lobe bearing more than 8 setae/
2. outer lobe bearing less than 8 setae/

#995. <Antenna endopod> actual segment 1 <location of the seta on outer lobe of the segment bilobate>/
1. distally/
2. proximally/

#996. <Antenna endopod> actual segment 1 <presence of seta>/
1. without seta/
2. with seta/

#997. <Antenna endopod> actual segment 1 <number of setae>/
1. one/
2. two/

#998. <Antenna endopod> actual segment 1 <location of setae>/
1. on inner margin/
2. on outer margin/

#999. <Antenna endopod> actual segment 1 <presence of spinules>/
1. without spinules/
2. with spinules/

#1000. <Antenna endopod> actual segment 1 <arrangement of spinules>/
1. as a row/
2. as a patch/

#1001. <Antenna endopod> actual segment 1 <position of spinules on the segment>/
1. obliquely/
2. not obliquely/

#1002. <Antenna endopod> actual segment 1 <location of spinules>/
1. on inner surface/
2. on outer surface/

#1003. <Antenna endopod> actual segment 1 <presence of pore>/
1. without pore/
2. with pore/

#1004. <Antenna endopod> Actual segment 2 <formation>/
1. not bilobate/
2. bilobate/

#1005. <Antenna endopod> Actual segment 2 <observation of discontinuity on the outer cuticle>/
1. with discontinuity on outer cuticle/
2. without discontinuity on outer cuticle/

#1006. <Antenna endopod> Actual segment 2 <condition of the discontinuity present>/
1. developed as a suture/
2. not developed as a suture/

#1007. <Antenna endopod> Actual segment 2 <condition of the development suture present>/

1. complete/
2. incomplete/

#1008. <Antenna endopod> Actual segment 2 <number of setae on the segment>/
1. inner lobe bearing 8 setae/
2. inner lobe bearing 9 setae/
3. inner lobe bearing 7 setae/

#1009. <Antenna endopod> Actual segment 2 <location of the setae on inner lobe of the segment>/
1. distally/
2. proximally/

#1010. <Antenna endopod> Actual segment 2 <number of setae on outer lobe of the segment>/
1. outer lobe bearing 6 setae/
2. outer lobe bearing 7 setae/

#1011. <Antenna endopod> Actual segment 2 <location of the setae on outer lobe of the segment>/
1. distally/
2. proximally/

#1012. <Antenna endopod> Actual segment 2 <presence of spinules>/
1. with spinules/
2. without spinules/

#1013. <Antenna endopod> Actual segment 2 <arrangement of spinules>/
1. as a row/
2. as a patch/

#1014. <Antenna endopod> Actual segment 2 <position of spinules on the segment>/
1. obliquely/
2. not obliquely/

#1015. <Antenna endopod> Actual segment 2 <location of spinules>/
1. on inner surface/
2. on outer surface/

#1016. <Antenna endopod> Actual segment 3 <number of setae on outer lobe of the segment>/
1. outer lobe bearing 7 setae/
2. outer lobe bearing 8 setae/

#1017. <Antenna endopod> Actual segment 3 <location of the setae on outer lobe of the segment>/
1. distally/
2. proximally/

#1018. <Antenna endopod> Actual segment 3 <presence of spinules>/
1. without spinules/
2. with spinules/

#1019. <Antenna endopod> Actual segment 3 <arrangement of spinules>/
1. as a row/
2. as a patch/

#1020. <Antenna endopod> Actual segment 3 <position of spinules on the segment>/
1. obliquely/
2. not obliquely/

#1021. <Antenna endopod> Actual segment 3 <location of spinules>/
1. on inner surface/
2. on outer surface/

#1022. Antenna exopod ancestral segment I and II <fusion>/
1. separated/
2. fused/

#1023. Antenna exopod ancestral segment II and III <fusion>/
1. separated/
2. fused/

#1024. Antenna exopod ancestral segment III and IV <fusion>/
1. separated/
2. fused/

#1025. Antenna exopod ancestral segment IV and V <fusion>/
1. separated/
2. fused/

#1026. Antenna exopod ancestral segment V and VI <fusion>/
1. separated/
2. fused/

#1027. Antenna exopod ancestral segment VI and VII <fusion>/
1. separated/
2. fused/

#1028. Antenna exopod ancestral segment VII and VIII <fusion>/
1. separated/
2. fused/

#1029. Antenna exopod ancestral segment VIII and IX <fusion>/
1. separated/
2. fused/

#1030. Antenna exopod ancestral segment IX and X <fusion>/

1. separated/
2. fused/

#1031. Antenna exopod actual <number of actual segments>/
1. 7-segmented/
2. 8-segmented/

#1032. <Antenna exopod> Actual segment 1 <composition segment>/
1. single/
2. compound/

#1033. <Antenna exopod> Actual segment 1 <relation of size of the segment>/
1. elongated (width-length, equal or larger ratio 2:1)/
2. not elongated (width-length, ratio 1:1)/

#1034. <Antenna exopod> Actual segment 1 <presence of seta>/
1. with seta/
2. without seta/

#1035. <Antenna exopod> Actual segment 1 <number of setae>/
1. one/
2. two/

#1036. <Antenna exopod> Actual segment 1 <location of setae>/
1. at inner surface/
2. at outer surface/

#1037. <Antenna exopod> Actual segment 2 <composition segment>/
1. single/
2. compound/

#1038. <Antenna exopod> Actual segment 2 <relation of size of the segment>/
1. elongated (larger width-length ratio 2:1)/
2. not elongated (lesser width-length ratio 2:1)/

#1039. <Antenna exopod> Actual segment 2 <presence of seta>/
1. with seta/
2. without seta/

#1040. <Antenna exopod> Actual segment 2 <number of setae>/
1. one/
2. three/

#1041. <Antenna exopod> Actual segment 2 <location of setae>/
1. at inner surface/
2. at outer surface/

#1042. <Antenna exopod> Actual segment 3 <composition segment>/
1. single/

2. compound/

#1043. <Antenna exopod> Actual segment 3 <relation of size of the segment>/
1. elongated (larger or equal width-length ratio 2:1)/
2. not elongated (lesser width-length ratio 2:1)/

#1044. <Antenna exopod> Actual segment 3 <presence of seta>/
1. with seta/
2. without seta/

#1045. <Antenna exopod> Actual segment 3 <number of setae>/
1. one/
2. two/

#1046. <Antenna exopod> Actual segment 3 <location of setae>/
1. at inner surface/
2. at outer surface/

#1047. <Antenna exopod> Actual segment 4 <composition segment>/
1. single/
2. compound/

#1048. <Antenna exopod> Actual segment 4 <relation of size of the segment>/
1. elongated (larger or equal width-length ratio 2:1)/
2. not elongated (lesser width-length ratio 2:1)/

#1049. <Antenna exopod> Actual segment 4 <presence of seta>/
1. with seta/
2. without seta/

#1050. <Antenna exopod> Actual segment 4 <number of setae>/
1. one/
2. two/

#1051. <Antenna exopod> Actual segment 4 <location of setae>/
1. at inner surface/
2. at outer surface/

#1052. <Antenna exopod> Actual segment 5 <composition segment>/
1. single/
2. compound/

#1053. <Antenna exopod> Actual segment 5 <relation of size of the segment>/
1. elongated (larger or equal width-length ratio 2:1)/
2. not elongated (lesser width-length ratio 2:1)/

#1054. <Antenna exopod> Actual segment 5 <presence of seta>/
1. with seta/
2. without seta/

#1055. <Antenna exopod> Actual segment 5 <number of setae>/
1. one/
2. two/

#1056. <Antenna exopod> Actual segment 5 <location of setae>/
1. at inner surface/
2. at outer surface/

#1057. <Antenna exopod> Actual segment 6 <composition segment>/
1. single/
2. compound/

#1058. <Antenna exopod> Actual segment 6 <relation of size of the segment>/
1. elongated (larger or equal width-length ratio 2:1)/
2. not elongated (lesser width-length ratio 2:1)/

#1059. <Antenna exopod> Actual segment 6 <presence of seta>/
1. with seta/
2. without seta/

#1060. <Antenna exopod> Actual segment 6 <number of setae>/
1. one/
2. two/

#1061. <Antenna exopod> Actual segment 6 <location of setae>/
1. at inner surface/
2. at outer surface/

#1062. <Antenna exopod> Actual segment 7 <composition segment>/
1. single/
2. compound/

#1063. <Antenna exopod> Actual segment 7 <relation of size of the segment>/
1. elongated (larger or equal width-length ratio 2:1)/
2. not elongated (lesser width-length ratio 2:1)/

#1064. <Antenna exopod> Actual segment 7 <presence of seta>/
1. with seta/
2. without seta/

#1065. <Antenna exopod> Actual segment 7 <number of setae>/
1. one/
2. two/

#1066. <Antenna exopod> Actual segment 7 <location of setae>/
1. at inner surface/
2. at outer surface/

#1067. <Antenna exopod> Actual segment 7 <number of setae>/
1. and two/
2. and three/

#1068. <Antenna exopod> Actual segment 7 <location of setae>/
1. at proximal surface/
2. at distal surface/

#1069. <Antenna exopod> Actual segment 8 <relation of size of the segment>/
1. not elongated (lesser width-length ratio 2:1)/
2. elongated (larger or equal width-length ratio 2:1)/

#1070. <Antenna exopod> Actual segment 8 <quantify of seta>/
1. with 3 setae/
2. with 4 setae/

#1071. <Antenna exopod> Actual segment 8 <location of setae>/
1. at inner surface/
2. at distal surface/

#1072. Mandible coxal gnathobase <composition>/
1. sclerotized/
2. not sclerotized/

#1073. Mandible coxal gnathobase <lobular form present>/
1. with lobe/
2. without lobe/

#1074. Mandible coxal gnathobase <condition of the protuberance>/
1. prominent/
2. not prominent/

#1075. Mandible coxal gnathobase <location of the ornamentation>/
1. on caudal margin/
2. on sub-caudal margin/

#1076. Mandible coxal gnathobase <presence of cutting blade>/
1. presence of cutting blade/
2. absence of cutting blade/

#1077. Mandible coxal gnathobase <composition of cutting blade>/
1. with tooth-like prominence/
2. without tooth-like prominence/

#1078. Mandible coxal gnathobase <number of prominences of cutting blade>/
1. one, distinctly/
2. two, distinctly/

#1079. Mandible coxal gnathobase <form of prominence 1 of cutting blade>/

1. 1 triangular/
2. 1 acute/

#1080. Mandible coxal gnathobase <location of prominence 1 of cutting blade>/
1. on caudal margin/
2. on sub-caudal margin/

#1081. Mandible coxal gnathobase <form of prominence 2 of cutting blade>/
1. and 1 triangular/
2. and 1 acute/

#1082. Mandible coxal gnathobase <location of prominence 2 of cutting blade>/
1. on caudal margin/
2. on sub-caudal margin/

#1083. Mandible coxal gnathobase <acute projection between the prominences>/
1. without acute projection between the prominences/
2. with acute projection between the prominences/

#1084. Mandible coxal gnathobase <presence of spinules on cutting blade>/
1. without additional spinules/
2. with additional spinules/

#1085. Mandible coxal gnathobase <arrangement of spinules on cutting blade>/
1. as a row/
2. as a patch/

#1086. Mandible coxal gnathobase <location of spinules on cutting blade>/
1. on dorsal surface/
2. on ventral surface/

#1087. Mandible coxal gnathobase <ornamentation of cutting blade>/
1. with seta/
2. without seta/

#1088. Mandible coxal gnathobase <number of setae of the cutting blade>/
1. 1/
2. 2/

#1089. Mandible coxal gnathobase <location of the seta 1 of the cutting blade>/
1. dorsally/
2. ventraly/

#1090. Mandible coxal gnathobase <location of the seta 1 of the cutting blade>/
1. on apical surface/
2. on caudal surface/

#1091. Mandible coxal gnathobase <presence of spinules on prominence 1 of cutting blade>/

1. without spinules/
2. with spinules/

#1092. Mandible coxal gnathobase <location of spinules on the cutting blade>/
1. apicalmost/
2. basalmost/

#1093. Mandible palps <number of rami>/
1. uniramous/
2. biramous/

#1094. Mandible palps <comprising the basis>/
1. comprising the basis/
2. not comprising the basis/

#1095. Mandible palps <presence of seta on basis>/
1. with seta/
2. without seta/

#1096. Mandible palps <number of setae on basis>/
1. three/
2. four/

#1097. Mandible palps <insertion of seta on basis>/
1. equally inserted/
2. differently inserted/

#1098. Mandible palps <location of setae equally inserted on basis>/
1. proximally/
2. medially/
3. distally/

#1099. Mandible palps <location of first seta differently inserted on basis>/
1. first medially/
2. first distally/

#1100. Mandible palps <reach of the first seta differently inserted on basis>/
1. reaching to beyond the endopod 1/
2. not reaching to beyond the endopod 1/

#1101. Mandible palps <location of second seta differently inserted on basis>/
1. second medially/
2. second distally/

#1102. Mandible palps <location of third seta differently inserted on basis>/
1. third medially/
2. third distally/

#1103. Mandible palps <location of fourth seta differently inserted on basis>/

1. fourth medially/
2. fourth distally/

#1104. Mandible palps <location of setae on basis>/
1. on inner margin/
2. on outer margin/

#1105. Mandible palps <ornamentation of the setae>/
1. none with setulose ornamentation/
2. all with setulose ornamentation/

#1106. Mandible endopod <number of segment>/
1. 1-segmented/
2. 2-segmented/

#1107. Mandible endopod 1 <presence of lobe>/
1. with lobe/
2. without lobe/

#1108. Mandible endopod 1 <presence of seta>/
1. bearing seta/
2. no bearing seta/

#1109. Mandible endopod 1 <number of setae>/
1. four/
2. five/

#1110. Mandible endopod 1 <insertion of seta>/
1. medially inserted/
2. distally inserted/

#1111. Mandible endopod 1 <presence of spinules>/
1. without spinules/
2. with spinules/

#1112. Mandible endopod 1 <arrangement of spinules>/
1. as a row/
2. as a patch/

#1113. Mandible endopod 1 <number of row of spinules>/
1. single/
2. double/

#1114. Mandible endopod 2 <presence of lobe>/
1. with lobe/
2. without lobe/

#1115. Mandible endopod 2 <presence of setae>/
1. bearing setae/

2. not bearing setae/

#1116. Mandible endopod 2 <total of setae>/
1. seven elements/
2. nine elements/
3. ten elements/

#1117. Mandible endopod 2 <insertion of setae>/
1. medially inserted/
2. distally inserted/

#1118. Mandible endopod 2 <presence of spinules>/
1. without spinules/
2. with spinules/

#1119. Mandible endopod 2 <arrangement of spinules>/
1. as a row/
2. as a patch/

#1120. Mandible endopod 2 <number of row of spinules>/
1. single/
2. double/

#1121. Mandible exopod <number of segment>/
1. 4-segmented/
2. 5-segmented/

#1122. Mandible exopod 1 <presence of seta>/
1. with seta/
2. without seta/

#1123. Mandible exopod 1 <total of seta>/
1. one element/
2. two elements/

#1124. Mandible exopod 1 <location of setae>/
1. proximally/
2. distally/

#1125. Mandible exopod 1 <location of setae>/
1. on inner margin/
2. on outer margin/

#1126. Mandible exopod 2 <presence of seta>/
1. with seta/
2. without seta/

#1127. Mandible exopod 2 <number of setae>/
1. one element/

2. two elements/

#1128. Mandible exopod 2 <location of setae>/
1. proximally/
2. distally/

#1129. Mandible exopod 2 <location of setae>/
1. on inner side/
2. on outer side/

#1130. Mandible exopod 3 <presence of seta>/
1. with seta/
2. without seta/

#1131. Mandible exopod 3 <number of setae>/
1. one element/
2. two elements/

#1132. Mandible exopod 3 <location of setae>/
1. proximally/
2. distally/

#1133. Mandible exopod 3 <location of setae>/
1. on inner side/
2. on outer side/

#1134. Mandible exopod 4 <presence of seta>/
1. with setae/
2. without setae/

#1135. Mandible exopod 4 <total of setae>/
1. one element/
2. two elements/
3. three elements/

#1136. Mandible exopod 4 <location of setae>/
1. on medial region/
2. on terminal region/

#1137. Mandible exopod 5 <number of seta>/
1. with 2 setae/
2. with 3 setae/

#1138. Maxillule <number of rami>/
1. uniramous/
2. biramous/

#1139. Maxillule <segmentation>/
1. 2-segmented/

2. 3-segmented/

#1140. Maxillule praecoxa <presence of praecoxal arthrite>/
1. with praecoxal arthrite/
2. without praecoxal arthrite/

#1141. Maxillule praecoxa <presence of spines of praecoxal arthrite>/
1. bearing spines/
2. not bearing spines/

#1142. Maxillule praecoxa <number of setae of praecoxal arthrite>/
1. fifteen elements/
2. sixteen elements/

#1143. Maxillule praecoxa <location of setae of praecoxal arthrite>/
1. ten marginally/
2. nine marginally/

#1144. Maxillule praecoxa <number and details>/
1. plus four sub-marginally/
2. plus five sub-marginally/

#1145. Maxillule praecoxa <presence of spinules on praecoxal arthrite>/
1. with spinules/
2. without spinules/

#1146. Maxillule praecoxa <arrangement of spinules on praecoxal arthrite>/
1. as a row/
2. as a patch/

#1147. Maxillule praecoxa <location of spinules on praecoxal arthrite>/
1. on marginal surface/
2. on sub-marginal surface/

#1148. Maxillule coxa <presence of coxal epipodite>/
1. with coxal epipodite/
2. without coxal epipodite/

#1149. Maxillule coxa <development of outer lobe of the coxal epipodite>/
1. with conspicuous outer lobe/
2. with non-conspicuous outer lobe/

#1150. Maxillule coxa <presence of seta on coxal epipodite>/
1. bearing setae/
2. not bearing setae/

#1151. Maxillule coxa <number of setae at coxal epipodite>/
1. nine elements/
2. eleven elements/

#1152. Maxillule coxa <presence of coxal endite>/
1. with coxal endite/
2. without coxal endite/

#1153. Maxillule coxa <relation of size of the coxal endite>/
1. elongated (larger or equal width-length ratio 2:1)/
2. not elongated (lesser width-length ratio 2:1)/

#1154. Maxillule coxa <presence of setae at coxal endite>/
1. bearing setae/
2. not bearing setae/

#1155. Maxillule coxa <number of setae at coxal endite>/
1. three elements/
2. four elements/

#1156. Maxillule basis <presence of basal endite>/
1. with basal endite/
2. without basal endite/

#1157. Maxillule basis <number of basal endite>/
1. single/
2. double/

#1158. Maxillule basis <relation of size of the basal endite>/
1. elongated (larger width-length ratio 2:1)/
2. not elongated (lesser width-length ratio 2:1)/

#1159. Maxillule basis <presence of setae at basal endite>/
1. with setae/
2. without setae/

#1160. Maxillule basis <number of setae at basal endite>/
1. six elements/
2. seven elements/

#1161. Maxillule basis <insertion of seta at basal endite>/
1. distally inserted/
2. proximally inserted/

#1162. Maxillule basis <location of first basal endite>/
1. first proximal/
2. first distal/

#1163. Maxillule basis <relation of size of the first basal endite>/
1. elongated (larger width-length ratio 2:1/
2. not elongated (lesser width-length ratio 2:1)/

#1164. Maxillule basis <fusion of the first basal endite to basis>/
1. separated from basis/
2. fused to basis/

#1165. Maxillule basis <presence of setae at first basal endite>/
1. with setae/
2. without setae/

#1166. Maxillule basis <number of setae at first basal endite>/
1. three elements/
2. four elements/

#1167. Maxillule basis <insertion of seta at first basal endite>/
1. distally inserted/
2. proximally inserted/

#1168. Maxillule basis <location of second basal endite>/
1. second proximal/
2. second distal/

#1169. Maxillule basis <fusion of the second basal endite>/
1. fused to basis/
2. separated from the basis/

#1170. Maxillule basis <relation of size of the second basal endite>/
1. elongated (larger width-length ratio 2:1)/
2. not elongated (lesser width-length ratio 2:1)/

#1171. Maxillule basis <presence of setae at second basal endite>/
1. with setae/
2. without setae/

#1172. Maxillule basis <number of setae at second basal endite>/
1. three elements/
2. four elements/

#1173. Maxillule basis <insertion of setae at second basal endite>/
1. distally inserted/
2. proximally inserted/

#1174. Maxillule basis <presense of setules at second basal endite>/
1. with setules/
2. without setules/

#1175. Maxillule basis <arrangement of ornamentation at second basal endite>/
1. as a row/
2. as a patch/

#1176. Maxillule basis <location of ornamentation at second basal endite>/

1. on inner side/
2. on outer side/

#1177. Maxillule basis <presence of basal exite>/
1. basal exite present/
2. basal exite absent/

#1178. Maxillule basis <presence of setae at basal exite>/
1. with setae/
2. without setae/

#1179. Maxillule basis <number of setae at basal exite>/
1. one element/
2. four elements/

#1180. Maxillule basis <location of setae at basal exite>/
1. on outer surface/
2. on inner surface/

#1181. Maxillule endopod <number of segment>/
1. 1-segmented/
2. 2-segmented/

#1182. Maxillule endopod 1 <arrangement of the segment>/
1. bilobate/
2. not bilobate/

#1183. Maxillule endopod 1 <location of the first lobule>/
1. first proximal/
2. first distal/

#1184. Maxillule endopod 1 <presence of setae>/
1. with setae/
2. without setae/

#1185. Maxillule endopod 1 <number of setae at first lobule>/
1. three elements/
2. four elements/

#1186. Maxillule endopod 1 <location of the second lobule>/
1. second proximal/
2. second distal/

#1187. Maxillule endopod 1 <presence of setae>/
1. with setae/
2. without setae/

#1188. Maxillule endopod 1 <number of setae>/
1. four elements/

2. five elements/

#1189. Maxillule exopod <number of segment>/
1. 1-segmented/
2. 2-segmented/

#1190. Maxillule exopod 1 <presence of setae>/
1. with setae/
2. without setae/

#1191. Maxillule exopod 1 <number of setae>/
1. four elements/
2. six elements/

#1192. Maxillule exopod 1 <presence of setules>/
1. without setules/
2. with setules/

#1193. Maxillule exopod 1 <arrangement of setules>/
1. as a row/
2. as a patch/

#1194. Maxillule exopod 1 <location of setules>/
1. on inner side/
2. on outer side/

#1195. Maxillule exopod 1 <presence of spinules>/
1. spinules present/
2. spinules absent/

#1196. Maxillule exopod 1 <location of the setules>/
1. on anterior surface/
2. on posterior surface/

#1197. Maxillule exopod 2 <presence of setae>/
1. with setae/
2. without setae/

#1198. Maxillule exopod 2 <total of setae>/
1. six elements/
2. eleven elements/

#1199. Maxillule exopod 2 <presence of setules>/
1. with setules/
2. without setules/

#1200. Maxillule exopod 2 <arrangement of setules>/
1. as a row/
2. as a patch/

#1201. Maxillule exopod 2 <location of setules>/
1. on inner margin/
2. on outer margin/

#1202. Maxilla <number of rami>/
1. uniramous/
2. biramous/

#1203. Maxilla <segmentation>/
1. 5-segmented/
2. 6-segmented/

#1204. Maxilla praecoxa <fusion>/
1. separated from the coxa/
2. fused to coxa/

#1205. Maxilla praecoxa <completeness of the fusion>/
1. completely/
2. incompletely/

#1206. Maxilla praecoxa <place of distinction of partial fusion>/
1. distinct externally/
2. distinct internally/

#1207. Maxilla praecoxa <presence of praecoxal endite>/
1. with praecoxal endite/
2. without praecoxal endite/

#1208. Maxilla praecoxa <number of praecoxal endite>/
1. single/
2. double/
3. triple/

#1209. Maxilla praecoxa <relation of size of the single praecoxal endite>/
1. elongated (larger width-length ratio 2:1)/
2. not elongated (lesser width-length ratio 2:1)/

#1210. Maxilla praecoxa <insertion of single praecoxal endite>/
1. proximally inserted/
2. distally inserted/

#1211. Maxilla praecoxa <relation of size of the first praecoxal endite>/
1. first elongated endite (larger or equal width length ratio 2:1)/
2. first non-elongated endite (lesser width length ratio 2:1)/

#1212. Maxilla praecoxa <insertion of first praecoxal endite>/
1. proximally inserted/
2. distally inserted/

#1213. Maxilla praecoxa <seta present on the praecoxal endite>/
1. with seta/
2. without seta/

#1214. Maxilla praecoxa <composition of seta on the praecoxal endite>/
1. straight/
2. plumose/

#1215. Maxilla praecoxa <number of straight seta of the praecoxal endite>/
1. 1 straight/
2. 3 straight/

#1216. Maxilla praecoxa <number of plumose seta of the praecoxal endite>/
1. 3 plumose/
2. 4 plumose/
3. 5 plumose/

#1217. Maxilla praecoxa <presence of spine at praecoxal endite>/
1. with spine/
2. without spine/

#1218. Maxilla praecoxa <number of spines at praecoxal endite>/
1. single/
2. double/

#1219. Maxilla praecoxa <presence of spinules at praecoxal endite>/
1. without spinules/
2. with spinules/

#1220. Maxilla praecoxa <arrangement of spinules at praecoxal endite>/
1. as a row/
2. as a patch/

#1221. Maxilla praecoxa <location of spinules at praecoxal endite>/
1. on proximal margin/
2. on distal margin/

#1222. Maxilla praecoxa <presence of setules at praecoxal endite>/
1. without setule/
2. with setule/

#1223. Maxilla praecoxa <arrangement of setules at praecoxal endite>/
1. as a row/
2. as a patch/

#1224. Maxilla praecoxa <insertion of setules at praecoxal endite>/
1. on proximal margin/
2. on distal margin/

#1225. Maxilla praecoxa <relation of size of the second praecoxal endite>/
1. second elongated endite (larger or equal width length ratio 2:1)/
2. second non-elongated endite (lesser width length ratio 2:1)/

#1226. Maxilla praecoxa <insertion of second praecoxal endite>/
1. medially inserted/
2. distally inserted/

#1227. Maxilla praecoxa <presence of seta on the second praecoxal endite>/
1. with seta/
2. without seta/

#1228. Maxilla praecoxa <composition of seta on the second praecoxal endite>/
1. straight/
2. plumose/

#1229. Maxilla praecoxa <number of straight seta of the second praecoxal endite>/
1. 1 straight/
2. 2 straight/

#1230. Maxilla praecoxa <number of plumose setae of the second praecoxal endite>/
1. 2 plumose/
2. 3 plumose/

#1231. Maxilla praecoxa <presence of spines at second praecoxal endite>/
1. with spine/
2. without spine/

#1232. Maxilla praecoxa <number of spines at second praecoxal endite>/
1. single/
2. double/

#1233. Maxilla praecoxa <presence of spinules at second praecoxal endite>/
1. with spinules/
2. without spinules/

#1234. Maxilla praecoxa <arrangement of spinules at second praecoxal endite>/
1. as a row/
2. as a patch/

#1235. Maxilla praecoxa <location of spinules at second praecoxal endite>/
1. on proximal margin/
2. on distal margin/

#1236. Maxilla praecoxa <presence of setules at second praecoxal endite>/
1. with setule/
2. without setule/

#1237. Maxilla praecoxa <arrangement of setules at second praecoxal endite>/
1. as a row/
2. as a patch/

#1238. Maxilla praecoxa <insertion of setules at second praecoxal endite>/
1. on proximal margin/
2. on distal margin/

#1239. Maxilla praecoxa <relation of size of the third praecoxal endite>/
1. third elongated endite (larger or equal width length ratio 2:1)/
2. third non-elongated endite (lesser width length ratio 2:1)/

#1240. Maxilla praecoxa <insertion of third praecoxal endite>/
1. medially inserted/
2. distally inserted/

#1241. Maxilla praecoxa <presence of seta on the third praecoxal endite>/
1. with seta/
2. without seta/

#1242. Maxilla praecoxa <number of straight seta of the third praecoxal endite>/
1. 1 straight/
2. 2 straight/

#1243. Maxilla praecoxa <number of plumose seta of the third praecoxal endite>/
1. 1 plumose/
2. 2 plumose/

#1244. Maxilla praecoxa <presence of outer seta at praecoxal>/
1. presence of outer seta/
2. absence of outer seta/

#1245. Maxilla coxa <presence of coxal endite>/
1. with coxal endite/
2. without coxal endite/

#1246. Maxilla coxa <number of coxal endite>/
1. double/
2. triple/

#1247. Maxilla coxa <relation of size of the first coxal endite>/
1. first elongated endite (larger or equal width)/
2. first non-elongated endite (lesser width-length ratio 2:1)/

#1248. Maxilla coxa <insertion of first coxal endite>/
1. proximally inserted/
2. distally inserted/

#1249. Maxilla coxa <presence of seta on the coxal endite>/
1. with seta/
2. without seta/

#1250. Maxilla coxa <composition of seta on the coxal endite>/
1. straight/
2. plumose/

#1251. Maxilla coxa <number of straight seta of the coxal endite>/
1. 1 straight/
2. 2 straight/

#1252. Maxilla coxa <number of plumose seta of the coxal endite>/
1. 1 plumose/
2. 3 plumose/

#1253. Maxilla coxa <presence of spine at coxal endite>/
1. with spine/
2. without spine/

#1254. Maxilla coxa <number of spines at coxal endite>/
1. single/
2. double/

#1255. Maxilla coxa <presence of spinules at coxal endite>/
1. with spinules/
2. without spinules/

#1256. Maxilla coxa <arrangement of spinules at coxal endite>/
1. as a row/
2. as a patch/

#1257. Maxilla coxa <location of spinules at coxal endite>/
1. on proximal margin/
2. on distal margin/

#1258. Maxilla coxa <presence of setules at coxal endite>/
1. with setules/
2. without setules/

#1259. Maxilla coxa <arrangement of setules at coxal endite>/
1. as a row/
2. as a patch/

#1260. Maxilla coxa <insertion of setules at coxal endite>/
1. on proximal margin/
2. on distal margin/

#1261. Maxilla coxa <relation of size of the second coxal endite>/

1. second elongated endite (larger or equal width)/
2. second non-elongated endite (lesser width-length ratio 2:1)/

#1262. Maxilla coxa <insertion of second coxal endite>/
1. medially inserted/
2. distally inserted/

#1263. Maxilla coxa <presence of seta on the second coxal endite>/
1. with seta/
2. without seta/

#1264. Maxilla coxa <composition of seta on the second coxal endite>/
1. straight/
2. plumose/

#1265. Maxilla coxa <number of straight setae of the second coxal endite>/
1. 2 straight/
2. 3 straight/

#1266. Maxilla coxa <number of plumose seta of the second coxal endite>/
1. 1 plumose/
2. 3 plumose/

#1267. Maxilla coxa <presence of spine at second coxal endite>/
1. with spine/
2. without spine/

#1268. Maxilla coxa <number of spines at second coxal endite>/
1. single/
2. double/

#1269. Maxilla coxa <presence of spinules at second coxal endite>/
1. with spinules/
2. without spinules/

#1270. Maxilla coxa <arrangement of spinules at second coxal endite>/
1. as a row/
2. as a patch/

#1271. Maxilla coxa <location of spinules at second coxal endite>/
1. on proximal margin/
2. on distal margin/

#1272. Maxilla coxa <presence of setules at second coxal endite>/
1. with setules/
2. without setules/

#1273. Maxilla coxa <arrangement of setules at second coxal endite>/
1. as a row/

2. as a patch/

#1274. Maxilla coxa <insertion of setules at second coxal endite>/
1. on proximal margin/
2. on distal margin/

#1275. Maxilla coxa <presence of outer seta at coxal endite>/
1. presence of outer seta/
2. absence of outer seta/

#1276. Maxilla coxa <relation of size of the triple coxal endite>/
1. third elongated (larger width length ratio 2:1)/
2. third not elongated (lesser width length ratio 2:1)/

#1277. Maxilla coxa <insertion of single coxal endite>/
1. proximally inserted/
2. distally inserted/

#1278. Maxilla coxa <presence of plumose seta on the third coxal endite>/
1. with one seta/
2. with three setae/

#1279. Maxilla coxa <composition of the seta on the third coxal endite>/
1. plumose/
2. straight/

#1280. Maxilla basis <presence of basal endite>/
1. with basal endite/
2. without basal endite/

#1281. Maxilla basis <number of basis endite>/
1. single/
2. double/

#1282. Maxilla basis <relation of size of the basal endite>/
1. elongated (larger or equal width-length ratio 2:1)/
2. not elongated (lesser width-length ratio 2:1)/

#1283. Maxilla basis <presence of seta on the basal endite>/
1. with seta/
2. without seta/

#1284. Maxilla basis <composition of seta on the basal endite>/
1. straight/
2. plumose/

#1285. Maxilla basis <number of straight seta of the basis endite>/
1. 1 straight/
2. 2 straight/

#1286. Maxilla basis <number of plumose seta of the basal endite>/
1. 3 plumose/
2. 4 plumose/

#1287. Maxilla basis <presence of spinules at basal endite>/
1. with spinules/
2. without spinules/

#1288. Maxilla basis <presence of outer seta at basal endite>/
1. presence of outer seta/
2. absence of outer seta/

#1289. Maxilla endopod <segmentation>/
1. 2-segmented/
2. 3-segmented/

#1290. Maxilla endopod 1 <presence of seta>/
1. with seta/
2. without seta/

#1291. Maxilla endopod 1 <composition of seta>/
1. <straight>/
2. <plumose>/

#1292. Maxilla endopod 1 <number of straight seta>/
1. 1 straight/
2. 2 straight/

#1293. Maxilla endopod 1 <number of plumose seta>/
1. 1 plumoses/
2. 2 plumoses/

#1294. Maxilla endopod 1 <presence of spine>/
1. with spine/
2. without spine/

#1295. Maxilla endopod 1 <number of spine>/
1. 1 element/
2. 2 elements/

#1296. Maxilla endopod 1 <presence of spinules>/
1. with spinules/
2. without spinules/

#1297. Maxilla endopod 1 <arrangement of spinules>/
1. as a row/
2. as a patch/

#1298. Maxilla endopod 1 <location of spinules>/
1. on proximal margin/
2. on distal margin/

#1299. Maxilla endopod 1 <presence of setules>/
1. with setules/
2. without setules/

#1300. Maxilla endopod 1 <arrangement of setules>/
1. as a row/
2. as a patch/

#1301. Maxilla endopod 1 <location of setules>/
1. on proximal margin/
2. on distal margin/

#1302. Maxilla endopod 2 <presence of seta>/
1. with seta/
2. without seta/

#1303. Maxilla endopod 2 <composition of seta>/
1. <straight>/
2. <plumose>/

#1304. Maxilla endopod 2 <number of straight seta>/
1. 1 straight/
2. 2 straight/

#1305. Maxilla endopod 2 <number of plumose seta>/
1. 1 plumose/
2. 2 plumose/

#1306. Maxilla endopod 2 <presence of spine>/
1. with spine/
2. without spine/

#1307. Maxilla endopod 2 <number of spine>/
1. 1 element/
2. 2 elements/

#1308. Maxilla endopod 3 <composition of the seta>/
1. plumose/
2. straight/

#1309. Maxilla endopod 2 <presence of spinules>/
1. with spinules/
2. without spinules/

#1310. Maxilla endopod 2 <arrangement of spinules>/

1. as a row/
2. as a patch/

#1311. Maxilla endopod 2 <location of spinules>/
1. on proximal margin/
2. on distal margin/

#1312. Maxilla endopod 2 <presence of setules>/
1. with setules/
2. without setules/

#1313. Maxilla endopod 2 <arrangement of setules>/
1. as a row/
2. as a patch/

#1314. Maxilla endopod 2 <location of setules>/
1. on proximal margin/
2. on distal margin/

#1315. Maxilla endopod 3 <quantify of seta>/
1. with 2 setae/
2. with 3 setae/

#1316. Maxilliped <number of rami>/
1. uniramous/
2. biramous/

#1317. Maxilliped <segmentation>/
1. 8-segmented/
2. 9-segmented/

#1318. Maxilliped praecoxa <fusion>/
1. separated from the coxa/
2. fused to coxa/

#1319. Maxilliped praecoxa <definition of the fusion>/
1. completely/
2. incompletely/

#1320. Maxilliped praecoxa <place of distinction of partial fusion>/
1. distinct externally/
2. distinct internally/

#1321. Maxilliped praecoxa <presence of praecoxal endite>/
1. with praecoxal endite/
2. without praecoxal endite/

#1322. Maxilliped praecoxa <relation of size of praecoxal endite>/
1. elongated (larger or equal width-length ratio 2:1)/

2. not elongated (lesser width-length ratio 2:1)/

#1323. Maxilliped praecoxa <insertion of the praecoxal endite>/
1. proximally inserted/
2. distally inserted/

#1324. Maxilliped praecoxa <presence of seta at praecoxal endite>/
1. with seta/
2. without seta/

#1325. Maxilliped praecoxa <composition of seta at praecoxal endite>/
1. <straight>/
2. <plumose>/

#1326. Maxilliped praecoxa <number of straight seta at praecoxal endite>/
1. 1 straight/
2. 2 straight/

#1327. Maxilliped praecoxa <number of plumose seta at praecoxal endite>/
1. 1 plumose/
2. 2 plumose/

#1328. Maxilliped praecoxa <presence of spinules at praecoxal endite>/
1. with spinules/
2. without spinules/

#1329. Maxilliped praecoxa <arrangement of spinules at praecoxal endite>/
1. as a row/
2. as a patch/

#1330. Maxilliped praecoxa <number of arrangement of spinules at praecoxal endite>/
1. single/
2. double/

#1331. Maxilliped praecoxa <location of spinules at praecoxal endite>/
1. on basal surface/
2. on apical surface/

#1332. Maxilliped praecoxa <presence of setules at praecoxal endite>/
1. with setules/
2. without setules/

#1333. Maxilliped praecoxa <arrangement of setules at praecoxal endite>/
1. as a row/
2. as a patch/

#1334. Maxilliped praecoxa <number of arrangment of setules at praecoxal endite>/

1. single/
2. double/

#1335. Maxilliped praecoxa <location of setules at praecoxal endite>/
1. on inner side/
2. on outer side/

#1336. Maxilliped coxa <presence of coxal endite>/
1. with coxal endite/
2. without coxal endite/

#1337. Maxilliped coxa <number of coxal endite>/
1. two coxal endite/
2. three coxal endite/

#1338. Maxilliped coxa <relation of size of coxal endite 1>/
1. first elongated (larger or equal width)/
2. first not elongated (lesser width-length ratio 2:1)/

#1339. Maxilliped coxa <insertion of the coxal endite 1>/
1. proximally inserted/
2. distally inserted/

#1340. Maxilliped coxa <presence of seta at coxal endite 1>/
1. with seta/
2. without seta/

#1341. Maxilliped coxa <composition of seta at coxal endite 1>/
1. <straight>/
2. <plumose>/

#1342. Maxilliped coxa <number of straight setae at coxal endite 1>/
1. 1 straight/
2. 2 straight/

#1343. Maxilliped coxa <number of plumose setae at coxal endite 1>/
1. 1 plumose/
2. 2 plumose/

#1344. Maxilliped coxa <presence of spinules at coxal endite 1>/
1. with spinules/
2. without spinules/

#1345. Maxilliped coxa <arrangement of spinules at coxal endite 1>/
1. as a row/
2. as a patch/

#1346. Maxilliped coxa <number of arrangement of spinules at coxal endite 1>/
1. single/

2. double/

#1347. Maxilliped coxa <location of spinules at coxal endite 1>/
1. on medial surface/
2. on apical surface/

#1348. Maxilliped coxa <presence of setules at coxal endite 1>/
1. with setules/
2. without setules/

#1349. Maxilliped coxa <arrangement of setules at coxal endite 1>/
1. as a row/
2. as a patch/

#1350. Maxilliped coxa <number of arrangment of setules at coxal endite 1>/
1. single/
2. double/

#1351. Maxilliped coxa <location of setules at coxal endite 1>/
1. on inner margin/
2. on outer margin/

#1352. Maxilliped coxa <relation of size of coxal endite 2>/
1. second elongated (larger or equal width)/
2. second not elongated (lesser width-length ratio 2:1)/

#1353. Maxilliped coxa <insertion of the coxal endite 2>/
1. medially inserted/
2. distally inserted/

#1354. Maxilliped coxa <presence of seta at coxal endite 2>/
1. with seta/
2. without seta/

#1355. Maxilliped coxa <composition of seta at coxal endite 2>/
1. <straight>/
2. <plumose>/

#1356. Maxilliped coxa <number of straight setae at coxal endite 2>/
1. 2 straight/
2. 3 straight/

#1357. Maxilliped coxa <number of plumose setae at coxal endite 2>/
1. 3 plumose/
2. 4 plumose/

#1358. Maxilliped coxa <presence of spinules at coxal endite 2>/
1. with spinules/
2. without spinules/

#1359. Maxilliped coxa <arrangement of spinules at coxal endite 2>/
1. as a row/
2. as a patch/

#1360. Maxilliped coxa <number of arrangement of spinules at coxal endite 2>/
1. single/
2. double/

#1361. Maxilliped coxa <location of spinules at coxal endite 2>/
1. on basal surface/
2. on medial surface/
3. on apical surface/

#1362. Maxilliped coxa <presence of setules at coxal endite 2>/
1. with setules/
2. without setules/

#1363. Maxilliped coxa <arrangement of setules at coxal endite 2>/
1. as a row/
2. as a patch/

#1364. Maxilliped coxa <number of arrangment of setules at coxal endite 2>/
1. single/
2. double/

#1365. Maxilliped coxa <location of setules at coxal endite 2>/
1. on inner margin/
2. on outer margin/

#1366. Maxilliped coxa <relation of size of coxal endite 3>/
1. third elongated (larger or equal width length ratio 2:1)/
2. third not elongated (lesser width-length ratio 2:1)/

#1367. Maxilliped coxa <insertion of the coxal endite 3>/
1. proximally inserted/
2. distally inserted/

#1368. Maxilliped coxa <presence of seta at coxal endite 3>/
1. with seta/
2. without seta/

#1369. Maxilliped coxa <composition of seta at coxal endite 3>/
1. <straight>/
2. <plumose>/

#1370. Maxilliped coxa <number of straight seta at coxal endite 3>/
1. 3 straight/
2. 4 straight/

#1371. Maxilliped coxa <number of plumose seta at coxal endite 3>/
1. 3 plumose/
2. 4 plumose/

#1372. Maxilliped coxa <reach of the plumose seta at coxal endite 3>/
1. at last one reaching to beyond of the basis/
2. none reaching to beyond of the basis/

#1373. Maxilliped coxa <presence of spinules at coxal endite 3>/
1. with spinules/
2. without spinules/

#1374. Maxilliped coxa <arrangement of spinules at coxal endite 3>/
1. as a row/
2. as a patch/

#1375. Maxilliped coxa <number of arrangement of spinules at coxal endite 3>/
1. single/
2. double/

#1376. Maxilliped coxa <location of spinules at coxal endite 3>/
1. on basal surface/
2. on apical surface/

#1377. Maxilliped coxa <presence of setules at coxal endite 3>/
1. with setules/
2. without setules/

#1378. Maxilliped coxa <arrangement of setules at coxal endite 3>/
1. as a row/
2. as a patch/

#1379. Maxilliped coxa <number of arrangment of setules at coxal endite 3>/
1. single/
2. double/

#1380. Maxilliped coxa <presence of lobe>/
1. with lobe/
2. without lobe/

#1381. Maxilliped coxa <development of the lobe>/
1. prominence/
2. not prominence/

#1382. Maxilliped coxa <location of the lobe>/
1. at inner distal angle/
2. at inner distal surface/

#1383. Maxilliped coxa <presence of ornamentation at lobe>/
1. ornamented/
2. not ornamented/

#1384. Maxilliped coxa <ornamentation of the lobe>/
1. with setules/
2. with spinules/

#1385. Maxilliped coxa <location of the ornamentation on lobe>/
1. continuously on margin/
2. discontinuously on margin/

#1386. Maxilliped basis <presence of basal endite>/
1. with basal endite/
2. without basal endite/

#1387. Maxilliped basis <relation of size of basal endite>/
1. not elongated (lesser width-length ratio 2:1)/
2. elongated (larger width-length ratio 2:1)/

#1388. Maxilliped basis <insertion of the basal endite>/
1. proximally inserted/
2. distally inserted/

#1389. Maxilliped basis <presence of seta>/
1. with seta/
2. without seta/

#1390. Maxilliped basis <composition of seta>/
1. <straight>/
2. <plumose>/

#1391. Maxilliped basis <number of straight setae>/
1. 2 straight/
2. 3 straight/

#1392. Maxilliped basis <number of plumose setae>/
1. 3 plumose/
2. 4 plumose/

#1393. Maxilliped basis <presence of spinules>/
1. with spinules/
2. without spinules/

#1394. Maxilliped basis <arrangement of spinules>/
1. as a row/
2. as a patch/

#1395. Maxilliped basis <number of arrangement of spinules>/

1. single/
2. double/

#1396. Maxilliped basis <location of spinules>/
1. on medial surface/
2. on apical surface/

#1397. Maxilliped basis <presence of setules>/
1. with setules/
2. without setules/

#1398. Maxilliped basis <arrangement of setules>/
1. as a row/
2. as a patch/

#1399. Maxilliped basis <number of arrangment of setules>/
1. single/
2. double/

#1400. Maxilliped basis <location of setules>/
1. on inner margin/
2. on outer margin/

#1401. Maxilliped endopod segment <segmentation>/
1. 6-segmented/
2. 5-segmented/

#1402. Maxilliped endopod segment 1 <presence of seta>/
1. with seta/
2. without seta/

#1403. Maxilliped endopod segment 1 <composition of seta>/
1. <straight>/
2. <plumose>/

#1404. Maxilliped endopod segment 1 <number of straight seta>/
1. 1 straight/
2. 2 straight/

#1405. Maxilliped endopod segment 1 <location of straight seta>/
1. on inner surface/
2. on outer surface/

#1406. Maxilliped endopod segment 1 <number of plumose seta>/
1. 1 plumose/
2. 2 plumose/

#1407. Maxilliped endopod segment 1 <location of plumose seta>/
1. on inner surface/

2. on outer surface/

#1408. Maxilliped endopod segment 2 <presence of seta>/
1. with seta/
2. without seta/

#1409. Maxilliped endopod segment 2 <composition of seta>/
1. <straight>/
2. <plumose>/

#1410. Maxilliped endopod segment 2 <number of straight seta>/
1. 3 straight/
2. 4 straight/

#1411. Maxilliped endopod segment 2 <location of straight seta>/
1. on inner surface/
2. on outer surface/

#1412. Maxilliped endopod segment 2 <number of plumose seta>/
1. 2 plumose/
2. 3 plumose/

#1413. Maxilliped endopod segment 2 <location of plumose seta>/
1. on inner surface/
2. on outer surface/

#1414. Maxilliped endopod segment 3 <presence of seta>/
1. with seta/
2. without seta/

#1415. Maxilliped endopod segment 3 <composition of seta>/
1. <straight>/
2. <plumose>/

#1416. Maxilliped endopod segment 3 <number of straight seta>/
1. 2 straight/
2. 3 straight/

#1417. Maxilliped endopod segment 3 <location of straight seta>/
1. on inner surface/
2. on outer surface/

#1418. Maxilliped endopod segment 3 <number of plumose seta>/
1. 1 plumose/
2. 2 plumose/

#1419. Maxilliped endopod segment 3 <location of plumose seta>/
1. on inner surface/
2. on outer surface/

#1420. Maxilliped endopod segment 4 <presence of seta>/
1. with seta/
2. without seta/

#1421. Maxilliped endopod segment 4 <composition of seta>/
1. <straight>/
2. <plumose>/

#1422. Maxilliped endopod segment 4 <number of straight seta>/
1. 2 straight/
2. 3 straight/

#1423. Maxilliped endopod segment 4 <location of straight seta>/
1. on inner surface/
2. on outer surface/

#1424. Maxilliped endopod segment 4 <number of plumose seta>/
1. 1 plumose/
2. 2 plumose/

#1425. Maxilliped endopod segment 4 <location of plumose seta>/
1. on inner surface/
2. on outer surface/

#1426. Maxilliped endopod segment 5 <presence of seta>/
1. with seta/
2. without seta/

#1427. Maxilliped endopod segment 5 <composition of seta>/
1. <straight>/
2. <plumose>/

#1428. Maxilliped endopod segment 5 <number of straight seta>/
1. 1 straight/
2. 2 straight/

#1429. Maxilliped endopod segment 5 <location of straight seta>/
1. on inner surface/
2. on outer surface/

#1430. Maxilliped endopod segment 5 <number of plumose seta>/
1. 2 plumose/
2. 4 plumose/

#1431. Maxilliped endopod segment 5 <location of plumose seta>/
1. on inner surface/
2. on outer surface/

#1432. Maxilliped endopod segment 5 <observation of outer seta>/
1. outer seta present/
2. outer seta absent/

#1433. Maxilliped endopod segment 6 <presence of seta>/
1. with seta/
2. without seta/

#1434. Maxilliped endopod segment 6 <composition of seta>/
1. <straight>/
2. <plumose>/

#1435. Maxilliped endopod segment 6 <number of straight seta>/
1. 3 straight/
2. 4 straight/

#1436. Maxilliped endopod segment 6 <location of straight seta>/
1. on inner surface/
2. on outer surface/

#1437. Maxilliped endopod segment 6 <number of plumose seta>/
1. 3 plumose/
2. 4 plumose/

#1438. Maxilliped endopod segment 6 <location of plumose seta>/
1. on inner surface/
2. on outer surface/

#1439. First swimming legs <symmetry>/
1. symmetrical/
2. asymmetrical/

#1440. First swimming legs <number of rami>/
1. uniramous/
2. biramous/

#1441. First swimming legs intercoxal plate <presence of seta>/
1. with seta/
2. without seta/

#1442. First swimming legs intercoxal plate <position of seta>/
1. centralized/
2. laterally/

#1443. First swimming legs intercoxal plate <location of setae>/
1. on anterior surface/
2. on posterior surface/

#1444. First swimming legs praecoxa <presence of praecoxa>/

1. present/
2. absent/

#1445. First swimming legs praecoxa <location of praecoxa>/
1. laterally located/
2. not laterally located/

#1446. First swimming legs coxa <presence of seta>/
1. with seta/
2. without seta/

#1447. First swimming legs coxa <number of setae>/
1. one/
2. two/

#1448. First swimming legs coxa <composition of the seta>/
1. plumose/
2. straight/

#1449. First swimming legs coxa <insertion of seta>/
1. proximally inserted/
2. distally inserted/

#1450. First swimming legs coxa <location of setae>/
1. on inner surface/
2. on outer surface/

#1451. First swimming legs coxa <relative length>/
1. surpassing to basal segment/
2. surpassing to first endopodal segment/

#1452. First swimming legs coxa <presence of setules>/
1. with setules/
2. without setules/

#1453. First swimming legs coxa <quantify of groups of setules>/
1. one group/
2. two group/

#1454. First swimming legs coxa <arrangement of setules>/
1. as a row/
2. as a patch/

#1455. First swimming legs coxa <continuity of insertion of setules>/
1. continuously/
2. discontinuously/

#1456. First swimming legs coxa <location of setules>/
1. on inner margin/

2. on outer margin/

#1457. First swimming legs coxa <arrangement of second group of setules>/
1. and as a row/
2. and as a patch/

#1458. First swimming legs coxa <number of arrangement of the second group setules>/
1. single/
2. double/

#1459. First swimming legs coxa <location of the second group setules>/
1. on anterior surface/
2. on posterior surface/

#1460. First swimming legs coxa <position of the second group setules>/
1. innerly/
2. outerly/

#1461. First swimming legs coxa <presence of spinules>/
1. with spinules/
2. without spinules/

#1462. First swimming legs coxa <arrangement of spinules>/
1. as a row/
2. as a patch/

#1463. First swimming legs coxa <number of spinules>/
1. single/
2. double/

#1464. First swimming legs coxa <inserted of spinules>/
1. medially inserted/
2. distally inserted/

#1465. First swimming legs coxa <location of spinules>/
1. at anterior surface/
2. at posterior surface/

#1466. First swimming legs coxa <presence of spine>/
1. with spine/
2. without spine/

#1467. First swimming legs basis <presence of seta>/
1. with seta/
2. without seta/

#1468. First swimming legs basis <number of setae>/
1. one/

2. two/

#1469. First swimming legs basis <insertion of seta>/
1. medially inserted/
2. distally inserted/

#1470. First swimming legs basis <location of setae>/
1. on inner surface/
2. on outer surface/

#1471. First swimming legs basis <relative length of the seta>/
1. surpassing to distal limit of the segment/
2. not surprassing to distal limit of the segment/

#1472. First swimming legs basis <presence of setules>/
1. with setules/
2. without setules/

#1473. First swimming legs basis <arrangement of setules>/
1. as a row/
2. as a patch/

#1474. First swimming legs basis <number of setules>/
1. single/
2. double/

#1475. First swimming legs basis <continuity of insertion of setules>/
1. continuously/
2. discontinuously/

#1476. First swimming legs basis <location of setules>/
1. on inner surface/
2. on outer surface/

#1477. First swimming legs basis <presence of spinules>/
1. with spinules/
2. without spinules/

#1478. First swimming legs basis <presence of spine>/
1. with spine/
2. without spine/

#1479. First swimming legs endopod <number of segmentation>/
1. 2-segmented/
2. 3-segmented/

#1480. First swimming legs endopod 1 <presence of seta>/
1. with seta/
2. without seta/

#1481. First swimming legs endopod 1 <seta composition>/
1. plumose/
2. straight/

#1482. First swimming legs endopod 1 <restriction or not of the seta>/
1. restricted/
2. unrestricted/

#1483. First swimming legs endopod 1 <location of the restricted seta>/
1. to inner surface/
2. to outer surface/

#1484. First swimming legs endopod 1 <number of setae on inner margin>/
1. one element/
2. two elements/

#1485. First swimming legs endopod 1 <presence of spine>/
1. with spine/
2. without spine/

#1486. First swimming legs endopod 1 <number of spine>/
1. 1/
2. 2/

#1487. First swimming legs endopod 1 <form of spine>/
1. stout/
2. slender/

#1488. First swimming legs endopod 1 <size of spine>/
1. equal size/
2. unequal size/

#1489. First swimming legs endopod 1 <relative length of the spine>/
1. longer 2x than origin segment/
2. not longer 2x than origin segment/

#1490. First swimming legs endopod 1 <relative length of the spine 1>/
1. one longer 2x than origin segment/
2. one not longer 2x than origin segment/

#1491. First swimming legs endopod 1 <relative length of the spine 2>/
1. and other longer 2x than origin segment/
2. and other not longer 2x than origin segment/

#1492. First swimming legs endopod 1 <type of edge of the spine>/
1. serrated/
2. not serrated/

#1493. First swimming legs endopod 1 <location of the composition of the spine>/
1. equally/
2. unequally/

#1494. First swimming legs endopod 1 <location of the composition of both spine>/
1. on inner edge/
2. on outer edge/

#1495. First swimming legs endopod 1 <location of the serrated spine 1>/
1. one on inner edge/
2. one on outer edge/

#1496. First swimming legs endopod 1 <location of the serrated spine 2>/
1. other on inner edge/
2. other on outer edge/

#1497. First swimming legs endopod 1 <ornamentation of the spine with non-serrated edge>/
1. with ornamentation on edge non-serrated/
2. without ornamentation on edge non-serrated/

#1498. First swimming legs endopod 1 <type pf ornamentation of the spine>/
1. of spinules/
2. of setules/

#1499. First swimming legs endopod 1 <presence of setules>/
1. with setules/
2. without setules/

#1500. First swimming legs endopod 1 <arrangement of setules>/
1. as a row/
2. as a patch/

#1501. First swimming legs endopod 1 <number of setules>/
1. single/
2. double/

#1502. First swimming legs endopod 1 <continuity of insertion of setules>/
1. continuously/
2. discontinuously/

#1503. First swimming legs endopod 1 <location of setules>/
1. on inner surface/
2. on outer surface/

#1504. First swimming legs endopod 1 <presence of spinules>/
1. with spinules/
2. without spinules/

#1505. First swimming legs endopod 1 <arrangement of spinules>/
1. as a row/
2. as a patch/

#1506. First swimming legs endopod 1 <number of spinules>/
1. single/
2. double/

#1507. First swimming legs endopod 1 <inserted of spinules>/
1. proximally inserted/
2. medially inserted/
3. distally inserted/

#1508. First swimming legs endopod 1 <location of spinules>/
1. at anterior surface/
2. at posterior surface/

#1509. First swimming legs endopod 1 <presence of Schmeil's organ>/
1. presence of Schmeil's organ/
2. absence of Schmeil's organ/

#1510. First swimming legs endopod 1 <location of Schmeil's organ>/
1. on anterior surface/
2. on posterior surface/

#1511. First swimming legs endopod 2 <presence of seta>/
1. with seta/
2. without seta/

#1512. First swimming legs endopod 2 <restriction or not of the seta>/
1. restricted/
2. unrestricted/

#1513. First swimming legs endopod 2 <insertion of the restricted seta>/
1. <to inner surface>/
2. <to outer surface>/
3. <to distal surface>/

#1514. First swimming legs endopod 2 <number of setae on inner margin>/
1. one on inner surface/
2. three on inner surface/

#1515. First swimming legs endopod 2 <number of setae on outer margin>/
1. one on outer surface/
2. two on outer surface/

#1516. First swimming legs endopod 2 <number of setae on distal margin>/
1. one on distal surface/

2. two on distal surface/

#1517. First swimming legs endopod 2 <setae composition>/
1. plumose/
2. straight/

#1518. First swimming legs endopod 2 <presence of spine>/
1. with spine/
2. without spine/

#1519. First swimming legs endopod 2 <number of spine>/
1. 1/
2. 2/

#1520. First swimming legs endopod 2 <form of spine>/
1. stout/
2. slender/

#1521. First swimming legs endopod 2 <size of spine>/
1. equal size/
2. unequal size/

#1522. First swimming legs endopod 2 <relative length of the spine>/
1. longer 2x than origin segment/
2. not longer 2x than origin segment/

#1523. First swimming legs endopod 2 <relative length of the spine 1>/
1. one longer 2x than origin segment/
2. one not longer 2x than origin segment/

#1524. First swimming legs endopod 2 <relative length of the spine 2>/
1. and other longer 2x than origin segment/
2. and other not longer 2x than origin segment/

#1525. First swimming legs endopod 2 <type of edge of the spine>/
1. serrated/
2. not serrated/

#1526. First swimming legs endopod 2 <location of the composition of the spine>/
1. equally/
2. unequally/

#1527. First swimming legs endopod 2 <location of the composition of both spine>/
1. on inner edge/
2. on outer edge/

#1528. First swimming legs endopod 2 <location of the serrated spine 1>/
1. one on inner edge/

2. one on outer edge/

#1529. First swimming legs endopod 2 <location of the serrated spine 2>/
1. other on inner edge/
2. other on outer edge/

#1530. First swimming legs endopod 2 <ornamentation of the spine with non-serrated edge>/
1. with ornamentation on edge non-serrated/
2. without ornamentation on edge non-serrated/

#1531. First swimming legs endopod 2 <type pf ornamentation of the spine>/
1. of spinules/
2. of setules/

#1532. First swimming legs endopod 2 <presence of setules>/
1. with setules/
2. without setules/

#1533. First swimming legs endopod 2 <arrangement of setules>/
1. as a row/
2. as a patch/

#1534. First swimming legs endopod 2 <number of setules>/
1. single/
2. double/

#1535. First swimming legs endopod 2 <continuity of insertion of setules>/
1. continuously/
2. discontinuously/

#1536. First swimming legs endopod 2 <location of setules>/
1. on inner surface/
2. on outer surface/

#1537. First swimming legs endopod 2 <presence of spinules>/
1. with spinules/
2. without spinules/

#1538. First swimming legs endopod 2 <arrangement of spinules>/
1. as a row/
2. as a patch/

#1539. First swimming legs endopod 2 <number of spinules>/
1. single/
2. double/

#1540. First swimming legs endopod 2 <inserted of spinules>/
1. proximally inserted/

2. medially inserted/
3. distally inserted/

#1541. First swimming legs endopod 2 <location of spinules>/
1. at anterior surface/
2. at posterior surface/

#1542. First swimming legs endopod 2 <presence of Schmeil's organ>/
1. presence of Schmeil's organ/
2. absence of Schmeil's organ/

#1543. First swimming legs endopod 2 <location of Schmeil's organ>/
1. on anterior surface/
2. on posterior surface/

#1544. First swimming legs endopod 3 <presence of the segment>/
1. presence/
2. absence/

#1545. First swimming legs exopod 1 <presence of seta>/
1. with seta/
2. without seta/

#1546. First swimming legs exopod 1 <restriction or not of the seta>/
1. restricted/
2. unrestricted/

#1547. First swimming legs exopod 1 <insertion of the restricted seta>/
1. <on inner surface>/
2. <on outer surface>/

#1548. First swimming legs exopod 1 <total of seta on inner margin>/
1. 1 on inner surface/
2. 2 on inner surface/

#1549. First swimming legs exopod 1 <total of seta on outer margin>/
1. 1 on outer surface/
2. 2 on outer surface/

#1550. First swimming legs exopod 1 <presence of spine>/
1. with spine/
2. without spine/

#1551. First swimming legs exopod 1 <total of spine>/
1. 1/
2. 2/

#1552. First swimming legs exopod 1 <form of spine>/
1. stout/

2. slender/

#1553. First swimming legs exopod 1 <comparative size between spine>/
1. equal size/
2. unequal size/

#1554. First swimming legs exopod 1 <relative length of the spine>/
1. larger than original segment/
2. smaller than original segment/

#1555. First swimming legs exopod 1 <reach of the spine>/
1. reaching to distal-third of the exopod 3/
2. not reaching to distal-third of the exopod 3/

#1556. First swimming legs exopod 1 <relative length of the spine 1>/
1. first longer 2x than origin segment/
2. first not longer 2x than origin segment/

#1557. First swimming legs exopod 1 <relative length of the spine 2>/
1. second longer 2x than origin segment/
2. second not longer 2x than origin segment/

#1558. First swimming legs exopod 1 <shape of the spine>/
1. serrated/
2. not serrated/

#1559. First swimming legs exopod 1 <location of the shape of the spine>/
1. on inner side/
2. on outer side/

#1560. First swimming legs exopod 1 <continuity of the serrated shape spine>/
1. continuously/
2. discontinuously/

#1561. First swimming legs exopod 1 <location of the serrated spine 1>/
1. first on inner side/
2. first on outer side/

#1562. First swimming legs exopod 1 <location of the serrated spine 2>/
1. second on inner side/
2. second on outer side/

#1563. First swimming legs exopod 1 <presence of setules>/
1. with setules/
2. without setules/

#1564. First swimming legs exopod 1 <arrangement of spinules>/
1. as a row/
2. as a patch/

#1565. First swimming legs exopod 1 <composition of the spinules>/
1. single/
2. double/

#1566. First swimming legs exopod 1 <arrangement of the ornamentation of the setules>/
1. as a row/
2. as a patch/

#1567. First swimming legs exopod 1 <location of setules>/
1. innerly/
2. outerly/

#1568. First swimming legs exopod 2 <presence of seta>/
1. with seta/
2. without seta/

#1569. First swimming legs exopod 2 <restriction or not of the seta>/
1. restricted/
2. unrestricted/

#1570. First swimming legs exopod 2 <insertion of the restricted seta>/
1. <to inner surface>/
2. <to outer surface>/

#1571. First swimming legs exopod 2 <total of seta on inner surface>/
1. 1 on inner surface/
2. 2 on inner surface/

#1572. First swimming legs exopod 2 <seta composition>/
1. plumose/
2. straight/

#1573. First swimming legs exopod 2 <total of seta on outer surface>/
1. 1 on outer surface/
2. 2 on outer surface/

#1574. First swimming legs exopod 2 <presence of spine>/
1. with spine/
2. without spine/

#1575. First swimming legs exopod 2 <total of spine>/
1. 1/
2. 2/

#1576. First swimming legs exopod 2 <form of spine>/
1. stout/
2. slender/

#1577. First swimming legs exopod 2 <comparative size of the spines>/
1. equal size/
2. unequal size/

#1578. First swimming legs exopod 2 <relative length of the spine>/
1. longer 2x than origin segment/
2. not longer 2x than origin segment/

#1579. First swimming legs exopod 2 <relative length of the spine 1>/
1. first longer 2x than origin segment/
2. first not longer 2x than origin segment/

#1580. First swimming legs exopod 2 <relative length of the spine 2>/
1. second longer 2x than origin segment/
2. second not longer 2x than origin segment/

#1581. First swimming legs exopod 2 <shape of the spine>/
1. serrated/
2. not serrated/

#1582. First swimming legs exopod 2 <location of the shape of the spine>/
1. on inner side/
2. on outer side/

#1583. First swimming legs exopod 2 <continuity of the serrated shape spine>/
1. continuously/
2. discontinuously/

#1584. First swimming legs exopod 2 <location of the serrated spine 1>/
1. first on inner side/
2. first on outer side/

#1585. First swimming legs exopod 2 <location of the serrated spine 2>/
1. second on inner side/
2. second on outer side/

#1586. First swimming legs exopod 2 <ornamentation of the spine with non-serrated side>/
1. without ornamentation on non-serrated side/
2. with ornamentation on non-serrated side/

#1587. First swimming legs exopod 2 <ornamentation of the spine>/
1. of spinules/
2. of setules/

#1588. First swimming legs exopod 2 <presence of setules>/
1. with setules/
2. without setules/

#1589. First swimming legs exopod 2 <arrangement of setules>/
1. as a row/
2. as a patch/

#1590. First swimming legs exopod 2 <number of setules>/
1. single/
2. double/

#1591. First swimming legs exopod 2 <continuity of insertion of setules>/
1. continuously/
2. discontinuously/

#1592. First swimming legs exopod 2 <location of setules>/
1. on inner margin/
2. on outer margin/

#1593. First swimming legs exopod 2 <presence of spinules>/
1. with spinules/
2. without spinules/

#1594. First swimming legs exopod 3 <presence of setules>/
1. with setule/
2. without setule/

#1595. First swimming legs exopod 3 <arrangement of setules>/
1. as a row/
2. as a patch/

#1596. First swimming legs exopod 3 <composition of setules>/
1. single/
2. double/

#1597. First swimming legs exopod 3 <continuity of insertion of setules>/
1. continuously/
2. discontinuously/

#1598. First swimming legs exopod 3 <location of setules>/
1. on inner surface/
2. on outer surface/

#1599. First swimming legs exopod 3 <presence of spinules>/
1. with spinules/
2. without spinules/

#1600. First swimming legs exopod 3 <presence of seta>/
1. with seta/
2. without seta/

#1601. First swimming legs exopod 3 <restriction or not of the seta>/
1. restricted/
2. unrestricted/

#1602. First swimming legs exopod 3 <insertion of the restricted seta>/
1. <to inner surface>/
2. <to outer surface>/
3. <to terminal surface>/

#1603. First swimming legs exopod 3 <total of setae on inner side>/
1. 1 on inner surface/
2. 2 on inner surface/

#1604. First swimming legs exopod 3 <total of setae on outer margin>/
1. 1 on outer surface/
2. 1 on outer surface/

#1605. First swimming legs exopod 3 <total of seta on terminal surface>/
1. 1 on terminal surface/
2. 2 on terminal surface/

#1606. First swimming legs exopod 3 <presence of spine>/
1. with spine/
2. without spine/

#1607. First swimming legs exopod 3 <total of spine>/
1. 1/
2. 2/

#1608. First swimming legs exopod 3 <comparative size between spines>/
1. equal size/
2. unequal size/

#1609. First swimming legs exopod 3 <relative length of the spine 1>/
1. first longer 2x than origin segment/
2. first not longer 2x than origin segment/

#1610. First swimming legs exopod 3 <form of spine 1>/
1. stout/
2. slender/

#1611. First swimming legs exopod 3 <shape of the spine 1>/
1. serrated/
2. not serrated/

#1612. First swimming legs exopod 3 <location of the shape of the spine 1>/
1. on inner side/
2. on outer side/

#1613. First swimming legs exopod 3 <continuity of the serrated shape spine 1>/
1. equally/
2. unequally/

#1614. First swimming legs exopod 3 <relative length of the spine 2>/
1. second longer 3x than origin segment/
2. second not longer 3x than origin segment/

#1615. First swimming legs exopod 3 <form of spine 2>/
1. stout/
2. slender/

#1616. First swimming legs exopod 3 <shape of the spine 2>/
1. serrated/
2. not serrated/

#1617. First swimming legs exopod 3 <location of the serrated spine 2>/
1. on inner side/
2. on outer side/

#1618. First swimming legs exopod 3 <ornamentation spine 2 with non-serrated side>/
1. with ornamentation on non-serrated side/
2. without ornamentation on non-serrated side/

#1619. First swimming legs exopod 3 <ornamentation type of the spine 2>/
1. by spinules/
2. by setules/

#1620. Second swimming legs <symmetry>/
1. symmetrical/
2. asymmetrical/

#1621. Second swimming legs <number of rami>/
1. uniramous/
2. biramous/

#1622. Second swimming legs intercoxal plate <presence of seta>/
1. with seta/
2. without seta/

#1623. Second swimming legs intercoxal plate <position of seta>/
1. centralized/
2. laterally/

#1624. Second swimming legs intercoxal plate <location of setae>/
1. on anterior surface/
2. on posterior surface/

#1625. Second swimming legs praecoxa <presence of praecoxa>/
1. present/
2. absent/

#1626. Second swimming legs praecoxa <location of praecoxa>/
1. located laterally/
2. not located laterally/

#1627. Second swimming legs coxa <presence of seta>/
1. with seta/
2. without seta/

#1628. Second swimming legs coxa <composition of the seta>/
1. plumose/
2. straight/

#1629. Second swimming legs coxa <insertion of seta>/
1. proximally inserted/
2. distally inserted/

#1630. Second swimming legs coxa <location of setae>/
1. on inner surface/
2. on outer surface/

#1631. Second swimming legs coxa <relative length>/
1. surpassing to basal segment/
2. surpassing to first endopodal segment/

#1632. Second swimming legs coxa <presence of setules>/
1. with setules/
2. without setules/

#1633. Second swimming legs coxa <arrangement of setules>/
1. as a row/
2. as a patch/

#1634. Second swimming legs coxa <number of setules>/
1. single/
2. double/

#1635. Second swimming legs coxa <continuity of insertion of setules>/
1. continuously/
2. discontinuously/

#1636. Second swimming legs coxa <location of setules>/
1. on inner margin/
2. on outer margin/

#1637. Second swimming legs coxa <presence of spinules>/

1. with spinules/
2. without spinules/

#1638. Second swimming legs coxa <arrangement of spinules>/
1. as a row/
2. as a patch/

#1639. Second swimming legs coxa <number of spinules>/
1. single/
2. double/

#1640. Second swimming legs coxa <inserted of spinules>/
1. proximally inserted/
2. distally inserted/

#1641. Second swimming legs coxa <location of spinules>/
1. at anterior surface/
2. at posterior surface/

#1642. Second swimming legs coxa <presence of spine>/
1. with spine/
2. without spine/

#1643. Second swimming legs basis <presence of seta>/
1. with seta/
2. without seta/

#1644. Second swimming legs basis <presence of setules>/
1. with setules/
2. without setules/

#1645. Second swimming legs basis <presence of spinules>/
1. with spinules/
2. without spinules/

#1646. Second swimming legs basis <presence of spine>/
1. with spine/
2. without spine/

#1647. Second swimming legs endopod <number of segmentation>/
1. 2-segmented/
2. 3-segmented/

#1648. Second swimming legs endopod 1 <presence of seta>/
1. with seta/
2. without seta/

#1649. Second swimming legs endopod 1 <seta composition>/
1. plumose/

2. straight/

#1650. Second swimming legs endopod 1 <restriction or not of the seta>/
1. restricted/
2. unrestricted/

#1651. Second swimming legs endopod 1 <insertion of the restricted seta>/
1. <to inner surface>/
2. <to outer surface>/

#1652. Second swimming legs endopod 1 <total of setae on inner surface>/
1. one on inner surface/
2. two on inner surface/

#1653. Second swimming legs endopod 1 <number of setae on outer surface>/
1. one on outer surface/
2. two on outer surface/

#1654. Second swimming legs endopod 1 <presence of spine>/
1. with spine/
2. without spine/

#1655. Second swimming legs endopod 1 <presence of setules>/
1. with setules/
2. without setules/

#1656. Second swimming legs endopod 1 <arrangement of setules>/
1. as a row/
2. as a patch/

#1657. Second swimming legs endopod 1 <number of setules>/
1. single/
2. double/

#1658. Second swimming legs endopod 1 <continuity of insertion of setules>/
1. continuously/
2. discontinuously/

#1659. Second swimming legs endopod 1 <location of setules>/
1. on inner surface/
2. on outer surface/

#1660. Second swimming legs endopod 1 <presence of spinules>/
1. with spinules/
2. without spinules/

#1661. Second swimming legs endopod 1 <arrangement of spinules>/
1. as a row/
2. as a patch/

#1662. Second swimming legs endopod 1 <number of spinules>/
1. single/
2. double/

#1663. Second swimming legs endopod 1 <inserted of spinules>/
1. proximally inserted/
2. distally inserted/

#1664. Second swimming legs endopod 1 <location of spinules>/
1. at anterior surfaces/
2. at posterior surfaces/

#1665. Second swimming legs endopod 1 <presence of Schmeil's organ>/
1. presence of Schmeil's organ/
2. absence of Schmeil's organ/

#1666. Second swimming legs endopod 1 <location of Schmeil's organ>/
1. on anterior surface/
2. on posterior surface/

#1667. Second swimming legs endopod 2 <presence of seta>/
1. with seta/
2. without seta/

#1668. Second swimming legs endopod 2 <setae composition>/
1. straight/
2. plumose/

#1669. Second swimming legs endopod 2 <restriction or not of the seta>/
1. restricted/
2. unrestricted/

#1670. Second swimming legs endopod 2 <insertion of the restricted seta>/
1. <to inner surface>/
2. <to outer surface>/

#1671. Second swimming legs endopod 2 <total of seta on inner surface>/
1. one on inner surface/
2. two on inner surface/

#1672. Second swimming legs endopod 2 <total of seta on outer surface>/
1. one on outer surface/
2. two on outer surface/

#1673. Second swimming legs endopod 2 <presence of spine>/
1. with spine/
2. without spine/

#1674. Second swimming legs endopod 2 <presence of setules>/
1. with setules/
2. without setules/

#1675. Second swimming legs endopod 2 <arrangement of setules>/
1. as a row/
2. as a patch/

#1676. Second swimming legs endopod 2 <number of setules>/
1. single/
2. double/

#1677. Second swimming legs endopod 2 <continuity of insertion of setules>/
1. continuously/
2. discontinuously/

#1678. Second swimming legs endopod 2 <location of setules>/
1. on inner side/
2. on outer side/

#1679. Second swimming legs endopod 2 <presence of spinules>/
1. with spinules/
2. without spinules/

#1680. Second swimming legs endopod 2 <arrangement of spinules>/
1. as a row/
2. as a patch/

#1681. Second swimming legs endopod 2 <number of spinules>/
1. single/
2. double/

#1682. Second swimming legs endopod 2 <inserted of spinules>/
1. innerly inserted/
2. outerly inserted/

#1683. Second swimming legs endopod 2 <presence of Schmeil's organ>/
1. presence of Schmeil's organ/
2. absence of Schmeil's organ/

#1684. Second swimming legs endopod 2 <location of Schmeil's organ>/
1. on anterior surface/
2. on posterior surface/

#1685. Second swimming legs endopod 3 <presence of seta>/
1. with seta/
2. without seta/

#1686. Second swimming legs endopod 3 <seta composition>/

1. straight/
2. plumose/

#1687. Second swimming legs endopod 3 <restriction or not of the seta>/
1. restricted/
2. unrestricted/

#1688. Second swimming legs endopod 3 <insertion of the restricted seta>/
1. <to inner surface>/
2. <to outer surface>/
3. <to terminal surface>/

#1689. Second swimming legs endopod 3 <number of setae on inner surface>/
1. two on inner surface/
2. three on inner surface/

#1690. Second swimming legs endopod 3 <number of setae on outer surface>/
1. two on outer surface/
2. three on outer surface/

#1691. Second swimming legs endopod 3 <number of setae on terminal surface>/
1. one on distal surface/
2. two on distal surface/

#1692. Second swimming legs endopod 3 <presence of spine>/
1. with spine/
2. without spine/

#1693. Second swimming legs endopod 3 <presence of setules>/
1. with setules/
2. without setules/

#1694. Second swimming legs endopod 3 <arrangement of setules>/
1. as a row/
2. as a patch/

#1695. Second swimming legs endopod 3 <number of setules>/
1. single/
2. double/

#1696. Second swimming legs endopod 3 <continuity of insertion of setules>/
1. continuously/
2. discontinuously/

#1697. Second swimming legs endopod 3 <location of setules>/
1. on inner surface/
2. on outer surface/

#1698. Second swimming legs endopod 3 <presence of spinules>/

1. with spinules/
2. without spinules/

#1699. Second swimming legs endopod 3 <arrangement of spinules>/
1. as a row/
2. as a patch/

#1700. Second swimming legs endopod 3 <number of spinules>/
1. single/
2. double/

#1701. Second swimming legs endopod 3 <inserted of spinules>/
1. proximally inserted/
2. distally inserted/

#1702. Second swimming legs endopod 3 <location of spinules>/
1. at anterior surface/
2. at posterior surface/

#1703. Second swimming legs endopod 3 <presence of Schmeil's organ>/
1. presence of Schmeil's organ/
2. absence of Schmeil's organ/

#1704. Second swimming legs endopod 3 <location of Schmeil's organ>/
1. on anterior surface/
2. on posterior surface/

#1705. Second swimming legs exopod 1 <presence of seta>/
1. with seta/
2. without seta/

#1706. Second swimming legs exopod 1 <restriction or not of the seta>/
1. restricted/
2. unrestricted/

#1707. Second swimming legs exopod 1 <insertion of the restricted seta>/
1. <to inner surface>/
2. <to outer surface>/

#1708. Second swimming legs exopod 1 <number of setae on inner surface>/
1. one on inner surface/
2. two on inner surface/

#1709. Second swimming legs exopod 1 <number of setae on outer surface>/
1. one on outer surface/
2. two on outer surface/

#1710. Second swimming legs exopod 1 <presence of spine>/
1. with spine/

2. without spine/

#1711. Second swimming legs exopod 1 <total of spine>/
1. 1/
2. 2/

#1712. Second swimming legs exopod 1 <form of spine>/
1. stout/
2. slender/

#1713. Second swimming legs exopod 1 <size of spine>/
1. equal size/
2. unequal size/

#1714. Second swimming legs exopod 1 <relative length of the spine>/
1. reaching to distal-third of the exopod 2/
2. not reaching to distal-third of the exopod 2/

#1715. Second swimming legs exopod 1 <shape of the spine>/
1. serrated/
2. not serrated/

#1716. Second swimming legs exopod 1 <location of the shape of the spine>/
1. on inner side/
2. on outer side/

#1717. Second swimming legs exopod 1 <presence of setules>/
1. with setules/
2. without setules/

#1718. Second swimming legs exopod 1 <arrangement of setules>/
1. as a row/
2. as a patch/

#1719. Second swimming legs exopod 1 <composition of the setules>/
1. single/
2. double/

#1720. Second swimming legs exopod 1 <continuity of insertion of setules>/
1. continuously/
2. discontinuously/

#1721. Second swimming legs exopod 1 <location of setules>/
1. on inner side/
2. on outer side/

#1722. Second swimming legs exopod 1 <presence of spinules>/
1. with spinules/
2. without spinules/

#1723. Second swimming legs exopod 1 <position of the Schmeil's organ>/
1. presence of Schmeil's organ/
2. absence of Schmeil's organ/

#1724. Second swimming legs exopod 2 <presence of seta>/
1. with seta/
2. without seta/

#1725. Second swimming legs exopod 2 <insertion of the restricted seta>/
1. <to inner surface>/
2. <to outer surface>/

#1726. Second swimming legs exopod 2 <restriction or not of the seta>/
1. unrestricted/
2. plurimarginal/

#1727. Second swimming legs exopod 2 <number of setae on inner surface>/
1. one on inner surface/
2. two on inner surface/

#1728. Second swimming legs exopod 2 <number of setae on outer surface>/
1. one on outer surface/
2. two on outer surface/

#1729. Second swimming legs exopod 2 <presence of spine>/
1. with spine/
2. without spine/

#1730. Second swimming legs exopod 2 <total of spine>/
1. 1/
2. 2/

#1731. Second swimming legs exopod 2 <form of spine>/
1. stout/
2. slender/

#1732. Second swimming legs exopod 2 <comparative size between spine>/
1. equal size/
2. unequal size/

#1733. Second swimming legs exopod 2 <relative length of the spine>/
1. not surpassing the exopod 3/
2. surpassing the exopod 3/

#1734. Second swimming legs exopod 2 <relative length of the spine 1>/
1. first longer 2x than origin segment/
2. first longer 2x than origin segment/

#1735. Second swimming legs exopod 2 <relative length of the spine 2>/
1. second longer 2x than origin segment/
2. second not longer 2x than origin segment/

#1736. Second swimming legs exopod 2 <shape of the spine>/
1. serrated/
2. not serrated/

#1737. Second swimming legs exopod 2 <location of the shape of the spine>/
1. on inner side/
2. on outer side/

#1738. Second swimming legs exopod 2 <presence of setules>/
1. with setules/
2. without setules/

#1739. Second swimming legs exopod 2 <arrangement of setules>/
1. as a row/
2. as a patch/

#1740. Second swimming legs exopod 2 <number of setules>/
1. single/
2. double/

#1741. Second swimming legs exopod 2 <continuity of insertion of setules>/
1. continuously/
2. discontinuously/

#1742. Second swimming legs exopod 2 <location of setules>/
1. on inner surface/
2. on outer surface/

#1743. Second swimming legs exopod 2 <presence of spinules>/
1. with spinules/
2. without spinules/

#1744. Second swimming legs exopod 2 <arrangement of spinules>/
1. as a row/
2. as a patch/

#1745. Second swimming legs exopod 2 <number of spinules>/
1. single/
2. double/

#1746. Second swimming legs exopod 2 <inserted of spinules>/
1. proximally inserted/
2. distally inserted/

#1747. Second swimming legs exopod 2 <location of spinules>/

1. at anterior surface/
2. at posterior surface/

#1748. Second swimming legs exopod 2 <presence of Schmeil's organ>/
1. presence of Schmeil's organ/
2. absence of Schmeil's organ/

#1749. Second swimming legs exopod 3 <presence of seta>/
1. with seta/
2. without seta/

#1750. Second swimming legs exopod 3 <restriction or not of the seta>/
1. restricted/
2. plurimarginal/

#1751. Second swimming legs exopod 3 <insertion of the restricted seta>/
1. <to inner surface>/
2. <to outer surface>/
3. <to terminal surface>/

#1752. Second swimming legs exopod 3 <number of setae on inner surface>/
1. two on inner surface/
2. three on inner surface/

#1753. Second swimming legs exopod 3 <number of setae on outer surface>/
1. one on outer surface/
2. two on outer surface/

#1754. Second swimming legs exopod 3 <number of setae on terminal margin>/
1. one on terminal surface/
2. two on terminal surface/

#1755. Second swimming legs exopod 3 <presence of spine>/
1. with spine/
2. without spine/

#1756. Second swimming legs exopod 3 <total of spine>/
1. 3/
2. 2/

#1757. Second swimming legs exopod 3 <comparative size between spines>/
1. equal size/
2. unequal size/

#1758. Second swimming legs exopod 3 <relative length outermost spine 1>/
1. first longer 2x than origin segment/
2. first not longer 2x than origin segment/

#1759. Second swimming legs exopod 3 <form of spine>/

1. stout/
2. slender/

#1760. Second swimming legs exopod 3 <shape of the spine 1>/
1. serrated/
2. not serrated/

#1761. Second swimming legs exopod 3 <location of the shape of the spine 1>/
1. on inner side/
2. on outer side/

#1762. Second swimming legs exopod 3 <continuity of the serrated shape spine 1>/
1. equally/
2. unequally/

#1763. Second swimming legs exopod 3 <relative length of the innermost spine 2>/
1. second longer 2x than origin segment/
2. second not longer 2x than origin segment/

#1764. Second swimming legs exopod 3 <form of spine 2>/
1. stout/
2. slender/

#1765. Second swimming legs exopod 3 <shape of the spine 2>/
1. serrated/
2. not serrated/

#1766. Second swimming legs exopod 3 <location of the serrated spine 2>/
1. on inner side/
2. on outer side/

#1767. Second swimming legs exopod 3 <ornamentation spine 2 with non-serrated side>/
1. with ornamentation on non-serrated side/
2. without ornamentation on non-serrated side/

#1768. Second swimming legs exopod 3 <ornamentation type of the spine 2>/
1. of spinules/
2. of setules/

#1769. Second swimming legs exopod 3 <presence of setules on the segment>/
1. no setules on outer surface/
2. setules on outer surface/

#1770. Second swimming legs exopod 3 <arrangement of setules>/
1. as a row/
2. as a patch/

#1771. Second swimming legs exopod 3 <number of setules>/

1. single/
2. double/

#1772. Second swimming legs exopod 3 <continuity of insertion of setules>/
1. continuously/
2. discontinuously/

#1773. Second swimming legs exopod 3 <location of setules>/
1. on inner surface/
2. on outer surface/

#1774. Second swimming legs exopod 3 <presence of spinules>/
1. with spinules/
2. without spinules/

#1775. Second swimming legs exopod 3 <arrangement of spinules>/
1. as a row/
2. as a patch/

#1776. Second swimming legs exopod 3 <number of spinules>/
1. single/
2. double/

#1777. Second swimming legs exopod 3 <inserted of spinules>/
1. proximally inserted/
2. distally inserted/

#1778. Second swimming legs exopod 3 <location of spinules>/
1. at anterior surface/
2. at posterior surface/

#1779. Second swimming legs exopod 3 <presence of Schmeil's organ>/
1. presence of Schmeil's organ/
2. absence of Schmeil's organ/

#1780. Third swimming legs <symmetry>/
1. symmetrical/
2. asymmetrical/

#1781. Third swimming legs <number of rami>/
1. uniramous/
2. biramous/

#1782. Third swimming legs intercoxal plate <presence of seta>/
1. with seta/
2. without seta/

#1783. Third swimming legs intercoxal plate <position of sensilla>/
1. centralized/

2. laterally/

#1784. Third swimming legs intercoxal plate <location of sensilla>/
1. on anterior surface/
2. on posterior surface/

#1785. Third swimming legs praecoxa <presence of praecoxa>/
1. present/
2. absent/

#1786. Third swimming legs praecoxa <location of praecoxa>/
1. laterally located/
2. not laterally located/

#1787. Third swimming legs coxa <presence of seta>/
1. with seta/
2. without seta/

#1788. Third swimming legs coxa <seta composition>/
1. plumose/
2. straight/

#1789. Third swimming legs coxa <insertion of seta>/
1. proximally inserted/
2. distally inserted/

#1790. Third swimming legs coxa <location of setae>/
1. on inner surface/
2. on outer surface/

#1791. Third swimming legs coxa <relative length>/
1. surpassing to basal segment/
2. surpassing to first endopodal segment/

#1792. Third swimming legs coxa <presence of setules>/
1. with setules/
2. without setules/

#1793. Third swimming legs coxa <arrangement of setules>/
1. as a row/
2. as a patch/

#1794. Third swimming legs coxa <number of setules>/
1. single/
2. double/

#1795. Third swimming legs coxa <continuity of insertion of setules>/
1. continuously/
2. discontinuously/

#1796. Third swimming legs coxa <location of setules>/
1. on inner margin/
2. on outer margin/

#1797. Third swimming legs coxa <presence of spinules>/
1. with spinules/
2. without spinules/

#1798. Third swimming legs coxa <arrangement of spinules>/
1. as a row/
2. as a patch/

#1799. Third swimming legs coxa <number of spinules>/
1. single/
2. double/

#1800. Third swimming legs coxa <inserted of spinules>/
1. proximally inserted/
2. distally inserted/

#1801. Third swimming legs coxa <location of spinules>/
1. at anterior surface/
2. at posterior surface/

#1802. Third swimming legs coxa <presence of spine>/
1. with spine/
2. without spine/

#1803. Third swimming legs basis <presence of seta>/
1. with seta/
2. without seta/

#1804. Third swimming legs basis <insertion of seta>/
1. proximally inserted/
2. distally inserted/

#1805. Third swimming legs basis <location of setae>/
1. on inner margin/
2. on outer margin/

#1806. Third swimming legs basis <relative length of the seta>/
1. surpassing to distal limit of the segment/
2. not surprassing to distal limit of the segment/

#1807. Third swimming legs basis <presence of setules>/
1. with setules/
2. without setules/

#1808. Third swimming legs basis <arrangement of setules>/
1. as a row/
2. as a patch/

#1809. Third swimming legs basis <number of setules>/
1. single/
2. double/

#1810. Third swimming legs basis <continuity of insertion of setules>/
1. continuously/
2. discontinuously/

#1811. Third swimming legs basis <location of setules>/
1. on inner surface/
2. on outer surface/

#1812. Third swimming legs basis <presence of spinules>/
1. with spinules/
2. without spinules/

#1813. Third swimming legs basis <arrangement of spinules>/
1. as a row/
2. as a patch/

#1814. Third swimming legs basis <number of spinules>/
1. single/
2. double/

#1815. Third swimming legs basis <inserted of spinules>/
1. proximally inserted/
2. distally inserted/

#1816. Third swimming legs basis <location of spinules>/
1. at anterior surface/
2. at posterior surface/

#1817. Third swimming legs basis <presence of spine>/
1. with spine/
2. without spine/

#1818. Third swimming legs endopod <number of segmentation>/
1. 2-segmented/
2. 3-segmented/

#1819. Third swimming legs endopod 1 <presence of seta>/
1. with seta/
2. without seta/

#1820. Third swimming legs endopod 1 <restriction or not of the seta>/

1. restricted/
2. unrestricted/

#1821. Third swimming legs endopod 1 <insertion of the restricted seta>/
1. <to inner surface>/
2. <to outer surface>/

#1822. Third swimming legs endopod 1 <number of setae on inner surface>/
1. one on inner surface/
2. two on inner surface/

#1823. Third swimming legs endopod 1 <number of setae on outer surface>/
1. one on outer surface/
2. two on outer surface/

#1824. Third swimming legs endopod 1 <presence of spine>/
1. with spine/
2. without spine/

#1825. Third swimming legs endopod 1 <presence of setules>/
1. with setules/
2. without setules/

#1826. Third swimming legs endopod 1 <arrangement of setules>/
1. as a row/
2. as a patch/

#1827. Third swimming legs endopod 1 <number of setules>/
1. single/
2. double/

#1828. Third swimming legs endopod 1 <continuity of insertion of setules>/
1. continuously/
2. discontinuously/

#1829. Third swimming legs endopod 1 <location of setules>/
1. on inner surface/
2. on outer surface/

#1830. Third swimming legs endopod 1 <presence of spinules>/
1. with spinules/
2. without spinules/

#1831. Third swimming legs endopod 1 <presence of Schmeil's organ>/
1. presence of Schmeil's organ/
2. absence of Schmeil's organ/

#1832. Third swimming legs endopod 2 <presence of seta>/
1. with seta/

2. without seta/

#1833. Third swimming legs endopod 2 <restriction or not of the seta>/
1. restricted/
2. unrestricted/

#1834. Third swimming legs endopod 2 <insertion of the restricted seta>/
1. <on inner surface>/
2. <on outer surface>/

#1835. Third swimming legs endopod 2 <number of setae on inner surface>/
1. one on inner surface/
2. two on inner surface/

#1836. Third swimming legs endopod 2 <seta composition>/
1. plumose/
2. straight/

#1837. Third swimming legs endopod 2 <number of setae on outer margin>/
1. one on outer margin/
2. two on outer margin/

#1838. Third swimming legs endopod 2 <presence of spine>/
1. with spine/
2. without spine/

#1839. Third swimming legs endopod 2 <presence of setules>/
1. with setules/
2. without setules/

#1840. Third swimming legs endopod 2 <arrangement of setules>/
1. as a row/
2. as a patch/

#1841. Third swimming legs endopod 2 <number of setules>/
1. single/
2. double/

#1842. Third swimming legs endopod 2 <continuity of insertion of setules>/
1. continuously/
2. discontinuously/

#1843. Third swimming legs endopod 2 <location of setules>/
1. on inner margin/
2. on outer margin/

#1844. Third swimming legs endopod 2 <presence of spinules>/
1. with spinules/
2. without spinules/

#1845. Third swimming legs endopod 2 <arrangement of spinules>/
1. as a row/
2. as a patch/

#1846. Third swimming legs endopod 2 <number of spinules>/
1. single/
2. double/

#1847. Third swimming legs endopod 2 <inserted of spinules>/
1. proximally inserted/
2. distally inserted/

#1848. Third swimming legs endopod 2 <location of spinules>/
1. at anterior surface/
2. at posterior surface/

#1849. Third swimming legs endopod 2 <presence of Schmeil's organ>/
1. absence of Schmeil's organ/
2. presence of Schmeil's organ/

#1850. Third swimming legs endopod 3 <presence of seta>/
1. with seta/
2. without seta/

#1851. Third swimming legs endopod 3 <seta composition>/
1. plumose/
2. straight/

#1852. Third swimming legs endopod 3 <restriction or not of the seta>/
1. unrestricted/
2. plurimarginal/

#1853. Third swimming legs endopod 3 <insertion of the restricted seta>/
1. <to inner surface>/
2. <to outer surface>/
3. <to terminal surface>/

#1854. Third swimming legs endopod 3 <number of setae on inner surface>/
1. two on inner surface/
2. three on inner surface/

#1855. Third swimming legs endopod 3 <number of setae on outer surface>/
1. one on outer surface/
2. two on outer surface/

#1856. Third swimming legs endopod 3 <number of setae on terminal surface>/
1. two on terminal surface/
2. three on terminal surface/

#1857. Third swimming legs endopod 3 <presence of spine>/
1. with spine/
2. without spine/

#1858. Third swimming legs endopod 3 <presence of setules>/
1. with setules/
2. without setules/

#1859. Third swimming legs endopod 3 <arrangement of setules>/
1. as a row/
2. as a patch/

#1860. Third swimming legs endopod 3 <number of setules>/
1. single/
2. double/

#1861. Third swimming legs endopod 3 <continuity of insertion of setules>/
1. continuously/
2. discontinuously/

#1862. Third swimming legs endopod 3 <location of setules>/
1. on inner surface/
2. on outer surface/

#1863. Third swimming legs endopod 3 <presence of spinules>/
1. with spinules/
2. without spinules/

#1864. Third swimming legs endopod 3 <arrangement of spinules>/
1. as a row/
2. as a patch/

#1865. Third swimming legs endopod 3 <inserted of spinules>/
1. proximally inserted/
2. distally inserted/

#1866. Third swimming legs endopod 3 <number of spinules>/
1. single/
2. double/

#1867. Third swimming legs endopod 3 <location of spinules>/
1. at anterior surface/
2. at posterior surface/

#1868. Third swimming legs endopod 3 <presence of Schmeil's organ>/
1. presence of Schmeil's organ/
2. absence of Schmeil's organ/

#1869. Third swimming legs exopod 1 <presence of seta>/
    1. with seta/
    2. without seta/

#1870. Third swimming legs exopod 1 <restriction or not of the seta>/
    1. restricted/
    2. unrestricted/

#1871. Third swimming legs exopod 1 <seta composition>/
    1. plumose/
    2. straight/

#1872. Third swimming legs exopod 1 <insertion of the restricted seta>/
    1. <to inner surface>/
    2. <to outer surface>/

#1873. Third swimming legs exopod 1 <number of setae on inner surface>/
    1. one on inner surface/
    2. two on inner surface/

#1874. Third swimming legs exopod 1 <number of setae on outer surface>/
    1. one on outer surface/
    2. two on outer surface/

#1875. Third swimming legs exopod 1 <presence of spine>/
    1. with spine/
    2. without spine/

#1876. Third swimming legs exopod 1 <total of spine>/
    1. 1/
    2. 2/

#1877. Third swimming legs exopod 1 <form of spine>/
    1. stout/
    2. slender/

#1878. Third swimming legs exopod 1 <relative length of the spine>/
    1. not reaching to the distal-third of the exopod 2/
    2. reaching to the distal-third of the exopod 2/

#1879. Third swimming legs exopod 1 <shape of the spine>/
    1. serrated/
    2. not serrated/

#1880. Third swimming legs exopod 1 <continuity of the serrated shape spine>/
    1. equally/
    2. unequally/

#1881. Third swimming legs exopod 1 <location of the shape of the spine>/

1. on inner surface/
2. on outer surface/

#1882. Third swimming legs exopod 1 <presence of setules>/
1. with setules/
2. without setules/

#1883. Third swimming legs exopod 1 <arrangement of setules>/
1. as a row/
2. as a patch/

#1884. Third swimming legs exopod 1 <number of setules>/
1. single/
2. double/

#1885. Third swimming legs exopod 1 <continuity of insertion of setules>/
1. continuously/
2. discontinuously/

#1886. Third swimming legs exopod 1 <location of setules>/
1. on inner surface/
2. on outer surface/

#1887. Third swimming legs exopod 1 <presence of spinules>/
1. with spinules/
2. without spinules/

#1888. Third swimming legs exopod 1 <arrangement of spinules>/
1. as a row/
2. as a patch/

#1889. Third swimming legs exopod 1 <number of spinules>/
1. single/
2. double/

#1890. Third swimming legs exopod 1 <inserted of spinules>/
1. proximally inserted/
2. distally inserted/

#1891. Third swimming legs exopod 1 <location of spinules>/
1. at anterior surface/
2. at posterior surface/

#1892. Third swimming legs exopod 1 <presence of Schmeil's organ>/
1. presence of Schmeil's organ/
2. absence of Schmeil's organ/

#1893. Third swimming legs exopod 2 <presence of seta>/
1. with seta/

2. without seta/

#1894. Third swimming legs exopod 2 <seta composition>/
1. plumose/
2. straight/

#1895. Third swimming legs exopod 2 <restriction or not of the seta>/
1. restricted/
2. unrestricted/

#1896. Third swimming legs exopod 2 <insertion of the restricted seta>/
1. <to inner surface>/
2. <to outer surface>/

#1897. Third swimming legs exopod 2 <number of setae on inner surface>/
1. one on inner surface/
2. two on inner surface/

#1898. Third swimming legs exopod 2 <number of setae on outer surface>/
1. one on outer surface/
2. two on outer surface/

#1899. Third swimming legs exopod 2 <presence of spine>/
1. with spine/
2. without spine/

#1900. Third swimming legs exopod 2 <number of spine>/
1. 1/
2. 2/

#1901. Third swimming legs exopod 2 <form of spine>/
1. stout/
2. slender/

#1902. Third swimming legs exopod 2 <relative length of the spine>/
1. reaching out to exopod 3/
2. not reaching out to exopod 3/

#1903. Third swimming legs exopod 2 <shape of the spine>/
1. serrated/
2. not serrated/

#1904. Third swimming legs exopod 2 <location of the shape of the spine>/
1. on inner side/
2. on outer side/

#1905. Third swimming legs exopod 2 <continuity of the serrated shape spine>/
1. equally/
2. unequally/

#1906. Third swimming legs exopod 2 <presence of setules>/
1. with setules/
2. without setules/

#1907. Third swimming legs exopod 2 <arrangement of setules>/
1. as a row/
2. as a patch/

#1908. Third swimming legs exopod 2 <number of setules>/
1. single/
2. double/

#1909. Third swimming legs exopod 2 <continuity of insertion of setules>/
1. continuously/
2. discontinuously/

#1910. Third swimming legs exopod 2 <location of setules>/
1. on inner side/
2. on outer side/

#1911. Third swimming legs exopod 2 <presence of spinules>/
1. with spinules/
2. without spinules/

#1912. Third swimming legs exopod 2 <arrangement of spinules>/
1. as a row/
2. as a patch/

#1913. Third swimming legs exopod 2 <number of spinules>/
1. single/
2. double/

#1914. Third swimming legs exopod 2 <inserted of spinules>/
1. proximally inserted/
2. distally inserted/

#1915. Third swimming legs exopod 2 <location of spinules>/
1. at anterior surface/
2. at posterior surface/

#1916. Third swimming legs exopod 2 <presence of Schmeil's organ>/
1. presence of Schmeil's organ/
2. absence of Schmeil's organ/

#1917. Third swimming legs exopod 3 <presence of setules>/
1. with setules/
2. without setules/

#1918. Third swimming legs exopod 3 <arrangement of setules>/
1. as a row/
2. as a patch/

#1919. Third swimming legs exopod 3 <number of setules>/
1. single/
2. double/

#1920. Third swimming legs exopod 3 <continuity of insertion of setules>/
1. continuously/
2. discontinuously/

#1921. Third swimming legs exopod 3 <location of setules>/
1. on inner margin/
2. on outer margin/

#1922. Third swimming legs exopod 3 <presence of spinules>/
1. with spinules/
2. without spinules/

#1923. Third swimming legs exopod 3 <arrangement of spinules>/
1. as a row/
2. as a patch/

#1924. Third swimming legs exopod 3 <number of spinules>/
1. single/
2. double/

#1925. Third swimming legs exopod 3 <inserted of spinules>/
1. proximally inserted/
2. distally inserted/

#1926. Third swimming legs exopod 3 <location of spinules>/
1. at anterior surface/
2. at posterior surface/

#1927. Third swimming legs exopod 3 <presence of seta>/
1. with seta/
2. without seta/

#1928. Third swimming legs exopod 3 <seta composition>/
1. plumose/
2. straight/

#1929. Third swimming legs exopod 3 <restriction or not of the seta>/
1. restricted/
2. unrestricted/

#1930. Third swimming legs exopod 3 <insertion of the restricted seta>/

1. <to inner surface>/
2. <to outer surface>/
3. <to terminal surface>/

#1931. Third swimming legs exopod 3 <number of setae on inner surface>/
1. four on inner surface/
2. three on inner surface/

#1932. Third swimming legs exopod 3 <number of setae on outer surface>/
1. one on outer surface/
2. two on outer surface/

#1933. Third swimming legs exopod 3 <number of setae on terminal surface>/
1. one on terminal surface/
2. two on terminal surface/

#1934. Third swimming legs exopod 3 <presence of spine>/
1. with spine/
2. without spine/

#1935. Third swimming legs exopod 3 <total of spine>/
1. 3/
2. 2/

#1936. Third swimming legs exopod 3 <comparative size between spines>/
1. equal size/
2. unequal size/

#1937. Third swimming legs exopod 3 <relative length outermost spine 1>/
1. first longer 2x than origin segment/
2. first not longer 2x than origin segment/

#1938. Third swimming legs exopod 3 <form of spine 1>/
1. stout/
2. slender/

#1939. Third swimming legs exopod 3 <shape of the spine 1>/
1. serrated/
2. not serrated/

#1940. Third swimming legs exopod 3 <location of the shape of the spine 1>/
1. on inner side/
2. on outer side/

#1941. Third swimming legs exopod 3 <continuity of the serrated shape spine 1>/
1. equally/
2. unequally/

#1942. Third swimming legs exopod 3 <relative length of the innermost spine 2>/

1. second longer 2x than origin segment/
2. second not longer 2x than origin segment/

#1943. Third swimming legs exopod 3 <form of spine 2>/
1. stout/
2. slender/

#1944. Third swimming legs exopod 3 <shape of the spine 2>/
1. serrated/
2. not serrated/

#1945. Third swimming legs exopod 3 <location of the serrated spine 2>/
1. on inner side/
2. on outer side/

#1946. Third swimming legs exopod 3 <ornamentation spine 2 with non-serrated side>/
1. with ornamentation on non-serrated side/
2. without ornamentation on non-serrated side/

#1947. Third swimming legs exopod 3 <ornamentation type of the spine 2>/
1. of spinules/
2. of setules/

#1948. Third swimming legs exopod 3 <presence of Schmeil's organ>/
1. presence of Schmeil's organ/
2. absence of Schmeil's organ/

#1949. Fourth swimming legs <symmetry>/
1. symmetrical/
2. asymmetrical/

#1950. Fourth swimming legs <number of rami>/
1. uniramous/
2. biramous/

#1951. Fourth swimming legs intercoxal plate <presence of sensilla>/
1. with sensilla/
2. without sensilla/

#1952. Fourth swimming legs praecoxa <presence of praecoxa>/
1. present/
2. absent/

#1953. Fourth swimming legs coxa <presence of seta>/
1. with seta/
2. without seta/

#1954. Fourth swimming legs coxa <insertion of seta>/

1. proximally inserted/
2. distally inserted/

#1955. Fourth swimming legs coxa <location of setae>/
1. on inner margin/
2. on outer margin/

#1956. Fourth swimming legs coxa <relative length>/
1. reaching out to endopod 1/
2. reaching out to the end of endopod 2/

#1957. Fourth swimming legs coxa <presence of spinules>/
1. with spinules/
2. without spinules/

#1958. Fourth swimming legs coxa <arrangement of the spinules>/
1. as a row/
2. as a patch/

#1959. Fourth swimming legs coxa <quantity of arrangement of the spinules>/
1. single/
2. double/

#1960. Fourth swimming legs coxa <presence of setules>/
1. setules present/
2. setules absent/

#1961. Fourth swimming legs coxa <arrangement of setules>/
1. as a patch innerly/
2. as a row innerly/

#1962. Fourth swimming legs basis <presence of seta>/
1. with seta/
2. without seta/

#1963. Fourth swimming legs basis <number of setae>/
1. one/
2. two/

#1964. Fourth swimming legs basis <insertion of seta>/
1. proximally inserted/
2. medially inserted/
3. distally inserted/

#1965. Fourth swimming legs basis <location of setae>/
1. on posterior surface/
2. on anterior surface/

#1966. Fourth swimming legs basis <relative length>/

1. larger than the original segment/
2. smaller than the original segment/

#1967. Fourth swimming legs basis <presence of setules>/
1. with setules/
2. without setules/

#1968. Fourth swimming legs basis <arrangement of setules>/
1. as a row/
2. as a patch/

#1969. Fourth swimming legs basis <number of setules>/
1. single/
2. double/

#1970. Fourth swimming legs basis <continuity of insertion of setules>/
1. continuously/
2. discontinuously/

#1971. Fourth swimming legs basis <location of setules>/
1. on inner side/
2. on outer side/

#1972. Fourth swimming legs basis <presence of spinules>/
1. with spinules/
2. without spinules/

#1973. Fourth swimming legs basis <arrangement of spinules>/
1. as a row/
2. as a patch/

#1974. Fourth swimming legs basis <number of spinules>/
1. single/
2. double/

#1975. Fourth swimming legs basis <inserted of spinules>/
1. medially inserted/
2. distally inserted/

#1976. Fourth swimming legs basis <location of spinules>/
1. at anterior surface/
2. at posterior surface/

#1977. Fourth swimming legs basis <presence of spine>/
1. with spine/
2. without spine/

#1978. Fourth swimming legs endopod <number of segmentation>/
1. 2-segmented/

2. 3-segmented/

#1979. Fourth swimming legs endopod 1 <presence of seta>/
1. with seta/
2. without seta/

#1980. Fourth swimming legs endopod 1 <number of setae>/
1. one/
2. two/

#1981. Fourth swimming legs endopod 1 <restriction or not of the seta>/
1. restricted/
2. unrestricted/

#1982. Fourth swimming legs endopod 1 <insertion of the restricted seta>/
1. on inner surface/
2. on outer surface/

#1983. Fourth swimming legs endopod 1 <presence of spine>/
1. with spine/
2. without spine/

#1984. Fourth swimming legs endopod 1 <presence of setules>/
1. with setules/
2. without setules/

#1985. Fourth swimming legs endopod 1 <arrangement of setules>/
1. as a row/
2. as a patch/

#1986. Fourth swimming legs endopod 1 <continuity of insertion of setules>/
1. continuously/
2. discontinuously/

#1987. Fourth swimming legs endopod 1 <location of setules>/
1. on inner surface/
2. on outer surface/

#1988. Fourth swimming legs endopod 1 <presence of spinules>/
1. with spinules/
2. without spinules/

#1989. Fourth swimming legs endopod 1 <presence of Schmeil's organ>/
1. absence of Schmeil's organ/
2. presence of Schmeil's organ/

#1990. Fourth swimming legs endopod 2 <presence of seta>/
1. with seta/
2. without seta/

#1991. Fourth swimming legs endopod 2 <restriction or not of the seta>/
1. restricted/
2. unrestricted/

#1992. Fourth swimming legs endopod 2 <insertion of the restricted seta>/
1. <to inner side>/
2. <to outer side>/

#1993. Fourth swimming legs endopod 2 <number of setae on inner margin>/
1. one on inner side/
2. two on inner side/

#1994. Fourth swimming legs endopod 2 <number of setae on outer margin>/
1. one on outer side/
2. two on outer side/

#1995. Fourth swimming legs endopod 2 <presence of spine>/
1. with spine/
2. without spine/

#1996. Fourth swimming legs endopod 2 <presence of setules>/
1. with setules/
2. without setules/

#1997. Fourth swimming legs endopod 2 <arrangement of setules>/
1. as a row/
2. as a patch/

#1998. Fourth swimming legs endopod 2 <number of setules>/
1. single/
2. double/

#1999. Fourth swimming legs endopod 2 <continuity of insertion of setules>/
1. continuously/
2. discontinuously/

#2000. Fourth swimming legs endopod 2 <location of setules>/
1. on inner surface/
2. on outer surface/

#2001. Fourth swimming legs endopod 2 <presence of spinules>/
1. with spinules/
2. without spinules/

#2002. Fourth swimming legs endopod 2 <presence of Schmeil's organ>/
1. absence of Schmeil's organ/
2. presence of Schmeil's organ/

#2003. Fourth swimming legs endopod 3 <presence of seta>/

1. with seta/
2. without seta/

#2004. Fourth swimming legs endopod 3 <restriction or not of the seta>/

1. restricted/
2. unrestricted/

#2005. Fourth swimming legs endopod 3 <insertion of the restricted seta>/

1. <to inner surface>/
2. <to outer surface>/
3. <to distal surface>/

#2006. Fourth swimming legs endopod 3 <number of setae on inner surface>/

1. three on inner surface/
2. two on inner surface/

#2007. Fourth swimming legs endopod 3 <number of setae on outer surface>/

1. one on outer surface/
2. two on outer surface/

#2008. Fourth swimming legs endopod 3 <number of setae on distal surface>/

1. two on distal surface/
2. three on distal surface/

#2009. Fourth swimming legs endopod 3 <presence of spine>/

1. with spine/
2. without spine/

#2010. Fourth swimming legs endopod 3 <presence of setules>/

1. with setules/
2. without setules/

#2011. Fourth swimming legs endopod 3 <arrangement of setules>/

1. as a row/
2. as a patch/

#2012. Fourth swimming legs endopod 3 <number of setules>/

1. single/
2. double/

#2013. Fourth swimming legs endopod 3 <continuity of insertion of setules>/

1. continuously/
2. discontinuously/

#2014. Fourth swimming legs endopod 3 <location of setules>/

1. on inner surface/
2. on outer surface/

#2015. Fourth swimming legs endopod 3 <presence of spinules>/
1. with spinules/
2. without spinules/

#2016. Fourth swimming legs endopod 3 <arrangement of spinules>/
1. as a row/
2. as a patch/

#2017. Fourth swimming legs endopod 3 <number of spinules>/
1. single/
2. double/

#2018. Fourth swimming legs endopod 3 <inserted of spinules>/
1. medially inserted/
2. distally inserted/

#2019. Fourth swimming legs endopod 3 <location of spinules>/
1. at anterior surface/
2. at posterior surface/

#2020. Fourth swimming legs endopod 3 <presence of Schmeil's organ>/
1. absence of Schmeil's organ/
2. presence of Schmeil's organ/

#2021. Fourth swimming legs exopod 1 <presence of seta>/
1. with seta/
2. without seta/

#2022. Fourth swimming legs exopod 1 <restriction or not of the seta>/
1. restricted/
2. unrestricted/

#2023. Fourth swimming legs exopod 1 <insertion of the restricted seta>/
1. <to inner surface>/
2. <to outer surface>/

#2024. Fourth swimming legs exopod 1 <number of setae on inner surface>/
1. one on inner surface/
2. two on inner surface/

#2025. Fourth swimming legs exopod 1 <number of setae on outer surface>/
1. one on outer surface/
2. two on outer surface/

#2026. Fourth swimming legs exopod 1 <presence of spine>/
1. with spine/
2. without spine/

#2027. Fourth swimming legs exopod 1 <total of spine>/

1. 1/
2. 2/

#2028. Fourth swimming legs exopod 1 <form of spine>/
1. stout/
2. slender/

#2029. Fourth swimming legs exopod 1 <relative length of the spine>/
1. reaching out to distal-third of the exopod 2/
2. not reaching out to distal-third of the exopod 2/

#2030. Fourth swimming legs exopod 1 <shape of the spine>/
1. serrated/
2. not serrated/

#2031. Fourth swimming legs exopod 1 <location of the shape of the spine>/
1. on inner side/
2. on outer side/

#2032. Fourth swimming legs exopod 1 <continuity of the serrated shape spine>/
1. equally/
2. unequally/

#2033. Fourth swimming legs exopod 1 <presence of setules>/
1. with setules/
2. without setules/

#2034. Fourth swimming legs exopod 1 <arrangement of setules>/
1. as a row/
2. as a patch/

#2035. Fourth swimming legs exopod 1 <number of setules>/
1. single/
2. double/

#2036. Fourth swimming legs exopod 1 <continuity of insertion of setules>/
1. continuously/
2. discontinuously/

#2037. Fourth swimming legs exopod 1 <location of setules>/
1. on inner surface/
2. on outer surface/

#2038. Fourth swimming legs exopod 1 <presence of spinules>/
1. with spinules/
2. without spinules/

#2039. Fourth swimming legs exopod 1 <arrangement of spinules>/
1. as a row/

2. as a patch/

#2040. Fourth swimming legs exopod 1 <number of spinules>/
1. single/
2. double/

#2041. Fourth swimming legs exopod 1 <inserted of spinules>/
1. proximally inserted/
2. medially inserted/
3. distally inserted/

#2042. Fourth swimming legs exopod 1 <location of spinules>/
1. at anterior surface/
2. at posterior surface/

#2043. Fourth swimming legs exopod 1 <presence of Schmeil's organ>/
1. absence of Schmeil's organ/
2. presence of Schmeil's organ/

#2044. Fourth swimming legs exopod 2 <presence of seta>/
1. with seta/
2. without seta/

#2045. Fourth swimming legs exopod 2 <restriction or not of the seta>/
1. restricted/
2. unrestricted/

#2046. Fourth swimming legs exopod 2 <insertion of the restricted seta>/
1. <to inner surface>/
2. <to outer surface>/

#2047. Fourth swimming legs exopod 2 <number of setae on inner surface>/
1. one on inner surface/
2. two on inner surface/

#2048. Fourth swimming legs exopod 2 <number of setae on outer surface>/
1. one on outer surface/
2. two on outer surface/

#2049. Fourth swimming legs exopod 2 <presence of spine>/
1. with spine/
2. without spine/

#2050. Fourth swimming legs exopod 2 <number of spine>/
1. 1/
2. 2/

#2051. Fourth swimming legs exopod 2 <form of spine>/
1. stout/

2. slender/

#2052. Fourth swimming legs exopod 2 <relative length of the spine>/
1. reaching the end of exopod 3/
2. not reaching the end of exopod 3/

#2053. Fourth swimming legs exopod 2 <shape of the spine>/
1. serrated/
2. not serrated/

#2054. Fourth swimming legs exopod 2 <location of the shape of the spine>/
1. on inner side/
2. on outer side/

#2055. Fourth swimming legs exopod 2 <continuity of the serrated shape spine>/
1. equally/
2. unequally/

#2056. Fourth swimming legs exopod 2 <presence of setules>/
1. with setules/
2. without setules/

#2057. Fourth swimming legs exopod 2 <arrangement of setules>/
1. as a row/
2. as a patch/

#2058. Fourth swimming legs exopod 2 <number of setules>/
1. single/
2. double/

#2059. Fourth swimming legs exopod 2 <continuity of insertion of setules>/
1. continuously/
2. discontinuously/

#2060. Fourth swimming legs exopod 2 <location of setules>/
1. on inner surface/
2. on outer surface/

#2061. Fourth swimming legs exopod 2 <presence of spinules>/
1. with spinules/
2. without spinules/

#2062. Fourth swimming legs exopod 2 <arrangement of spinules>/
1. as a row/
2. as a patch/

#2063. Fourth swimming legs exopod 2 <number of spinules>/
1. single/
2. double/

#2064. Fourth swimming legs exopod 2 <inserted of spinules>/
1. proximally inserted/
2. distally inserted/

#2065. Fourth swimming legs exopod 2 <location of spinules>/
1. at anterior surface/
2. at posterior surface/

#2066. Fourth swimming legs exopod 2 <presence of Schmeil's organ>/
1. absence of Schmeil's organ/
2. presence of Schmeil's organ/

#2067. Fourth swimming legs exopod 3 <presence of setules>/
1. with setules/
2. without setules/

#2068. Fourth swimming legs exopod 3 <arrangement of setules>/
1. as a row/
2. as a patch/

#2069. Fourth swimming legs exopod 3 <number of setules>/
1. single/
2. double/

#2070. Fourth swimming legs exopod 3 <continuity of insertion of setules>/
1. continuously/
2. discontinuously/

#2071. Fourth swimming legs exopod 3 <location of setules>/
1. on inner margin/
2. on outer margin/

#2072. Fourth swimming legs exopod 3 <presence of spinules>/
1. with spinules/
2. without spinules/

#2073. Fourth swimming legs exopod 3 <arrangement of spinules>/
1. as a row/
2. as a patch/

#2074. Fourth swimming legs exopod 3 <number of spinules>/
1. single/
2. double/

#2075. Fourth swimming legs exopod 3 <inserted of spinules>/
1. proximally inserted/
2. distally inserted/

#2076. Fourth swimming legs exopod 3 <location of spinules>/
1. at anterior surface/
2. at posterior surface/

#2077. Fourth swimming legs exopod 3 <presence of seta>/
1. with seta/
2. without seta/

#2078. Fourth swimming legs exopod 3 <restriction or not of the seta>/
1. restricted/
2. unrestricted/

#2079. Fourth swimming legs exopod 3 <insertion of the restricted seta>/
1. <to inner surface>/
2. <to outer surface>/
3. <to distal surface>/

#2080. Fourth swimming legs exopod 3 <number of setae on inner surface>/
1. four on inner surface/
2. three on inner surface/

#2081. Fourth swimming legs exopod 3 <number of setae on outer surface>/
1. one on outer surface/
2. two on outer surface/

#2082. Fourth swimming legs exopod 3 <number of setae on distal surface>/
1. one on distal surface/
2. two on distal surface/

#2083. Fourth swimming legs exopod 3 <presence of spine>/
1. with spine/
2. without spine/

#2084. Fourth swimming legs exopod 3 <total of spine>/
1. 2/
2. 3/

#2085. Fourth swimming legs exopod 3 <comparative size between spine>/
1. equal size/
2. unequal size/

#2086. Fourth swimming legs exopod 3 <relative length outermost spine 1>/
1. first longer 2x than origin segment/
2. first not longer 2x than origin segment/

#2087. Fourth swimming legs exopod 3 <form of spine 1>/
1. stout/
2. slender/

#2088. Fourth swimming legs exopod 3 <shape of the spine 1>/
1. serrated/
2. not serrated/

#2089. Fourth swimming legs exopod 3 <location of the shape of the spine 1>/
1. on inner side/
2. on outer side/

#2090. Fourth swimming legs exopod 3 <continuity of the serrated shape spine 1>/
1. equally/
2. unequally/

#2091. Fourth swimming legs exopod 3 <relative length of the innermost spine 2>/
1. second longer 2x than origin segment/
2. second not longer 2x than origin segment/

#2092. Fourth swimming legs exopod 3 <form of spine 2>/
1. stout/
2. slender/

#2093. Fourth swimming legs exopod 3 <shape of the spine 2>/
1. serrated/
2. not serrated/

#2094. Fourth swimming legs exopod 3 <location of the shape of the spine 2>/
1. on inner side/
2. on outer side/

#2095. Fourth swimming legs exopod 3 <ornamentation spine 2 with non-serrated side>/
1. with ornamentation on non-serrated side/
2. without ornamentation on non-serrated side/

#2096. Fourth swimming legs exopod 3 <type of ornamentation on non-serrated side>/
1. of spinules/
2. of setules/

#2097. Fourth swimming legs exopod 3 <presence of Schmeil's organ>/
1. absence of Schmeil's organ/
2. presence of Schmeil's organ/

#2098. <male> fifth swimming legs <symmetry>/
1. symmetrical/
2. asymmetrical/

#2099. <male> fifth swimming legs intercoxal plate <relative size of the intercoxal plate>/
1. with length equal or greater than width on 1.5x/
2. with length not equal or greater than width on 1.5x/

#2100. <male> fifth swimming legs intercoxal plate <pattern of the anterior margin of the intercoxal plate>/
1. with regular proximal margin/
2. with irregular proximal margin/

#2101. <male> fifth swimming legs intercoxal plate <pattern of the anterior margin of the intercoxal plate>/
1. continuous to/
2. discontinuous to/

#2102. <male> fifth swimming legs intercoxal plate <comparison of pattern of the anterior margin of the intercoxal plate>/
1. the anterior margin of the left coxa/
2. the anterior margin of the right coxa/

#2103. <male> fifth swimming legs intercoxal plate <sensilla on the posterior margin of the right lateral>/
1. posterior sensilla on the right lateral absent/
2. posterior sensilla on the right lateral present/

#2104. <male> fifth left swimming leg <number of rami>/
1. uniramous/
2. biramous/

#2105. <male> fifth left swimming leg <reach of the left leg compared to right leg>/
1. reaching first right exopod segment/
2. surpassing first right exopod segment/

#2106. <male> fifth left swimming leg <reaching of the size of the left leg compared to right leg>/
1. proximally/
2. medially/
3. distally/

#2107. <male> fifth left swimming leg praecoxa <presence of praecoxa>/
1. present/
2. absent/

#2108. <male> fifth left swimming leg praecoxa <development>/
1. developed/
2. rudimentary/

#2109. <male> fifth left swimming leg praecoxa <fusion>/
1. fused to coxa/
2. separated from the coxa/

#2110. <male> fifth left swimming leg praecoxa <condition of the fusion>/
1. completely/
2. incompletely/

#2111. <male> fifth left swimming leg praecoxa <ornamentation>/
1. without ornamentation/

2. with ornamentation/

#2112. <male> fifth left swimming leg coxa <form of the inner side>/
1. concave inner side/
2. convex inner side/

#2113. <male> fifth left swimming leg coxa <presence of like-teeth on inner border>/
1. without teeth-like structures/
2. with teeth-like structures/

#2114. <male> fifth left swimming leg coxa <condition of the like-teeth on inner border>/
1. continuously/
2. discontinuously/

#2115. <male> fifth left swimming leg coxa <presence of process>/
1. without process/
2. with process/

#2116. <male> fifth left swimming leg coxa <form of the process>/
1. triangulated/
2. conical/

#2117. <male> fifth left swimming leg coxa <location of the process>/
1. on posterior surface/
2. on marginal surface/

#2118. <male> fifth left swimming leg coxa <location of the process>/
1. outer side/
2. inner side/

#2119. <male> fifth left swimming leg coxa <inserted of the process>/
1. proximally inserted/
2. distally inserted/

#2120. <male> fifth left swimming leg coxa <projection of the process>/
1. not projecting over basis/
2. projecting over basis/

#2121. <male> fifth left swimming leg coxa <projection reach on the basis>/
1. reaching to the proximal surface/
2. reaching to the medial surface/

#2122. <male> fifth left swimming leg coxa <presence of sensilla on process>/
1. with sensilla/
2. without sensilla/

#2123. <male> fifth left swimming leg coxa <form of the sensilla on process>/

1. stout/
2. slender/

#2124. <male> fifth left swimming leg coxa <shape of the sensilla on process>/
1. triangular/
2. rounded/

#2125. <male> fifth left swimming leg coxa <location of the sensilla on process>/
1. at apex/
2. at nadir/

#2126. <male> fifth left swimming leg coxa <relative length of the sensilla on process>/
1. not longer 2x than insertion basis/
2. longer 2x than insertion basis/

#2127. <male> fifth left swimming leg coxa <presence of internal extension of the segment>/
1. without marginal extension/
2. with marginal extension/

#2128. <male> fifth left swimming leg coxa <form of extension of the segment>/
1. triangular/
2. sub-triangular/

#2129. <male> fifth left swimming leg coxa <location of extension of the segment>/
1. at outer side/
2. at inner side/

#2130. <male> fifth left swimming leg coxa <insertion of extension of the segment>/
1. distally inserted/
2. medially inserted/

#2131. <male> fifth left swimming leg coxa <presence of intumescence>/
1. without swelling/
2. with swelling/

#2132. <male> fifth left swimming leg coxa <location of the intumescence>/
1. on inner side/
2. on outer side/

#2133. <male> fifth left swimming leg coxa <position of the intumescence>/
1. medially/
2. distally/

#2134. <male> fifth left swimming leg coxa <presence of seta>/
1. without seta/
2. with seta/

#2135. <male> fifth left swimming leg coxa <location of the seta>/
1. on inner side/

2. on outer side/

#2136. <male> fifth left swimming leg coxa <presence of spinules>/
1. without spinules/
2. with spinules/

#2137. <male> fifth left swimming leg coxa <arrangement of spinules>/
1. as a row/
2. as a patch/

#2138. <male> fifth left swimming leg basis <general form>/
1. sub-cylindrical/
2. sub-triangular/

#2139. <male> fifth left swimming leg basis <comparison of size between inner and outer side>/
1. equal size between inner and outer side/
2. unequal size between inner and outer side/

#2140. <male> fifth left swimming leg basis <relation of size between inner and outer side>/
1. shorter outer than inner side/
2. shorter inner than outer side/

#2141. <male> fifth left swimming leg basis <form of the inner side>/
1. with rectilinear inner side/
2. with concave inner side/
3. with convex inner side/

#2142. <male> fifth left swimming leg basis <observation of rounded inner expansion>/
1. rounded internal proximal expansion absent/
2. rounded internal proximal expansion present/

#2143. <male> fifth left swimming leg basis <presence of outgrowth>/
1. without outgrowth/
2. with outgrowth/

#2144. <male> fifth left swimming leg basis <location of outgrowth>/
1. on anterior surface/
2. on posterior surface/

#2145. <male> fifth left swimming leg basis <presence of groove>/
1. without groove/
2. with groove/

#2146. <male> fifth left swimming leg basis <depth of groove>/
1. deep/
2. shallow/

#2147. <male> fifth left swimming leg basis <depth of groove>/

1. obliquely/
2. longitudinally/

#2148. <male> fifth left swimming leg basis <insertion of groove>/
1. on anterior surface/
2. on posterior surface/

#2149. <male> fifth left swimming leg basis <reach of groove>/
1. not reaching the endopodal lobe/
2. reaching the endopodal lobe/

#2150. <male> fifth left swimming leg basis <ornamentation of the groove>/
1. not ornamented/
2. ornamented/

#2151. <male> fifth left swimming leg basis <type of ornamentation of the groove>/
1. with tubercles/
2. without tubercles/

#2152. <male> fifth left swimming leg basis <location of ornamentation of the groove>/
1. throughout of the inner margin/
2. throughout of the outer margin/

#2153. <male> fifth left swimming leg basis <presence of protuberance>/
1. absence of protuberance/
2. presence of protuberance/

#2154. <male> fifth left swimming leg basis <quantify of protuberances>/
1. single/
2. double/

#2155. <male> fifth left swimming leg basis <form of the protuberance>/
1. semicircular/
2. circular/

#2156. <male> fifth left swimming leg basis <location of the protuberance>/
1. on inner margin/
2. on outer margin/

#2157. <male> fifth left swimming leg basis <ornamentation of the protuberance>/
1. ornamented/
2. not ornamented/

#2158. <male> fifth left swimming leg basis <size of the ornamentation on the protuberance>/
1. with tubercles/
2. with denticles/

#2159. <male> fifth left swimming leg basis <arrangement of the ornamentation on the protuberance>/
1. as a row/
2. as a patch/

#2160. <male> fifth left swimming leg basis <form of the arrangement of the ornamentation on the protuberance>/
1. minutely granular/
2. not minutely granular/

#2161. <male> fifth left swimming leg basis <coverage of the arrangement of the ornamentation on the protuberance>/
1. covering the element/
2. not covering the element/

#2162. <male> fifth left swimming leg basis <extension of the arrangement of the ornamentation on the protuberance>/
1. until to adjacent surface/
2. not until to adjacent surface/

#2163. <male> fifth left swimming leg basis <presence of seta>/
1. without seta/
2. with seta/

#2164. <male> fifth left swimming leg basis <insertion of the seta>/
1. innerly inserted/
2. outerly inserted/

#2165. <male> fifth left swimming leg basis <reach of the seta>/
1. not longer 2x than origin segment/
2. longer 2x than origin segment/

#2166. <male> fifth left swimming leg basis <presence of the minutely granules>/
1. absence of minutely granular/
2. presence of minutely granular/

#2167. <male> fifth left swimming leg basis <arrangement of the minutely granules>/
1. as a row/
2. as a patch/

#2168. <male> fifth left swimming leg basis <location of the minutely granular>/
1. innerly/
2. outerly/

#2169. <male> fifth left swimming leg endopod <fusion between segments 1 and 2>/
1. segments 1 and 2 separated/
2. segments 1 and 2 fused/

#2170. <male> fifth left swimming leg endopod <fusion between segment 2 and 3>/
1. segments 2 and 3 separated/
2. segments 2 and 3 fused/

#2171. <male> fifth left swimming leg endopod <fusion condition between segment 2 and 3>/
1. complete suture line/
2. incomplete suture line/

#2172. <male> fifth left swimming leg endopod <segmentation>/
1. 1-segmented/
2. 2-segmented/

#2173. <male> fifth left swimming leg endopod <general form>/
1. stout/
2. slender/

#2174. <male> fifth left swimming leg endopod <fusion>/
1. separated from the basis/
2. fused to basis/

#2175. <male> fifth left swimming leg endopod <condition of the fusion>/
1. completely/
2. incompletely/

#2176. <male> fifth left swimming leg endopod <location of the fusion>/
1. through distal inner rim/
2. through distal inner margin/

#2177. <male> fifth left swimming leg endopod <presence of suture line>/
1. without suture line/
2. with suture line/

#2178. <male> fifth left swimming leg endopod <condition of suture line>/
1. completely/
2. incompletely/

#2179. <male> fifth left swimming leg endopod <location of suture line>/
1. on posterior surface/
2. on anterior surface/

#2180. <male> fifth left swimming leg endopod <ornamentation present>/
1. not ornamented/
2. ornamented/

#2181. <male> fifth left swimming leg endopod <ornamented segment>/
1. ornamented on segment 1/
2. ornamented on segment 2/

#2182. <male> fifth left swimming leg endopod <location of the ornamentation>/

1. on inner side/
2. on outer side/

#2183. <male> fifth left swimming leg endopod <spinules present>/
1. with spinules/
2. without spinules/

#2184. <male> fifth left swimming leg endopod <quantity of spinules>/
1. four elements/
2. more than four elements/

#2185. <male> fifth left swimming leg endopod <arrangement of the ornamentation>/
1. as a row/
2. as a patch/

#2186. <male> fifth left swimming leg endopod <insertion of ornamentation>/
1. terminally/
2. sub-terminally/

#2187. <male> fifth left swimming leg endopod <presence of setules>/
1. row of setules absent/
2. row of setules present/

#2188. <male> fifth left swimming leg endopod <presence of seta>/
1. without seta/
2. with seta/

#2189. <male> fifth left swimming leg exopod <fusion between segment 1 and 2>/
1. segments 1 and 2 separated/
2. segments 1 and 2 fused/

#2190. <male> fifth left swimming leg exopod <fusion between segments 2 and 3>/
1. segments 2 and 3 separated/
2. segments 2 and 3 fused/

#2191. <male> fifth left swimming leg exopod <segmentation>/
1. 1-segmented/
2. 2-segmented/

#2192. <male> fifth left swimming leg exopod <general form>/
1. stout/
2. slender/

#2193. <male> fifth left swimming leg exopod <fusion>/
1. separated from the basis/
2. fused to basis/

#2194. <male> fifth left swimming leg exopod <condition of the fusion>/
1. completely/

2. incompletely/

#2195. <male> fifth left swimming leg exopod 1 <general form of the segment>/
1. sub-triangular/
2. sub-cylindrical/

#2196. <male> fifth left swimming leg exopod 1 <size relation of the segment>/
1. longer than broad/
2. broader than long/

#2197. <male> fifth left swimming leg exopod 1 <comparison of size between inner and outer side>/
1. equal size between inner and outer side/
2. unequal size between inner and outer side/

#2198. <male> fifth left swimming leg exopod 1 <relation of size between inner and outer side>/
1. shorter outer than inner side/
2. shorter inner than outer side/

#2199. <male> fifth left swimming leg exopod 1 <form of the inner side>/
1. rectilinear inner side/
2. concave inner side/
3. convex inner side/

#2200. <male> fifth left swimming leg exopod 1 <form of the outer side>/
1. concave outer side/
2. rectilinear outer side/
3. convex outer side/

#2201. <male> fifth left swimming leg exopod 1 <presence of intumescence>/
1. without swelling/
2. with swelling/

#2202. <male> fifth left swimming leg exopod 1 <form of the intumescence>/
1. triangular/
2. semicircular/

#2203. <male> fifth left swimming leg exopod 1 <position of the intumescence>/
1. distally/
2. medially/
3. proximally/

#2204. <male> fifth left swimming leg exopod 1 <location of the intumescence>/
1. on inner margin/
2. on outer margin/

#2205. <male> fifth left swimming leg exopod 1 <presence of extension of the segment>/

1. without marginal extension/
2. with marginal extension/

#2206. <male> fifth left swimming leg exopod 1 <form of the extension of the segment>/
1. sub-triangular/
2. rounded/

#2207. <male> fifth left swimming leg exopod 1 <location of the extension of the segment>/
1. at inner side/
2. at outer side/

#2208. <male> fifth left swimming leg exopod 1 <insertion of the extension of the segment>/
1. distally inserted/
2. medially inserted/

#2209. <male> fifth left swimming leg exopod 1 <presence of process>/
1. without process/
2. with process/

#2210. <male> fifth left swimming leg exopod 1 <quantify of process>/
1. single/
2. double/

#2211. <male> fifth left swimming leg exopod 1 <local of the single process>/
1. proximal/
2. medial/

#2212. <male> fifth left swimming leg exopod 1 <local of the double process>/
1. and other medial/
2. and other distal/

#2213. <male> fifth left swimming leg exopod 1 <position of the process>/
1. posteriorly/
2. marginally/

#2214. <male> fifth left swimming leg exopod 1 <presence of lobe>/
1. without lobe/
2. with lobe/

#2215. <male> fifth left swimming leg exopod 1 <number of lobe>/
1. single/
2. double/

#2216. <male> fifth left swimming leg exopod 1 <form of lobe>/
1. semicircular/
2. circular/

#2217. <male> fifth left swimming leg exopod 1 <insertion of lobe>/
1. medially inserted/
2. distally inserted/

#2218. <male> fifth left swimming leg exopod 1 <location of lobe>/
1. on inner side/
2. on outer side/

#2219. <male> fifth left swimming leg exopod 1 <presence of coverage on lobe>/
1. covered/
2. not covered/

#2220. <male> fifth left swimming leg exopod 1 <type of ornamentation on lobe>/
1. by setules/
2. by tubercles/

#2221. <male> fifth left swimming leg exopod 1 <presence of outer spine>/
1. without outer spine/
2. with outer spine/

#2222. <male> fifth left swimming leg exopod 1 <presence of seta>/
1. absence seta/
2. presence seta/

#2223. <male> fifth left swimming leg exopod 2 <general form of the segment>/
1. sub-triangular/
2. digitiform/

#2224. <male> fifth left swimming leg exopod 2 <size relation of the segment>/
1. longer than broad/
2. broader than long/

#2225. <male> fifth left swimming leg exopod 2 <comparison of size between inner and outer side>/
1. equal size between inner and outer side/
2. unequal size between inner and outer side/

#2226. <male> fifth left swimming leg exopod 2 <relation of size between inner and outer side>/
1. shorter outer than inner side/
2. shorter inner than outer side/

#2227. <male> fifth left swimming leg exopod 2 <form of the inner side>/
1. disform inner side/
2. uniform inner side/

#2228. <male> fifth left swimming leg exopod 2 <form of the outer side>/
1. with rectilinear outer side/

2. with convex outer side/

#2229. <male> fifth left swimming leg exopod 2 <presence of setulose pad>/
1. setulose pad absent/
2. setulose pad present/

#2230. <male> fifth left swimming leg exopod 2 <condition of the setulose pad>/
1. rounded prominently/
2. not rounded prominently/

#2231. <male> fifth left swimming leg exopod 2 <position of the prominently pad>/
1. proximally/
2. medially/

#2232. <male> fifth left swimming leg exopod 2 <location of the intumescence>/
1. on inner side/
2. on outer side/

#2233. <male> fifth left swimming leg exopod 2 <observation of inflated medial region on anterior surface>/
1. inflated medial region absent/
2. inflated medial region present/

#2234. <male> fifth left swimming leg exopod 2 <ornamentation of the inflated medial region>/
1. setulose/
2. spinulese/

#2235. <male> fifth left swimming leg exopod 2 <insertion of the inflated medial region>/
1. anteriorly/
2. posterioly/

#2236. <male> fifth left swimming leg exopod 2 <presence of distal process>/
1. distal process absent/
2. distal process present/

#2237. <male> fifth left swimming leg exopod 2 <presence of digitiform distal process>/
1. digitiform/
2. acute-form/

#2238. <male> fifth left swimming leg exopod 2 <denticular form present on the process>/
1. non denticulate/
2. denticulate/

#2239. <male> fifth left swimming leg exopod 2 <condition of the denticular form>/
1. bicuspidate/
2. not bicuspidate/

#2240. <male> fifth left swimming leg exopod 2 <ornamentation on the distal process>/

1. without tranverse row of denticles/
2. with tranverse row of denticles/

#2241. <male> fifth left swimming leg exopod 2 <addictional ornamentation on the distal process present>/
1. plus oblique row of 5 denticles/
2. none oblique row of 5 denticles/

#2242. <male> fifth left swimming leg exopod 2 <position of insertion of the process>/
1. at anterior surface/
2. at posterior surface/

#2243. <male> fifth left swimming leg exopod 2 <process direction>/
1. innerly directed/
2. not innerly directed/

#2244. <male> fifth left swimming leg exopod 2 <presence of the seta>/
1. with seta/
2. without seta/

#2245. <male> fifth left swimming leg exopod 2 <shape seta>/
1. spiniform/
2. non spiniform/

#2246. <male> fifth left swimming leg exopod 2 <ornamentation by spinules on spiniform seta>/
1. not ornamented by spinules/
2. ornamented by spinules/

#2247. <male> fifth left swimming leg exopod 2 <reach of the seta>/
1. not surpassing the distal-point of the segment/
2. surpassing the distal-point of the segment/

#2248. <male> fifth left swimming leg exopod 2 <presence of spine leg>/
1. without outer spine/
2. with outer spine/

#2249. <male> fifth left swimming leg exopod 2 <observation of terminal claw>/
1. terminal claw absent/
2. terminal claw present/

#2250. <male> fifth left swimming leg exopod 2 <comparison size to right leg's terminal claw>/
1. 1.5x longer than the right terminal claw/
2. 3x longer than the right terminal claw/

#2251. <male> fifth left swimming leg exopod 2 <rigidity of the terminal claw>/
1. not sclerotized/
2. sclerotized/

#2252. <male> fifth left swimming leg exopod 2 <general form of the terminal claw>/
1. arched/
2. sinuous/

#2253. <male> fifth left swimming leg exopod 2 <direction of the terminal claw>/
1. inward/
2. outward/

#2254. <male> fifth left swimming leg exopod 2 <presence of outer spine on terminal claw>/
1. without outer spine absent/
2. with outer spine present/

#2255. <male> fifth right swimming leg <number of rami>/
1. uniramous/
2. biramous/

#2256. <male> fifth right swimming leg praecoxa <presence of praecoxa>/
1. absent/
2. present/

#2257. <male> fifth right swimming leg praecoxa <fusion>/
1. separated from the coxa/
2. fused to coxa/

#2258. <male> fifth right swimming leg praecoxa <condition of the fusion>/
1. completely/
2. incompletely/

#2259. <male> fifth right swimming leg praecoxa <ornamentation>/
1. without ornamentation/
2. with ornamentation/

#2260. <male> fifth right swimming leg coxa <form of the inner side>/
1. concave inner side/
2. convex inner side/

#2261. <male> fifth right swimming leg coxa <presence of like-teeth on inner side>/
1. without teeth-like structures/
2. with teeth-like structures/

#2262. <male> fifth right swimming leg coxa <condition of the like-teeth on inner side>/
1. continuously/
2. discontinuously/

#2263. <male> fifth right swimming leg coxa <presence of process>/
1. without process/
2. with process/

#2264. <male> fifth right swimming leg coxa <form of the process>/
1. rounded/
2. conical/

#2265. <male> fifth right swimming leg coxa <inserted of the process>/
1. proximally inserted/
2. distally inserted/

#2266. <male> fifth right swimming leg coxa <location of the process>/
1. on posterior surface/
2. on lateral surface/

#2267. <male> fifth right swimming leg coxa <location of the process>/
1. closest to the outer rim/
2. closest to the inner rim/

#2268. <male> fifth right swimming leg coxa <projection of the process>/
1. not projecting over basis/
2. projecting over basis/

#2269. <male> fifth right swimming leg coxa <reach of the projection of the process>/
1. not beyond the first third/
2. beyond the first third/

#2270. <male> fifth right swimming leg coxa <projection of the process beyond of the first third>/
1. until the medial surface/
2. until the distal surface/

#2271. <male> fifth right swimming leg coxa <presence of triangular protuberance on the process >/
1. without triangular protuberance innerly/
2. with triangular protuberance innerly/

#2272. <male> fifth right swimming leg coxa <presence of sensilla on process>/
1. without sensilla/
2. with sensilla/

#2273. <male> fifth right swimming leg coxa <form of the sensilla on process>/
1. stout/
2. slender/

#2274. <male> fifth right swimming leg coxa <location of the sensilla on process>/

1. at apex/
2. at nadir/

#2275. <male> fifth right swimming leg coxa <relative length of the sensilla on process>/
1. not longer 2x than basal insertion/
2. longer 2x than basal insertion/

#2276. <male> fifth right swimming leg coxa <presence of internal extension of the segment>/
1. without marginal extension/
2. with marginal extension/

#2277. <male> fifth right swimming leg coxa <presence of seta>/
1. without seta/
2. with seta/

#2278. <male> fifth right swimming leg coxa <presence of spinules>/
1. without spinules/
2. with spinules/

#2279. <male> fifth right swimming leg coxa <arrangement of spinules>/
1. as a row/
2. as a patch/

#2280. <male> fifth right swimming leg basis <general form>/
1. trapezoidal/
2. cylindrical/

#2281. <male> fifth right swimming leg basis <comparison of size between inner and outer side>/
1. equal size between inner and outer side/
2. unequal size between inner and outer side/

#2282. <male> fifth right swimming leg basis <relation of size between inner and outer side>/
1. shorter outer than inner side/
2. shorter inner than outer side/

#2283. <male> fifth right swimming leg basis <form of the inner side>/
1. concave inner side/
2. rectilinear inner side/
3. convex inner side/

#2284. <male> fifth right swimming leg basis <presence of tumescence>/
1. tumescence absent/
2. tumescence present/

#2285. <male> fifth right swimming leg basis <form of the tumescence>/
1. not inflated/
2. inflated/

#2286. <male> fifth right swimming leg basis <inflated process condition>/
1. bilobed/
2. not bilobed/

#2287. <male> fifth right swimming leg basis <tumescence position condition>/
1. restricted on inner surface/
2. unrestricted on inner surface/

#2288. <male> fifth right swimming leg basis <position of the tumescence>/
1. proximally/
2. medially/

#2289. <male> fifth right swimming leg basis <observation of protuberance leg>/
1. without protuberance/
2. with protuberance/

#2290. <male> fifth right swimming leg basis <quantity of protuberance leg>/
1. single/
2. double/

#2291. <male> fifth right swimming leg basis <form of the protuberance leg>/
1. semicircular/
2. triangular/

#2292. <male> fifth right swimming leg basis <location of the protuberance>/
1. on inner side/
2. on outer side/

#2293. <male> fifth right swimming leg basis <position of the protuberance>/
1. proximally/
2. medially/
3. distally/

#2294. <male> fifth right swimming leg basis <ornamentation of the protuberance>/
1. not ornamented/
2. ornamented/

#2295. <male> fifth right swimming leg basis <size of the ornamentation on the protuberance>/
1. with tubercles/
2. with denticles/

#2296. <male> fifth right swimming leg basis <arrangement of the ornamentation on the protuberance>/
1. as a row/
2. as a patch/

#2297. <male> fifth right swimming leg basis <arrangement of the ornamentation on the protuberance>/
1. minutely granular/

2. not minutely granular/

#2298. <male> fifth right swimming leg basis <coverage of the arrangement of the ornamentation on the protuberance>/
1. covering the element/
2. not covering the element/

#2299. <male> fifth right swimming leg basis <extension of the arrangement of the ornamentation on the protuberance>/
1. until to adjacent surface/
2. not until to adjacent surface/

#2300. <male> fifth right swimming leg basis <presence of the minutely granules>/
1. absence of distinct minutely granular/
2. presence of distinct minutely granular/

#2301. <male> fifth right swimming leg basis <arrangement of the minutely granules>/
1. as a row/
2. as a patch/

#2302. <male> fifth right swimming leg basis <location of the minutely granules>/
1. proximally/
2. medially/
3. distally/

#2303. <male> fifth right swimming leg basis <position of the minutely granules>/
1. on inner side/
2. on outer side/

#2304. <male> fifth right swimming leg basis <observation of additional internal process>/
1. additional inner process absent/
2. additional inner process present/

#2305. <male> fifth right swimming leg basis <form of the process present>/
1. digitiform/
2. triangular/

#2306. <male> fifth right swimming leg basis <position of the inner process>/
1. medially/
2. distally/

#2307. <male> fifth right swimming leg basis <presence of posterior groove>/
1. without posterior groove/
2. with posterior groove/

#2308. <male> fifth right swimming leg basis <depth of groove>/
1. deep/
2. shallow/

#2309. <male> fifth right swimming leg basis <depth of groove>/
1. obliquely/
2. latitudinally/
3. longitudinally/

#2310. <male> fifth right swimming leg basis <reach of groove>/
1. reaching the endopodal lobe/
2. not reaching the endopodal lobe/

#2311. <male> fifth right swimming leg basis <ornamentation of the groove>/
1. ornamented/
2. not ornamented/

#2312. <male> fifth right swimming leg basis <type of ornamentation of the groove>/
1. with tubercles/
2. with spinules/

#2313. <male> fifth right swimming leg basis <location of ornamentation of the groove>/
1. throughout of the inner border/
2. throughout of the outer border/

#2314. <male> fifth right swimming leg basis <presence of seta>/
1. without seta/
2. with seta/

#2315. <male> fifth right swimming leg basis <insertion of the seta leg>/
1. innerly inserted/
2. outerly inserted/

#2316. <male> fifth right swimming leg basis <location of the seta leg>/
1. on anterior surface/
2. on posterior surface/

#2317. <male> fifth right swimming leg basis <reach of the seta>/
1. not longer 2x than origin segment/
2. longer 2x than origin segment/

#2318. <male> fifth right swimming leg basis <observation of posterior protrusion>/
1. posterior protrusion absent/
2. posterior protrusion present/

#2319. <male> fifth right swimming leg basis <observation of distal tegument expansion>/
1. distal tegument expansion absent/
2. distal tegument expansion present/

#2320. <male> fifth right swimming leg basis <form of the distal process>/
1. triangular/
2. rounded/

#2321. <male> fifth right swimming leg basis <insertion surface of the distal triangular process>/
1. anteriorly/
2. posteriorly/

#2322. <male> fifth right swimming leg endopod <presence of endopodite>/
1. present/
2. absent/

#2323. <male> fifth right swimming leg endopod <fusion>/
1. separated from the basis/
2. fused to basis/

#2324. <male> fifth right swimming leg endopod <location of insertion>/
1. on posterior surface/
2. on anterior surface/

#2325. <male> fifth right swimming leg endopod <fusion between segments 1 and 2>/
1. segments 1 and 2 separated/
2. segments 1 and 2 fused/

#2326. <male> fifth right swimming leg endopod <fusion between segment 2 and 3>/
1. segments 2 and 3 separated/
2. segments 2 and 3 fused/

#2327. <male> fifth right swimming leg endopod <segmentation>/
1. 1-segmented/
2. 2-segmented/

#2328. <male> fifth right swimming leg endopod <general form leg>/
1. stout/
2. slender/

#2329. <male> fifth right swimming leg endopod <ornamentation leg>/
1. not ornamented/
2. ornamented/

#2330. <male> fifth right swimming leg endopod <type of ornamentation leg>/
1. with spinules/
2. with setules/

#2331. <male> fifth right swimming leg endopod <arrangement of the ornamentation>/
1. as a row/

2. as a patch/

#2332. <male> fifth right swimming leg endopod <location of the ornamentation>/
1. on inner side/
2. on outer side/

#2333. <male> fifth right swimming leg endopod <insertion of ornamentation>/
1. terminally/
2. sub-terminally/

#2334. <male> fifth right swimming leg endopod <presence of seta leg>/
1. without seta/
2. with seta/

#2335. <male> fifth right swimming leg exopod <fusion between segment 1 and 2>/
1. segments 1 and 2 separated/
2. segments 1 and 2 fused/

#2336. <male> fifth right swimming leg exopod <fusion between segments 2 and 3>/
1. segments 2 and 3 separated/
2. segments 2 and 3 fused/

#2337. <male> fifth right swimming leg exopod <segmentation>/
1. 1-segmented/
2. 2-segmented/

#2338. <male> fifth right swimming leg exopod <general form>/
1. stout/
2. slender/

#2339. <male> fifth right swimming leg exopod <fusion>/
1. separated from the basis/
2. fused to basis/

#2340. <male> fifth right swimming leg exopod <condition of the fusion leg>/
1. completely/
2. incompletely/

#2341. <male> fifth right swimming leg exopod 1 <general form of the segment>/
1. sub-cylindrical/
2. trapezium/

#2342. <male> fifth right swimming leg exopod 1 <size relation of the segment>/
1. longer than broad/
2. broader than long/

#2343. <male> fifth right swimming leg exopod 1 <size proportion of the segment>/
1. nearly 1.25 times/

2. nearly 2 times/

#2344. <male> fifth right swimming leg exopod 1 <comparison of size between inner and outer side>/
1. equal size between both sides/
2. unequal size between both sides/

#2345. <male> fifth right swimming leg exopod 1 <relation of size between inner and outer side>/
1. shorter outer than inner side/
2. shorter inner than outer side/

#2346. <male> fifth right swimming leg exopod 1 <form of the inner side>/
1. rectilinear inner side/
2. concave inner side/
3. convex inner side/

#2347. <male> fifth right swimming leg exopod 1 <form of the outer side>/
1. rectilinear outer side/
2. concave outer side/
3. convex outer side/

#2348. <male> fifth right swimming leg exopod 1 <presence of extension of the segment>/
1. without marginal extension/
2. with marginal extension/

#2349. <male> fifth right swimming leg exopod 1 <form of the extension of the segment>/
1. sub-triangular/
2. acute/

#2350. <male> fifth right swimming leg exopod 1 <insertion of the extension of the segment>/
1. distally inserted/
2. medially inserted/

#2351. <male> fifth right swimming leg exopod 1 <location of the extension of the segment>/
1. at inner rim/
2. at outer rim/

#2352. <male> fifth right swimming leg exopod 1 <presence of ornamentation on extension of the segment>/
1. spinules absent/
2. spinules present/

#2353. <male> fifth right swimming leg exopod 1 <presence of process>/
1. without process/
2. with process/

#2354. <male> fifth right swimming leg exopod 1 <general form of the process>/
1. triangular/
2. rounded/

#2355. <male> fifth right swimming leg exopod 1 <form of the process>/

1. arched/
2. rectilinear/

#2356. <male> fifth right swimming leg exopod 1 <direction of the arched process>/
1. externally directed/
2. internally directed/

#2357. <male> fifth right swimming leg exopod 1 <form tip of the process>/
1. sharp tip/
2. blunt tip/

#2358. <male> fifth right swimming leg exopod 1 <rigidity of the process>/
1. sclerotized/
2. not sclerotized/

#2359. <male> fifth right swimming leg exopod 1 <ornamentation of the tip of the process>/
1. without ornamentation/
2. with ornamentation/

#2360. <male> fifth right swimming leg exopod 1 <insertion of the process>/
1. proximally inserted/
2. distally inserted/

#2361. <male> fifth right swimming leg exopod 1 <location of the process>/
1. at marginal surface/
2. at posterior surface/

#2362. <male> fifth right swimming leg exopod 1 <projection of the process>/
1. not projecting over next segment/
2. projecting over next segment/

#2363. <male> fifth right swimming leg exopod 1 <presence of outer spine leg>/
1. without outer spine/
2. with outer spine/

#2364. <male> fifth right swimming leg exopod 1 <presence of seta leg>/
1. without seta/
2. with seta/

#2365. <male> fifth right swimming leg exopod 1 <number of setae>/
1. single/
2. double/

#2366. <male> fifth right swimming leg exopod 1 <form of the seta leg>/
1. spiniform/
2. non-spiniform/

#2367. <male> fifth right swimming leg exopod 1 <comparison size of the seta>/
1. equal size to origin segment/

2. unequal size to origin segment/

#2368. <male> fifth right swimming leg exopod 1 <comparison size of the seta>/
1. larger/
2. smaller/

#2369. <male> fifth right swimming leg exopod 1 <observation of internal prominence>/
1. internal prominence absent/
2. internal prominence present/

#2370. <male> fifth right swimming leg exopod 1 <form of the internal prominence>/
1. acute/
2. digitiform/

#2371. <male> fifth right swimming leg exopod 1 <observation of posterior lamella>/
1. lamella on posterior surface absent/
2. lamella on posterior surface present/

#2372. <male> fifth right swimming leg exopod 1 <shape of the posterior lamella>/
1. trapezoidal form/
2. lobular form/

#2373. <male> fifth right swimming leg exopod 1 <reach of the lamella on posterior surface>/
1. surpassing to margin/
2. not surpassing to margin/

#2374. <male> fifth right swimming leg exopod 1 <lamella surpassing to lateral margin>/
1. outerly/
2. distally/

#2375. <male> fifth right swimming leg exopod 2 <general form of the segment>/
1. cylindrical/
2. elliptical/

#2376. <male> fifth right swimming leg exopod 2 <size relation of the segment>/
1. longer than broad/
2. broader than long/

#2377. <male> fifth right swimming leg exopod 2 <size proportion of the segment>/
1. nearly 2 times/
2. nearly 2.5 times/

#2378. <male> fifth right swimming leg exopod 2 <comparison of size between sides>/
1. equal size between both sides/
2. unequal size between both sides/

#2379. <male> fifth right swimming leg exopod 2 <form of the inner side>/
1. disform inner side/
2. uniform inner side/

#2380. <male> fifth right swimming leg exopod 2 <form of the outer side>/
1. concave outer side/
2. convex outer side/

#2381. <male> fifth right swimming leg exopod 2 <presence of posterior proximal swelling>/
1. without posterior proximal swelling/
2. with posterior proximal swelling/

#2382. <male> fifth right swimming leg exopod 2 <presence of inner-posterior process>/
1. inner-posterior process absent/
2. inner-posterior process present/

#2383. <male> fifth right swimming leg exopod 2 <form of the inner process>/
1. tubercular/
2. sub-triangular/

#2384. <male> fifth right swimming leg exopod 2 <position of process>/
1. medially/
2. distally/

#2385. <male> fifth right swimming leg exopod 2 <presence of expansion of the segment>/
1. without marginal expansion/
2. with marginal expansion/

#2386. <male> fifth right swimming leg exopod 2 <position of the marginal extension>/
1. medial/
2. distal/

#2387. <male> fifth right swimming leg exopod 2 <location of the marginal extension>/
1. innerly/
2. outerly/

#2388. <male> fifth right swimming leg exopod 2 <observation of curved ridge on distal posterior surface>/
1. curved ridge on distal posterior surface absent/
2. curved ridge on distal posterior surface present/

#2389. <male> fifth right swimming leg exopod 2 <presence of chitinous knobs>/
1. chitinous knobs absent/
2. chitinous knobs present/

#2390. <male> fifth right swimming leg exopod 2 <quantify of chitinous knobs>/
1. with 1-2 posteriorly/
2. with 3-6 posteriorly/

#2391. <male> fifth right swimming leg exopod 2 <presence of spine leg>/
1. without outer spine/
2. with outer spine/

#2392. <male> fifth right swimming leg exopod 2 <insertion of spine leg>/
1. inserted premedially/
2. inserted medially/
3. inserted sub-distally/

#2393. <male> fifth right swimming leg exopod 2 <form of the spine leg>/
1. arched/
2. rectilinear/

#2394. <male> fifth right swimming leg exopod 2 <direction of the spine leg>/
1. externally directed/
2. internally directed/

#2395. <male> fifth right swimming leg exopod 2 <ornamentation on the inner side>/
1. not ornamented innerly/
2. ornamented innerly/

#2396. <male> fifth right swimming leg exopod 2 <type of ornamentation on the spine>/
1. by denticles/
2. by spinules/

#2397. <male> fifth right swimming leg exopod 2 <arrangement of the ornamentation on the spine>/
1. as a row/
2. as a patch/

#2398. <male> fifth right swimming leg exopod 2 <presence of ornamentation on the outer side>/
1. not ornamented outerly/
2. ornamented outerly/

#2399. <male> fifth right swimming leg exopod 2 <type of ornamentation on the spine>/
1. by denticles/
2. by spinules/

#2400. <male> fifth right swimming leg exopod 2 <arrangement of the ornamentation on the spine>/
1. as a row/
2. as a patch/

#2401. <male> fifth right swimming leg exopod 2 <type of the tip of the spine>/
1. blunt tip/
2. sharp tip/

#2402. <male> fifth right swimming leg exopod 2 <form of the tip of the spine>/
1. without apparent curve/
2. with apparent curve/

#2403. <male> fifth right swimming leg exopod 2 <direction of the curvature of the tip of the spine>/
1. innerly directed/
2. outerly directed/

#2404. <male> fifth right swimming leg exopod 2 <comparison size of the spine to insertion segment>/
1. lesser than the length of the exopod 2/
2. larger than or equal to the length of the exopod 2/

#2405. <male> fifth right swimming leg exopod 2 <Spine size condition smaller than insertion segment>/
1. until to 2 times its size/
2. beyond to 2 times its size/

#2406. <male> fifth right swimming leg exopod 2 <Relative size of spine smaller up to 2 times>/
1. 3x/
2. 6x/

#2407. <male> fifth right swimming leg exopod 2 <Relative size of spine smaller beyond to 2 times>/
1. 1.5x/
2. 2x/

#2408. <male> fifth right swimming leg exopod 2 <presence of sensilla leg>/
1. sensilla absent/
2. sensilla present/

#2409. <male> fifth right swimming leg exopod 2 <form of sensilla leg>/
1. spiniform/
2. slender/

#2410. <male> fifth right swimming leg exopod 2 <position of sensilla leg>/
1. proximally/
2. distally/

#2411. <male> fifth right swimming leg exopod 2 <insertion surface of sensilla leg>/
1. on inner surface/
2. on outer surface/

#2412. <male> fifth right swimming leg exopod 2 <observation of terminal claw>/
1. terminal claw absent/
2. terminal claw present/

#2413. <male> fifth right swimming leg exopod 2 <comparison size of the terminal claw to insertion segment>/
1. not equal or longer 1.5 times than insertion segment/
2. equal or longer 1.5 times than insertion segment/

#2414. <male> fifth right swimming leg exopod 2 <rigidity of the terminal claw>/
1. sclerotized/
2. not sclerotized/

#2415. <male> fifth right swimming leg exopod 2 <general form of the terminal claw>/
1. arched/
2. rectilinear/

#2416. <male> fifth right swimming leg exopod 2 <direction of the terminal claw>/

1. inward/
2. outward/

#2417. <male> fifth right swimming leg exopod 2 <presence of curvature at the terminal claw>/
1. without conspicuous curve/
2. with conspicuous curve/

#2418. <male> fifth right swimming leg exopod 2 <location of the enlargement at the terminal claw>/
1. proximally/
2. medially/
3. distally/

#2419. <male> fifth right swimming leg exopod 2 <presence of ornamentation on the terminal claw>/
1. not ornamented innerly/
2. ornamented innerly/

#2420. <male> fifth right swimming leg exopod 2 <type of ornamentation on the terminal claw>/
1. by spinules/
2. by denticles/

#2421. <male> fifth right swimming leg exopod 2 <arrangement of the ornamentation on the terminal claw>/
1. as a row/
2. as a patch/

#2422. <male> fifth right swimming leg exopod 2 <distribution of the ornamentation on the terminal claw>/
1. throughout extension/
2. partially on extension/

#2423. <male> fifth right swimming leg exopod 2 <location of the ornamentation on the terminal claw>/
1. proximally/
2. medially/
3. distally/

#2424. <male> fifth right swimming leg exopod 2 <form of the outer side of the terminal claw>/
1. not ornamented outerly/
2. ornamented outerly/

#2425. <male> fifth right swimming leg exopod 2 <type of the tip of the terminal claw >/
1. blunt tip/
2. sharp tip/

#2426. <male> fifth right swimming leg exopod 2 <form of the tip of the terminal claw >/
1. not curved tip/
2. curved tip/

#2427. <male> fifth right swimming leg exopod 2 <direction of the tip of the terminal claw>/
1. inwards/
2. outwards/

#2428. <male> fifth right swimming leg exopod 2 <presence of constriction at the terminal claw>/
1. without medial constriction/
2. with medial constriction/

#2429. <male> fifth right swimming leg exopod 2 <presence of hyaline process on terminal claw>/
1. hyaline process absent/
2. hyaline process present/

#2430. <male> fifth right swimming leg exopod 2 <insertion surface of the hyaline process on terminal claw>/
1. on proximal third/
2. on medial third/

#2431. <male> fifth right swimming leg exopod 2 <position of hyaline process on terminal claw>/
1. outerlly/
2. innerly/

#2432. <female> body <size compared to male in dorsal view>/
1. smaller and slenderer than male/
2. longer and wider than male/

#2433. <female> prosome <widest region, in dorsal view>/
1. widest at first metasome segment/
2. widest at posterior cephalosome/

#2434. <female> distal margin of the prosomal segments < line of setules presente at posterior margin>/
1. without one line of setules at posterior margin/
2. with one line of setules at posterior margin/

#2435. <female> prosome segments <presence of spinules at posterior margin of prosome segments>/
1. without spinules at prosomal segments/
2. with spinules at least at one prosomal segment/

#2436. <female> Fourth metasome segment <presence of dorsal protuberance>/
1. absence of dorsal protuberance/
2. presence of dorsal protuberance/

#2437. <female> Fourth metasome segment <form of the dorsal protuberance>/
1. digitform/
2. rounded/

#2438. <female> Fourth metasome segment <relative length of the dorsal protuberance>/
1. 2x longer than wide/
2. 2x wider than long/

#2439. <female> Fourth metasome segment <position of dorsal protuberance>/
1. inserted medially/
2. inserted distally/

#2440. <female> Fourth metasome segment <presence of posterior process on dorsal protuberance>/

1. without posterior process/
2. with posterior process/

#2441. <female> Fourth metasome segment <form of the posterior process on protuberance>/
1. acute-form/
2. stout-form/

#2442. <female> Fourth metasome segment <position of the posterior process on protuberance>/
1. proximally/
2. medially/

#2443. <female> Fourth metasome segment <presence of anterior process on dorsal protuberance>/
1. without anterior process/
2. with anterior process/

#2444. <female> Fourth metasome segment <form of the anterior process on protuberance>/
1. acute-form/
2. stout-form/

#2445. <female> Fourth metasome segment <position of the anterior process on protuberance>/
1. proximally/
2. medially/

#2446. <female> fourth metasome segment <presence of sensilla on the dorsal protuberance>/
1. without proximal sensillae present/
2. with proximal sensillae present/

#2447. <female> fourth and fifth metasome segments <fusion or separation indicated by suture presence, absence>/
1. separated/
2. fused/

#2448. <female> fourth and fifth metasome segments <state of the fusion>/
1. totally/
2. partially/

#2449. <female> fourth and fifth metasome segments <location of the fusion>/
1. on lateral surface/
2. on dorsal surface/

#2450. <female> limit between fourth and fifth metasome segments <presence of ornamentation>/
1. without ornamentation/
2. ornamented/

#2451. <female> limit between fourth and fifth metasome segments <type of ornamentation>/
1. with setules/
2. with spinules/

#2452. <female> limit between fourth and fifth metasome segments <arrangement of the spinules>/
1. as a row/

2. as a patch/

#2453. <female> limit between fourth and fifth metasome segments <number of rows>/
1. single/
2. double/

#2454. <female> limit between fourth and fifth metasome segments <row of spinules complete or incomplete>/
1. complete/
2. incomplete/

#2455. <female> limit between fourth and fifth metasome segments <size of spinules>/
1. same size/
2. different size/

#2456. <female> limit between fourth and fifth metasome segments <spinule row arrangement>/
1. entirely over limit (lateral, dorsal)/
2. partially over limit/

#2457. <female> limit between fourth and fifth metasome segments <position of spinules>/
1. dorsally/
2. laterally/

#2458. <female> fifth metasome segment <observation of dorsal setules row medially>/
1. ornamented with dorsal setules row medially/
2. not ornamented with dorsal setules row medially/

#2459. <female> fifth metasome segment <presence of sensillae>/
1. without sensilla/
2. with sensilla/

#2460. <female> fifth metasome segment <sensillae position>/
1. dorsally/
2. laterally/

#2461. <female> fifth metasome segment <presence of epimeral plates>/
1. without epimeral plates/
2. with epimeral plates/

#2462. <female> epimeral plates <symmetry>/
1. asymmetrical/
2. symmetrical/

#2463. <female> right epimeral plates <form>/
1. reduced, as rounded distal corner segment limit/
2. prominent, as projections/

#2464. <female> right epimeral plates <wide>/
1. thinner than the left/
2. not thinner than the left/

#2465. <female> right epimeral plate <projections present>/
1. with projection/
2. without projection/

#2466. <female> right epimeral plate <number of projections>/
1. one each side/
2. two each side/

#2467. <female> right epimeral plate <direction>/
1. posterior-laterally directed/
2. posterior-dorsally directed/

#2468. <female> right epimeral plate <size of projection in relation to genital segment>/
1. not reaching half length of the genital segment/
2. reaching half length of the genital segment/

#2469. <female> right epimeral plate <presence of sensillae>/
1. without sensilla at the apex/
2. with sensilla at the apex/

#2470. <female> right epimeral plate <observation of dorsal-posterior sensillae>/
1. dorsal-posterior sensilla absent/
2. dorsal-posterior sensilla present/

#2471. <female> right epimeral plate <condition of the medial inner sensillae>/
1. slender/
2. stout/

#2472. <female> right epimeral plate <presence of ornamentation>/
1. without ornamentation/
2. ornamented/

#2473. <female> right epimeral plate <type of ornamentation>/
1. with spines/
2. with spinules/

#2474. <female> right epimeral plate <arrangement of ornamentation>/
1. as a row/
2. as a patch/

#2475. <female> right epimeral plate <location of the ornamentation>/
1. innerly/
2. outerlly/

#2476. <female> right epimeral plate <location of the ornamentation>/
1. on dorsal surface/
2. on ventral surface/

#2477. <female> left epimeral plate <presence of expansion>/
1. without expansion/
2. with expansion/

#2478. <female> left epimeral plate <shape of expansion>/
1. semicircular/
2. conical/

#2479. <female> left epimeral plate <position of the expansion>/
1. on posterior surface/
2. on medial surface/

#2480. <female> left epimeral plate <location of the expansion>/
1. ventrally/
2. dorsally/

#2481. <female> left epimeral plate <presence of sensilla on the expansion>/
1. with sensilla/
2. without sensilla/

#2482. <female> left epimeral plate <insertion of sensilla on the expansion>/
1. at tip/
2. at base/

#2483. <female> Urosome <number of segments>/
1. 2-segmented/
2. 3-segmented/
3. 4-segmented/

#2484. <female> Genital double-somite <symmetry in dorsal view>/
1. asymmetrical in dorsal view/
2. symmetrical in dorsal view/

#2485. <female> Genital double-somite <size proportions>/
1. longer than broad/
2. broader than long/

#2486. <female> Genital double-somite <size comparison>/
1. shorter than other urosomites combined/
2. longer than other urosomites combined/

#2487. <female> Genital double-somite <observation of dorsal suture>/
1. dorsal suture at mid-length absent/
2. dorsal suture at mid-length present/

#2488. <female> Genital double-somite <presence of spinules covering the segment>/
1. not covered by spinules/
2. covered by spinules/

#2489. <female> Genital double-somite <swollen or expansion>/
1. with swelling/
2. with expansion/

#2490. <female> Genital double-somite <form of the swollen>/
1. conical/
2. rounded/

#2491. <female> Genital double-somite <form of the lateral expansion>/
1. like a bulging dilation/
2. like an ovoid ventrolateral hypertrophy/

#2492. <female> Genital double-somite <similarity of swollen>/
1. equal size/
2. unequal size/

#2493. <female> Genital double-somite <comparison between left and right swollen>/
1. greater left than right/
2. greater right than left/

#2494. <female> Genital double-somite <position of swollen>/
1. anteriorly/
2. posteriorly/

#2495. <female> Genital double-somite <presence of sensilla>/
1. with sensillae/
2. without sensillae/

#2496. <female> Genital double-somite <total of sensillae on left side>/
1. on both sides/
2. on single side/

#2497. <female> Genital double-somite <location sensillae on single side>/
1. <left side>/
2. <right side>/

#2498. <female> Genital double-somite <number of sensillae on left side>/
1. one/
2. two/

#2499. <female> Genital double-somite <composition of the sensillae on left side>/
1. slender/
2. stout/

#2500. <female> Genital double-somite <shape tip of the sensilla on left side>/
1. with robust apex/
2. with bifid apex/

#2501. <female> Genital double-somite <location of sensillae on left side>/

1. at left rim/
2. at left lateral/

#2502. <female> Genital double-somite <sensilla on lobular base on left side>/
1. not on lobular base/
2. on lobular base/

#2503. <female> Genital double-somite <position of sensillae on left side>/
1. anteriorly/
2. medially/

#2504. <female> Genital double-somite <number of sensillae on right side>/
1. one/
2. two/

#2505. <female> Genital double-somite <composition of the sensillae on right side>/
1. stout/
2. thin/

#2506. <female> Genital double-somite <location of sensillae on right side>/
1. at right rim/
2. at right lateral/

#2507. <female> Genital double-somite <sensilla on lobular base on right side>/
1. not on lobular base/
2. on lobular base/

#2508. <female> Genital double-somite <position of sensillae on right side>/
1. anteriorly/
2. medially/

#2509. <female> Genital double-somite <shape tip of the sensilla on right side>/
1. with robust apex/
2. with bifid apex/

#2510. <female> Genital double-somite <size comparison between sensillae>/
1. of equal size between then/
2. of unequal size between then/

#2511. <female> Genital double-somite <size comparison between sensillae>/
1. left bigger than right/
2. right bigger than left/

#2512. <female> Genital double-somite <lateral protuberance present>/
1. lateral protuberance absent/
2. lateral protuberance present/

#2513. <female> Genital double-somite <shape of the protuberance present>/
1. rounded/

2. wrinkled/

#2514. <female> Genital double-somite <location of the protuberance present>/
1. on the left side/
2. on the right side/

#2515. <female> Genital double-somite <presence of expansion, in dorsal view>/
1. without right posterior rim expanded/
2. with right posterior rim expanded/

#2516. <female> Genital double-somite <projection of expansion>/
1. over next segment/
2. not over next segment/

#2517. <female> Genital double-somite <observation of addictional sensillae on lateral- posterior portion>/
1. without slender sensilla on each posterior rim/
2. with slender sensilla on each posterior rim/

#2518. <female> Genital double-somite <presence of posterior-dorsal process>/
1. without posterior-dorsal process/
2. with posterior-dorsal process/

#2519. <female> Genital double-somite <location of the posterior-dorsal process>/
1. on either side/
2. in a single side/

#2520. <female> Genital double-somite <location of the process in a single side>/
1. <on the right>/
2. <on the left>/

#2521. <female> Genital double-somite <location of the posterior-dorsal process on either side>/
1. single on the left/
2. double on the left/

#2522. <female> Genital double-somite <location of the posterior-dorsal process on either side>/
1. single on the right/
2. double on the right/

#2523. <female> Genital double-somite <observation of genital operculum>/
1. opercular pad absent/
2. opercular pad present/

#2524. <female> Genital double-somite <composition of the genital operculum>/
1. broader than longer/
2. longer than broader/

#2525. <female> Genital double-somite <symmetry of the opercular pad>/
1. symmetrical/
2. asymmetrical/

#2526. <female> Genital double-somite <lateral development of the opercular pad>/
1. developmented laterally/
2. not developmented laterally/

#2527. <female> Genital double-somite <expansion of the opercular pad>/
1. not expanded posteriorly/
2. expanded posteriorly/

#2528. <female> Genital double-somite <coverage of genital operculum>/
1. covering totally/
2. covering partially/

#2529. <female> Genital double-somite <number of genital aperture>/
1. single gonoporal slit/
2. double gonoporal slit/

#2530. <female> Genital double-somite <location of the posterior-central projection>/
1. located ventrally/
2. located laterally/

#2531. <female> Genital double-somite <presence of membrane artrodial>/
1. without arthrodial membrane/
2. with arthrodial membrane/

#2532. <female> Genital double-somite <location of the membrane artrodial>/
1. inserted anteriorly/
2. inserted laterally/

#2533. <female> Genital double-somite <presence of posterior-ventral process in lateral view>/
1. post-genital process absent/
2. post-genital process present/

#2534. <female> Genital double-somite <quantify of posterior-ventral process in lateral view>/
1. single/
2. double/

#2535. <female> Genital double-somite <presence of distal-ventral tumescence, in lateral view>/
1. disto-ventral tumescence absent/
2. disto-ventral tumescence present/

#2536. <female> Genital double-somite <appearance of the ventral integument of the double genital segment>/
1. ventral vertical folds absent/
2. ventral vertical folds present/

#2537. <female> Genital double-somite <observation of dorsal sensilla>/
1. dorsal sensilla absent/
2. dorsal sensilla present/

#2538. <female> Second urosome segment <observation of ventral fusion to anal segment>/
1. without ventral fusion to anal segment/
2. with ventral fusion to anal segment/

#2539. <female> Second urosome segment <observation of elongated spiniform process>/
1. rigth distal process absent/
2. right distal process present/

#2540. <female> Second urosome segment <form of the right distal process present>/
1. spiniform/
2. lobed form/

#2541. <female> Second urosome segment <expansion of right distal process present>/
1. elongated/
2. not elongated/

#2542. <female> Caudal rami <presence of patch of setules on outer surface>/
1. patch of setules on outer surface absent/
2. patch of setules on outer surface present/

#2543. <female> Caudal rami <presence of patch of spinules on outer surface>/
1. patch of spinules on outer surface absent/
2. patch of spinules on outer surface present/

#2544. <female> Rostrum <observation of basal process>/
1. basal process absent/
2. basal process present/

#2545. <female> Rostrum <localization of the basal process>/
1. on right side ventrally/
2. on left side ventrally/

#2546. <female> Antennules <symmetry>/
1. symmetrical/
2. asymmetrical/

#2547. <female> Right antennule <reach of the antennules>/
1. surpassing to genital double-segment/
2. not surpassing to genital double-segment/

#2548. <female> Right antennule <extension of the reach of the antennules>/
1. extending beyond caudal rami/
2. not extending beyond caudal rami/

#2549. <female> Right antennule <limit of the reach of the antennules>/
1. not exceeding the caudal setae/
2. exceeding the caudal setae/

#2550. <female> Right antennule <similarity to male left antennule>/

1. ornamentation pattern equal to male left antennule/
2. ornamentation pattern unequal to male left antennule/

#2551. <female> Right antennule <similarity to ornamentation pattern male left antennule>/
1. mostly/
2. fully/

#2552. <female> Right antennule actual segment 13 <presence of seta>/
1. with seta/
2. without seta/

#2553. <female> Right antennule actual segment 13 <total of setae>/
1. 1/
2. 2/

#2554. <female> Right antennule actual segment 13 <presence of aesthetasc>/
1. with aesthetasc/
2. without aesthetasc/

#2555. <female> Right antennule actual segment 14 <presence of seta>/
1. with seta/
2. without seta/

#2556. <female> Right antennule actual segment 14 <total of seta>/
1. 1/
2. 2/

#2557. <female> Right antennule actual segment 14 <presence of aesthetasc>/
1. with aesthetasc/
2. without aesthetasc/

#2558. <female> Right antennule actual segment 14 <total of aesthetasc>/
1. 1/
2. 2/

#2559. <female> Right antennule actual segment 15 <presence of seta>/
1. with seta/
2. without seta/

#2560. <female> Right antennule actual segment 15 <presence of aesthetasc>/
1. with aesthetasc/
2. without aesthetasc/

#2561. <female> Right antennule actual segment 16 <presence of seta>/
1. with seta/
2. without seta/

#2562. <female> Right antennule actual segment 16 <total of setae>/
1. 1/

2. 2/

#2563. <female> Right antennule actual segment 16 <presence of aesthetasc>/
1. with aesthetasc/
2. without aesthetasc/

#2564. <female> Right antennule actual segment 16 <presence of aesthetasc>/
1. 1/
2. 2/

#2565. <female> Right antennule actual segment 17 <presence of seta>/
1. with seta/
2. without seta/

#2566. <female> Right antennule actual segment 17 <presence of seta>/
1. 1/
2. 2/

#2567. <female> Right antennule actual segment 18 <presence of seta>/
1. with seta/
2. without seta/

#2568. <female> Right antennule actual segment 18 <total of seta>/
1. 1/
2. 2/

#2569. <female> Fifth swimming legs <symmetry>/
1. symmetrical/
2. asymmetrical/

#2570. <female> Fifth swimming legs <number of rami to symmetrical legs>/
1. uniramous/
2. biramous/

#2571. <female> Fifth swimming legs intercoxal plate <relative size of the legs>/
1. wider than long/
2. longer than wide/

#2572. <female> Fifth swimming legs intercoxal plate <fusion state to legs>/
1. fused to legs/
2. separated from the legs/

#2573. <female> Fifth swimming legs praecoxa <presence of praecoxa to symmetrical legs>/
1. with sclerite praecoxal/
2. without sclerite praecoxal/

#2574. <female> Fifth swimming legs praecoxa <fusion to symmetrical legs>/
1. separated from the coxa/
2. fused to coxa/

#2575. <female> Fifth swimming legs praecoxa <condition of the fusion to symmetrical legs>/
1. completely/
2. incompletely/

#2576. <female> Fifth swimming legs praecoxa <ornamentation to symmetrical legs>/
1. without ornamentation/
2. with ornamentation/

#2577. <female> Fifth swimming legs praecoxa <type of ornamentation to symmetrical legs>/
1. of denticles/
2. of spinules/

#2578. <female> Fifth swimming legs praecoxa <location of ornamentation to symmetrical legs>/
1. on anterior surface/
2. on posterior surface/

#2579. <female> Fifth swimming legs coxa <presence of process to symmetrical legs>/
1. without process/
2. with process/

#2580. <female> Fifth swimming legs coxa <form of the process to symmetrical legs>/
1. conical/
2. rounded/

#2581. <female> Fifth swimming legs coxa <location of the process to symmetrical legs>/
1. at the outer rim/
2. at the inner rim/

#2582. <female> Fifth swimming legs coxa <inserted of the process to symmetrical legs>/
1. proximally/
2. medially/
3. distally/

#2583. <female> Fifth swimming legs coxa <presence of sensilla on process to symmetrical legs>/
1. sensilla present/
2. sensilla absent/

#2584. <female> Fifth swimming legs coxa <form of the sensilla on process to symmetrical legs>/
1. stout/
2. slender/

#2585. <female> Fifth swimming legs coxa <location of the sensilla on process to symmetrical legs>/
1. at apex/
2. at nadir/

#2586. <female> Fifth swimming legs coxa <projection of the process to symmetrical legs>/
1. not projecting over basal segment/
2. projecting over basal segment/

#2587. <female> Fifth swimming legs coxa <relative length of the sensilla to insertion basis>/
1. not longer 2x than basal insertion/
2. longer 2x than basal insertion/

#2588. <female> Fifth swimming legs coxa <observation of internal extension of the segment>/
1. marginal extension absent/
2. marginal extension present/

#2589. <female> Fifth swimming legs coxa <form of extension of the segment to symmetrical legs>/
1. triangular/
2. sub-triangular/

#2590. <female> Fifth swimming legs coxa <location of extension of the segment to symmetrical legs>/
1. at inner rim/
2. at inner lateral/

#2591. <female> Fifth swimming legs coxa <insertion of extension of the segment to symmetrical legs>/
1. distally inserted/
2. medially inserted/

#2592. <female> Fifth swimming legs coxa <presence of intumescence to symmetrical legs>/
1. without swelling/
2. with swelling/

#2593. <female> Fifth swimming legs coxa <presence of seta to syimmetrical legs>/
1. without seta/
2. with seta/

#2594. <female> Fifth swimming legs coxa <presence of spinules to syimmetrical legs>/
1. without spinules/
2. with spinules/

#2595. <female> Fifth swimming legs basis <general form to symmetrical legs>/
1. sub-triangular/
2. sub-cylindrical/

#2596. <female> Fifth swimming legs basis <comparison of size between sides>/
1. equal size between inner and outer sides/
2. unequal size between inner and outer sides/

#2597. <female> Fifth swimming legs basis <relation of size between inner and outer side>/
1. shorter outer than inner side/
2. shorter inner than outer side/

#2598. <female> Fifth swimming legs basis <form of the inner side>/
1. with concave inner side/
2. with convex inner side/

#2599. <female> Fifth swimming legs basis <presence of proximal inner outgrowth to symmetrical legs>/
1. without proximal inner outgrowth/
2. with proximal inner outgrowth/

#2600. <female> Fifth swimming legs basis <present of groove>/
1. without groove/
2. with groove/

#2601. <female> Fifth swimming legs basis <ornamentation of the groove>/
1. ornamented/
2. not ornamented/

#2602. <female> Fifth swimming legs basis <presence of distal extension>/
1. without distal extension/
2. with distal extension/

#2603. <female> Fifth swimming legs basis <location of the distal extension>/
1. on lateral surface/
2. on posterior surface/

#2604. <female> Fifth swimming legs basis <presence of seta to symmetrical legs>/
1. without seta/
2. with seta/

#2605. <female> Fifth swimming legs basis <insertion of the seta to symmetrical legs>/
1. outerly inserted/
2. innerly inserted/

#2606. <female> Fifth swimming legs basis <location of the seta to symmetrical legs>/
1. on anterior surface/
2. on posterior surface/

#2607. <female> Fifth swimming legs basis <reach of the seta to symmetrical legs>/
1. not longer 2x than origin segment/
2. longer 2x than origin segment/

#2608. <female> Fifth swimming legs basis <reach of the seta to symmetrical legs>/
1. reaching to exopod 1 distally/
2. not reaching to exopod 1 distally/

#2609. <female> Fifth swimming legs basis <observation of ornamentation with spinules posteriorly>/
1. absence of spinules patch posteriorly/
2. presence of spinules patch posteriorly/

#2610. <female> fifth swimming legs endopod <fusion between segments 1 and 2 to symmetrical legs>/
1. segments 1 and 2 separated/
2. segments 1 and 2 fused/

#2611. <female> fifth swimming legs endopod <fusion between segment 2 and 3 to symmetrical legs>/
1. segments 2 and 3 separated/
2. segments 2 and 3 fused/

#2612. <female> fifth swimming legs endopod <segmentation to symmetrical legs>/
1. 1-segmented/
2. 2-segmented/

#2613. <female> fifth swimming legs endopod <condition of the suture between the segmentation>/
1. with complete suture/
2. with incomplete suture/

#2614. <female> fifth swimming legs endopod <general form to symmetrical legs>/
1. stout/
2. slender/

#2615. <female> fifth swimming legs endopod <fusion to basis of the legs>/
1. separated from the basis/
2. fused to basis/

#2616. <female> fifth swimming legs endopod <fusion condition to basis of the legs>/
1. completely/
2. incompletely/

#2617. <female> fifth swimming legs endopod <cuticle condition at the symmetrical legs>/
1. absent discontinuity cuticle/
2. present discontinuity cuticle/

#2618. <female> fifth swimming legs endopod <location of the condition of the cuticle>/
1. on inner side/
2. on outer side/

#2619. <female> fifth swimming legs endopod <ornamented segment>/
1. ornamentation on segment 1/
2. ornamentation on segment 2/

#2620. <female> fifth swimming legs endopod <type of ornamentation to symmetrical legs>/
1. with spinules/
2. with setules/

#2621. <female> fifth swimming legs endopod <arrangement of the ornamentation to symmetrical legs>/
1. as a row/
2. as a patch/

#2622. <female> fifth swimming legs endopod <number of ornamentation to symmetrical legs>/
1. single/
2. double/

#2623. <female> fifth swimming legs endopod <condition of the spinules or setules row>/

1. non-oblique/
2. oblique/

#2624. <female> fifth swimming legs endopod <insertion of ornamentation to symmetrical legs>/
1. terminally/
2. sub-terminally/

#2625. <female> fifth swimming legs endopod <location of the ornamentation to symmetrical legs>/
1. at anterior surface/
2. at posterior surface/

#2626. <female> fifth swimming legs endopod <presence of setae to symmetrical legs>/
1. with seta/
2. without seta/

#2627. <female> fifth swimming legs endopod <number of setae to symmetrical legs>/
1. single/
2. double/

#2628. <female> fifth swimming legs endopod <insertion of seta 1 to symmetrical legs>/
1. one medially/
2. one distally/

#2629. <female> fifth swimming legs endopod <location of setae 1 to symmetrical legs>/
1. on anterior surface/
2. on posterior surface/

#2630. <female> fifth swimming legs endopod <form of seta 1 to symmetrical legs>/
1. arched/
2. rectilinear/

#2631. <female> fifth swimming legs endopod <insertion of seta 2 to symmetrical legs>/
1. one medially/
2. one distally/

#2632. <female> fifth swimming legs endopod <location of setae 2 to symmetrical legs>/
1. on anterior surface/
2. on posterior surface/

#2633. <female> fifth swimming legs endopod <insertion of seta 2 to symmetrical legs>/
1. arched/
2. rectilinear/

#2634. <female> fifth swimming legs endopod <size between setae to symmetrical legs>/
1. of equal size/
2. of unequal size/

#2635. <female> fifth swimming legs endopod <comparison of size between setae>/
1. distal seta longer than medial seta/

2. proximal seta longer than distal seta/

#2636. <female> fifth swimming legs exopod <fusion between segment 1 and 2 to symmetrical legs>/
1. segments 1 and 2 separated/
2. segments 1 and 2 fused/

#2637. <female> fifth swimming legs exopod <fusion between segments 2 and 3 to symmetrical legs>/
1. segments 2 and 3 separated/
2. segments 2 and 3 fused/

#2638. <female> fifth swimming legs exopod <segmentation to symmetrical legs>/
1. 2-segmented/
2. 3-segmented/

#2639. <female> fifth swimming legs exopod <fusion to symmetrical legs>/
1. separated from the basis/
2. fused to basis/

#2640. <female> fifth swimming legs exopod <condition of the fusion to symmetrical legs>/
1. completely/
2. incompletely/

#2641. <female> fifth swimming legs exopod 1 <general form of the segment to symmetrical legs>/
1. sub-cylindrical/
2. eliptical/

#2642. <female> fifth swimming legs exopod 1 <size relation of the segment to symmetrical legs>/
1. longer than wide/
2. wider than long/

#2643. <female> fifth swimming legs exopod 1 <size proportion of the segment>/
1. not longer or equal than 2 times/
2. longer or equal than 2 times/

#2644. <female> fifth swimming legs exopod 1 <comparison of size between inner and outer sides>/
1. with equal size between inner and outer sides/
2. with unequal size between inner and outer side/

#2645. <female> fifth swimming legs exopod 1 <relation of size between inner and outer sides>/
1. shorter outer than inner side/
2. shorter inner than outer side/

#2646. <female> fifth swimming legs exopod 1 <form of the inner side to symmetrical legs>/
1. with convex inner side/
2. with rectilinear inner side/

#2647. <female> fifth swimming legs exopod 1 <form of the outer side to symmetrical legs>/
1. with convex outer side/
2. with concave outer side/

3. with rectilinear outer side/

#2648. <female> fifth swimming legs exopod 1 <presence of intumescence to symmetrical legs>/
1. without swelling/
2. with swelling/

#2649. <female> fifth swimming legs exopod 1 <presence of extension of the segment to symmetrical legs>/
1. without marginal extension/
2. with marginal extension/

#2650. <female> fifth swimming legs exopod 1 <presence of process in posterior view>/
1. without posterior process/
2. with posterior process/

#2651. <female> fifth swimming legs exopod 1 <local of insertion of the process>/
1. distally/
2. proximally/

#2652. <female> fifth swimming legs exopod 1 <presence of spine to symmetrical legs>/
1. without outer spine posteriorly/
2. with outer spine posteriorly/

#2653. <female> fifth swimming legs exopod 1 <presence of seta to symmetrical legs>/
1. without seta/
2. with seta/

#2654. <female> fifth swimming legs exopod 2 <general form of the segment to symmetrical legs>/
1. sub-triangular/
2. sub-cylindrical/

#2655. <female> fifth swimming legs exopod 2 <size relation of the segment to symmetrical legs>/
1. longer than broad/
2. broader than long/

#2656. <female> fifth swimming legs exopod 2 <size proportion of the segment>/
1. longer or equal than 2 times/
2. no longer or equal than 2 times/

#2657. <female> fifth swimming legs exopod 2 <presence of intumescence to symmetrical legs>/
1. without swelling/
2. with swelling/

#2658. <female> fifth swimming legs exopod 2 <presence of extension of the segment to symmetrical legs>/
1. without marginal extension/
2. with marginal extension/

#2659. <female> fifth swimming legs exopod 2 <presence of process to symmetrical legs>/
1. without process/
2. with process/

#2660. <female> fifth swimming legs exopod 2 <presence of lobe>/
1. without lobe/
2. with lobe/

#2661. <female> fifth swimming legs exopod 2 <presence of spine>/
1. without spine/
2. with spine/

#2662. <female> fifth swimming legs exopod 2 <insertion of spine>/
1. inserted laterally/
2. inserted ventrally/

#2663. <female> fifth swimming legs exopod 2 <form of the spine>/
1. arched/
2. rectilinear/

#2664. <female> fifth swimming legs exopod 2 <direction of the spine>/
1. externally directed/
2. internally directed/

#2665. <female> fifth swimming legs exopod 2 <presence of ornamentation on the spine>/
1. without ornamentation/
2. with ornamentation/

#2666. <female> fifth swimming legs exopod 2 <type of the tip of the spine>/
1. blunt tip/
2. sharp tip/

#2667. <female> fifth swimming legs exopod 2 <relative size of the spine to next segment>/
1. equal size or larger than next segment/
2. smaller than next segment/

#2668. <female> fifth swimming legs exopod 2 <presence of seta to symmetrical legs>/
1. without seta/
2. with seta/

#2669. <female> fifth swimming legs exopod 3 <general form of the segment to symmetrical legs>/
1. cylindrical/
2. sub-cylindrical/

#2670. <female> fifth swimming legs exopod 3 <size relation of the segment to symmetrical legs>/
1. longer than wide/
2. wider than long/

#2671. <female> fifth swimming legs exopod 3 <presence of intumescence to symmetrical legs>/
1. without swelling/
2. with swelling/

#2672. <female> fifth swimming legs exopod 3 <presence of process to symmetrical legs>/
1. without process/
2. with process/

#2673. <female> fifth swimming legs exopod 3 <presence of lobe to symmetrical legs>/
1. without lobe/
2. with lobe/

#2674. <female> fifth swimming legs exopod 3 <presence of spine to symmetrical legs>/
1. without spine/
2. with spine/

#2675. <female> fifth swimming legs exopod 3 <presence of seta to symmetrical legs>/
1. without seta/
2. with seta/

#2676. <female> fifth swimming legs exopod 3 <number of setae to symmetrical legs>/
1. single/
2. double/

#2677. <female> fifth swimming legs exopod 3 <position of the seta to symmetrical legs>/
1. inserted terminally/
2. inserted sub-terminally/

#2678. <female> fifth swimming legs exopod 3 <comparison size of the double seta>/
1. equal size between them/
2. unequal size between them/

#2679. <female> fifth swimming legs exopod 3 <comparison size of the double seta to symmetrical legs>/
1. outer seta smaller than inner/
2. outer seta longer than inner/

#2680. <female> fifth swimming legs exopod 3 <size proportion of the double seta to symmetrical legs>/
1. nearly 1 time/
2. nearly 3 times/

#2681. <female> fifth swimming legs exopod 3 <ornamentation of the external seta>/
1. outer seta not ornamented by setules/
2. outer seta ornamented by setules/

#2682. <female> fifth swimming legs exopod 3 <ornamentation of the seta to symmetrical legs>/
1. without ornamentation/
2. with ornamentation/

#2683. <female> fifth swimming legs exopod 3 <presence of terminal claw to symmetrical legs>/
1. absence of terminal claw/
2. presence of terminal claw/

#2684. <female> fifth swimming legs exopod 3 <rigidity of the terminal claw to symmetrical legs>/

1. not sclerotized/
2. sclerotized/

#2685. <female> fifth swimming legs exopod 3 <general form of the terminal claw in posterior view>/
1. arched/
2. rectilinear/

#2686. <female> fifth swimming legs exopod 3 <direction of the terminal claw in posterior view>/
1. externally directed/
2. internally directed/

#2687. <female> fifth swimming legs exopod 3 <form of the inner side of the terminal claw>/
1. concave inner side/
2. convex inner side/

#2688. <female> fifth swimming legs exopod 3 <presence of ornamentation on the terminal claw>/
1. without ornamentation/
2. with ornamentation/

#2689. <female> fifth swimming legs exopod 3 <type of ornamentation on the terminal claw>/
1. of spinules/
2. of denticles/

#2690. <female> fifth swimming legs exopod 3 <arrangement of the ornamentation on the terminal claw>/
1. as a row/
2. as a patch/

#2691. <female> fifth swimming legs exopod 3 <location of the ornamentation on the terminal claw>/
1. on surface fully/
2. on surface partially/

#2692. <female> fifth swimming legs exopod 3 <location of the ornamentation on the terminal claw>/
1. at medial region/
2. at distal region/

#2693. <female> fifth swimming legs exopod 3 <form of the outer border of the terminal claw>/
1. concave outer side/
2. rectilinear outer side/

#2694. <female> fifth swimming legs exopod 3 <presence of ornamentation on the terminal claw>/
1. without ornamentation/
2. with ornamentation/

#2695. <female> fifth swimming legs exopod 3 <type of ornamentation on the terminal claw>/
1. of spinules/
2. of denticles/

#2696. <female> fifth swimming legs exopod 3 <arrangement of the ornamentation on the terminal claw>/
1. as a row/

2. as a patch/

#2697. <female> fifth swimming legs exopod 3 <ornamentation on the terminal claw>/
1. on surface fully/
2. on surface partially/

#2698. <female> fifth swimming legs exopod 3 <location of the ornamentation on the terminal claw>/
1. at medial region/
2. at distal region/

#2699. <female> fifth swimming legs exopod 3 <type of the tip of the terminal claw>/
1. blunt tip/
2. sharp tip/

#2700. <female> fifth swimming legs exopod 3 <relative size of the terminal claw to origin segment>/
1. 6 times longer than origin segment/
2. not 6 times longer origin segment/
